# Somatic mosaicism in the mature brain reveals clonal cellular distributions during cortical development

**DOI:** 10.1101/2020.08.10.244814

**Authors:** Martin W. Breuss, Xiaoxu Yang, Danny Antaki, Johannes C. M. Schlachetzki, Addison J. Lana, Xin Xu, Guoliang Chai, Valentina Stanley, Qiong Song, Traci Fang Newmeyer, An Nguyen, Beibei Cao, Alexi Nott, Jennifer McEvoy-Venneri, Martina P. Pasillas, Shareef Nahas, Lucitia Van Der Kraan, Yan Ding, NIMH Brain Somatic Mosaicism Network, Christopher K. Glass, Joseph G. Gleeson

**Affiliations:** Department of Neurosciences, University of California, San Diego, La Jolla, CA 92093, USA; Rady Children’s Institute for Genomic Medicine, San Diego, CA 92025, USA; Department of Cellular and Molecular Medicine, University of California, San Diego, La Jolla, CA 92093, USA; Department of Medicine, University of California, San Diego, La Jolla, CA 92093, USA

## Abstract

The structure of the human neocortex underlies species-specific features and is a reflection of intricate developmental programs. Here we analyzed neocortical cellular lineages through a comprehensive assessment of brain somatic mosaicism—which acts as a neutral recorder of lineage history. We employed deep whole genome and variant sequencing in a single *postmortem* neurotypical human brain across 25 anatomic regions and three distinct modalities: bulk geographies, sorted cell types, and single nuclei. We identified 259 mosaic variants, revealing remarkable differences in localization, clonal abundance, cell type specificity, and clade distribution. We identified a set of hierarchical cellular diffusion barriers, whereby the left-right axis separation of the neocortex occurs prior to anterior-posterior and dorsal-ventral axis separation. We also found that stochastic distribution is a driver of clonal dispersion, and that rules regarding cellular lineages and anatomical boundaries are often ignored. Our data provides a comprehensive analysis of brain somatic mosaicism across the human cerebral cortex, deconvolving clonal distributions and migration patterns in the human embryo.

**One Sentence Summary:** Comprehensive evaluation of brain somatic mosaicism in the adult human identifies rules governing cellular distribution during embryogenesis.

---

Somatic mosaicism is a natural phenomenon by which some but not all cells in a tissue harbor a distinguishing mutational variant (*1-3*). These variants often arise in progenitor cells, and consequently are shared by the entire daughter lineage in what is called ‘clonal mosaicism’ (*3*). Although best known for their impact in human diseases, such as in cancers or epilepsy, disease-relevant mutations represent a small fraction of total mosaicism, given that most are neutral ‘passengers’ inscribed during DNA replication (*1, 4*). Sparse and randomly distributed, these variants can serve as labels of clonal expansion to reveal cellular histories and shared lineages in disease, for instance, in cancers (*3, 5, 6*). Increasingly, similar approaches have been applied to a wide range of healthy tissues (*7, 8*), but the sparsity of these neutral somatic variants necessitates deep whole genome sequencing (WGS) and specialized computational approaches to resolve lineage relationships.

The brain is an attractive target for discovery, with a complex developmental history, and largely immobile, non-proliferative cellular structures conferring anatomically distinct cognitive functions (*2*). Indeed, prior animal model studies using engineered mosaicism helped to deconvolve cellular lineages (*9, 10*), but an understanding of cellular lineages in humans requires analysis of natural mosaicism. Recent studies have applied either bulk or single cell sequencing to identify mosaicism as a function of aging or cellular spread across different tissues (*8, 11-13*), but the potential to understand cellular linages in a single human, across the entire neocortex and other organs, has not been appreciated.

Here, using a top-down (i.e. bulk WGS) approach, we studied a single 70-year neurotypical old female who died suddenly of natural causes. The cadaver was accessed with minimal *postmortem* interval to collect the brain, heart, liver, and kidneys. The cerebral cortex was removed and separated into 10 ‘lobes’: 5 for each hemisphere, including prefrontal (PF), frontal (F), parietal (P), occipital (O), and temporal (T) cortex using standard landmarks (*14*). Each lobe was biopsied with 8 mm punches at 13 different locations of defined geometric inter-biopsy distance. The central of these was denoted as ‘Sml’, and the remaining lobar tissue was homogenized and denoted as ‘Lrg’ (Fig. 1A). Cerebellum, heart, liver and both kidneys were also sampled with 8 mm punches.

**Fig. 1.**
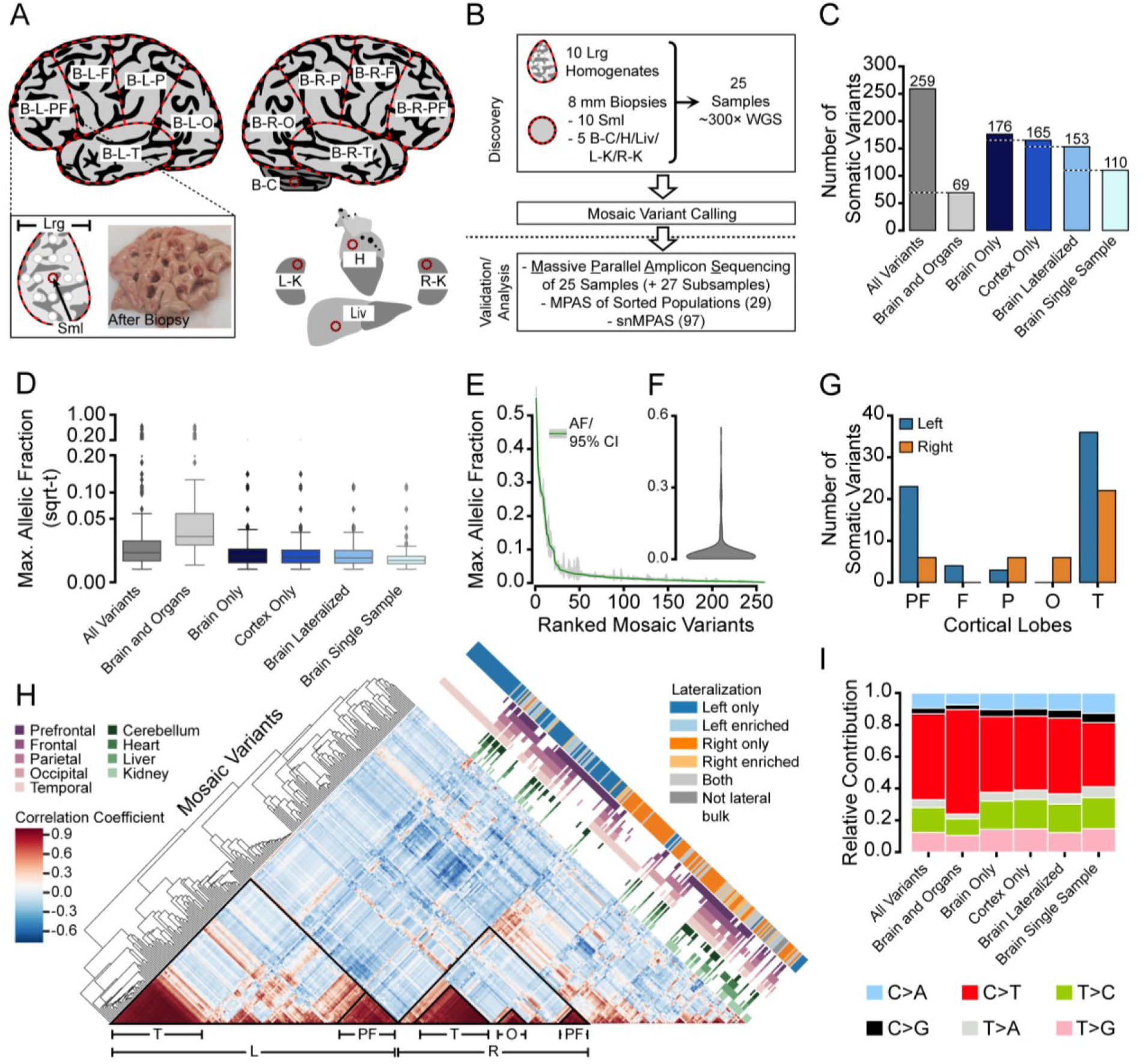
Mosaic variants across a human neocortex are mostly lateralized and region-specific. (**A**) Schematic of the cortical and non-cortical sampled regions. B: brain was divided into 10 lobes for sampling: L: left; R: right; PF: prefrontal cortex; F: frontal cortex; P: parietal cortex; O: occipital cortex; T: temporal cortex. Additionally, C: cerebellum; H: heart; K: kidney; Liv: liver were sampled. Box: schematic and 8 mm punch-biopsy post-sampling of L-PF tissue. Central punch is the Sml and remaining tissue is the Lrg sample. (**B**) Sampling and sequencing strategy. DNA from 8 mm punch from each Sml sample, together with the remaining Lrg sample, and 8 mm punches from other organs underwent 300× deep whole genome sequencing (WGS). Workflow was separated into ‘Discovery’ and ‘Validation/Analysis’ phases. MPAS: massive parallel amplicon sequencing; snMPAS: single nuclei MPAS.(**C**) Number of confirmed and quantified somatic variants in the 25 sequenced tissues. (**D**) Square root transformed (sqrt-t) maximal allelic fraction (AF_max_) across sampled tissues for all 259 mosaic variants. Variants with lower AF_max_ showed more restricted distributions. (**E**-**F**) Ranked (E) and violin (F) plots of the 259 AF_max_ values, with individual values and 95% confidence intervals in E. (**G**) Number of variants found exclusively in Sml biopsies each lobe (total n=106). (**H**) Hierarchical clustering plot showing variant distributions across 25 samples, and categorization based on the lateralization, showing lateralization as the main driver of variant spread, with further clustering into anatomical regions. (**I**) Relative contribution of the six possible base substitutions for variants showing C>T predominance.

For the variant discovery phase, we performed target depth 300× WGS on 25 samples: the Sml and Lrg samples from each of the 10 lobes (Fig. 1B, Fig. S1, A and B), as well as the 5 non-cortical tissues. State-of-the-art mosaicism detection methods for single nucleotide and insertion/deletion variants allowed for identification of candidate variants from WGS (*15, 16*) (Fig. S1C, Data S1). For the variant validation phase, we performed massive parallel amplicon sequencing (MPAS; average depth 7,000×, see Materials and Methods), designed for each variant, to remove false calls and to accurately assess allelic fractions (AF) in each of the original 25 samples (Fig. S2). We subsequently measured each validated variant in a subset (27) of the separate additional lobar tissue biopsies to determine distributions. We also performed MPAS on neuronal and glial populations defined by fluorescence-activated nuclei sorting (FANS) and isolated single nuclei (snMPAS) from lobar tissues to assess cell types and clades (Fig. 1B, Data S2).

Following this detection pipeline, 259 variants were confirmed as mosaic (i.e. significantly above the ∼0.2% AF background noise; 5% FDR) in at least one of the 25 initial samples (see Materials and Methods) (Fig. 1C), and were roughly equally distributed across the genome (Fig. S3). A quarter (69; 26.6%) were detected in the brain and at least one other organ; and consistent with oversampling of the brain, the majority were specific to the neocortex (165; 63.7%). Almost half were found in only one single biopsy (110; 42.5%)—and all but 4 of these in Sml only. The average maximal allelic fraction (across 25 tissues; AF_max_) of the mosaic variants was 3.1% (range: 0.2-55%) (Fig. 1D). In agreement with models in which variants shared in multiple tissues occurred earlier in development (*8*), these variants exhibited a higher mean AF_max_ than those restricted to neocortex (7.8% vs. 1.2%, respectively). Overall, the AF_max_ distribution was strongly biased towards variants of lower AFs (90^th^ percentile: 4.5%; 50^th^ percentile: 1.1%) (Fig. 1, E and F).

There were 106 of the 259 variants that were detected only in a single Sml biopsy (Fig. 1G). Of the 10 Sml biopsies, 7 harbored 6 or fewer mosaic variants; however, L-PF, L-T, and R-T all had in excess of 20. The increased number detected in both temporal biopsies—likely arising independently and relatively late in development—may suggest increased clonal expansion in this region. The large number of variants in these specific 8 mm biopsies was reflected in correlation analysis and hierarchical clustering across all 259 variants; in addition, lateralization to one hemisphere defined mosaic variants present in more than one sample (Fig. 1H).

Mutational signatures—the relative occurrence of the different base-pair substitutions—were similar for all types of brain-restricted variants (Fig. 1I); however, those shared across the body exhibited a higher rate of C>T substitutions. This class of mutation has been correlated with DNA oxidative deamination related to methylation, consistent with an early embryonic origin (*17*).

Next, we assessed the overlap of variants with different genomic annotations, such as epigenetic marks or gene features (Fig. S4). Interestingly, similar to the mutational signatures, putative earlier and brain-organ shared variants exhibited distinct patterns from brain-restricted mutations. For instance, the latter were depleted in areas of H3K27Ac found in stem cells. Together, this combined analysis suggests that most variants are present at AFs < 5% with highly variable distributions across tissues, and further highlights that early tissue-shared variants likely arise through distinct genetic and epigenetic events.

To assess variant ‘geography ‘(i.e. localization across the sampled tissues), we employed ‘geography of clone’ (‘geoclones’) analysis to project MPAS data onto the body plan (Fig. 2A, Data S3). We found that variants of similar AF exhibited distinct geographies, such as those distributed widely or those restricted to the brain, neocortex, one hemisphere, or even a single biopsy (Fig. 2, B to G). Thus variants of similar AF from a Sml biopsy often had dramatically distinct distributions across other tissues (Fig. 2, B to H, Fig. S5A). Variants also often exhibited differences between neighboring tissues, even when widespread across the brain or body; and several of the mutations exhibited ‘discontinuous’ appearances across the hemispheres, cerebellum and other organs (Fig. 2C, Fig. S5B). This suggested that the stochastic distribution of low level, early-occurring clones through partition events may significantly amplify or suppress clonal abundance, acting as a developmental ‘bottleneck’. However, we could not exclude an effect of variants on cellular fitness.

**Fig. 2.**
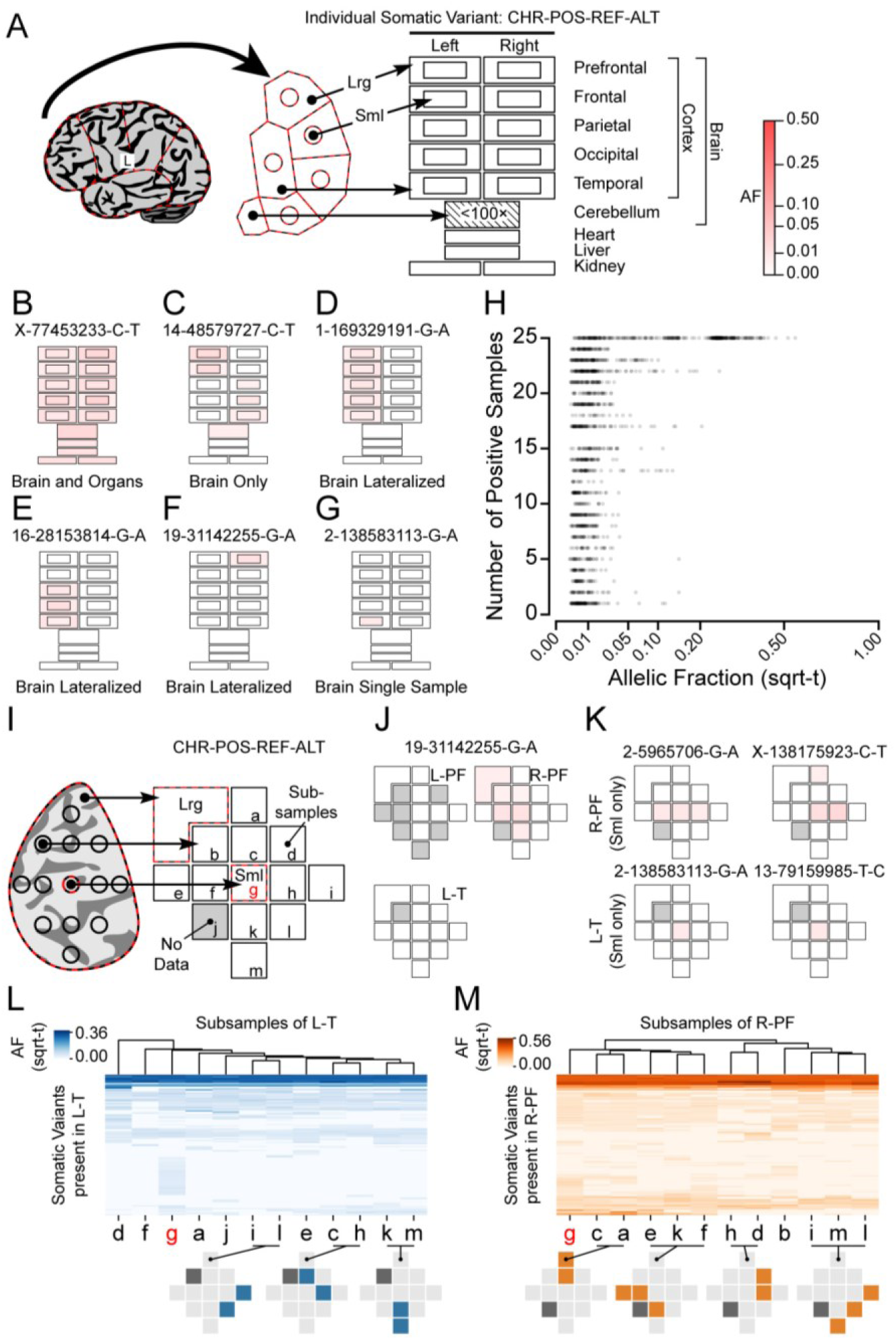
Geography of mosaic variants reveals distributions driven by stochastic and clonal dynamics. (**A**) Projection of sampled regions onto a ‘geography of clone’, or ‘geoclone’. CHR: chromosome; POS: genome position (hg19); REF: reference base; ALT: alternative mosaic base, with intensity representing AF in shaded red. (**B** to **G**) Geoclone examples. Note some geoclones show distributions over the entire body plan while others are highly restricted. (**H**) Scatter plot of all 259 variant and 25 sample mosaic pairs. Note that variants >5% AF typically show wide distributions whereas <5% AF can show restricted or wide distributions. (**I**) Regional subsampling of AF across small lobar biopsy sites. No Data: sample not subjected to MPAS. (**J**) Local spread of one variant restricted to R-PF. (**K**) Local spread of variants within one lobe or just the central punch. (**L** and **M**) AF-based hierarchical clustering of variants and tissues in subsamples in L-T (L) and R-PF (M). Note that adjacent samples tend to cluster together, and that ‘g’, representing the center punch in both lobes, shows the most unique variants, because it was used for ‘discovery’. Below: Locations of variants within the anatomic regions (blue/orange: present; light grey: not detected; dark grey: No Data). For instance, in L, i/l, c/h, k/m cluster together.

To extend the geoclone analysis, we quantified AFs in a subset of additional lobar biopsies (Fig. 2I); for instance, a variant detected only in the Lrg and Sml tissues of R-PF (Fig. 2F) spread exclusively and unevenly in this region (Fig. 2J). This information allowed us to distinguish two scenarios for detected variants specific to only Sml, and absent in the corresponding Lrg biopsies: 1) clones restricted only to the Sml area; 2) clones spread across entire lobes but not represented at sufficient AFs to be detected in the Lrg biopsy by MPAS. Both explanations appear to apply to different clones (Fig. 2K, Fig. S6A).

Stochastic partition and distribution was a likely explanation for the discontinuous evidence for variants across the 25 samples. Hierarchical clustering of variants within these clonal ‘micro-geographies’, however, showed that neighboring small biopsies were more similar to each other than those that were distanced (Fig. 2, L and M, Fig. S6B). This finding is consistent with local spreads—or migration—of clones rather than a combination of multiple bottlenecks. Together, the analysis of broad and local geography of clones assessed through mosaicism revealed geographic differences despite similar AFs. Moreover, these patterns are likely a reflection of both local spread of clones as well as of multiple developmental partitions with resultant stochastic cellular distribution prior to extensive cellular proliferation.

Although analysis of bulk tissue DNA revealed clonal geography, it limited insights into cellular subtypes and lineages. We thus performed FANS on nuclear homogenates of Lrg biopsies (Fig. 3A, Fig. S7) (*18*), employing labeling and sorting strategies enriching for populations of neurons (NeuN^+^), oligodendrocytes (OLIG2^+^), and astrocytes (NeuN^-^/LHX2^+^) for most of the targeted samples (Fig. 3B, Fig. S8-10). In addition to these brain-derived cells, for two regions (R-PF and R-T) we also obtained samples enriched for microglia (PU.1^+^), which have a distinct developmental origin (Fig. 3B, Fig. S8-10) (*19*).

**Fig. 3.**
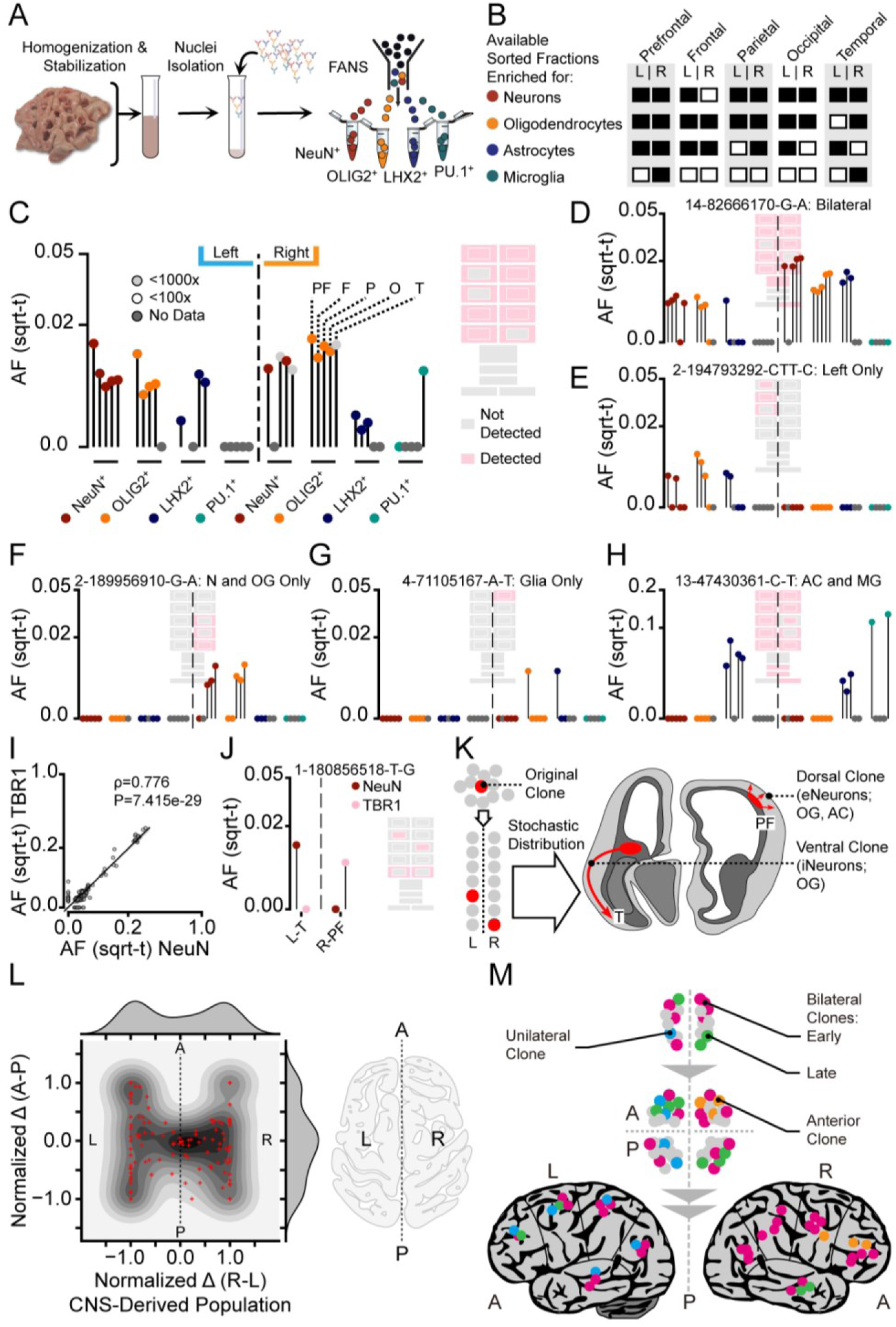
Analysis of sorted cell types suggests stochastic distribution of clones across the body axes and within brain. (**A**) Workflow of fluorescence-activated nuclei sorting (FANS) to isolate nuclei from homogenates (Lrg) and to enrich for populations of neurons (NeuN^+^), oligodendrocytes (OLIG2^+^), astrocytes (NeuN^-^/LHX2^+^), or microglia (PU.1^+^). (**B**) Available sorted populations from cortical areas. Black: available; White: quantity/quality not sufficient. (**C**) Hypothetical variant in ‘lolliplot’, showing AF across anatomic regions (PF, F, P, O, T) and cell types (at bottom) in left vs. right of sampled tissues. (**D**-**H**) Individual lolliplots with distinct anatomic distributions and cellular contributions, with a geoclone representation projected behind. For instance, H shows a bilateral astrocyte clone that is also present in the tested microglia populations. N: neurons; OG: oligodendrocytes; AC: astrocytes; MG: microglia. (**I**) Correlation of AFs for NeuN^+^ and TBR1^+^ populations in L-T and R-PF. Spearman correlation’s ρ and P-value document that the two populations show high correlation overall, but not for all variants. (**J**) Lolliplots of the AF in NeuN^+^ and TBR1^+^ cells and an absence/presence geoclone for 1-180856518-T-G in L-T and R-PF. The two regions show inverse patterns for neuronal markers. (**K**) Proposed mechanisms of stochastic distribution, resulting in ventral clone placement in the left T anlage and dorsal placement in right PF anlage, with a concomitant enrichment for the respective neuronal cell types. i/eNeurons: inhibitory/excitatory neurons. (**L**) Contour plot with individual data points and two kernel density estimation plots for normalized difference of mosaic variants between average AF of sorted brain-derived cells (i.e. non-PU.1+) on the left and right (L, R) hemispheres (Normalized Δ; see Materials and Methods) and between anterior (PF, F) and posterior (P, O, T) brain regions (A, P). The ‘H’ shape of the contour supports a clonal diffusion barrier (CDB) occurring between L-R tissue prior to A-P tissue in the neocortex. (**M**) Proposed order of bilateral and anterior-posterior CDBs during development. Unilateral and bilateral clones can be distinguished, but anterior (or posterior) clones mostly arise secondarily.

Overall MPAS results on sorted population resembled those from bulk tissues; and most signal originated from brain-derived cell types (Fig. 3, C to G, Fig. S11, A to C, Data S3). In addition, sorted population analysis distinguished distinct clonal patterns: for instance, through variants absent versus present in the astrocyte-enriched population. Based on current models of human brain development, these differences might represent ventral and dorsal cortical clones, respectively, as only the latter are thought to produce local astrocytes (*20*). These observations were supported by the overall correlation of AFs within brain regions for these populations (Fig. S11, D to F). Here, neurons showed higher correlations with putative oligodendrocytes than astrocytes. For multiple variants, populations of astrocytes with bilateral representation correlated with microglia (Fig. 3H, Data S3). This observed pattern was reflected in the overall marker correlation: of all brain-derived cells, the microglia-enriched population correlated best with the astrocyte-enriched fraction (Fig. S11, G to I). As there are no known biological populations that overlap these markers, this result might suggest a low-level clone under positive selection in two proliferative populations; although we cannot exclude that the mutation arose separately in two lineages, or that there was death of sister clones.

Analysis of inter-variant correlation based on the sorted populations revealed distinct but overlapping patterns when compared to the correlations obtained from the 25 bulk tissues (Fig. 1H, Fig. S12). Notably, the similarities were mainly derived from L-R asymmetries, and, indeed, many of the variants shared across the hemispheres or with the cerebellum or other organs (n=82) showed a visibly asymmetric bilateral distribution (Fig. 3D, Fig. S8A, Data S3). A combined analysis revealed that 31 variants were significantly enriched by at least 2-fold in either hemisphere (L: 15; R: 16) (Fig. S13A). As expected from a stochastic distribution of mosaic populations, asymmetric variants were more likely to be of lower AF and did not show an overall preference for either hemisphere (Fig. S13B). Assuming that clones were randomly restricted to the separated halves of the progenitors before left-right partition in the neural anlage, we calculated a maximal population size that would support the observed, stochastic differences (see Materials and Methods; Fig. S13, C and D): this approach estimated that the population destined to form the neocortical areas prior bilateral partition is at most ∼160-211 cells (Fig. S13, F to G). Using the highest average AF of hemisphere-specific variants, we also estimate that the population consists of at least ∼86 cells (Fig. S13, E and H). Thus, prior to the terminal left-right split the neocortical anlage comprises an estimated ∼ 90-200 cells.

A combined analysis of variants through mosaic abundance in bulk tissues and sorted populations by UMAP confirmed that the left-right split was the major determinant of variant and sample clustering (Fig. S14, A to I), and, moreover, subsamples of the same lobe clustered together (Fig. S14, H and I). Overall, assessment across bulk and sorted populations confirmed the results derived from the individual analyses.

The presence of certain variants in both neurons and oligodendrocytes but not astrocytes suggested the detection of both ventral and dorsal clones. To test this we performed additional FANS in two brain regions (L-T and R-PF), employing antibodies against TBR1, a marker for excitatory neurons, exclusively derived from dorsal radial clones (*21*). Overall, AFs of NeuN^+^ and TBR1^+^ populations exhibited positive correlation (Spearman’s ρ=0.776; P=7.415e-29), suggesting that most clones were present in ventral and dorsal progenitors (Fig. 3I). Consistent with an additional, independent clonal source of oligodendrocytes, they showed lower correlation than the putative astrocytes with TBR1^+^ cells (Fig. S9-10, J and K); and the latter correlation was higher than the one with the NeuN^+^ population.

Several mosaic variants exhibited absence in one of the two neuronal marker populations, possibly due to locally related clones that contribute predominantly to ventrally- or radially-derived neurons (Data S3). An intriguing example was a cortically-restricted, bilateral geoclone (1-180856518-T-G) that was only positive for NeuN^+^ but not TBR1^+^ in L-T, but showed the inverse pattern in R-PF (Fig. 3J). This finding suggested a low-abundance clone at the time of left-right partition that was stochastically restricted onto the ventral area of the left temporal lobe anlage and onto the dorsal area of the right prefrontal lobe anlage (Fig. 3K). Together, our results suggested that L-R boundary formation occurs first, followed by the anterior-posterior (A-P) and then the dorsal-ventral (D-V). Indeed, a combined analysis confirmed this hypothesis, as A-P asymmetries were dependent on a prior L-R asymmetry break (Fig. 3L-M).

To complement the data derived from bulk geographies and FANS-based genotyping, we additionally performed single nuclear genotyping, extracted from a subsample of the Sml L-T biopsy, using single nuclei whole genome amplification followed by snMPAS on 48 NeuN^+^/DAPI^+^ and 47 NeuN^-^/DAPI^+^ nuclei (Fig. 4A, Fig S15-16, Data S4, Materials and Methods). A double-ranked plot of high-confidence variants (∼1% FDR, Materials and Methods) revealed three widespread, major clonal clades with peak bulk tissue AFs between ∼9-13% (Fig. 4B-C). Based on their abundance, the founder variants of clades I and II likely occurred during the four-cell stage, whereas clade III occurred either at the four-cell stage with relative depletion or at the eight-cell stage with relative amplification (Fig. 4D). We favor the latter hypothesis, as clade I and II are robustly negatively correlated, whereas clade III is poorly correlated with Clade I (Fig. S17). An algorithm-based lineage analysis using the observed variants as input (Fig. 4B) largely agreed with this classification (Fig. S18A) (*22*). Additional manual analysis and placement of the distinct clades onto the UMAP variant plot suggested that these early clones showed little bias towards their later geographical localization and that stochastic distribution during partition acted independently of clonal lineage (Fig. S18B and S19). Indeed, a separate lineage reconstruction and relative contribution analysis for these clades supported their widespread distribution across bulk tissues and sorted populations (Fig. 4E, Fig. S20).

**Fig. 4.**
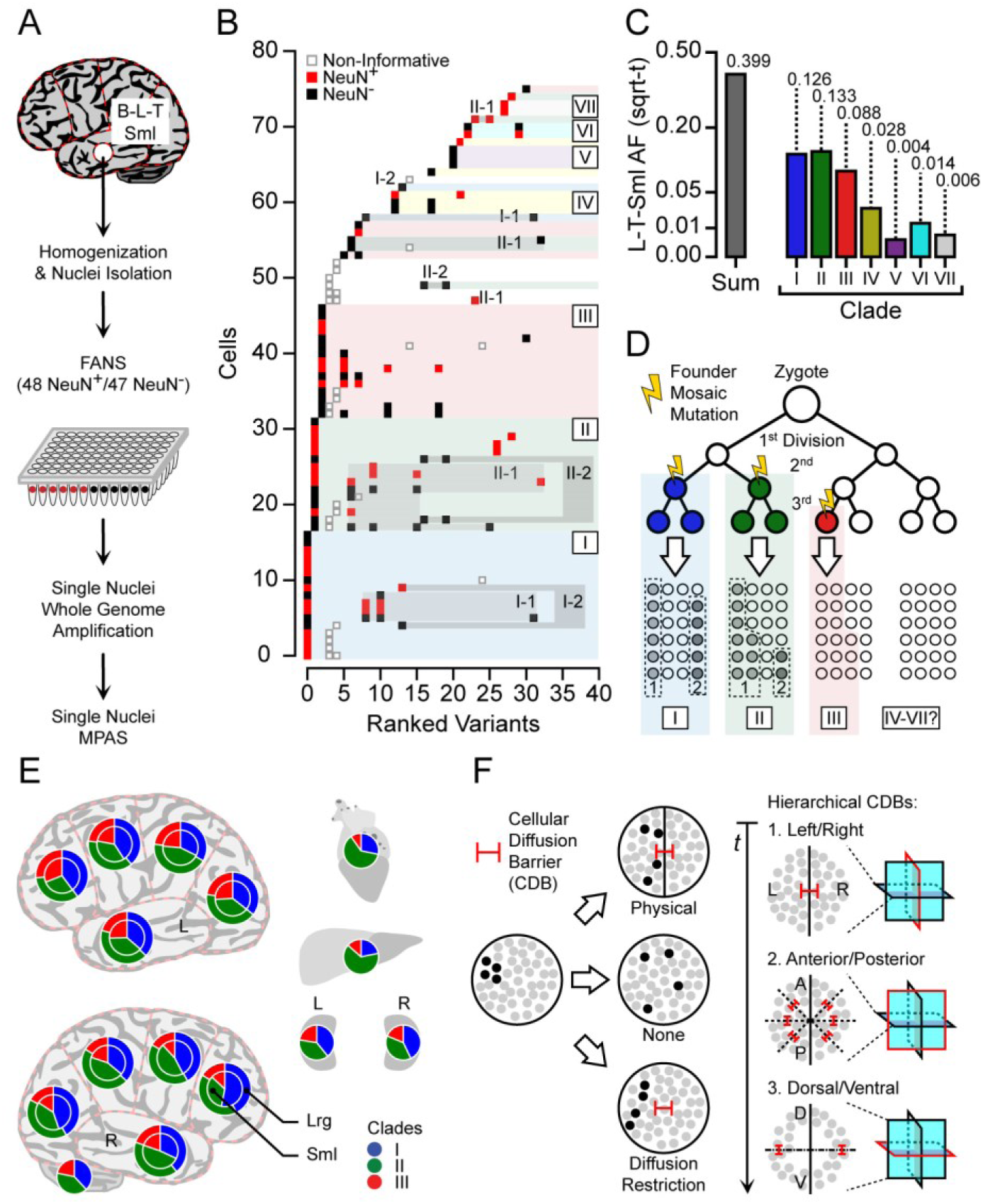
Single nuclei genotyping of mosaic variants resolves cellular lineage and its interaction with stochastic distribution. (**A**) Single nuclear MPAS workflow: single nuclei from L-T (Sml) were sorted into a 96-well plate, then DNA was amplified and subjected to MPAS. (**B**) Ranked plot of filtered, mosaic variants (n=33) and the cells in which they were detected (n=76). Grey edges: non-informative variants; their placement across clades was inconsistent with other clades and their own AF across tissues; likely caused by genotyping errors. Seven clades (I-VII) and likely sub-clades in I and II (I-1, I-2, II-1, and II-2) were detected. The majority of cells belonged to three major clades: I-III. (**C**) Observed AF in L-T-Sml for each of the founder variants in the clades I-VII (i.e. the left-most variant in each clade) was consistent with detection in single nuclei. (**D**) Proposed origin of the three major clades (and their founder mutations) during early embryonic divisions based on the observed AF, placing the founder variants of clades I and II at ∼4-cell stage, and clade III at ∼8-cell stage. (**E**) Relative contribution of the three major clades (I-III) using nested pie charts (Sml: inner; Lrg: outer). The major clades were found in all tissues, but show variation in the relative contributions. (**F**) Model for cellular diffusion barriers (CDBs). Without any CDB (‘None’) cells diffuse or migrate freely. The presence of a CDB restricts clonal exchange and leads to differential clonal abundance. Model of hierarchical CDBs, following in order: left-right, anterior-posterior, and dorsal-ventral. t: time. Red lines: orientation of CDB. Dashed lines: visual plane in the schematic.

Together, our approach provides a comprehensive analysis of brain somatic mosaicism across the human cerebral cortex. The experimental design, by its nature, identified early mosaic mutational events present in many cells within a region, while still capturing events that were exclusively present within single 8 mm biopsies estimated to contain ∼50M cells (*23*). We propose that cellular diffusion barriers (CDBs) are the main driver of the stochastic distribution; and that these form in a hierarchical fashion where a left-right CDB occurs prior to anterior-posterior and dorsal-ventral CDBs (Fig. 4F).

While analyses of clones in model organisms allowed the dissection of cellular distributions in the cortex, prior studies were limited to time points following hemispheric partition (*9, 10, 24*); and thus the consideration of clonal relationships occurring prior to hemispheric partitioning, especially in human brain, represents a conceptual advance. Many of our observations confirm insights derived from prior animal studies, such as the distinction between ventral and dorsal clones, and shared lineage between cortical cells.

Our data raise interesting questions that could be addressed through the evaluation of additional samples using our methods. For instance, it is unclear whether the bilateral enrichment of local mosaic variants in both temporal lobes was related to developmental dynamic expansion as has been proposed (*25*). Similarly, the peculiar correlation of bilateral astrocyte-enriched cell fraction with putative microglia may be of significance beyond stochastic clonal amplification. Additional samples would further add to catalogue of clonal distributions that we present, which could aid in the interpretation of clinically-relevant mosaic mutations. Nevertheless, our comprehensive assessment of mosaic mutations across the neocortex provides a framework with which current analyses of brain mosaicism resulting from brain tumors and other pathological conditions may be compared.

## Acknowledgments

The authors wish to thank the individual donor for tissues for this project, and to thank the BSMN, Dr. Sangmoon Lee, Dr. Changuk Chung, and Isaac Tang (UCSD) for feedback and suggestions. Sequencing was provided by the Rady Children’s Institute for Genomic Medicine. We thank R. Sinkovits, A. Majumdar, S. Strande at the San Diego Supercomputer Center;

## Funding

M.W.B. was supported by an EMBO Long-Term Fellowship (no. ALTF 174-2015), co-funded by the Marie Curie Actions of the European Commission (nos. LTFCOFUND2013 and GA-2013-609409), and an Erwin Schrödinger Fellowship by the Austrian Science Fund (no. J 4197-B30). This study was supported by grants to J.G.G. from NIMH 1U01 MH108898 parent and supplement grant, and to CKG from NIA RF1 AG061060-02, R01 AG056511-02, R01 NS096170-04);

## Author contributions

M.W.B., X.Y., D.A., J.C.M.S., and J.G.G. conceived the project and designed the experiments. M.W.B., X.Y., J.C.M.S., A.J.L., G.C., Q.S., T.F.N., A. Nott, and M.P.P. performed the experiments. X.Y., D.A., X.X., M.W.B., J.C.M.S., A. Nguyen, and B.C. performed the bioinformatics and data analyses. M.W.B., X.Y., V.S., J.M-V., S.N., L.V.D.K., and Y.D. organized, handled, and sequenced human samples. J.G.G. and C.K.G. provided financial and laboratory resources and supervised the project. M.W.B., X.Y., D.A., J.C.M.S., and J.G.G. wrote the manuscript. All authors saw and commented on the manuscript before submission.

## Competing interests

Authors declare no competing interests;

## Data and materials availability

Raw whole genome sequencing and massive parallel amplicon sequencing data is available through NDA (NDA study #919). Summary tables of the data are included as supplementary data. Critical processing and analysis pipelines, as well as custom scripts are available online (https://github.com/shishenyxx/Adult_brain_somatic_mosaicism). Other materials or software are available through the authors upon reasonable request.

## Notes

### Competing Interest Statement

The authors have declared no competing interest.

