## Supplementary material for "Somatic mosaicism in the mature brain reveals clonal cellular distributions during cortical development": Data S3

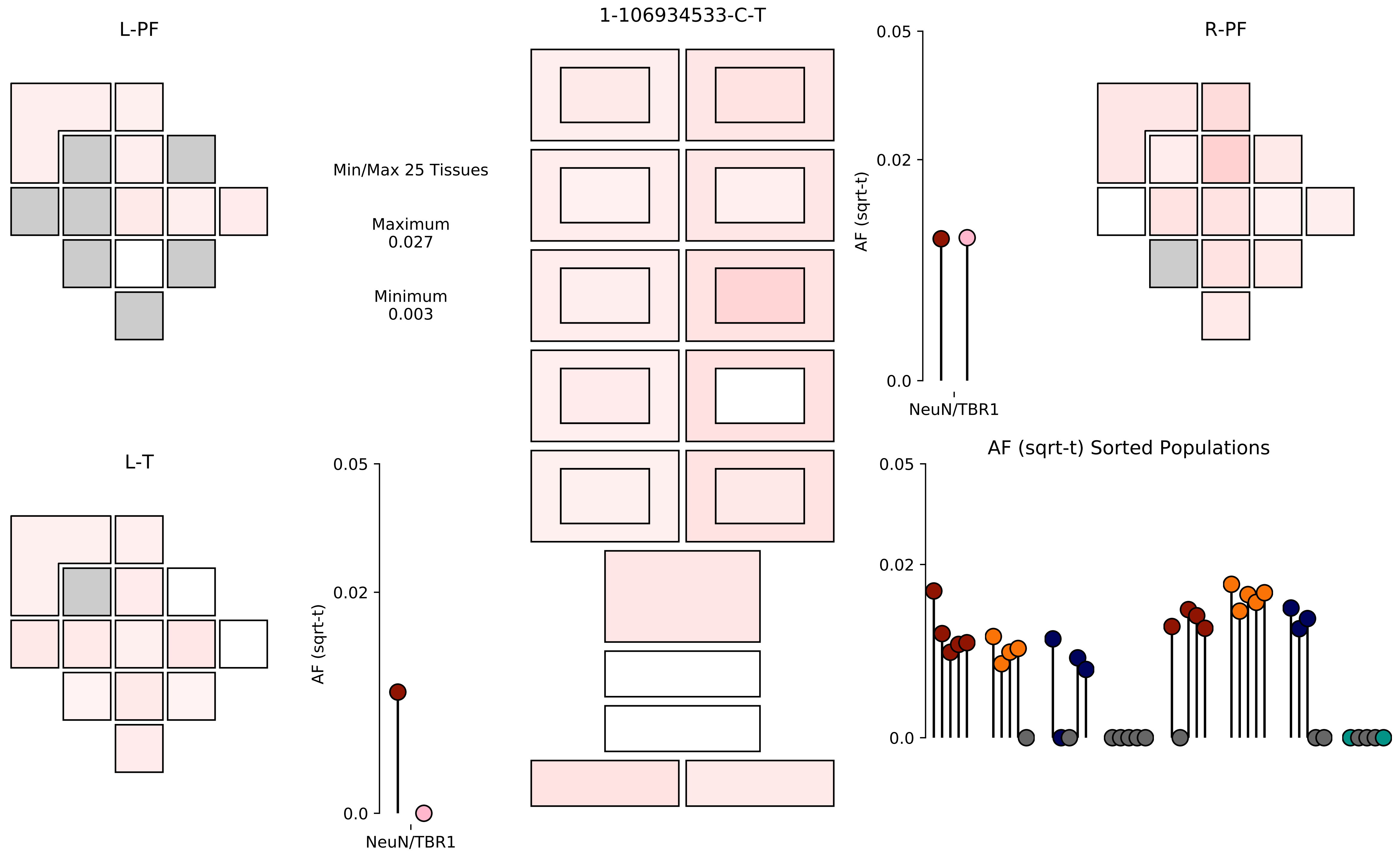
1-106934533-C-T
L-PF
R-PF
L-T
Min/Max 25 Tissues
Maximum
0.027
Minimum
0.003
AF (sqrt-t)
0.05
0.02
0.0
NeuN/TBR1
AF (sqrt-t) Sorted Populations

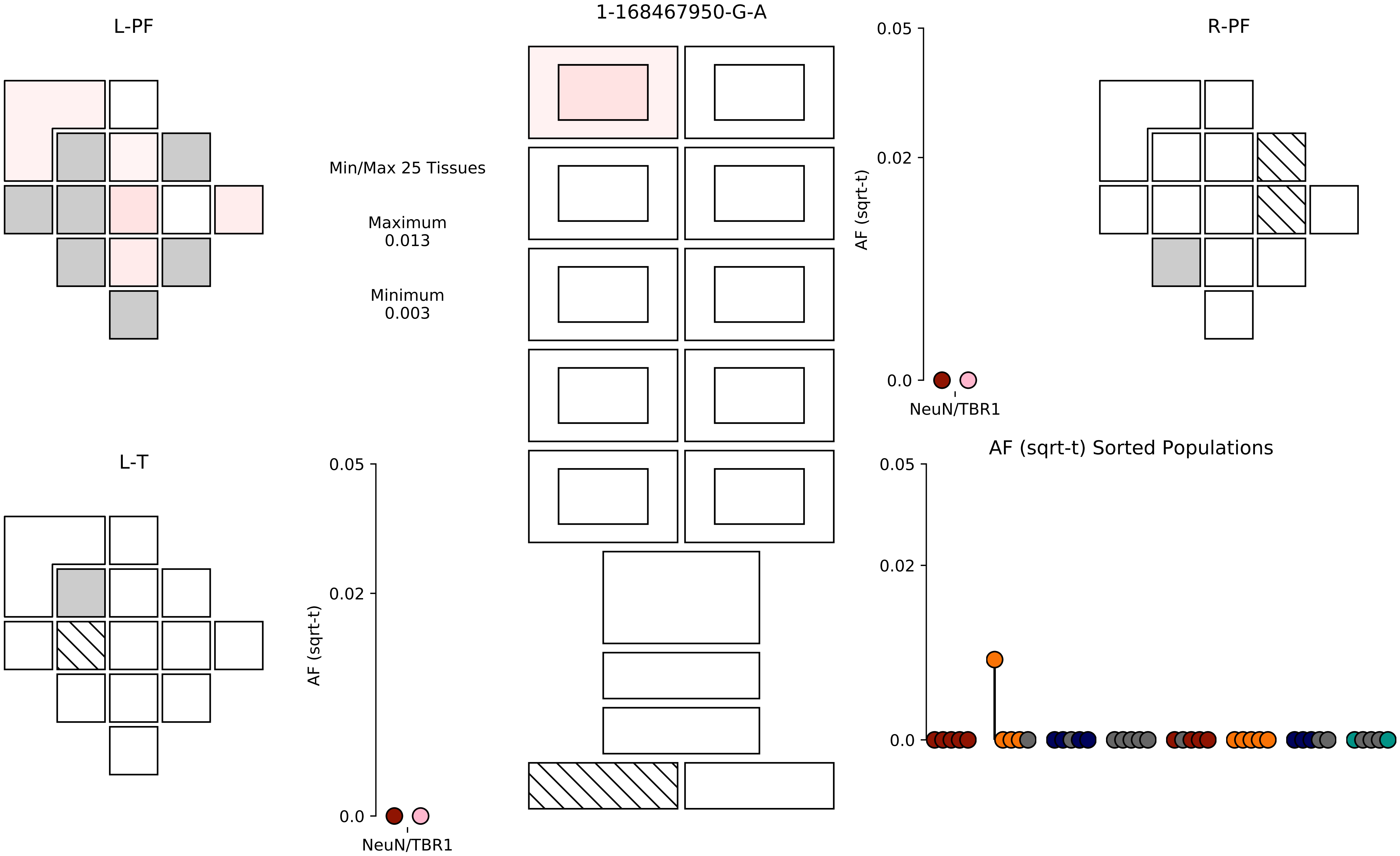
L-PF
1-168467950-G-A
R-PF
Min/Max 25 Tissues
Maximum
0.013
Minimum
0.003
0.05
0.02
0.0
AF (sqrt-t)
NeuN/TBR1
L-T
0.05
0.02
0.0
AF (sqrt-t)
NeuN/TBR1
AF (sqrt-t) Sorted Populations
0.05
0.02
0.0

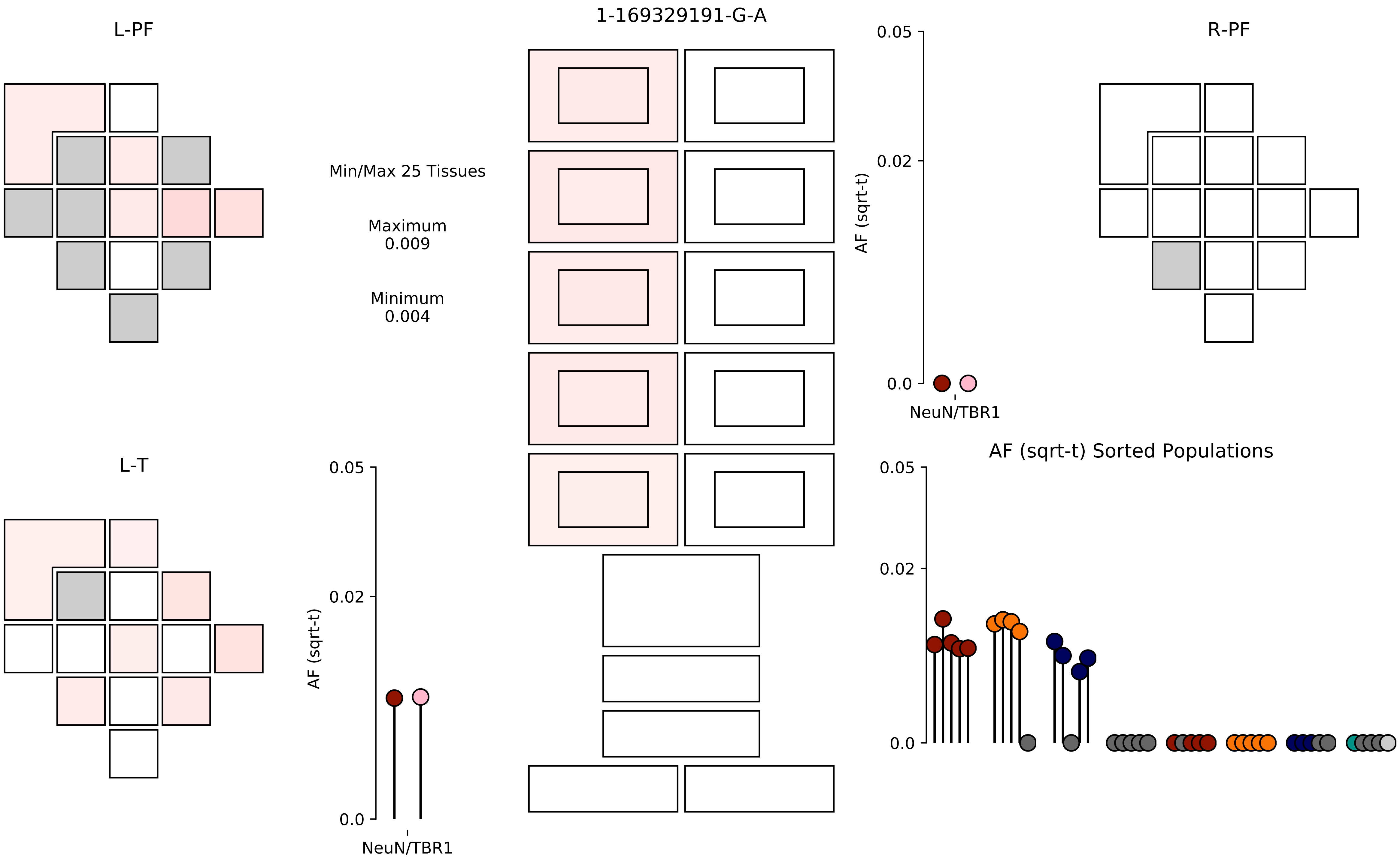

1-169329191-G-A
L-PF
R-PF
L-T
Min/Max 25 Tissues
Maximum
0.009
Minimum
0.004
AF (sqrt-t)
0.05
0.02
0.0
NeuN/TBR1
AF (sqrt-t) Sorted Populations

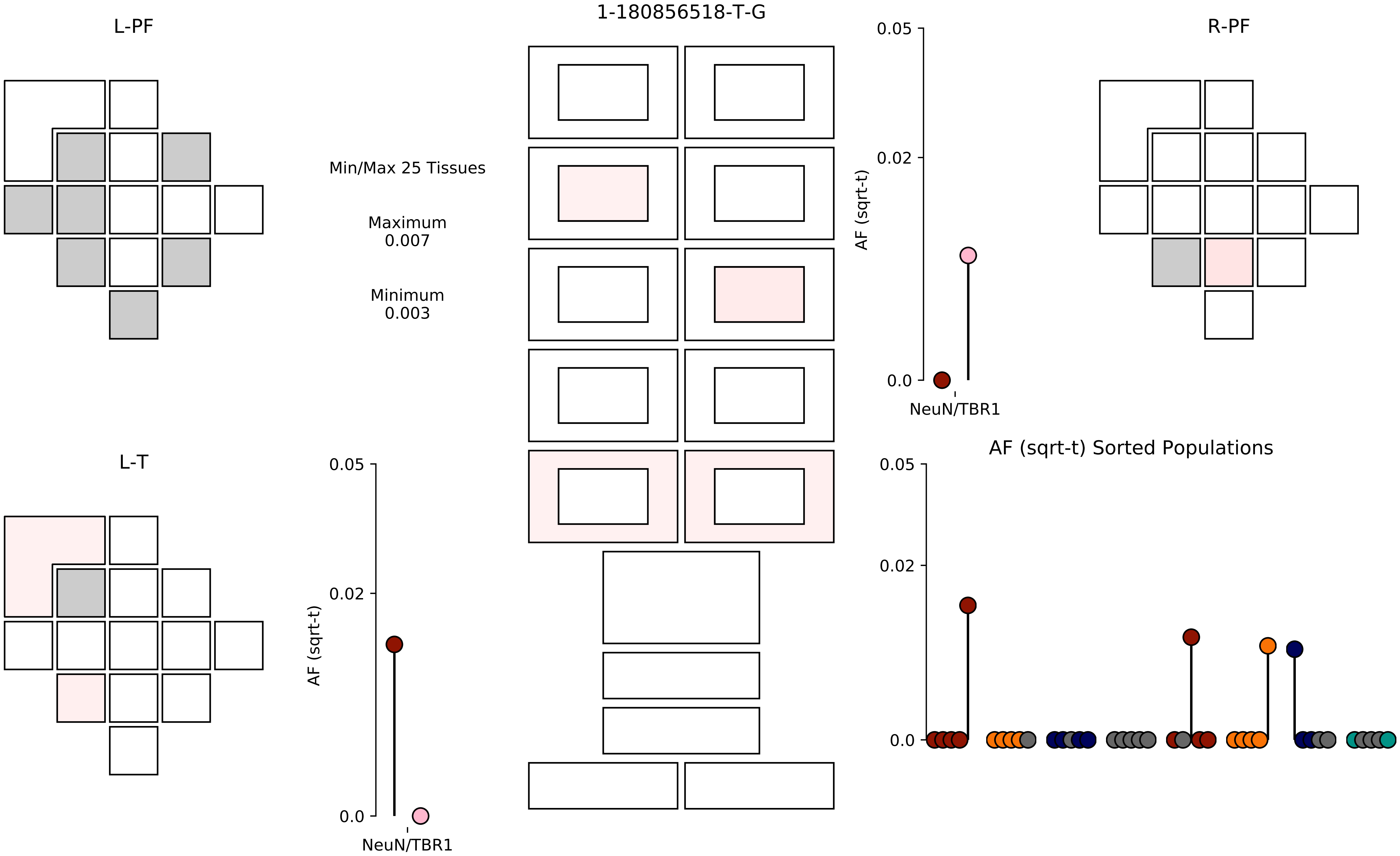

1-180856518-T-G
L-PF
R-PF
L-T
Min/Max 25 Tissues
Maximum
0.007
Minimum
0.003
AF (sqrt-t)
0.05
0.02
0.0
NeuN/TBR1
AF (sqrt-t) Sorted Populations

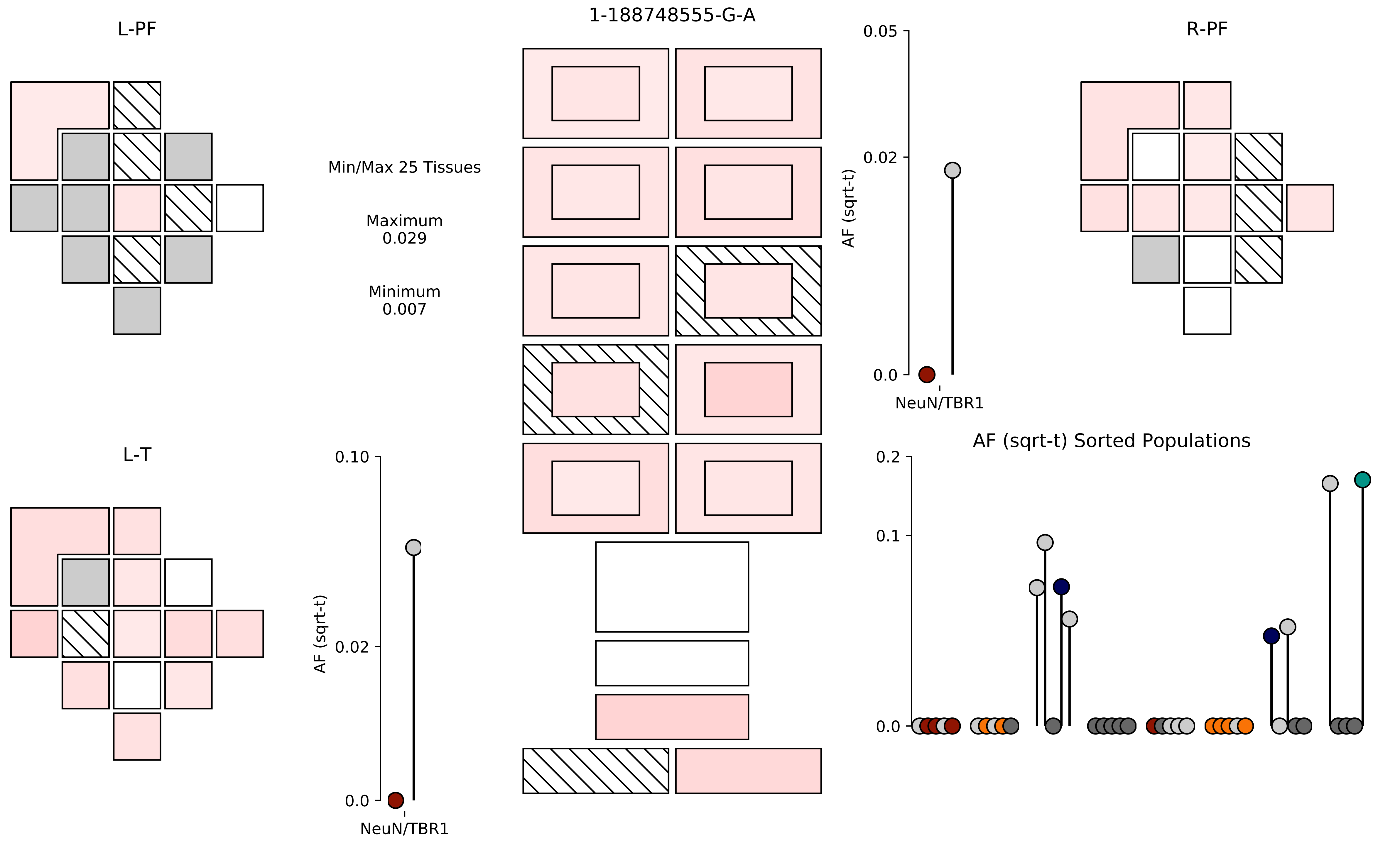
L-PF
1-188748555-G-A
R-PF
Min/Max 25 Tissues
Maximum
0.029
Minimum
0.007
0.05
0.02
0.0
AF (sqrt-t)
NeuN/TBR1
L-T
0.10
0.02
0.0
AF (sqrt-t)
NeuN/TBR1
AF (sqrt-t) Sorted Populations
0.2
0.1
0.0

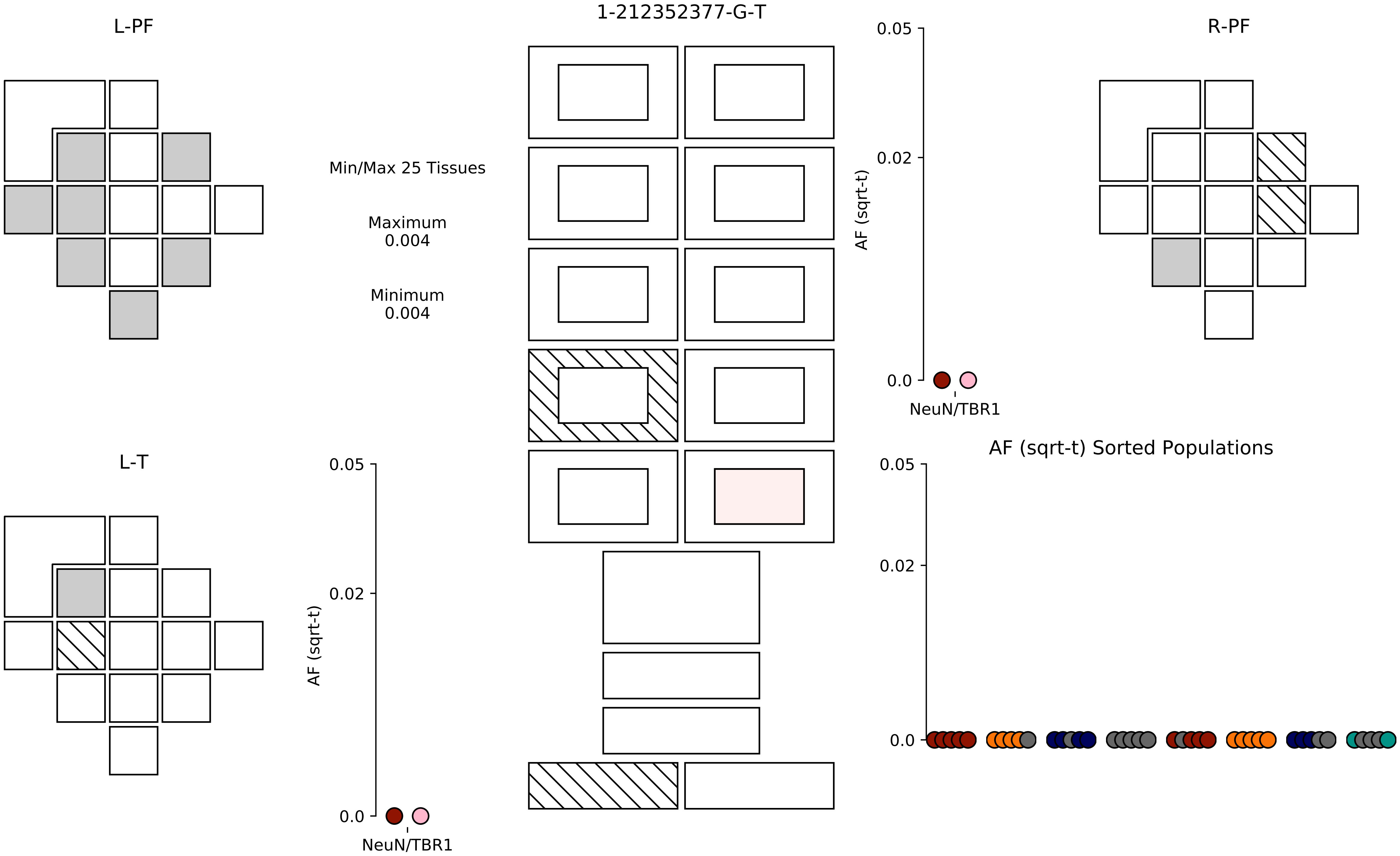
L-PF
1-212352377-G-T
R-PF
Min/Max 25 Tissues
Maximum
0.004
Minimum
0.004
0.05
0.02
0.0
AF (sqrt-t)
NeuN/TBR1
L-T
0.05
0.02
0.0
AF (sqrt-t)
NeuN/TBR1
AF (sqrt-t) Sorted Populations
0.05
0.02
0.0

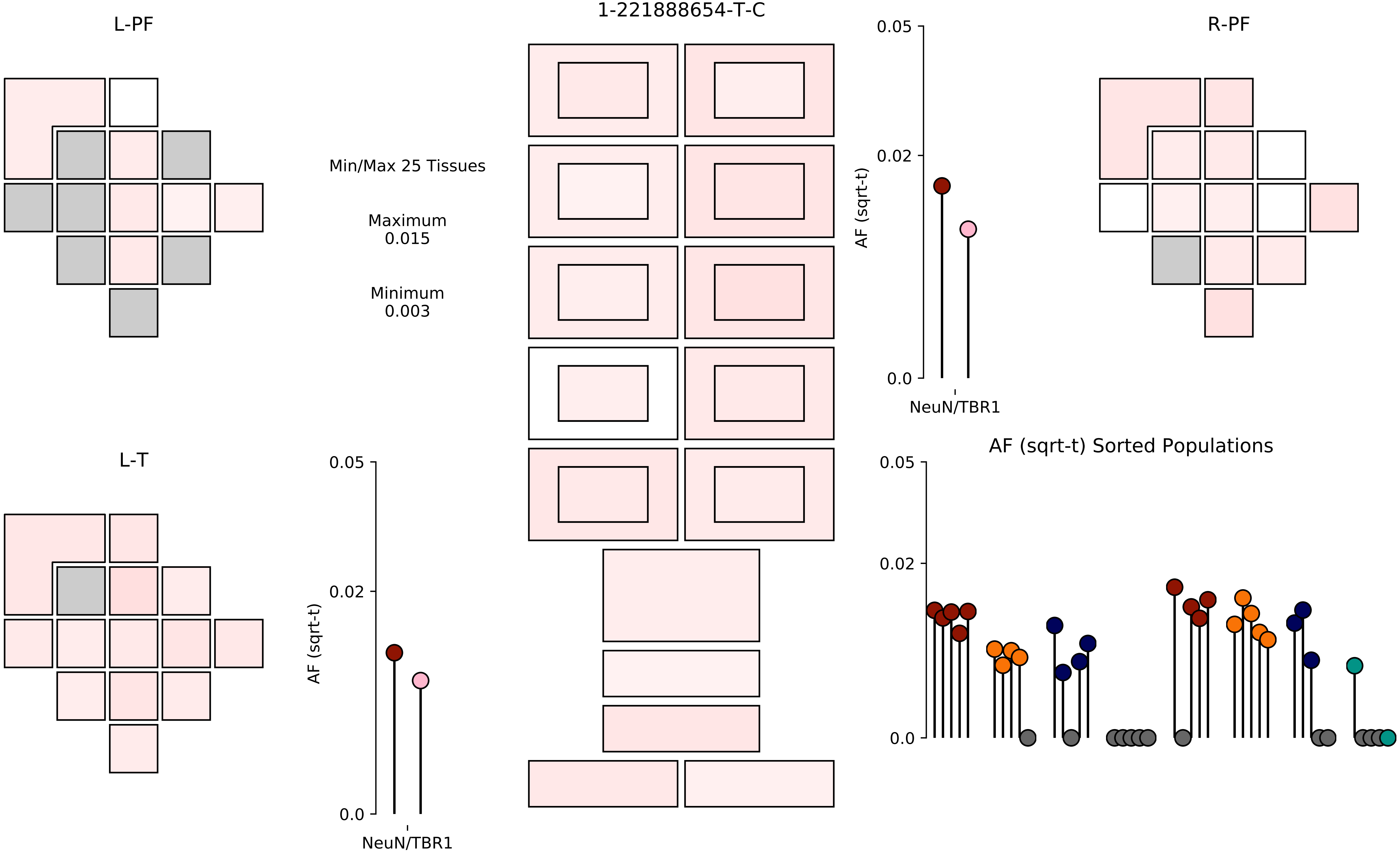

L-PF
1-221888654-T-C
R-PF
Min/Max 25 Tissues
Maximum
0.015
Minimum
0.003
0.05
0.02
0.0
AF (sqrt-t)
NeuN/TBR1
L-T
0.05
0.02
0.0
AF (sqrt-t)
NeuN/TBR1
AF (sqrt-t) Sorted Populations
0.05
0.02
0.0

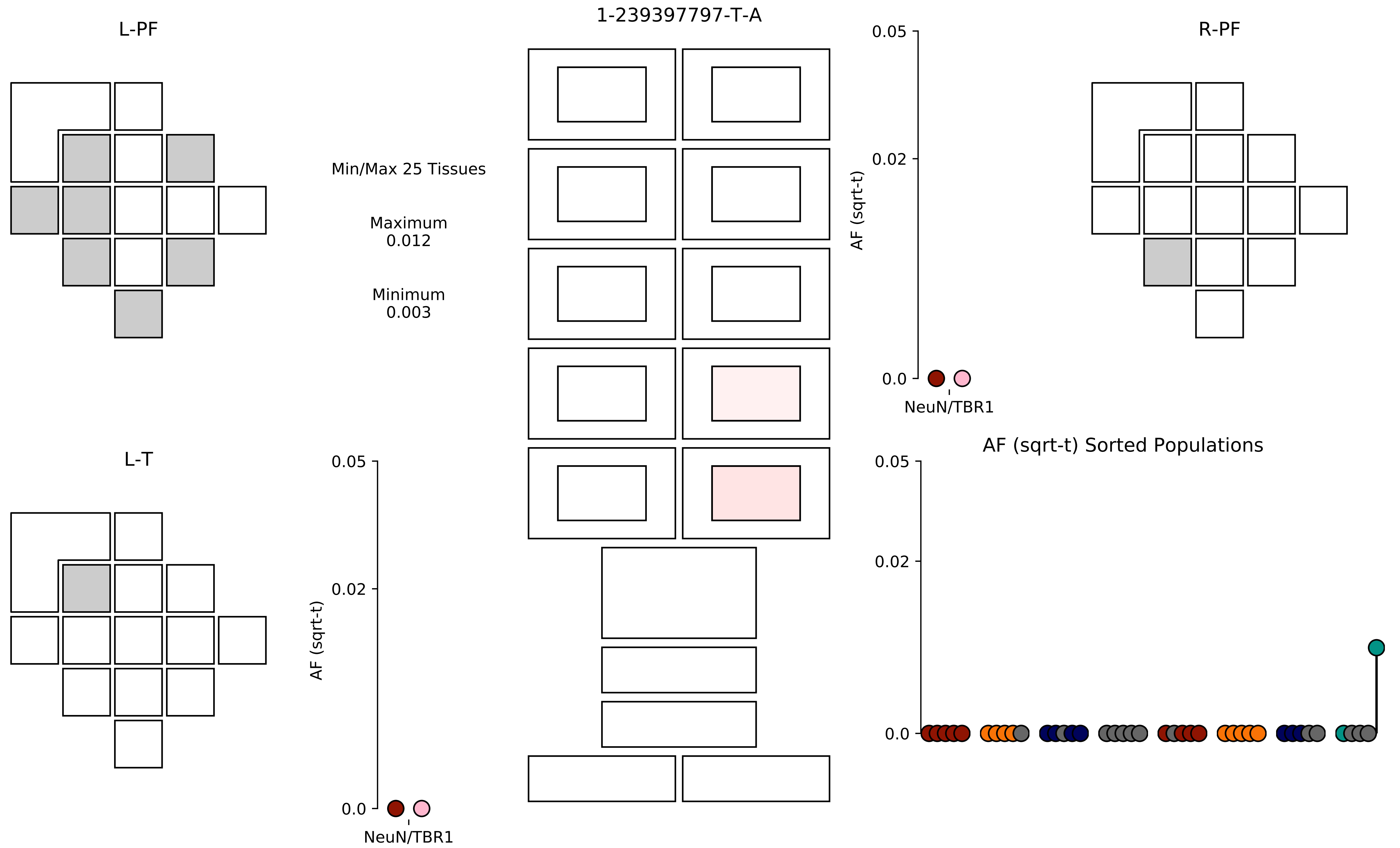
L-PF
1-239397797-T-A
R-PF
Min/Max 25 Tissues
Maximum
0.012
Minimum
0.003
0.05
0.02
0.0
AF (sqrt-t)
NeuN/TBR1
L-T
AF (sqrt-t) Sorted Populations

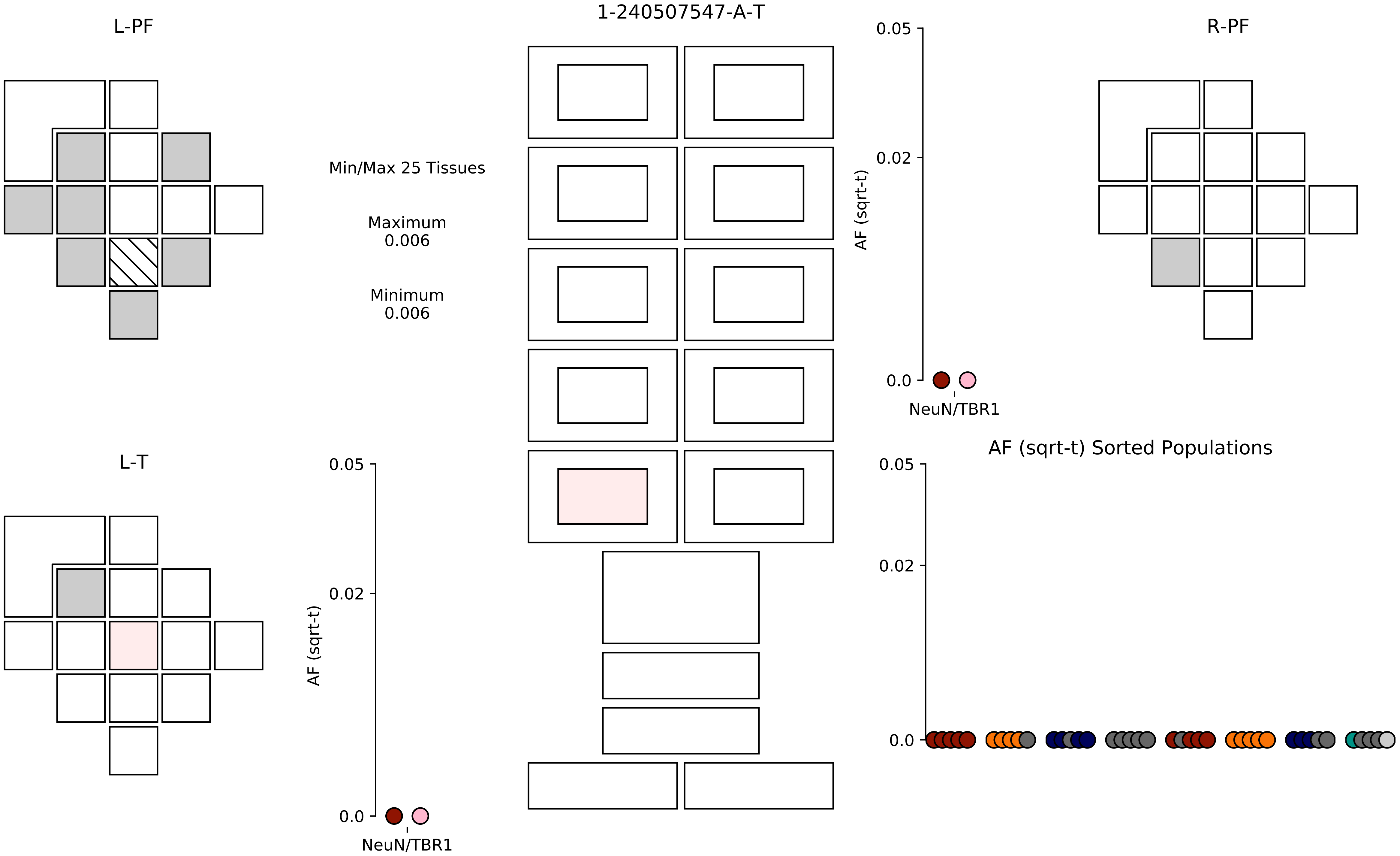

1-240507547-A-T
L-PF
R-PF
L-T
Min/Max 25 Tissues
Maximum
0.006
Minimum
0.006
AF (sqrt-t)
0.05
0.02
0.0
NeuN/TBR1
AF (sqrt-t)
0.05
0.02
0.0
NeuN/TBR1
AF (sqrt-t) Sorted Populations
0.05
0.02
0.0

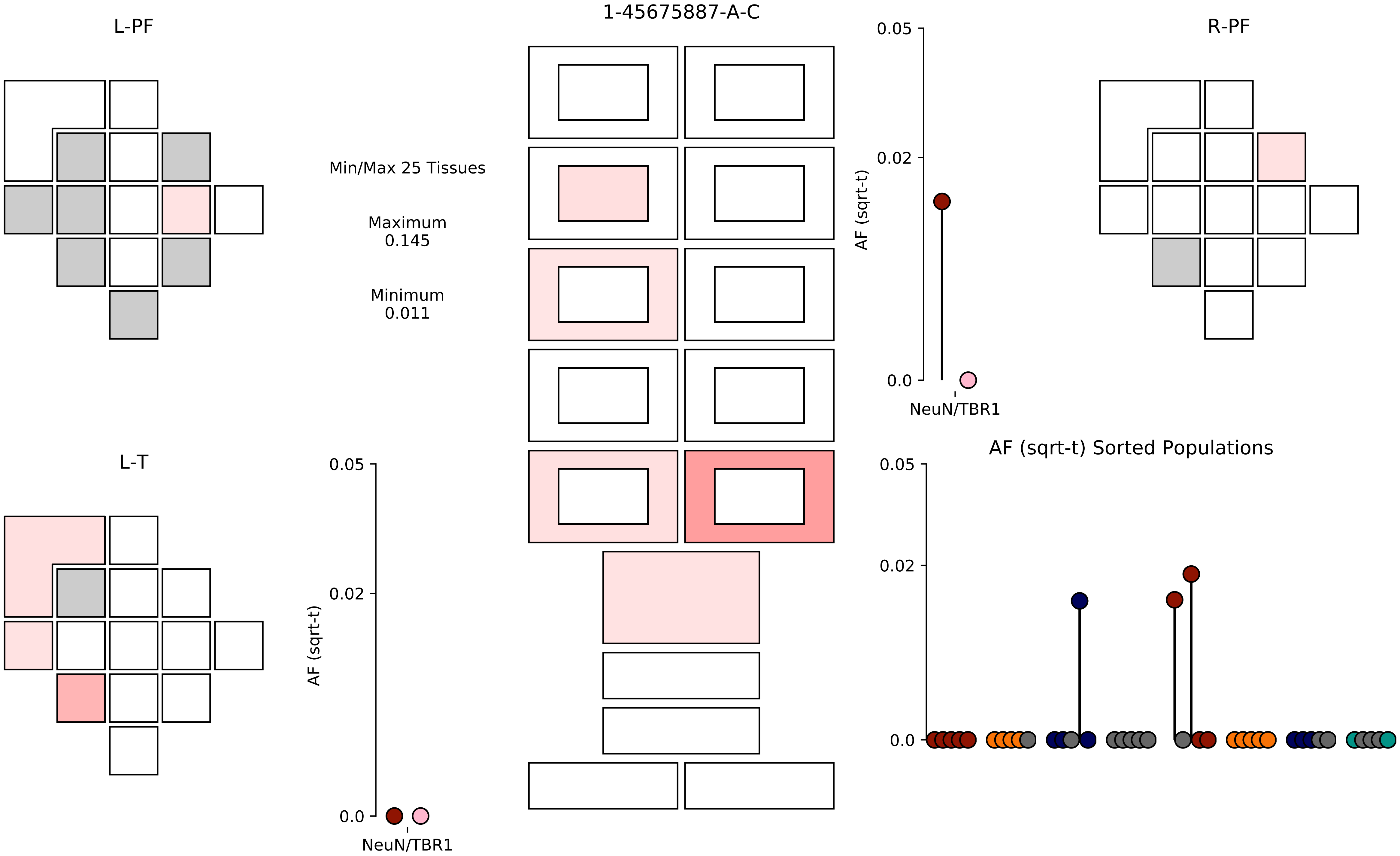
L-PF
1-45675887-A-C
R-PF
Min/Max 25 Tissues
Maximum
0.145
Minimum
0.011
0.05
0.02
0.0
AF (sqrt-t)
NeuN/TBR1
L-T
AF (sqrt-t) Sorted Populations

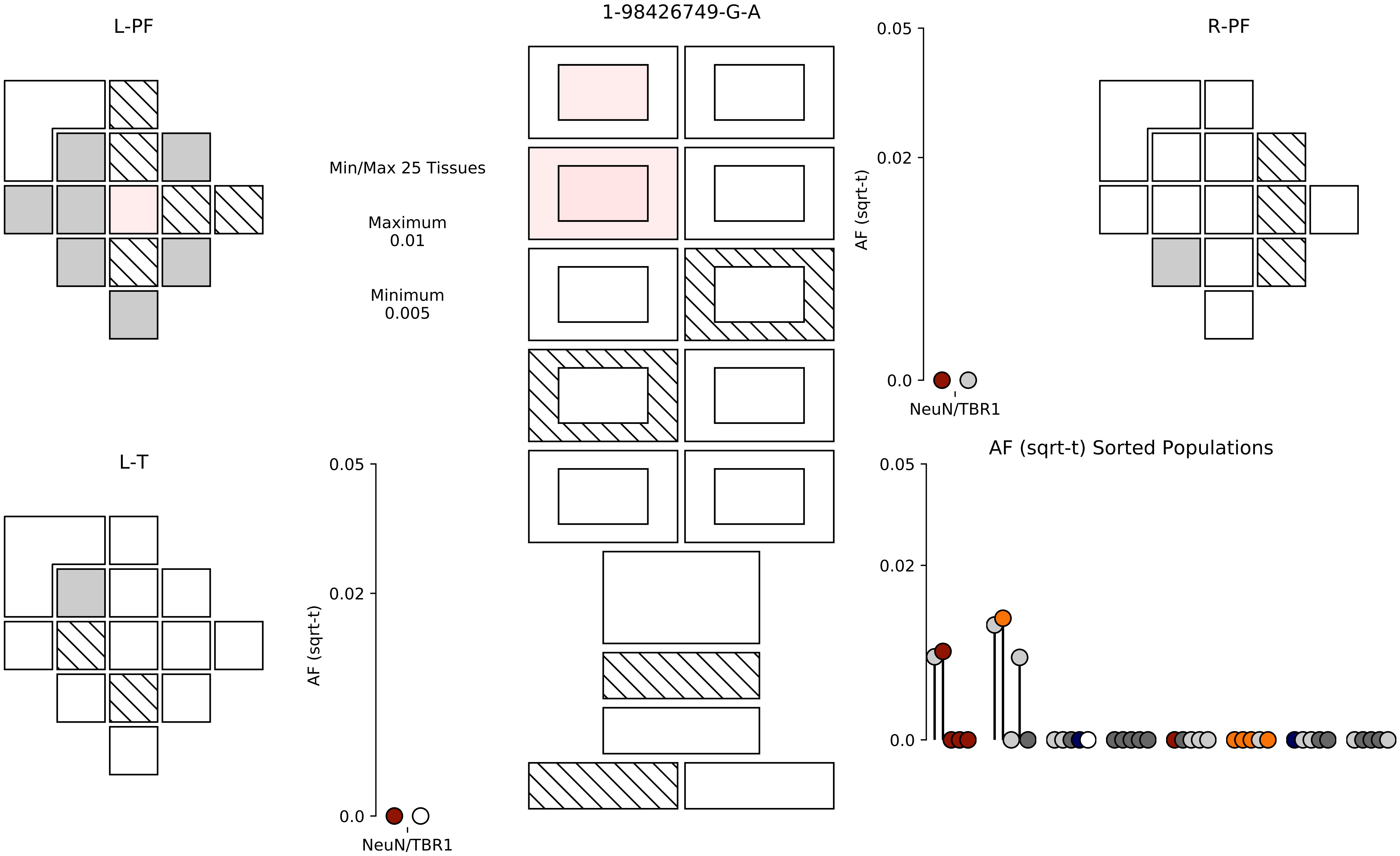

1-98426749-G-A
L-PF
R-PF
L-T
Min/Max 25 Tissues
Maximum
0.01
Minimum
0.005
AF (sqrt-t)
0.05
0.02
0.0
NeuN/TBR1
AF (sqrt-t) Sorted Populations

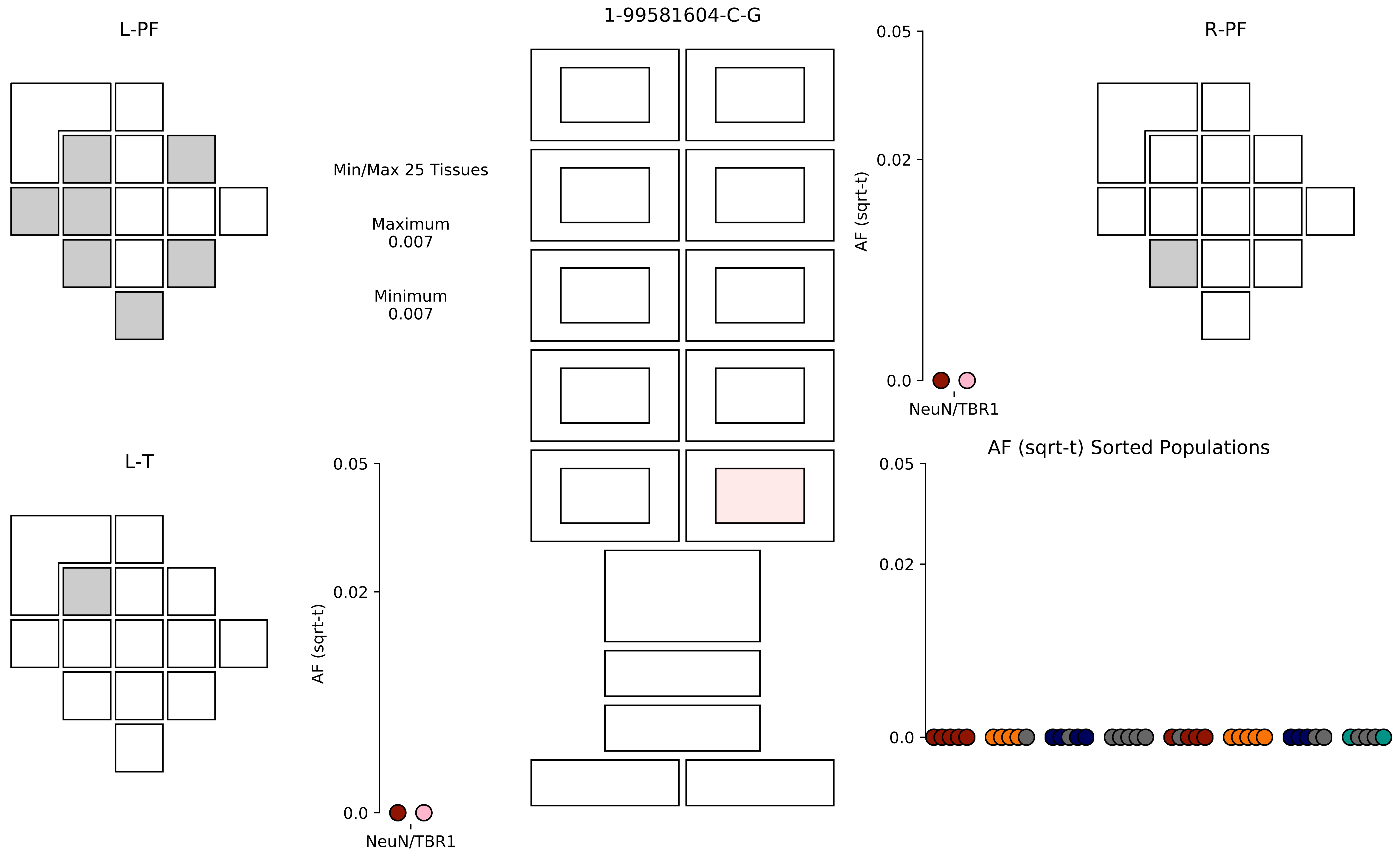

L-PF
1-99581604-C-G
R-PF
Min/Max 25 Tissues
Maximum
0.007
Minimum
0.007
0.05
0.02
0.0
AF (sqrt-t)
NeuN/TBR1
L-T
0.05
0.02
0.0
AF (sqrt-t)
NeuN/TBR1
AF (sqrt-t) Sorted Populations
0.05
0.02
0.0

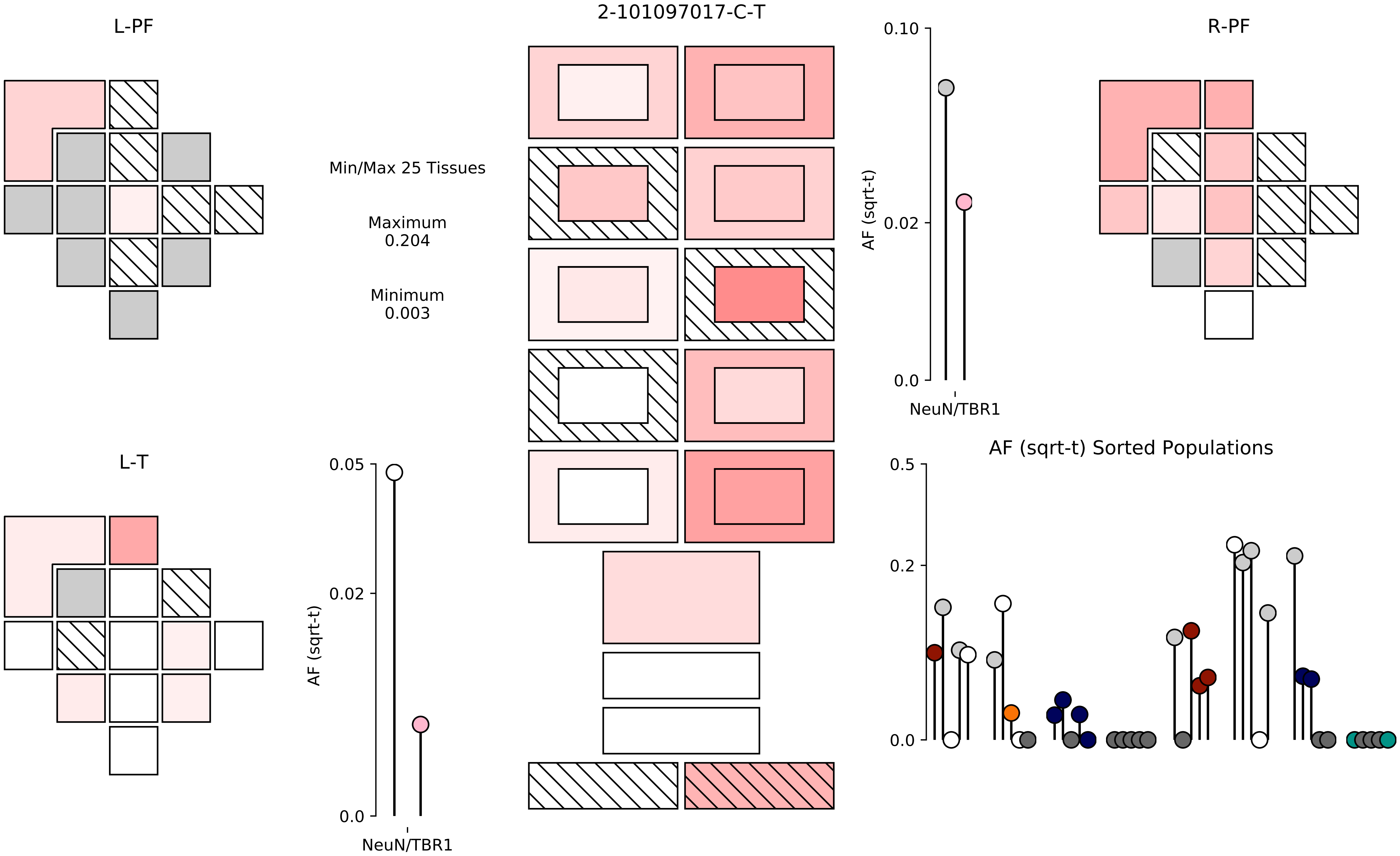

2-101097017-C-T
L-PF
R-PF
L-T
Min/Max 25 Tissues
Maximum
0.204
Minimum
0.003
0.10
0.02
0.0
AF (sqrt-t)
NeuN/TBR1
0.05
0.02
0.0
AF (sqrt-t)
NeuN/TBR1
AF (sqrt-t) Sorted Populations
0.5
0.2
0.0

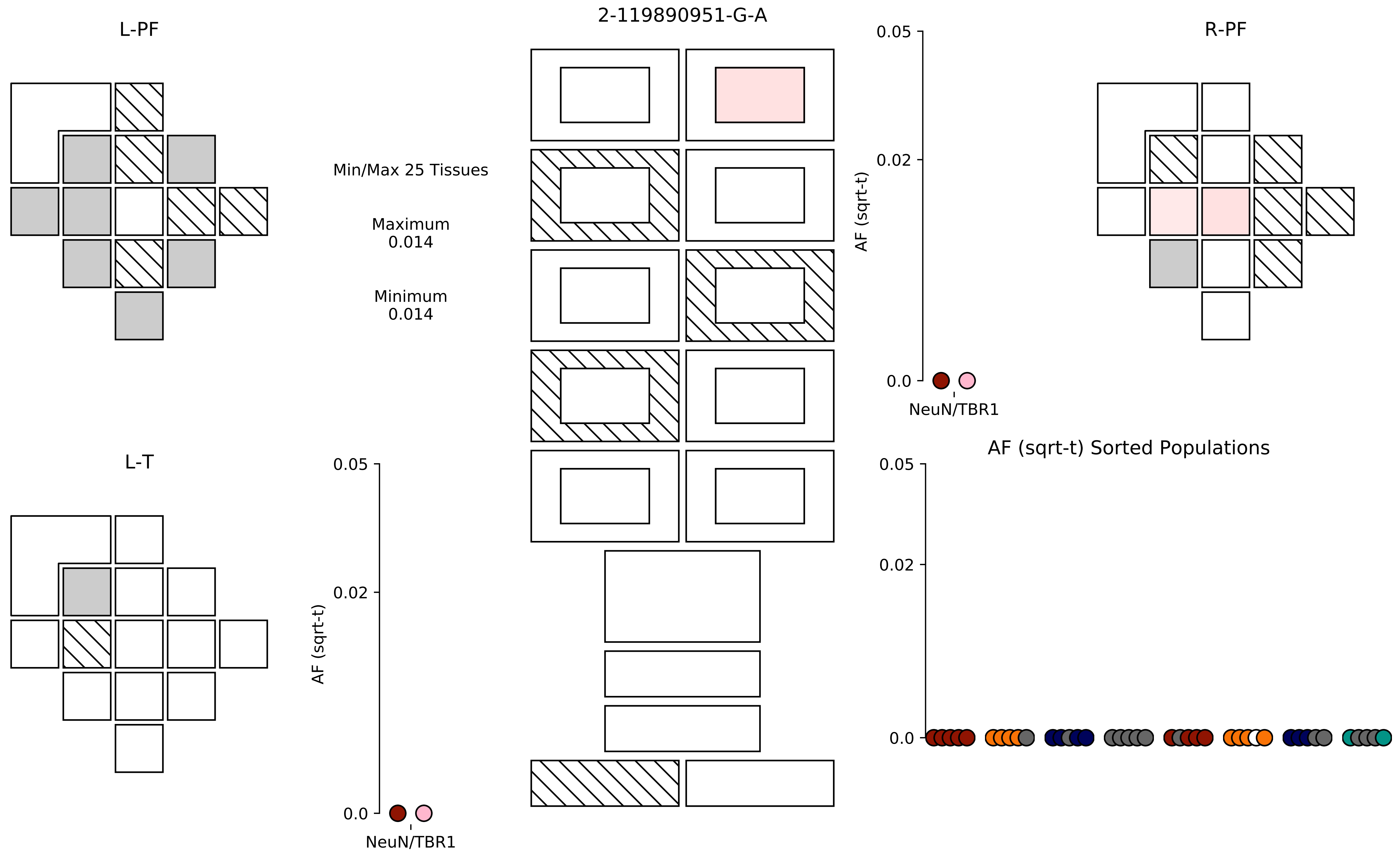

L-PF
2-119890951-G-A
R-PF
Min/Max 25 Tissues
Maximum
0.014
Minimum
0.014
0.05
0.02
0.0
AF (sqrt-t)
NeuN/TBR1
L-T
0.05
0.02
0.0
AF (sqrt-t)
NeuN/TBR1
AF (sqrt-t) Sorted Populations
0.05
0.02
0.0

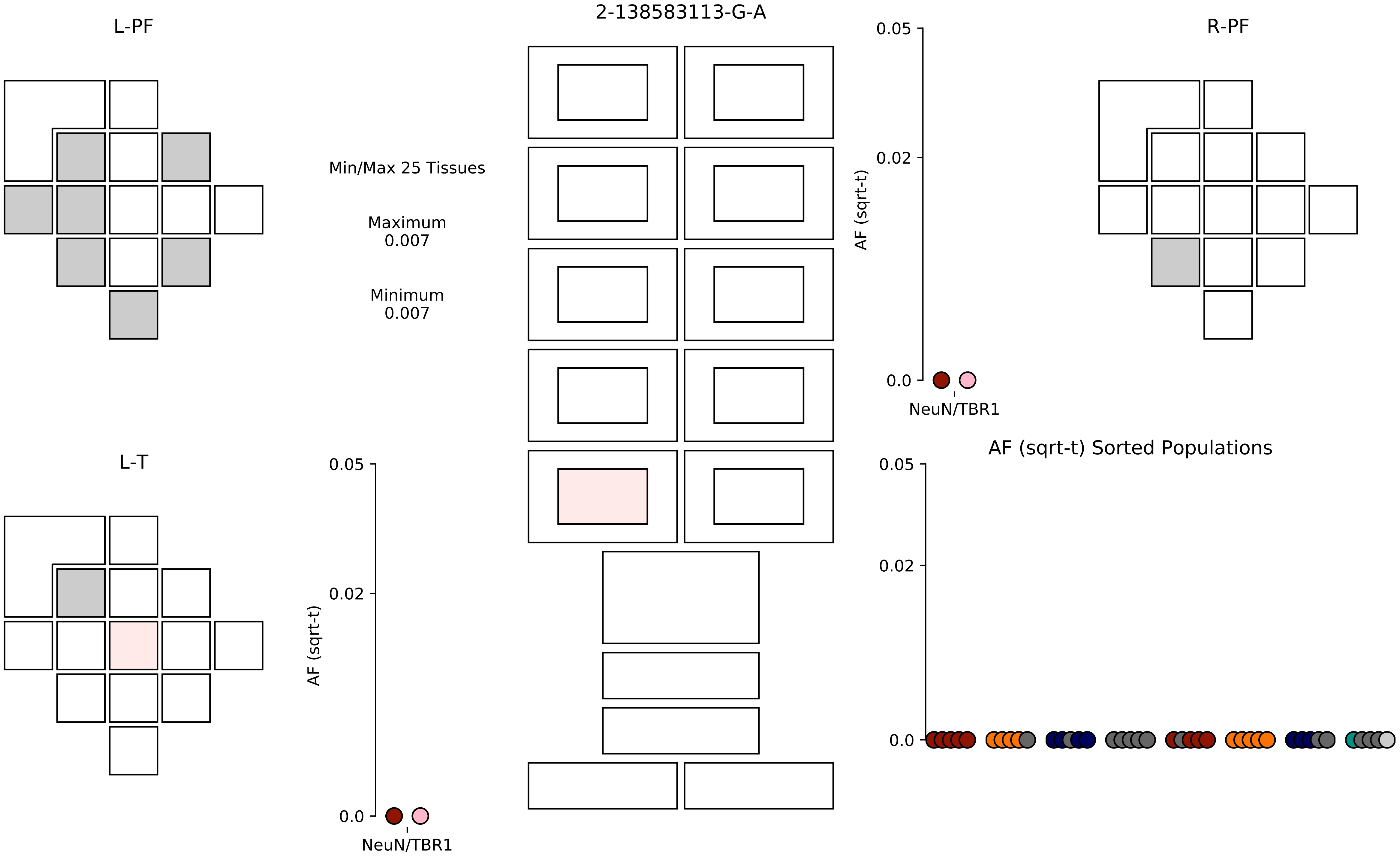
2-138583113-G-A
L-PF
R-PF
L-T
Min/Max 25 Tissues
Maximum
0.007
Minimum
0.007
0.05
0.02
0.0
AF (sqrt-t)
NeuN/TBR1
0.05
0.02
0.0
AF (sqrt-t)
NeuN/TBR1
AF (sqrt-t) Sorted Populations
0.05
0.02
0.0

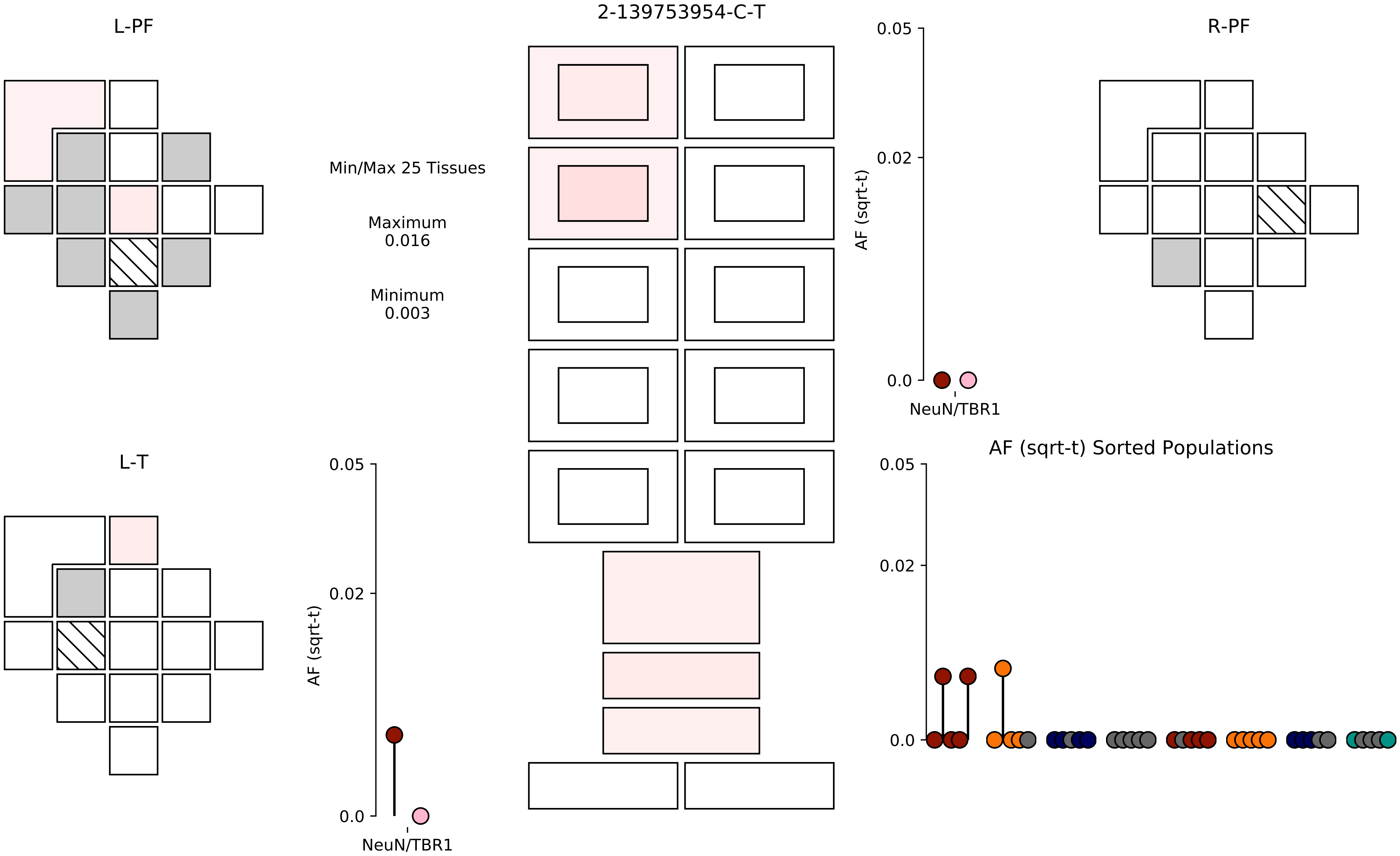

2-139753954-C-T
L-PF
R-PF
L-T
Min/Max 25 Tissues
Maximum
0.016
Minimum
0.003
AF (sqrt-t)
0.05
0.02
0.0
NeuN/TBR1
AF (sqrt-t) Sorted Populations

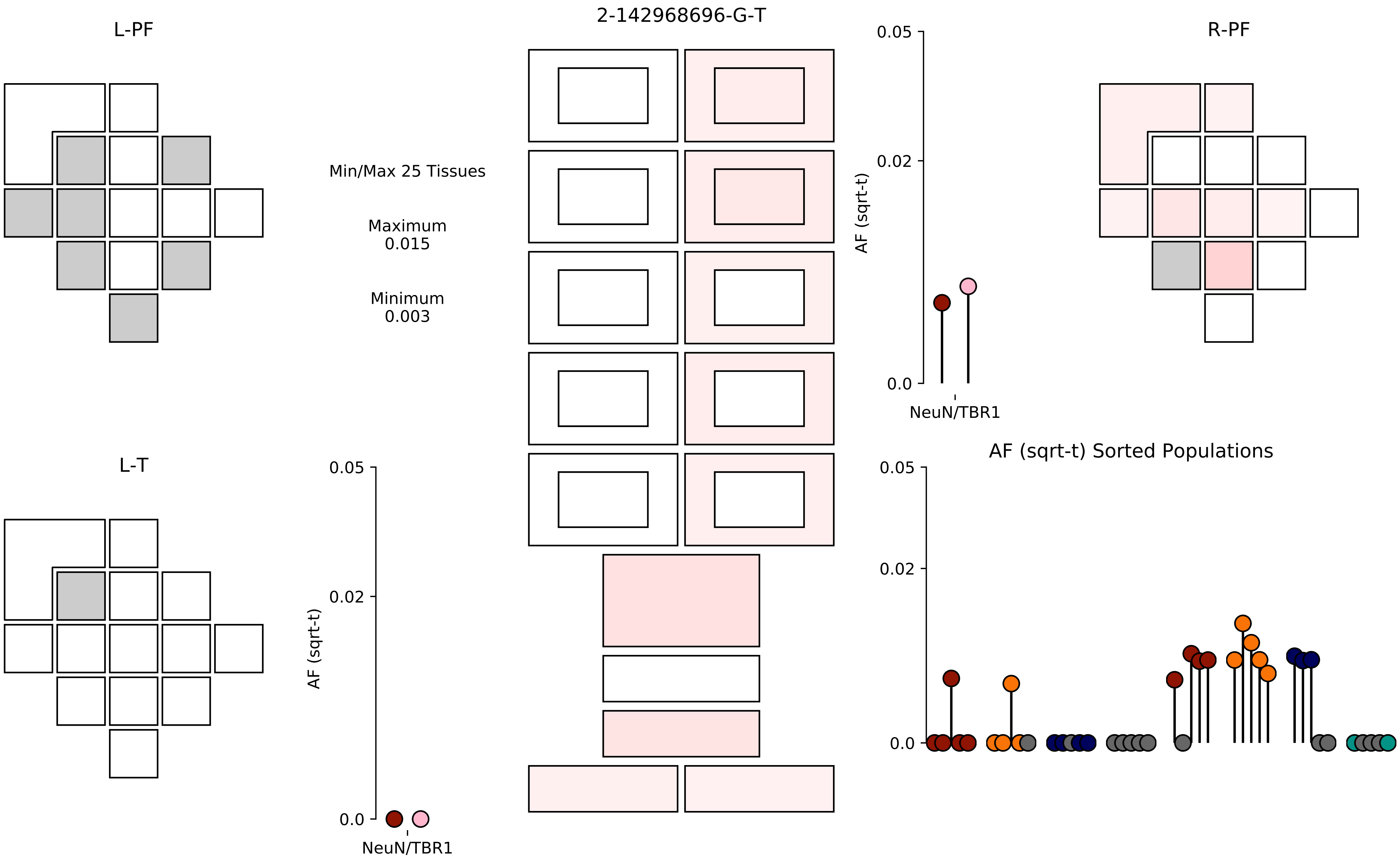

2-142968696-G-T
L-PF
R-PF
L-T
Min/Max 25 Tissues
Maximum
0.015
Minimum
0.003
AF (sqrt-t)
0.05
0.02
0.0
NeuN/TBR1
AF (sqrt-t) Sorted Populations

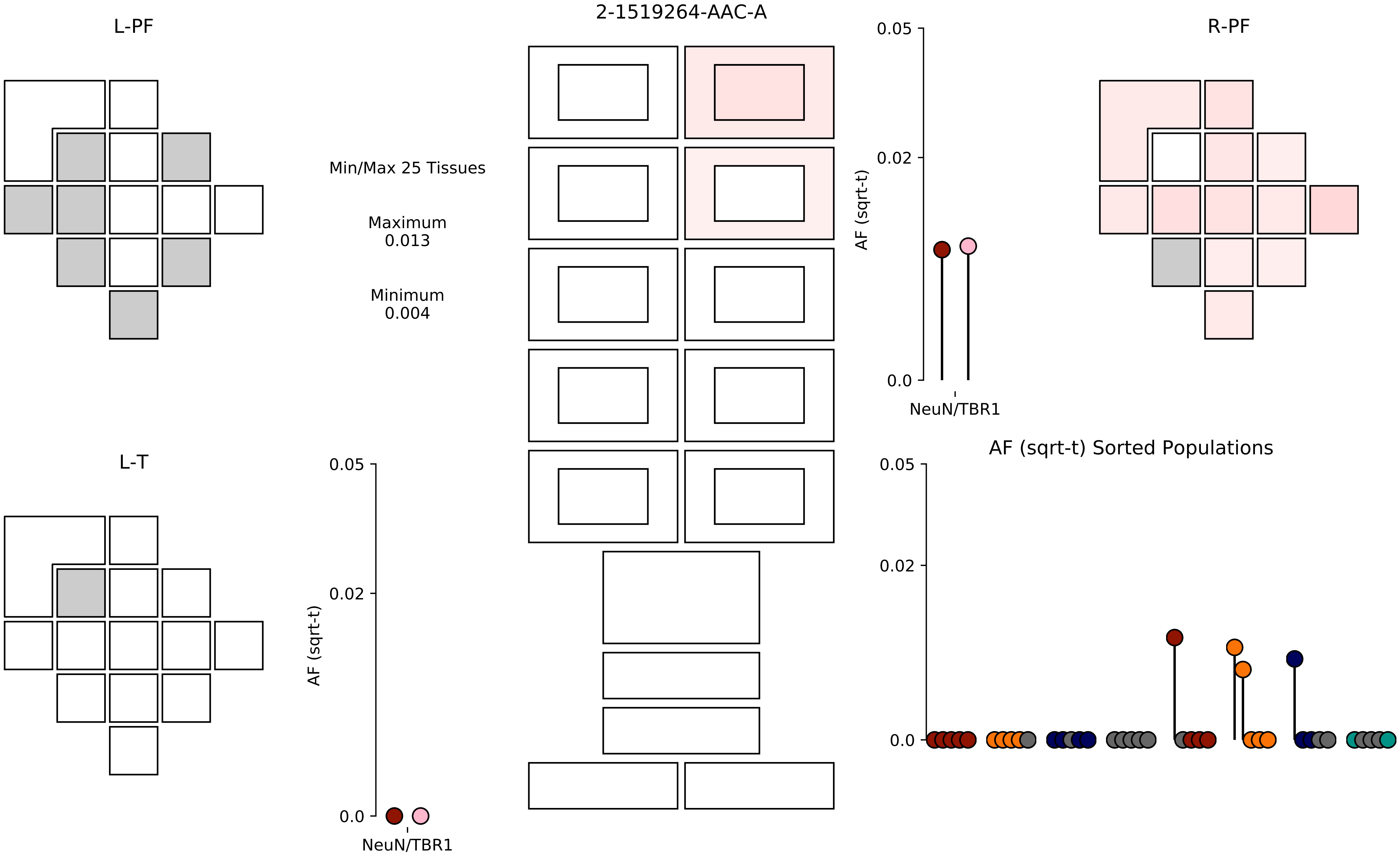

2-1519264-AAC-A
L-PF
R-PF
L-T
Min/Max 25 Tissues
Maximum
0.013
Minimum
0.004
AF (sqrt-t)
0.05
0.02
0.0
NeuN/TBR1
AF (sqrt-t) Sorted Populations

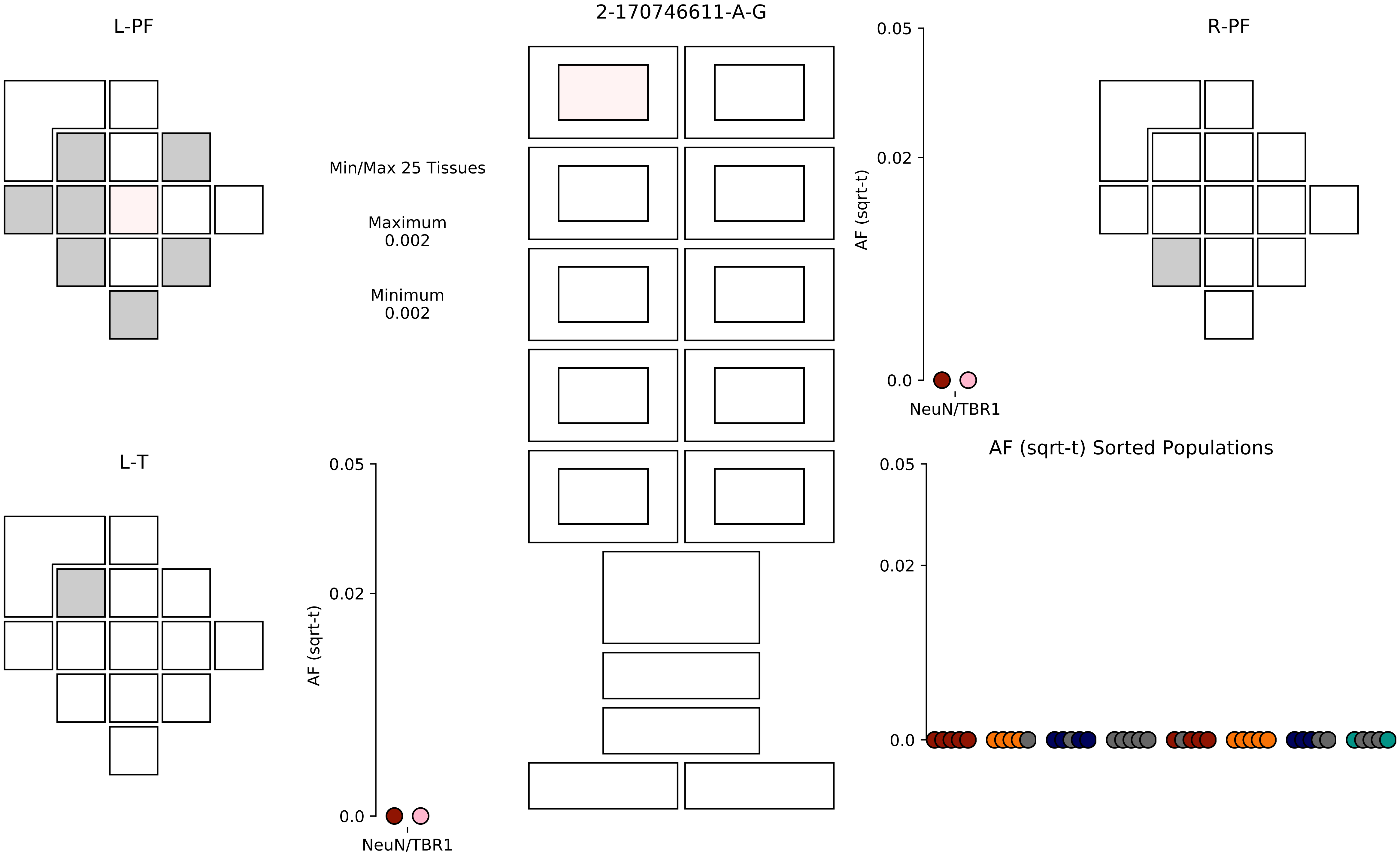
L-PF
2-170746611-A-G
R-PF
Min/Max 25 Tissues
Maximum
0.002
Minimum
0.002
0.05
0.02
0.0
AF (sqrt-t)
NeuN/TBR1
L-T
AF (sqrt-t) Sorted Populations

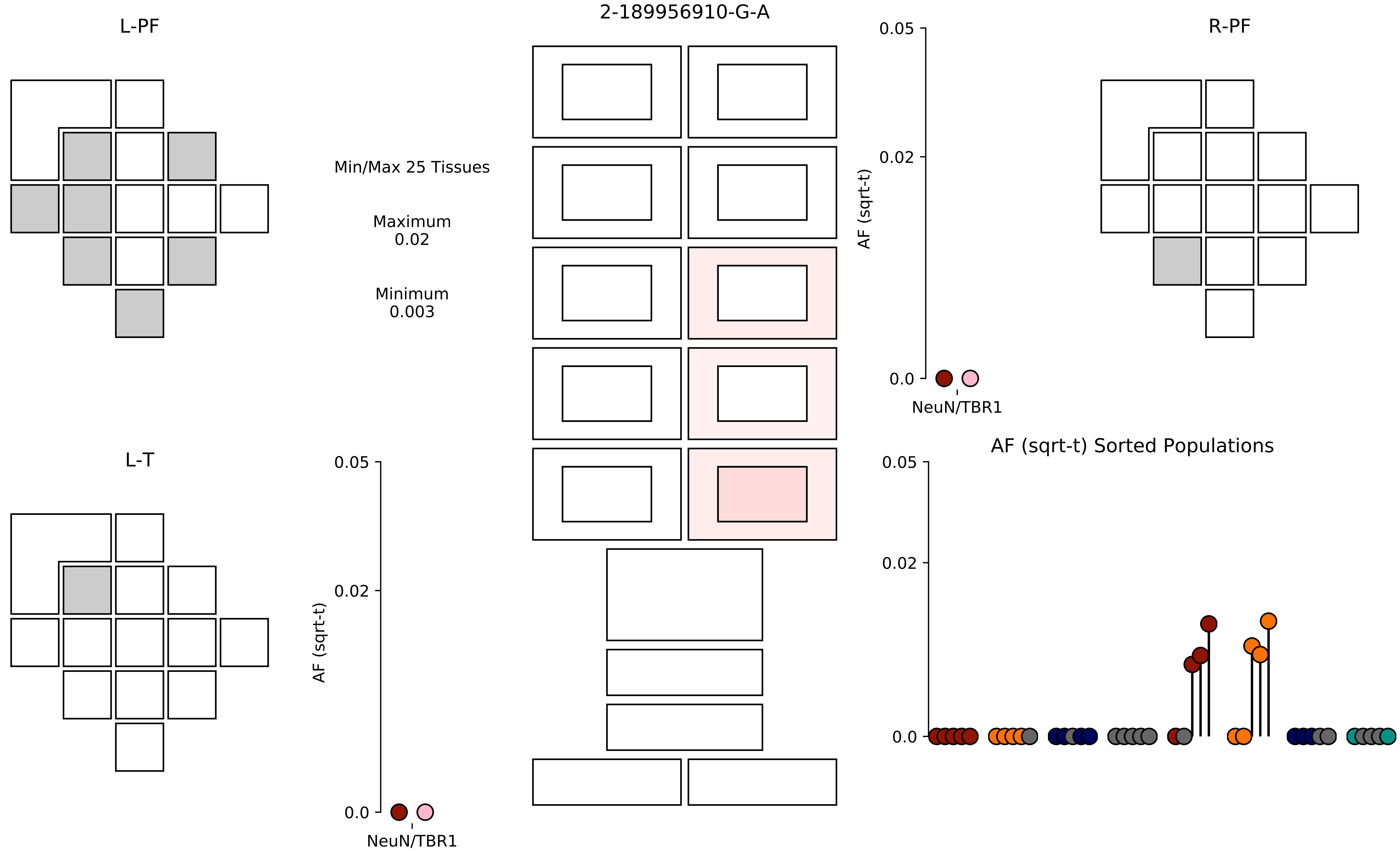

2-189956910-G-A
L-PF
R-PF
L-T
Min/Max 25 Tissues
Maximum
0.02
Minimum
0.003
AF (sqrt-t)
0.05
0.02
0.0
NeuN/TBR1
AF (sqrt-t) Sorted Populations

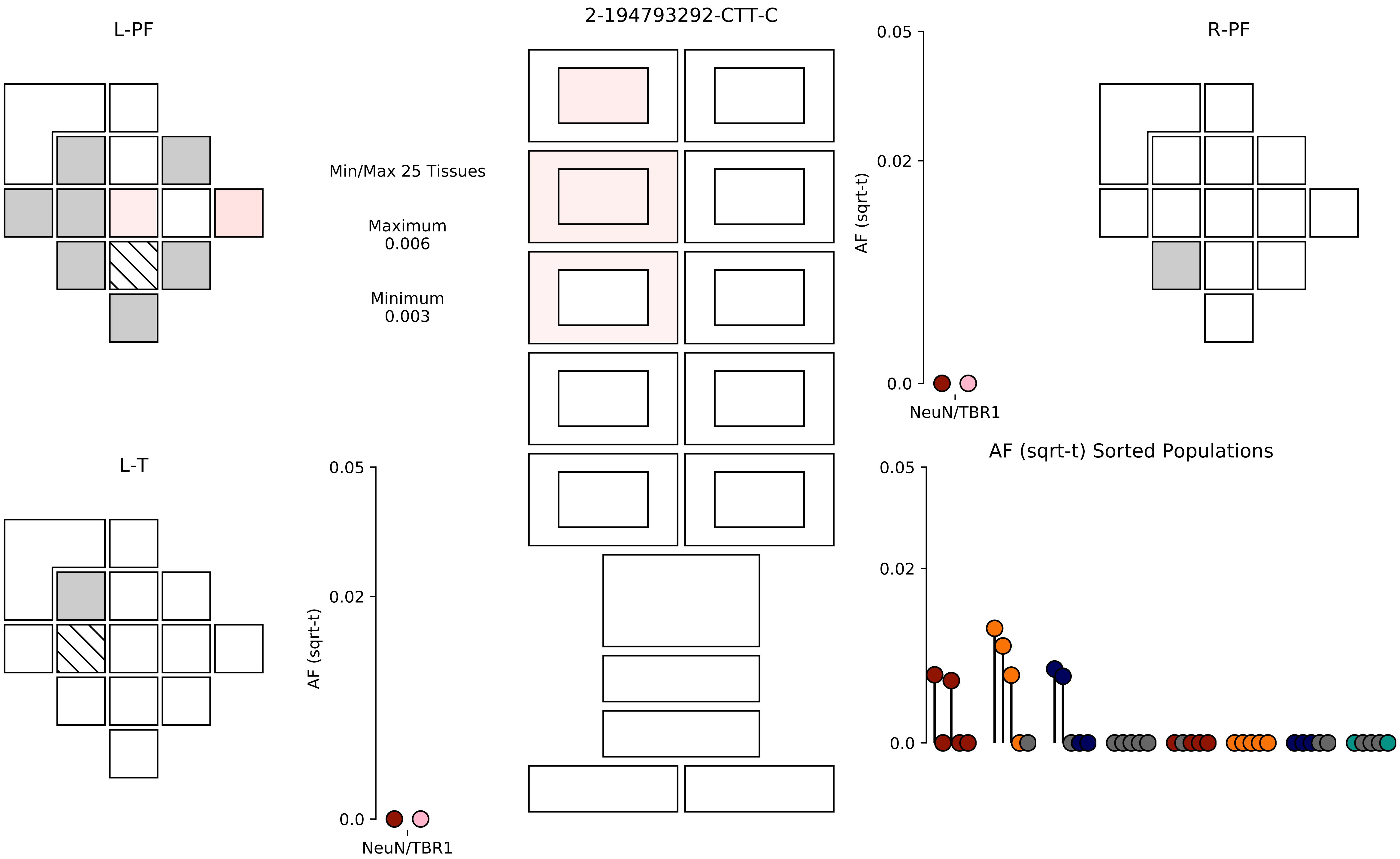

2-194793292-CTT-C
L-PF
R-PF
L-T
Min/Max 25 Tissues
Maximum
0.006
Minimum
0.003
AF (sqrt-t)
0.05
0.02
0.0
NeuN/TBR1
AF (sqrt-t) Sorted Populations

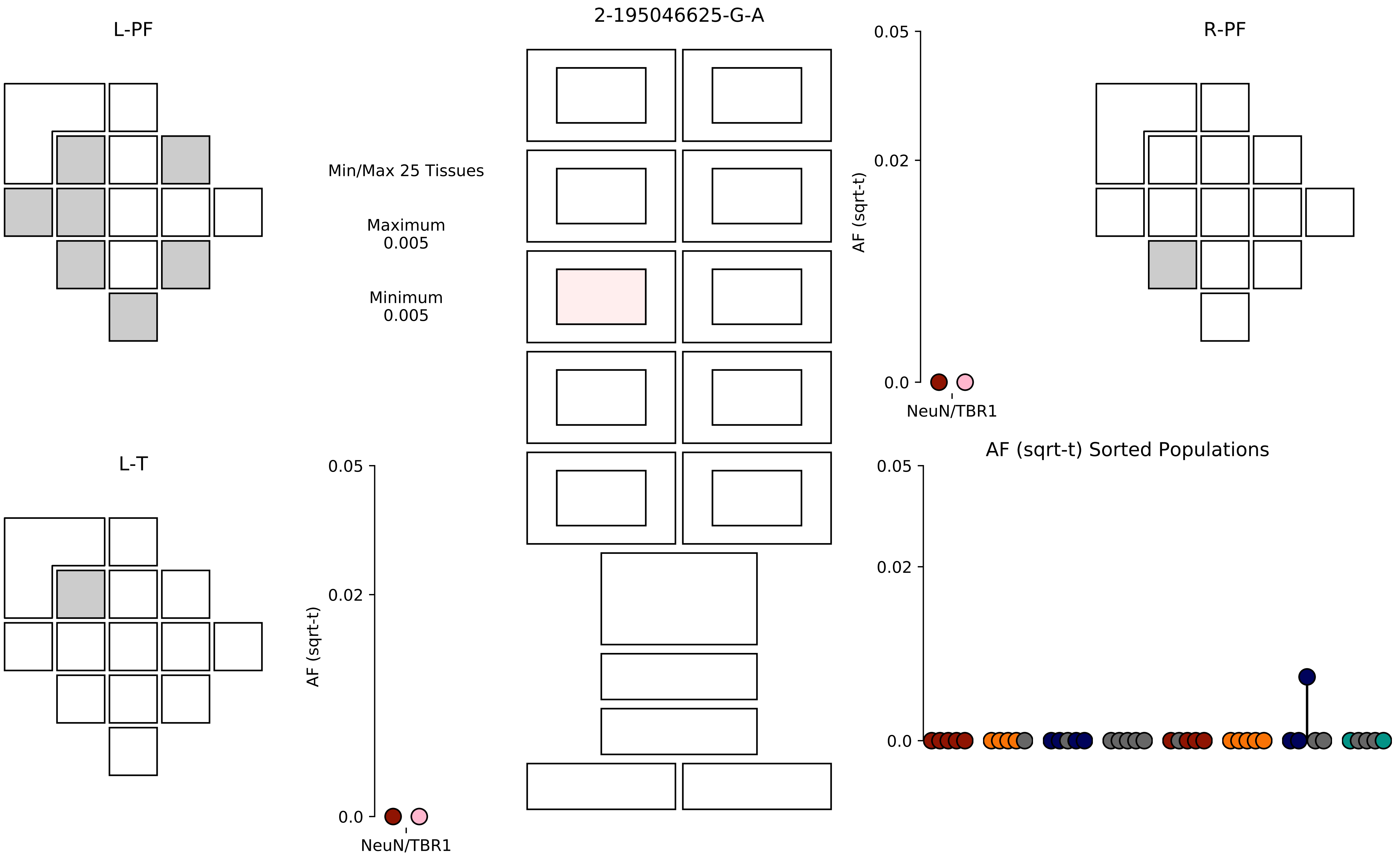

2-195046625-G-A
L-PF
R-PF
L-T
Min/Max 25 Tissues
Maximum
0.005
Minimum
0.005
AF (sqrt-t)
0.05
0.02
0.0
NeuN/TBR1
AF (sqrt-t) Sorted Populations

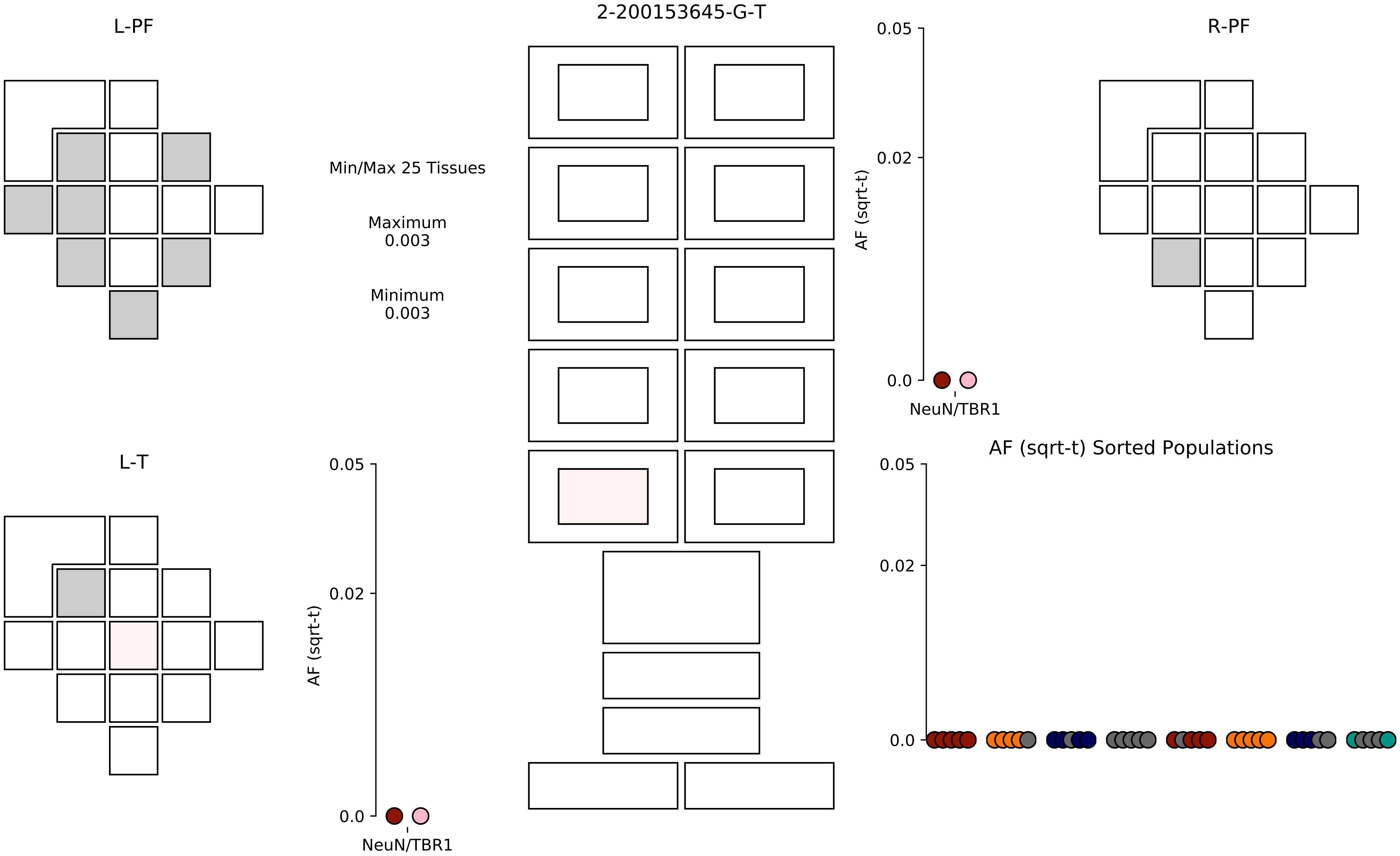

2-200153645-G-T
L-PF
R-PF
L-T
Min/Max 25 Tissues
Maximum
0.003
Minimum
0.003
0.05
0.02
0.0
AF (sqrt-t)
NeuN/TBR1
AF (sqrt-t) Sorted Populations

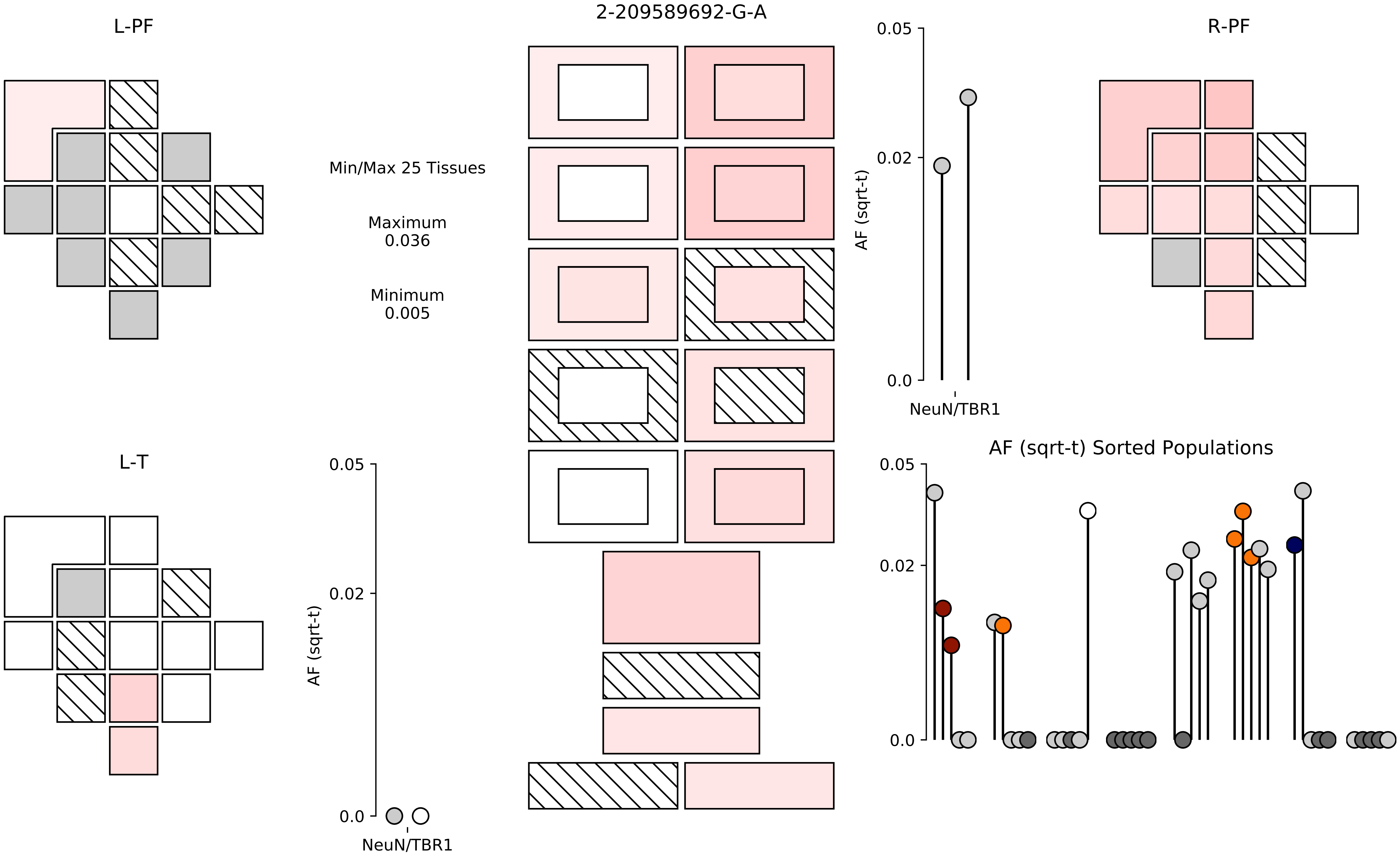

2-209589692-G-A
L-PF
R-PF
L-T
Min/Max 25 Tissues
Maximum
0.036
Minimum
0.005
AF (sqrt-t)
0.05
0.02
0.0
NeuN/TBR1
AF (sqrt-t) Sorted Populations

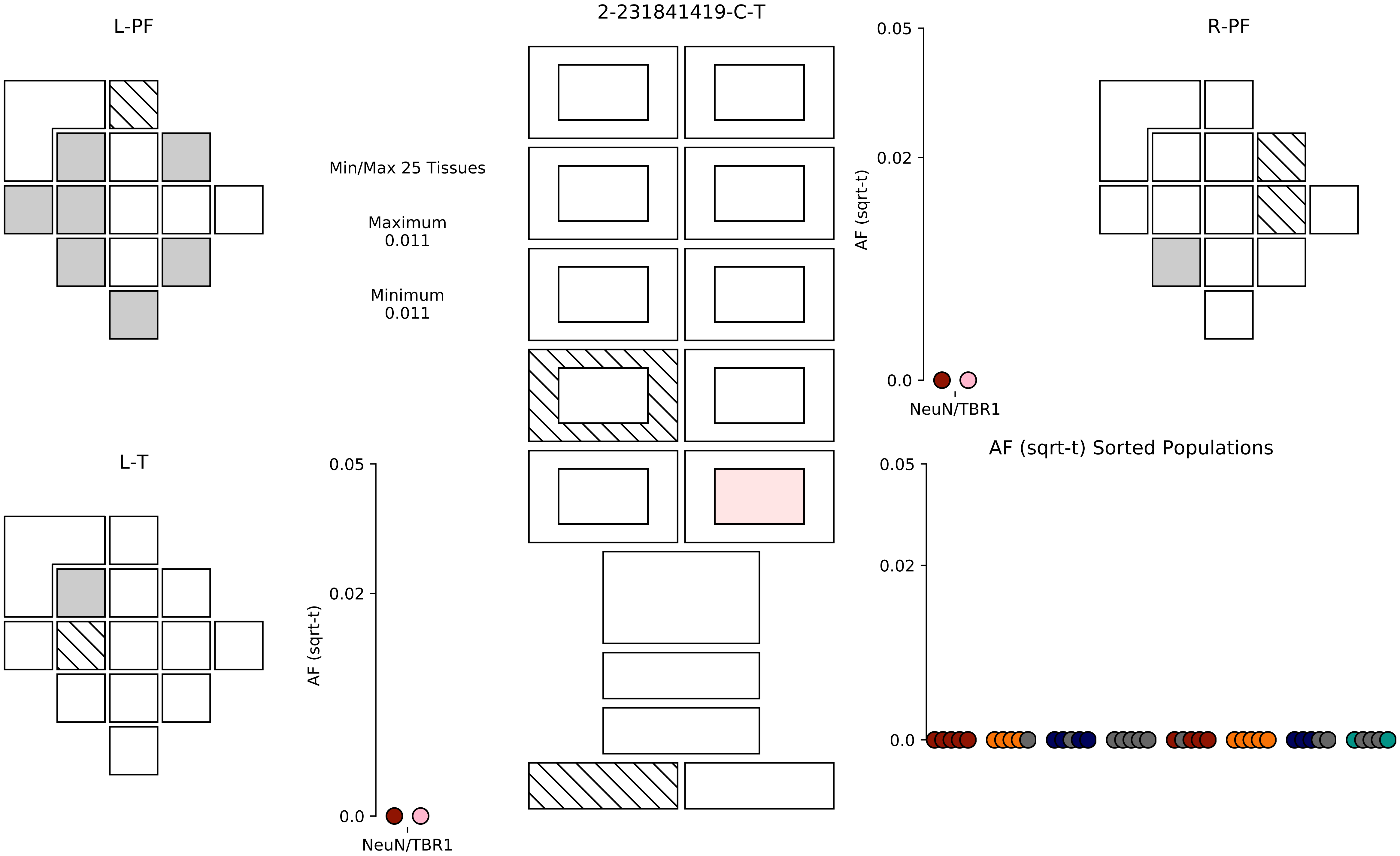
2-231841419-C-T
L-PF
R-PF
L-T
Min/Max 25 Tissues
Maximum
0.011
Minimum
0.011
AF (sqrt-t)
0.05
0.02
0.0
NeuN/TBR1
AF (sqrt-t)
0.05
0.02
0.0
NeuN/TBR1
AF (sqrt-t) Sorted Populations
0.05
0.02
0.0

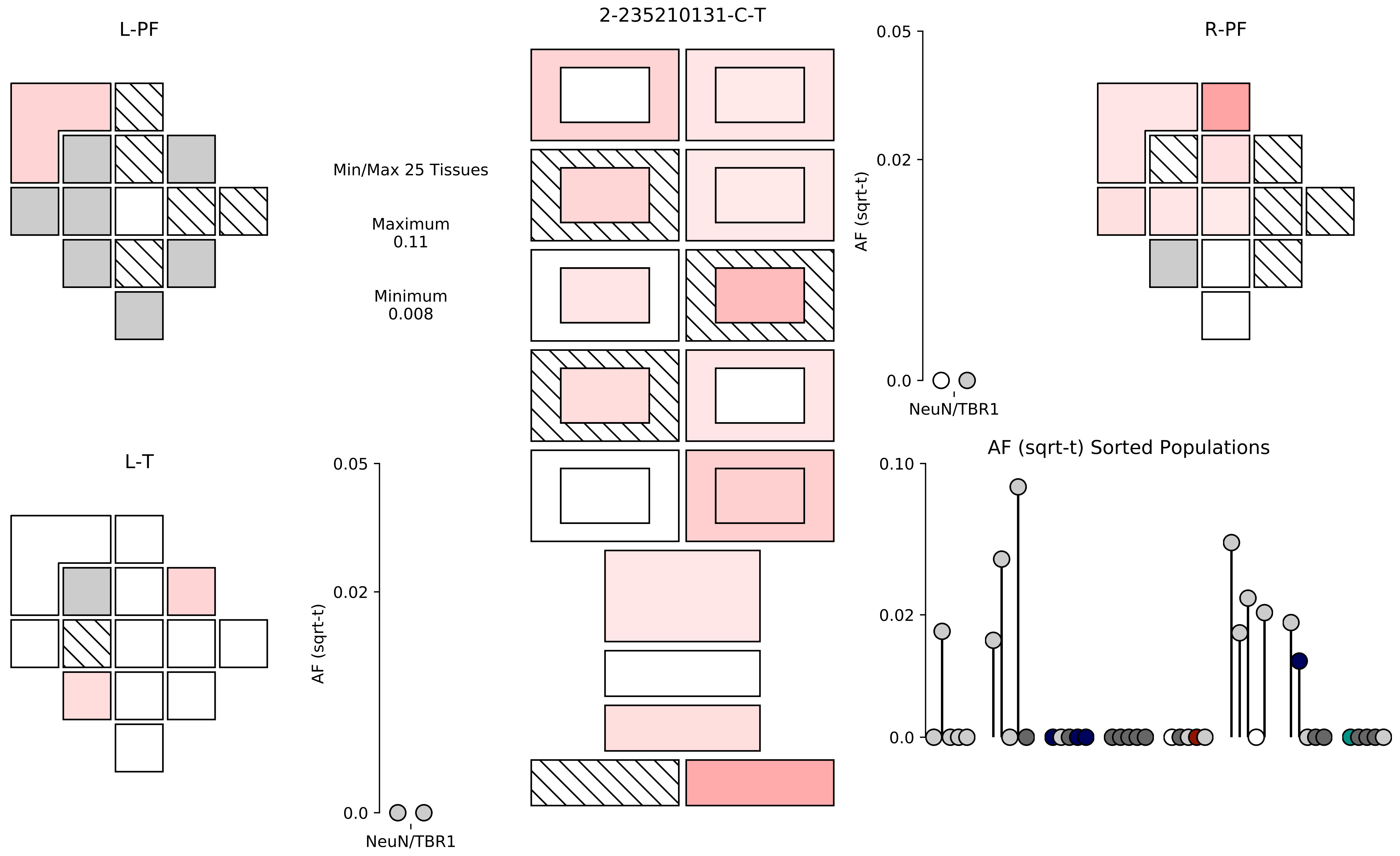

L-PF
2-235210131-C-T
R-PF
Min/Max 25 Tissues
Maximum
0.11
Minimum
0.008
0.05
0.02
0.0
AF (sqrt-t)
NeuN/TBR1
L-T
0.05
0.02
0.0
AF (sqrt-t)
NeuN/TBR1
AF (sqrt-t) Sorted Populations
0.10
0.02
0.0

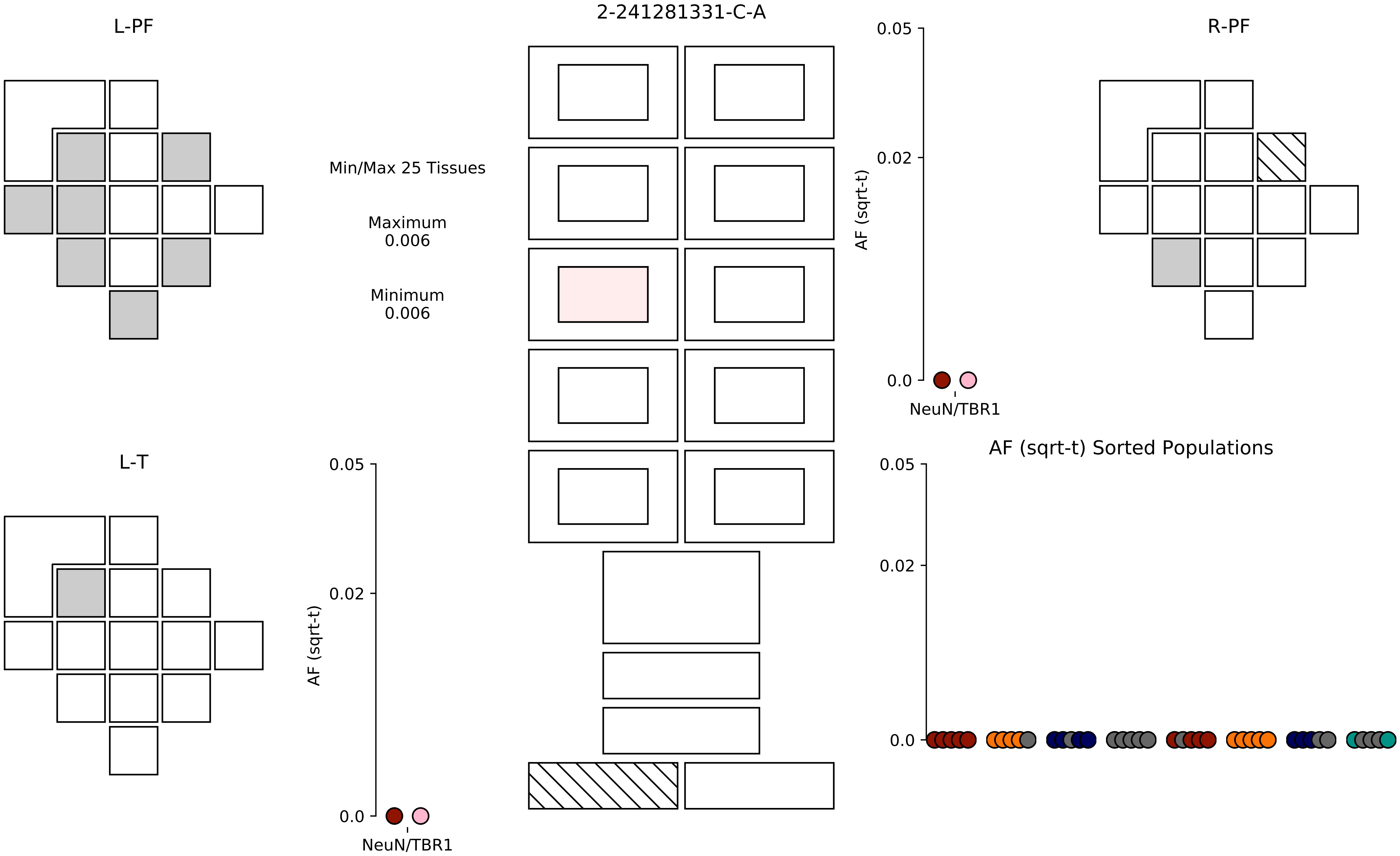

2-241281331-C-A
L-PF
R-PF
L-T
Min/Max 25 Tissues
Maximum
0.006
Minimum
0.006
0.05
0.02
0.0
AF (sqrt-t)
NeuN/TBR1
AF (sqrt-t) Sorted Populations

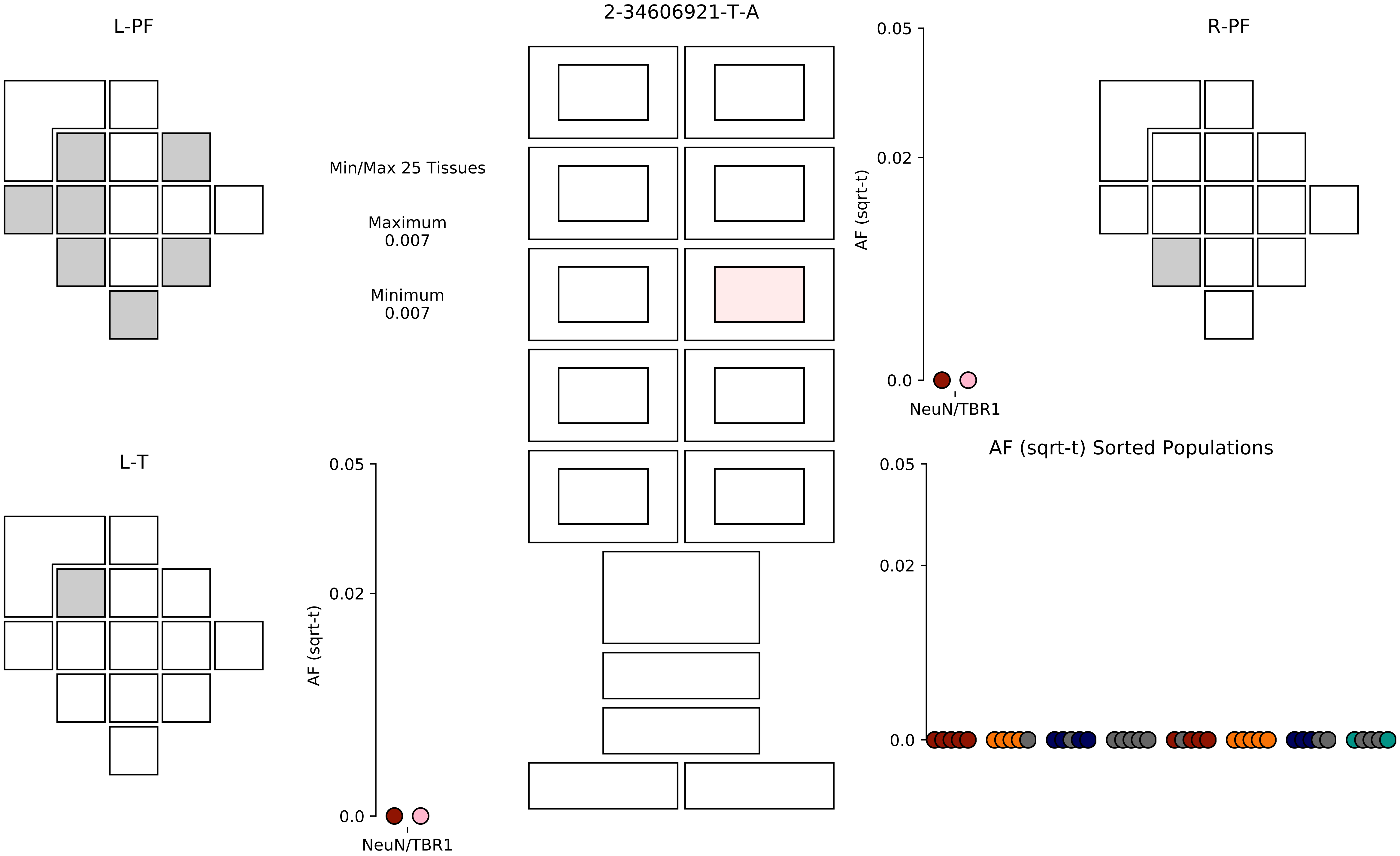

2-34606921-T-A
L-PF
R-PF
L-T
Min/Max 25 Tissues
Maximum
0.007
Minimum
0.007
AF (sqrt-t)
0.05
0.02
0.0
NeuN/TBR1
AF (sqrt-t) Sorted Populations

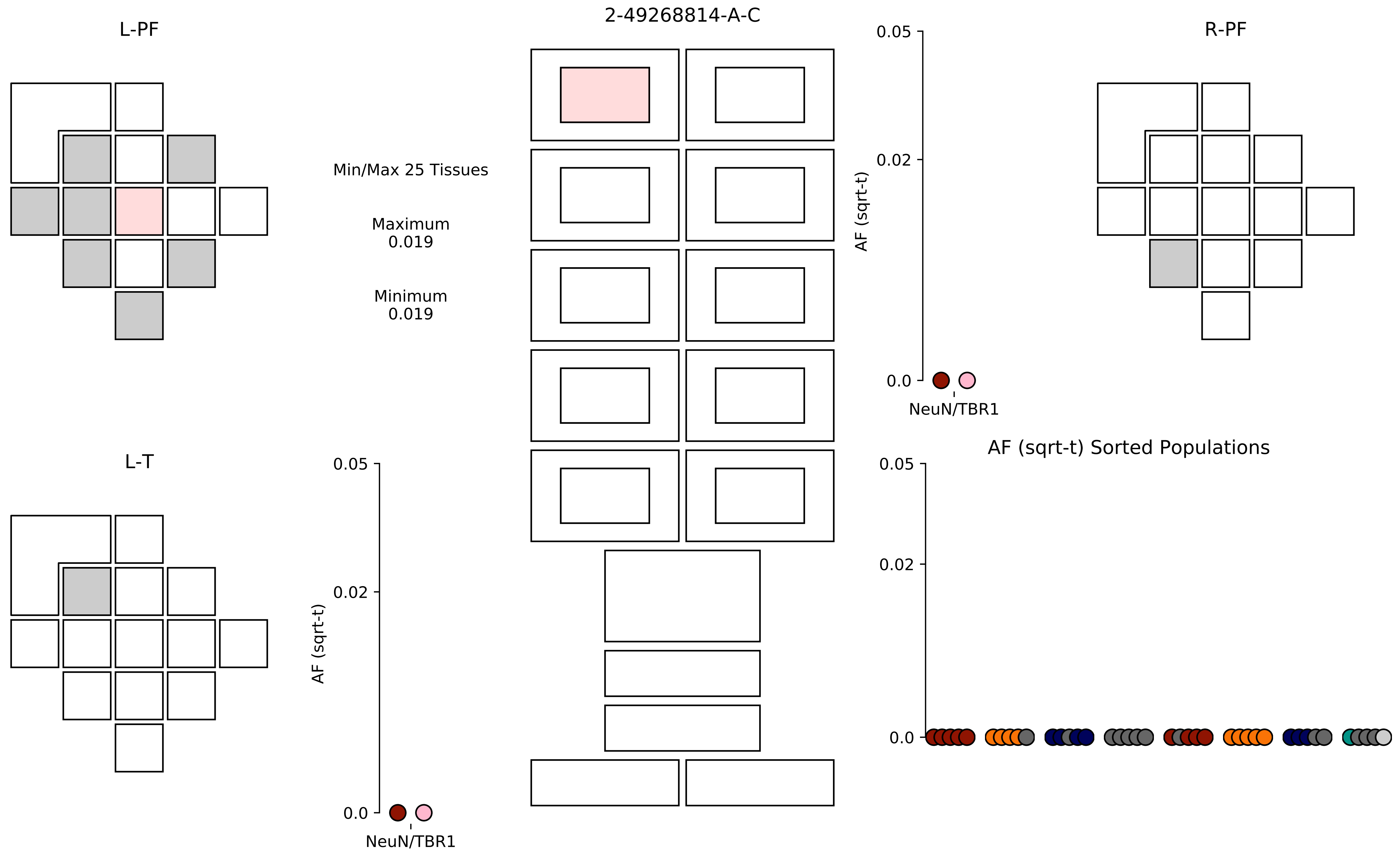

L-PF
2-49268814-A-C
R-PF
Min/Max 25 Tissues
Maximum
0.019
Minimum
0.019
0.05
0.02
0.0
AF (sqrt-t)
NeuN/TBR1
L-T
0.05
0.02
0.0
AF (sqrt-t)
NeuN/TBR1
AF (sqrt-t) Sorted Populations
0.05
0.02
0.0

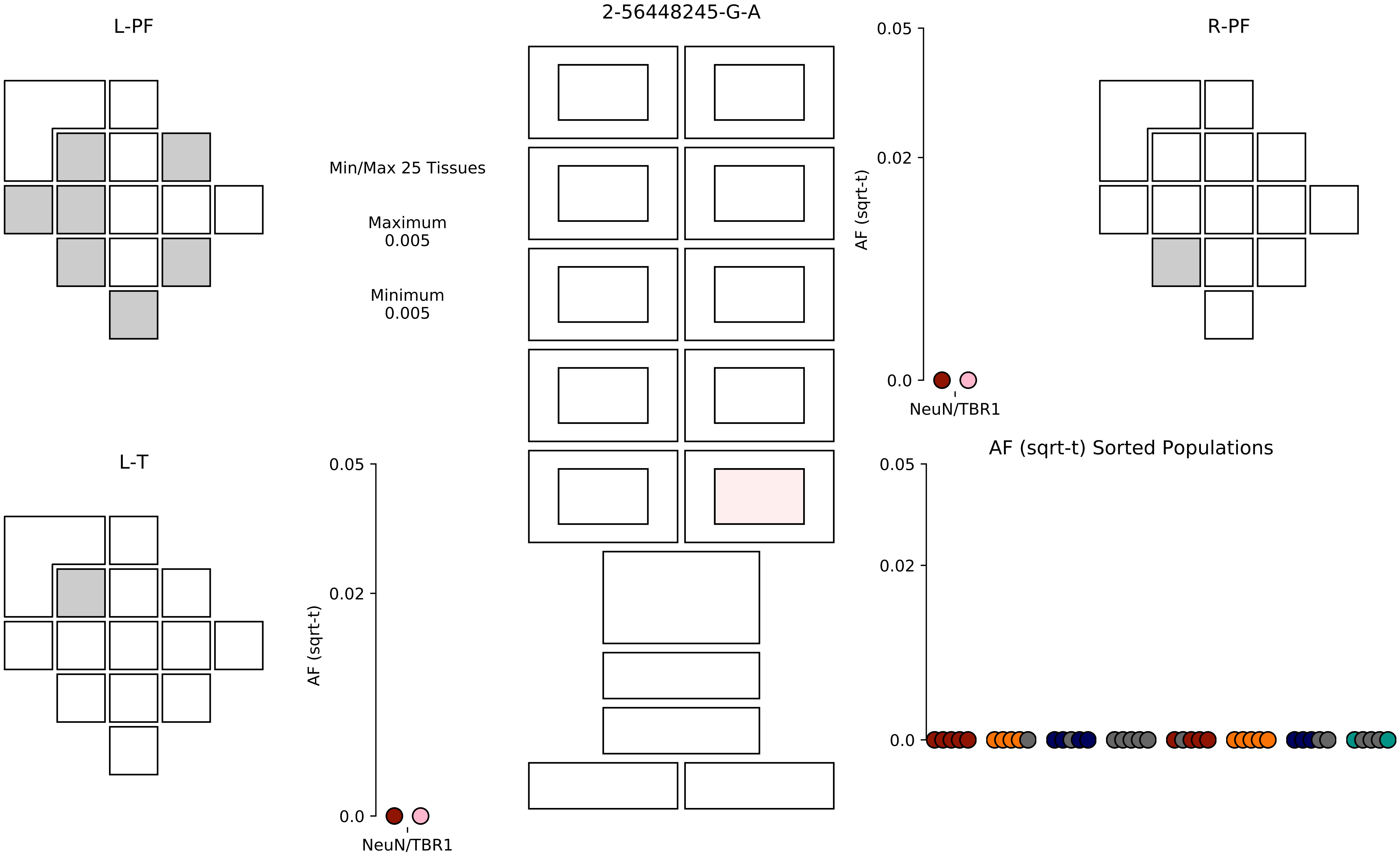
2-56448245-G-A
L-PF
R-PF
L-T
Min/Max 25 Tissues
Maximum
0.005
Minimum
0.005
AF (sqrt-t)
0.05
0.02
0.0
NeuN/TBR1
AF (sqrt-t)
0.05
0.02
0.0
NeuN/TBR1
AF (sqrt-t) Sorted Populations
0.05
0.02
0.0

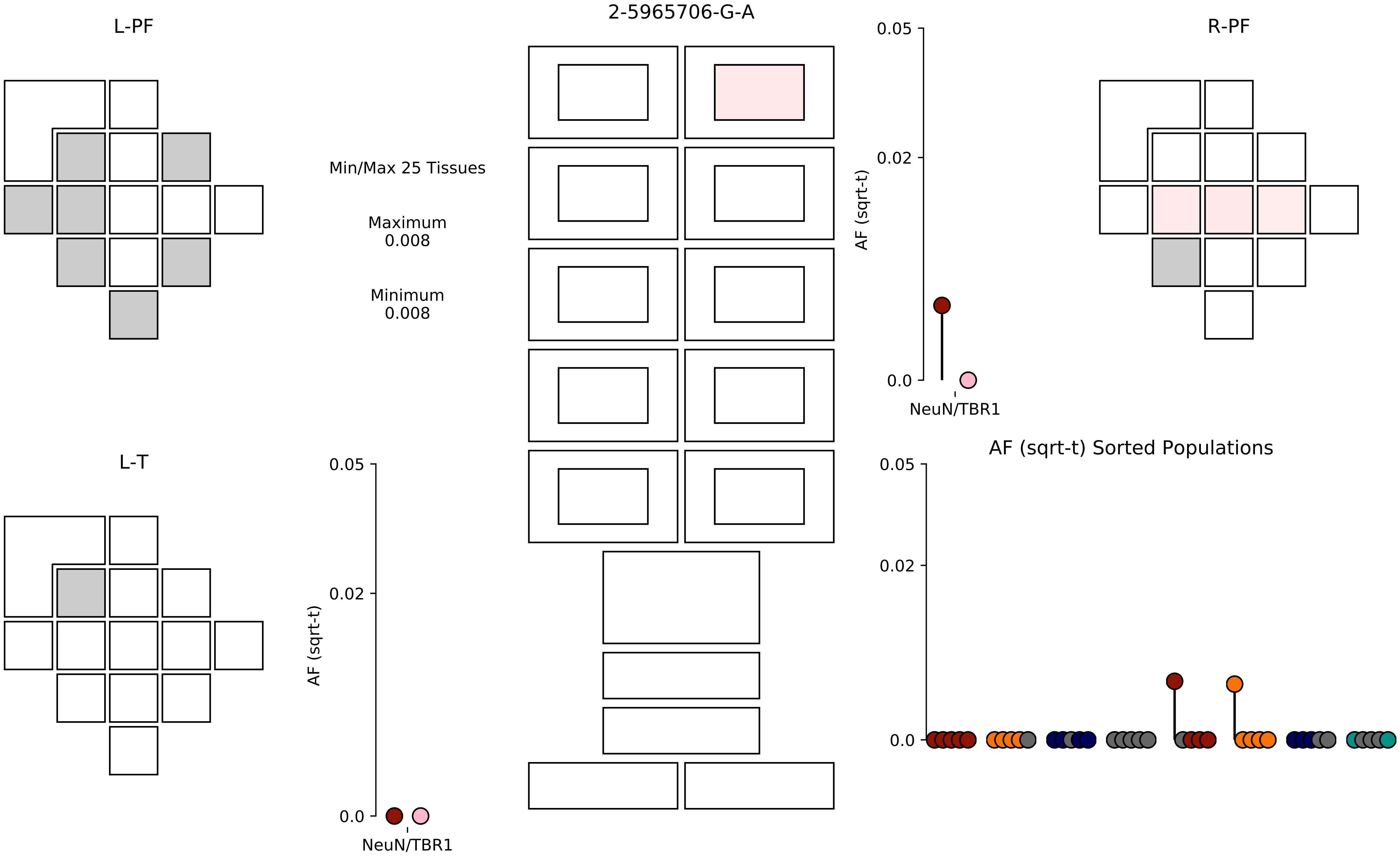

2-5965706-G-A
L-PF
R-PF
L-T
Min/Max 25 Tissues
Maximum
0.008
Minimum
0.008
AF (sqrt-t)
0.05
0.02
0.0
NeuN/TBR1
AF (sqrt-t) Sorted Populations

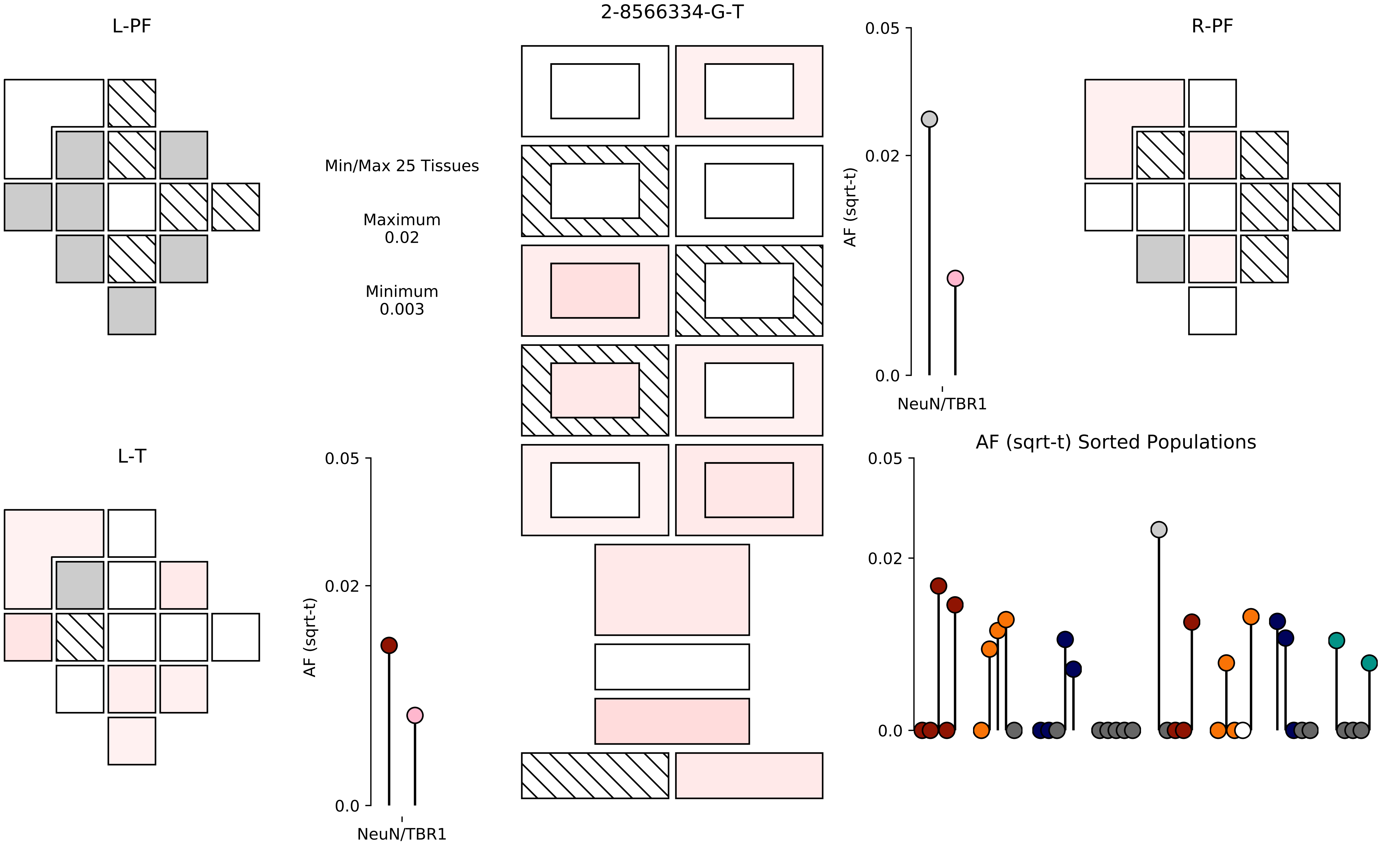

L-PF
2-8566334-G-T
R-PF
Min/Max 25 Tissues
Maximum
0.02
Minimum
0.003
0.05
0.02
0.0
AF (sqrt-t)
NeuN/TBR1
L-T
0.05
0.02
0.0
AF (sqrt-t)
NeuN/TBR1
AF (sqrt-t) Sorted Populations
0.05
0.02
0.0

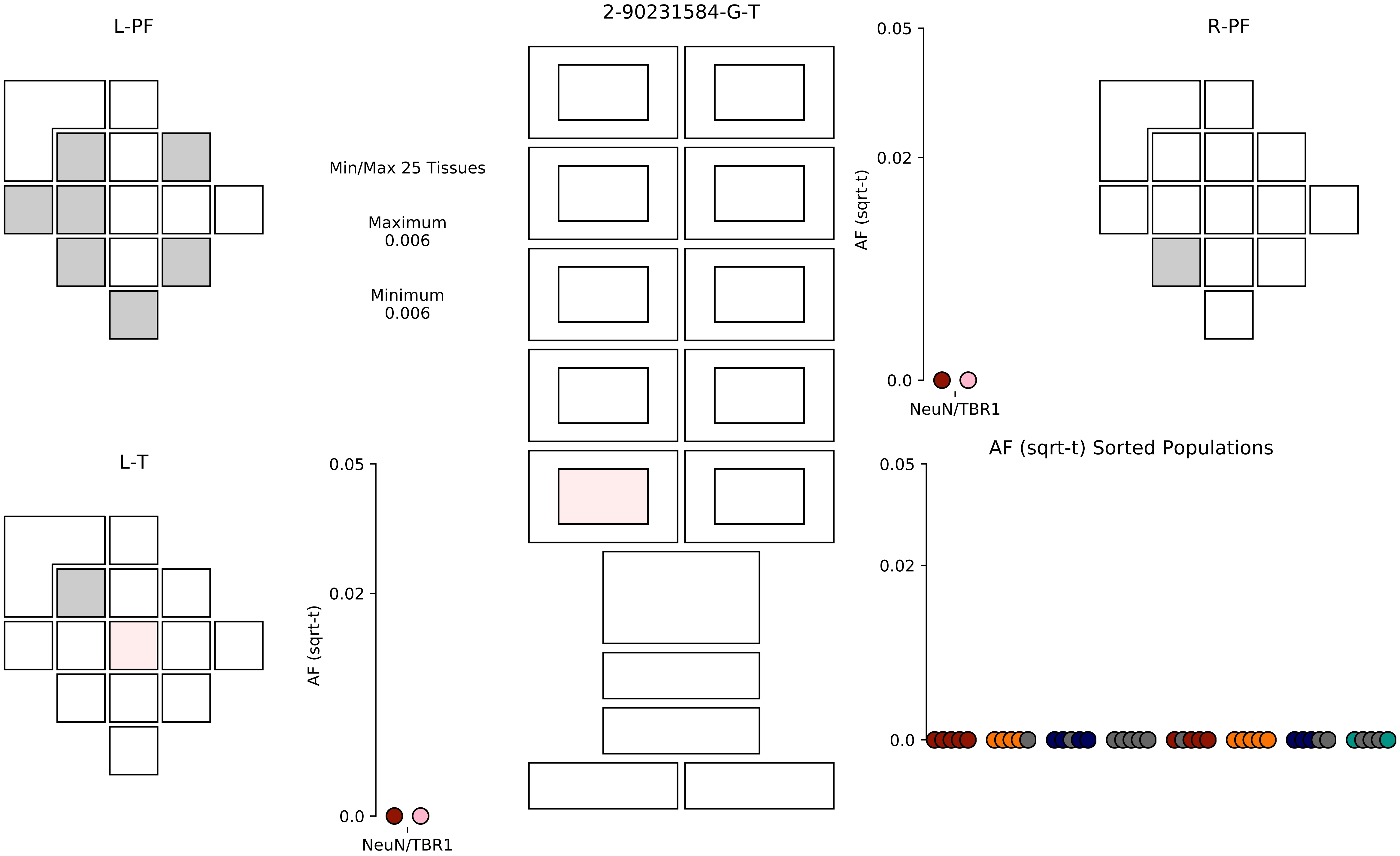

2-90231584-G-T
L-PF
R-PF
L-T
Min/Max 25 Tissues
Maximum
0.006
Minimum
0.006
AF (sqrt-t)
0.05
0.02
0.0
NeuN/TBR1
AF (sqrt-t) Sorted Populations

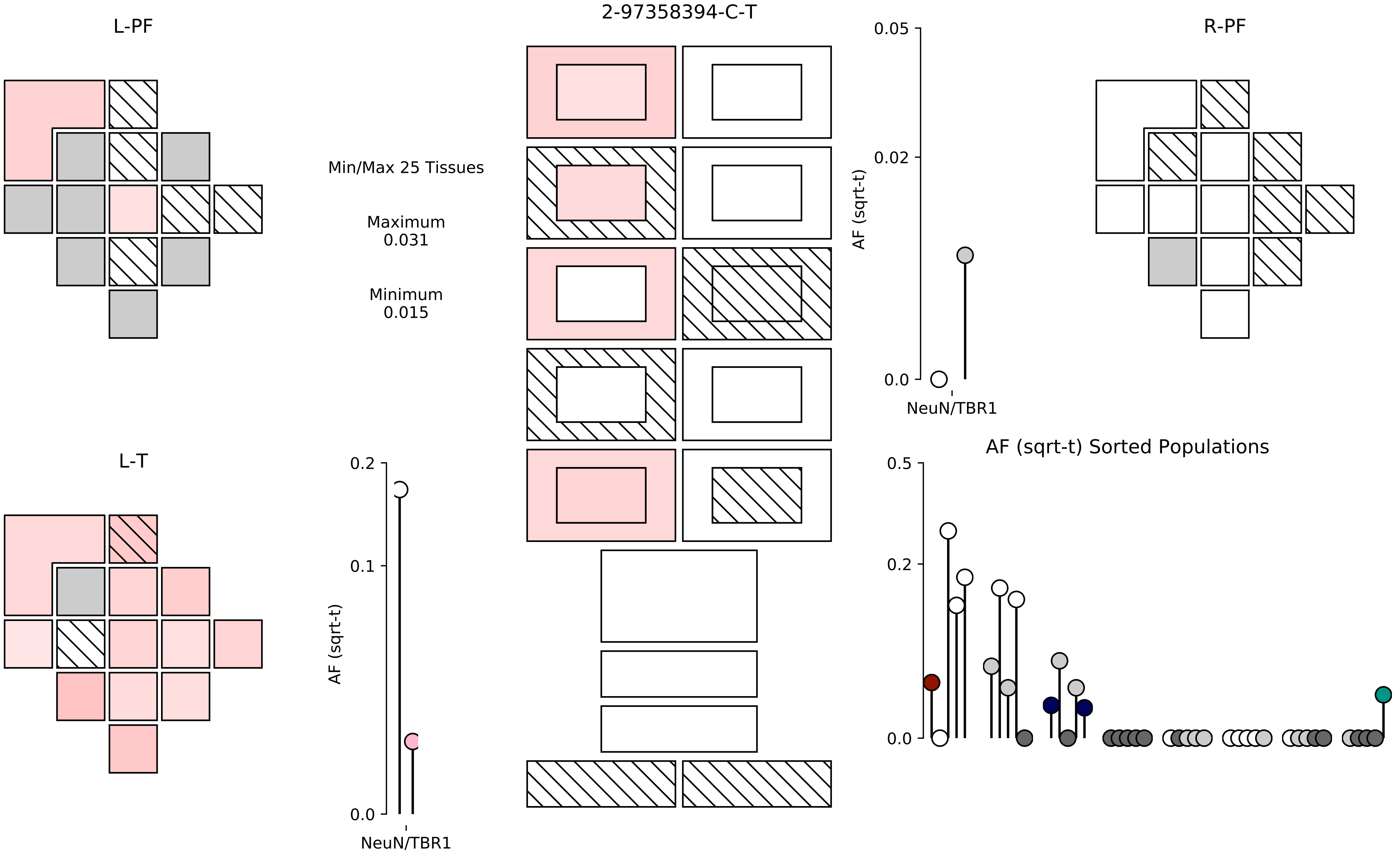

L-PF
2-97358394-C-T
R-PF
Min/Max 25 Tissues
Maximum
0.031
Minimum
0.015
0.05
0.02
0.0
AF (sqrt-t)
NeuN/TBR1
L-T
0.2
0.1
0.0
AF (sqrt-t)
NeuN/TBR1
AF (sqrt-t) Sorted Populations
0.5
0.2
0.0

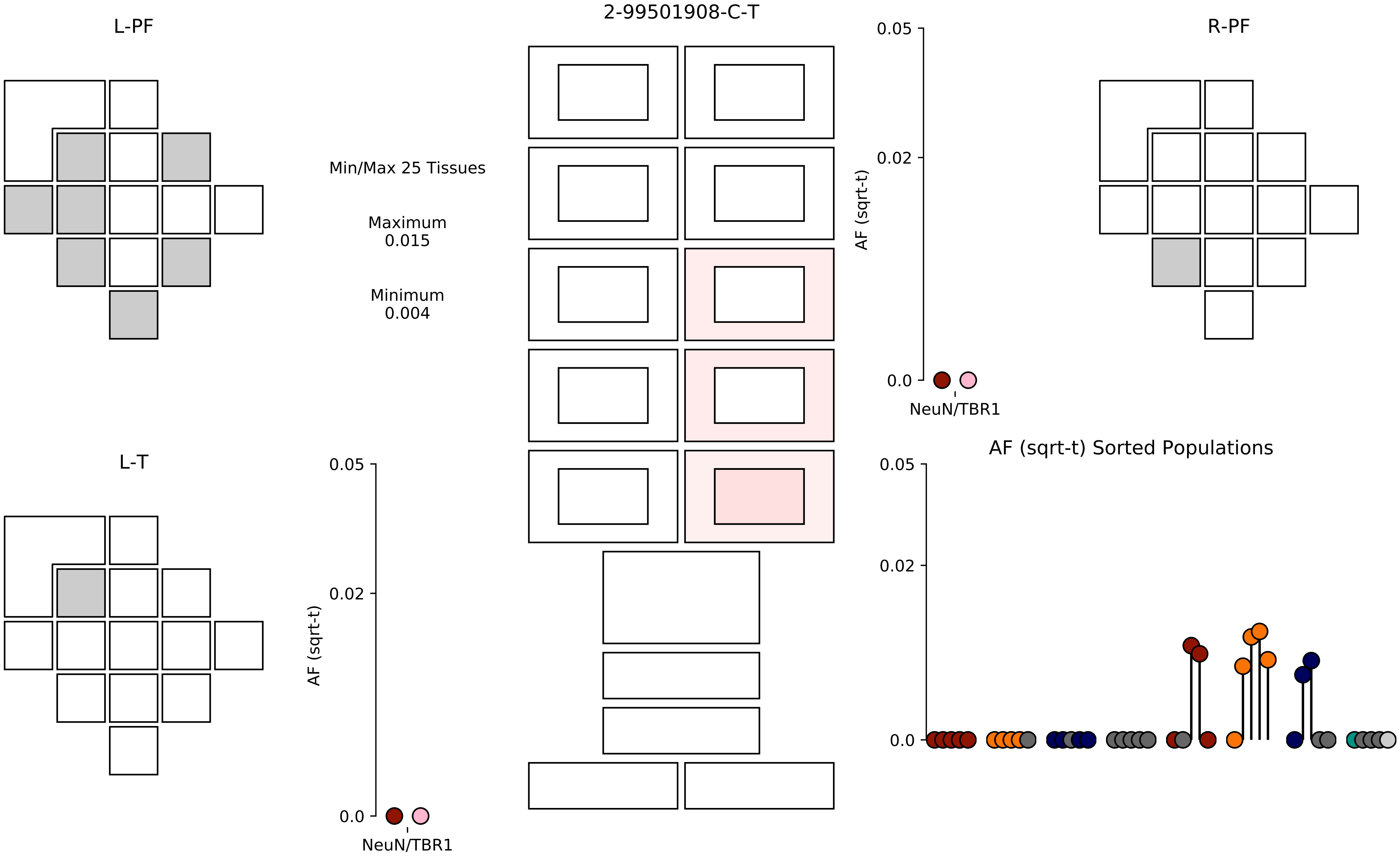
2-99501908-C-T
L-PF
R-PF
L-T
Min/Max 25 Tissues
Maximum
0.015
Minimum
0.004
AF (sqrt-t)
0.05
0.02
0.0
NeuN/TBR1
AF (sqrt-t) Sorted Populations

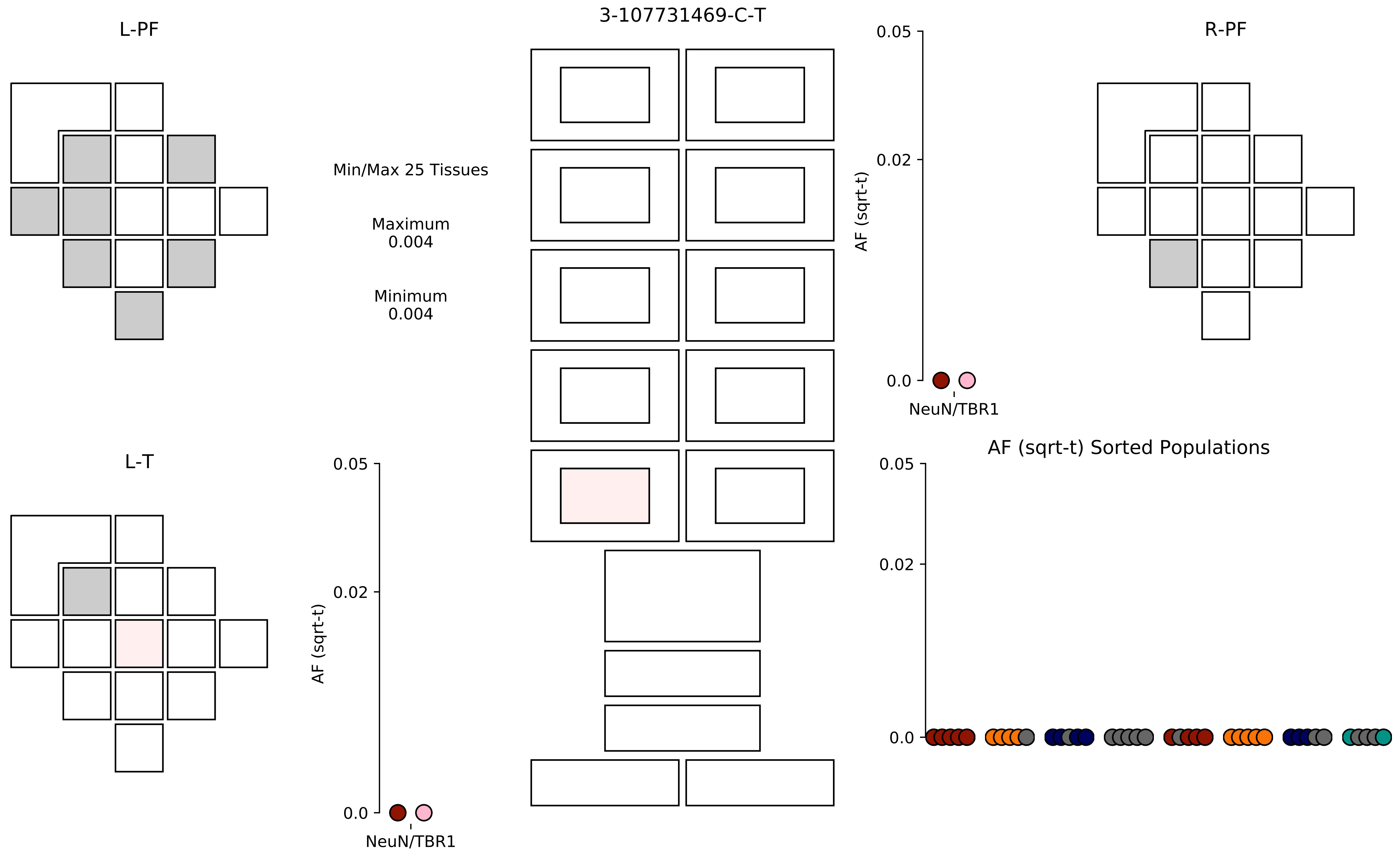
L-PF
3-107731469-C-T
R-PF
Min/Max 25 Tissues
Maximum
0.004
Minimum
0.004
AF (sqrt-t)
0.05
0.02
0.0
NeuN/TBR1
L-T
AF (sqrt-t) Sorted Populations

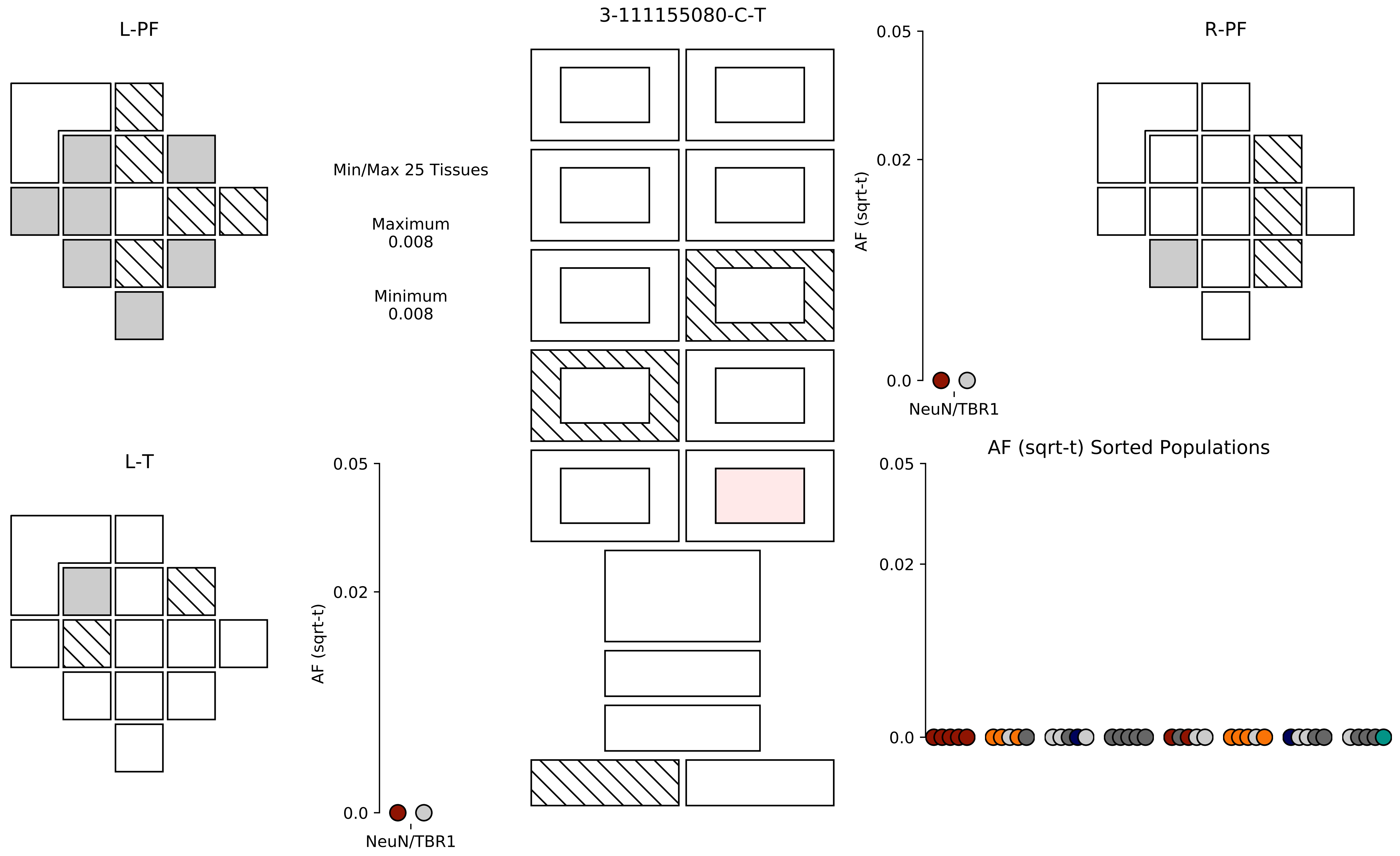

3-111155080-C-T
L-PF
R-PF
L-T
Min/Max 25 Tissues
Maximum
0.008
Minimum
0.008
AF (sqrt-t)
0.05
0.02
0.0
NeuN/TBR1
AF (sqrt-t)
0.05
0.02
0.0
NeuN/TBR1
AF (sqrt-t) Sorted Populations
0.05
0.02
0.0

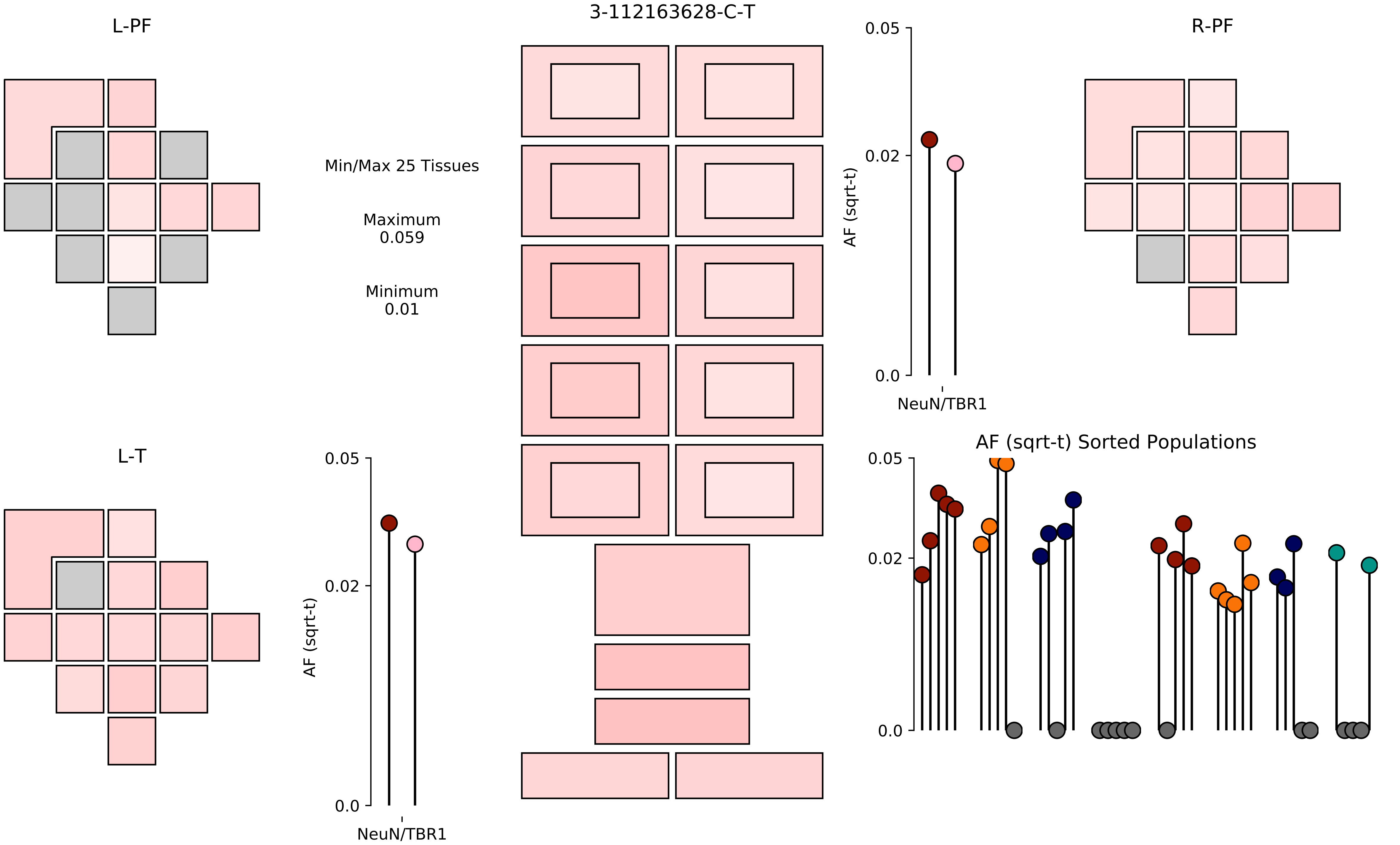

3-112163628-C-T
L-PF
R-PF
L-T
Min/Max 25 Tissues
Maximum
0.059
Minimum
0.01
0.05
0.02
0.0
AF (sqrt-t)
NeuN/TBR1
AF (sqrt-t) Sorted Populations

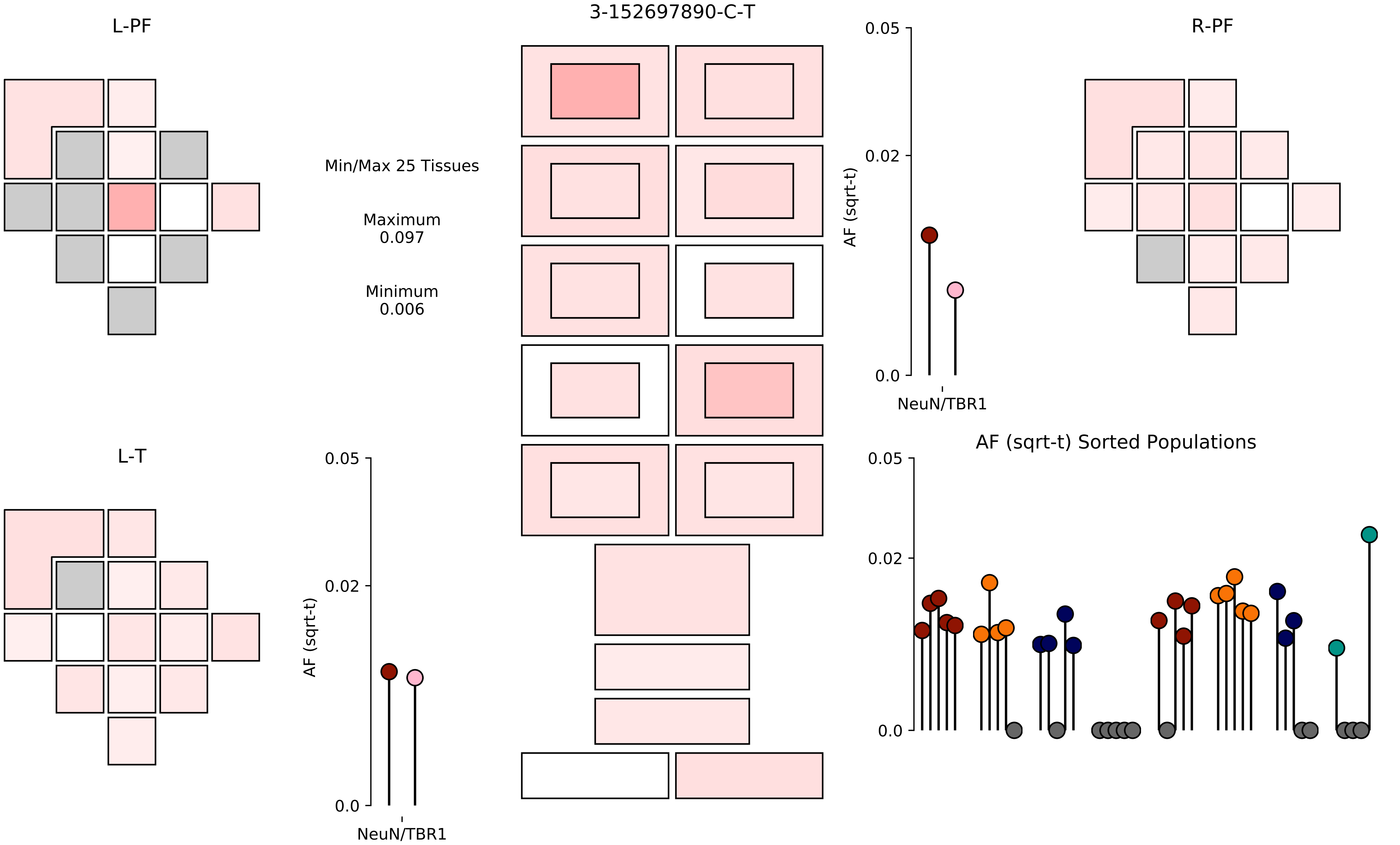

3-152697890-C-T
L-PF
R-PF
L-T
Min/Max 25 Tissues
Maximum
0.097
Minimum
0.006
AF (sqrt-t)
0.05
0.02
0.0
NeuN/TBR1
AF (sqrt-t)
0.05
0.02
0.0
NeuN/TBR1
AF (sqrt-t) Sorted Populations
0.05
0.02
0.0

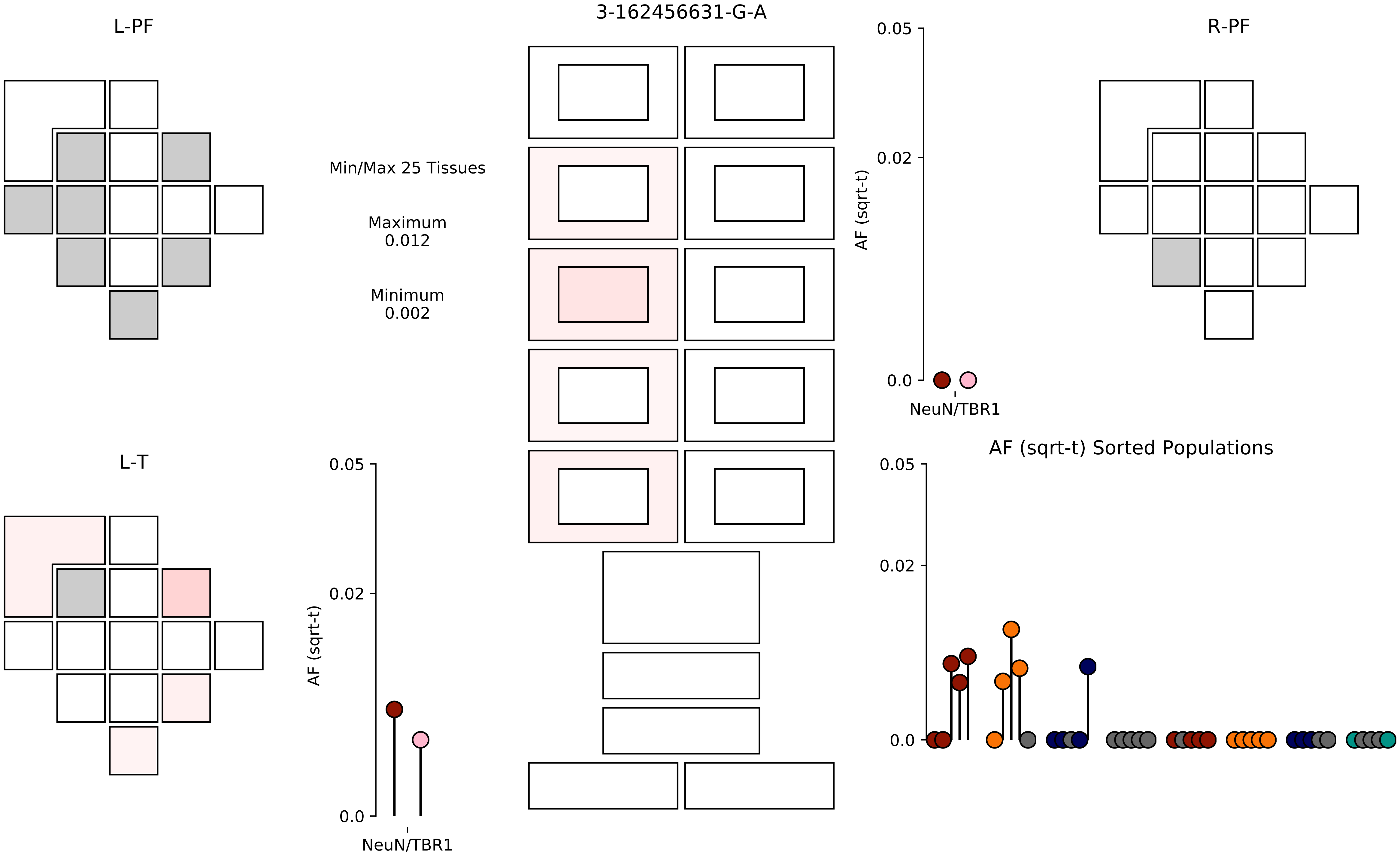
3-162456631-G-A
L-PF
R-PF
L-T
Min/Max 25 Tissues
Maximum
0.012
Minimum
0.002
AF (sqrt-t)
0.05
0.02
0.0
NeuN/TBR1
AF (sqrt-t) Sorted Populations

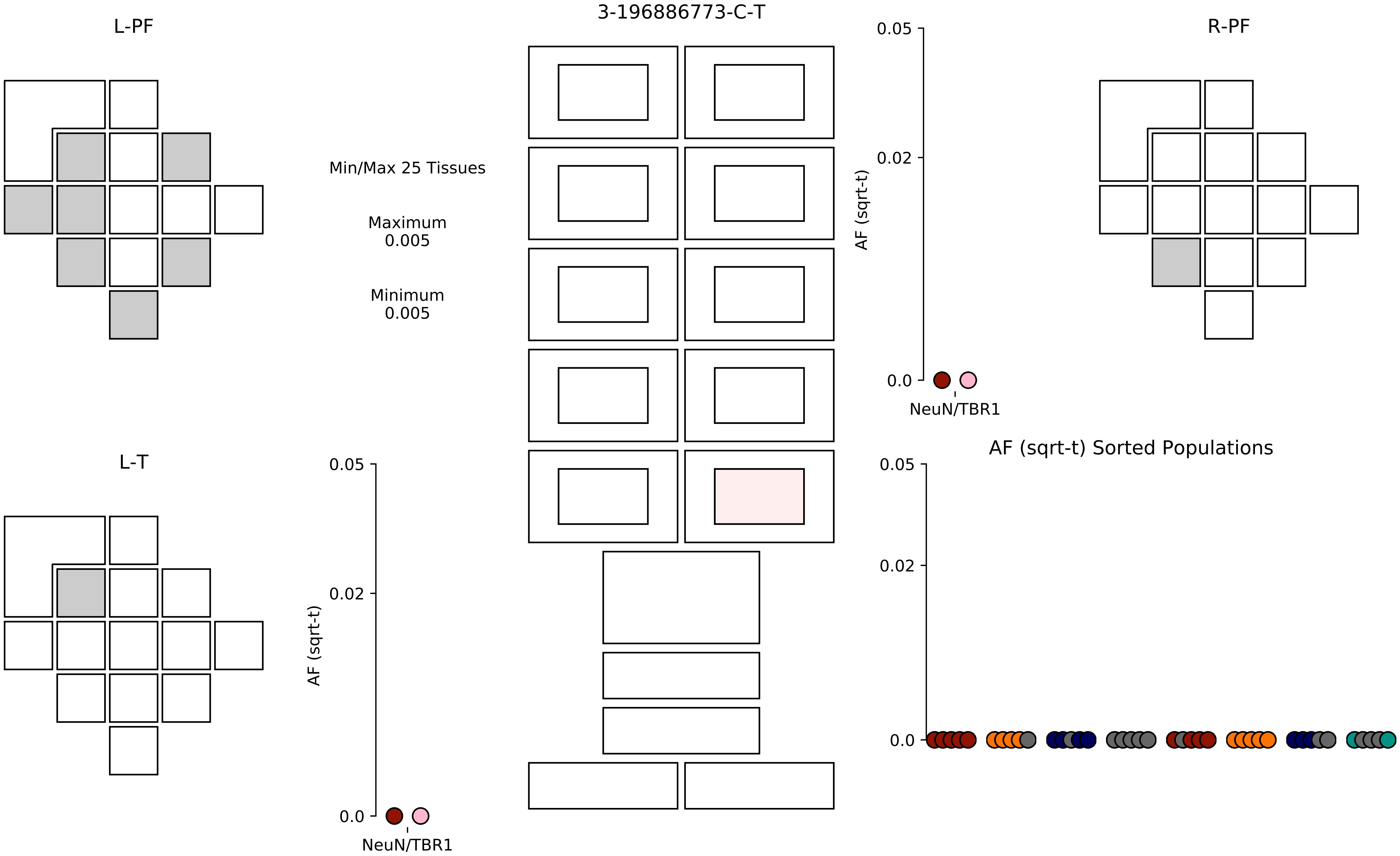

3-196886773-C-T
L-PF
R-PF
L-T
Min/Max 25 Tissues
Maximum
0.005
Minimum
0.005
0.05
0.02
0.0
AF (sqrt-t)
NeuN/TBR1
AF (sqrt-t) Sorted Populations

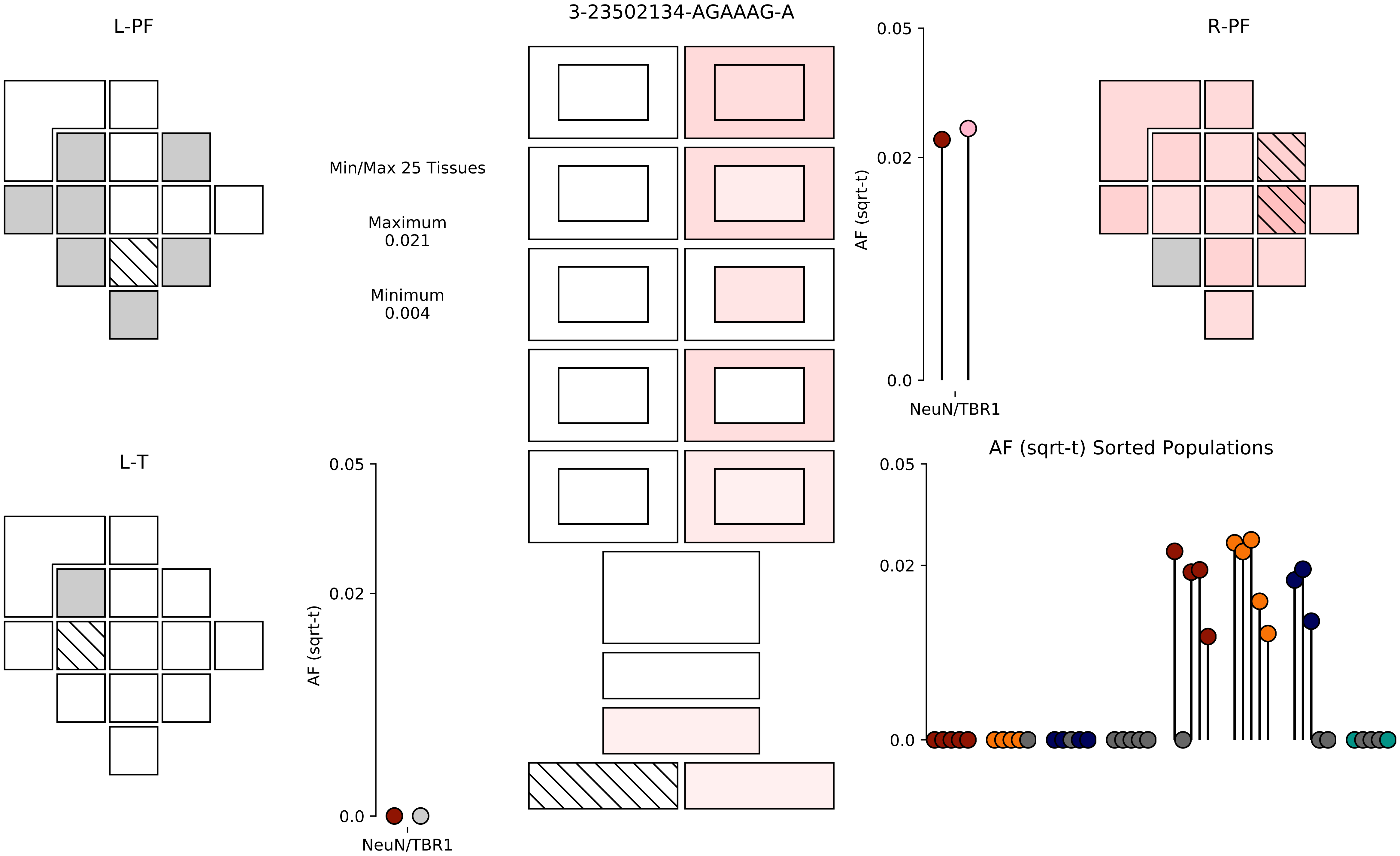
3-23502134-AGAAAG-A
L-PF
R-PF
L-T
Min/Max 25 Tissues
Maximum
0.021
Minimum
0.004
AF (sqrt-t)
0.05
0.02
0.0
NeuN/TBR1
AF (sqrt-t) Sorted Populations

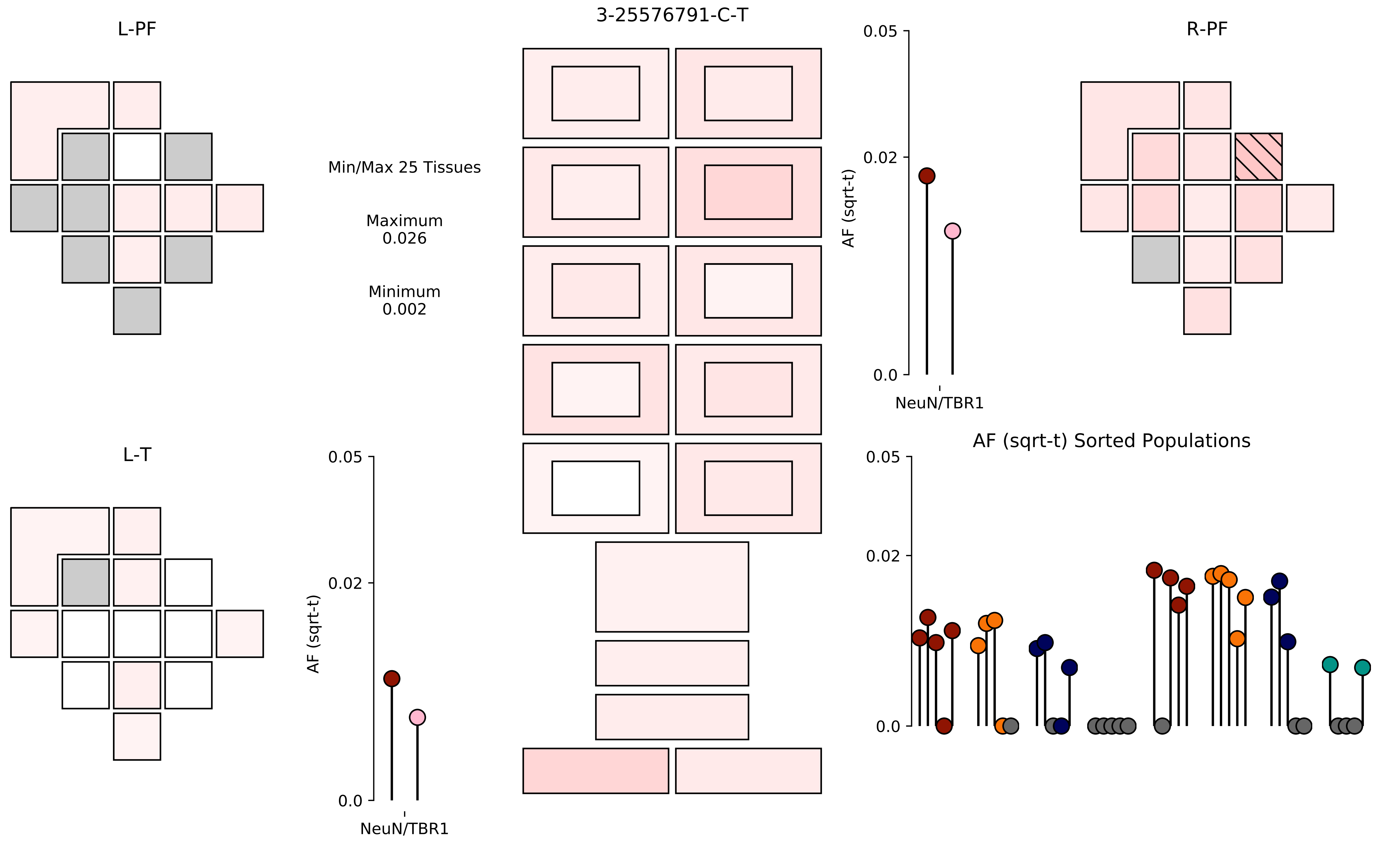

L-PF
3-25576791-C-T
R-PF
Min/Max 25 Tissues
Maximum
0.026
Minimum
0.002
0.05
0.02
0.0
AF (sqrt-t)
NeuN/TBR1
L-T
0.05
0.02
0.0
AF (sqrt-t)
NeuN/TBR1
AF (sqrt-t) Sorted Populations
0.05
0.02
0.0

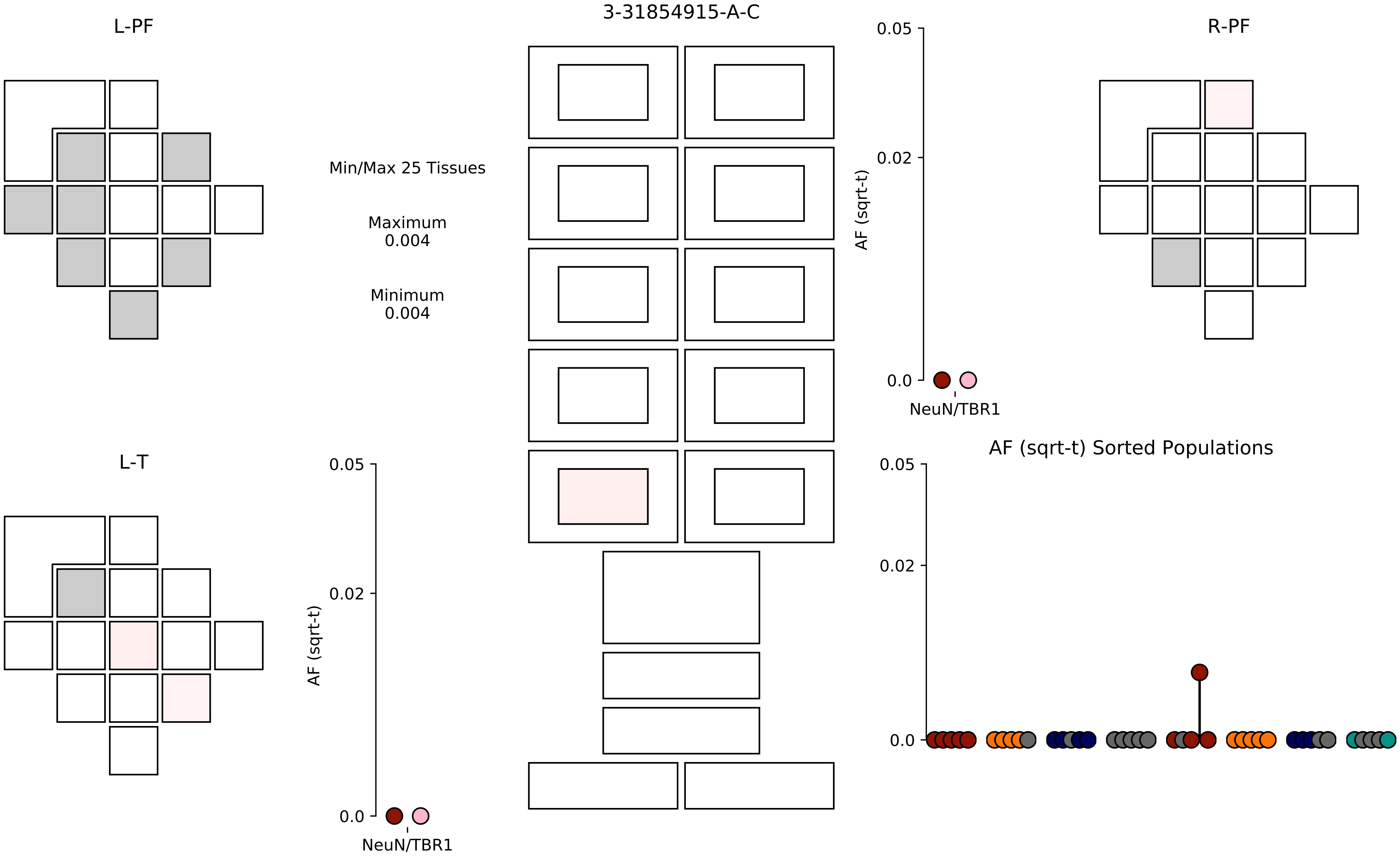
3-31854915-A-C
L-PF
R-PF
L-T
Min/Max 25 Tissues
Maximum
0.004
Minimum
0.004
AF (sqrt-t)
0.05
0.02
0.0
NeuN/TBR1
AF (sqrt-t) Sorted Populations

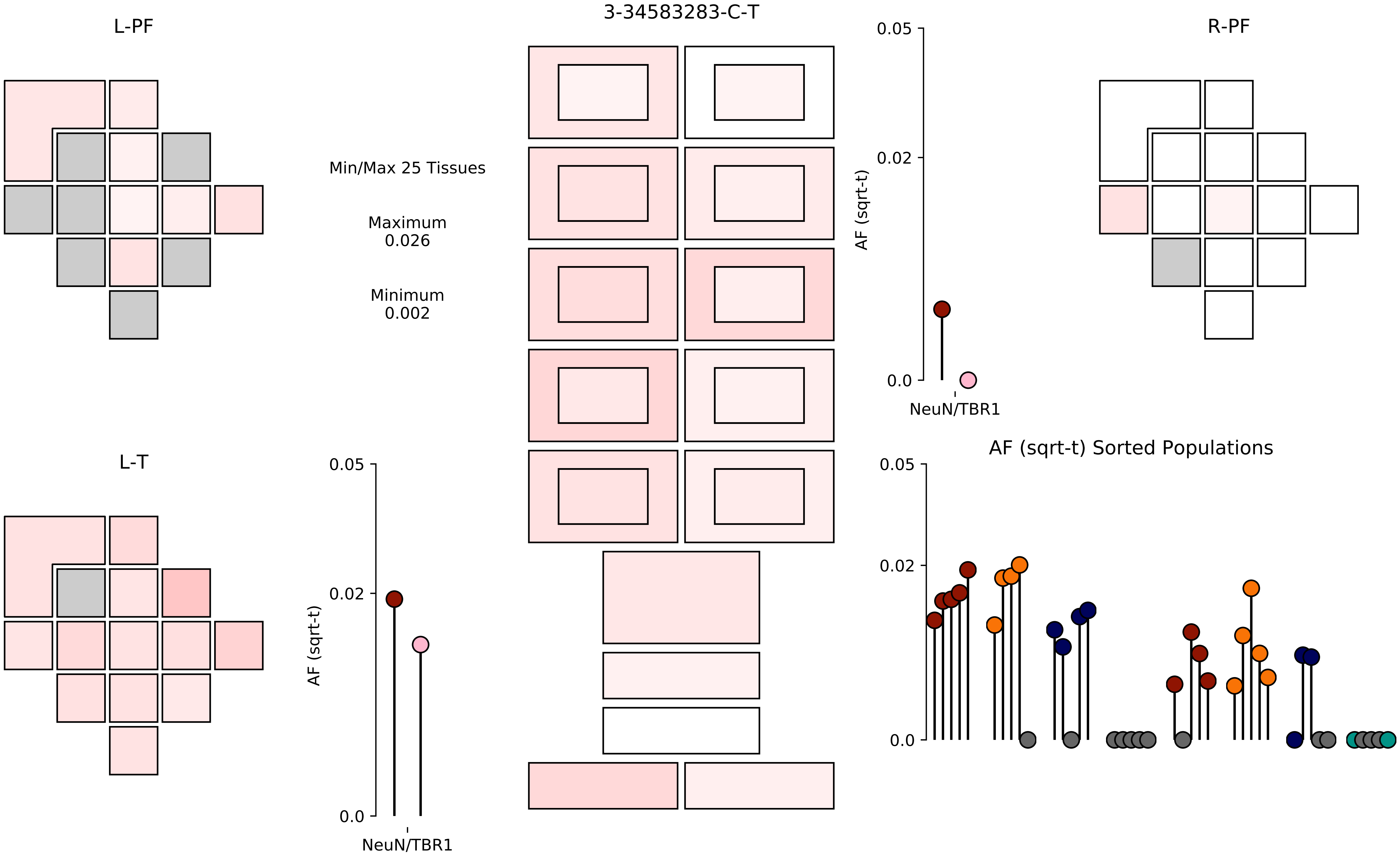
L-PF
3-34583283-C-T
0.05
0.02
0.0
AF (sqrt-t)
NeuN/TBR1
R-PF
Min/Max 25 Tissues
Maximum
0.026
Minimum
0.002
L-T
0.05
0.02
0.0
AF (sqrt-t)
NeuN/TBR1
AF (sqrt-t) Sorted Populations
0.05
0.02
0.0

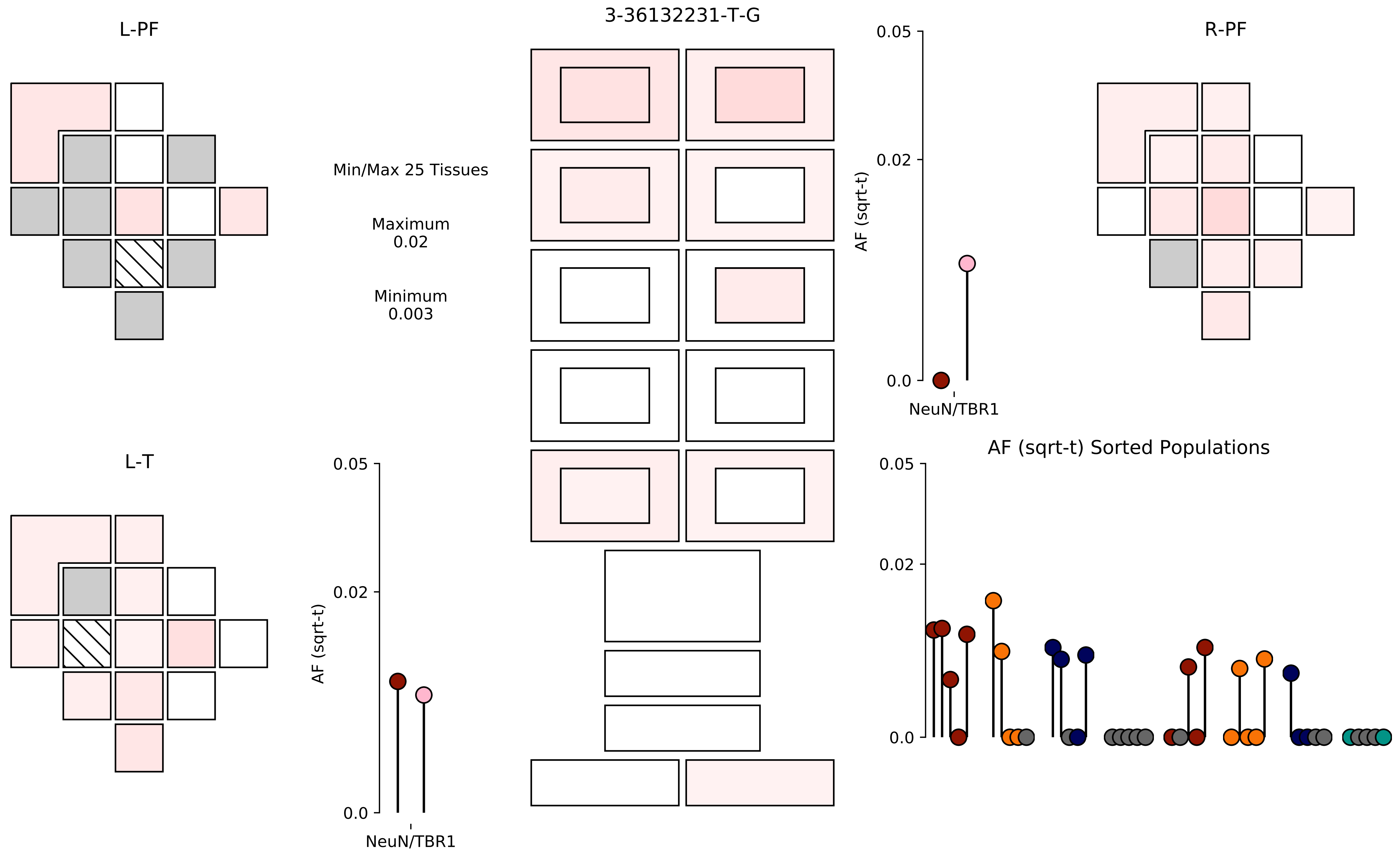
3-36132231-T-G
L-PF
R-PF
L-T
Min/Max 25 Tissues
Maximum
0.02
Minimum
0.003
AF (sqrt-t)
0.05
0.02
0.0
NeuN/TBR1
AF (sqrt-t) Sorted Populations

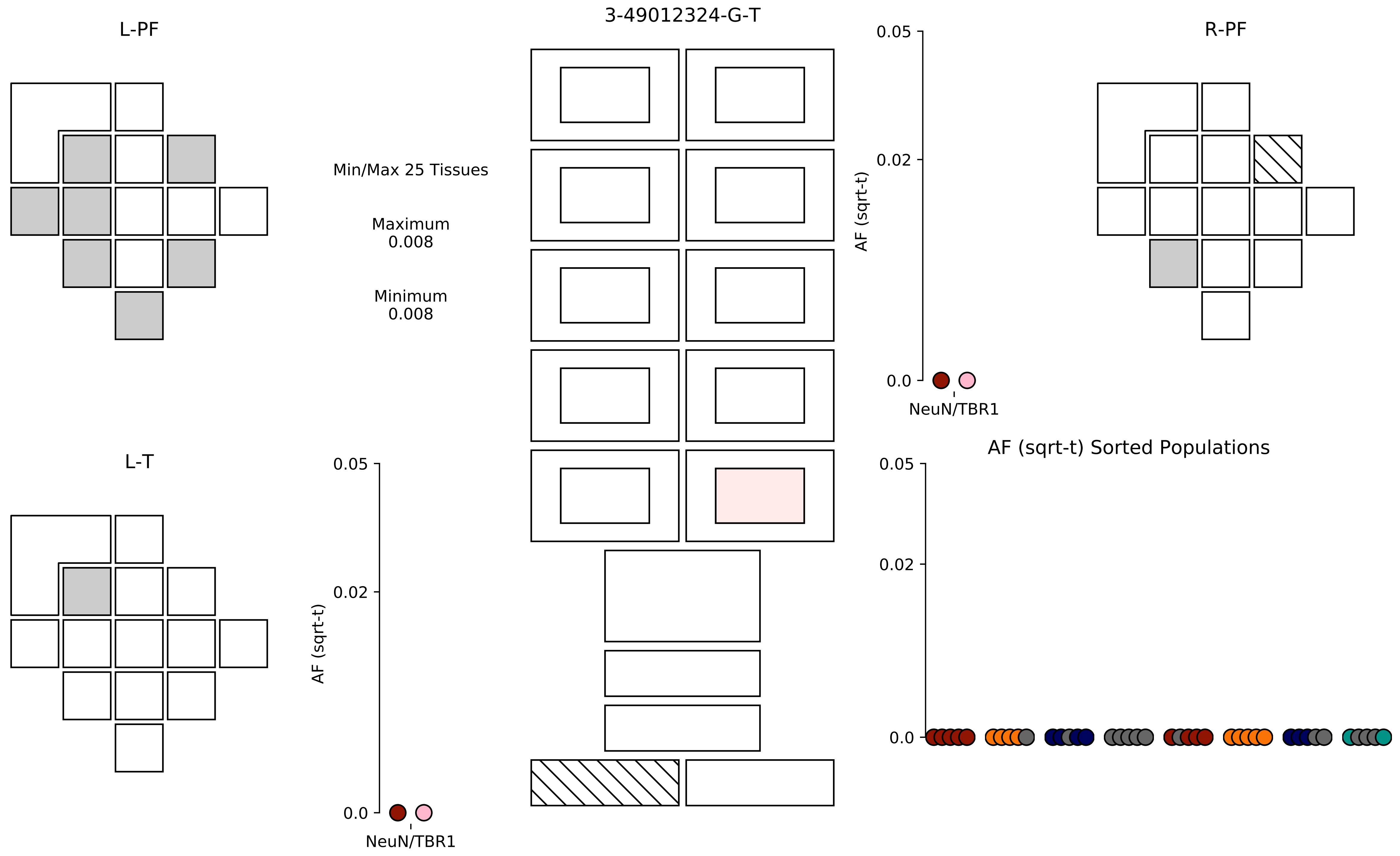

3-49012324-G-T
L-PF
R-PF
L-T
Min/Max 25 Tissues
Maximum
0.008
Minimum
0.008
AF (sqrt-t)
0.05
0.02
0.0
NeuN/TBR1
AF (sqrt-t) Sorted Populations

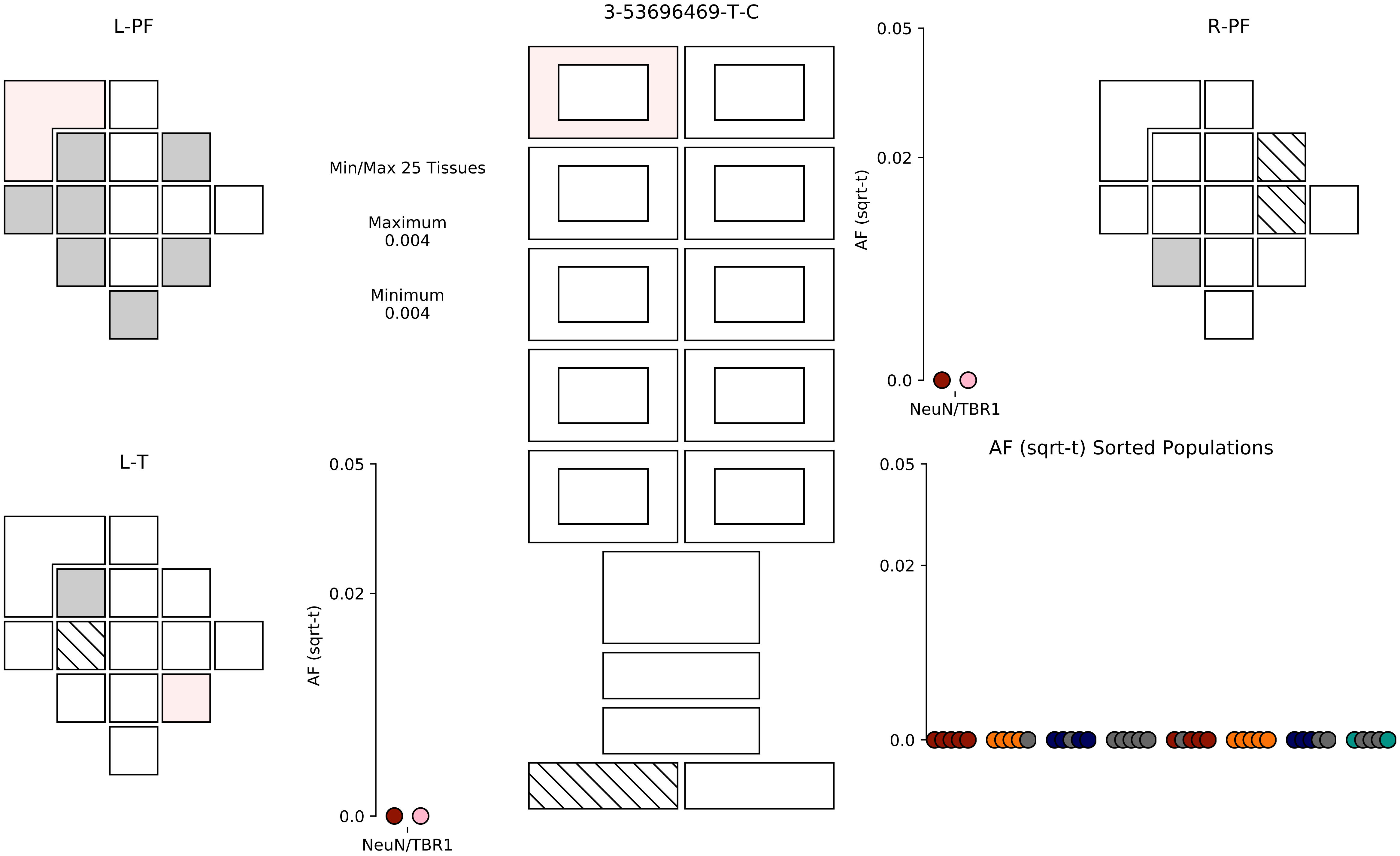
3-53696469-T-C
L-PF
R-PF
L-T
Min/Max 25 Tissues
Maximum
0.004
Minimum
0.004
AF (sqrt-t)
0.05
0.02
0.0
NeuN/TBR1
AF (sqrt-t) Sorted Populations

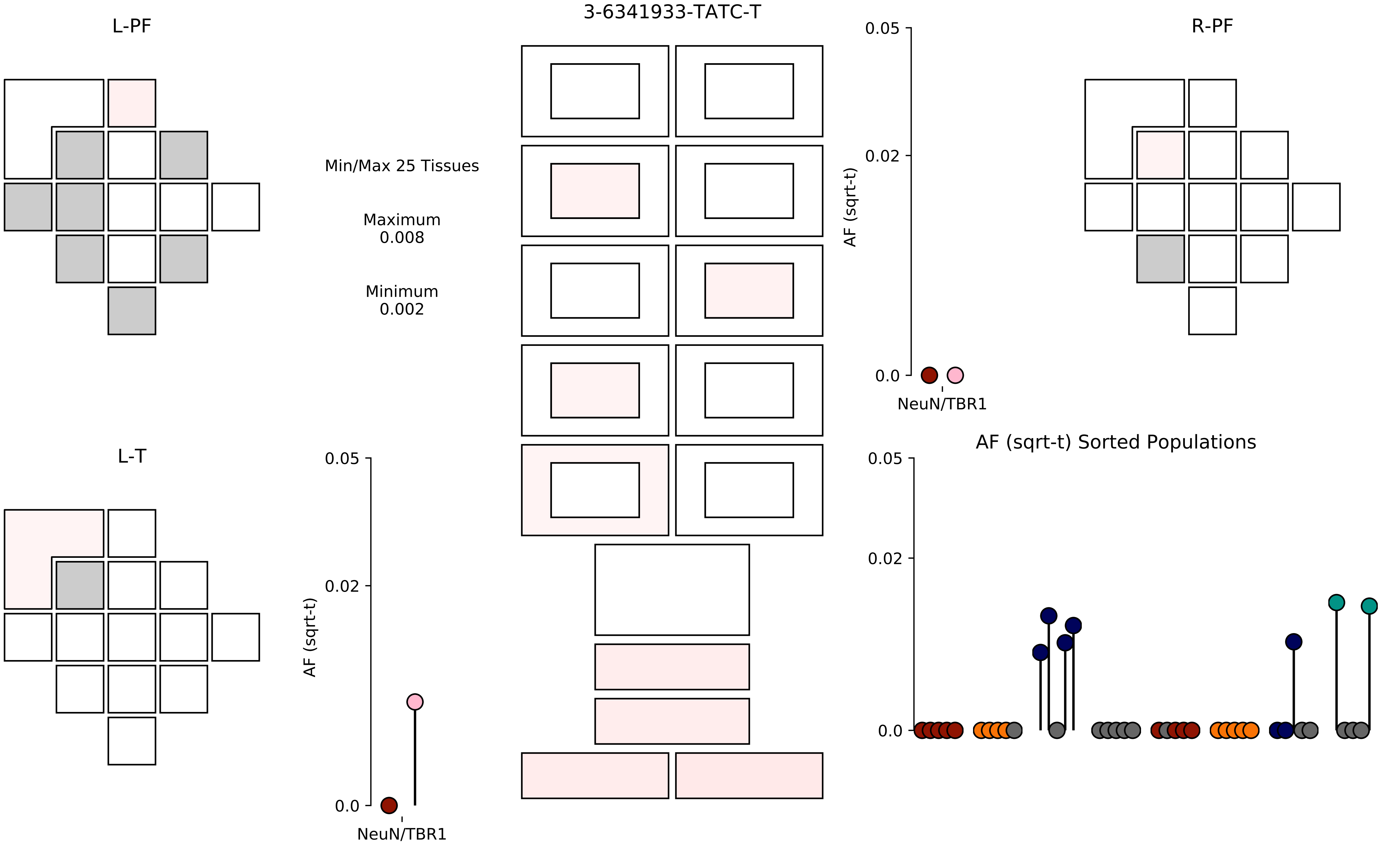

L-PF
3-6341933-TATC-T
R-PF
Min/Max 25 Tissues
Maximum
0.008
Minimum
0.002
0.05
0.02
0.0
AF (sqrt-t)
NeuN/TBR1
L-T
0.05
0.02
0.0
AF (sqrt-t)
NeuN/TBR1
AF (sqrt-t) Sorted Populations
0.05
0.02
0.0

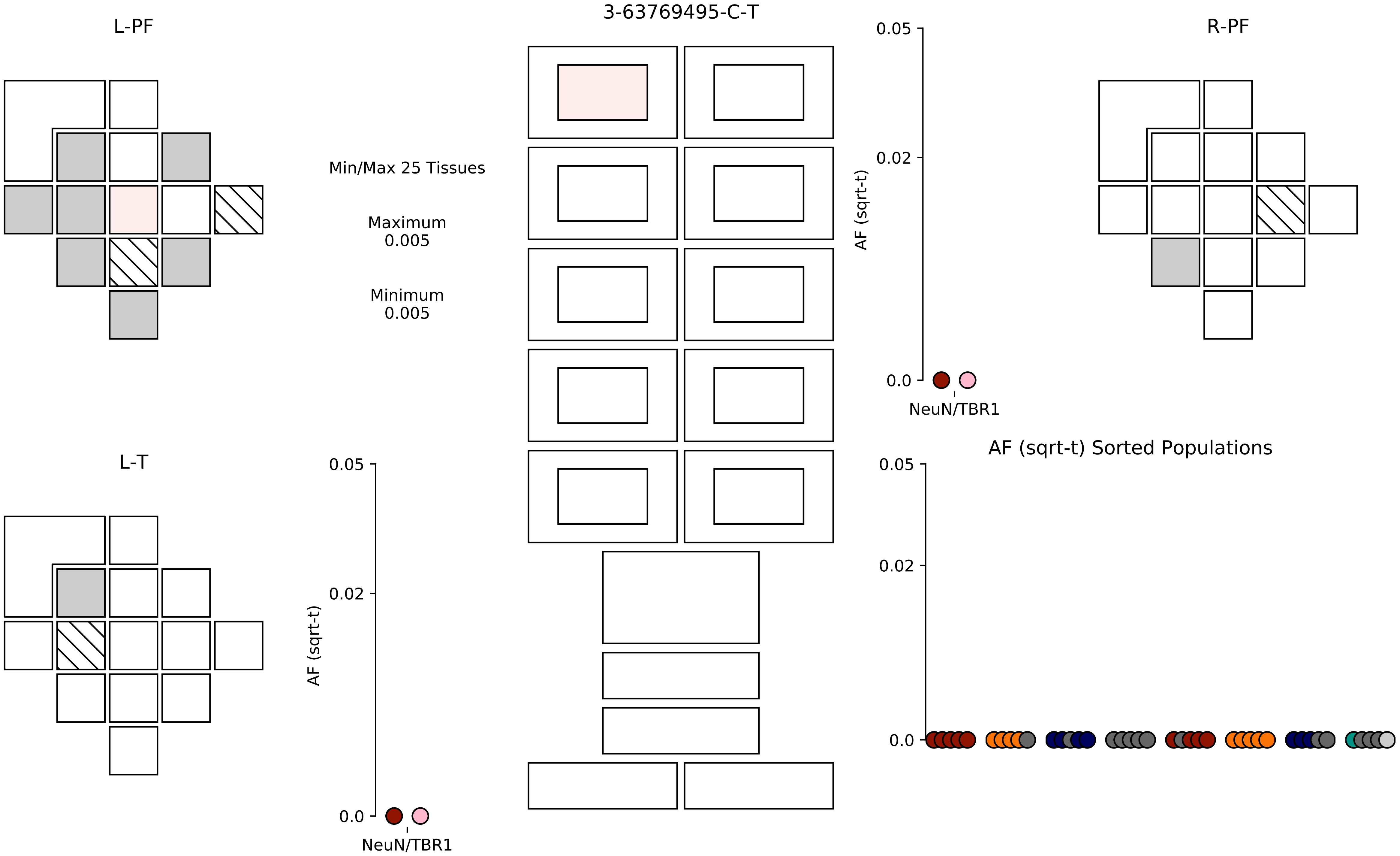

3-63769495-C-T
L-PF
R-PF
L-T
Min/Max 25 Tissues
Maximum
0.005
Minimum
0.005
AF (sqrt-t)
0.05
0.02
0.0
NeuN/TBR1
AF (sqrt-t) Sorted Populations

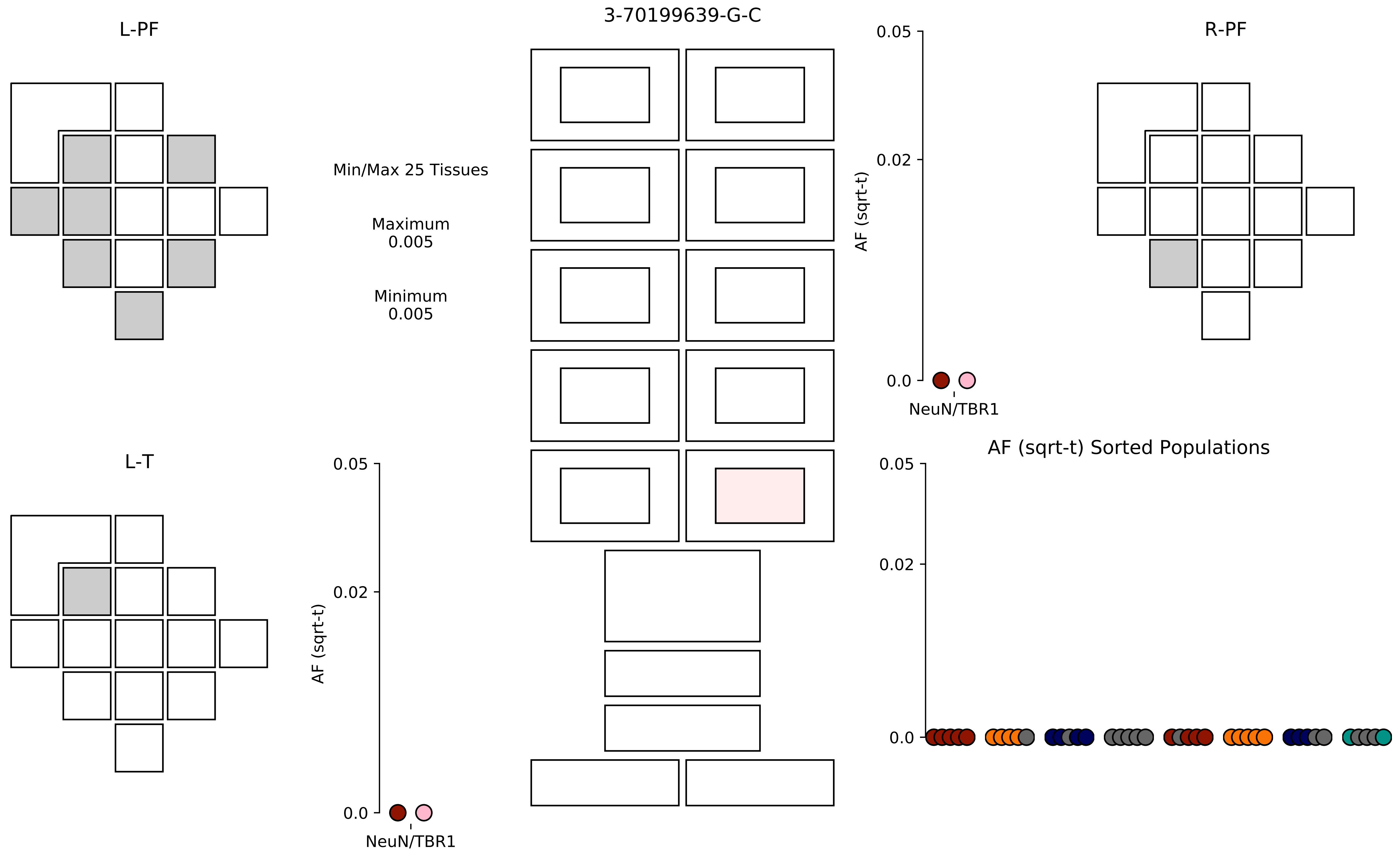

L-PF
3-70199639-G-C
R-PF
Min/Max 25 Tissues
Maximum
0.005
Minimum
0.005
0.05
0.02
0.0
AF (sqrt-t)
NeuN/TBR1
L-T
AF (sqrt-t) Sorted Populations

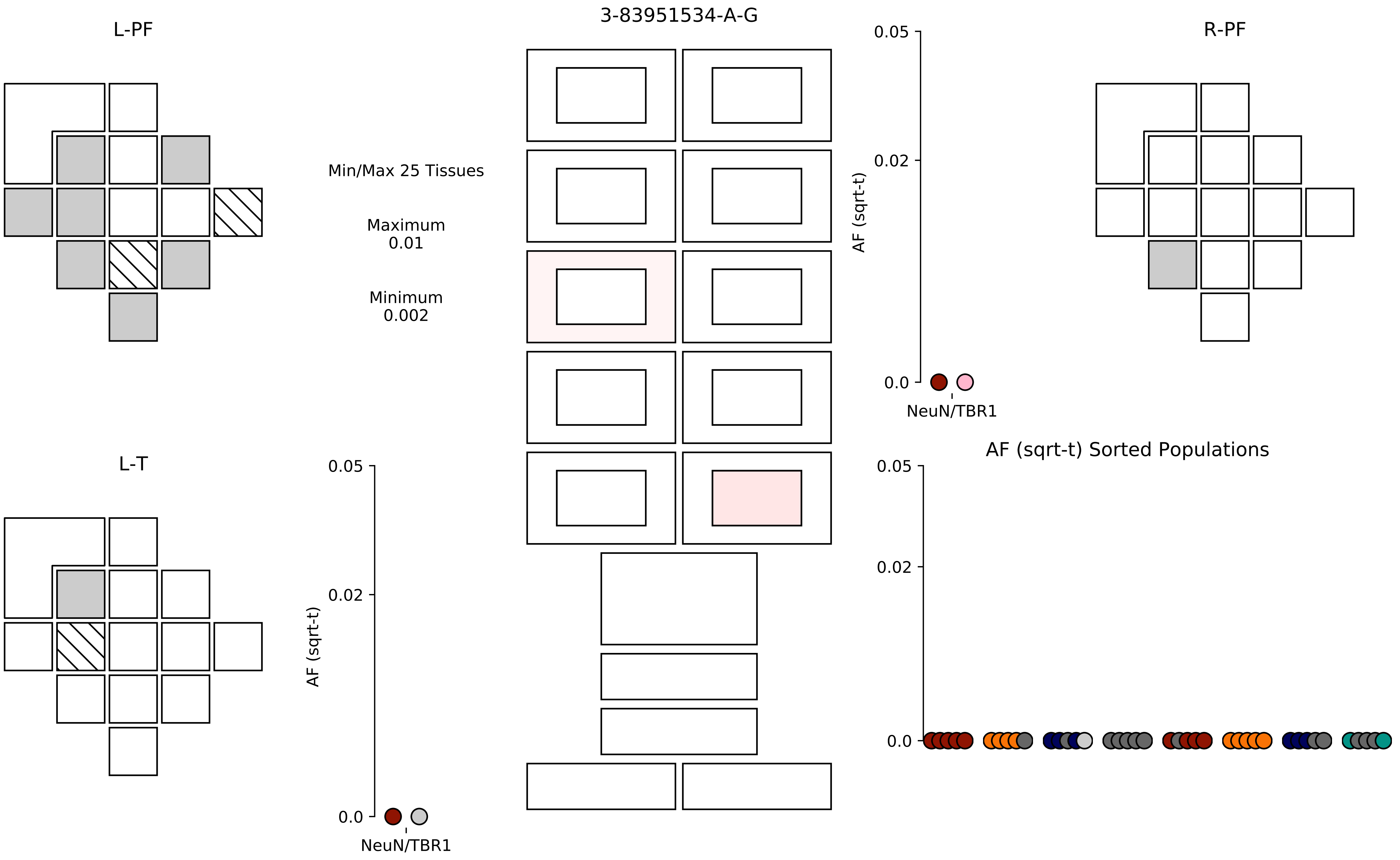

3-83951534-A-G
L-PF
R-PF
L-T
Min/Max 25 Tissues
Maximum
0.01
Minimum
0.002
AF (sqrt-t)
0.05
0.02
0.0
NeuN/TBR1
AF (sqrt-t) Sorted Populations

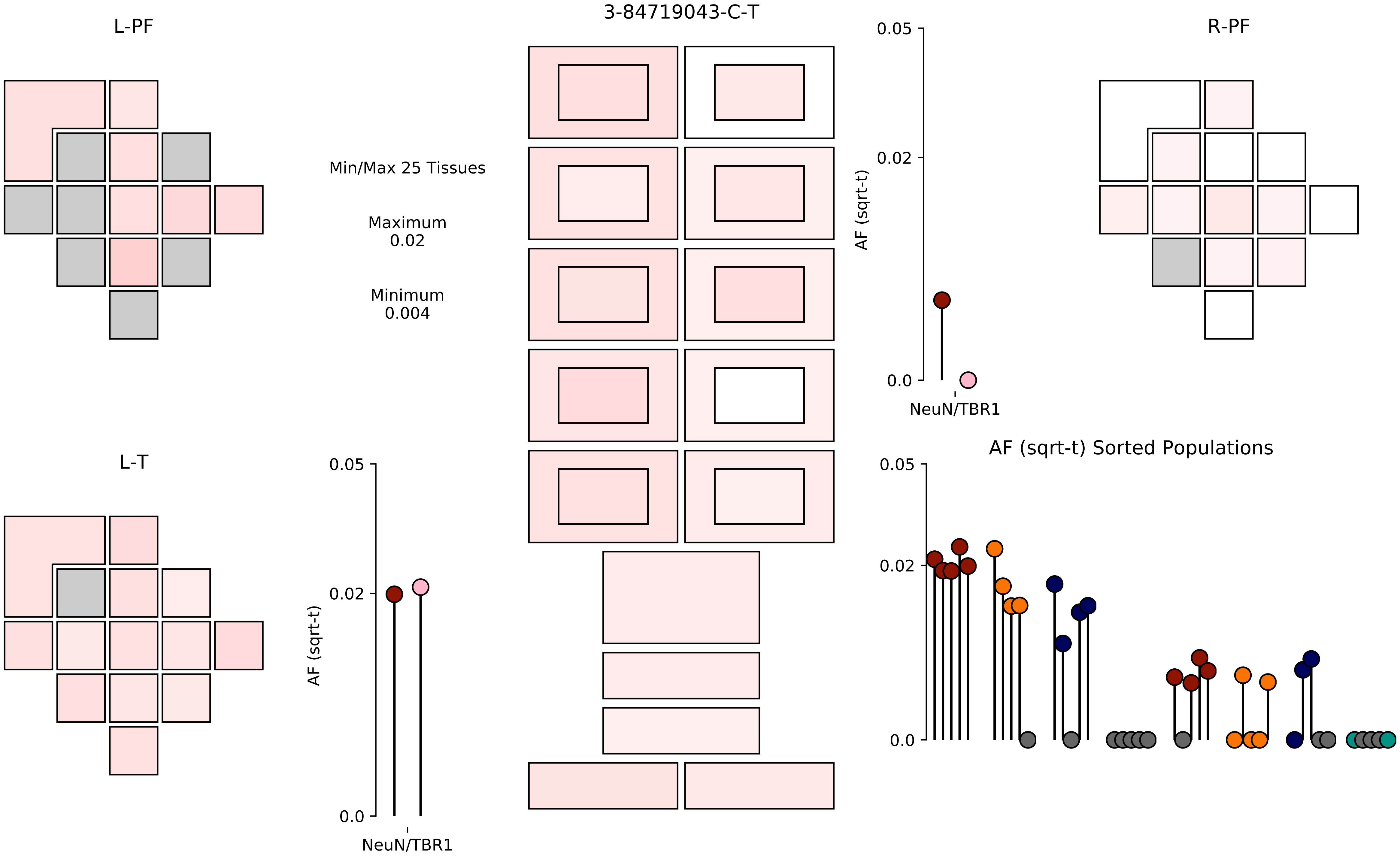
3-84719043-C-T
L-PF
R-PF
L-T
Min/Max 25 Tissues
Maximum
0.02
Minimum
0.004
0.05
0.02
0.0
AF (sqrt-t)
NeuN/TBR1
AF (sqrt-t) Sorted Populations

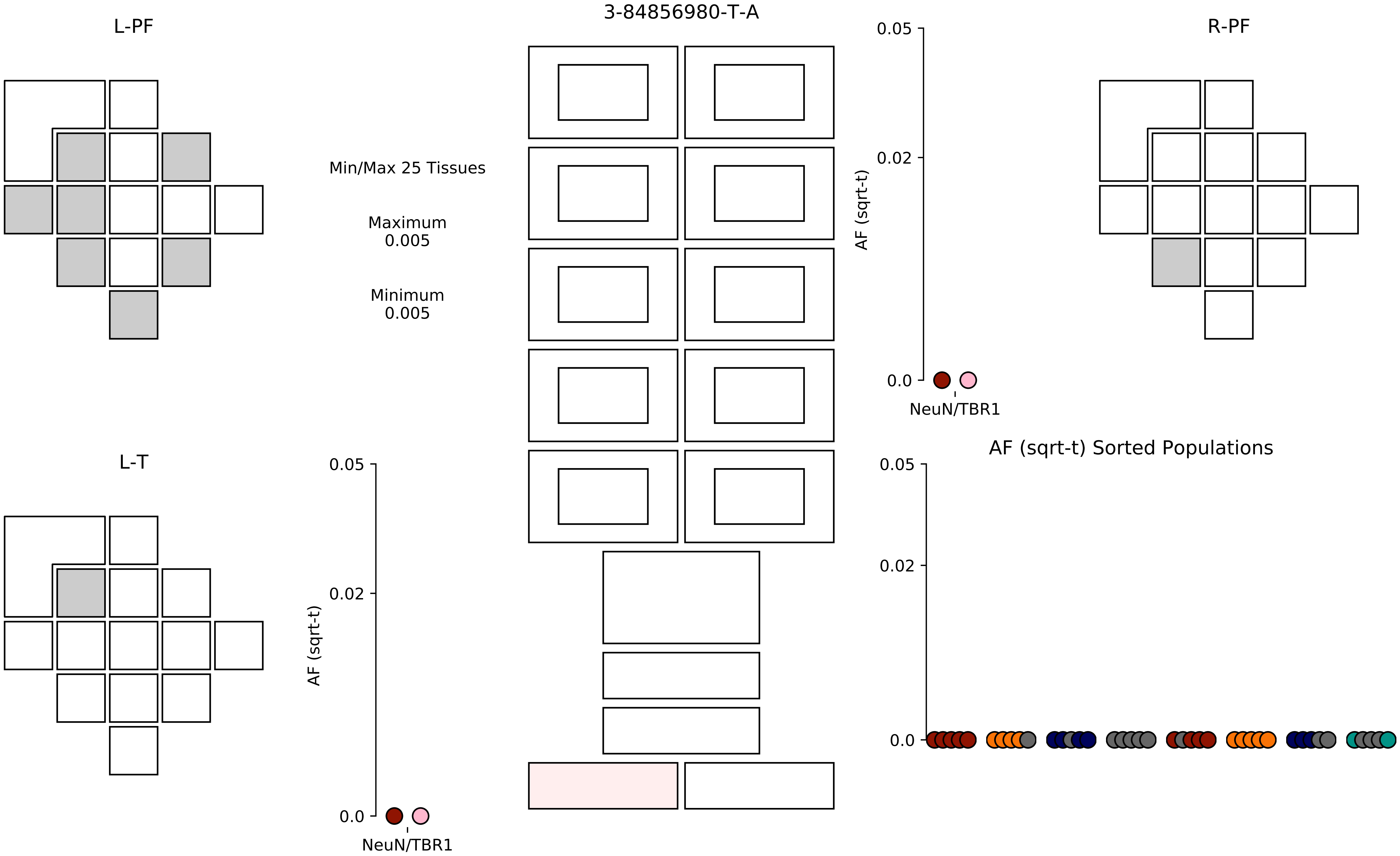
3-84856980-T-A
L-PF
R-PF
L-T
Min/Max 25 Tissues
Maximum
0.005
Minimum
0.005
0.05
0.02
0.0
AF (sqrt-t)
NeuN/TBR1
AF (sqrt-t) Sorted Populations

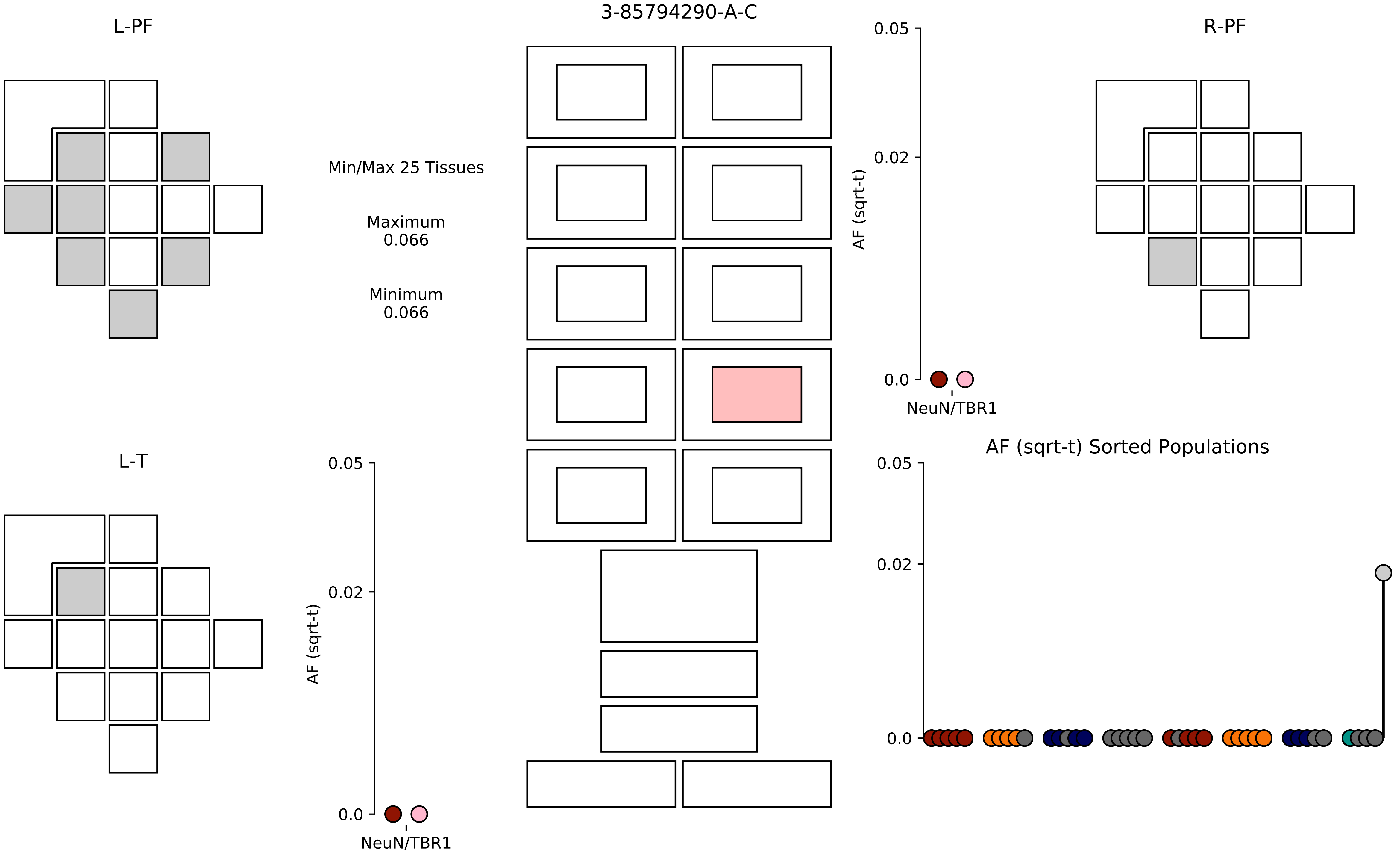

3-85794290-A-C
L-PF
R-PF
L-T
Min/Max 25 Tissues
Maximum
0.066
Minimum
0.066
AF (sqrt-t)
0.05
0.02
0.0
NeuN/TBR1
AF (sqrt-t) Sorted Populations

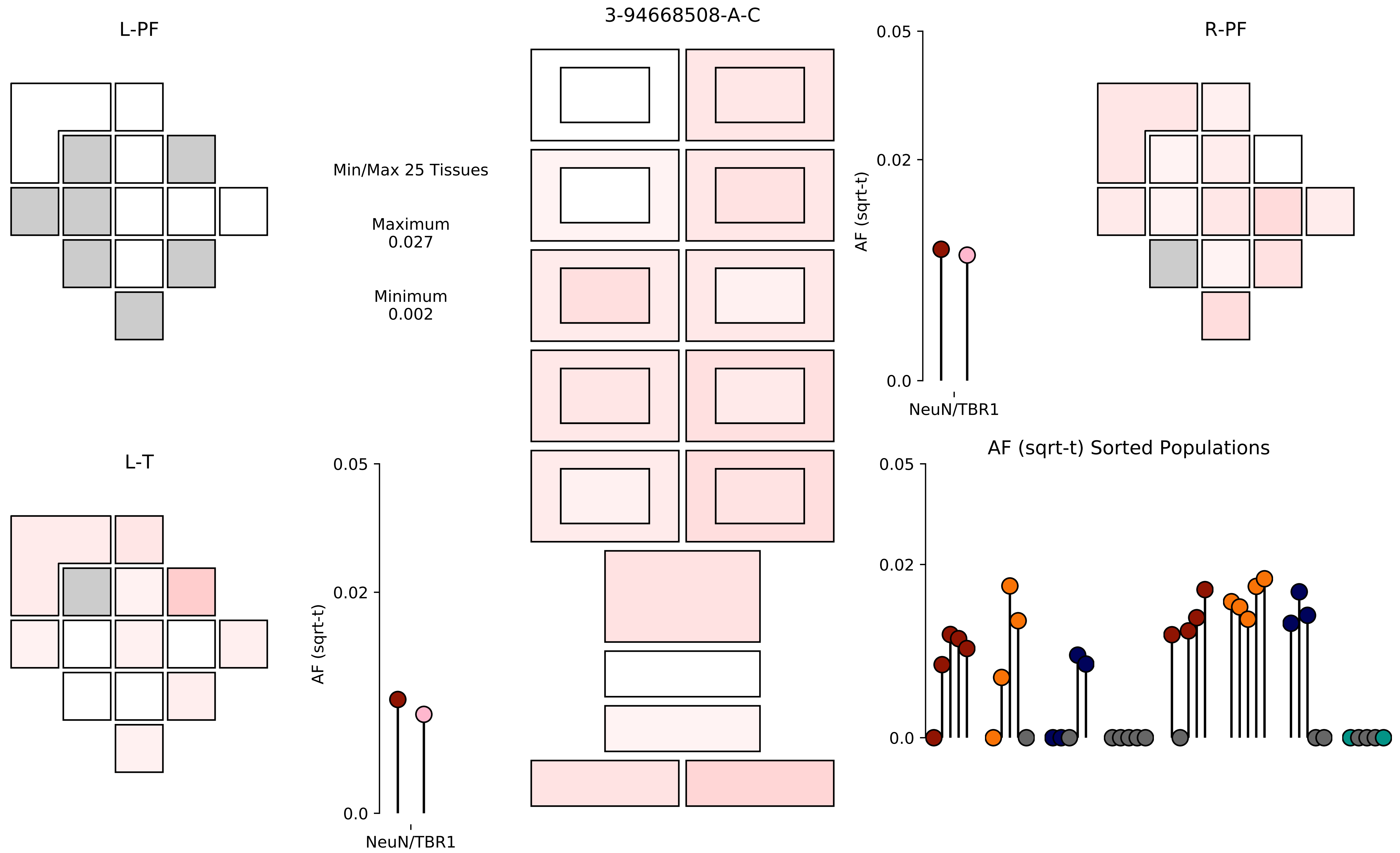

3-94668508-A-C
L-PF
R-PF
L-T
Min/Max 25 Tissues
Maximum
0.027
Minimum
0.002
AF (sqrt-t)
NeuN/TBR1
0.05
0.02
0.0
AF (sqrt-t) Sorted Populations

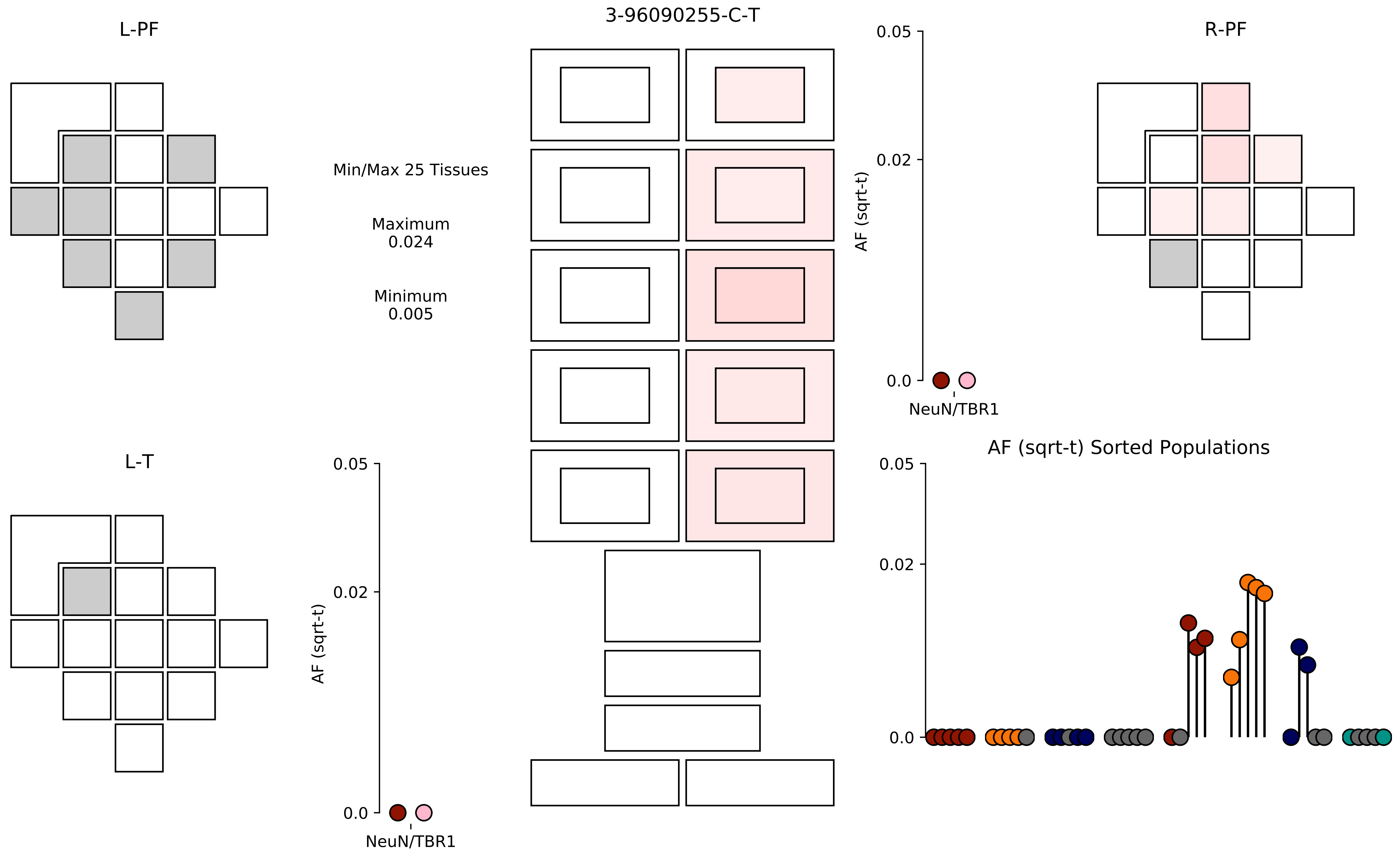

L-PF
3-96090255-C-T
R-PF
Min/Max 25 Tissues
Maximum
0.024
Minimum
0.005
0.05
0.02
0.0
AF (sqrt-t)
NeuN/TBR1
L-T
AF (sqrt-t) Sorted Populations

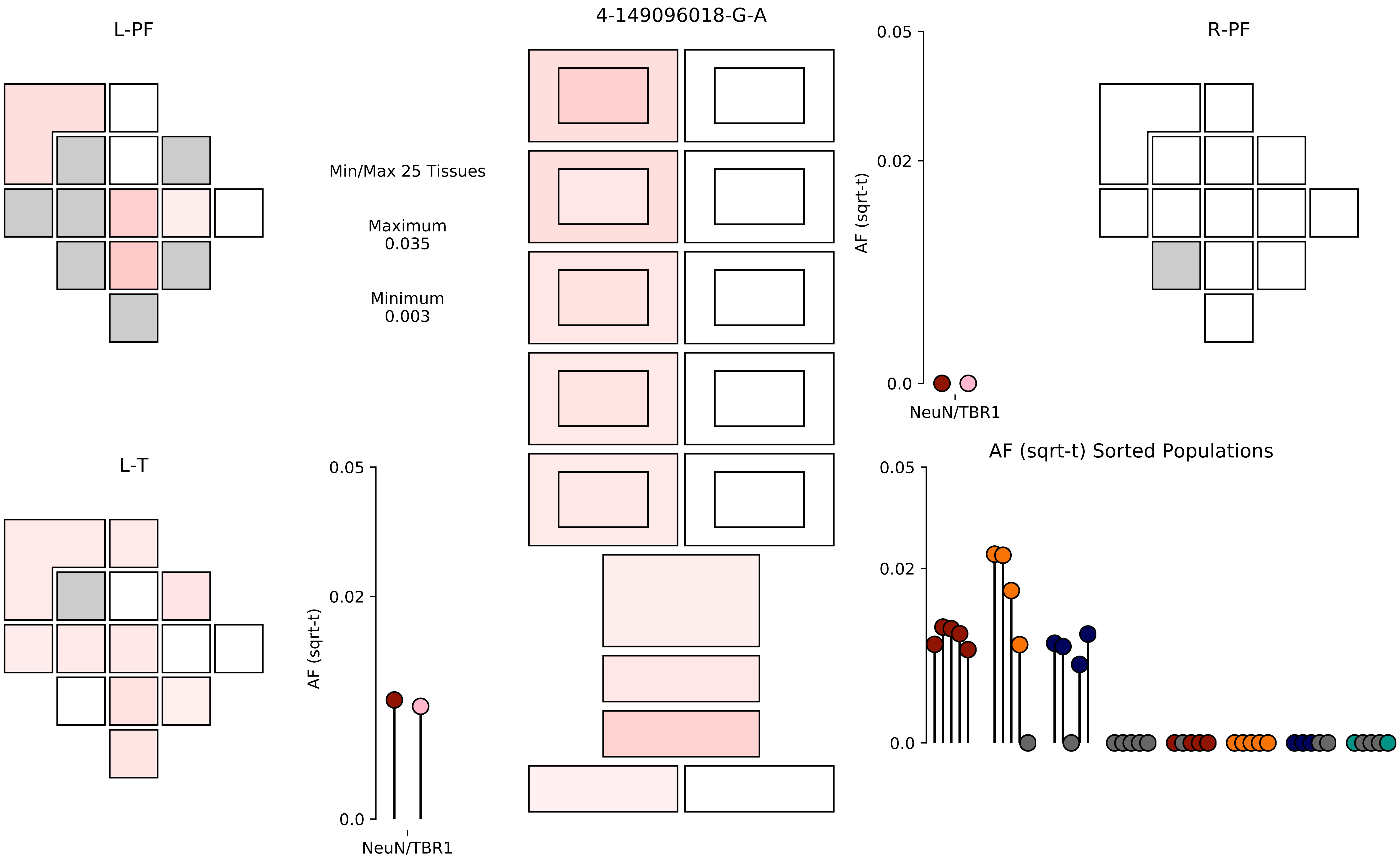

4-149096018-G-A
L-PF
R-PF
L-T
Min/Max 25 Tissues
Maximum
0.035
Minimum
0.003
AF (sqrt-t)
0.05
0.02
0.0
NeuN/TBR1
AF (sqrt-t) Sorted Populations

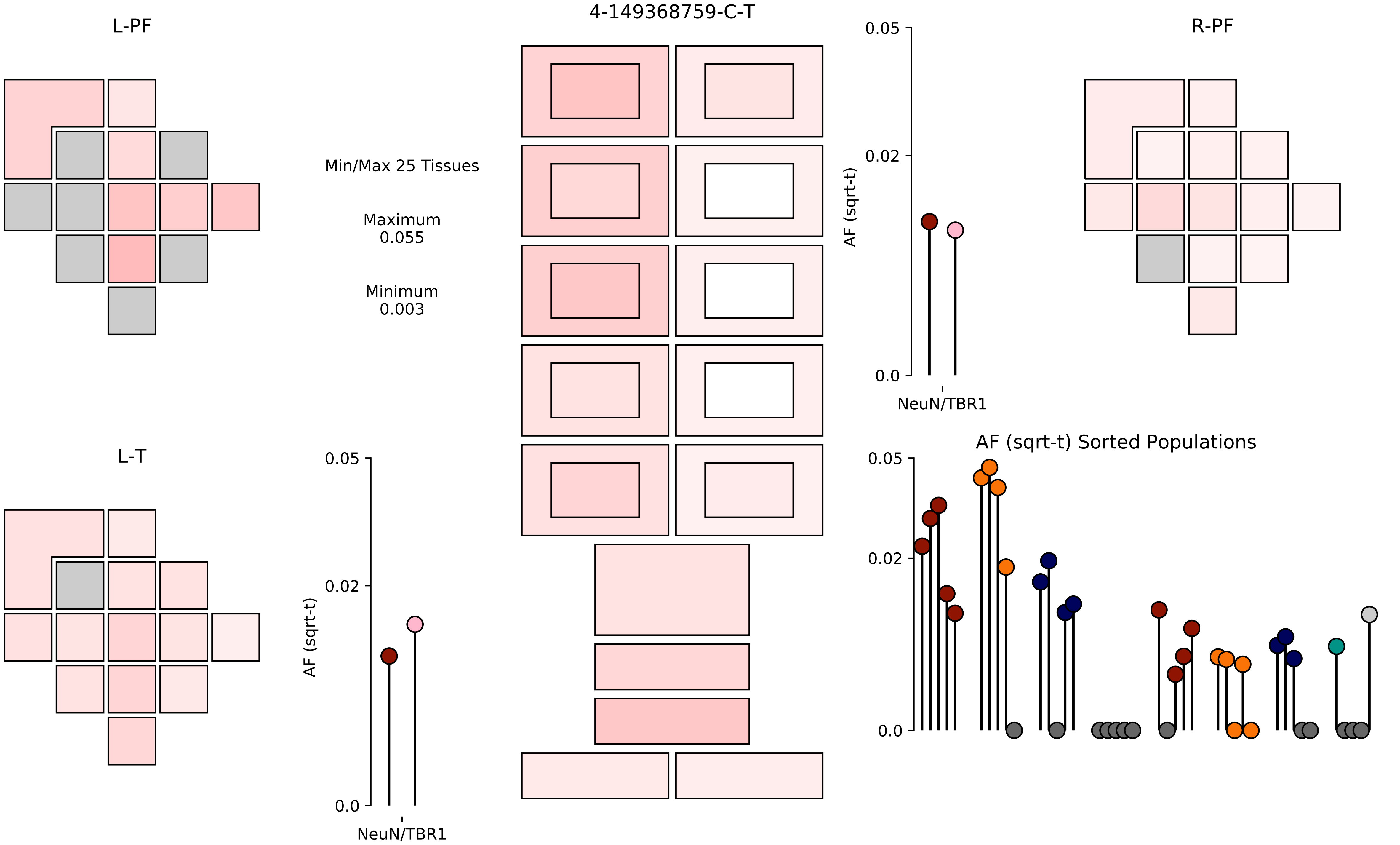

4-149368759-C-T
L-PF
R-PF
L-T
Min/Max 25 Tissues
Maximum
0.055
Minimum
0.003
AF (sqrt-t)
0.05
0.02
0.0
NeuN/TBR1
AF (sqrt-t) Sorted Populations

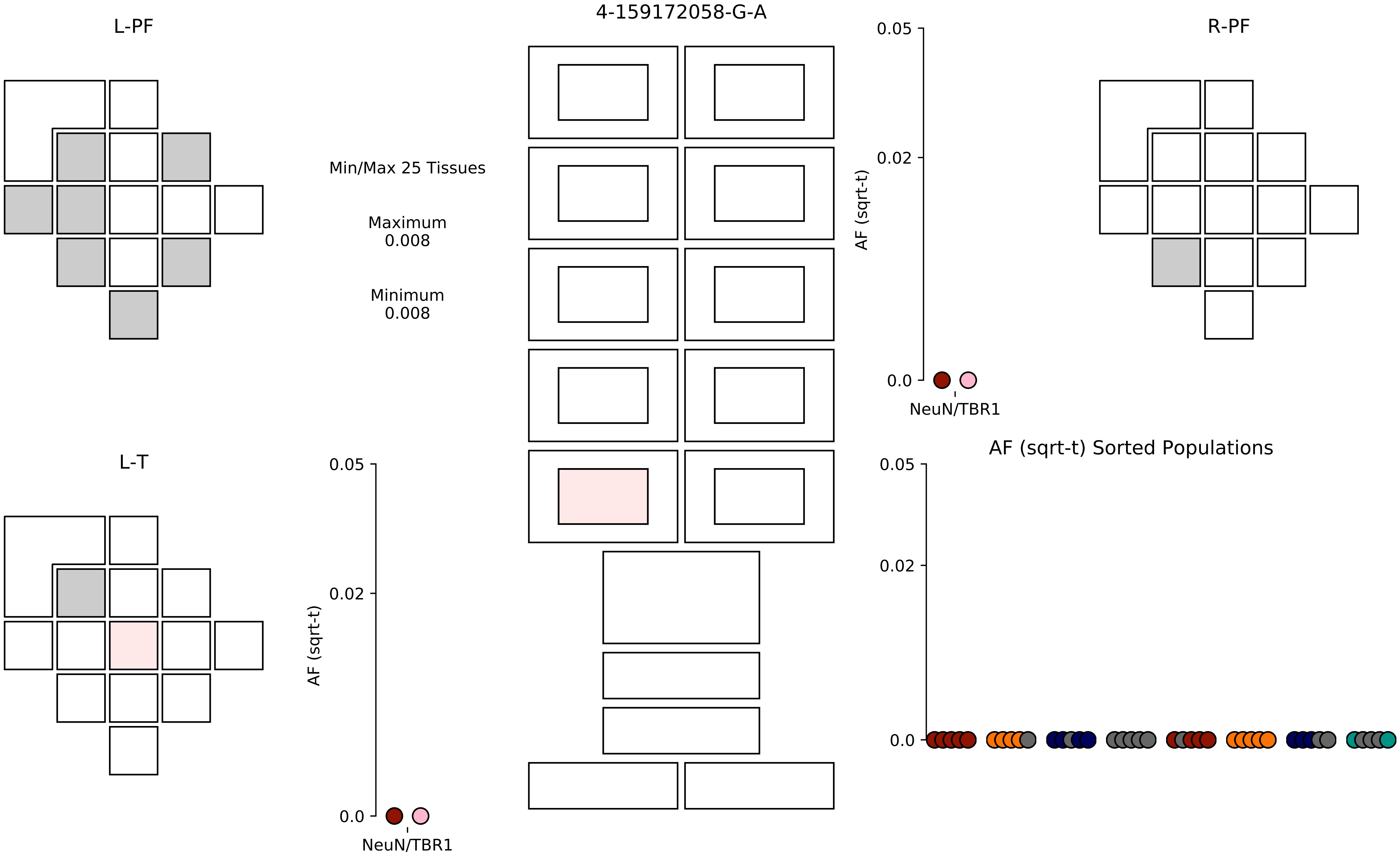

4-159172058-G-A
L-PF
R-PF
L-T
Min/Max 25 Tissues
Maximum
0.008
Minimum
0.008
AF (sqrt-t)
0.05
0.02
0.0
NeuN/TBR1
AF (sqrt-t) Sorted Populations

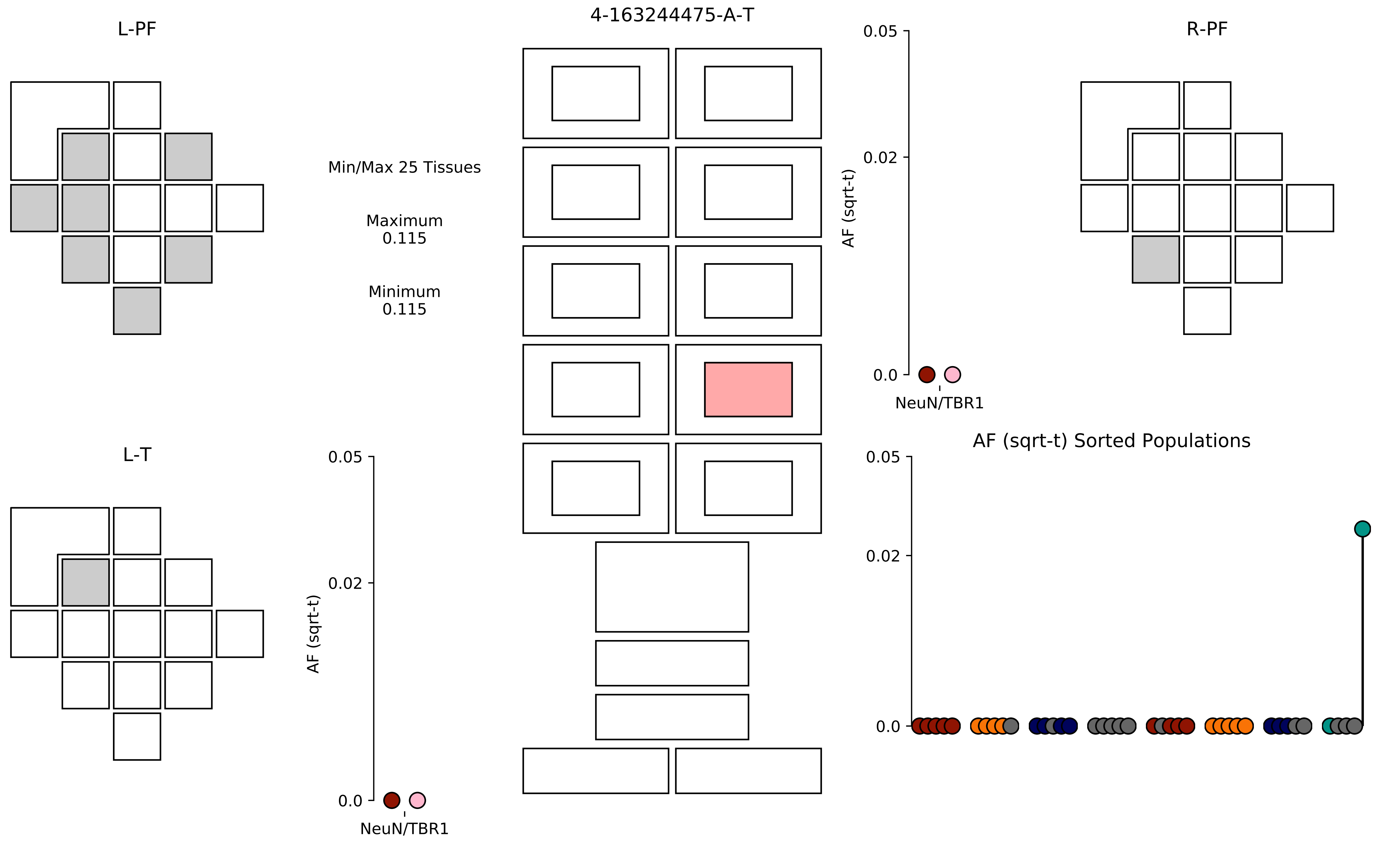

4-163244475-A-T
L-PF
R-PF
L-T
Min/Max 25 Tissues
Maximum
0.115
Minimum
0.115
AF (sqrt-t)
0.05
0.02
0.0
NeuN/TBR1
AF (sqrt-t) Sorted Populations

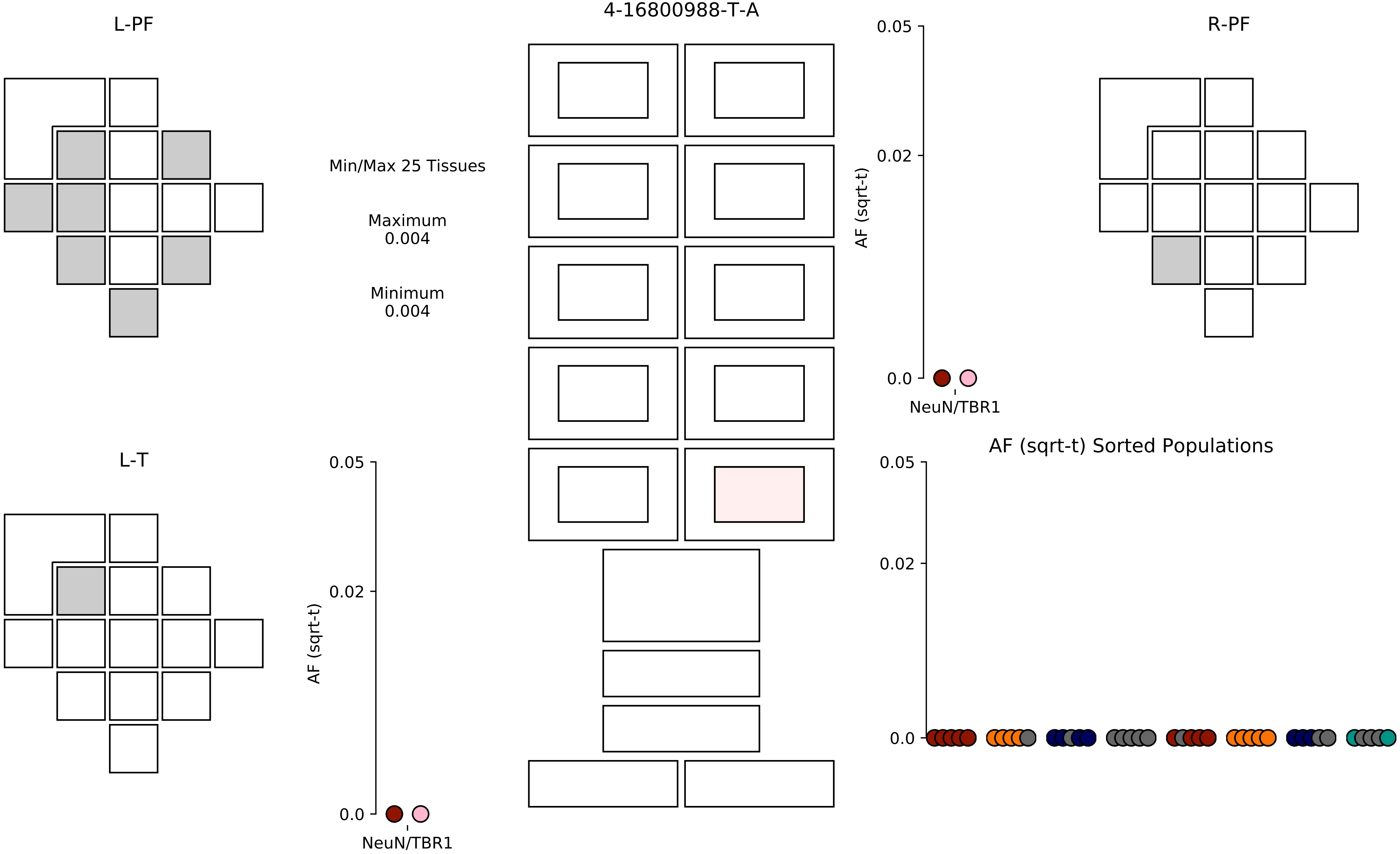
4-16800988-T-A
L-PF
R-PF
L-T
Min/Max 25 Tissues
Maximum
0.004
Minimum
0.004
AF (sqrt-t)
0.05
0.02
0.0
NeuN/TBR1
AF (sqrt-t)
0.05
0.02
0.0
NeuN/TBR1
AF (sqrt-t) Sorted Populations
0.05
0.02
0.0

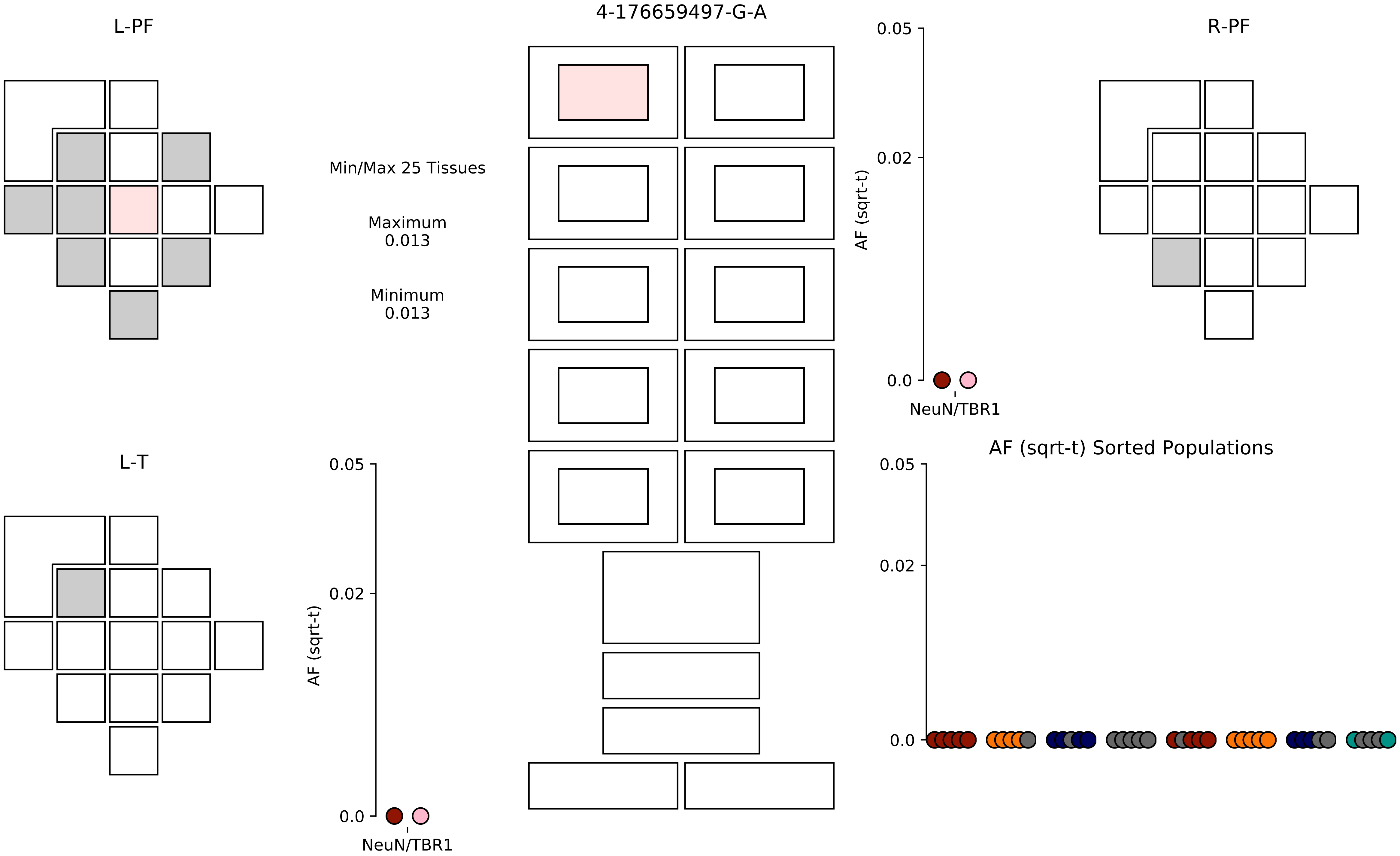

4-176659497-G-A
L-PF
R-PF
L-T
Min/Max 25 Tissues
Maximum
0.013
Minimum
0.013
AF (sqrt-t)
0.05
0.02
0.0
NeuN/TBR1
AF (sqrt-t)
0.05
0.02
0.0
NeuN/TBR1
AF (sqrt-t) Sorted Populations
0.05
0.02
0.0

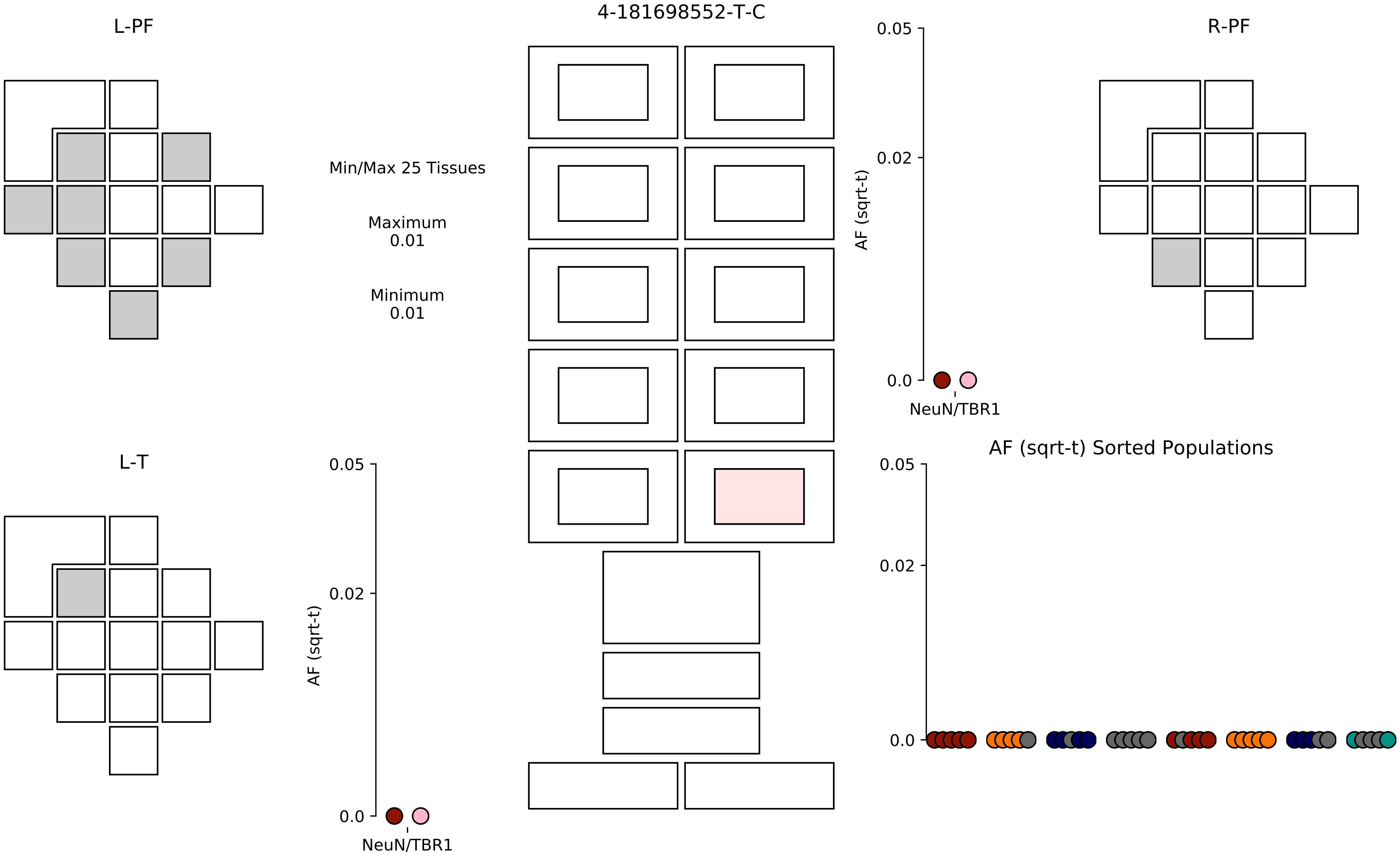

4-181698552-T-C
L-PF
R-PF
L-T
Min/Max 25 Tissues
Maximum
0.01
Minimum
0.01
AF (sqrt-t)
0.05
0.02
0.0
NeuN/TBR1
AF (sqrt-t) Sorted Populations

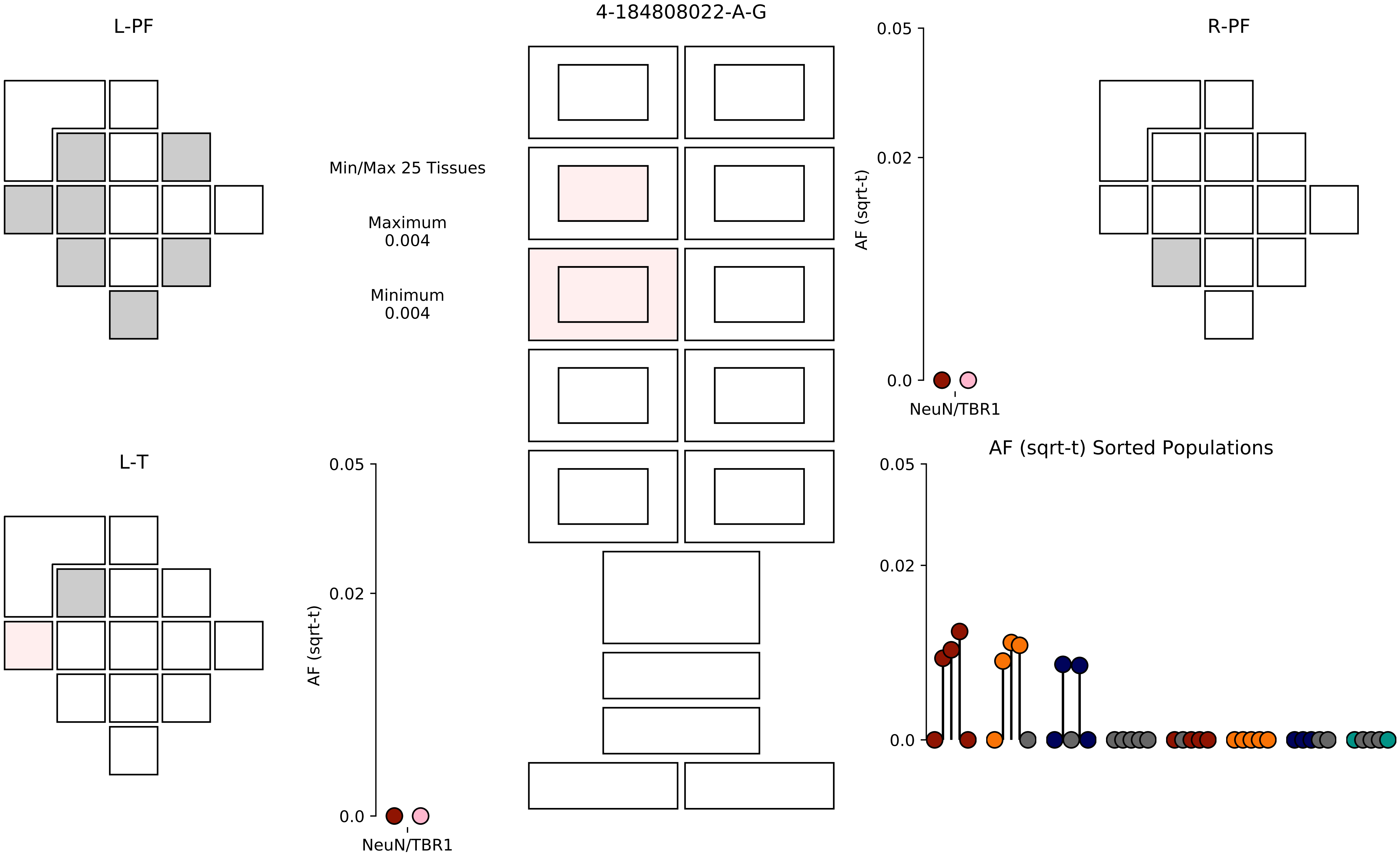

L-PF
4-184808022-A-G
R-PF
Min/Max 25 Tissues
Maximum
0.004
Minimum
0.004
AF (sqrt-t)
0.05
0.02
0.0
NeuN/TBR1
L-T
AF (sqrt-t) Sorted Populations

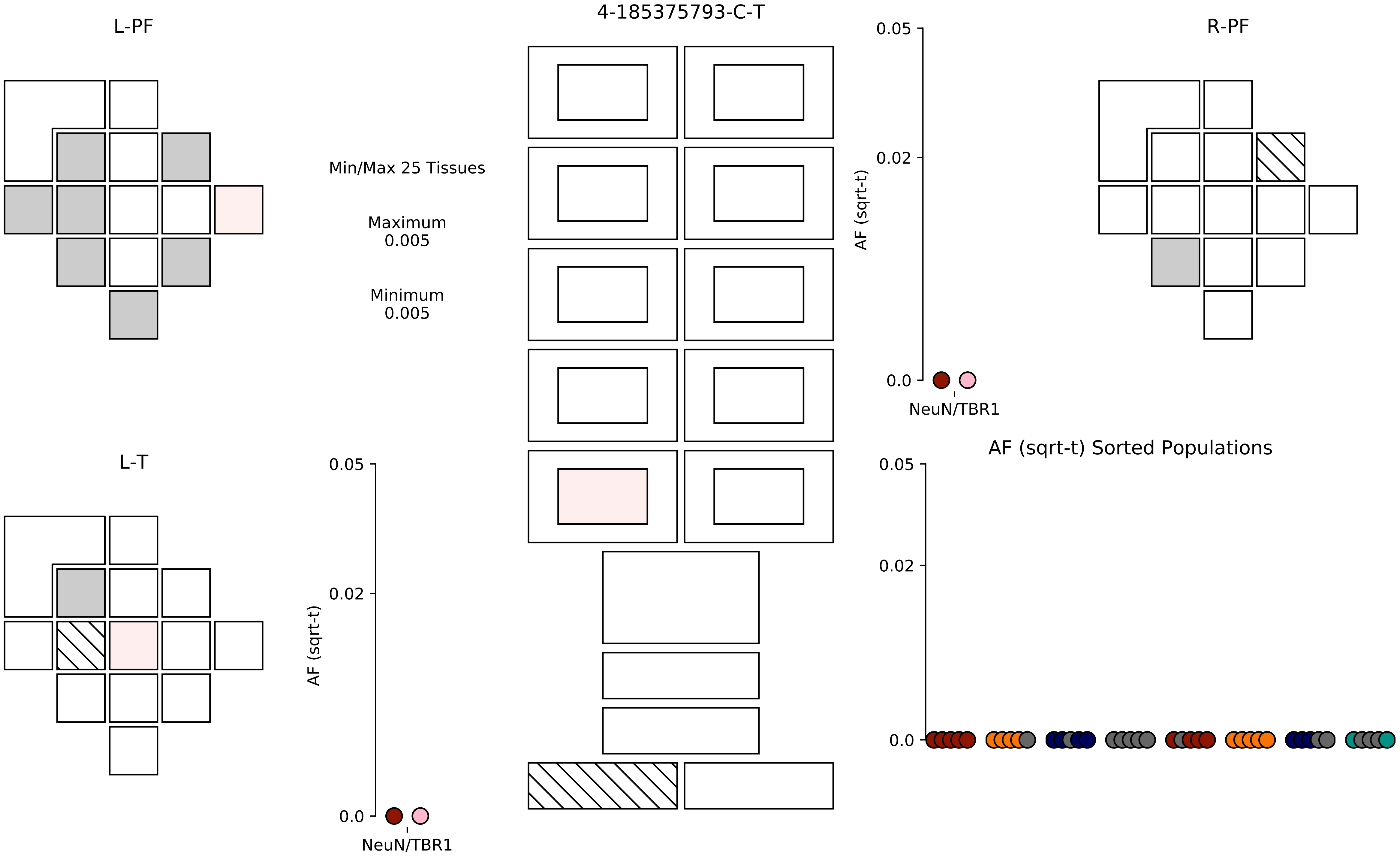

4-185375793-C-T
L-PF
R-PF
L-T
Min/Max 25 Tissues
Maximum
0.005
Minimum
0.005
AF (sqrt-t)
0.05
0.02
0.0
NeuN/TBR1
AF (sqrt-t) Sorted Populations

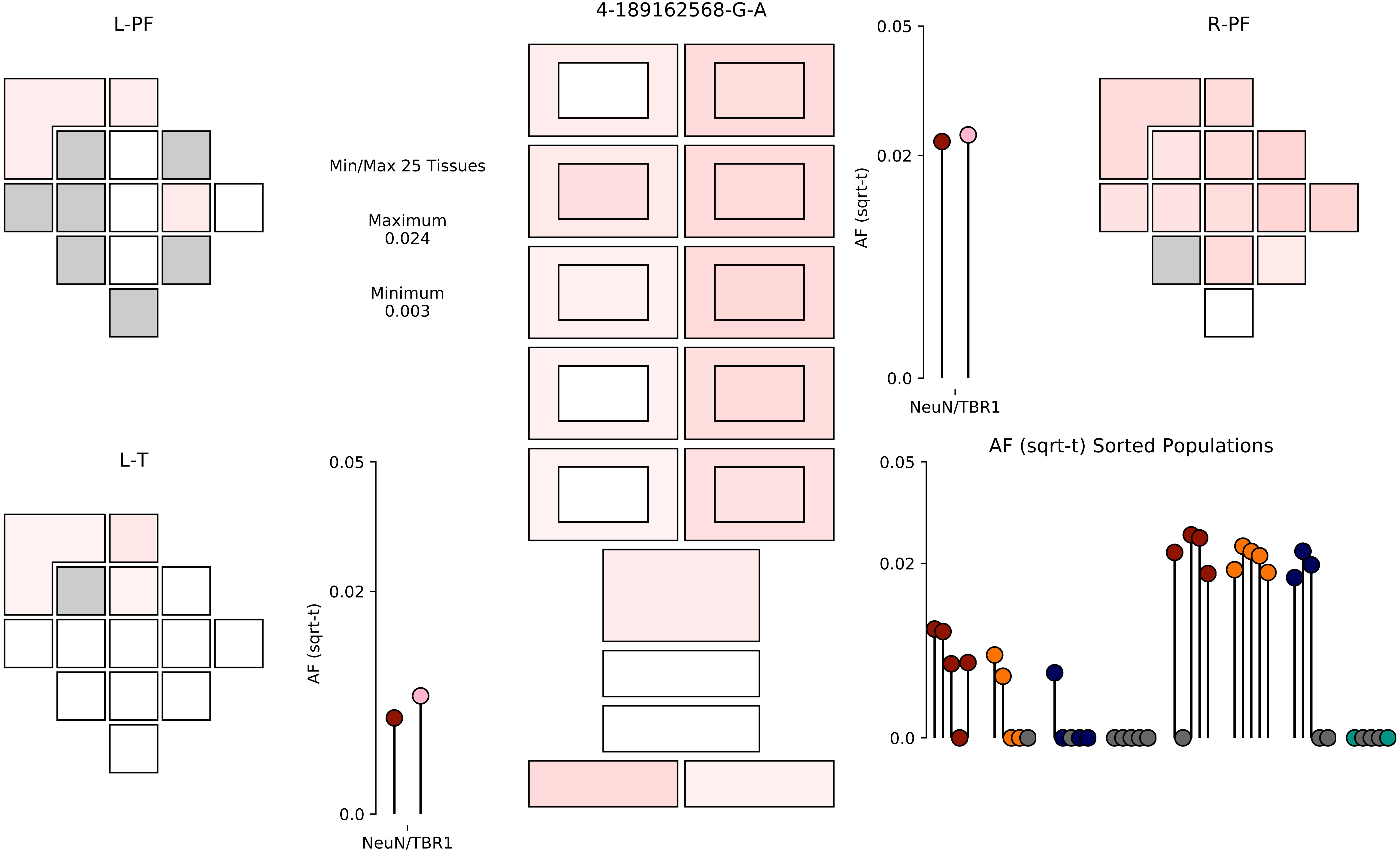
4-189162568-G-A
L-PF
R-PF
L-T
Min/Max 25 Tissues
Maximum
0.024
Minimum
0.003
AF (sqrt-t)
0.05
0.02
0.0
NeuN/TBR1
AF (sqrt-t) Sorted Populations

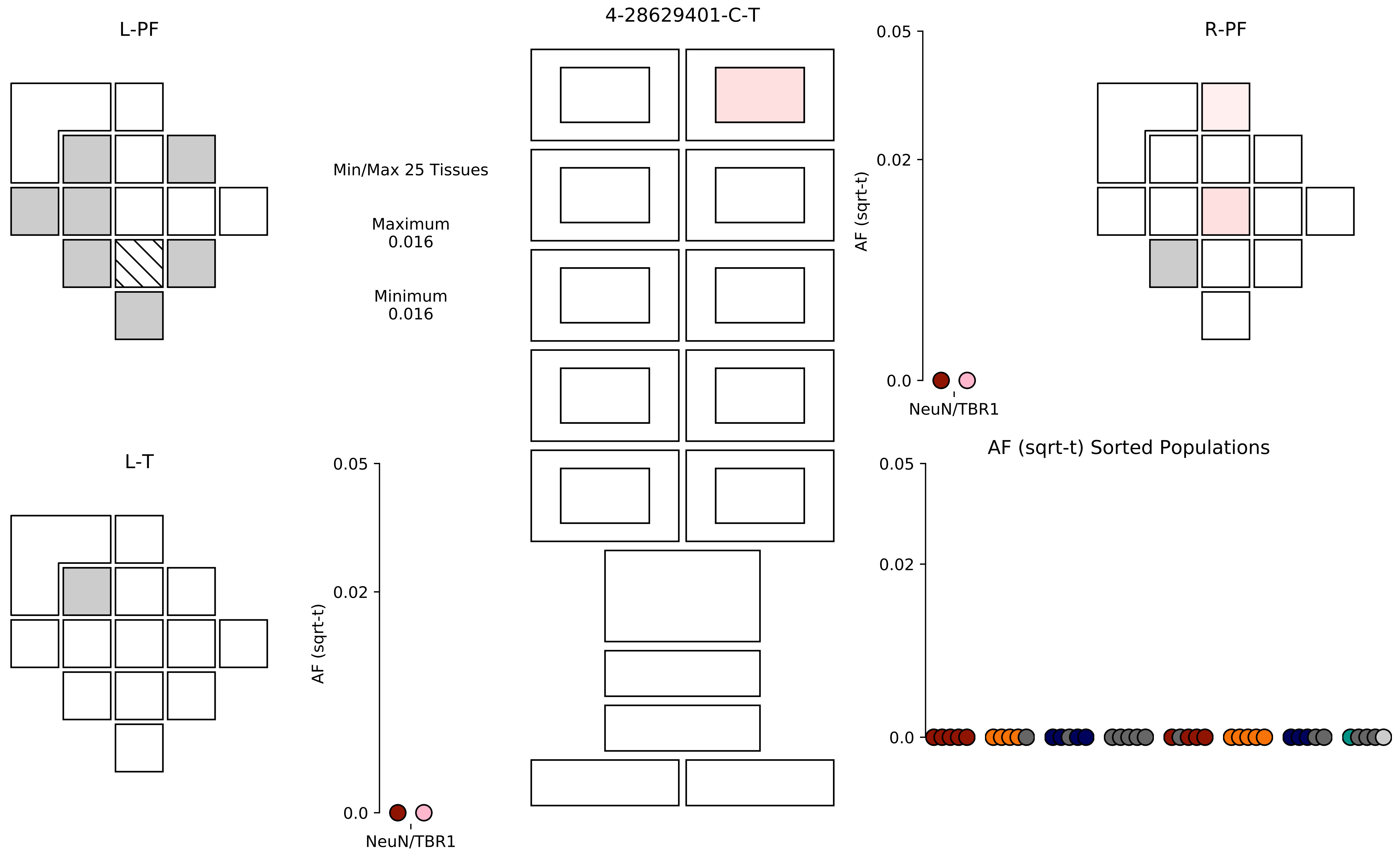

L-PF
4-28629401-C-T
R-PF
Min/Max 25 Tissues
Maximum
0.016
Minimum
0.016
0.05
0.02
0.0
AF (sqrt-t)
NeuN/TBR1
L-T
AF (sqrt-t) Sorted Populations

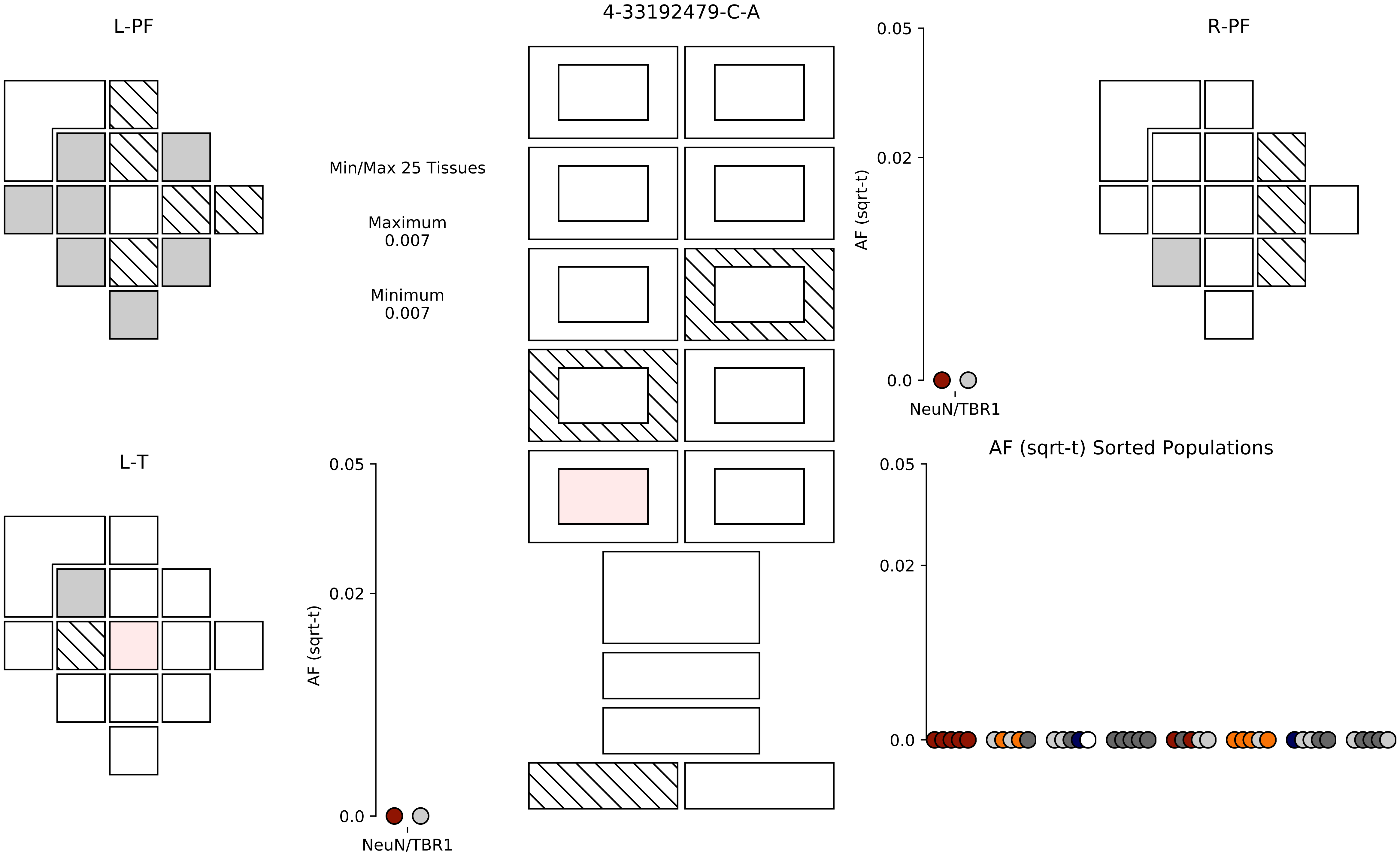
L-PF
4-33192479-C-A
R-PF
Min/Max 25 Tissues
Maximum
0.007
Minimum
0.007
0.05
0.02
0.0
AF (sqrt-t)
NeuN/TBR1
L-T
0.05
0.02
0.0
AF (sqrt-t)
NeuN/TBR1
AF (sqrt-t) Sorted Populations
0.05
0.02
0.0

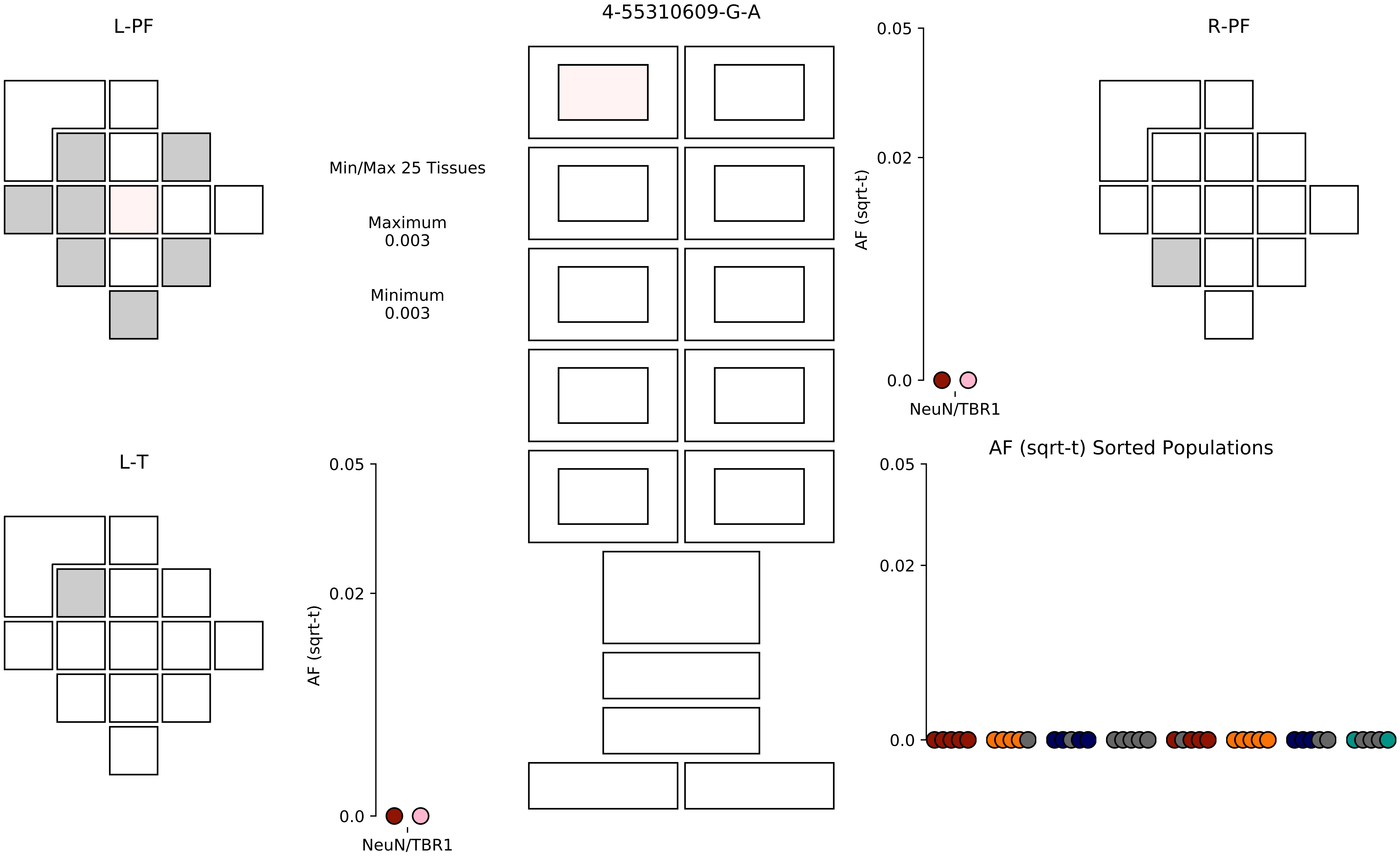
4-55310609-G-A
L-PF
R-PF
L-T
Min/Max 25 Tissues
Maximum
0.003
Minimum
0.003
AF (sqrt-t)
0.05
0.02
0.0
NeuN/TBR1
AF (sqrt-t) Sorted Populations

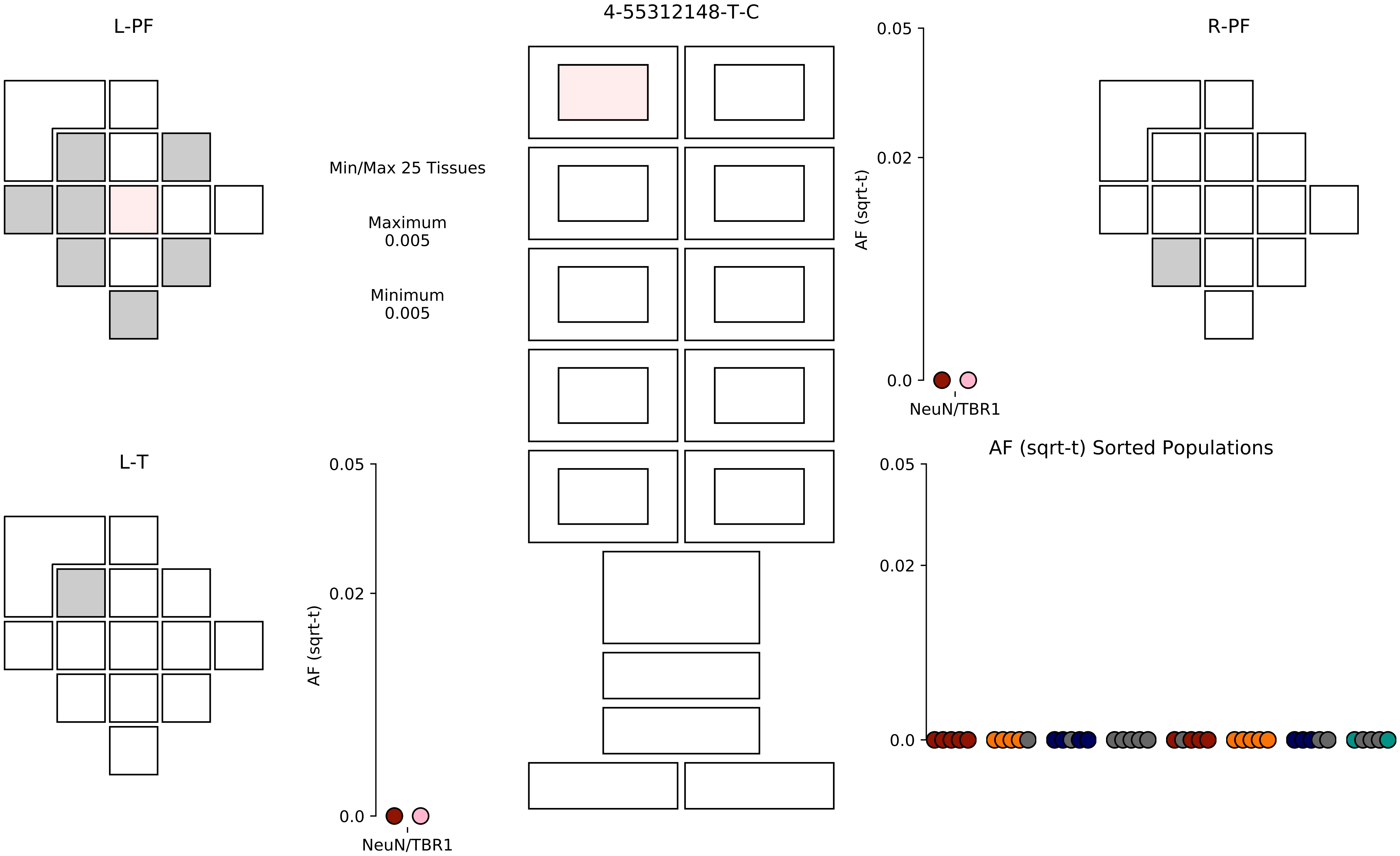
4-55312148-T-C
L-PF
R-PF
L-T
Min/Max 25 Tissues
Maximum
0.005
Minimum
0.005
AF (sqrt-t)
0.05
0.02
0.0
NeuN/TBR1
AF (sqrt-t) Sorted Populations

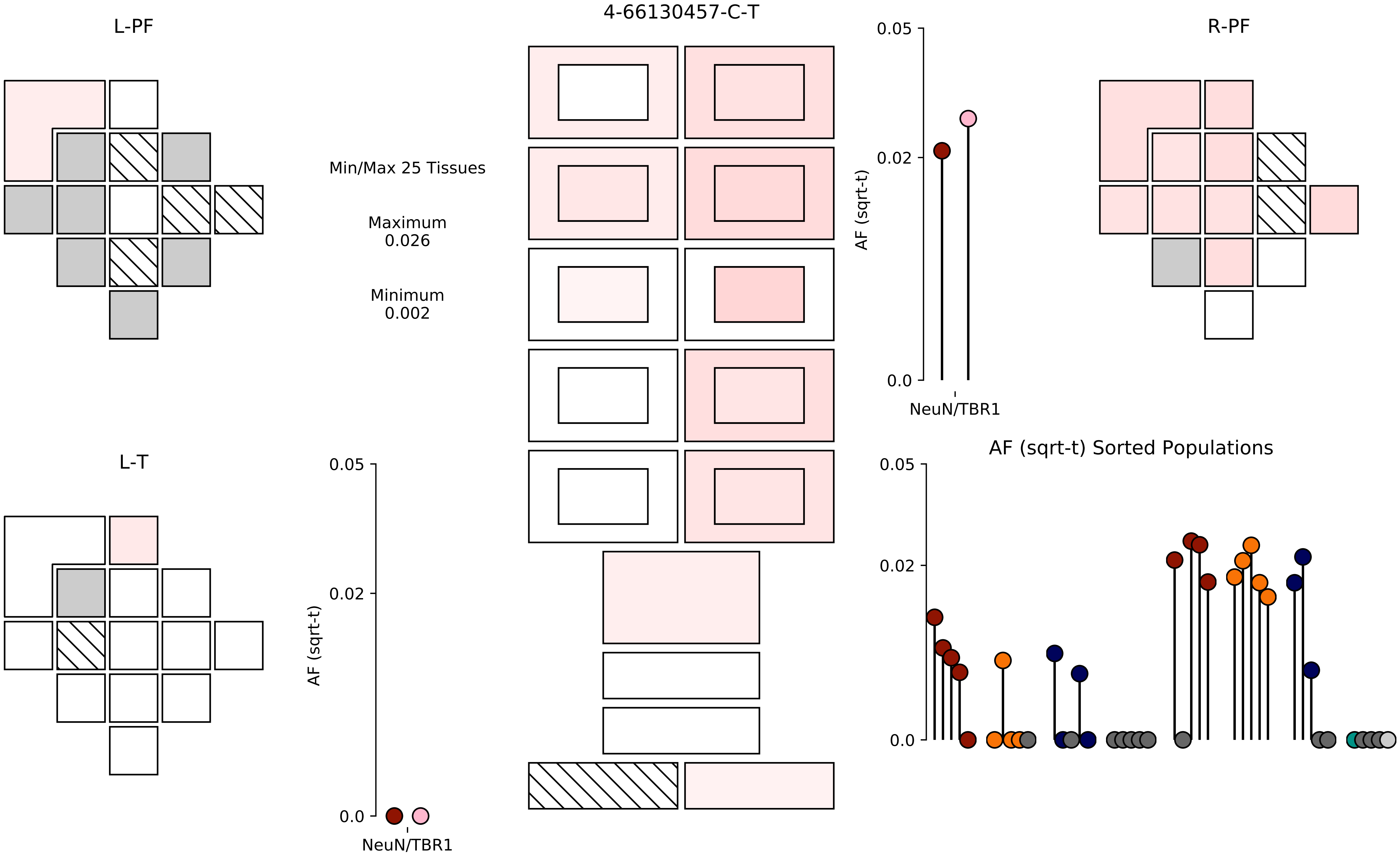

L-PF
4-66130457-C-T
R-PF
Min/Max 25 Tissues
Maximum
0.026
Minimum
0.002
0.05
0.02
0.0
AF (sqrt-t)
NeuN/TBR1
L-T
AF (sqrt-t) Sorted Populations

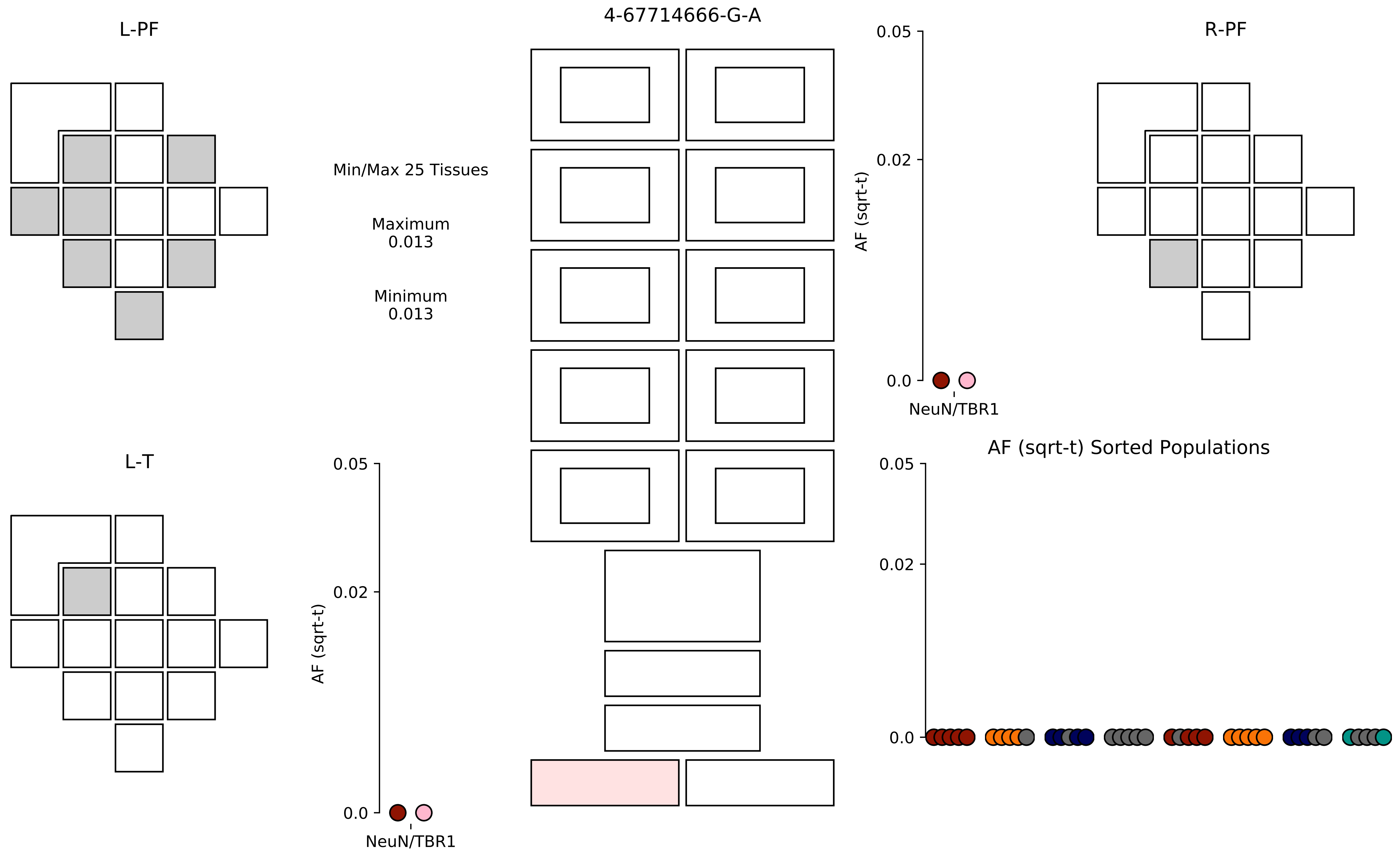
4-67714666-G-A
L-PF
R-PF
L-T
Min/Max 25 Tissues
Maximum
0.013
Minimum
0.013
AF (sqrt-t)
0.05
0.02
0.0
NeuN/TBR1
AF (sqrt-t) Sorted Populations

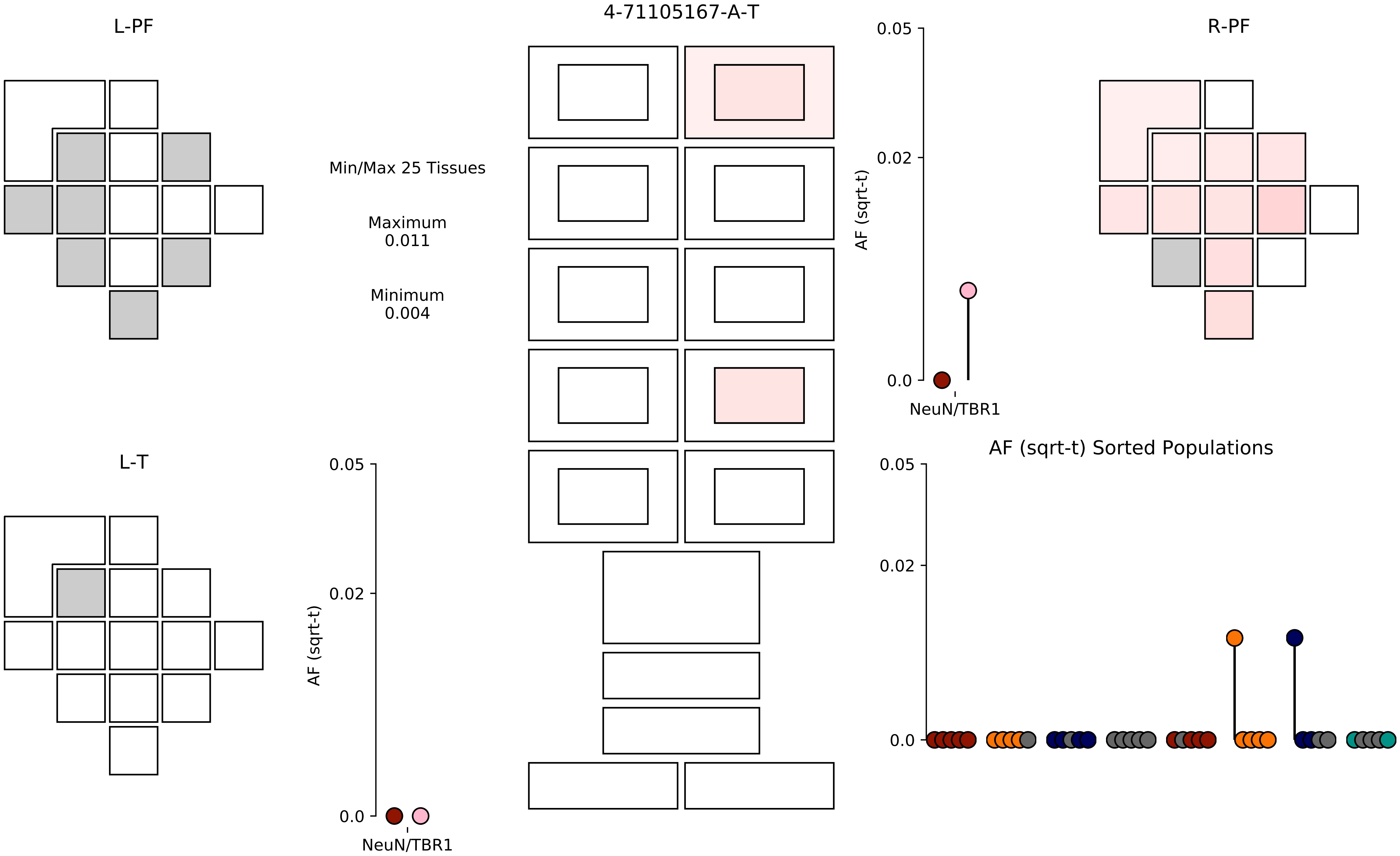

4-71105167-A-T
L-PF
R-PF
L-T
Min/Max 25 Tissues
Maximum
0.011
Minimum
0.004
0.05
0.02
0.0
AF (sqrt-t)
NeuN/TBR1
AF (sqrt-t) Sorted Populations

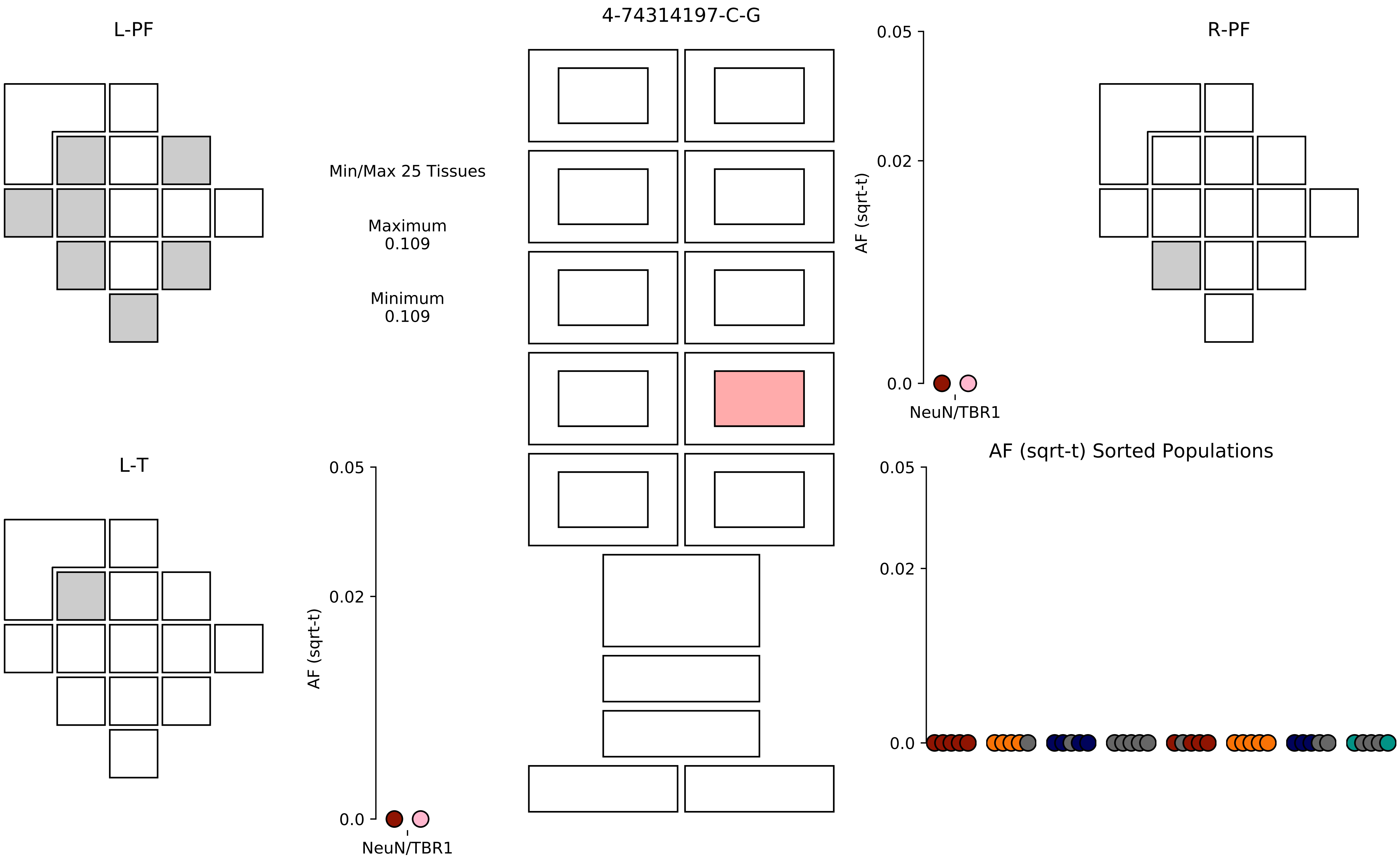
4-74314197-C-G
L-PF
R-PF
L-T
Min/Max 25 Tissues
Maximum
0.109
Minimum
0.109
AF (sqrt-t)
0.05
0.02
0.0
NeuN/TBR1
AF (sqrt-t) Sorted Populations

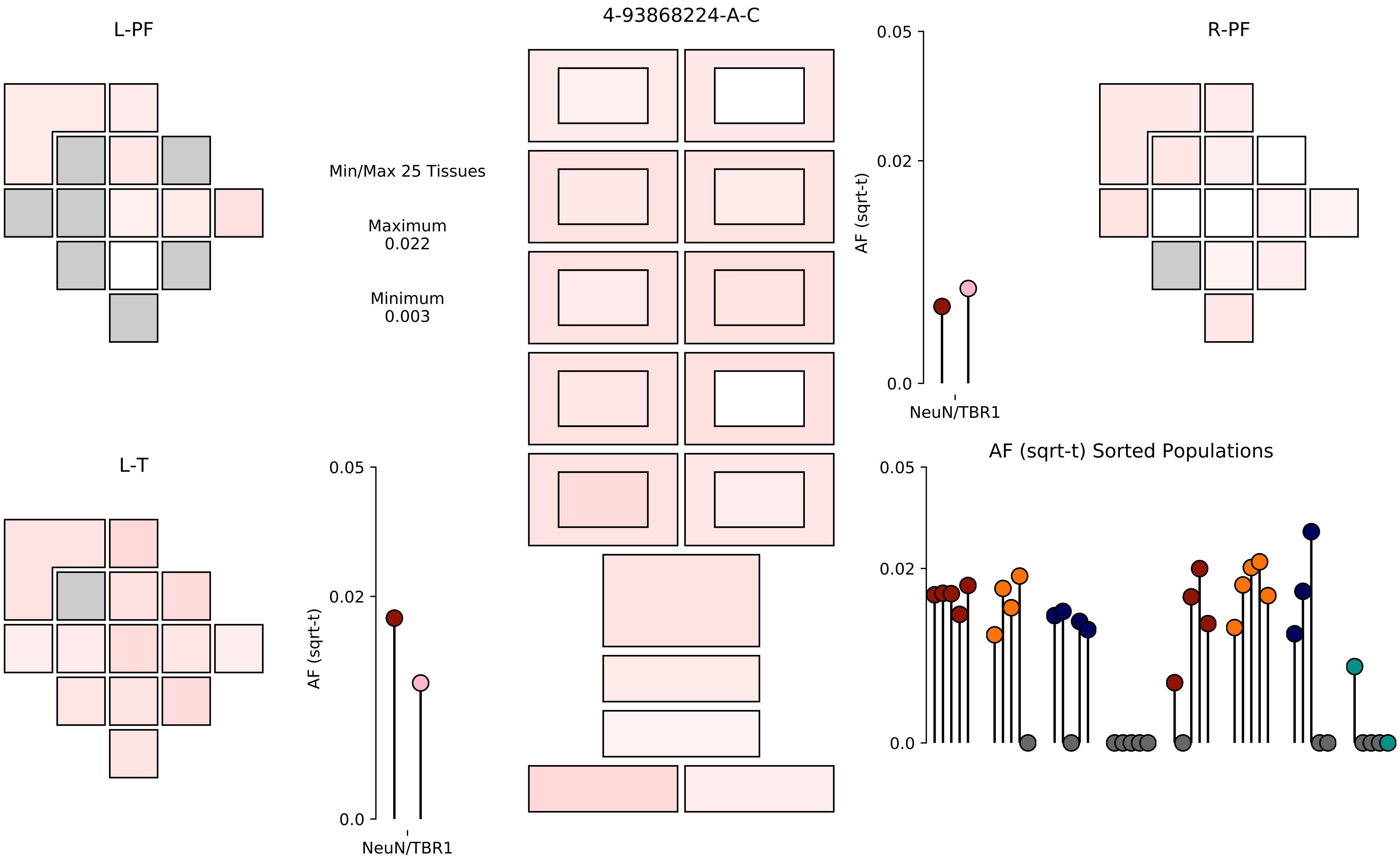
L-PF
4-93868224-A-C
R-PF
Min/Max 25 Tissues
Maximum
0.022
Minimum
0.003
0.05
0.02
0.0
AF (sqrt-t)
NeuN/TBR1
L-T
0.05
0.02
0.0
AF (sqrt-t)
NeuN/TBR1
AF (sqrt-t) Sorted Populations
0.05
0.02
0.0

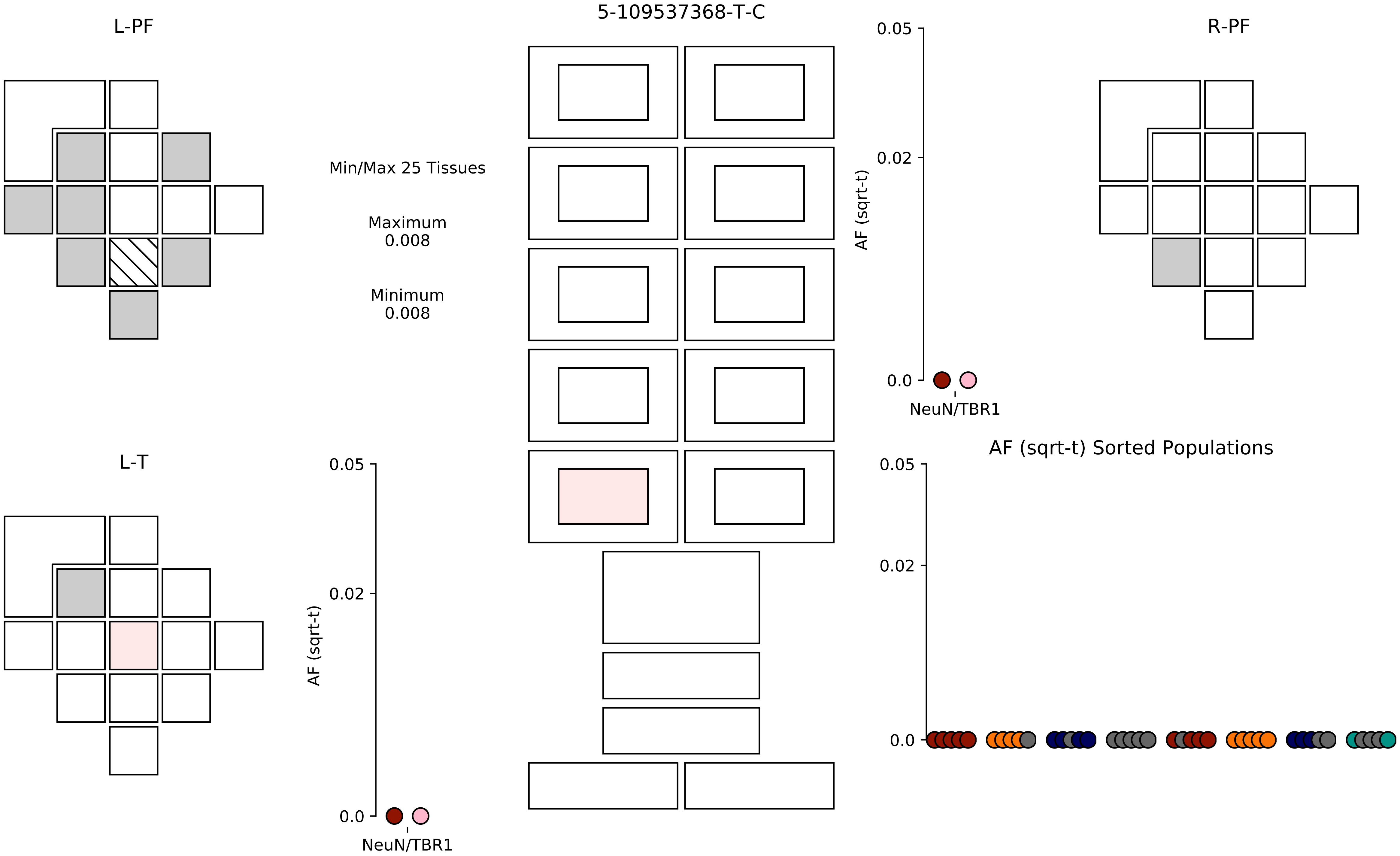

5-109537368-T-C
L-PF
R-PF
L-T
Min/Max 25 Tissues
Maximum
0.008
Minimum
0.008
AF (sqrt-t)
0.05
0.02
0.0
NeuN/TBR1
AF (sqrt-t)
0.05
0.02
0.0
NeuN/TBR1
AF (sqrt-t) Sorted Populations
0.05
0.02
0.0

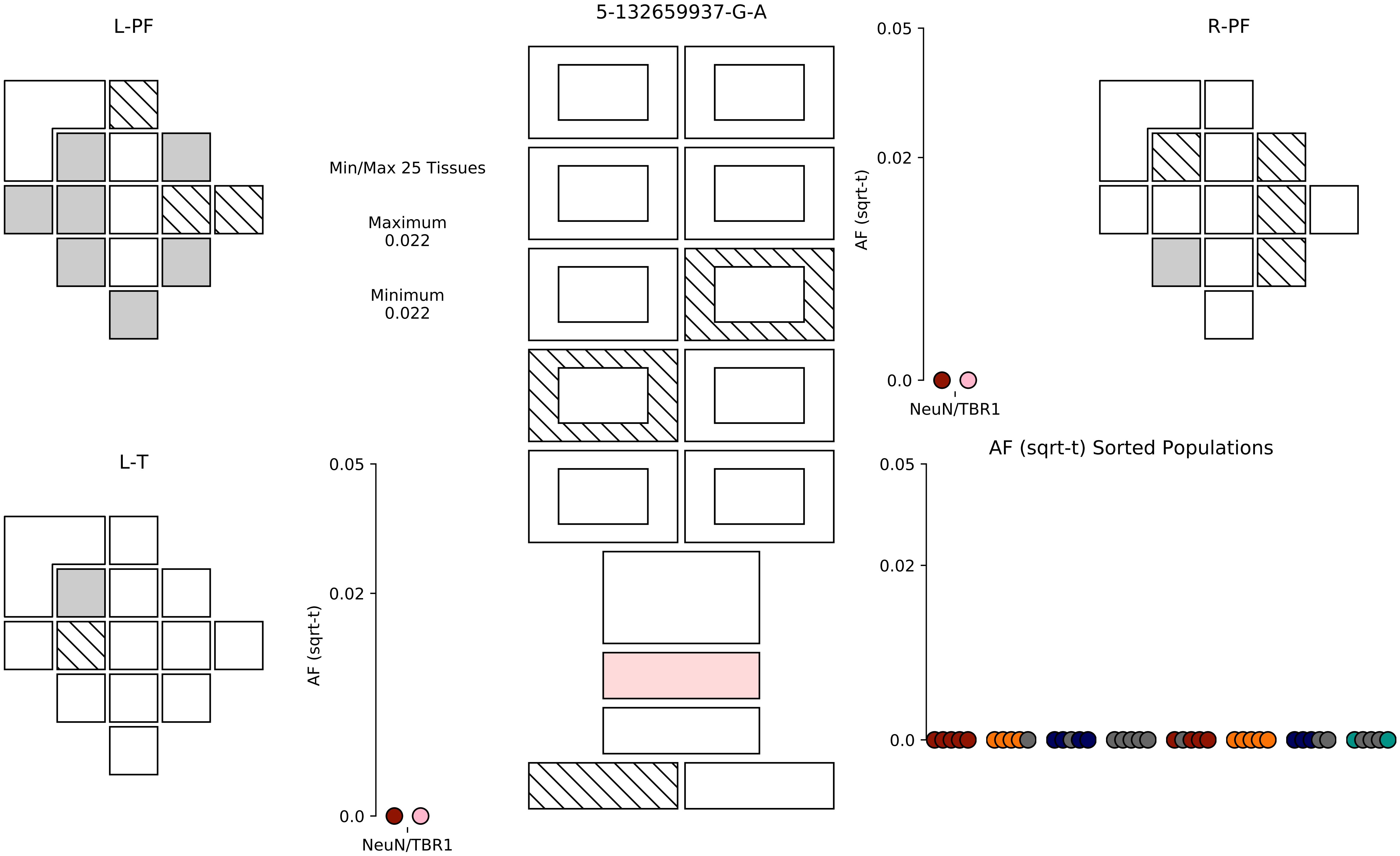

5-132659937-G-A
L-PF
R-PF
L-T
Min/Max 25 Tissues
Maximum
0.022
Minimum
0.022
AF (sqrt-t)
0.05
0.02
0.0
NeuN/TBR1
AF (sqrt-t) Sorted Populations

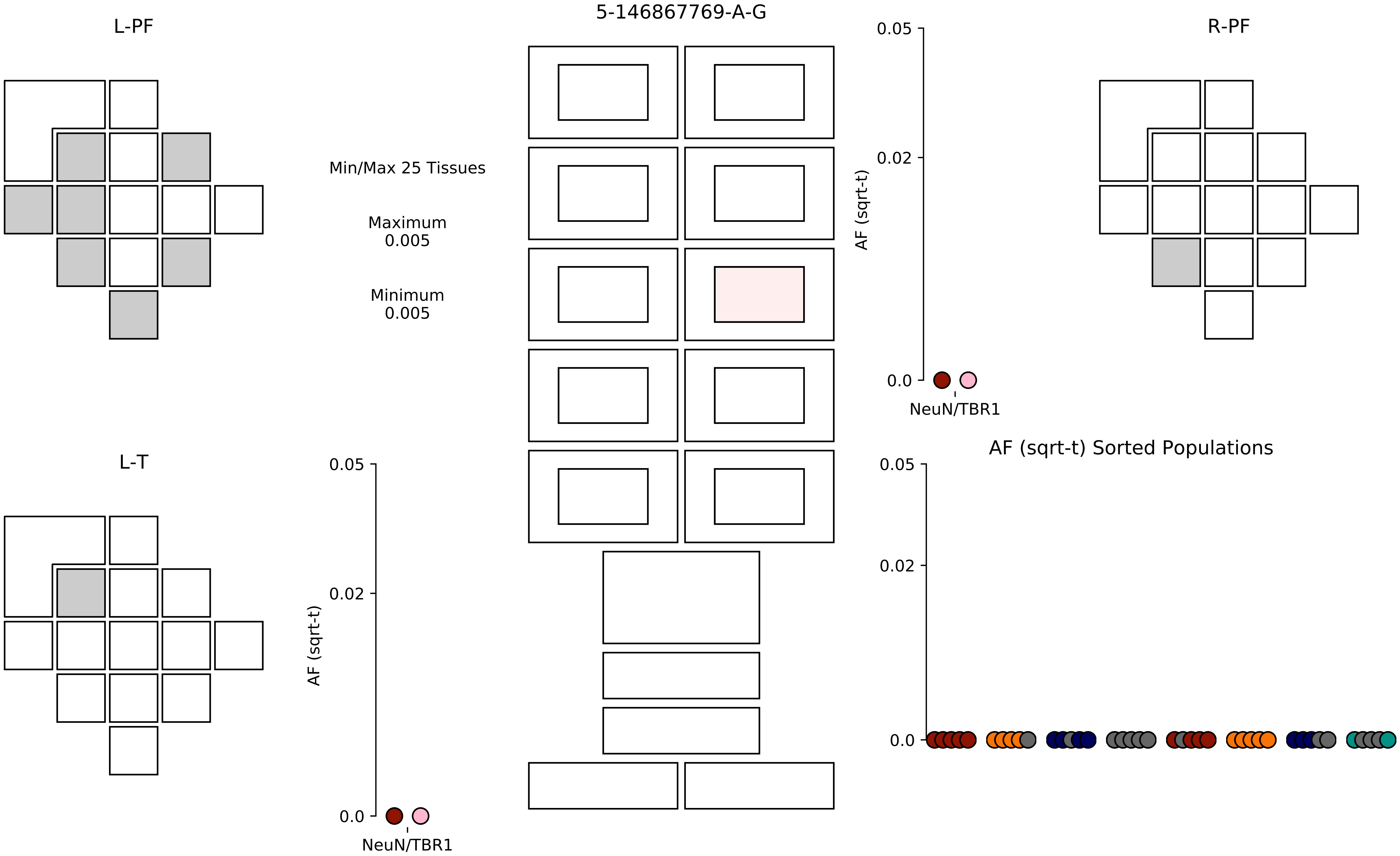

5-146867769-A-G
L-PF
R-PF
L-T
Min/Max 25 Tissues
Maximum
0.005
Minimum
0.005
0.05
0.02
0.0
AF (sqrt-t)
NeuN/TBR1
AF (sqrt-t) Sorted Populations

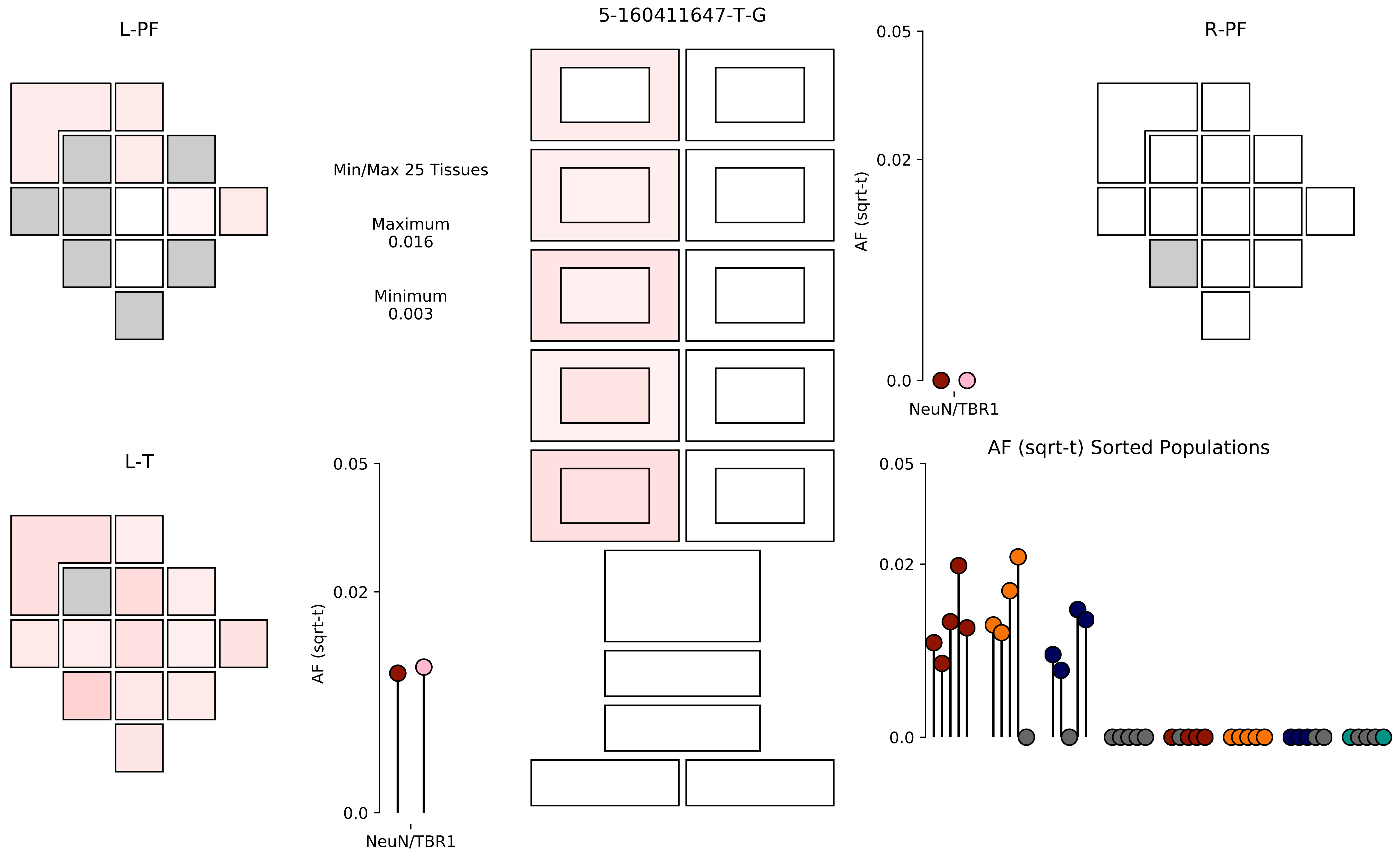

5-160411647-T-G
L-PF
R-PF
L-T
Min/Max 25 Tissues
Maximum
0.016
Minimum
0.003
0.05
0.02
0.0
AF (sqrt-t)
NeuN/TBR1
AF (sqrt-t) Sorted Populations

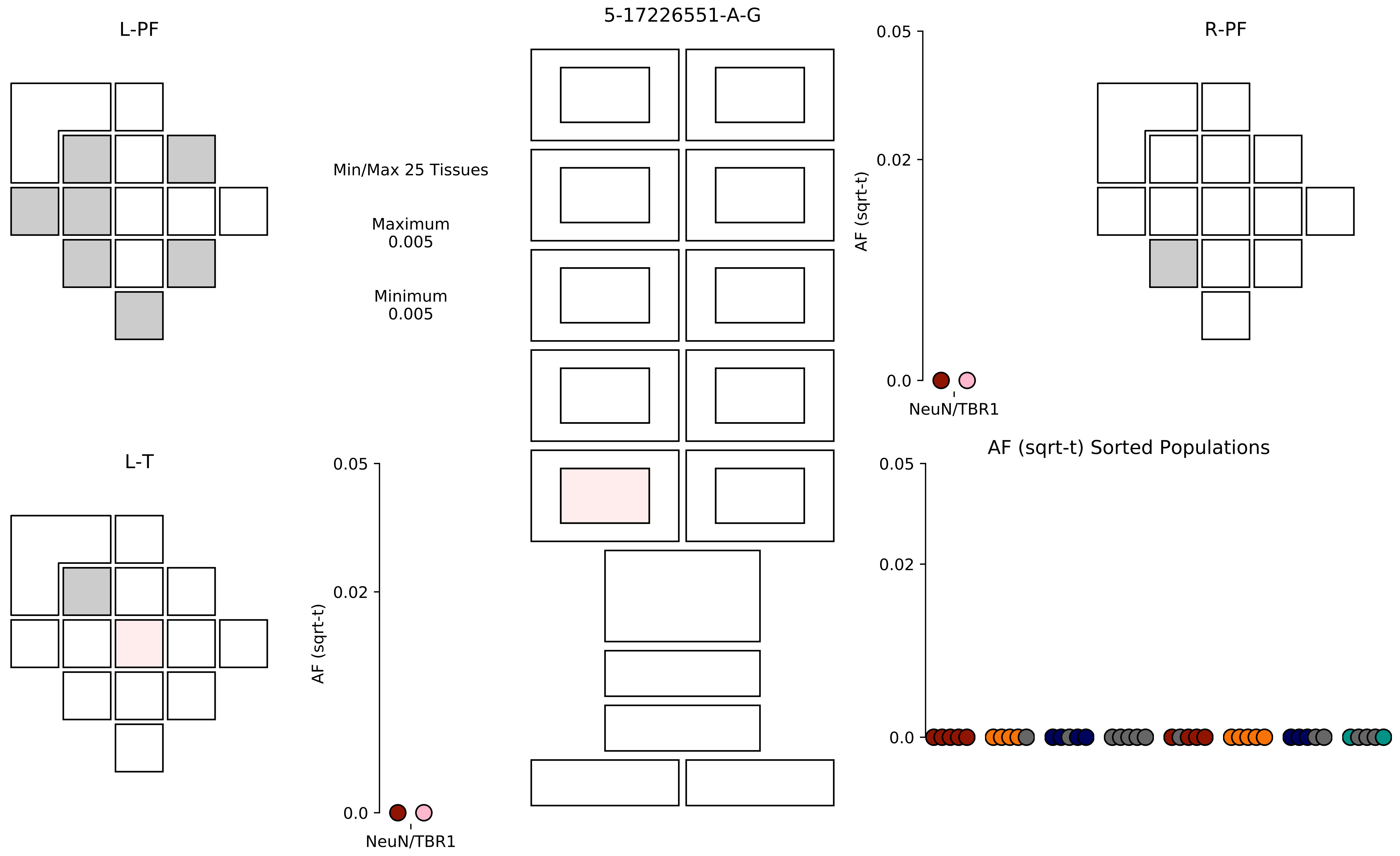

5-17226551-A-G
L-PF
R-PF
L-T
Min/Max 25 Tissues
Maximum
0.005
Minimum
0.005
AF (sqrt-t)
0.05
0.02
0.0
NeuN/TBR1
AF (sqrt-t) Sorted Populations

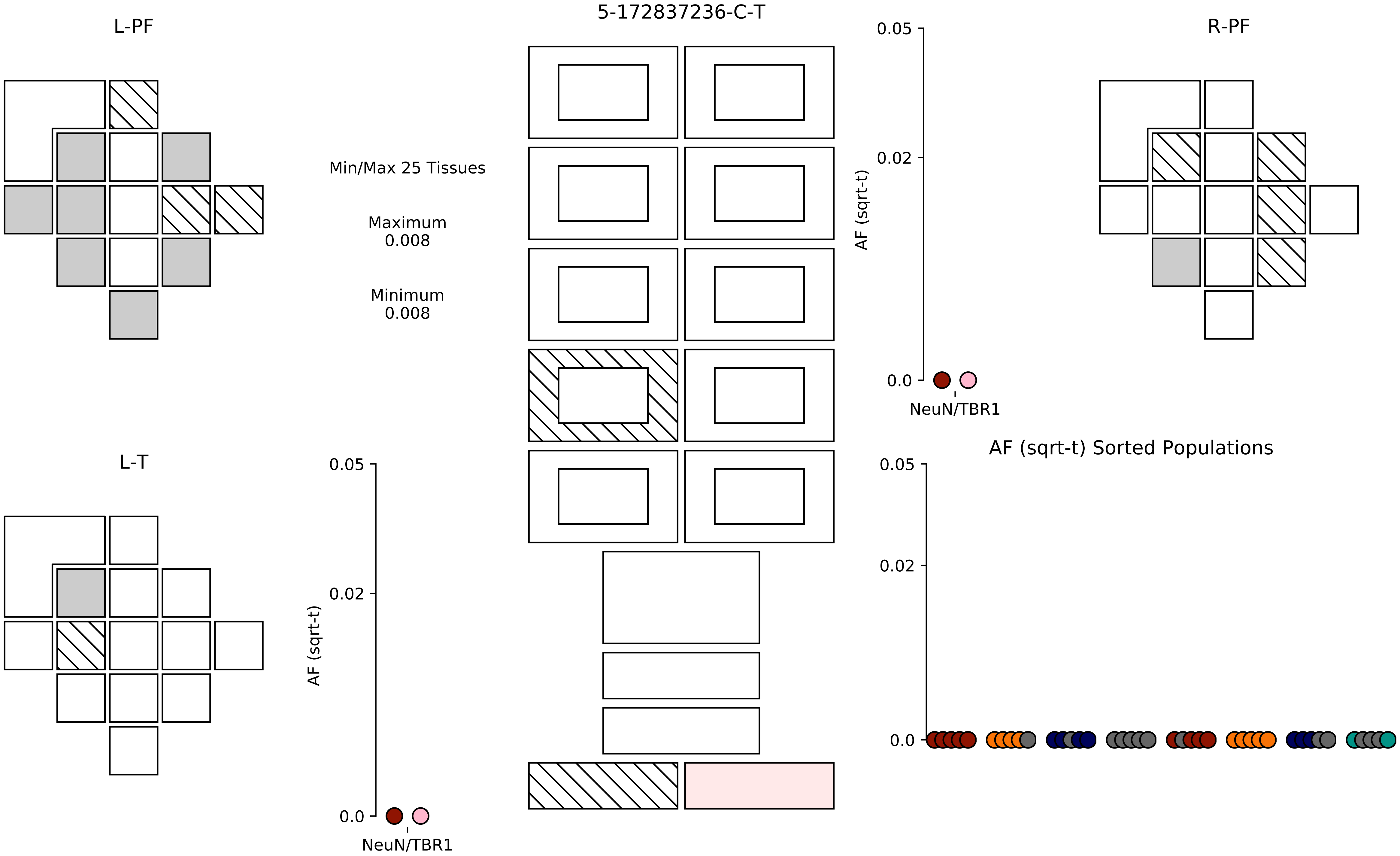

5-172837236-C-T
L-PF
R-PF
L-T
Min/Max 25 Tissues
Maximum
0.008
Minimum
0.008
AF (sqrt-t)
0.05
0.02
0.0
NeuN/TBR1
AF (sqrt-t) Sorted Populations

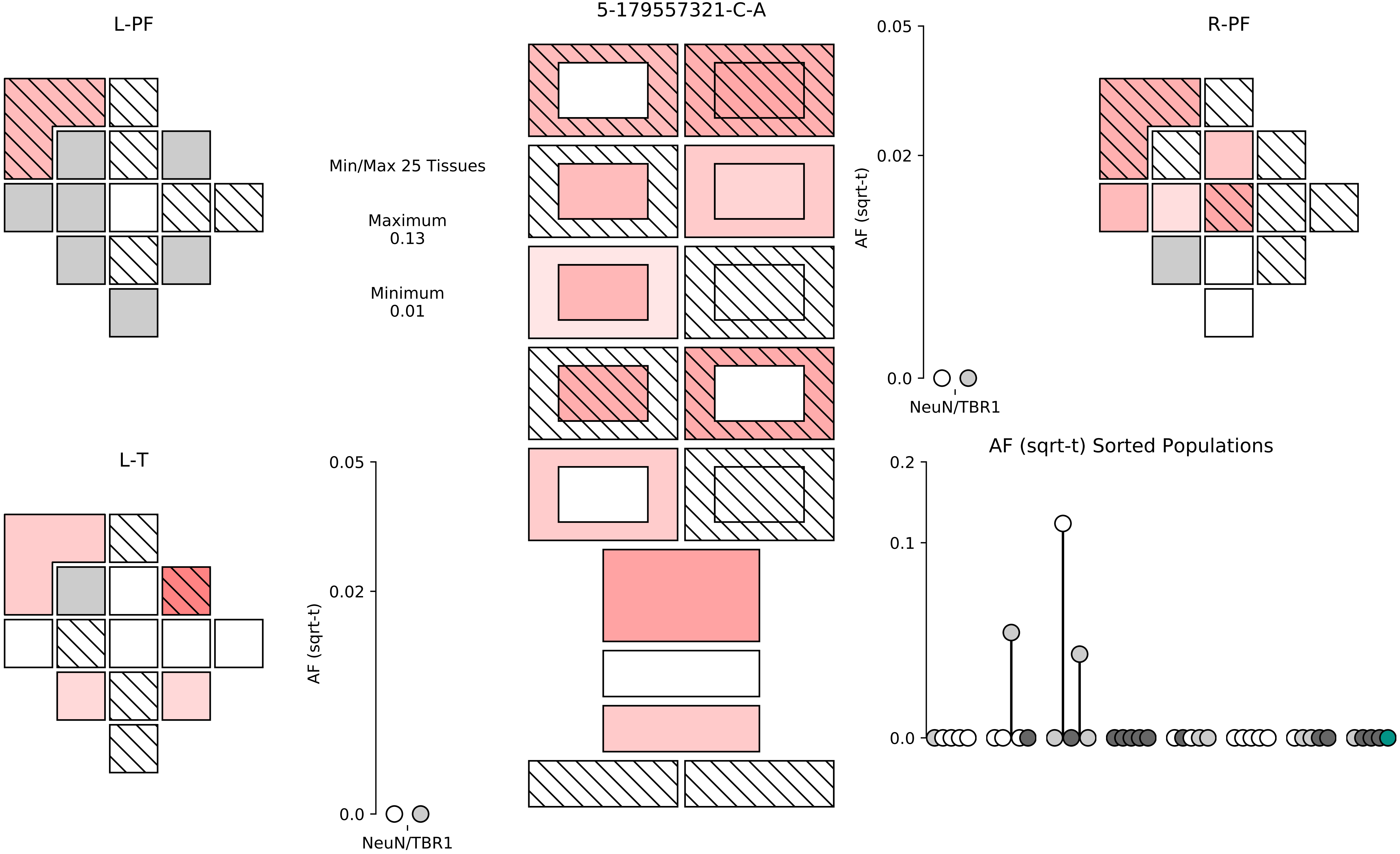
5-179557321-C-A
L-PF
R-PF
L-T
Min/Max 25 Tissues
Maximum
0.13
Minimum
0.01
AF (sqrt-t)
0.05
0.02
0.0
NeuN/TBR1
AF (sqrt-t) Sorted Populations
0.2
0.1
0.0

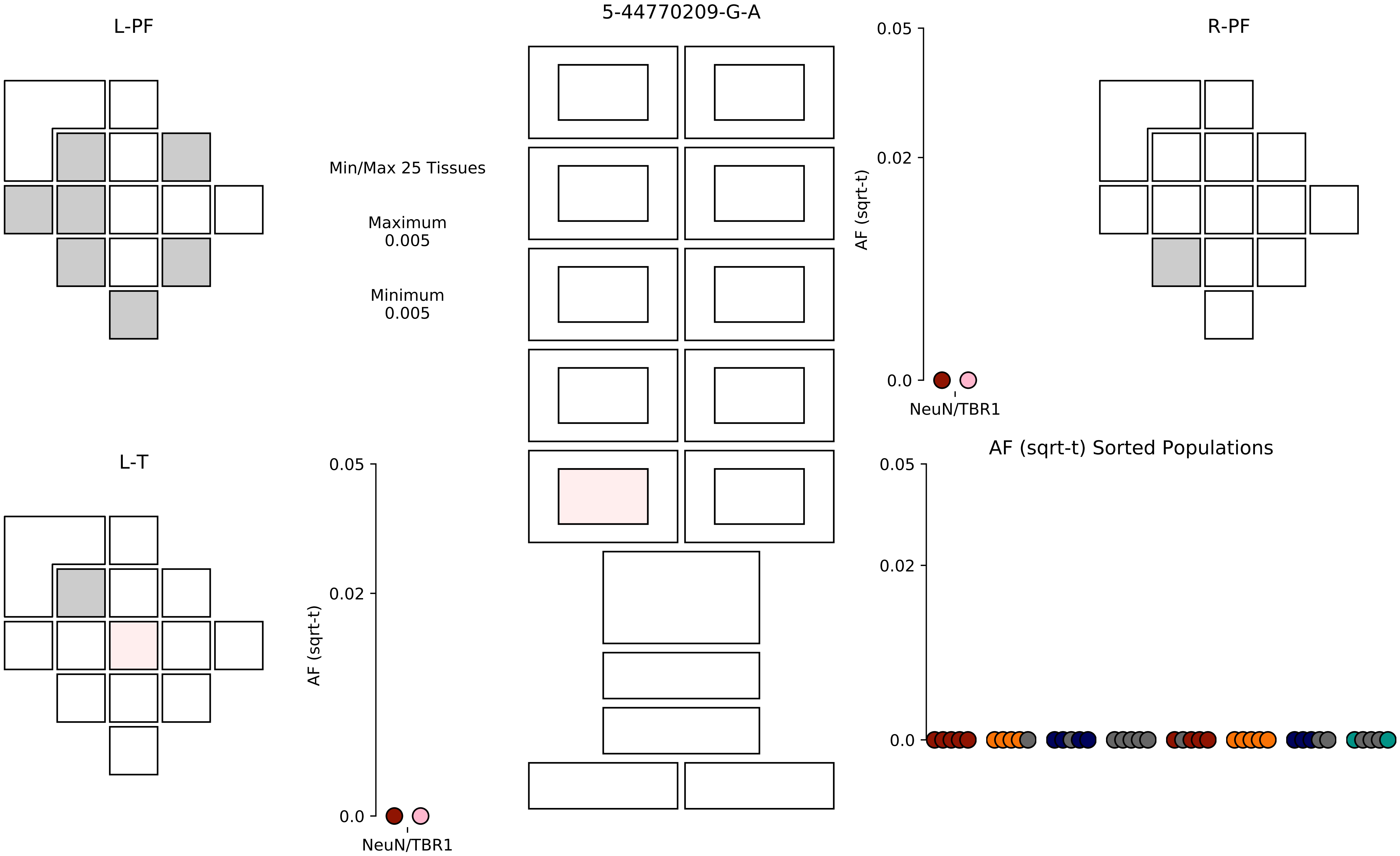

L-PF
5-44770209-G-A
R-PF
0.05
0.02
0.0
AF (sqrt-t)
NeuN/TBR1
Min/Max 25 Tissues
Maximum
0.005
Minimum
0.005
L-T
AF (sqrt-t) Sorted Populations

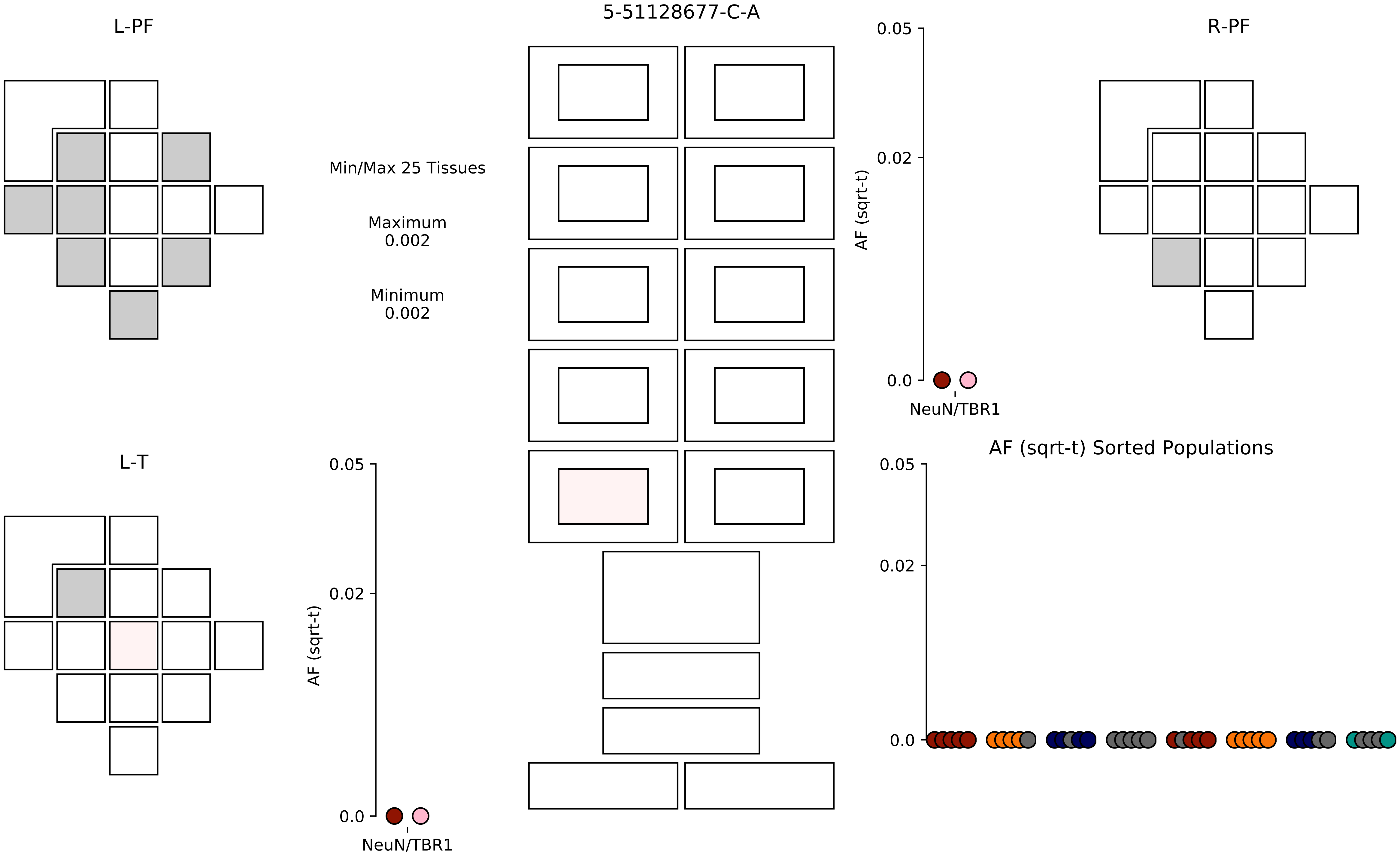
5-51128677-C-A
L-PF
R-PF
L-T
Min/Max 25 Tissues
Maximum
0.002
Minimum
0.002
AF (sqrt-t)
0.05
0.02
0.0
NeuN/TBR1
AF (sqrt-t)
0.05
0.02
0.0
NeuN/TBR1
AF (sqrt-t) Sorted Populations
0.05
0.02
0.0

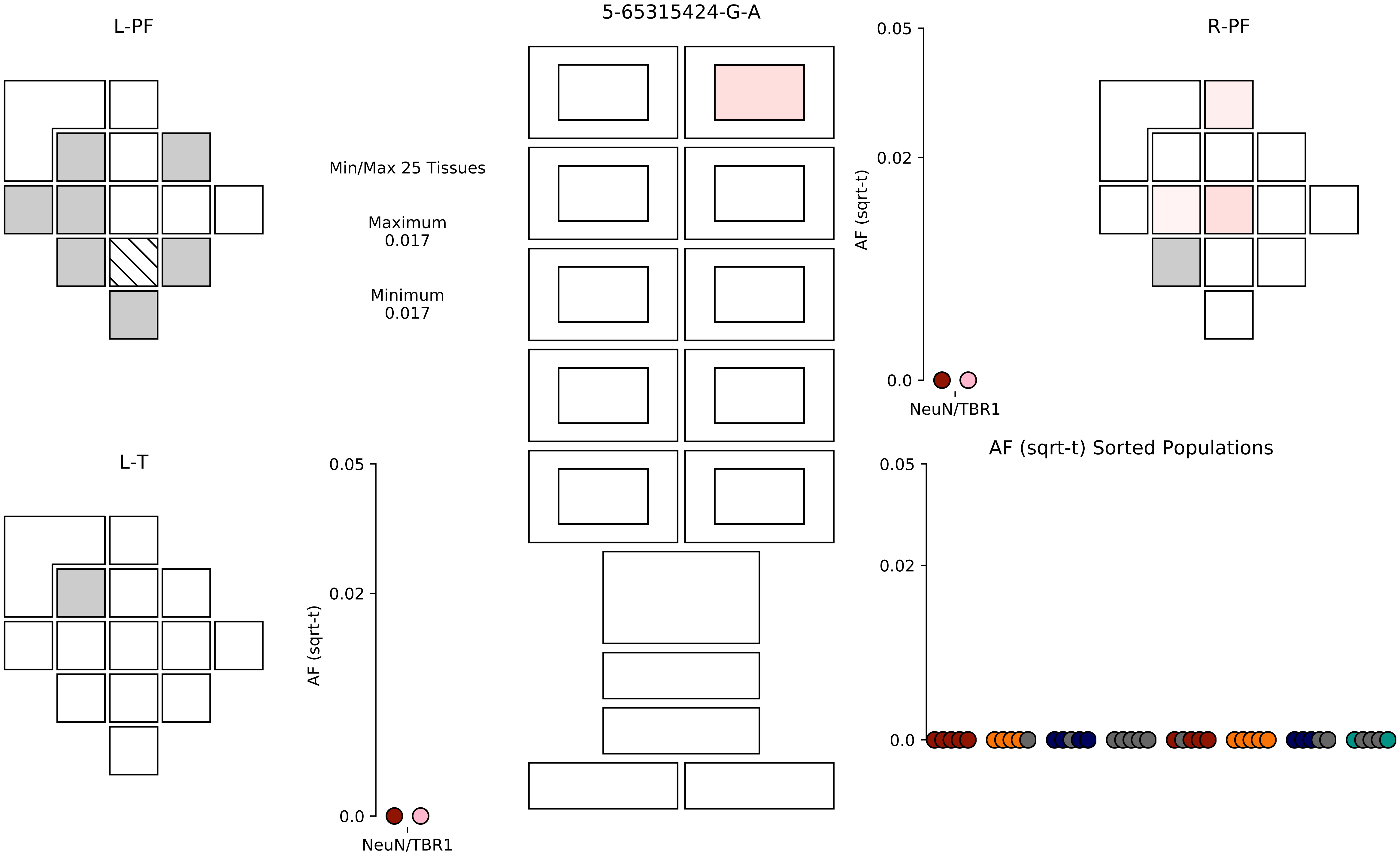
5-65315424-G-A
L-PF
R-PF
L-T
Min/Max 25 Tissues
Maximum
0.017
Minimum
0.017
AF (sqrt-t)
0.05
0.02
0.0
NeuN/TBR1
AF (sqrt-t)
0.05
0.02
0.0
NeuN/TBR1
AF (sqrt-t) Sorted Populations
0.05
0.02
0.0

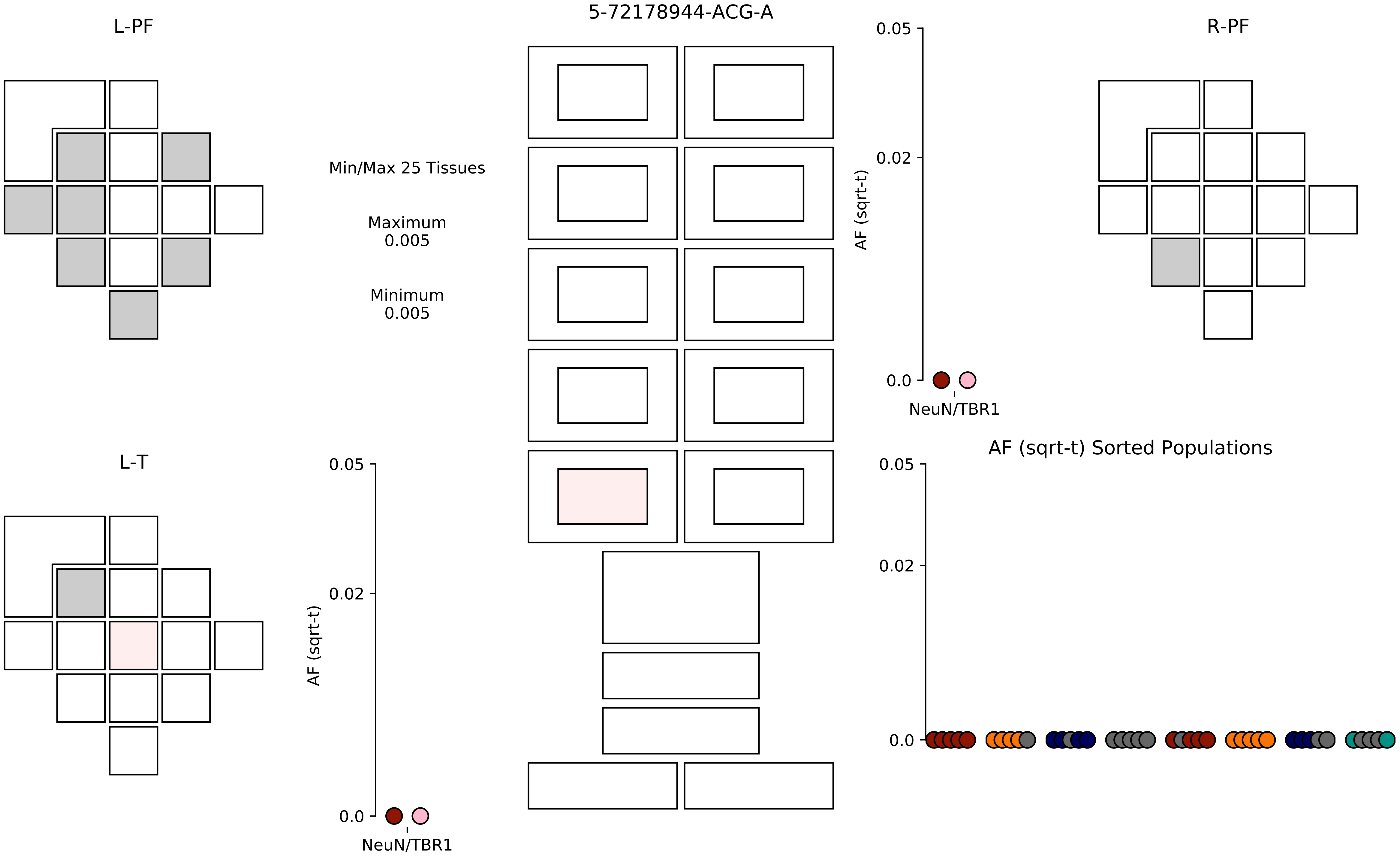

5-72178944-ACG-A
L-PF
R-PF
L-T
Min/Max 25 Tissues
Maximum
0.005
Minimum
0.005
AF (sqrt-t)
0.05
0.02
0.0
NeuN/TBR1
AF (sqrt-t) Sorted Populations

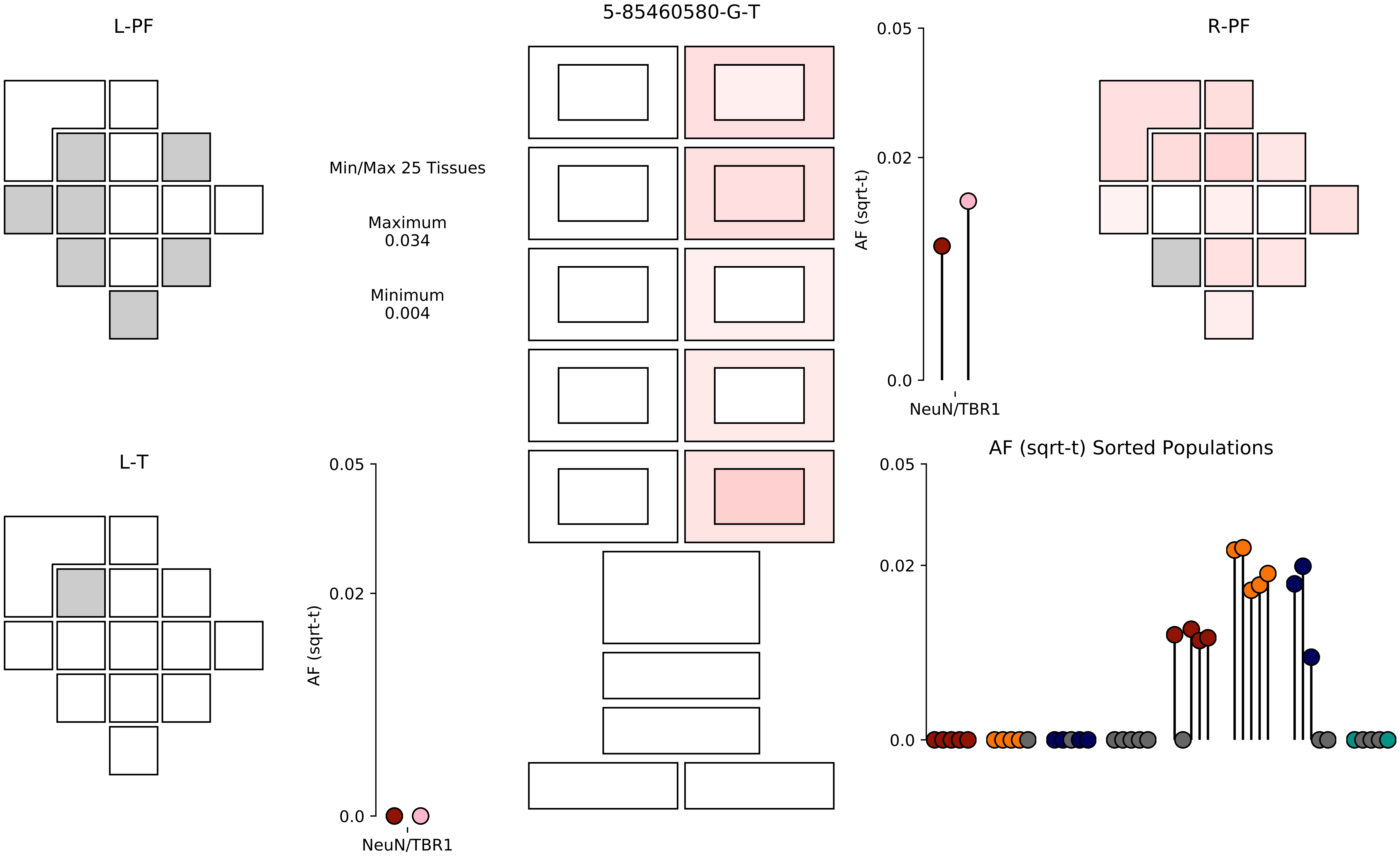

5-85460580-G-T
L-PF
R-PF
L-T
Min/Max 25 Tissues
Maximum
0.034
Minimum
0.004
AF (sqrt-t)
0.05
0.02
0.0
NeuN/TBR1
AF (sqrt-t) Sorted Populations

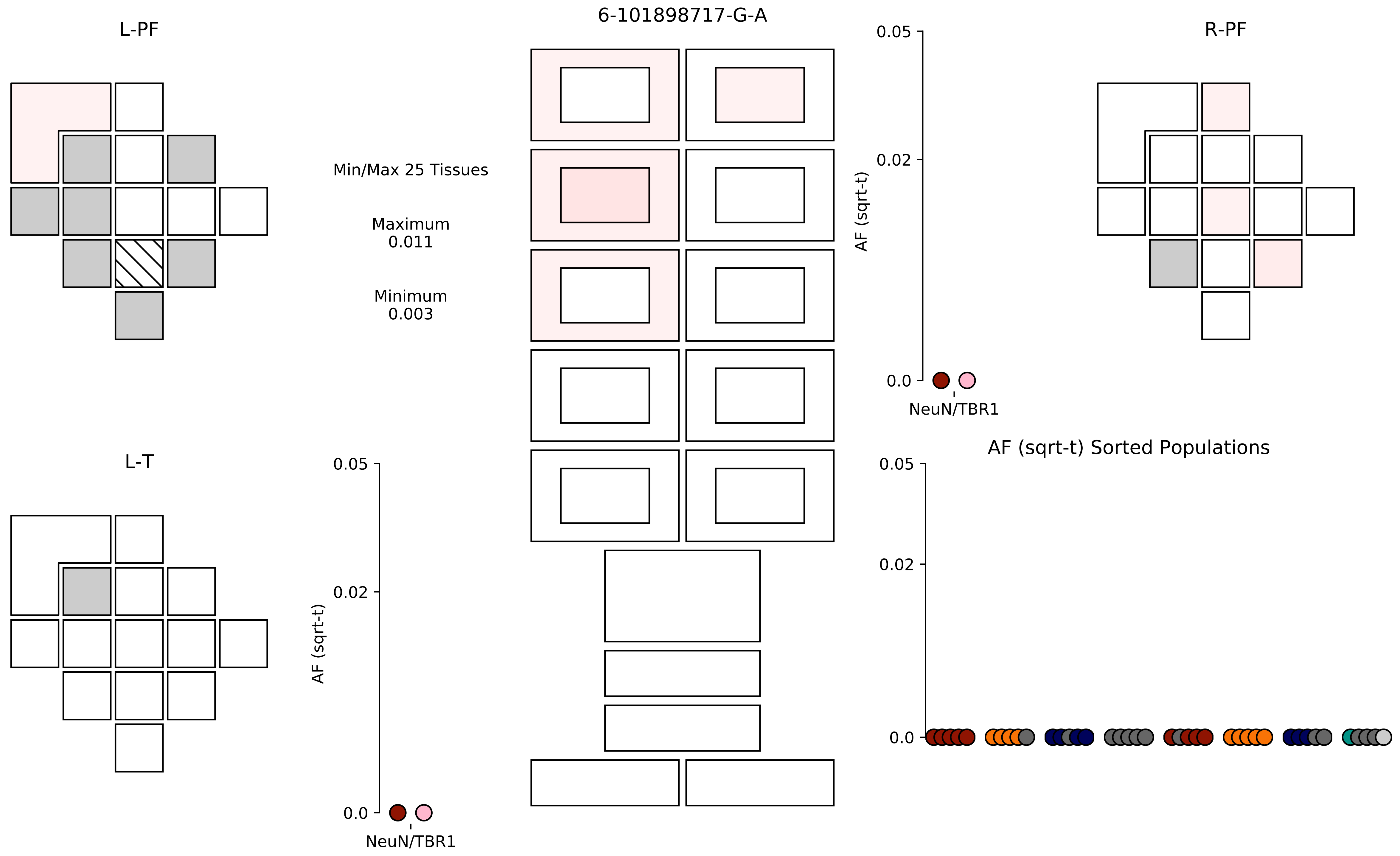

6-101898717-G-A
L-PF
R-PF
L-T
Min/Max 25 Tissues
Maximum
0.011
Minimum
0.003
AF (sqrt-t)
0.05
0.02
0.0
NeuN/TBR1
AF (sqrt-t) Sorted Populations

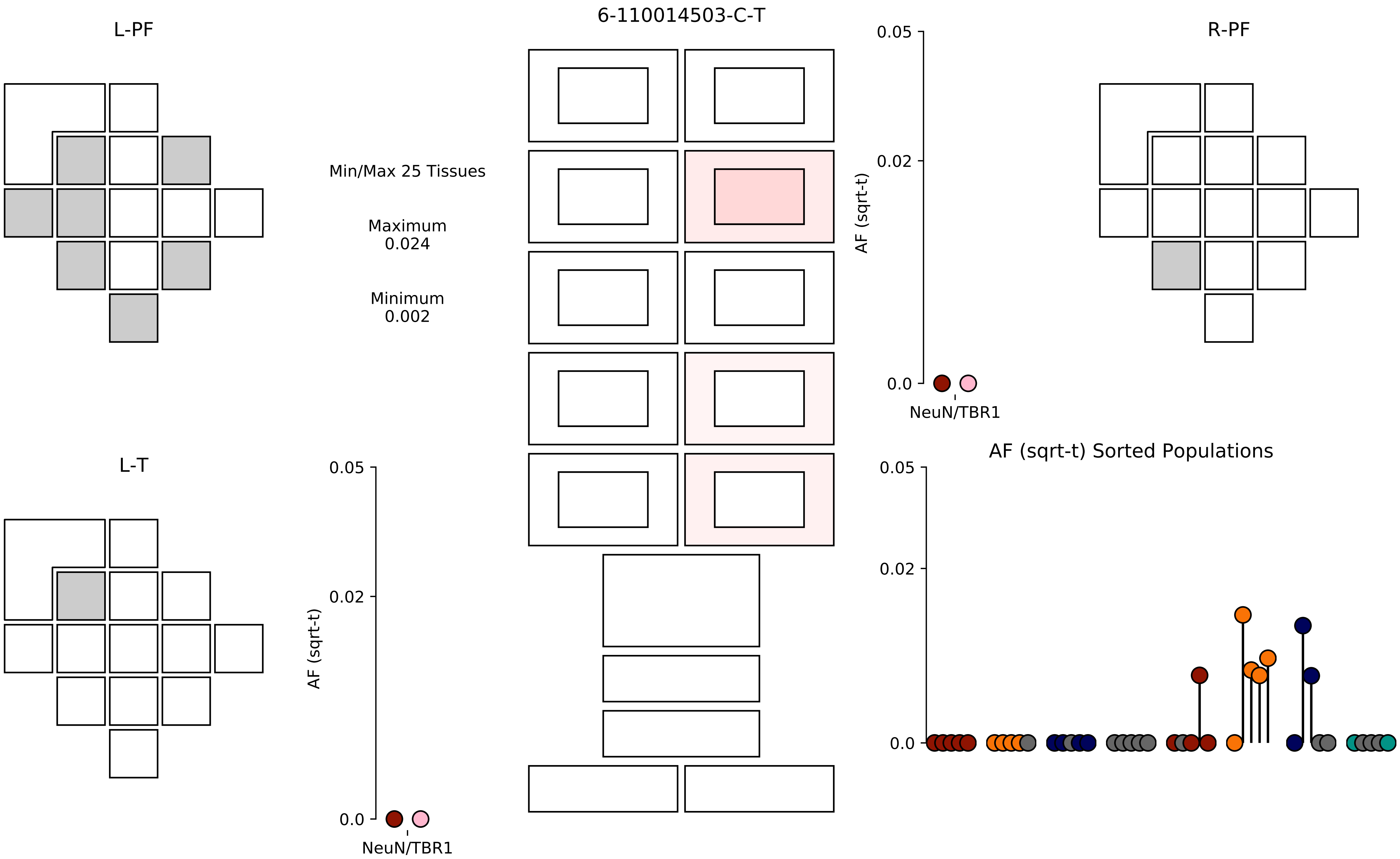
6-110014503-C-T
L-PF
R-PF
L-T
Min/Max 25 Tissues
Maximum
0.024
Minimum
0.002
AF (sqrt-t)
0.05
0.02
0.0
NeuN/TBR1
AF (sqrt-t) Sorted Populations

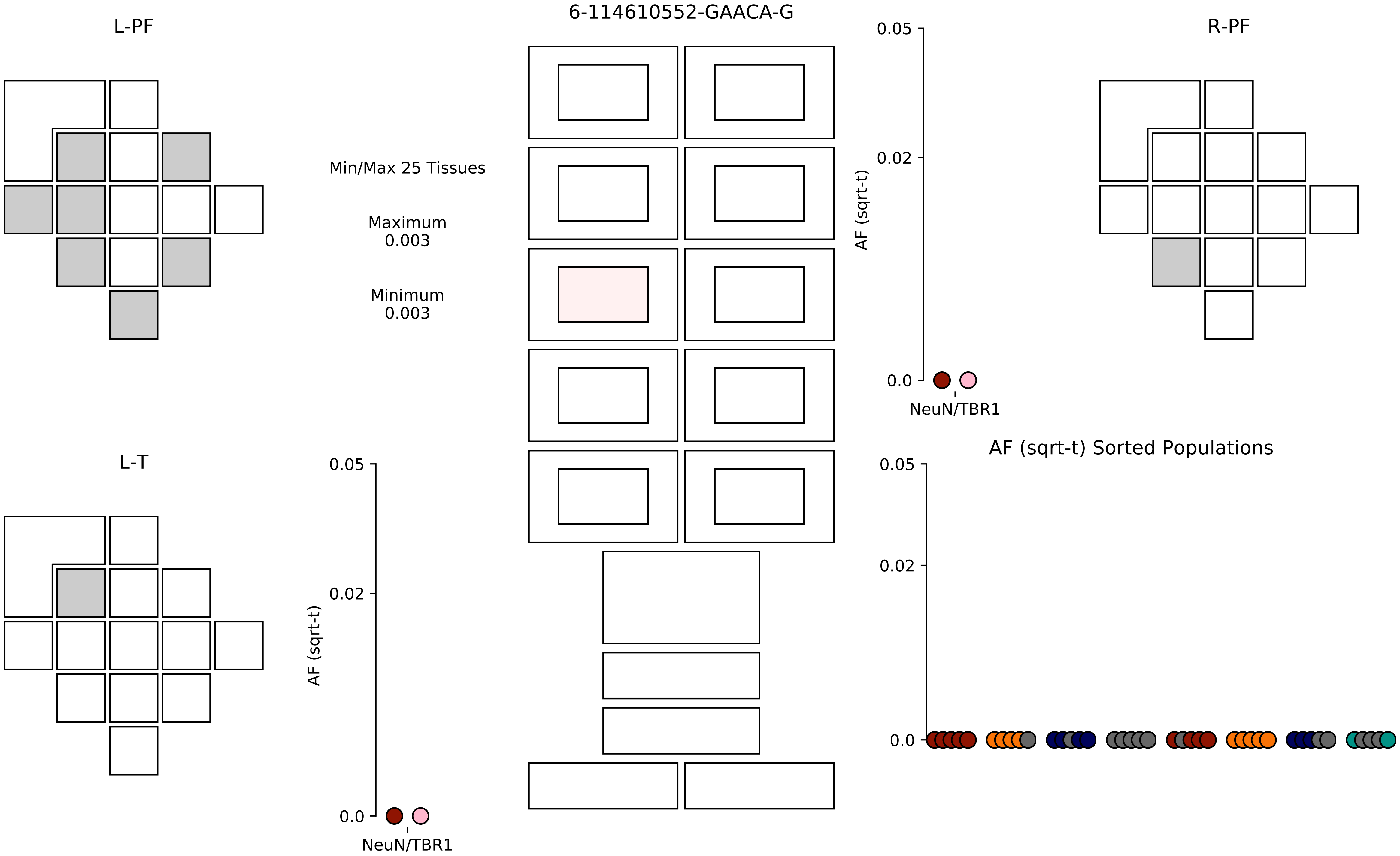

6-114610552-GAACA-G
L-PF
R-PF
L-T
Min/Max 25 Tissues
Maximum
0.003
Minimum
0.003
0.05
0.02
0.0
AF (sqrt-t)
NeuN/TBR1
AF (sqrt-t) Sorted Populations

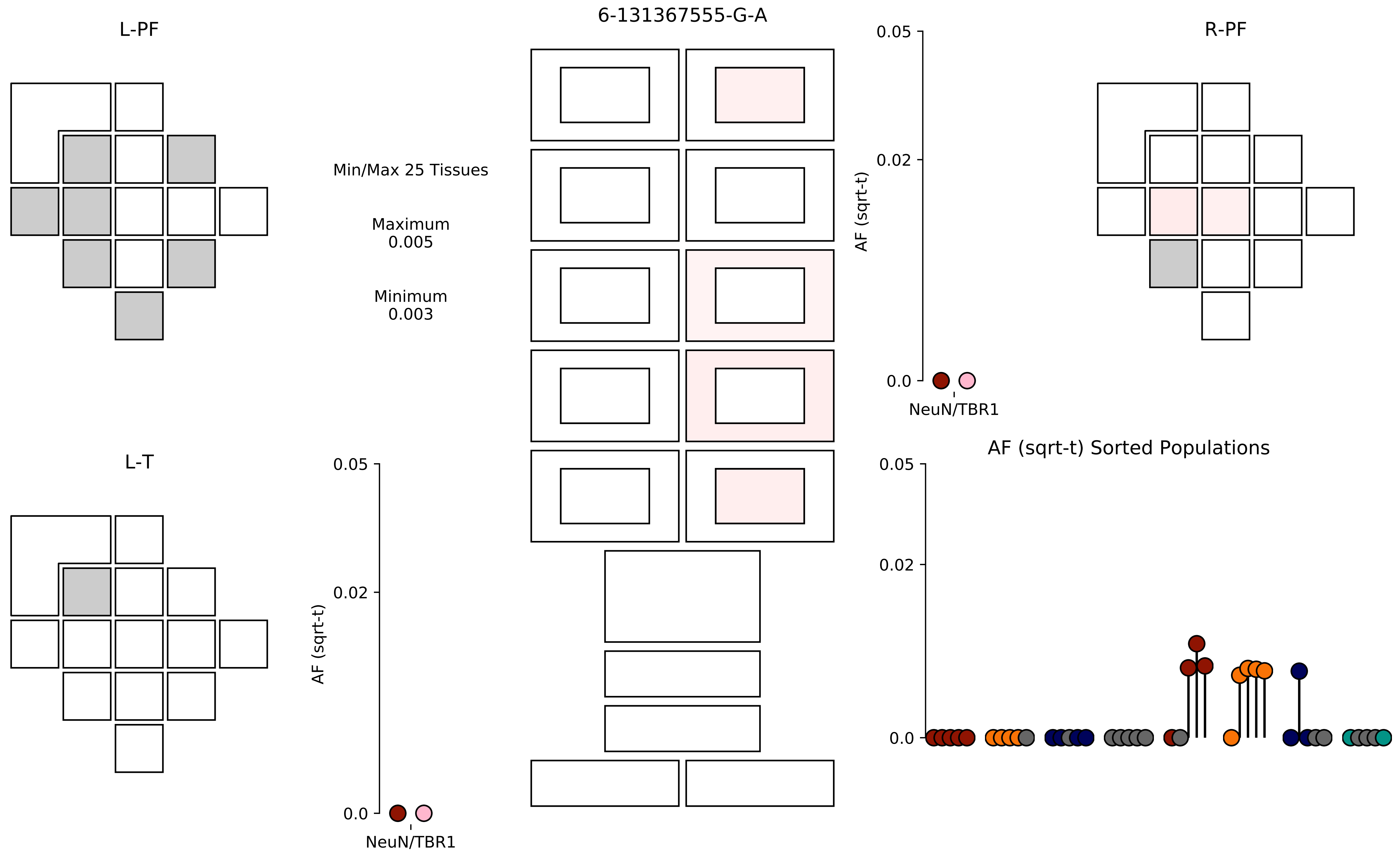

6-131367555-G-A
L-PF
R-PF
L-T
Min/Max 25 Tissues
Maximum
0.005
Minimum
0.003
AF (sqrt-t)
0.05
0.02
0.0
NeuN/TBR1
AF (sqrt-t)
0.05
0.02
0.0
NeuN/TBR1
AF (sqrt-t) Sorted Populations
0.05
0.02
0.0

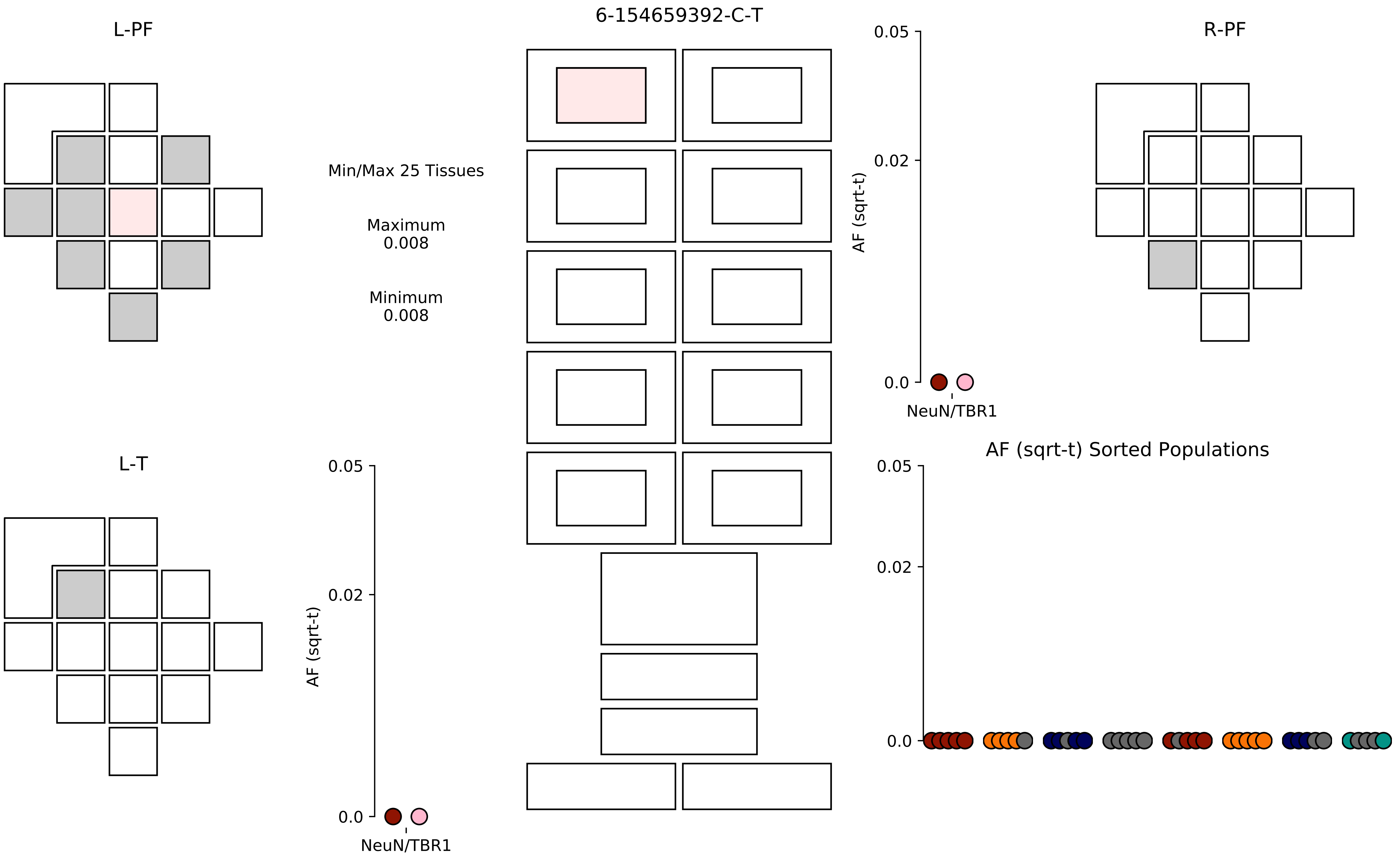

6-154659392-C-T
L-PF
R-PF
L-T
Min/Max 25 Tissues
Maximum
0.008
Minimum
0.008
AF (sqrt-t)
0.05
0.02
0.0
NeuN/TBR1
AF (sqrt-t) Sorted Populations

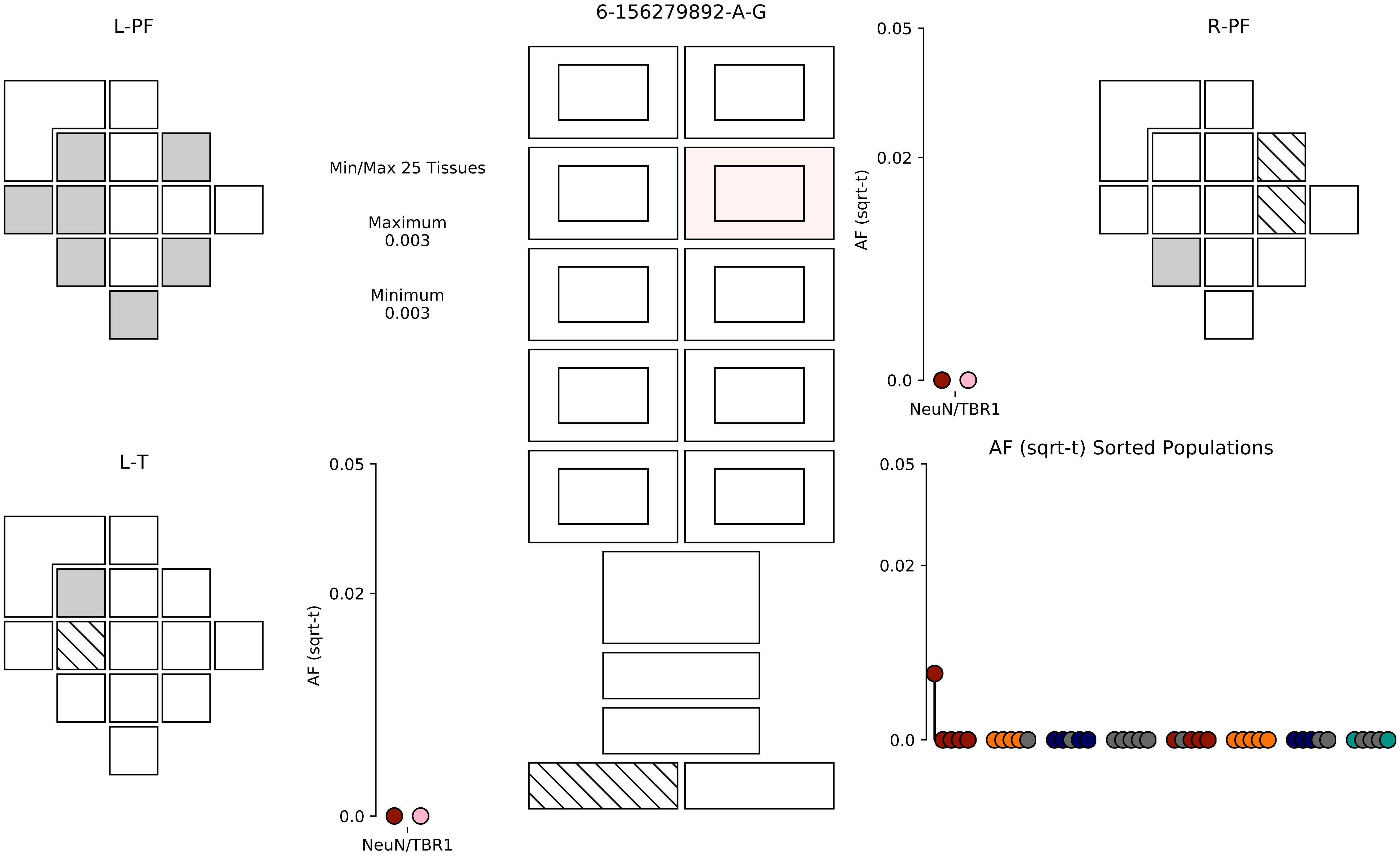
L-PF
6-156279892-A-G
R-PF
Min/Max 25 Tissues
Maximum
0.003
Minimum
0.003
AF (sqrt-t)
0.05
0.02
0.0
NeuN/TBR1
L-T
AF (sqrt-t) Sorted Populations

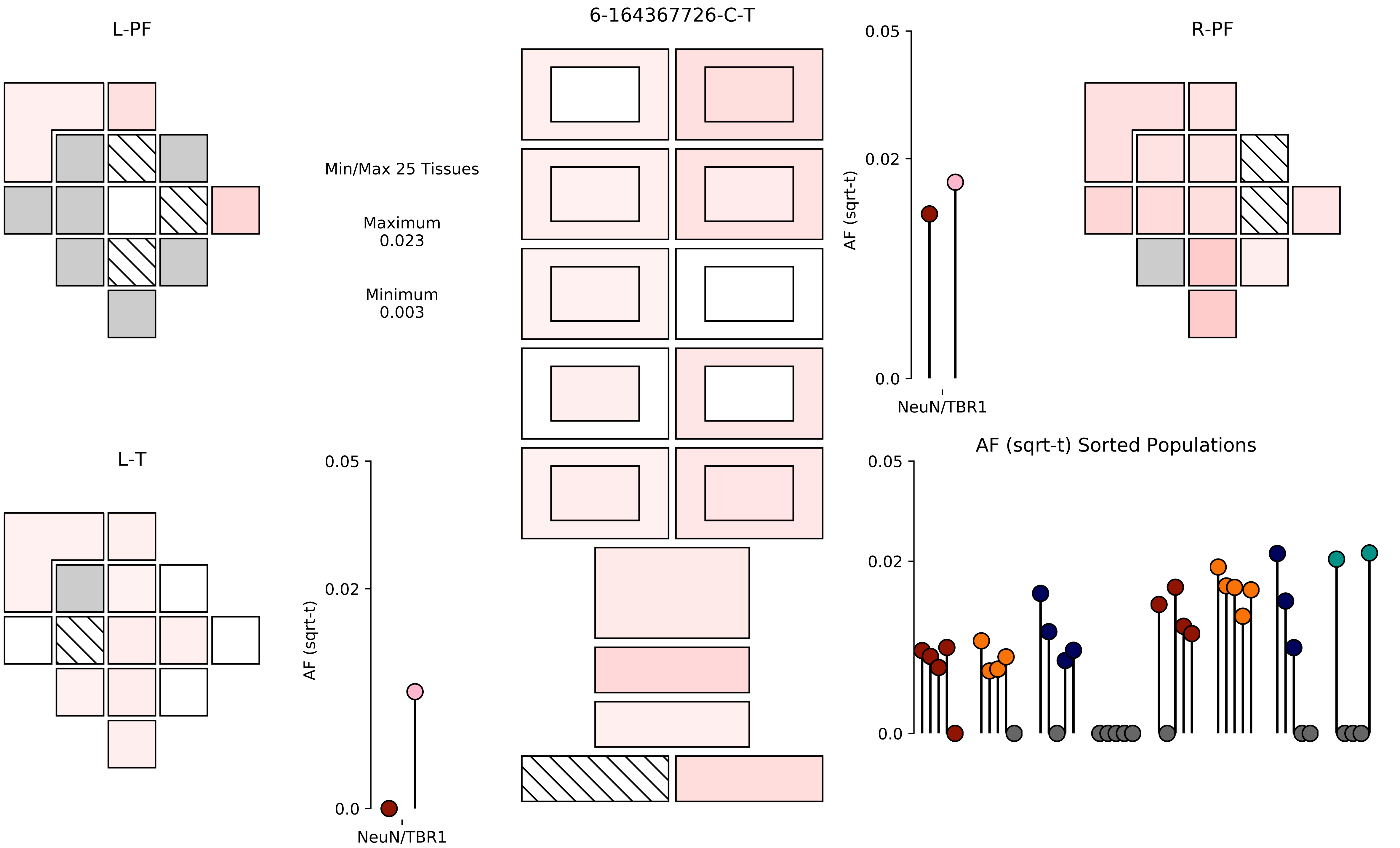

6-164367726-C-T
L-PF
R-PF
L-T
Min/Max 25 Tissues
Maximum
0.023
Minimum
0.003
AF (sqrt-t)
NeuN/TBR1
0.05
0.02
0.0
AF (sqrt-t) Sorted Populations

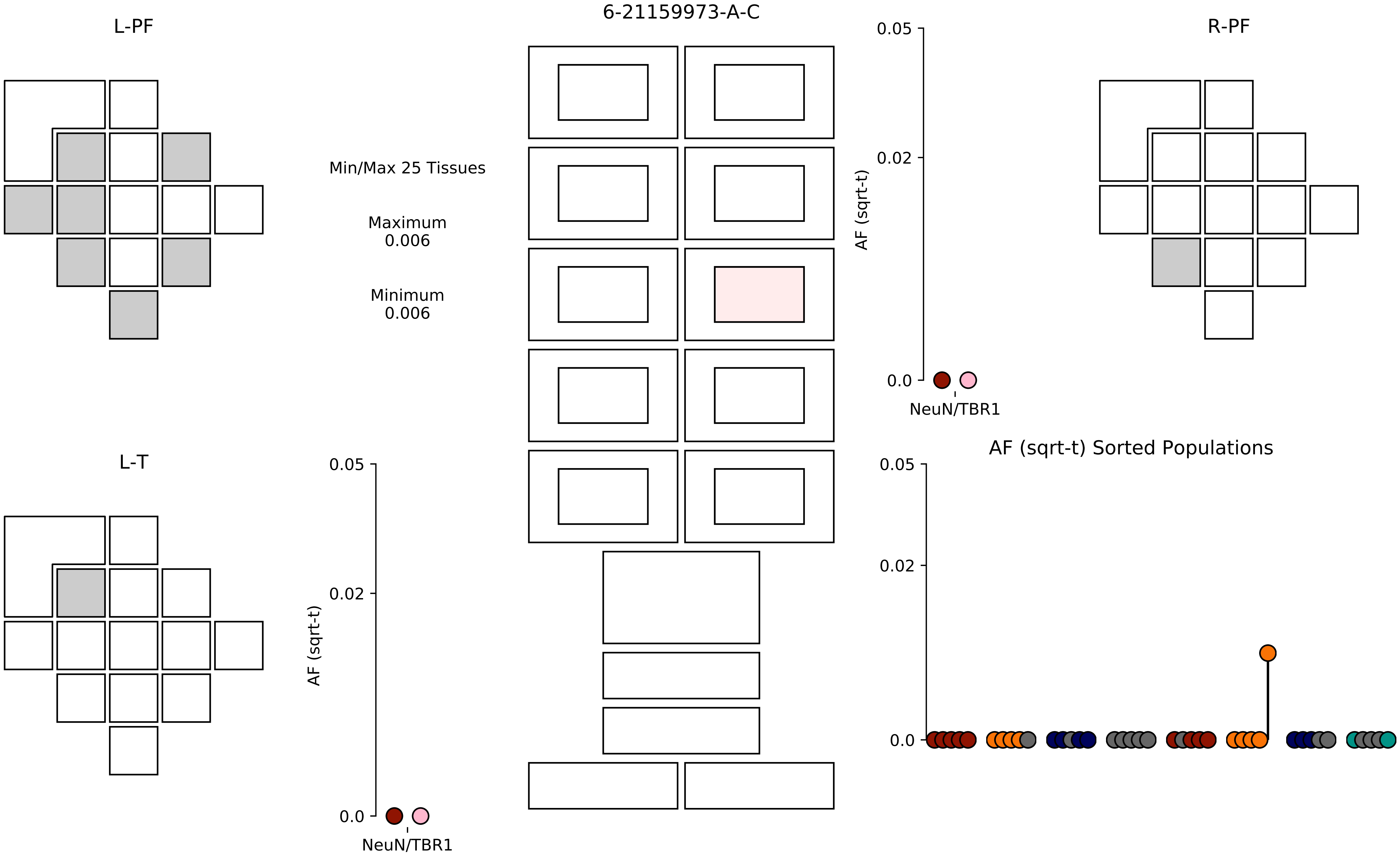

6-21159973-A-C
L-PF
R-PF
L-T
Min/Max 25 Tissues
Maximum
0.006
Minimum
0.006
AF (sqrt-t)
0.05
0.02
0.0
NeuN/TBR1
AF (sqrt-t) Sorted Populations

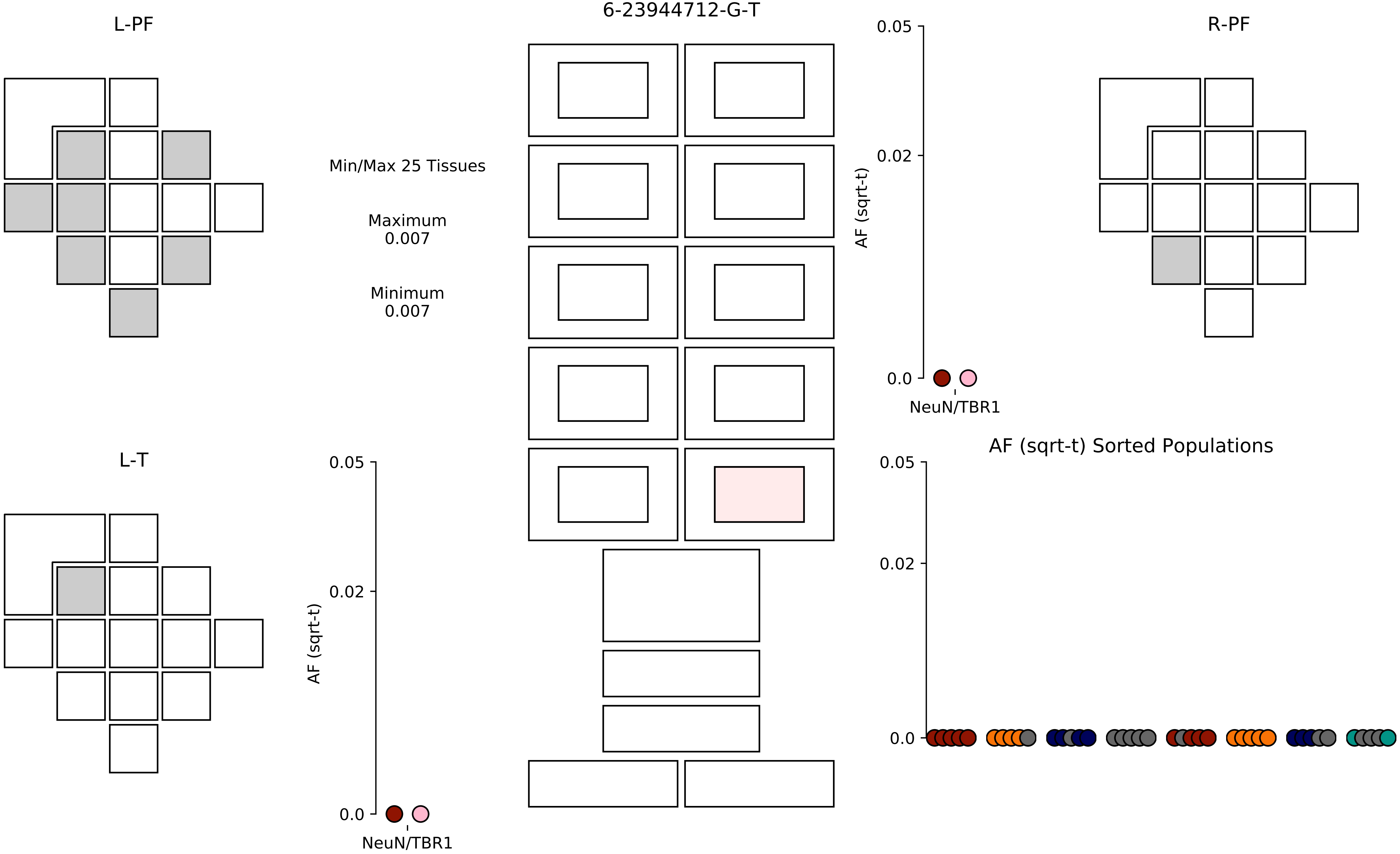

6-23944712-G-T
L-PF
R-PF
L-T
Min/Max 25 Tissues
Maximum
0.007
Minimum
0.007
AF (sqrt-t)
0.05
0.02
0.0
NeuN/TBR1
AF (sqrt-t)
0.05
0.02
0.0
NeuN/TBR1
AF (sqrt-t) Sorted Populations
0.05
0.02
0.0

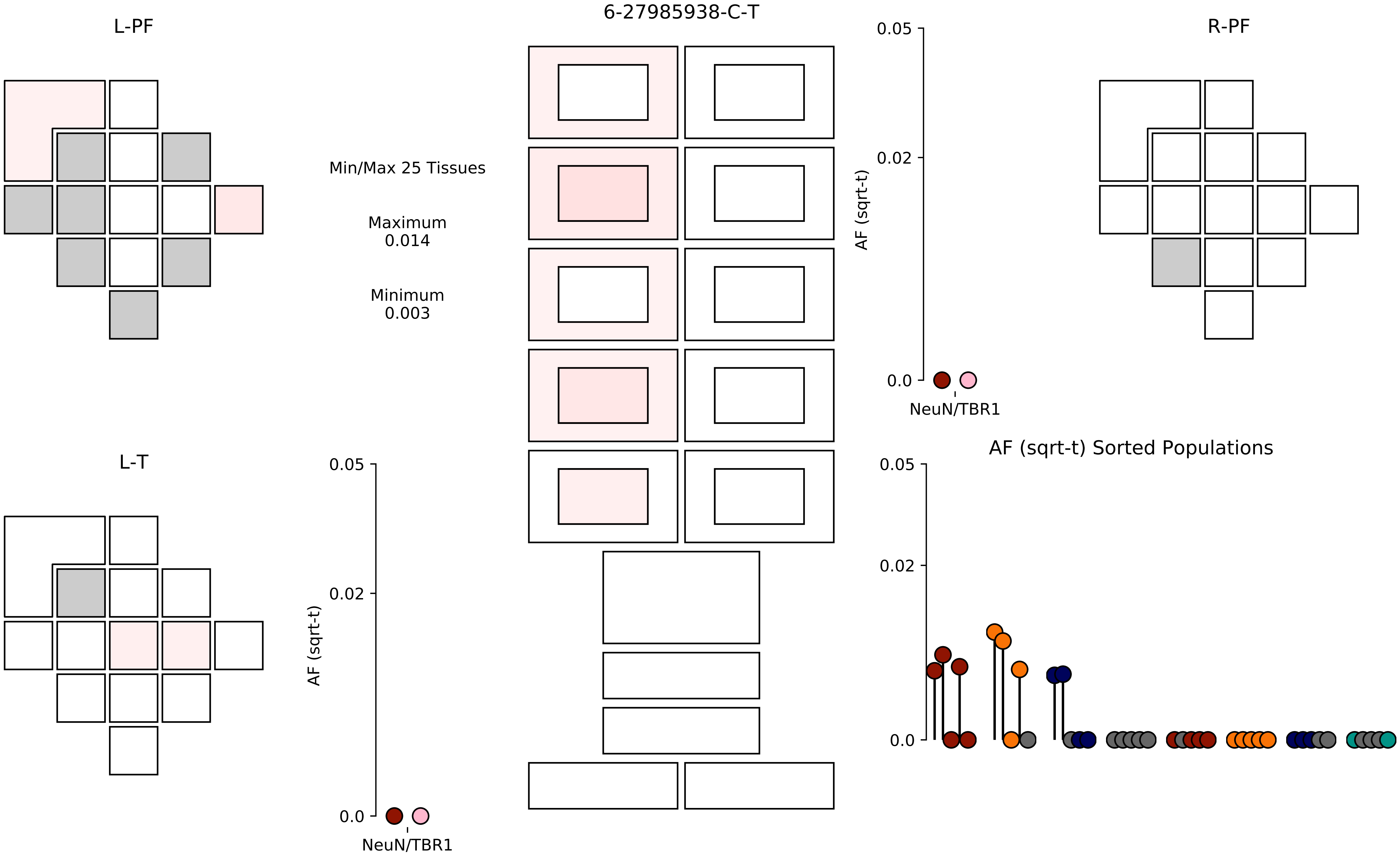

6-27985938-C-T
L-PF
R-PF
L-T
Min/Max 25 Tissues
Maximum
0.014
Minimum
0.003
AF (sqrt-t)
0.05
0.02
0.0
NeuN/TBR1
AF (sqrt-t) Sorted Populations

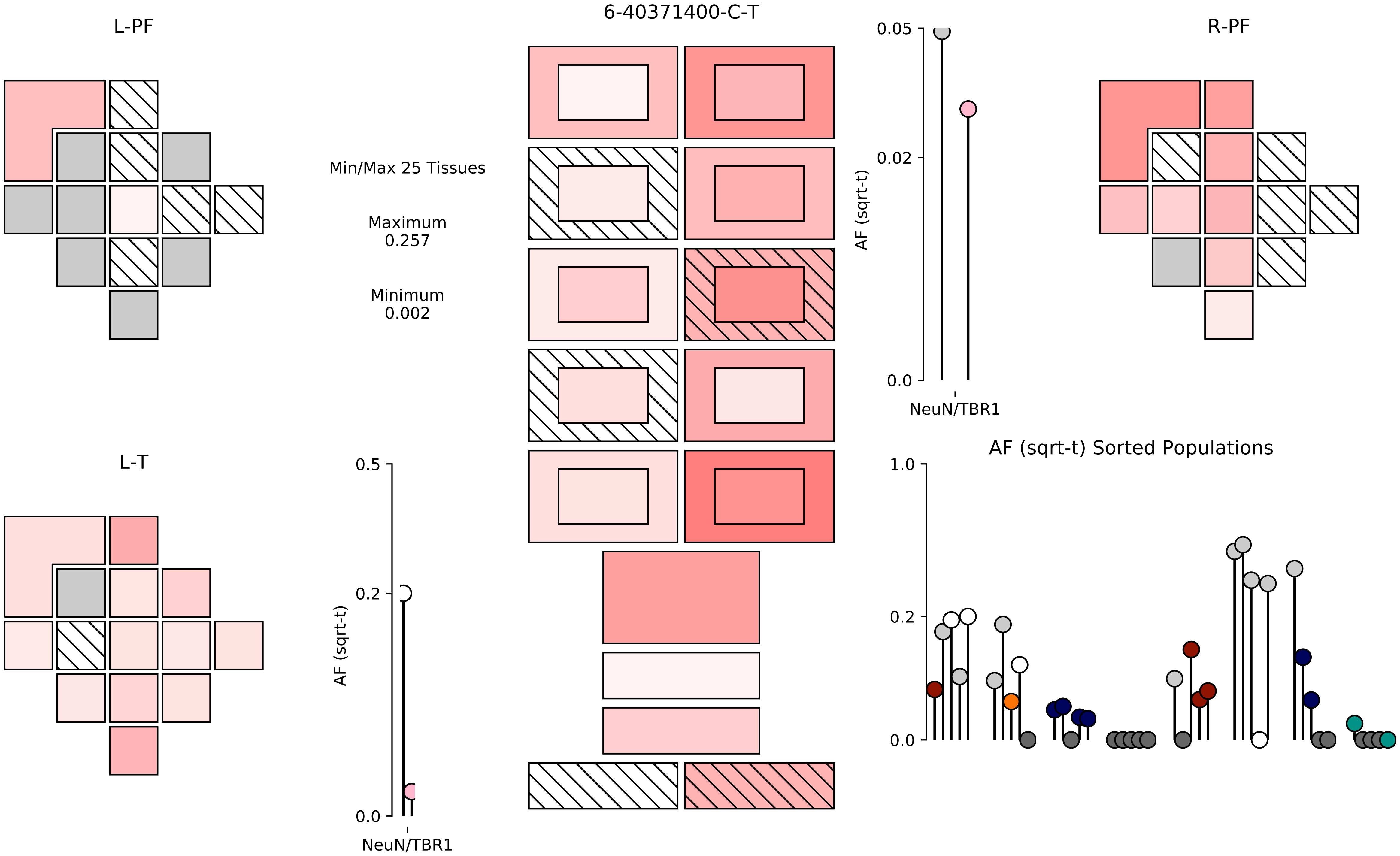

L-PF
6-40371400-C-T
R-PF
Min/Max 25 Tissues
Maximum
0.257
Minimum
0.002
0.05
0.02
0.0
AF (sqrt-t)
NeuN/TBR1
L-T
0.5
0.2
0.0
AF (sqrt-t)
NeuN/TBR1
AF (sqrt-t) Sorted Populations
1.0
0.2
0.0

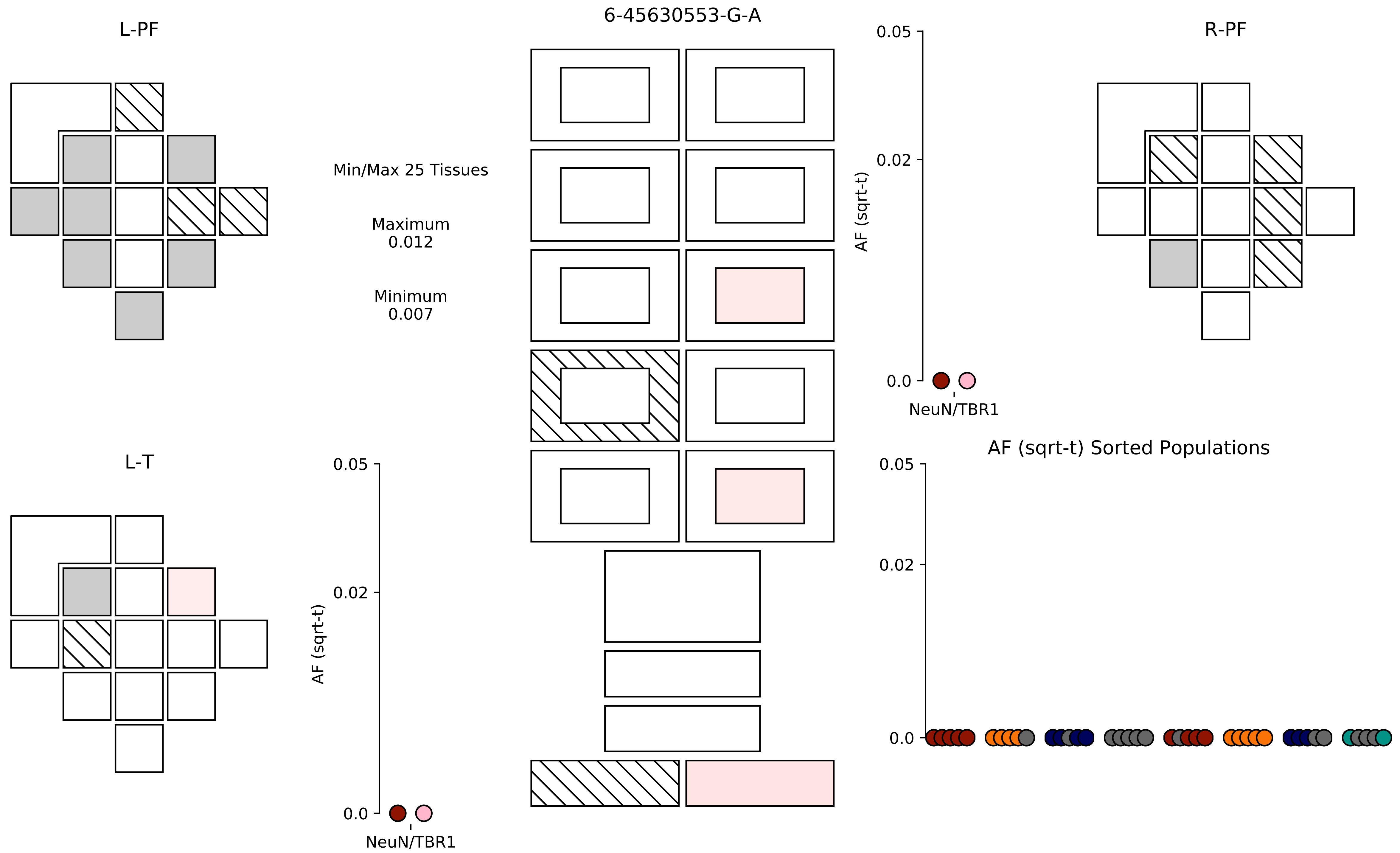

L-PF
6-45630553-G-A
R-PF
Min/Max 25 Tissues
Maximum
0.012
Minimum
0.007
0.05
0.02
0.0
AF (sqrt-t)
NeuN/TBR1
L-T
0.05
0.02
0.0
AF (sqrt-t)
NeuN/TBR1
AF (sqrt-t) Sorted Populations
0.05
0.02
0.0

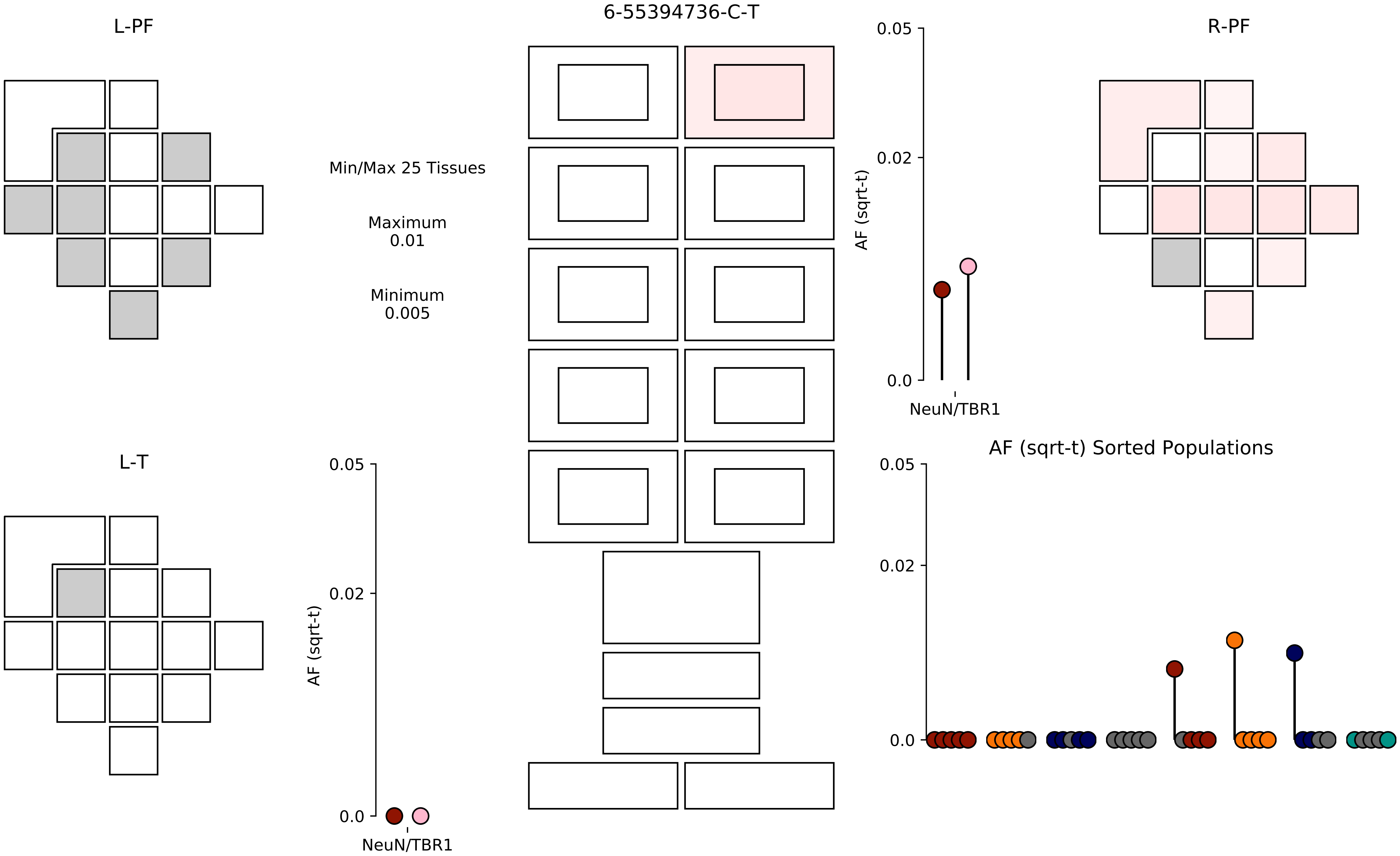

6-55394736-C-T
L-PF
R-PF
L-T
Min/Max 25 Tissues
Maximum
0.01
Minimum
0.005
AF (sqrt-t)
0.05
0.02
0.0
NeuN/TBR1
AF (sqrt-t) Sorted Populations

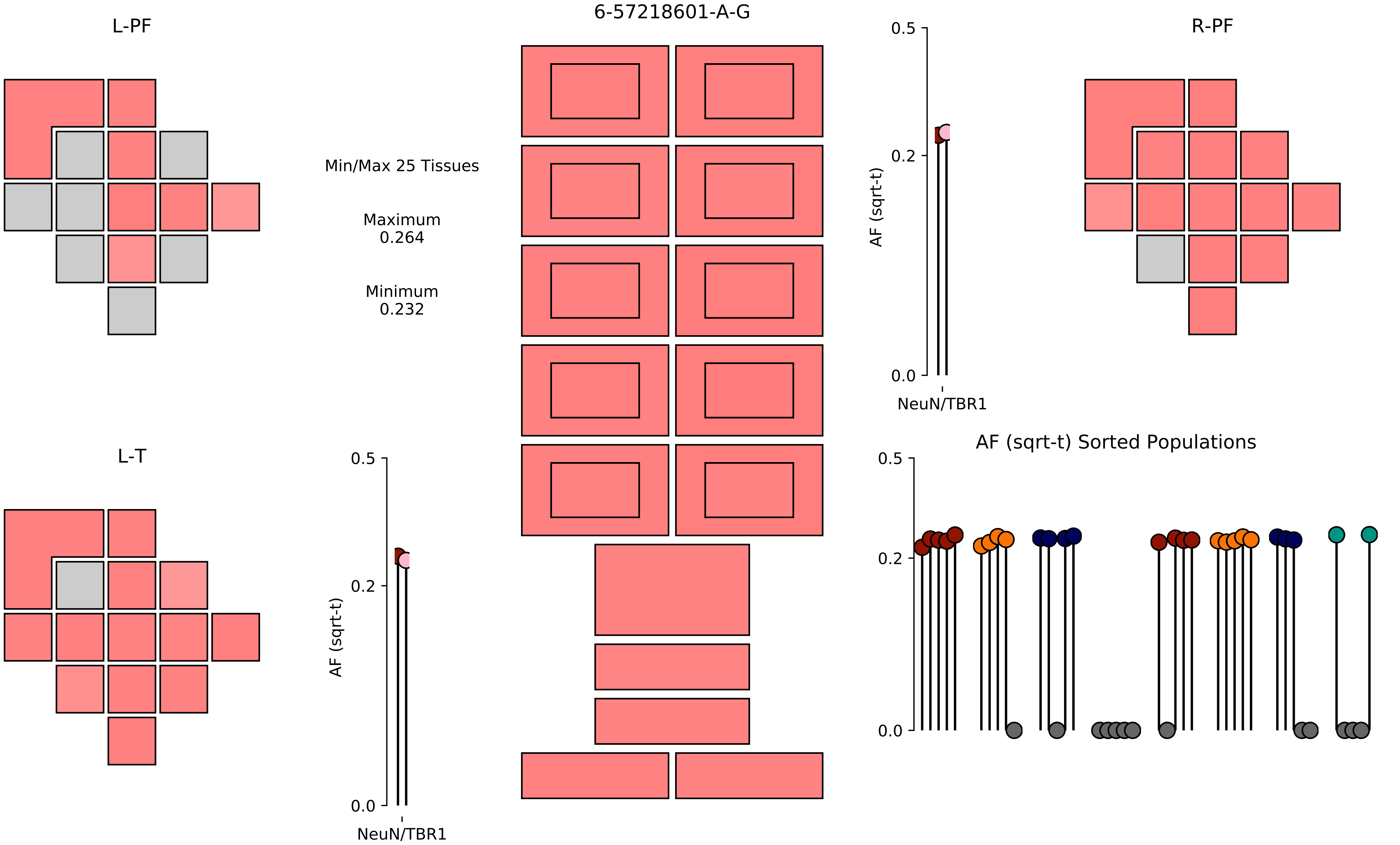

6-57218601-A-G
L-PF
R-PF
L-T
Min/Max 25 Tissues
Maximum
0.264
Minimum
0.232
AF (sqrt-t)
0.5
0.2
0.0
NeuN/TBR1
AF (sqrt-t) Sorted Populations

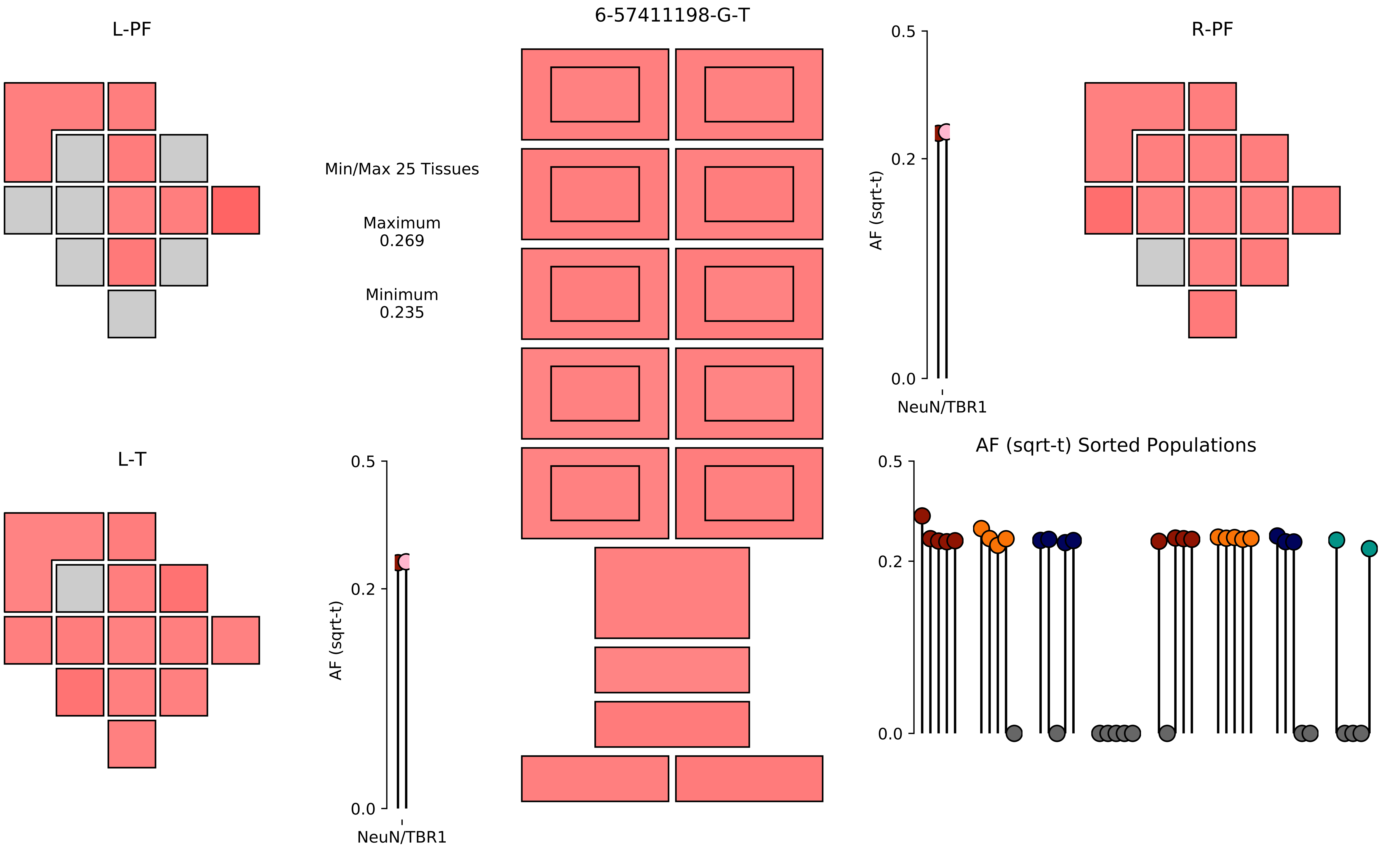
6-57411198-G-T
L-PF
R-PF
L-T
Min/Max 25 Tissues
Maximum
0.269
Minimum
0.235
AF (sqrt-t)
0.5
0.2
0.0
NeuN/TBR1
AF (sqrt-t)
0.5
0.2
0.0
NeuN/TBR1
AF (sqrt-t) Sorted Populations
0.5
0.2
0.0

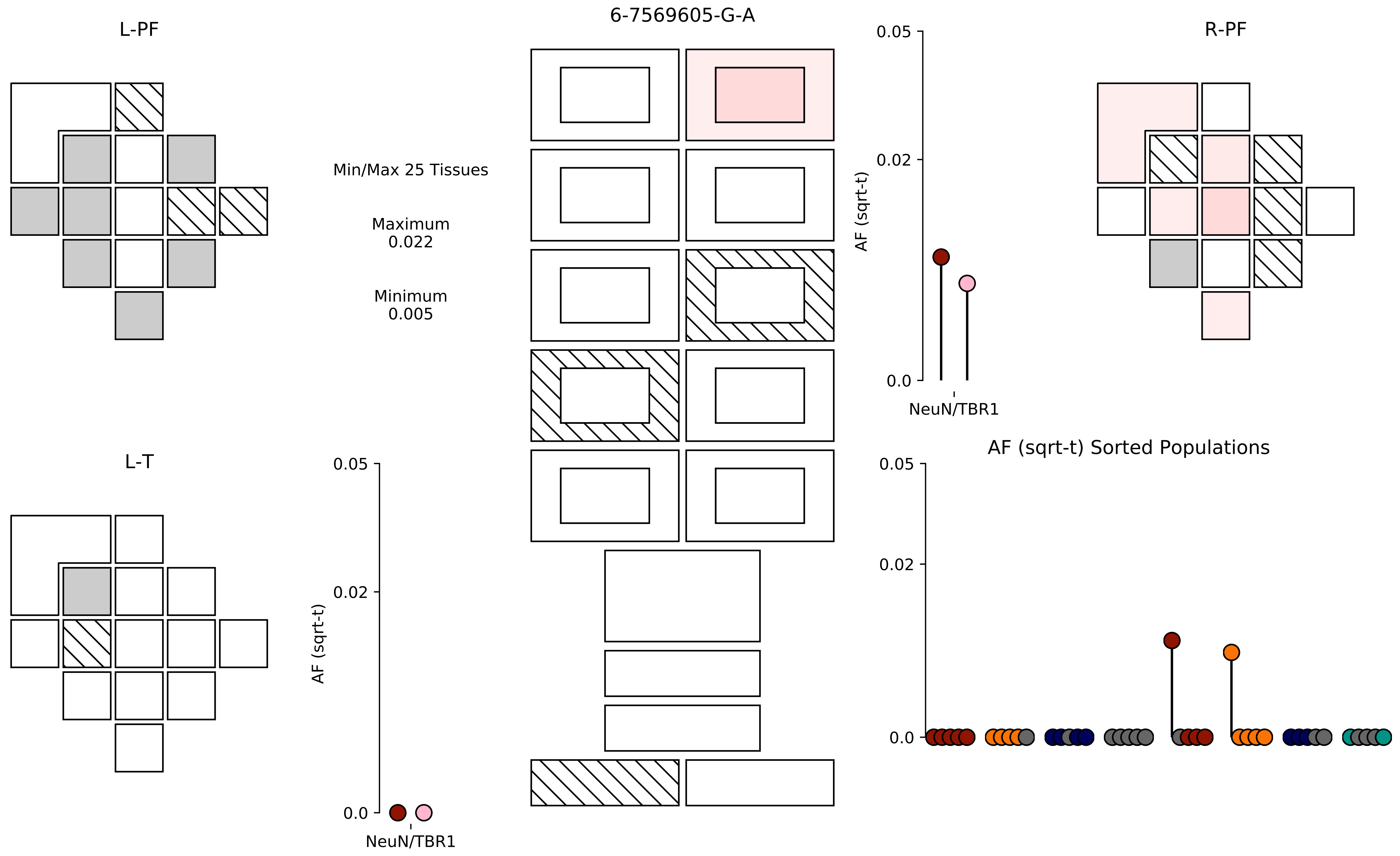

6-7569605-G-A
L-PF
Min/Max 25 Tissues
Maximum
0.022
Minimum
0.005
R-PF
0.05
0.02
0.0
AF (sqrt-t)
NeuN/TBR1
L-T
0.05
0.02
0.0
AF (sqrt-t)
NeuN/TBR1
AF (sqrt-t) Sorted Populations
0.05
0.02
0.0

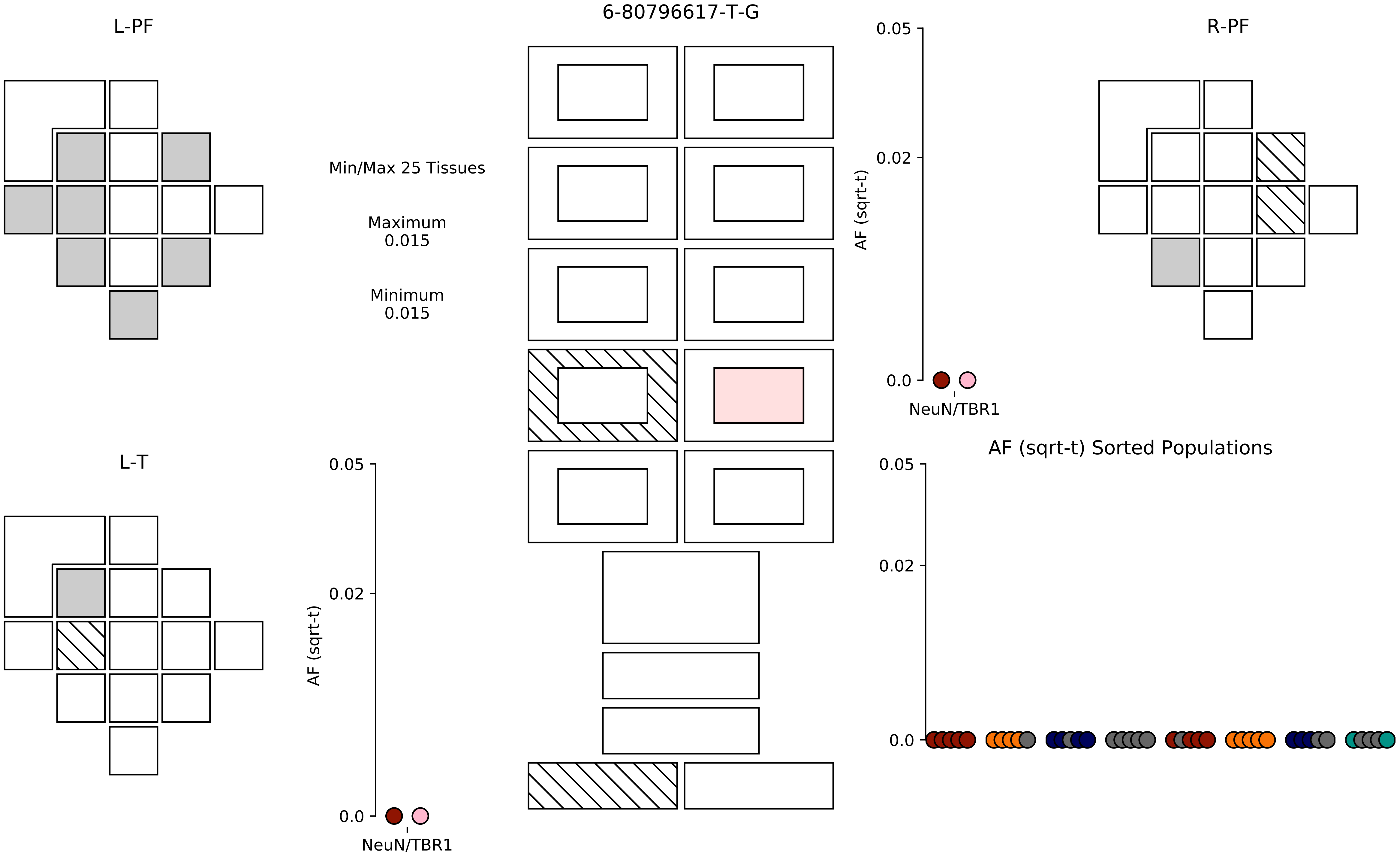

6-80796617-T-G
L-PF
R-PF
L-T
Min/Max 25 Tissues
Maximum
0.015
Minimum
0.015
AF (sqrt-t)
0.05
0.02
0.0
NeuN/TBR1
AF (sqrt-t) Sorted Populations

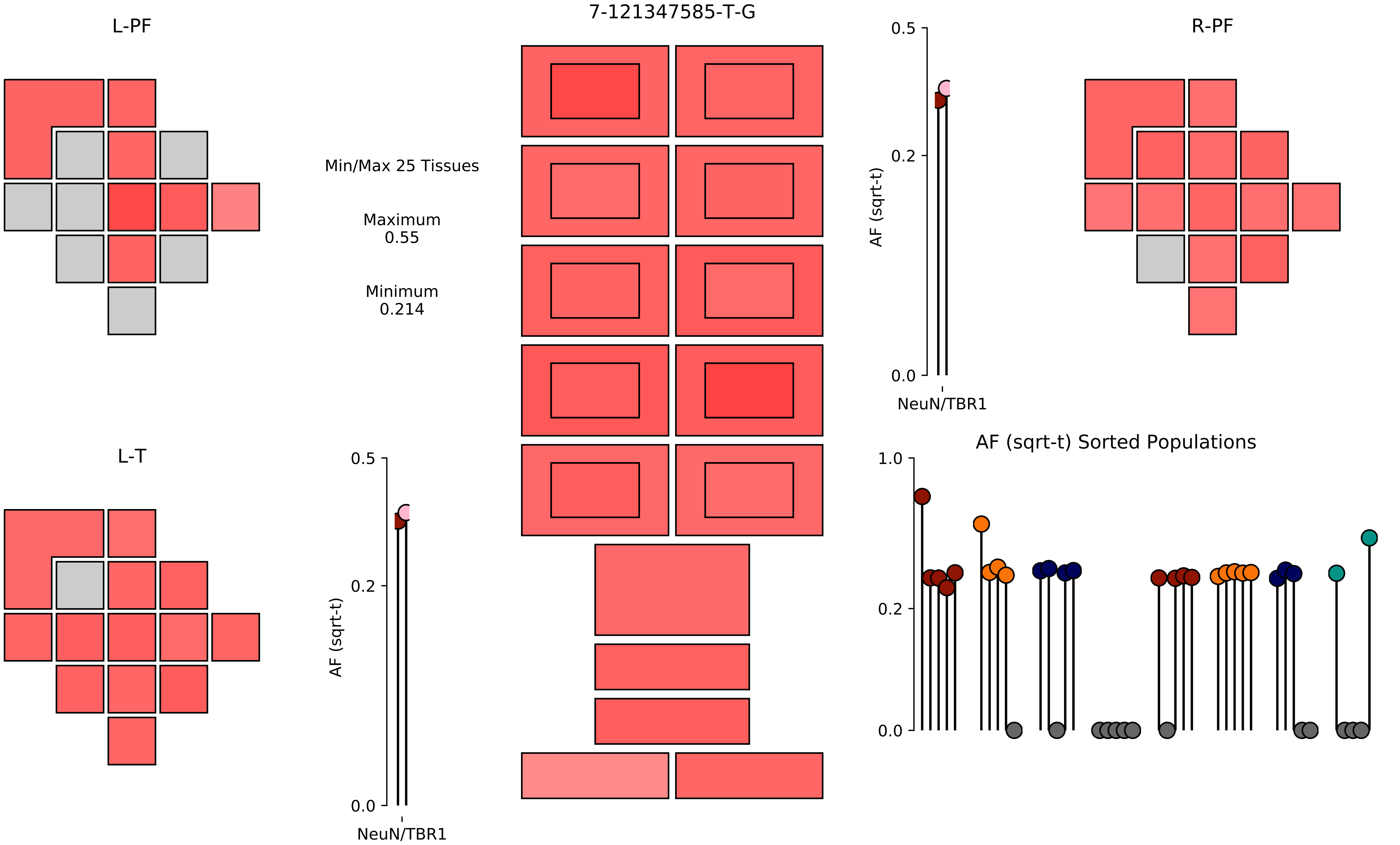
L-PF
7-121347585-T-G
R-PF
Min/Max 25 Tissues
Maximum
0.55
Minimum
0.214
AF (sqrt-t)
0.5
0.2
0.0
NeuN/TBR1
L-T
AF (sqrt-t)
0.5
0.2
0.0
NeuN/TBR1
AF (sqrt-t) Sorted Populations
1.0
0.2
0.0

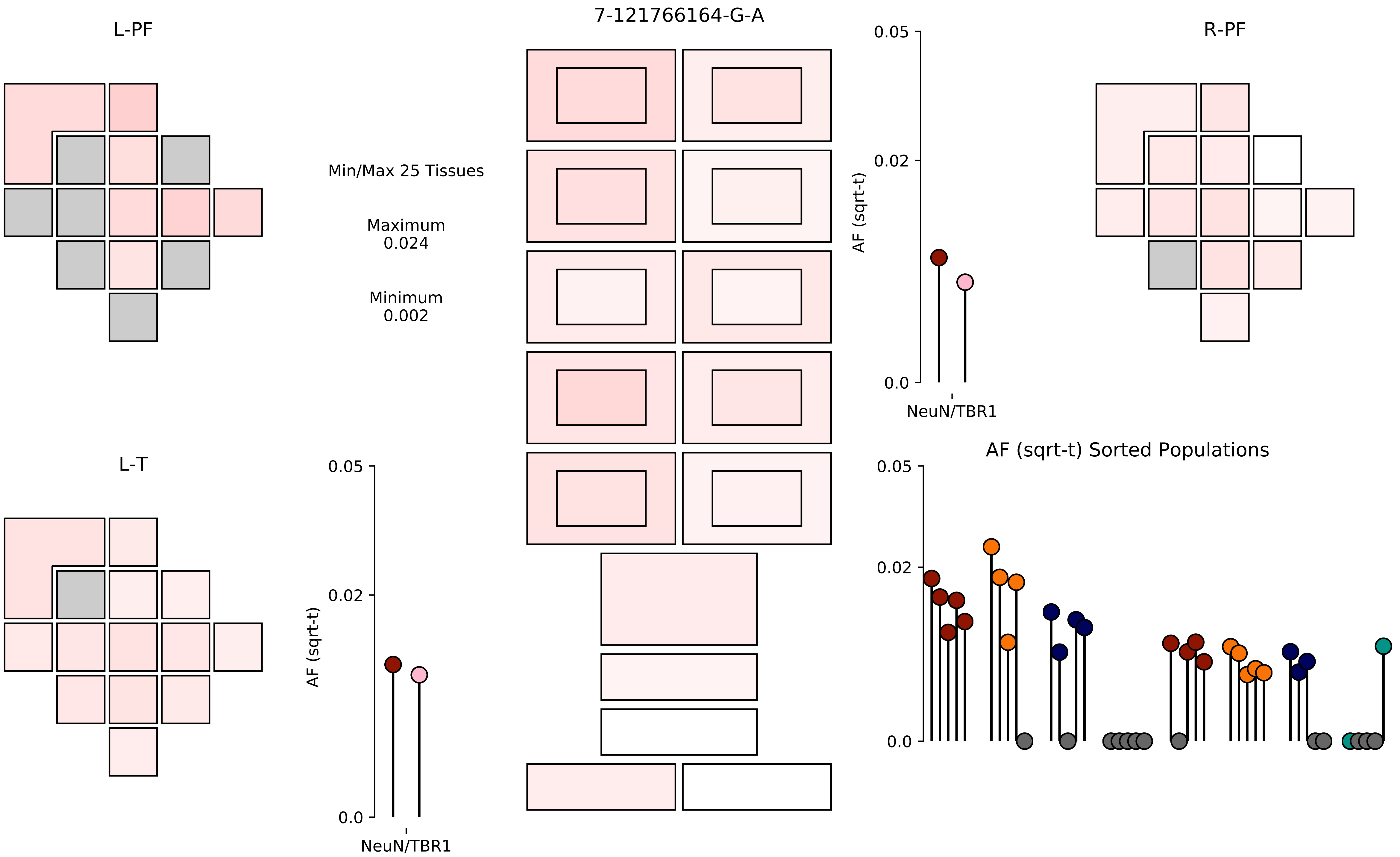

L-PF
7-121766164-G-A
R-PF
Min/Max 25 Tissues
Maximum
0.024
Minimum
0.002
0.05
0.02
0.0
AF (sqrt-t)
NeuN/TBR1
L-T
0.05
0.02
0.0
AF (sqrt-t)
NeuN/TBR1
AF (sqrt-t) Sorted Populations
0.05
0.02
0.0

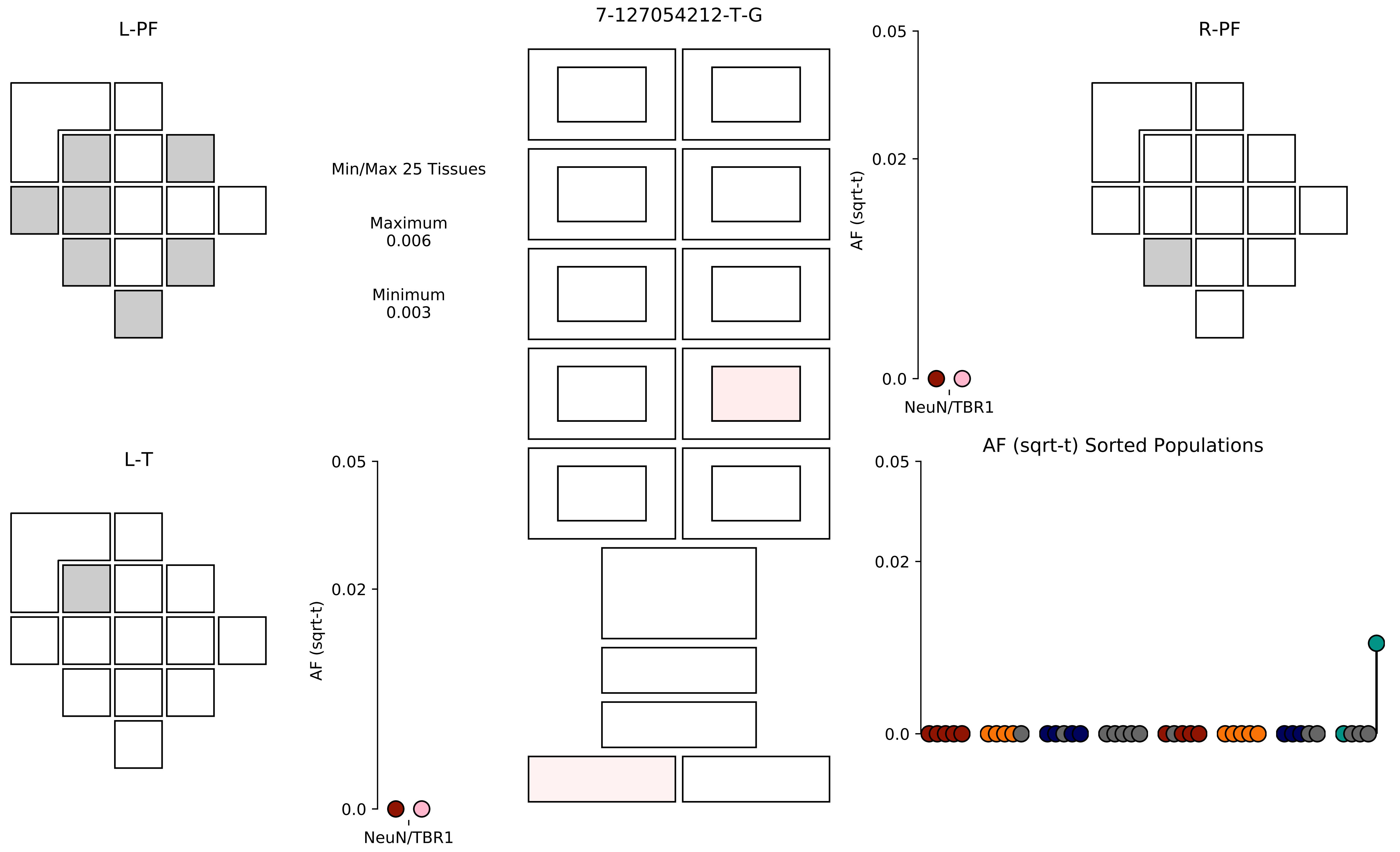

7-127054212-T-G
L-PF
R-PF
L-T
Min/Max 25 Tissues
Maximum
0.006
Minimum
0.003
AF (sqrt-t)
0.05
0.02
0.0
NeuN/TBR1
AF (sqrt-t) Sorted Populations

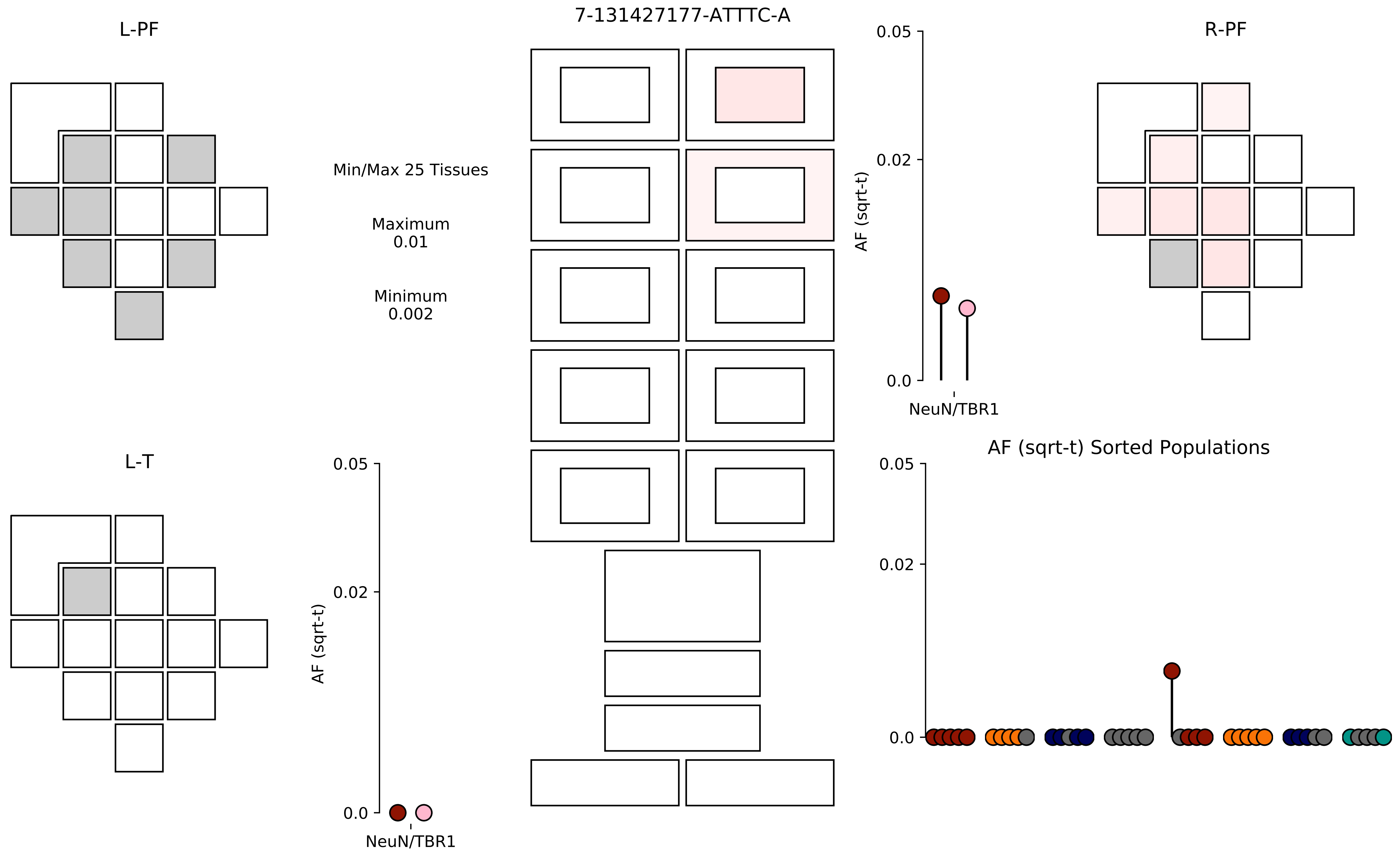
7-131427177-ATTTC-A
L-PF
R-PF
L-T
Min/Max 25 Tissues
Maximum
0.01
Minimum
0.002
AF (sqrt-t)
0.05
0.02
0.0
NeuN/TBR1
AF (sqrt-t) Sorted Populations

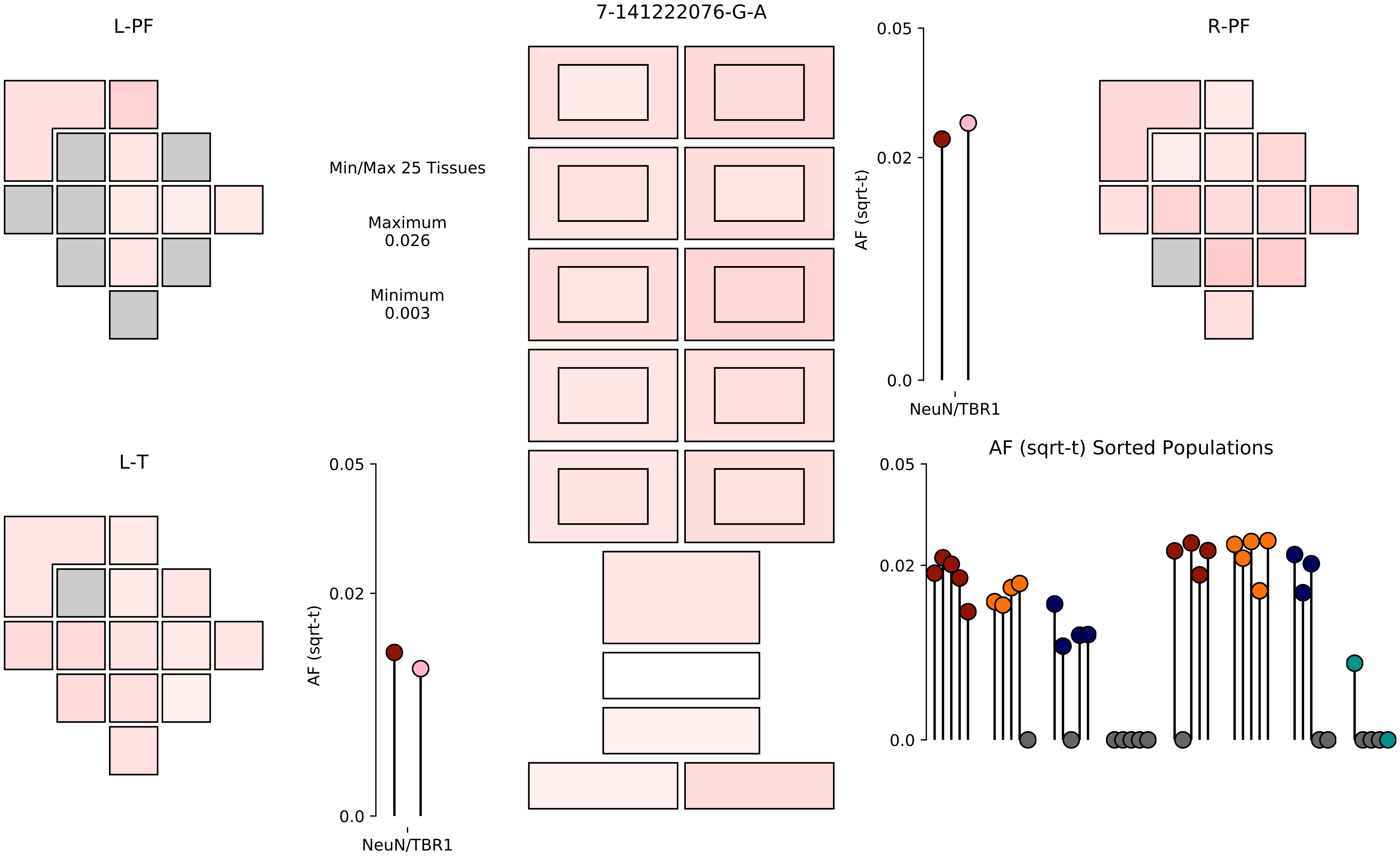
7-141222076-G-A
L-PF
R-PF
L-T
Min/Max 25 Tissues
Maximum
0.026
Minimum
0.003
0.05
0.02
0.0
AF (sqrt-t)
NeuN/TBR1
AF (sqrt-t) Sorted Populations

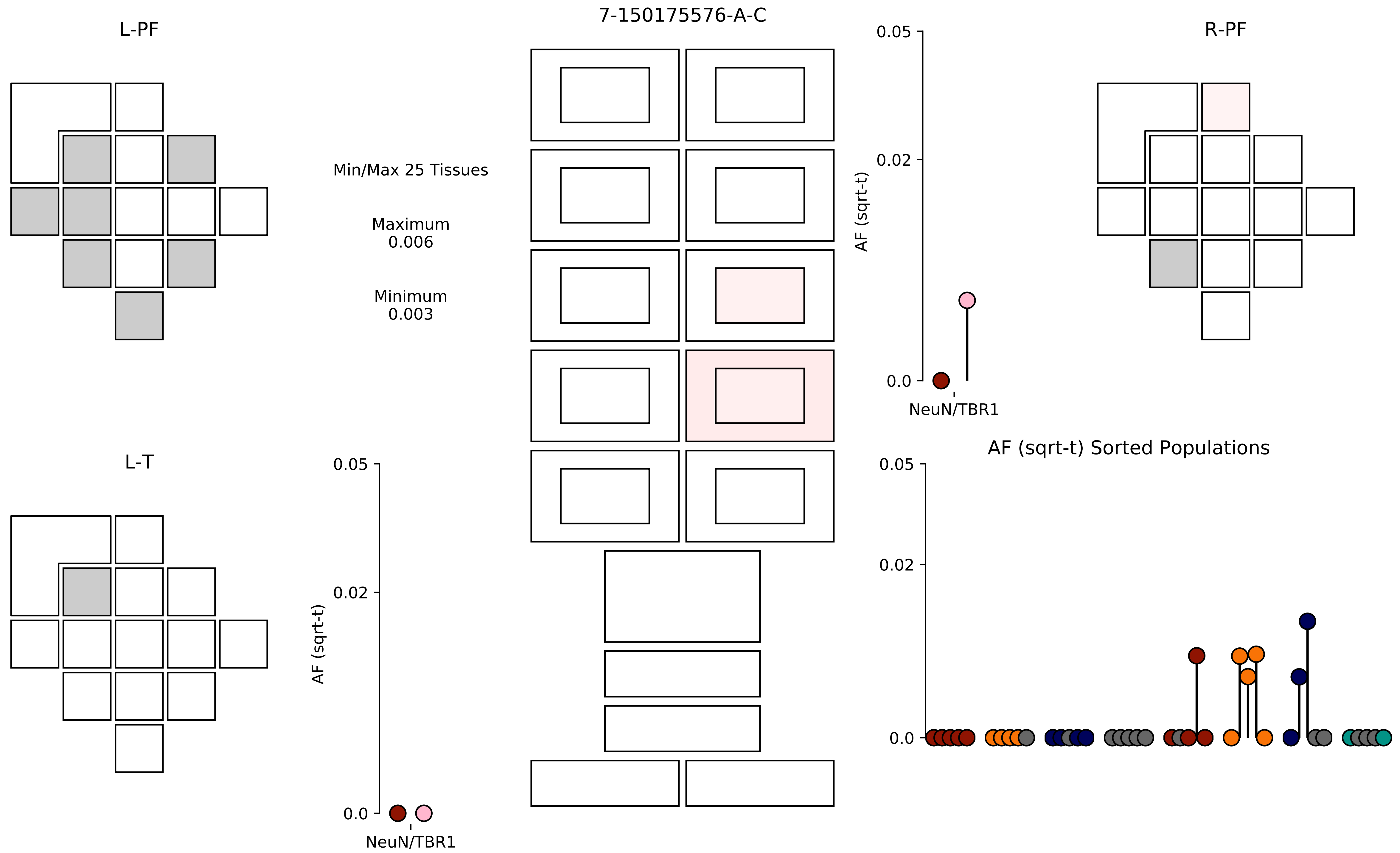

7-150175576-A-C
L-PF
R-PF
L-T
Min/Max 25 Tissues
Maximum
0.006
Minimum
0.003
AF (sqrt-t)
0.05
0.02
0.0
NeuN/TBR1
AF (sqrt-t)
0.05
0.02
0.0
NeuN/TBR1
AF (sqrt-t) Sorted Populations
0.05
0.02
0.0

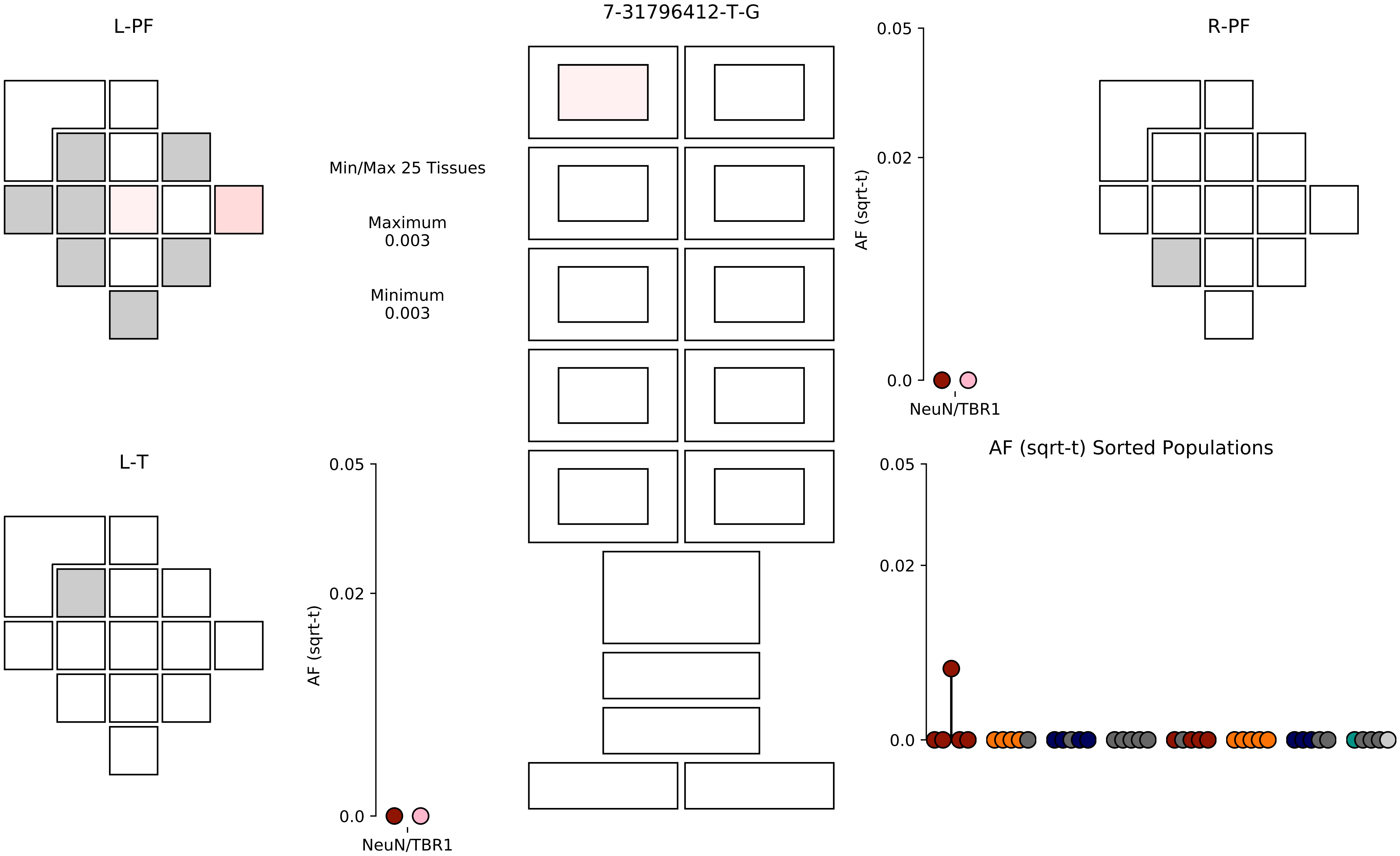
L-PF
7-31796412-T-G
R-PF
Min/Max 25 Tissues
Maximum
0.003
Minimum
0.003
0.05
0.02
0.0
AF (sqrt-t)
NeuN/TBR1
L-T
AF (sqrt-t) Sorted Populations

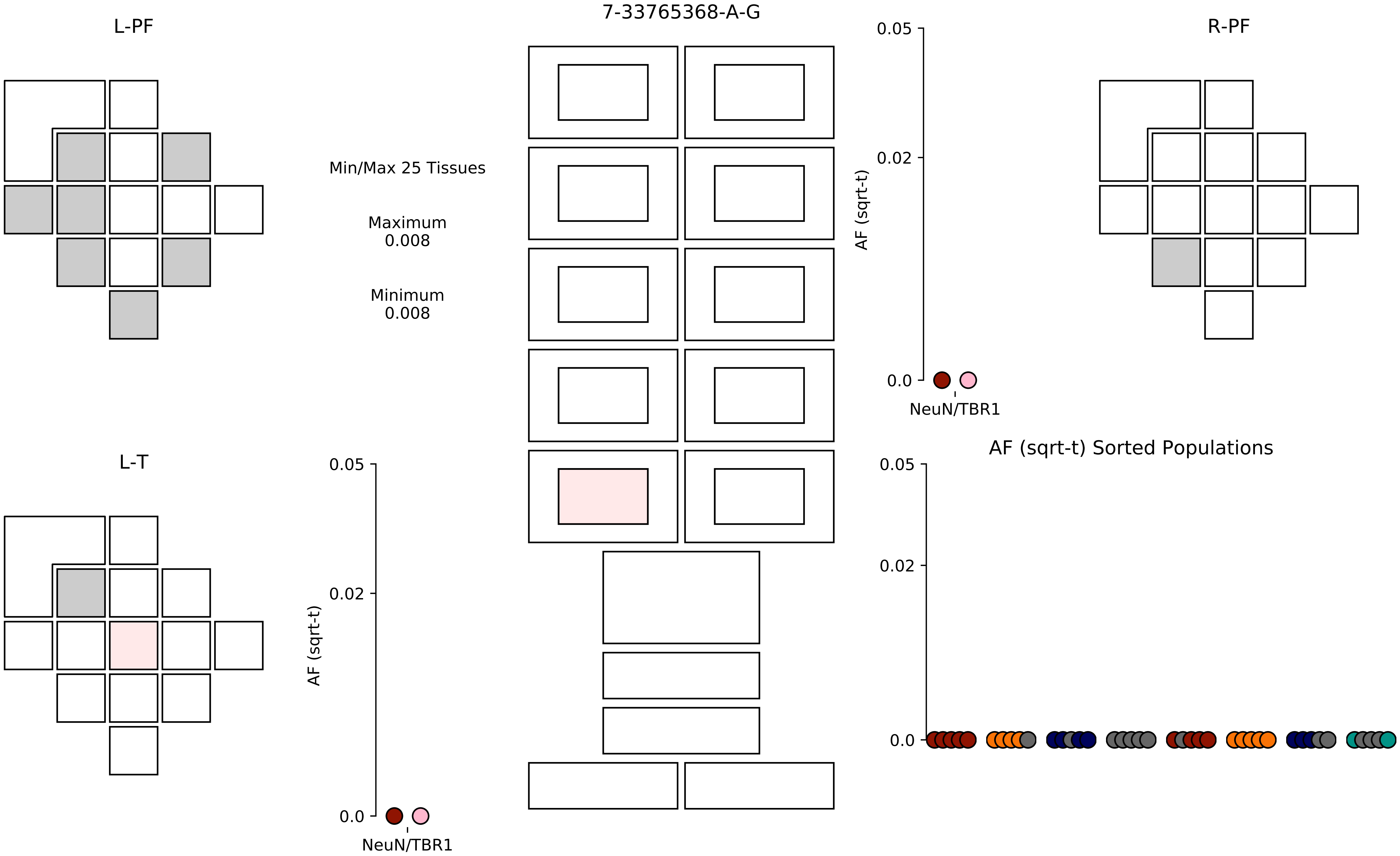
7-33765368-A-G
L-PF
R-PF
L-T
Min/Max 25 Tissues
Maximum
0.008
Minimum
0.008
AF (sqrt-t)
0.05
0.02
0.0
NeuN/TBR1
AF (sqrt-t)
0.05
0.02
0.0
NeuN/TBR1
AF (sqrt-t) Sorted Populations
0.05
0.02
0.0

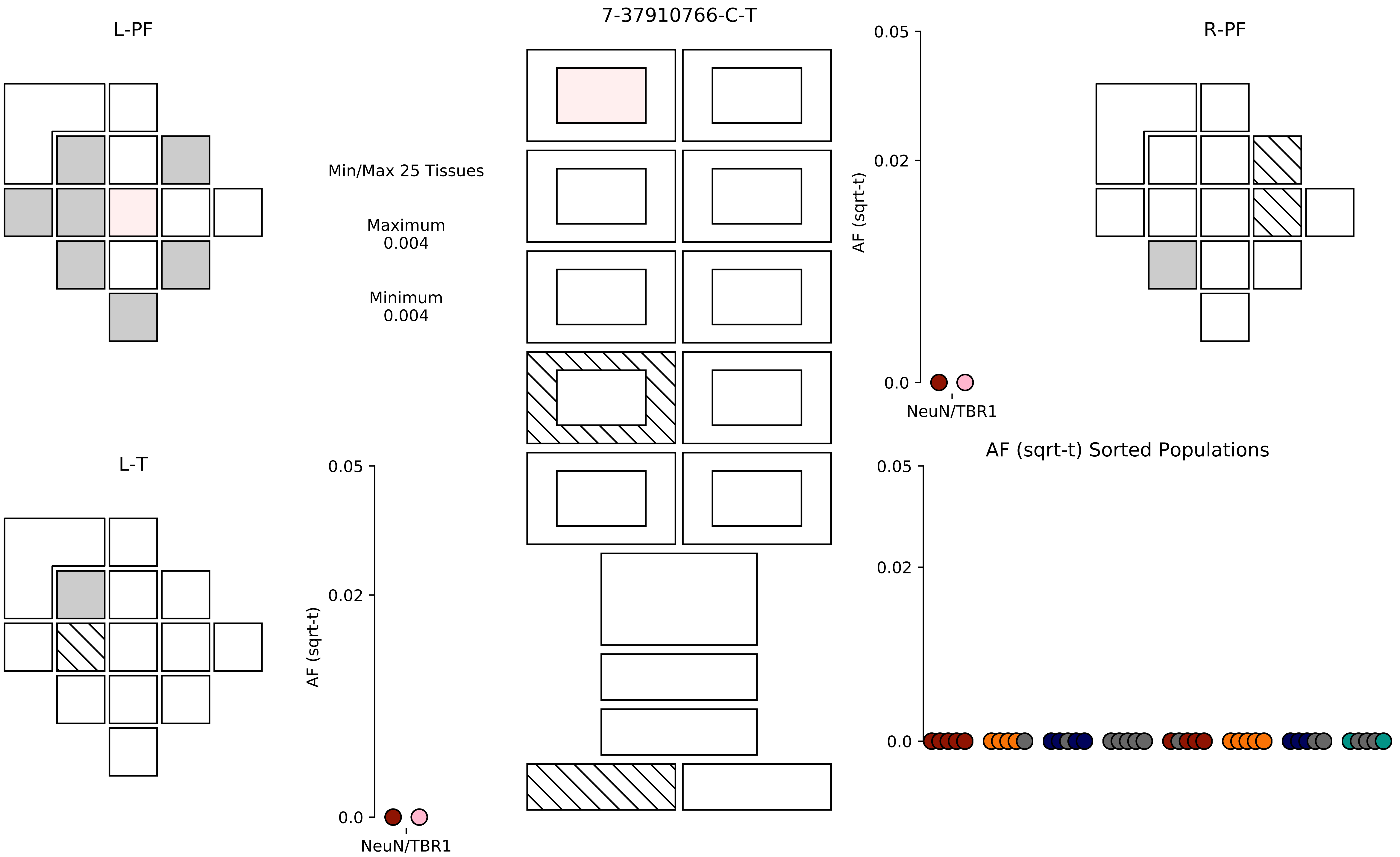
7-37910766-C-T
L-PF
R-PF
L-T
Min/Max 25 Tissues
Maximum
0.004
Minimum
0.004
AF (sqrt-t)
0.05
0.02
0.0
NeuN/TBR1
AF (sqrt-t)
0.05
0.02
0.0
NeuN/TBR1
AF (sqrt-t) Sorted Populations
0.05
0.02
0.0

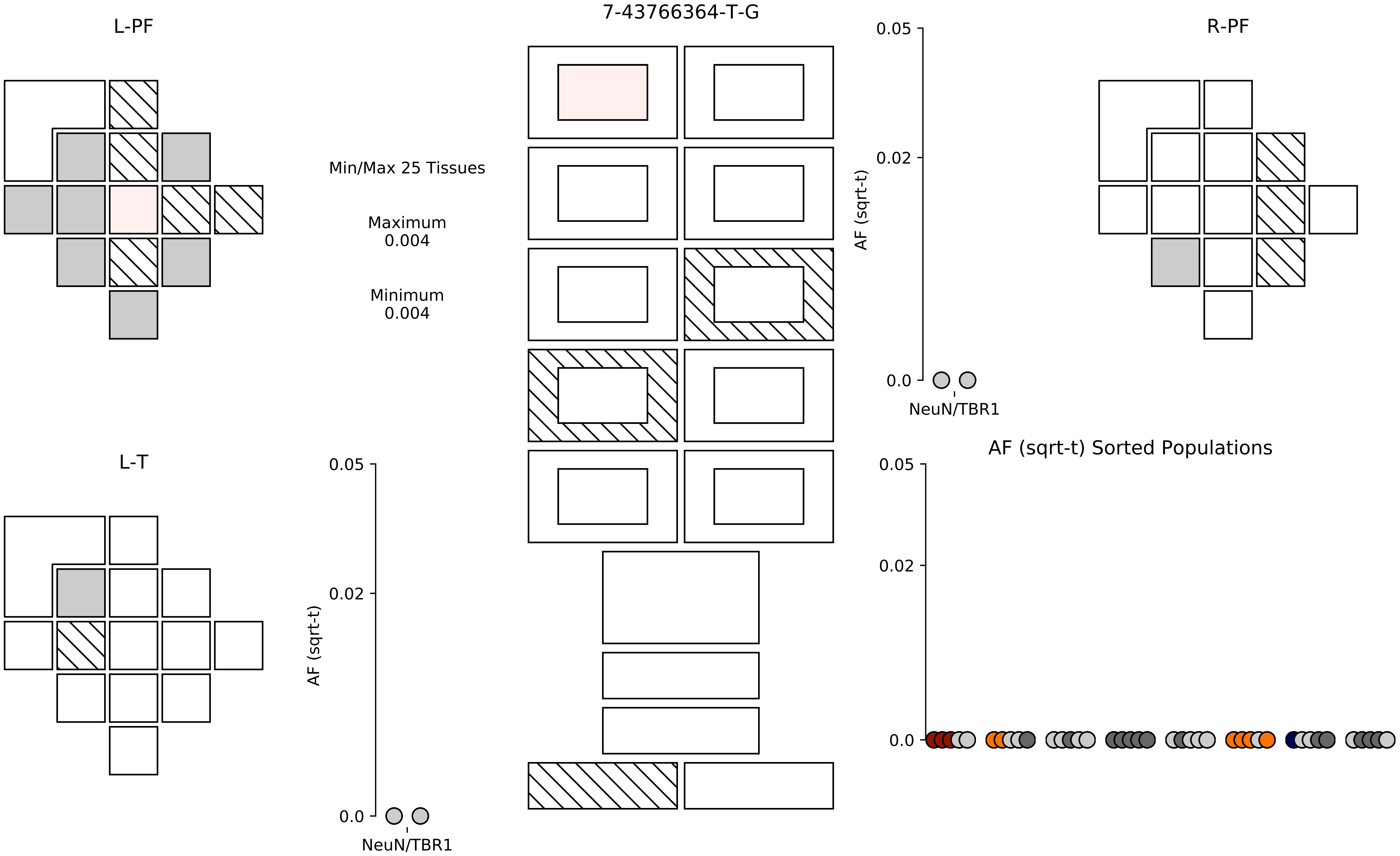
L-PF
7-43766364-T-G
R-PF
Min/Max 25 Tissues
Maximum
0.004
Minimum
0.004
0.05
0.02
0.0
AF (sqrt-t)
NeuN/TBR1
L-T
0.05
0.02
0.0
AF (sqrt-t)
NeuN/TBR1
AF (sqrt-t) Sorted Populations
0.05
0.02
0.0

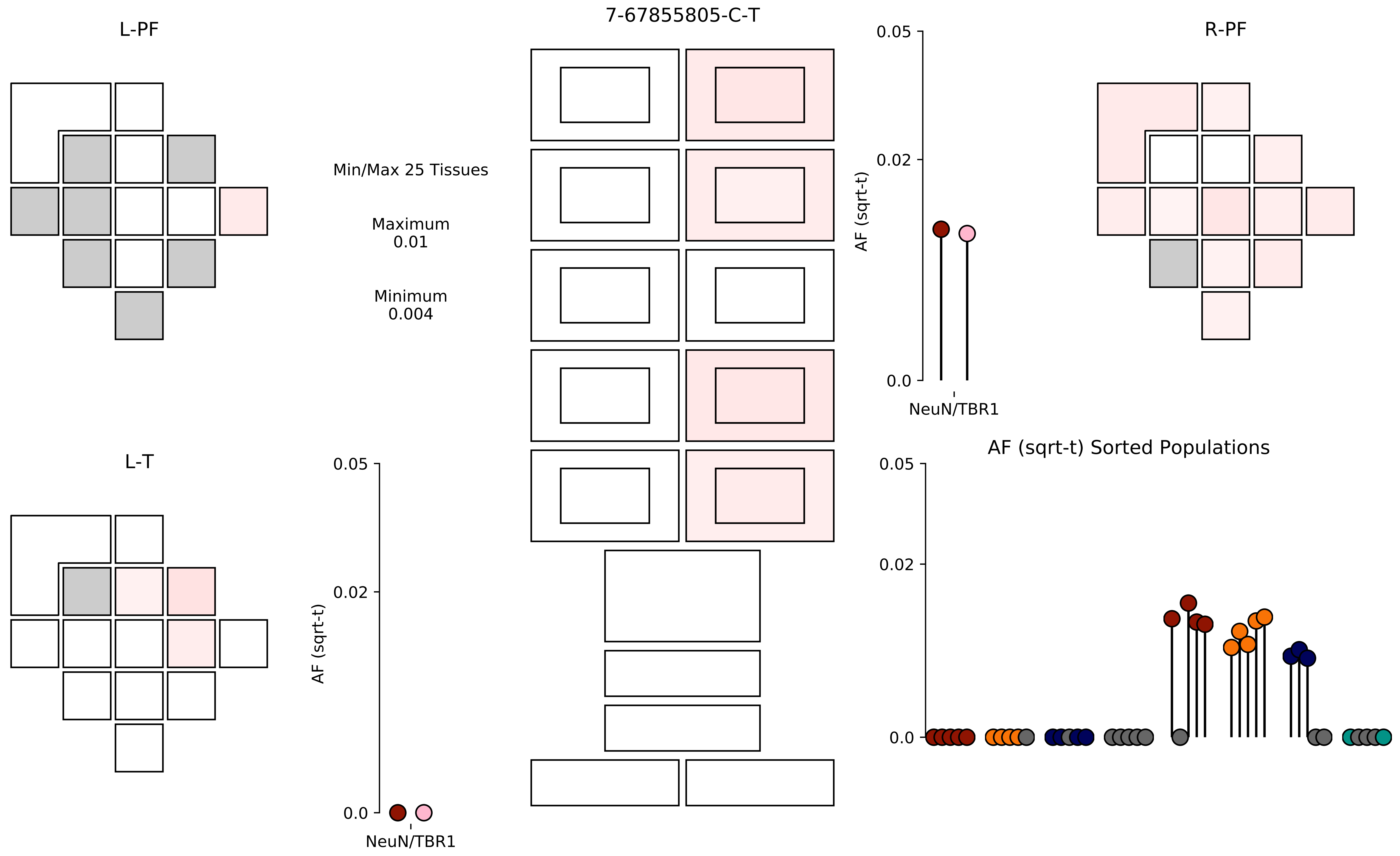

L-PF
7-67855805-C-T
R-PF
Min/Max 25 Tissues
Maximum
0.01
Minimum
0.004
0.05
0.02
0.0
AF (sqrt-t)
NeuN/TBR1
L-T
AF (sqrt-t) Sorted Populations

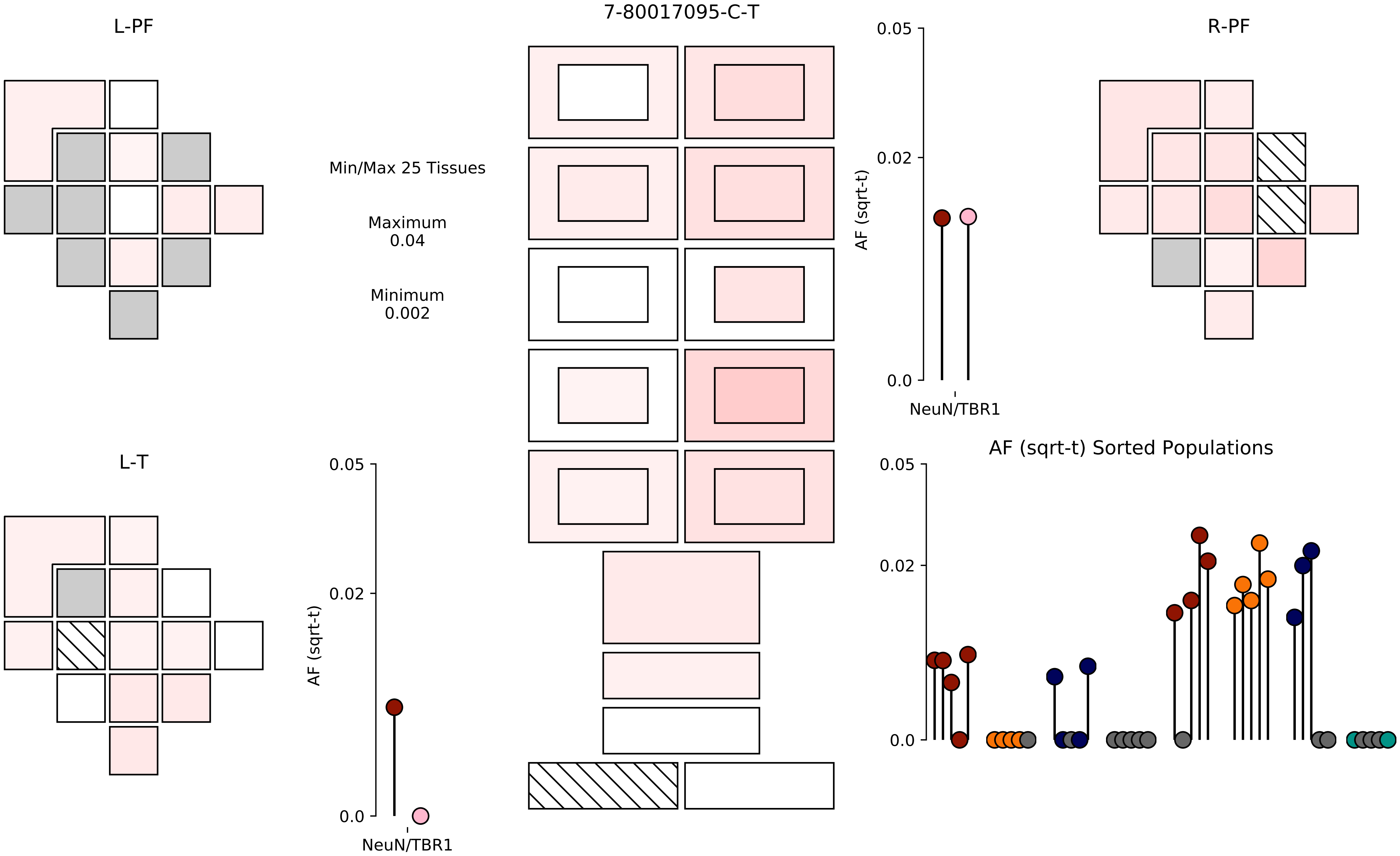
L-PF
7-80017095-C-T
R-PF
Min/Max 25 Tissues
Maximum
0.04
Minimum
0.002
0.05
0.02
0.0
AF (sqrt-t)
NeuN/TBR1
L-T
0.05
0.02
0.0
AF (sqrt-t)
NeuN/TBR1
AF (sqrt-t) Sorted Populations
0.05
0.02
0.0

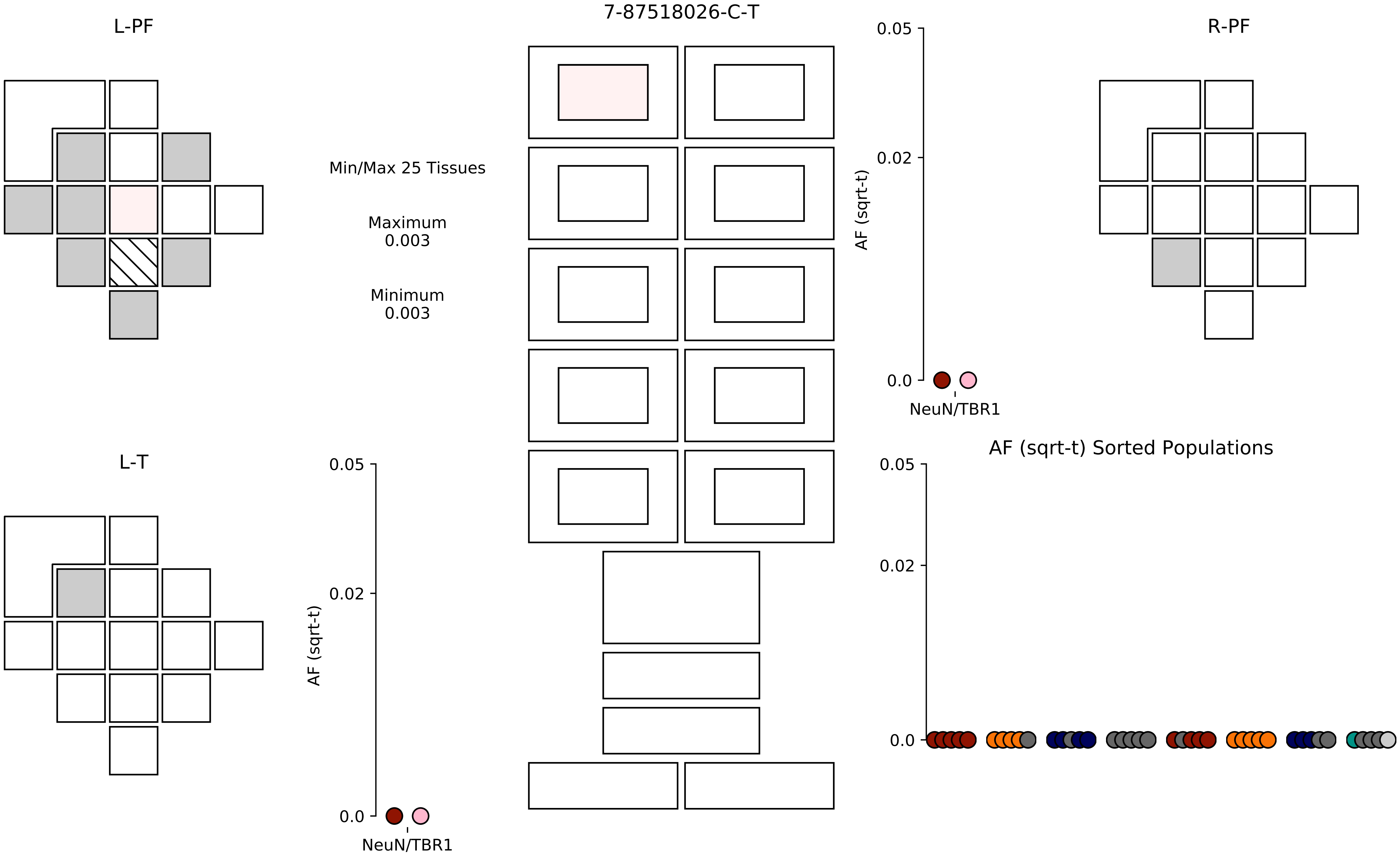

7-87518026-C-T
L-PF
R-PF
L-T
Min/Max 25 Tissues
Maximum
0.003
Minimum
0.003
AF (sqrt-t)
0.05
0.02
0.0
NeuN/TBR1
AF (sqrt-t) Sorted Populations

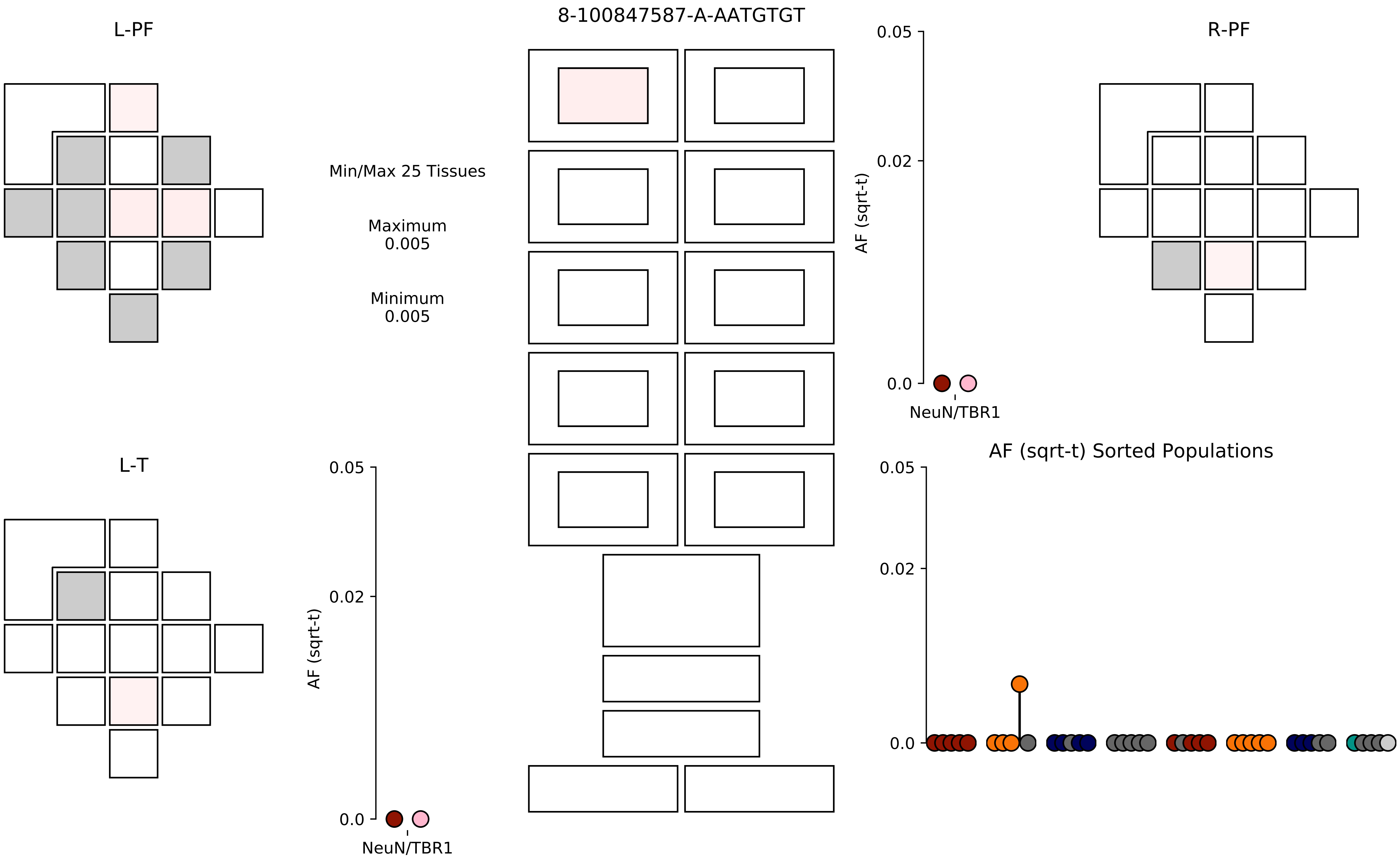

8-100847587-A-AATGTGT
L-PF
R-PF
L-T
Min/Max 25 Tissues
Maximum
0.005
Minimum
0.005
AF (sqrt-t)
0.05
0.02
0.0
NeuN/TBR1
AF (sqrt-t) Sorted Populations

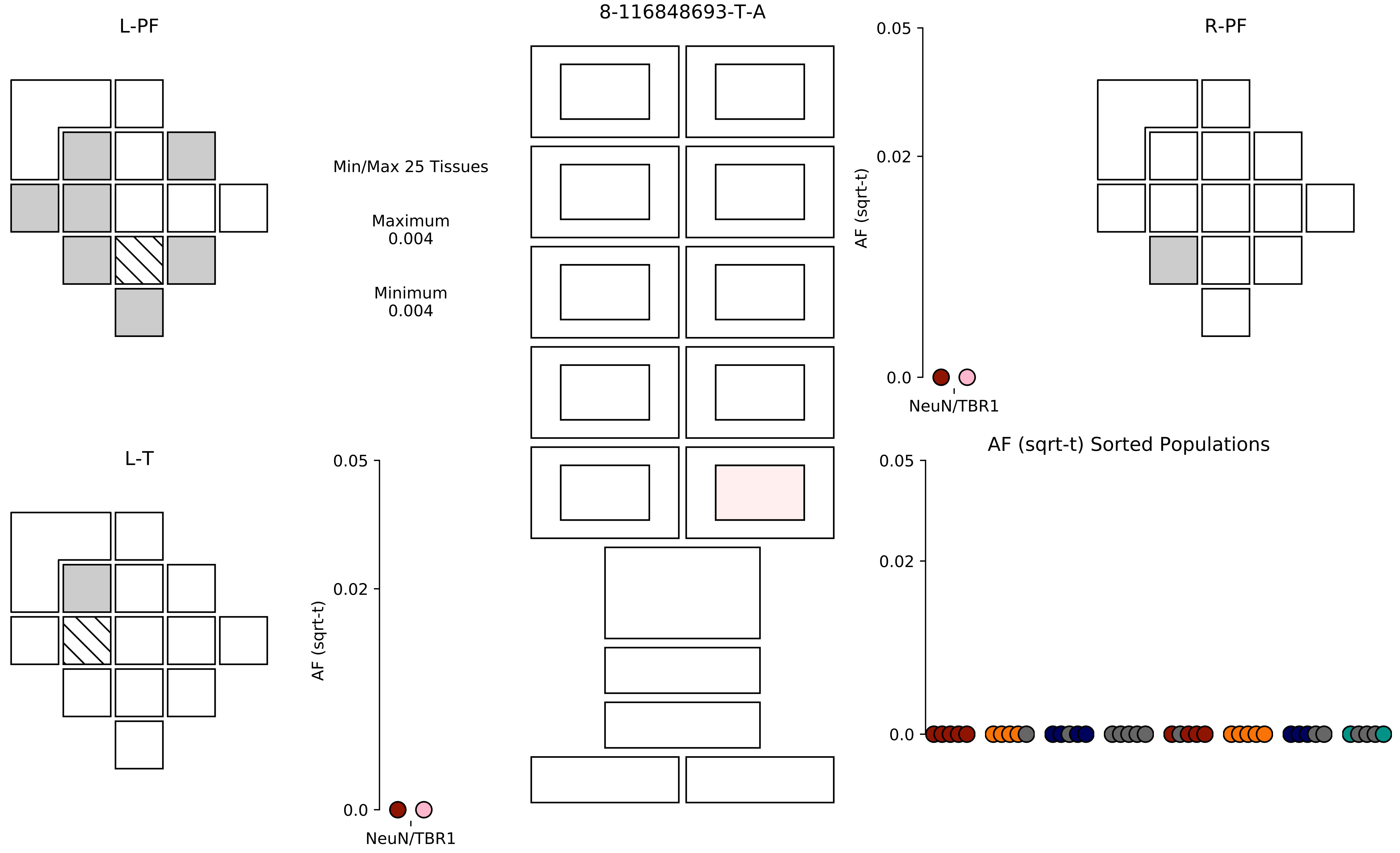

8-116848693-T-A
L-PF
R-PF
L-T
Min/Max 25 Tissues
Maximum
0.004
Minimum
0.004
AF (sqrt-t)
0.05
0.02
0.0
NeuN/TBR1
AF (sqrt-t)
0.05
0.02
0.0
NeuN/TBR1
AF (sqrt-t) Sorted Populations
0.05
0.02
0.0

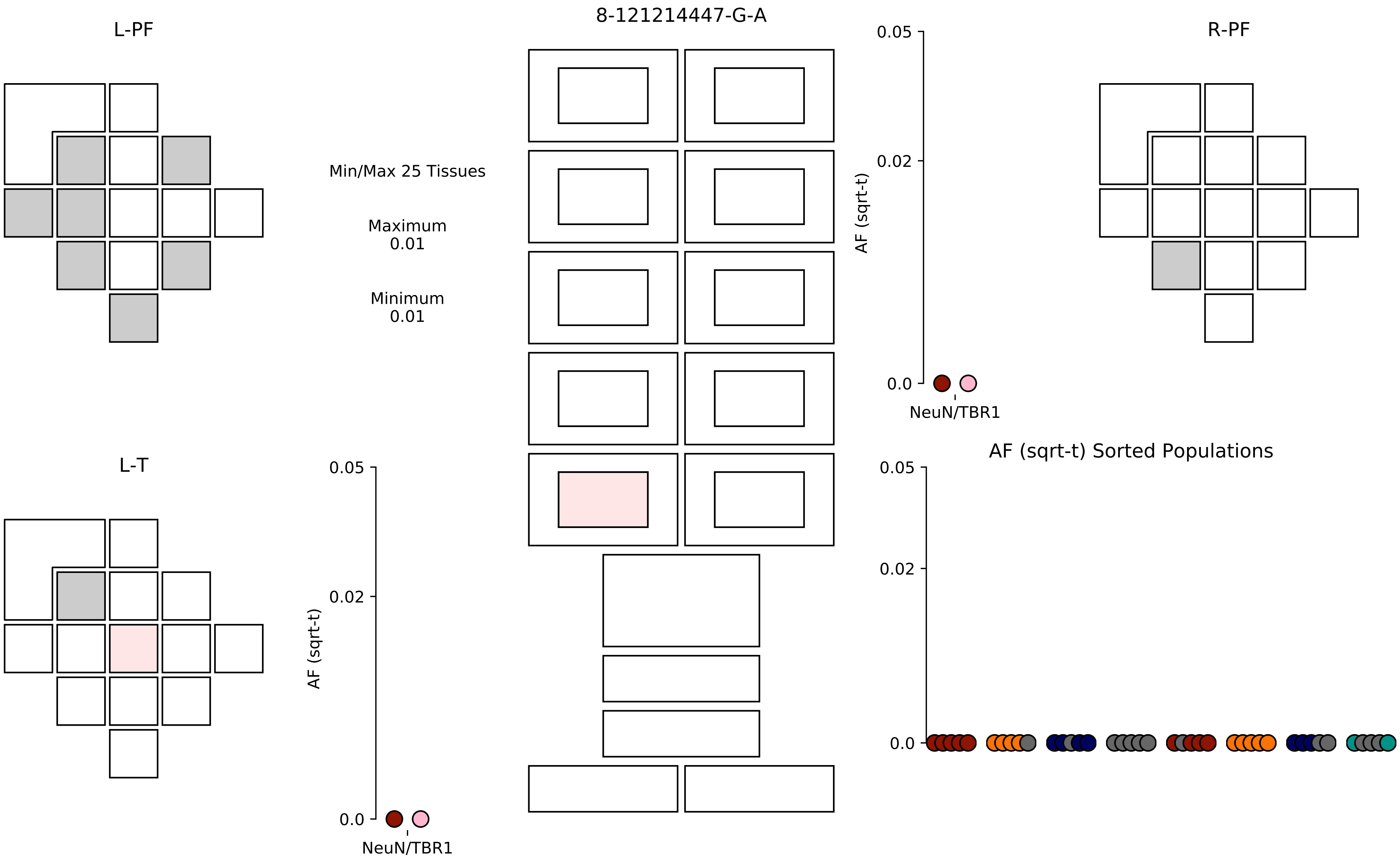
8-121214447-G-A
L-PF
R-PF
L-T
Min/Max 25 Tissues
Maximum
0.01
Minimum
0.01
AF (sqrt-t)
0.05
0.02
0.0
NeuN/TBR1
AF (sqrt-t) Sorted Populations

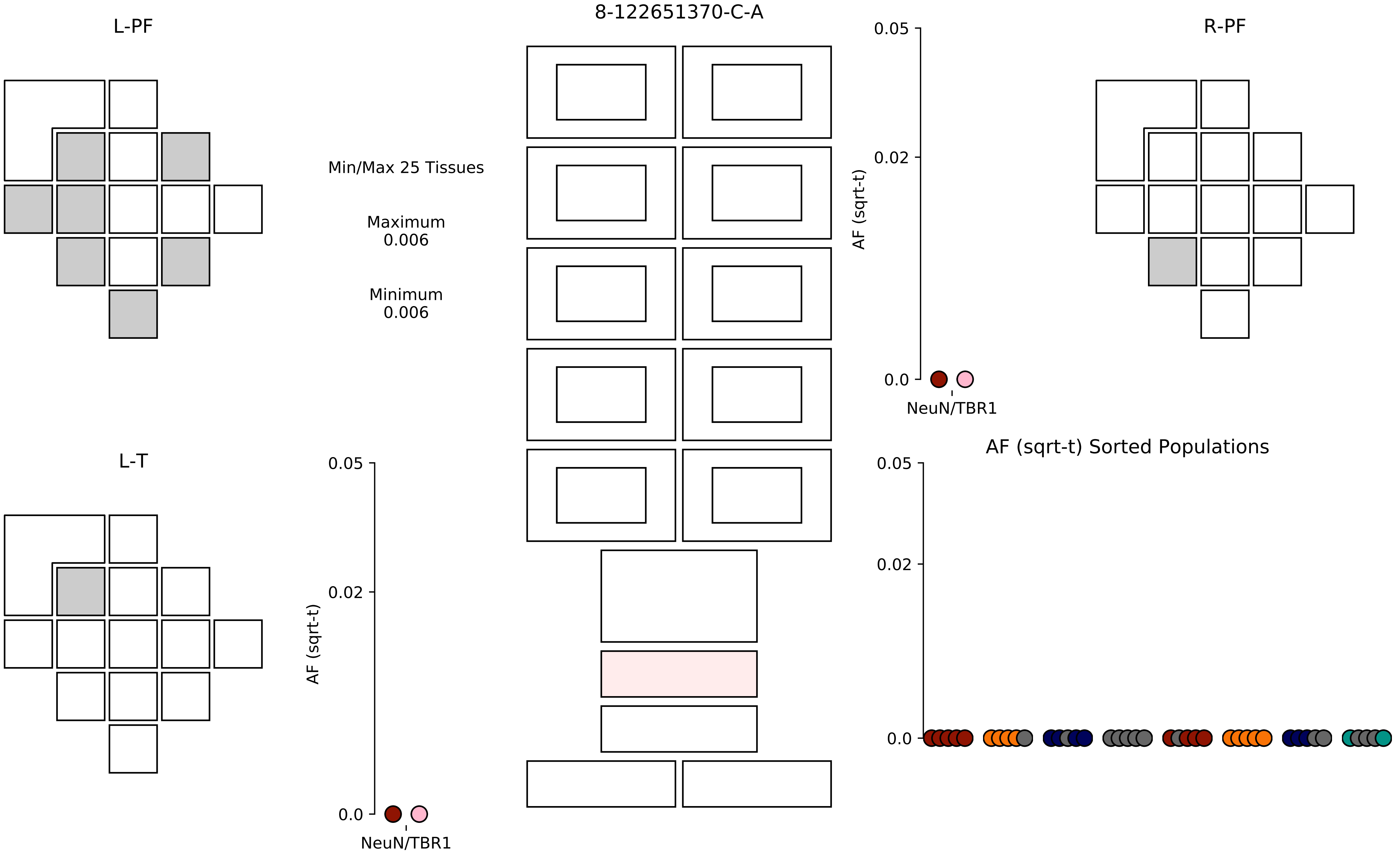

8-122651370-C-A
L-PF
R-PF
L-T
Min/Max 25 Tissues
Maximum
0.006
Minimum
0.006
0.05
0.02
0.0
AF (sqrt-t)
NeuN/TBR1
AF (sqrt-t) Sorted Populations

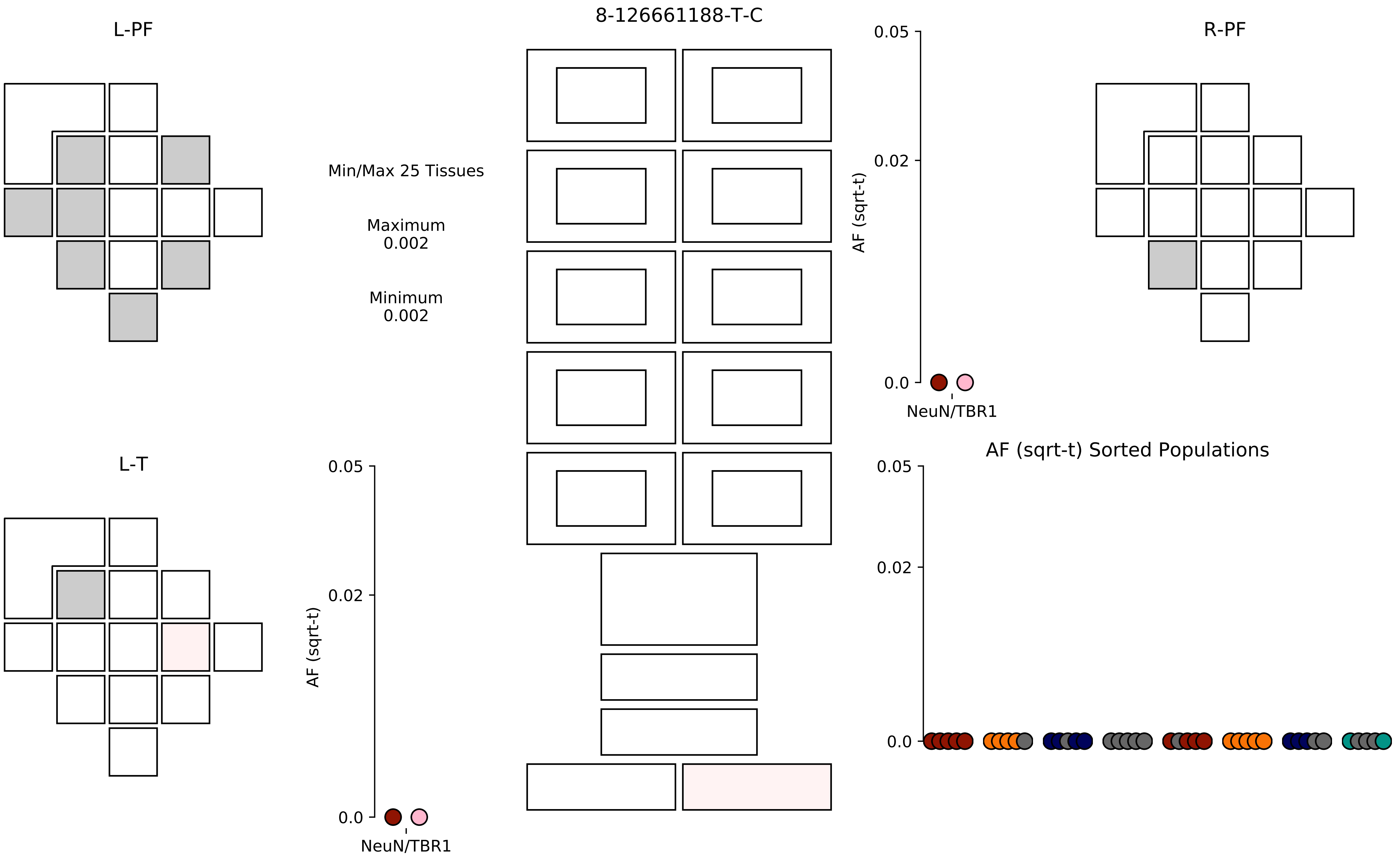

8-126661188-T-C
L-PF
R-PF
L-T
Min/Max 25 Tissues
Maximum
0.002
Minimum
0.002
AF (sqrt-t)
0.05
0.02
0.0
NeuN/TBR1
AF (sqrt-t)
0.05
0.02
0.0
NeuN/TBR1
AF (sqrt-t) Sorted Populations
0.05
0.02
0.0

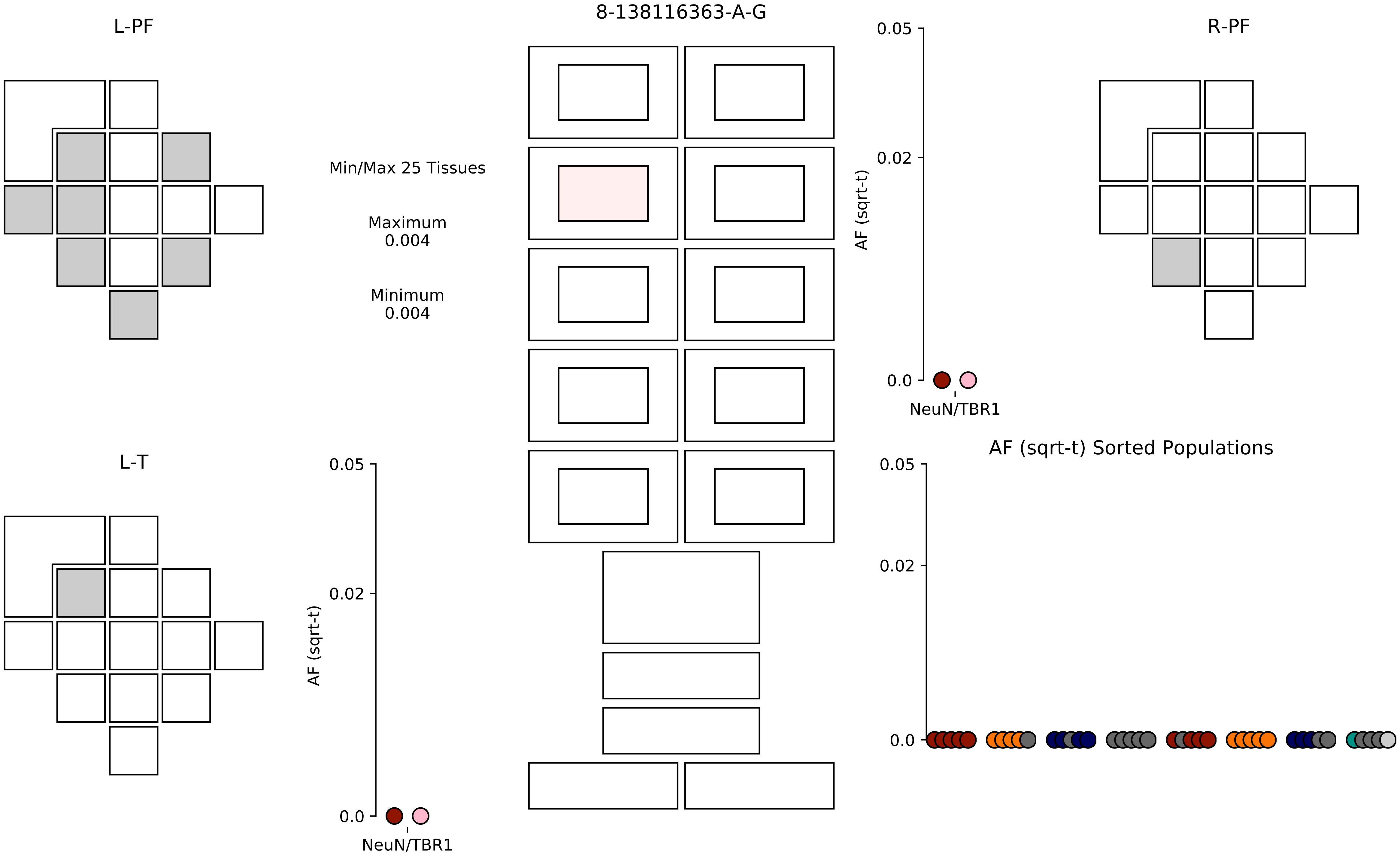
8-138116363-A-G
L-PF
R-PF
L-T
Min/Max 25 Tissues
Maximum
0.004
Minimum
0.004
AF (sqrt-t)
0.05
0.02
0.0
NeuN/TBR1
AF (sqrt-t)
0.05
0.02
0.0
NeuN/TBR1
AF (sqrt-t) Sorted Populations
0.05
0.02
0.0

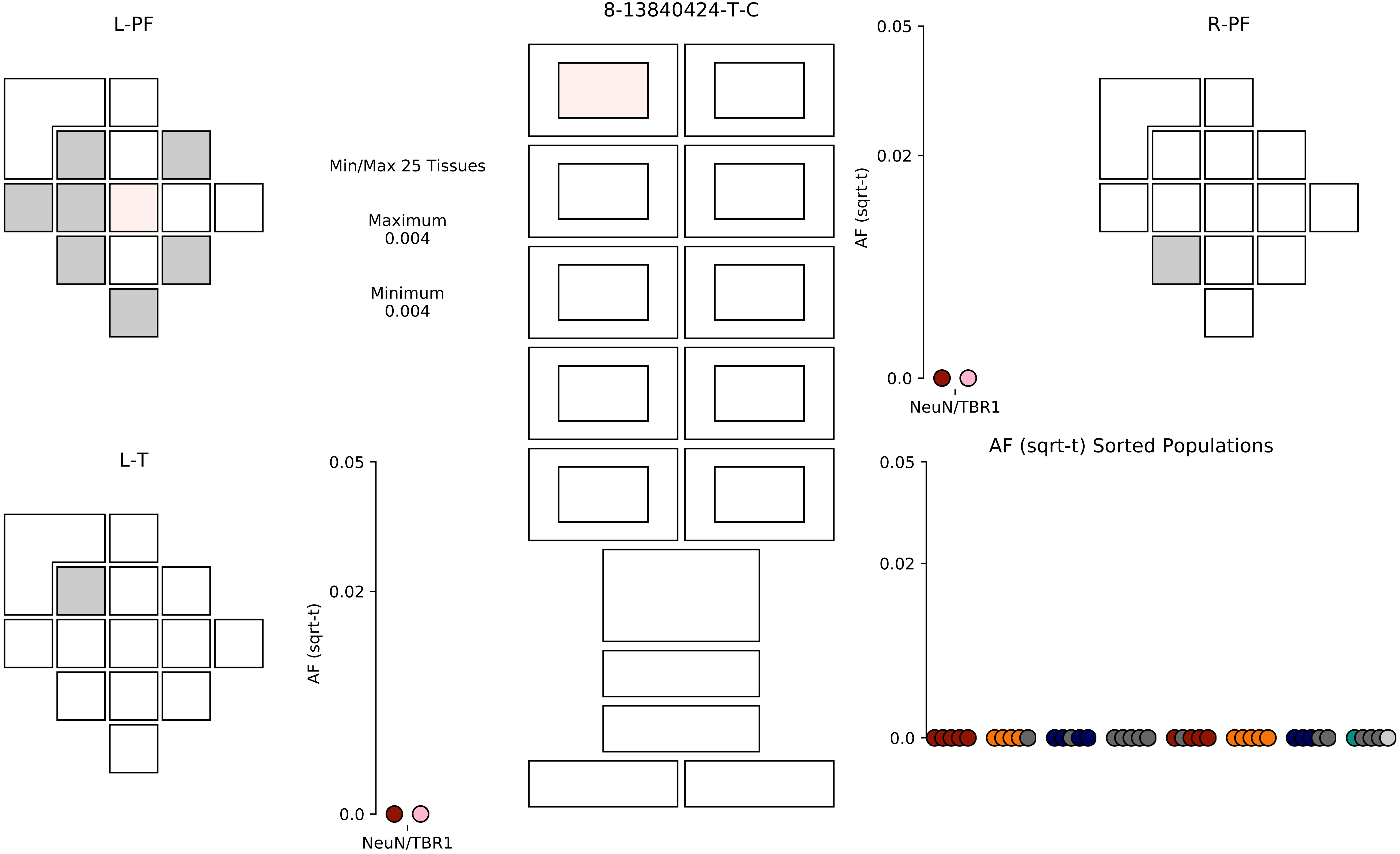

L-PF
8-13840424-T-C
R-PF
Min/Max 25 Tissues
Maximum
0.004
Minimum
0.004
0.05
0.02
0.0
AF (sqrt-t)
NeuN/TBR1
L-T
AF (sqrt-t) Sorted Populations

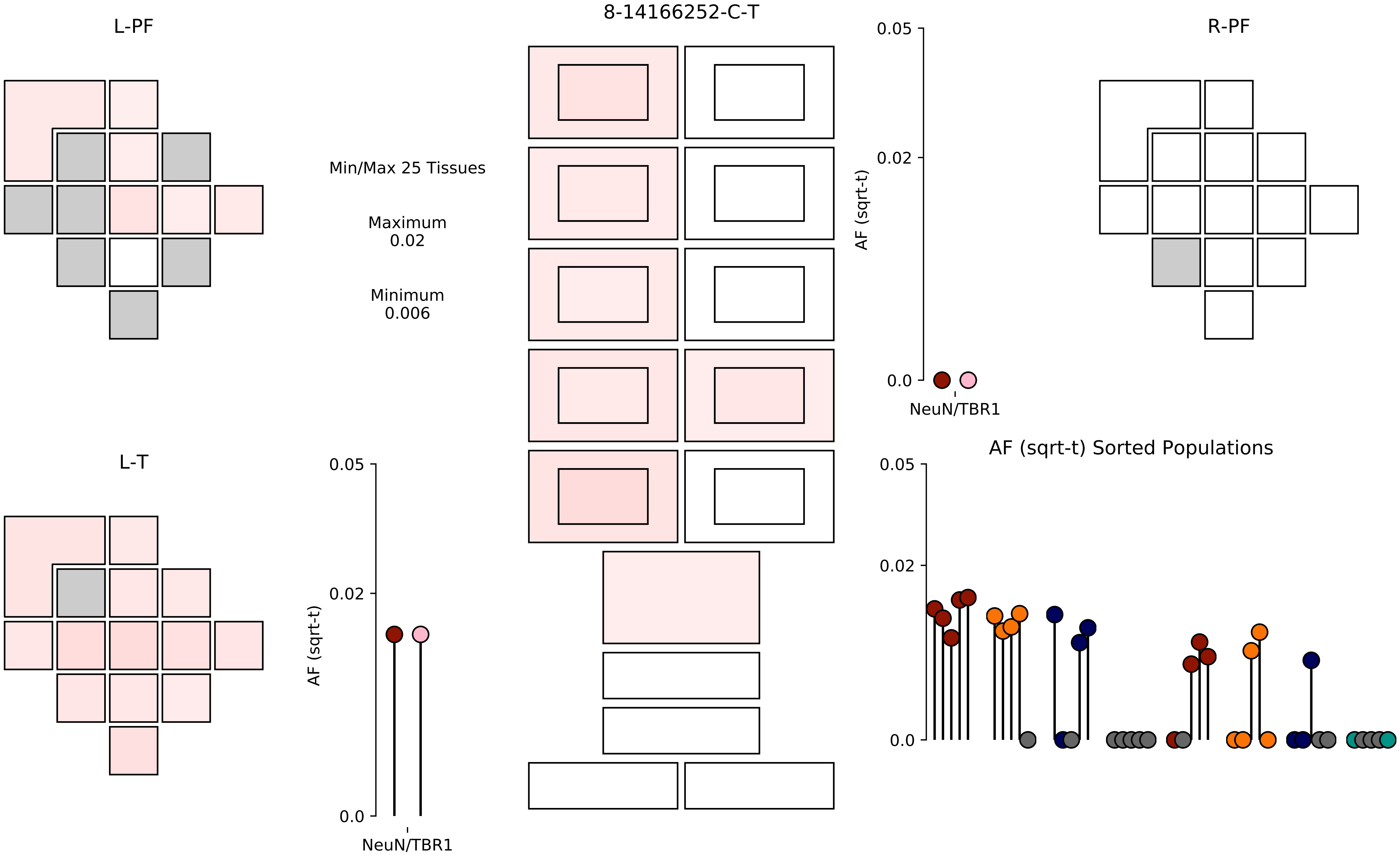

L-PF
8-14166252-C-T
R-PF
Min/Max 25 Tissues
Maximum
0.02
Minimum
0.006
0.05
0.02
0.0
AF (sqrt-t)
NeuN/TBR1
L-T
0.05
0.02
0.0
AF (sqrt-t)
NeuN/TBR1
AF (sqrt-t) Sorted Populations
0.05
0.02
0.0

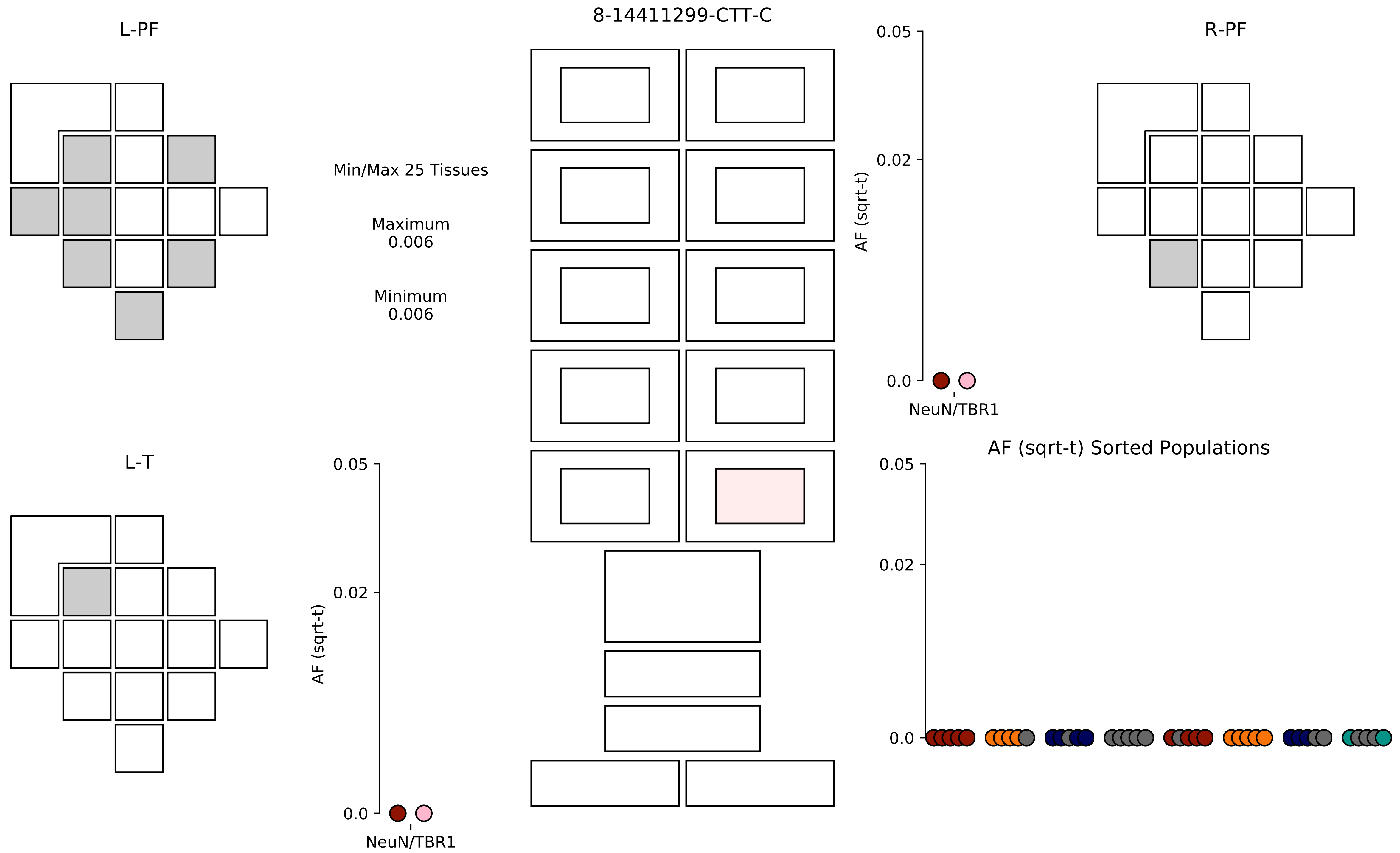
8-14411299-CTT-C
L-PF
R-PF
L-T
Min/Max 25 Tissues
Maximum
0.006
Minimum
0.006
AF (sqrt-t)
0.05
0.02
0.0
NeuN/TBR1
AF (sqrt-t) Sorted Populations

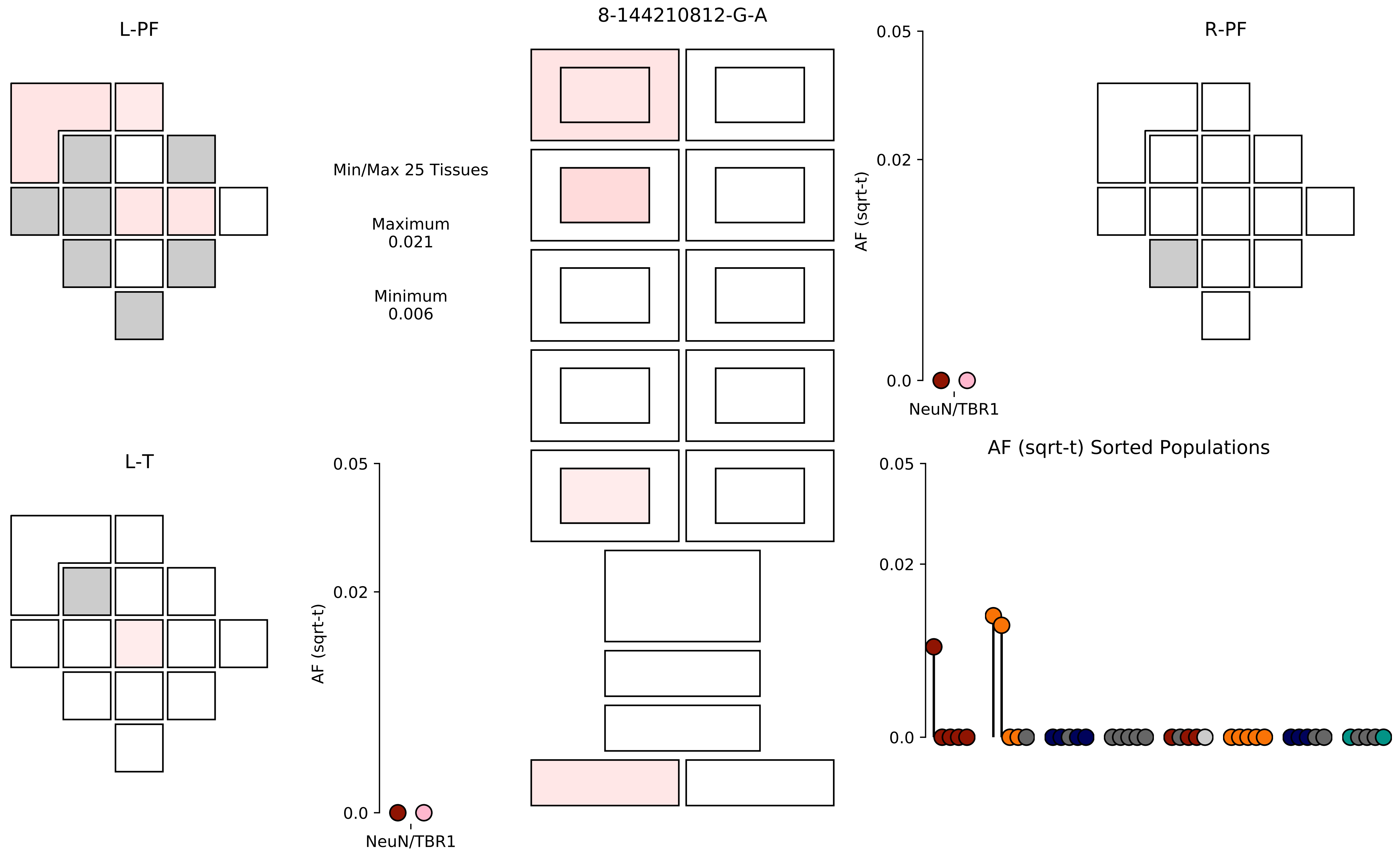
8-144210812-G-A
L-PF
R-PF
L-T
Min/Max 25 Tissues
Maximum
0.021
Minimum
0.006
AF (sqrt-t)
0.05
0.02
0.0
NeuN/TBR1
AF (sqrt-t)
0.05
0.02
0.0
NeuN/TBR1
AF (sqrt-t) Sorted Populations
0.05
0.02
0.0

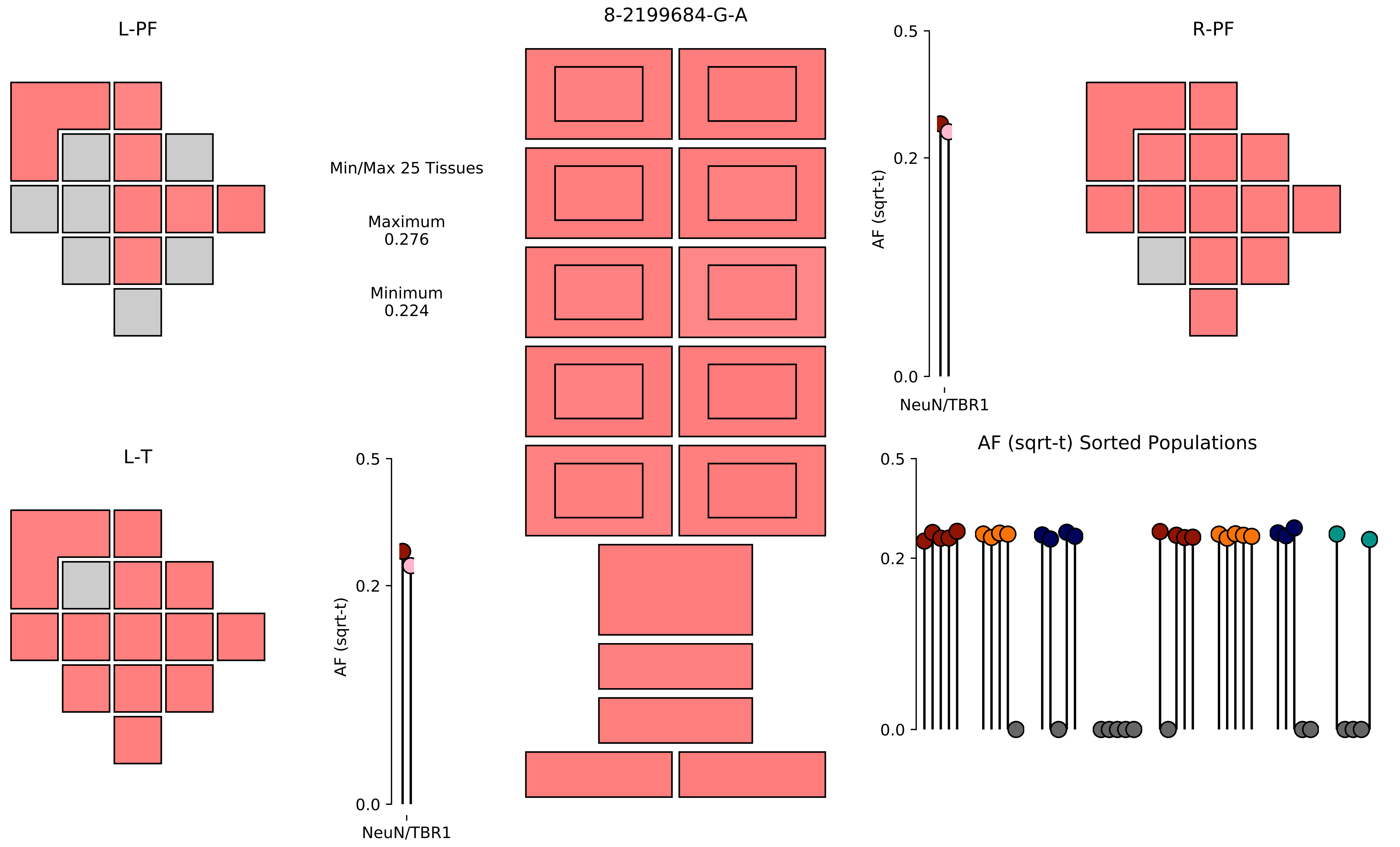
8-2199684-G-A
L-PF
R-PF
L-T
Min/Max 25 Tissues
Maximum
0.276
Minimum
0.224
AF (sqrt-t)
0.5
0.2
0.0
NeuN/TBR1
AF (sqrt-t) Sorted Populations

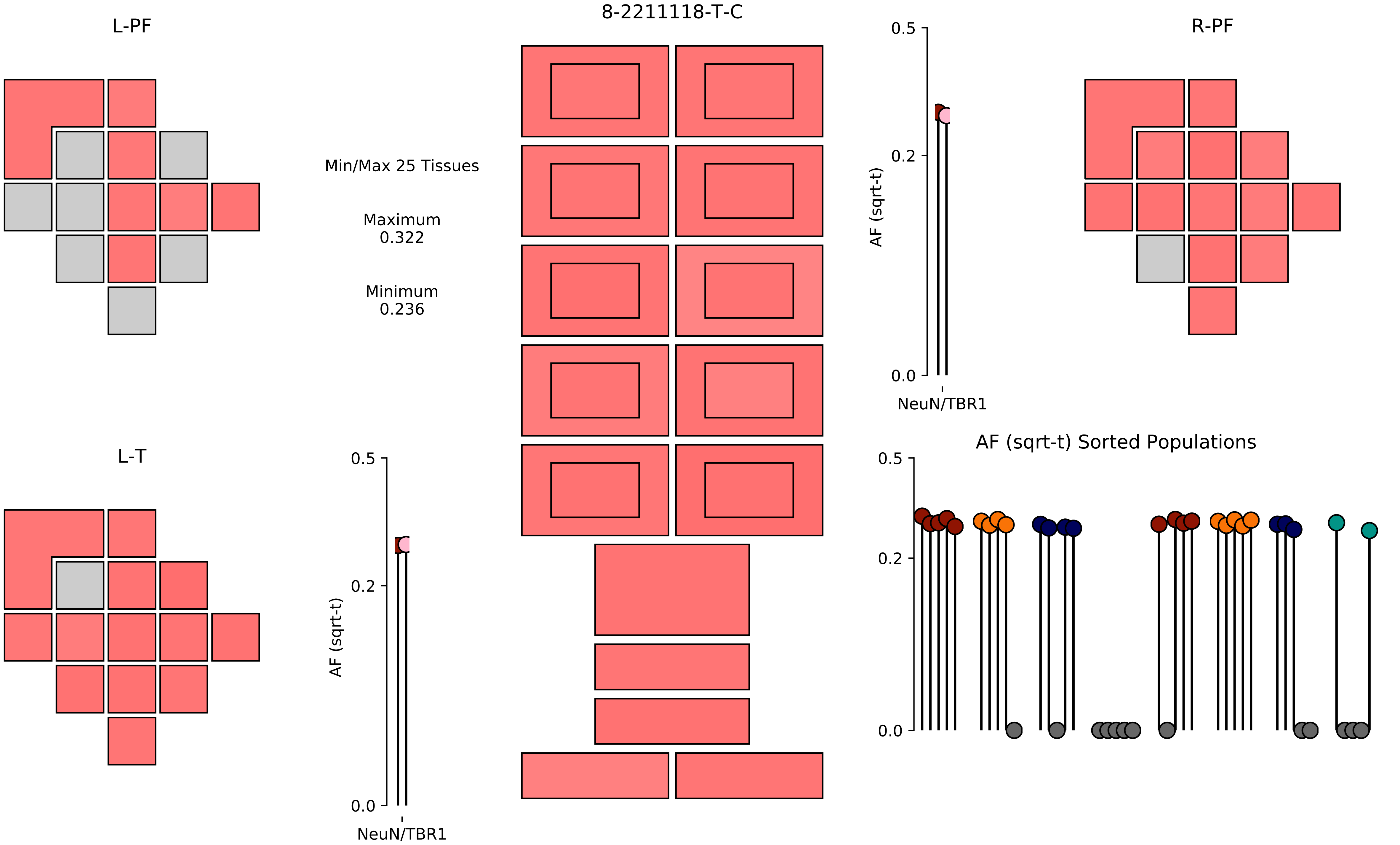
8-2211118-T-C
L-PF
R-PF
L-T
Min/Max 25 Tissues
Maximum
0.322
Minimum
0.236
AF (sqrt-t)
0.5
0.2
0.0
NeuN/TBR1
AF (sqrt-t) Sorted Populations

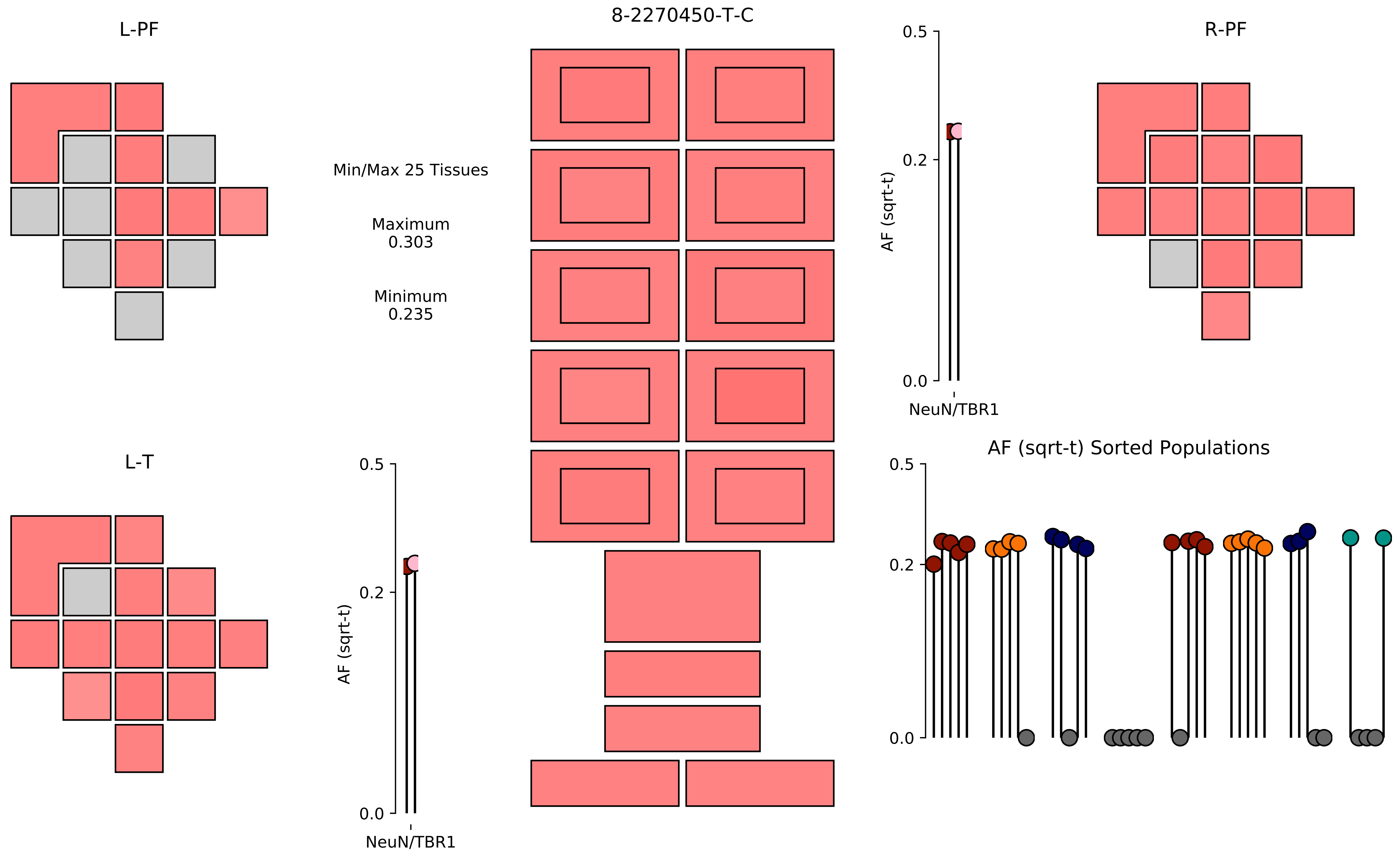

8-2270450-T-C
L-PF
R-PF
L-T
Min/Max 25 Tissues
Maximum
0.303
Minimum
0.235
AF (sqrt-t)
0.5
0.2
0.0
NeuN/TBR1
AF (sqrt-t) Sorted Populations

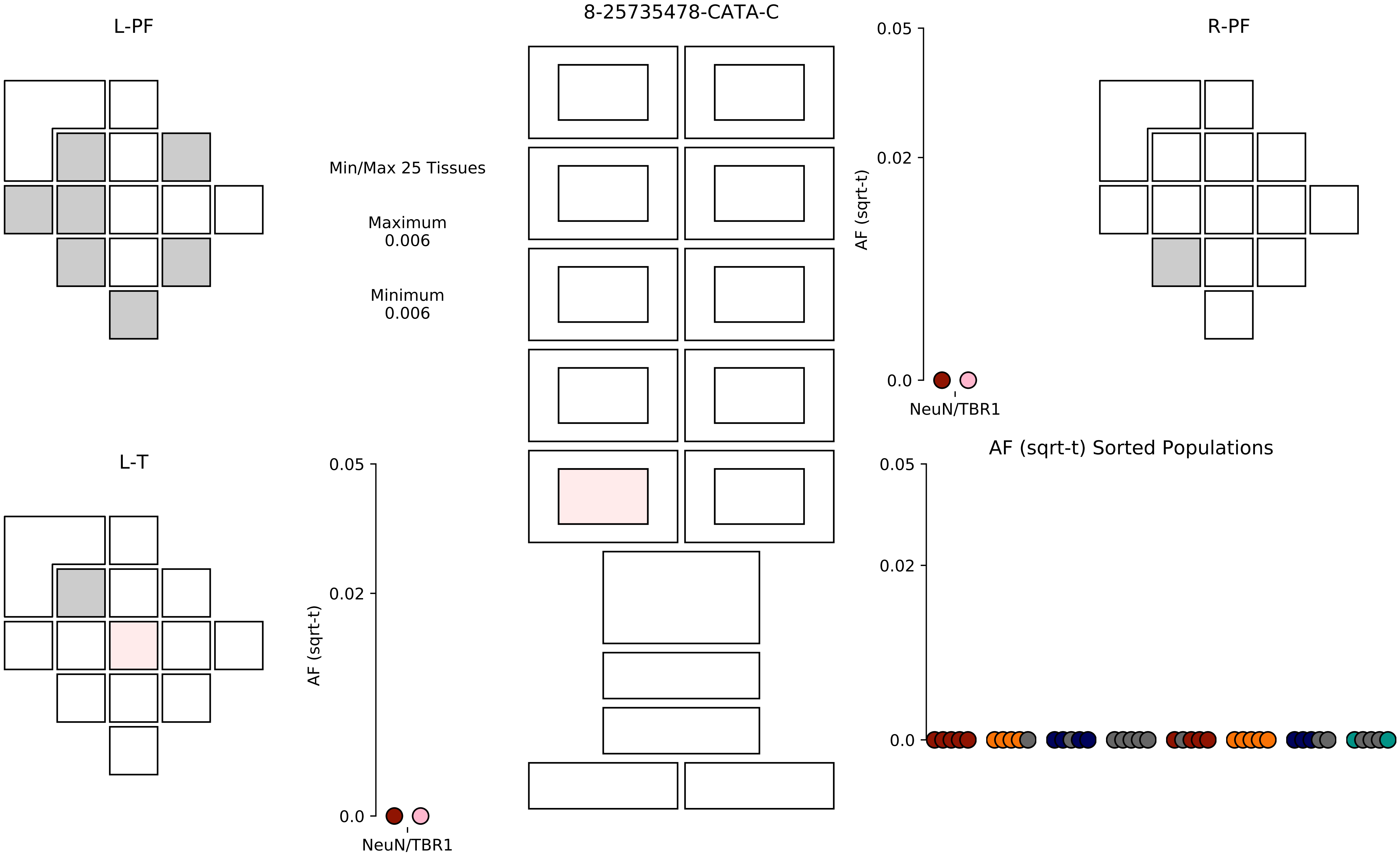

8-25735478-CATA-C
L-PF
R-PF
L-T
Min/Max 25 Tissues
Maximum
0.006
Minimum
0.006
AF (sqrt-t)
0.05
0.02
0.0
NeuN/TBR1
AF (sqrt-t)
0.05
0.02
0.0
NeuN/TBR1
AF (sqrt-t) Sorted Populations
0.05
0.02
0.0

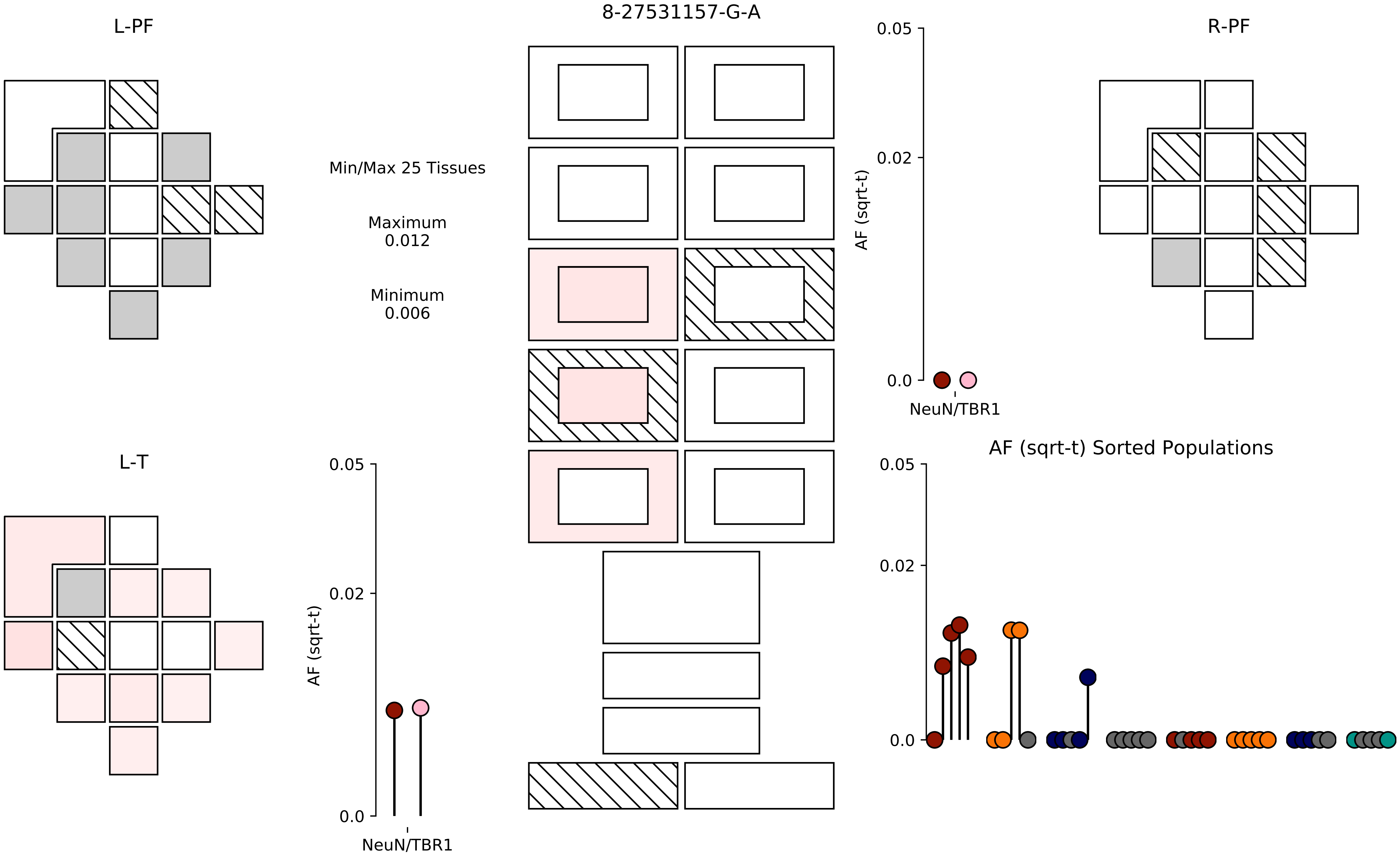

L-PF
8-27531157-G-A
R-PF
Min/Max 25 Tissues
Maximum
0.012
Minimum
0.006
0.05
0.02
0.0
AF (sqrt-t)
NeuN/TBR1
L-T
AF (sqrt-t) Sorted Populations

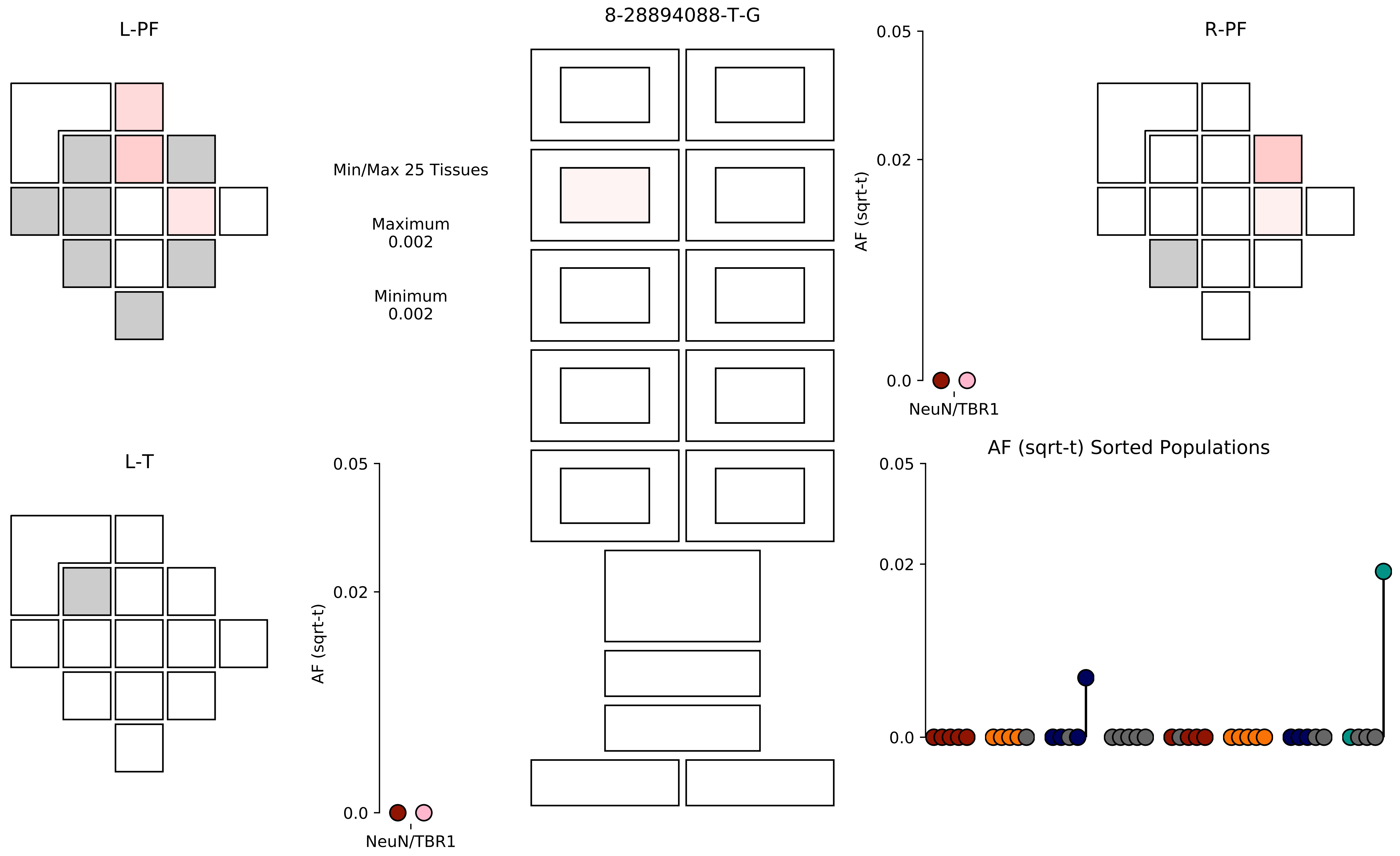

8-28894088-T-G
L-PF
R-PF
L-T
Min/Max 25 Tissues
Maximum
0.002
Minimum
0.002
AF (sqrt-t)
0.05
0.02
0.0
NeuN/TBR1
AF (sqrt-t) Sorted Populations

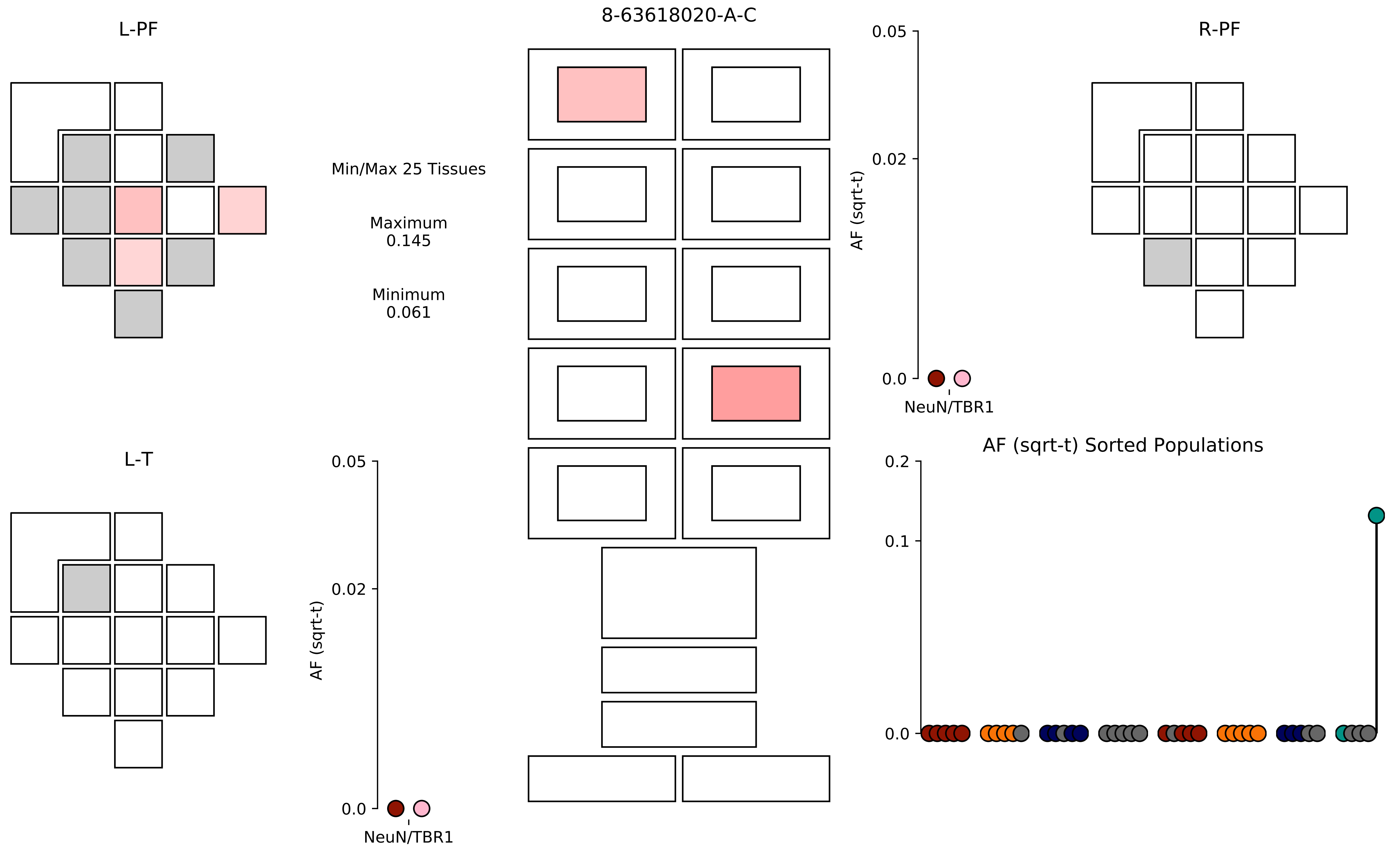

8-63618020-A-C
L-PF
R-PF
L-T
Min/Max 25 Tissues
Maximum
0.145
Minimum
0.061
AF (sqrt-t)
0.05
0.02
0.0
NeuN/TBR1
AF (sqrt-t) Sorted Populations
0.2
0.1
0.0

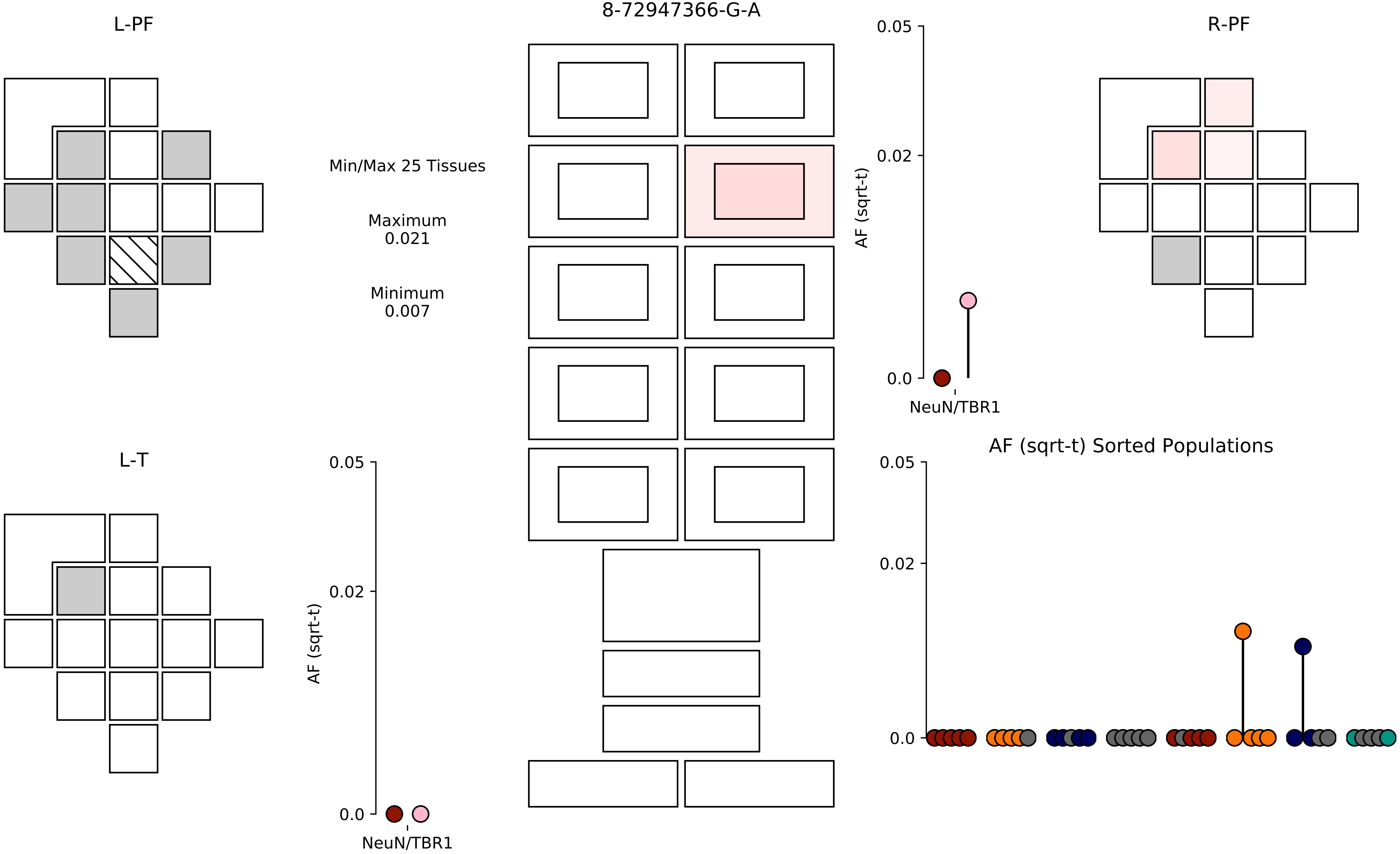

8-72947366-G-A
L-PF
R-PF
L-T
Min/Max 25 Tissues
Maximum
0.021
Minimum
0.007
AF (sqrt-t)
0.05
0.02
0.0
NeuN/TBR1
AF (sqrt-t)
0.05
0.02
0.0
NeuN/TBR1
AF (sqrt-t) Sorted Populations
0.05
0.02
0.0

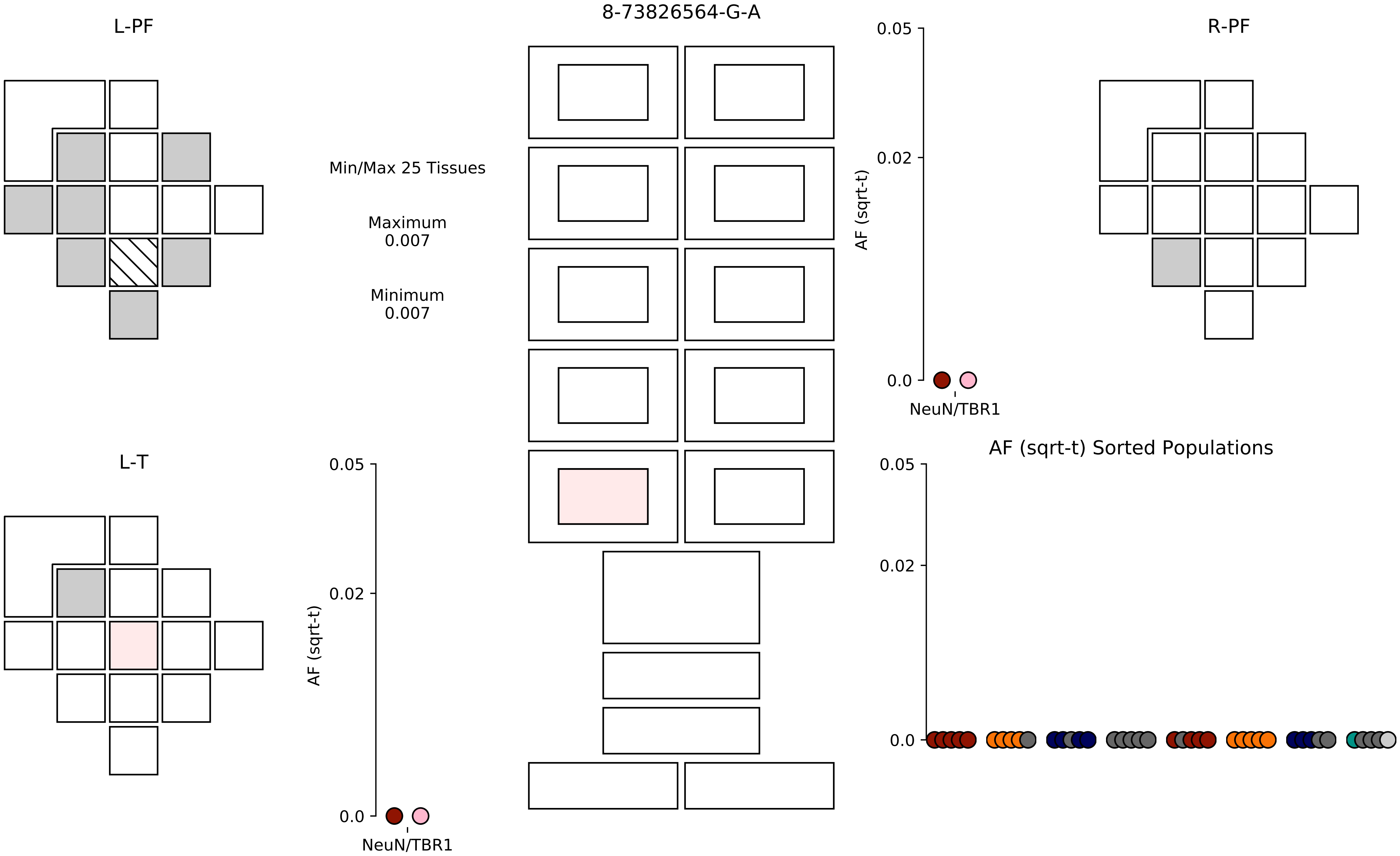

8-73826564-G-A
L-PF
R-PF
L-T
Min/Max 25 Tissues
Maximum
0.007
Minimum
0.007
AF (sqrt-t)
0.05
0.02
0.0
NeuN/TBR1
AF (sqrt-t) Sorted Populations

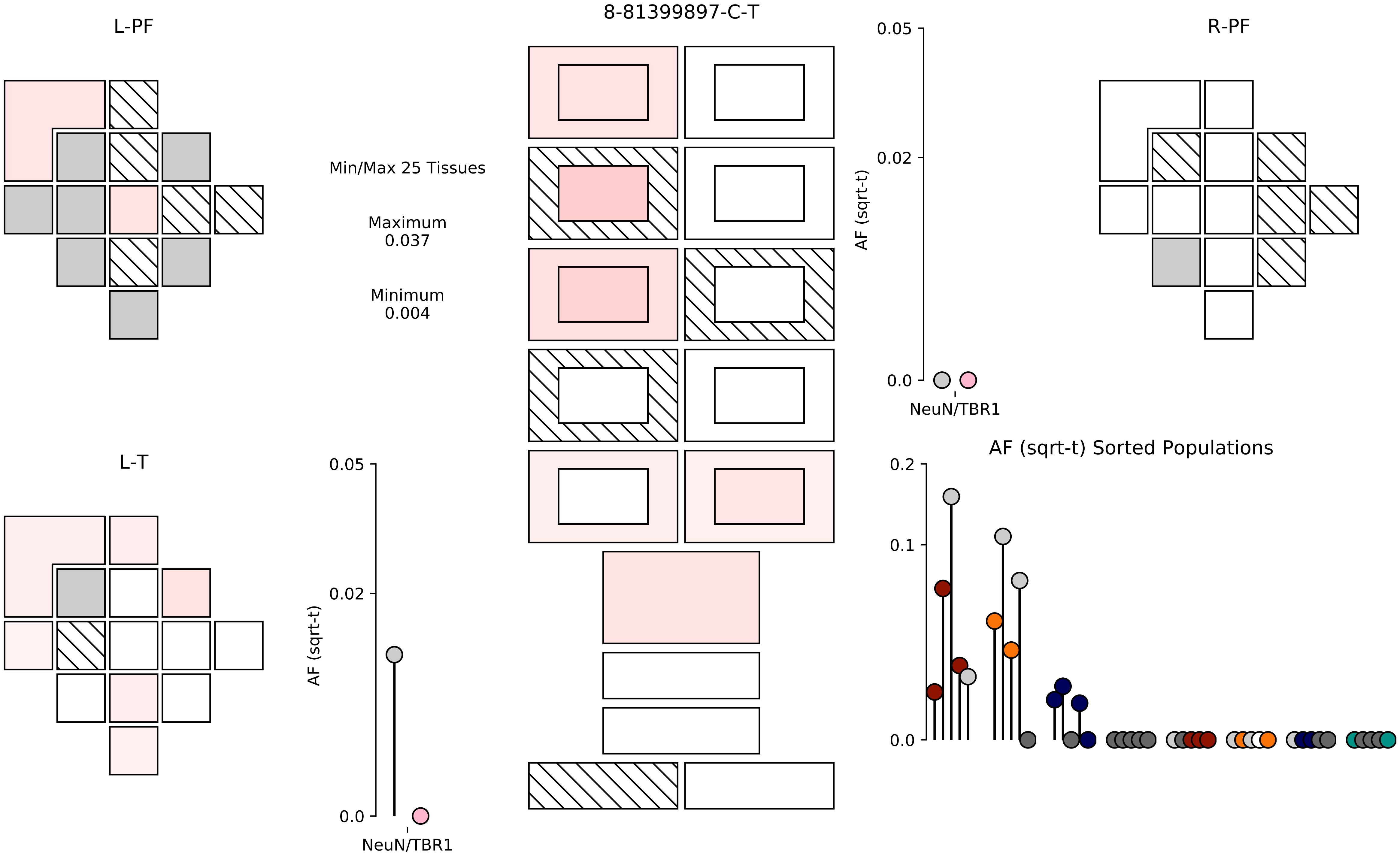

8-81399897-C-T
L-PF
R-PF
L-T
Min/Max 25 Tissues
Maximum
0.037
Minimum
0.004
AF (sqrt-t)
0.05
0.02
0.0
NeuN/TBR1
AF (sqrt-t) Sorted Populations
0.2
0.1
0.0

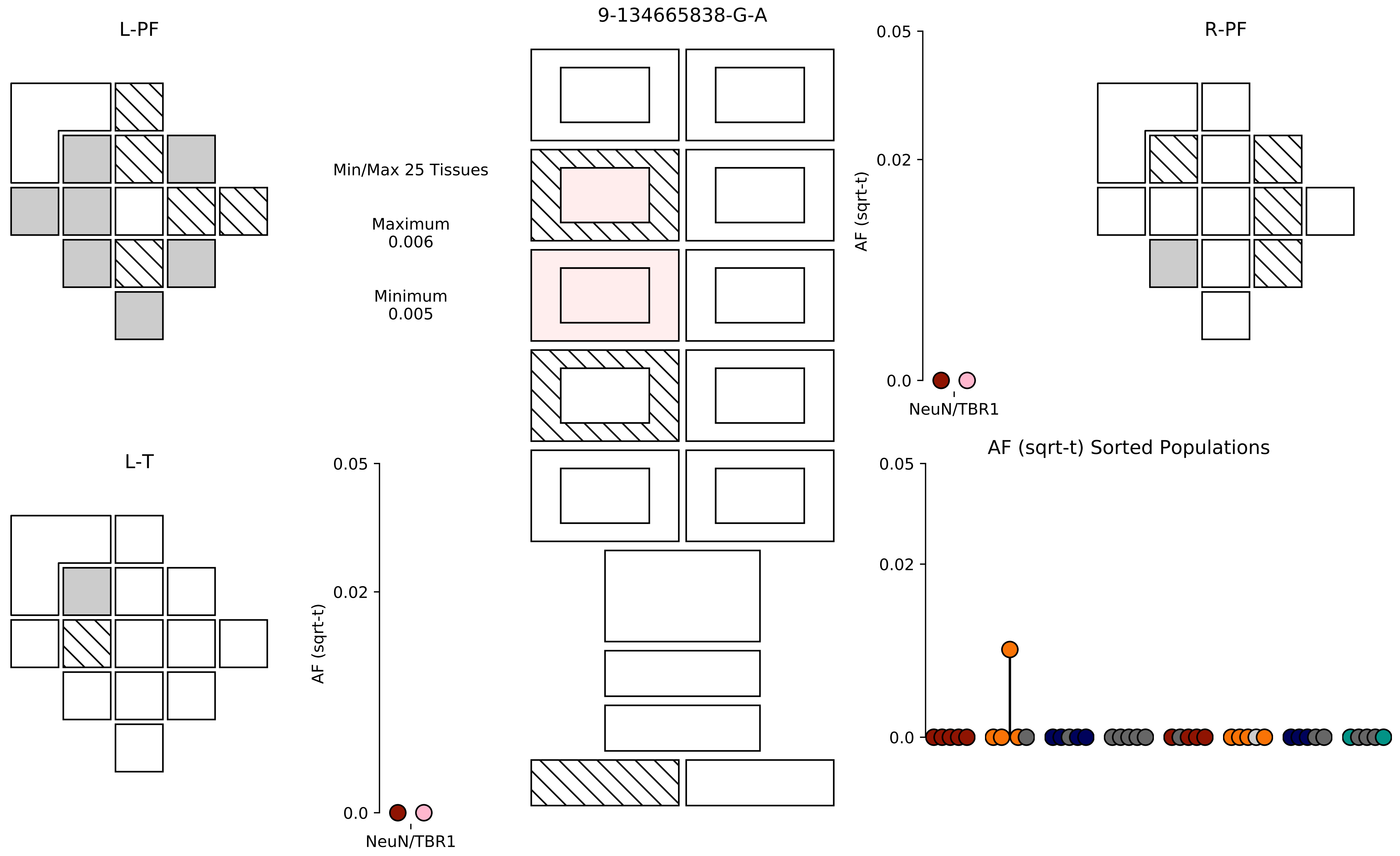
L-PF
9-134665838-G-A
R-PF
Min/Max 25 Tissues
Maximum
0.006
Minimum
0.005
0.05
0.02
0.0
AF (sqrt-t)
NeuN/TBR1
L-T
0.05
0.02
0.0
AF (sqrt-t)
NeuN/TBR1
AF (sqrt-t) Sorted Populations
0.05
0.02
0.0

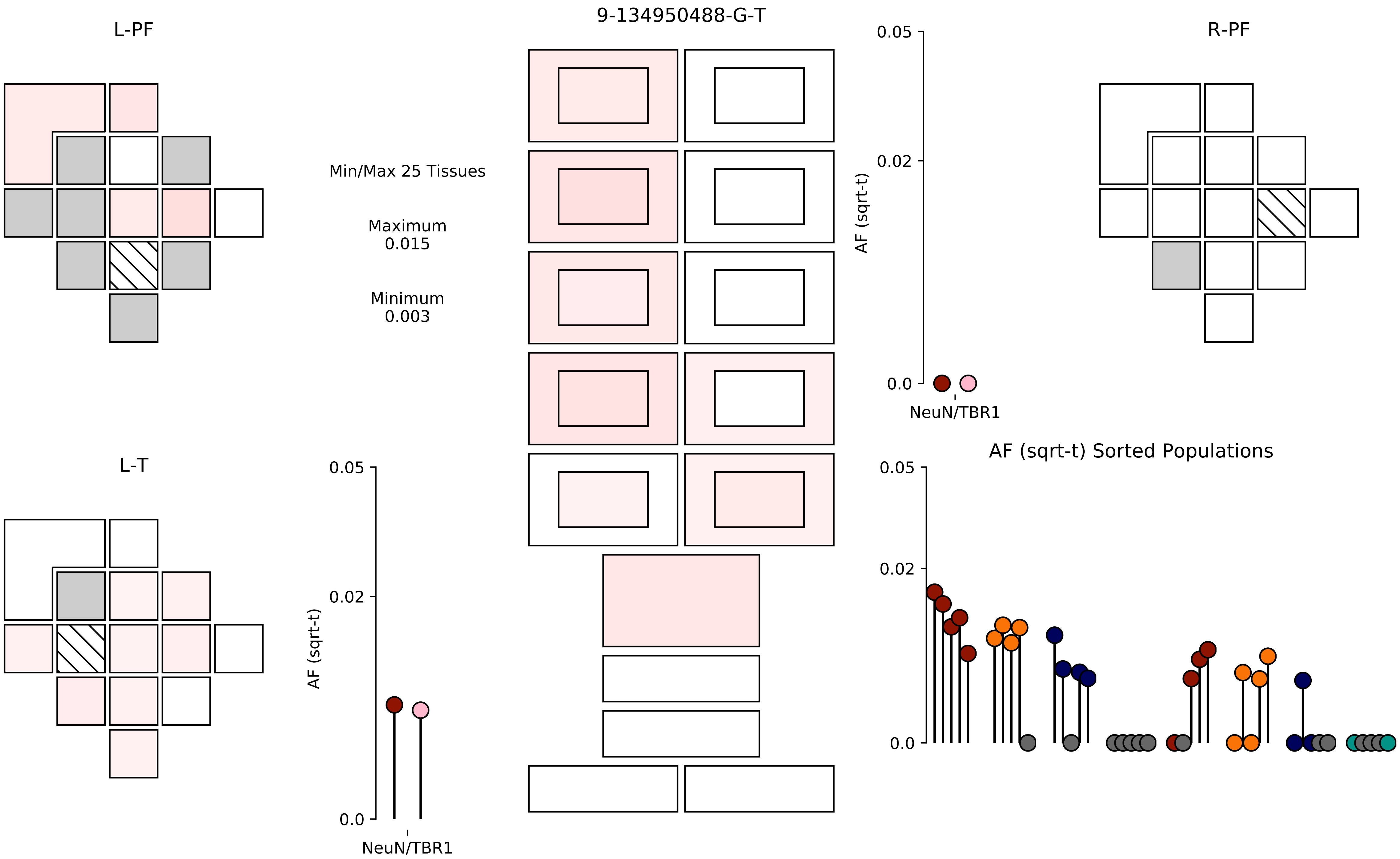
9-134950488-G-T
L-PF
R-PF
L-T
Min/Max 25 Tissues
Maximum
0.015
Minimum
0.003
AF (sqrt-t)
0.05
0.02
0.0
NeuN/TBR1
AF (sqrt-t) Sorted Populations

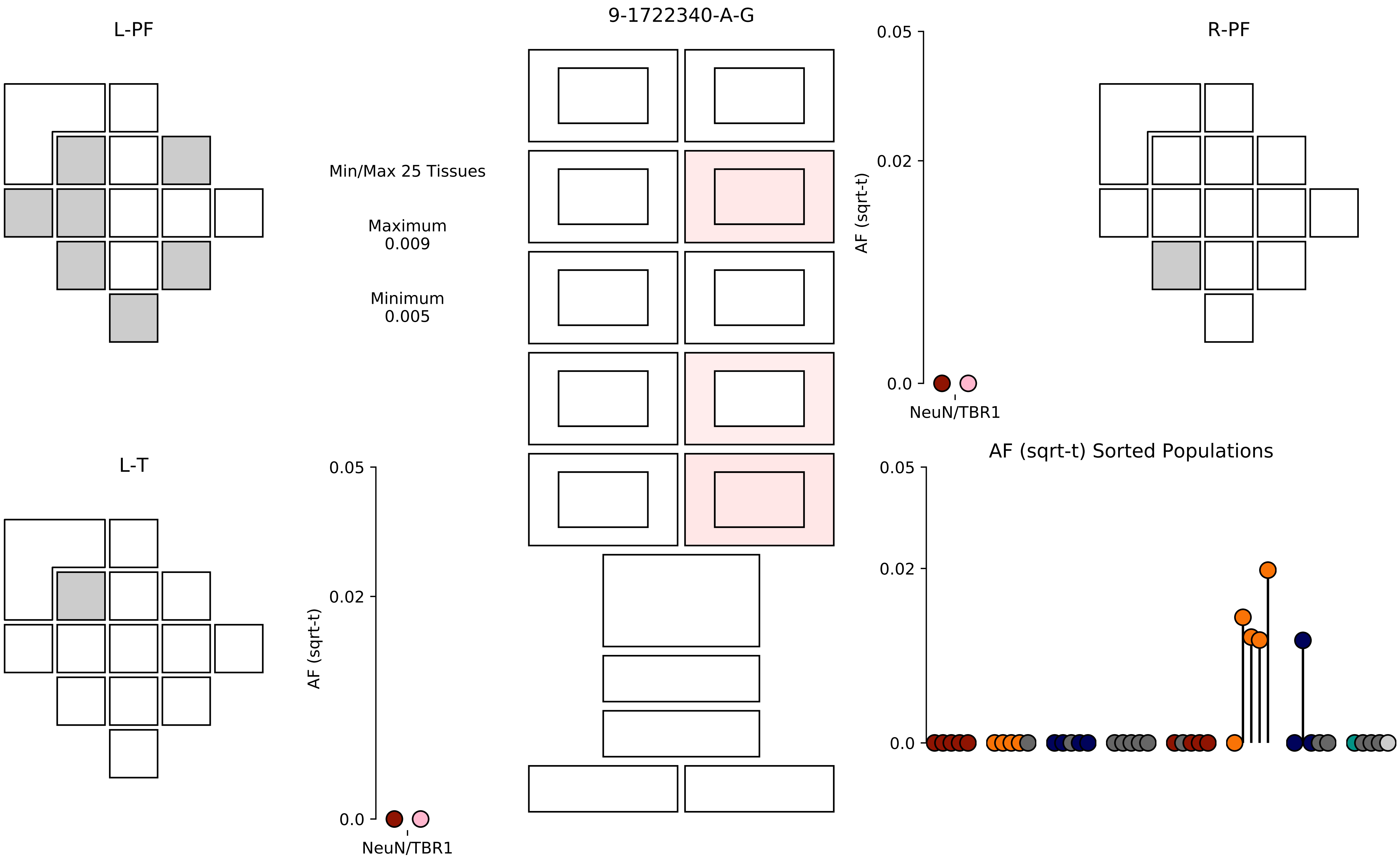
9-1722340-A-G
L-PF
R-PF
L-T
Min/Max 25 Tissues
Maximum
0.009
Minimum
0.005
AF (sqrt-t)
0.05
0.02
0.0
NeuN/TBR1
AF (sqrt-t)
0.05
0.02
0.0
NeuN/TBR1
AF (sqrt-t) Sorted Populations
0.05
0.02
0.0

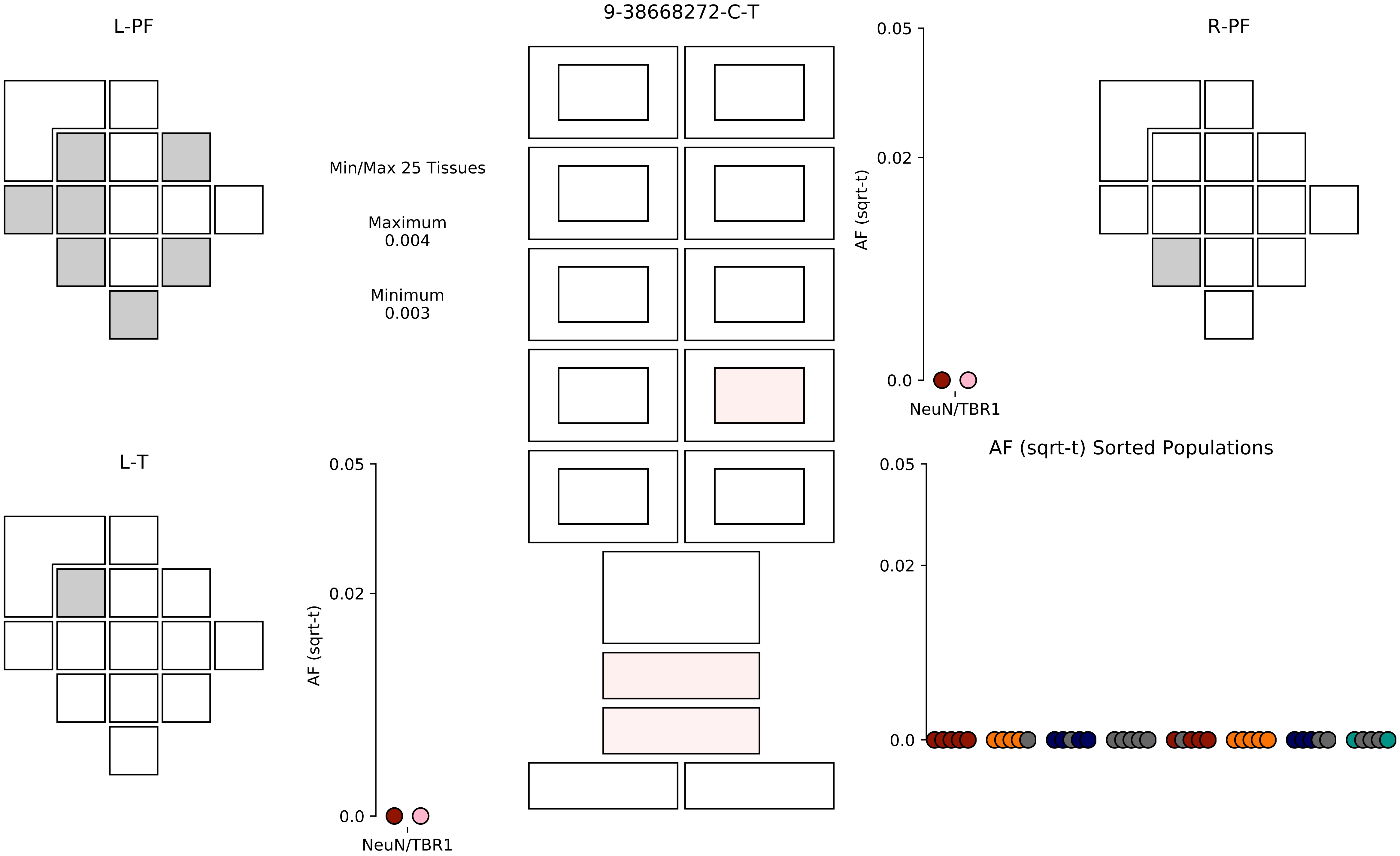

9-38668272-C-T
L-PF
R-PF
L-T
Min/Max 25 Tissues
Maximum
0.004
Minimum
0.003
AF (sqrt-t)
0.05
0.02
0.0
NeuN/TBR1
AF (sqrt-t) Sorted Populations

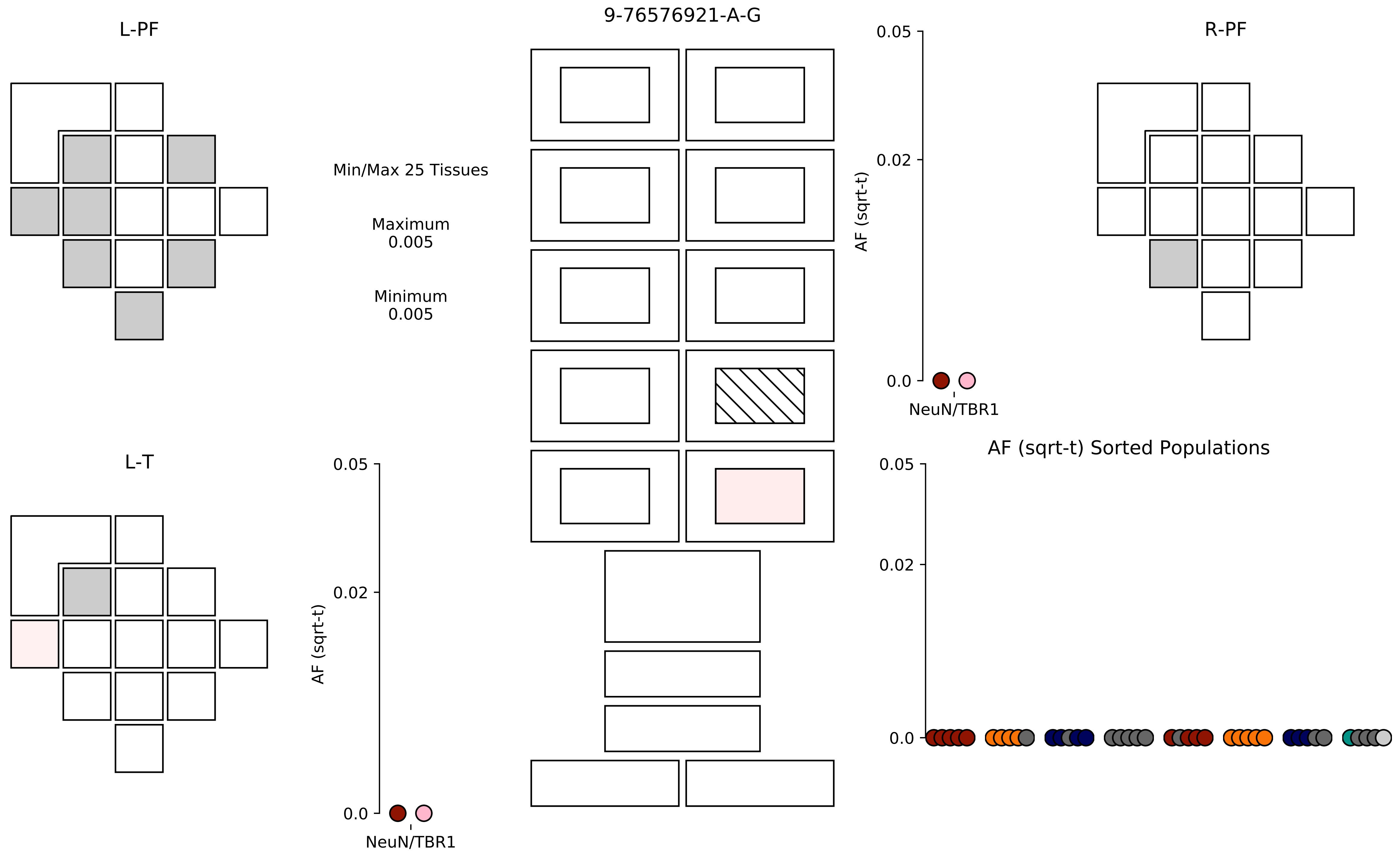
9-76576921-A-G
L-PF
R-PF
L-T
Min/Max 25 Tissues
Maximum
0.005
Minimum
0.005
AF (sqrt-t)
0.05
0.02
0.0
NeuN/TBR1
AF (sqrt-t) Sorted Populations

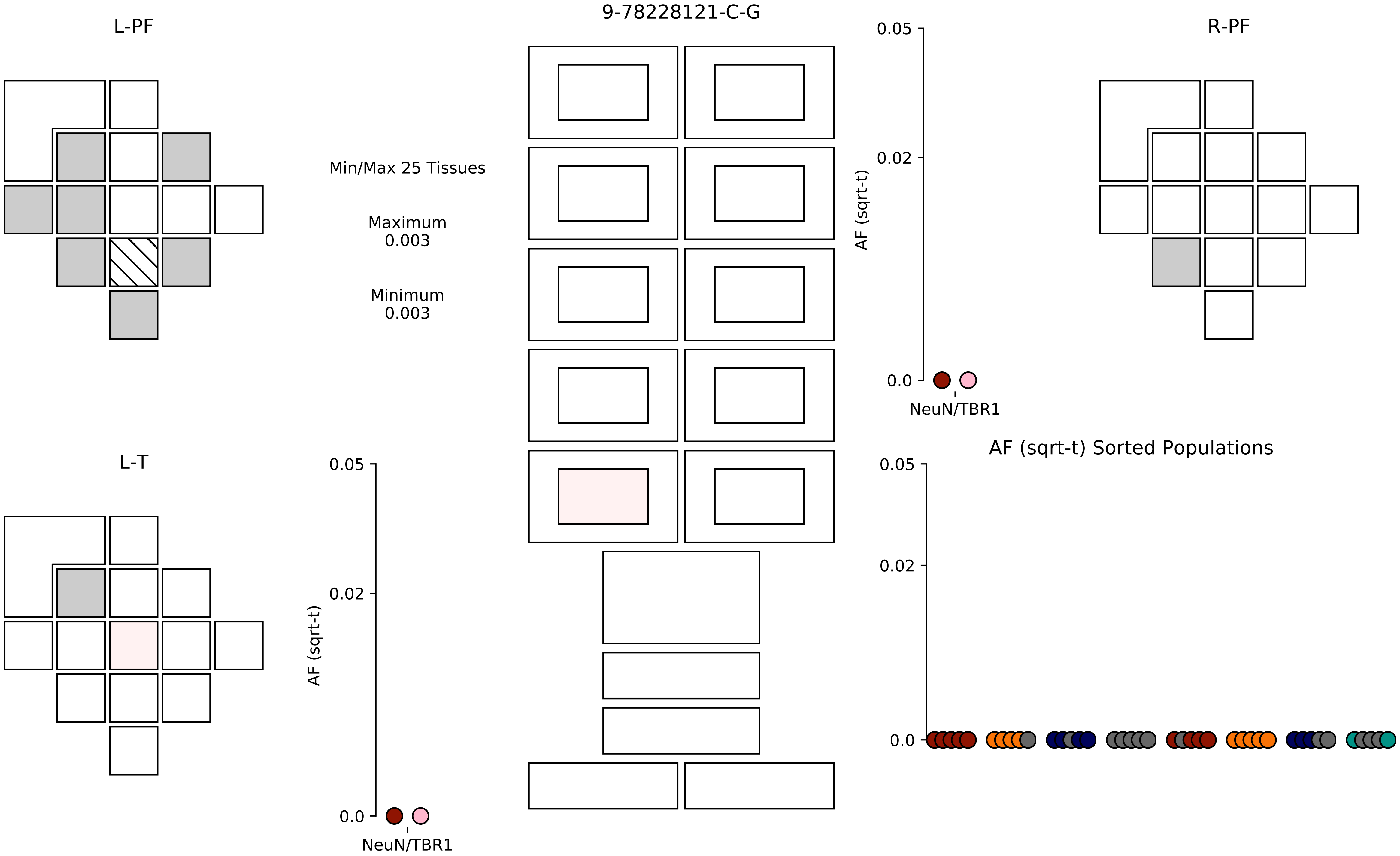

9-78228121-C-G
L-PF
R-PF
L-T
Min/Max 25 Tissues
Maximum
0.003
Minimum
0.003
AF (sqrt-t)
0.05
0.02
0.0
NeuN/TBR1
AF (sqrt-t) Sorted Populations

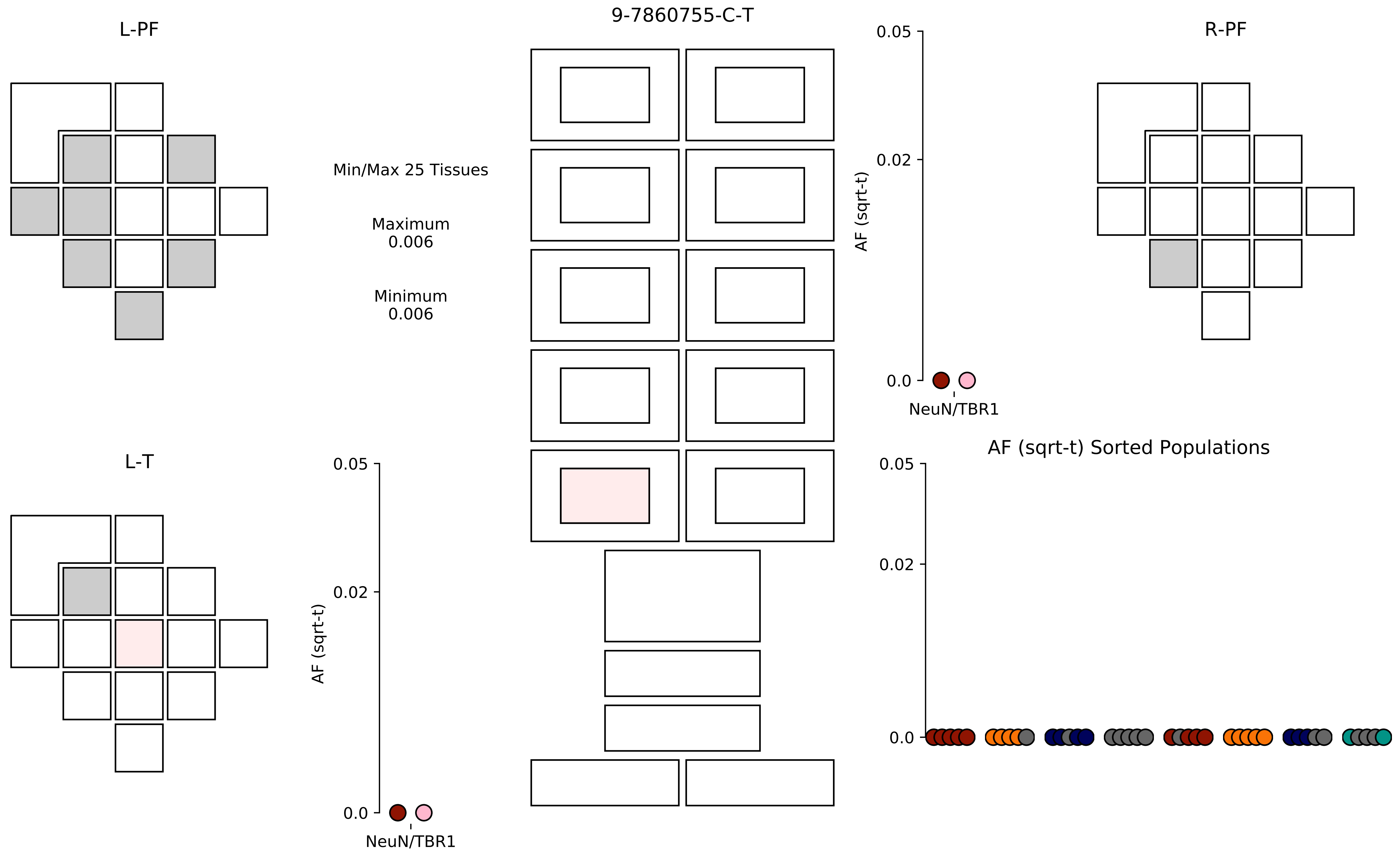

9-7860755-C-T
L-PF
R-PF
L-T
Min/Max 25 Tissues
Maximum
0.006
Minimum
0.006
AF (sqrt-t)
0.05
0.02
0.0
NeuN/TBR1
AF (sqrt-t) Sorted Populations

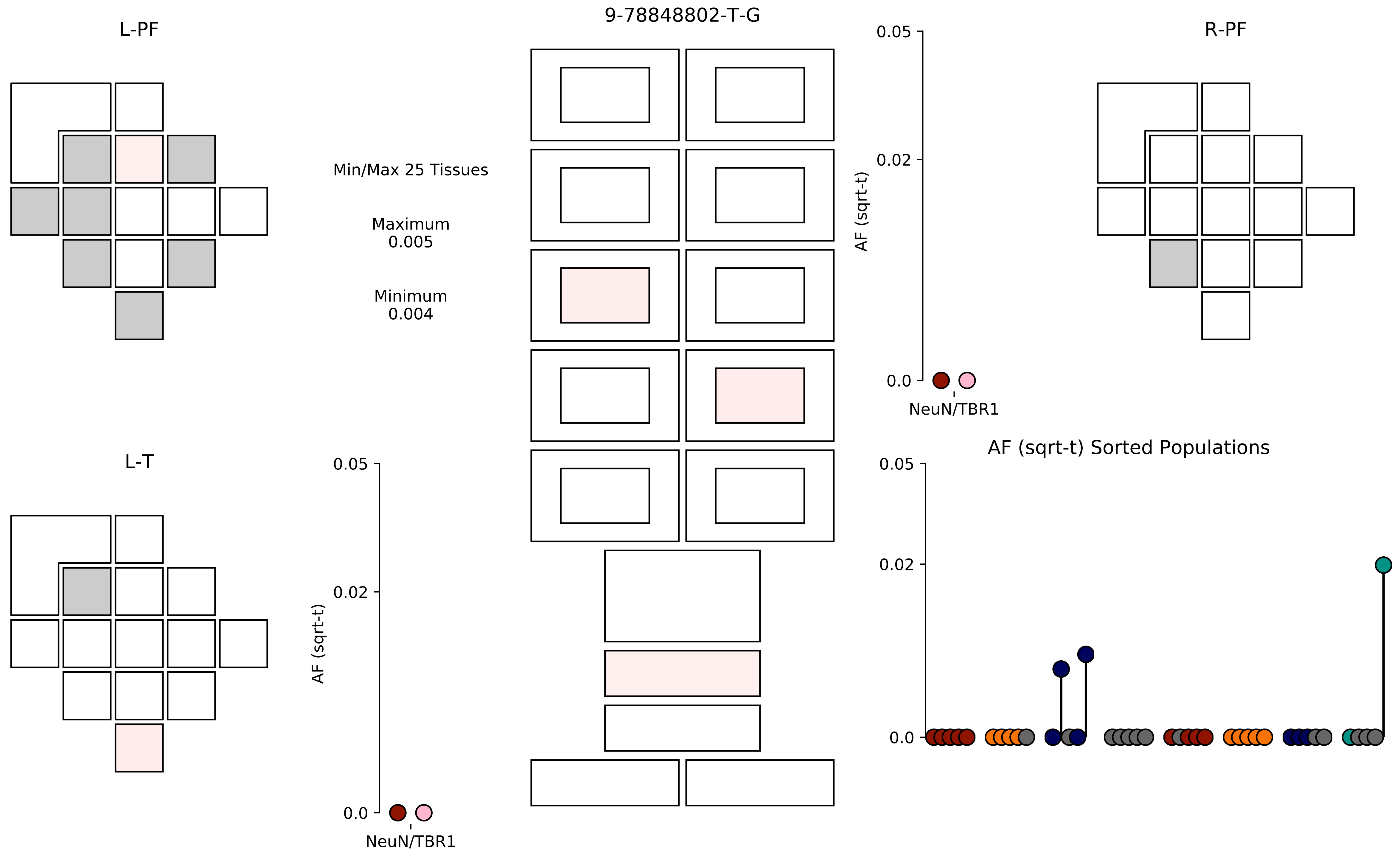
9-78848802-T-G
L-PF
R-PF
L-T
Min/Max 25 Tissues
Maximum
0.005
Minimum
0.004
AF (sqrt-t)
0.05
0.02
0.0
NeuN/TBR1
AF (sqrt-t) Sorted Populations

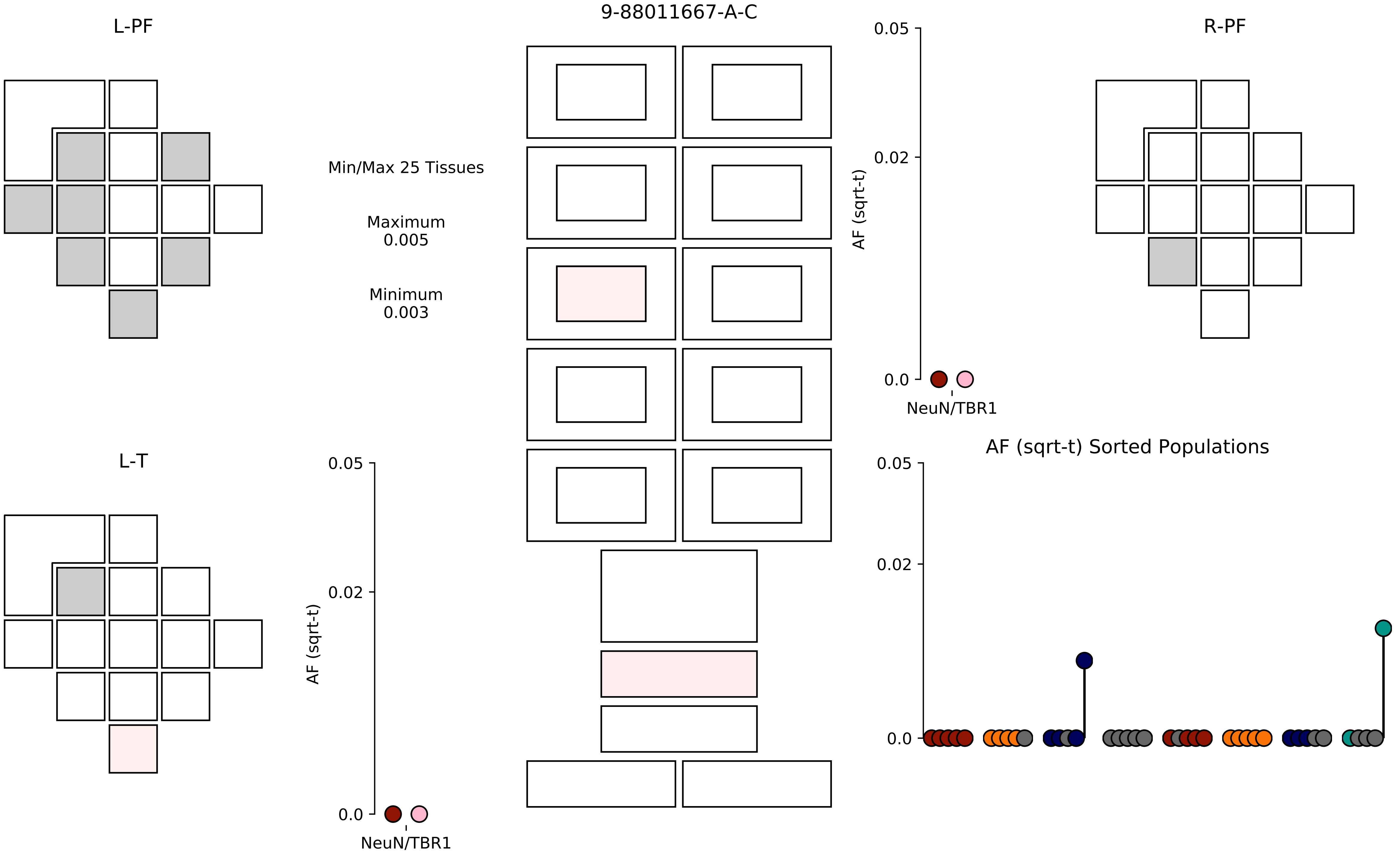

9-88011667-A-C
L-PF
R-PF
L-T
Min/Max 25 Tissues
Maximum
0.005
Minimum
0.003
AF (sqrt-t)
0.05
0.02
0.0
NeuN/TBR1
AF (sqrt-t) Sorted Populations

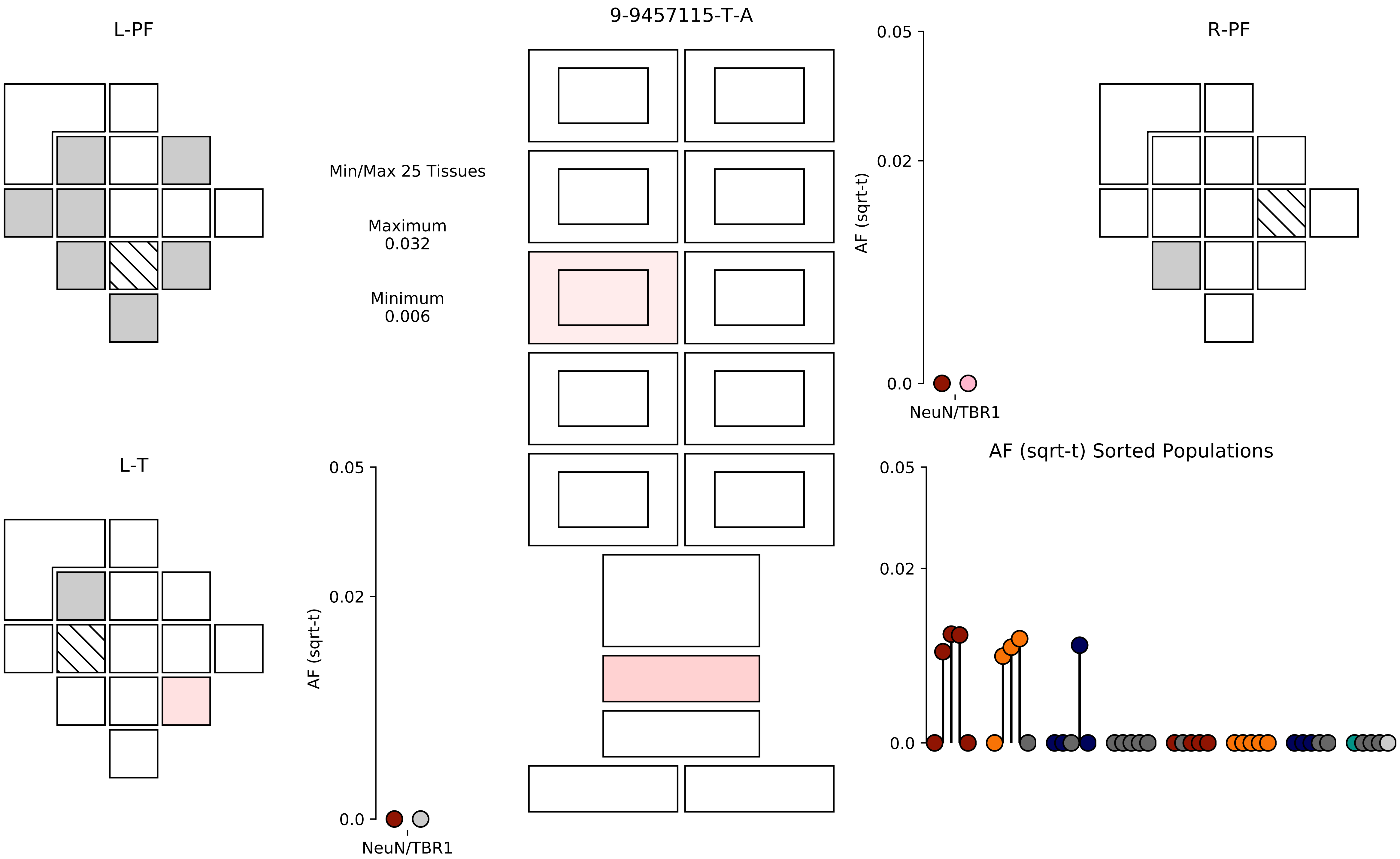
L-PF
9-9457115-T-A
R-PF
Min/Max 25 Tissues
Maximum
0.032
Minimum
0.006
AF (sqrt-t)
0.05
0.02
0.0
NeuN/TBR1
L-T
AF (sqrt-t) Sorted Populations

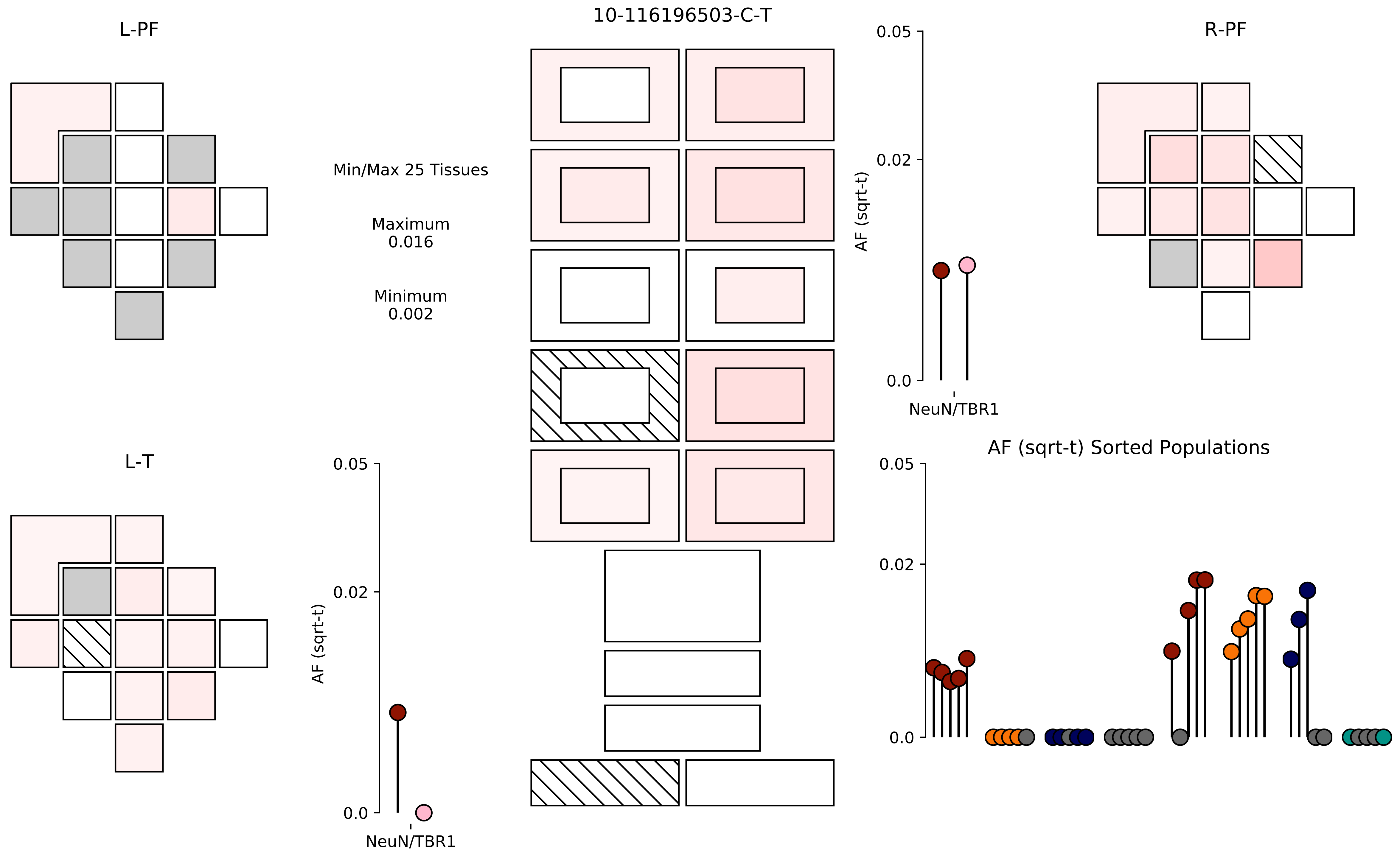

10-116196503-C-T
L-PF
R-PF
L-T
Min/Max 25 Tissues
Maximum
0.016
Minimum
0.002
AF (sqrt-t)
0.05
0.02
0.0
NeuN/TBR1
AF (sqrt-t)
0.05
0.02
0.0
NeuN/TBR1
AF (sqrt-t) Sorted Populations
0.05
0.02
0.0

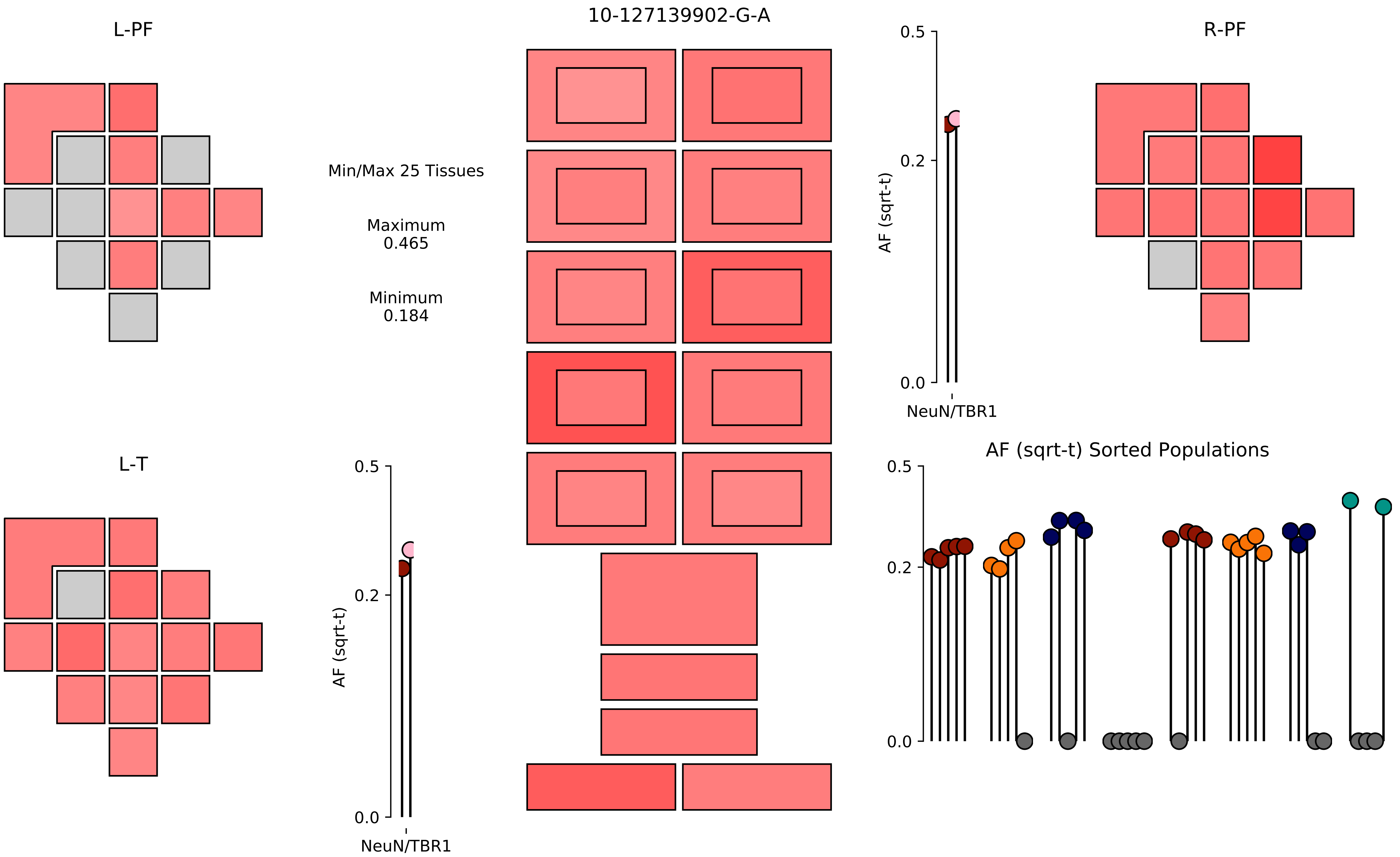

10-127139902-G-A
L-PF
R-PF
L-T
Min/Max 25 Tissues
Maximum
0.465
Minimum
0.184
AF (sqrt-t)
NeuN/TBR1
0.5
0.2
0.0
AF (sqrt-t) Sorted Populations

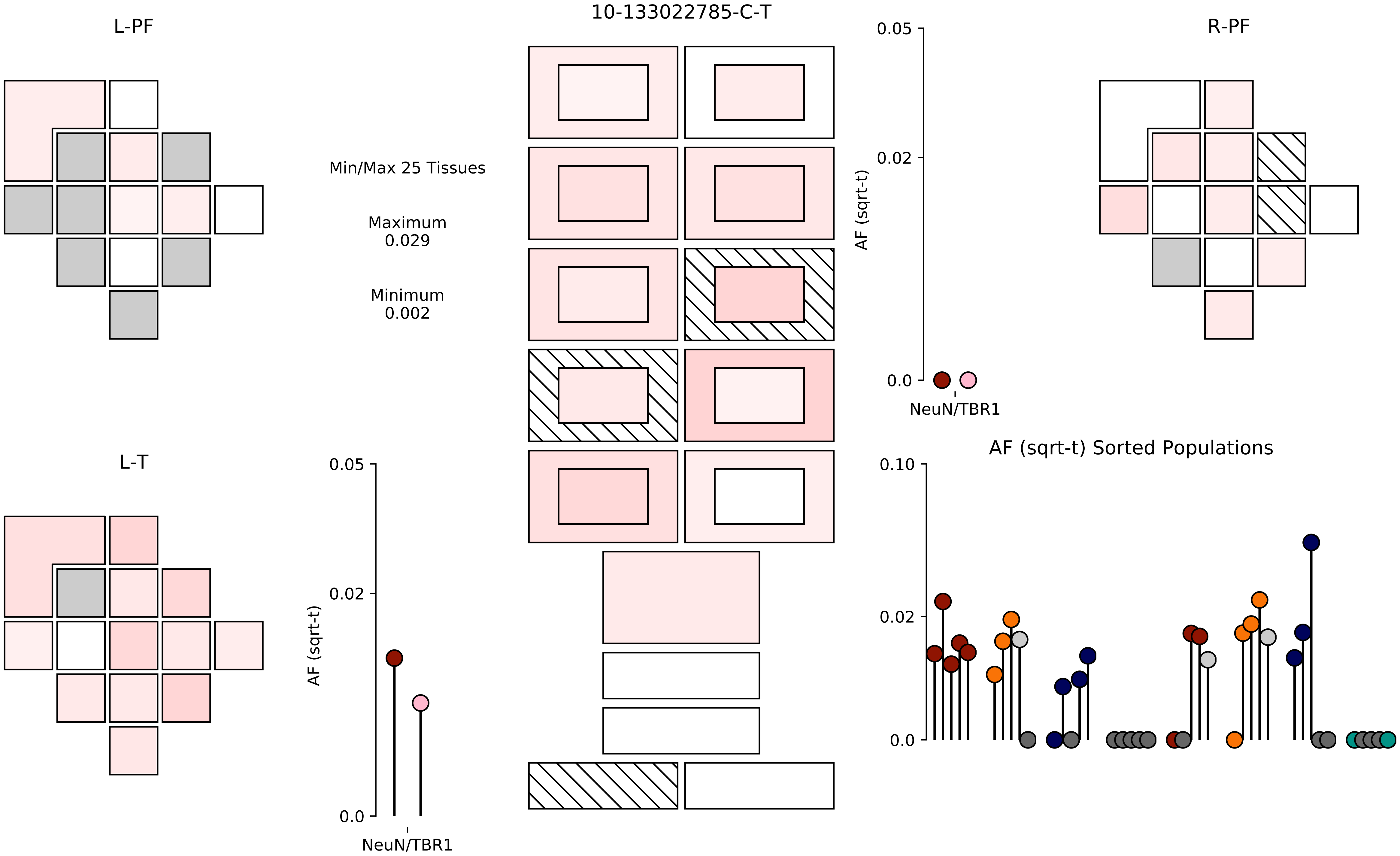

L-PF
10-133022785-C-T
R-PF
Min/Max 25 Tissues
Maximum
0.029
Minimum
0.002
0.05
0.02
0.0
AF (sqrt-t)
NeuN/TBR1
L-T
0.05
0.02
0.0
AF (sqrt-t)
NeuN/TBR1
AF (sqrt-t) Sorted Populations
0.10
0.02
0.0

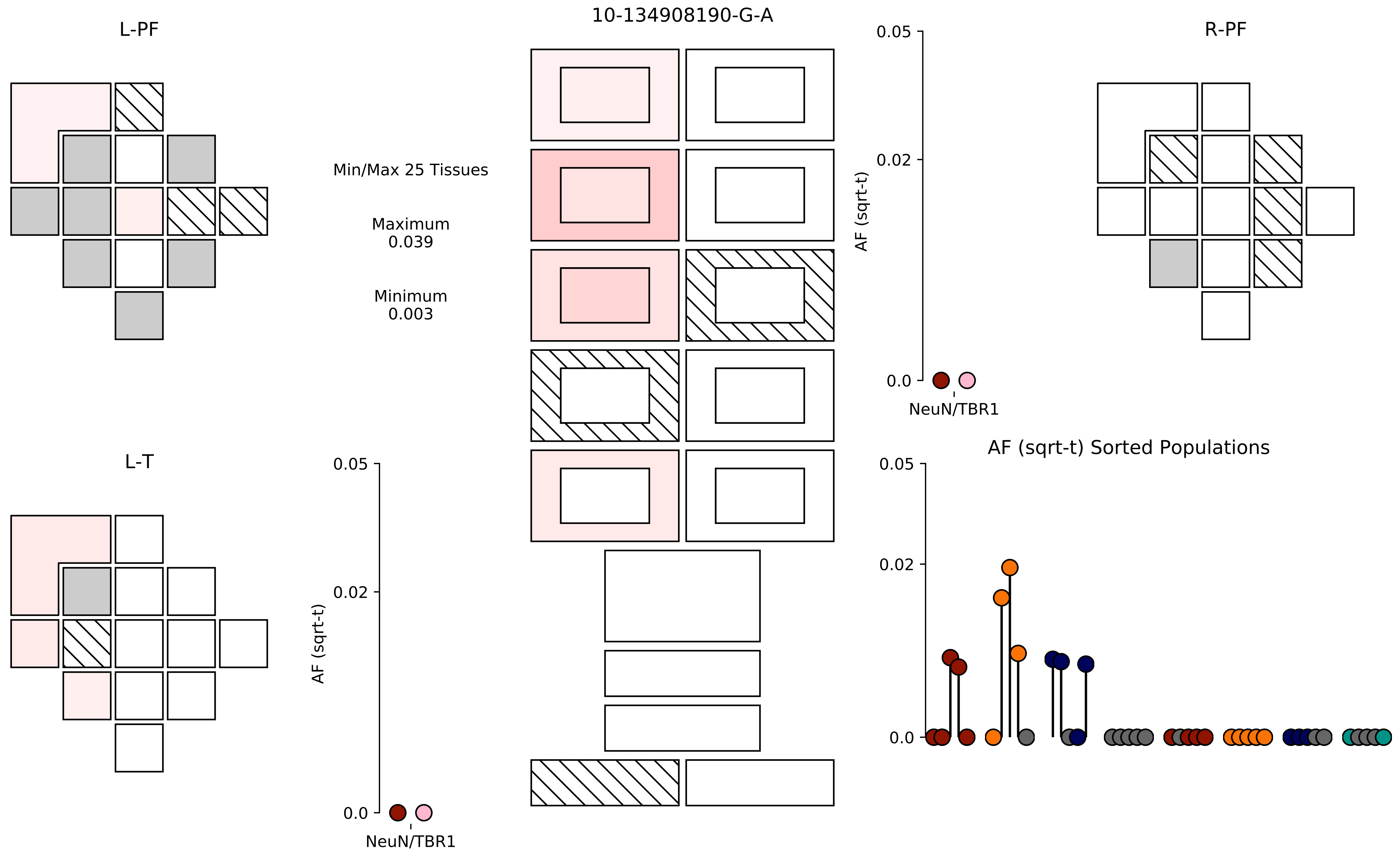

10-134908190-G-A
L-PF
R-PF
L-T
Min/Max 25 Tissues
Maximum
0.039
Minimum
0.003
AF (sqrt-t)
0.05
0.02
0.0
NeuN/TBR1
AF (sqrt-t) Sorted Populations

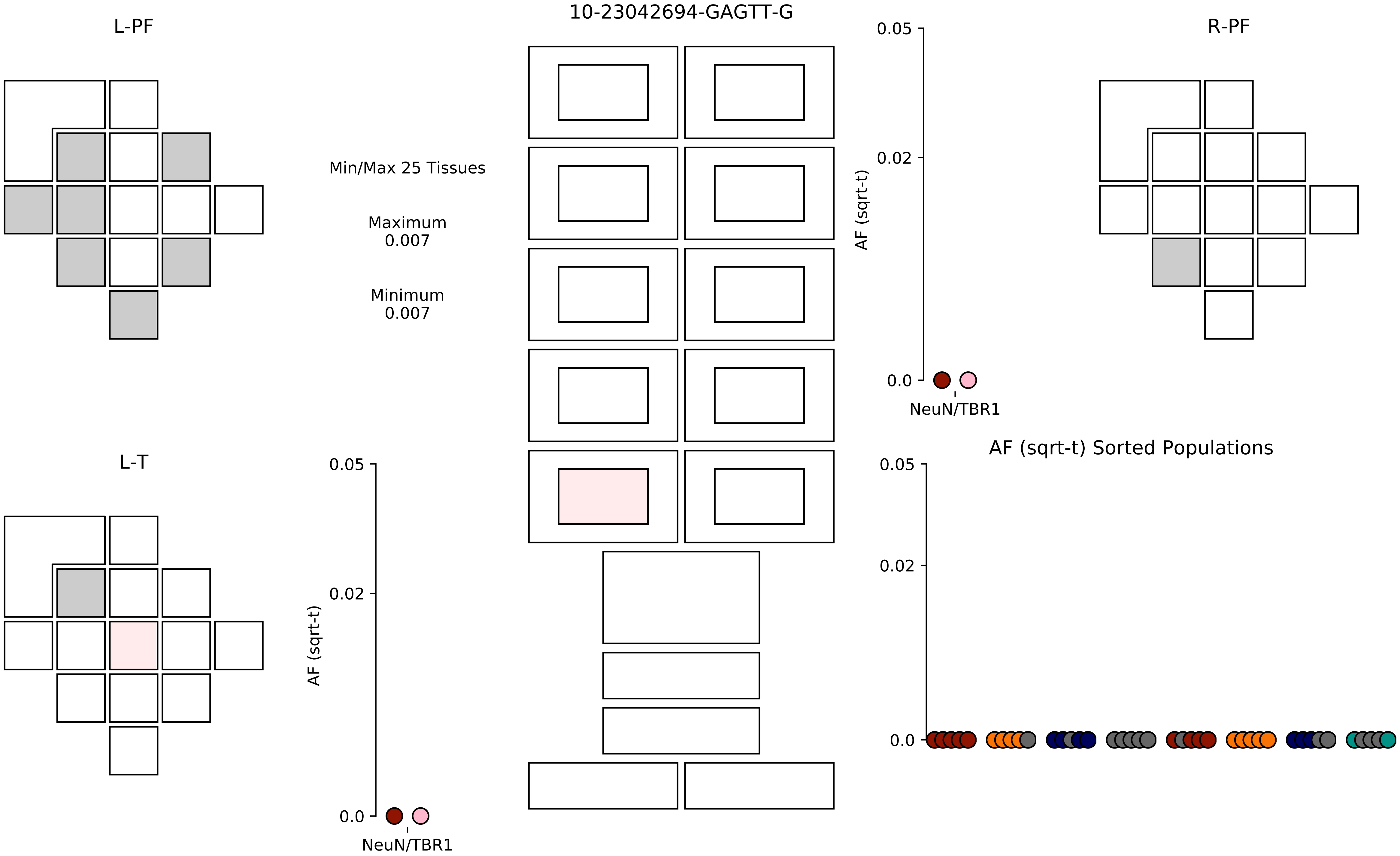

10-23042694-GAGTT-G
L-PF
R-PF
L-T
Min/Max 25 Tissues
Maximum
0.007
Minimum
0.007
AF (sqrt-t)
0.05
0.02
0.0
NeuN/TBR1
AF (sqrt-t) Sorted Populations

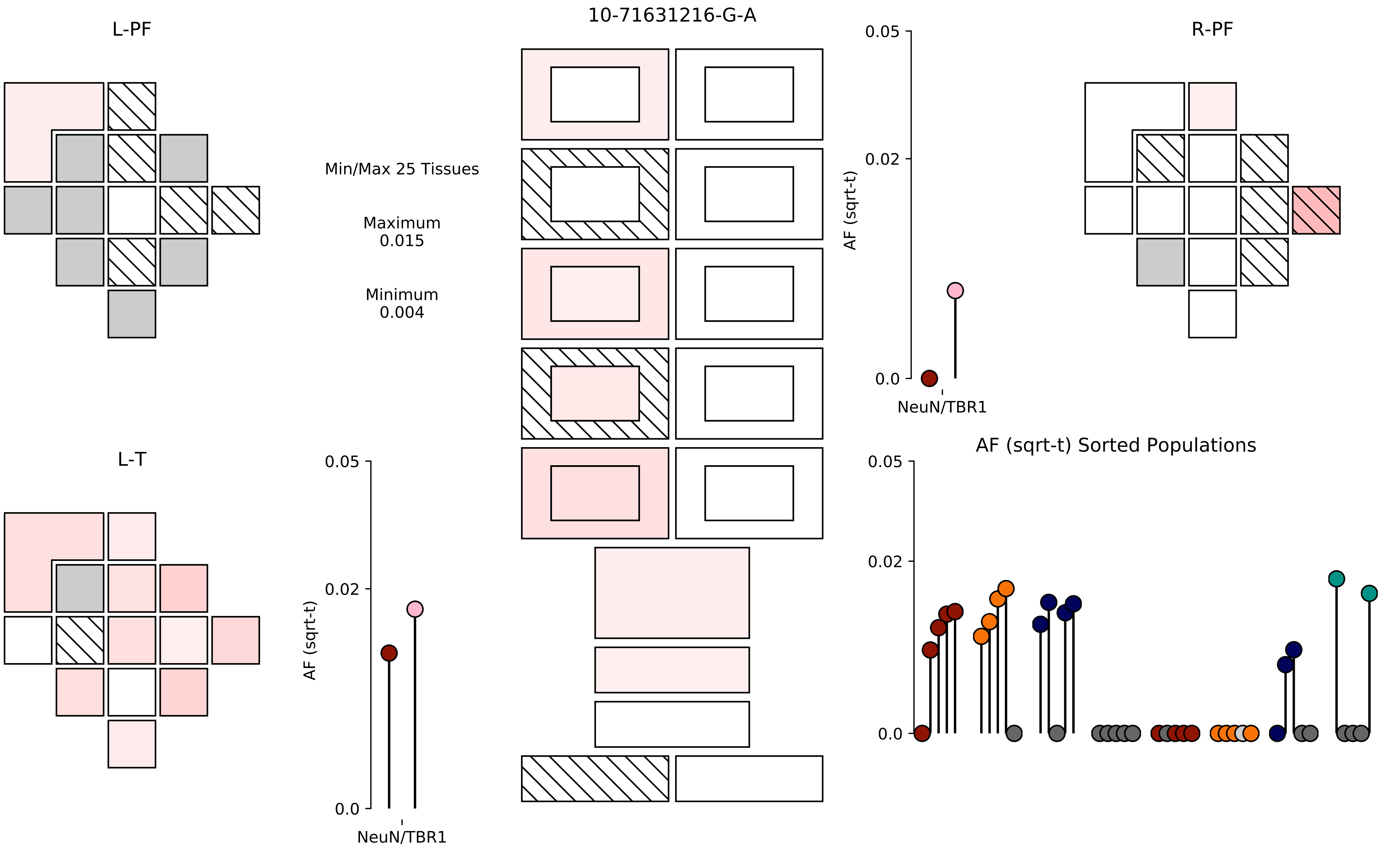

10-71631216-G-A
L-PF
R-PF
L-T
Min/Max 25 Tissues
Maximum
0.015
Minimum
0.004
AF (sqrt-t)
0.05
0.02
0.0
NeuN/TBR1
AF (sqrt-t) Sorted Populations

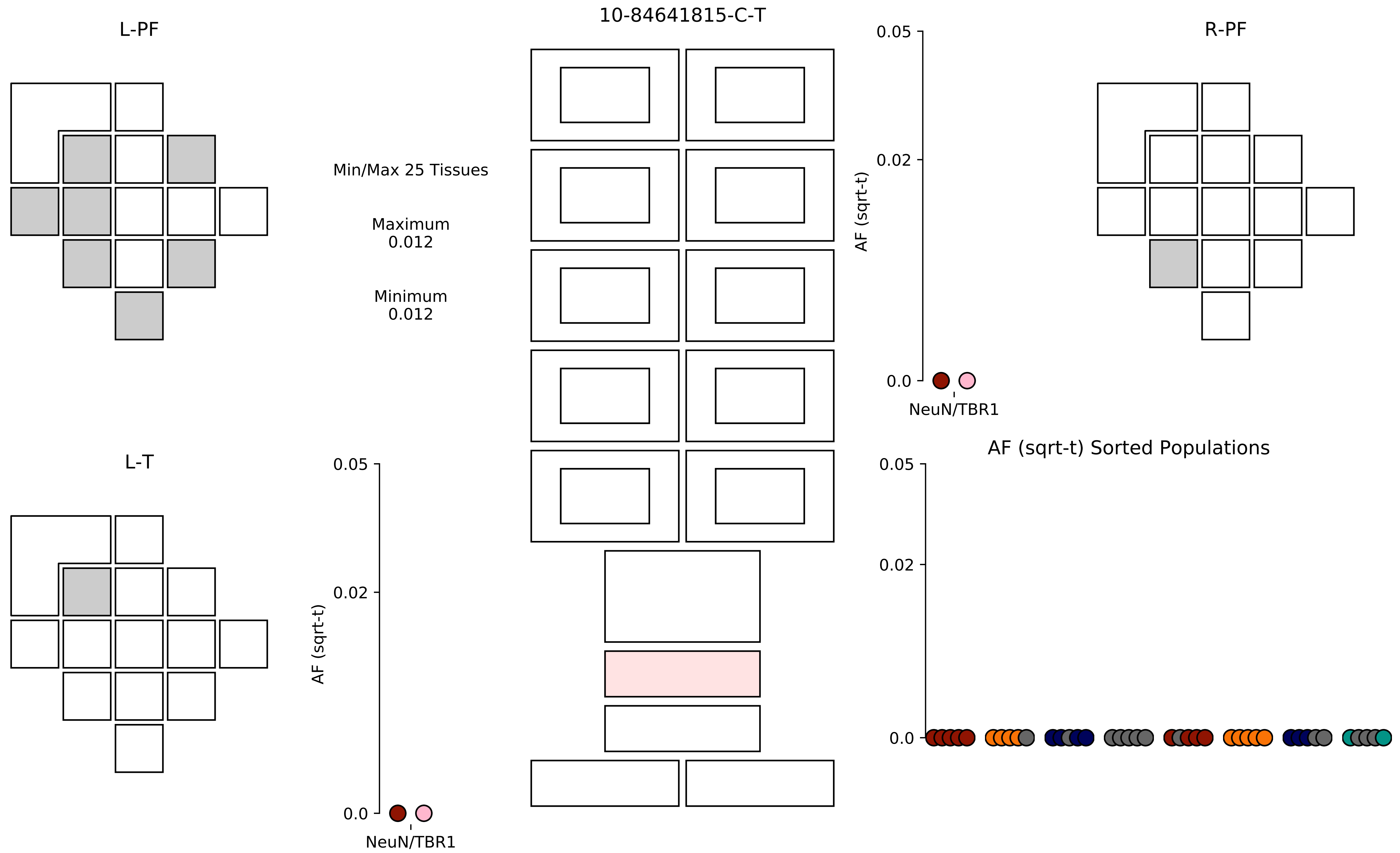

10-84641815-C-T
L-PF
R-PF
L-T
Min/Max 25 Tissues
Maximum
0.012
Minimum
0.012
AF (sqrt-t)
0.05
0.02
0.0
NeuN/TBR1
AF (sqrt-t) Sorted Populations

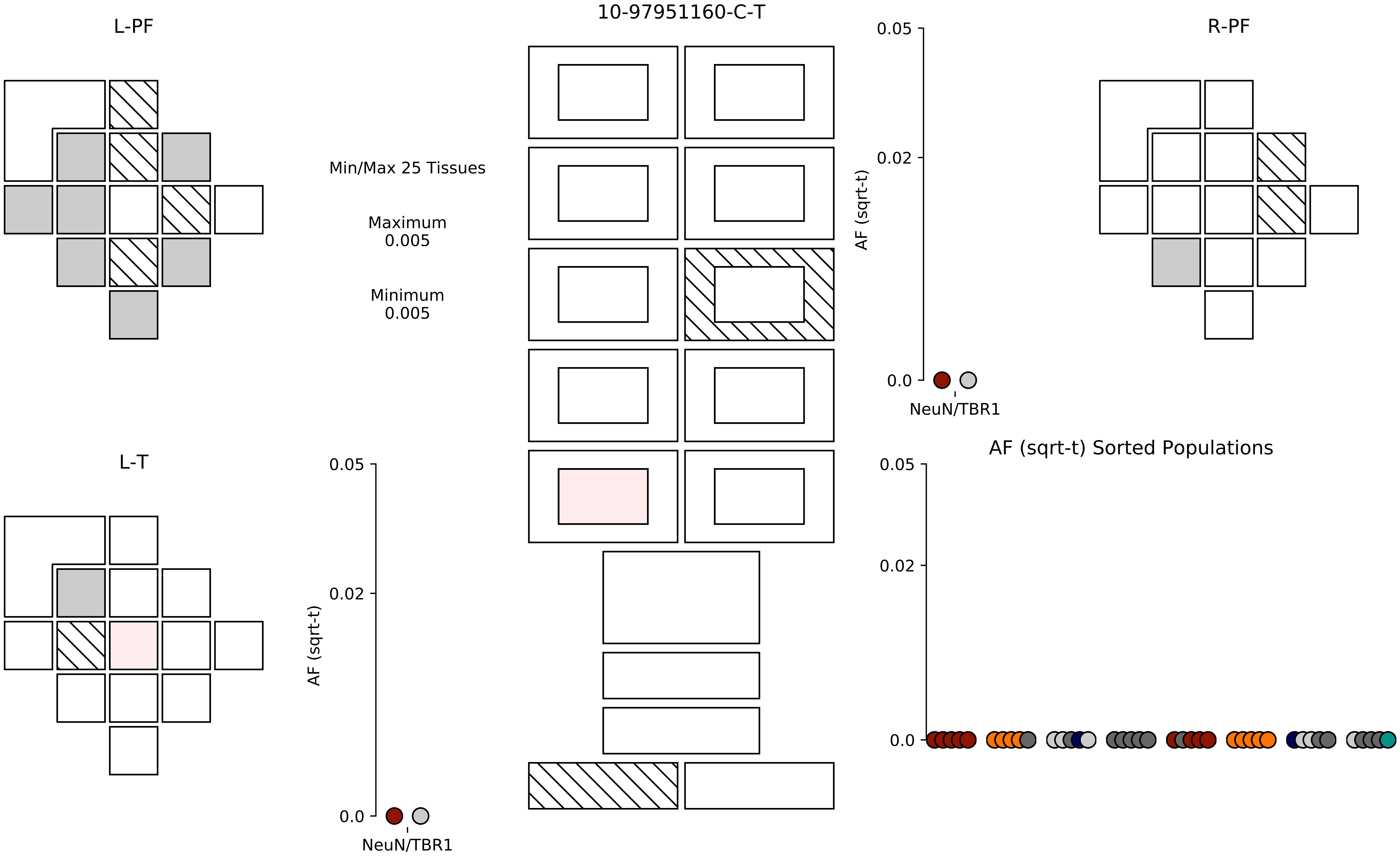
10-97951160-C-T
L-PF
R-PF
L-T
Min/Max 25 Tissues
Maximum
0.005
Minimum
0.005
AF (sqrt-t)
0.05
0.02
0.0
NeuN/TBR1
AF (sqrt-t)
0.05
0.02
0.0
NeuN/TBR1
AF (sqrt-t) Sorted Populations
0.05
0.02
0.0

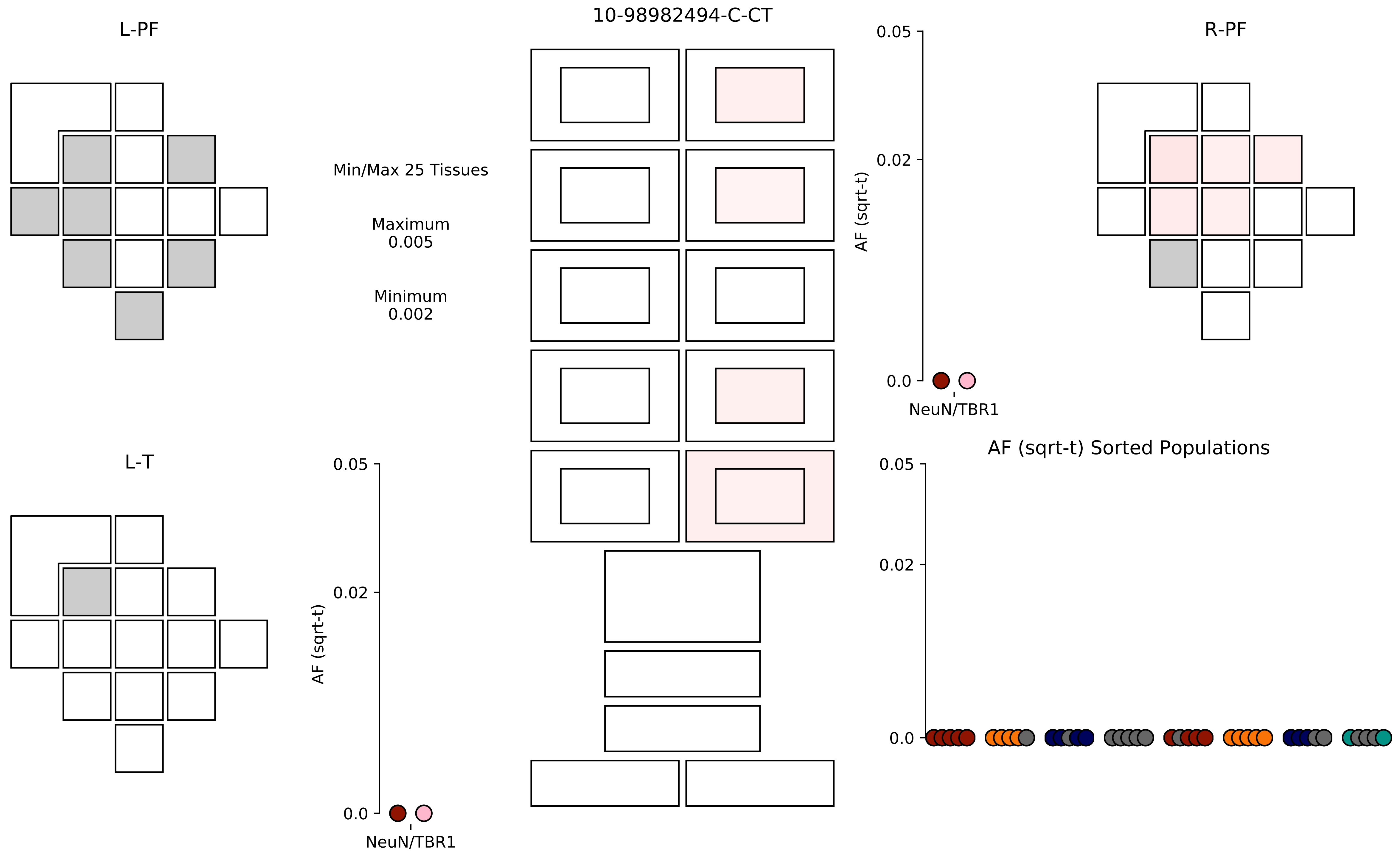

L-PF
10-98982494-C-CT
R-PF
Min/Max 25 Tissues
Maximum
0.005
Minimum
0.002
0.05
0.02
0.0
AF (sqrt-t)
NeuN/TBR1
L-T
0.05
0.02
0.0
AF (sqrt-t)
NeuN/TBR1
AF (sqrt-t) Sorted Populations
0.05
0.02
0.0

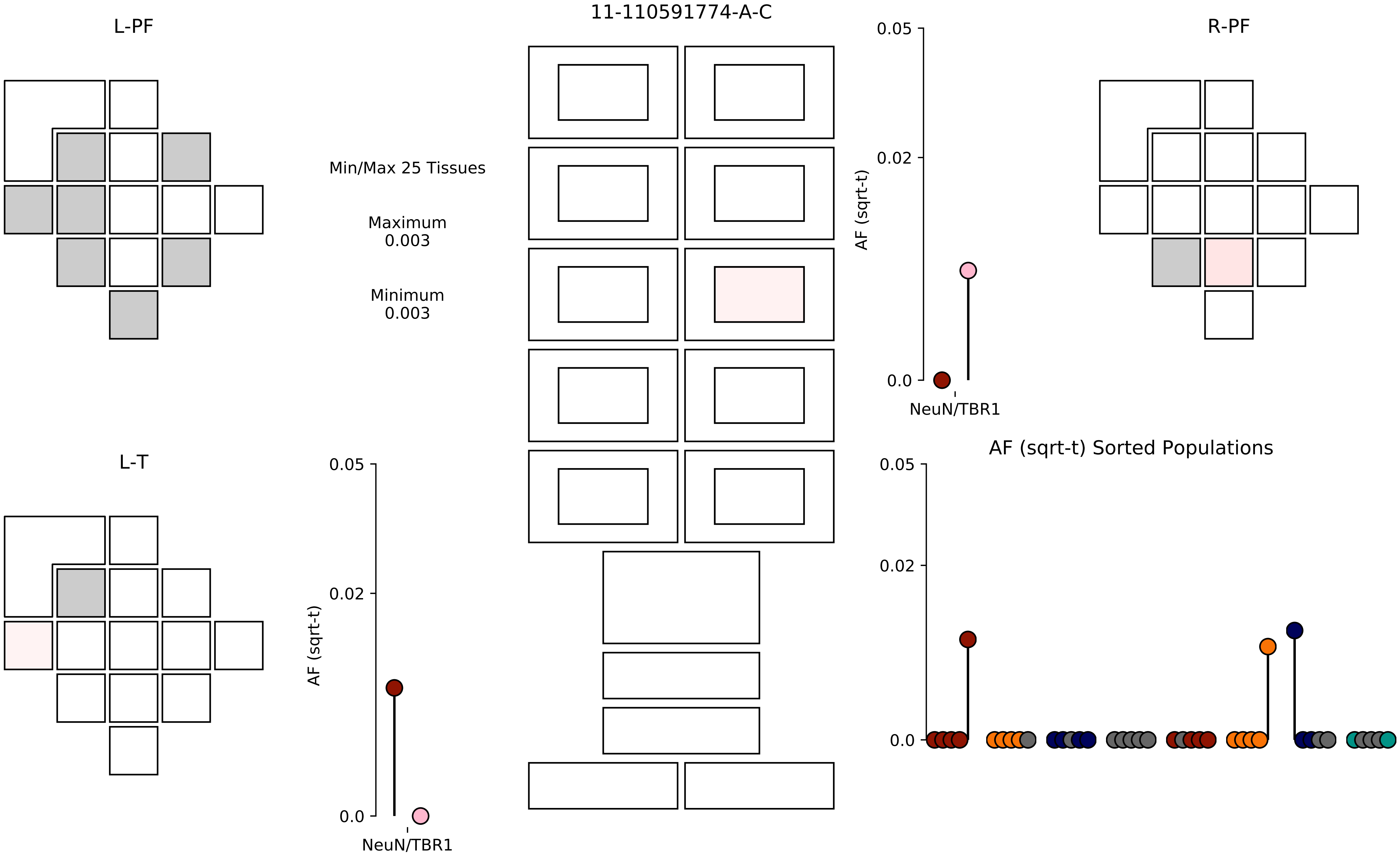

11-110591774-A-C
L-PF
R-PF
L-T
Min/Max 25 Tissues
Maximum
0.003
Minimum
0.003
AF (sqrt-t)
0.05
0.02
0.0
NeuN/TBR1
AF (sqrt-t) Sorted Populations

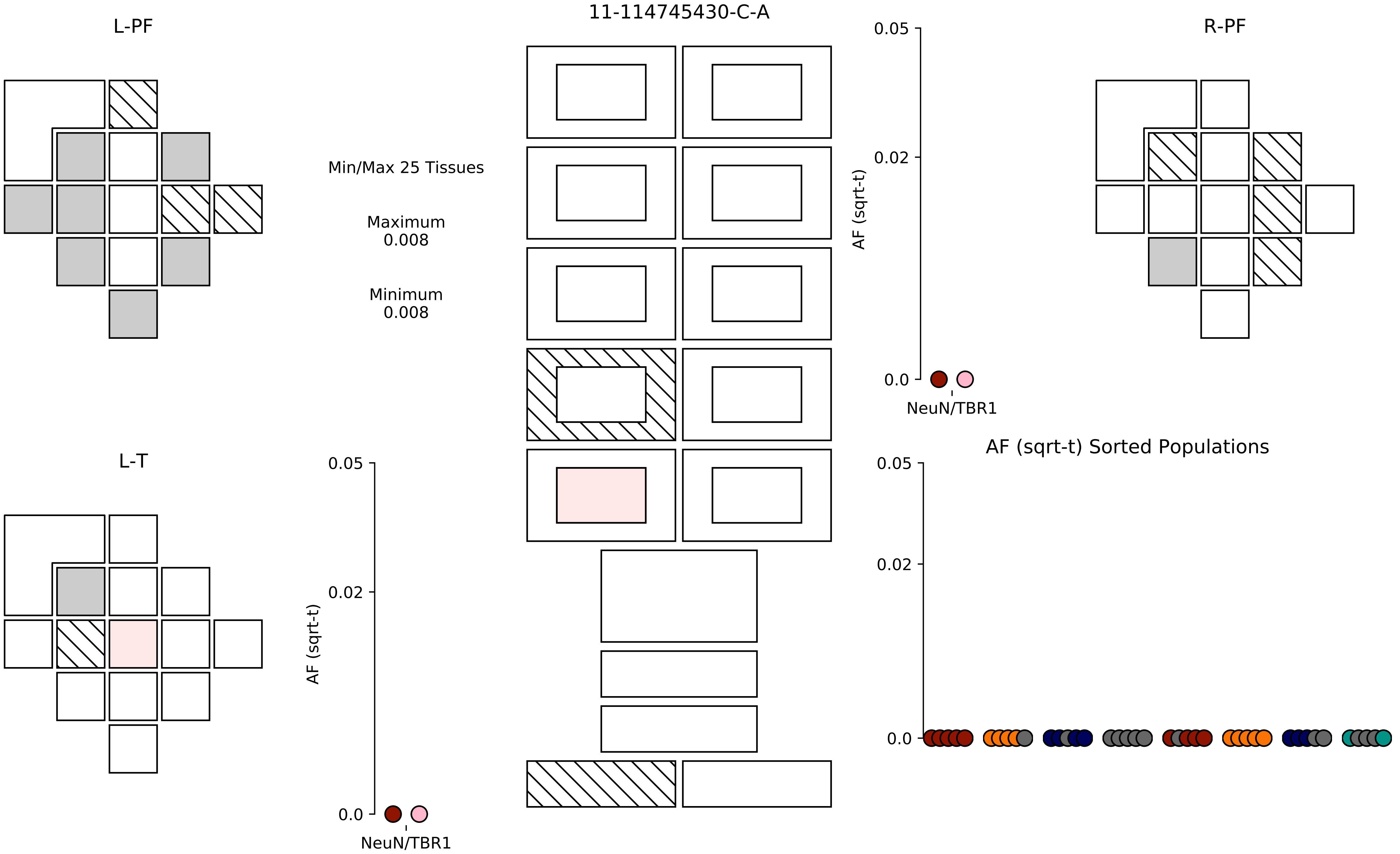

11-114745430-C-A
L-PF
R-PF
L-T
Min/Max 25 Tissues
Maximum
0.008
Minimum
0.008
AF (sqrt-t)
0.05
0.02
0.0
NeuN/TBR1
AF (sqrt-t)
0.05
0.02
0.0
NeuN/TBR1
AF (sqrt-t) Sorted Populations
0.05
0.02
0.0

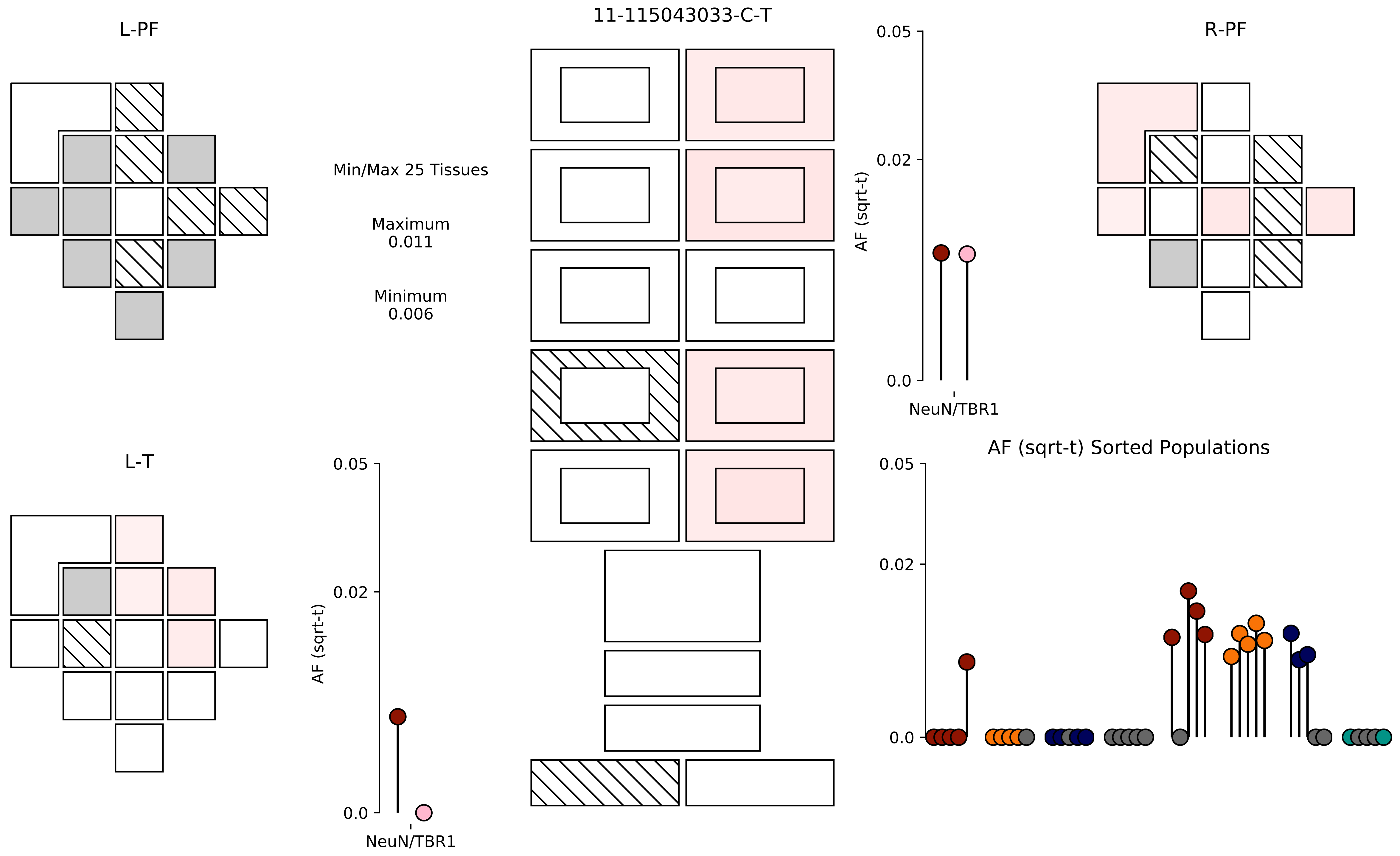
11-115043033-C-T
L-PF
R-PF
L-T
Min/Max 25 Tissues
Maximum
0.011
Minimum
0.006
AF (sqrt-t)
0.05
0.02
0.0
NeuN/TBR1
AF (sqrt-t) Sorted Populations

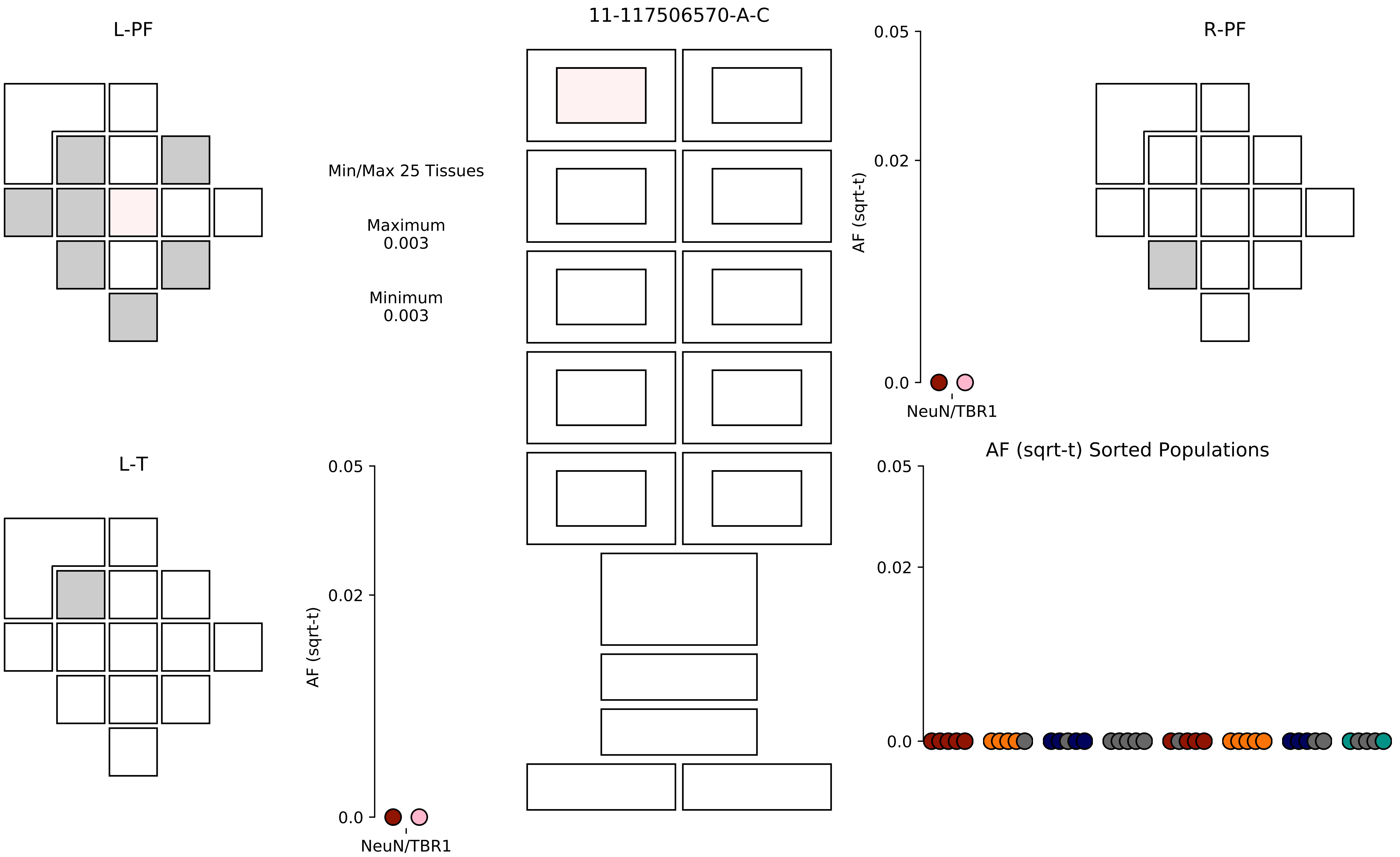
11-117506570-A-C
L-PF
R-PF
L-T
Min/Max 25 Tissues
Maximum
0.003
Minimum
0.003
AF (sqrt-t)
0.05
0.02
0.0
NeuN/TBR1
AF (sqrt-t) Sorted Populations

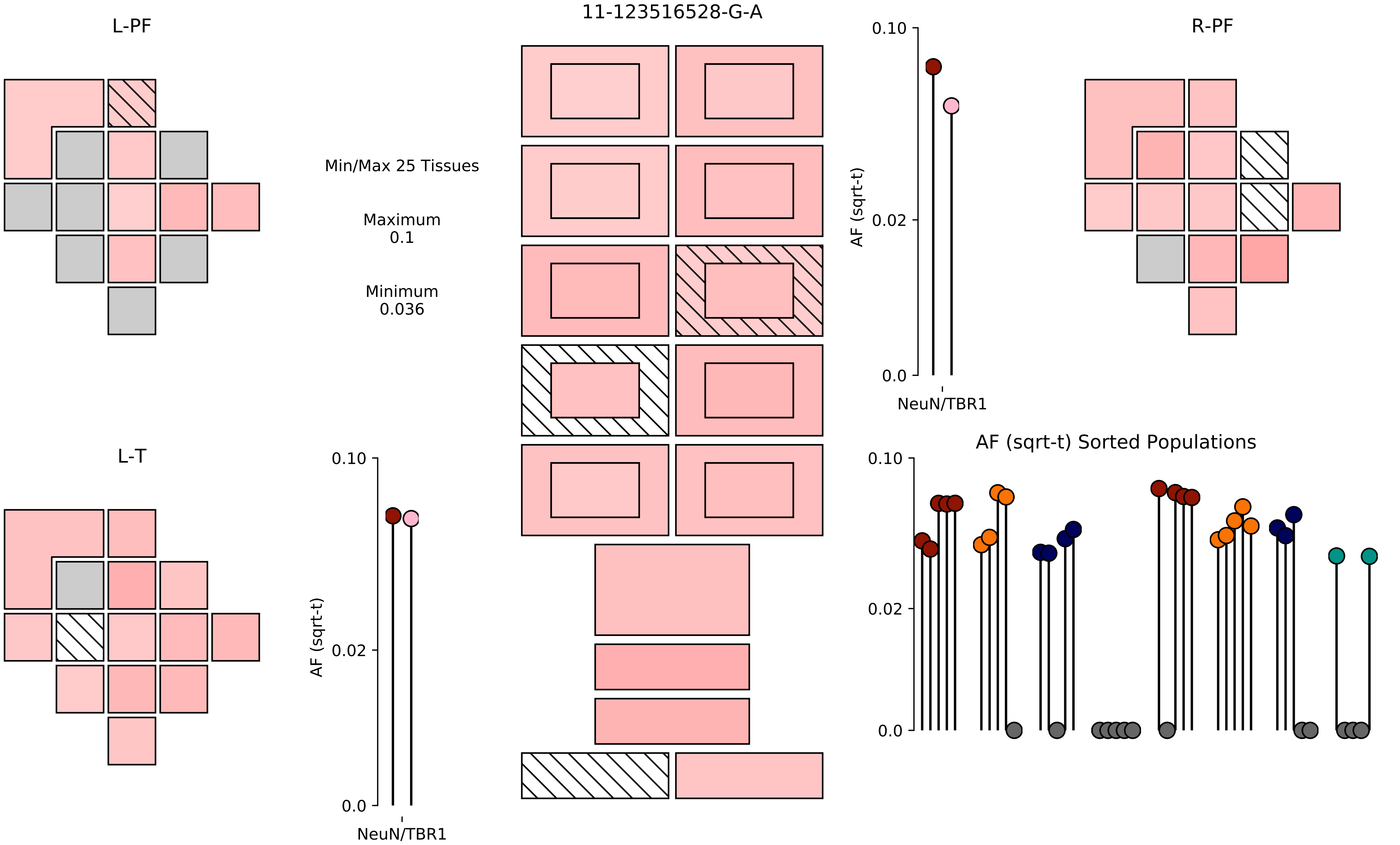

11-123516528-G-A
L-PF
R-PF
L-T
Min/Max 25 Tissues
Maximum
0.1
Minimum
0.036
AF (sqrt-t)
0.10
0.02
0.0
NeuN/TBR1
AF (sqrt-t) Sorted Populations

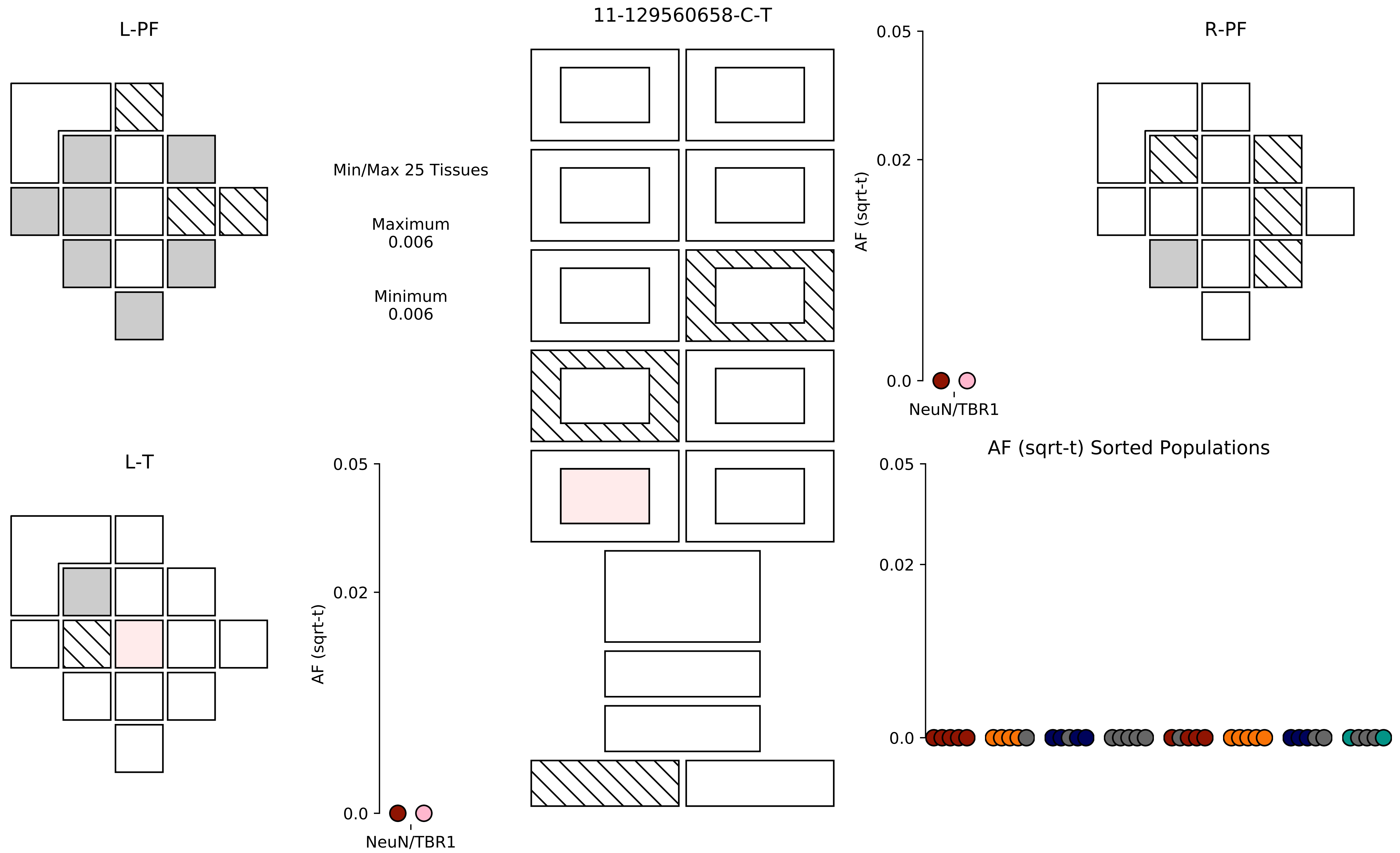

11-129560658-C-T
L-PF
R-PF
L-T
Min/Max 25 Tissues
Maximum
0.006
Minimum
0.006
0.05
0.02
0.0
AF (sqrt-t)
NeuN/TBR1
AF (sqrt-t) Sorted Populations

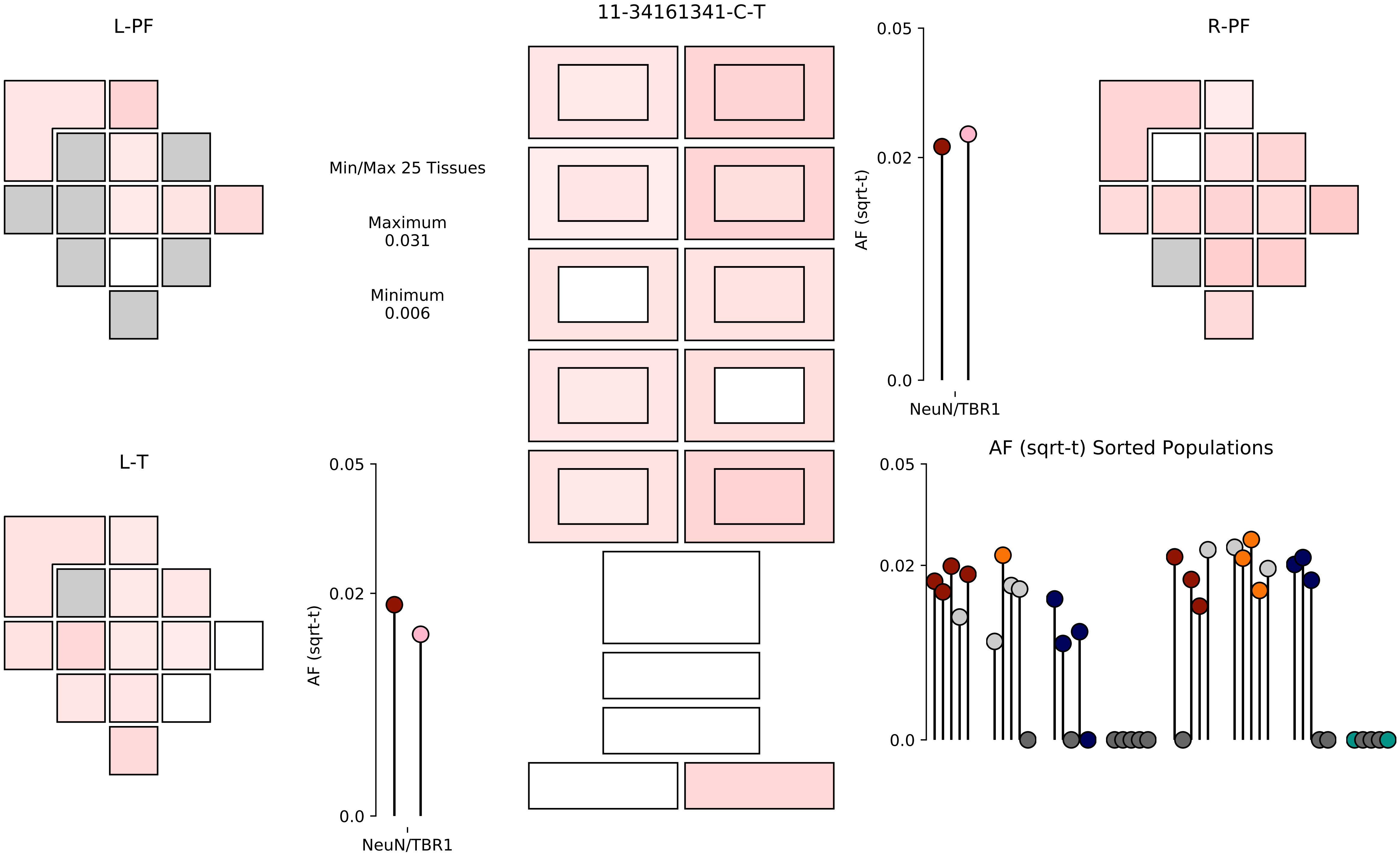

11-34161341-C-T
L-PF
R-PF
L-T
Min/Max 25 Tissues
Maximum
0.031
Minimum
0.006
0.05
0.02
0.0
AF (sqrt-t)
NeuN/TBR1
AF (sqrt-t) Sorted Populations

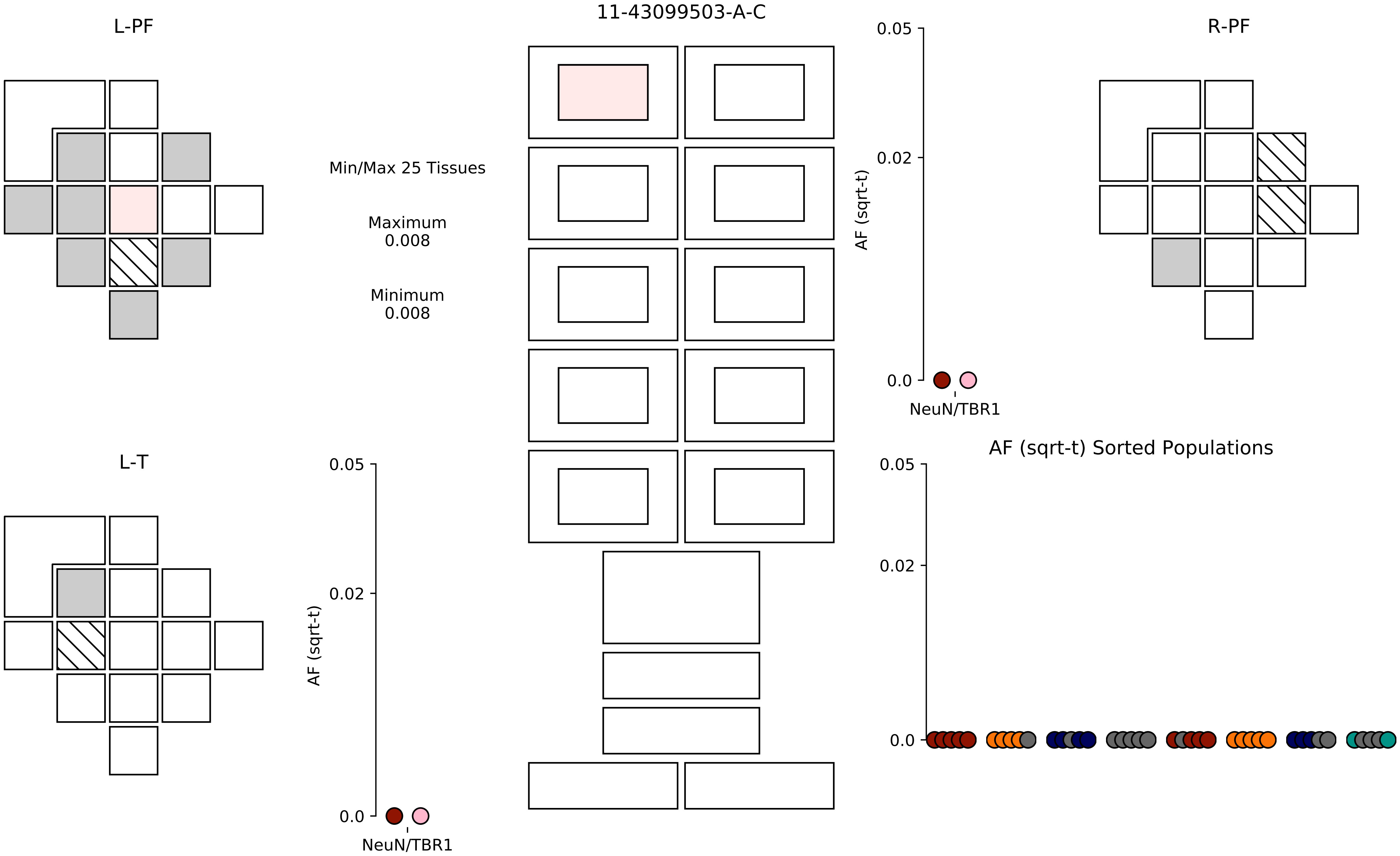
L-PF
11-43099503-A-C
R-PF
Min/Max 25 Tissues
Maximum
0.008
Minimum
0.008
0.05
0.02
0.0
AF (sqrt-t)
NeuN/TBR1
L-T
0.05
0.02
0.0
AF (sqrt-t)
NeuN/TBR1
AF (sqrt-t) Sorted Populations
0.05
0.02
0.0

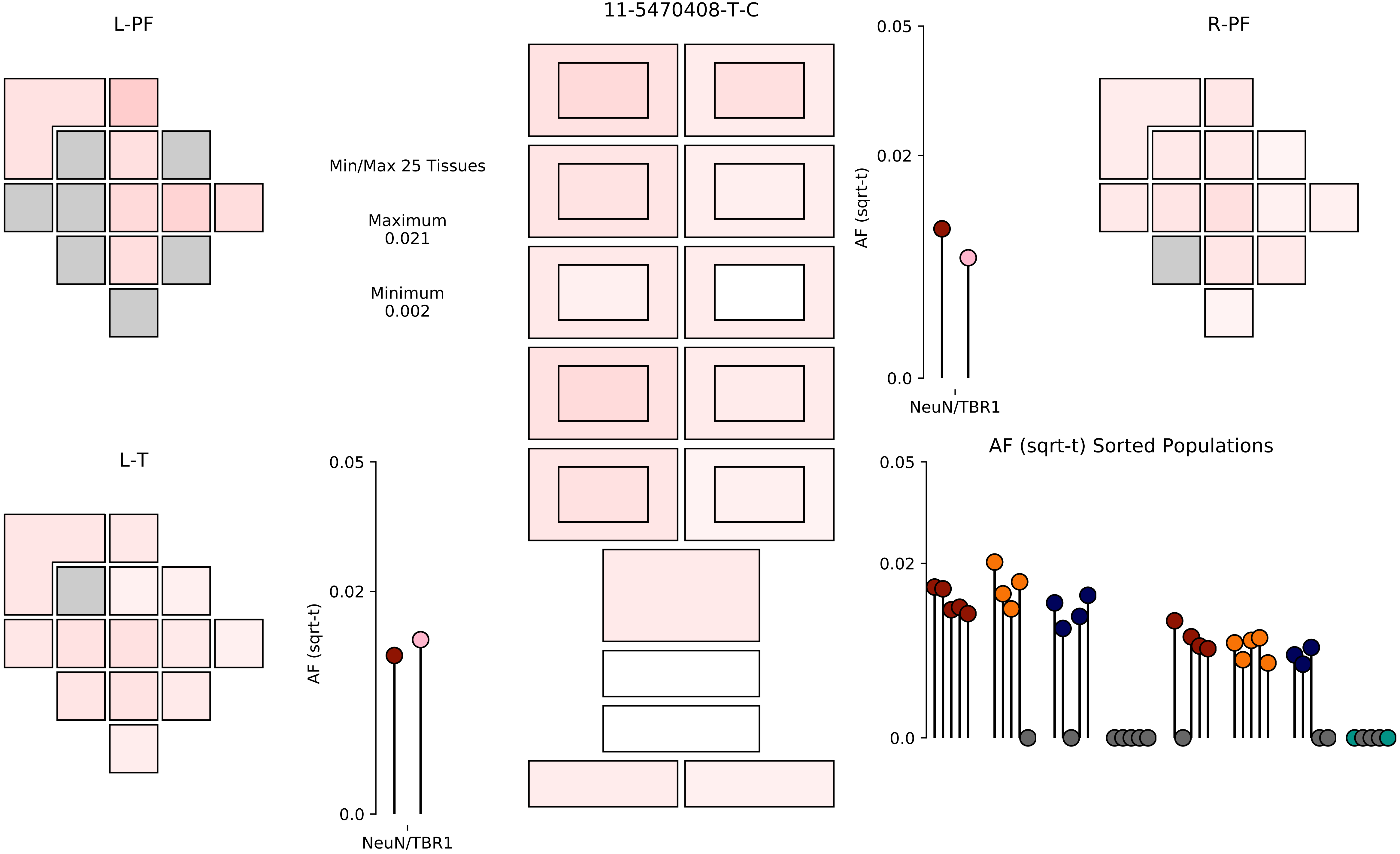
11-5470408-T-C
L-PF
R-PF
L-T
Min/Max 25 Tissues
Maximum
0.021
Minimum
0.002
AF (sqrt-t)
0.05
0.02
0.0
NeuN/TBR1
AF (sqrt-t) Sorted Populations

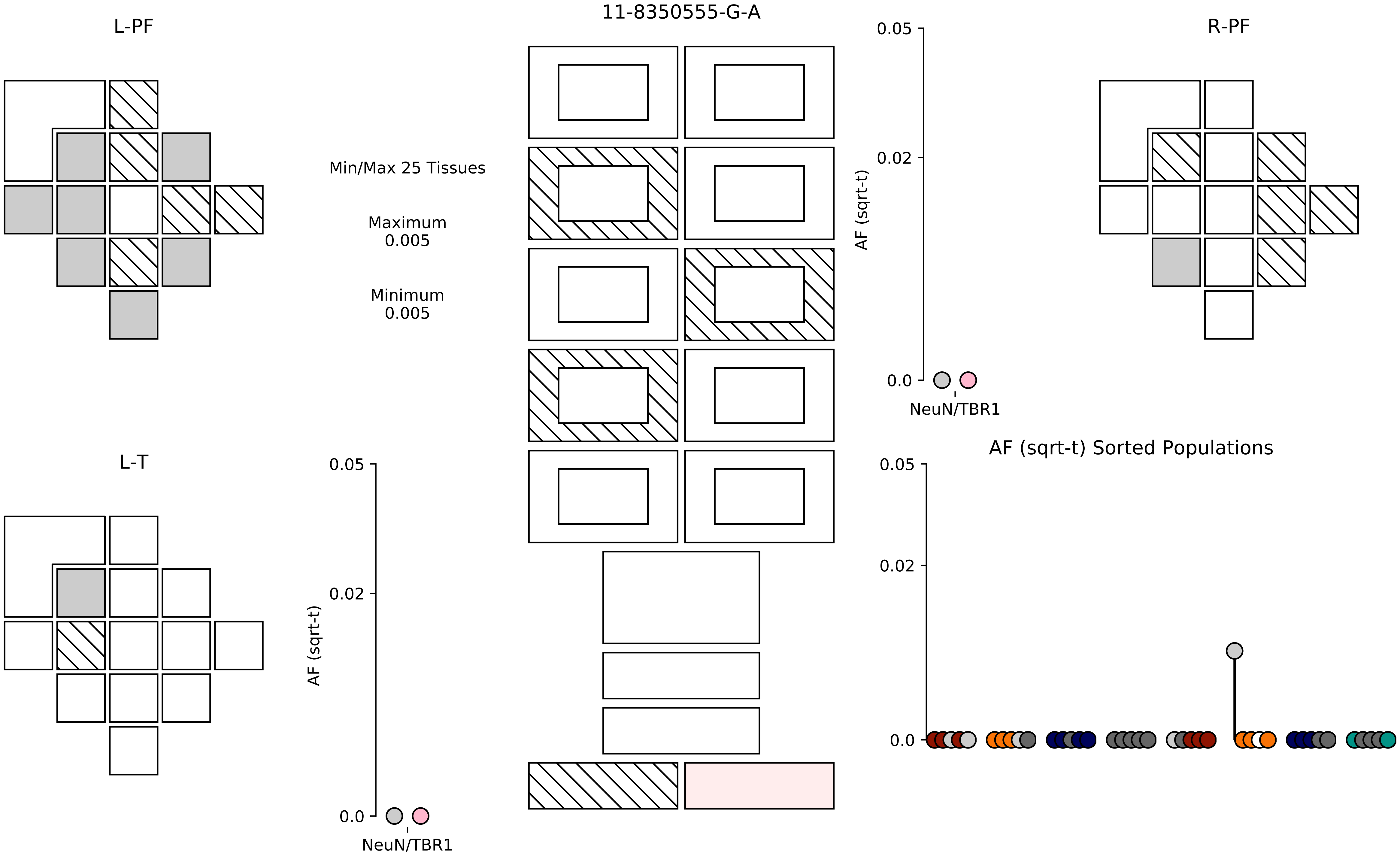
L-PF
11-8350555-G-A
R-PF
Min/Max 25 Tissues
Maximum
0.005
Minimum
0.005
AF (sqrt-t)
0.05
0.02
0.0
NeuN/TBR1
L-T
AF (sqrt-t) Sorted Populations

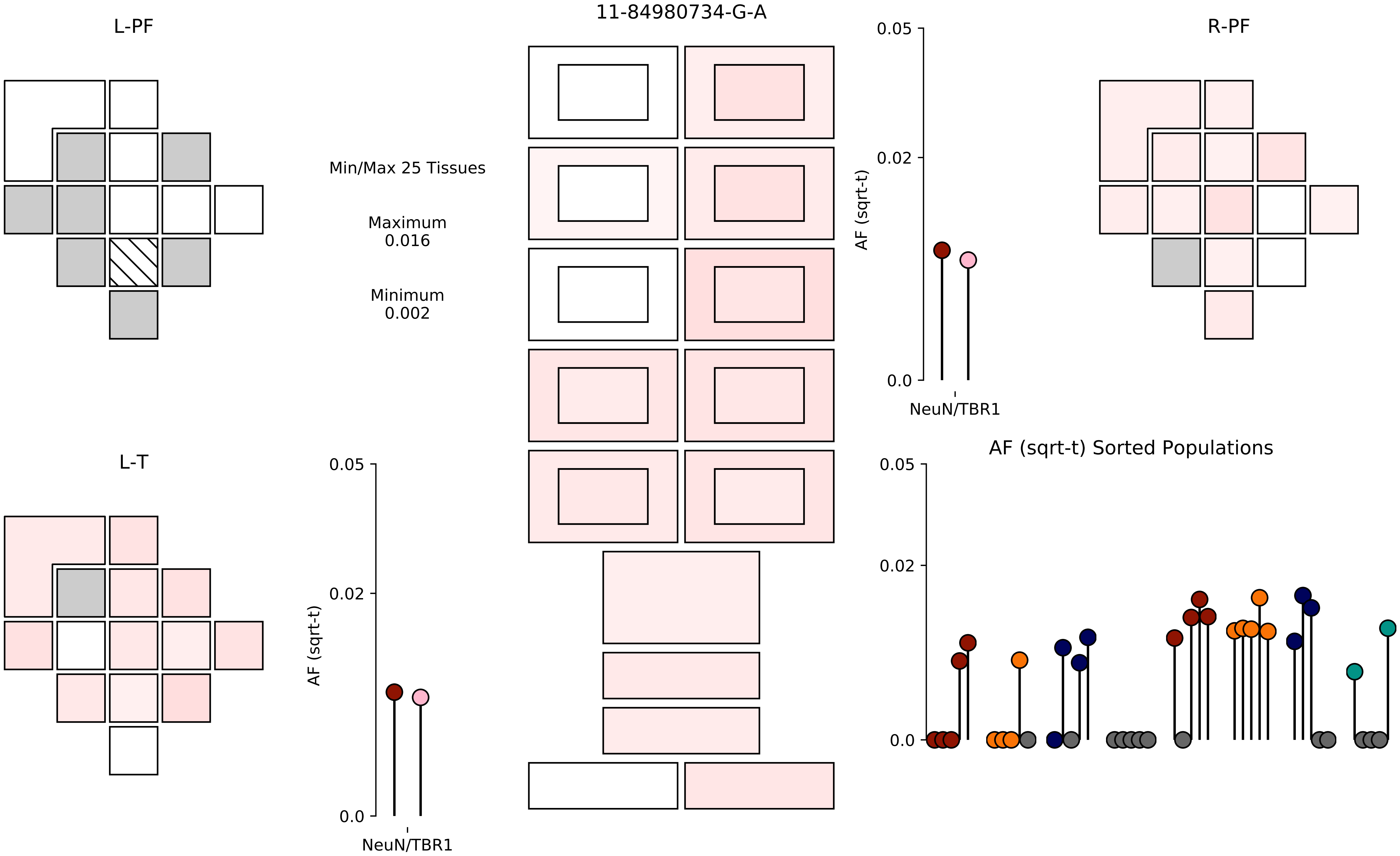

11-84980734-G-A
L-PF
R-PF
L-T
Min/Max 25 Tissues
Maximum
0.016
Minimum
0.002
AF (sqrt-t)
0.05
0.02
0.0
NeuN/TBR1
AF (sqrt-t)
0.05
0.02
0.0
NeuN/TBR1
AF (sqrt-t) Sorted Populations
0.05
0.02
0.0

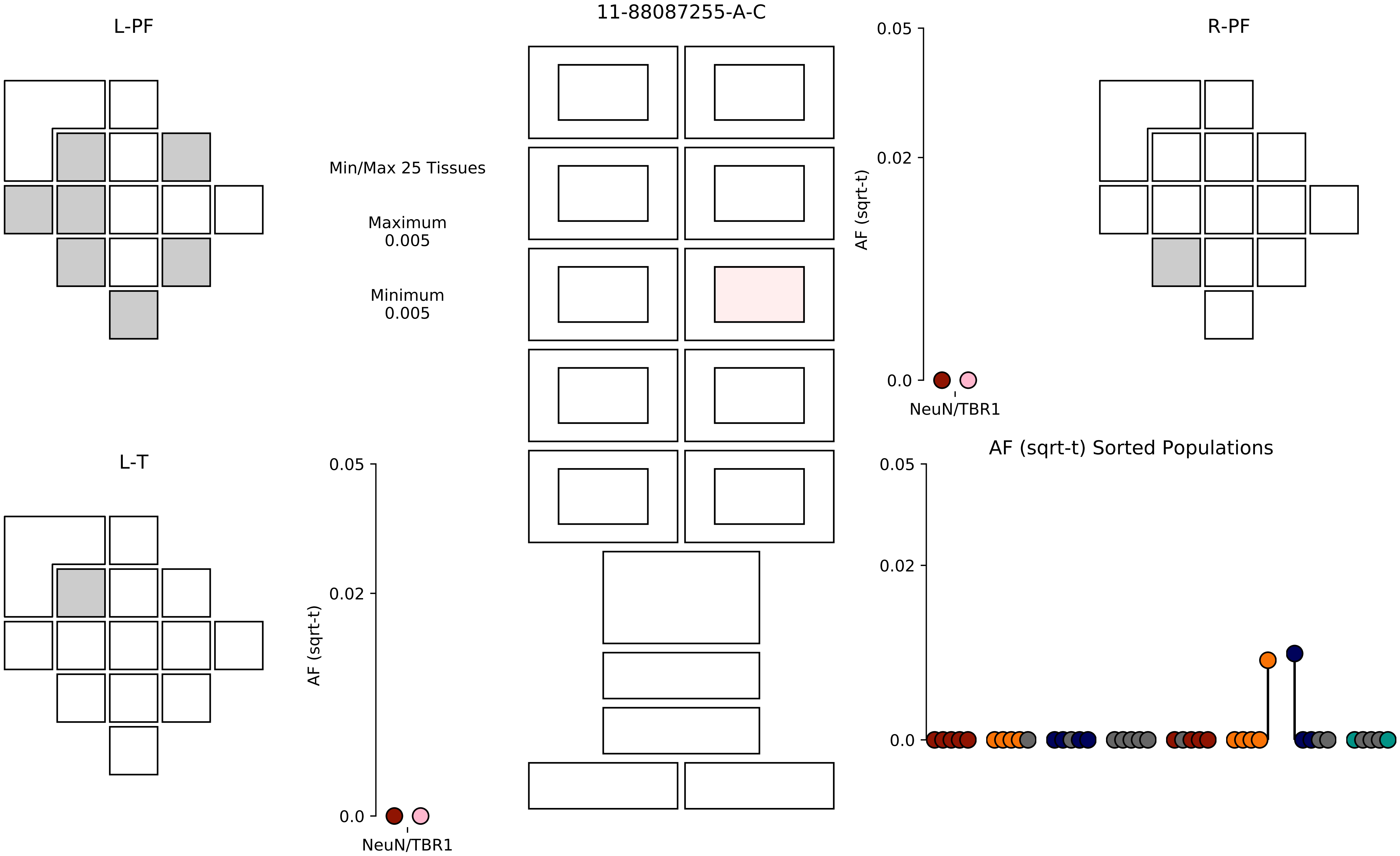

11-88087255-A-C
L-PF
R-PF
L-T
Min/Max 25 Tissues
Maximum
0.005
Minimum
0.005
AF (sqrt-t)
0.05
0.02
0.0
NeuN/TBR1
AF (sqrt-t) Sorted Populations

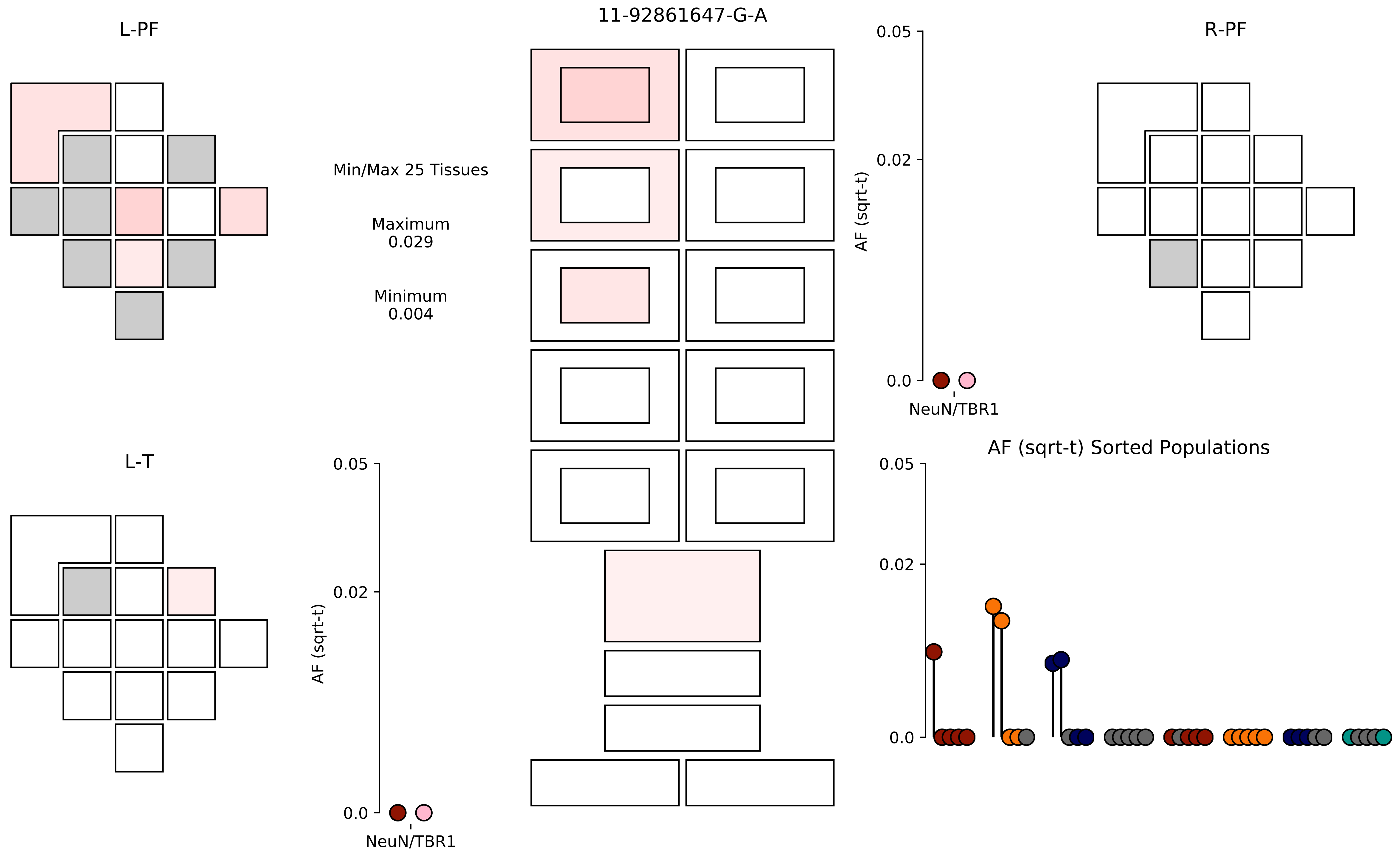
L-PF
11-92861647-G-A
R-PF
Min/Max 25 Tissues
Maximum
0.029
Minimum
0.004
0.05
0.02
0.0
AF (sqrt-t)
NeuN/TBR1
L-T
0.05
0.02
0.0
AF (sqrt-t)
NeuN/TBR1
AF (sqrt-t) Sorted Populations
0.05
0.02
0.0

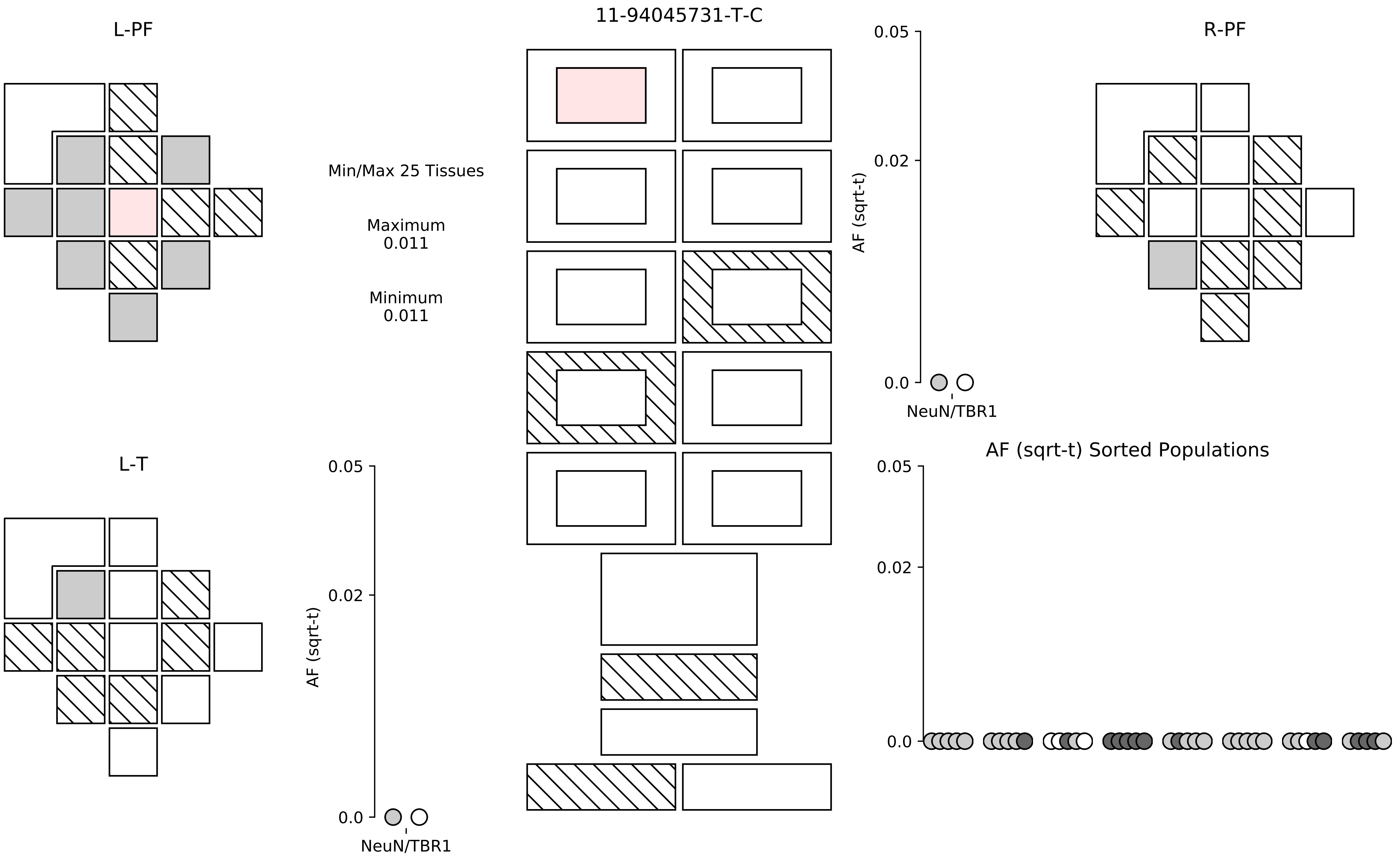

11-94045731-T-C
L-PF
R-PF
L-T
Min/Max 25 Tissues
Maximum
0.011
Minimum
0.011
AF (sqrt-t)
0.05
0.02
0.0
NeuN/TBR1
AF (sqrt-t)
0.05
0.02
0.0
NeuN/TBR1
AF (sqrt-t) Sorted Populations
0.05
0.02
0.0

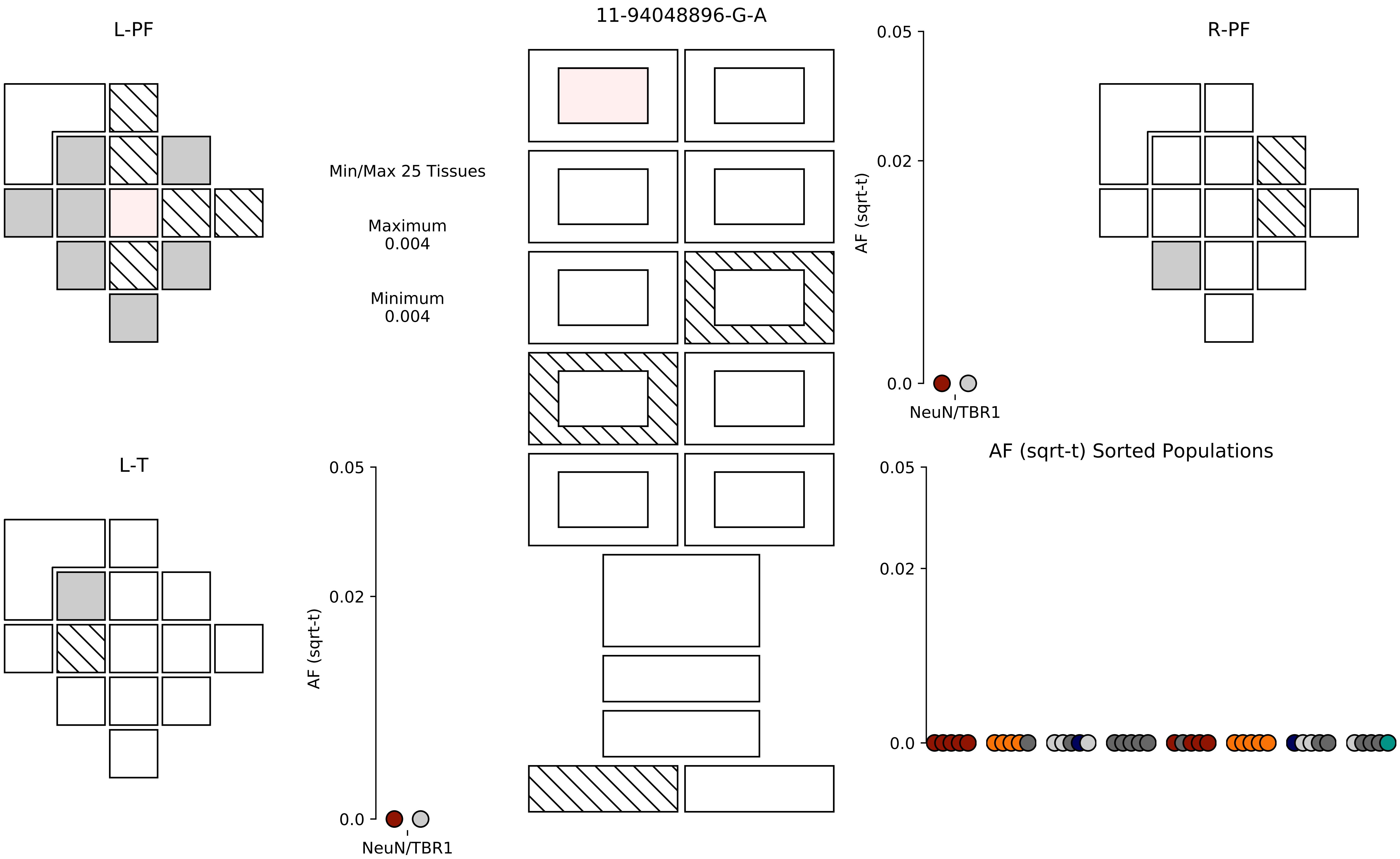
11-94048896-G-A
L-PF
R-PF
L-T
Min/Max 25 Tissues
Maximum
0.004
Minimum
0.004
AF (sqrt-t)
0.05
0.02
0.0
NeuN/TBR1
AF (sqrt-t)
0.05
0.02
0.0
NeuN/TBR1
AF (sqrt-t) Sorted Populations
0.05
0.02
0.0

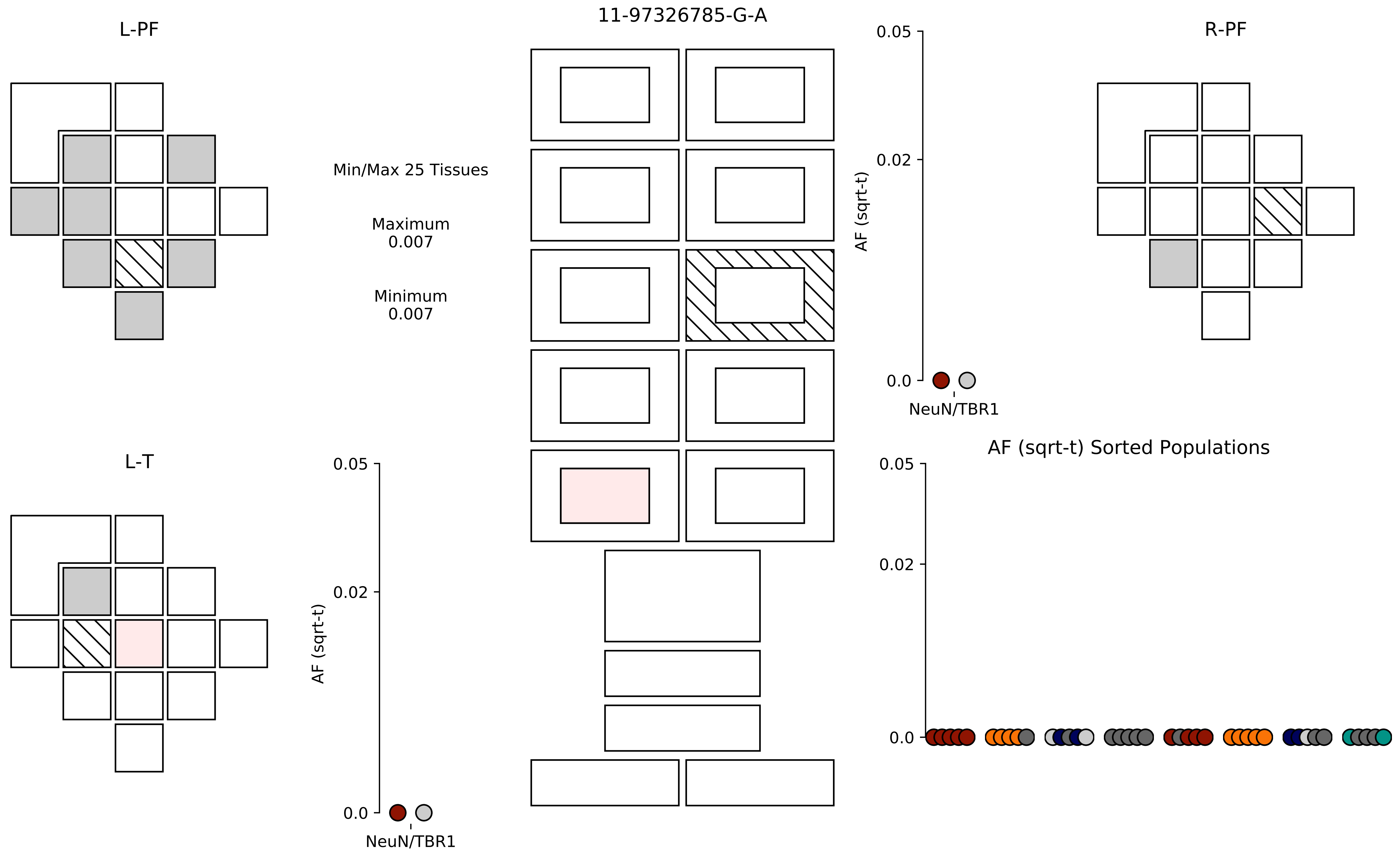
L-PF
11-97326785-G-A
R-PF
Min/Max 25 Tissues
Maximum
0.007
Minimum
0.007
0.05
0.02
0.0
AF (sqrt-t)
NeuN/TBR1
L-T
AF (sqrt-t) Sorted Populations
0.05
0.02
0.0
AF (sqrt-t)
NeuN/TBR1
0.05
0.02
0.0

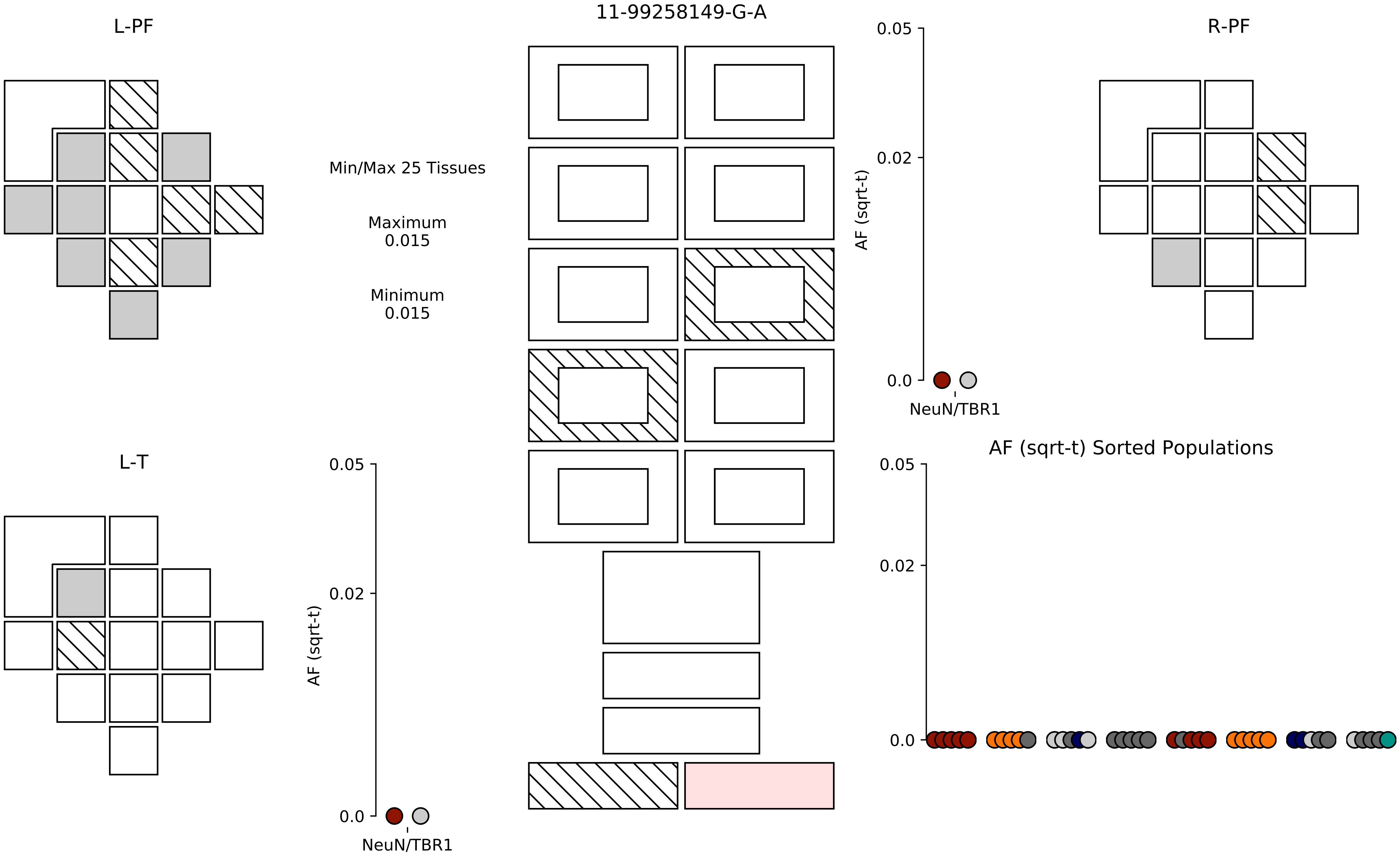

L-PF
11-99258149-G-A
R-PF
Min/Max 25 Tissues
Maximum
0.015
Minimum
0.015
0.05
0.02
0.0
AF (sqrt-t)
NeuN/TBR1
L-T
0.05
0.02
0.0
AF (sqrt-t)
NeuN/TBR1
AF (sqrt-t) Sorted Populations
0.05
0.02
0.0

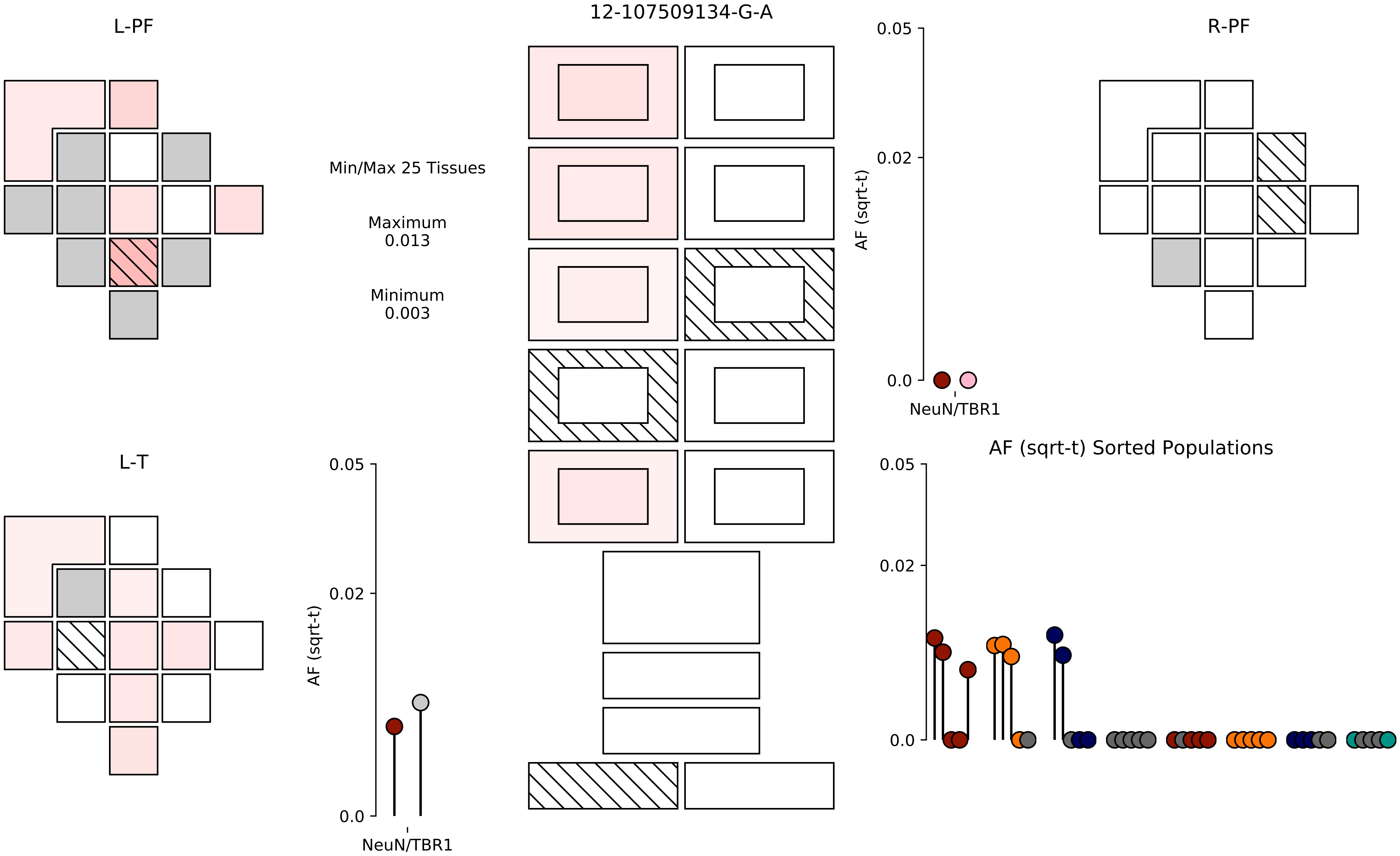

12-107509134-G-A
L-PF
R-PF
L-T
Min/Max 25 Tissues
Maximum
0.013
Minimum
0.003
0.05
0.02
0.0
AF (sqrt-t)
NeuN/TBR1
AF (sqrt-t) Sorted Populations

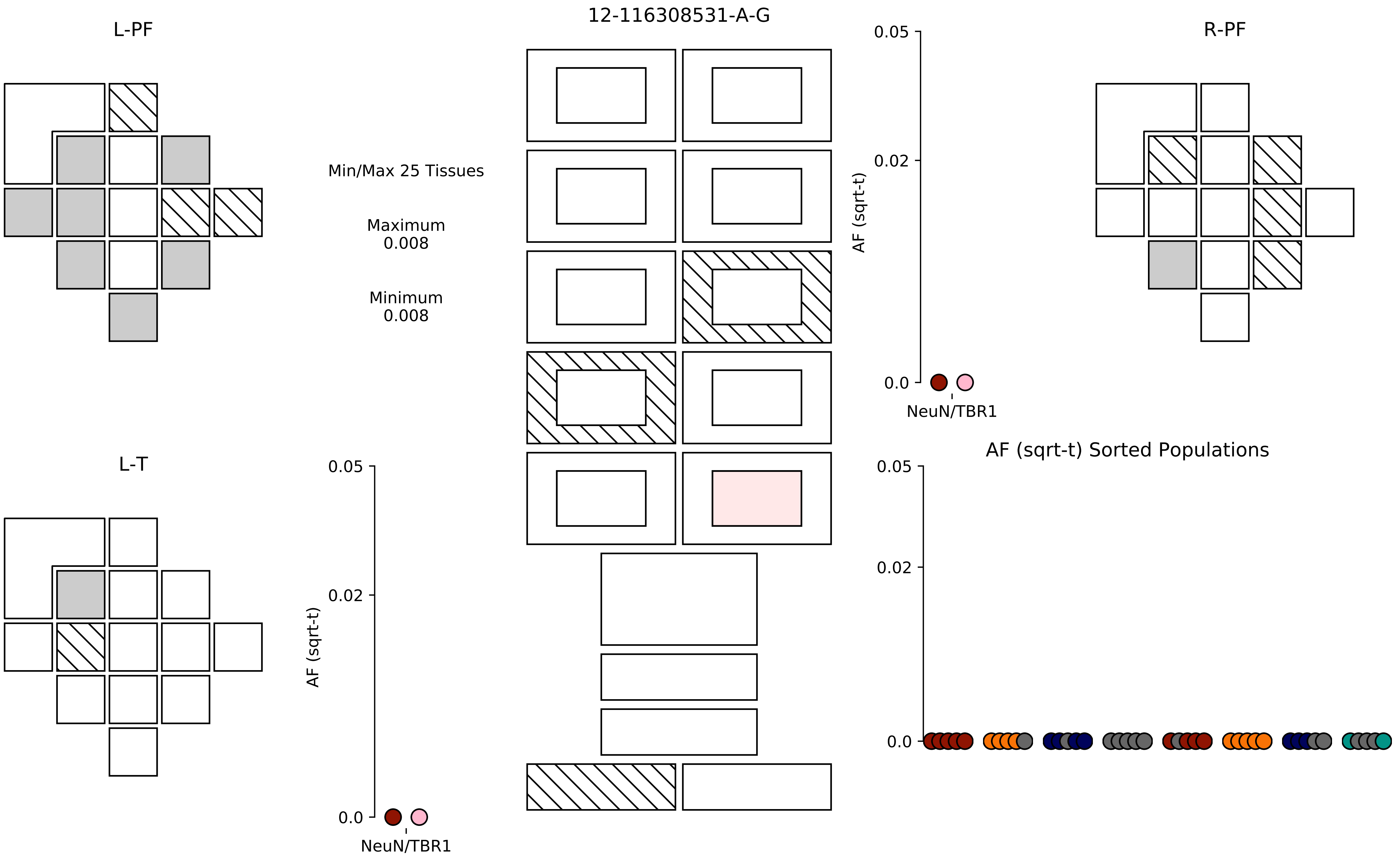

L-PF
12-116308531-A-G
R-PF
Min/Max 25 Tissues
Maximum
0.008
Minimum
0.008
0.05
0.02
0.0
AF (sqrt-t)
NeuN/TBR1
L-T
0.05
0.02
0.0
AF (sqrt-t)
NeuN/TBR1
AF (sqrt-t) Sorted Populations
0.05
0.02
0.0

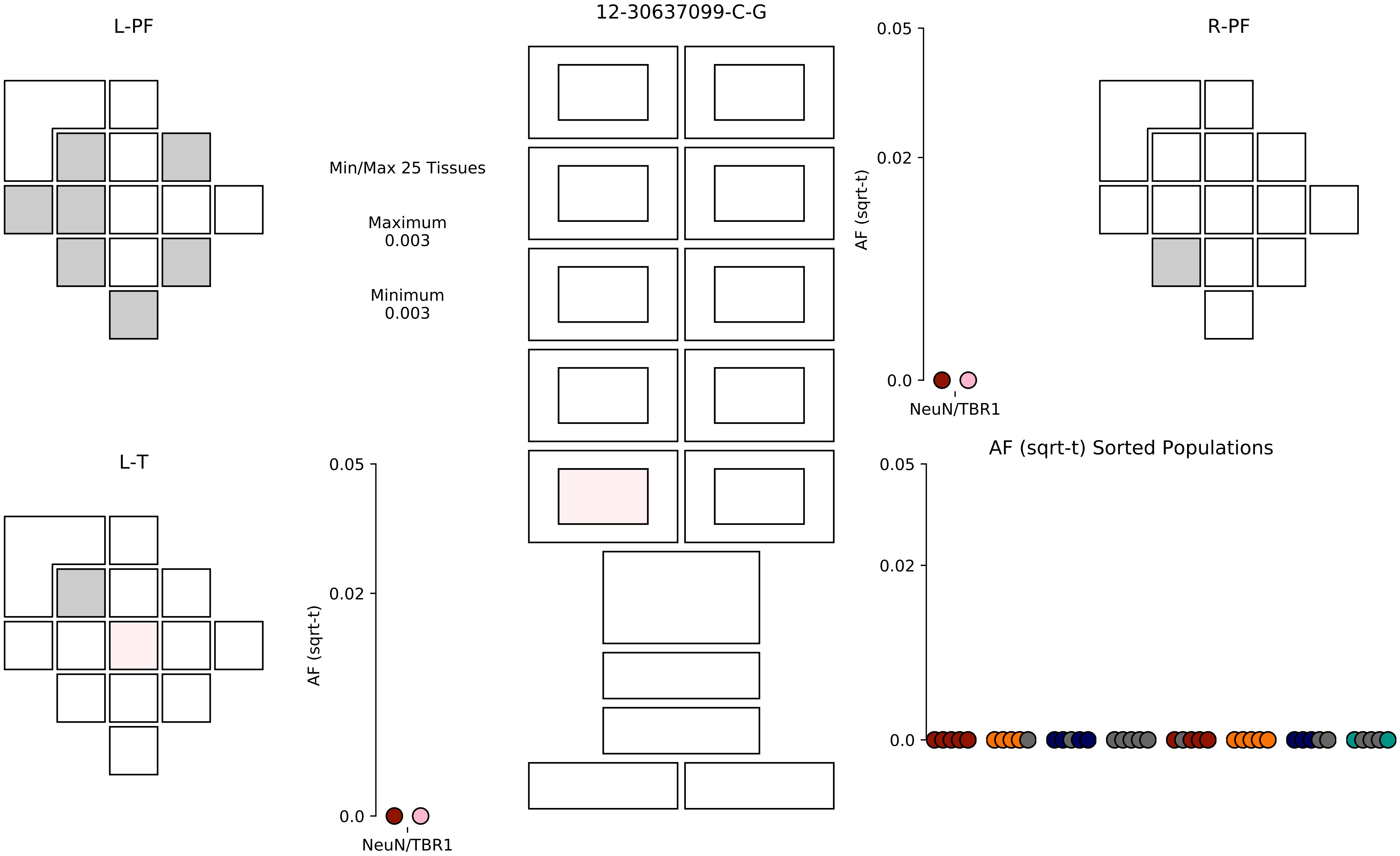

12-30637099-C-G
L-PF
R-PF
L-T
Min/Max 25 Tissues
Maximum
0.003
Minimum
0.003
AF (sqrt-t)
0.05
0.02
0.0
NeuN/TBR1
AF (sqrt-t) Sorted Populations

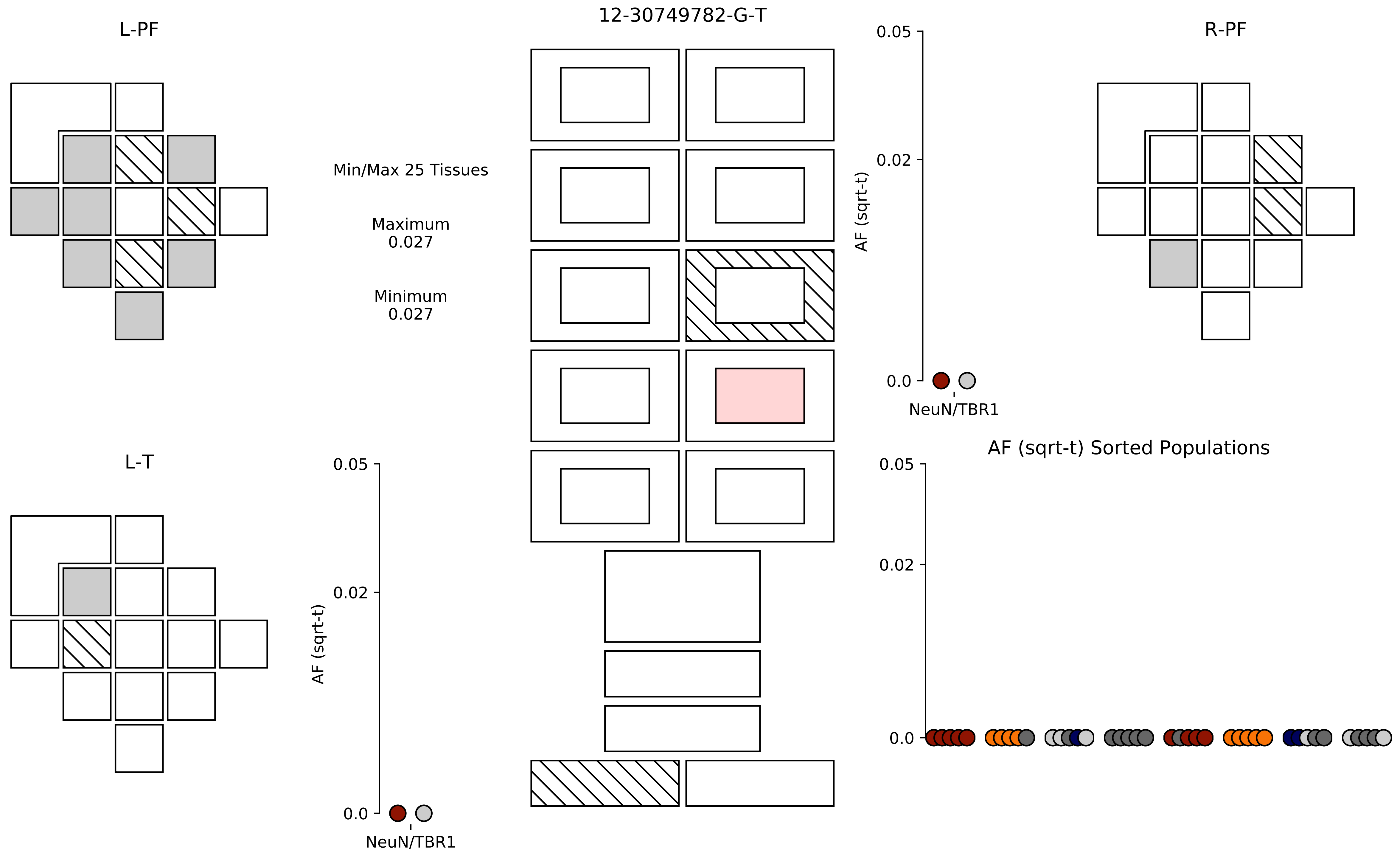

12-30749782-G-T
L-PF
R-PF
L-T
Min/Max 25 Tissues
Maximum
0.027
Minimum
0.027
AF (sqrt-t)
0.05
0.02
0.0
NeuN/TBR1
AF (sqrt-t) Sorted Populations

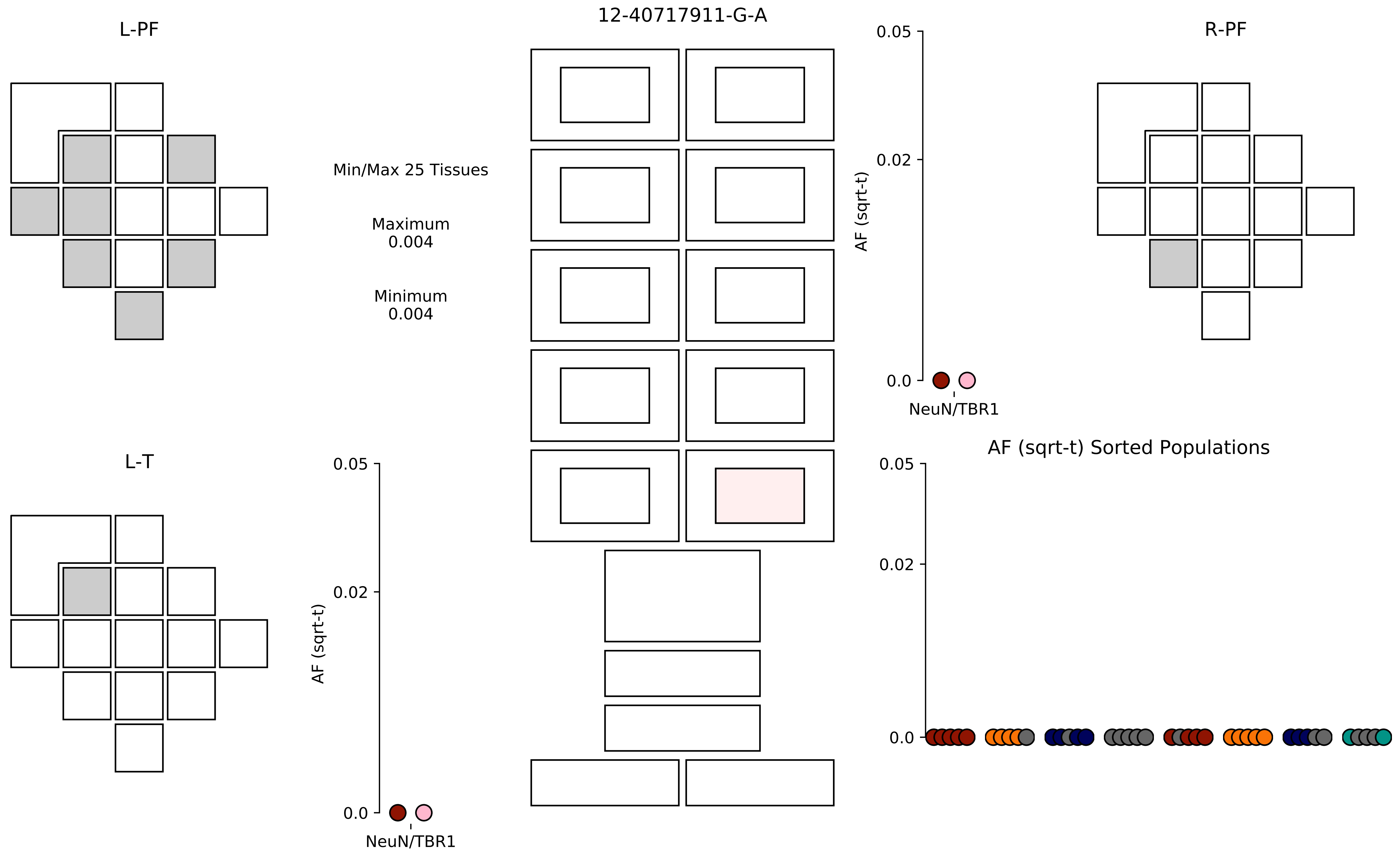

12-40717911-G-A
L-PF
R-PF
L-T
Min/Max 25 Tissues
Maximum
0.004
Minimum
0.004
AF (sqrt-t)
0.05
0.02
0.0
NeuN/TBR1
AF (sqrt-t) Sorted Populations

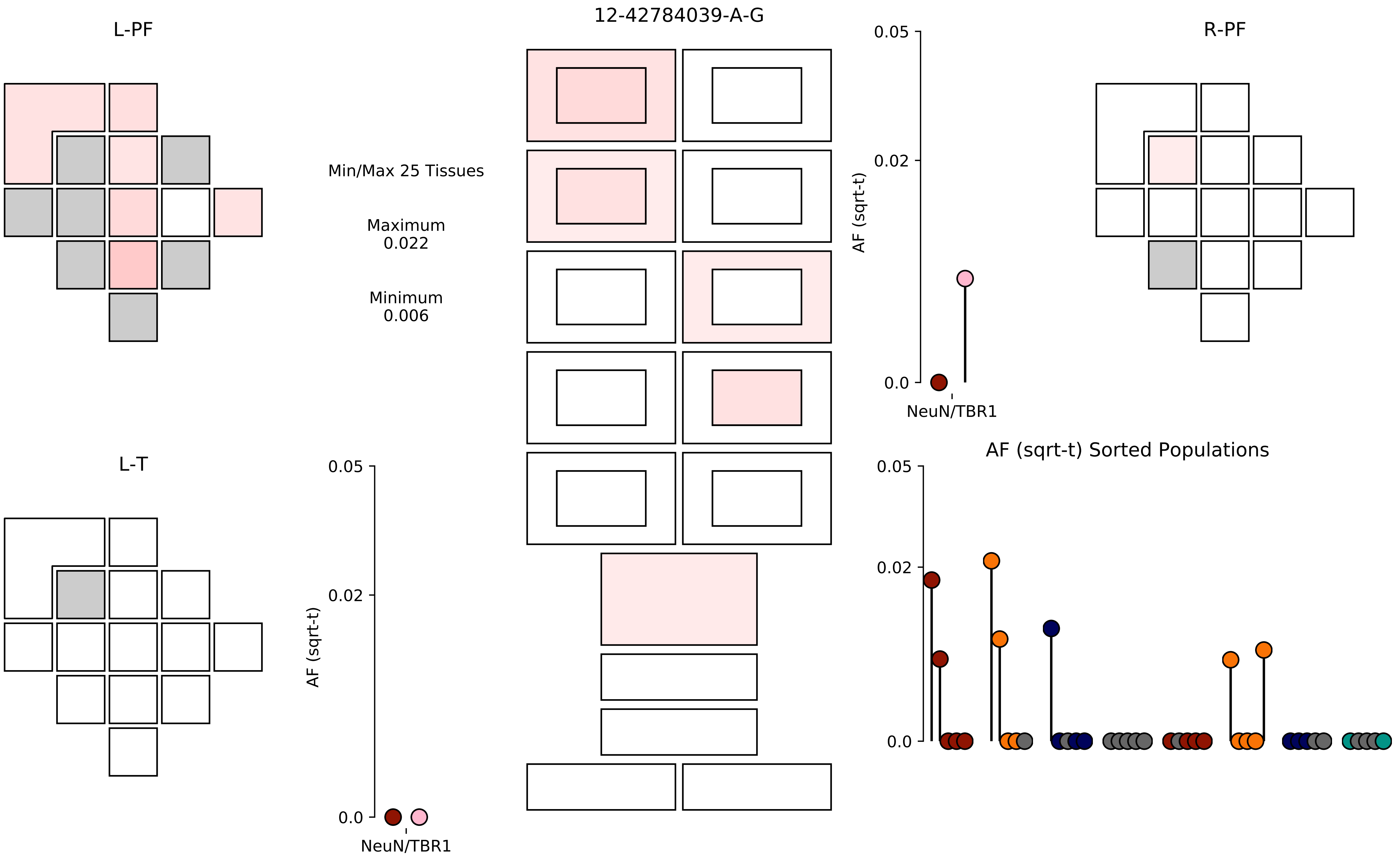
12-42784039-A-G
L-PF
R-PF
L-T
Min/Max 25 Tissues
Maximum
0.022
Minimum
0.006
AF (sqrt-t)
0.05
0.02
0.0
NeuN/TBR1
AF (sqrt-t) Sorted Populations

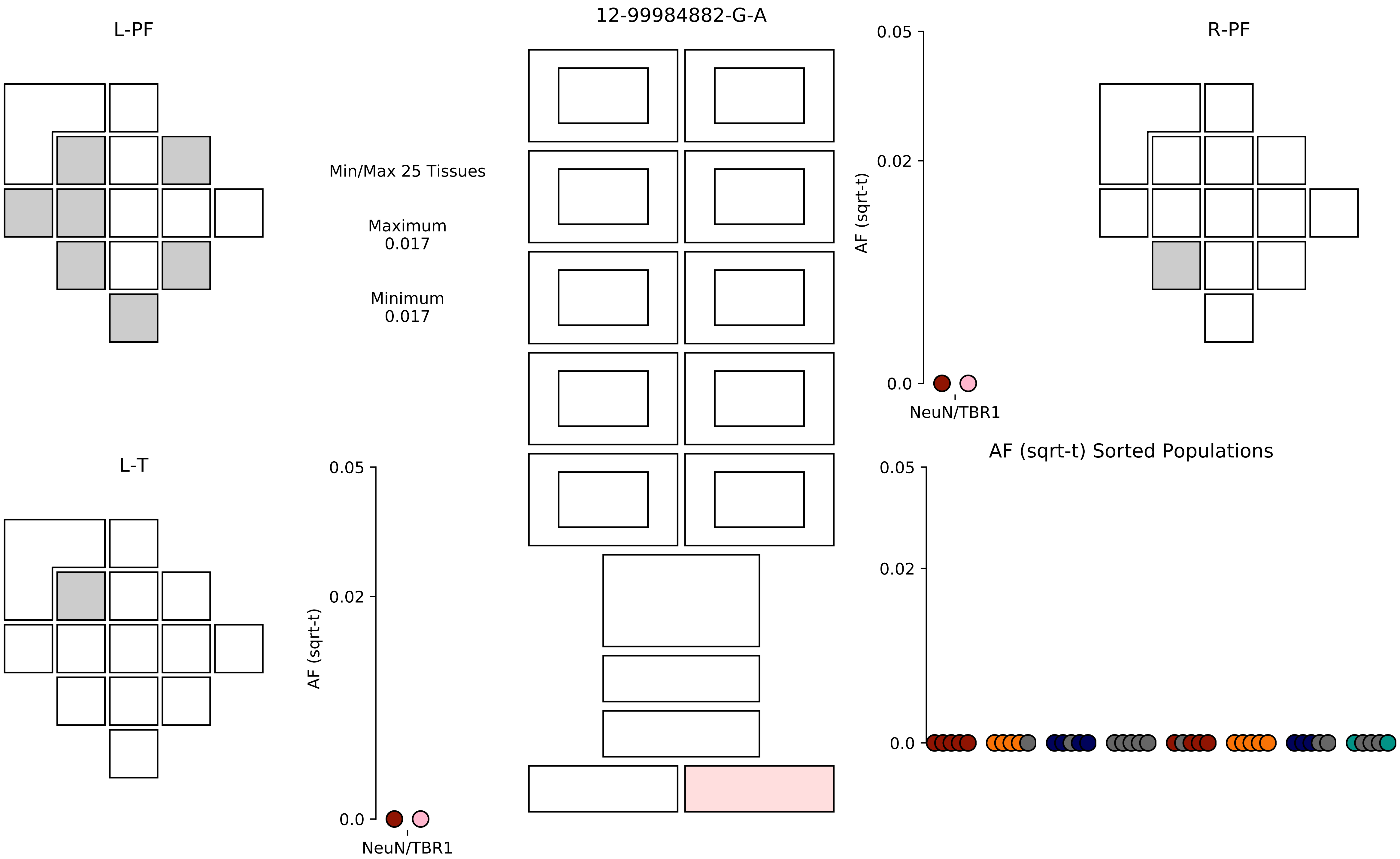

12-99984882-G-A
L-PF
R-PF
L-T
Min/Max 25 Tissues
Maximum
0.017
Minimum
0.017
AF (sqrt-t)
0.05
0.02
0.0
NeuN/TBR1
AF (sqrt-t) Sorted Populations

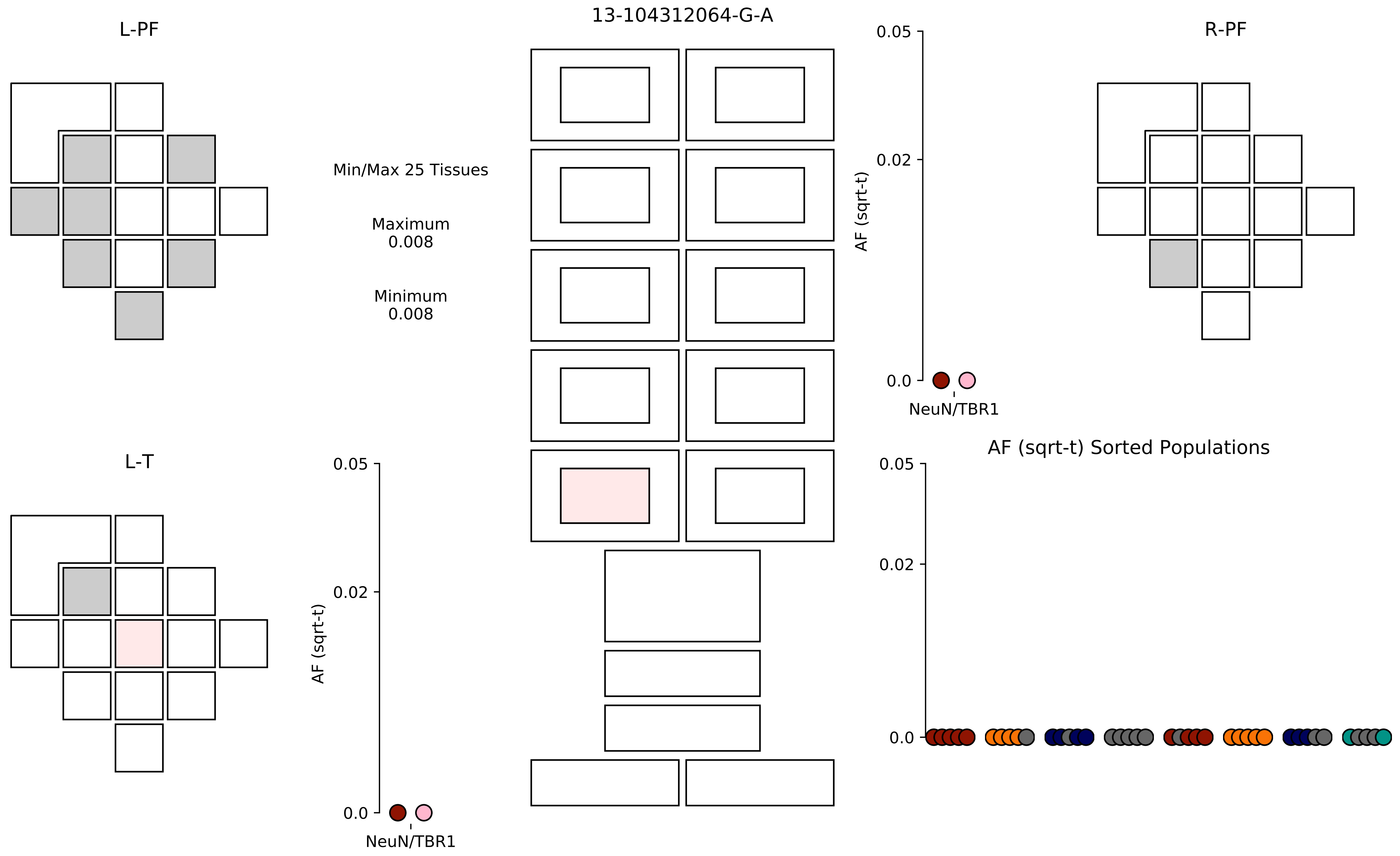

13-104312064-G-A
L-PF
R-PF
L-T
Min/Max 25 Tissues
Maximum
0.008
Minimum
0.008
AF (sqrt-t)
0.05
0.02
0.0
NeuN/TBR1
AF (sqrt-t)
0.05
0.02
0.0
NeuN/TBR1
AF (sqrt-t) Sorted Populations
0.05
0.02
0.0

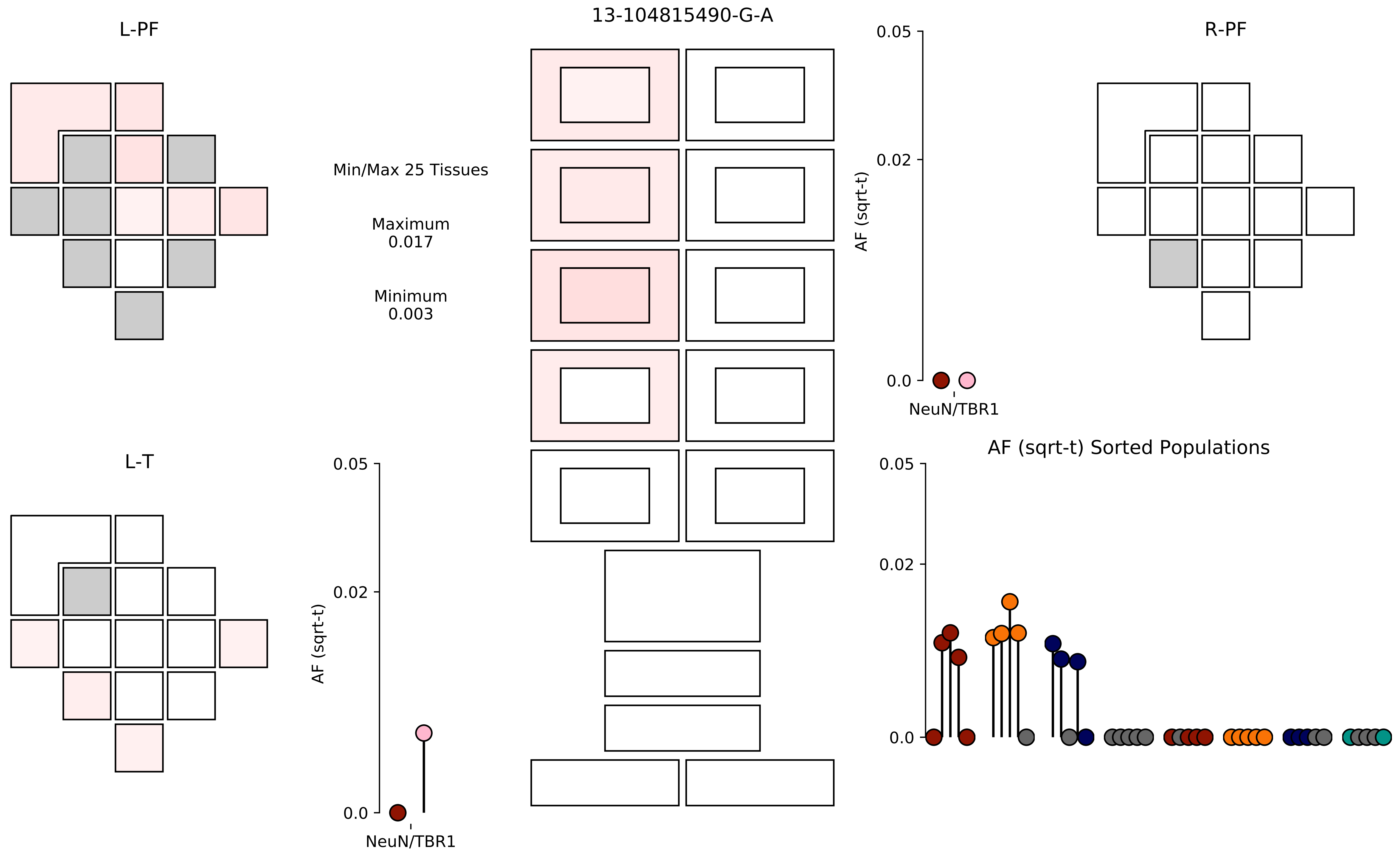

13-104815490-G-A
L-PF
R-PF
L-T
Min/Max 25 Tissues
Maximum
0.017
Minimum
0.003
0.05
0.02
0.0
AF (sqrt-t)
NeuN/TBR1
AF (sqrt-t) Sorted Populations

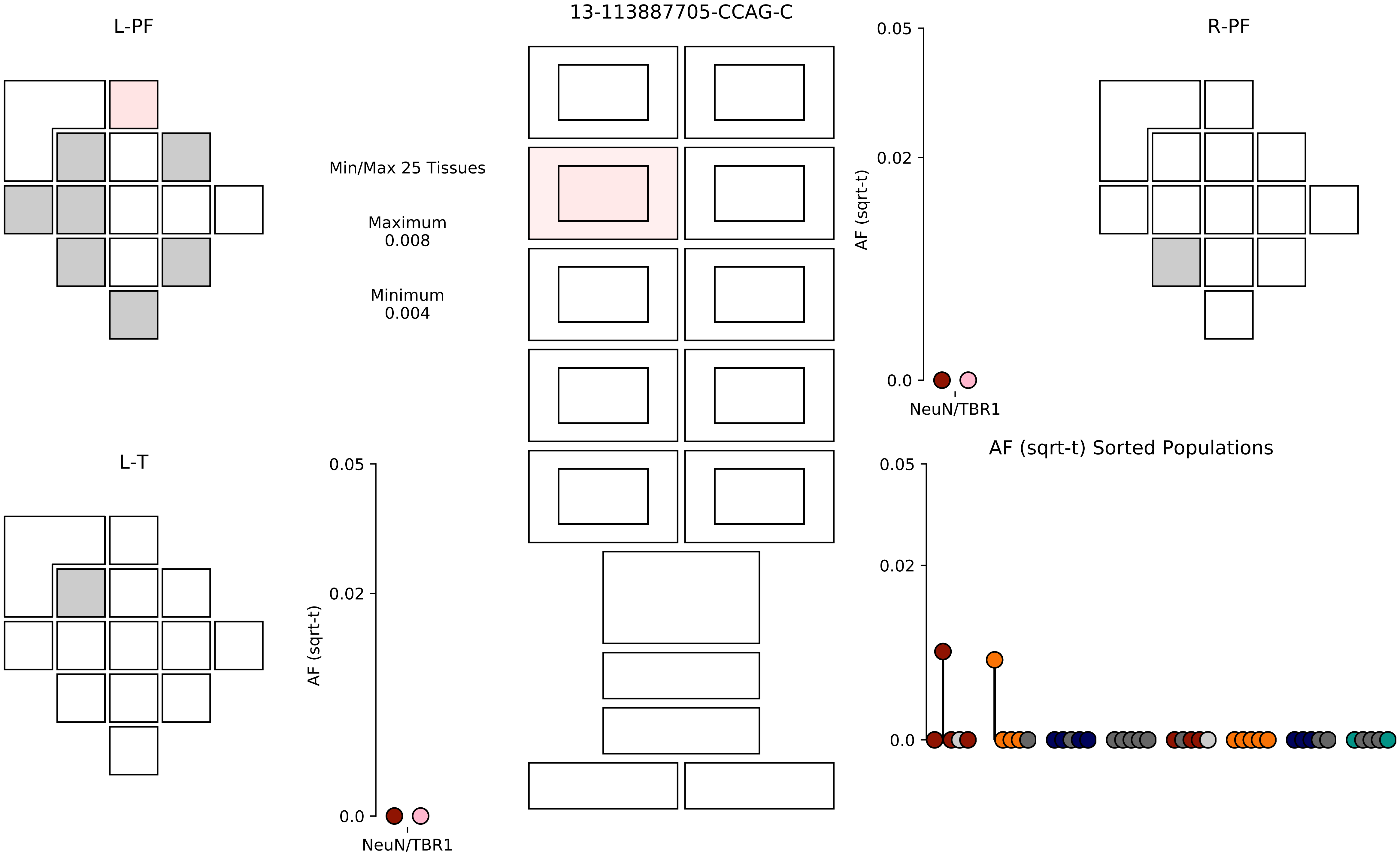
L-PF
13-113887705-CCAG-C
R-PF
Min/Max 25 Tissues
Maximum
0.008
Minimum
0.004
0.05
0.02
0.0
AF (sqrt-t)
NeuN/TBR1
L-T
AF (sqrt-t) Sorted Populations

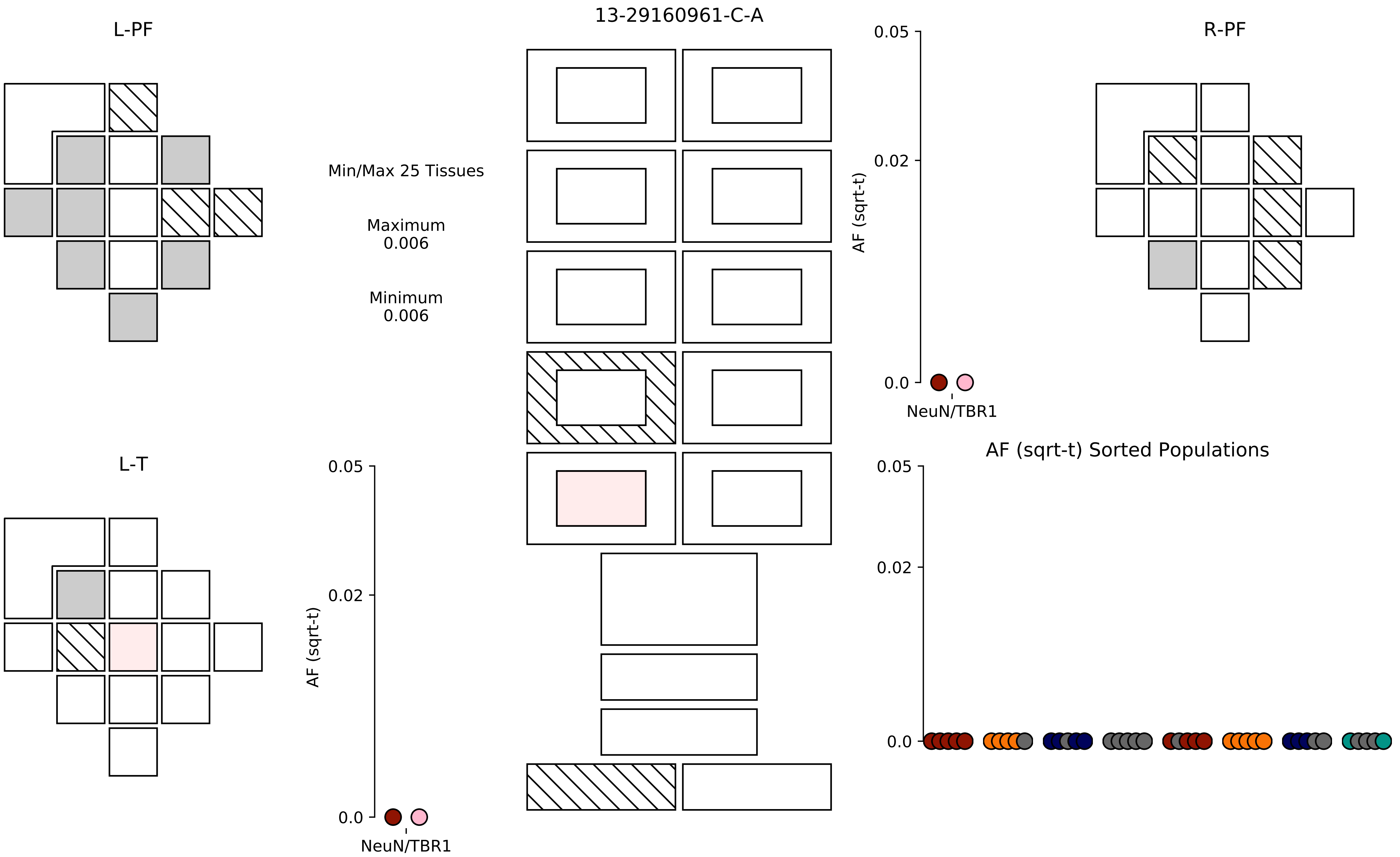

13-29160961-C-A
L-PF
R-PF
L-T
Min/Max 25 Tissues
Maximum
0.006
Minimum
0.006
AF (sqrt-t)
0.05
0.02
0.0
NeuN/TBR1
AF (sqrt-t) Sorted Populations

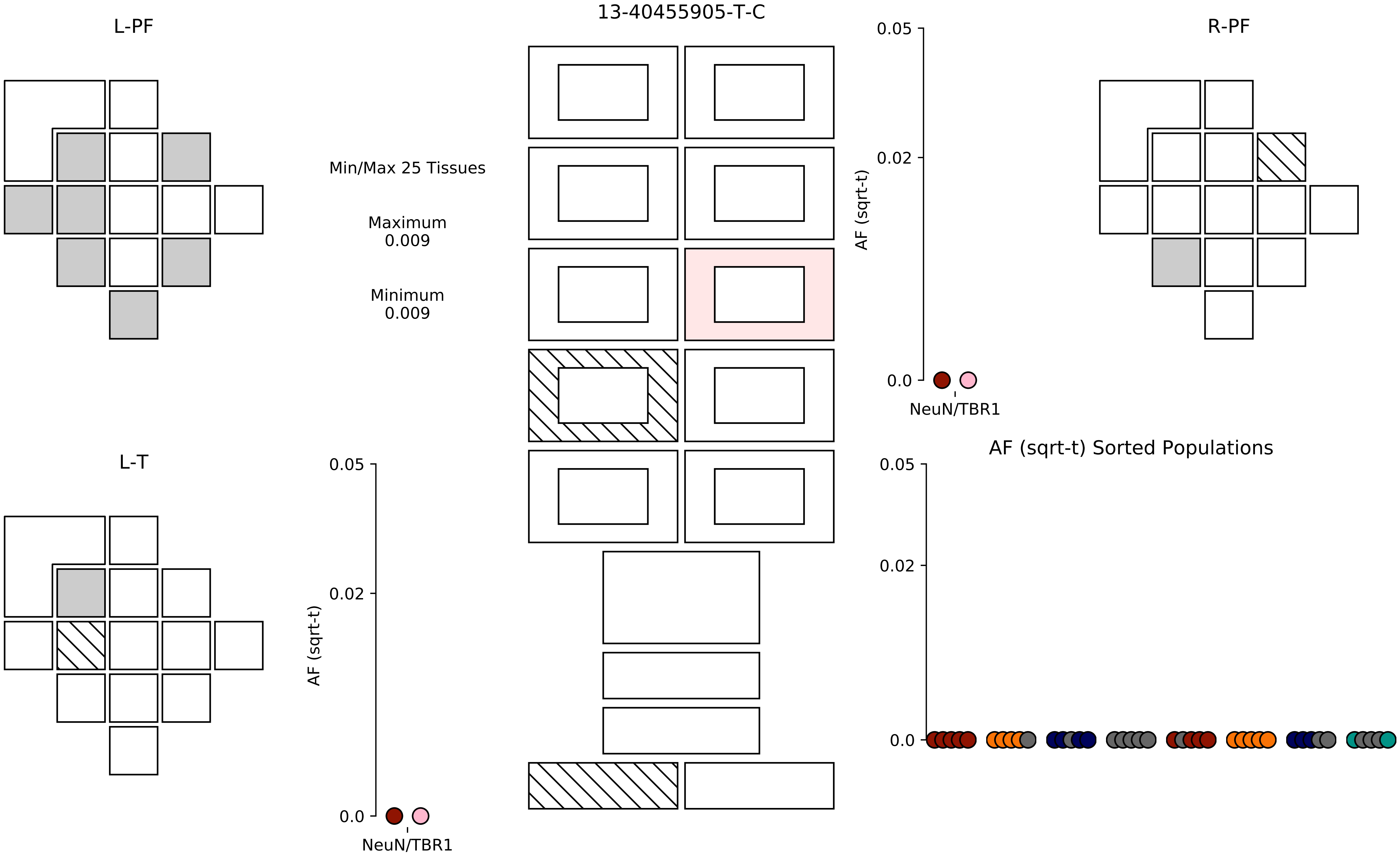
L-PF
13-40455905-T-C
R-PF
Min/Max 25 Tissues
Maximum
0.009
Minimum
0.009
AF (sqrt-t)
0.05
0.02
0.0
NeuN/TBR1
L-T
AF (sqrt-t)
0.05
0.02
0.0
NeuN/TBR1
AF (sqrt-t) Sorted Populations
0.05
0.02
0.0

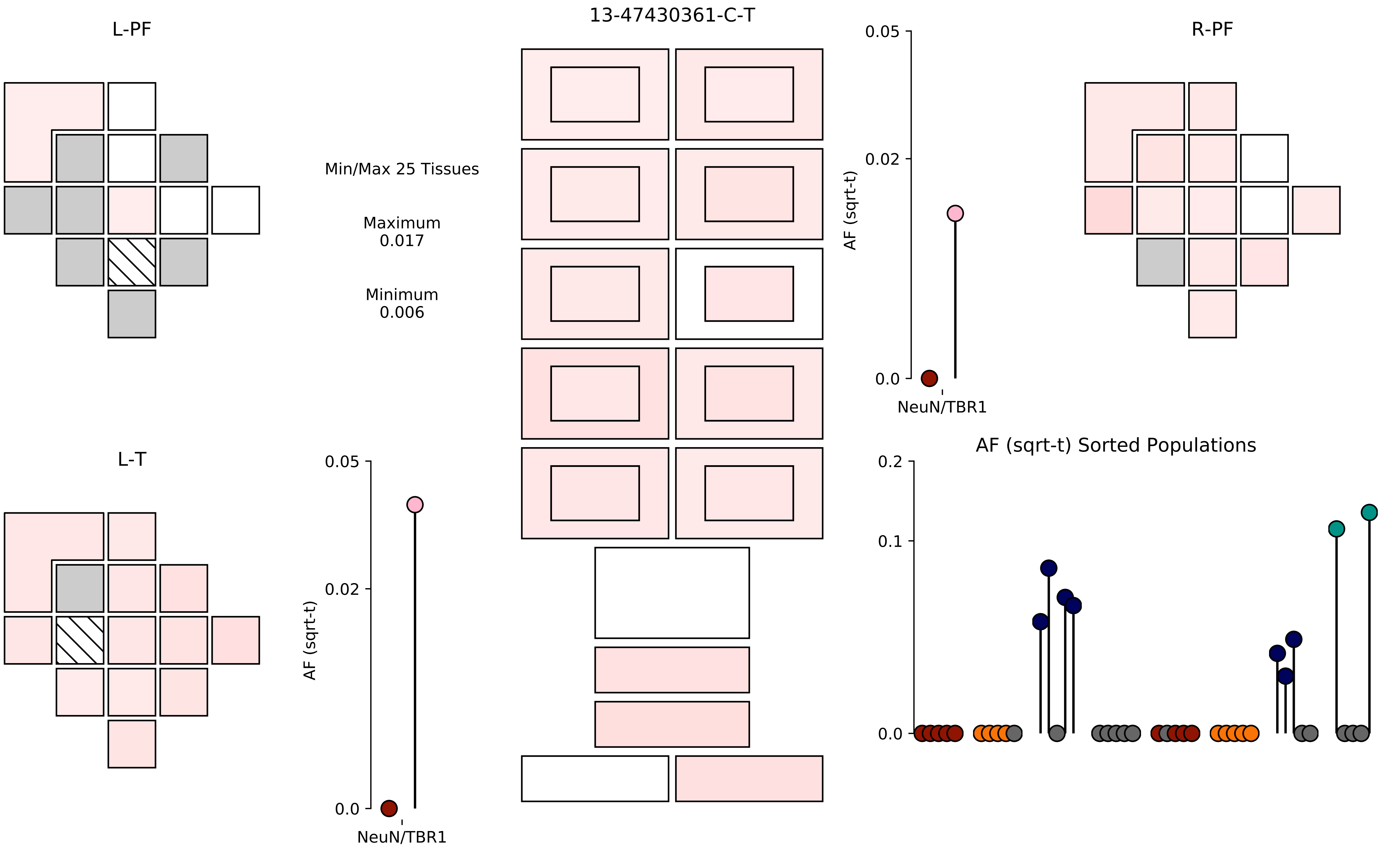

13-47430361-C-T
L-PF
R-PF
L-T
Min/Max 25 Tissues
Maximum
0.017
Minimum
0.006
0.05
0.02
0.0
AF (sqrt-t)
NeuN/TBR1
0.05
0.02
0.0
AF (sqrt-t)
NeuN/TBR1
AF (sqrt-t) Sorted Populations
0.2
0.1
0.0

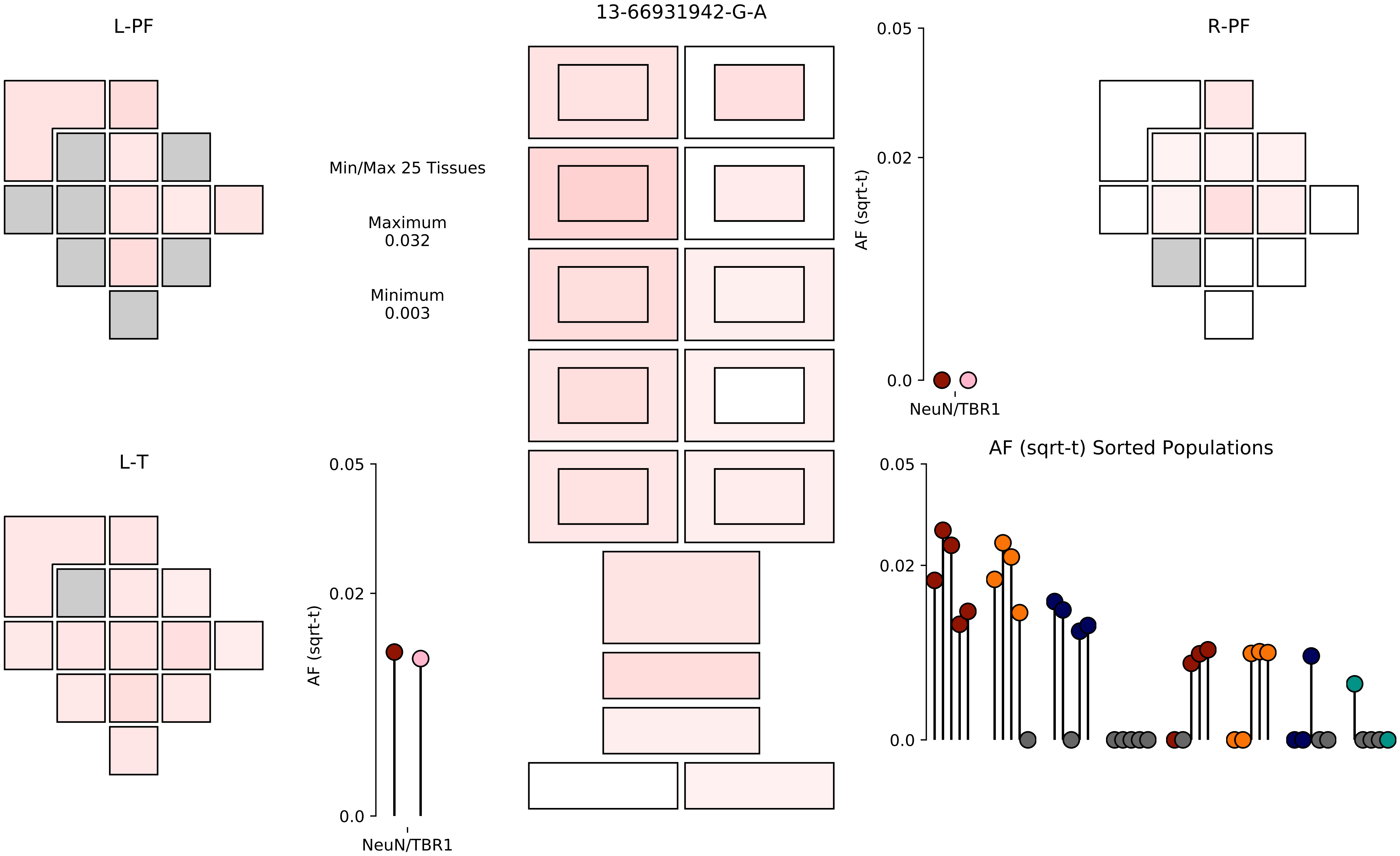

13-66931942-G-A
L-PF
R-PF
L-T
Min/Max 25 Tissues
Maximum
0.032
Minimum
0.003
AF (sqrt-t)
0.05
0.02
0.0
NeuN/TBR1
AF (sqrt-t) Sorted Populations

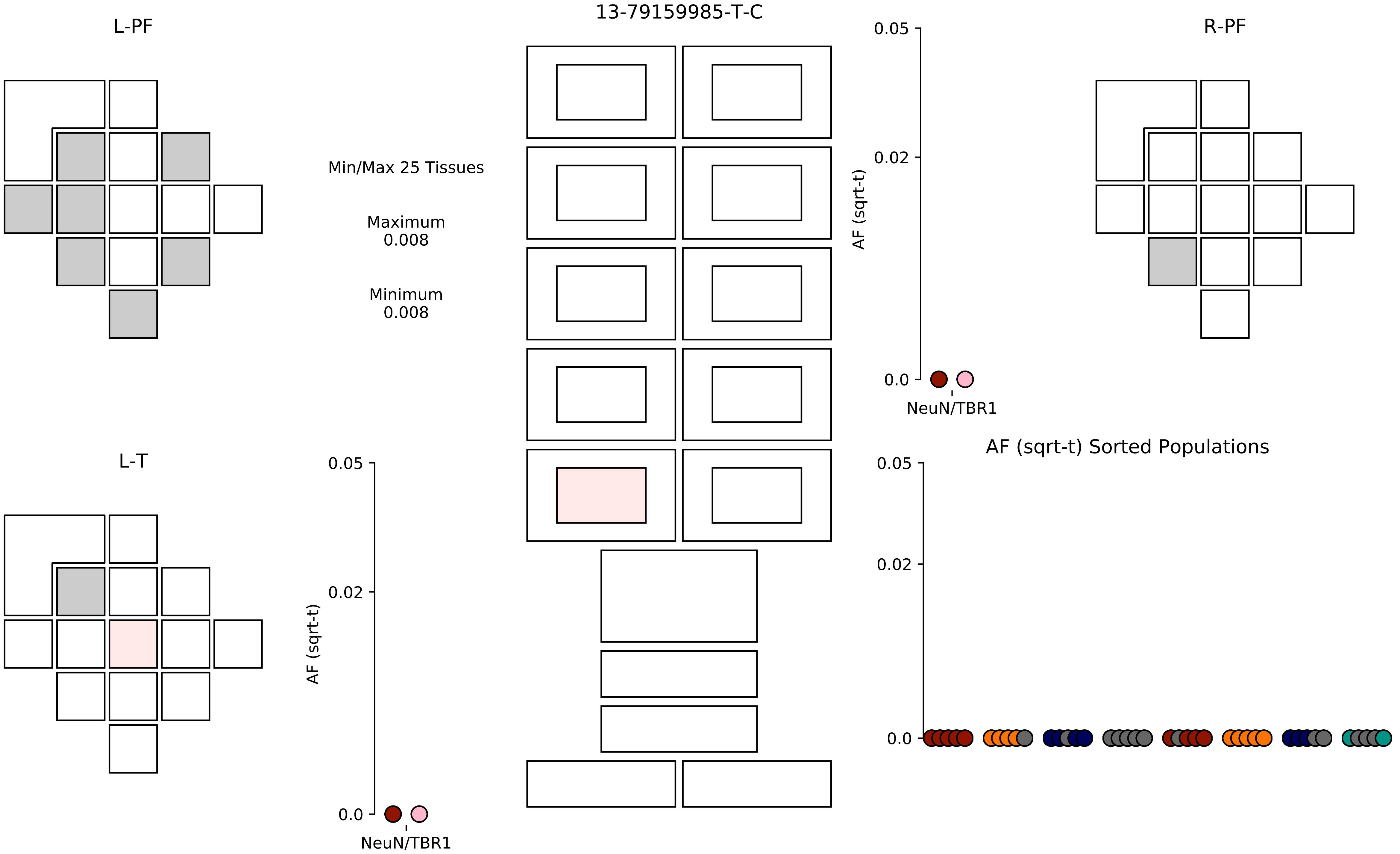

13-79159985-T-C
L-PF
R-PF
L-T
Min/Max 25 Tissues
Maximum
0.008
Minimum
0.008
AF (sqrt-t)
0.05
0.02
0.0
NeuN/TBR1
AF (sqrt-t)
0.05
0.02
0.0
NeuN/TBR1
AF (sqrt-t) Sorted Populations
0.05
0.02
0.0

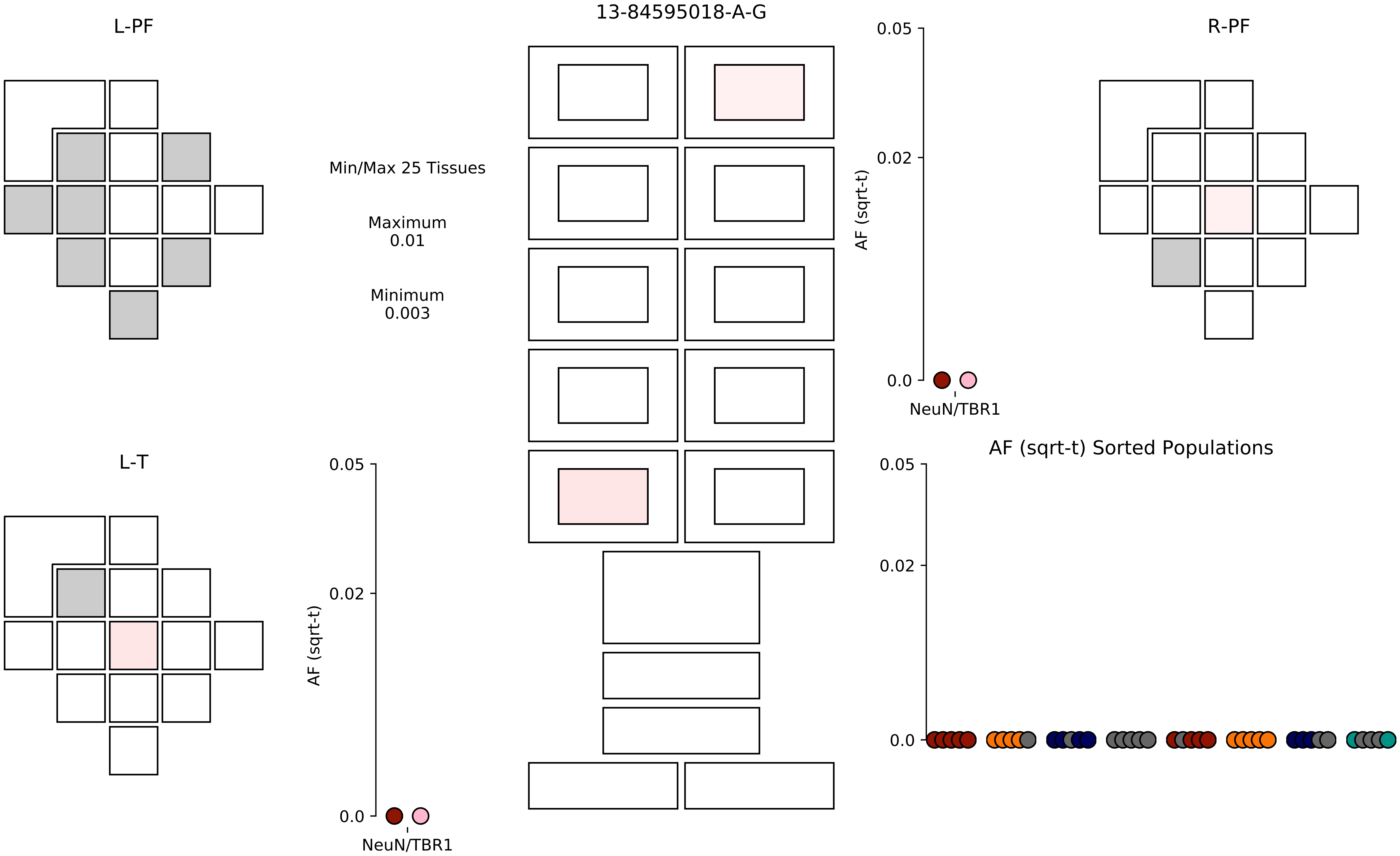

13-84595018-A-G
L-PF
R-PF
L-T
Min/Max 25 Tissues
Maximum
0.01
Minimum
0.003
AF (sqrt-t)
0.05
0.02
0.0
NeuN/TBR1
AF (sqrt-t) Sorted Populations

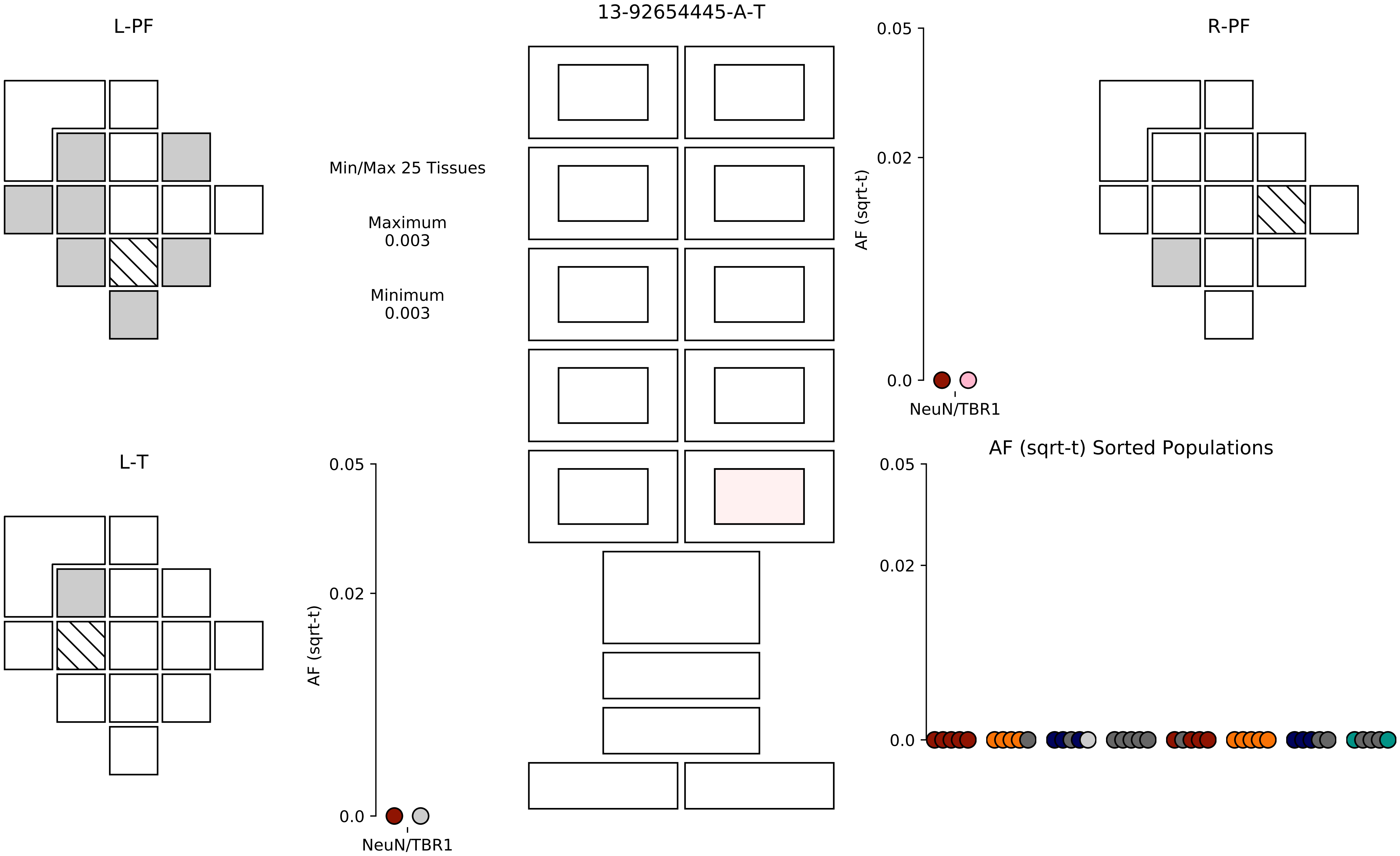

13-92654445-A-T
L-PF
R-PF
L-T
Min/Max 25 Tissues
Maximum
0.003
Minimum
0.003
AF (sqrt-t)
0.05
0.02
0.0
NeuN/TBR1
AF (sqrt-t)
0.05
0.02
0.0
NeuN/TBR1
AF (sqrt-t) Sorted Populations
0.05
0.02
0.0

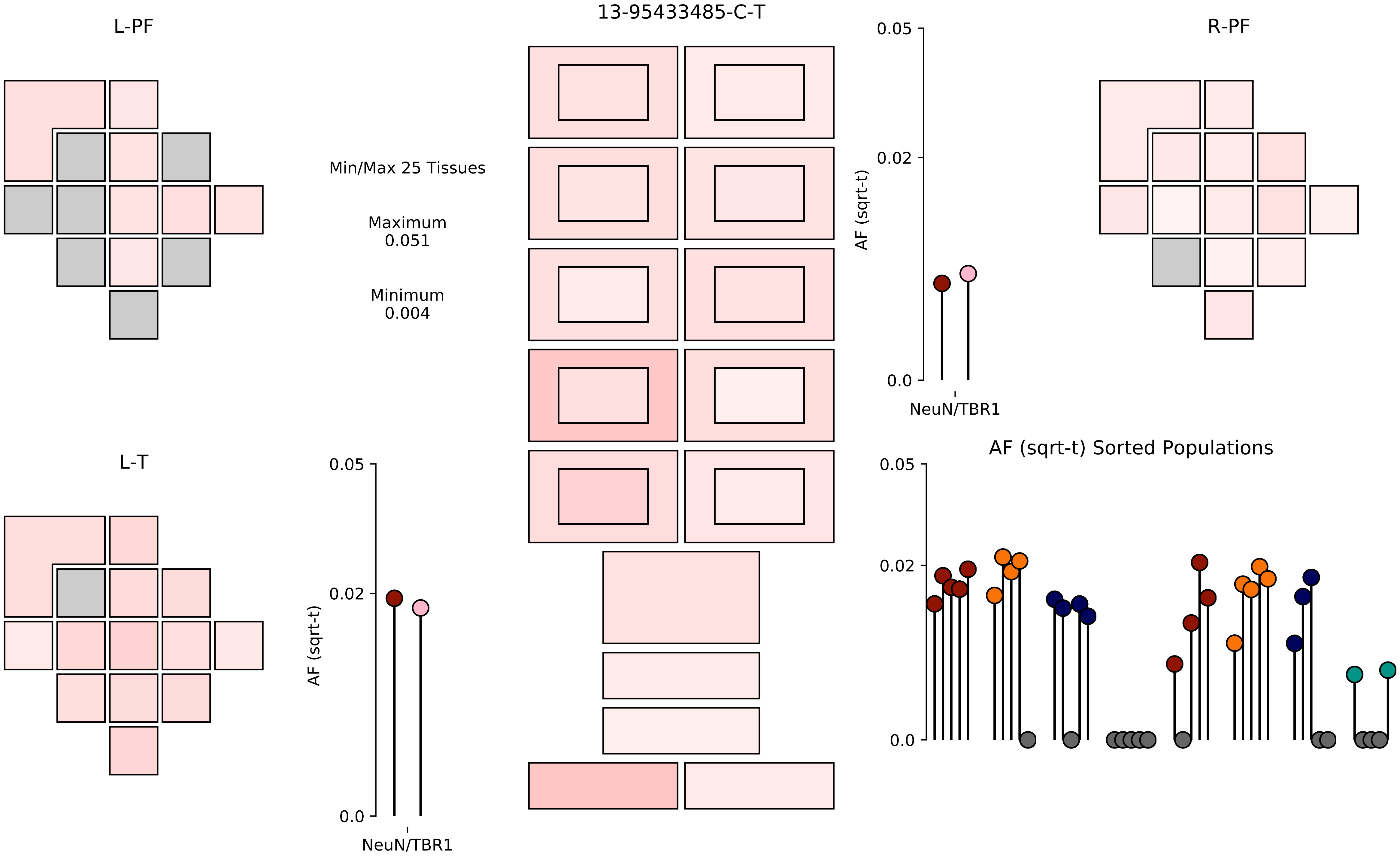

13-95433485-C-T
L-PF
R-PF
L-T
Min/Max 25 Tissues
Maximum
0.051
Minimum
0.004
AF (sqrt-t)
0.05
0.02
0.0
NeuN/TBR1
AF (sqrt-t) Sorted Populations

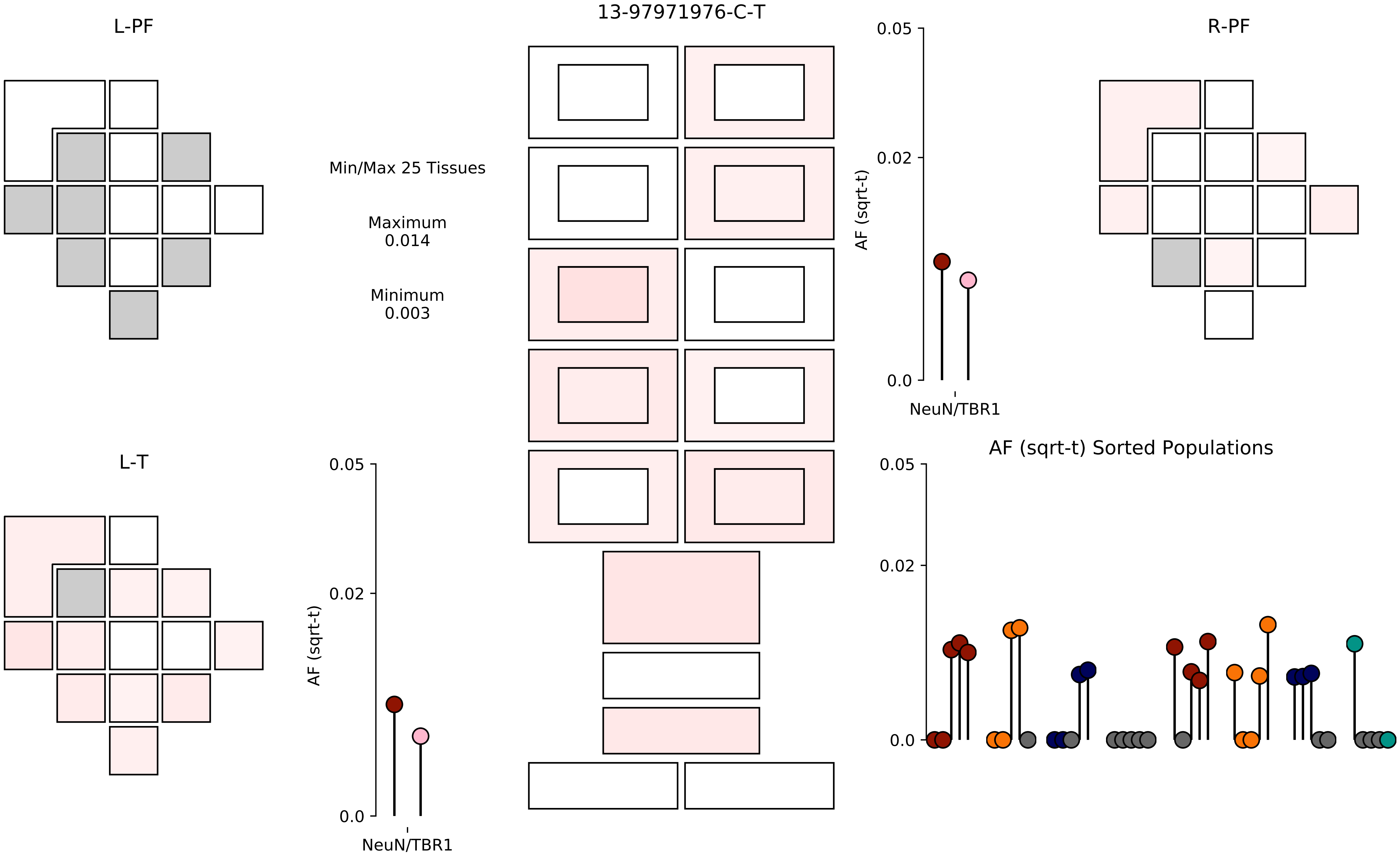

13-97971976-C-T
L-PF
R-PF
L-T
Min/Max 25 Tissues
Maximum
0.014
Minimum
0.003
AF (sqrt-t)
0.05
0.02
0.0
NeuN/TBR1
AF (sqrt-t) Sorted Populations

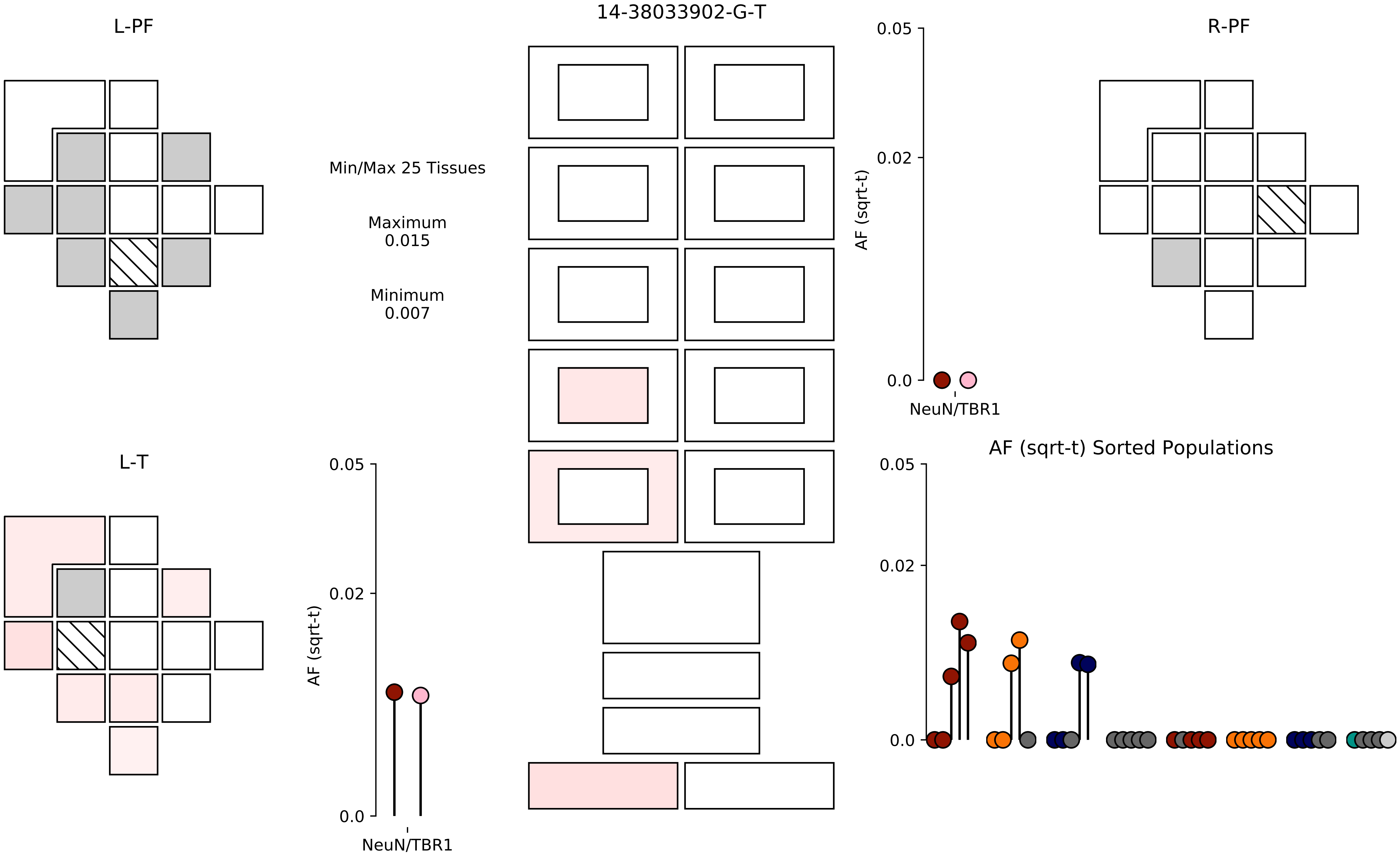

14-38033902-G-T
L-PF
R-PF
L-T
Min/Max 25 Tissues
Maximum
0.015
Minimum
0.007
AF (sqrt-t)
0.05
0.02
0.0
NeuN/TBR1
AF (sqrt-t) Sorted Populations

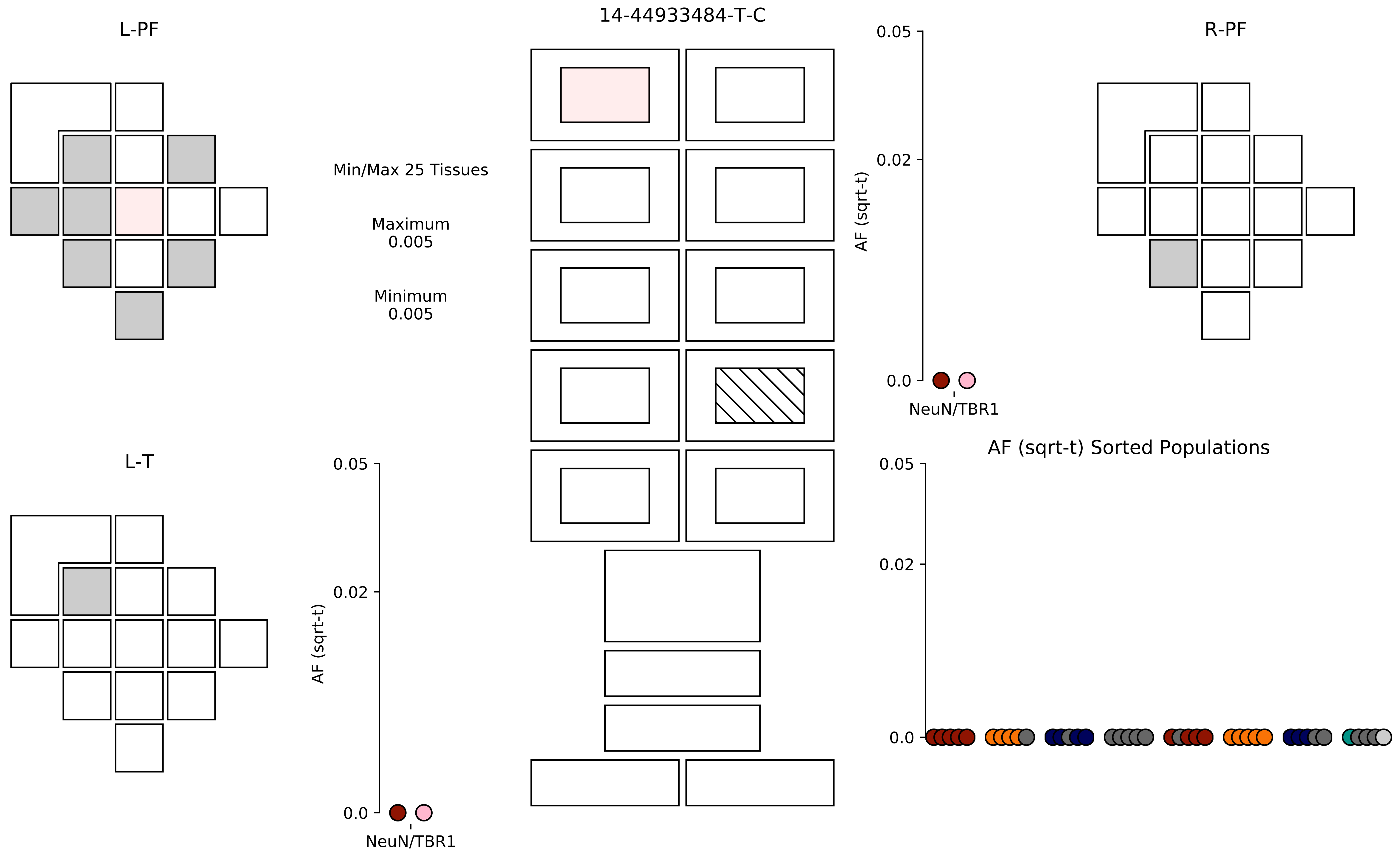

L-PF
14-44933484-T-C
R-PF
Min/Max 25 Tissues
Maximum
0.005
Minimum
0.005
0.05
0.02
0.0
AF (sqrt-t)
NeuN/TBR1
L-T
0.05
0.02
0.0
AF (sqrt-t)
NeuN/TBR1
AF (sqrt-t) Sorted Populations
0.05
0.02
0.0

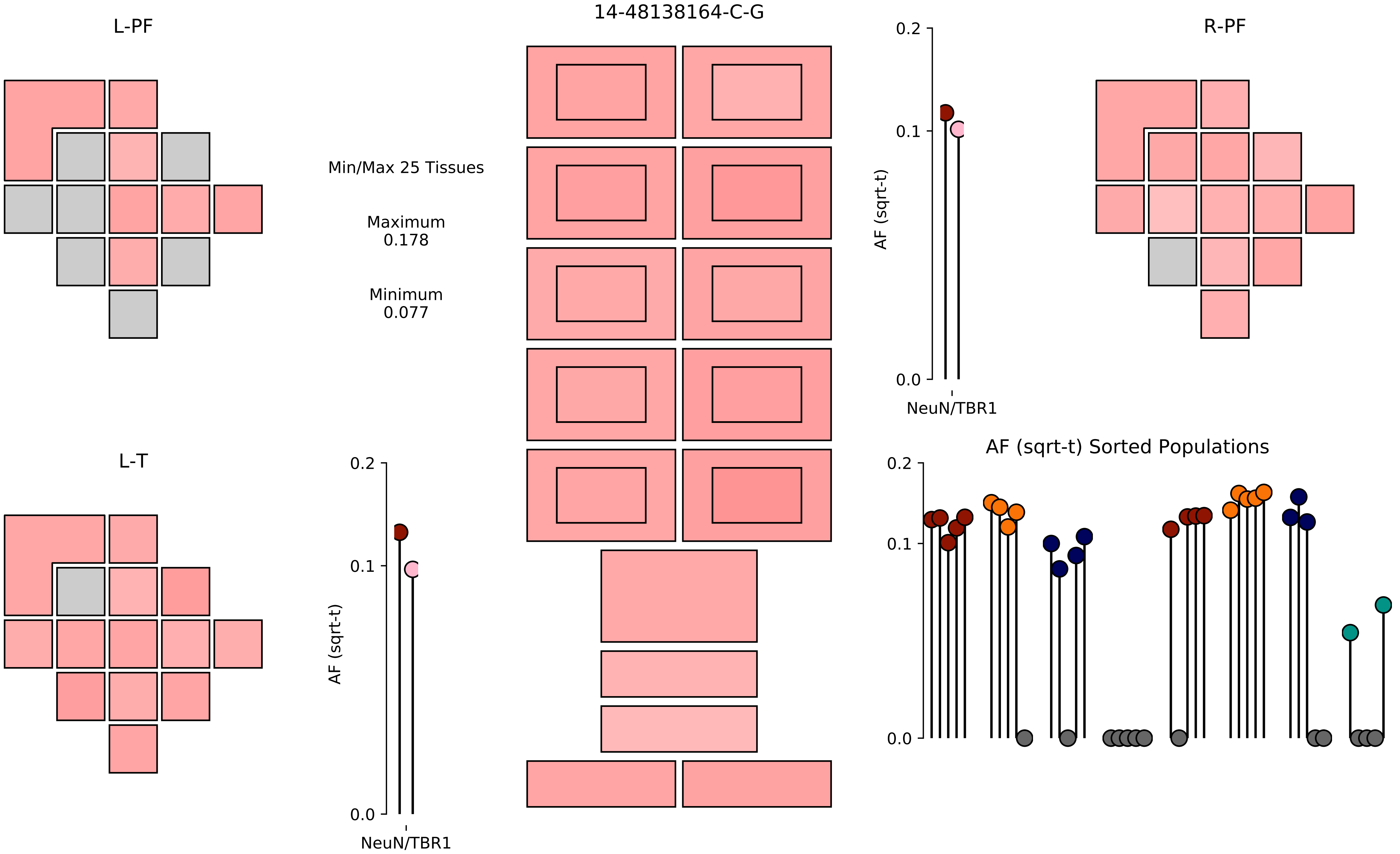

14-48138164-C-G
L-PF
R-PF
L-T
Min/Max 25 Tissues
Maximum
0.178
Minimum
0.077
AF (sqrt-t)
NeuN/TBR1
0.0
0.1
0.2
AF (sqrt-t) Sorted Populations

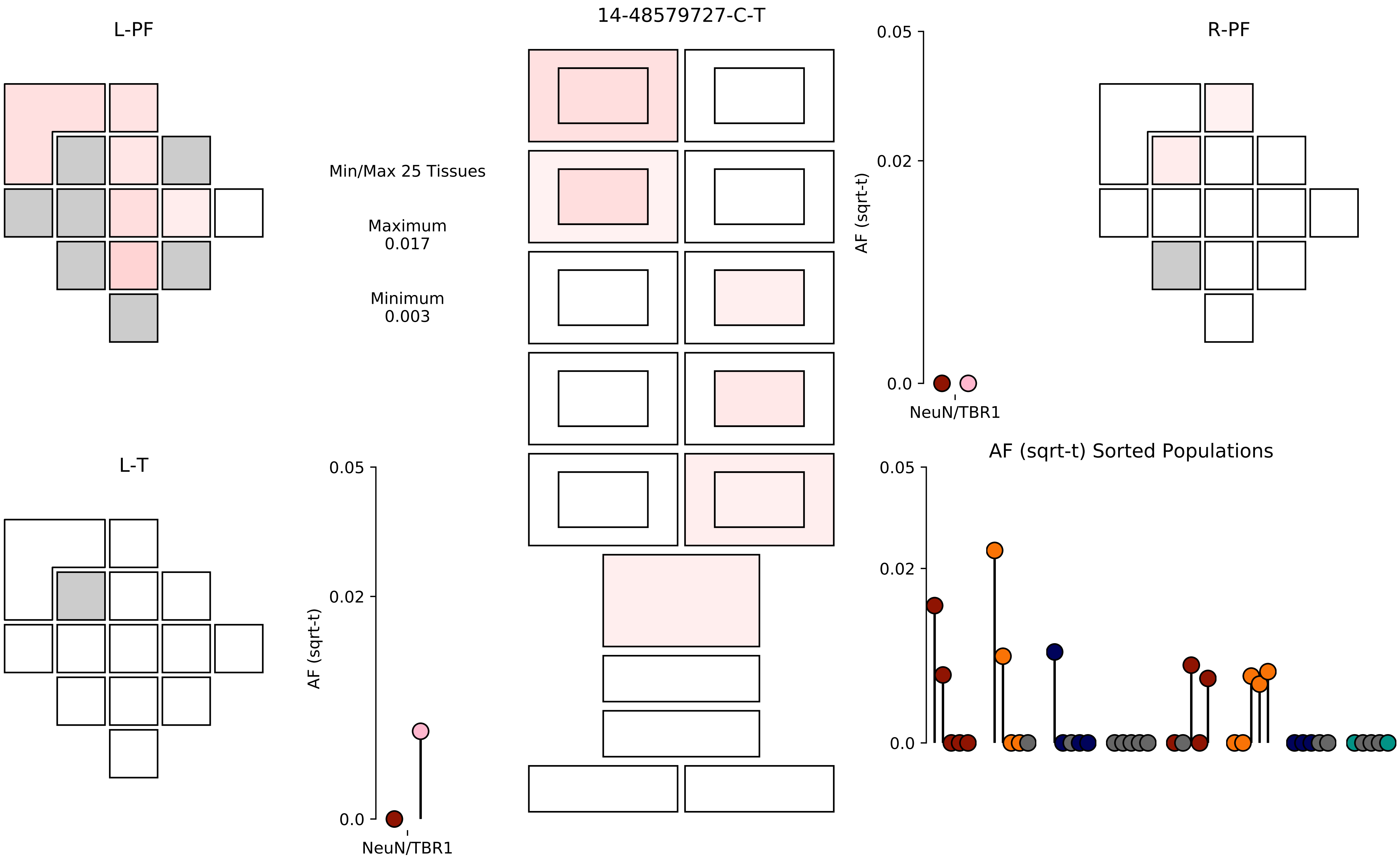

14-48579727-C-T
L-PF
R-PF
L-T
Min/Max 25 Tissues
Maximum
0.017
Minimum
0.003
AF (sqrt-t)
NeuN/TBR1
0.05
0.02
0.0
AF (sqrt-t) Sorted Populations

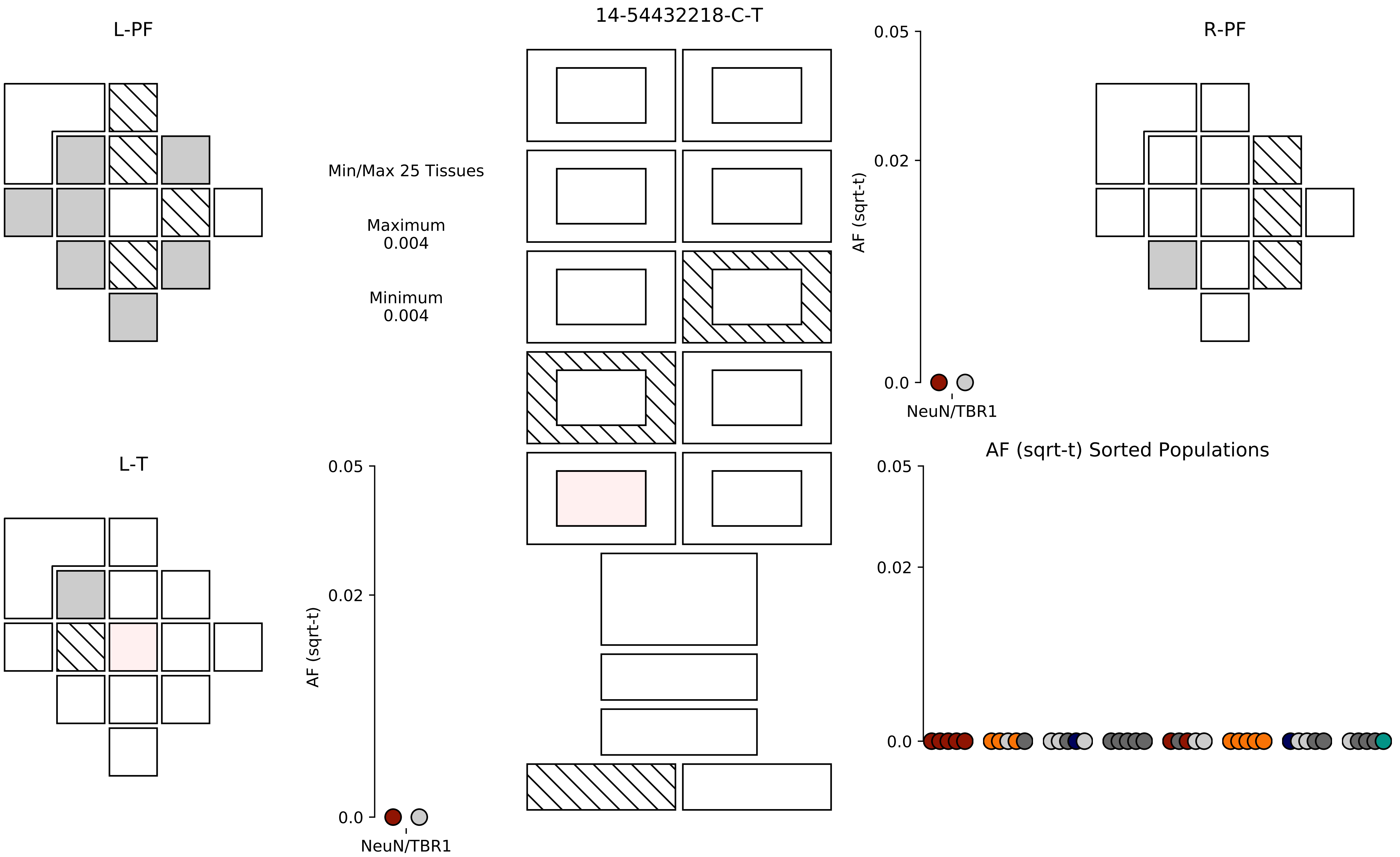

14-54432218-C-T
L-PF
R-PF
L-T
Min/Max 25 Tissues
Maximum
0.004
Minimum
0.004
0.05
0.02
0.0
AF (sqrt-t)
NeuN/TBR1
AF (sqrt-t) Sorted Populations

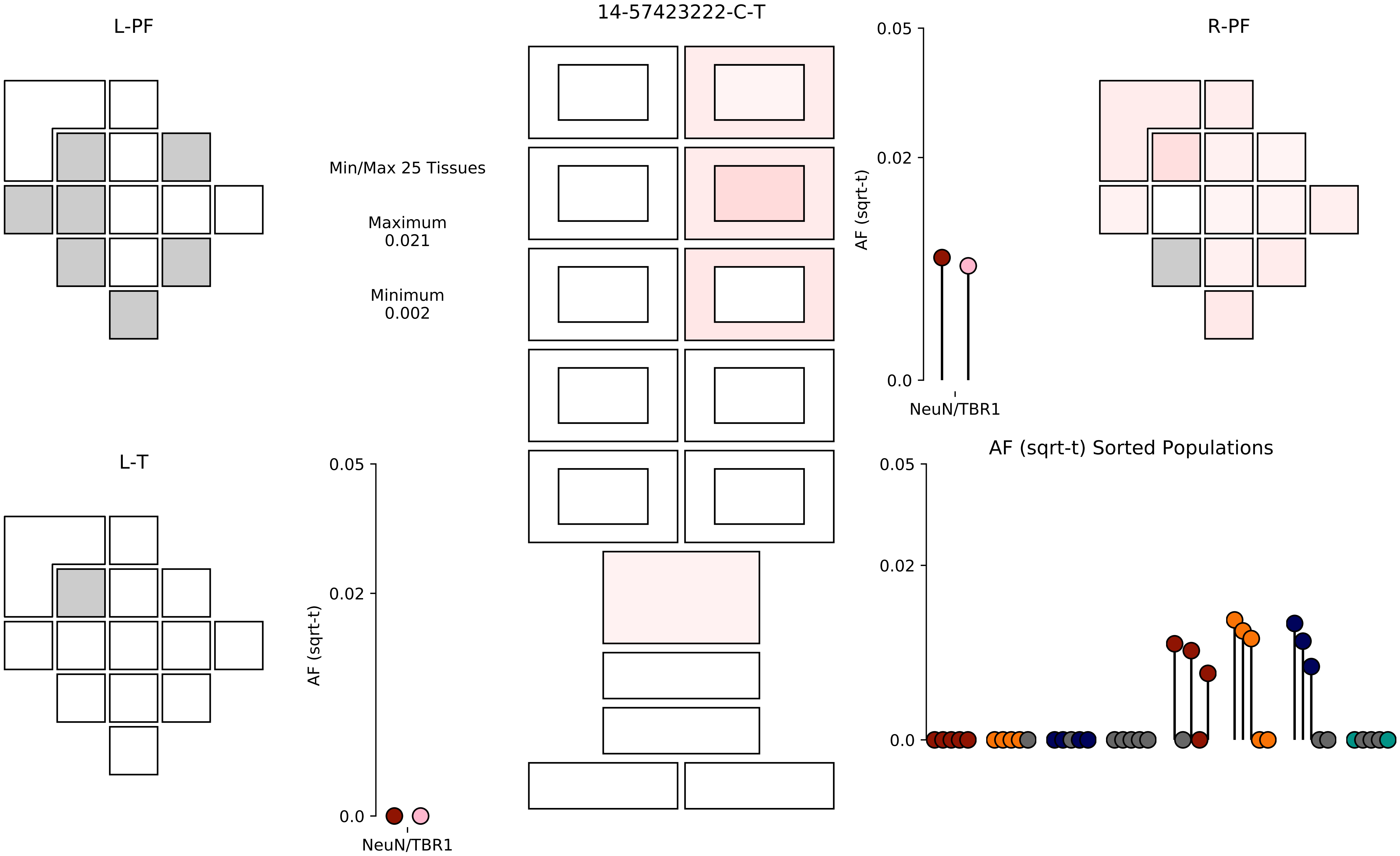

14-57423222-C-T
L-PF
R-PF
L-T
Min/Max 25 Tissues
Maximum
0.021
Minimum
0.002
AF (sqrt-t)
0.05
0.02
0.0
NeuN/TBR1
AF (sqrt-t) Sorted Populations

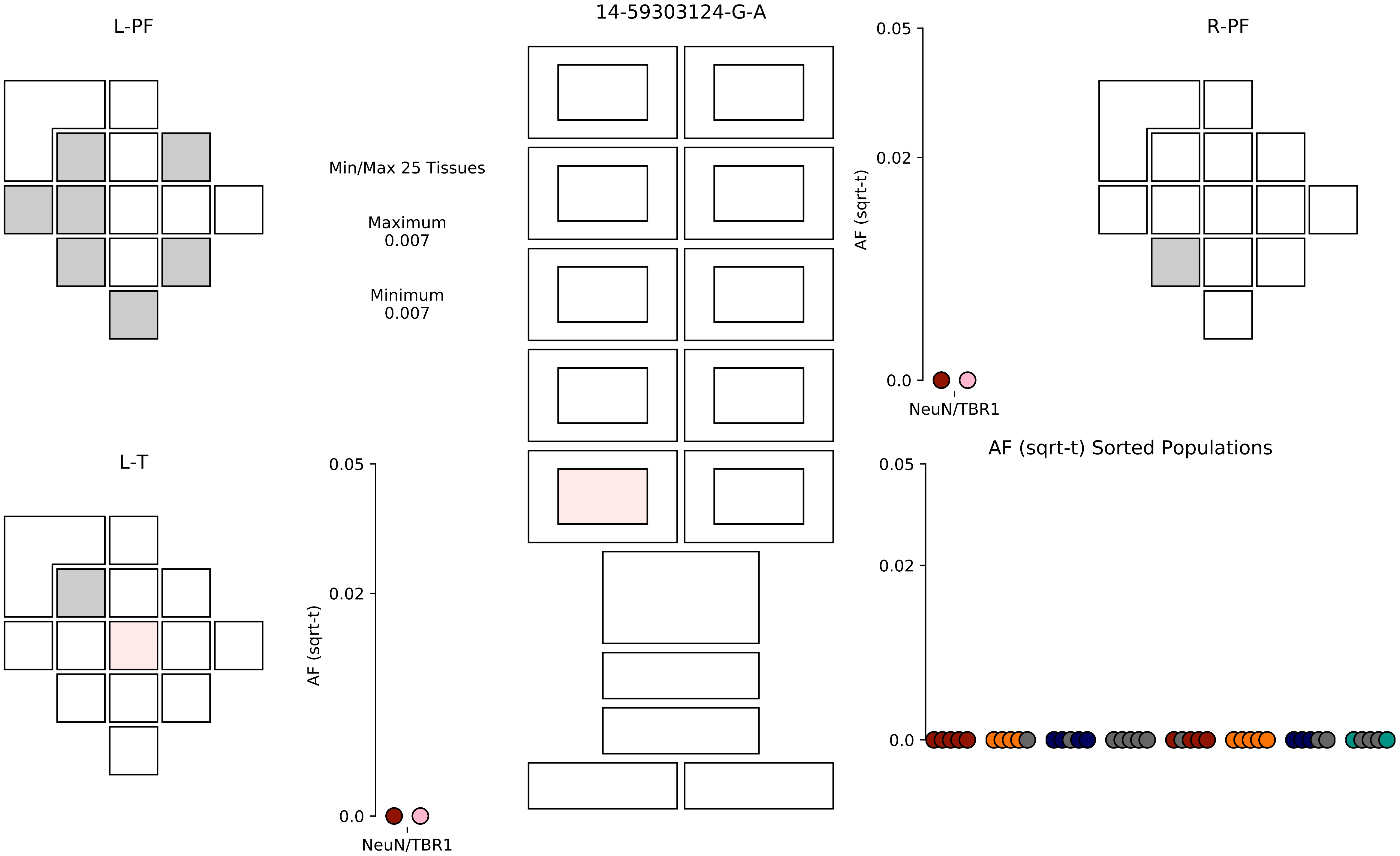

14-59303124-G-A
L-PF
R-PF
L-T
Min/Max 25 Tissues
Maximum
0.007
Minimum
0.007
AF (sqrt-t)
0.05
0.02
0.0
NeuN/TBR1
AF (sqrt-t) Sorted Populations

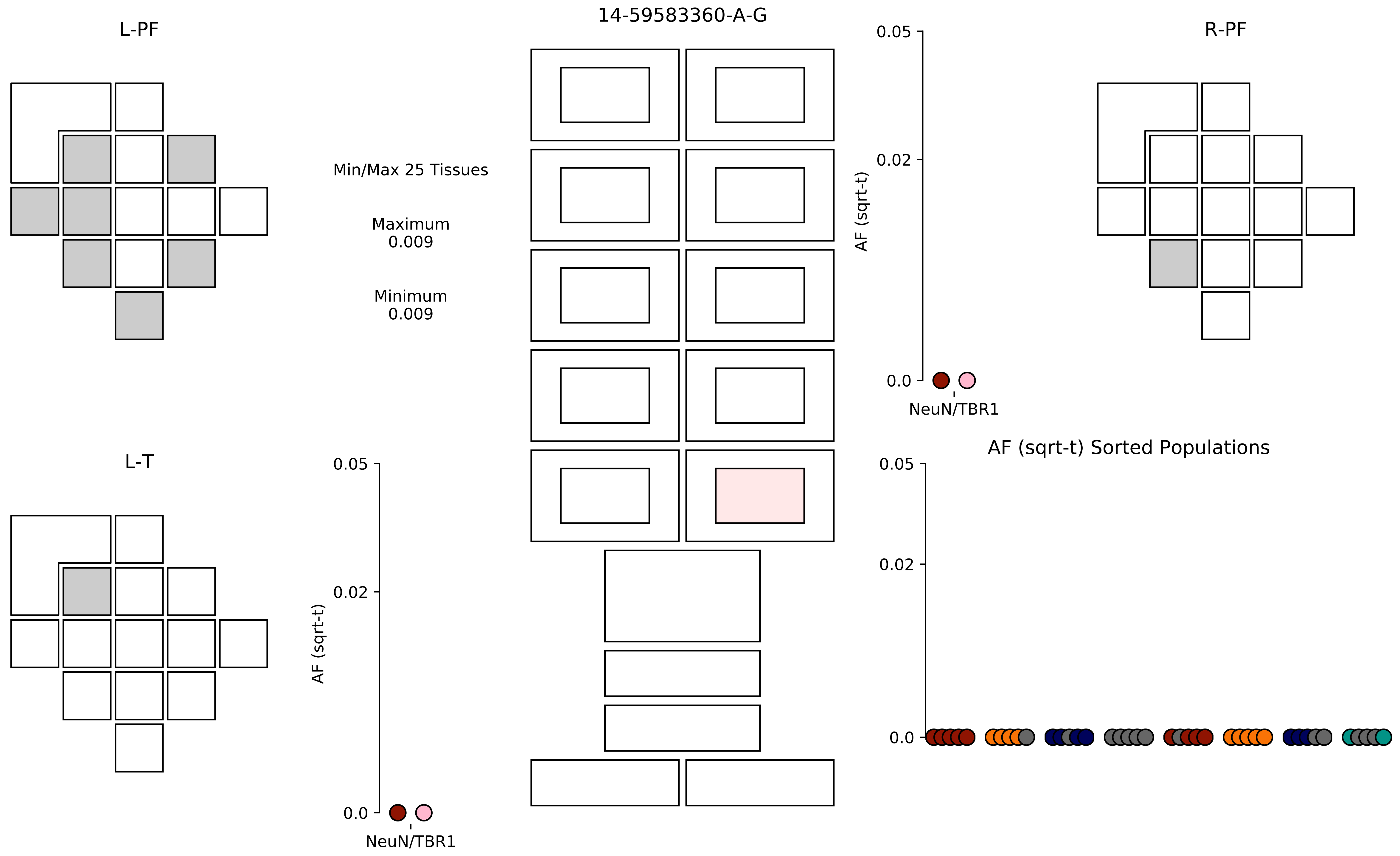
14-59583360-A-G
L-PF
R-PF
L-T
Min/Max 25 Tissues
Maximum
0.009
Minimum
0.009
AF (sqrt-t)
0.05
0.02
0.0
NeuN/TBR1
AF (sqrt-t)
0.05
0.02
0.0
NeuN/TBR1
AF (sqrt-t) Sorted Populations
0.05
0.02
0.0

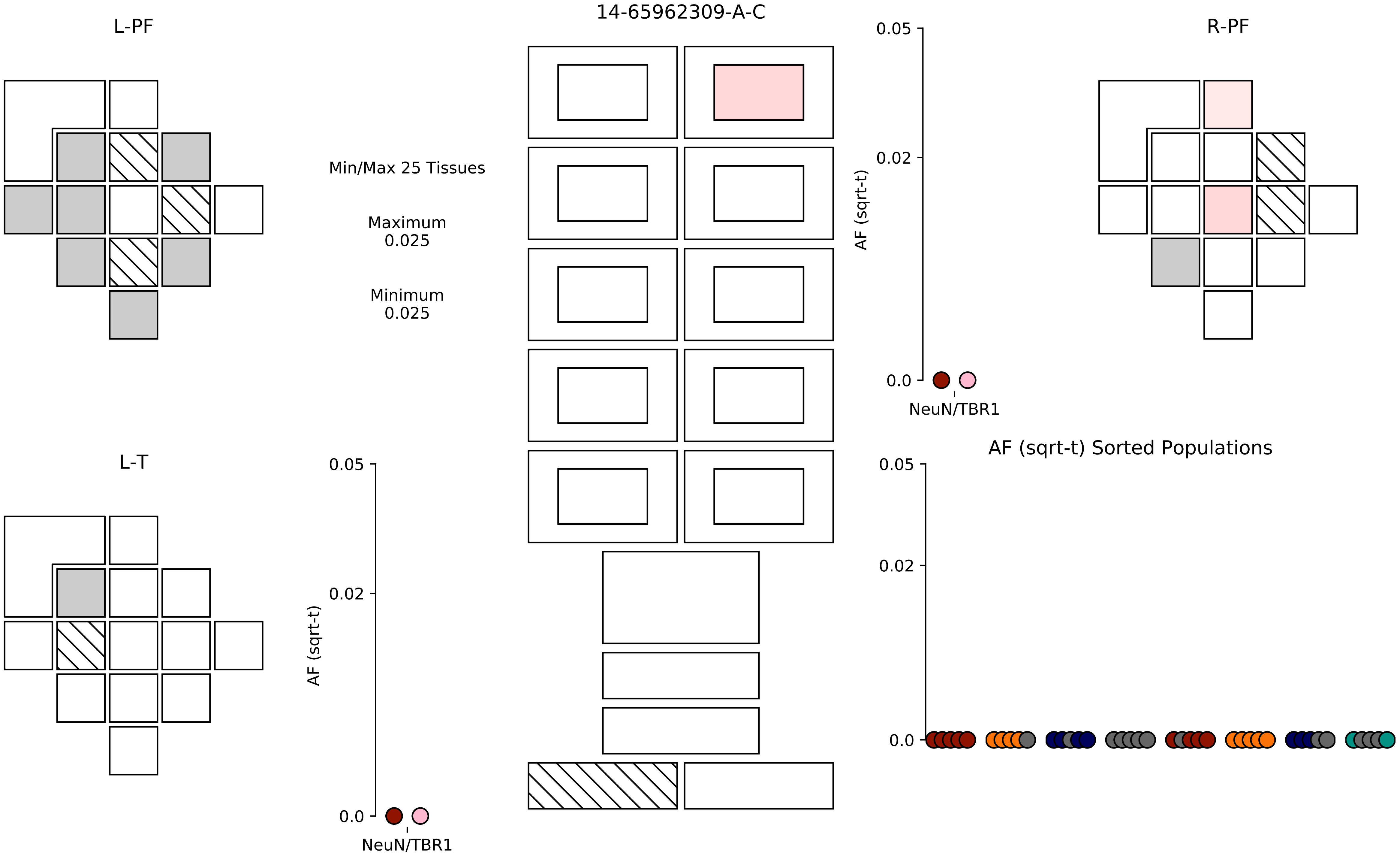
14-65962309-A-C
L-PF
R-PF
L-T
Min/Max 25 Tissues
Maximum
0.025
Minimum
0.025
AF (sqrt-t)
0.05
0.02
0.0
NeuN/TBR1
AF (sqrt-t) Sorted Populations

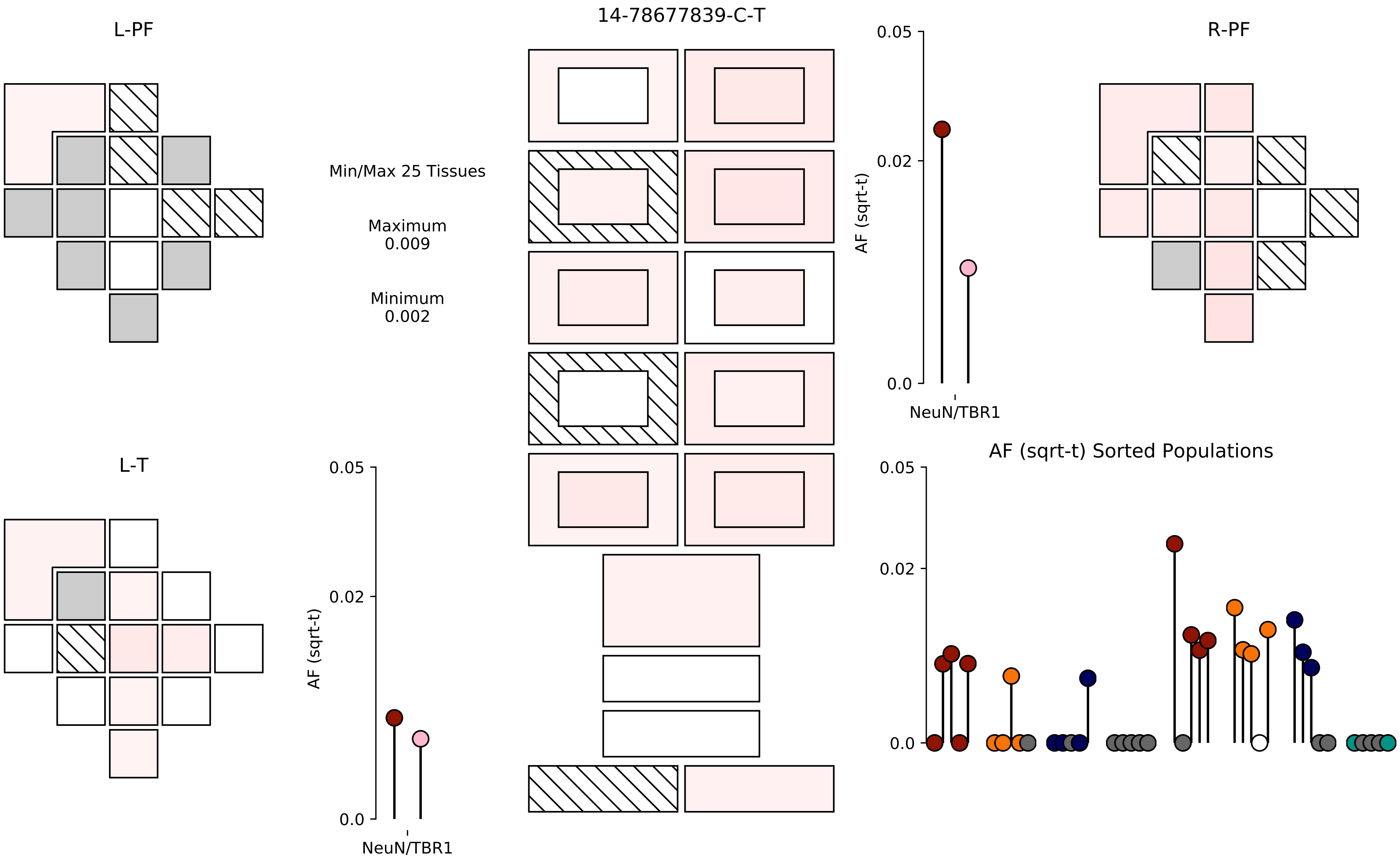

14-78677839-C-T
L-PF
R-PF
L-T
Min/Max 25 Tissues
Maximum
0.009
Minimum
0.002
AF (sqrt-t)
0.05
0.02
0.0
NeuN/TBR1
AF (sqrt-t) Sorted Populations

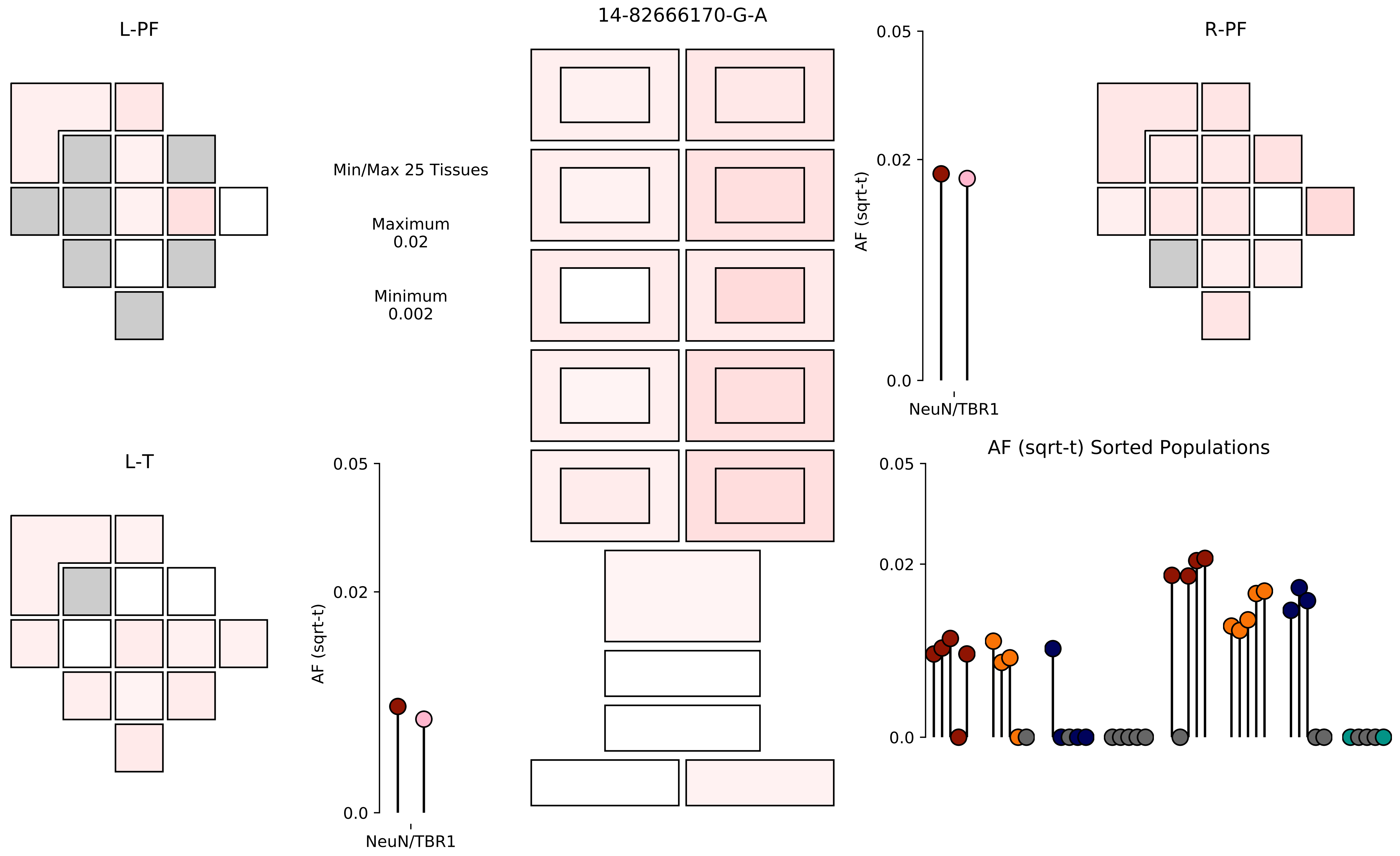

14-82666170-G-A
L-PF
R-PF
L-T
Min/Max 25 Tissues
Maximum
0.02
Minimum
0.002
0.05
0.02
0.0
AF (sqrt-t)
NeuN/TBR1
AF (sqrt-t) Sorted Populations

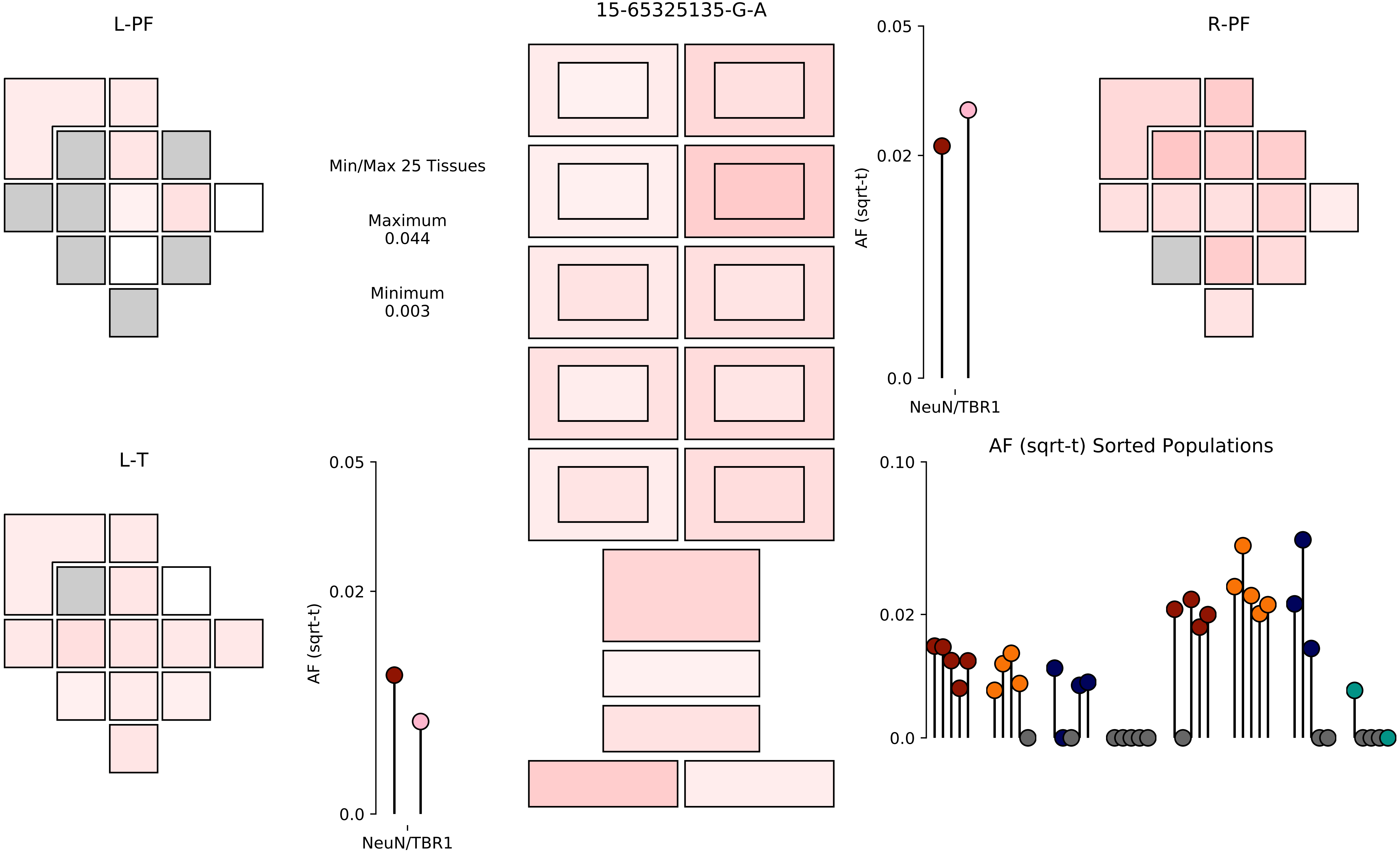

15-65325135-G-A
L-PF
R-PF
L-T
Min/Max 25 Tissues
Maximum
0.044
Minimum
0.003
AF (sqrt-t)
0.05
0.02
0.0
NeuN/TBR1
AF (sqrt-t) Sorted Populations
0.10
0.02
0.0

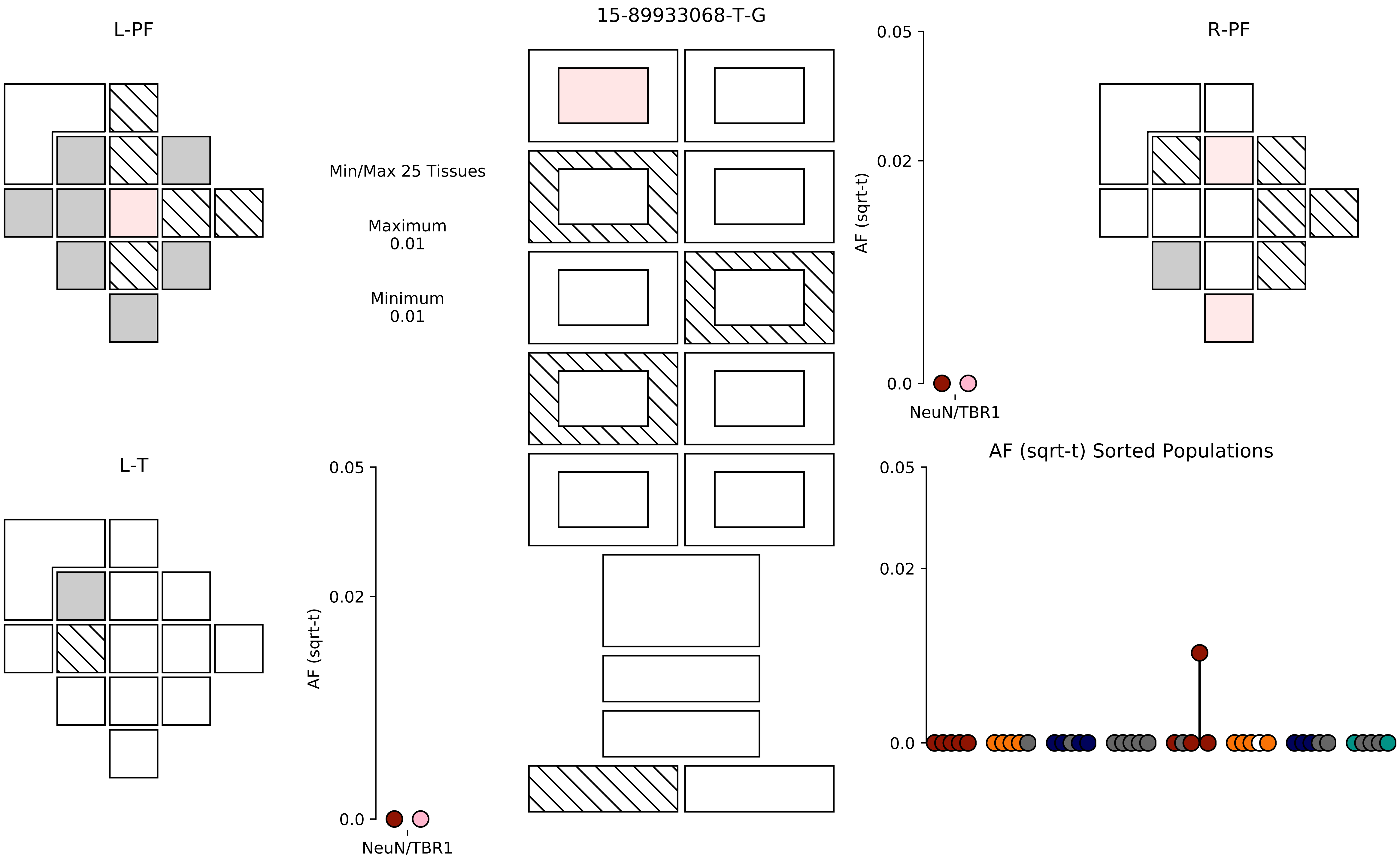

15-89933068-T-G
L-PF
R-PF
L-T
Min/Max 25 Tissues
Maximum
0.01
Minimum
0.01
AF (sqrt-t)
0.05
0.02
0.0
NeuN/TBR1
AF (sqrt-t) Sorted Populations

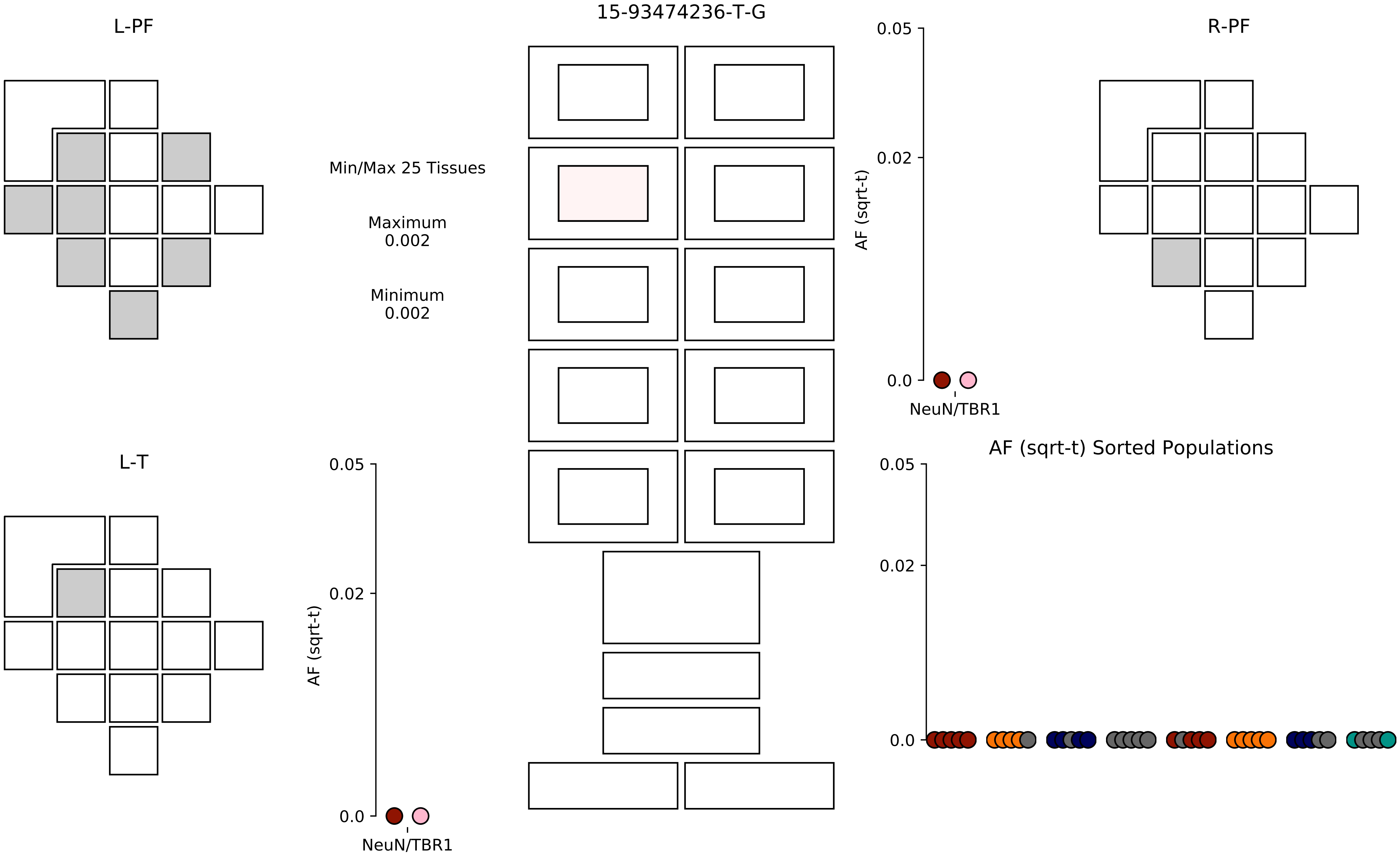
15-93474236-T-G
L-PF
R-PF
L-T
Min/Max 25 Tissues
Maximum
0.002
Minimum
0.002
AF (sqrt-t)
0.05
0.02
0.0
NeuN/TBR1
AF (sqrt-t) Sorted Populations

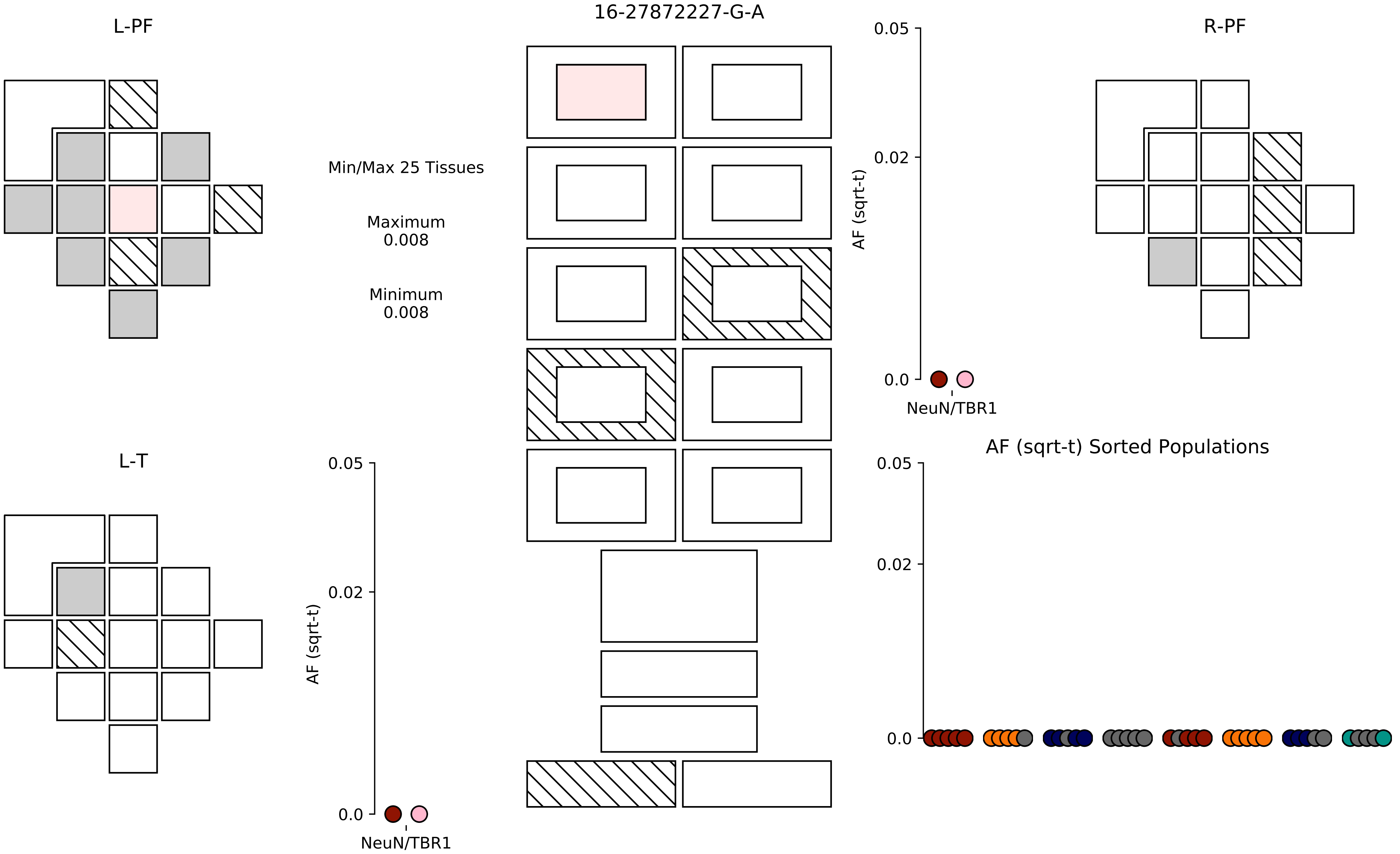

L-PF
16-27872227-G-A
R-PF
Min/Max 25 Tissues
Maximum
0.008
Minimum
0.008
0.05
0.02
0.0
AF (sqrt-t)
NeuN/TBR1
L-T
0.05
0.02
0.0
AF (sqrt-t)
NeuN/TBR1
AF (sqrt-t) Sorted Populations
0.05
0.02
0.0

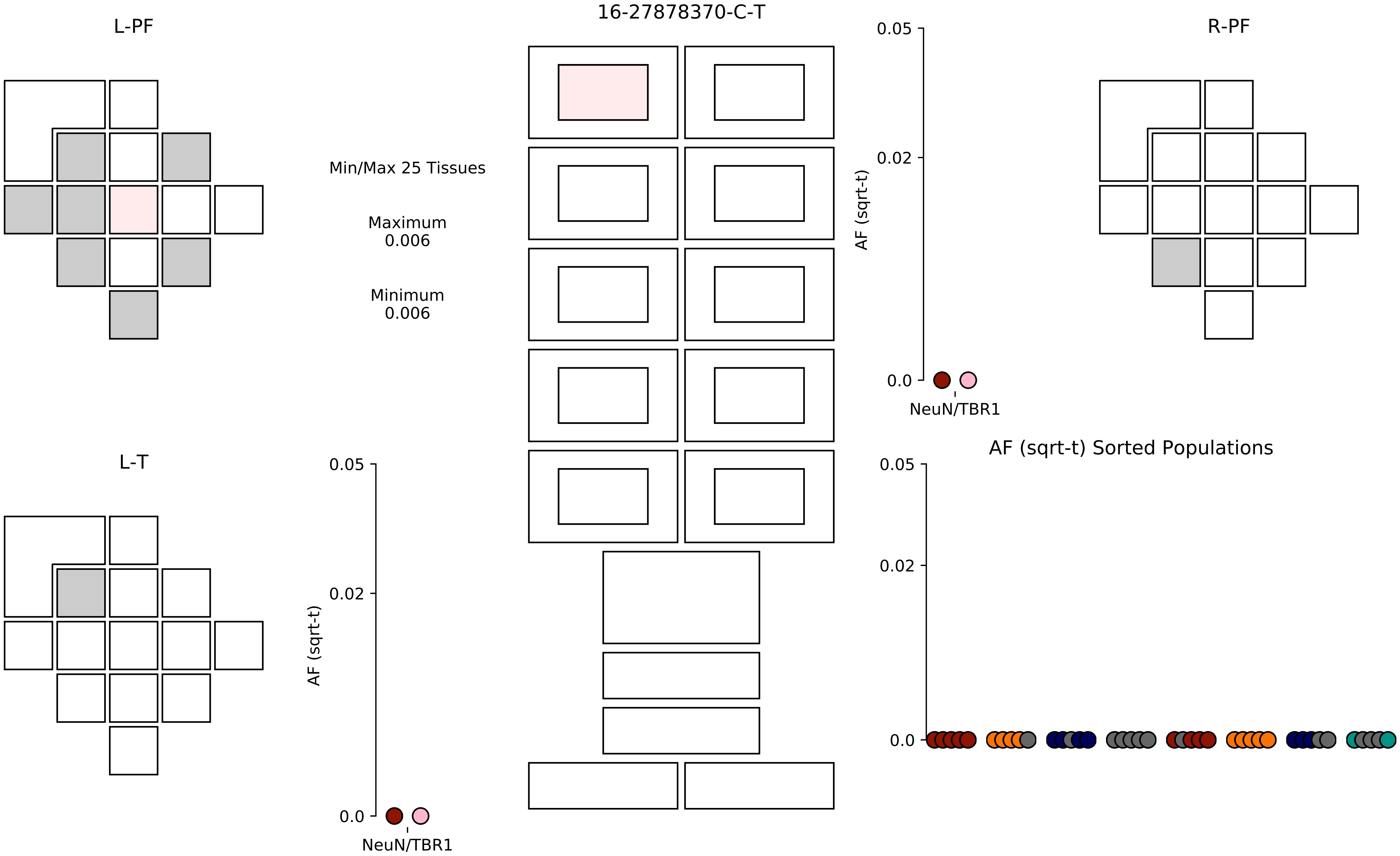
16-27878370-C-T
L-PF
R-PF
L-T
Min/Max 25 Tissues
Maximum
0.006
Minimum
0.006
AF (sqrt-t)
0.05
0.02
0.0
NeuN/TBR1
AF (sqrt-t)
0.05
0.02
0.0
NeuN/TBR1
AF (sqrt-t) Sorted Populations
0.05
0.02
0.0

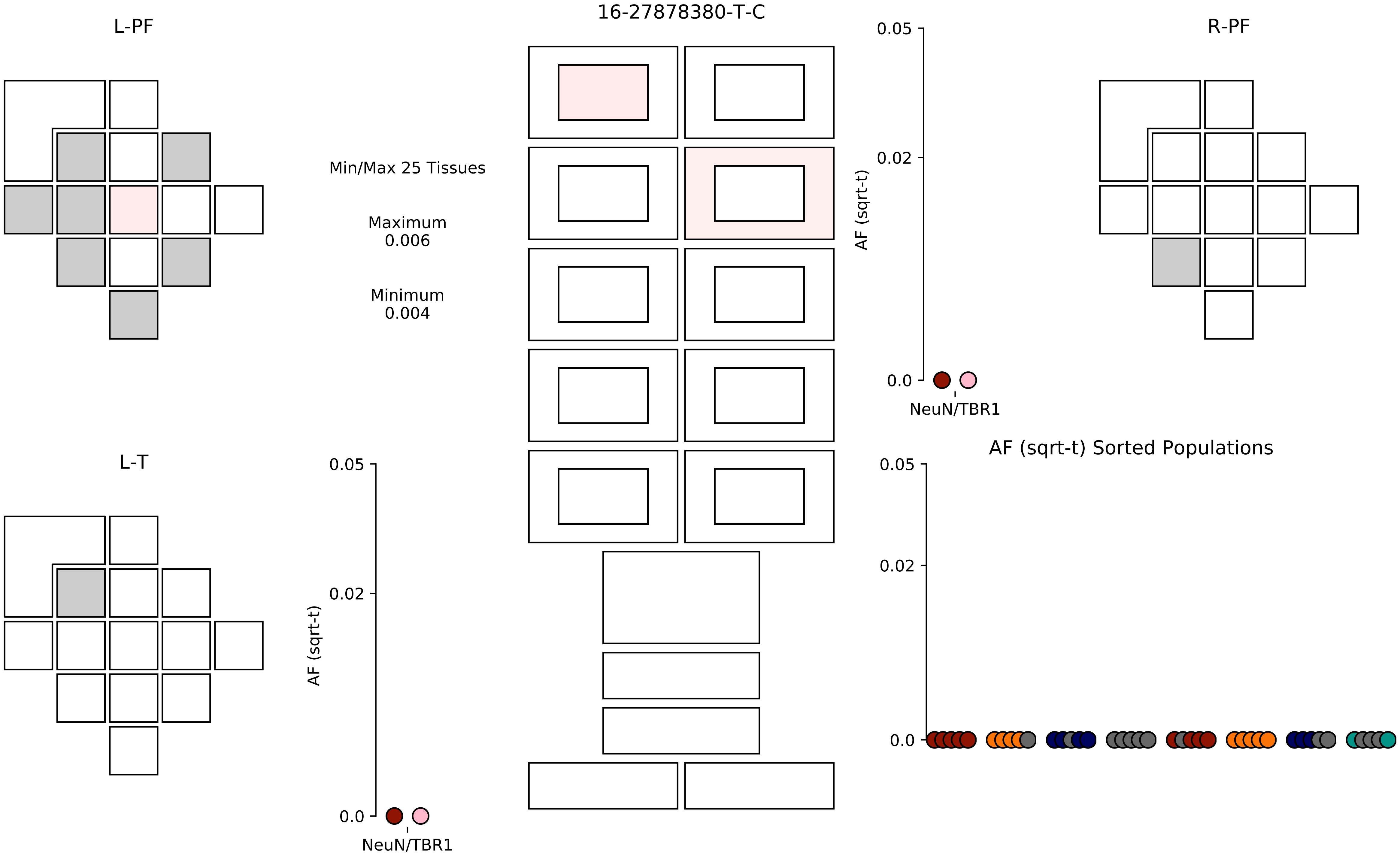

16-27878380-T-C
L-PF
R-PF
L-T
Min/Max 25 Tissues
Maximum
0.006
Minimum
0.004
0.05
0.02
0.0
AF (sqrt-t)
NeuN/TBR1
AF (sqrt-t) Sorted Populations

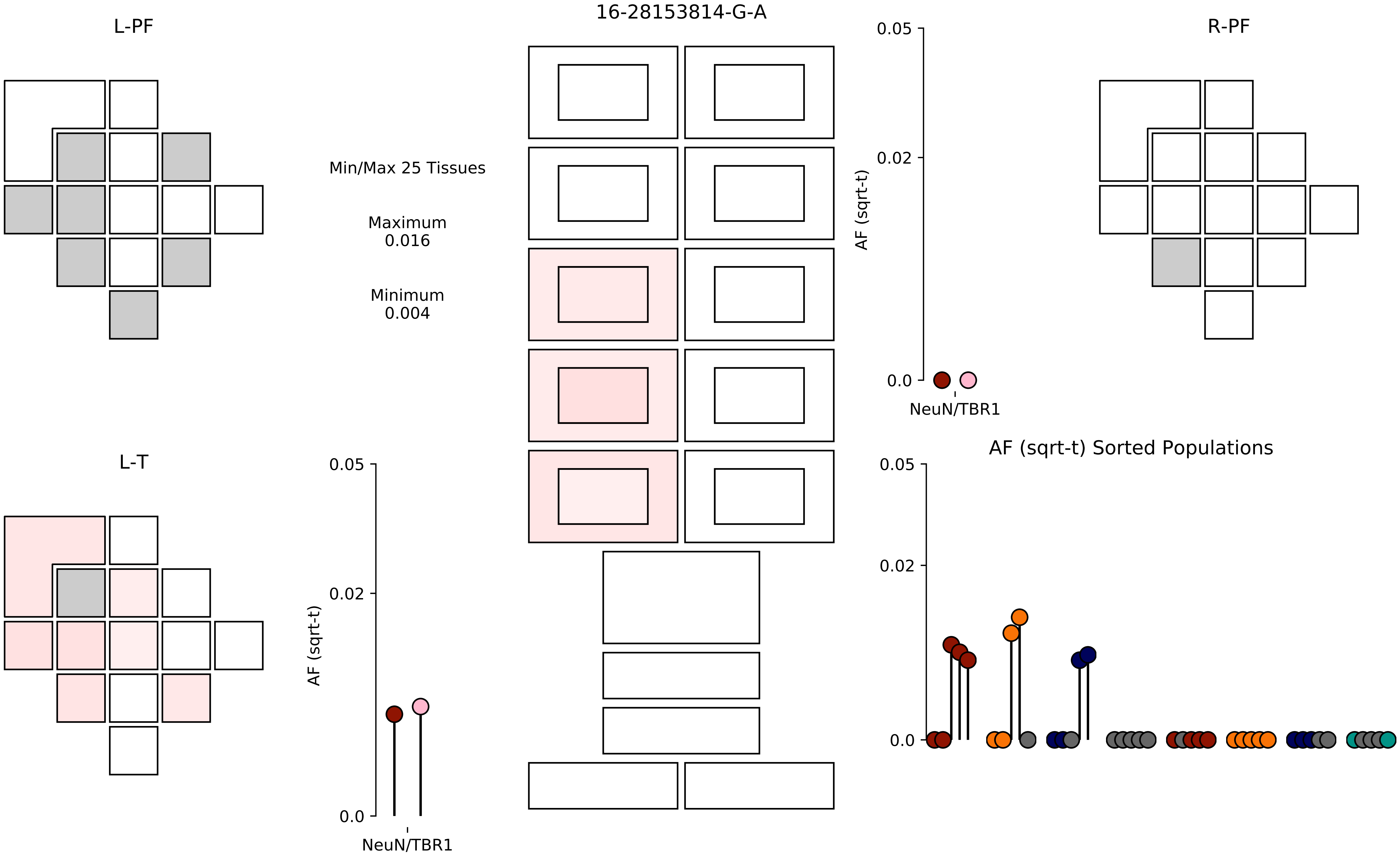
16-28153814-G-A
L-PF
R-PF
L-T
Min/Max 25 Tissues
Maximum
0.016
Minimum
0.004
0.05
0.02
0.0
AF (sqrt-t)
NeuN/TBR1
AF (sqrt-t) Sorted Populations

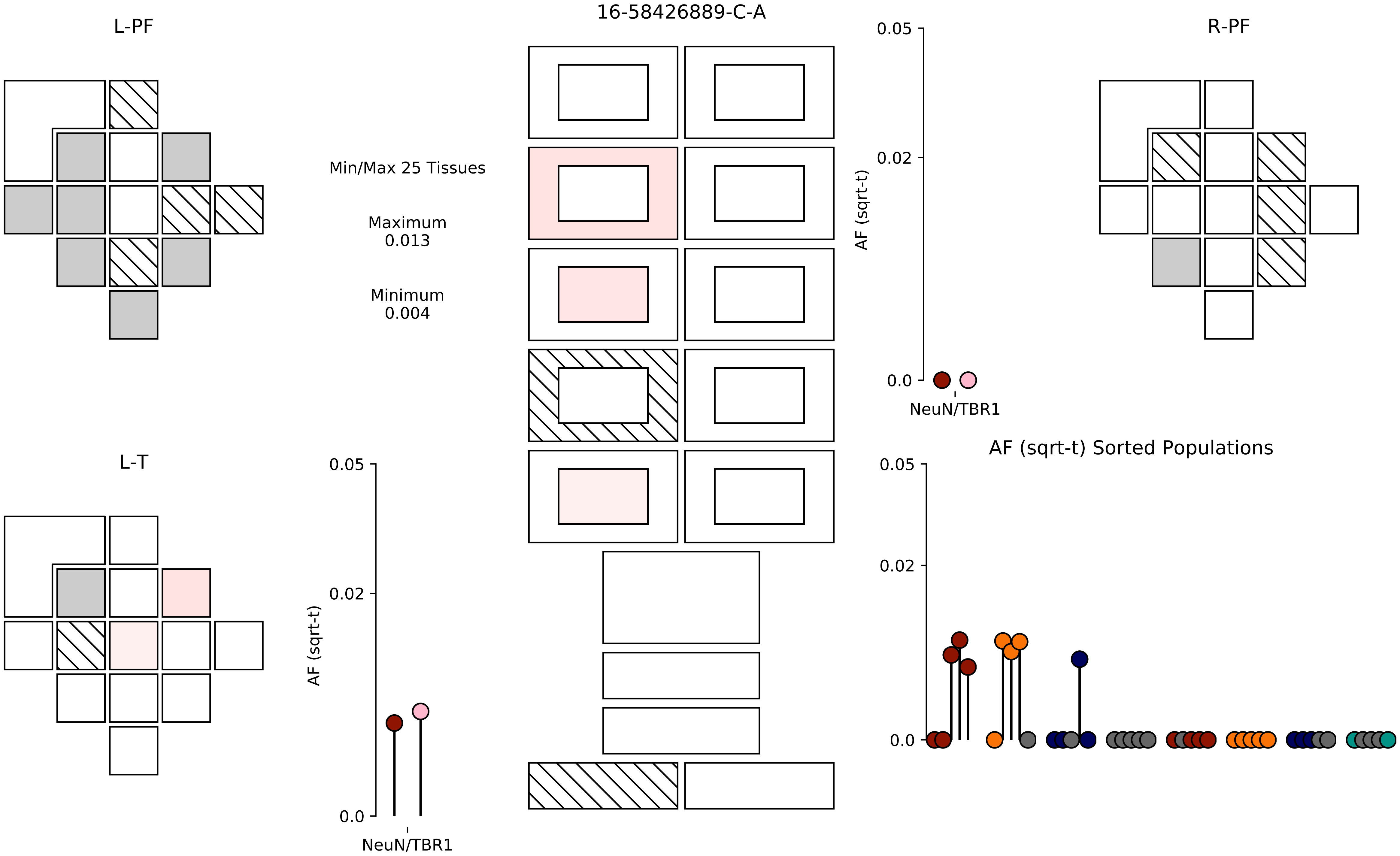

16-58426889-C-A
L-PF
R-PF
L-T
Min/Max 25 Tissues
Maximum
0.013
Minimum
0.004
AF (sqrt-t)
0.05
0.02
0.0
NeuN/TBR1
AF (sqrt-t) Sorted Populations

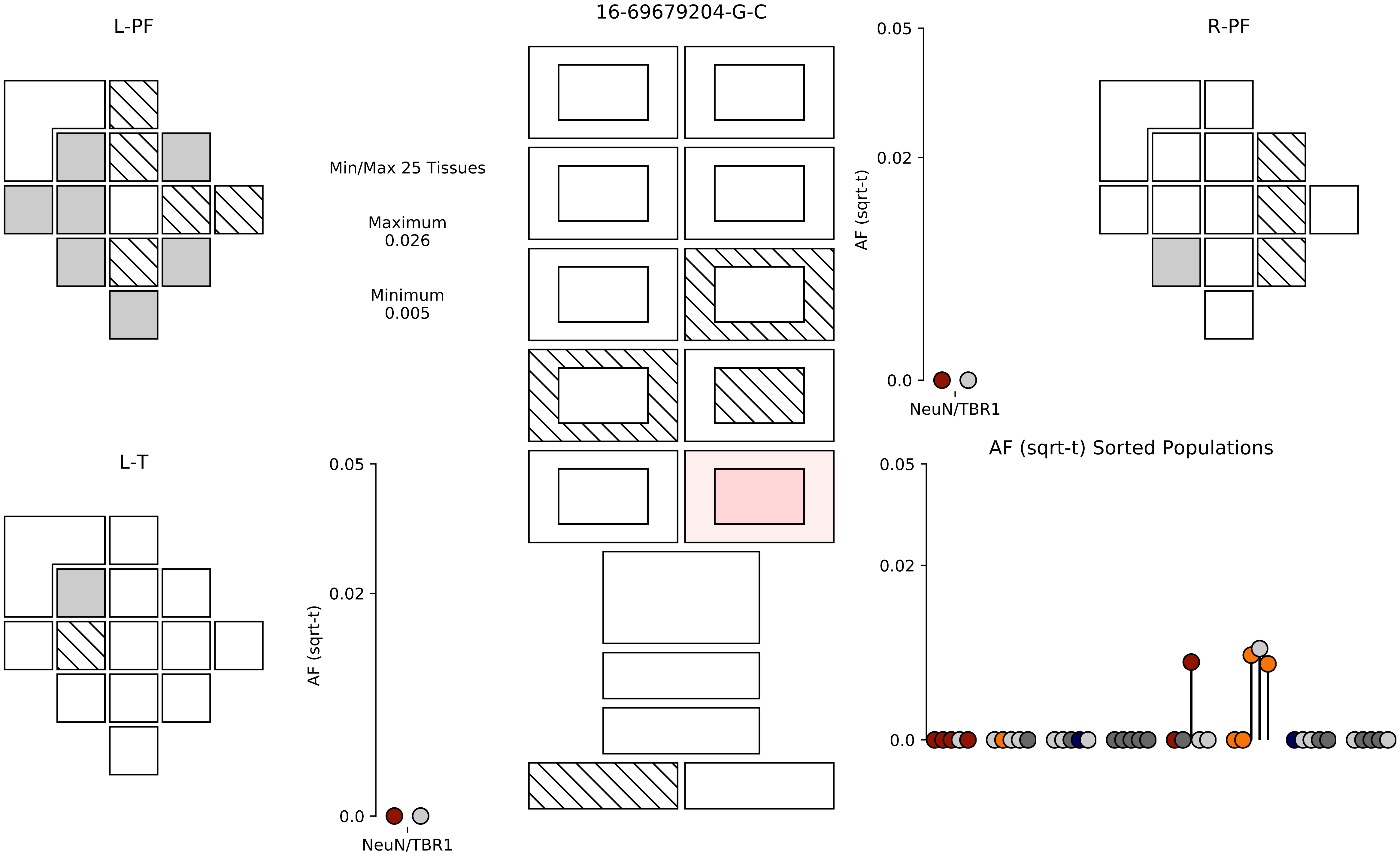
16-69679204-G-C
L-PF
R-PF
L-T
Min/Max 25 Tissues
Maximum
0.026
Minimum
0.005
0.05
0.02
0.0
AF (sqrt-t)
NeuN/TBR1
AF (sqrt-t) Sorted Populations

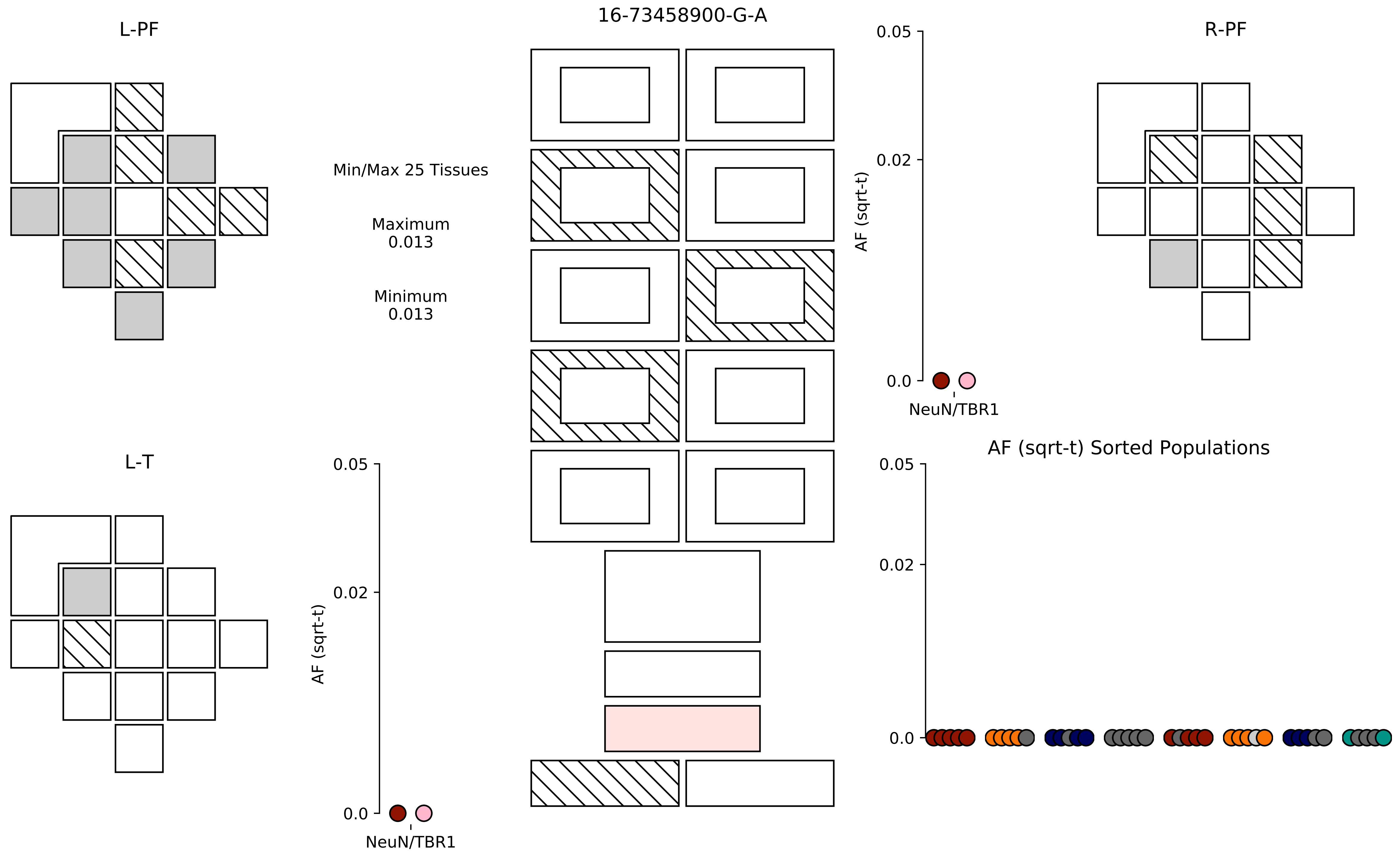
16-73458900-G-A
L-PF
R-PF
L-T
Min/Max 25 Tissues
Maximum
0.013
Minimum
0.013
AF (sqrt-t)
0.05
0.02
0.0
NeuN/TBR1
AF (sqrt-t) Sorted Populations

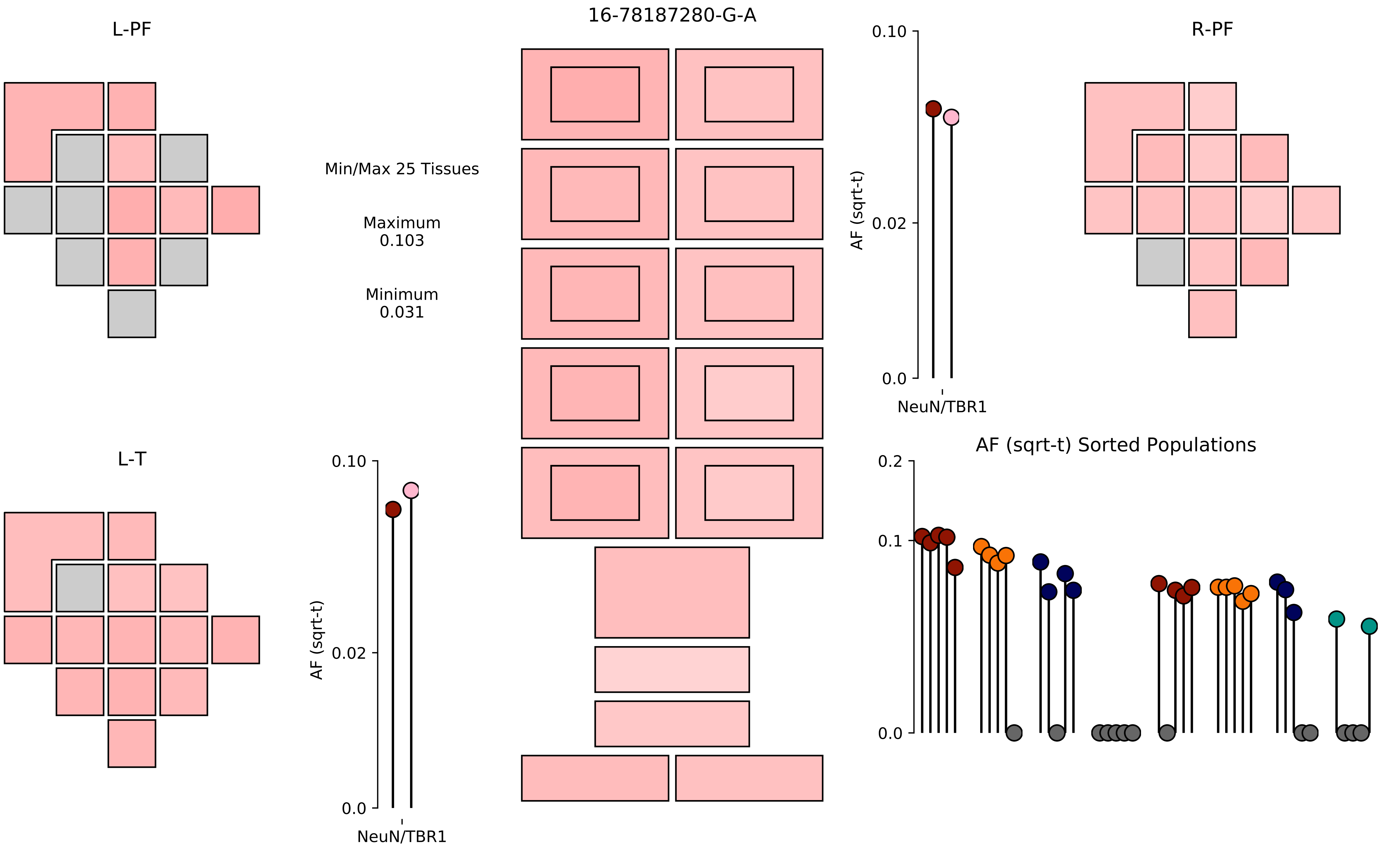
16-78187280-G-A
L-PF
R-PF
L-T
Min/Max 25 Tissues
Maximum
0.103
Minimum
0.031
AF (sqrt-t)
0.10
0.02
0.0
NeuN/TBR1
AF (sqrt-t)
0.10
0.02
0.0
NeuN/TBR1
AF (sqrt-t) Sorted Populations
0.2
0.1
0.0

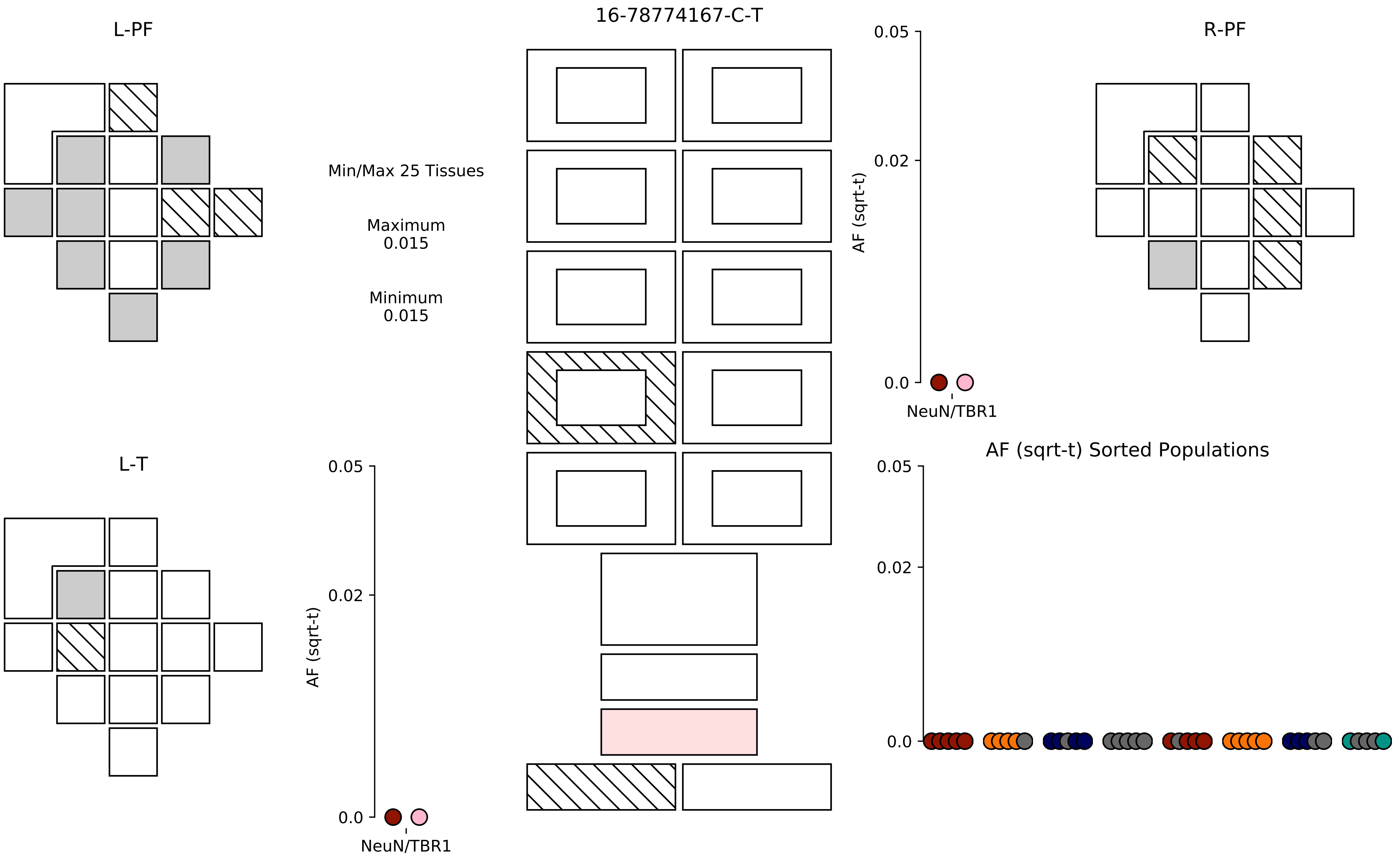
16-78774167-C-T
L-PF
R-PF
L-T
Min/Max 25 Tissues
Maximum
0.015
Minimum
0.015
AF (sqrt-t)
0.05
0.02
0.0
NeuN/TBR1
AF (sqrt-t)
0.05
0.02
0.0
NeuN/TBR1
AF (sqrt-t) Sorted Populations
0.05
0.02
0.0

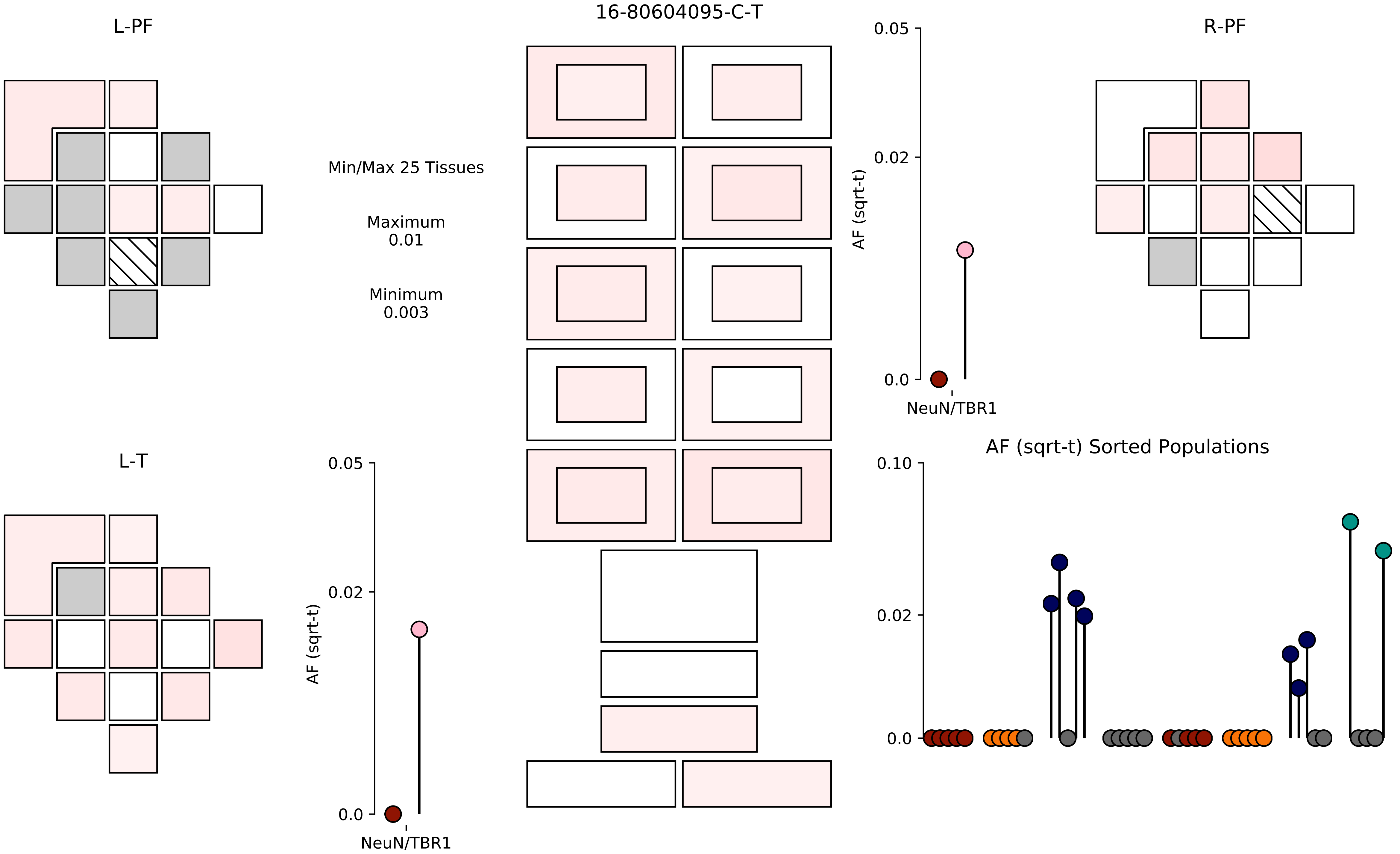
16-80604095-C-T
L-PF
R-PF
L-T
Min/Max 25 Tissues
Maximum
0.01
Minimum
0.003
AF (sqrt-t)
0.05
0.02
0.0
NeuN/TBR1
AF (sqrt-t) Sorted Populations
0.10
0.02
0.0

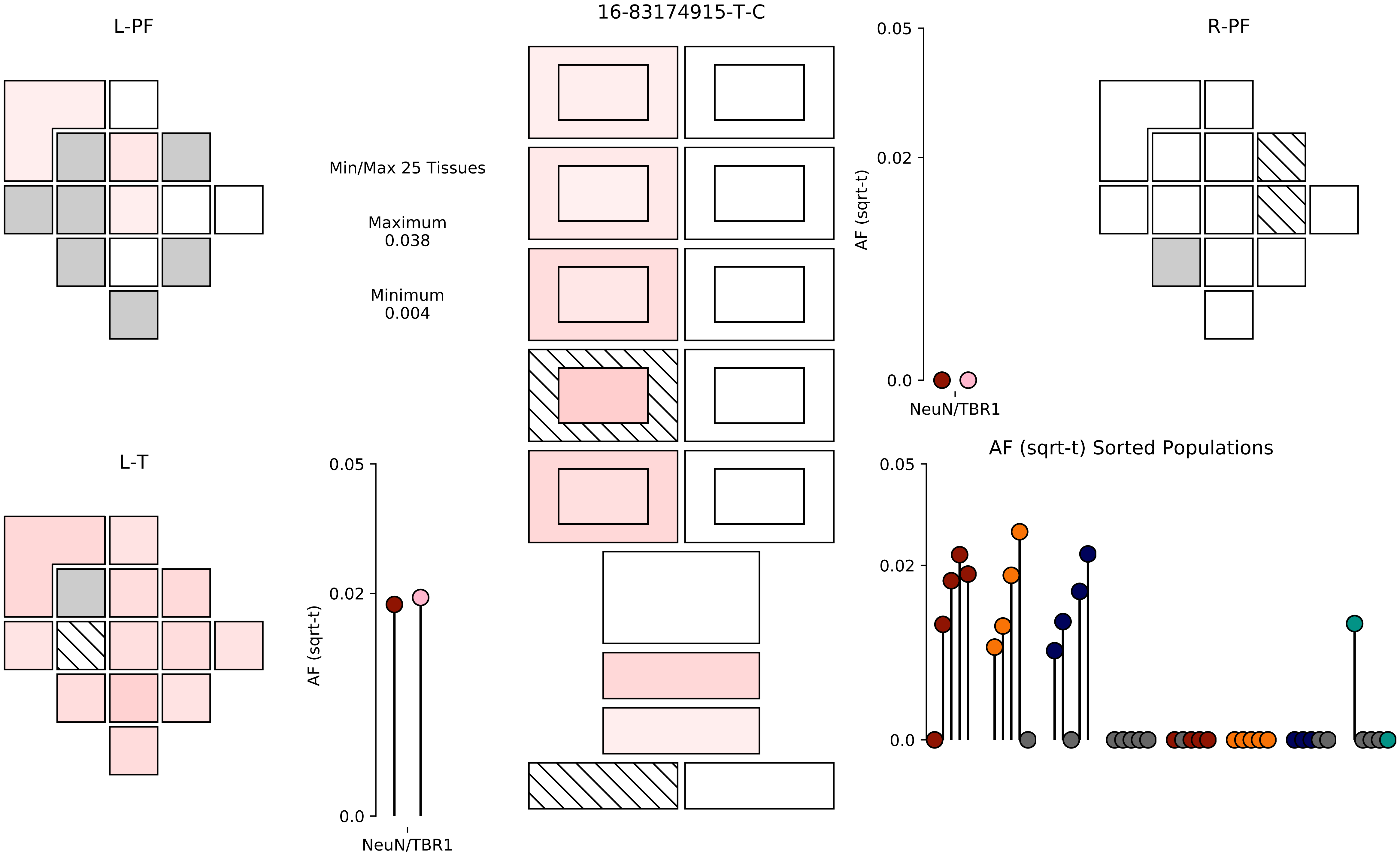
16-83174915-T-C
L-PF
R-PF
L-T
Min/Max 25 Tissues
Maximum
0.038
Minimum
0.004
AF (sqrt-t)
0.05
0.02
0.0
NeuN/TBR1
AF (sqrt-t) Sorted Populations

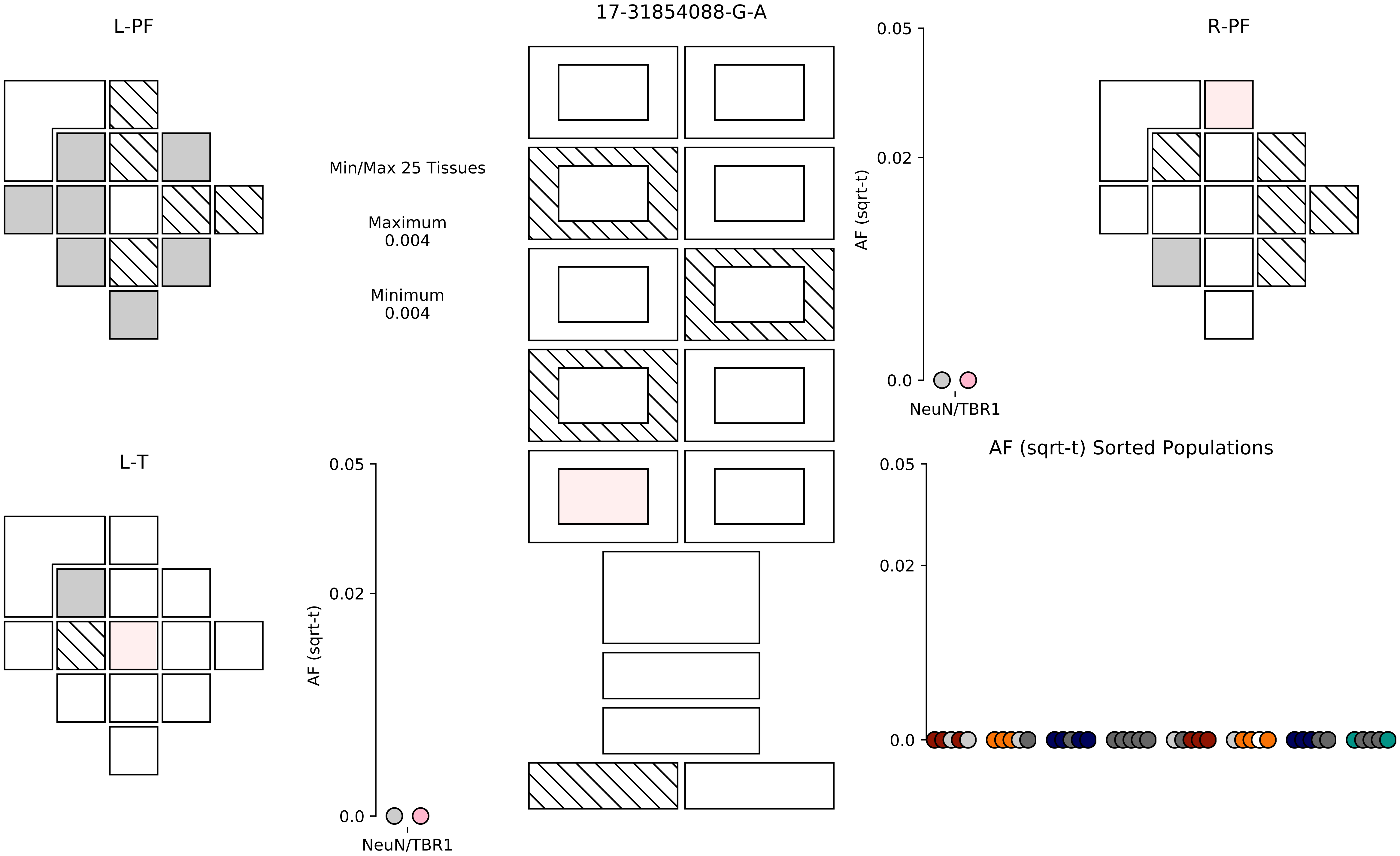

17-31854088-G-A
L-PF
R-PF
L-T
Min/Max 25 Tissues
Maximum
0.004
Minimum
0.004
AF (sqrt-t)
0.05
0.02
0.0
NeuN/TBR1
AF (sqrt-t) Sorted Populations

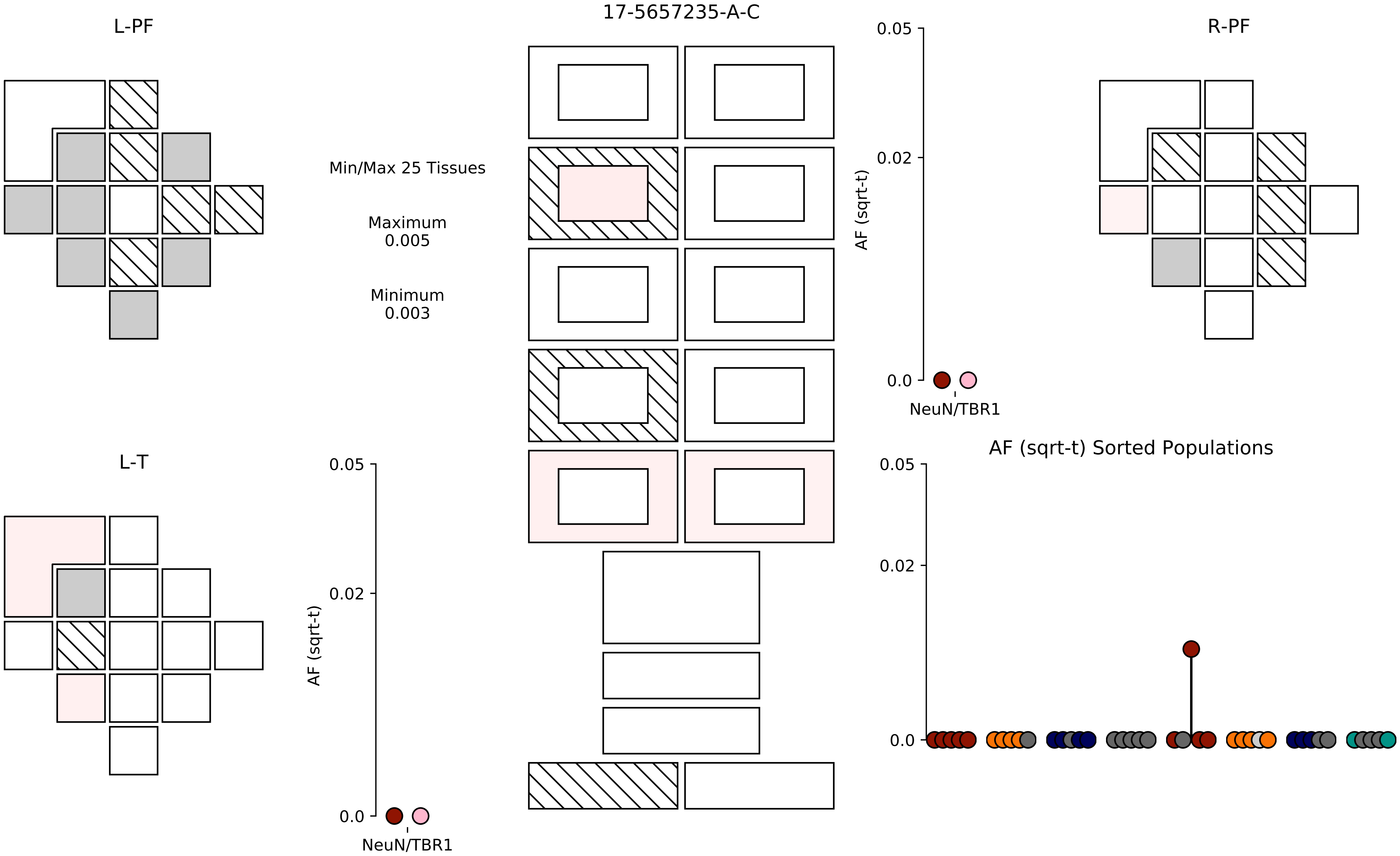

17-5657235-A-C
L-PF
R-PF
L-T
Min/Max 25 Tissues
Maximum
0.005
Minimum
0.003
AF (sqrt-t)
0.05
0.02
0.0
NeuN/TBR1
AF (sqrt-t) Sorted Populations

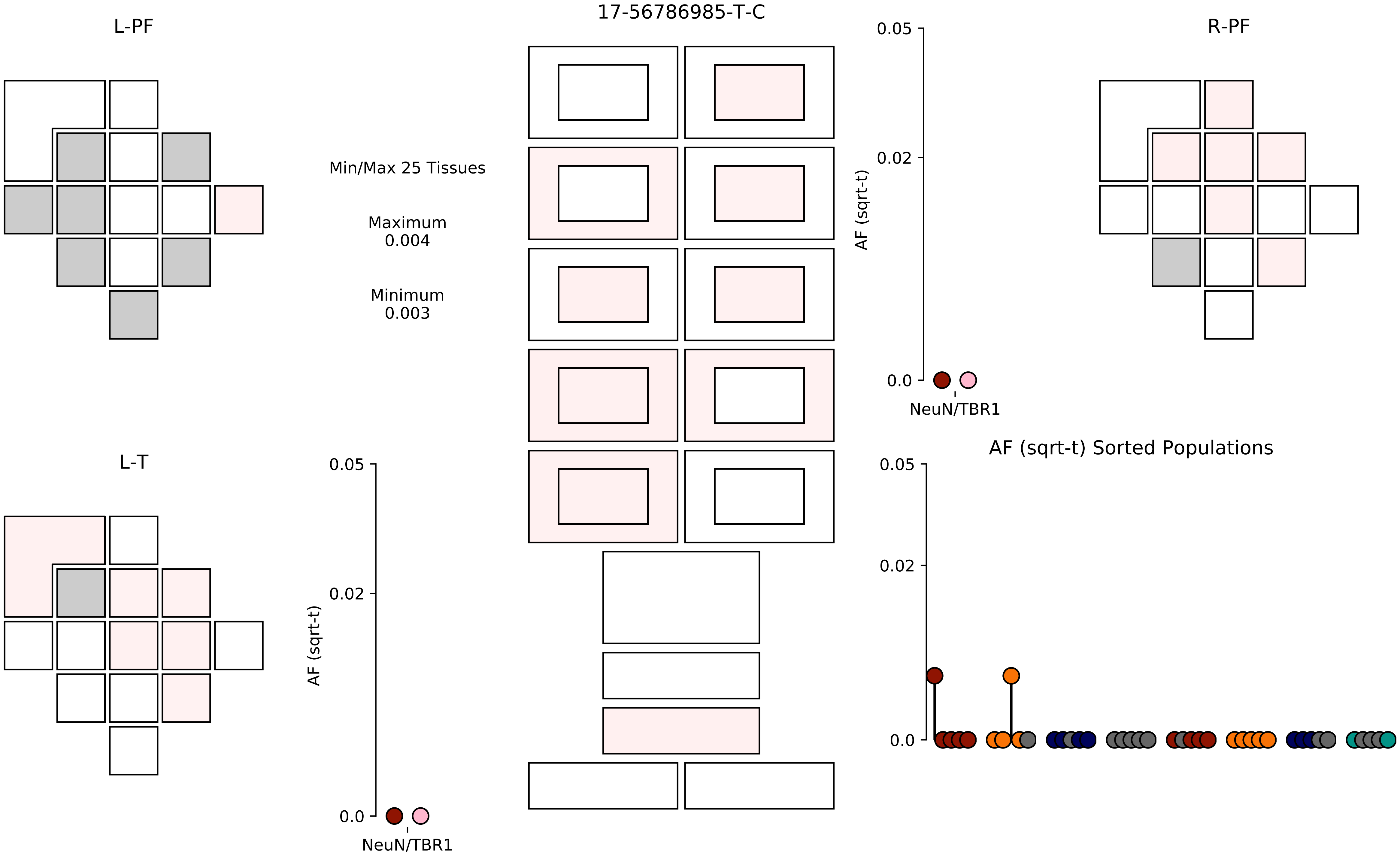

17-56786985-T-C
L-PF
R-PF
L-T
Min/Max 25 Tissues
Maximum
0.004
Minimum
0.003
AF (sqrt-t)
0.05
0.02
0.0
NeuN/TBR1
AF (sqrt-t) Sorted Populations

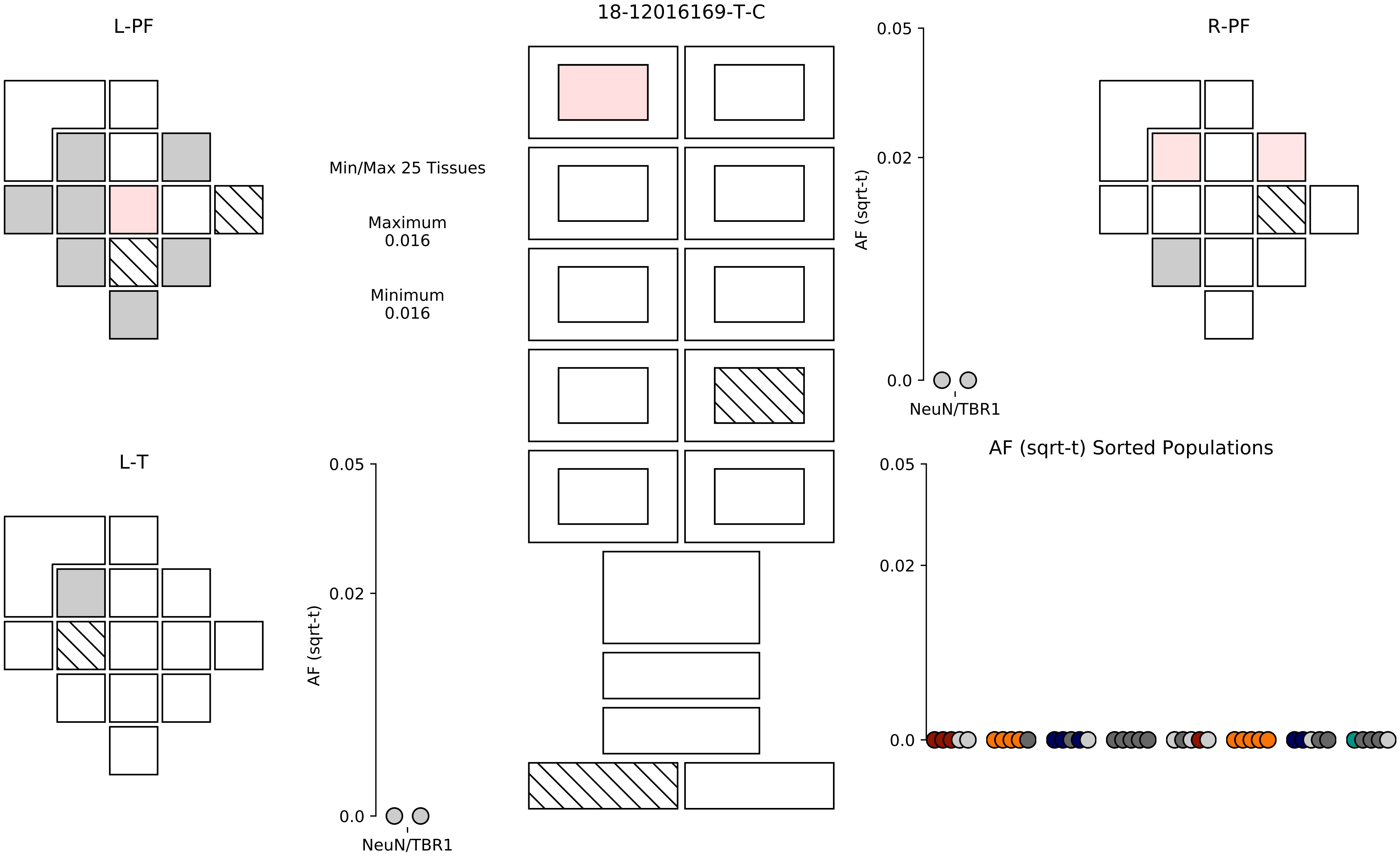

18-12016169-T-C
L-PF
R-PF
L-T
Min/Max 25 Tissues
Maximum
0.016
Minimum
0.016
AF (sqrt-t)
0.05
0.02
0.0
NeuN/TBR1
AF (sqrt-t) Sorted Populations

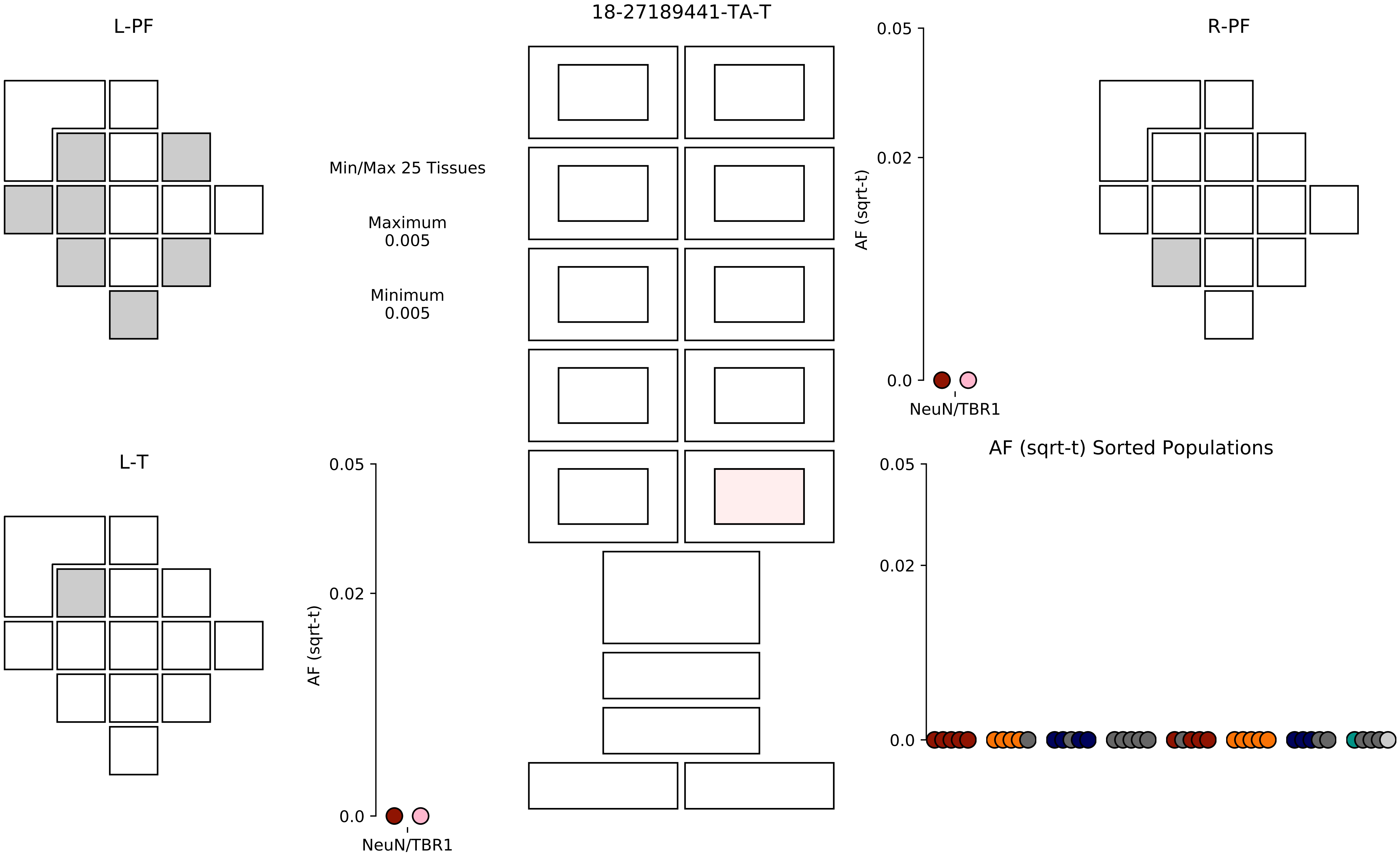

18-27189441-TA-T
L-PF
R-PF
L-T
Min/Max 25 Tissues
Maximum
0.005
Minimum
0.005
AF (sqrt-t)
0.05
0.02
0.0
NeuN/TBR1
AF (sqrt-t) Sorted Populations

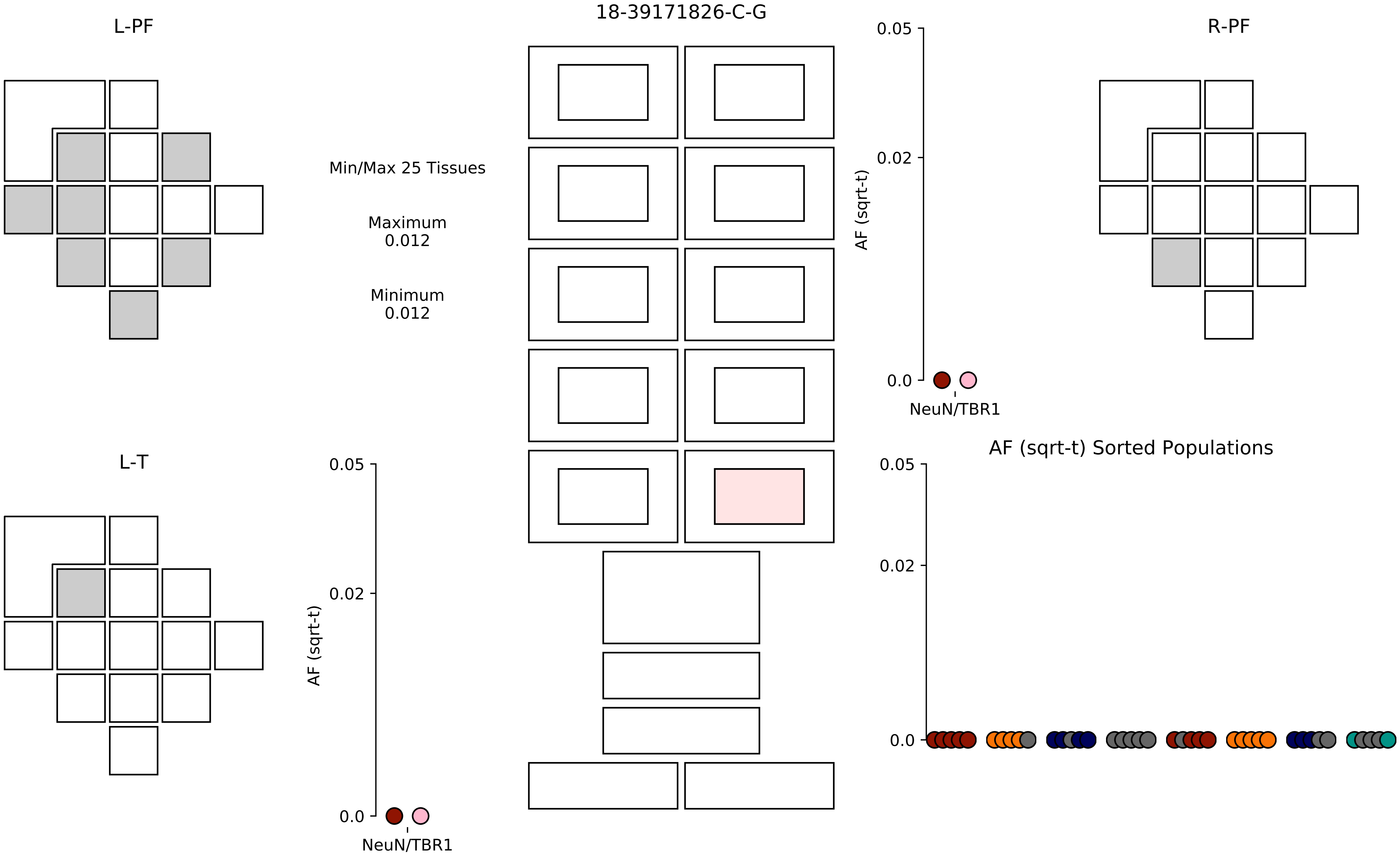

18-39171826-C-G
L-PF
R-PF
L-T
Min/Max 25 Tissues
Maximum
0.012
Minimum
0.012
AF (sqrt-t)
0.05
0.02
0.0
NeuN/TBR1
AF (sqrt-t) Sorted Populations

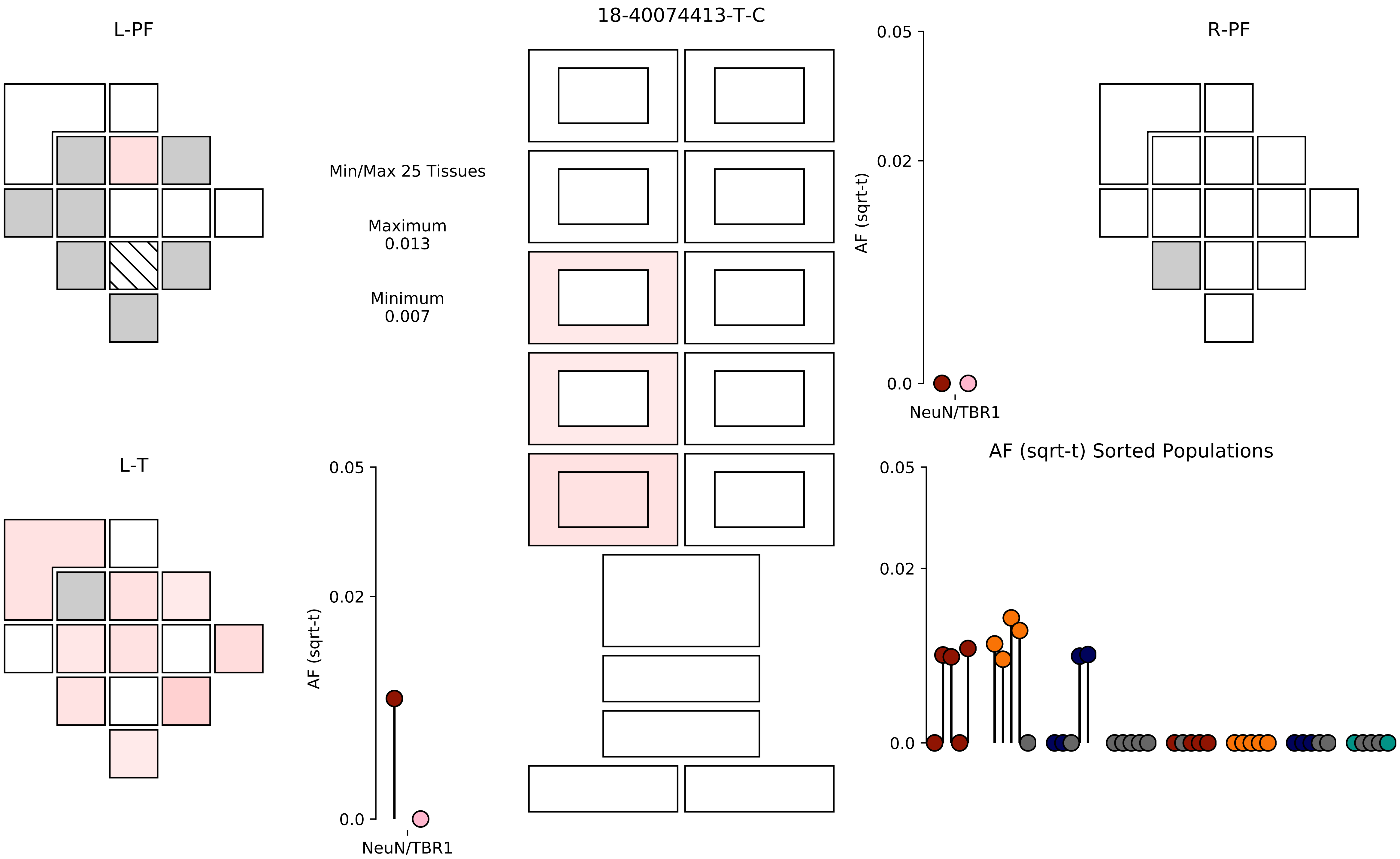

18-40074413-T-C
L-PF
R-PF
L-T
Min/Max 25 Tissues
Maximum
0.013
Minimum
0.007
AF (sqrt-t)
0.05
0.02
0.0
NeuN/TBR1
AF (sqrt-t) Sorted Populations

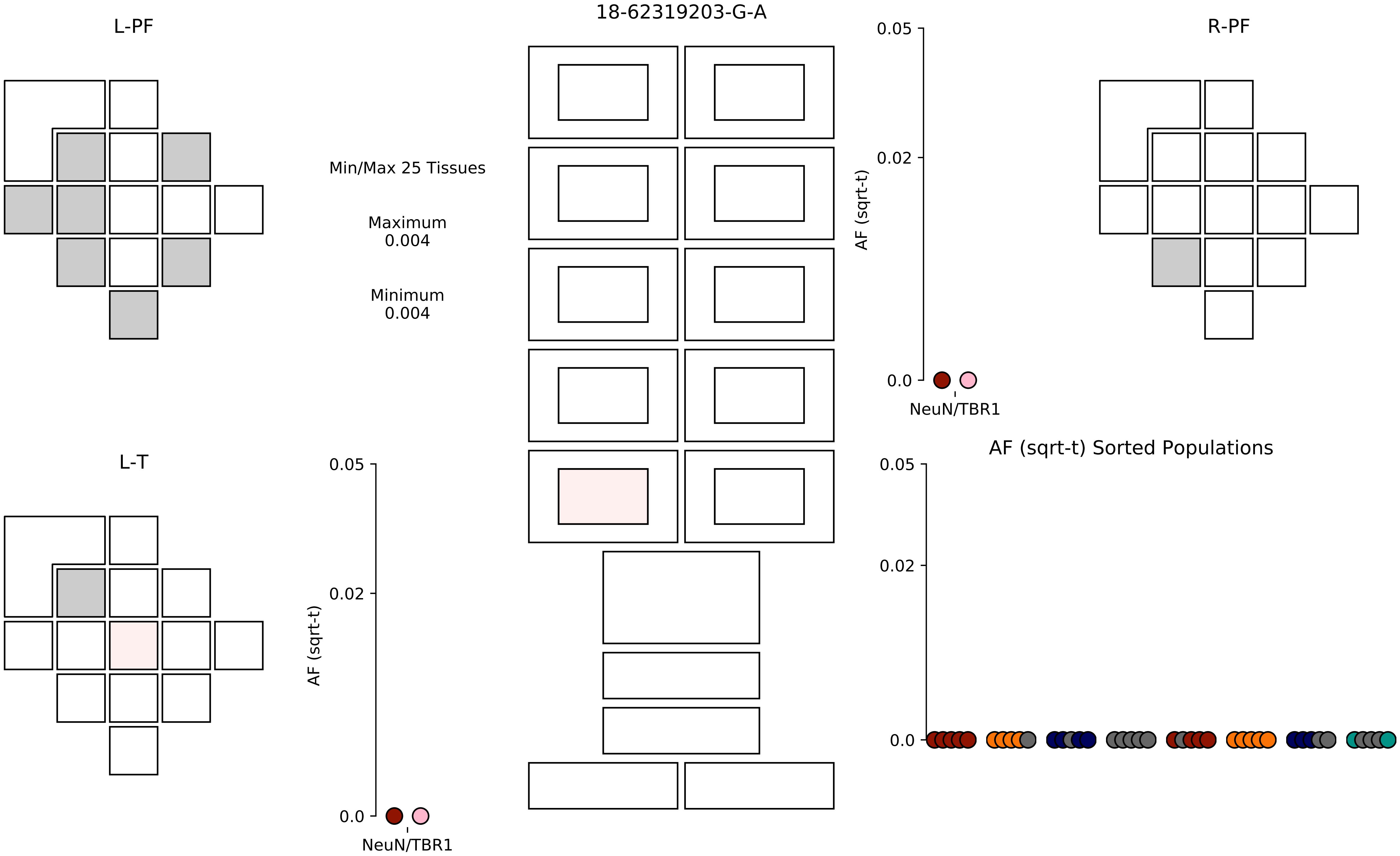

18-62319203-G-A
L-PF
R-PF
L-T
Min/Max 25 Tissues
Maximum
0.004
Minimum
0.004
AF (sqrt-t)
0.05
0.02
0.0
NeuN/TBR1
AF (sqrt-t) Sorted Populations

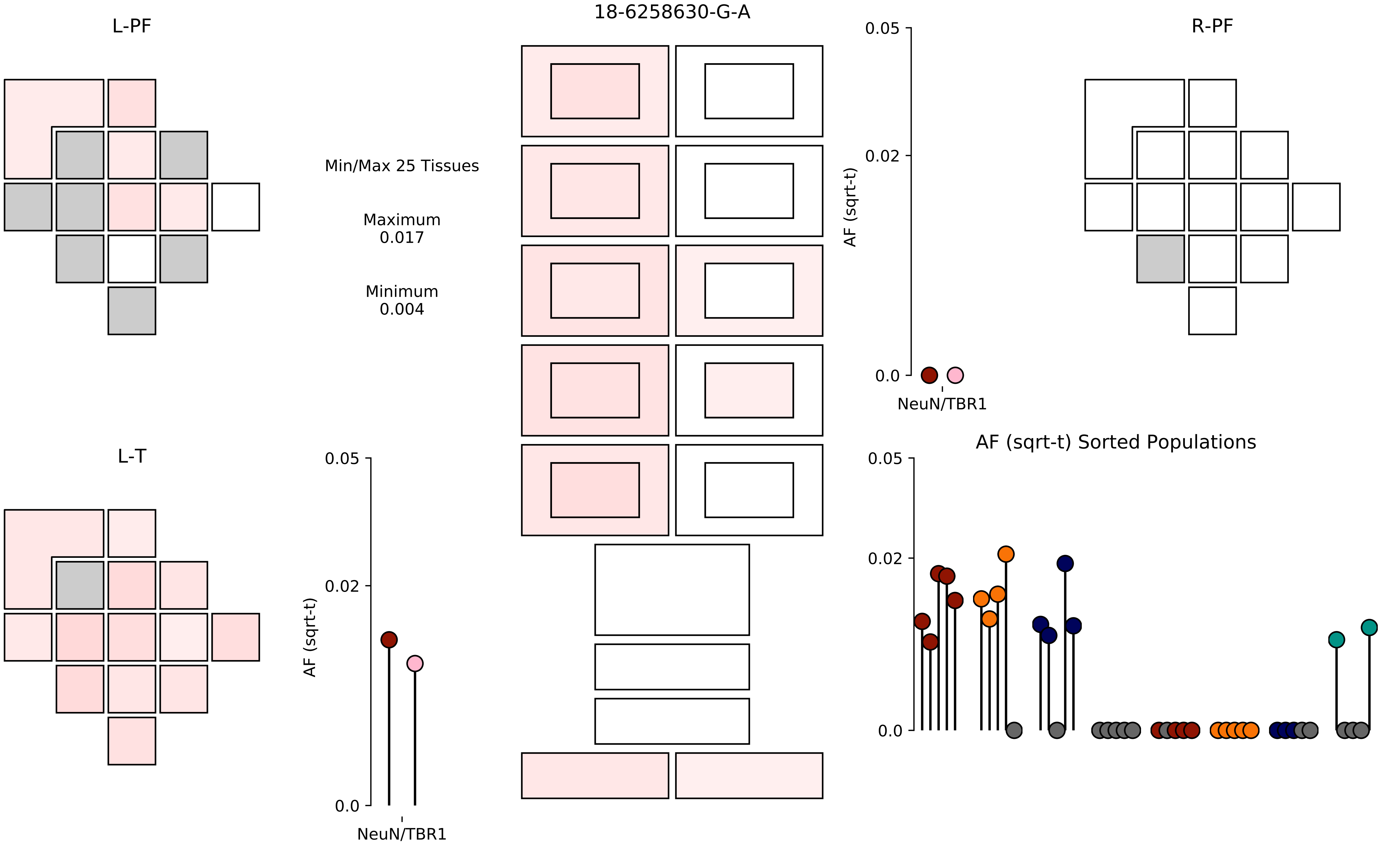

18-6258630-G-A
L-PF
R-PF
L-T
Min/Max 25 Tissues
Maximum
0.017
Minimum
0.004
AF (sqrt-t)
0.05
0.02
0.0
NeuN/TBR1
AF (sqrt-t) Sorted Populations

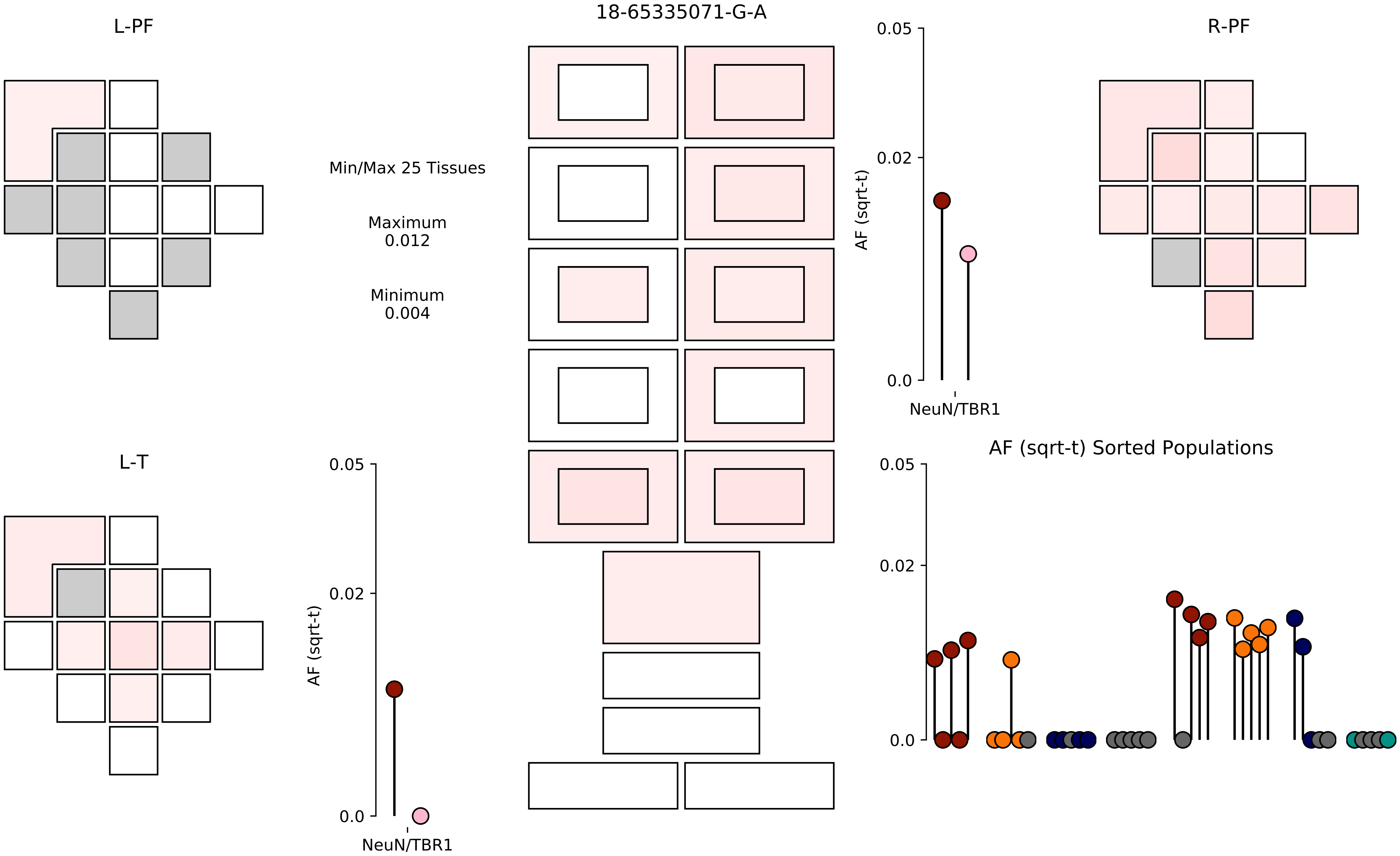

18-65335071-G-A
L-PF
R-PF
L-T
Min/Max 25 Tissues
Maximum
0.012
Minimum
0.004
AF (sqrt-t)
0.05
0.02
0.0
NeuN/TBR1
AF (sqrt-t) Sorted Populations

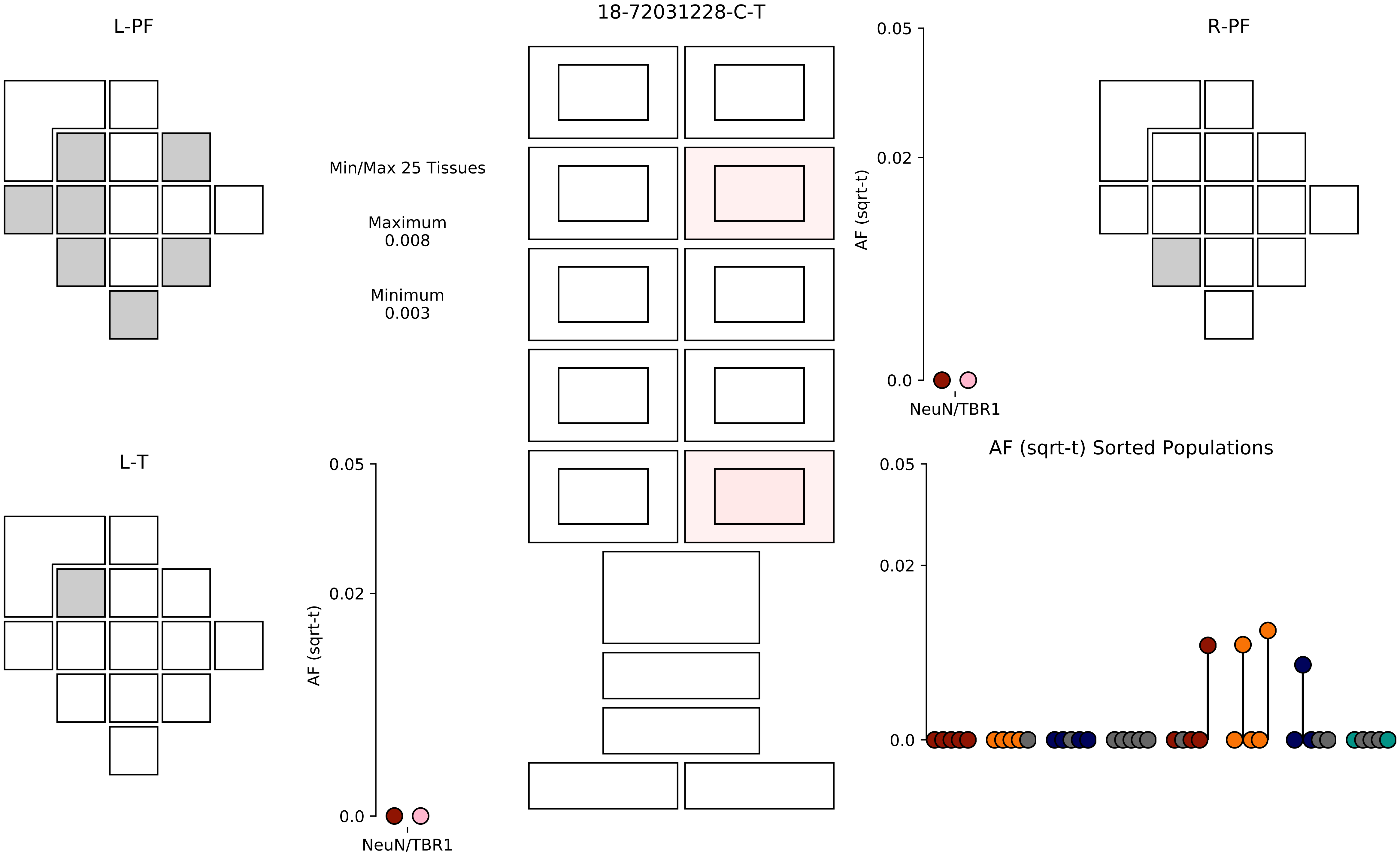

18-72031228-C-T
L-PF
R-PF
L-T
Min/Max 25 Tissues
Maximum
0.008
Minimum
0.003
AF (sqrt-t)
0.05
0.02
0.0
NeuN/TBR1
AF (sqrt-t)
0.05
0.02
0.0
NeuN/TBR1
AF (sqrt-t) Sorted Populations
0.05
0.02
0.0

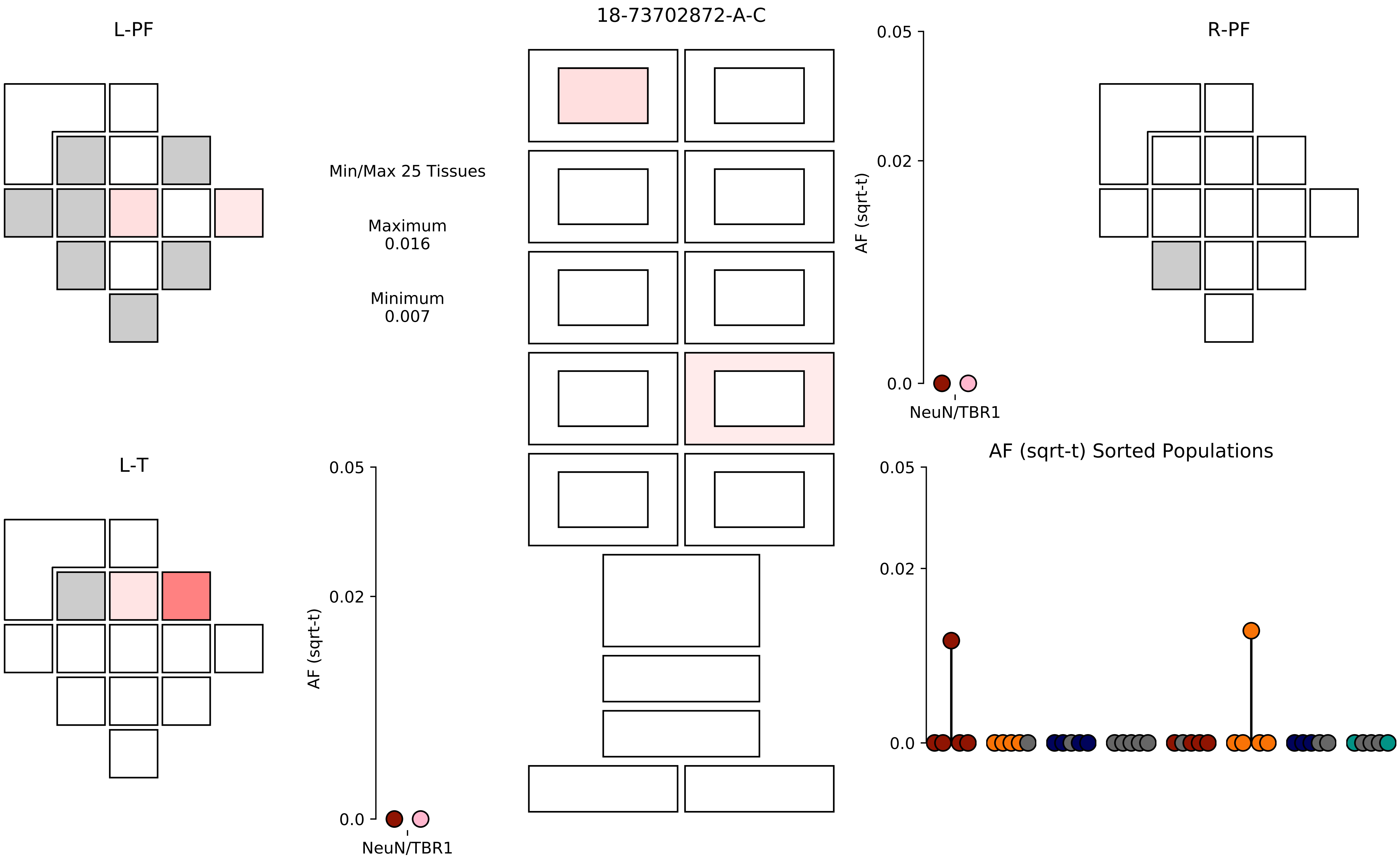

18-73702872-A-C
L-PF
R-PF
L-T
Min/Max 25 Tissues
Maximum
0.016
Minimum
0.007
AF (sqrt-t)
0.05
0.02
0.0
NeuN/TBR1
AF (sqrt-t) Sorted Populations

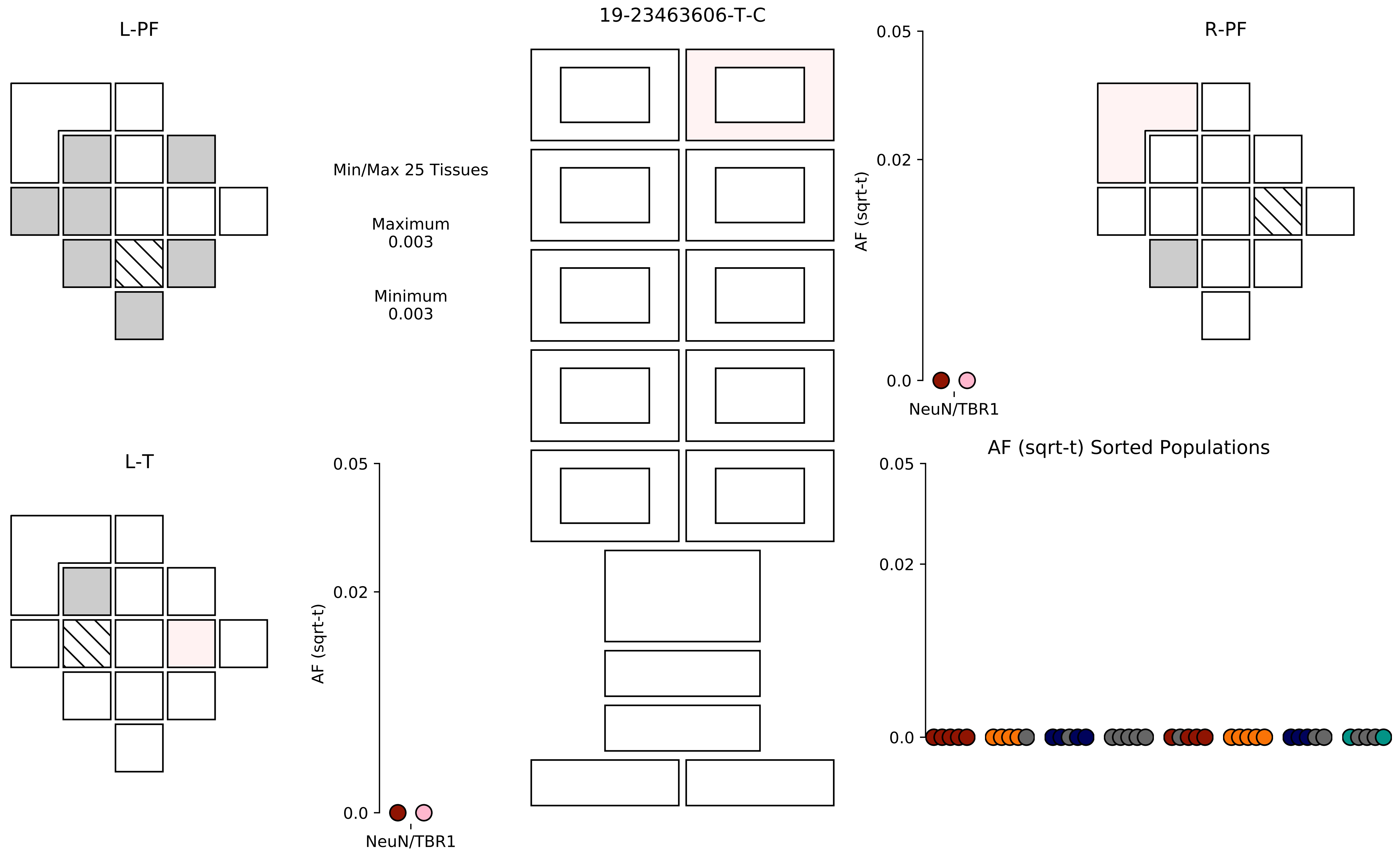
19-23463606-T-C
L-PF
R-PF
L-T
Min/Max 25 Tissues
Maximum
0.003
Minimum
0.003
AF (sqrt-t)
0.05
0.02
0.0
NeuN/TBR1
AF (sqrt-t) Sorted Populations

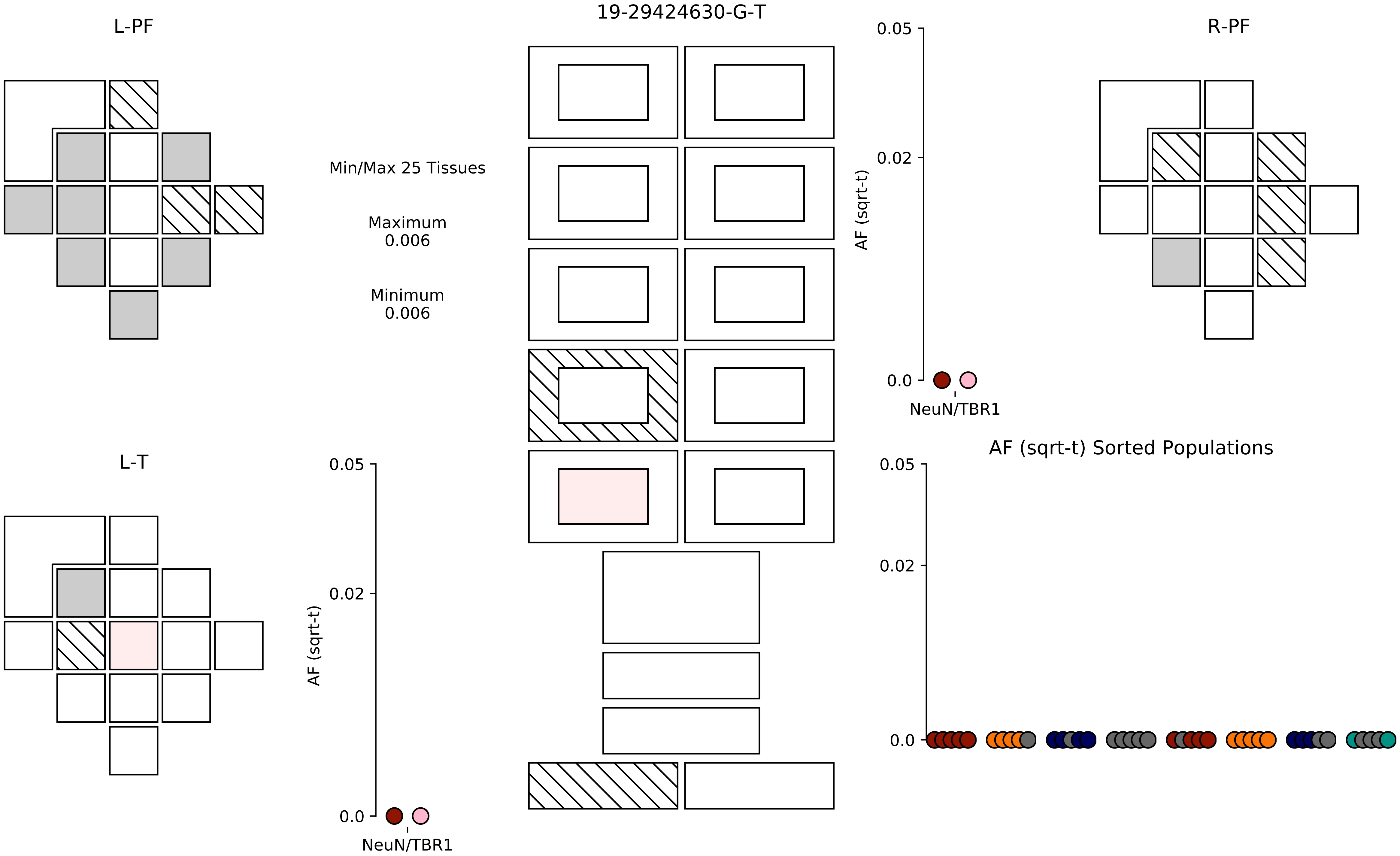
19-29424630-G-T
L-PF
R-PF
L-T
Min/Max 25 Tissues
Maximum
0.006
Minimum
0.006
AF (sqrt-t)
0.05
0.02
0.0
NeuN/TBR1
AF (sqrt-t)
0.05
0.02
0.0
NeuN/TBR1
AF (sqrt-t) Sorted Populations
0.05
0.02
0.0

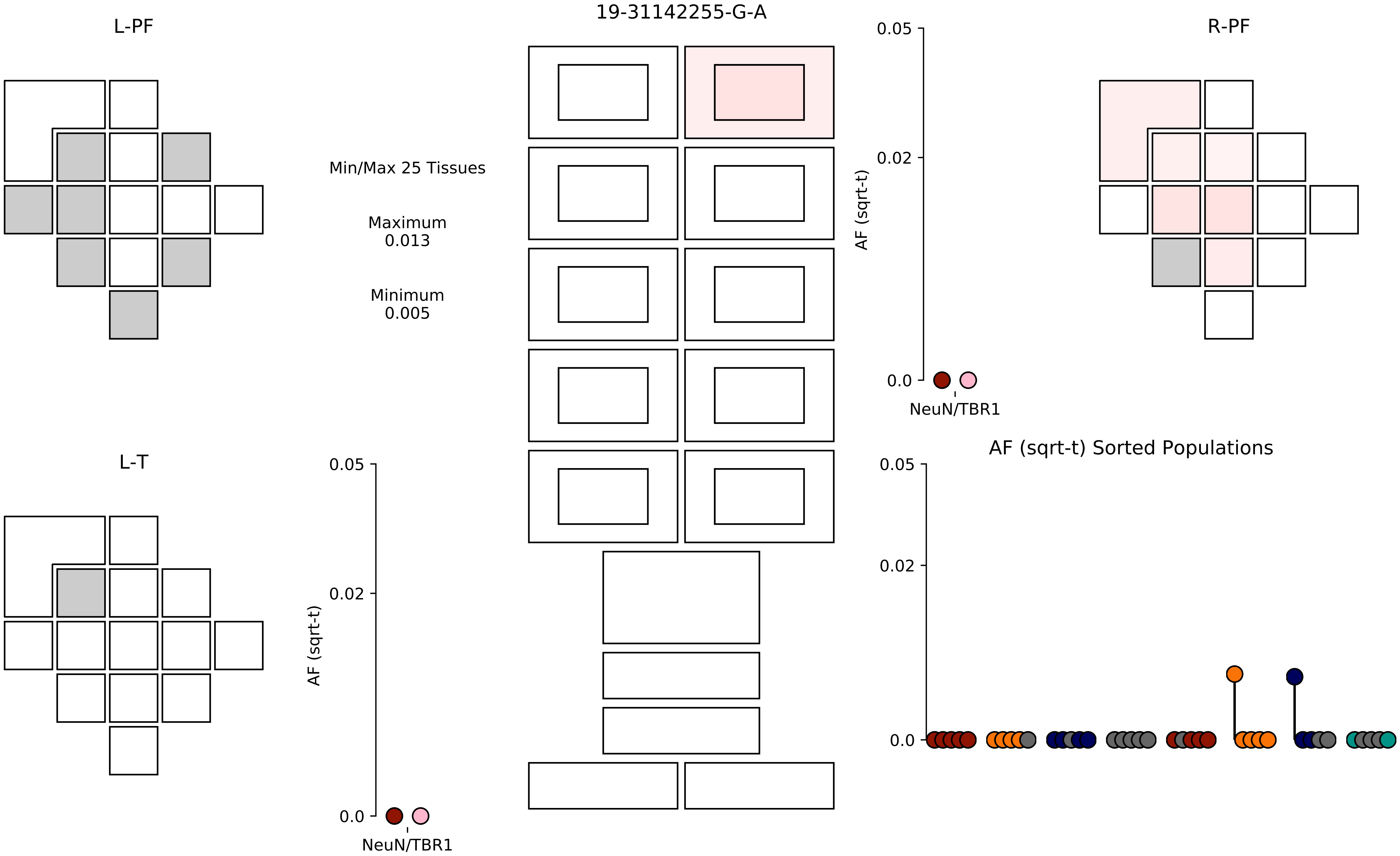

19-31142255-G-A
L-PF
R-PF
L-T
Min/Max 25 Tissues
Maximum
0.013
Minimum
0.005
AF (sqrt-t)
0.05
0.02
0.0
NeuN/TBR1
AF (sqrt-t) Sorted Populations

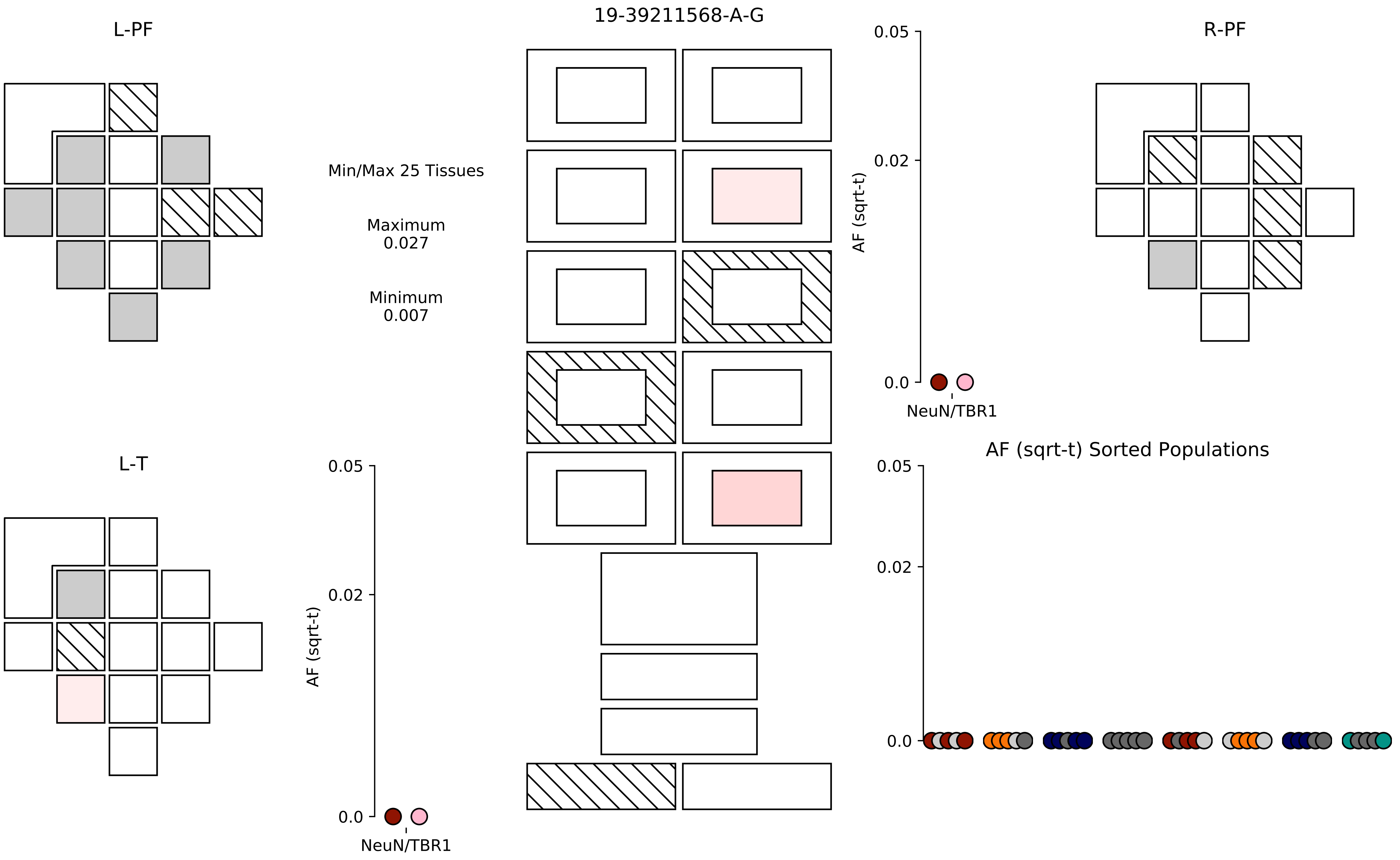

19-39211568-A-G
L-PF
R-PF
L-T
Min/Max 25 Tissues
Maximum
0.027
Minimum
0.007
AF (sqrt-t)
0.05
0.02
0.0
NeuN/TBR1
AF (sqrt-t) Sorted Populations

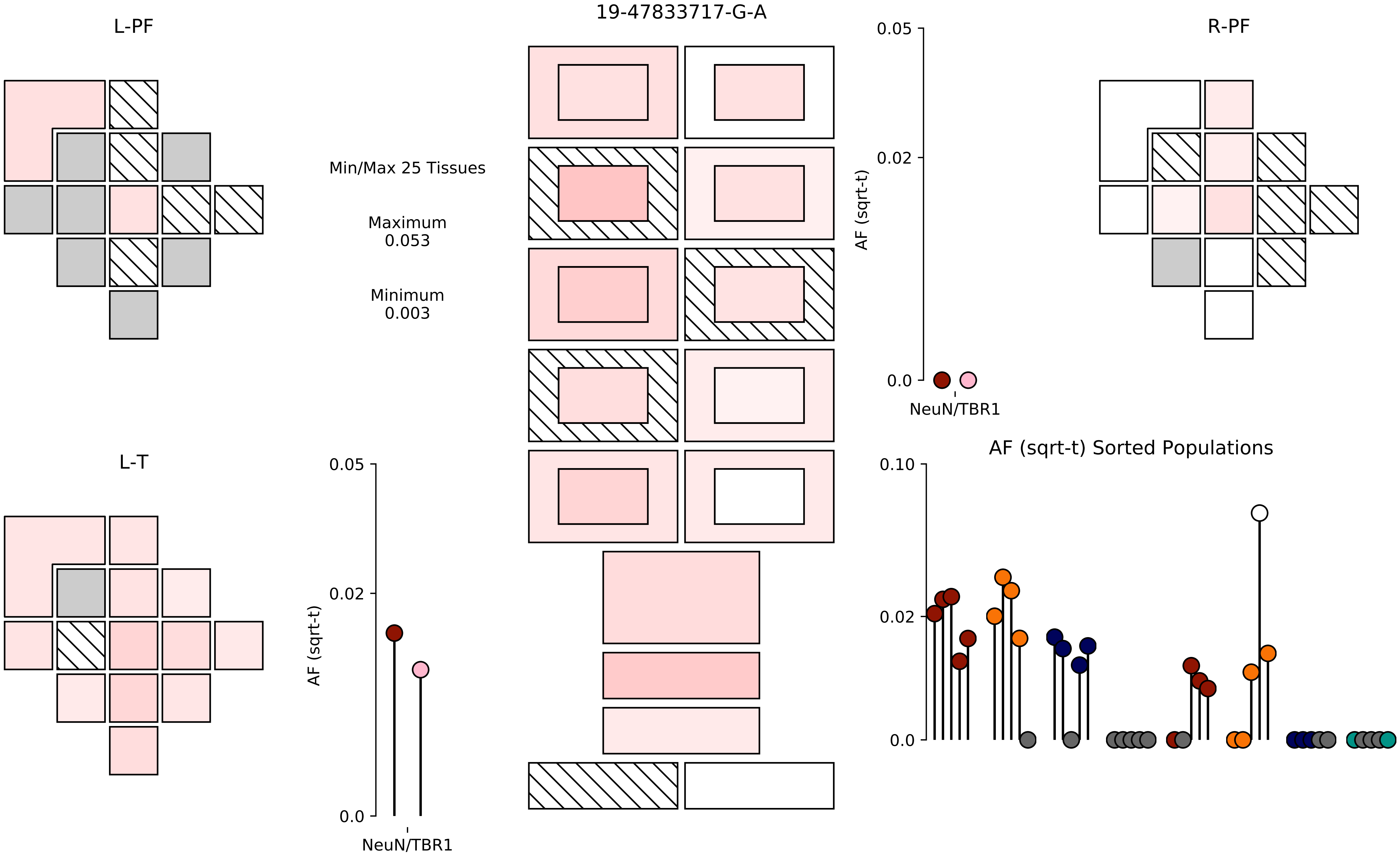

L-PF
19-47833717-G-A
R-PF
Min/Max 25 Tissues
Maximum
0.053
Minimum
0.003
0.05
0.02
0.0
AF (sqrt-t)
NeuN/TBR1
L-T
0.05
0.02
0.0
AF (sqrt-t)
NeuN/TBR1
AF (sqrt-t) Sorted Populations
0.10
0.02
0.0

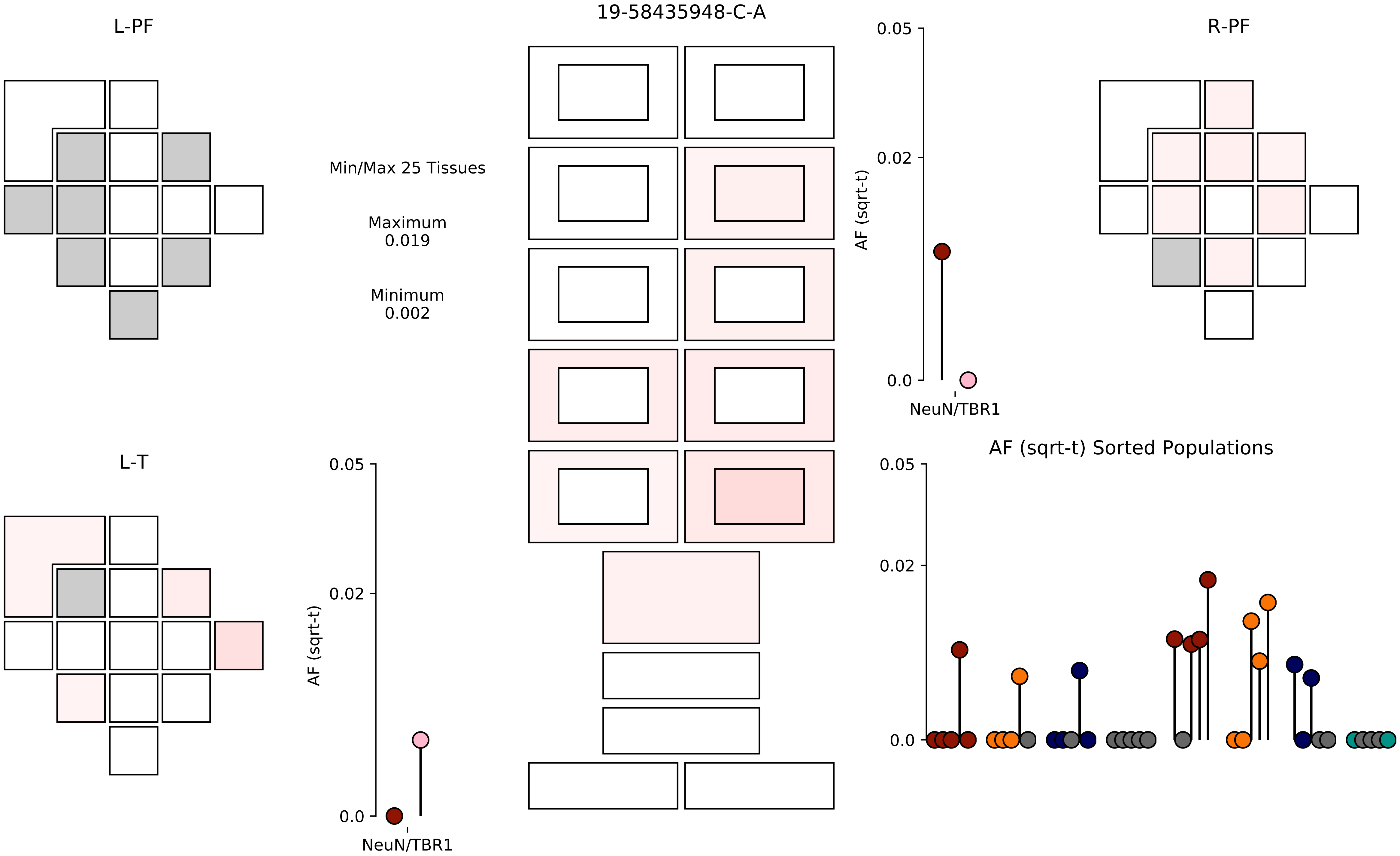

19-58435948-C-A
L-PF
R-PF
L-T
Min/Max 25 Tissues
Maximum
0.019
Minimum
0.002
AF (sqrt-t)
0.05
0.02
0.0
NeuN/TBR1
AF (sqrt-t) Sorted Populations

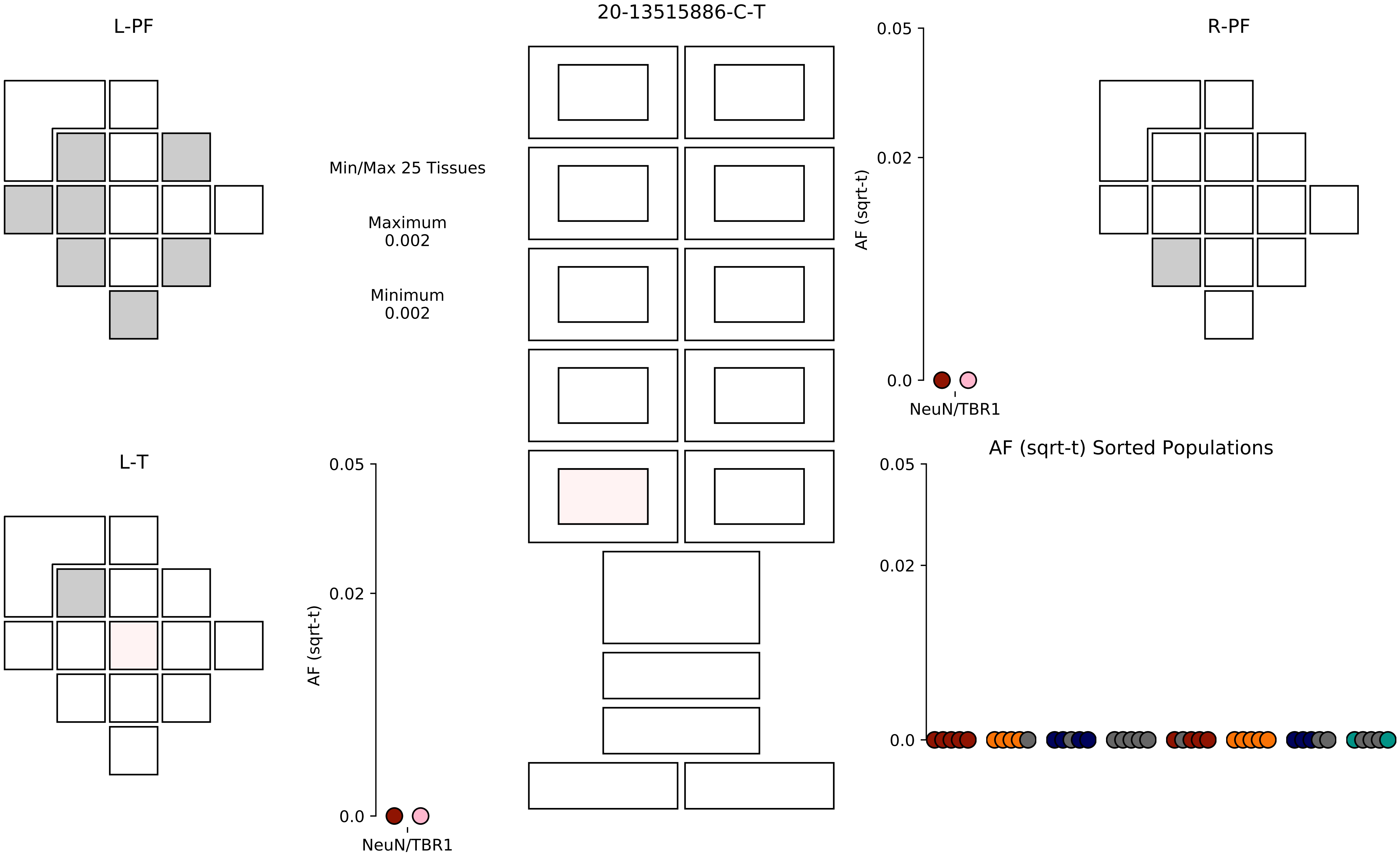
20-13515886-C-T
L-PF
R-PF
L-T
Min/Max 25 Tissues
Maximum
0.002
Minimum
0.002
AF (sqrt-t)
0.05
0.02
0.0
NeuN/TBR1
AF (sqrt-t) Sorted Populations

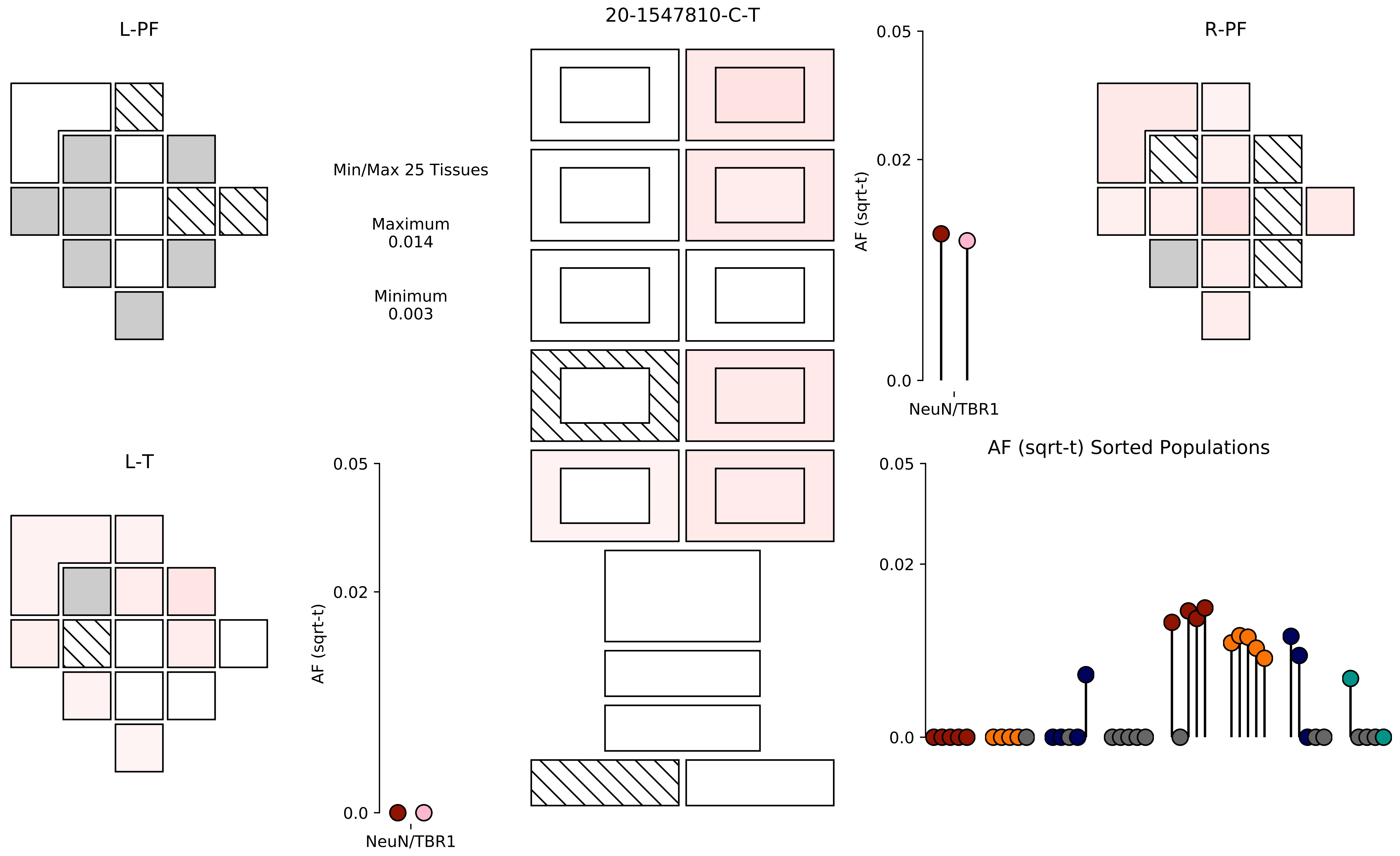

20-1547810-C-T
L-PF
R-PF
L-T
Min/Max 25 Tissues
Maximum
0.014
Minimum
0.003
AF (sqrt-t)
0.05
0.02
0.0
NeuN/TBR1
AF (sqrt-t)
0.05
0.02
0.0
NeuN/TBR1
AF (sqrt-t) Sorted Populations
0.05
0.02
0.0

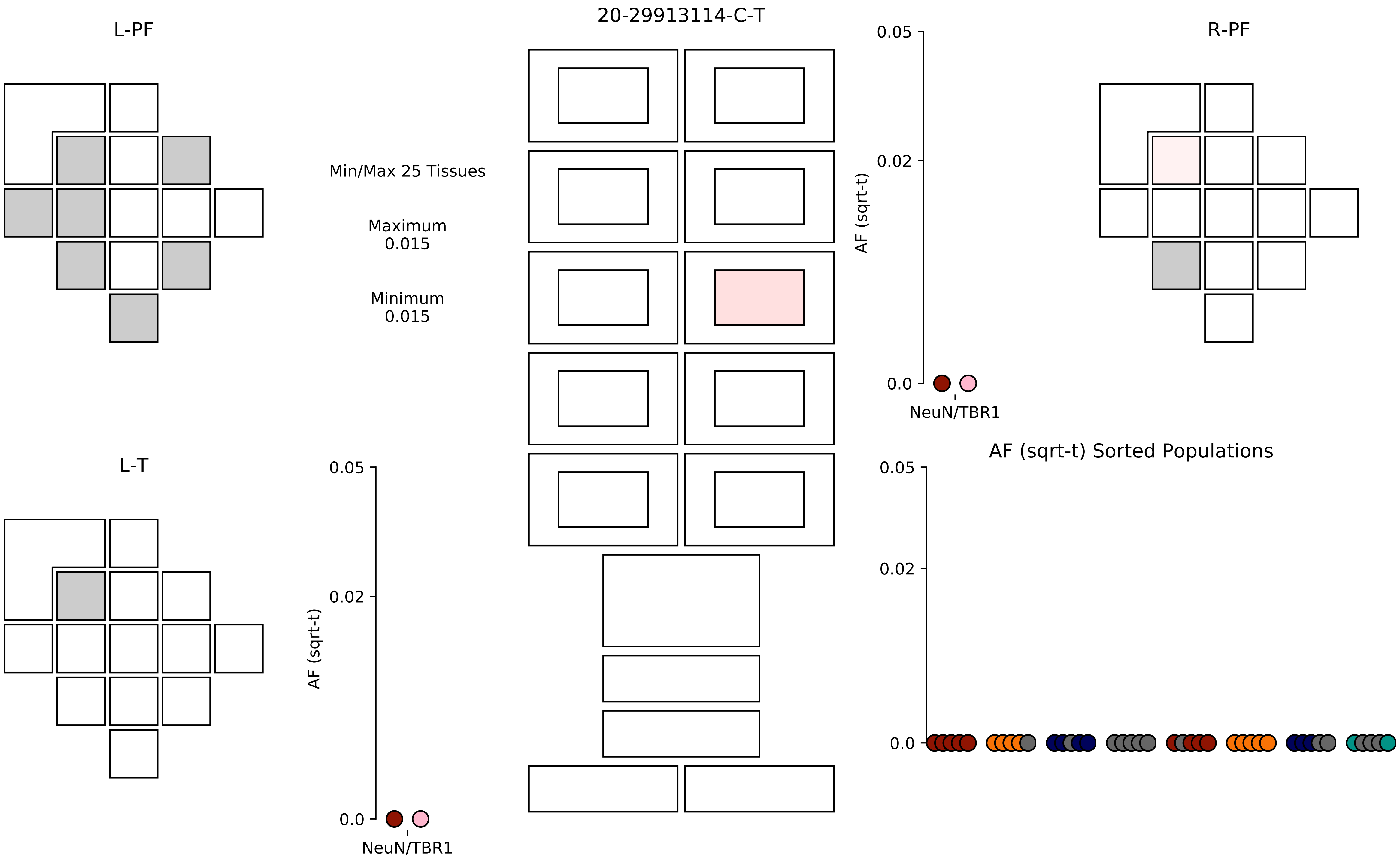

20-29913114-C-T
L-PF
R-PF
L-T
Min/Max 25 Tissues
Maximum
0.015
Minimum
0.015
0.05
0.02
0.0
AF (sqrt-t)
NeuN/TBR1
AF (sqrt-t) Sorted Populations

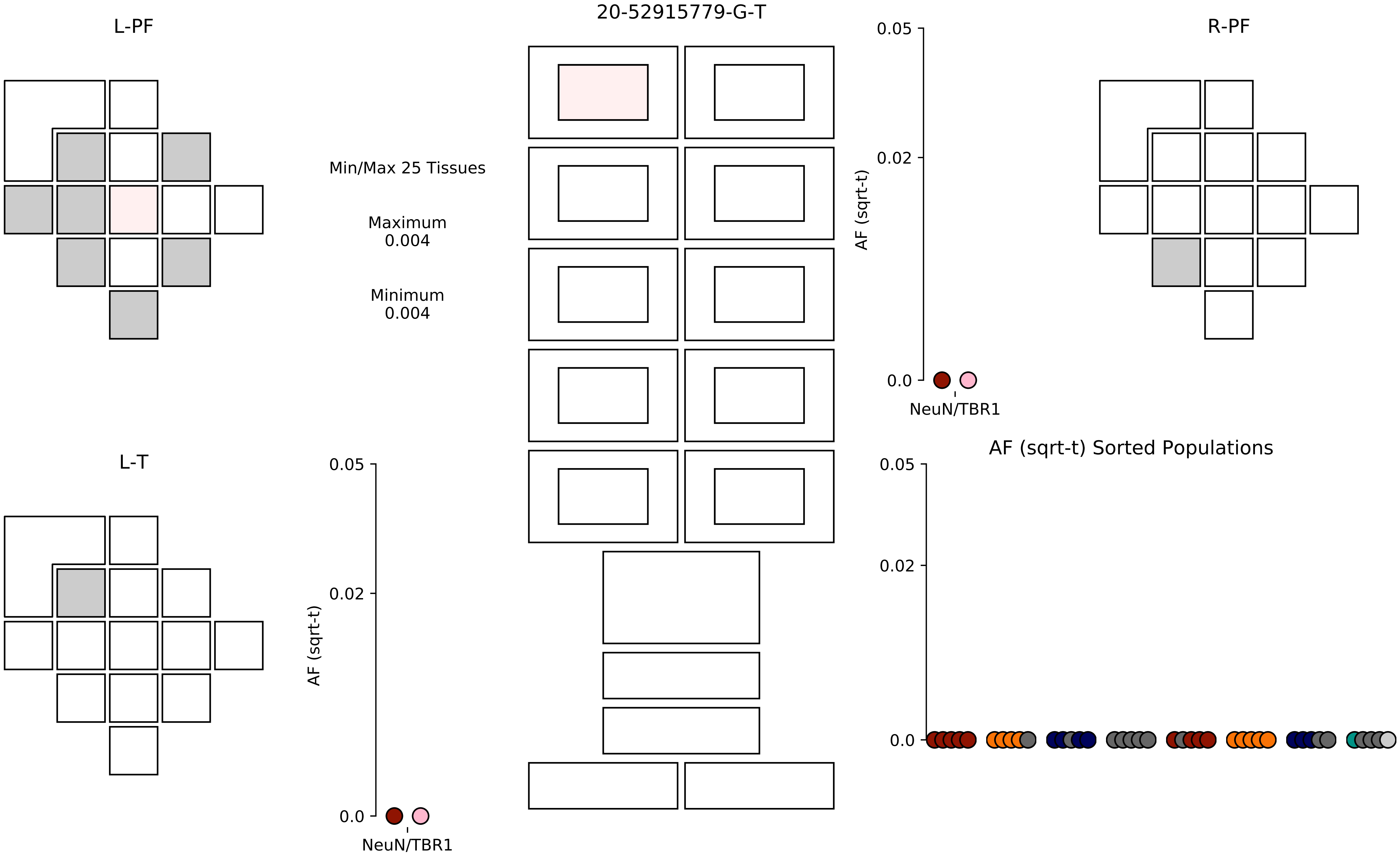

20-52915779-G-T
L-PF
R-PF
L-T
Min/Max 25 Tissues
Maximum
0.004
Minimum
0.004
AF (sqrt-t)
0.05
0.02
0.0
NeuN/TBR1
AF (sqrt-t) Sorted Populations

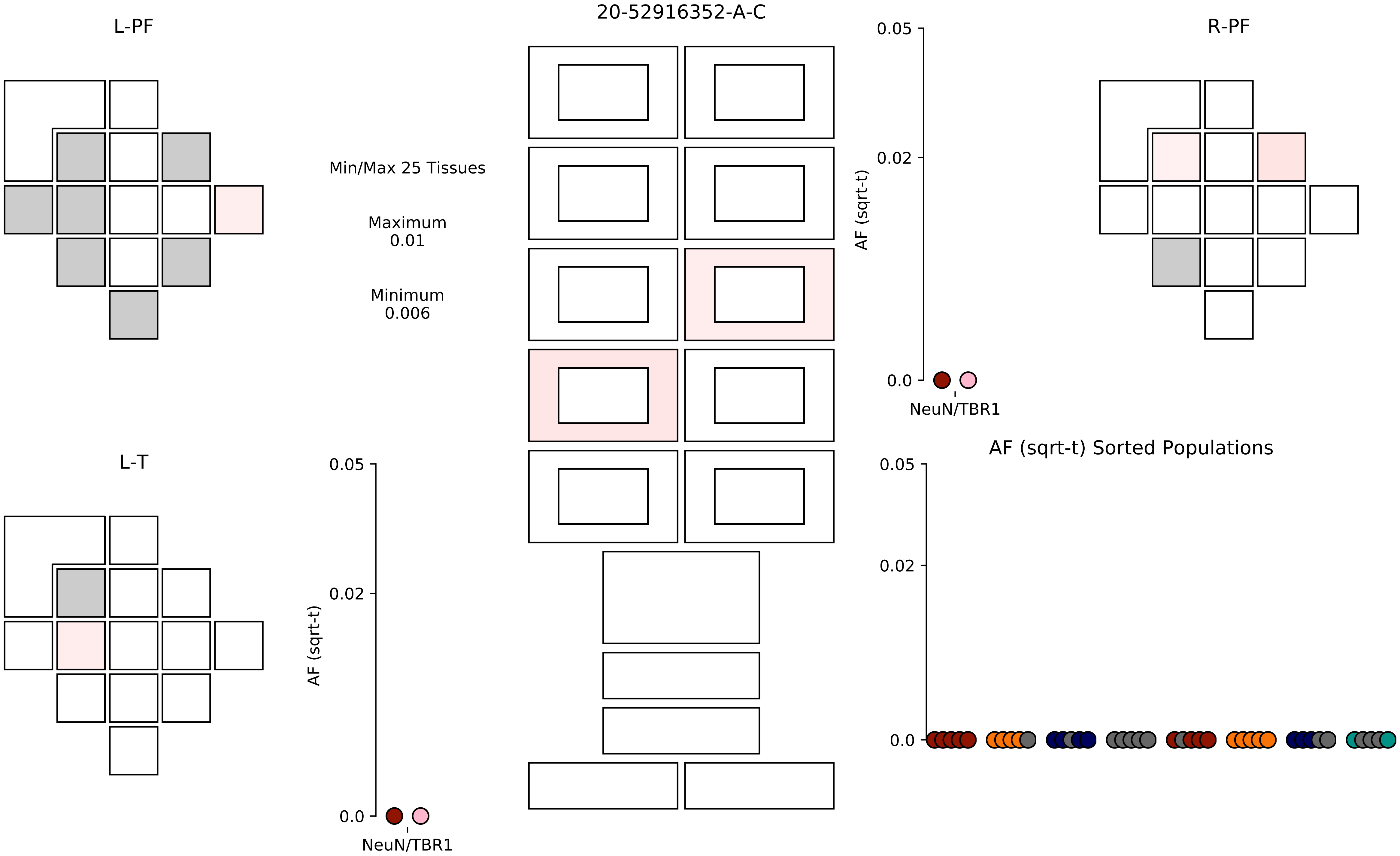

20-52916352-A-C
L-PF
R-PF
L-T
Min/Max 25 Tissues
Maximum
0.01
Minimum
0.006
AF (sqrt-t)
0.05
0.02
0.0
NeuN/TBR1
AF (sqrt-t)
0.05
0.02
0.0
NeuN/TBR1
AF (sqrt-t) Sorted Populations
0.05
0.02
0.0

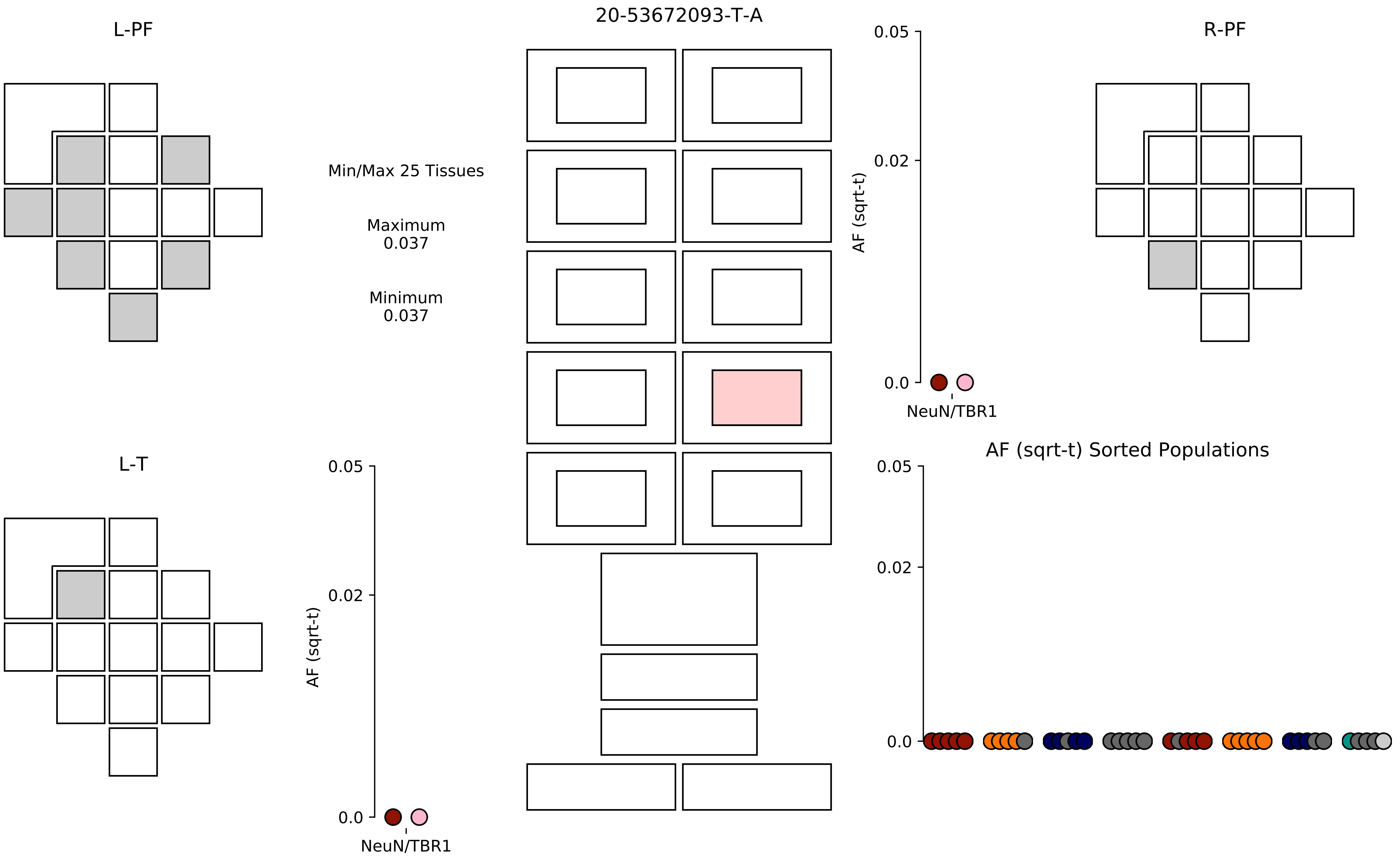

20-53672093-T-A
L-PF
R-PF
L-T
Min/Max 25 Tissues
Maximum
0.037
Minimum
0.037
AF (sqrt-t)
0.05
0.02
0.0
NeuN/TBR1
AF (sqrt-t) Sorted Populations

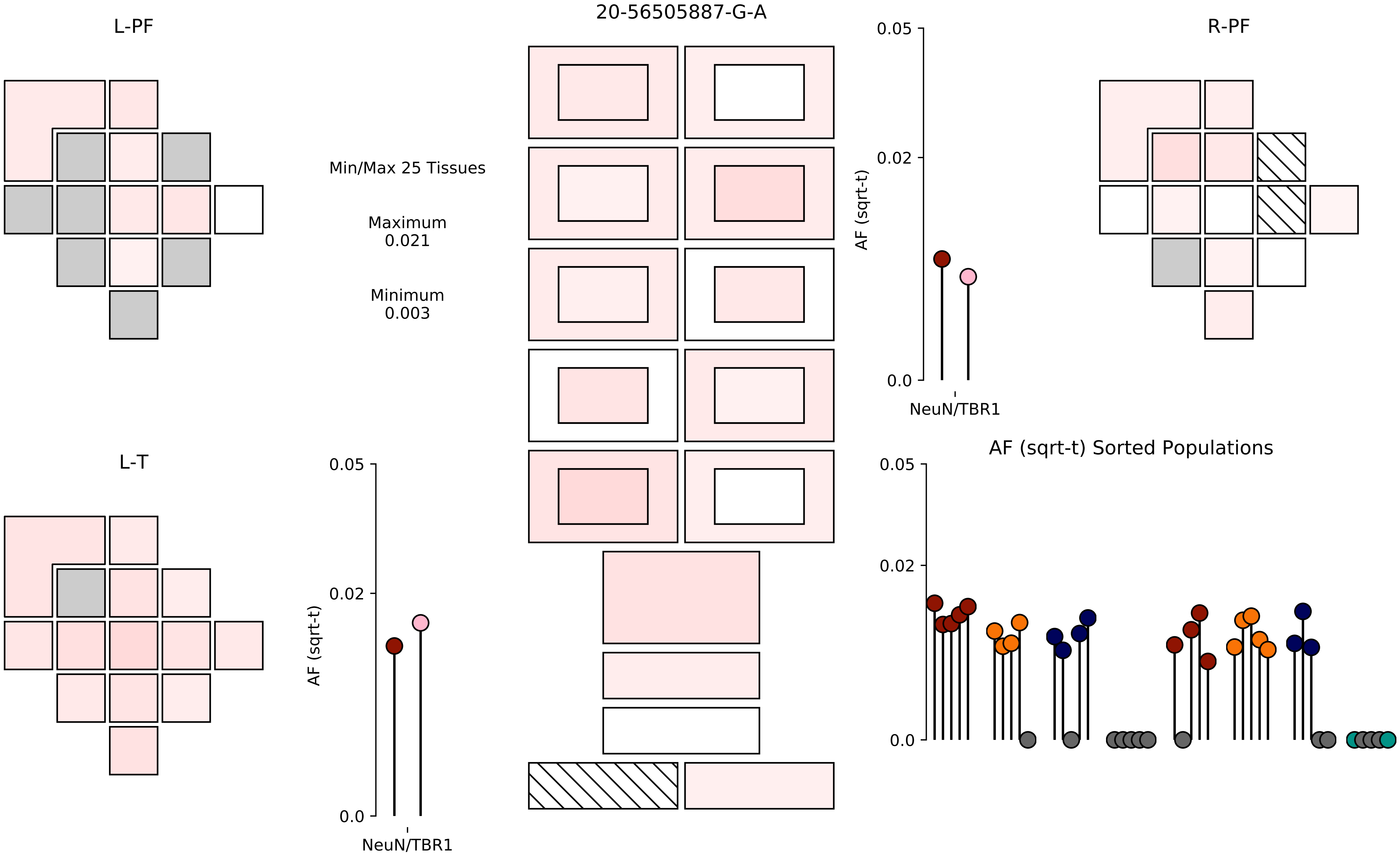

20-56505887-G-A
L-PF
R-PF
L-T
Min/Max 25 Tissues
Maximum
0.021
Minimum
0.003
AF (sqrt-t)
0.05
0.02
0.0
NeuN/TBR1
AF (sqrt-t) Sorted Populations

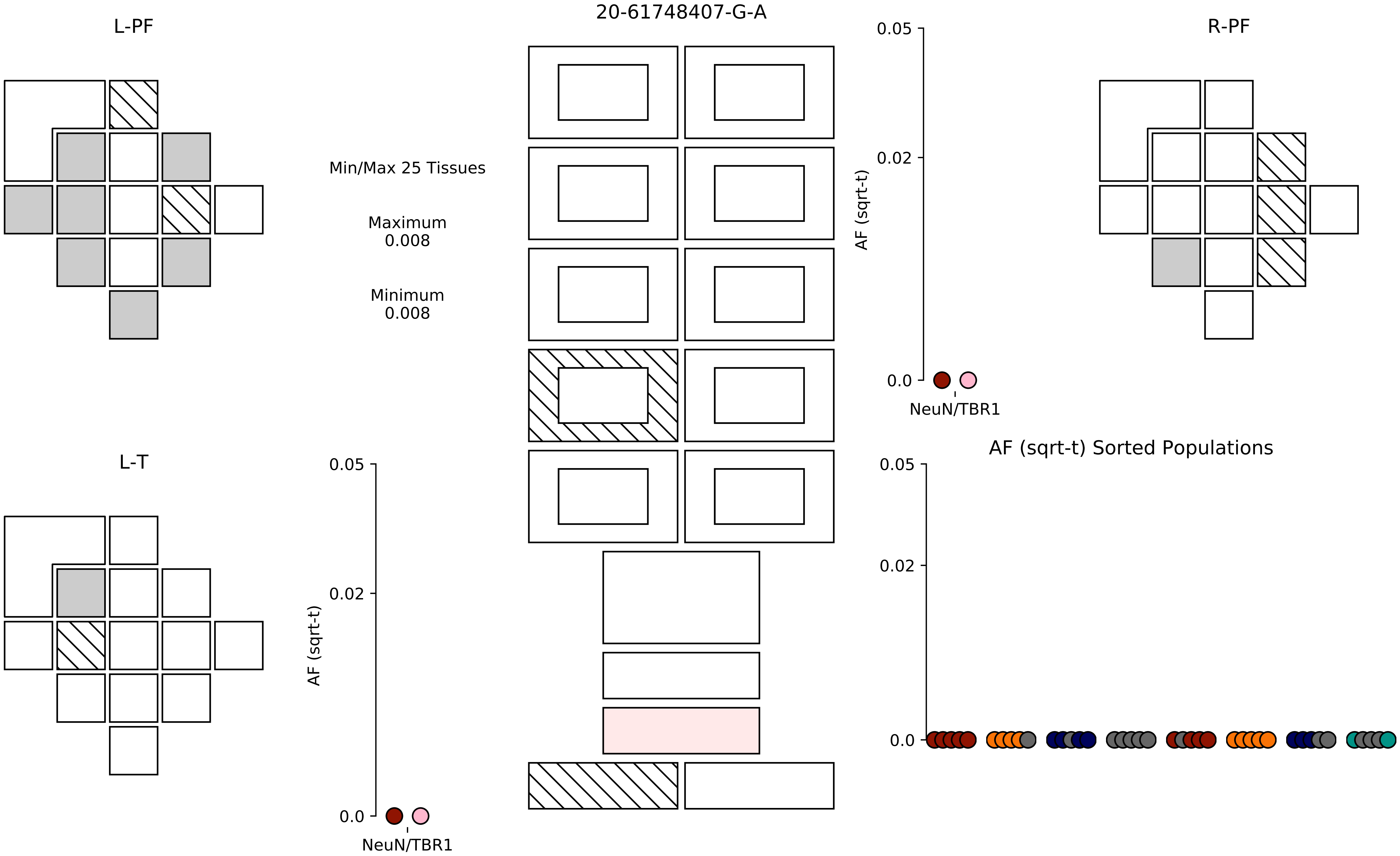

20-61748407-G-A
L-PF
R-PF
L-T
Min/Max 25 Tissues
Maximum
0.008
Minimum
0.008
AF (sqrt-t)
0.05
0.02
0.0
NeuN/TBR1
AF (sqrt-t) Sorted Populations

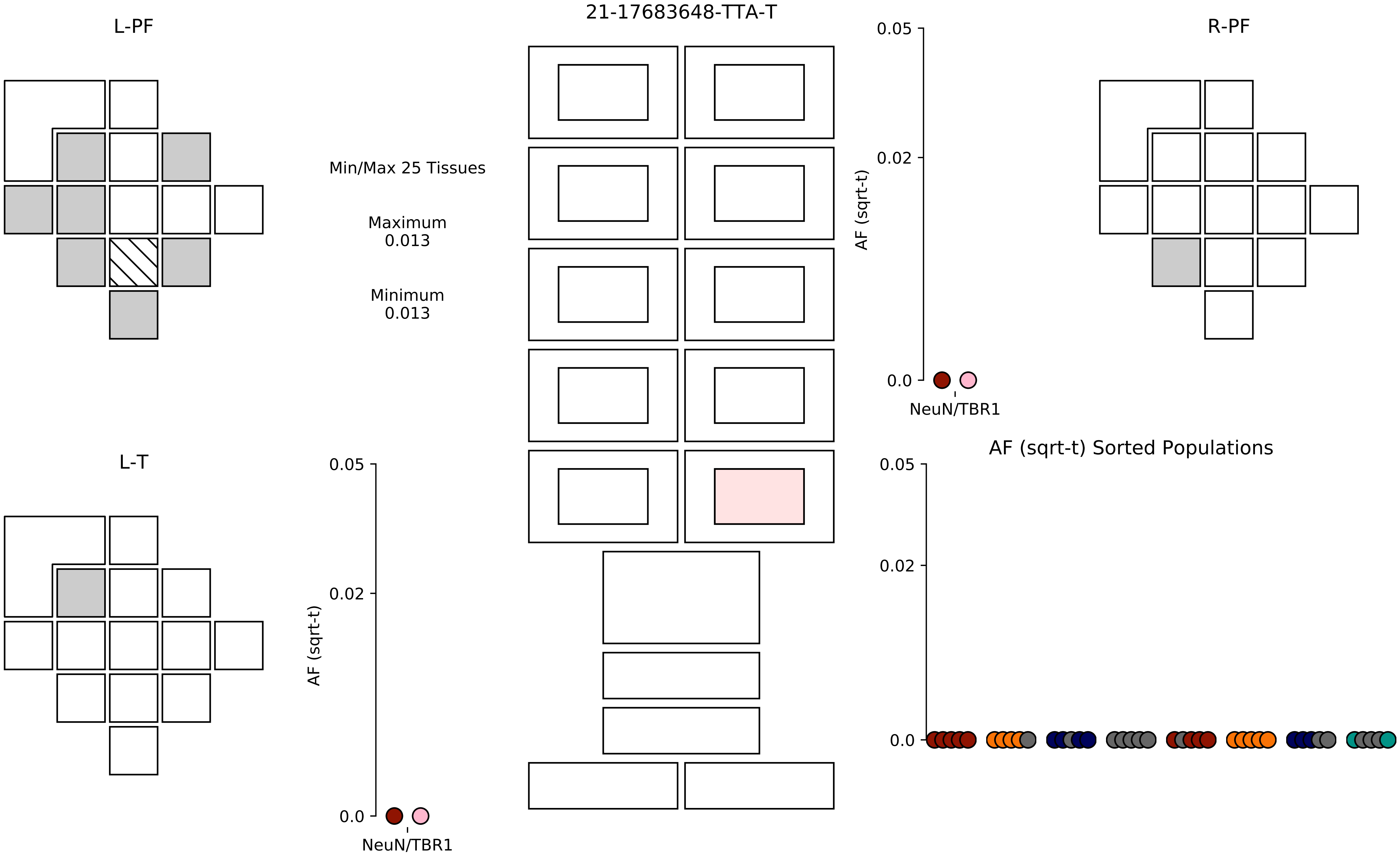
21-17683648-TTA-T
L-PF
R-PF
L-T
Min/Max 25 Tissues
Maximum
0.013
Minimum
0.013
AF (sqrt-t)
0.05
0.02
0.0
NeuN/TBR1
AF (sqrt-t) Sorted Populations

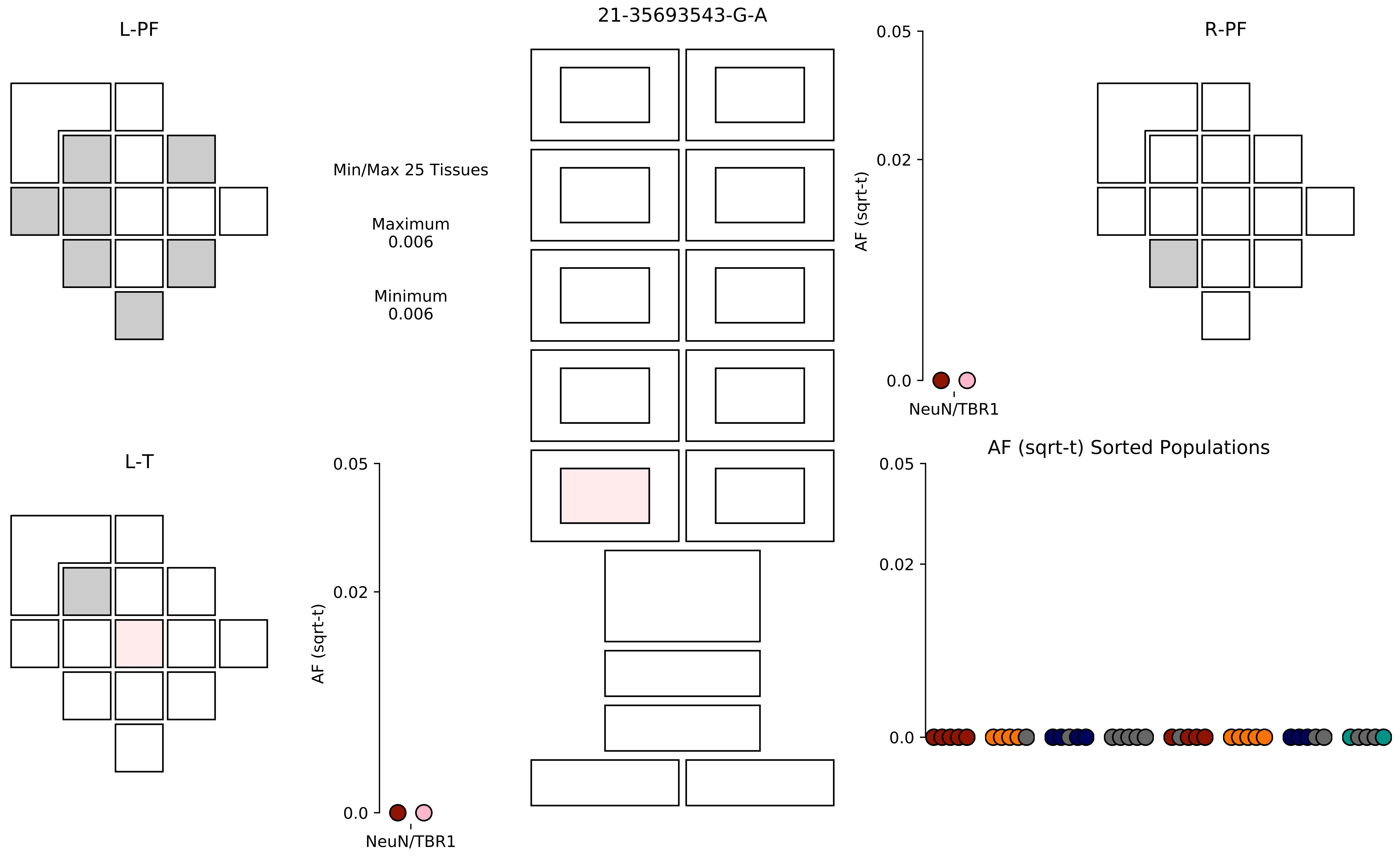

21-35693543-G-A
L-PF
R-PF
L-T
Min/Max 25 Tissues
Maximum
0.006
Minimum
0.006
AF (sqrt-t)
0.05
0.02
0.0
NeuN/TBR1
AF (sqrt-t)
0.05
0.02
0.0
NeuN/TBR1
AF (sqrt-t) Sorted Populations
0.05
0.02
0.0

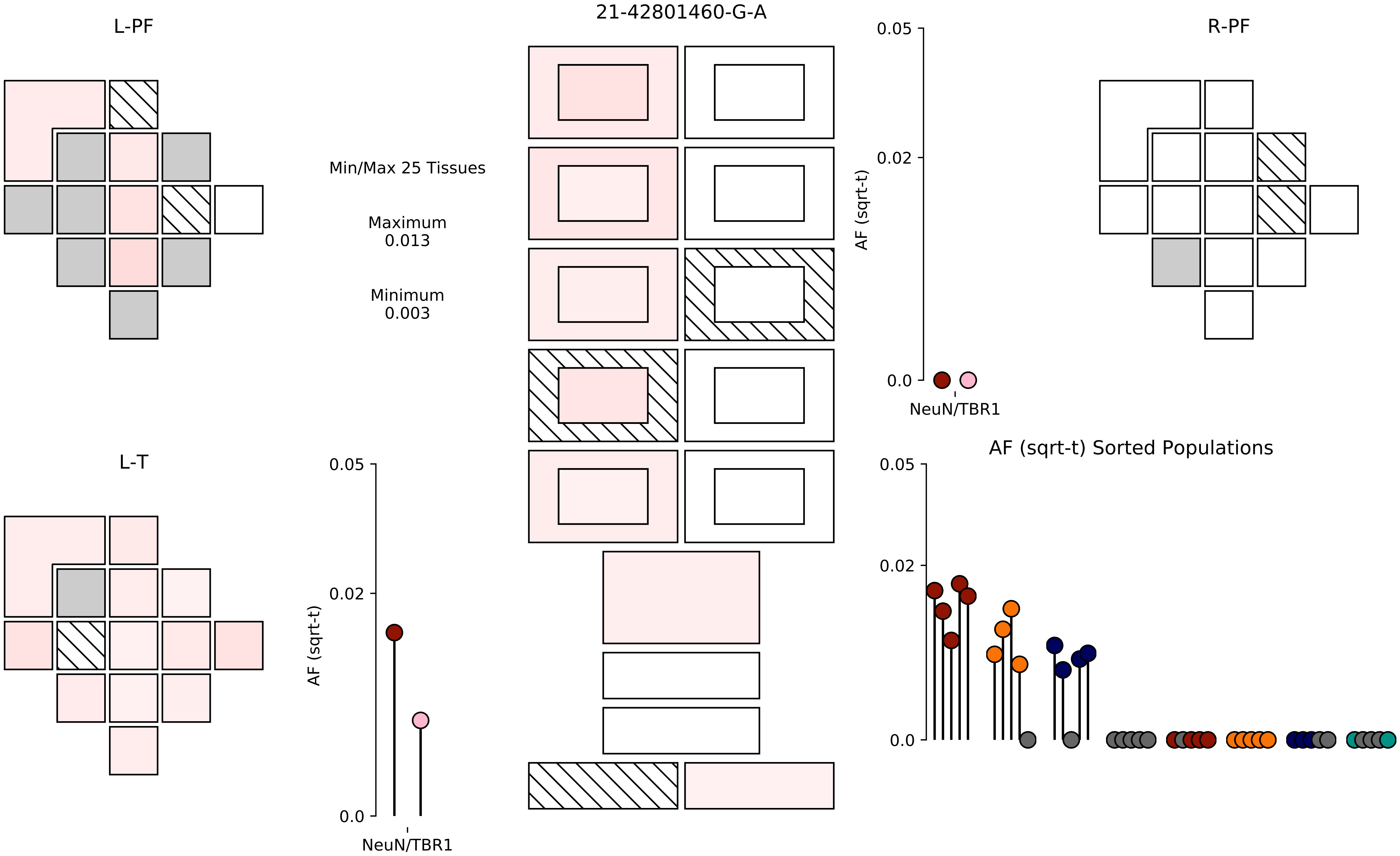
21-42801460-G-A
L-PF
R-PF
L-T
Min/Max 25 Tissues
Maximum
0.013
Minimum
0.003
AF (sqrt-t)
0.05
0.02
0.0
NeuN/TBR1
AF (sqrt-t)
0.05
0.02
0.0
NeuN/TBR1
AF (sqrt-t) Sorted Populations
0.05
0.02
0.0

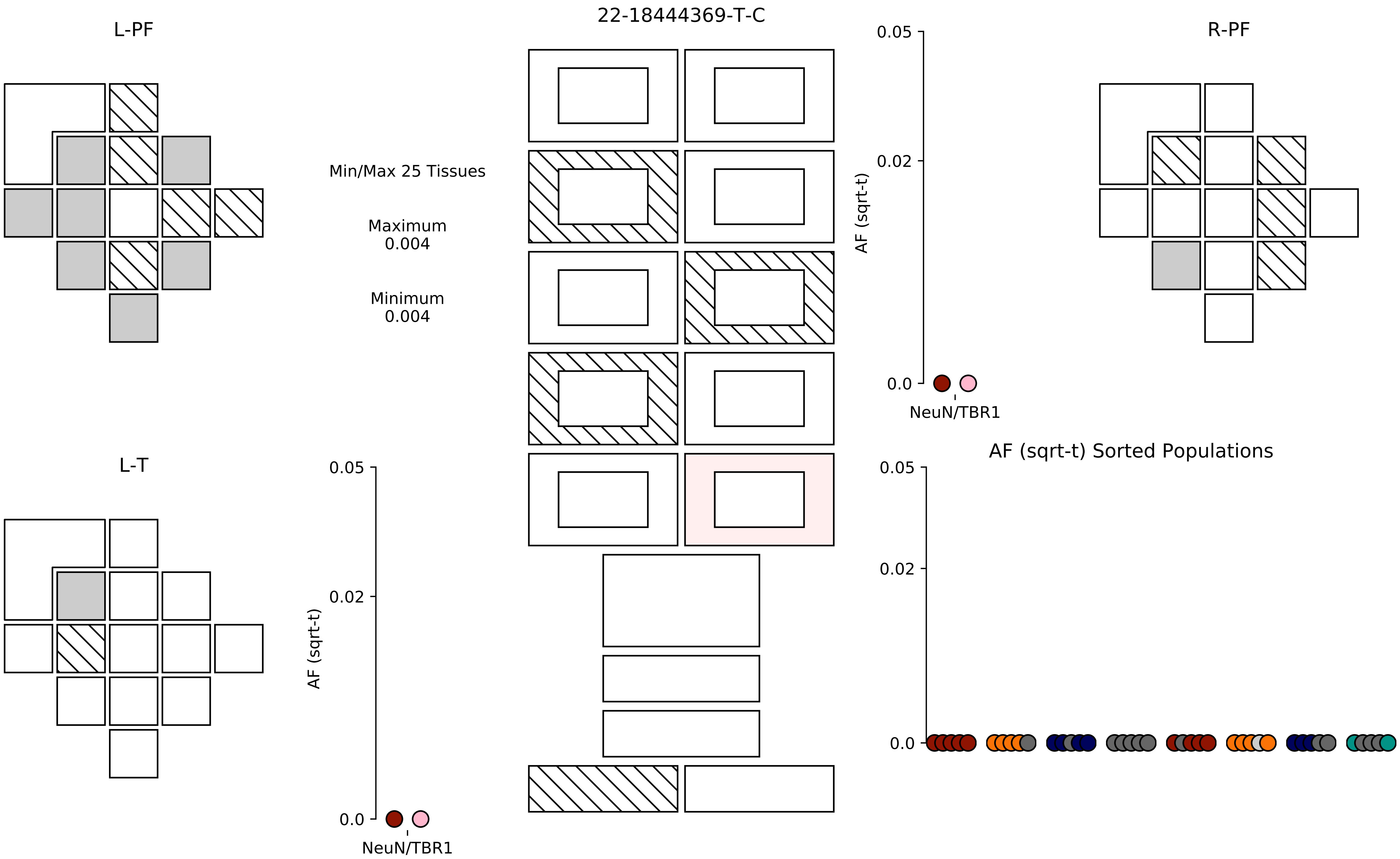

22-18444369-T-C
L-PF
R-PF
L-T
Min/Max 25 Tissues
Maximum
0.004
Minimum
0.004
AF (sqrt-t)
0.05
0.02
0.0
NeuN/TBR1
AF (sqrt-t) Sorted Populations

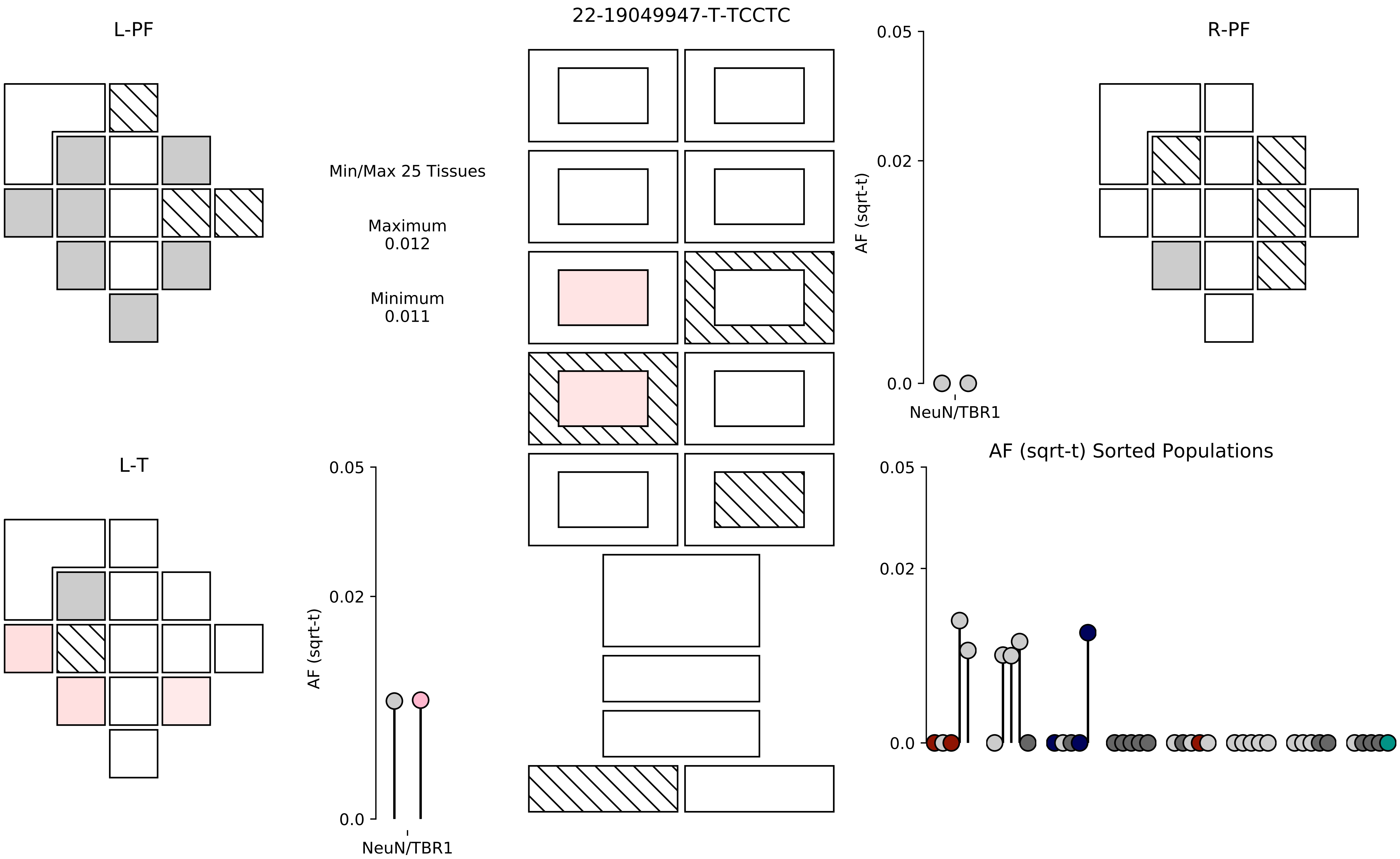

22-19049947-T-TCCTC
L-PF
R-PF
L-T
Min/Max 25 Tissues
Maximum
0.012
Minimum
0.011
AF (sqrt-t)
0.05
0.02
0.0
NeuN/TBR1
AF (sqrt-t) Sorted Populations

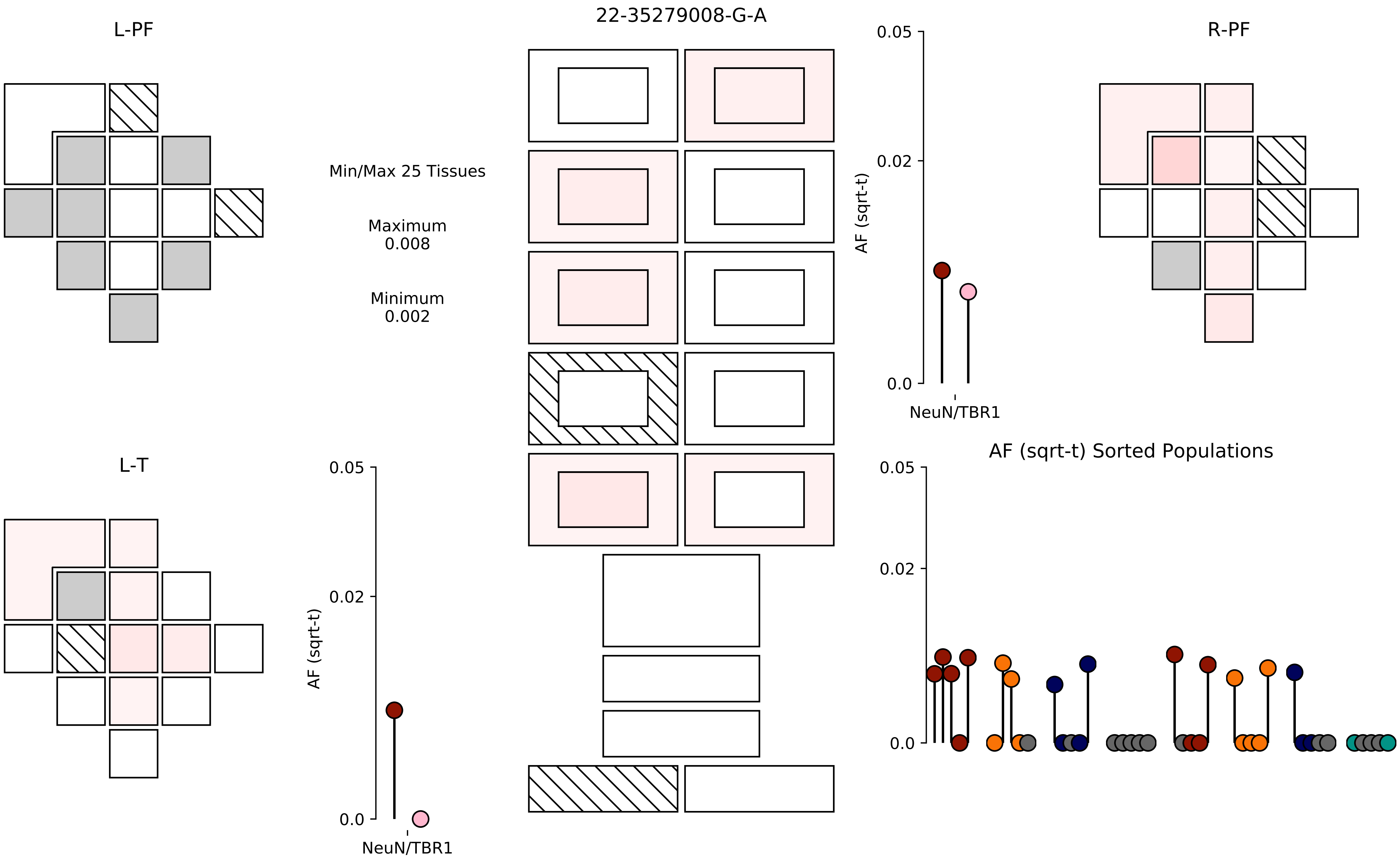
22-35279008-G-A
L-PF
R-PF
L-T
Min/Max 25 Tissues
Maximum
0.008
Minimum
0.002
AF (sqrt-t)
0.05
0.02
0.0
NeuN/TBR1
AF (sqrt-t) Sorted Populations

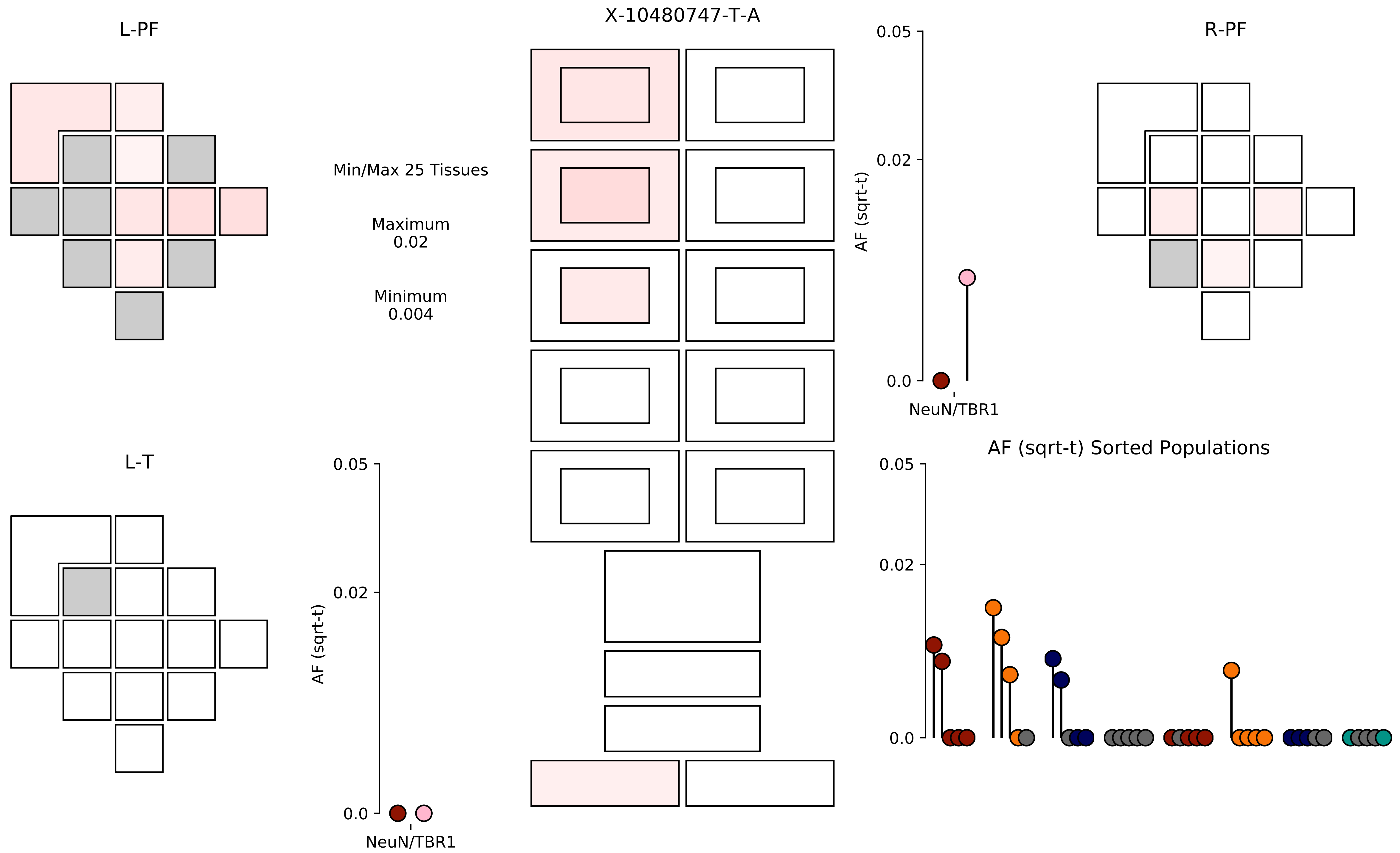
L-PF
X-10480747-T-A
R-PF
Min/Max 25 Tissues
Maximum
0.02
Minimum
0.004
L-T
AF (sqrt-t)
0.05
0.02
0.0
NeuN/TBR1
AF (sqrt-t) Sorted Populations

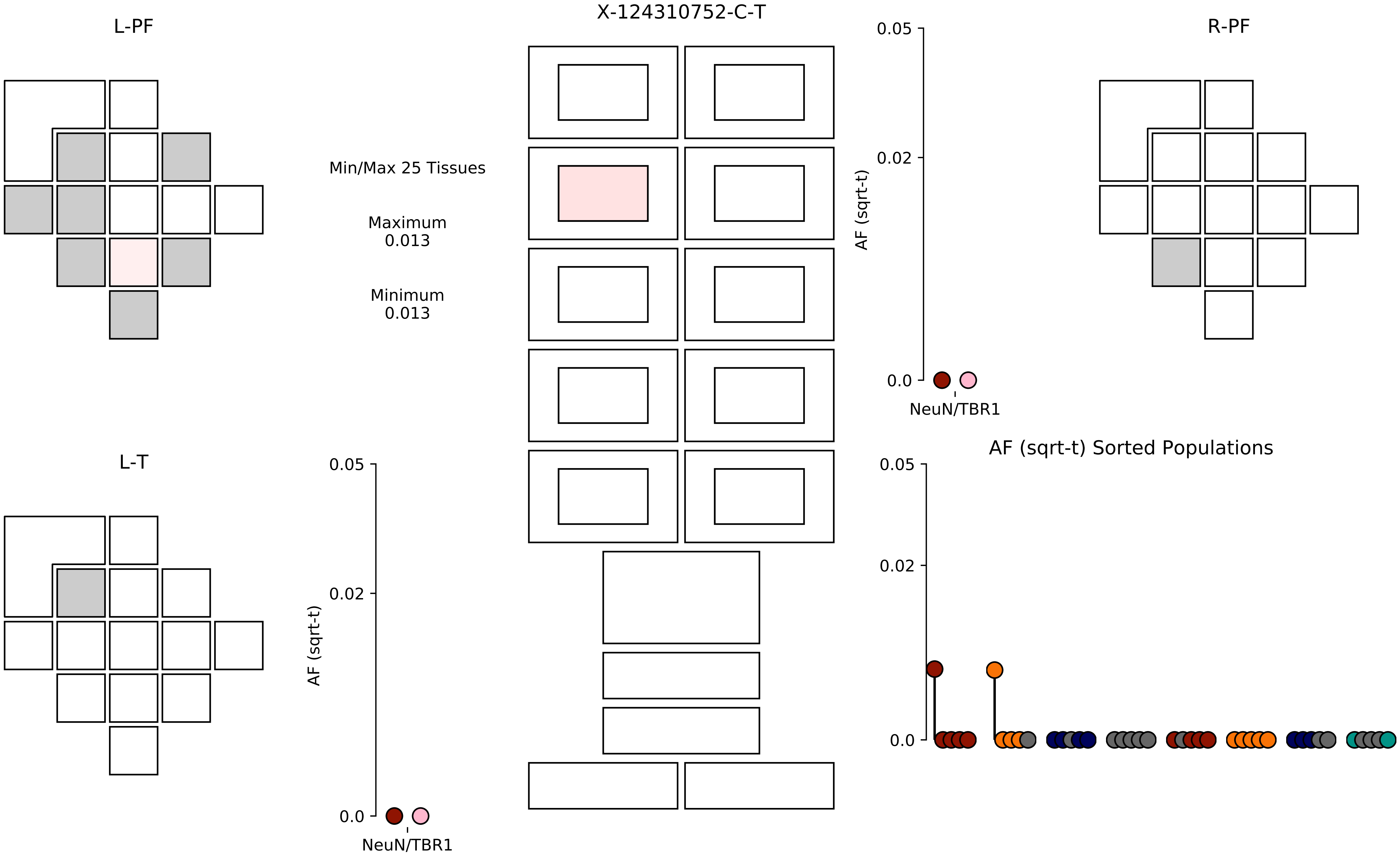
X-124310752-C-T
L-PF
R-PF
L-T
Min/Max 25 Tissues
Maximum
0.013
Minimum
0.013
AF (sqrt-t)
0.05
0.02
0.0
NeuN/TBR1
AF (sqrt-t) Sorted Populations

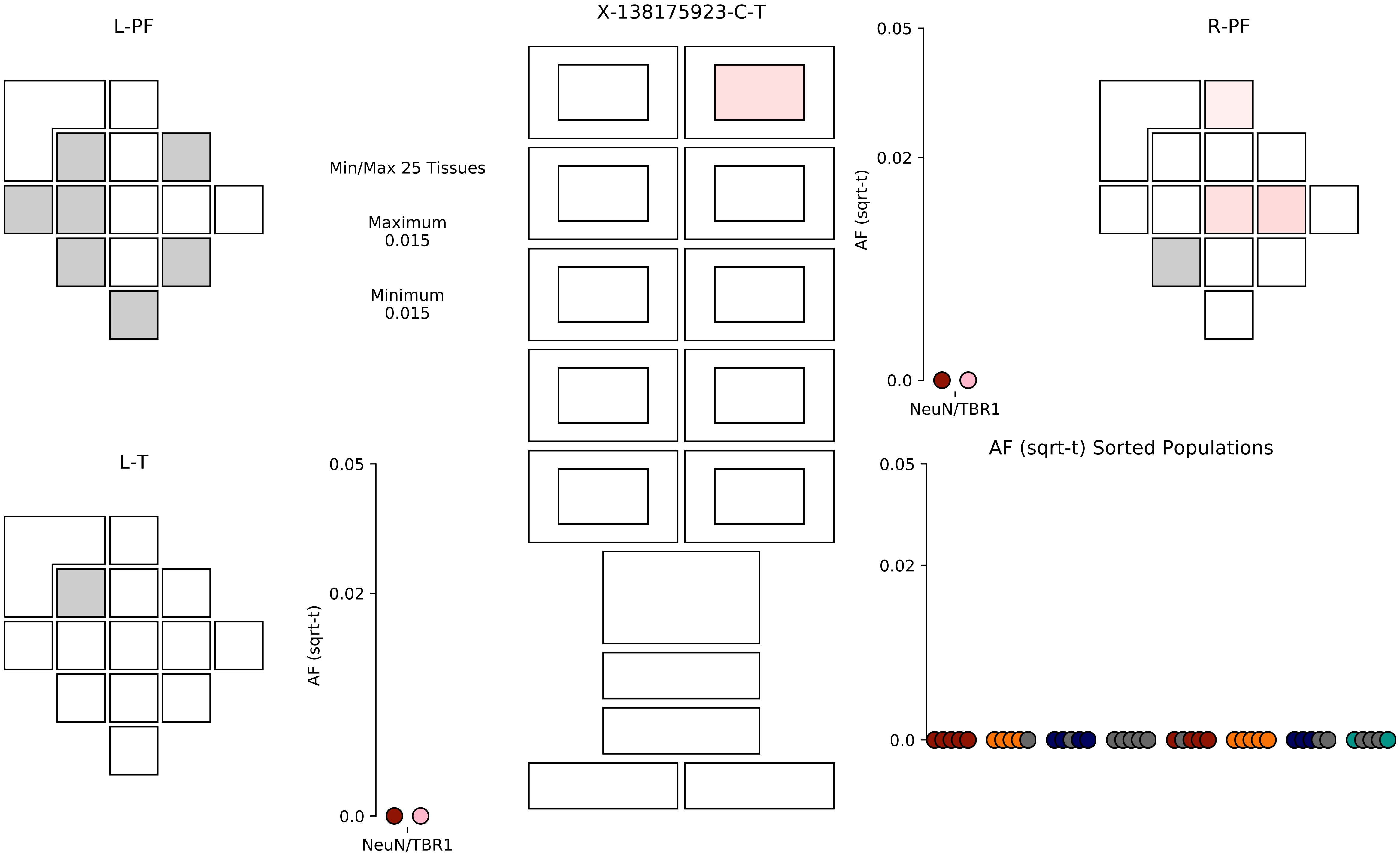

X-138175923-C-T
L-PF
R-PF
L-T
Min/Max 25 Tissues
Maximum
0.015
Minimum
0.015
0.05
0.02
0.0
AF (sqrt-t)
NeuN/TBR1
AF (sqrt-t) Sorted Populations

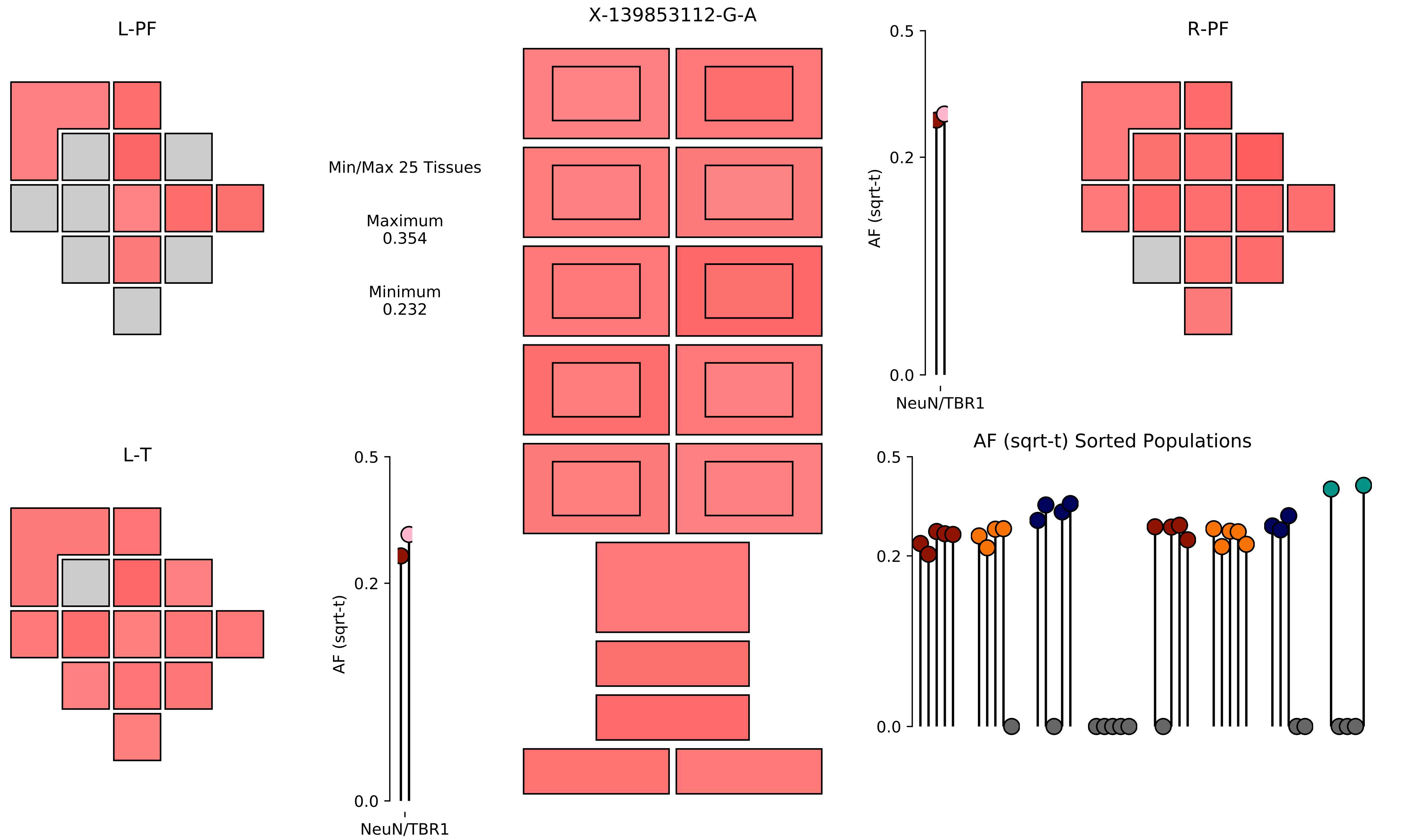

X-139853112-G-A
L-PF
R-PF
L-T
Min/Max 25 Tissues
Maximum
0.354
Minimum
0.232
AF (sqrt-t)
NeuN/TBR1
0.5
0.2
0.0
AF (sqrt-t) Sorted Populations

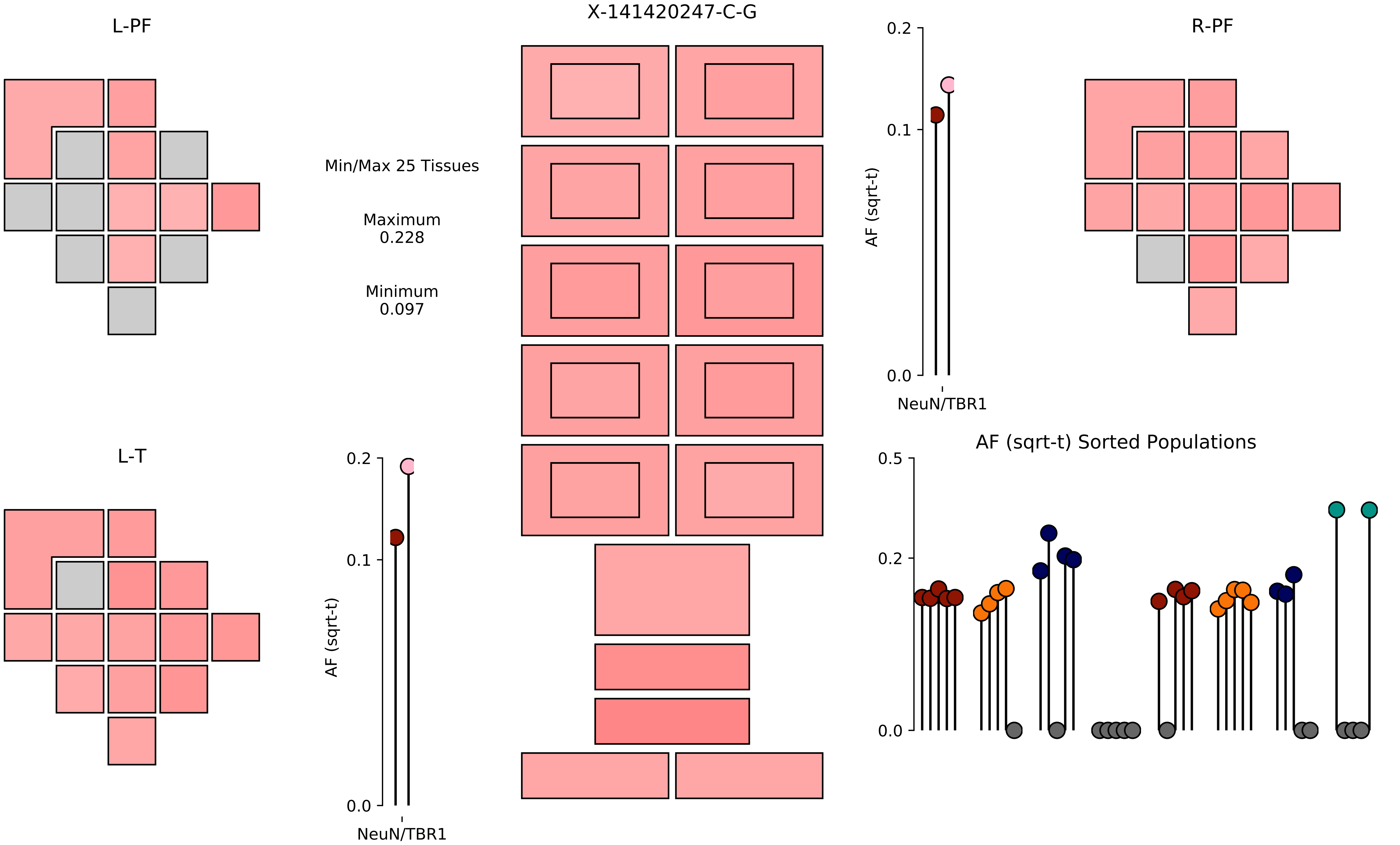
X-141420247-C-G
L-PF
R-PF
L-T
Min/Max 25 Tissues
Maximum
0.228
Minimum
0.097
AF (sqrt-t)
NeuN/TBR1
0.0
0.1
0.2
AF (sqrt-t) Sorted Populations
0.5

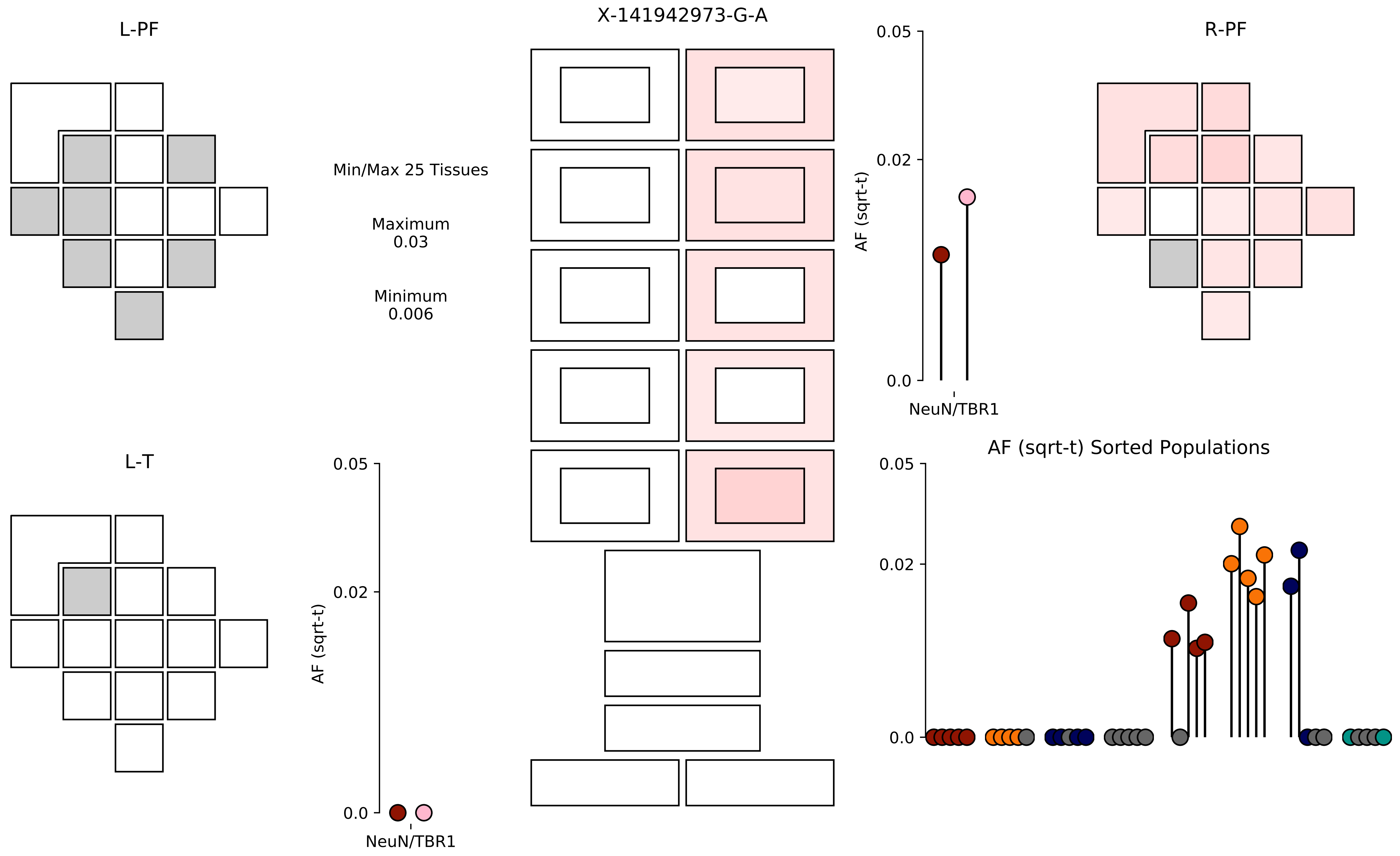

X-141942973-G-A
L-PF
R-PF
L-T
Min/Max 25 Tissues
Maximum
0.03
Minimum
0.006
AF (sqrt-t)
0.05
0.02
0.0
NeuN/TBR1
AF (sqrt-t) Sorted Populations

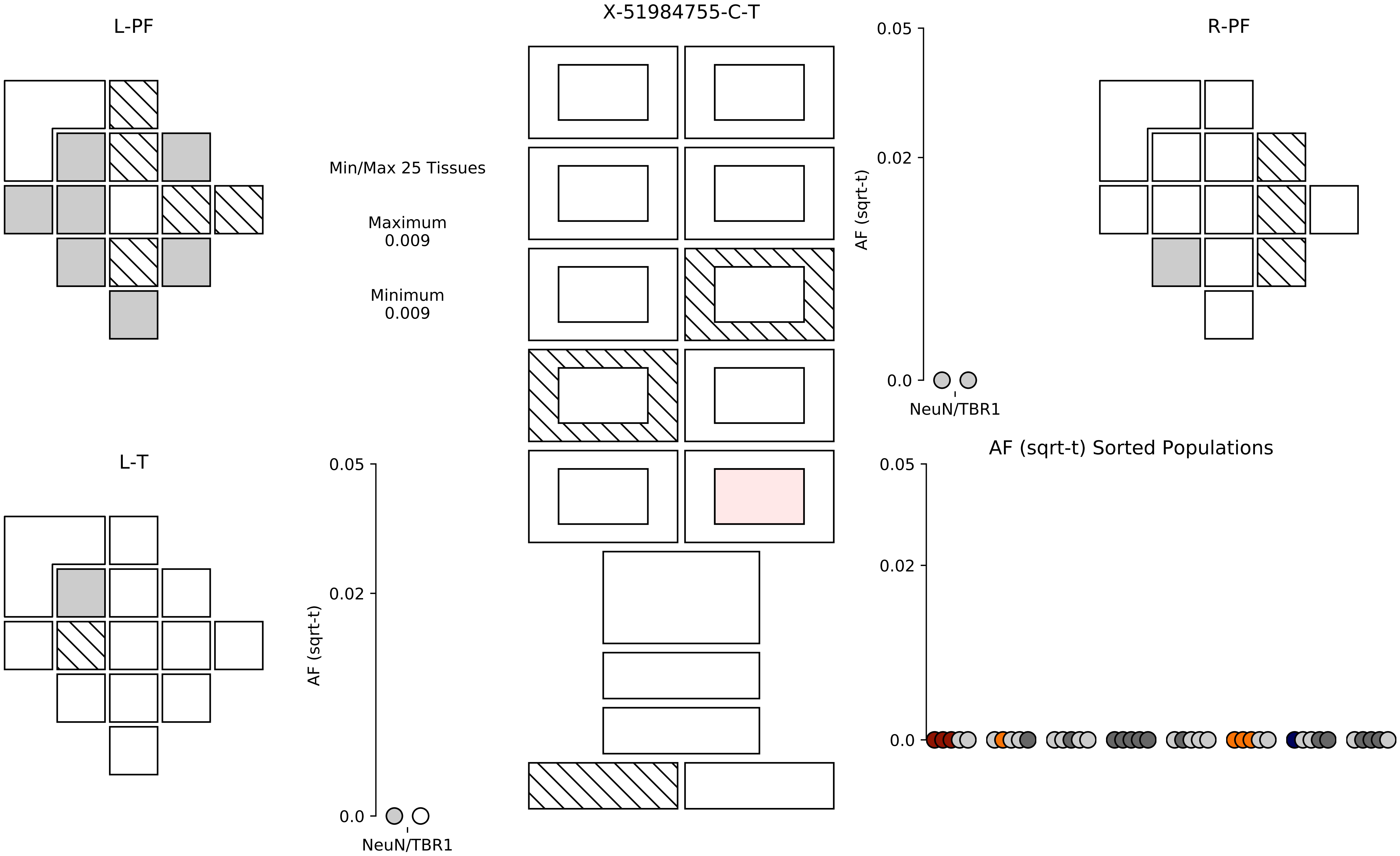

L-PF
X-51984755-C-T
R-PF
Min/Max 25 Tissues
Maximum
0.009
Minimum
0.009
0.05
0.02
0.0
AF (sqrt-t)
NeuN/TBR1
L-T
0.05
0.02
0.0
AF (sqrt-t)
NeuN/TBR1
AF (sqrt-t) Sorted Populations
0.05
0.02
0.0

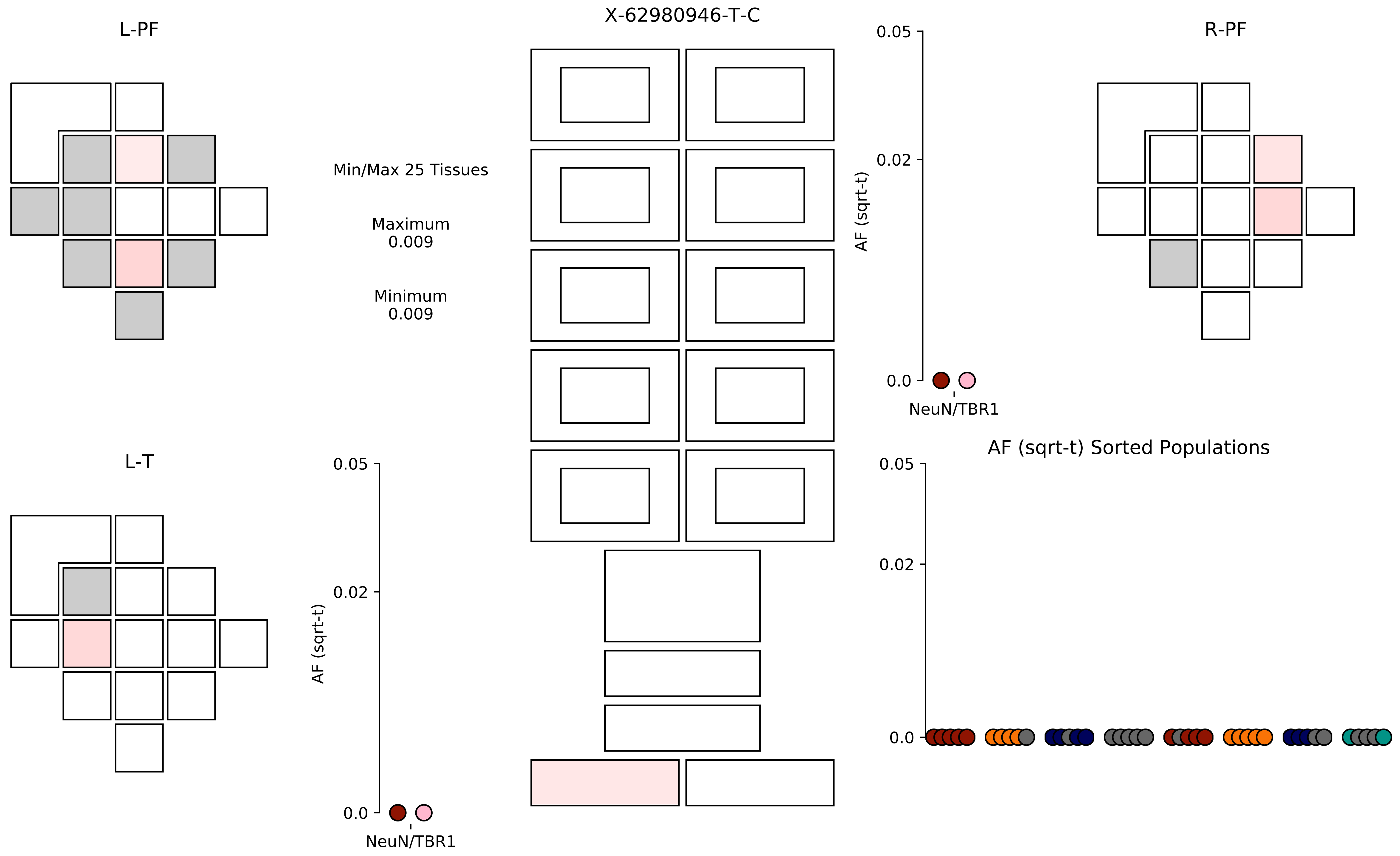

X-62980946-T-C
L-PF
R-PF
L-T
Min/Max 25 Tissues
Maximum
0.009
Minimum
0.009
AF (sqrt-t)
0.05
0.02
0.0
NeuN/TBR1
AF (sqrt-t) Sorted Populations

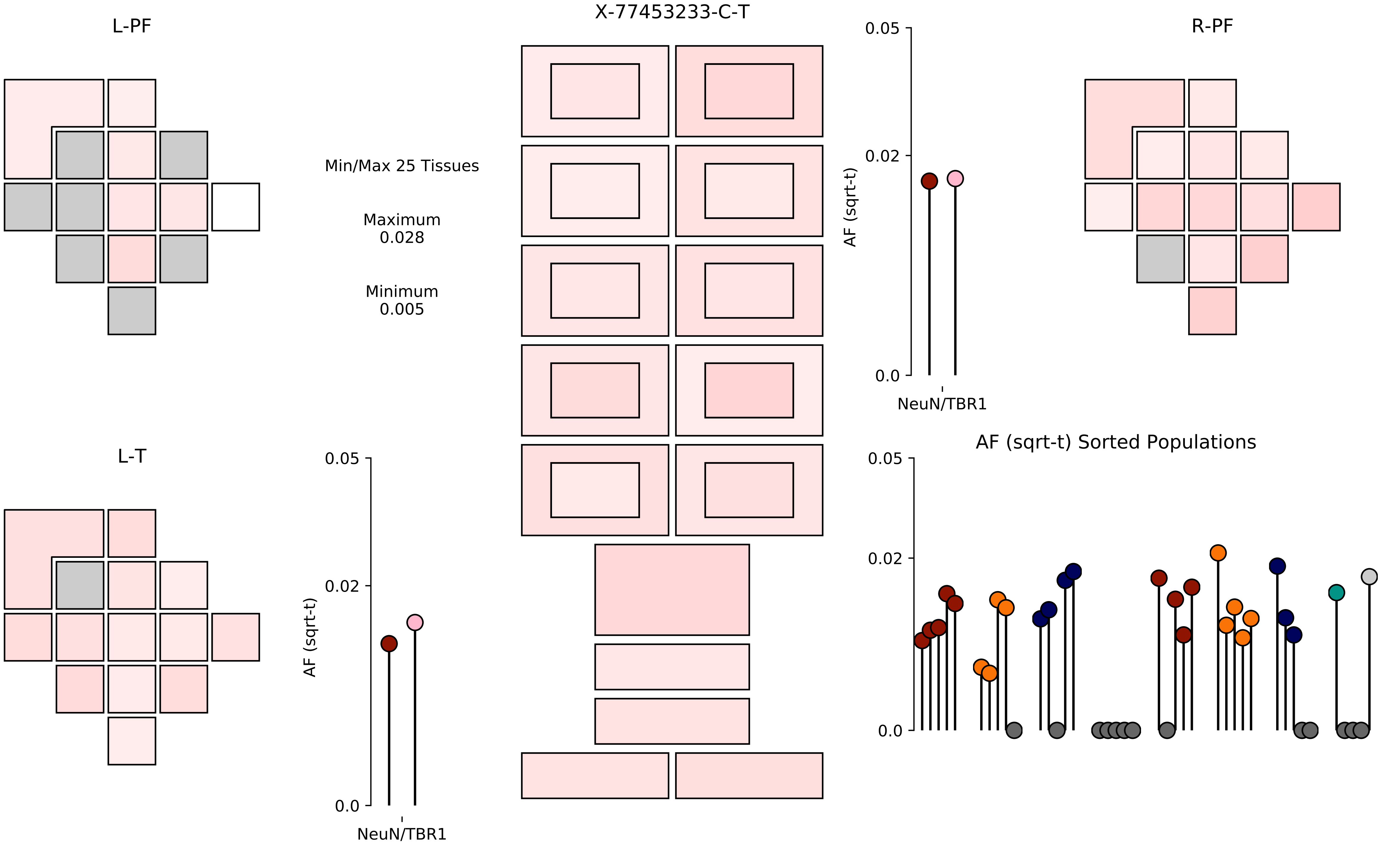

X-77453233-C-T
L-PF
R-PF
L-T
Min/Max 25 Tissues
Maximum
0.028
Minimum
0.005
AF (sqrt-t)
0.05
0.02
0.0
NeuN/TBR1
AF (sqrt-t) Sorted Populations
