## Supplementary material for "Somatic mosaicism in the mature brain reveals clonal cellular distributions during cortical development": Data S4

1−106934533−C−T

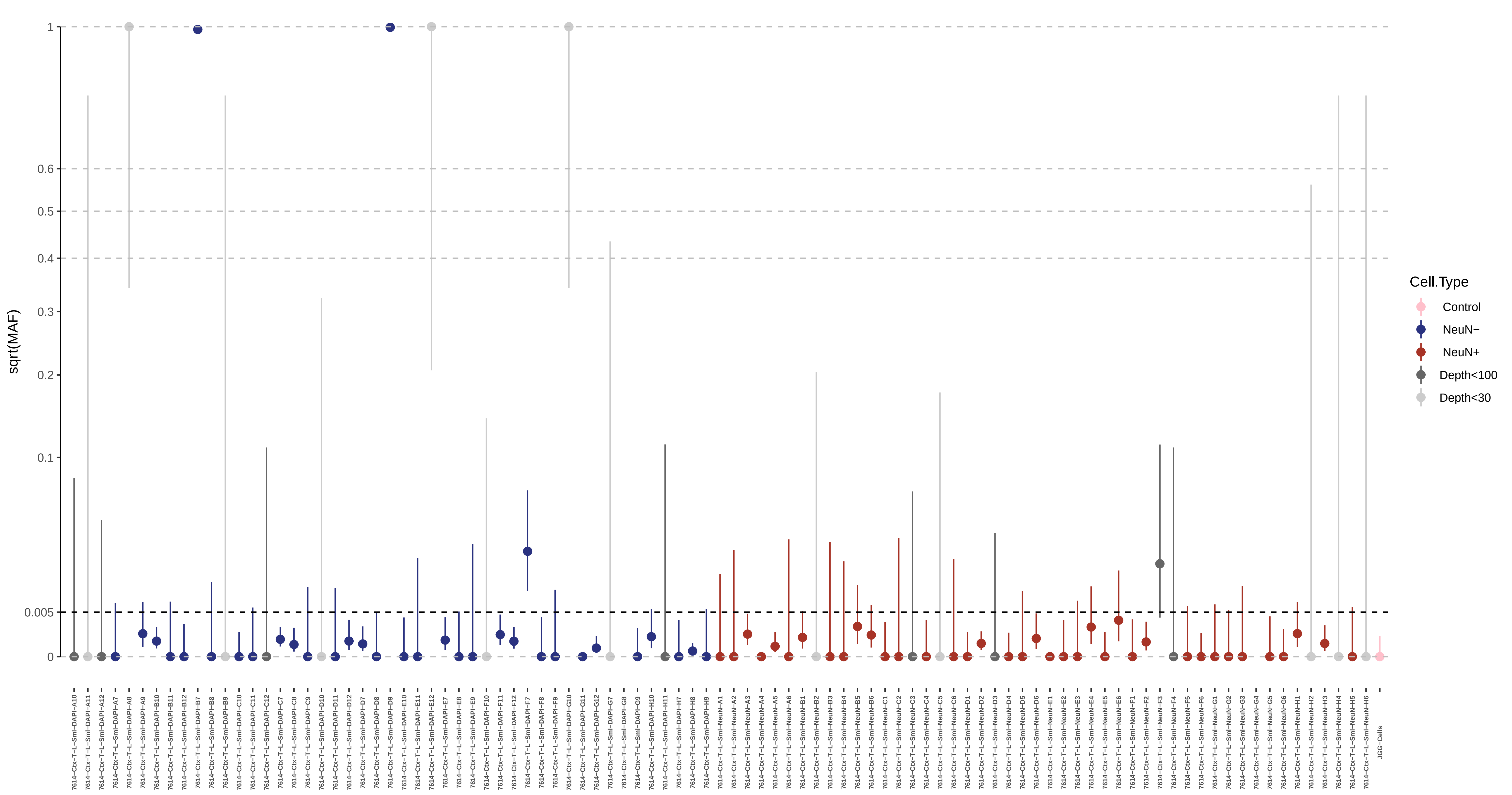

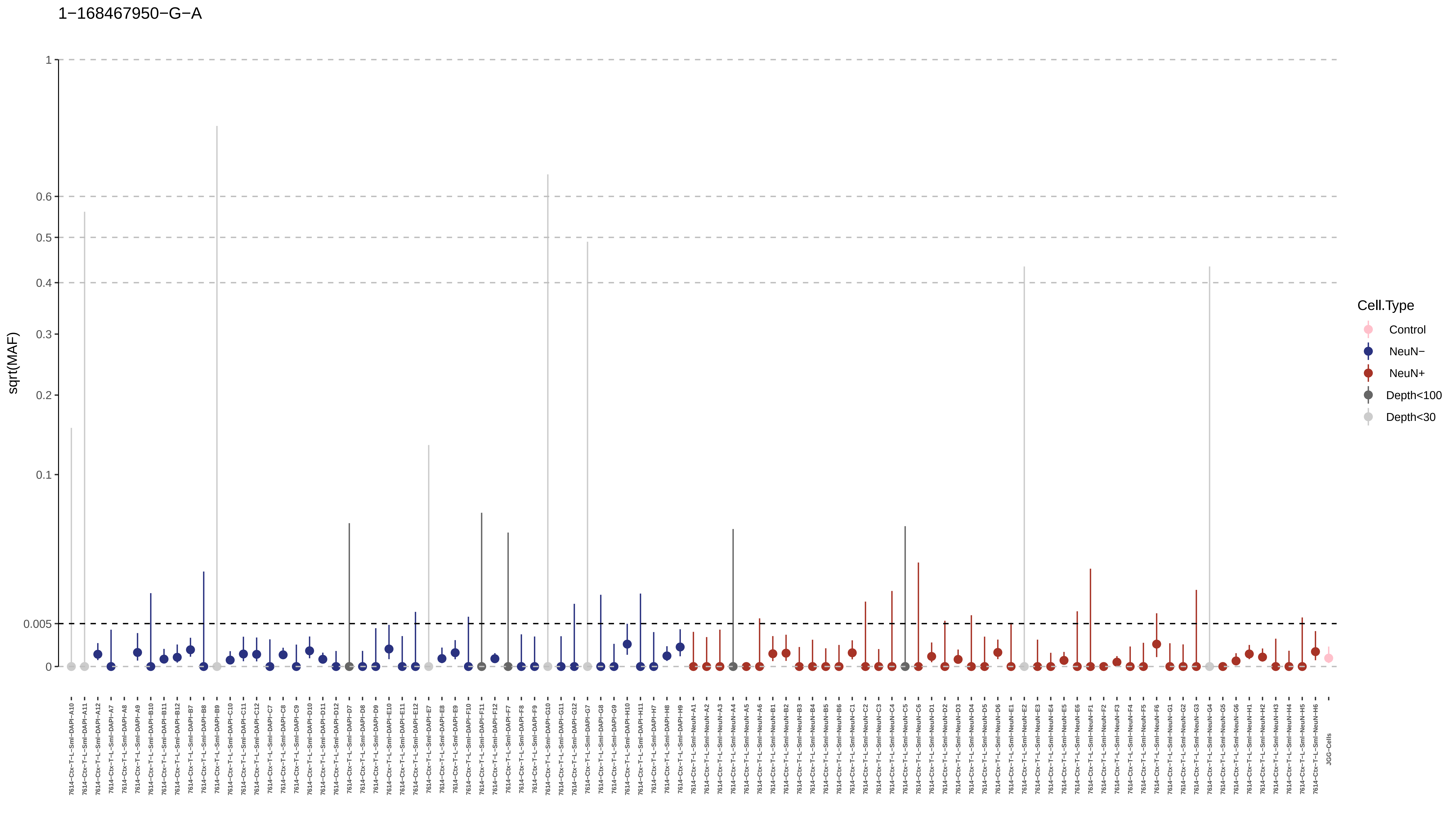
1−168467950−G−A
sqrt(MAF)
1
0.6
0.5
0.4
0.3
0.2
0.1
0.005
0
Cell.Type
Control
NeuN−
NeuN+
Depth<100
Depth<30
7614−Ctx−T−L−Sml−DAPI−A10
7614−Ctx−T−L−Sml−DAPI−A11
7614−Ctx−T−L−Sml−DAPI−A12
7614−Ctx−T−L−Sml−DAPI−A7
7614−Ctx−T−L−Sml−DAPI−A8
7614−Ctx−T−L−Sml−DAPI−A9
7614−Ctx−T−L−Sml−DAPI−B10
7614−Ctx−T−L−Sml−DAPI−B11
7614−Ctx−T−L−Sml−DAPI−B12
7614−Ctx−T−L−Sml−DAPI−B7
7614−Ctx−T−L−Sml−DAPI−B8
7614−Ctx−T−L−Sml−DAPI−B9
7614−Ctx−T−L−Sml−DAPI−C10
7614−Ctx−T−L−Sml−DAPI−C11
7614−Ctx−T−L−Sml−DAPI−C12
7614−Ctx−T−L−Sml−DAPI−C7
7614−Ctx−T−L−Sml−DAPI−C8
7614−Ctx−T−L−Sml−DAPI−C9
7614−Ctx−T−L−Sml−DAPI−D10
7614−Ctx−T−L−Sml−DAPI−D11
7614−Ctx−T−L−Sml−DAPI−D12
7614−Ctx−T−L−Sml−DAPI−D7
7614−Ctx−T−L−Sml−DAPI−D8
7614−Ctx−T−L−Sml−DAPI−D9
7614−Ctx−T−L−Sml−DAPI−E10
7614−Ctx−T−L−Sml−DAPI−E11
7614−Ctx−T−L−Sml−DAPI−E12
7614−Ctx−T−L−Sml−DAPI−E7
7614−Ctx−T−L−Sml−DAPI−E8
7614−Ctx−T−L−Sml−DAPI−E9
7614−Ctx−T−L−Sml−DAPI−F10
7614−Ctx−T−L−Sml−DAPI−F11
7614−Ctx−T−L−Sml−DAPI−F12
7614−Ctx−T−L−Sml−DAPI−F7
7614−Ctx−T−L−Sml−DAPI−F8
7614−Ctx−T−L−Sml−DAPI−F9
7614−Ctx−T−L−Sml−DAPI−G10
7614−Ctx−T−L−Sml−DAPI−G11
7614−Ctx−T−L−Sml−DAPI−G12
7614−Ctx−T−L−Sml−DAPI−G7
7614−Ctx−T−L−Sml−DAPI−G8
7614−Ctx−T−L−Sml−DAPI−G9
7614−Ctx−T−L−Sml−DAPI−H10
7614−Ctx−T−L−Sml−DAPI−H11
7614−Ctx−T−L−Sml−DAPI−H7
7614−Ctx−T−L−Sml−DAPI−H8
7614−Ctx−T−L−Sml−DAPI−H9
7614−Ctx−T−L−Sml−NeuN−A1
7614−Ctx−T−L−Sml−NeuN−A2
7614−Ctx−T−L−Sml−NeuN−A3
7614−Ctx−T−L−Sml−NeuN−A4
7614−Ctx−T−L−Sml−NeuN−A5
7614−Ctx−T−L−Sml−NeuN−A6
7614−Ctx−T−L−Sml−NeuN−B1
7614−Ctx−T−L−Sml−NeuN−B2
7614−Ctx−T−L−Sml−NeuN−B3
7614−Ctx−T−L−Sml−NeuN−B4
7614−Ctx−T−L−Sml−NeuN−B5
7614−Ctx−T−L−Sml−NeuN−B6
7614−Ctx−T−L−Sml−NeuN−C1
7614−Ctx−T−L−Sml−NeuN−C2
7614−Ctx−T−L−Sml−NeuN−C3
7614−Ctx−T−L−Sml−NeuN−C4
7614−Ctx−T−L−Sml−NeuN−C5
7614−Ctx−T−L−Sml−NeuN−C6
7614−Ctx−T−L−Sml−NeuN−D1
7614−Ctx−T−L−Sml−NeuN−D2
7614−Ctx−T−L−Sml−NeuN−D3
7614−Ctx−T−L−Sml−NeuN−D4
7614−Ctx−T−L−Sml−NeuN−D5
7614−Ctx−T−L−Sml−NeuN−D6
7614−Ctx−T−L−Sml−NeuN−E1
7614−Ctx−T−L−Sml−NeuN−E2
7614−Ctx−T−L−Sml−NeuN−E3
7614−Ctx−T−L−Sml−NeuN−E4
7614−Ctx−T−L−Sml−NeuN−E5
7614−Ctx−T−L−Sml−NeuN−E6
7614−Ctx−T−L−Sml−NeuN−F1
7614−Ctx−T−L−Sml−NeuN−F2
7614−Ctx−T−L−Sml−NeuN−F3
7614−Ctx−T−L−Sml−NeuN−F4
7614−Ctx−T−L−Sml−NeuN−F5
7614−Ctx−T−L−Sml−NeuN−F6
7614−Ctx−T−L−Sml−NeuN−G1
7614−Ctx−T−L−Sml−NeuN−G2
7614−Ctx−T−L−Sml−NeuN−G3
7614−Ctx−T−L−Sml−NeuN−G4
7614−Ctx−T−L−Sml−NeuN−G5
7614−Ctx−T−L−Sml−NeuN−G6
7614−Ctx−T−L−Sml−NeuN−H1
7614−Ctx−T−L−Sml−NeuN−H2
7614−Ctx−T−L−Sml−NeuN−H3
7614−Ctx−T−L−Sml−NeuN−H4
7614−Ctx−T−L−Sml−NeuN−H5
7614−Ctx−T−L−Sml−NeuN−H6
JGG-Cells

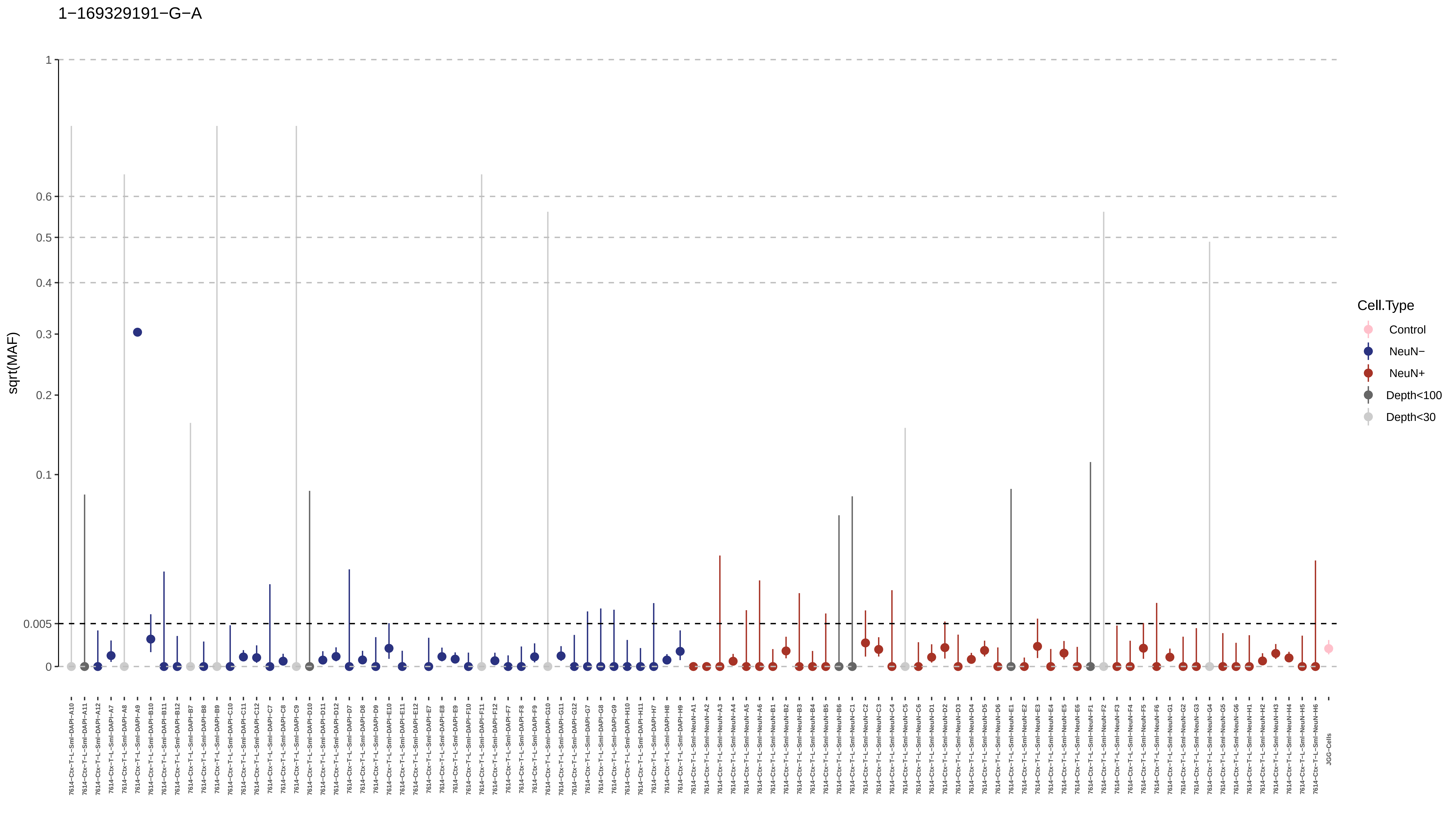
1−169329191−G−A
sqrt(MAF)
1
0.6
0.5
0.4
0.3
0.2
0.1
0.005
0
Cell.Type
Control
NeuN−
NeuN+
Depth<100
Depth<30
7614−Ctx−T−L−Sml−DAPI−A10
7614−Ctx−T−L−Sml−DAPI−A11
7614−Ctx−T−L−Sml−DAPI−A12
7614−Ctx−T−L−Sml−DAPI−A7
7614−Ctx−T−L−Sml−DAPI−A8
7614−Ctx−T−L−Sml−DAPI−A9
7614−Ctx−T−L−Sml−DAPI−B10
7614−Ctx−T−L−Sml−DAPI−B11
7614−Ctx−T−L−Sml−DAPI−B12
7614−Ctx−T−L−Sml−DAPI−B7
7614−Ctx−T−L−Sml−DAPI−B8
7614−Ctx−T−L−Sml−DAPI−B9
7614−Ctx−T−L−Sml−DAPI−C10
7614−Ctx−T−L−Sml−DAPI−C11
7614−Ctx−T−L−Sml−DAPI−C12
7614−Ctx−T−L−Sml−DAPI−C7
7614−Ctx−T−L−Sml−DAPI−C8
7614−Ctx−T−L−Sml−DAPI−C9
7614−Ctx−T−L−Sml−DAPI−D10
7614−Ctx−T−L−Sml−DAPI−D11
7614−Ctx−T−L−Sml−DAPI−D12
7614−Ctx−T−L−Sml−DAPI−D7
7614−Ctx−T−L−Sml−DAPI−D8
7614−Ctx−T−L−Sml−DAPI−D9
7614−Ctx−T−L−Sml−DAPI−E10
7614−Ctx−T−L−Sml−DAPI−E11
7614−Ctx−T−L−Sml−DAPI−E12
7614−Ctx−T−L−Sml−DAPI−E7
7614−Ctx−T−L−Sml−DAPI−E8
7614−Ctx−T−L−Sml−DAPI−E9
7614−Ctx−T−L−Sml−DAPI−F10
7614−Ctx−T−L−Sml−DAPI−F11
7614−Ctx−T−L−Sml−DAPI−F12
7614−Ctx−T−L−Sml−DAPI−F7
7614−Ctx−T−L−Sml−DAPI−F8
7614−Ctx−T−L−Sml−DAPI−F9
7614−Ctx−T−L−Sml−DAPI−G10
7614−Ctx−T−L−Sml−DAPI−G11
7614−Ctx−T−L−Sml−DAPI−G12
7614−Ctx−T−L−Sml−DAPI−G7
7614−Ctx−T−L−Sml−DAPI−G8
7614−Ctx−T−L−Sml−DAPI−G9
7614−Ctx−T−L−Sml−DAPI−H10
7614−Ctx−T−L−Sml−DAPI−H11
7614−Ctx−T−L−Sml−DAPI−H7
7614−Ctx−T−L−Sml−DAPI−H8
7614−Ctx−T−L−Sml−DAPI−H9
7614−Ctx−T−L−Sml−NeuN−A1
7614−Ctx−T−L−Sml−NeuN−A2
7614−Ctx−T−L−Sml−NeuN−A3
7614−Ctx−T−L−Sml−NeuN−A4
7614−Ctx−T−L−Sml−NeuN−A5
7614−Ctx−T−L−Sml−NeuN−A6
7614−Ctx−T−L−Sml−NeuN−B1
7614−Ctx−T−L−Sml−NeuN−B2
7614−Ctx−T−L−Sml−NeuN−B3
7614−Ctx−T−L−Sml−NeuN−B4
7614−Ctx−T−L−Sml−NeuN−B5
7614−Ctx−T−L−Sml−NeuN−B6
7614−Ctx−T−L−Sml−NeuN−C1
7614−Ctx−T−L−Sml−NeuN−C2
7614−Ctx−T−L−Sml−NeuN−C3
7614−Ctx−T−L−Sml−NeuN−C4
7614−Ctx−T−L−Sml−NeuN−C5
7614−Ctx−T−L−Sml−NeuN−C6
7614−Ctx−T−L−Sml−NeuN−D1
7614−Ctx−T−L−Sml−NeuN−D2
7614−Ctx−T−L−Sml−NeuN−D3
7614−Ctx−T−L−Sml−NeuN−D4
7614−Ctx−T−L−Sml−NeuN−D5
7614−Ctx−T−L−Sml−NeuN−D6
7614−Ctx−T−L−Sml−NeuN−E1
7614−Ctx−T−L−Sml−NeuN−E2
7614−Ctx−T−L−Sml−NeuN−E3
7614−Ctx−T−L−Sml−NeuN−E4
7614−Ctx−T−L−Sml−NeuN−E5
7614−Ctx−T−L−Sml−NeuN−E6
7614−Ctx−T−L−Sml−NeuN−F1
7614−Ctx−T−L−Sml−NeuN−F2
7614−Ctx−T−L−Sml−NeuN−F3
7614−Ctx−T−L−Sml−NeuN−F4
7614−Ctx−T−L−Sml−NeuN−F5
7614−Ctx−T−L−Sml−NeuN−F6
7614−Ctx−T−L−Sml−NeuN−G1
7614−Ctx−T−L−Sml−NeuN−G2
7614−Ctx−T−L−Sml−NeuN−G3
7614−Ctx−T−L−Sml−NeuN−G4
7614−Ctx−T−L−Sml−NeuN−G5
7614−Ctx−T−L−Sml−NeuN−G6
7614−Ctx−T−L−Sml−NeuN−H1
7614−Ctx−T−L−Sml−NeuN−H2
7614−Ctx−T−L−Sml−NeuN−H3
7614−Ctx−T−L−Sml−NeuN−H4
7614−Ctx−T−L−Sml−NeuN−H5
7614−Ctx−T−L−Sml−NeuN−H6
JGG-Cells

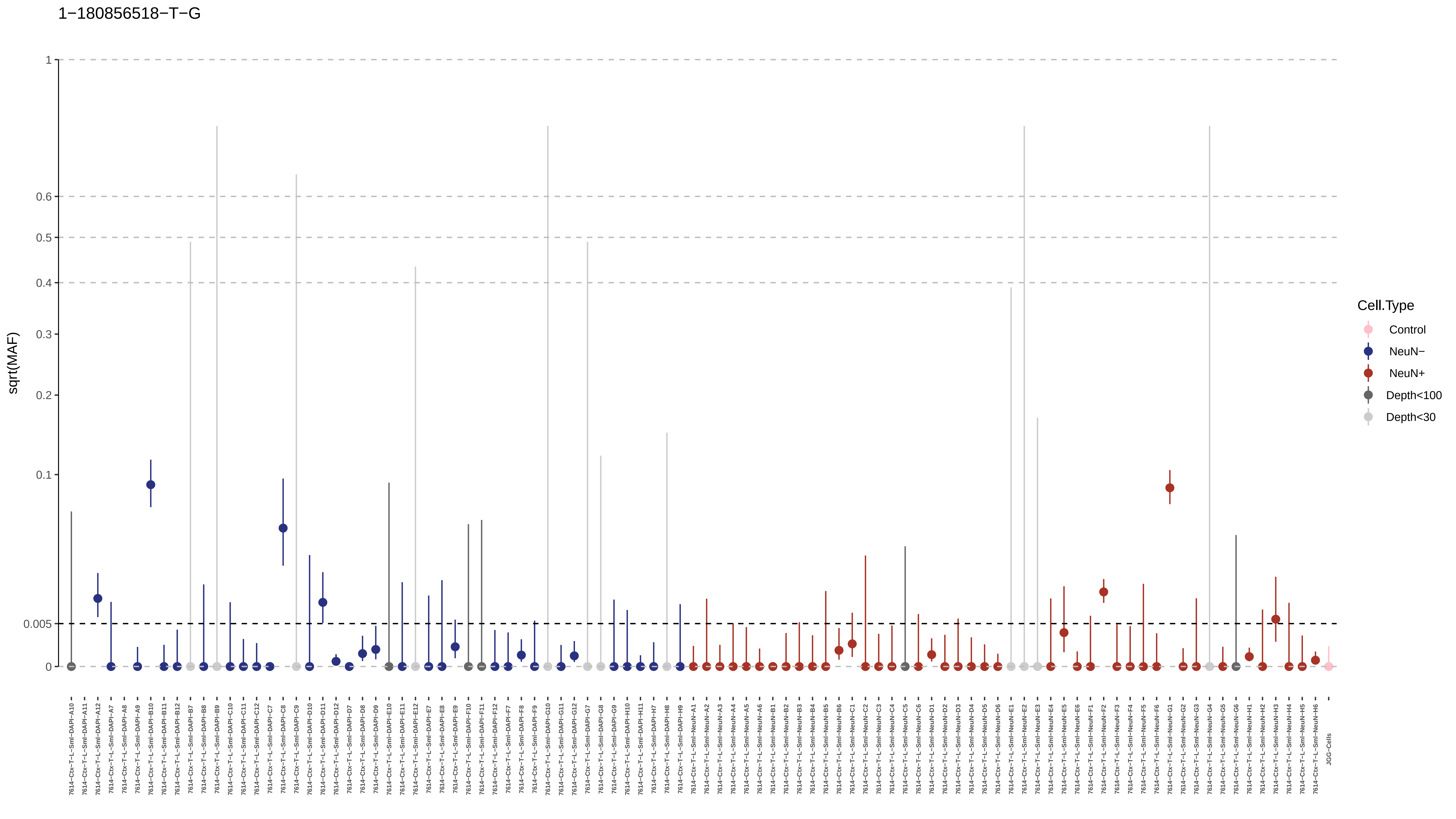
1−180856518−T−G
sqrt(MAF)
1
0.6
0.5
0.4
0.3
0.2
0.1
0.005
0
Cell.Type
Control
NeuN−
NeuN+
Depth<100
Depth<30

1−188748555−G−A

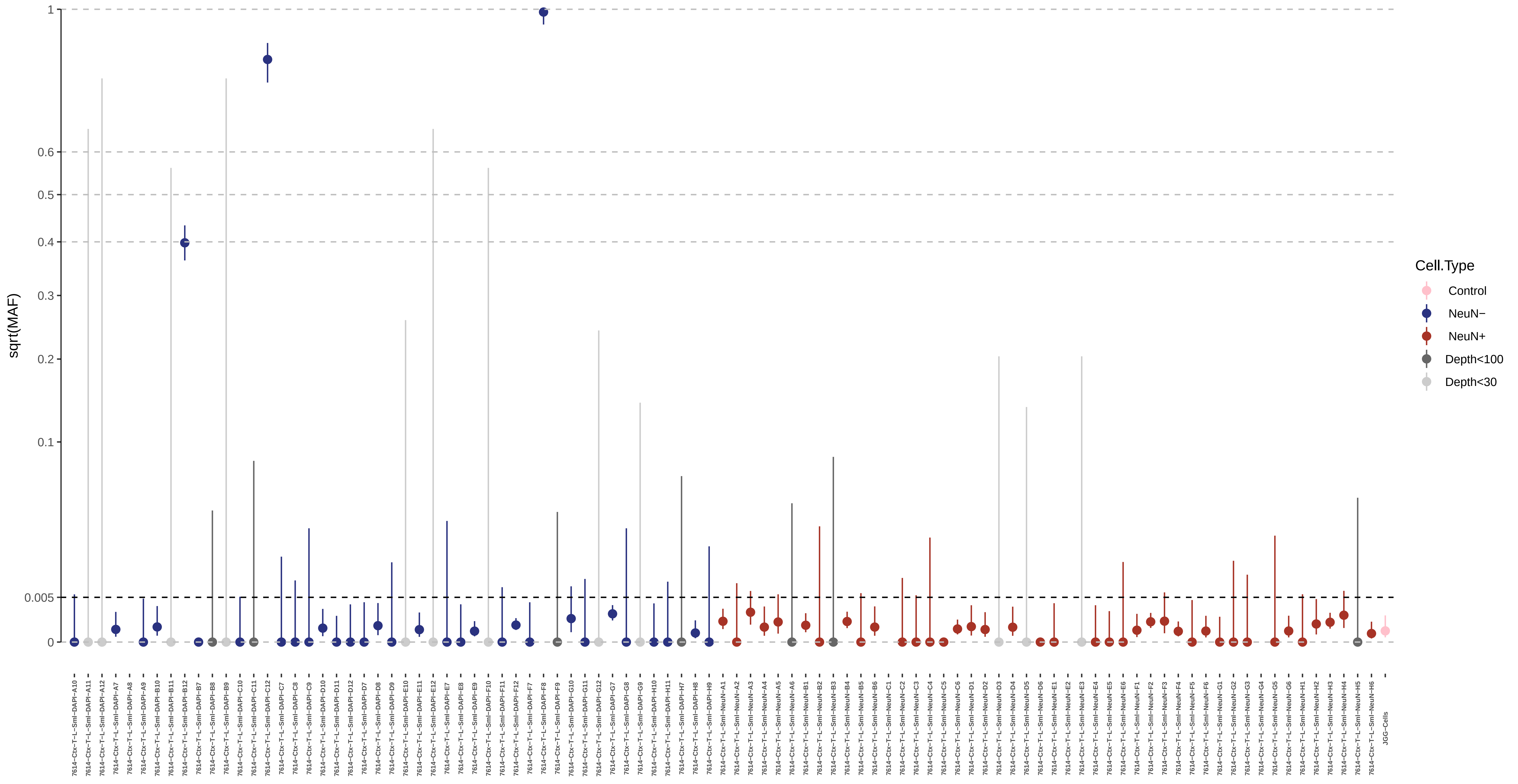

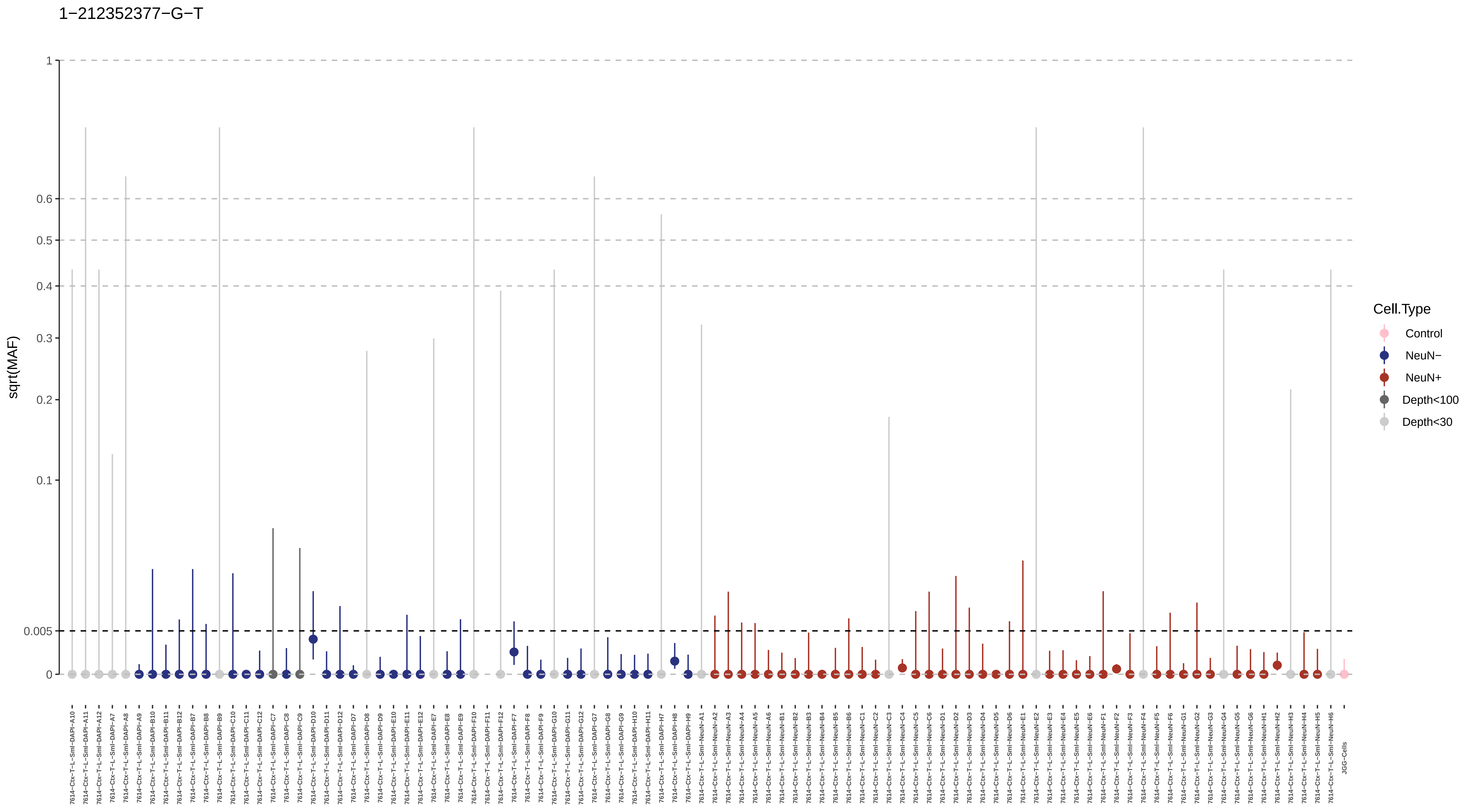
1−212352377−G−T
sqrt(MAF)
1
0.6
0.5
0.4
0.3
0.2
0.1
0.005
0
Cell.Type
Control
NeuN−
NeuN+
Depth<100
Depth<30
JGG-Cells

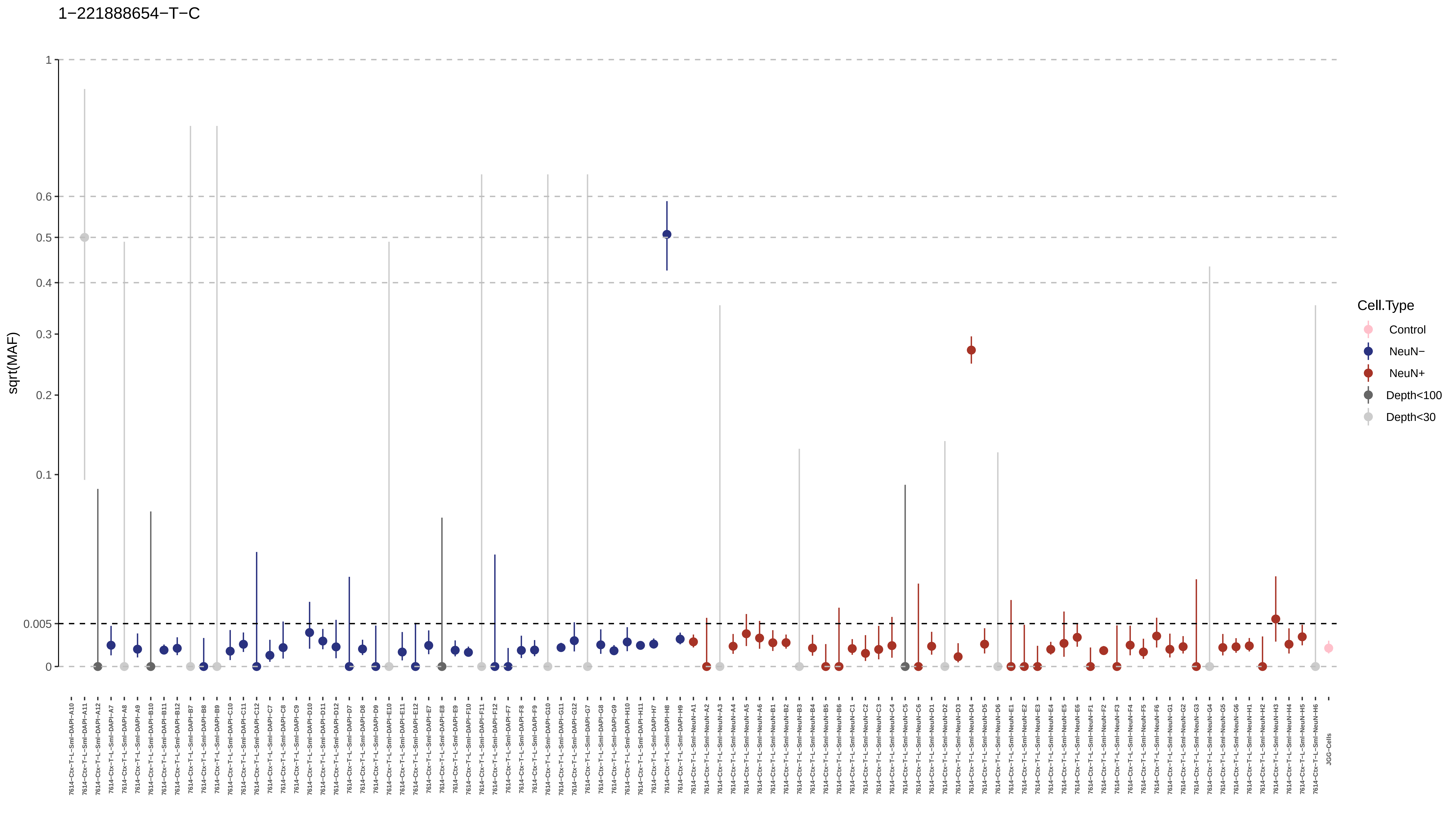
1−221888654−T−C
sqrt(MAF)
1
0.6
0.5
0.4
0.3
0.2
0.1
0.005
0
Cell.Type
Control
NeuN−
NeuN+
Depth<100
Depth<30
JGG-Cells

## 1−239397797−T−A

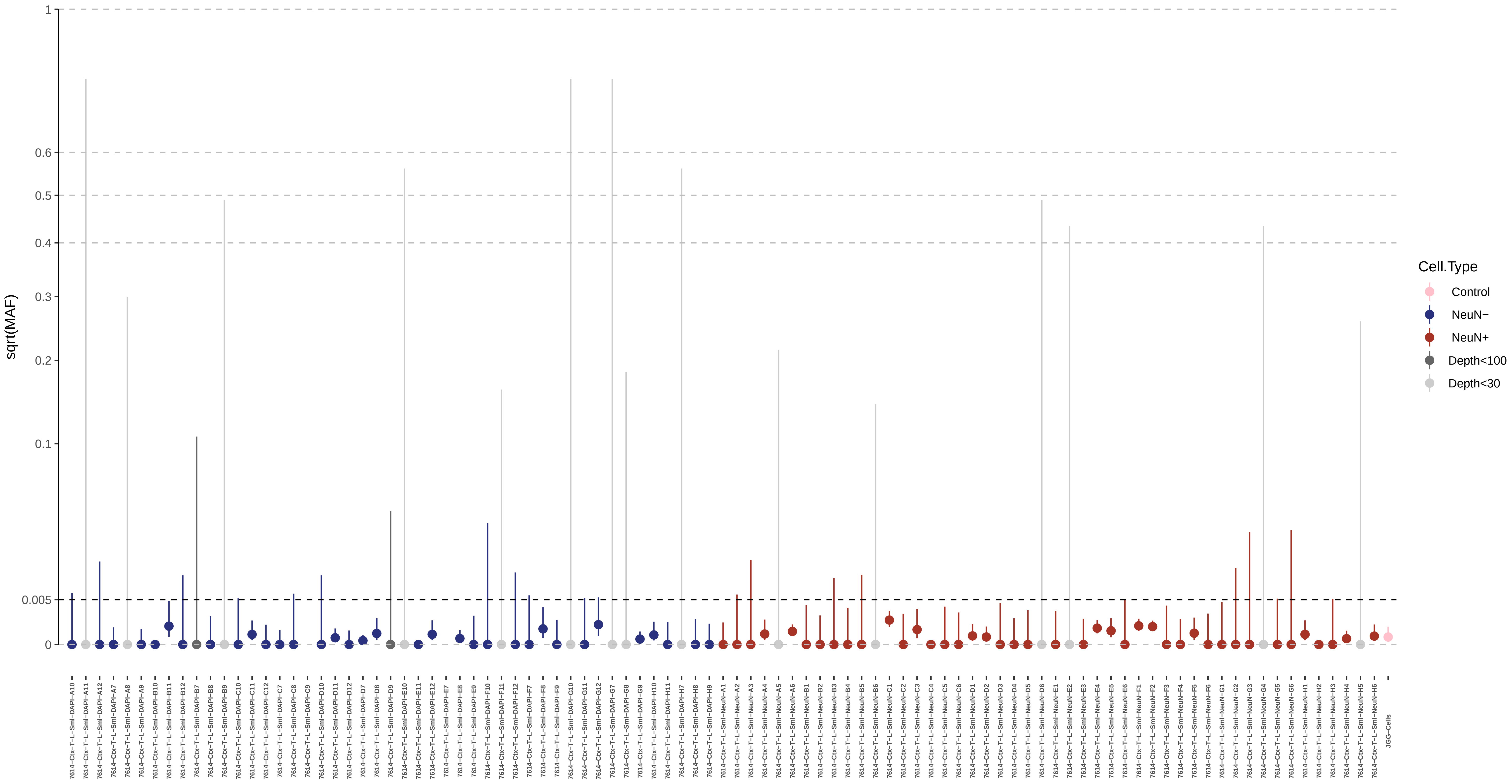

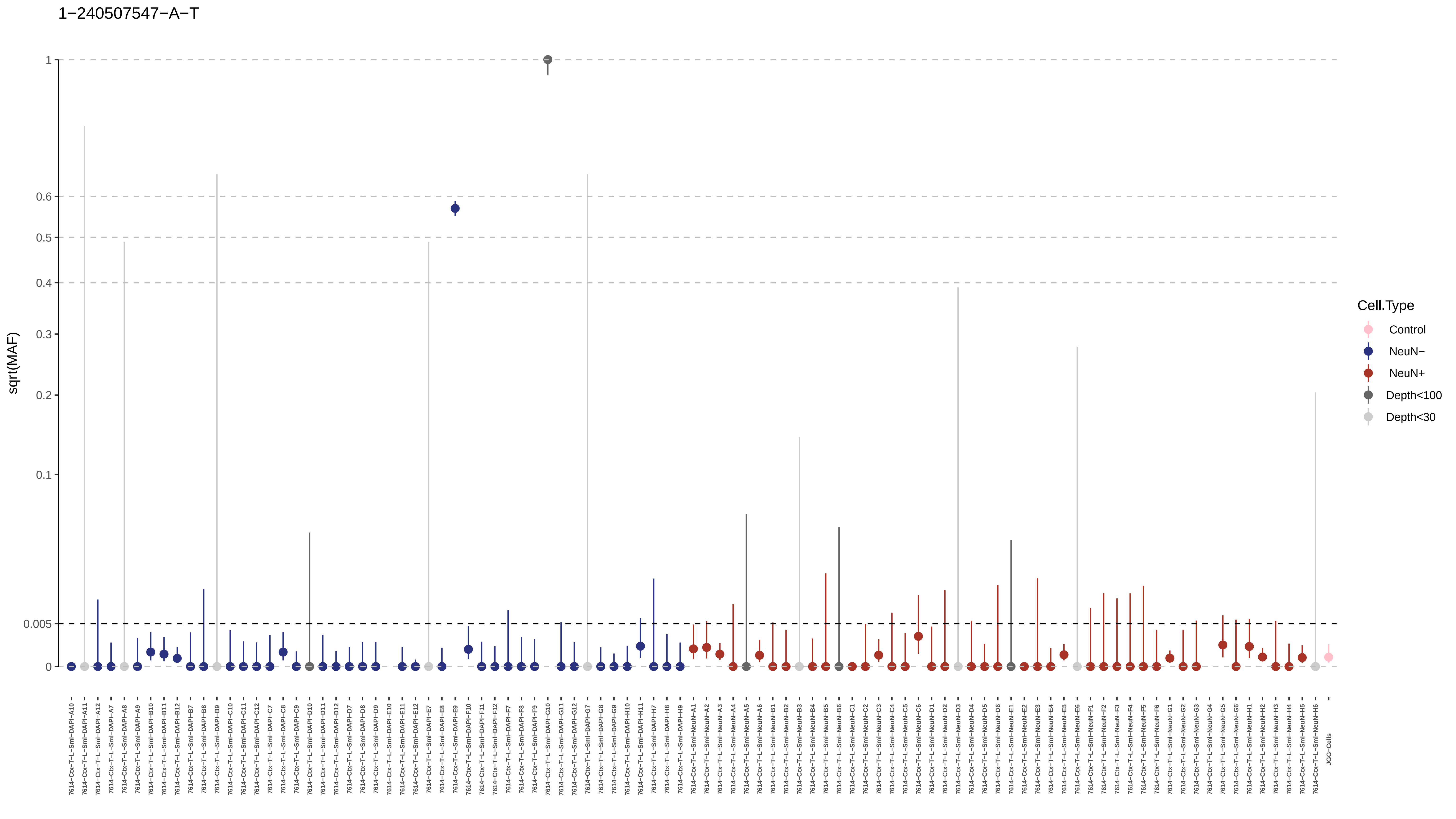
1−240507547−A−T
sqrt(MAF)
1
0.6
0.5
0.4
0.3
0.2
0.1
0.005
0
Cell.Type
Control
NeuN−
NeuN+
Depth<100
Depth<30
7614−Ctx−T−L−Sml−DAPI−A10
7614−Ctx−T−L−Sml−DAPI−A11
7614−Ctx−T−L−Sml−DAPI−A12
7614−Ctx−T−L−Sml−DAPI−A7
7614−Ctx−T−L−Sml−DAPI−A8
7614−Ctx−T−L−Sml−DAPI−A9
7614−Ctx−T−L−Sml−DAPI−B10
7614−Ctx−T−L−Sml−DAPI−B11
7614−Ctx−T−L−Sml−DAPI−B12
7614−Ctx−T−L−Sml−DAPI−B7
7614−Ctx−T−L−Sml−DAPI−B8
7614−Ctx−T−L−Sml−DAPI−B9
7614−Ctx−T−L−Sml−DAPI−C10
7614−Ctx−T−L−Sml−DAPI−C11
7614−Ctx−T−L−Sml−DAPI−C12
7614−Ctx−T−L−Sml−DAPI−C7
7614−Ctx−T−L−Sml−DAPI−C8
7614−Ctx−T−L−Sml−DAPI−C9
7614−Ctx−T−L−Sml−DAPI−D10
7614−Ctx−T−L−Sml−DAPI−D11
7614−Ctx−T−L−Sml−DAPI−D12
7614−Ctx−T−L−Sml−DAPI−D7
7614−Ctx−T−L−Sml−DAPI−D8
7614−Ctx−T−L−Sml−DAPI−D9
7614−Ctx−T−L−Sml−DAPI−E10
7614−Ctx−T−L−Sml−DAPI−E11
7614−Ctx−T−L−Sml−DAPI−E12
7614−Ctx−T−L−Sml−DAPI−E7
7614−Ctx−T−L−Sml−DAPI−E8
7614−Ctx−T−L−Sml−DAPI−E9
7614−Ctx−T−L−Sml−DAPI−F10
7614−Ctx−T−L−Sml−DAPI−F11
7614−Ctx−T−L−Sml−DAPI−F12
7614−Ctx−T−L−Sml−DAPI−F7
7614−Ctx−T−L−Sml−DAPI−F8
7614−Ctx−T−L−Sml−DAPI−F9
7614−Ctx−T−L−Sml−DAPI−G10
7614−Ctx−T−L−Sml−DAPI−G11
7614−Ctx−T−L−Sml−DAPI−G12
7614−Ctx−T−L−Sml−DAPI−G7
7614−Ctx−T−L−Sml−DAPI−G8
7614−Ctx−T−L−Sml−DAPI−G9
7614−Ctx−T−L−Sml−DAPI−H10
7614−Ctx−T−L−Sml−DAPI−H11
7614−Ctx−T−L−Sml−DAPI−H7
7614−Ctx−T−L−Sml−DAPI−H8
7614−Ctx−T−L−Sml−DAPI−H9
7614−Ctx−T−L−Sml−NeuN−A1
7614−Ctx−T−L−Sml−NeuN−A2
7614−Ctx−T−L−Sml−NeuN−A3
7614−Ctx−T−L−Sml−NeuN−A4
7614−Ctx−T−L−Sml−NeuN−A5
7614−Ctx−T−L−Sml−NeuN−A6
7614−Ctx−T−L−Sml−NeuN−B1
7614−Ctx−T−L−Sml−NeuN−B2
7614−Ctx−T−L−Sml−NeuN−B3
7614−Ctx−T−L−Sml−NeuN−B4
7614−Ctx−T−L−Sml−NeuN−B5
7614−Ctx−T−L−Sml−NeuN−B6
7614−Ctx−T−L−Sml−NeuN−C1
7614−Ctx−T−L−Sml−NeuN−C2
7614−Ctx−T−L−Sml−NeuN−C3
7614−Ctx−T−L−Sml−NeuN−C4
7614−Ctx−T−L−Sml−NeuN−C5
7614−Ctx−T−L−Sml−NeuN−C6
7614−Ctx−T−L−Sml−NeuN−D1
7614−Ctx−T−L−Sml−NeuN−D2
7614−Ctx−T−L−Sml−NeuN−D3
7614−Ctx−T−L−Sml−NeuN−D4
7614−Ctx−T−L−Sml−NeuN−D5
7614−Ctx−T−L−Sml−NeuN−D6
7614−Ctx−T−L−Sml−NeuN−E1
7614−Ctx−T−L−Sml−NeuN−E2
7614−Ctx−T−L−Sml−NeuN−E3
7614−Ctx−T−L−Sml−NeuN−E4
7614−Ctx−T−L−Sml−NeuN−E5
7614−Ctx−T−L−Sml−NeuN−E6
7614−Ctx−T−L−Sml−NeuN−F1
7614−Ctx−T−L−Sml−NeuN−F2
7614−Ctx−T−L−Sml−NeuN−F3
7614−Ctx−T−L−Sml−NeuN−F4
7614−Ctx−T−L−Sml−NeuN−F5
7614−Ctx−T−L−Sml−NeuN−F6
7614−Ctx−T−L−Sml−NeuN−G1
7614−Ctx−T−L−Sml−NeuN−G2
7614−Ctx−T−L−Sml−NeuN−G3
7614−Ctx−T−L−Sml−NeuN−G4
7614−Ctx−T−L−Sml−NeuN−G5
7614−Ctx−T−L−Sml−NeuN−G6
7614−Ctx−T−L−Sml−NeuN−H1
7614−Ctx−T−L−Sml−NeuN−H2
7614−Ctx−T−L−Sml−NeuN−H3
7614−Ctx−T−L−Sml−NeuN−H4
7614−Ctx−T−L−Sml−NeuN−H5
7614−Ctx−T−L−Sml−NeuN−H6
JGG-Cells

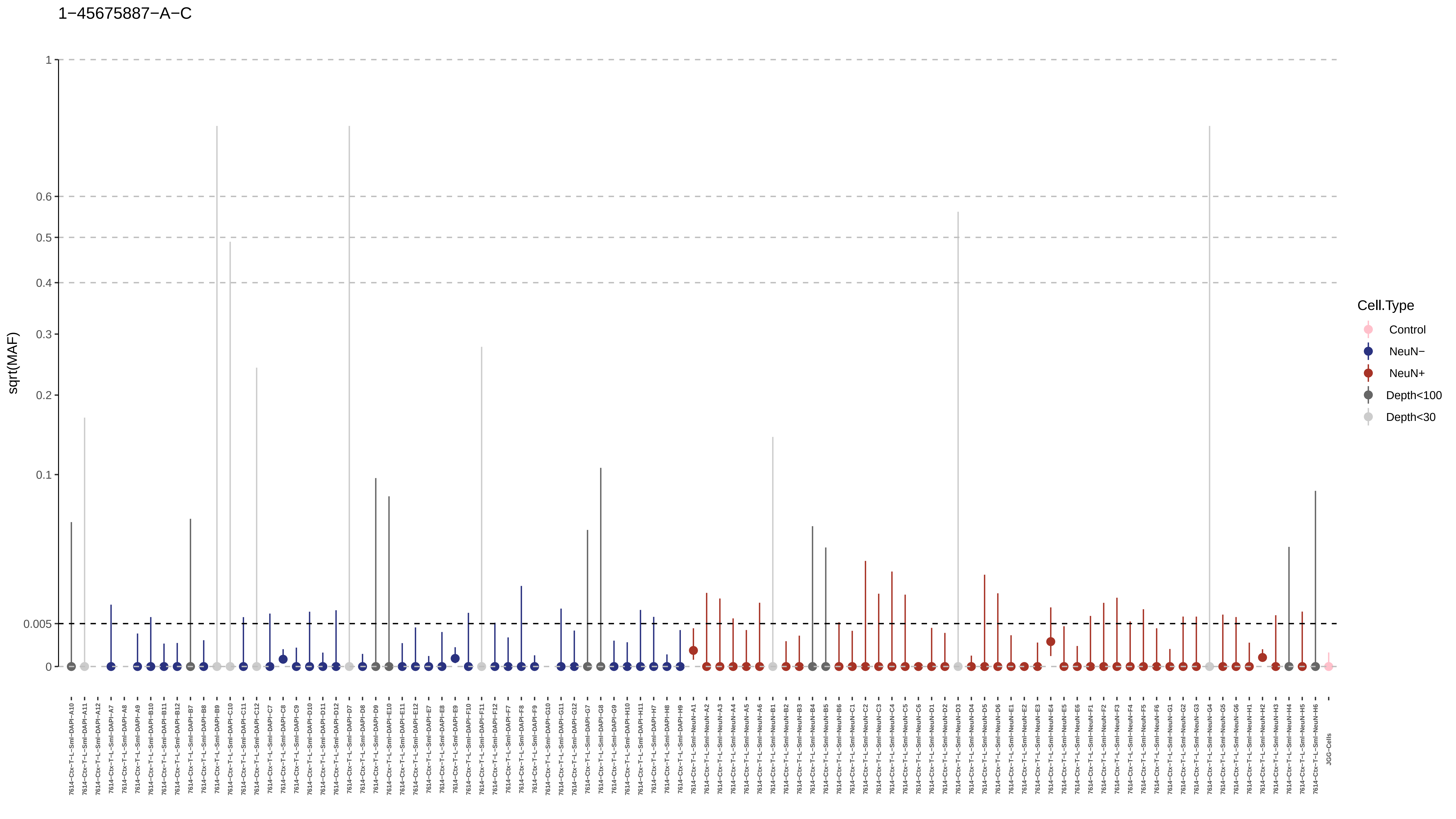
1−45675887−A−C
sqrt(MAF)
1
0.6
0.5
0.4
0.3
0.2
0.1
0.005
0
Cell.Type
Control
NeuN−
NeuN+
Depth<100
Depth<30

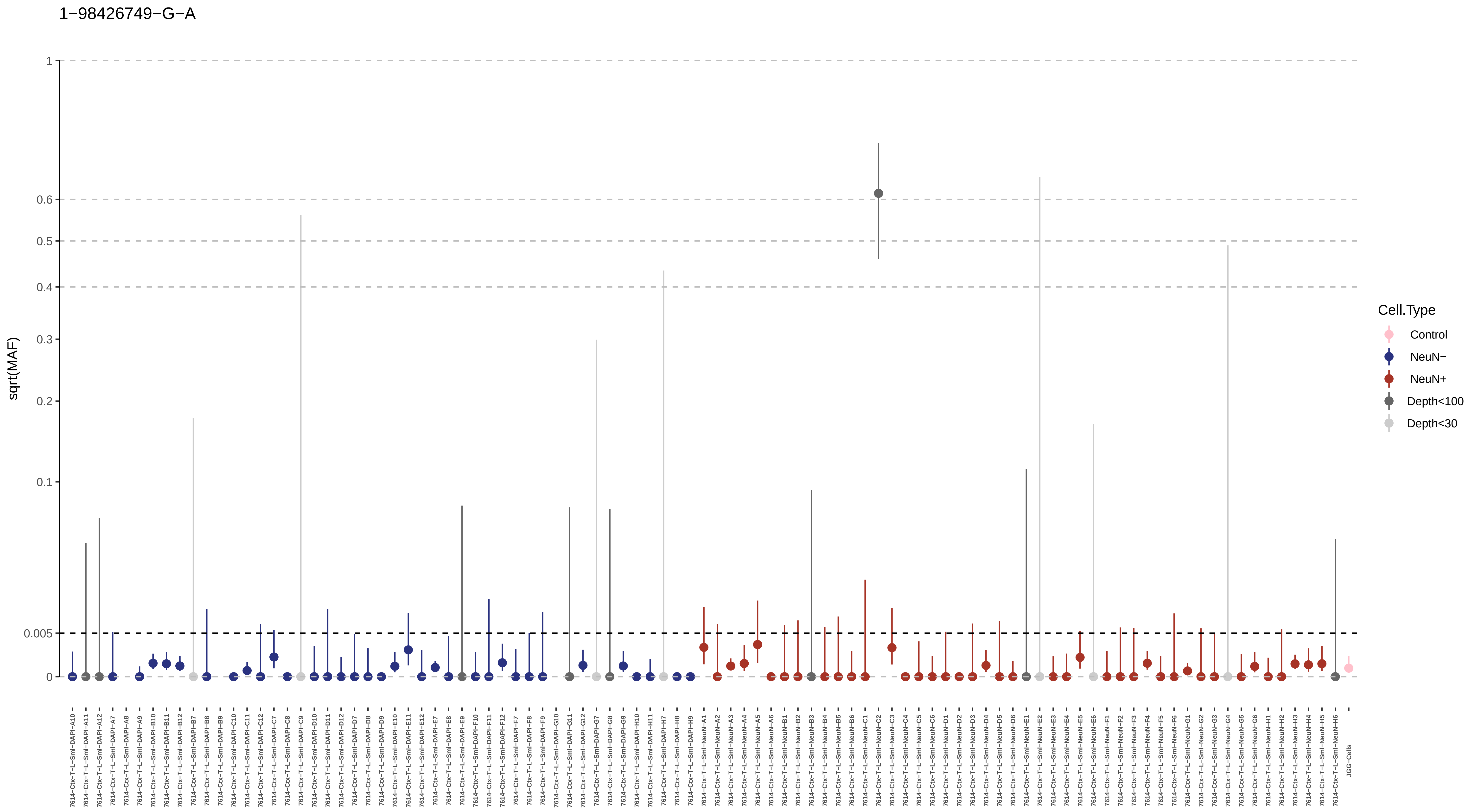

1−98426749−G−A
sqrt(MAF)
1
0.6
0.5
0.4
0.3
0.2
0.1
0.005
0
Cell.Type
Control
NeuN−
NeuN+
Depth<100
Depth<30
7614−Ctx−T−L−Sml−DAPI−A10
7614−Ctx−T−L−Sml−DAPI−A11
7614−Ctx−T−L−Sml−DAPI−A12
7614−Ctx−T−L−Sml−DAPI−A7
7614−Ctx−T−L−Sml−DAPI−A8
7614−Ctx−T−L−Sml−DAPI−A9
7614−Ctx−T−L−Sml−DAPI−B10
7614−Ctx−T−L−Sml−DAPI−B11
7614−Ctx−T−L−Sml−DAPI−B12
7614−Ctx−T−L−Sml−DAPI−B7
7614−Ctx−T−L−Sml−DAPI−B8
7614−Ctx−T−L−Sml−DAPI−B9
7614−Ctx−T−L−Sml−DAPI−C10
7614−Ctx−T−L−Sml−DAPI−C11
7614−Ctx−T−L−Sml−DAPI−C12
7614−Ctx−T−L−Sml−DAPI−C7
7614−Ctx−T−L−Sml−DAPI−C8
7614−Ctx−T−L−Sml−DAPI−C9
7614−Ctx−T−L−Sml−DAPI−D10
7614−Ctx−T−L−Sml−DAPI−D11
7614−Ctx−T−L−Sml−DAPI−D12
7614−Ctx−T−L−Sml−DAPI−D7
7614−Ctx−T−L−Sml−DAPI−D8
7614−Ctx−T−L−Sml−DAPI−D9
7614−Ctx−T−L−Sml−DAPI−E10
7614−Ctx−T−L−Sml−DAPI−E11
7614−Ctx−T−L−Sml−DAPI−E12
7614−Ctx−T−L−Sml−DAPI−E7
7614−Ctx−T−L−Sml−DAPI−E8
7614−Ctx−T−L−Sml−DAPI−E9
7614−Ctx−T−L−Sml−DAPI−F10
7614−Ctx−T−L−Sml−DAPI−F11
7614−Ctx−T−L−Sml−DAPI−F12
7614−Ctx−T−L−Sml−DAPI−F7
7614−Ctx−T−L−Sml−DAPI−F8
7614−Ctx−T−L−Sml−DAPI−F9
7614−Ctx−T−L−Sml−DAPI−G10
7614−Ctx−T−L−Sml−DAPI−G11
7614−Ctx−T−L−Sml−DAPI−G12
7614−Ctx−T−L−Sml−DAPI−G7
7614−Ctx−T−L−Sml−DAPI−G8
7614−Ctx−T−L−Sml−DAPI−G9
7614−Ctx−T−L−Sml−DAPI−H10
7614−Ctx−T−L−Sml−DAPI−H11
7614−Ctx−T−L−Sml−DAPI−H7
7614−Ctx−T−L−Sml−DAPI−H8
7614−Ctx−T−L−Sml−DAPI−H9
7614−Ctx−T−L−Sml−NeuN−A1
7614−Ctx−T−L−Sml−NeuN−A2
7614−Ctx−T−L−Sml−NeuN−A3
7614−Ctx−T−L−Sml−NeuN−A4
7614−Ctx−T−L−Sml−NeuN−A5
7614−Ctx−T−L−Sml−NeuN−A6
7614−Ctx−T−L−Sml−NeuN−B1
7614−Ctx−T−L−Sml−NeuN−B2
7614−Ctx−T−L−Sml−NeuN−B3
7614−Ctx−T−L−Sml−NeuN−B4
7614−Ctx−T−L−Sml−NeuN−B5
7614−Ctx−T−L−Sml−NeuN−B6
7614−Ctx−T−L−Sml−NeuN−C1
7614−Ctx−T−L−Sml−NeuN−C2
7614−Ctx−T−L−Sml−NeuN−C3
7614−Ctx−T−L−Sml−NeuN−C4
7614−Ctx−T−L−Sml−NeuN−C5
7614−Ctx−T−L−Sml−NeuN−C6
7614−Ctx−T−L−Sml−NeuN−D1
7614−Ctx−T−L−Sml−NeuN−D2
7614−Ctx−T−L−Sml−NeuN−D3
7614−Ctx−T−L−Sml−NeuN−D4
7614−Ctx−T−L−Sml−NeuN−D5
7614−Ctx−T−L−Sml−NeuN−D6
7614−Ctx−T−L−Sml−NeuN−E1
7614−Ctx−T−L−Sml−NeuN−E2
7614−Ctx−T−L−Sml−NeuN−E3
7614−Ctx−T−L−Sml−NeuN−E4
7614−Ctx−T−L−Sml−NeuN−E5
7614−Ctx−T−L−Sml−NeuN−E6
7614−Ctx−T−L−Sml−NeuN−F1
7614−Ctx−T−L−Sml−NeuN−F2
7614−Ctx−T−L−Sml−NeuN−F3
7614−Ctx−T−L−Sml−NeuN−F4
7614−Ctx−T−L−Sml−NeuN−F5
7614−Ctx−T−L−Sml−NeuN−F6
7614−Ctx−T−L−Sml−NeuN−G1
7614−Ctx−T−L−Sml−NeuN−G2
7614−Ctx−T−L−Sml−NeuN−G3
7614−Ctx−T−L−Sml−NeuN−G4
7614−Ctx−T−L−Sml−NeuN−G5
7614−Ctx−T−L−Sml−NeuN−G6
7614−Ctx−T−L−Sml−NeuN−H1
7614−Ctx−T−L−Sml−NeuN−H2
7614−Ctx−T−L−Sml−NeuN−H3
7614−Ctx−T−L−Sml−NeuN−H4
7614−Ctx−T−L−Sml−NeuN−H5
7614−Ctx−T−L−Sml−NeuN−H6
JGG−Cells

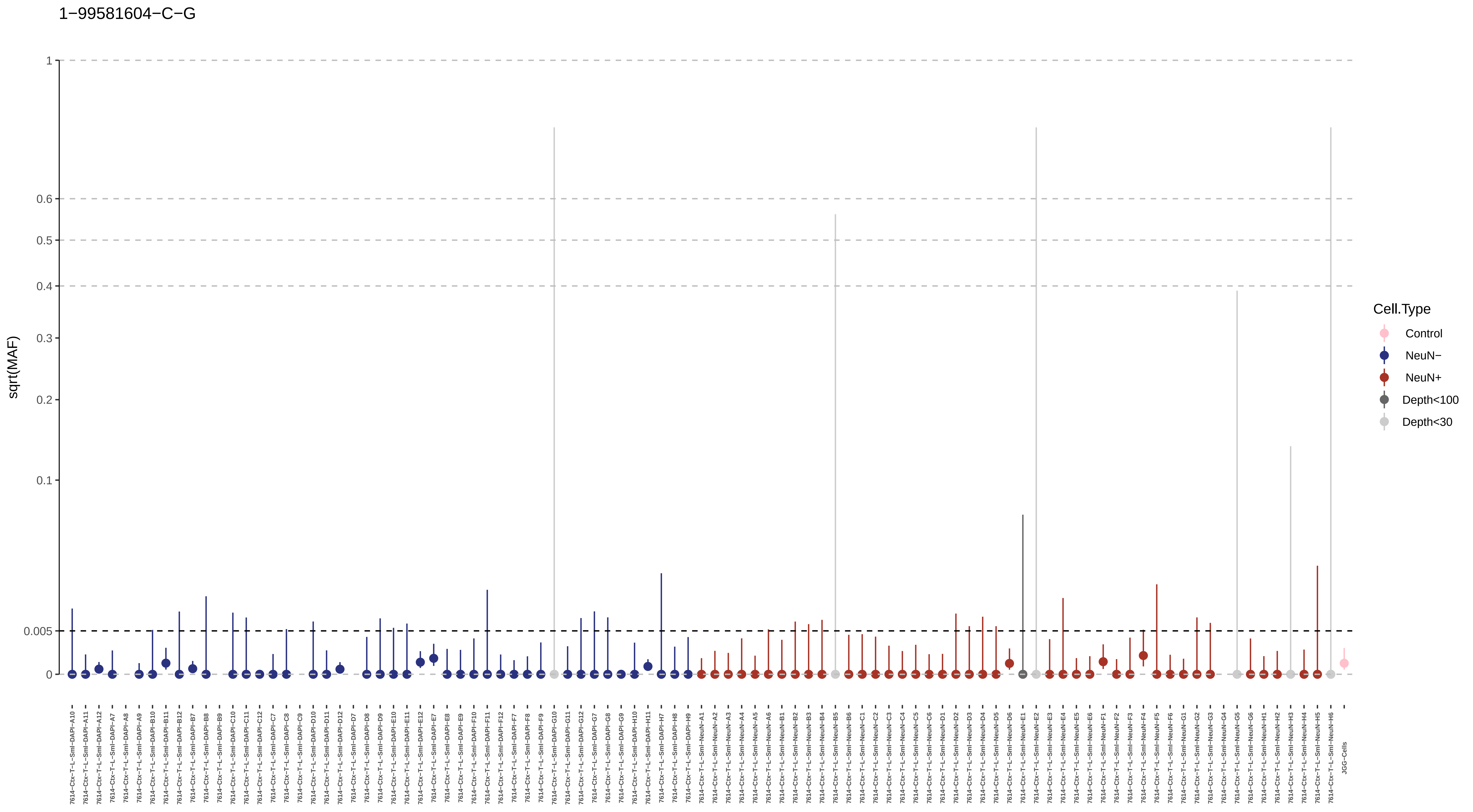
1−99581604−C−G
sqrt(MAF)
1
0.6
0.5
0.4
0.3
0.2
0.1
0.005
0
Cell.Type
Control
NeuN−
NeuN+
Depth<100
Depth<30
JGG-Cells

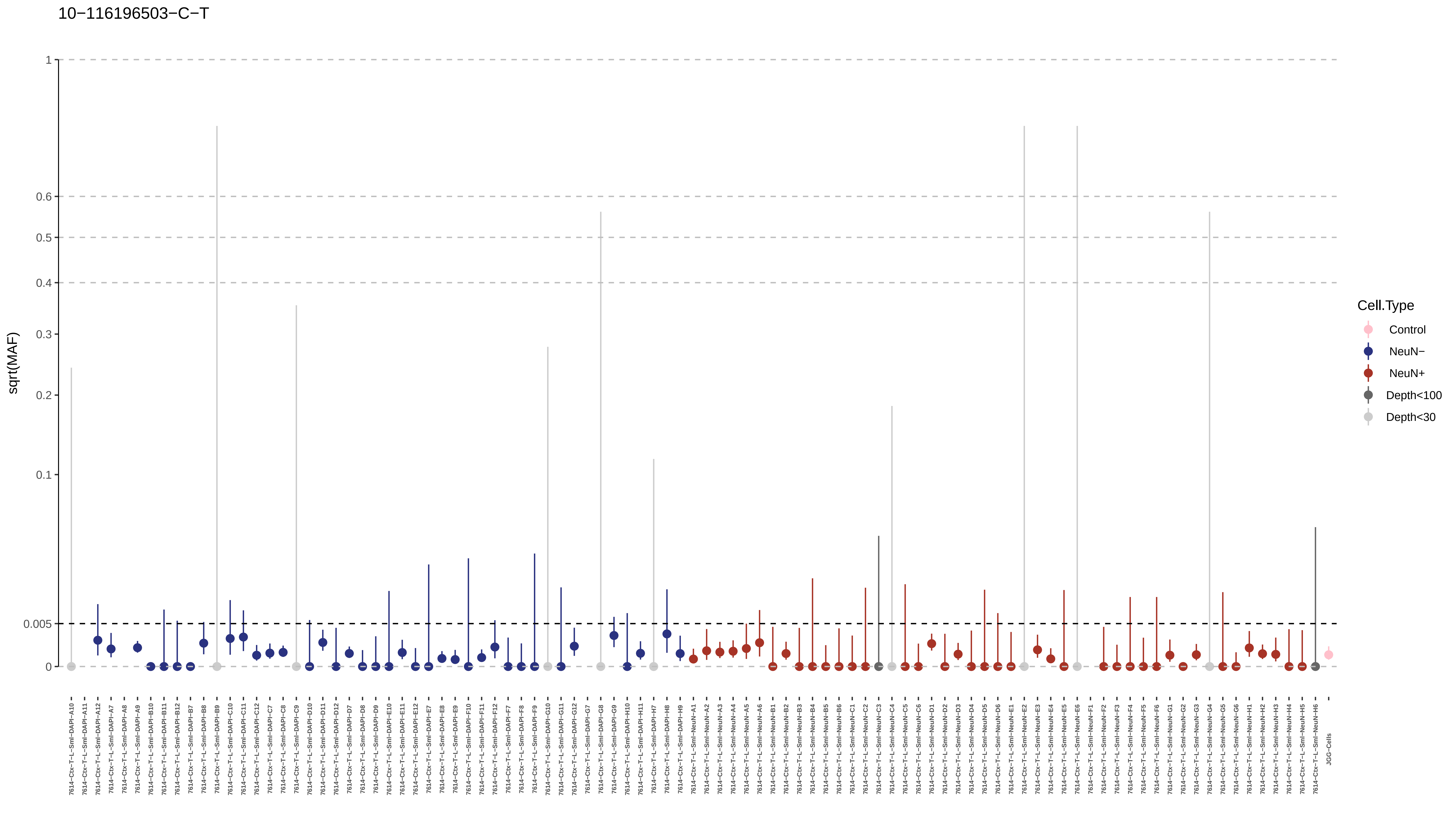

10−116196503−C−T
sqrt(MAF)
1
0.6
0.5
0.4
0.3
0.2
0.1
0.005
0
Cell.Type
Control
NeuN−
NeuN+
Depth<100
Depth<30
7614−Ctx−T−L−Sml−DAPI−A10
7614−Ctx−T−L−Sml−DAPI−A11
7614−Ctx−T−L−Sml−DAPI−A12
7614−Ctx−T−L−Sml−DAPI−A7
7614−Ctx−T−L−Sml−DAPI−A8
7614−Ctx−T−L−Sml−DAPI−A9
7614−Ctx−T−L−Sml−DAPI−B10
7614−Ctx−T−L−Sml−DAPI−B11
7614−Ctx−T−L−Sml−DAPI−B12
7614−Ctx−T−L−Sml−DAPI−B7
7614−Ctx−T−L−Sml−DAPI−B8
7614−Ctx−T−L−Sml−DAPI−B9
7614−Ctx−T−L−Sml−DAPI−C10
7614−Ctx−T−L−Sml−DAPI−C11
7614−Ctx−T−L−Sml−DAPI−C12
7614−Ctx−T−L−Sml−DAPI−C7
7614−Ctx−T−L−Sml−DAPI−C8
7614−Ctx−T−L−Sml−DAPI−C9
7614−Ctx−T−L−Sml−DAPI−D10
7614−Ctx−T−L−Sml−DAPI−D11
7614−Ctx−T−L−Sml−DAPI−D12
7614−Ctx−T−L−Sml−DAPI−D7
7614−Ctx−T−L−Sml−DAPI−D8
7614−Ctx−T−L−Sml−DAPI−D9
7614−Ctx−T−L−Sml−DAPI−E10
7614−Ctx−T−L−Sml−DAPI−E11
7614−Ctx−T−L−Sml−DAPI−E12
7614−Ctx−T−L−Sml−DAPI−E7
7614−Ctx−T−L−Sml−DAPI−E8
7614−Ctx−T−L−Sml−DAPI−E9
7614−Ctx−T−L−Sml−DAPI−F10
7614−Ctx−T−L−Sml−DAPI−F11
7614−Ctx−T−L−Sml−DAPI−F12
7614−Ctx−T−L−Sml−DAPI−F7
7614−Ctx−T−L−Sml−DAPI−F8
7614−Ctx−T−L−Sml−DAPI−F9
7614−Ctx−T−L−Sml−DAPI−G10
7614−Ctx−T−L−Sml−DAPI−G11
7614−Ctx−T−L−Sml−DAPI−G12
7614−Ctx−T−L−Sml−DAPI−G7
7614−Ctx−T−L−Sml−DAPI−G8
7614−Ctx−T−L−Sml−DAPI−G9
7614−Ctx−T−L−Sml−DAPI−H10
7614−Ctx−T−L−Sml−DAPI−H11
7614−Ctx−T−L−Sml−DAPI−H7
7614−Ctx−T−L−Sml−DAPI−H8
7614−Ctx−T−L−Sml−DAPI−H9
7614−Ctx−T−L−Sml−NeuN−A1
7614−Ctx−T−L−Sml−NeuN−A2
7614−Ctx−T−L−Sml−NeuN−A3
7614−Ctx−T−L−Sml−NeuN−A4
7614−Ctx−T−L−Sml−NeuN−A5
7614−Ctx−T−L−Sml−NeuN−A6
7614−Ctx−T−L−Sml−NeuN−B1
7614−Ctx−T−L−Sml−NeuN−B2
7614−Ctx−T−L−Sml−NeuN−B3
7614−Ctx−T−L−Sml−NeuN−B4
7614−Ctx−T−L−Sml−NeuN−B5
7614−Ctx−T−L−Sml−NeuN−B6
7614−Ctx−T−L−Sml−NeuN−C1
7614−Ctx−T−L−Sml−NeuN−C2
7614−Ctx−T−L−Sml−NeuN−C3
7614−Ctx−T−L−Sml−NeuN−C4
7614−Ctx−T−L−Sml−NeuN−C5
7614−Ctx−T−L−Sml−NeuN−C6
7614−Ctx−T−L−Sml−NeuN−D1
7614−Ctx−T−L−Sml−NeuN−D2
7614−Ctx−T−L−Sml−NeuN−D3
7614−Ctx−T−L−Sml−NeuN−D4
7614−Ctx−T−L−Sml−NeuN−D5
7614−Ctx−T−L−Sml−NeuN−D6
7614−Ctx−T−L−Sml−NeuN−E1
7614−Ctx−T−L−Sml−NeuN−E2
7614−Ctx−T−L−Sml−NeuN−E3
7614−Ctx−T−L−Sml−NeuN−E4
7614−Ctx−T−L−Sml−NeuN−E5
7614−Ctx−T−L−Sml−NeuN−E6
7614−Ctx−T−L−Sml−NeuN−F1
7614−Ctx−T−L−Sml−NeuN−F2
7614−Ctx−T−L−Sml−NeuN−F3
7614−Ctx−T−L−Sml−NeuN−F4
7614−Ctx−T−L−Sml−NeuN−F5
7614−Ctx−T−L−Sml−NeuN−F6
7614−Ctx−T−L−Sml−NeuN−G1
7614−Ctx−T−L−Sml−NeuN−G2
7614−Ctx−T−L−Sml−NeuN−G3
7614−Ctx−T−L−Sml−NeuN−G4
7614−Ctx−T−L−Sml−NeuN−G5
7614−Ctx−T−L−Sml−NeuN−G6
7614−Ctx−T−L−Sml−NeuN−H1
7614−Ctx−T−L−Sml−NeuN−H2
7614−Ctx−T−L−Sml−NeuN−H3
7614−Ctx−T−L−Sml−NeuN−H4
7614−Ctx−T−L−Sml−NeuN−H5
7614−Ctx−T−L−Sml−NeuN−H6
JGG−Cells

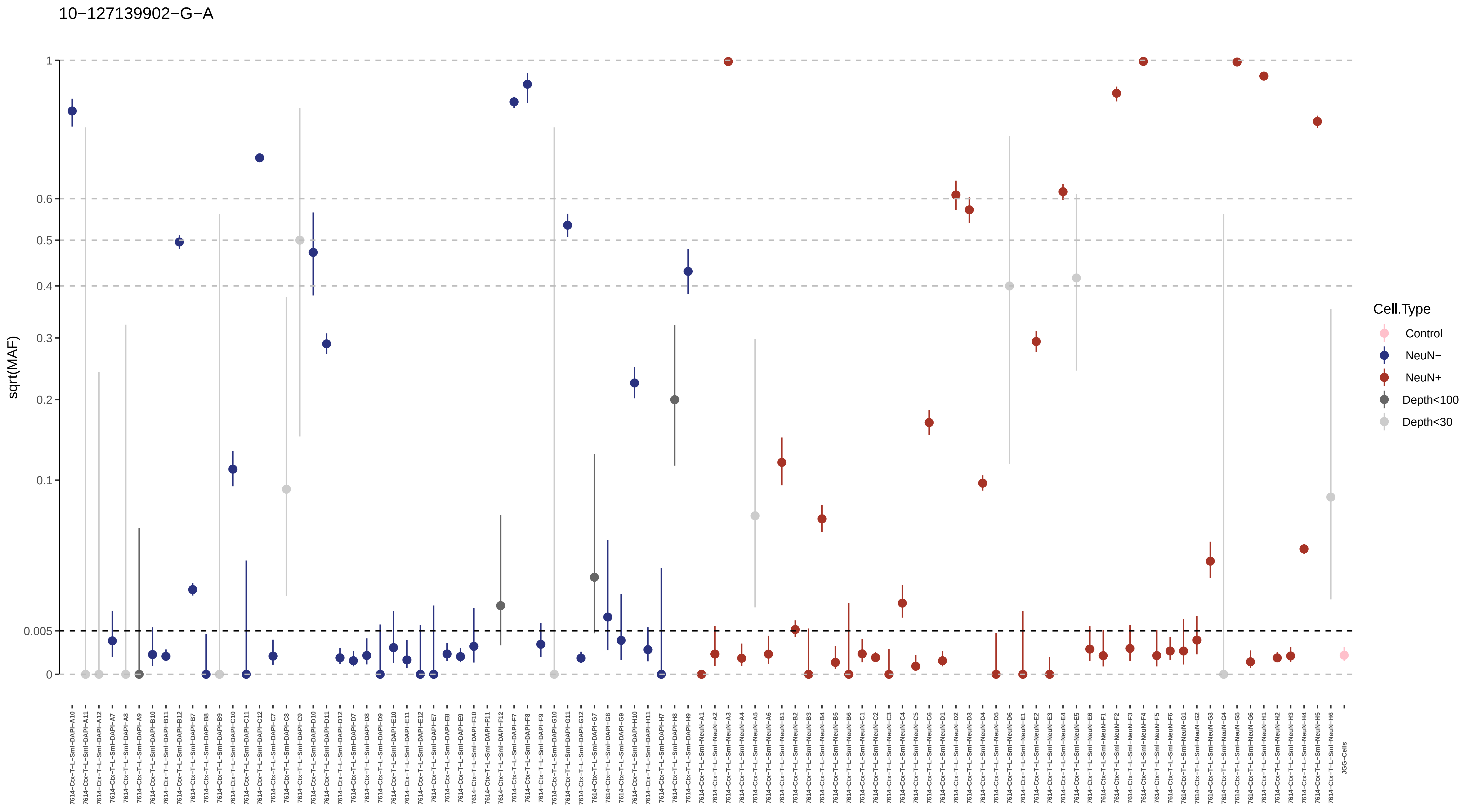
10−127139902−G−A
sqrt(MAF)
1
0.6
0.5
0.4
0.3
0.2
0.1
0.005
0
Cell.Type
Control
NeuN−
NeuN+
Depth<100
Depth<30
JGG-Cells

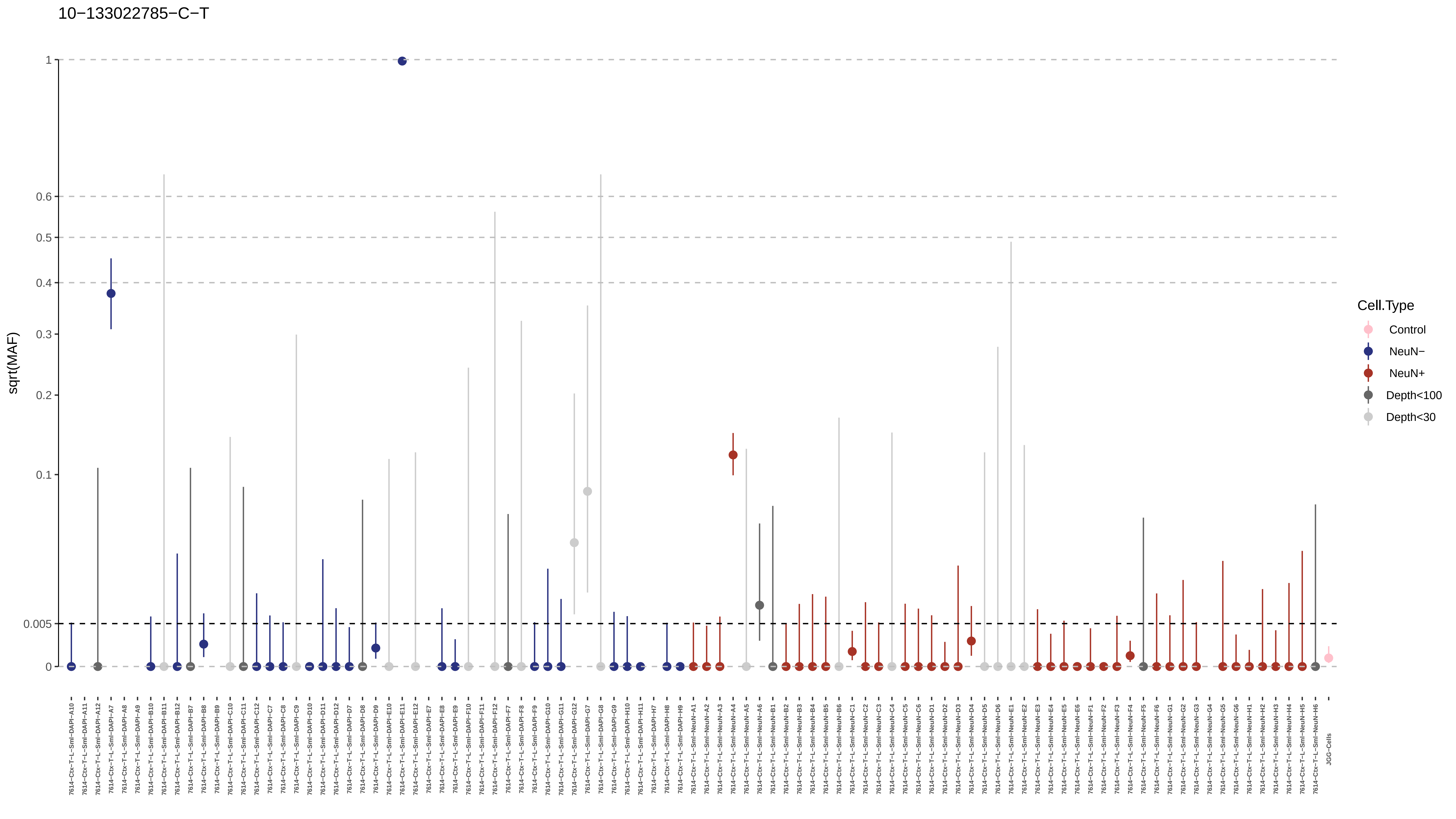
10−133022785−C−T
sqrt(MAF)
1
0.6
0.5
0.4
0.3
0.2
0.1
0.005
0
Cell.Type
Control
NeuN−
NeuN+
Depth<100
Depth<30

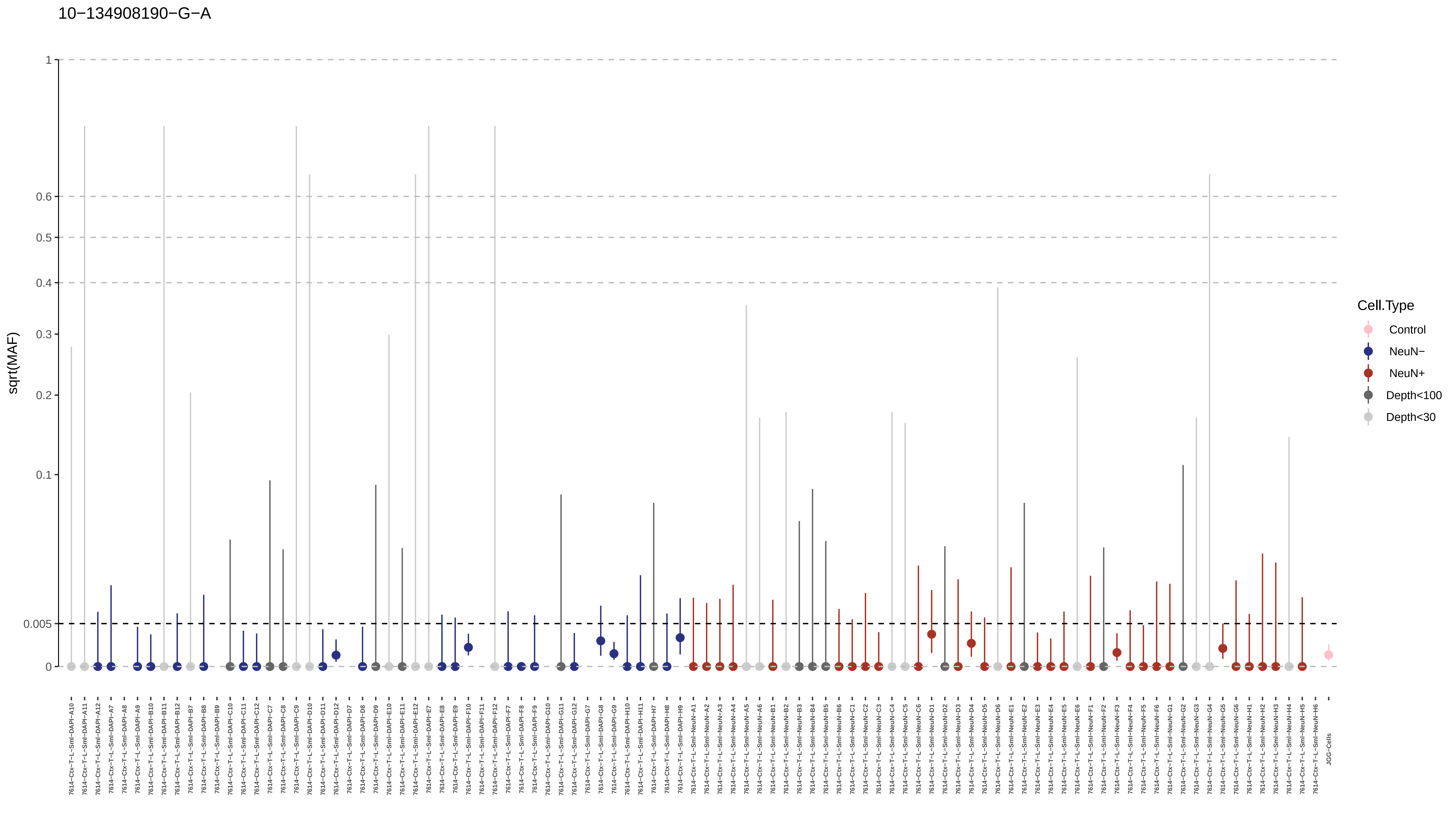
10−134908190−G−A
sqrt(MAF)
1
0.6
0.5
0.4
0.3
0.2
0.1
0.005
0
Cell.Type
Control
NeuN−
NeuN+
Depth<100
Depth<30

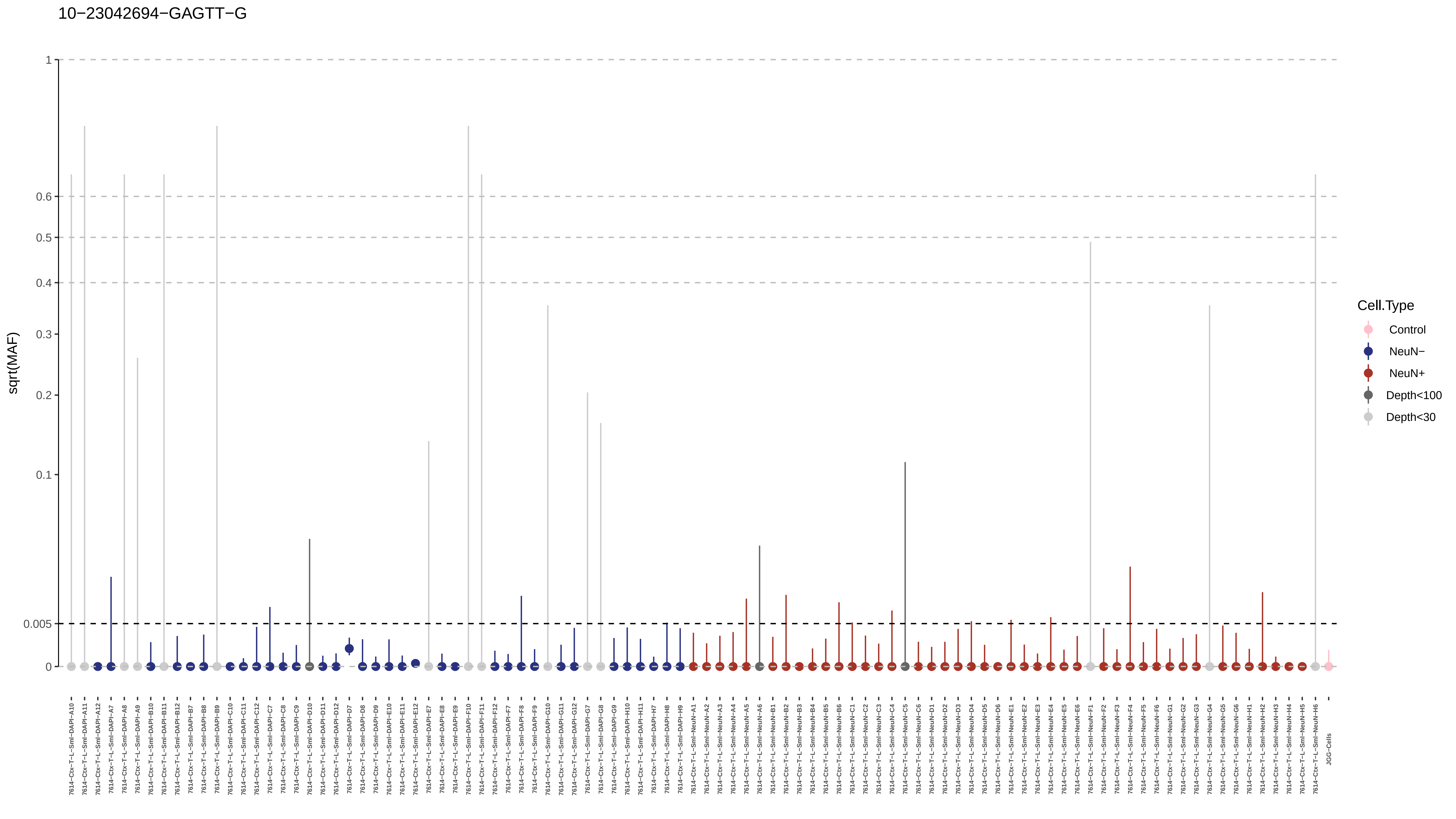
10−23042694−GAGTT−G
sqrt(MAF)
1
0.6
0.5
0.4
0.3
0.2
0.1
0.005
0
Cell.Type
Control
NeuN−
NeuN+
Depth<100
Depth<30
7614−Ctx−T−L−Sml−DAPI−A10
7614−Ctx−T−L−Sml−DAPI−A11
7614−Ctx−T−L−Sml−DAPI−A12
7614−Ctx−T−L−Sml−DAPI−A7
7614−Ctx−T−L−Sml−DAPI−A8
7614−Ctx−T−L−Sml−DAPI−A9
7614−Ctx−T−L−Sml−DAPI−B10
7614−Ctx−T−L−Sml−DAPI−B11
7614−Ctx−T−L−Sml−DAPI−B12
7614−Ctx−T−L−Sml−DAPI−B7
7614−Ctx−T−L−Sml−DAPI−B8
7614−Ctx−T−L−Sml−DAPI−B9
7614−Ctx−T−L−Sml−DAPI−C10
7614−Ctx−T−L−Sml−DAPI−C11
7614−Ctx−T−L−Sml−DAPI−C12
7614−Ctx−T−L−Sml−DAPI−C7
7614−Ctx−T−L−Sml−DAPI−C8
7614−Ctx−T−L−Sml−DAPI−C9
7614−Ctx−T−L−Sml−DAPI−D10
7614−Ctx−T−L−Sml−DAPI−D11
7614−Ctx−T−L−Sml−DAPI−D12
7614−Ctx−T−L−Sml−DAPI−D7
7614−Ctx−T−L−Sml−DAPI−D8
7614−Ctx−T−L−Sml−DAPI−D9
7614−Ctx−T−L−Sml−DAPI−E10
7614−Ctx−T−L−Sml−DAPI−E11
7614−Ctx−T−L−Sml−DAPI−E12
7614−Ctx−T−L−Sml−DAPI−E7
7614−Ctx−T−L−Sml−DAPI−E8
7614−Ctx−T−L−Sml−DAPI−E9
7614−Ctx−T−L−Sml−DAPI−F10
7614−Ctx−T−L−Sml−DAPI−F11
7614−Ctx−T−L−Sml−DAPI−F12
7614−Ctx−T−L−Sml−DAPI−F7
7614−Ctx−T−L−Sml−DAPI−F8
7614−Ctx−T−L−Sml−DAPI−F9
7614−Ctx−T−L−Sml−DAPI−G10
7614−Ctx−T−L−Sml−DAPI−G11
7614−Ctx−T−L−Sml−DAPI−G12
7614−Ctx−T−L−Sml−DAPI−G7
7614−Ctx−T−L−Sml−DAPI−G8
7614−Ctx−T−L−Sml−DAPI−G9
7614−Ctx−T−L−Sml−DAPI−H10
7614−Ctx−T−L−Sml−DAPI−H11
7614−Ctx−T−L−Sml−DAPI−H7
7614−Ctx−T−L−Sml−DAPI−H8
7614−Ctx−T−L−Sml−DAPI−H9
7614−Ctx−T−L−Sml−NeuN−A1
7614−Ctx−T−L−Sml−NeuN−A2
7614−Ctx−T−L−Sml−NeuN−A3
7614−Ctx−T−L−Sml−NeuN−A4
7614−Ctx−T−L−Sml−NeuN−A5
7614−Ctx−T−L−Sml−NeuN−A6
7614−Ctx−T−L−Sml−NeuN−B1
7614−Ctx−T−L−Sml−NeuN−B2
7614−Ctx−T−L−Sml−NeuN−B3
7614−Ctx−T−L−Sml−NeuN−B4
7614−Ctx−T−L−Sml−NeuN−B5
7614−Ctx−T−L−Sml−NeuN−B6
7614−Ctx−T−L−Sml−NeuN−C1
7614−Ctx−T−L−Sml−NeuN−C2
7614−Ctx−T−L−Sml−NeuN−C3
7614−Ctx−T−L−Sml−NeuN−C4
7614−Ctx−T−L−Sml−NeuN−C5
7614−Ctx−T−L−Sml−NeuN−C6
7614−Ctx−T−L−Sml−NeuN−D1
7614−Ctx−T−L−Sml−NeuN−D2
7614−Ctx−T−L−Sml−NeuN−D3
7614−Ctx−T−L−Sml−NeuN−D4
7614−Ctx−T−L−Sml−NeuN−D5
7614−Ctx−T−L−Sml−NeuN−D6
7614−Ctx−T−L−Sml−NeuN−E1
7614−Ctx−T−L−Sml−NeuN−E2
7614−Ctx−T−L−Sml−NeuN−E3
7614−Ctx−T−L−Sml−NeuN−E4
7614−Ctx−T−L−Sml−NeuN−E5
7614−Ctx−T−L−Sml−NeuN−E6
7614−Ctx−T−L−Sml−NeuN−F1
7614−Ctx−T−L−Sml−NeuN−F2
7614−Ctx−T−L−Sml−NeuN−F3
7614−Ctx−T−L−Sml−NeuN−F4
7614−Ctx−T−L−Sml−NeuN−F5
7614−Ctx−T−L−Sml−NeuN−F6
7614−Ctx−T−L−Sml−NeuN−G1
7614−Ctx−T−L−Sml−NeuN−G2
7614−Ctx−T−L−Sml−NeuN−G3
7614−Ctx−T−L−Sml−NeuN−G4
7614−Ctx−T−L−Sml−NeuN−G5
7614−Ctx−T−L−Sml−NeuN−G6
7614−Ctx−T−L−Sml−NeuN−H1
7614−Ctx−T−L−Sml−NeuN−H2
7614−Ctx−T−L−Sml−NeuN−H3
7614−Ctx−T−L−Sml−NeuN−H4
7614−Ctx−T−L−Sml−NeuN−H5
7614−Ctx−T−L−Sml−NeuN−H6
JGG-Cells

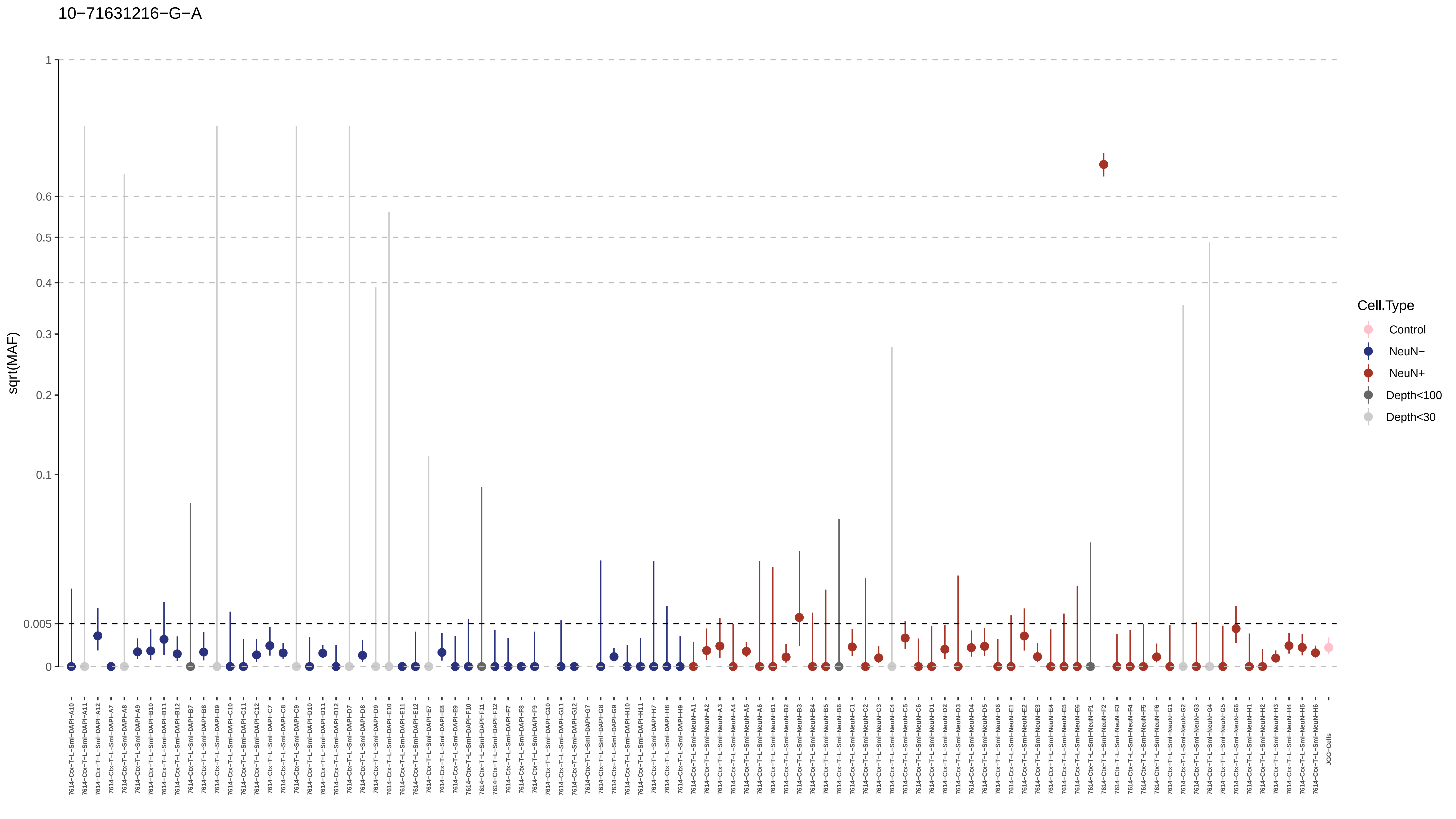

10−71631216−G−A
sqrt(MAF)
1
0.6
0.5
0.4
0.3
0.2
0.1
0.005
0
Cell.Type
Control
NeuN−
NeuN+
Depth<100
Depth<30

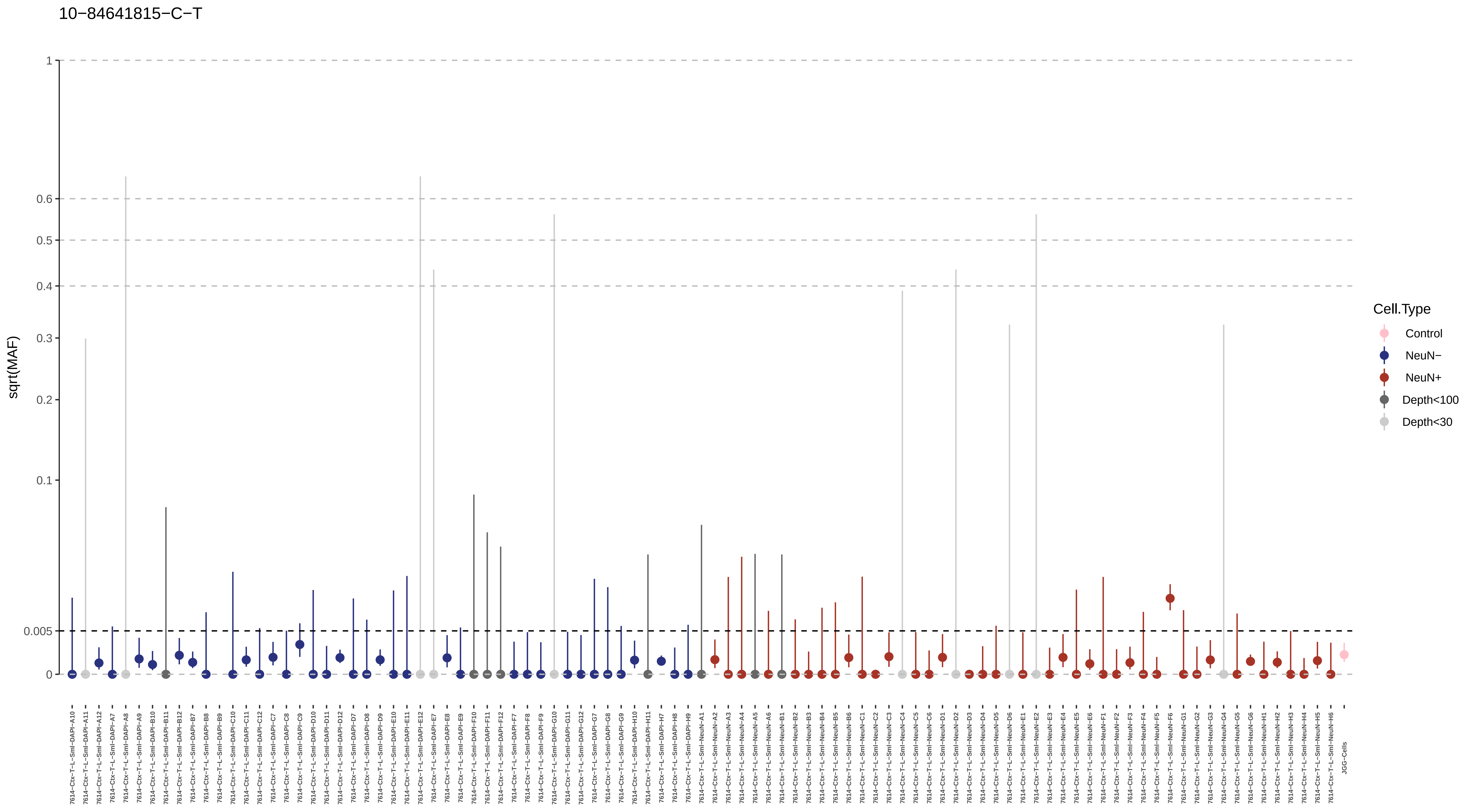

10−84641815−C−T
sqrt(MAF)
1
0.6
0.5
0.4
0.3
0.2
0.1
0.005
0
Cell.Type
Control
NeuN−
NeuN+
Depth<100
Depth<30

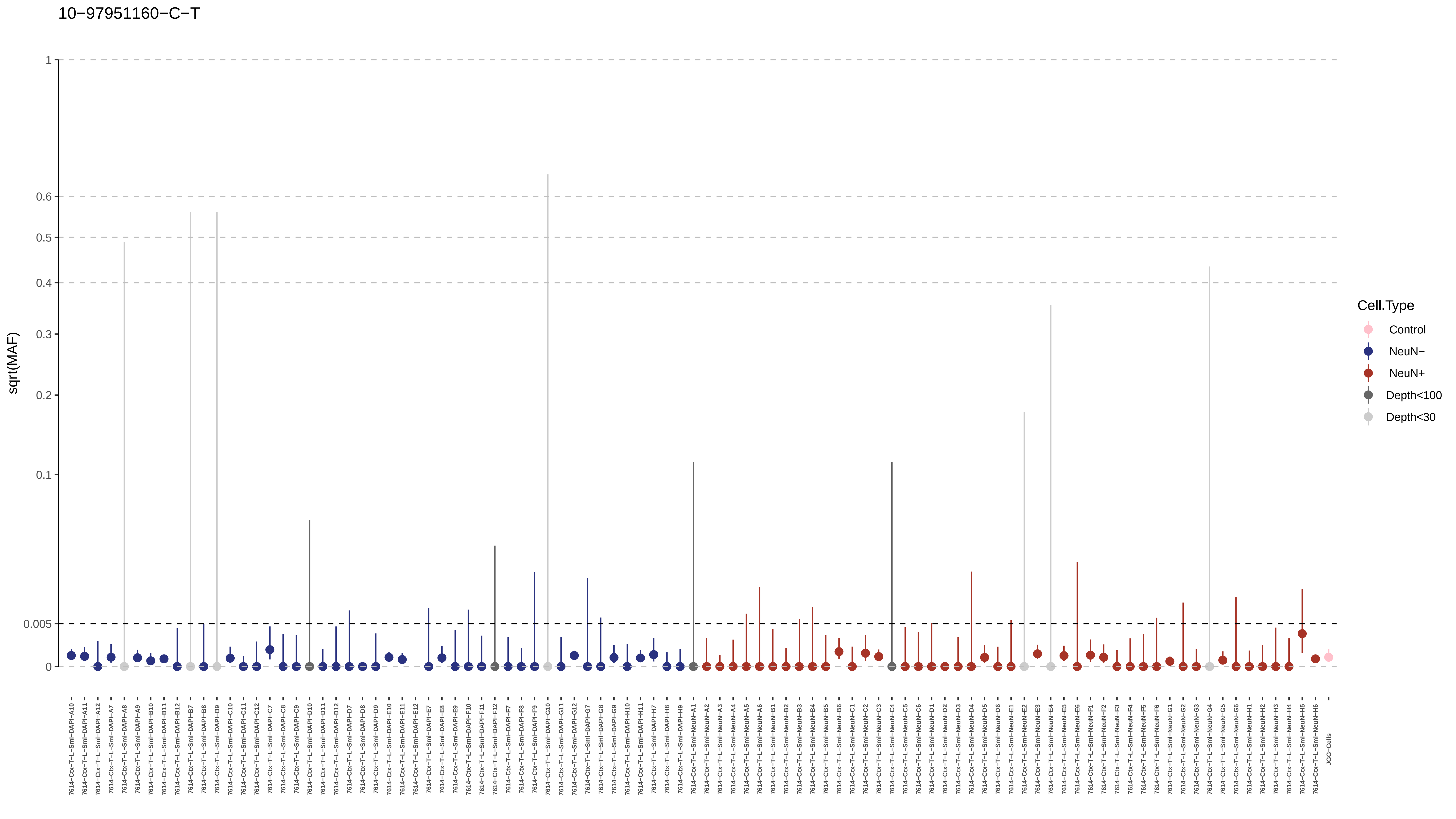
10−97951160−C−T
sqrt(MAF)
1
0.6
0.5
0.4
0.3
0.2
0.1
0.005
0
Cell.Type
Control
NeuN−
NeuN+
Depth<100
Depth<30

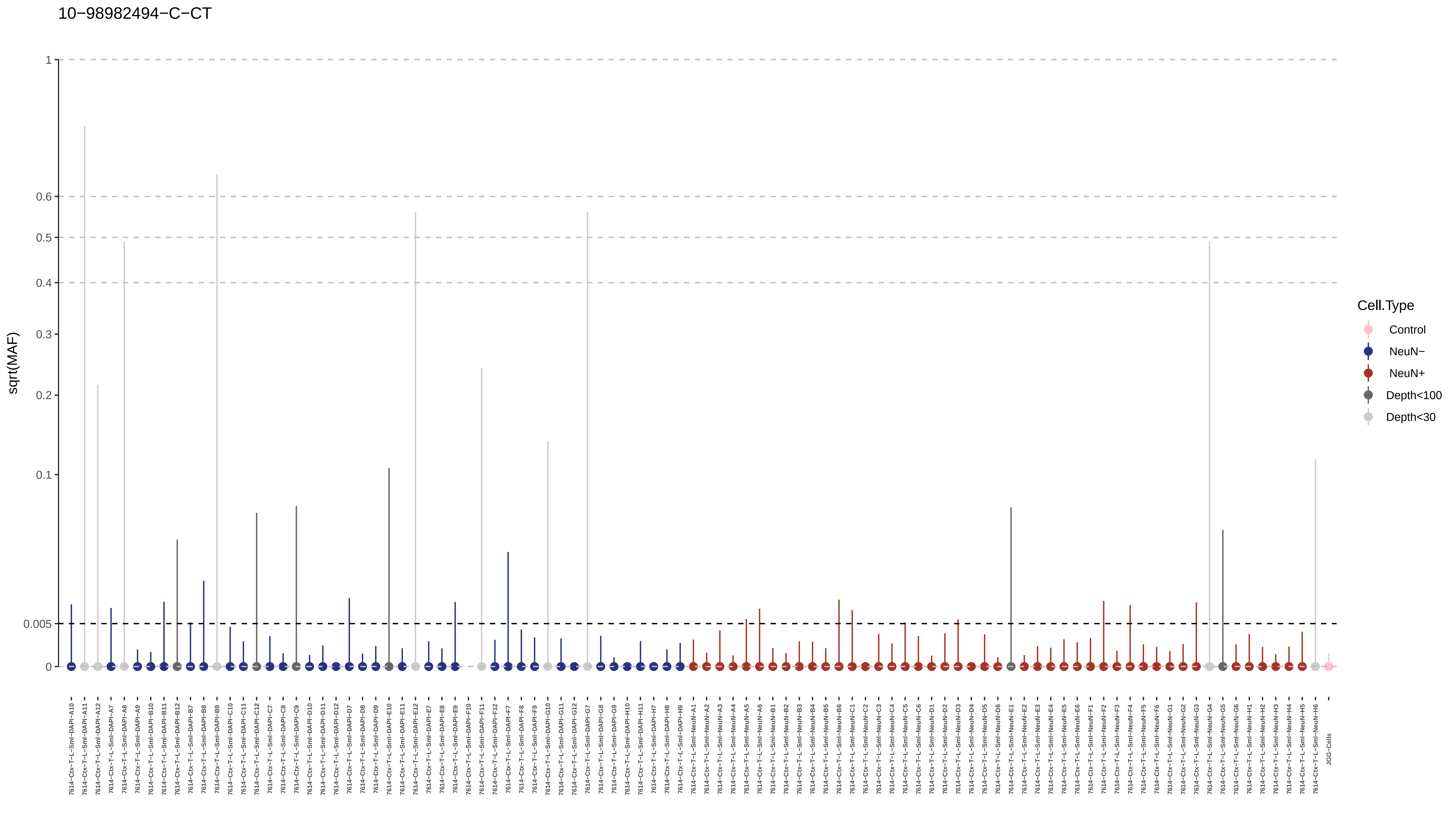
10−98982494−C−CT
sqrt(MAF)
1
0.6
0.5
0.4
0.3
0.2
0.1
0.005
0
Cell.Type
Control
NeuN−
NeuN+
Depth<100
Depth<30
JGG-Cells

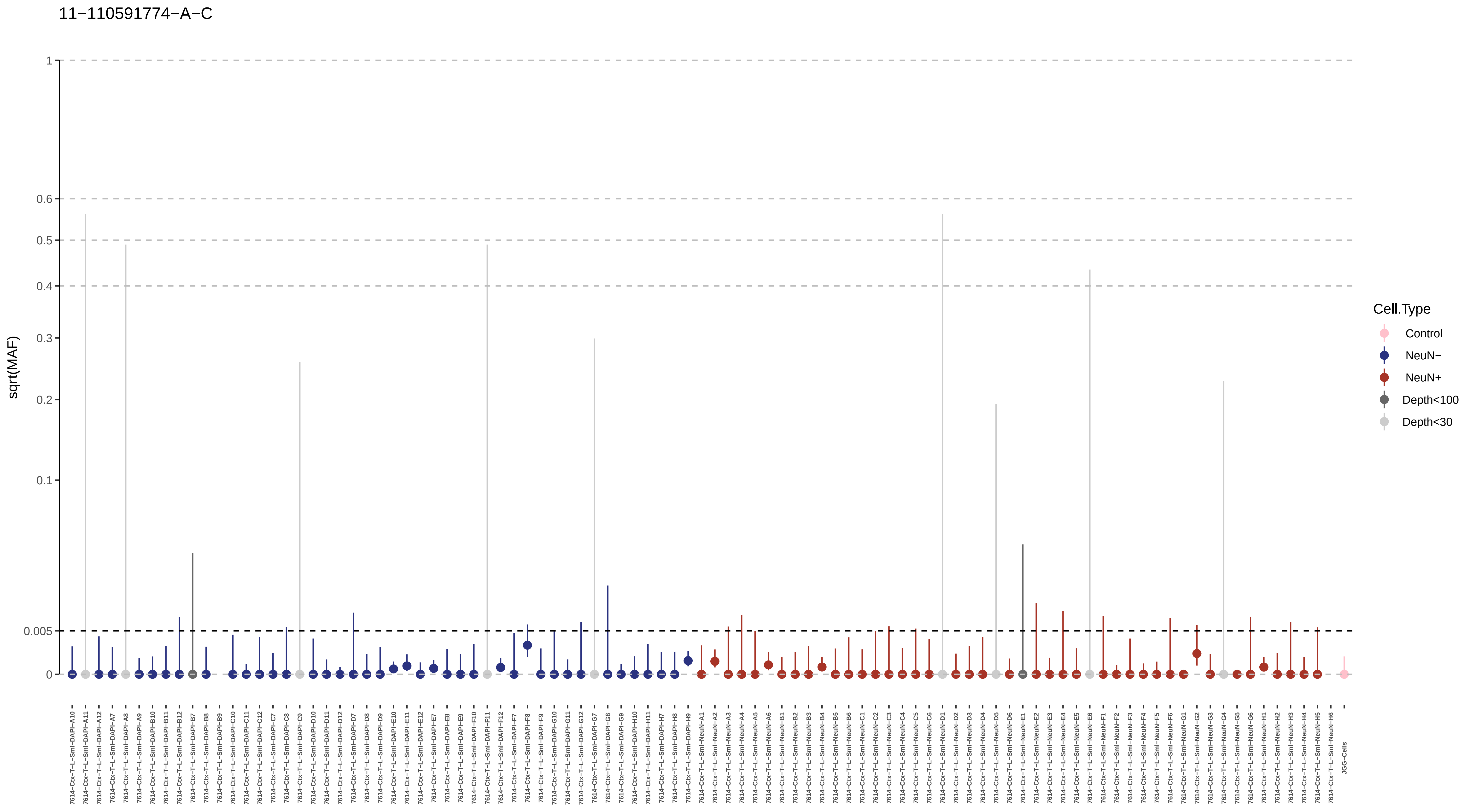
11−110591774−A−C
sqrt(MAF)
1
0.6
0.5
0.4
0.3
0.2
0.1
0.005
0
Cell.Type
Control
NeuN−
NeuN+
Depth<100
Depth<30
JGG-Cells

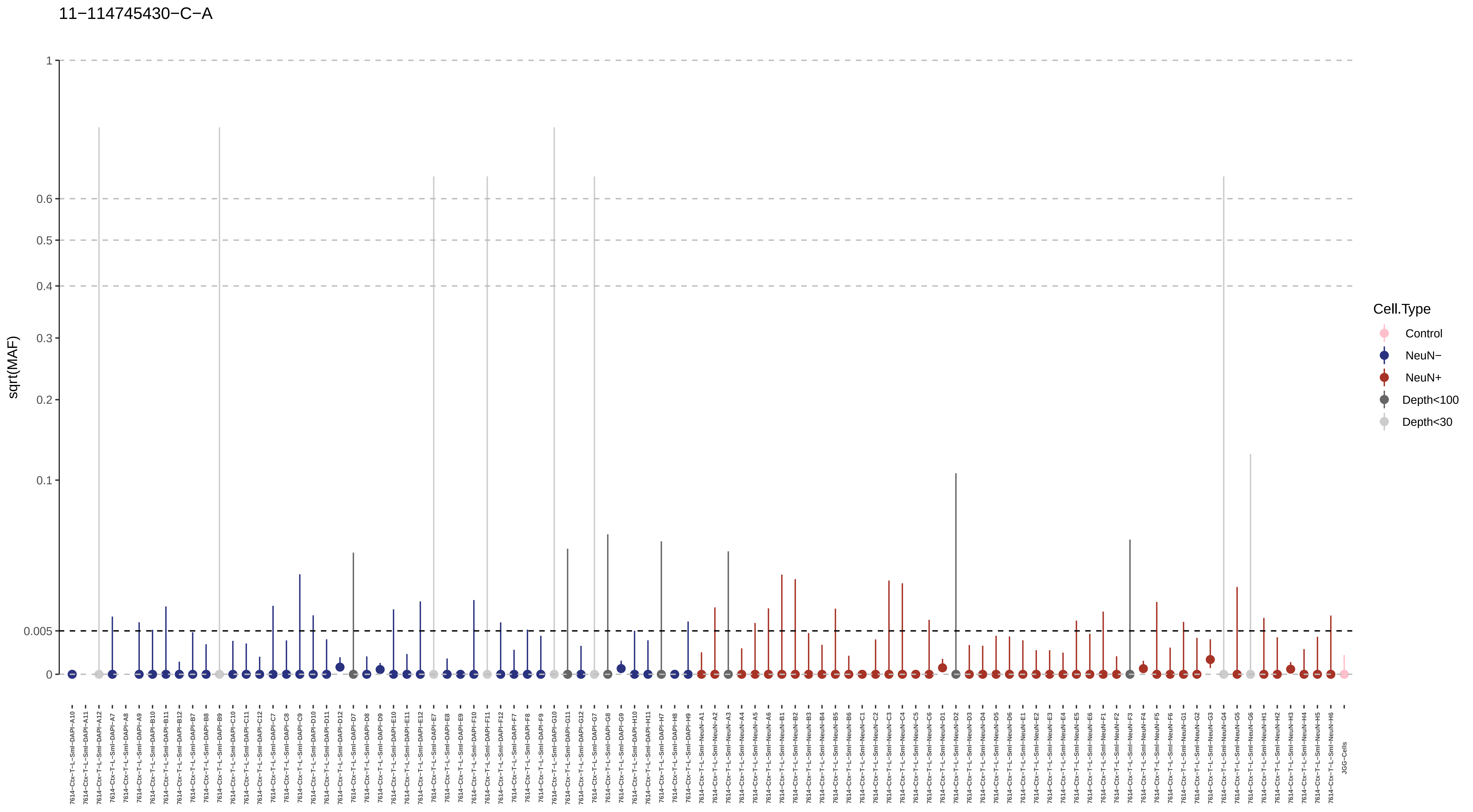

11−114745430−C−A
sqrt(MAF)
1
0.6
0.5
0.4
0.3
0.2
0.1
0.005
0
Cell.Type
Control
NeuN−
NeuN+
Depth<100
Depth<30

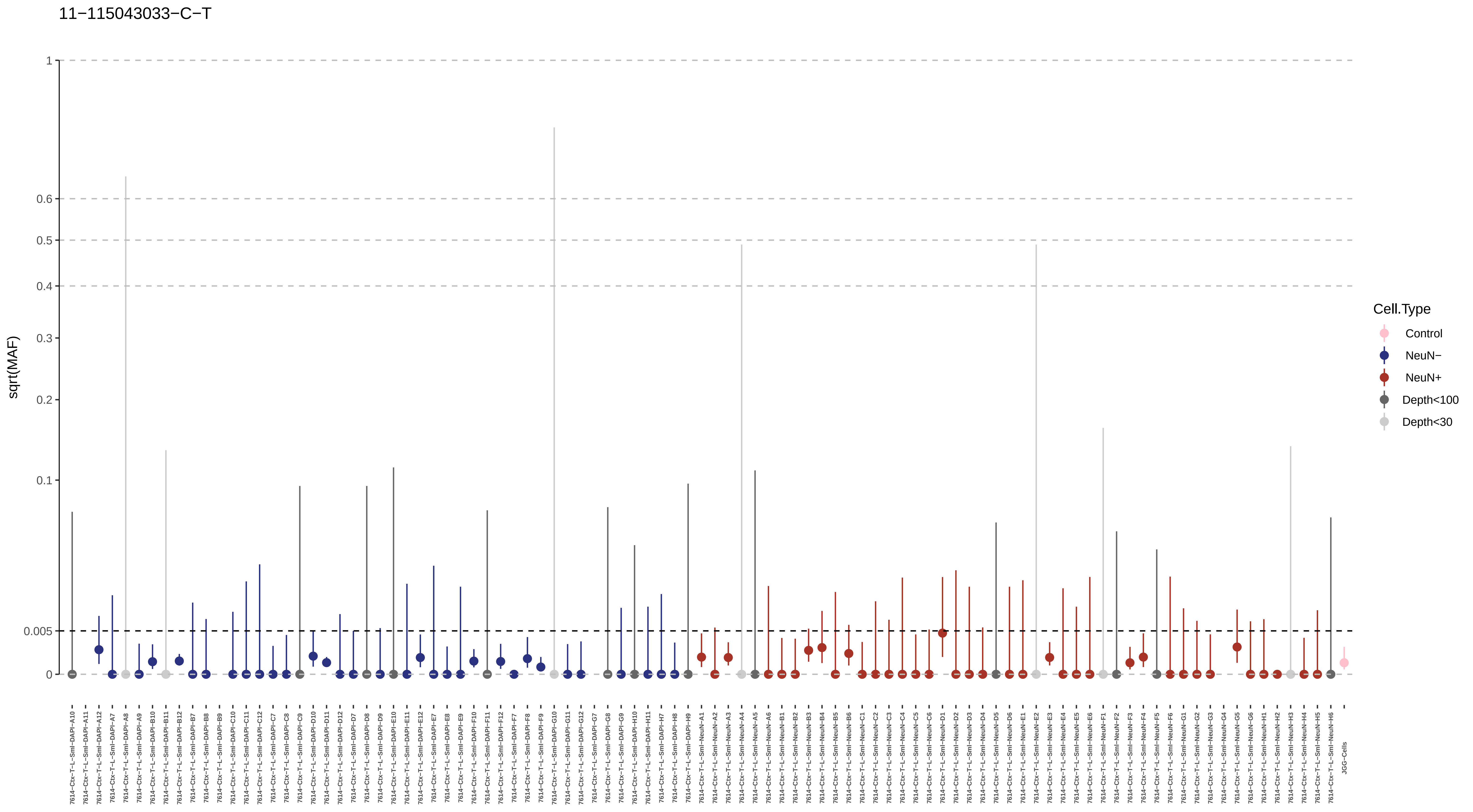

11−115043033−C−T
sqrt(MAF)
1
0.6
0.5
0.4
0.3
0.2
0.1
0.005
0
Cell.Type
Control
NeuN−
NeuN+
Depth<100
Depth<30
7614−Ctx−T−L−Sml−DAPI−A10
7614−Ctx−T−L−Sml−DAPI−A11
7614−Ctx−T−L−Sml−DAPI−A12
7614−Ctx−T−L−Sml−DAPI−A7
7614−Ctx−T−L−Sml−DAPI−A8
7614−Ctx−T−L−Sml−DAPI−A9
7614−Ctx−T−L−Sml−DAPI−B10
7614−Ctx−T−L−Sml−DAPI−B11
7614−Ctx−T−L−Sml−DAPI−B12
7614−Ctx−T−L−Sml−DAPI−B7
7614−Ctx−T−L−Sml−DAPI−B8
7614−Ctx−T−L−Sml−DAPI−B9
7614−Ctx−T−L−Sml−DAPI−C10
7614−Ctx−T−L−Sml−DAPI−C11
7614−Ctx−T−L−Sml−DAPI−C12
7614−Ctx−T−L−Sml−DAPI−C7
7614−Ctx−T−L−Sml−DAPI−C8
7614−Ctx−T−L−Sml−DAPI−C9
7614−Ctx−T−L−Sml−DAPI−D10
7614−Ctx−T−L−Sml−DAPI−D11
7614−Ctx−T−L−Sml−DAPI−D12
7614−Ctx−T−L−Sml−DAPI−D7
7614−Ctx−T−L−Sml−DAPI−D8
7614−Ctx−T−L−Sml−DAPI−D9
7614−Ctx−T−L−Sml−DAPI−E10
7614−Ctx−T−L−Sml−DAPI−E11
7614−Ctx−T−L−Sml−DAPI−E12
7614−Ctx−T−L−Sml−DAPI−E7
7614−Ctx−T−L−Sml−DAPI−E8
7614−Ctx−T−L−Sml−DAPI−E9
7614−Ctx−T−L−Sml−DAPI−F10
7614−Ctx−T−L−Sml−DAPI−F11
7614−Ctx−T−L−Sml−DAPI−F12
7614−Ctx−T−L−Sml−DAPI−F7
7614−Ctx−T−L−Sml−DAPI−F8
7614−Ctx−T−L−Sml−DAPI−F9
7614−Ctx−T−L−Sml−DAPI−G10
7614−Ctx−T−L−Sml−DAPI−G11
7614−Ctx−T−L−Sml−DAPI−G12
7614−Ctx−T−L−Sml−DAPI−G7
7614−Ctx−T−L−Sml−DAPI−G8
7614−Ctx−T−L−Sml−DAPI−G9
7614−Ctx−T−L−Sml−DAPI−H10
7614−Ctx−T−L−Sml−DAPI−H11
7614−Ctx−T−L−Sml−DAPI−H7
7614−Ctx−T−L−Sml−DAPI−H8
7614−Ctx−T−L−Sml−DAPI−H9
7614−Ctx−T−L−Sml−NeuN−A1
7614−Ctx−T−L−Sml−NeuN−A2
7614−Ctx−T−L−Sml−NeuN−A3
7614−Ctx−T−L−Sml−NeuN−A4
7614−Ctx−T−L−Sml−NeuN−A5
7614−Ctx−T−L−Sml−NeuN−A6
7614−Ctx−T−L−Sml−NeuN−B1
7614−Ctx−T−L−Sml−NeuN−B2
7614−Ctx−T−L−Sml−NeuN−B3
7614−Ctx−T−L−Sml−NeuN−B4
7614−Ctx−T−L−Sml−NeuN−B5
7614−Ctx−T−L−Sml−NeuN−B6
7614−Ctx−T−L−Sml−NeuN−C1
7614−Ctx−T−L−Sml−NeuN−C2
7614−Ctx−T−L−Sml−NeuN−C3
7614−Ctx−T−L−Sml−NeuN−C4
7614−Ctx−T−L−Sml−NeuN−C5
7614−Ctx−T−L−Sml−NeuN−C6
7614−Ctx−T−L−Sml−NeuN−D1
7614−Ctx−T−L−Sml−NeuN−D2
7614−Ctx−T−L−Sml−NeuN−D3
7614−Ctx−T−L−Sml−NeuN−D4
7614−Ctx−T−L−Sml−NeuN−D5
7614−Ctx−T−L−Sml−NeuN−D6
7614−Ctx−T−L−Sml−NeuN−E1
7614−Ctx−T−L−Sml−NeuN−E2
7614−Ctx−T−L−Sml−NeuN−E3
7614−Ctx−T−L−Sml−NeuN−E4
7614−Ctx−T−L−Sml−NeuN−E5
7614−Ctx−T−L−Sml−NeuN−E6
7614−Ctx−T−L−Sml−NeuN−F1
7614−Ctx−T−L−Sml−NeuN−F2
7614−Ctx−T−L−Sml−NeuN−F3
7614−Ctx−T−L−Sml−NeuN−F4
7614−Ctx−T−L−Sml−NeuN−F5
7614−Ctx−T−L−Sml−NeuN−F6
7614−Ctx−T−L−Sml−NeuN−G1
7614−Ctx−T−L−Sml−NeuN−G2
7614−Ctx−T−L−Sml−NeuN−G3
7614−Ctx−T−L−Sml−NeuN−G4
7614−Ctx−T−L−Sml−NeuN−G5
7614−Ctx−T−L−Sml−NeuN−G6
7614−Ctx−T−L−Sml−NeuN−H1
7614−Ctx−T−L−Sml−NeuN−H2
7614−Ctx−T−L−Sml−NeuN−H3
7614−Ctx−T−L−Sml−NeuN−H4
7614−Ctx−T−L−Sml−NeuN−H5
7614−Ctx−T−L−Sml−NeuN−H6
JGG-Cells

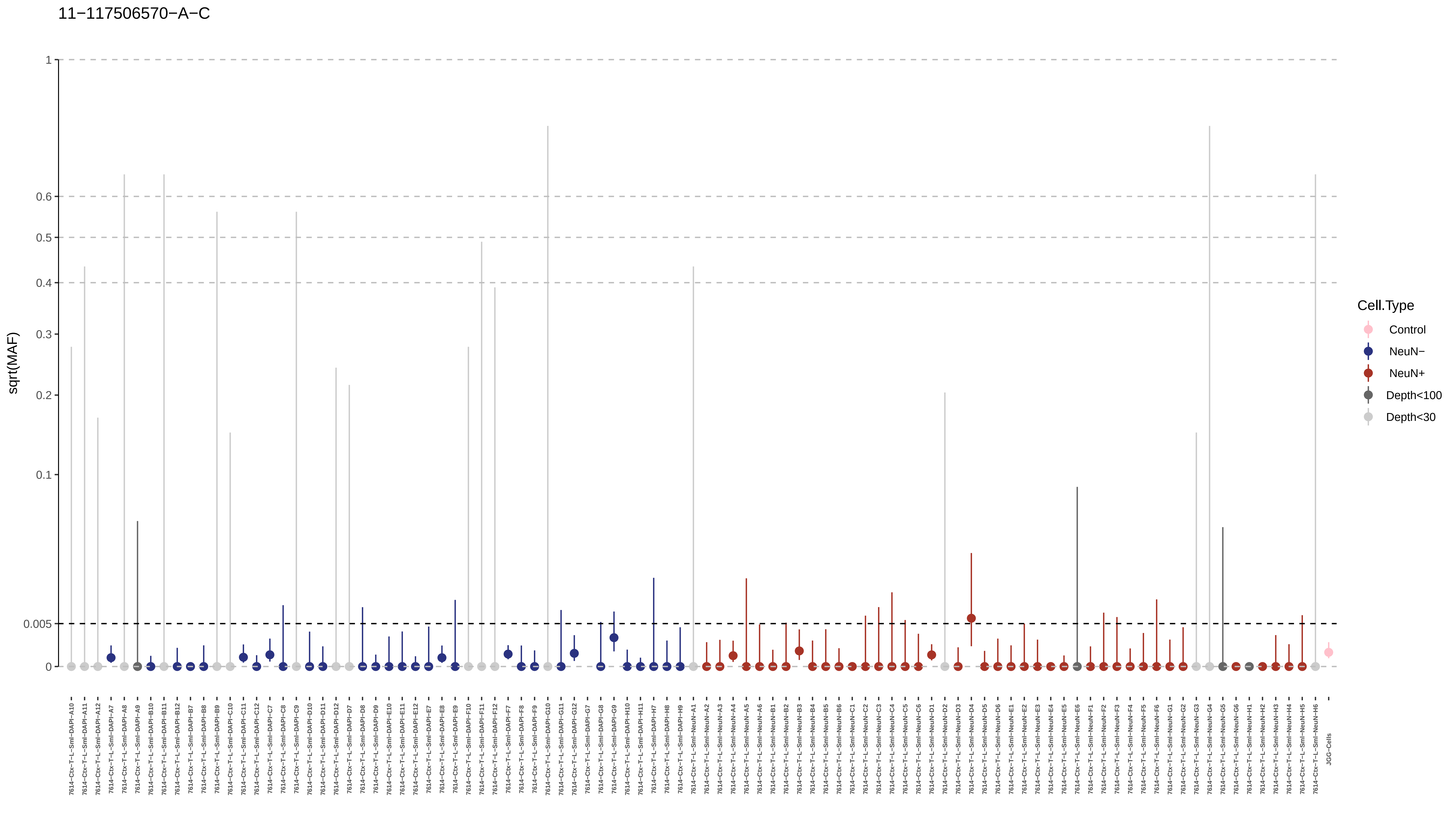

11−117506570−A−C
sqrt(MAF)
1
0.6
0.5
0.4
0.3
0.2
0.1
0.005
0
Cell.Type
Control
NeuN−
NeuN+
Depth<100
Depth<30
JGG-Cells

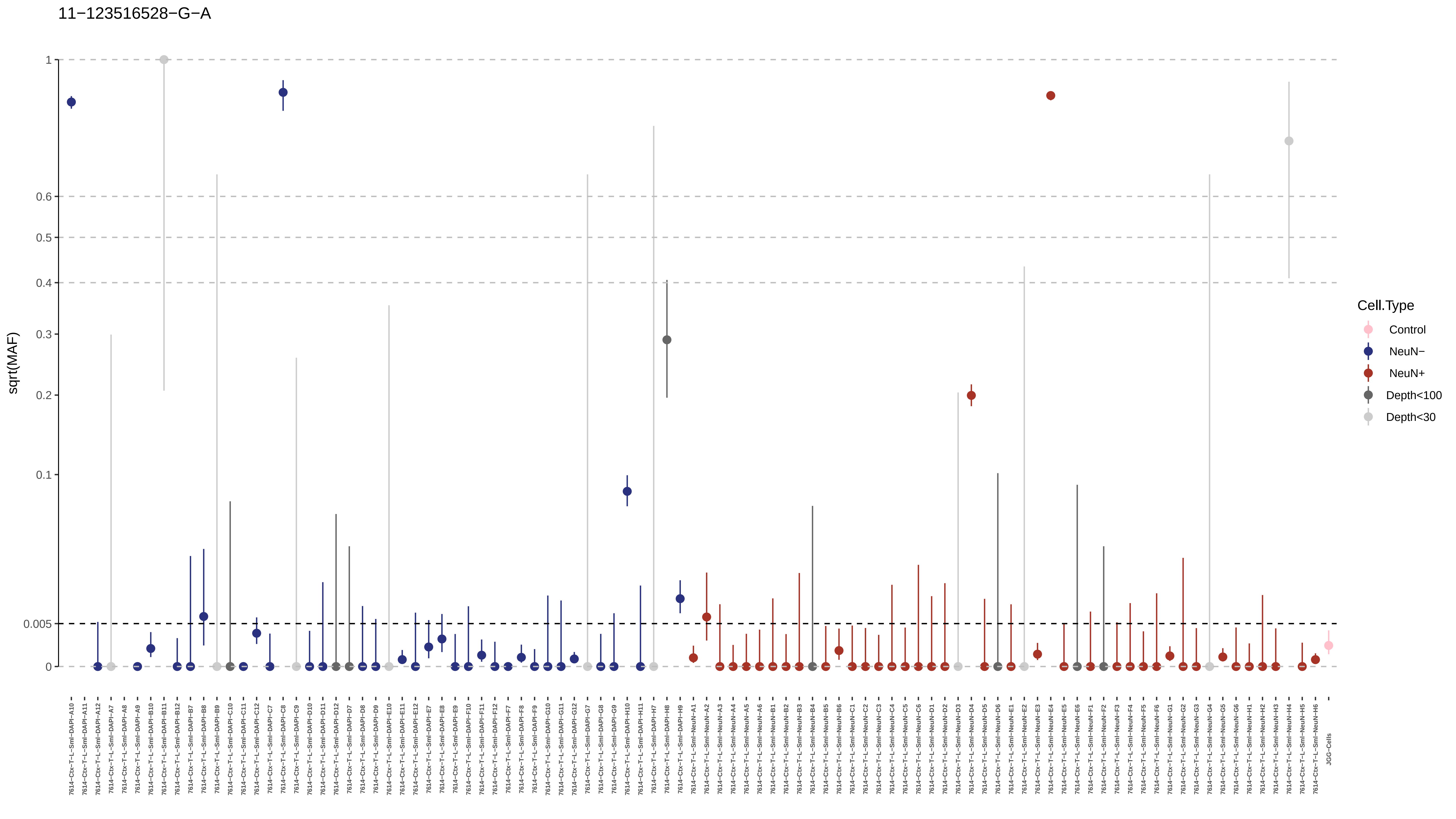

11−123516528−G−A
sqrt(MAF)
1
0.6
0.5
0.4
0.3
0.2
0.1
0.005
0
Cell.Type
Control
NeuN−
NeuN+
Depth<100
Depth<30
JGG-Cells

11−129560658−C−T

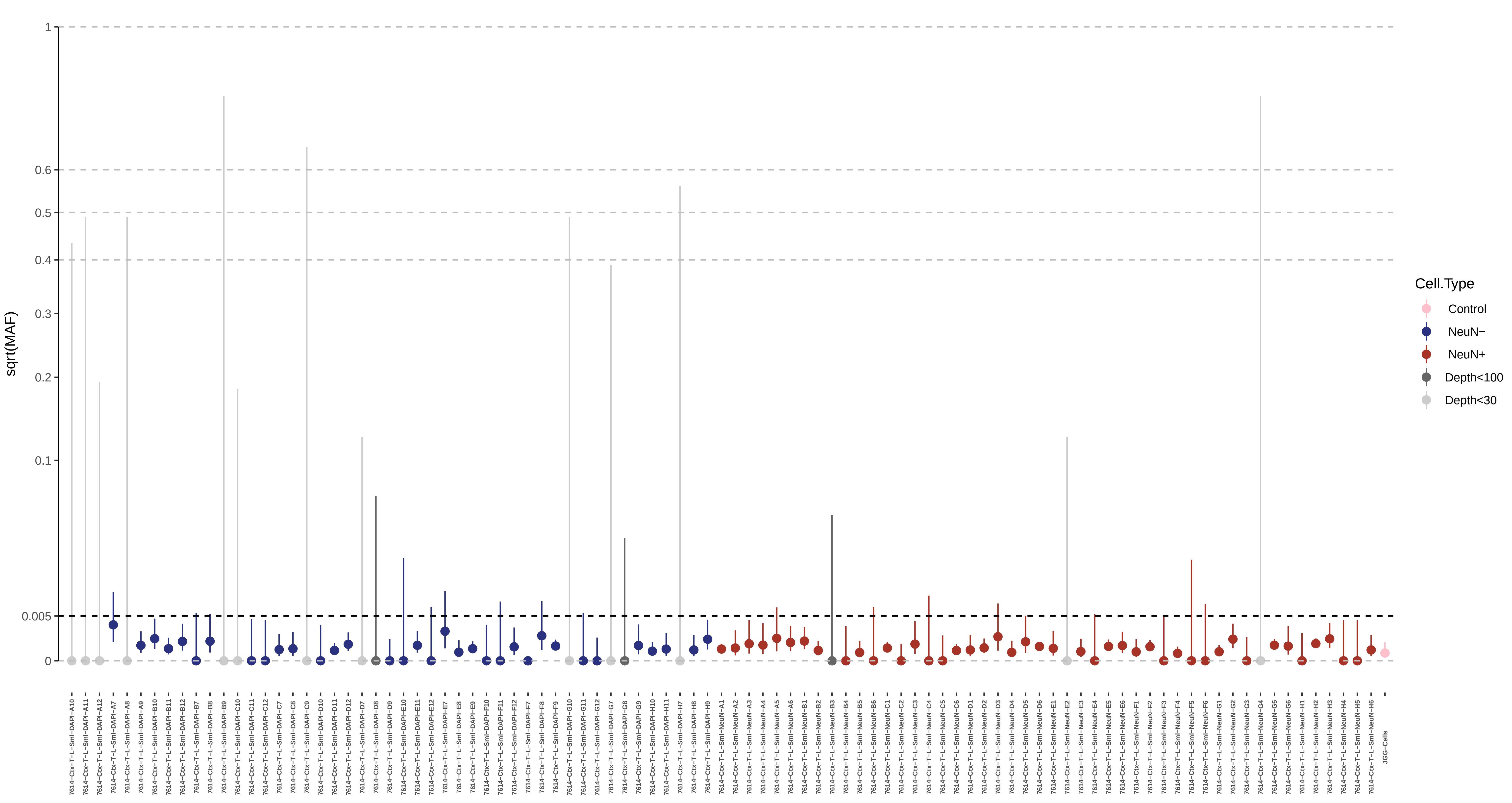

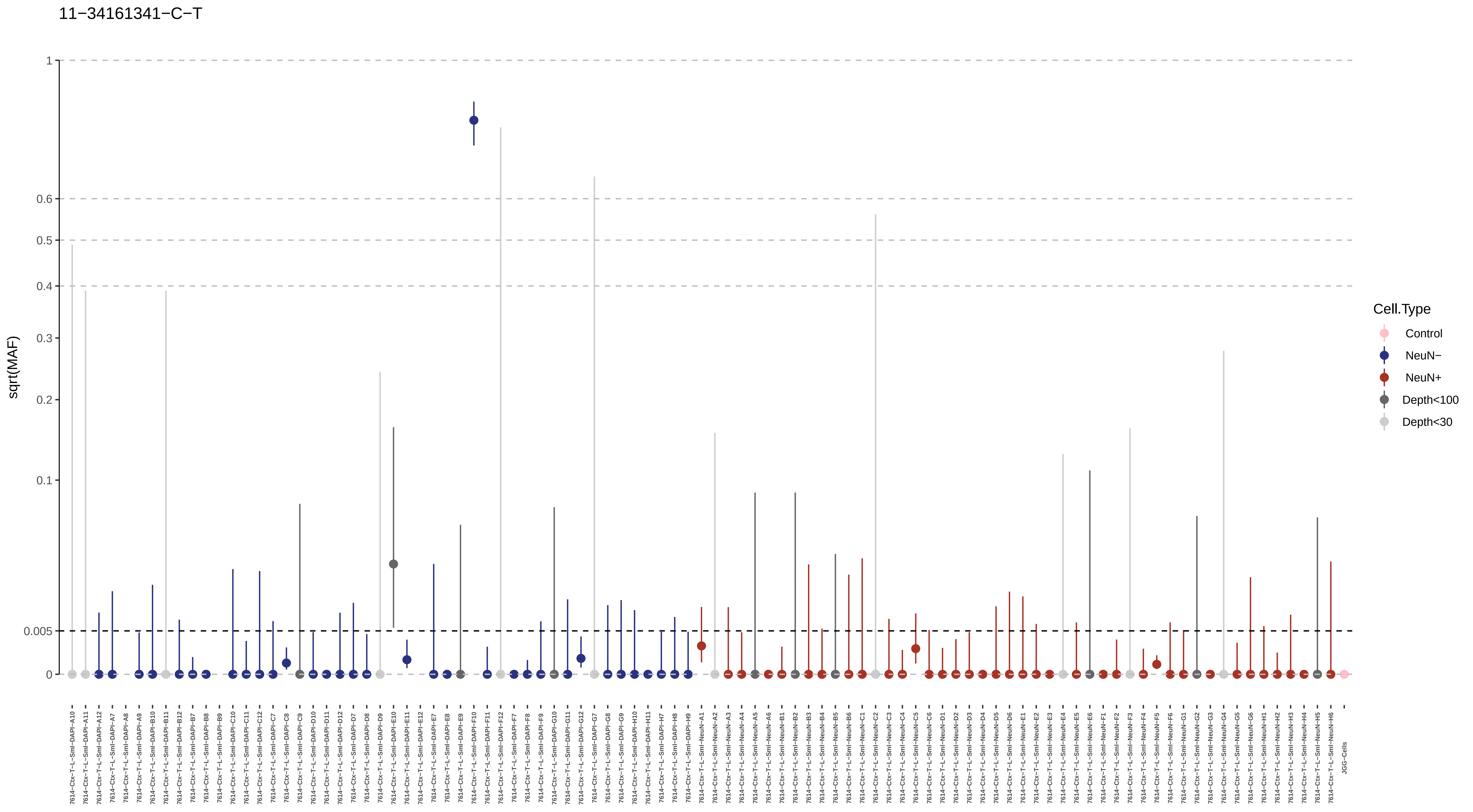
11−34161341−C−T
sqrt(MAF)
1
0.6
0.5
0.4
0.3
0.2
0.1
0.005
0
Cell.Type
Control
NeuN−
NeuN+
Depth<100
Depth<30

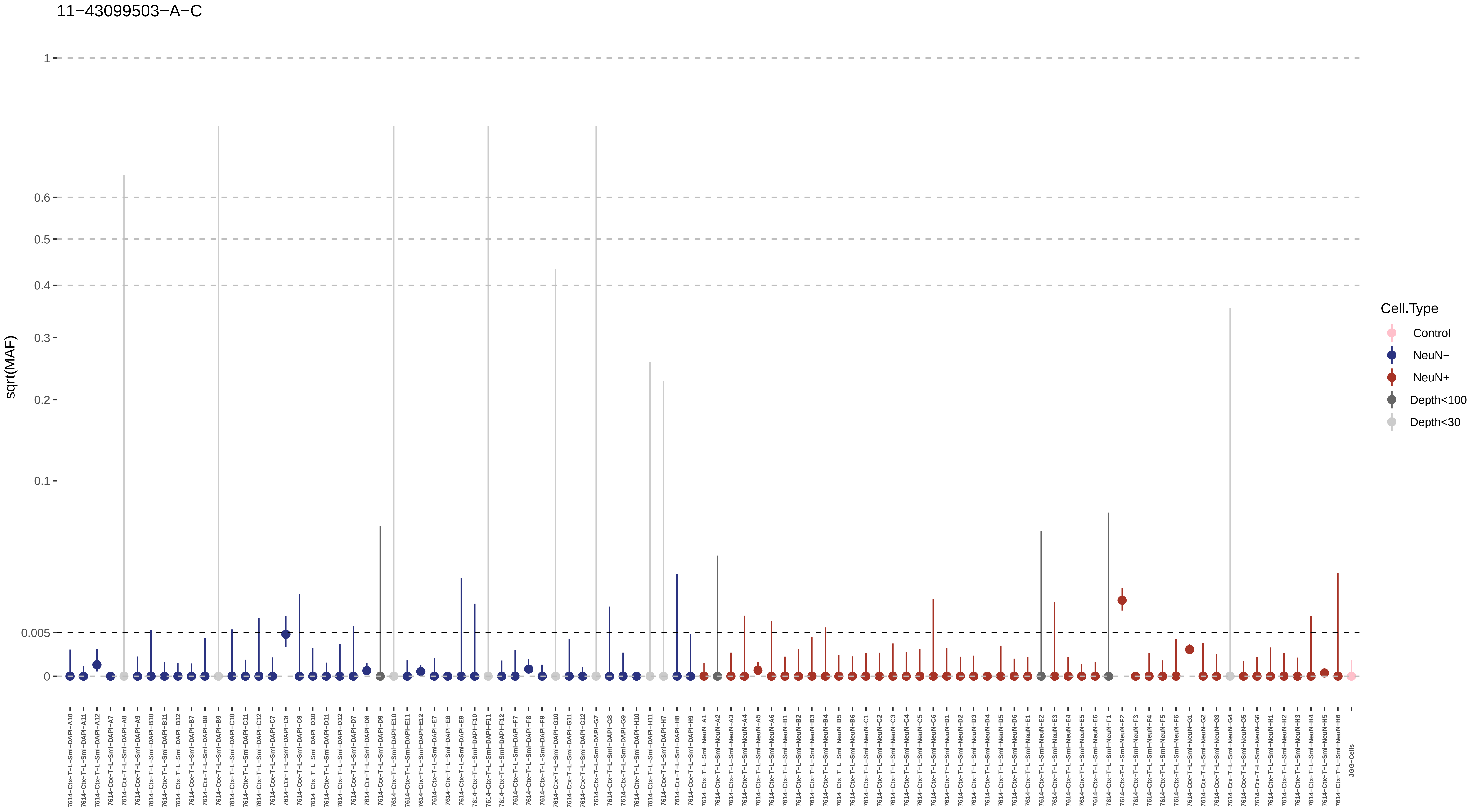

11−43099503−A−C
sqrt(MAF)
1
0.6
0.5
0.4
0.3
0.2
0.1
0.005
0
Cell.Type
Control
NeuN−
NeuN+
Depth<100
Depth<30
7614−Ctx−T−L−Sml−DAPI−A10
7614−Ctx−T−L−Sml−DAPI−A11
7614−Ctx−T−L−Sml−DAPI−A12
7614−Ctx−T−L−Sml−DAPI−A7
7614−Ctx−T−L−Sml−DAPI−A8
7614−Ctx−T−L−Sml−DAPI−A9
7614−Ctx−T−L−Sml−DAPI−B10
7614−Ctx−T−L−Sml−DAPI−B11
7614−Ctx−T−L−Sml−DAPI−B12
7614−Ctx−T−L−Sml−DAPI−B7
7614−Ctx−T−L−Sml−DAPI−B8
7614−Ctx−T−L−Sml−DAPI−B9
7614−Ctx−T−L−Sml−DAPI−C10
7614−Ctx−T−L−Sml−DAPI−C11
7614−Ctx−T−L−Sml−DAPI−C12
7614−Ctx−T−L−Sml−DAPI−C7
7614−Ctx−T−L−Sml−DAPI−C8
7614−Ctx−T−L−Sml−DAPI−C9
7614−Ctx−T−L−Sml−DAPI−D10
7614−Ctx−T−L−Sml−DAPI−D11
7614−Ctx−T−L−Sml−DAPI−D12
7614−Ctx−T−L−Sml−DAPI−D7
7614−Ctx−T−L−Sml−DAPI−D8
7614−Ctx−T−L−Sml−DAPI−D9
7614−Ctx−T−L−Sml−DAPI−E10
7614−Ctx−T−L−Sml−DAPI−E11
7614−Ctx−T−L−Sml−DAPI−E12
7614−Ctx−T−L−Sml−DAPI−E7
7614−Ctx−T−L−Sml−DAPI−E8
7614−Ctx−T−L−Sml−DAPI−E9
7614−Ctx−T−L−Sml−DAPI−F10
7614−Ctx−T−L−Sml−DAPI−F11
7614−Ctx−T−L−Sml−DAPI−F12
7614−Ctx−T−L−Sml−DAPI−F7
7614−Ctx−T−L−Sml−DAPI−F8
7614−Ctx−T−L−Sml−DAPI−F9
7614−Ctx−T−L−Sml−DAPI−G10
7614−Ctx−T−L−Sml−DAPI−G11
7614−Ctx−T−L−Sml−DAPI−G12
7614−Ctx−T−L−Sml−DAPI−G7
7614−Ctx−T−L−Sml−DAPI−G8
7614−Ctx−T−L−Sml−DAPI−G9
7614−Ctx−T−L−Sml−DAPI−H10
7614−Ctx−T−L−Sml−DAPI−H11
7614−Ctx−T−L−Sml−DAPI−H7
7614−Ctx−T−L−Sml−DAPI−H8
7614−Ctx−T−L−Sml−DAPI−H9
7614−Ctx−T−L−Sml−NeuN−A1
7614−Ctx−T−L−Sml−NeuN−A2
7614−Ctx−T−L−Sml−NeuN−A3
7614−Ctx−T−L−Sml−NeuN−A4
7614−Ctx−T−L−Sml−NeuN−A5
7614−Ctx−T−L−Sml−NeuN−A6
7614−Ctx−T−L−Sml−NeuN−B1
7614−Ctx−T−L−Sml−NeuN−B2
7614−Ctx−T−L−Sml−NeuN−B3
7614−Ctx−T−L−Sml−NeuN−B4
7614−Ctx−T−L−Sml−NeuN−B5
7614−Ctx−T−L−Sml−NeuN−B6
7614−Ctx−T−L−Sml−NeuN−C1
7614−Ctx−T−L−Sml−NeuN−C2
7614−Ctx−T−L−Sml−NeuN−C3
7614−Ctx−T−L−Sml−NeuN−C4
7614−Ctx−T−L−Sml−NeuN−C5
7614−Ctx−T−L−Sml−NeuN−C6
7614−Ctx−T−L−Sml−NeuN−D1
7614−Ctx−T−L−Sml−NeuN−D2
7614−Ctx−T−L−Sml−NeuN−D3
7614−Ctx−T−L−Sml−NeuN−D4
7614−Ctx−T−L−Sml−NeuN−D5
7614−Ctx−T−L−Sml−NeuN−D6
7614−Ctx−T−L−Sml−NeuN−E1
7614−Ctx−T−L−Sml−NeuN−E2
7614−Ctx−T−L−Sml−NeuN−E3
7614−Ctx−T−L−Sml−NeuN−E4
7614−Ctx−T−L−Sml−NeuN−E5
7614−Ctx−T−L−Sml−NeuN−E6
7614−Ctx−T−L−Sml−NeuN−F1
7614−Ctx−T−L−Sml−NeuN−F2
7614−Ctx−T−L−Sml−NeuN−F3
7614−Ctx−T−L−Sml−NeuN−F4
7614−Ctx−T−L−Sml−NeuN−F5
7614−Ctx−T−L−Sml−NeuN−F6
7614−Ctx−T−L−Sml−NeuN−G1
7614−Ctx−T−L−Sml−NeuN−G2
7614−Ctx−T−L−Sml−NeuN−G3
7614−Ctx−T−L−Sml−NeuN−G4
7614−Ctx−T−L−Sml−NeuN−G5
7614−Ctx−T−L−Sml−NeuN−G6
7614−Ctx−T−L−Sml−NeuN−H1
7614−Ctx−T−L−Sml−NeuN−H2
7614−Ctx−T−L−Sml−NeuN−H3
7614−Ctx−T−L−Sml−NeuN−H4
7614−Ctx−T−L−Sml−NeuN−H5
7614−Ctx−T−L−Sml−NeuN−H6
JGG-Cells

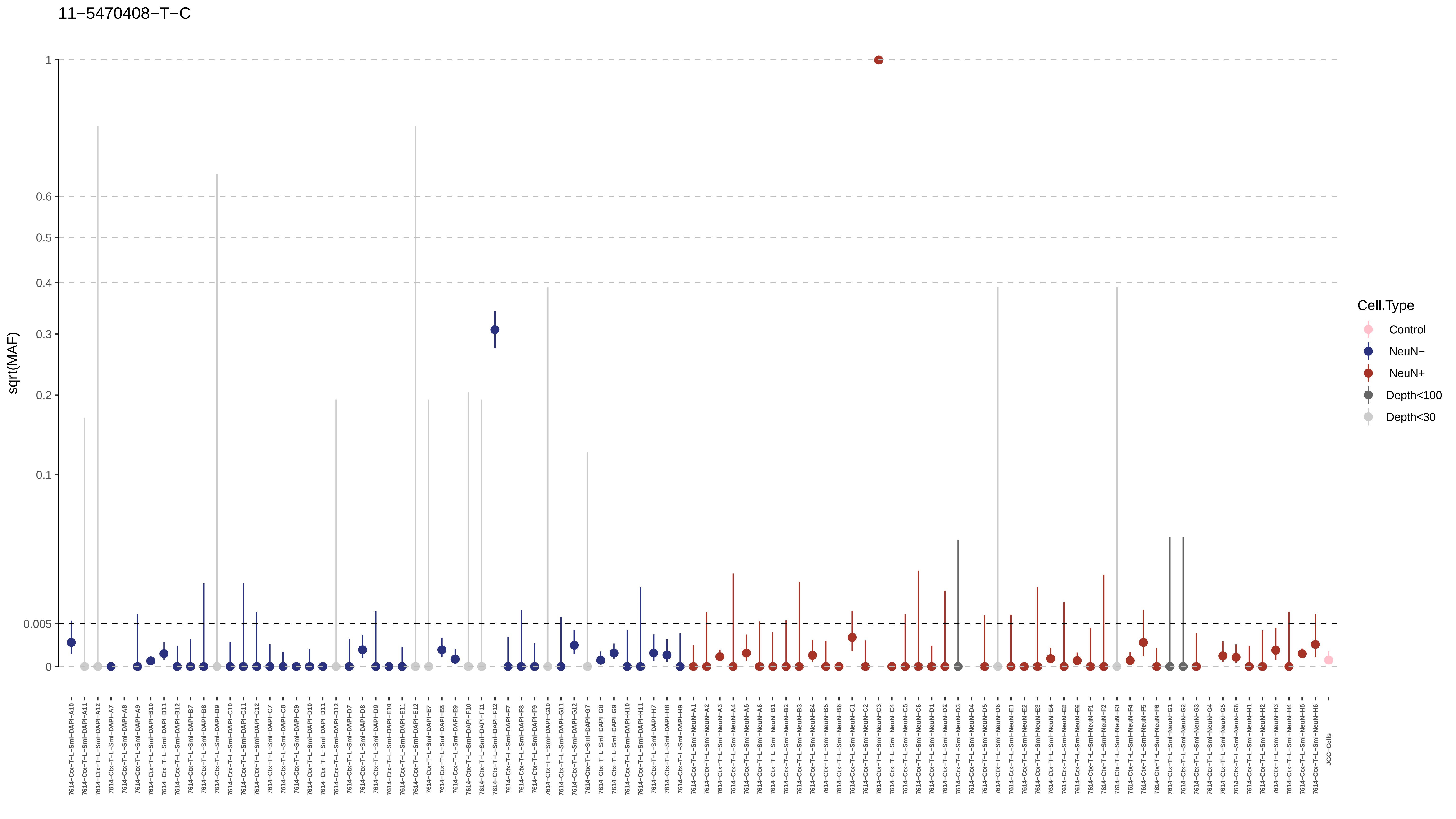
11−5470408−T−C
sqrt(MAF)
1
0.6
0.5
0.4
0.3
0.2
0.1
0.005
0
Cell.Type
Control
NeuN−
NeuN+
Depth<100
Depth<30

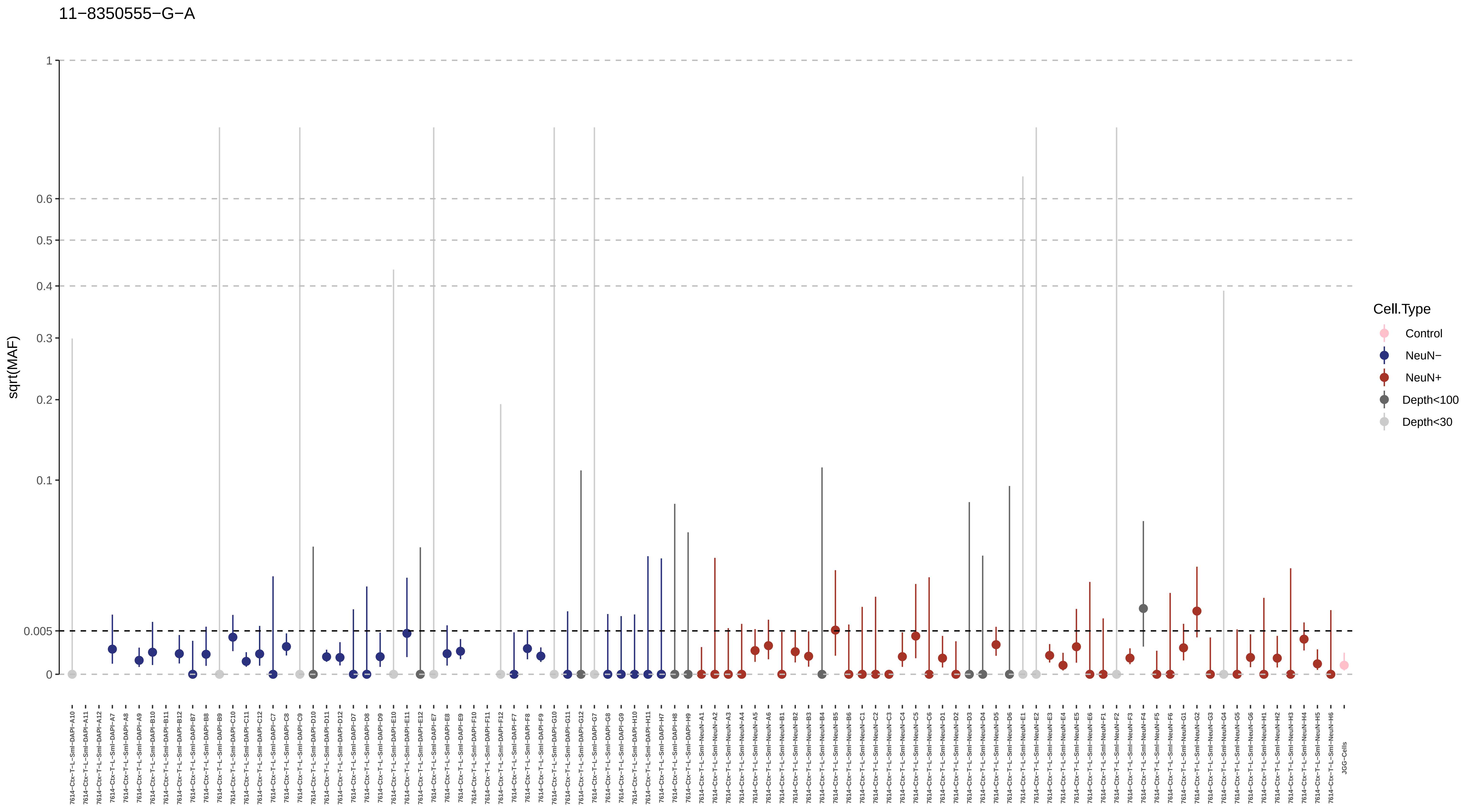

11−8350555−G−A
sqrt(MAF)
1
0.6
0.5
0.4
0.3
0.2
0.1
0.005
0
Cell.Type
Control
NeuN−
NeuN+
Depth<100
Depth<30
7614−Ctx−T−L−Sml−DAPI−A10
7614−Ctx−T−L−Sml−DAPI−A11
7614−Ctx−T−L−Sml−DAPI−A12
7614−Ctx−T−L−Sml−DAPI−A7
7614−Ctx−T−L−Sml−DAPI−A8
7614−Ctx−T−L−Sml−DAPI−A9
7614−Ctx−T−L−Sml−DAPI−B10
7614−Ctx−T−L−Sml−DAPI−B11
7614−Ctx−T−L−Sml−DAPI−B12
7614−Ctx−T−L−Sml−DAPI−B7
7614−Ctx−T−L−Sml−DAPI−B8
7614−Ctx−T−L−Sml−DAPI−B9
7614−Ctx−T−L−Sml−DAPI−C10
7614−Ctx−T−L−Sml−DAPI−C11
7614−Ctx−T−L−Sml−DAPI−C12
7614−Ctx−T−L−Sml−DAPI−C7
7614−Ctx−T−L−Sml−DAPI−C8
7614−Ctx−T−L−Sml−DAPI−C9
7614−Ctx−T−L−Sml−DAPI−D10
7614−Ctx−T−L−Sml−DAPI−D11
7614−Ctx−T−L−Sml−DAPI−D12
7614−Ctx−T−L−Sml−DAPI−D7
7614−Ctx−T−L−Sml−DAPI−D8
7614−Ctx−T−L−Sml−DAPI−D9
7614−Ctx−T−L−Sml−DAPI−E10
7614−Ctx−T−L−Sml−DAPI−E11
7614−Ctx−T−L−Sml−DAPI−E12
7614−Ctx−T−L−Sml−DAPI−E7
7614−Ctx−T−L−Sml−DAPI−E8
7614−Ctx−T−L−Sml−DAPI−E9
7614−Ctx−T−L−Sml−DAPI−F10
7614−Ctx−T−L−Sml−DAPI−F11
7614−Ctx−T−L−Sml−DAPI−F12
7614−Ctx−T−L−Sml−DAPI−F7
7614−Ctx−T−L−Sml−DAPI−F8
7614−Ctx−T−L−Sml−DAPI−F9
7614−Ctx−T−L−Sml−DAPI−G10
7614−Ctx−T−L−Sml−DAPI−G11
7614−Ctx−T−L−Sml−DAPI−G12
7614−Ctx−T−L−Sml−DAPI−G7
7614−Ctx−T−L−Sml−DAPI−G8
7614−Ctx−T−L−Sml−DAPI−G9
7614−Ctx−T−L−Sml−DAPI−H10
7614−Ctx−T−L−Sml−DAPI−H11
7614−Ctx−T−L−Sml−DAPI−H7
7614−Ctx−T−L−Sml−DAPI−H8
7614−Ctx−T−L−Sml−DAPI−H9
7614−Ctx−T−L−Sml−NeuN−A1
7614−Ctx−T−L−Sml−NeuN−A2
7614−Ctx−T−L−Sml−NeuN−A3
7614−Ctx−T−L−Sml−NeuN−A4
7614−Ctx−T−L−Sml−NeuN−A5
7614−Ctx−T−L−Sml−NeuN−A6
7614−Ctx−T−L−Sml−NeuN−B1
7614−Ctx−T−L−Sml−NeuN−B2
7614−Ctx−T−L−Sml−NeuN−B3
7614−Ctx−T−L−Sml−NeuN−B4
7614−Ctx−T−L−Sml−NeuN−B5
7614−Ctx−T−L−Sml−NeuN−B6
7614−Ctx−T−L−Sml−NeuN−C1
7614−Ctx−T−L−Sml−NeuN−C2
7614−Ctx−T−L−Sml−NeuN−C3
7614−Ctx−T−L−Sml−NeuN−C4
7614−Ctx−T−L−Sml−NeuN−C5
7614−Ctx−T−L−Sml−NeuN−C6
7614−Ctx−T−L−Sml−NeuN−D1
7614−Ctx−T−L−Sml−NeuN−D2
7614−Ctx−T−L−Sml−NeuN−D3
7614−Ctx−T−L−Sml−NeuN−D4
7614−Ctx−T−L−Sml−NeuN−D5
7614−Ctx−T−L−Sml−NeuN−D6
7614−Ctx−T−L−Sml−NeuN−E1
7614−Ctx−T−L−Sml−NeuN−E2
7614−Ctx−T−L−Sml−NeuN−E3
7614−Ctx−T−L−Sml−NeuN−E4
7614−Ctx−T−L−Sml−NeuN−E5
7614−Ctx−T−L−Sml−NeuN−E6
7614−Ctx−T−L−Sml−NeuN−F1
7614−Ctx−T−L−Sml−NeuN−F2
7614−Ctx−T−L−Sml−NeuN−F3
7614−Ctx−T−L−Sml−NeuN−F4
7614−Ctx−T−L−Sml−NeuN−F5
7614−Ctx−T−L−Sml−NeuN−F6
7614−Ctx−T−L−Sml−NeuN−G1
7614−Ctx−T−L−Sml−NeuN−G2
7614−Ctx−T−L−Sml−NeuN−G3
7614−Ctx−T−L−Sml−NeuN−G4
7614−Ctx−T−L−Sml−NeuN−G5
7614−Ctx−T−L−Sml−NeuN−G6
7614−Ctx−T−L−Sml−NeuN−H1
7614−Ctx−T−L−Sml−NeuN−H2
7614−Ctx−T−L−Sml−NeuN−H3
7614−Ctx−T−L−Sml−NeuN−H4
7614−Ctx−T−L−Sml−NeuN−H5
7614−Ctx−T−L−Sml−NeuN−H6
JGG−Cells

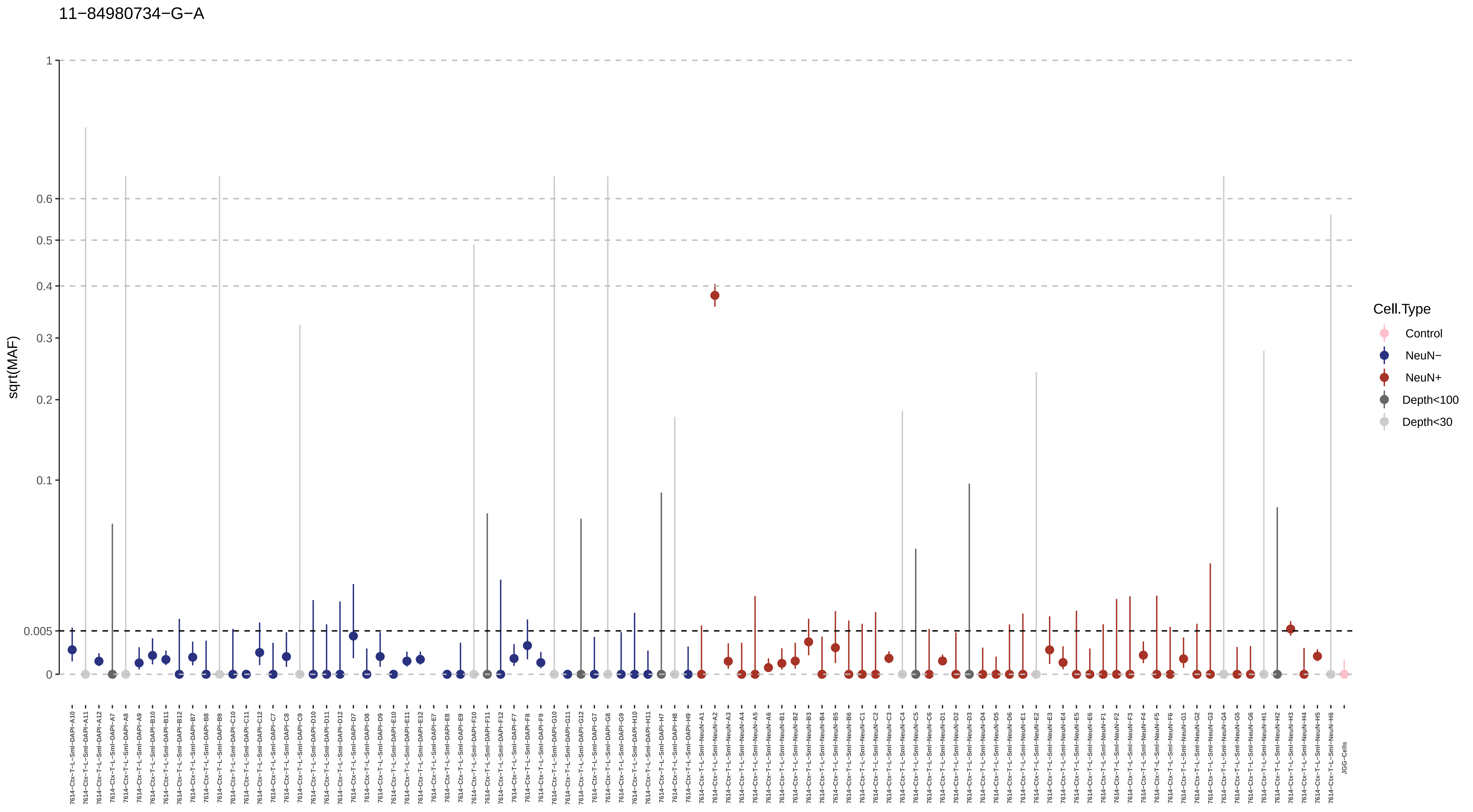

11−84980734−G−A
sqrt(MAF)
1
0.6
0.5
0.4
0.3
0.2
0.1
0.005
0
Cell.Type
Control
NeuN−
NeuN+
Depth<100
Depth<30

11−88087255−A−C

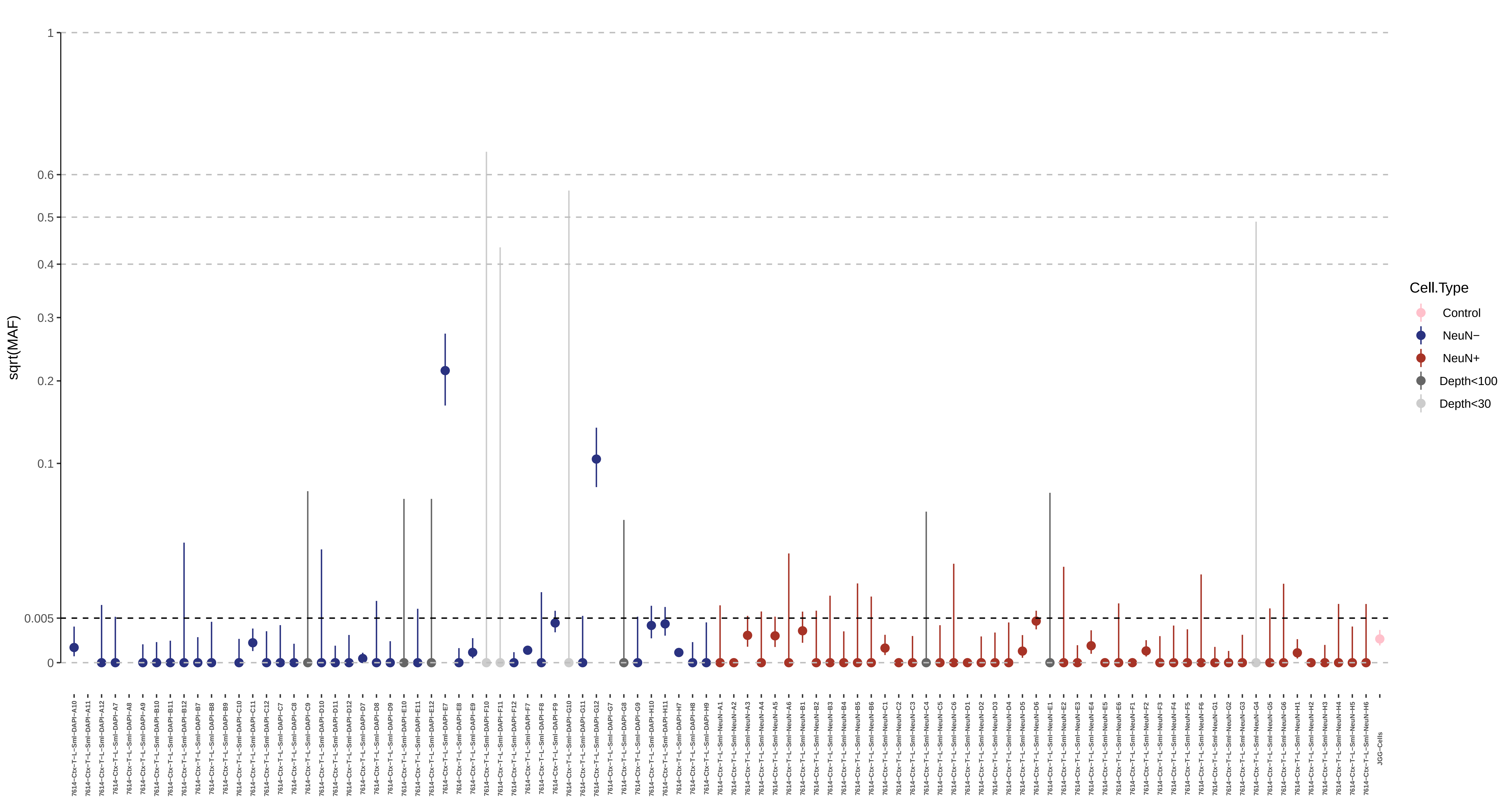

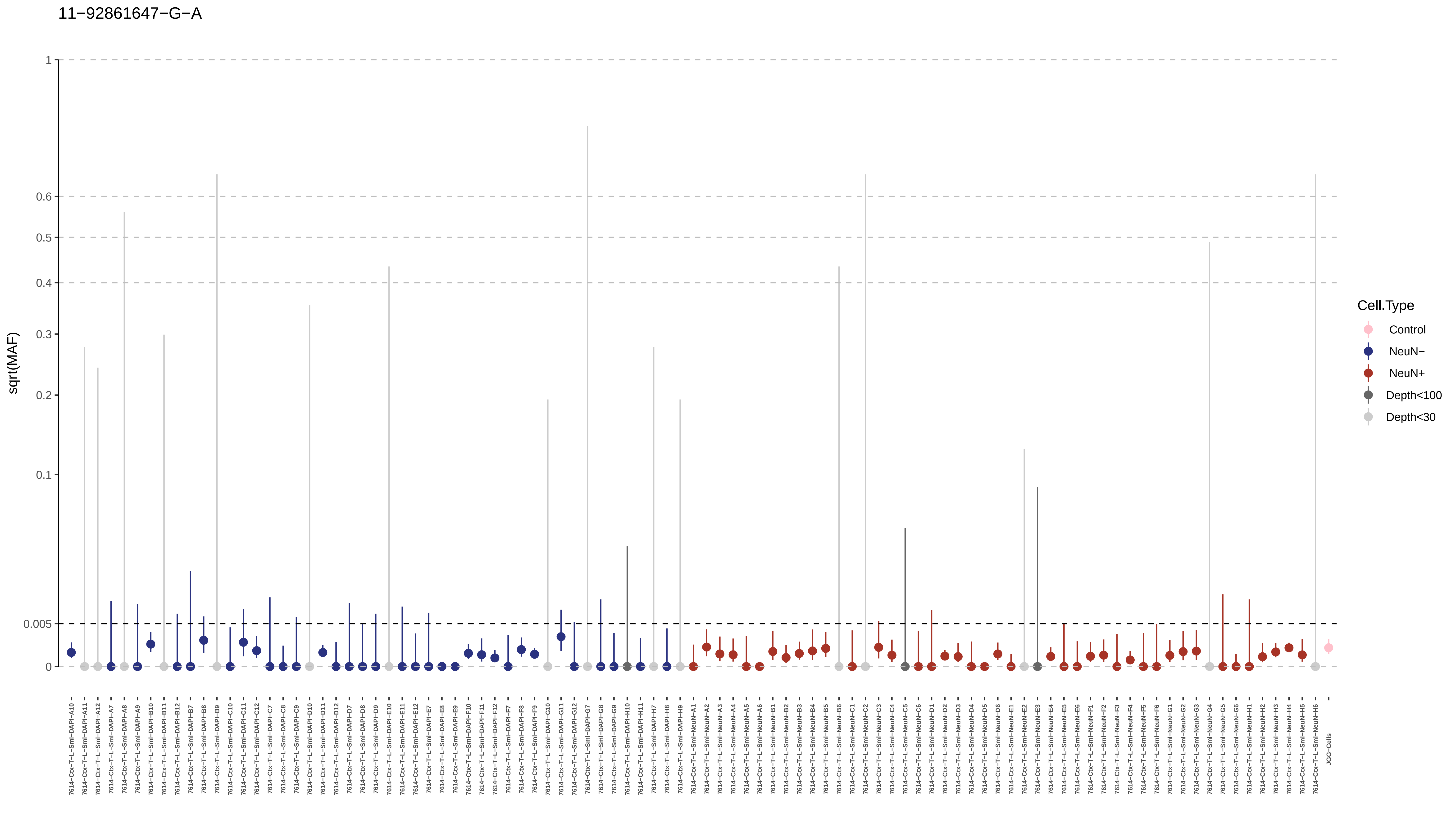
11−92861647−G−A
sqrt(MAF)
1
0.6
0.5
0.4
0.3
0.2
0.1
0.005
0
Cell.Type
Control
NeuN−
NeuN+
Depth<100
Depth<30
JGG-Cells

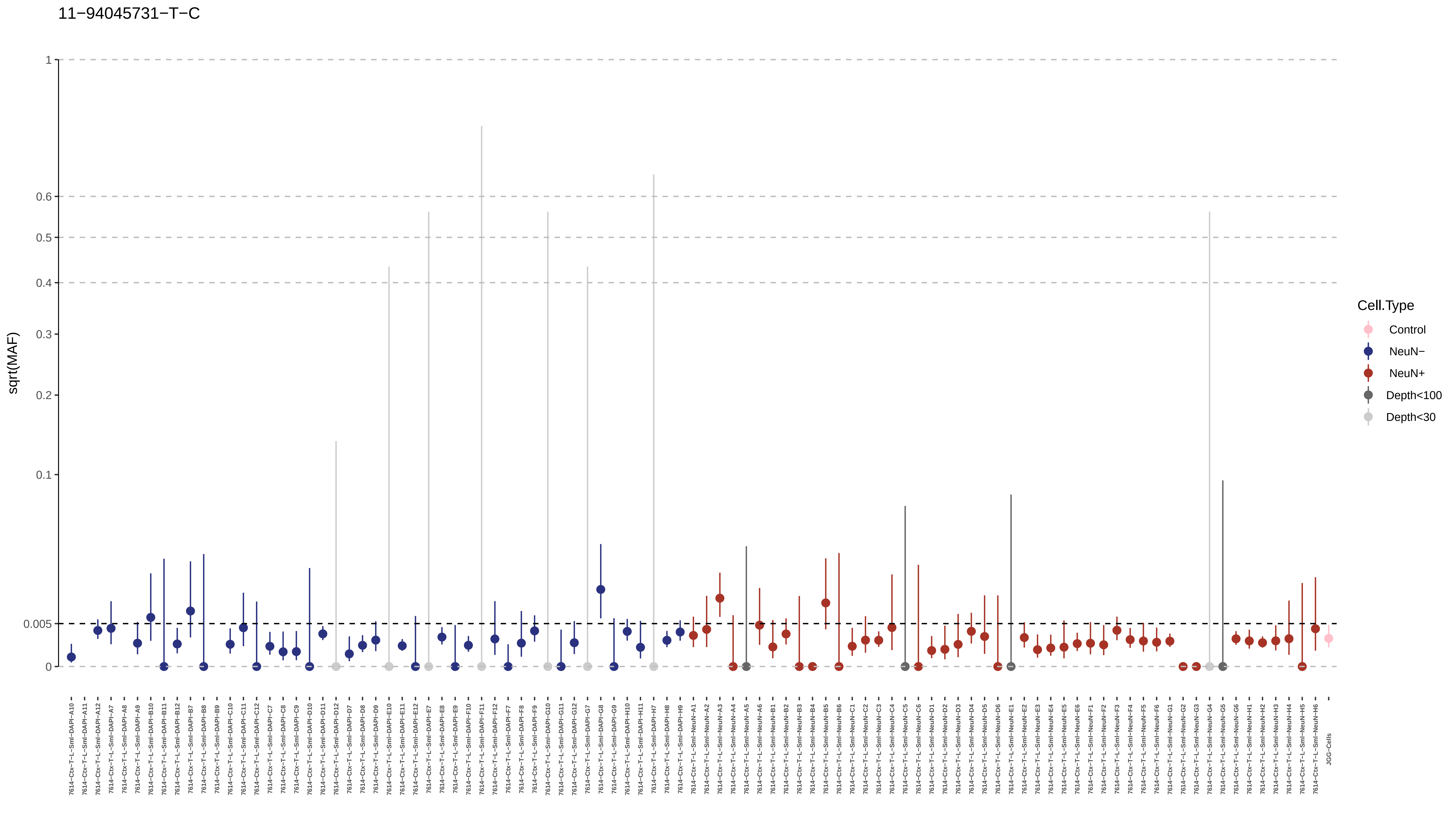

11−94045731−T−C
sqrt(MAF)
1
0.6
0.5
0.4
0.3
0.2
0.1
0.005
0
Cell.Type
Control
NeuN−
NeuN+
Depth<100
Depth<30

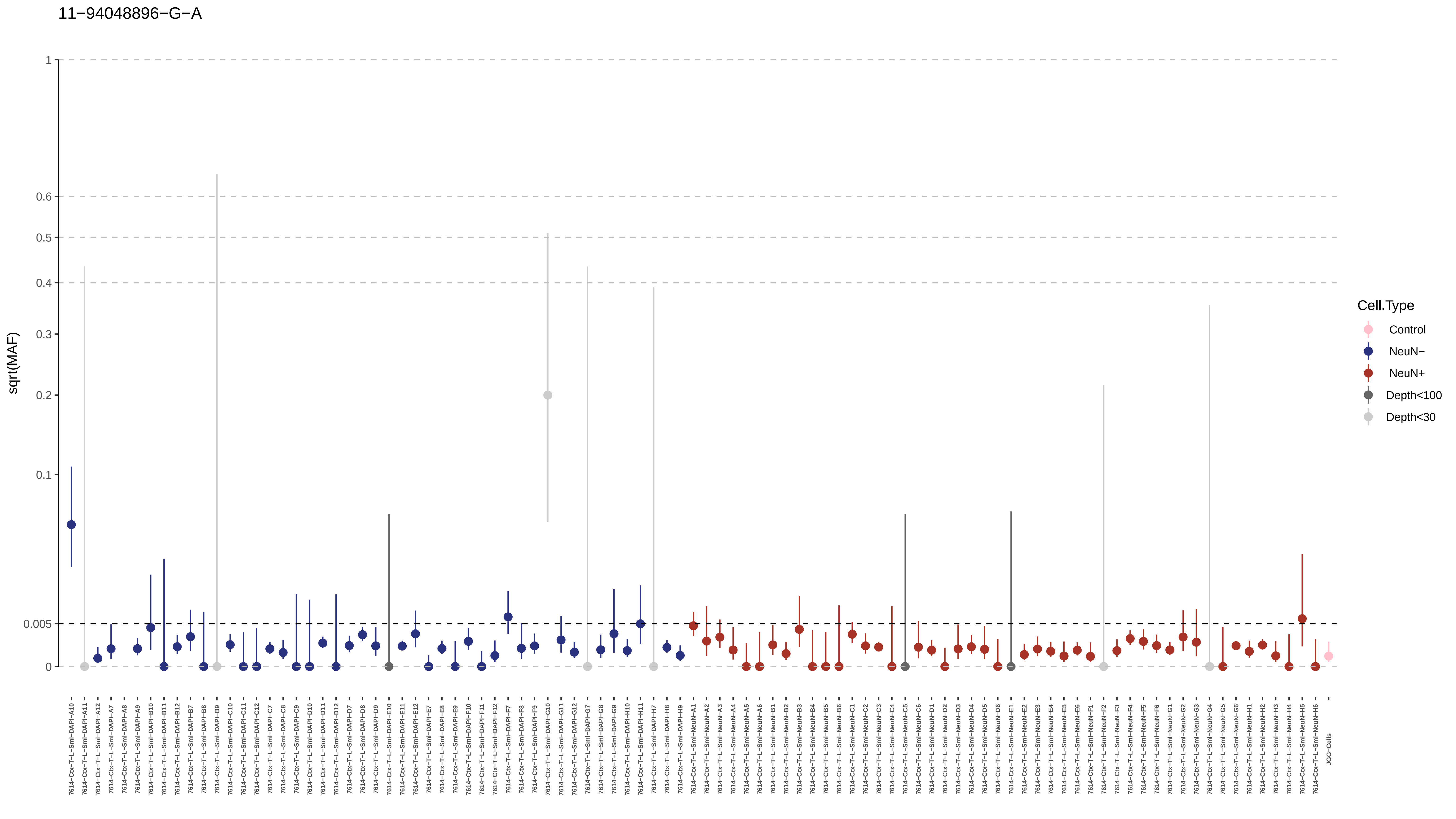

11−94048896−G−A
sqrt(MAF)
1
0.6
0.5
0.4
0.3
0.2
0.1
0.005
0
Cell.Type
Control
NeuN−
NeuN+
Depth<100
Depth<30

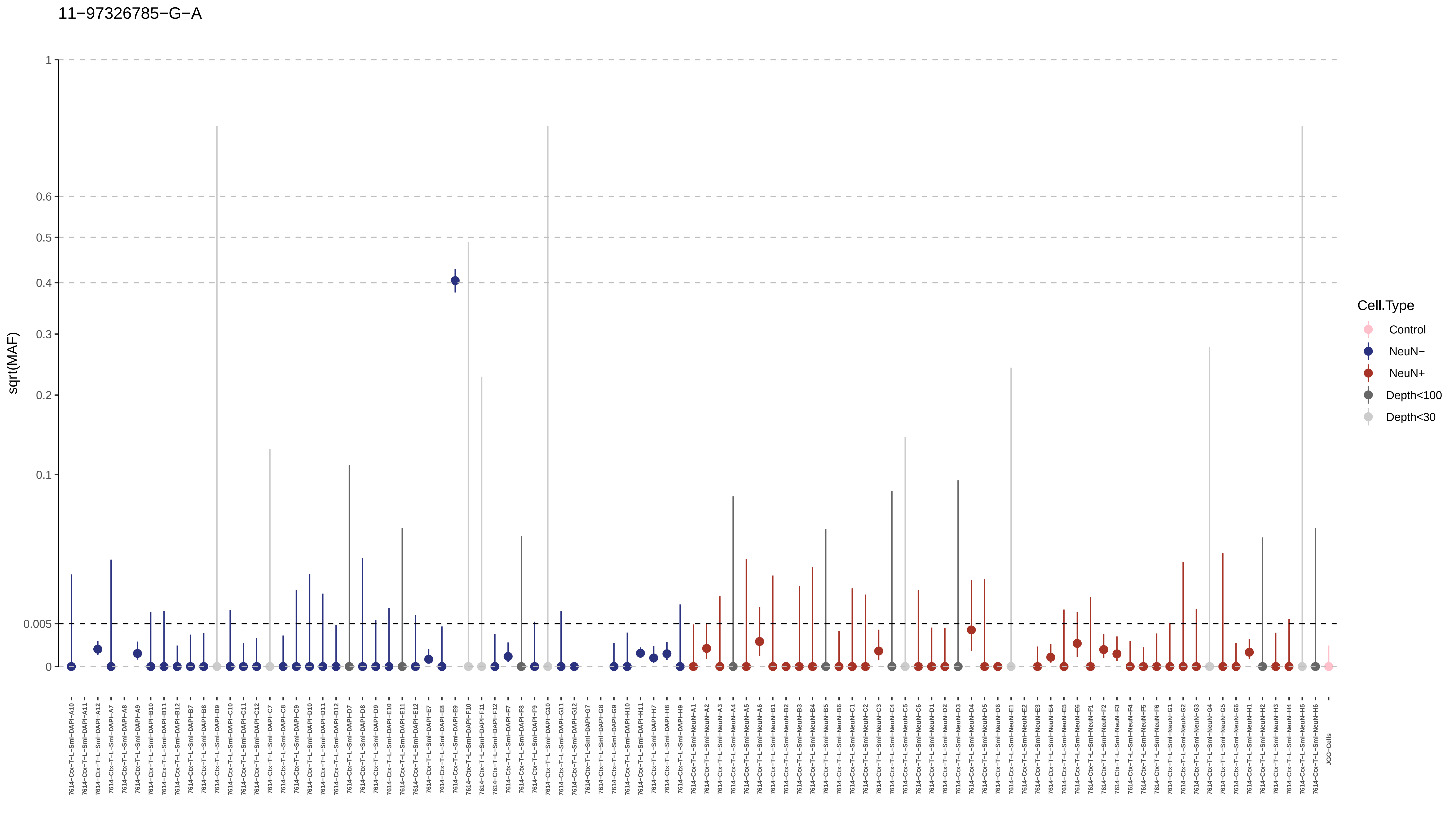
11−97326785−G−A
sqrt(MAF)
1
0.6
0.5
0.4
0.3
0.2
0.1
0.005
0
Cell.Type
Control
NeuN−
NeuN+
Depth<100
Depth<30

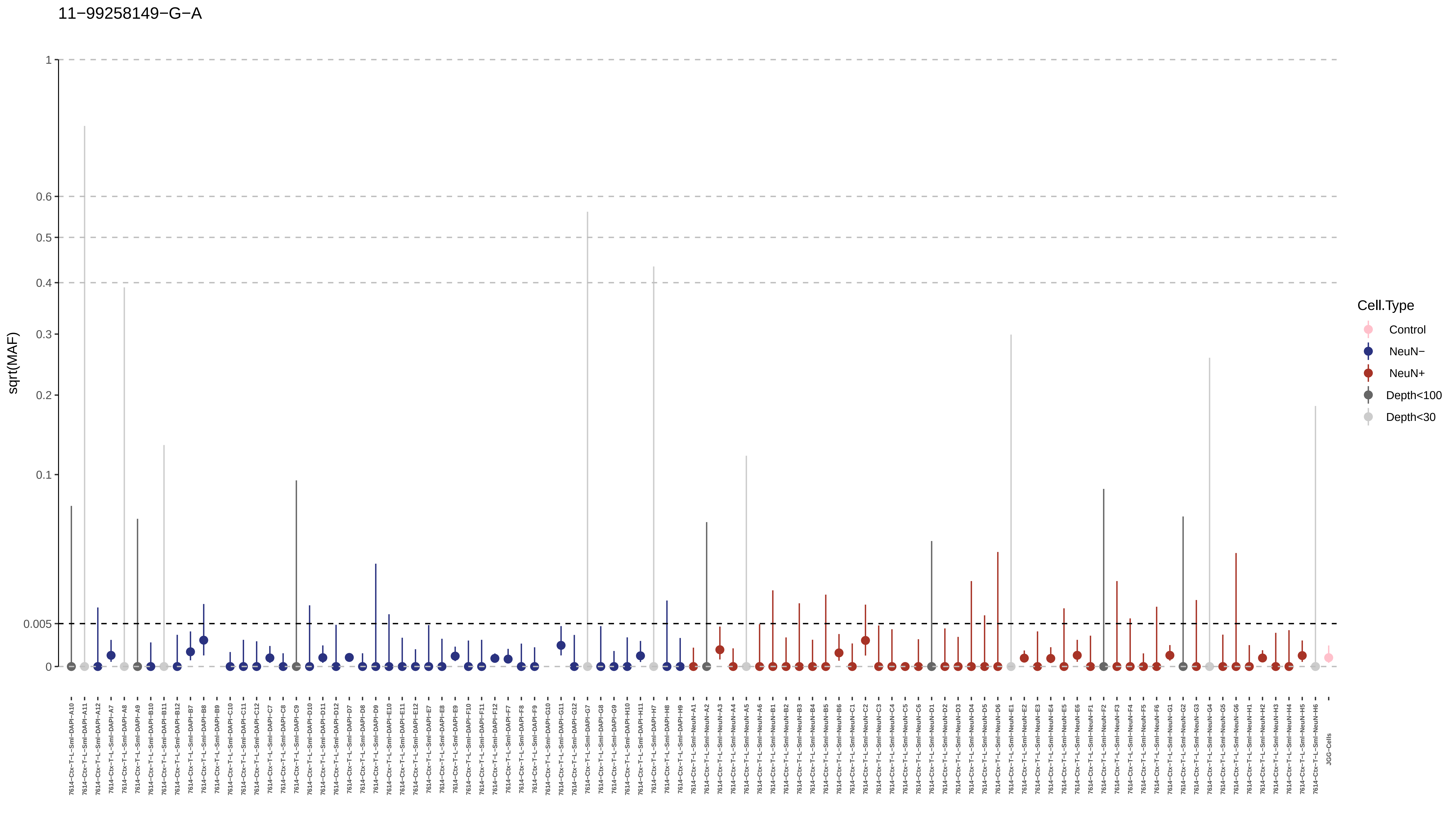
11−99258149−G−A
sqrt(MAF)
1
0.6
0.5
0.4
0.3
0.2
0.1
0.005
0
Cell.Type
Control
NeuN−
NeuN+
Depth<100
Depth<30
JGG-Cells

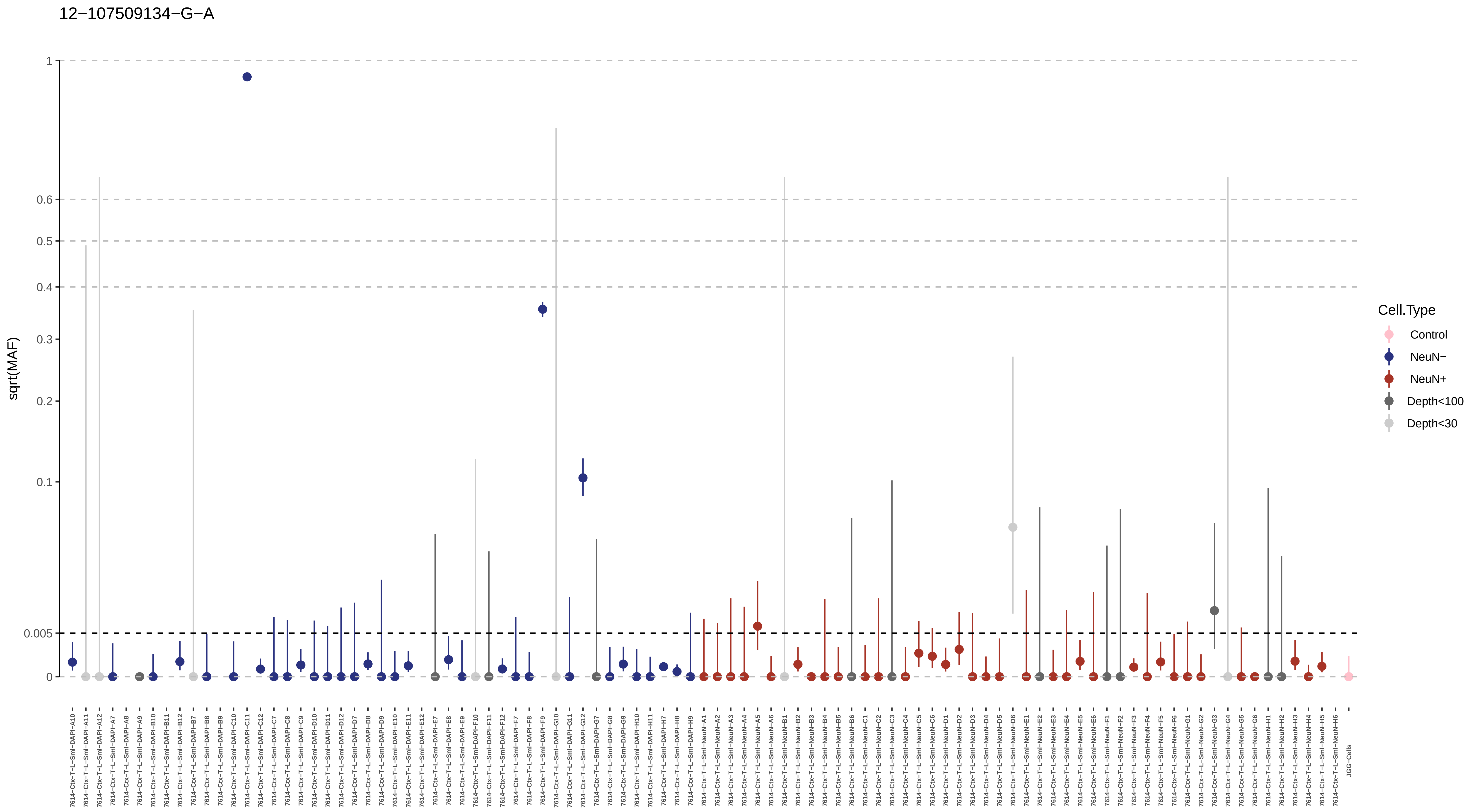
12−107509134−G−A
sqrt(MAF)
1
0.6
0.5
0.4
0.3
0.2
0.1
0.005
0
Cell.Type
Control
NeuN−
NeuN+
Depth<100
Depth<30
7614−Ctx−T−L−Sml−DAPI−A10
7614−Ctx−T−L−Sml−DAPI−A11
7614−Ctx−T−L−Sml−DAPI−A12
7614−Ctx−T−L−Sml−DAPI−A7
7614−Ctx−T−L−Sml−DAPI−A8
7614−Ctx−T−L−Sml−DAPI−A9
7614−Ctx−T−L−Sml−DAPI−B10
7614−Ctx−T−L−Sml−DAPI−B11
7614−Ctx−T−L−Sml−DAPI−B12
7614−Ctx−T−L−Sml−DAPI−B7
7614−Ctx−T−L−Sml−DAPI−B8
7614−Ctx−T−L−Sml−DAPI−B9
7614−Ctx−T−L−Sml−DAPI−C10
7614−Ctx−T−L−Sml−DAPI−C11
7614−Ctx−T−L−Sml−DAPI−C12
7614−Ctx−T−L−Sml−DAPI−C7
7614−Ctx−T−L−Sml−DAPI−C8
7614−Ctx−T−L−Sml−DAPI−C9
7614−Ctx−T−L−Sml−DAPI−D10
7614−Ctx−T−L−Sml−DAPI−D11
7614−Ctx−T−L−Sml−DAPI−D12
7614−Ctx−T−L−Sml−DAPI−D7
7614−Ctx−T−L−Sml−DAPI−D8
7614−Ctx−T−L−Sml−DAPI−D9
7614−Ctx−T−L−Sml−DAPI−E10
7614−Ctx−T−L−Sml−DAPI−E11
7614−Ctx−T−L−Sml−DAPI−E12
7614−Ctx−T−L−Sml−DAPI−E7
7614−Ctx−T−L−Sml−DAPI−E8
7614−Ctx−T−L−Sml−DAPI−E9
7614−Ctx−T−L−Sml−DAPI−F10
7614−Ctx−T−L−Sml−DAPI−F11
7614−Ctx−T−L−Sml−DAPI−F12
7614−Ctx−T−L−Sml−DAPI−F7
7614−Ctx−T−L−Sml−DAPI−F8
7614−Ctx−T−L−Sml−DAPI−F9
7614−Ctx−T−L−Sml−DAPI−G10
7614−Ctx−T−L−Sml−DAPI−G11
7614−Ctx−T−L−Sml−DAPI−G12
7614−Ctx−T−L−Sml−DAPI−G7
7614−Ctx−T−L−Sml−DAPI−G8
7614−Ctx−T−L−Sml−DAPI−G9
7614−Ctx−T−L−Sml−DAPI−H10
7614−Ctx−T−L−Sml−DAPI−H11
7614−Ctx−T−L−Sml−DAPI−H7
7614−Ctx−T−L−Sml−DAPI−H8
7614−Ctx−T−L−Sml−DAPI−H9
7614−Ctx−T−L−Sml−NeuN−A1
7614−Ctx−T−L−Sml−NeuN−A2
7614−Ctx−T−L−Sml−NeuN−A3
7614−Ctx−T−L−Sml−NeuN−A4
7614−Ctx−T−L−Sml−NeuN−A5
7614−Ctx−T−L−Sml−NeuN−A6
7614−Ctx−T−L−Sml−NeuN−B1
7614−Ctx−T−L−Sml−NeuN−B2
7614−Ctx−T−L−Sml−NeuN−B3
7614−Ctx−T−L−Sml−NeuN−B4
7614−Ctx−T−L−Sml−NeuN−B5
7614−Ctx−T−L−Sml−NeuN−B6
7614−Ctx−T−L−Sml−NeuN−C1
7614−Ctx−T−L−Sml−NeuN−C2
7614−Ctx−T−L−Sml−NeuN−C3
7614−Ctx−T−L−Sml−NeuN−C4
7614−Ctx−T−L−Sml−NeuN−C5
7614−Ctx−T−L−Sml−NeuN−C6
7614−Ctx−T−L−Sml−NeuN−D1
7614−Ctx−T−L−Sml−NeuN−D2
7614−Ctx−T−L−Sml−NeuN−D3
7614−Ctx−T−L−Sml−NeuN−D4
7614−Ctx−T−L−Sml−NeuN−D5
7614−Ctx−T−L−Sml−NeuN−D6
7614−Ctx−T−L−Sml−NeuN−E1
7614−Ctx−T−L−Sml−NeuN−E2
7614−Ctx−T−L−Sml−NeuN−E3
7614−Ctx−T−L−Sml−NeuN−E4
7614−Ctx−T−L−Sml−NeuN−E5
7614−Ctx−T−L−Sml−NeuN−E6
7614−Ctx−T−L−Sml−NeuN−F1
7614−Ctx−T−L−Sml−NeuN−F2
7614−Ctx−T−L−Sml−NeuN−F3
7614−Ctx−T−L−Sml−NeuN−F4
7614−Ctx−T−L−Sml−NeuN−F5
7614−Ctx−T−L−Sml−NeuN−F6
7614−Ctx−T−L−Sml−NeuN−G1
7614−Ctx−T−L−Sml−NeuN−G2
7614−Ctx−T−L−Sml−NeuN−G3
7614−Ctx−T−L−Sml−NeuN−G4
7614−Ctx−T−L−Sml−NeuN−G5
7614−Ctx−T−L−Sml−NeuN−G6
7614−Ctx−T−L−Sml−NeuN−H1
7614−Ctx−T−L−Sml−NeuN−H2
7614−Ctx−T−L−Sml−NeuN−H3
7614−Ctx−T−L−Sml−NeuN−H4
7614−Ctx−T−L−Sml−NeuN−H5
7614−Ctx−T−L−Sml−NeuN−H6
JGG-Cells

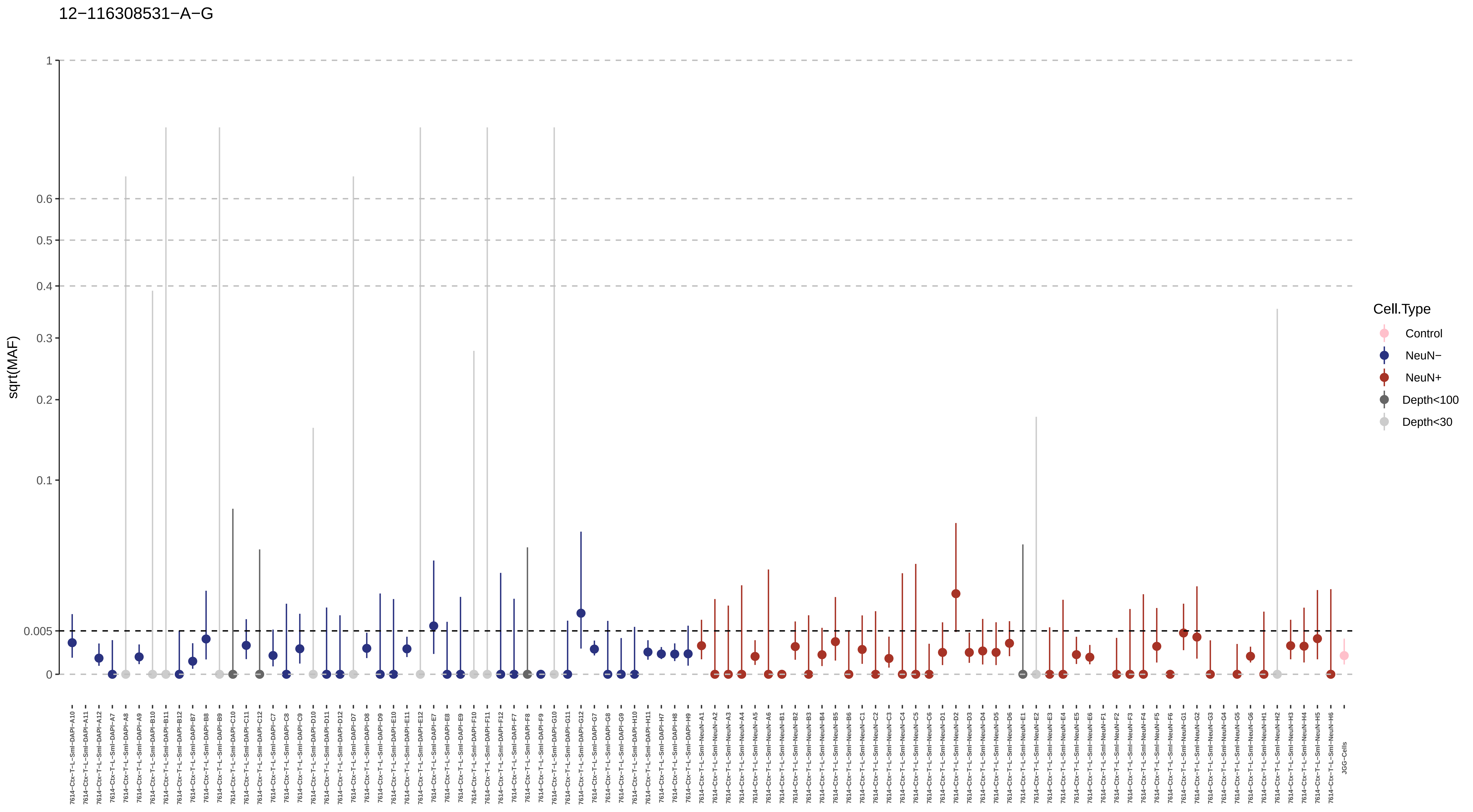
12−116308531−A−G
sqrt(MAF)
1
0.6
0.5
0.4
0.3
0.2
0.1
0.005
0
Cell.Type
Control
NeuN−
NeuN+
Depth<100
Depth<30
JGG-Cells

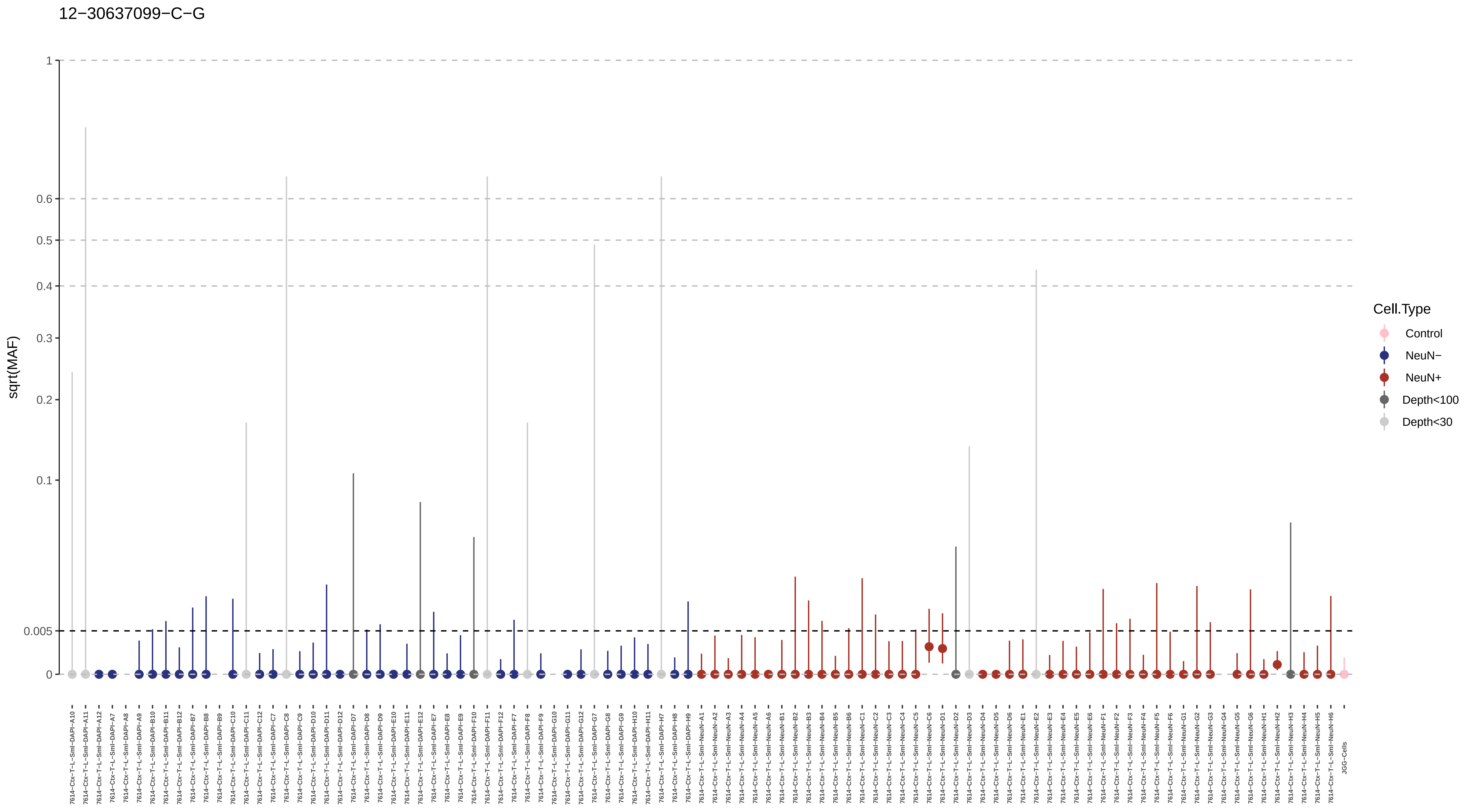

12−30637099−C−G
sqrt(MAF)
1
0.6
0.5
0.4
0.3
0.2
0.1
0.005
0
Cell.Type
Control
NeuN−
NeuN+
Depth<100
Depth<30

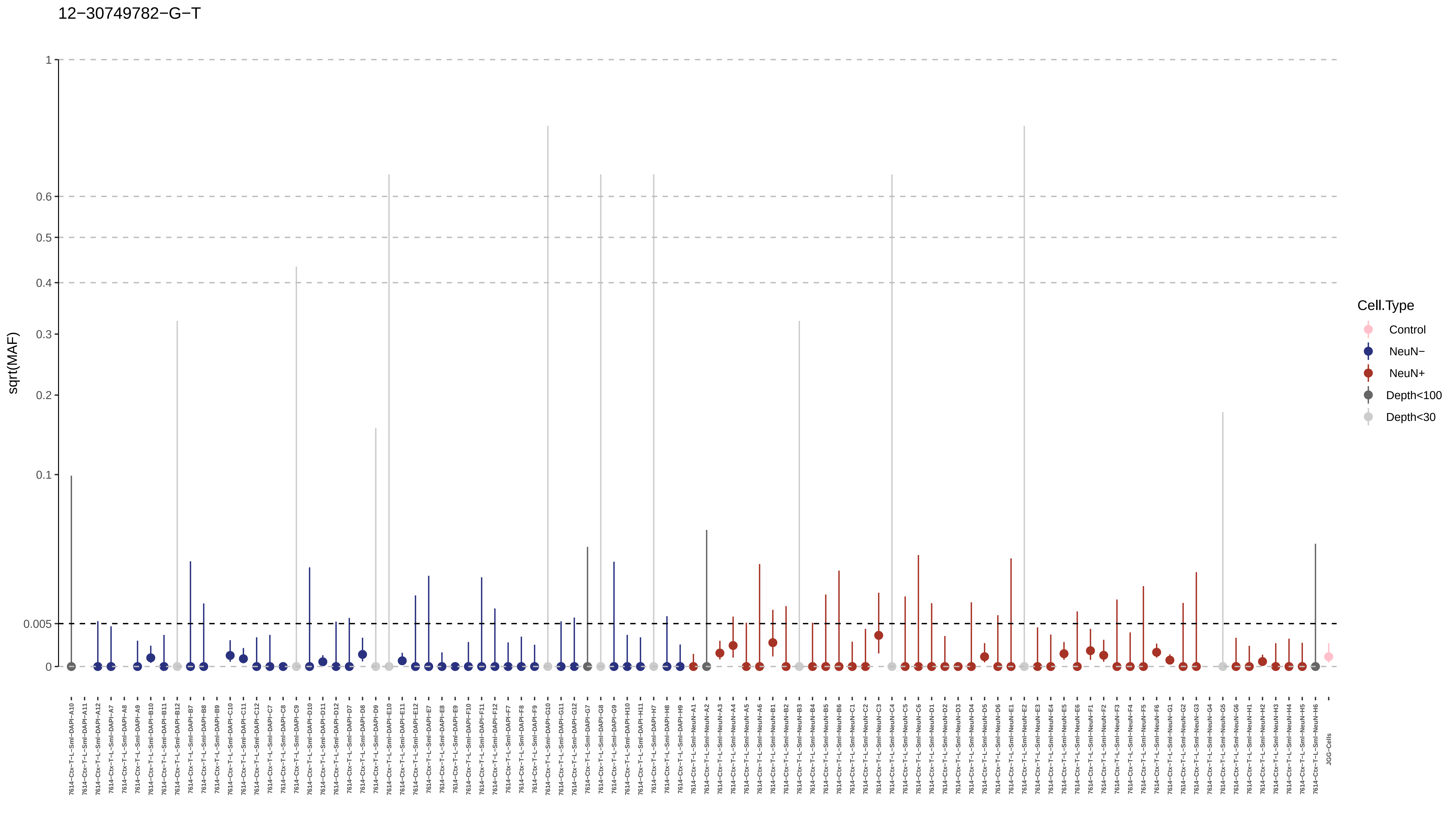
12−30749782−G−T
sqrt(MAF)
1
0.6
0.5
0.4
0.3
0.2
0.1
0.005
0
Cell.Type
Control
NeuN−
NeuN+
Depth<100
Depth<30
JGG-Cells

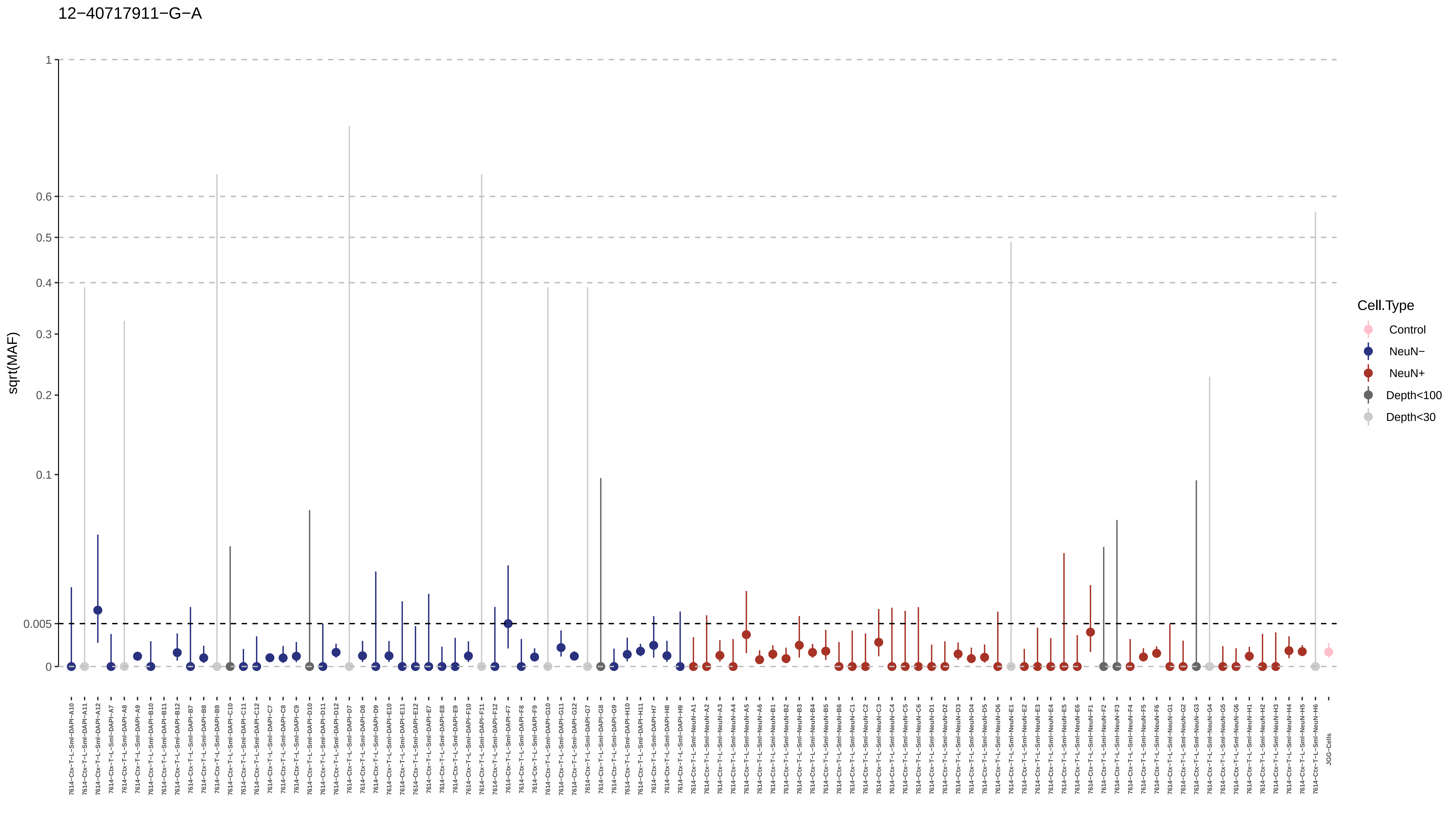

12−40717911−G−A
sqrt(MAF)
1
0.6
0.5
0.4
0.3
0.2
0.1
0.005
0
Cell.Type
Control
NeuN−
NeuN+
Depth<100
Depth<30
JGG-Cells

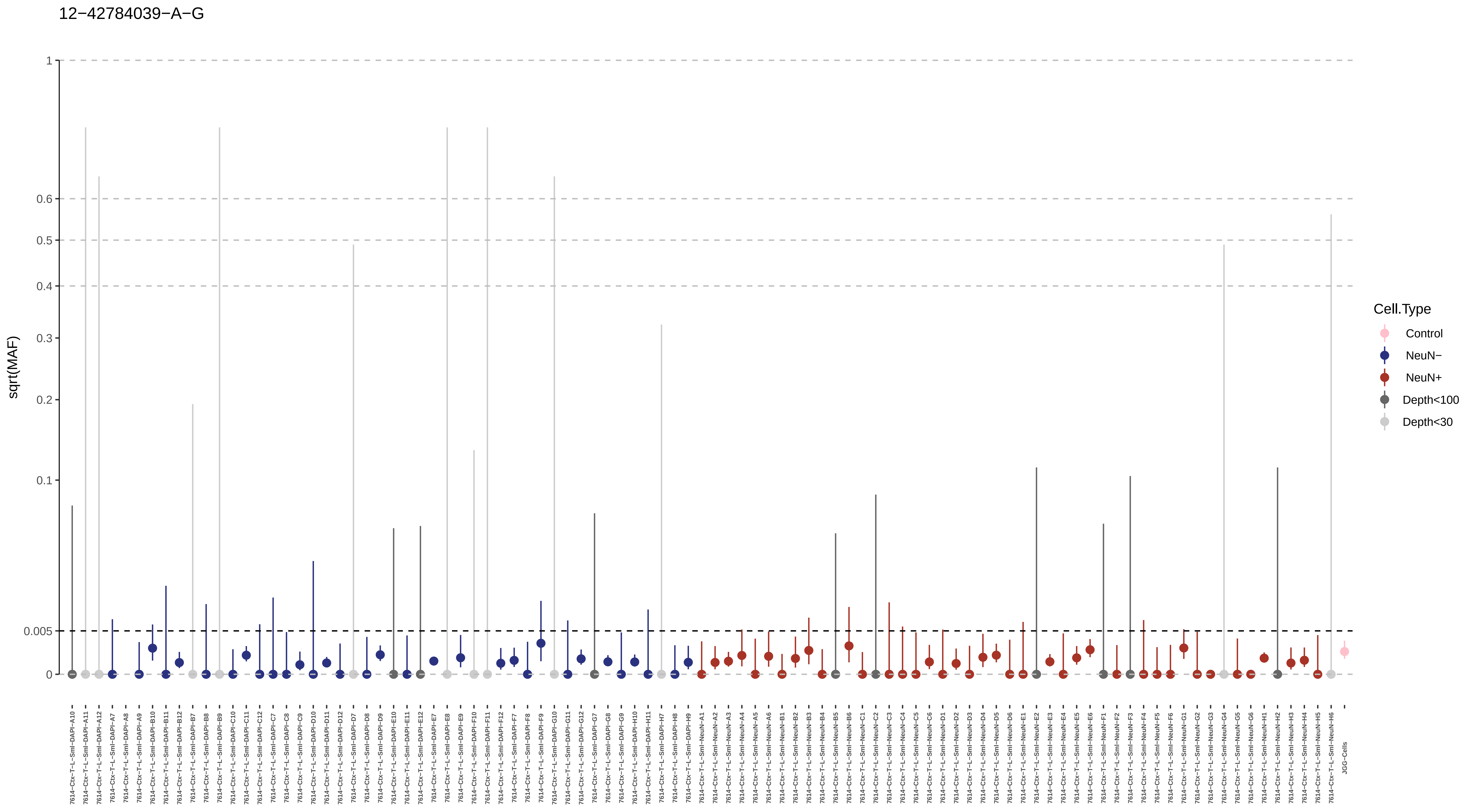

12−42784039−A−G
sqrt(MAF)
1
0.6
0.5
0.4
0.3
0.2
0.1
0.005
0
Cell.Type
Control
NeuN−
NeuN+
Depth<100
Depth<30
7614−Ctx−T−L−Sml−DAPI−A10
7614−Ctx−T−L−Sml−DAPI−A11
7614−Ctx−T−L−Sml−DAPI−A12
7614−Ctx−T−L−Sml−DAPI−A7
7614−Ctx−T−L−Sml−DAPI−A8
7614−Ctx−T−L−Sml−DAPI−A9
7614−Ctx−T−L−Sml−DAPI−B10
7614−Ctx−T−L−Sml−DAPI−B11
7614−Ctx−T−L−Sml−DAPI−B12
7614−Ctx−T−L−Sml−DAPI−B7
7614−Ctx−T−L−Sml−DAPI−B8
7614−Ctx−T−L−Sml−DAPI−B9
7614−Ctx−T−L−Sml−DAPI−C10
7614−Ctx−T−L−Sml−DAPI−C11
7614−Ctx−T−L−Sml−DAPI−C12
7614−Ctx−T−L−Sml−DAPI−C7
7614−Ctx−T−L−Sml−DAPI−C8
7614−Ctx−T−L−Sml−DAPI−C9
7614−Ctx−T−L−Sml−DAPI−D10
7614−Ctx−T−L−Sml−DAPI−D11
7614−Ctx−T−L−Sml−DAPI−D12
7614−Ctx−T−L−Sml−DAPI−D7
7614−Ctx−T−L−Sml−DAPI−D8
7614−Ctx−T−L−Sml−DAPI−D9
7614−Ctx−T−L−Sml−DAPI−E10
7614−Ctx−T−L−Sml−DAPI−E11
7614−Ctx−T−L−Sml−DAPI−E12
7614−Ctx−T−L−Sml−DAPI−E7
7614−Ctx−T−L−Sml−DAPI−E8
7614−Ctx−T−L−Sml−DAPI−E9
7614−Ctx−T−L−Sml−DAPI−F10
7614−Ctx−T−L−Sml−DAPI−F11
7614−Ctx−T−L−Sml−DAPI−F12
7614−Ctx−T−L−Sml−DAPI−F7
7614−Ctx−T−L−Sml−DAPI−F8
7614−Ctx−T−L−Sml−DAPI−F9
7614−Ctx−T−L−Sml−DAPI−G10
7614−Ctx−T−L−Sml−DAPI−G11
7614−Ctx−T−L−Sml−DAPI−G12
7614−Ctx−T−L−Sml−DAPI−G7
7614−Ctx−T−L−Sml−DAPI−G8
7614−Ctx−T−L−Sml−DAPI−G9
7614−Ctx−T−L−Sml−DAPI−H10
7614−Ctx−T−L−Sml−DAPI−H11
7614−Ctx−T−L−Sml−DAPI−H7
7614−Ctx−T−L−Sml−DAPI−H8
7614−Ctx−T−L−Sml−DAPI−H9
7614−Ctx−T−L−Sml−NeuN−A1
7614−Ctx−T−L−Sml−NeuN−A2
7614−Ctx−T−L−Sml−NeuN−A3
7614−Ctx−T−L−Sml−NeuN−A4
7614−Ctx−T−L−Sml−NeuN−A5
7614−Ctx−T−L−Sml−NeuN−A6
7614−Ctx−T−L−Sml−NeuN−B1
7614−Ctx−T−L−Sml−NeuN−B2
7614−Ctx−T−L−Sml−NeuN−B3
7614−Ctx−T−L−Sml−NeuN−B4
7614−Ctx−T−L−Sml−NeuN−B5
7614−Ctx−T−L−Sml−NeuN−B6
7614−Ctx−T−L−Sml−NeuN−C1
7614−Ctx−T−L−Sml−NeuN−C2
7614−Ctx−T−L−Sml−NeuN−C3
7614−Ctx−T−L−Sml−NeuN−C4
7614−Ctx−T−L−Sml−NeuN−C5
7614−Ctx−T−L−Sml−NeuN−C6
7614−Ctx−T−L−Sml−NeuN−D1
7614−Ctx−T−L−Sml−NeuN−D2
7614−Ctx−T−L−Sml−NeuN−D3
7614−Ctx−T−L−Sml−NeuN−D4
7614−Ctx−T−L−Sml−NeuN−D5
7614−Ctx−T−L−Sml−NeuN−D6
7614−Ctx−T−L−Sml−NeuN−E1
7614−Ctx−T−L−Sml−NeuN−E2
7614−Ctx−T−L−Sml−NeuN−E3
7614−Ctx−T−L−Sml−NeuN−E4
7614−Ctx−T−L−Sml−NeuN−E5
7614−Ctx−T−L−Sml−NeuN−E6
7614−Ctx−T−L−Sml−NeuN−F1
7614−Ctx−T−L−Sml−NeuN−F2
7614−Ctx−T−L−Sml−NeuN−F3
7614−Ctx−T−L−Sml−NeuN−F4
7614−Ctx−T−L−Sml−NeuN−F5
7614−Ctx−T−L−Sml−NeuN−F6
7614−Ctx−T−L−Sml−NeuN−G1
7614−Ctx−T−L−Sml−NeuN−G2
7614−Ctx−T−L−Sml−NeuN−G3
7614−Ctx−T−L−Sml−NeuN−G4
7614−Ctx−T−L−Sml−NeuN−G5
7614−Ctx−T−L−Sml−NeuN−G6
7614−Ctx−T−L−Sml−NeuN−H1
7614−Ctx−T−L−Sml−NeuN−H2
7614−Ctx−T−L−Sml−NeuN−H3
7614−Ctx−T−L−Sml−NeuN−H4
7614−Ctx−T−L−Sml−NeuN−H5
7614−Ctx−T−L−Sml−NeuN−H6
JGG−Cells

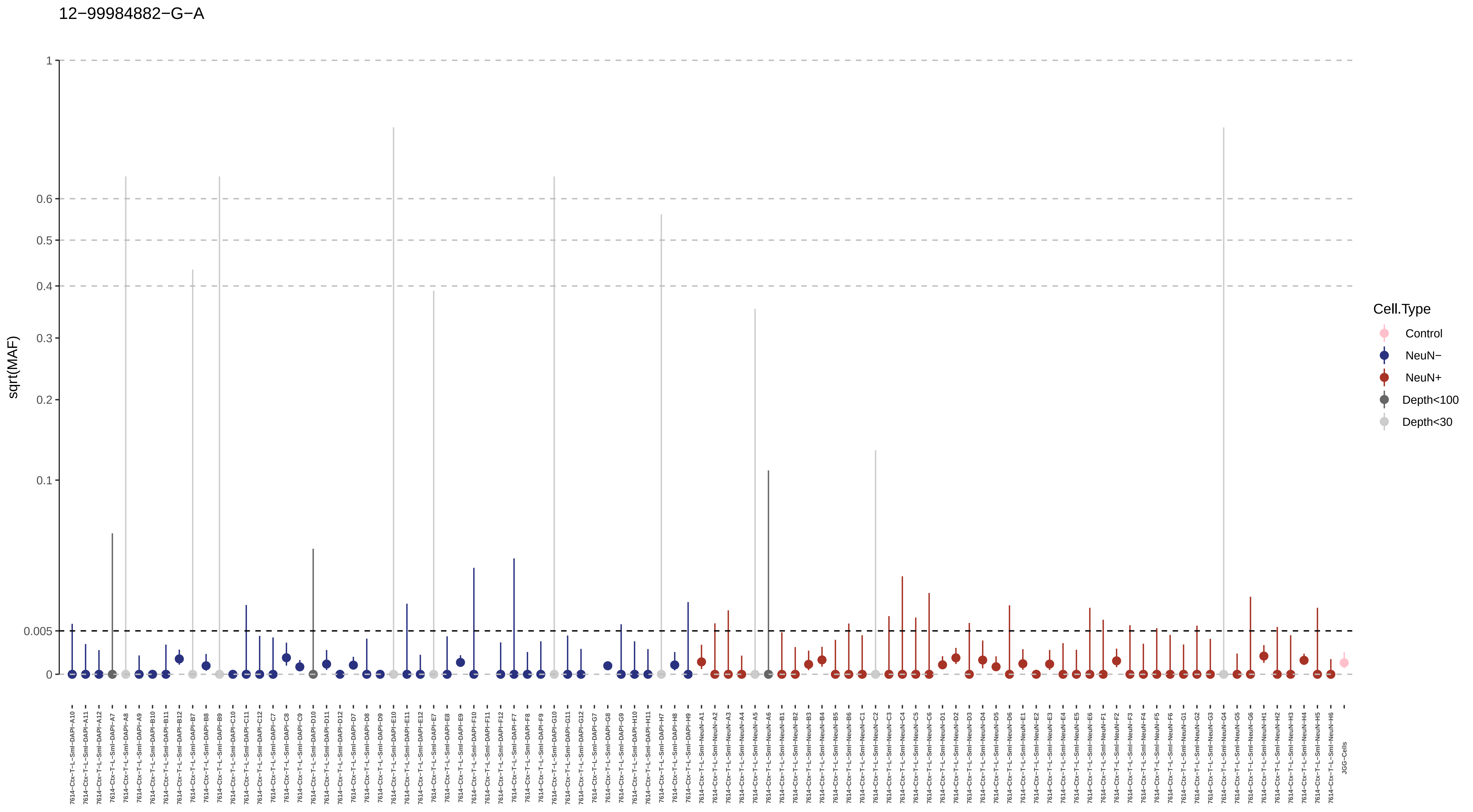
12−99984882−G−A
sqrt(MAF)
1
0.6
0.5
0.4
0.3
0.2
0.1
0.005
0
Cell.Type
Control
NeuN−
NeuN+
Depth<100
Depth<30
7614−Ctx−T−L−Sml−DAPI−A10
7614−Ctx−T−L−Sml−DAPI−A11
7614−Ctx−T−L−Sml−DAPI−A12
7614−Ctx−T−L−Sml−DAPI−A7
7614−Ctx−T−L−Sml−DAPI−A8
7614−Ctx−T−L−Sml−DAPI−A9
7614−Ctx−T−L−Sml−DAPI−B10
7614−Ctx−T−L−Sml−DAPI−B11
7614−Ctx−T−L−Sml−DAPI−B12
7614−Ctx−T−L−Sml−DAPI−B7
7614−Ctx−T−L−Sml−DAPI−B8
7614−Ctx−T−L−Sml−DAPI−B9
7614−Ctx−T−L−Sml−DAPI−C10
7614−Ctx−T−L−Sml−DAPI−C11
7614−Ctx−T−L−Sml−DAPI−C12
7614−Ctx−T−L−Sml−DAPI−C7
7614−Ctx−T−L−Sml−DAPI−C8
7614−Ctx−T−L−Sml−DAPI−C9
7614−Ctx−T−L−Sml−DAPI−D10
7614−Ctx−T−L−Sml−DAPI−D11
7614−Ctx−T−L−Sml−DAPI−D12
7614−Ctx−T−L−Sml−DAPI−D7
7614−Ctx−T−L−Sml−DAPI−D8
7614−Ctx−T−L−Sml−DAPI−D9
7614−Ctx−T−L−Sml−DAPI−E10
7614−Ctx−T−L−Sml−DAPI−E11
7614−Ctx−T−L−Sml−DAPI−E12
7614−Ctx−T−L−Sml−DAPI−E7
7614−Ctx−T−L−Sml−DAPI−E8
7614−Ctx−T−L−Sml−DAPI−E9
7614−Ctx−T−L−Sml−DAPI−F10
7614−Ctx−T−L−Sml−DAPI−F11
7614−Ctx−T−L−Sml−DAPI−F12
7614−Ctx−T−L−Sml−DAPI−F7
7614−Ctx−T−L−Sml−DAPI−F8
7614−Ctx−T−L−Sml−DAPI−F9
7614−Ctx−T−L−Sml−DAPI−G10
7614−Ctx−T−L−Sml−DAPI−G11
7614−Ctx−T−L−Sml−DAPI−G12
7614−Ctx−T−L−Sml−DAPI−G7
7614−Ctx−T−L−Sml−DAPI−G8
7614−Ctx−T−L−Sml−DAPI−G9
7614−Ctx−T−L−Sml−DAPI−H10
7614−Ctx−T−L−Sml−DAPI−H11
7614−Ctx−T−L−Sml−DAPI−H7
7614−Ctx−T−L−Sml−DAPI−H8
7614−Ctx−T−L−Sml−DAPI−H9
7614−Ctx−T−L−Sml−NeuN−A1
7614−Ctx−T−L−Sml−NeuN−A2
7614−Ctx−T−L−Sml−NeuN−A3
7614−Ctx−T−L−Sml−NeuN−A4
7614−Ctx−T−L−Sml−NeuN−A5
7614−Ctx−T−L−Sml−NeuN−A6
7614−Ctx−T−L−Sml−NeuN−B1
7614−Ctx−T−L−Sml−NeuN−B2
7614−Ctx−T−L−Sml−NeuN−B3
7614−Ctx−T−L−Sml−NeuN−B4
7614−Ctx−T−L−Sml−NeuN−B5
7614−Ctx−T−L−Sml−NeuN−B6
7614−Ctx−T−L−Sml−NeuN−C1
7614−Ctx−T−L−Sml−NeuN−C2
7614−Ctx−T−L−Sml−NeuN−C3
7614−Ctx−T−L−Sml−NeuN−C4
7614−Ctx−T−L−Sml−NeuN−C5
7614−Ctx−T−L−Sml−NeuN−C6
7614−Ctx−T−L−Sml−NeuN−D1
7614−Ctx−T−L−Sml−NeuN−D2
7614−Ctx−T−L−Sml−NeuN−D3
7614−Ctx−T−L−Sml−NeuN−D4
7614−Ctx−T−L−Sml−NeuN−D5
7614−Ctx−T−L−Sml−NeuN−D6
7614−Ctx−T−L−Sml−NeuN−E1
7614−Ctx−T−L−Sml−NeuN−E2
7614−Ctx−T−L−Sml−NeuN−E3
7614−Ctx−T−L−Sml−NeuN−E4
7614−Ctx−T−L−Sml−NeuN−E5
7614−Ctx−T−L−Sml−NeuN−E6
7614−Ctx−T−L−Sml−NeuN−F1
7614−Ctx−T−L−Sml−NeuN−F2
7614−Ctx−T−L−Sml−NeuN−F3
7614−Ctx−T−L−Sml−NeuN−F4
7614−Ctx−T−L−Sml−NeuN−F5
7614−Ctx−T−L−Sml−NeuN−F6
7614−Ctx−T−L−Sml−NeuN−G1
7614−Ctx−T−L−Sml−NeuN−G2
7614−Ctx−T−L−Sml−NeuN−G3
7614−Ctx−T−L−Sml−NeuN−G4
7614−Ctx−T−L−Sml−NeuN−G5
7614−Ctx−T−L−Sml−NeuN−G6
7614−Ctx−T−L−Sml−NeuN−H1
7614−Ctx−T−L−Sml−NeuN−H2
7614−Ctx−T−L−Sml−NeuN−H3
7614−Ctx−T−L−Sml−NeuN−H4
7614−Ctx−T−L−Sml−NeuN−H5
7614−Ctx−T−L−Sml−NeuN−H6
JGG-Cells

13−104312064−G−A

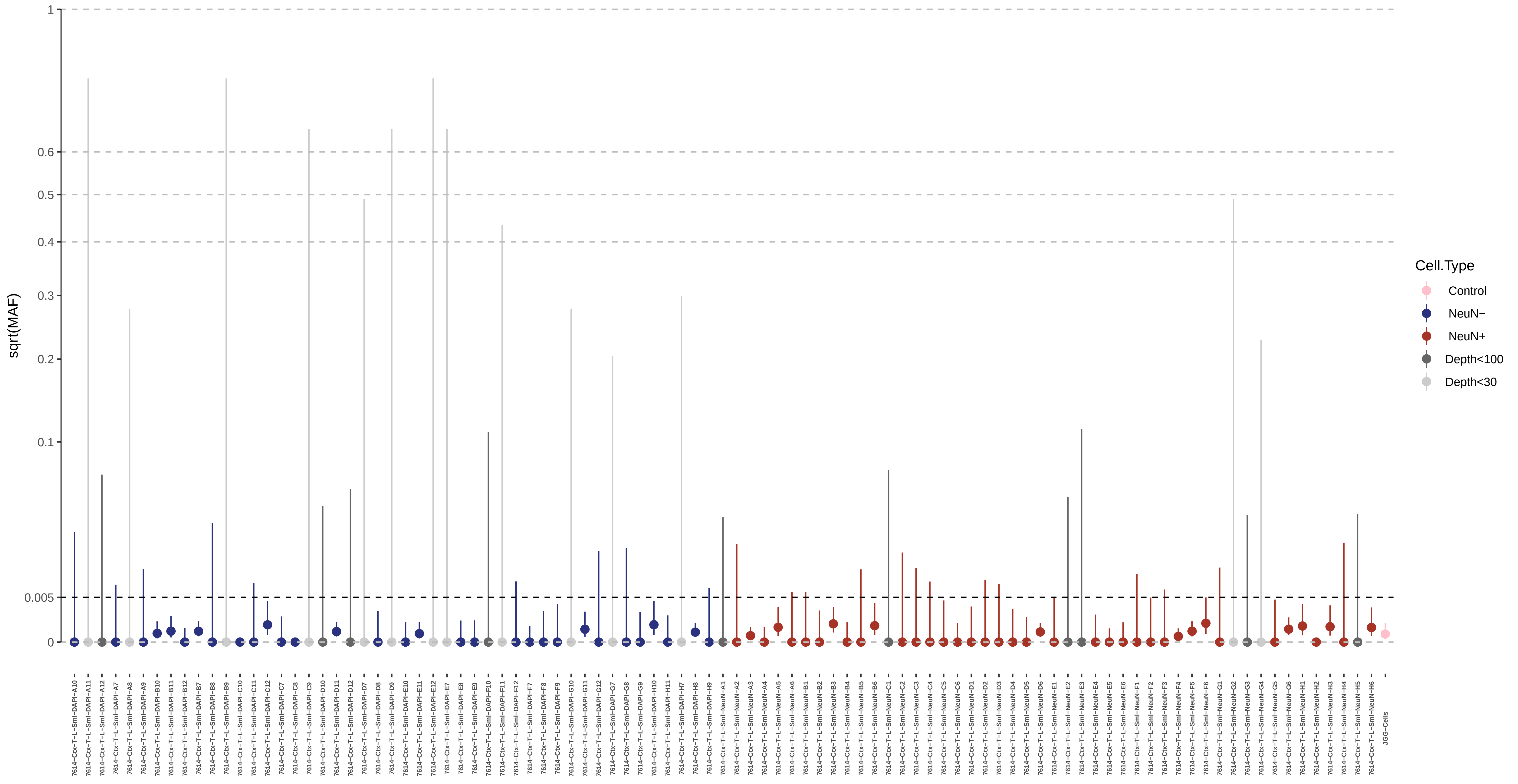

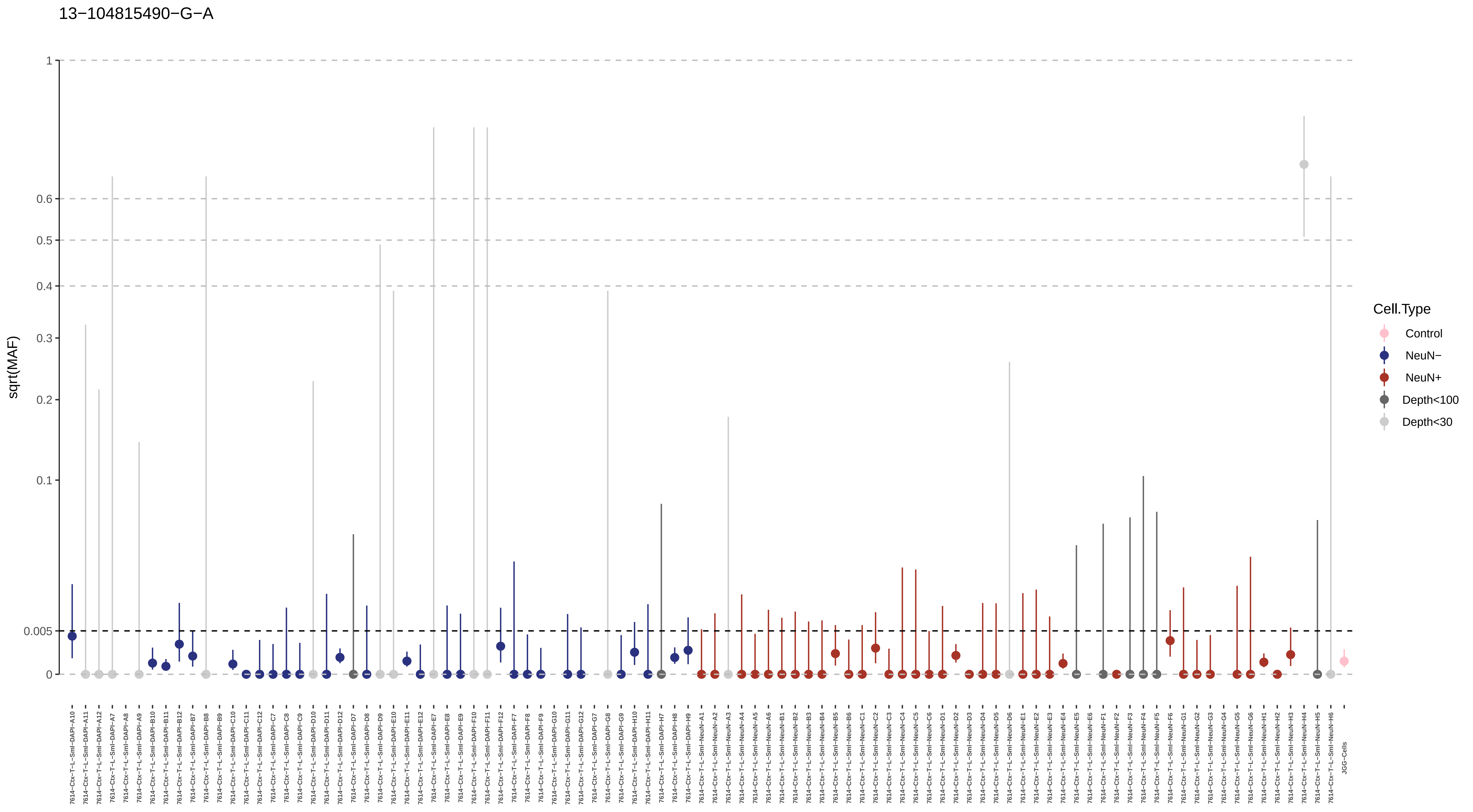
13−104815490−G−A
sqrt(MAF)
1
0.6
0.5
0.4
0.3
0.2
0.1
0.005
0
Cell.Type
Control
NeuN−
NeuN+
Depth<100
Depth<30

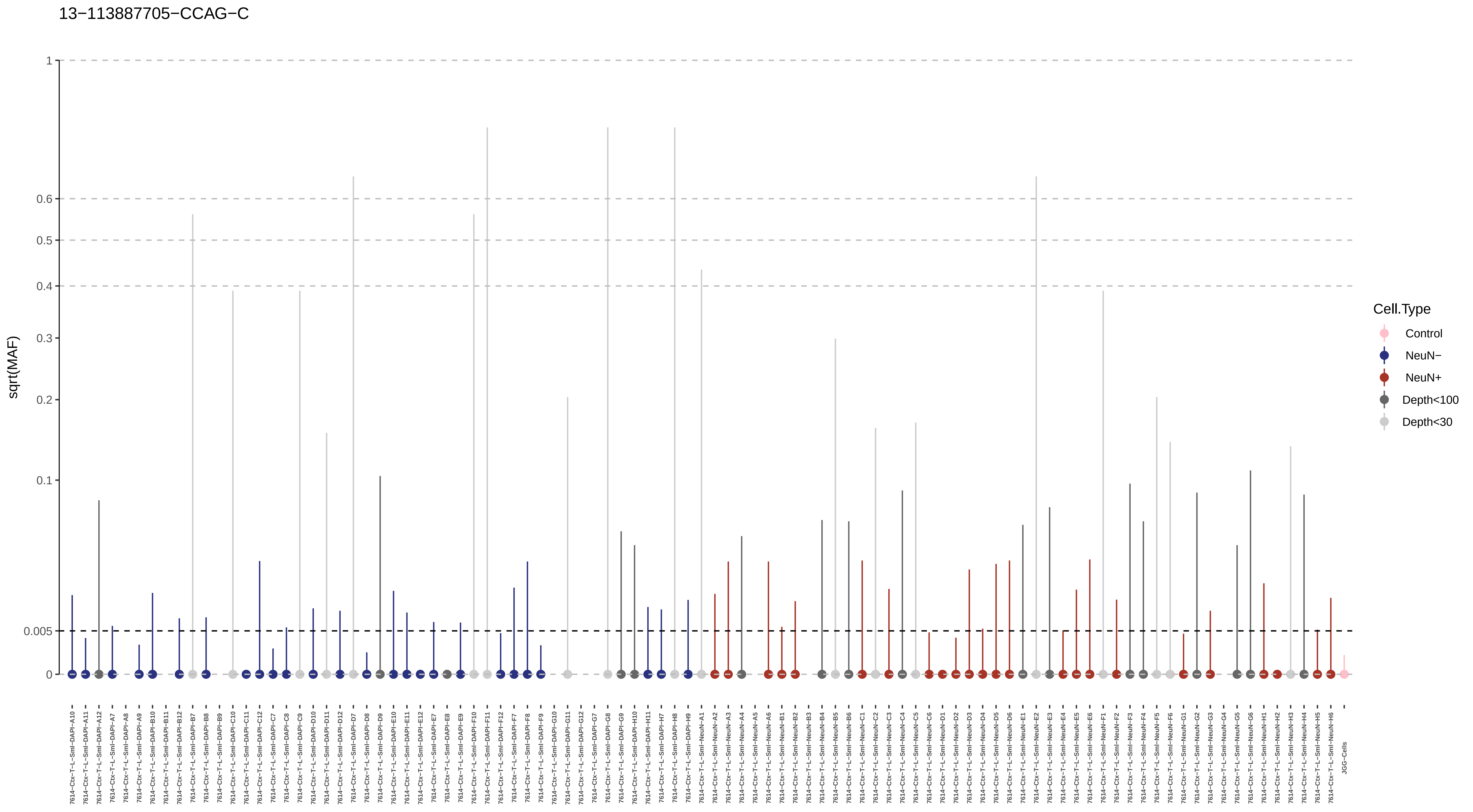
13−113887705−CCAG−C
sqrt(MAF)
1
0.6
0.5
0.4
0.3
0.2
0.1
0.005
0
Cell.Type
Control
NeuN−
NeuN+
Depth<100
Depth<30
7614−Ctx−T−L−Sml−DAPI−A10
7614−Ctx−T−L−Sml−DAPI−A11
7614−Ctx−T−L−Sml−DAPI−A12
7614−Ctx−T−L−Sml−DAPI−A7
7614−Ctx−T−L−Sml−DAPI−A8
7614−Ctx−T−L−Sml−DAPI−A9
7614−Ctx−T−L−Sml−DAPI−B10
7614−Ctx−T−L−Sml−DAPI−B11
7614−Ctx−T−L−Sml−DAPI−B12
7614−Ctx−T−L−Sml−DAPI−B7
7614−Ctx−T−L−Sml−DAPI−B8
7614−Ctx−T−L−Sml−DAPI−B9
7614−Ctx−T−L−Sml−DAPI−C10
7614−Ctx−T−L−Sml−DAPI−C11
7614−Ctx−T−L−Sml−DAPI−C12
7614−Ctx−T−L−Sml−DAPI−C7
7614−Ctx−T−L−Sml−DAPI−C8
7614−Ctx−T−L−Sml−DAPI−C9
7614−Ctx−T−L−Sml−DAPI−D10
7614−Ctx−T−L−Sml−DAPI−D11
7614−Ctx−T−L−Sml−DAPI−D12
7614−Ctx−T−L−Sml−DAPI−D7
7614−Ctx−T−L−Sml−DAPI−D8
7614−Ctx−T−L−Sml−DAPI−D9
7614−Ctx−T−L−Sml−DAPI−E10
7614−Ctx−T−L−Sml−DAPI−E11
7614−Ctx−T−L−Sml−DAPI−E12
7614−Ctx−T−L−Sml−DAPI−E7
7614−Ctx−T−L−Sml−DAPI−E8
7614−Ctx−T−L−Sml−DAPI−E9
7614−Ctx−T−L−Sml−DAPI−F10
7614−Ctx−T−L−Sml−DAPI−F11
7614−Ctx−T−L−Sml−DAPI−F12
7614−Ctx−T−L−Sml−DAPI−F7
7614−Ctx−T−L−Sml−DAPI−F8
7614−Ctx−T−L−Sml−DAPI−F9
7614−Ctx−T−L−Sml−DAPI−G10
7614−Ctx−T−L−Sml−DAPI−G11
7614−Ctx−T−L−Sml−DAPI−G12
7614−Ctx−T−L−Sml−DAPI−G7
7614−Ctx−T−L−Sml−DAPI−G8
7614−Ctx−T−L−Sml−DAPI−G9
7614−Ctx−T−L−Sml−DAPI−H10
7614−Ctx−T−L−Sml−DAPI−H11
7614−Ctx−T−L−Sml−DAPI−H7
7614−Ctx−T−L−Sml−DAPI−H8
7614−Ctx−T−L−Sml−DAPI−H9
7614−Ctx−T−L−Sml−NeuN−A1
7614−Ctx−T−L−Sml−NeuN−A2
7614−Ctx−T−L−Sml−NeuN−A3
7614−Ctx−T−L−Sml−NeuN−A4
7614−Ctx−T−L−Sml−NeuN−A5
7614−Ctx−T−L−Sml−NeuN−A6
7614−Ctx−T−L−Sml−NeuN−B1
7614−Ctx−T−L−Sml−NeuN−B2
7614−Ctx−T−L−Sml−NeuN−B3
7614−Ctx−T−L−Sml−NeuN−B4
7614−Ctx−T−L−Sml−NeuN−B5
7614−Ctx−T−L−Sml−NeuN−B6
7614−Ctx−T−L−Sml−NeuN−C1
7614−Ctx−T−L−Sml−NeuN−C2
7614−Ctx−T−L−Sml−NeuN−C3
7614−Ctx−T−L−Sml−NeuN−C4
7614−Ctx−T−L−Sml−NeuN−C5
7614−Ctx−T−L−Sml−NeuN−C6
7614−Ctx−T−L−Sml−NeuN−D1
7614−Ctx−T−L−Sml−NeuN−D2
7614−Ctx−T−L−Sml−NeuN−D3
7614−Ctx−T−L−Sml−NeuN−D4
7614−Ctx−T−L−Sml−NeuN−D5
7614−Ctx−T−L−Sml−NeuN−D6
7614−Ctx−T−L−Sml−NeuN−E1
7614−Ctx−T−L−Sml−NeuN−E2
7614−Ctx−T−L−Sml−NeuN−E3
7614−Ctx−T−L−Sml−NeuN−E4
7614−Ctx−T−L−Sml−NeuN−E5
7614−Ctx−T−L−Sml−NeuN−E6
7614−Ctx−T−L−Sml−NeuN−F1
7614−Ctx−T−L−Sml−NeuN−F2
7614−Ctx−T−L−Sml−NeuN−F3
7614−Ctx−T−L−Sml−NeuN−F4
7614−Ctx−T−L−Sml−NeuN−F5
7614−Ctx−T−L−Sml−NeuN−F6
7614−Ctx−T−L−Sml−NeuN−G1
7614−Ctx−T−L−Sml−NeuN−G2
7614−Ctx−T−L−Sml−NeuN−G3
7614−Ctx−T−L−Sml−NeuN−G4
7614−Ctx−T−L−Sml−NeuN−G5
7614−Ctx−T−L−Sml−NeuN−G6
7614−Ctx−T−L−Sml−NeuN−H1
7614−Ctx−T−L−Sml−NeuN−H2
7614−Ctx−T−L−Sml−NeuN−H3
7614−Ctx−T−L−Sml−NeuN−H4
7614−Ctx−T−L−Sml−NeuN−H5
7614−Ctx−T−L−Sml−NeuN−H6
JGG−Cells

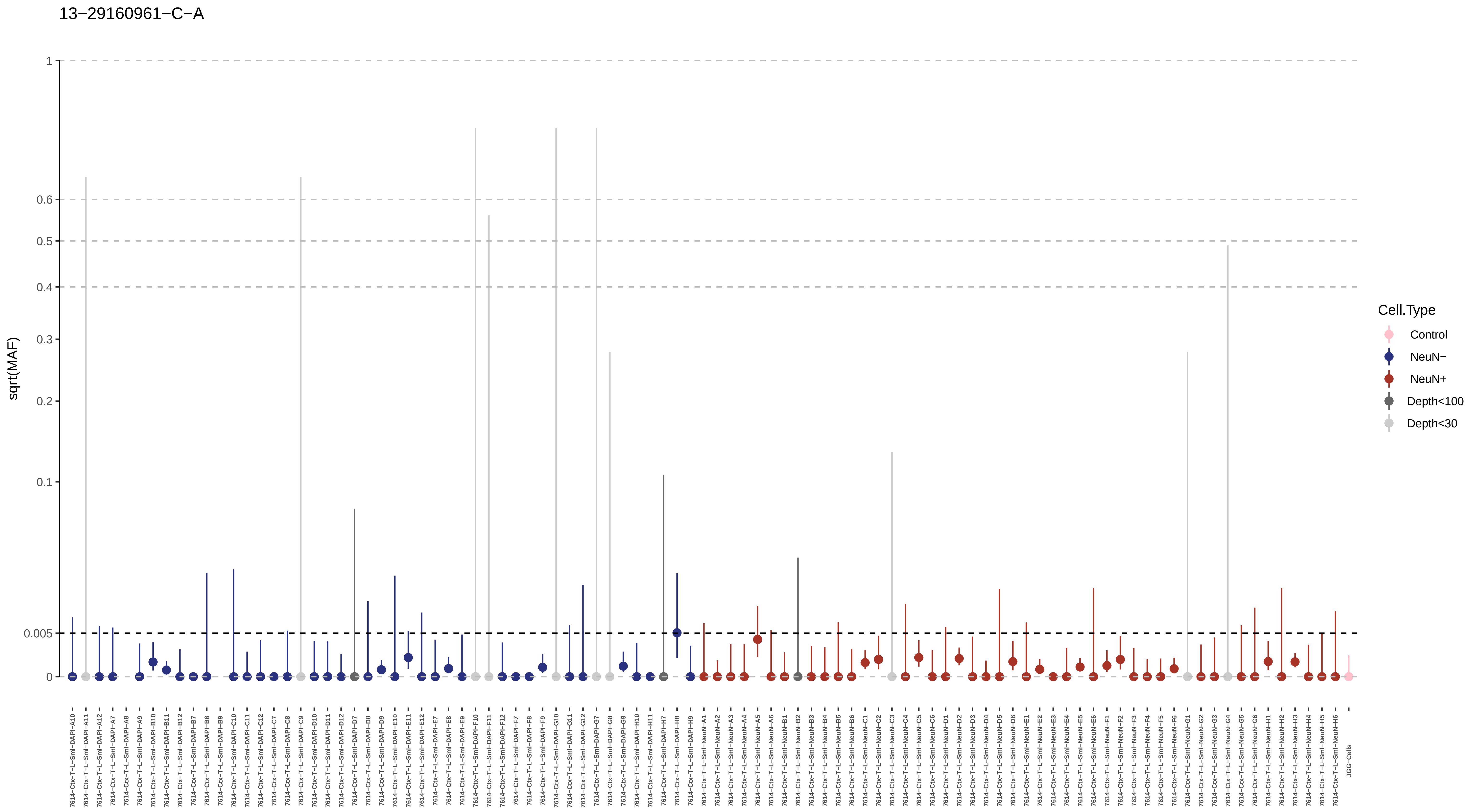

13−29160961−C−A
sqrt(MAF)
1
0.6
0.5
0.4
0.3
0.2
0.1
0.005
0
Cell.Type
Control
NeuN−
NeuN+
Depth<100
Depth<30
7614−Ctx−T−L−Sml−DAPI−A10
7614−Ctx−T−L−Sml−DAPI−A11
7614−Ctx−T−L−Sml−DAPI−A12
7614−Ctx−T−L−Sml−DAPI−A7
7614−Ctx−T−L−Sml−DAPI−A8
7614−Ctx−T−L−Sml−DAPI−A9
7614−Ctx−T−L−Sml−DAPI−B10
7614−Ctx−T−L−Sml−DAPI−B11
7614−Ctx−T−L−Sml−DAPI−B12
7614−Ctx−T−L−Sml−DAPI−B7
7614−Ctx−T−L−Sml−DAPI−B8
7614−Ctx−T−L−Sml−DAPI−B9
7614−Ctx−T−L−Sml−DAPI−C10
7614−Ctx−T−L−Sml−DAPI−C11
7614−Ctx−T−L−Sml−DAPI−C12
7614−Ctx−T−L−Sml−DAPI−C7
7614−Ctx−T−L−Sml−DAPI−C8
7614−Ctx−T−L−Sml−DAPI−C9
7614−Ctx−T−L−Sml−DAPI−D10
7614−Ctx−T−L−Sml−DAPI−D11
7614−Ctx−T−L−Sml−DAPI−D12
7614−Ctx−T−L−Sml−DAPI−D7
7614−Ctx−T−L−Sml−DAPI−D8
7614−Ctx−T−L−Sml−DAPI−D9
7614−Ctx−T−L−Sml−DAPI−E10
7614−Ctx−T−L−Sml−DAPI−E11
7614−Ctx−T−L−Sml−DAPI−E12
7614−Ctx−T−L−Sml−DAPI−E7
7614−Ctx−T−L−Sml−DAPI−E8
7614−Ctx−T−L−Sml−DAPI−E9
7614−Ctx−T−L−Sml−DAPI−F10
7614−Ctx−T−L−Sml−DAPI−F11
7614−Ctx−T−L−Sml−DAPI−F12
7614−Ctx−T−L−Sml−DAPI−F7
7614−Ctx−T−L−Sml−DAPI−F8
7614−Ctx−T−L−Sml−DAPI−F9
7614−Ctx−T−L−Sml−DAPI−G10
7614−Ctx−T−L−Sml−DAPI−G11
7614−Ctx−T−L−Sml−DAPI−G12
7614−Ctx−T−L−Sml−DAPI−G7
7614−Ctx−T−L−Sml−DAPI−G8
7614−Ctx−T−L−Sml−DAPI−G9
7614−Ctx−T−L−Sml−DAPI−H10
7614−Ctx−T−L−Sml−DAPI−H11
7614−Ctx−T−L−Sml−DAPI−H7
7614−Ctx−T−L−Sml−DAPI−H8
7614−Ctx−T−L−Sml−DAPI−H9
7614−Ctx−T−L−Sml−NeuN−A1
7614−Ctx−T−L−Sml−NeuN−A2
7614−Ctx−T−L−Sml−NeuN−A3
7614−Ctx−T−L−Sml−NeuN−A4
7614−Ctx−T−L−Sml−NeuN−A5
7614−Ctx−T−L−Sml−NeuN−A6
7614−Ctx−T−L−Sml−NeuN−B1
7614−Ctx−T−L−Sml−NeuN−B2
7614−Ctx−T−L−Sml−NeuN−B3
7614−Ctx−T−L−Sml−NeuN−B4
7614−Ctx−T−L−Sml−NeuN−B5
7614−Ctx−T−L−Sml−NeuN−B6
7614−Ctx−T−L−Sml−NeuN−C1
7614−Ctx−T−L−Sml−NeuN−C2
7614−Ctx−T−L−Sml−NeuN−C3
7614−Ctx−T−L−Sml−NeuN−C4
7614−Ctx−T−L−Sml−NeuN−C5
7614−Ctx−T−L−Sml−NeuN−C6
7614−Ctx−T−L−Sml−NeuN−D1
7614−Ctx−T−L−Sml−NeuN−D2
7614−Ctx−T−L−Sml−NeuN−D3
7614−Ctx−T−L−Sml−NeuN−D4
7614−Ctx−T−L−Sml−NeuN−D5
7614−Ctx−T−L−Sml−NeuN−D6
7614−Ctx−T−L−Sml−NeuN−E1
7614−Ctx−T−L−Sml−NeuN−E2
7614−Ctx−T−L−Sml−NeuN−E3
7614−Ctx−T−L−Sml−NeuN−E4
7614−Ctx−T−L−Sml−NeuN−E5
7614−Ctx−T−L−Sml−NeuN−E6
7614−Ctx−T−L−Sml−NeuN−F1
7614−Ctx−T−L−Sml−NeuN−F2
7614−Ctx−T−L−Sml−NeuN−F3
7614−Ctx−T−L−Sml−NeuN−F4
7614−Ctx−T−L−Sml−NeuN−F5
7614−Ctx−T−L−Sml−NeuN−F6
7614−Ctx−T−L−Sml−NeuN−G1
7614−Ctx−T−L−Sml−NeuN−G2
7614−Ctx−T−L−Sml−NeuN−G3
7614−Ctx−T−L−Sml−NeuN−G4
7614−Ctx−T−L−Sml−NeuN−G5
7614−Ctx−T−L−Sml−NeuN−G6
7614−Ctx−T−L−Sml−NeuN−H1
7614−Ctx−T−L−Sml−NeuN−H2
7614−Ctx−T−L−Sml−NeuN−H3
7614−Ctx−T−L−Sml−NeuN−H4
7614−Ctx−T−L−Sml−NeuN−H5
7614−Ctx−T−L−Sml−NeuN−H6
JGG−Cells

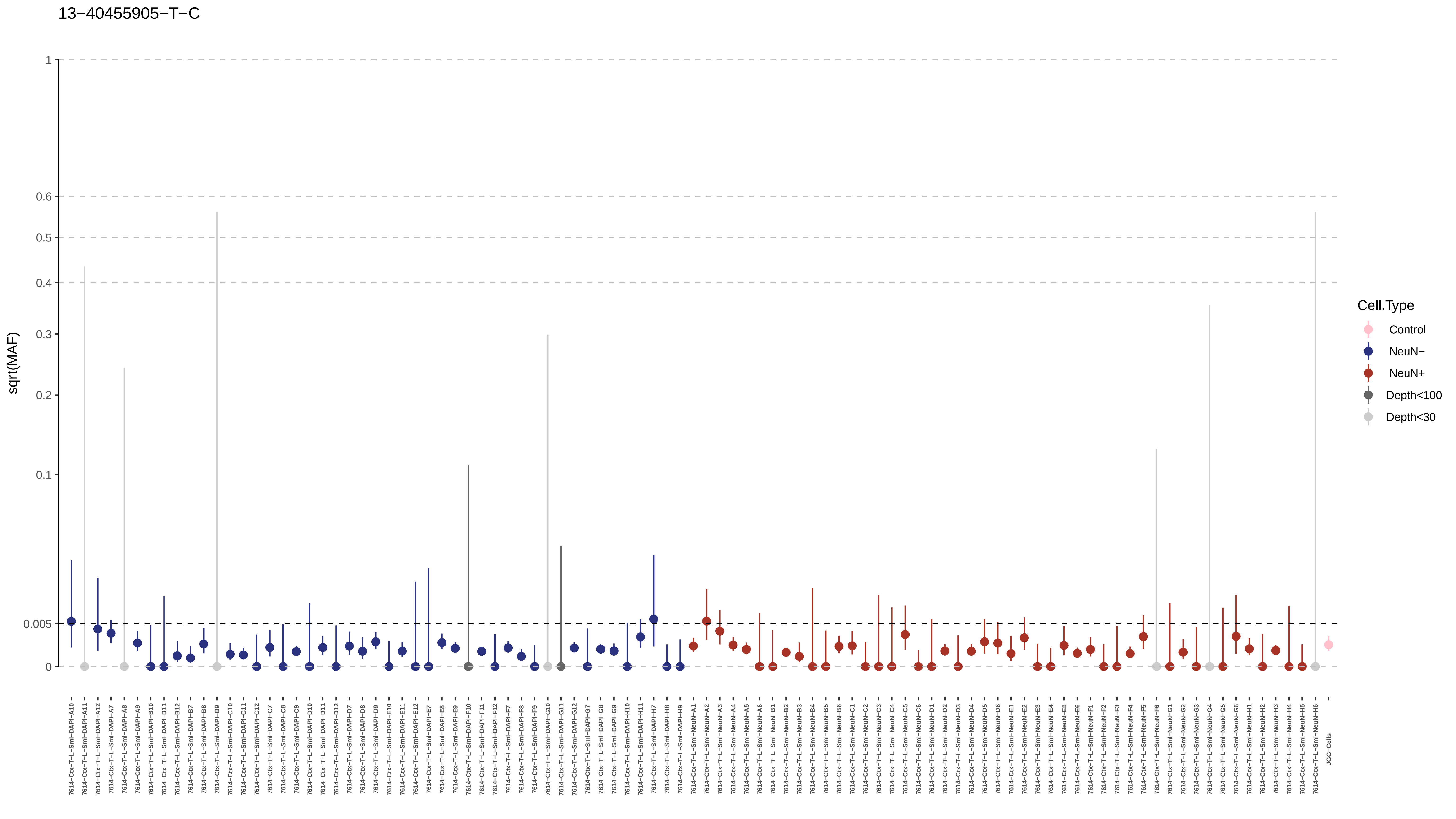
13−40455905−T−C
sqrt(MAF)
1
0.6
0.5
0.4
0.3
0.2
0.1
0.005
0
Cell.Type
Control
NeuN−
NeuN+
Depth<100
Depth<30
7614−Ctx−T−L−Sml−DAPI−A10
7614−Ctx−T−L−Sml−DAPI−A11
7614−Ctx−T−L−Sml−DAPI−A12
7614−Ctx−T−L−Sml−DAPI−A7
7614−Ctx−T−L−Sml−DAPI−A8
7614−Ctx−T−L−Sml−DAPI−A9
7614−Ctx−T−L−Sml−DAPI−B10
7614−Ctx−T−L−Sml−DAPI−B11
7614−Ctx−T−L−Sml−DAPI−B12
7614−Ctx−T−L−Sml−DAPI−B7
7614−Ctx−T−L−Sml−DAPI−B8
7614−Ctx−T−L−Sml−DAPI−B9
7614−Ctx−T−L−Sml−DAPI−C10
7614−Ctx−T−L−Sml−DAPI−C11
7614−Ctx−T−L−Sml−DAPI−C12
7614−Ctx−T−L−Sml−DAPI−C7
7614−Ctx−T−L−Sml−DAPI−C8
7614−Ctx−T−L−Sml−DAPI−C9
7614−Ctx−T−L−Sml−DAPI−D10
7614−Ctx−T−L−Sml−DAPI−D11
7614−Ctx−T−L−Sml−DAPI−D12
7614−Ctx−T−L−Sml−DAPI−D7
7614−Ctx−T−L−Sml−DAPI−D8
7614−Ctx−T−L−Sml−DAPI−D9
7614−Ctx−T−L−Sml−DAPI−E10
7614−Ctx−T−L−Sml−DAPI−E11
7614−Ctx−T−L−Sml−DAPI−E12
7614−Ctx−T−L−Sml−DAPI−E7
7614−Ctx−T−L−Sml−DAPI−E8
7614−Ctx−T−L−Sml−DAPI−E9
7614−Ctx−T−L−Sml−DAPI−F10
7614−Ctx−T−L−Sml−DAPI−F11
7614−Ctx−T−L−Sml−DAPI−F12
7614−Ctx−T−L−Sml−DAPI−F7
7614−Ctx−T−L−Sml−DAPI−F8
7614−Ctx−T−L−Sml−DAPI−F9
7614−Ctx−T−L−Sml−DAPI−G10
7614−Ctx−T−L−Sml−DAPI−G11
7614−Ctx−T−L−Sml−DAPI−G12
7614−Ctx−T−L−Sml−DAPI−G7
7614−Ctx−T−L−Sml−DAPI−G8
7614−Ctx−T−L−Sml−DAPI−G9
7614−Ctx−T−L−Sml−DAPI−H10
7614−Ctx−T−L−Sml−DAPI−H11
7614−Ctx−T−L−Sml−DAPI−H7
7614−Ctx−T−L−Sml−DAPI−H8
7614−Ctx−T−L−Sml−DAPI−H9
7614−Ctx−T−L−Sml−NeuN−A1
7614−Ctx−T−L−Sml−NeuN−A2
7614−Ctx−T−L−Sml−NeuN−A3
7614−Ctx−T−L−Sml−NeuN−A4
7614−Ctx−T−L−Sml−NeuN−A5
7614−Ctx−T−L−Sml−NeuN−A6
7614−Ctx−T−L−Sml−NeuN−B1
7614−Ctx−T−L−Sml−NeuN−B2
7614−Ctx−T−L−Sml−NeuN−B3
7614−Ctx−T−L−Sml−NeuN−B4
7614−Ctx−T−L−Sml−NeuN−B5
7614−Ctx−T−L−Sml−NeuN−B6
7614−Ctx−T−L−Sml−NeuN−C1
7614−Ctx−T−L−Sml−NeuN−C2
7614−Ctx−T−L−Sml−NeuN−C3
7614−Ctx−T−L−Sml−NeuN−C4
7614−Ctx−T−L−Sml−NeuN−C5
7614−Ctx−T−L−Sml−NeuN−C6
7614−Ctx−T−L−Sml−NeuN−D1
7614−Ctx−T−L−Sml−NeuN−D2
7614−Ctx−T−L−Sml−NeuN−D3
7614−Ctx−T−L−Sml−NeuN−D4
7614−Ctx−T−L−Sml−NeuN−D5
7614−Ctx−T−L−Sml−NeuN−D6
7614−Ctx−T−L−Sml−NeuN−E1
7614−Ctx−T−L−Sml−NeuN−E2
7614−Ctx−T−L−Sml−NeuN−E3
7614−Ctx−T−L−Sml−NeuN−E4
7614−Ctx−T−L−Sml−NeuN−E5
7614−Ctx−T−L−Sml−NeuN−E6
7614−Ctx−T−L−Sml−NeuN−F1
7614−Ctx−T−L−Sml−NeuN−F2
7614−Ctx−T−L−Sml−NeuN−F3
7614−Ctx−T−L−Sml−NeuN−F4
7614−Ctx−T−L−Sml−NeuN−F5
7614−Ctx−T−L−Sml−NeuN−F6
7614−Ctx−T−L−Sml−NeuN−G1
7614−Ctx−T−L−Sml−NeuN−G2
7614−Ctx−T−L−Sml−NeuN−G3
7614−Ctx−T−L−Sml−NeuN−G4
7614−Ctx−T−L−Sml−NeuN−G5
7614−Ctx−T−L−Sml−NeuN−G6
7614−Ctx−T−L−Sml−NeuN−H1
7614−Ctx−T−L−Sml−NeuN−H2
7614−Ctx−T−L−Sml−NeuN−H3
7614−Ctx−T−L−Sml−NeuN−H4
7614−Ctx−T−L−Sml−NeuN−H5
7614−Ctx−T−L−Sml−NeuN−H6
JGG−Cells

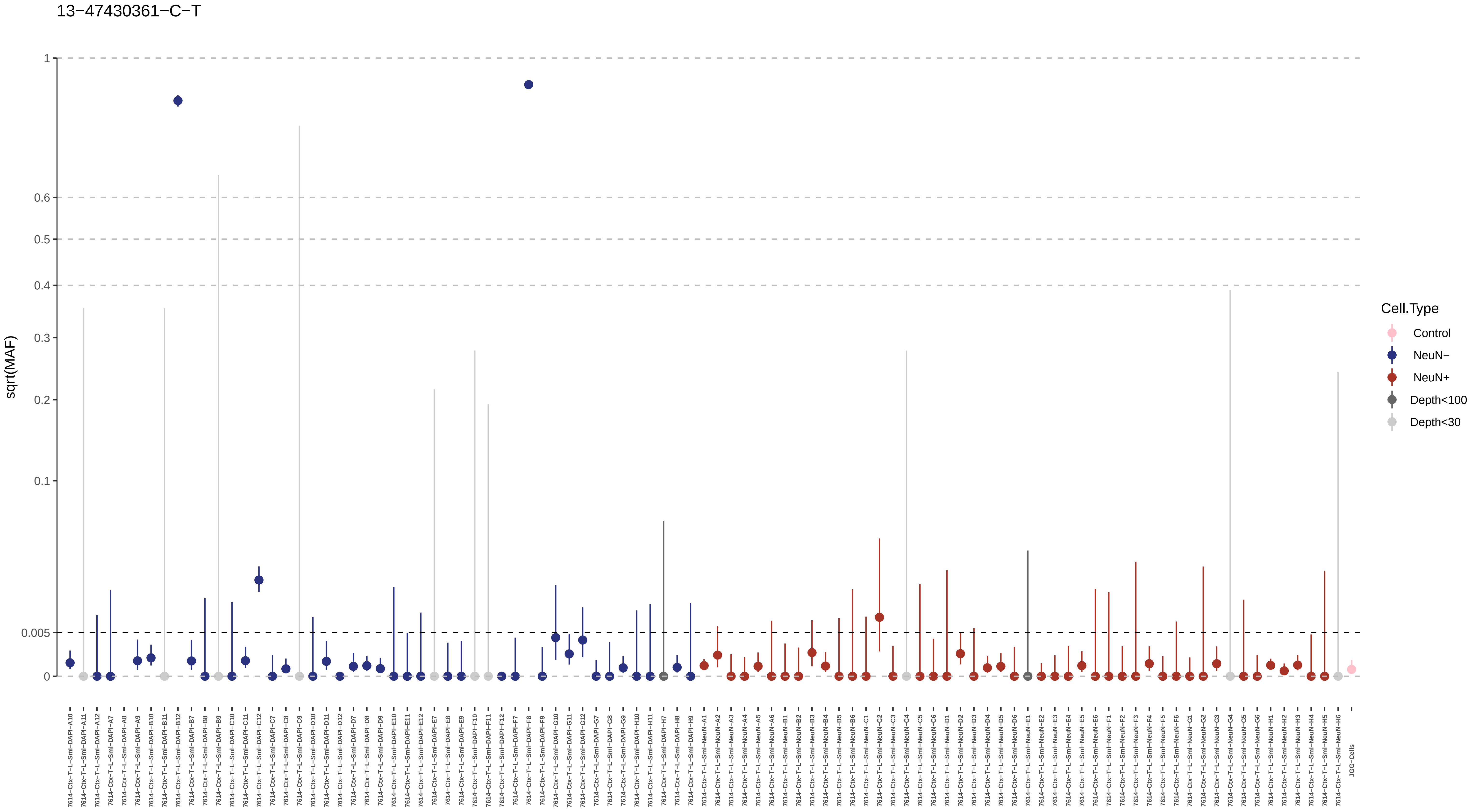
13−47430361−C−T
sqrt(MAF)
1
0.6
0.5
0.4
0.3
0.2
0.1
0.005
0
Cell.Type
Control
NeuN−
NeuN+
Depth<100
Depth<30
7614−Ctx−T−L−Sml−DAPI−A10
7614−Ctx−T−L−Sml−DAPI−A11
7614−Ctx−T−L−Sml−DAPI−A12
7614−Ctx−T−L−Sml−DAPI−A7
7614−Ctx−T−L−Sml−DAPI−A8
7614−Ctx−T−L−Sml−DAPI−A9
7614−Ctx−T−L−Sml−DAPI−B10
7614−Ctx−T−L−Sml−DAPI−B11
7614−Ctx−T−L−Sml−DAPI−B12
7614−Ctx−T−L−Sml−DAPI−B7
7614−Ctx−T−L−Sml−DAPI−B8
7614−Ctx−T−L−Sml−DAPI−B9
7614−Ctx−T−L−Sml−DAPI−C10
7614−Ctx−T−L−Sml−DAPI−C11
7614−Ctx−T−L−Sml−DAPI−C12
7614−Ctx−T−L−Sml−DAPI−C7
7614−Ctx−T−L−Sml−DAPI−C8
7614−Ctx−T−L−Sml−DAPI−C9
7614−Ctx−T−L−Sml−DAPI−D10
7614−Ctx−T−L−Sml−DAPI−D11
7614−Ctx−T−L−Sml−DAPI−D12
7614−Ctx−T−L−Sml−DAPI−D7
7614−Ctx−T−L−Sml−DAPI−D8
7614−Ctx−T−L−Sml−DAPI−D9
7614−Ctx−T−L−Sml−DAPI−E10
7614−Ctx−T−L−Sml−DAPI−E11
7614−Ctx−T−L−Sml−DAPI−E12
7614−Ctx−T−L−Sml−DAPI−E7
7614−Ctx−T−L−Sml−DAPI−E8
7614−Ctx−T−L−Sml−DAPI−E9
7614−Ctx−T−L−Sml−DAPI−F10
7614−Ctx−T−L−Sml−DAPI−F11
7614−Ctx−T−L−Sml−DAPI−F12
7614−Ctx−T−L−Sml−DAPI−F7
7614−Ctx−T−L−Sml−DAPI−F8
7614−Ctx−T−L−Sml−DAPI−F9
7614−Ctx−T−L−Sml−DAPI−G10
7614−Ctx−T−L−Sml−DAPI−G11
7614−Ctx−T−L−Sml−DAPI−G12
7614−Ctx−T−L−Sml−DAPI−G7
7614−Ctx−T−L−Sml−DAPI−G8
7614−Ctx−T−L−Sml−DAPI−G9
7614−Ctx−T−L−Sml−DAPI−H10
7614−Ctx−T−L−Sml−DAPI−H11
7614−Ctx−T−L−Sml−DAPI−H7
7614−Ctx−T−L−Sml−DAPI−H8
7614−Ctx−T−L−Sml−DAPI−H9
7614−Ctx−T−L−Sml−NeuN−A1
7614−Ctx−T−L−Sml−NeuN−A2
7614−Ctx−T−L−Sml−NeuN−A3
7614−Ctx−T−L−Sml−NeuN−A4
7614−Ctx−T−L−Sml−NeuN−A5
7614−Ctx−T−L−Sml−NeuN−A6
7614−Ctx−T−L−Sml−NeuN−B1
7614−Ctx−T−L−Sml−NeuN−B2
7614−Ctx−T−L−Sml−NeuN−B3
7614−Ctx−T−L−Sml−NeuN−B4
7614−Ctx−T−L−Sml−NeuN−B5
7614−Ctx−T−L−Sml−NeuN−B6
7614−Ctx−T−L−Sml−NeuN−C1
7614−Ctx−T−L−Sml−NeuN−C2
7614−Ctx−T−L−Sml−NeuN−C3
7614−Ctx−T−L−Sml−NeuN−C4
7614−Ctx−T−L−Sml−NeuN−C5
7614−Ctx−T−L−Sml−NeuN−C6
7614−Ctx−T−L−Sml−NeuN−D1
7614−Ctx−T−L−Sml−NeuN−D2
7614−Ctx−T−L−Sml−NeuN−D3
7614−Ctx−T−L−Sml−NeuN−D4
7614−Ctx−T−L−Sml−NeuN−D5
7614−Ctx−T−L−Sml−NeuN−D6
7614−Ctx−T−L−Sml−NeuN−E1
7614−Ctx−T−L−Sml−NeuN−E2
7614−Ctx−T−L−Sml−NeuN−E3
7614−Ctx−T−L−Sml−NeuN−E4
7614−Ctx−T−L−Sml−NeuN−E5
7614−Ctx−T−L−Sml−NeuN−E6
7614−Ctx−T−L−Sml−NeuN−F1
7614−Ctx−T−L−Sml−NeuN−F2
7614−Ctx−T−L−Sml−NeuN−F3
7614−Ctx−T−L−Sml−NeuN−F4
7614−Ctx−T−L−Sml−NeuN−F5
7614−Ctx−T−L−Sml−NeuN−F6
7614−Ctx−T−L−Sml−NeuN−G1
7614−Ctx−T−L−Sml−NeuN−G2
7614−Ctx−T−L−Sml−NeuN−G3
7614−Ctx−T−L−Sml−NeuN−G4
7614−Ctx−T−L−Sml−NeuN−G5
7614−Ctx−T−L−Sml−NeuN−G6
7614−Ctx−T−L−Sml−NeuN−H1
7614−Ctx−T−L−Sml−NeuN−H2
7614−Ctx−T−L−Sml−NeuN−H3
7614−Ctx−T−L−Sml−NeuN−H4
7614−Ctx−T−L−Sml−NeuN−H5
7614−Ctx−T−L−Sml−NeuN−H6
JGG-Cells

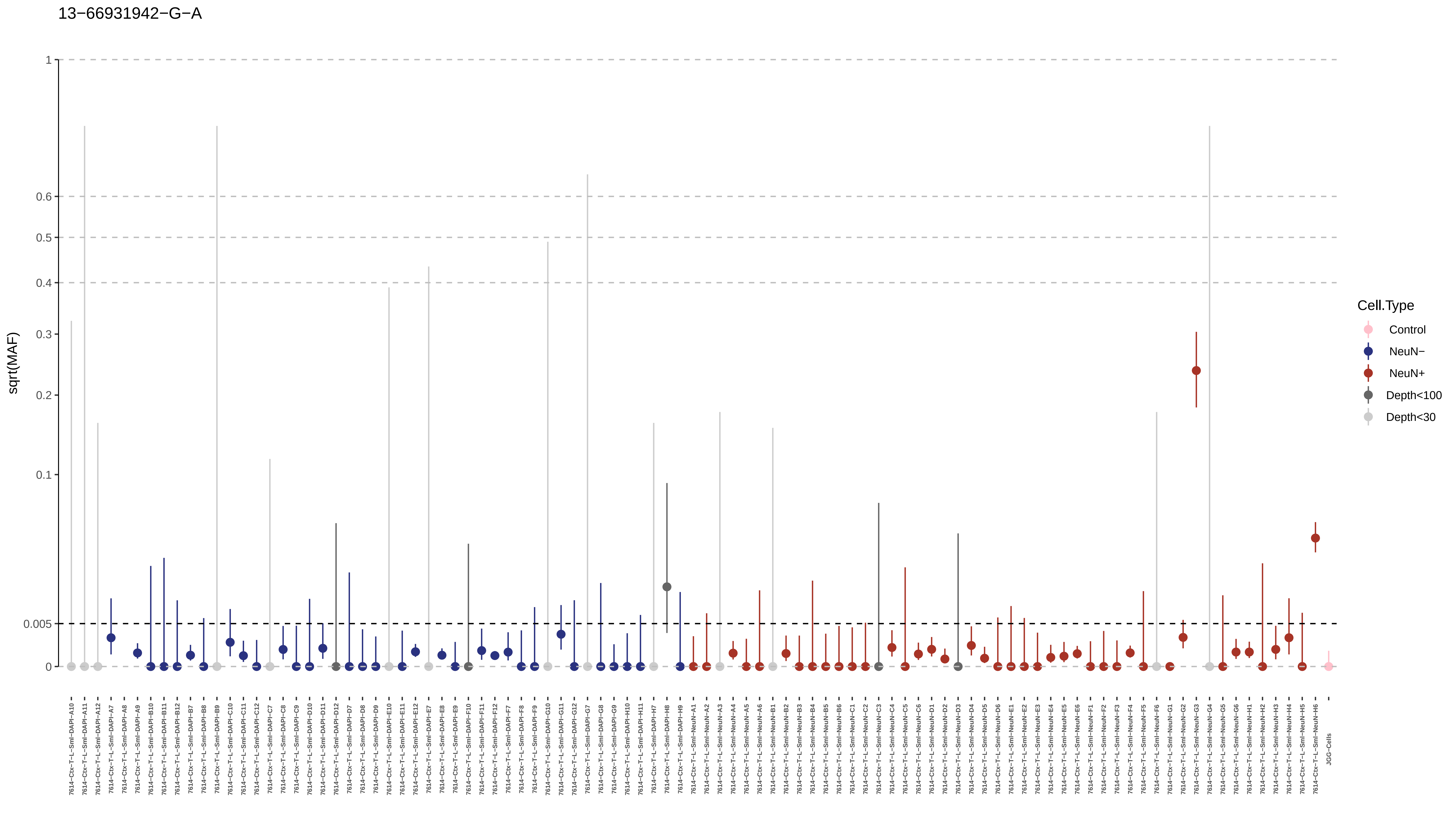
13−66931942−G−A
sqrt(MAF)
1
0.6
0.5
0.4
0.3
0.2
0.1
0.005
0
Cell.Type
Control
NeuN−
NeuN+
Depth<100
Depth<30

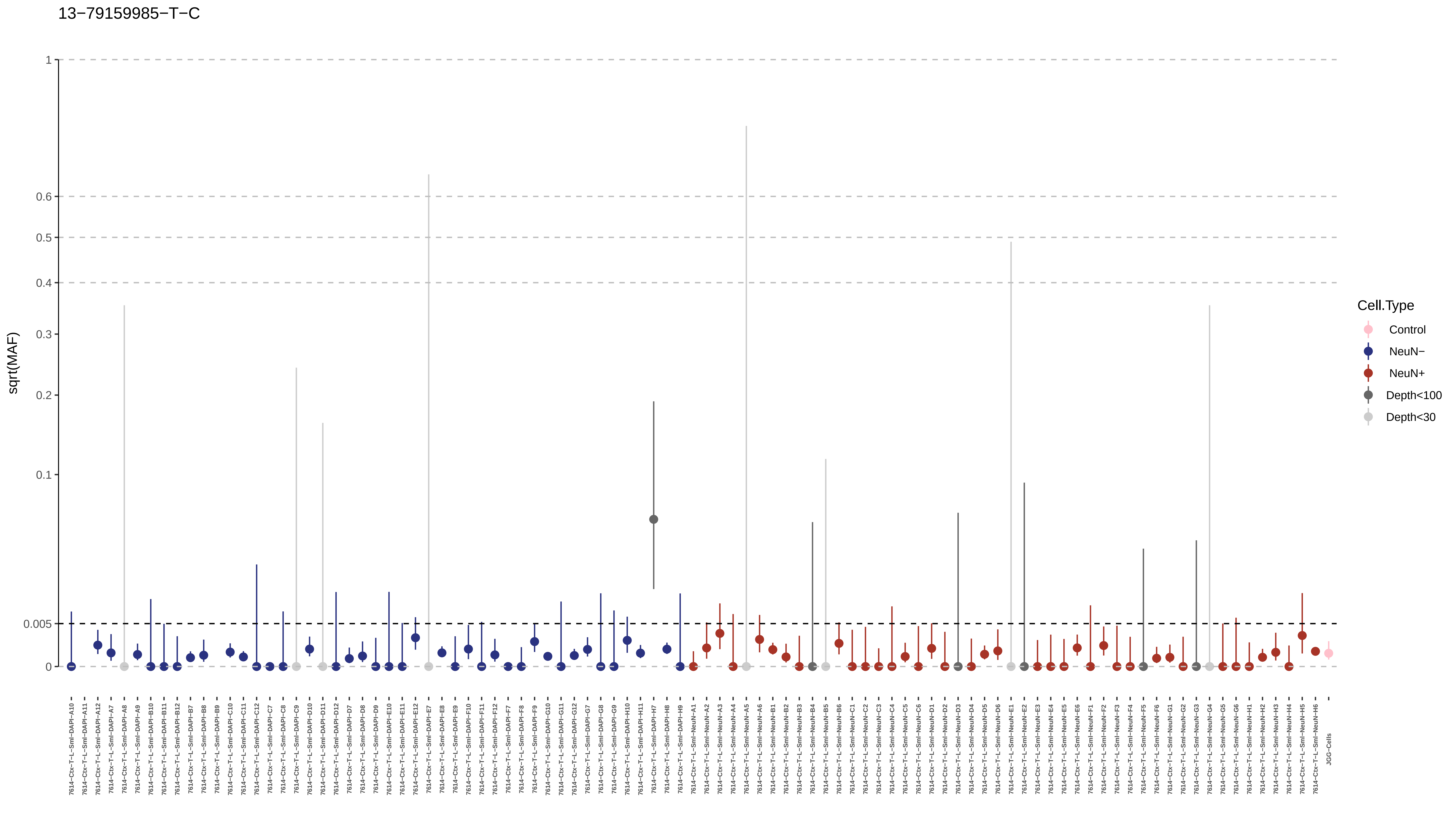

13−79159985−T−C
sqrt(MAF)
1
0.6
0.5
0.4
0.3
0.2
0.1
0.005
0
Cell.Type
Control
NeuN−
NeuN+
Depth<100
Depth<30
7614−Ctx−T−L−Sml−DAPI−A10
7614−Ctx−T−L−Sml−DAPI−A11
7614−Ctx−T−L−Sml−DAPI−A12
7614−Ctx−T−L−Sml−DAPI−A7
7614−Ctx−T−L−Sml−DAPI−A8
7614−Ctx−T−L−Sml−DAPI−A9
7614−Ctx−T−L−Sml−DAPI−B10
7614−Ctx−T−L−Sml−DAPI−B11
7614−Ctx−T−L−Sml−DAPI−B12
7614−Ctx−T−L−Sml−DAPI−B7
7614−Ctx−T−L−Sml−DAPI−B8
7614−Ctx−T−L−Sml−DAPI−B9
7614−Ctx−T−L−Sml−DAPI−C10
7614−Ctx−T−L−Sml−DAPI−C11
7614−Ctx−T−L−Sml−DAPI−C12
7614−Ctx−T−L−Sml−DAPI−C7
7614−Ctx−T−L−Sml−DAPI−C8
7614−Ctx−T−L−Sml−DAPI−C9
7614−Ctx−T−L−Sml−DAPI−D10
7614−Ctx−T−L−Sml−DAPI−D11
7614−Ctx−T−L−Sml−DAPI−D12
7614−Ctx−T−L−Sml−DAPI−D7
7614−Ctx−T−L−Sml−DAPI−D8
7614−Ctx−T−L−Sml−DAPI−D9
7614−Ctx−T−L−Sml−DAPI−E10
7614−Ctx−T−L−Sml−DAPI−E11
7614−Ctx−T−L−Sml−DAPI−E12
7614−Ctx−T−L−Sml−DAPI−E7
7614−Ctx−T−L−Sml−DAPI−E8
7614−Ctx−T−L−Sml−DAPI−E9
7614−Ctx−T−L−Sml−DAPI−F10
7614−Ctx−T−L−Sml−DAPI−F11
7614−Ctx−T−L−Sml−DAPI−F12
7614−Ctx−T−L−Sml−DAPI−F7
7614−Ctx−T−L−Sml−DAPI−F8
7614−Ctx−T−L−Sml−DAPI−F9
7614−Ctx−T−L−Sml−DAPI−G10
7614−Ctx−T−L−Sml−DAPI−G11
7614−Ctx−T−L−Sml−DAPI−G12
7614−Ctx−T−L−Sml−DAPI−G7
7614−Ctx−T−L−Sml−DAPI−G8
7614−Ctx−T−L−Sml−DAPI−G9
7614−Ctx−T−L−Sml−DAPI−H10
7614−Ctx−T−L−Sml−DAPI−H11
7614−Ctx−T−L−Sml−DAPI−H7
7614−Ctx−T−L−Sml−DAPI−H8
7614−Ctx−T−L−Sml−DAPI−H9
7614−Ctx−T−L−Sml−NeuN−A1
7614−Ctx−T−L−Sml−NeuN−A2
7614−Ctx−T−L−Sml−NeuN−A3
7614−Ctx−T−L−Sml−NeuN−A4
7614−Ctx−T−L−Sml−NeuN−A5
7614−Ctx−T−L−Sml−NeuN−A6
7614−Ctx−T−L−Sml−NeuN−B1
7614−Ctx−T−L−Sml−NeuN−B2
7614−Ctx−T−L−Sml−NeuN−B3
7614−Ctx−T−L−Sml−NeuN−B4
7614−Ctx−T−L−Sml−NeuN−B5
7614−Ctx−T−L−Sml−NeuN−B6
7614−Ctx−T−L−Sml−NeuN−C1
7614−Ctx−T−L−Sml−NeuN−C2
7614−Ctx−T−L−Sml−NeuN−C3
7614−Ctx−T−L−Sml−NeuN−C4
7614−Ctx−T−L−Sml−NeuN−C5
7614−Ctx−T−L−Sml−NeuN−C6
7614−Ctx−T−L−Sml−NeuN−D1
7614−Ctx−T−L−Sml−NeuN−D2
7614−Ctx−T−L−Sml−NeuN−D3
7614−Ctx−T−L−Sml−NeuN−D4
7614−Ctx−T−L−Sml−NeuN−D5
7614−Ctx−T−L−Sml−NeuN−D6
7614−Ctx−T−L−Sml−NeuN−E1
7614−Ctx−T−L−Sml−NeuN−E2
7614−Ctx−T−L−Sml−NeuN−E3
7614−Ctx−T−L−Sml−NeuN−E4
7614−Ctx−T−L−Sml−NeuN−E5
7614−Ctx−T−L−Sml−NeuN−E6
7614−Ctx−T−L−Sml−NeuN−F1
7614−Ctx−T−L−Sml−NeuN−F2
7614−Ctx−T−L−Sml−NeuN−F3
7614−Ctx−T−L−Sml−NeuN−F4
7614−Ctx−T−L−Sml−NeuN−F5
7614−Ctx−T−L−Sml−NeuN−F6
7614−Ctx−T−L−Sml−NeuN−G1
7614−Ctx−T−L−Sml−NeuN−G2
7614−Ctx−T−L−Sml−NeuN−G3
7614−Ctx−T−L−Sml−NeuN−G4
7614−Ctx−T−L−Sml−NeuN−G5
7614−Ctx−T−L−Sml−NeuN−G6
7614−Ctx−T−L−Sml−NeuN−H1
7614−Ctx−T−L−Sml−NeuN−H2
7614−Ctx−T−L−Sml−NeuN−H3
7614−Ctx−T−L−Sml−NeuN−H4
7614−Ctx−T−L−Sml−NeuN−H5
7614−Ctx−T−L−Sml−NeuN−H6
JGG-Cells

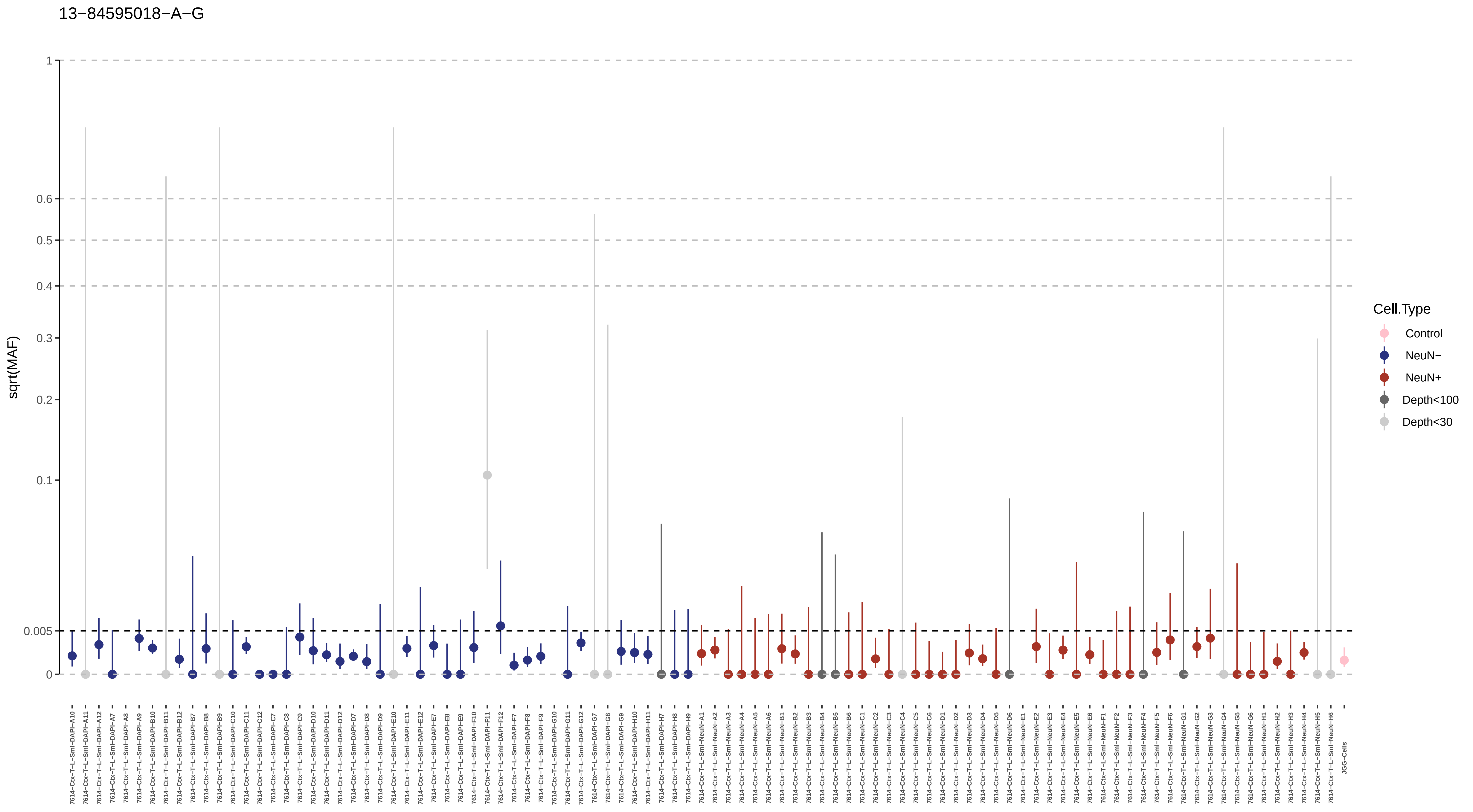
13−84595018−A−G
sqrt(MAF)
1
0.6
0.5
0.4
0.3
0.2
0.1
0.005
0
Cell.Type
Control
NeuN−
NeuN+
Depth<100
Depth<30
7614−Ctx−T−L−Sml−DAPI−A10
7614−Ctx−T−L−Sml−DAPI−A11
7614−Ctx−T−L−Sml−DAPI−A12
7614−Ctx−T−L−Sml−DAPI−A7
7614−Ctx−T−L−Sml−DAPI−A8
7614−Ctx−T−L−Sml−DAPI−A9
7614−Ctx−T−L−Sml−DAPI−B10
7614−Ctx−T−L−Sml−DAPI−B11
7614−Ctx−T−L−Sml−DAPI−B12
7614−Ctx−T−L−Sml−DAPI−B7
7614−Ctx−T−L−Sml−DAPI−B8
7614−Ctx−T−L−Sml−DAPI−B9
7614−Ctx−T−L−Sml−DAPI−C10
7614−Ctx−T−L−Sml−DAPI−C11
7614−Ctx−T−L−Sml−DAPI−C12
7614−Ctx−T−L−Sml−DAPI−C7
7614−Ctx−T−L−Sml−DAPI−C8
7614−Ctx−T−L−Sml−DAPI−C9
7614−Ctx−T−L−Sml−DAPI−D10
7614−Ctx−T−L−Sml−DAPI−D11
7614−Ctx−T−L−Sml−DAPI−D12
7614−Ctx−T−L−Sml−DAPI−D7
7614−Ctx−T−L−Sml−DAPI−D8
7614−Ctx−T−L−Sml−DAPI−D9
7614−Ctx−T−L−Sml−DAPI−E10
7614−Ctx−T−L−Sml−DAPI−E11
7614−Ctx−T−L−Sml−DAPI−E12
7614−Ctx−T−L−Sml−DAPI−E7
7614−Ctx−T−L−Sml−DAPI−E8
7614−Ctx−T−L−Sml−DAPI−E9
7614−Ctx−T−L−Sml−DAPI−F10
7614−Ctx−T−L−Sml−DAPI−F11
7614−Ctx−T−L−Sml−DAPI−F12
7614−Ctx−T−L−Sml−DAPI−F7
7614−Ctx−T−L−Sml−DAPI−F8
7614−Ctx−T−L−Sml−DAPI−F9
7614−Ctx−T−L−Sml−DAPI−G10
7614−Ctx−T−L−Sml−DAPI−G11
7614−Ctx−T−L−Sml−DAPI−G12
7614−Ctx−T−L−Sml−DAPI−G7
7614−Ctx−T−L−Sml−DAPI−G8
7614−Ctx−T−L−Sml−DAPI−G9
7614−Ctx−T−L−Sml−DAPI−H10
7614−Ctx−T−L−Sml−DAPI−H11
7614−Ctx−T−L−Sml−DAPI−H7
7614−Ctx−T−L−Sml−DAPI−H8
7614−Ctx−T−L−Sml−DAPI−H9
7614−Ctx−T−L−Sml−NeuN−A1
7614−Ctx−T−L−Sml−NeuN−A2
7614−Ctx−T−L−Sml−NeuN−A3
7614−Ctx−T−L−Sml−NeuN−A4
7614−Ctx−T−L−Sml−NeuN−A5
7614−Ctx−T−L−Sml−NeuN−A6
7614−Ctx−T−L−Sml−NeuN−B1
7614−Ctx−T−L−Sml−NeuN−B2
7614−Ctx−T−L−Sml−NeuN−B3
7614−Ctx−T−L−Sml−NeuN−B4
7614−Ctx−T−L−Sml−NeuN−B5
7614−Ctx−T−L−Sml−NeuN−B6
7614−Ctx−T−L−Sml−NeuN−C1
7614−Ctx−T−L−Sml−NeuN−C2
7614−Ctx−T−L−Sml−NeuN−C3
7614−Ctx−T−L−Sml−NeuN−C4
7614−Ctx−T−L−Sml−NeuN−C5
7614−Ctx−T−L−Sml−NeuN−C6
7614−Ctx−T−L−Sml−NeuN−D1
7614−Ctx−T−L−Sml−NeuN−D2
7614−Ctx−T−L−Sml−NeuN−D3
7614−Ctx−T−L−Sml−NeuN−D4
7614−Ctx−T−L−Sml−NeuN−D5
7614−Ctx−T−L−Sml−NeuN−D6
7614−Ctx−T−L−Sml−NeuN−E1
7614−Ctx−T−L−Sml−NeuN−E2
7614−Ctx−T−L−Sml−NeuN−E3
7614−Ctx−T−L−Sml−NeuN−E4
7614−Ctx−T−L−Sml−NeuN−E5
7614−Ctx−T−L−Sml−NeuN−E6
7614−Ctx−T−L−Sml−NeuN−F1
7614−Ctx−T−L−Sml−NeuN−F2
7614−Ctx−T−L−Sml−NeuN−F3
7614−Ctx−T−L−Sml−NeuN−F4
7614−Ctx−T−L−Sml−NeuN−F5
7614−Ctx−T−L−Sml−NeuN−F6
7614−Ctx−T−L−Sml−NeuN−G1
7614−Ctx−T−L−Sml−NeuN−G2
7614−Ctx−T−L−Sml−NeuN−G3
7614−Ctx−T−L−Sml−NeuN−G4
7614−Ctx−T−L−Sml−NeuN−G5
7614−Ctx−T−L−Sml−NeuN−G6
7614−Ctx−T−L−Sml−NeuN−H1
7614−Ctx−T−L−Sml−NeuN−H2
7614−Ctx−T−L−Sml−NeuN−H3
7614−Ctx−T−L−Sml−NeuN−H4
7614−Ctx−T−L−Sml−NeuN−H5
7614−Ctx−T−L−Sml−NeuN−H6
JGG−Cells

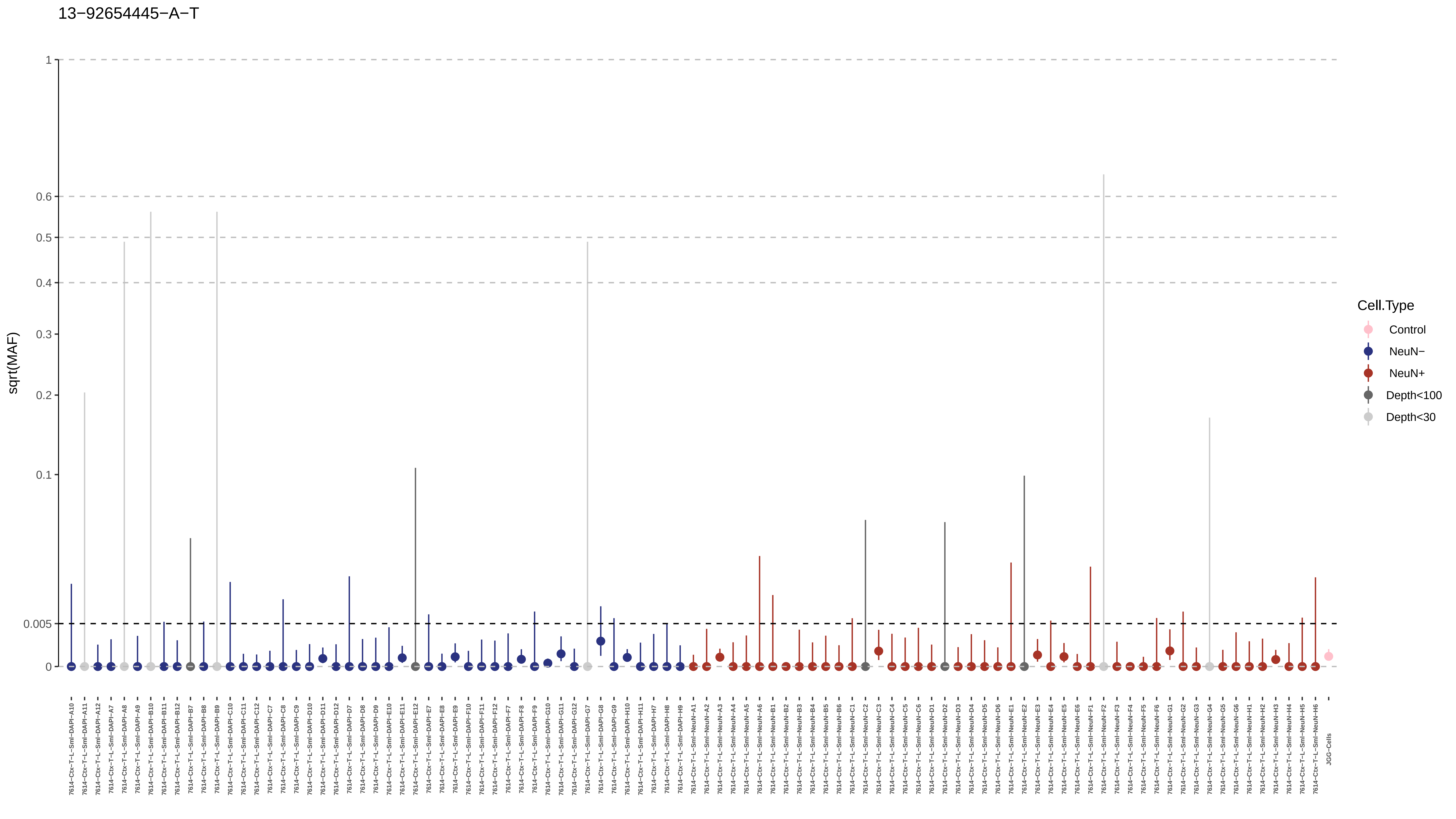
13−92654445−A−T
sqrt(MAF)
1
0.6
0.5
0.4
0.3
0.2
0.1
0.005
0
Cell.Type
Control
NeuN−
NeuN+
Depth<100
Depth<30
7614−Ctx−T−L−Sml−DAPI−A10
7614−Ctx−T−L−Sml−DAPI−A11
7614−Ctx−T−L−Sml−DAPI−A12
7614−Ctx−T−L−Sml−DAPI−A7
7614−Ctx−T−L−Sml−DAPI−A8
7614−Ctx−T−L−Sml−DAPI−A9
7614−Ctx−T−L−Sml−DAPI−B10
7614−Ctx−T−L−Sml−DAPI−B11
7614−Ctx−T−L−Sml−DAPI−B12
7614−Ctx−T−L−Sml−DAPI−B7
7614−Ctx−T−L−Sml−DAPI−B8
7614−Ctx−T−L−Sml−DAPI−B9
7614−Ctx−T−L−Sml−DAPI−C10
7614−Ctx−T−L−Sml−DAPI−C11
7614−Ctx−T−L−Sml−DAPI−C12
7614−Ctx−T−L−Sml−DAPI−C7
7614−Ctx−T−L−Sml−DAPI−C8
7614−Ctx−T−L−Sml−DAPI−C9
7614−Ctx−T−L−Sml−DAPI−D10
7614−Ctx−T−L−Sml−DAPI−D11
7614−Ctx−T−L−Sml−DAPI−D12
7614−Ctx−T−L−Sml−DAPI−D7
7614−Ctx−T−L−Sml−DAPI−D8
7614−Ctx−T−L−Sml−DAPI−D9
7614−Ctx−T−L−Sml−DAPI−E10
7614−Ctx−T−L−Sml−DAPI−E11
7614−Ctx−T−L−Sml−DAPI−E12
7614−Ctx−T−L−Sml−DAPI−E7
7614−Ctx−T−L−Sml−DAPI−E8
7614−Ctx−T−L−Sml−DAPI−E9
7614−Ctx−T−L−Sml−DAPI−F10
7614−Ctx−T−L−Sml−DAPI−F11
7614−Ctx−T−L−Sml−DAPI−F12
7614−Ctx−T−L−Sml−DAPI−F7
7614−Ctx−T−L−Sml−DAPI−F8
7614−Ctx−T−L−Sml−DAPI−F9
7614−Ctx−T−L−Sml−DAPI−G10
7614−Ctx−T−L−Sml−DAPI−G11
7614−Ctx−T−L−Sml−DAPI−G12
7614−Ctx−T−L−Sml−DAPI−G7
7614−Ctx−T−L−Sml−DAPI−G8
7614−Ctx−T−L−Sml−DAPI−G9
7614−Ctx−T−L−Sml−DAPI−H10
7614−Ctx−T−L−Sml−DAPI−H11
7614−Ctx−T−L−Sml−DAPI−H7
7614−Ctx−T−L−Sml−DAPI−H8
7614−Ctx−T−L−Sml−DAPI−H9
7614−Ctx−T−L−Sml−NeuN−A1
7614−Ctx−T−L−Sml−NeuN−A2
7614−Ctx−T−L−Sml−NeuN−A3
7614−Ctx−T−L−Sml−NeuN−A4
7614−Ctx−T−L−Sml−NeuN−A5
7614−Ctx−T−L−Sml−NeuN−A6
7614−Ctx−T−L−Sml−NeuN−B1
7614−Ctx−T−L−Sml−NeuN−B2
7614−Ctx−T−L−Sml−NeuN−B3
7614−Ctx−T−L−Sml−NeuN−B4
7614−Ctx−T−L−Sml−NeuN−B5
7614−Ctx−T−L−Sml−NeuN−B6
7614−Ctx−T−L−Sml−NeuN−C1
7614−Ctx−T−L−Sml−NeuN−C2
7614−Ctx−T−L−Sml−NeuN−C3
7614−Ctx−T−L−Sml−NeuN−C4
7614−Ctx−T−L−Sml−NeuN−C5
7614−Ctx−T−L−Sml−NeuN−C6
7614−Ctx−T−L−Sml−NeuN−D1
7614−Ctx−T−L−Sml−NeuN−D2
7614−Ctx−T−L−Sml−NeuN−D3
7614−Ctx−T−L−Sml−NeuN−D4
7614−Ctx−T−L−Sml−NeuN−D5
7614−Ctx−T−L−Sml−NeuN−D6
7614−Ctx−T−L−Sml−NeuN−E1
7614−Ctx−T−L−Sml−NeuN−E2
7614−Ctx−T−L−Sml−NeuN−E3
7614−Ctx−T−L−Sml−NeuN−E4
7614−Ctx−T−L−Sml−NeuN−E5
7614−Ctx−T−L−Sml−NeuN−E6
7614−Ctx−T−L−Sml−NeuN−F1
7614−Ctx−T−L−Sml−NeuN−F2
7614−Ctx−T−L−Sml−NeuN−F3
7614−Ctx−T−L−Sml−NeuN−F4
7614−Ctx−T−L−Sml−NeuN−F5
7614−Ctx−T−L−Sml−NeuN−F6
7614−Ctx−T−L−Sml−NeuN−G1
7614−Ctx−T−L−Sml−NeuN−G2
7614−Ctx−T−L−Sml−NeuN−G3
7614−Ctx−T−L−Sml−NeuN−G4
7614−Ctx−T−L−Sml−NeuN−G5
7614−Ctx−T−L−Sml−NeuN−G6
7614−Ctx−T−L−Sml−NeuN−H1
7614−Ctx−T−L−Sml−NeuN−H2
7614−Ctx−T−L−Sml−NeuN−H3
7614−Ctx−T−L−Sml−NeuN−H4
7614−Ctx−T−L−Sml−NeuN−H5
7614−Ctx−T−L−Sml−NeuN−H6
JGG-Cells

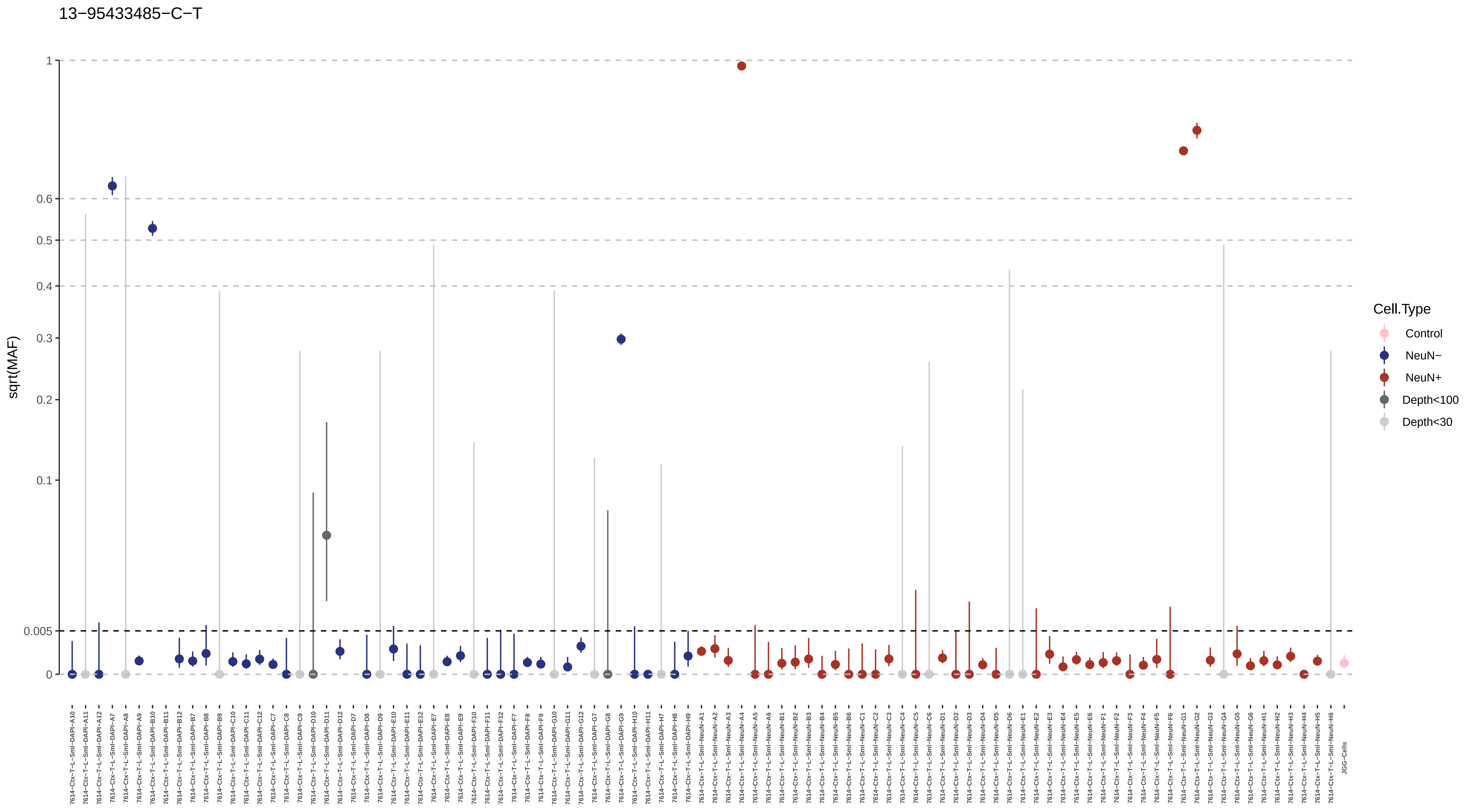

13−95433485−C−T
sqrt(MAF)
1
0.6
0.5
0.4
0.3
0.2
0.1
0.005
0
Cell.Type
Control
NeuN−
NeuN+
Depth<100
Depth<30

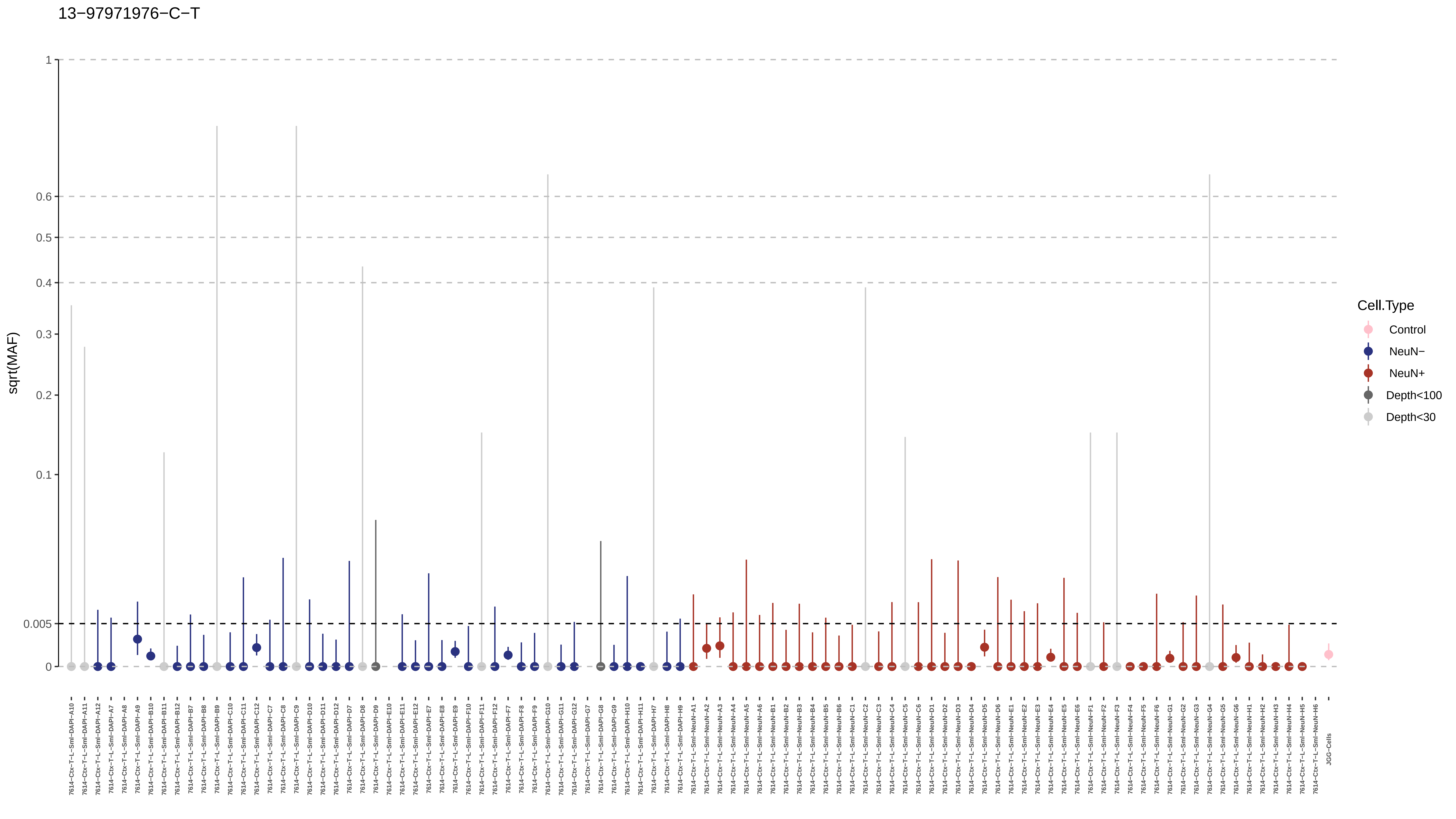

13−97971976−C−T
sqrt(MAF)
1
0.6
0.5
0.4
0.3
0.2
0.1
0.005
0
Cell.Type
Control
NeuN−
NeuN+
Depth<100
Depth<30
7614−Ctx−T−L−Sml−DAPI−A10
7614−Ctx−T−L−Sml−DAPI−A11
7614−Ctx−T−L−Sml−DAPI−A12
7614−Ctx−T−L−Sml−DAPI−A7
7614−Ctx−T−L−Sml−DAPI−A8
7614−Ctx−T−L−Sml−DAPI−A9
7614−Ctx−T−L−Sml−DAPI−B10
7614−Ctx−T−L−Sml−DAPI−B11
7614−Ctx−T−L−Sml−DAPI−B12
7614−Ctx−T−L−Sml−DAPI−B7
7614−Ctx−T−L−Sml−DAPI−B8
7614−Ctx−T−L−Sml−DAPI−B9
7614−Ctx−T−L−Sml−DAPI−C10
7614−Ctx−T−L−Sml−DAPI−C11
7614−Ctx−T−L−Sml−DAPI−C12
7614−Ctx−T−L−Sml−DAPI−C7
7614−Ctx−T−L−Sml−DAPI−C8
7614−Ctx−T−L−Sml−DAPI−C9
7614−Ctx−T−L−Sml−DAPI−D10
7614−Ctx−T−L−Sml−DAPI−D11
7614−Ctx−T−L−Sml−DAPI−D12
7614−Ctx−T−L−Sml−DAPI−D7
7614−Ctx−T−L−Sml−DAPI−D8
7614−Ctx−T−L−Sml−DAPI−D9
7614−Ctx−T−L−Sml−DAPI−E10
7614−Ctx−T−L−Sml−DAPI−E11
7614−Ctx−T−L−Sml−DAPI−E12
7614−Ctx−T−L−Sml−DAPI−E7
7614−Ctx−T−L−Sml−DAPI−E8
7614−Ctx−T−L−Sml−DAPI−E9
7614−Ctx−T−L−Sml−DAPI−F10
7614−Ctx−T−L−Sml−DAPI−F11
7614−Ctx−T−L−Sml−DAPI−F12
7614−Ctx−T−L−Sml−DAPI−F7
7614−Ctx−T−L−Sml−DAPI−F8
7614−Ctx−T−L−Sml−DAPI−F9
7614−Ctx−T−L−Sml−DAPI−G10
7614−Ctx−T−L−Sml−DAPI−G11
7614−Ctx−T−L−Sml−DAPI−G12
7614−Ctx−T−L−Sml−DAPI−G7
7614−Ctx−T−L−Sml−DAPI−G8
7614−Ctx−T−L−Sml−DAPI−G9
7614−Ctx−T−L−Sml−DAPI−H10
7614−Ctx−T−L−Sml−DAPI−H11
7614−Ctx−T−L−Sml−DAPI−H7
7614−Ctx−T−L−Sml−DAPI−H8
7614−Ctx−T−L−Sml−DAPI−H9
7614−Ctx−T−L−Sml−NeuN−A1
7614−Ctx−T−L−Sml−NeuN−A2
7614−Ctx−T−L−Sml−NeuN−A3
7614−Ctx−T−L−Sml−NeuN−A4
7614−Ctx−T−L−Sml−NeuN−A5
7614−Ctx−T−L−Sml−NeuN−A6
7614−Ctx−T−L−Sml−NeuN−B1
7614−Ctx−T−L−Sml−NeuN−B2
7614−Ctx−T−L−Sml−NeuN−B3
7614−Ctx−T−L−Sml−NeuN−B4
7614−Ctx−T−L−Sml−NeuN−B5
7614−Ctx−T−L−Sml−NeuN−B6
7614−Ctx−T−L−Sml−NeuN−C1
7614−Ctx−T−L−Sml−NeuN−C2
7614−Ctx−T−L−Sml−NeuN−C3
7614−Ctx−T−L−Sml−NeuN−C4
7614−Ctx−T−L−Sml−NeuN−C5
7614−Ctx−T−L−Sml−NeuN−C6
7614−Ctx−T−L−Sml−NeuN−D1
7614−Ctx−T−L−Sml−NeuN−D2
7614−Ctx−T−L−Sml−NeuN−D3
7614−Ctx−T−L−Sml−NeuN−D4
7614−Ctx−T−L−Sml−NeuN−D5
7614−Ctx−T−L−Sml−NeuN−D6
7614−Ctx−T−L−Sml−NeuN−E1
7614−Ctx−T−L−Sml−NeuN−E2
7614−Ctx−T−L−Sml−NeuN−E3
7614−Ctx−T−L−Sml−NeuN−E4
7614−Ctx−T−L−Sml−NeuN−E5
7614−Ctx−T−L−Sml−NeuN−E6
7614−Ctx−T−L−Sml−NeuN−F1
7614−Ctx−T−L−Sml−NeuN−F2
7614−Ctx−T−L−Sml−NeuN−F3
7614−Ctx−T−L−Sml−NeuN−F4
7614−Ctx−T−L−Sml−NeuN−F5
7614−Ctx−T−L−Sml−NeuN−F6
7614−Ctx−T−L−Sml−NeuN−G1
7614−Ctx−T−L−Sml−NeuN−G2
7614−Ctx−T−L−Sml−NeuN−G3
7614−Ctx−T−L−Sml−NeuN−G4
7614−Ctx−T−L−Sml−NeuN−G5
7614−Ctx−T−L−Sml−NeuN−G6
7614−Ctx−T−L−Sml−NeuN−H1
7614−Ctx−T−L−Sml−NeuN−H2
7614−Ctx−T−L−Sml−NeuN−H3
7614−Ctx−T−L−Sml−NeuN−H4
7614−Ctx−T−L−Sml−NeuN−H5
7614−Ctx−T−L−Sml−NeuN−H6
JGG-Cells

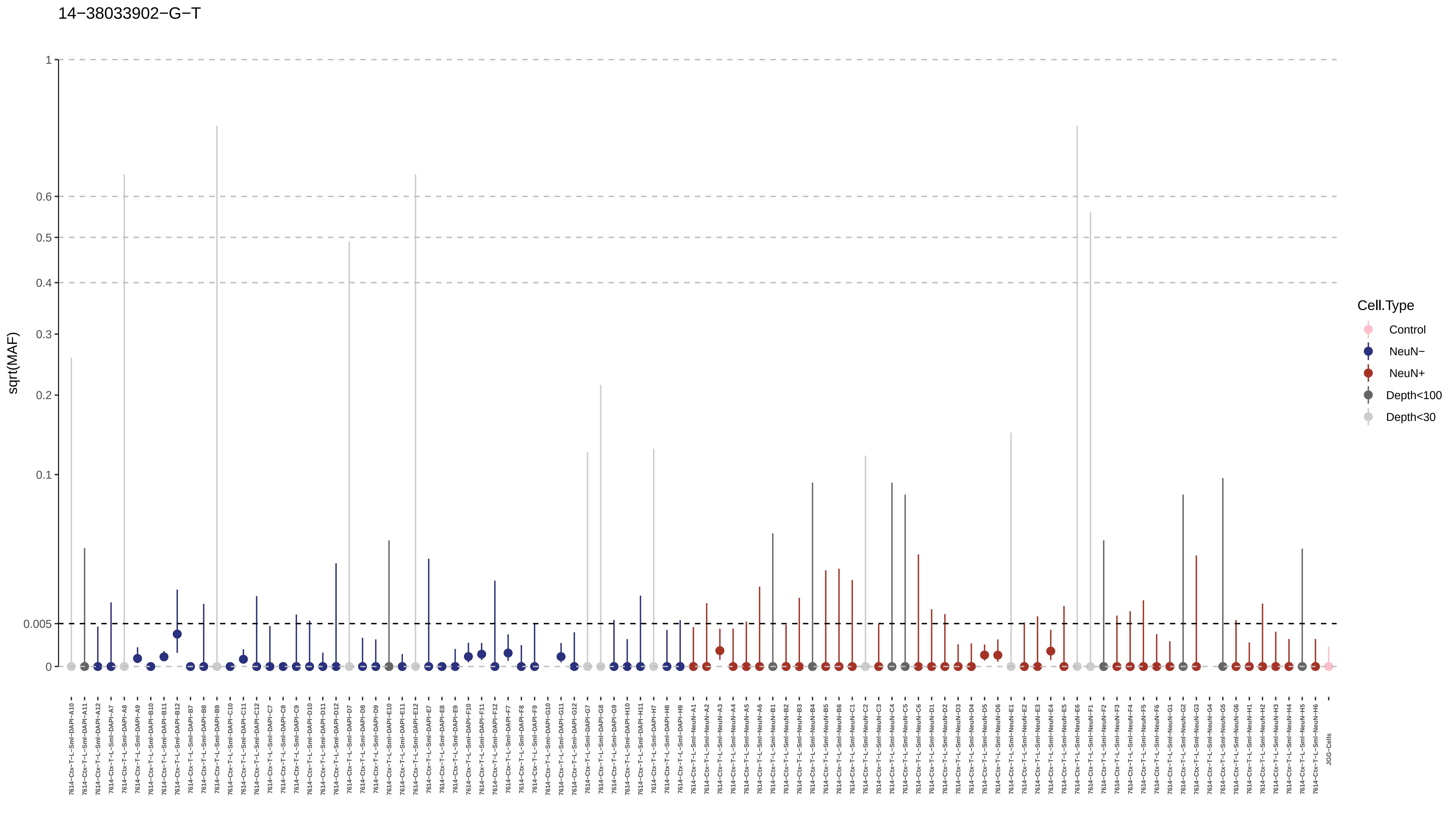
14−38033902−G−T
sqrt(MAF)
1
0.6
0.5
0.4
0.3
0.2
0.1
0.005
0
Cell.Type
Control
NeuN−
NeuN+
Depth<100
Depth<30

14−44933484−T−C

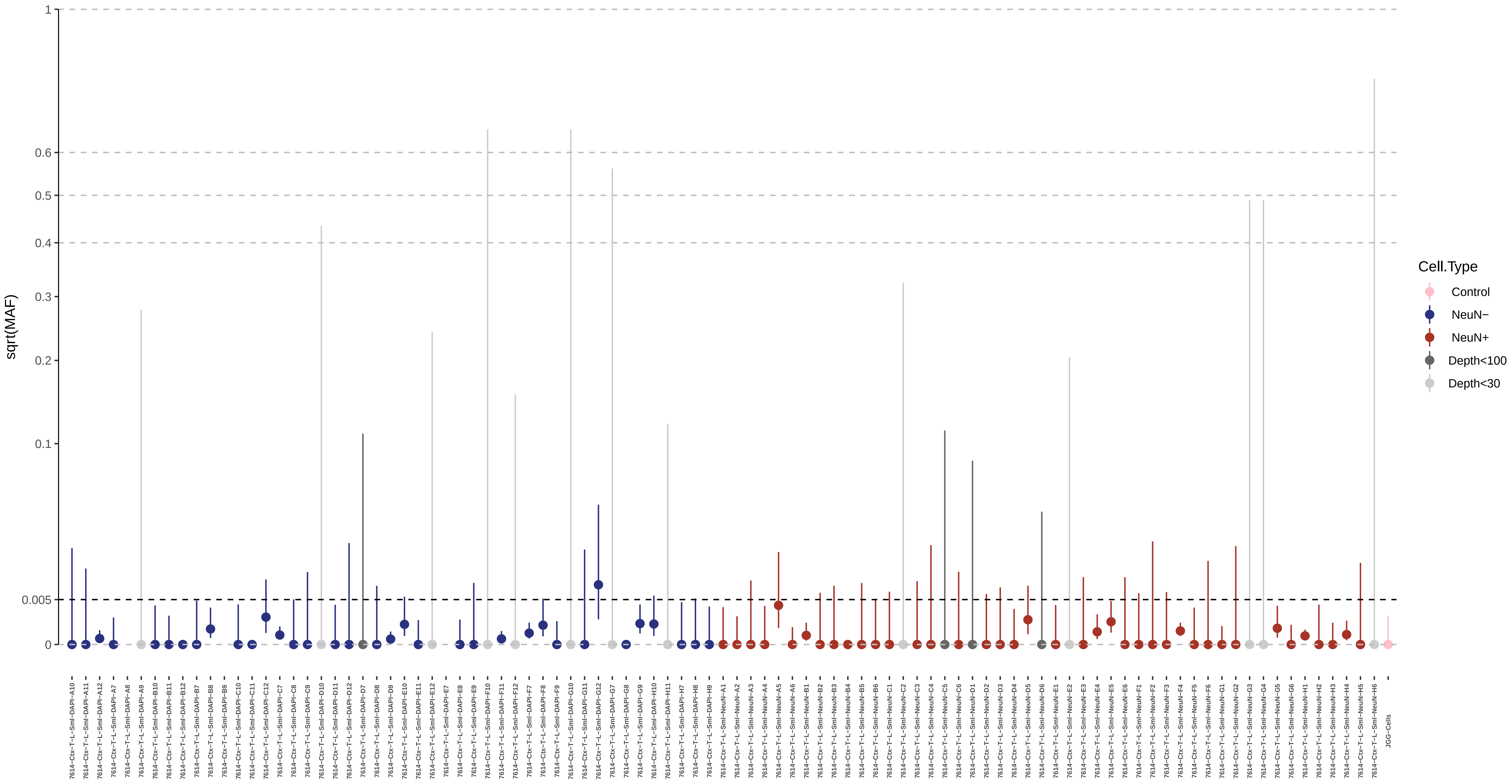

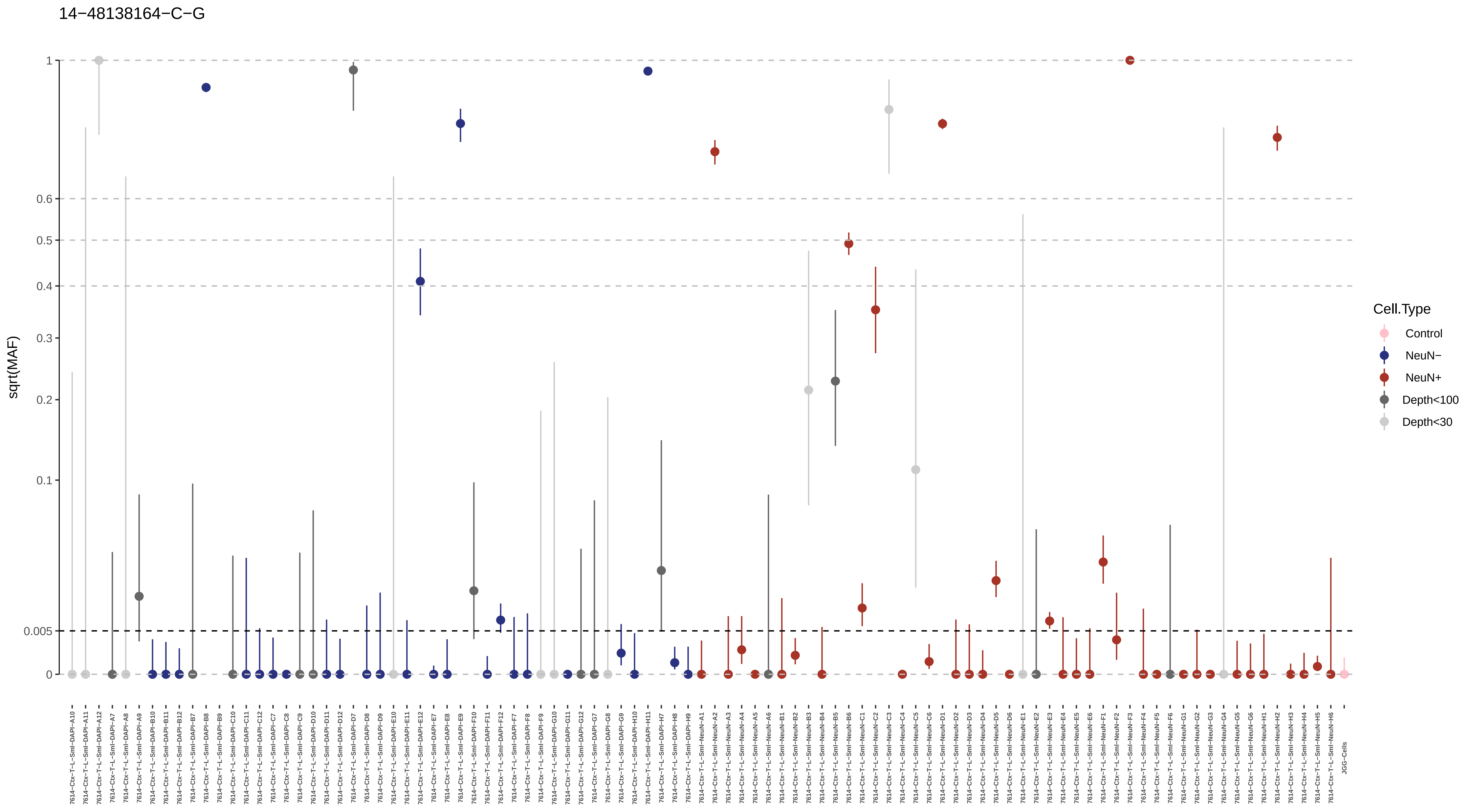
14−48138164−C−G
sqrt(MAF)
1
0.6
0.5
0.4
0.3
0.2
0.1
0.005
0
Cell.Type
Control
NeuN−
NeuN+
Depth<100
Depth<30

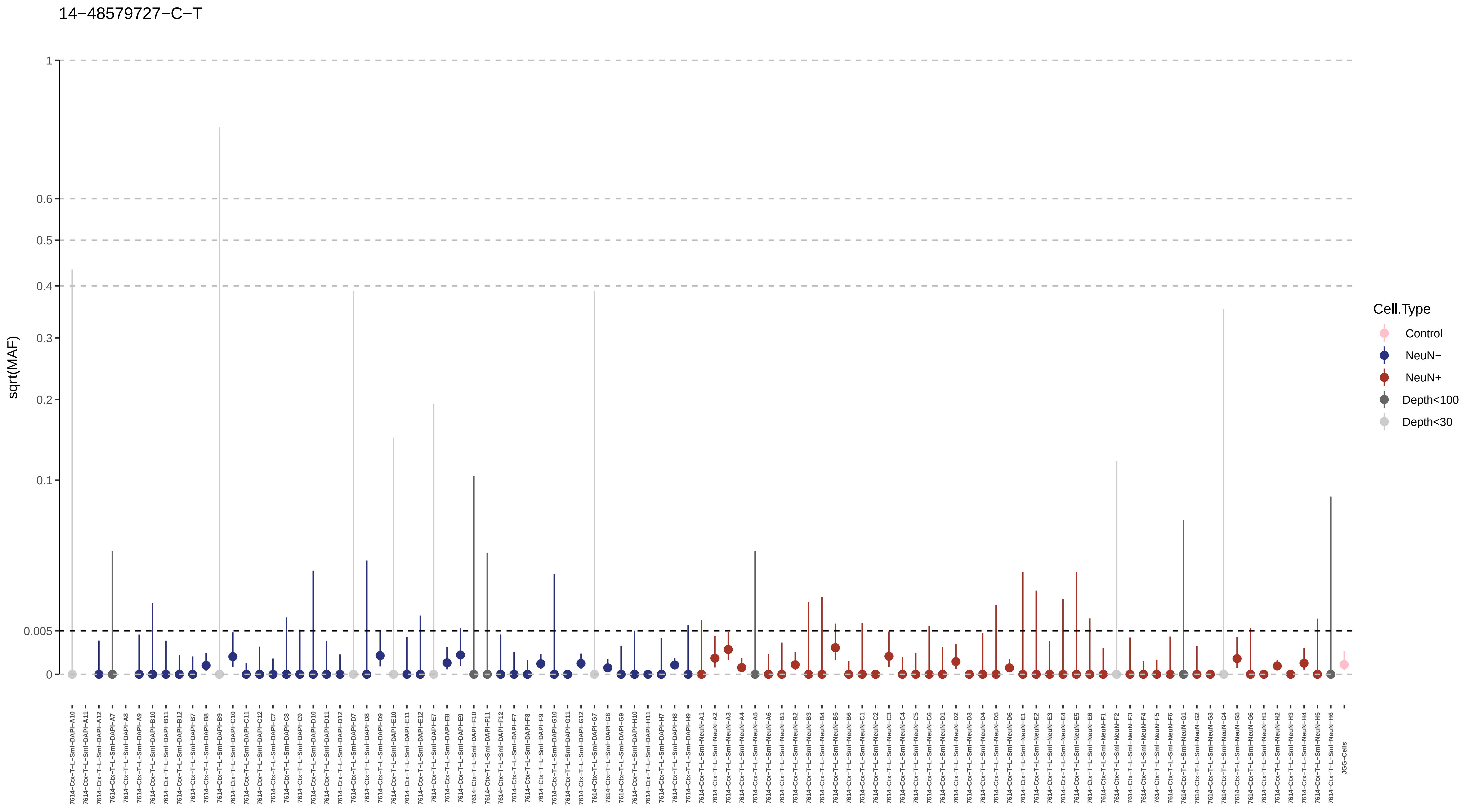
14−48579727−C−T
sqrt(MAF)
1
0.6
0.5
0.4
0.3
0.2
0.1
0.005
0
Cell.Type
Control
NeuN−
NeuN+
Depth<100
Depth<30
JGG-Cells

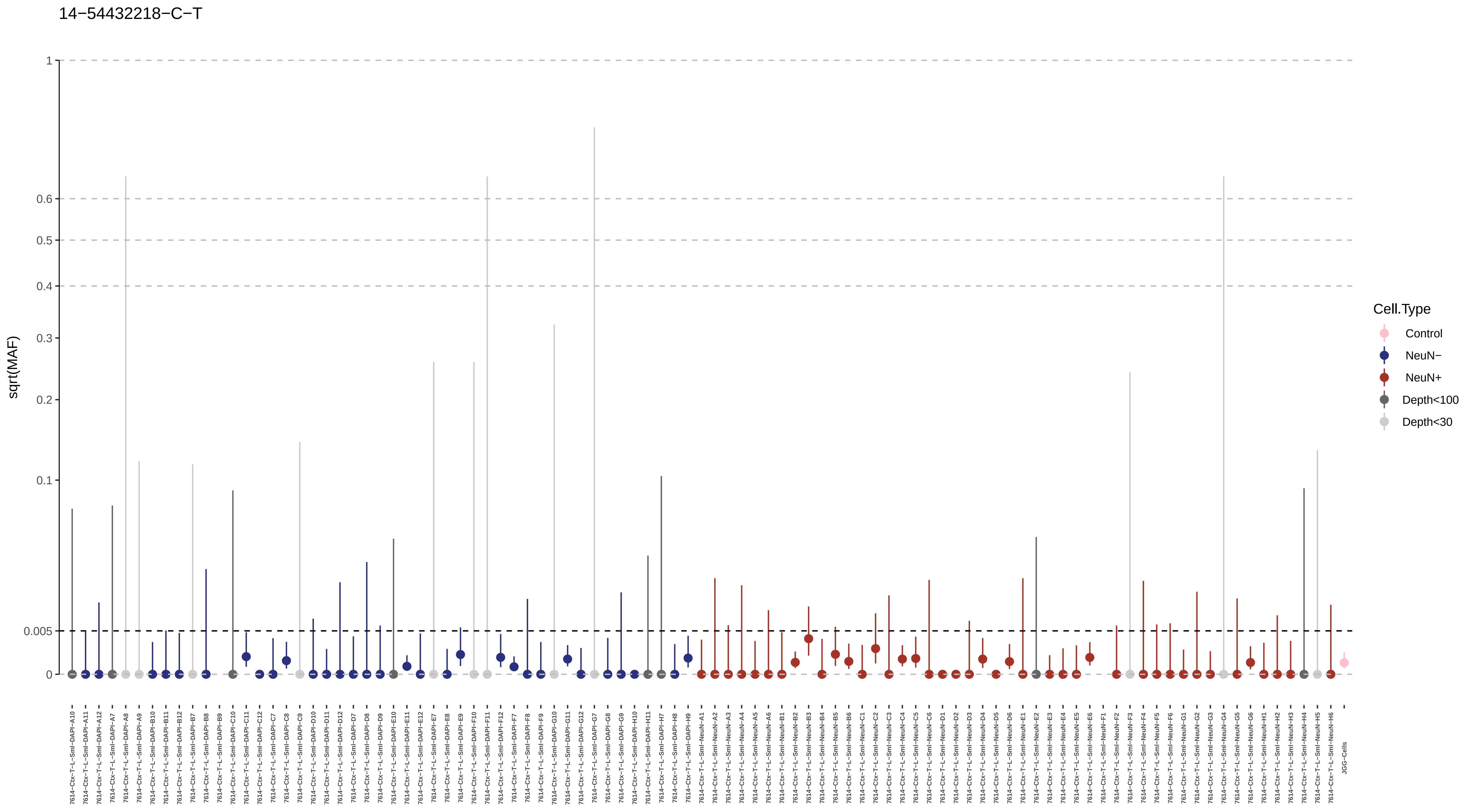

14−54432218−C−T
sqrt(MAF)
1
0.6
0.5
0.4
0.3
0.2
0.1
0.005
0
Cell.Type
Control
NeuN−
NeuN+
Depth<100
Depth<30
JGG-Cells

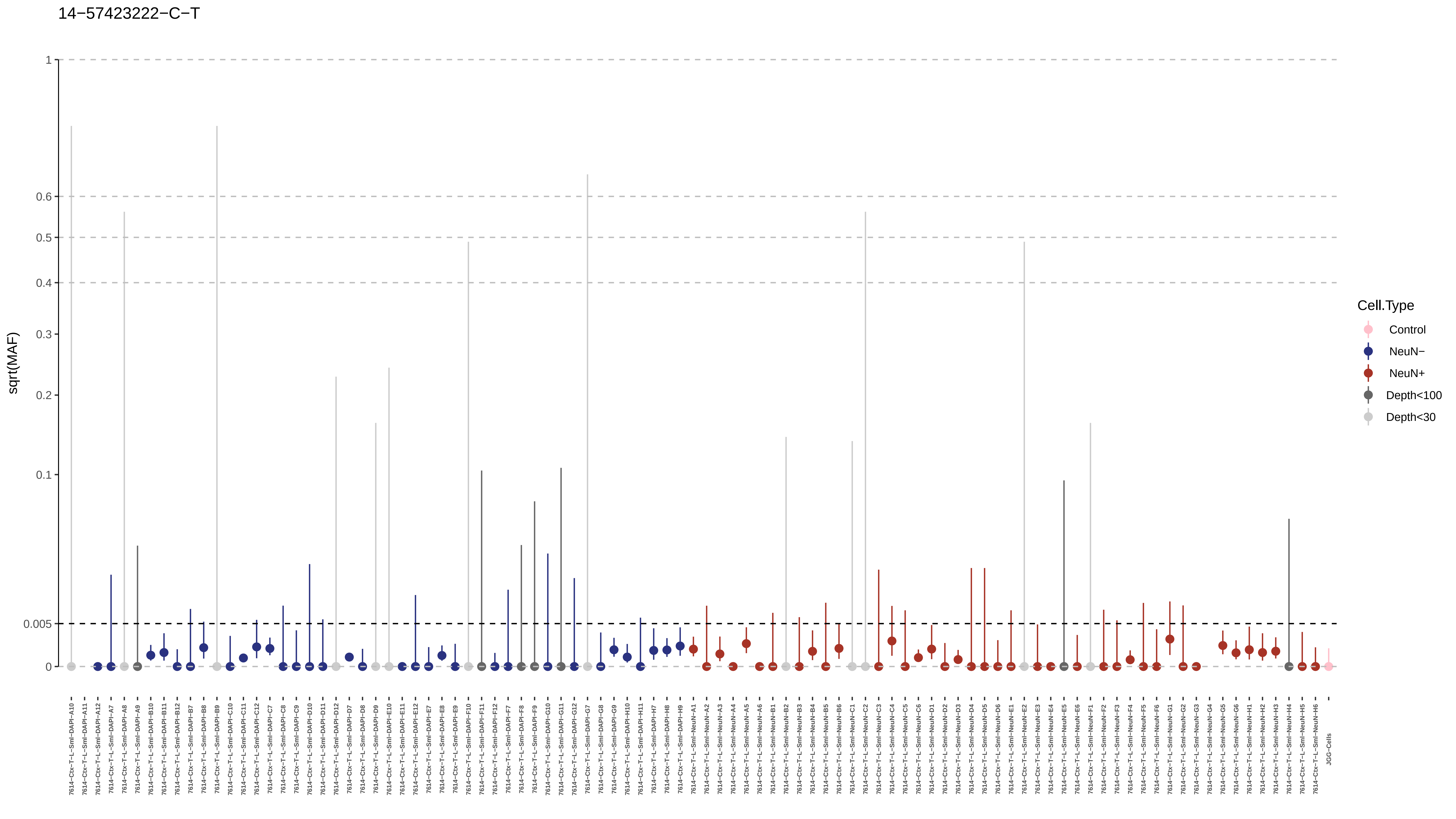

14−57423222−C−T
sqrt(MAF)
1
0.6
0.5
0.4
0.3
0.2
0.1
0.005
0
Cell.Type
Control
NeuN−
NeuN+
Depth<100
Depth<30
JGG-Cells

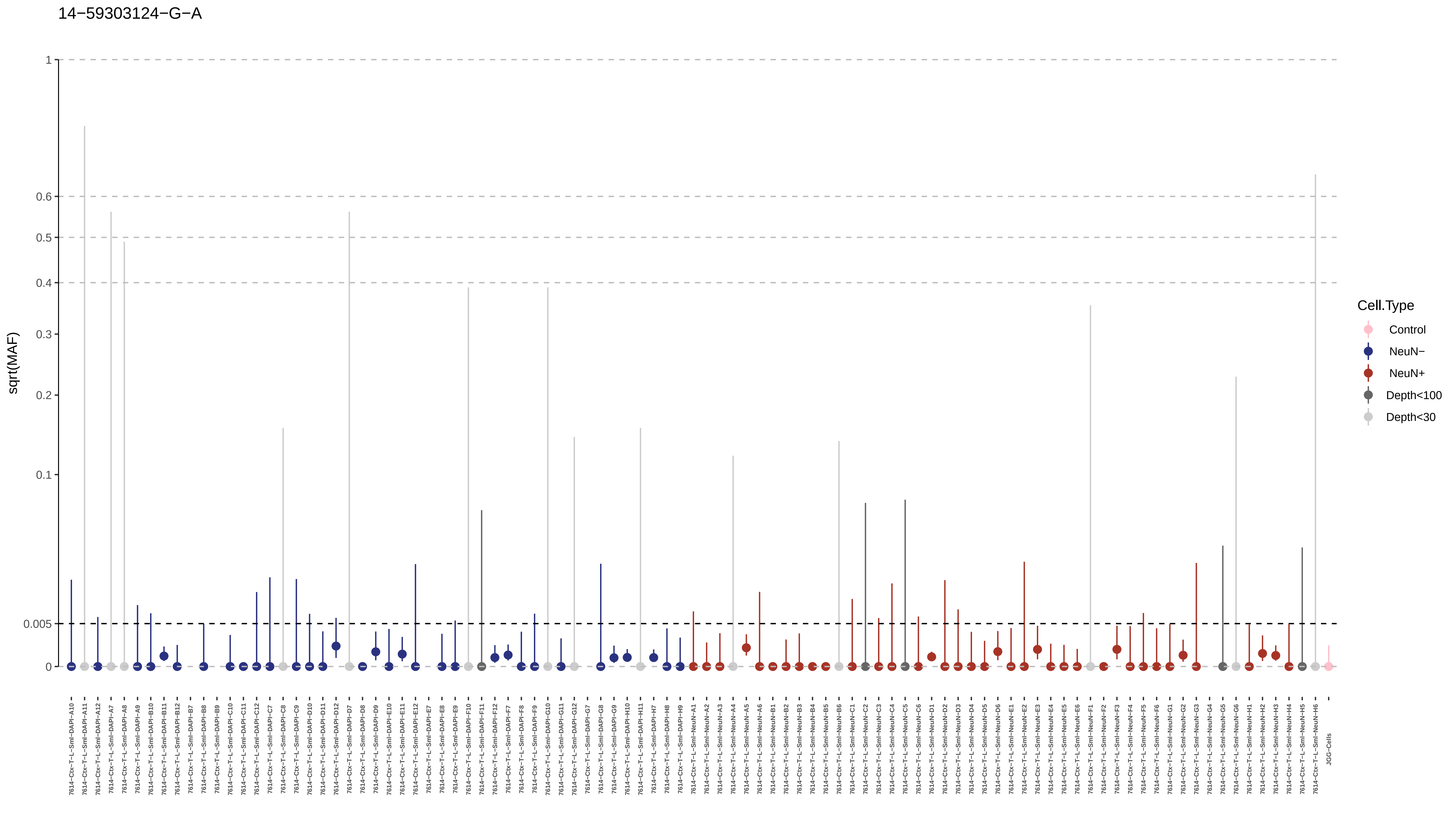

14−59303124−G−A
sqrt(MAF)
1
0.6
0.5
0.4
0.3
0.2
0.1
0.005
0
Cell.Type
Control
NeuN−
NeuN+
Depth<100
Depth<30

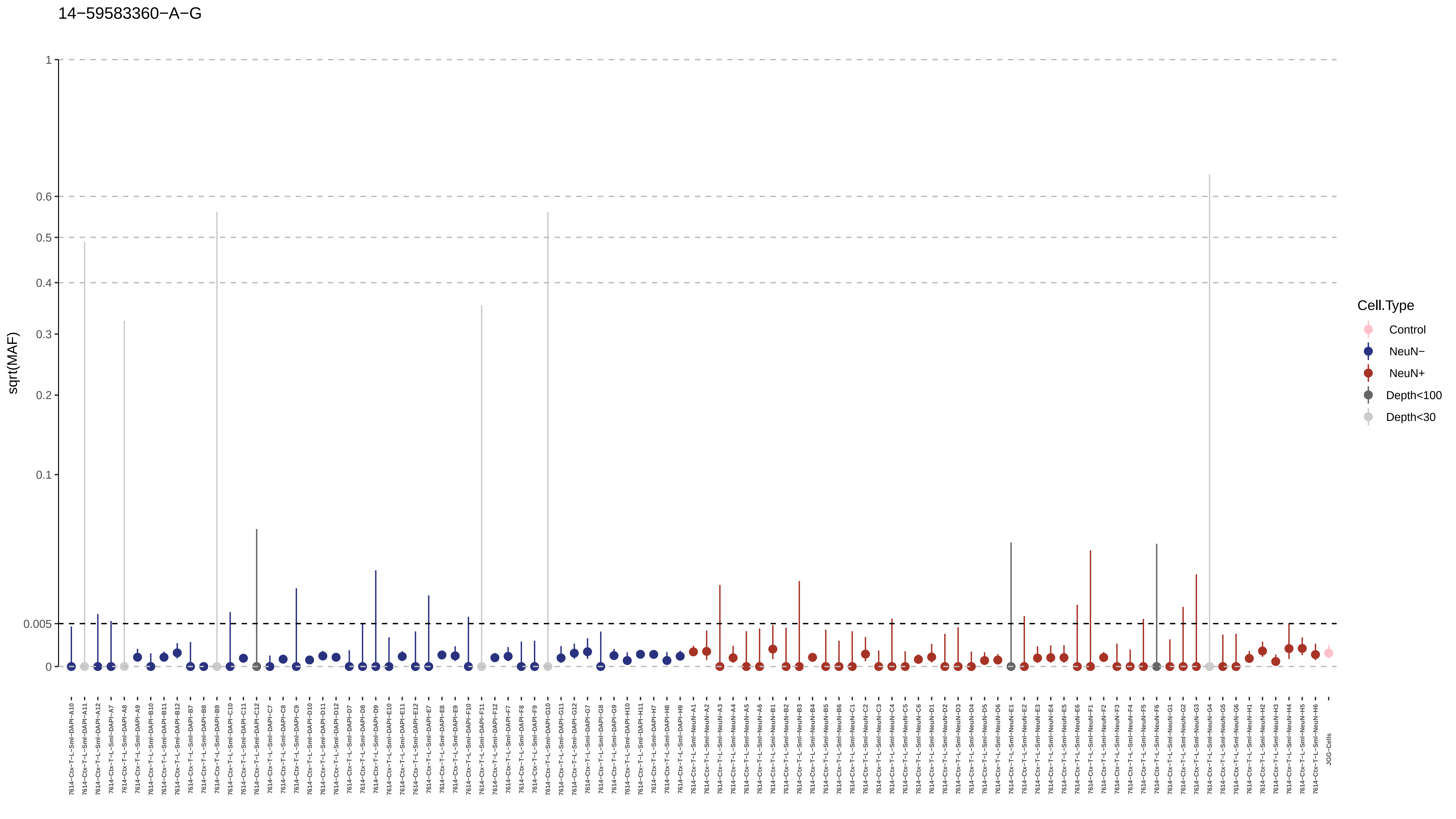
14−59583360−A−G
sqrt(MAF)
1
0.6
0.5
0.4
0.3
0.2
0.1
0.005
0
Cell.Type
Control
NeuN−
NeuN+
Depth<100
Depth<30

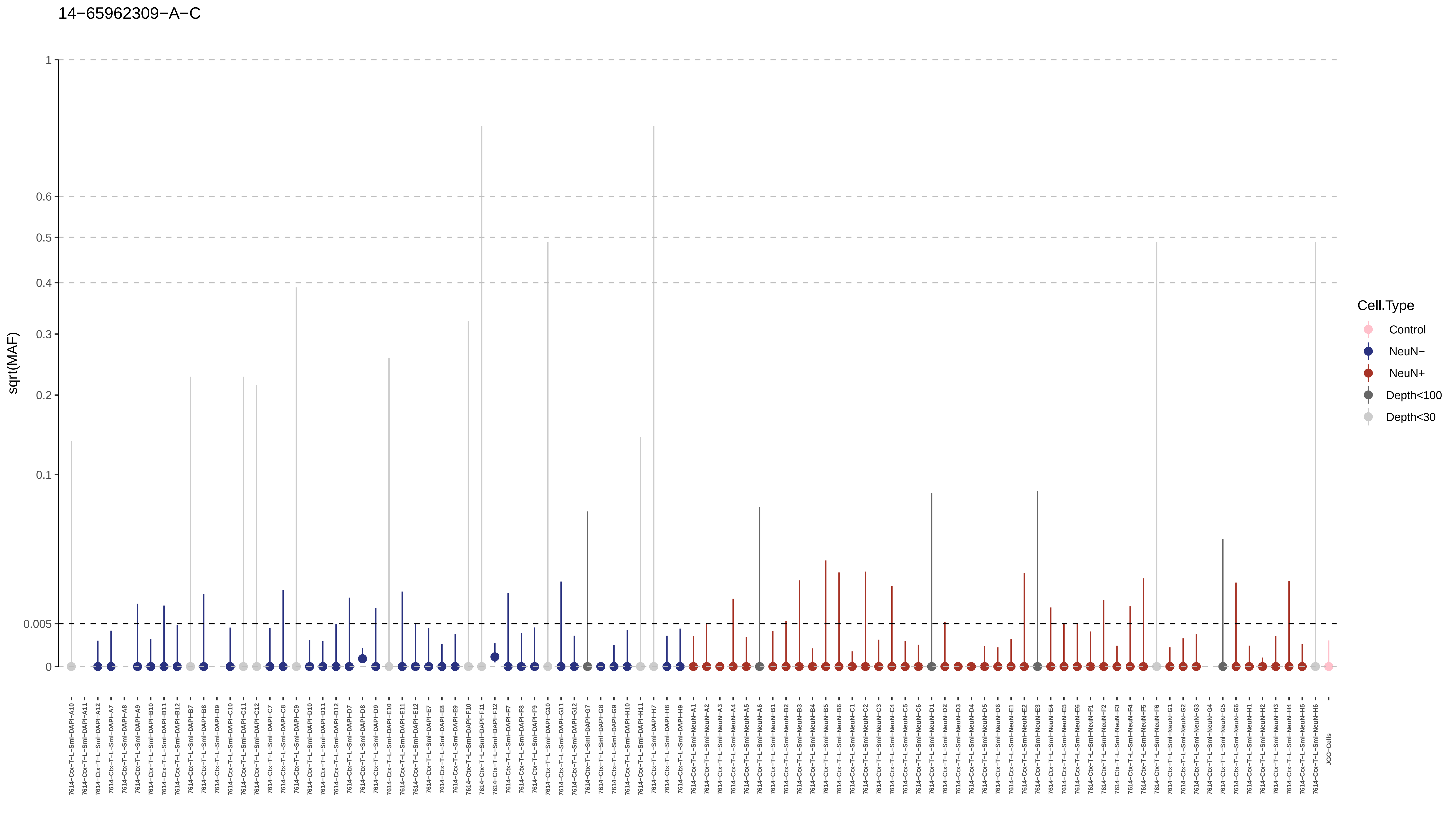
14−65962309−A−C
sqrt(MAF)
1
0.6
0.5
0.4
0.3
0.2
0.1
0.005
0
Cell.Type
Control
NeuN−
NeuN+
Depth<100
Depth<30

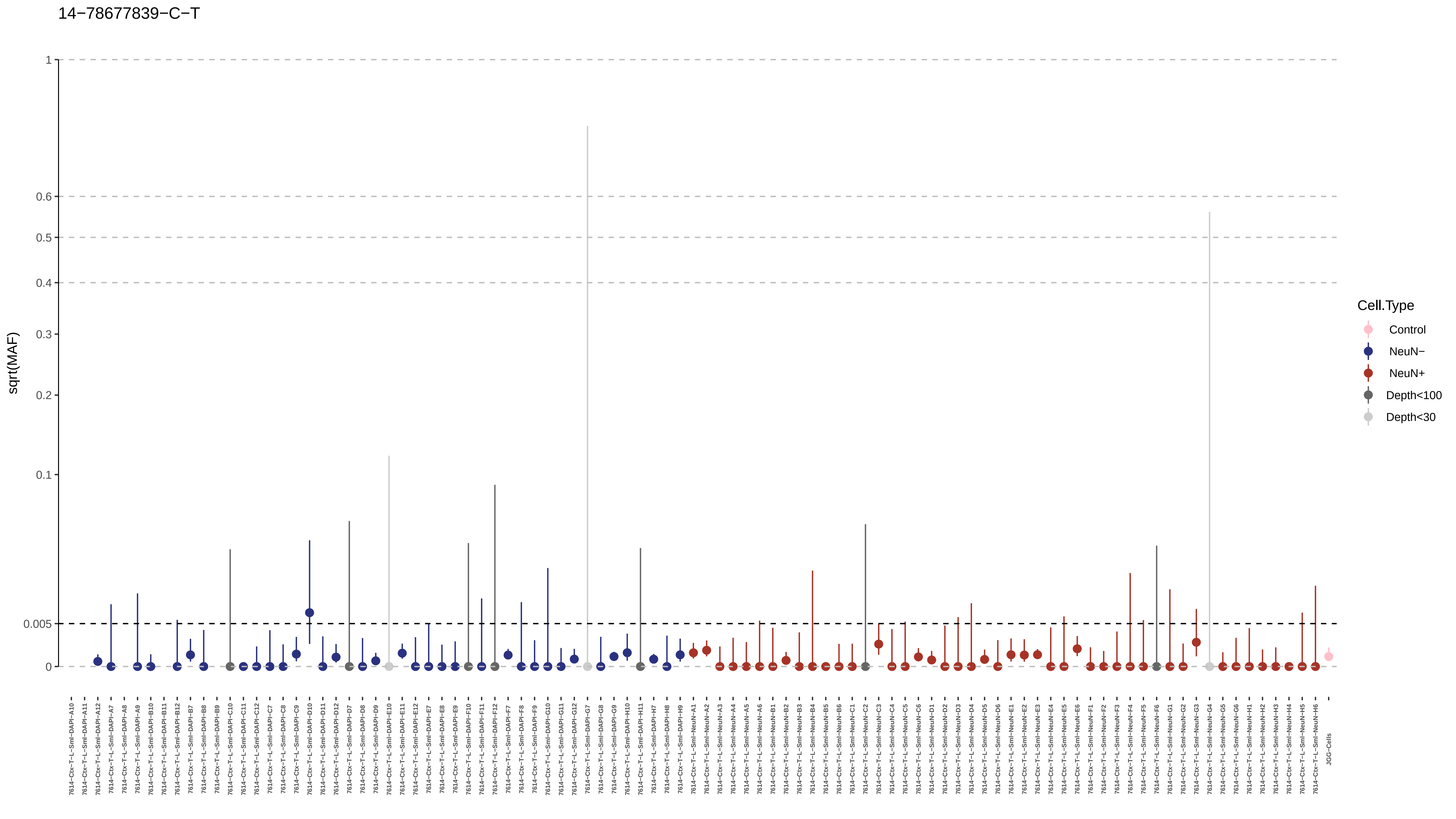

14−78677839−C−T
sqrt(MAF)
1
0.6
0.5
0.4
0.3
0.2
0.1
0.005
0
Cell.Type
Control
NeuN−
NeuN+
Depth<100
Depth<30
JGG-Cells

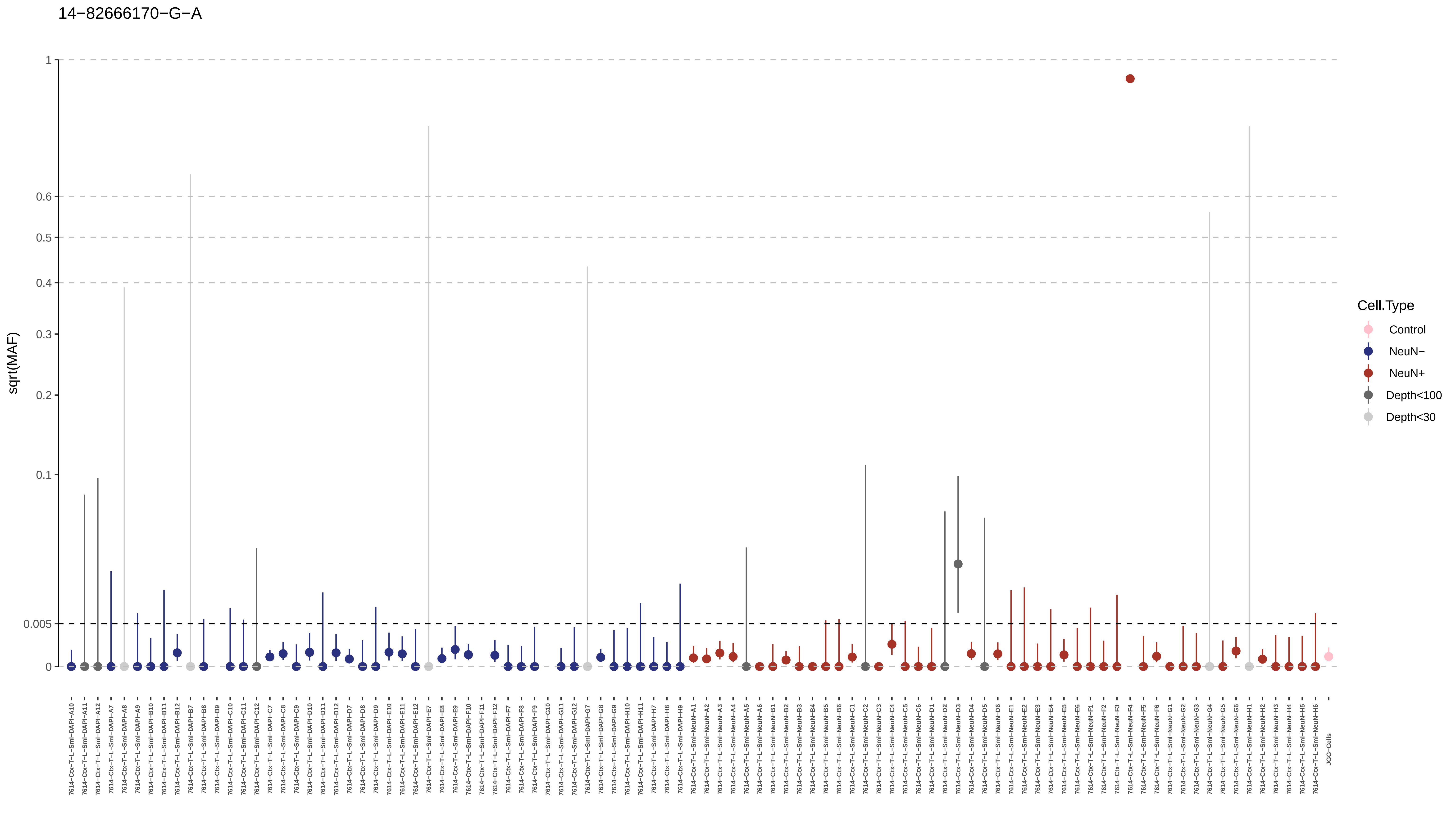
14−82666170−G−A
sqrt(MAF)
1
0.6
0.5
0.4
0.3
0.2
0.1
0.005
0
Cell.Type
Control
NeuN−
NeuN+
Depth<100
Depth<30

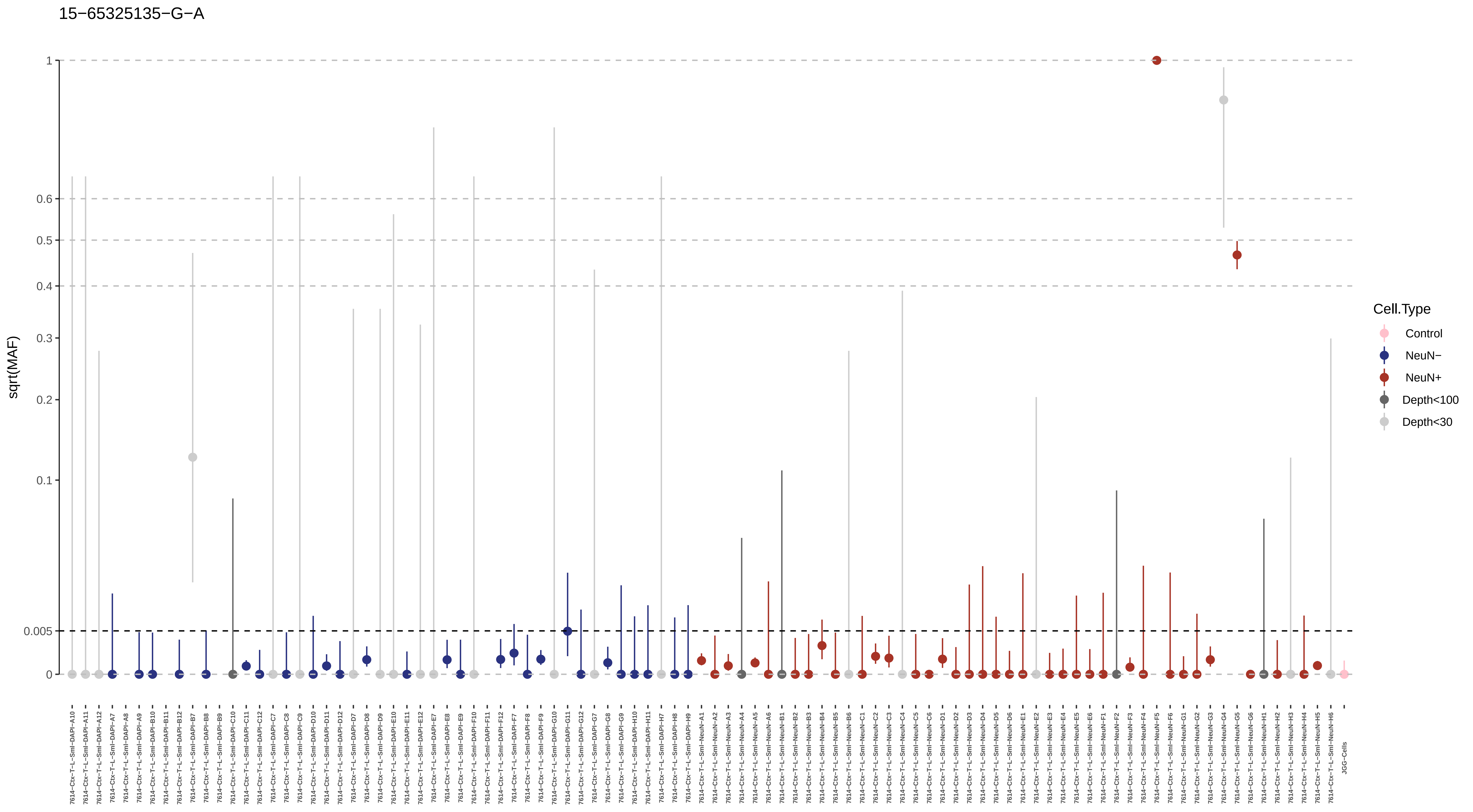

15−65325135−G−A
sqrt(MAF)
1
0.6
0.5
0.4
0.3
0.2
0.1
0.005
0
Cell.Type
Control
NeuN−
NeuN+
Depth<100
Depth<30
7614−Ctx−T−L−Sml−DAPI−A10
7614−Ctx−T−L−Sml−DAPI−A11
7614−Ctx−T−L−Sml−DAPI−A12
7614−Ctx−T−L−Sml−DAPI−A7
7614−Ctx−T−L−Sml−DAPI−A8
7614−Ctx−T−L−Sml−DAPI−A9
7614−Ctx−T−L−Sml−DAPI−B10
7614−Ctx−T−L−Sml−DAPI−B11
7614−Ctx−T−L−Sml−DAPI−B12
7614−Ctx−T−L−Sml−DAPI−B7
7614−Ctx−T−L−Sml−DAPI−B8
7614−Ctx−T−L−Sml−DAPI−B9
7614−Ctx−T−L−Sml−DAPI−C10
7614−Ctx−T−L−Sml−DAPI−C11
7614−Ctx−T−L−Sml−DAPI−C12
7614−Ctx−T−L−Sml−DAPI−C7
7614−Ctx−T−L−Sml−DAPI−C8
7614−Ctx−T−L−Sml−DAPI−C9
7614−Ctx−T−L−Sml−DAPI−D10
7614−Ctx−T−L−Sml−DAPI−D11
7614−Ctx−T−L−Sml−DAPI−D12
7614−Ctx−T−L−Sml−DAPI−D7
7614−Ctx−T−L−Sml−DAPI−D8
7614−Ctx−T−L−Sml−DAPI−D9
7614−Ctx−T−L−Sml−DAPI−E10
7614−Ctx−T−L−Sml−DAPI−E11
7614−Ctx−T−L−Sml−DAPI−E12
7614−Ctx−T−L−Sml−DAPI−E7
7614−Ctx−T−L−Sml−DAPI−E8
7614−Ctx−T−L−Sml−DAPI−E9
7614−Ctx−T−L−Sml−DAPI−F10
7614−Ctx−T−L−Sml−DAPI−F11
7614−Ctx−T−L−Sml−DAPI−F12
7614−Ctx−T−L−Sml−DAPI−F7
7614−Ctx−T−L−Sml−DAPI−F8
7614−Ctx−T−L−Sml−DAPI−F9
7614−Ctx−T−L−Sml−DAPI−G10
7614−Ctx−T−L−Sml−DAPI−G11
7614−Ctx−T−L−Sml−DAPI−G12
7614−Ctx−T−L−Sml−DAPI−G7
7614−Ctx−T−L−Sml−DAPI−G8
7614−Ctx−T−L−Sml−DAPI−G9
7614−Ctx−T−L−Sml−DAPI−H10
7614−Ctx−T−L−Sml−DAPI−H11
7614−Ctx−T−L−Sml−DAPI−H7
7614−Ctx−T−L−Sml−DAPI−H8
7614−Ctx−T−L−Sml−DAPI−H9
7614−Ctx−T−L−Sml−NeuN−A1
7614−Ctx−T−L−Sml−NeuN−A2
7614−Ctx−T−L−Sml−NeuN−A3
7614−Ctx−T−L−Sml−NeuN−A4
7614−Ctx−T−L−Sml−NeuN−A5
7614−Ctx−T−L−Sml−NeuN−A6
7614−Ctx−T−L−Sml−NeuN−B1
7614−Ctx−T−L−Sml−NeuN−B2
7614−Ctx−T−L−Sml−NeuN−B3
7614−Ctx−T−L−Sml−NeuN−B4
7614−Ctx−T−L−Sml−NeuN−B5
7614−Ctx−T−L−Sml−NeuN−B6
7614−Ctx−T−L−Sml−NeuN−C1
7614−Ctx−T−L−Sml−NeuN−C2
7614−Ctx−T−L−Sml−NeuN−C3
7614−Ctx−T−L−Sml−NeuN−C4
7614−Ctx−T−L−Sml−NeuN−C5
7614−Ctx−T−L−Sml−NeuN−C6
7614−Ctx−T−L−Sml−NeuN−D1
7614−Ctx−T−L−Sml−NeuN−D2
7614−Ctx−T−L−Sml−NeuN−D3
7614−Ctx−T−L−Sml−NeuN−D4
7614−Ctx−T−L−Sml−NeuN−D5
7614−Ctx−T−L−Sml−NeuN−D6
7614−Ctx−T−L−Sml−NeuN−E1
7614−Ctx−T−L−Sml−NeuN−E2
7614−Ctx−T−L−Sml−NeuN−E3
7614−Ctx−T−L−Sml−NeuN−E4
7614−Ctx−T−L−Sml−NeuN−E5
7614−Ctx−T−L−Sml−NeuN−E6
7614−Ctx−T−L−Sml−NeuN−F1
7614−Ctx−T−L−Sml−NeuN−F2
7614−Ctx−T−L−Sml−NeuN−F3
7614−Ctx−T−L−Sml−NeuN−F4
7614−Ctx−T−L−Sml−NeuN−F5
7614−Ctx−T−L−Sml−NeuN−F6
7614−Ctx−T−L−Sml−NeuN−G1
7614−Ctx−T−L−Sml−NeuN−G2
7614−Ctx−T−L−Sml−NeuN−G3
7614−Ctx−T−L−Sml−NeuN−G4
7614−Ctx−T−L−Sml−NeuN−G5
7614−Ctx−T−L−Sml−NeuN−G6
7614−Ctx−T−L−Sml−NeuN−H1
7614−Ctx−T−L−Sml−NeuN−H2
7614−Ctx−T−L−Sml−NeuN−H3
7614−Ctx−T−L−Sml−NeuN−H4
7614−Ctx−T−L−Sml−NeuN−H5
7614−Ctx−T−L−Sml−NeuN−H6
JGG−Cells

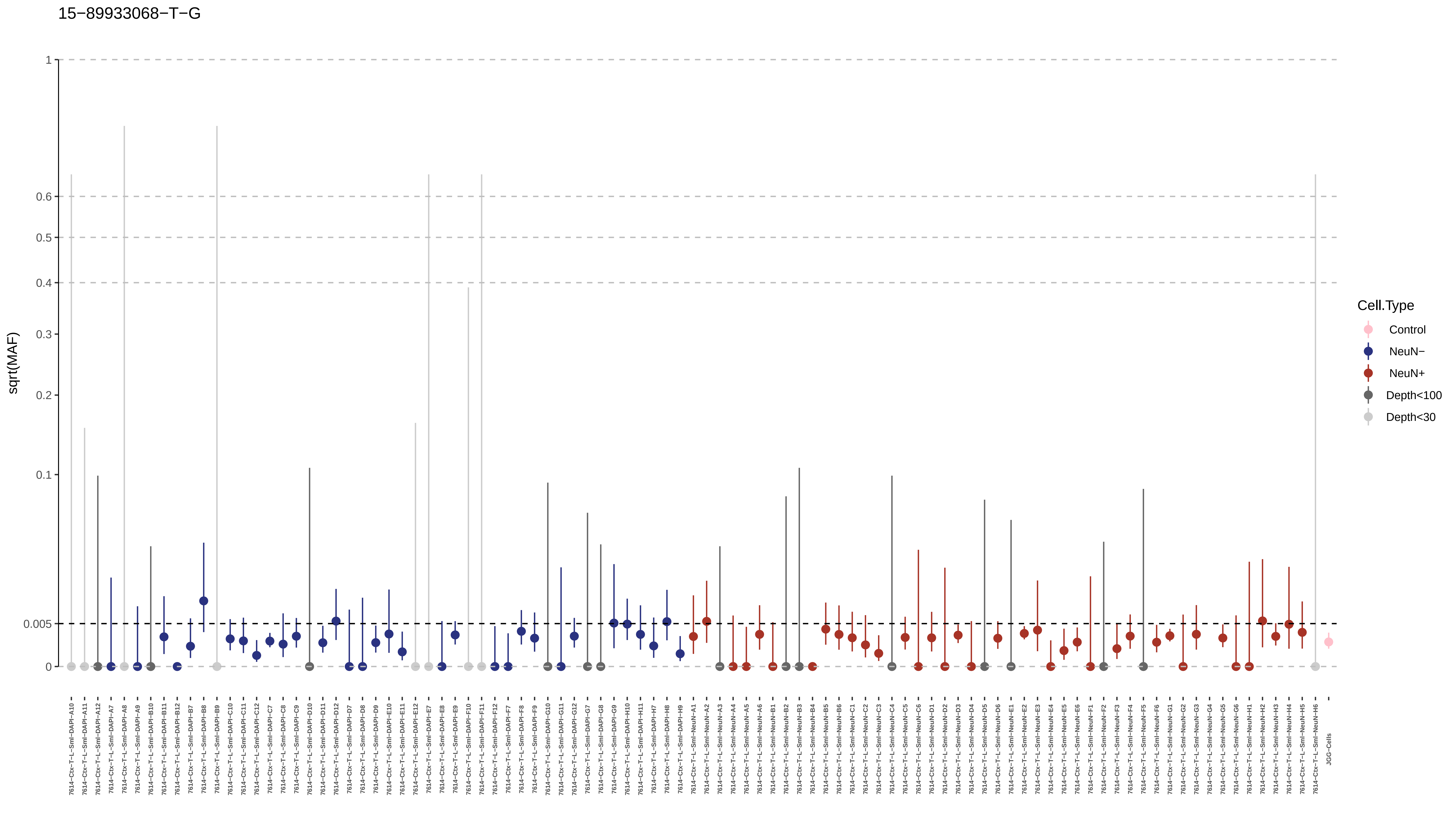
15−89933068−T−G
sqrt(MAF)
1
0.6
0.5
0.4
0.3
0.2
0.1
0.005
0
Cell.Type
Control
NeuN−
NeuN+
Depth<100
Depth<30
JGG-Cells

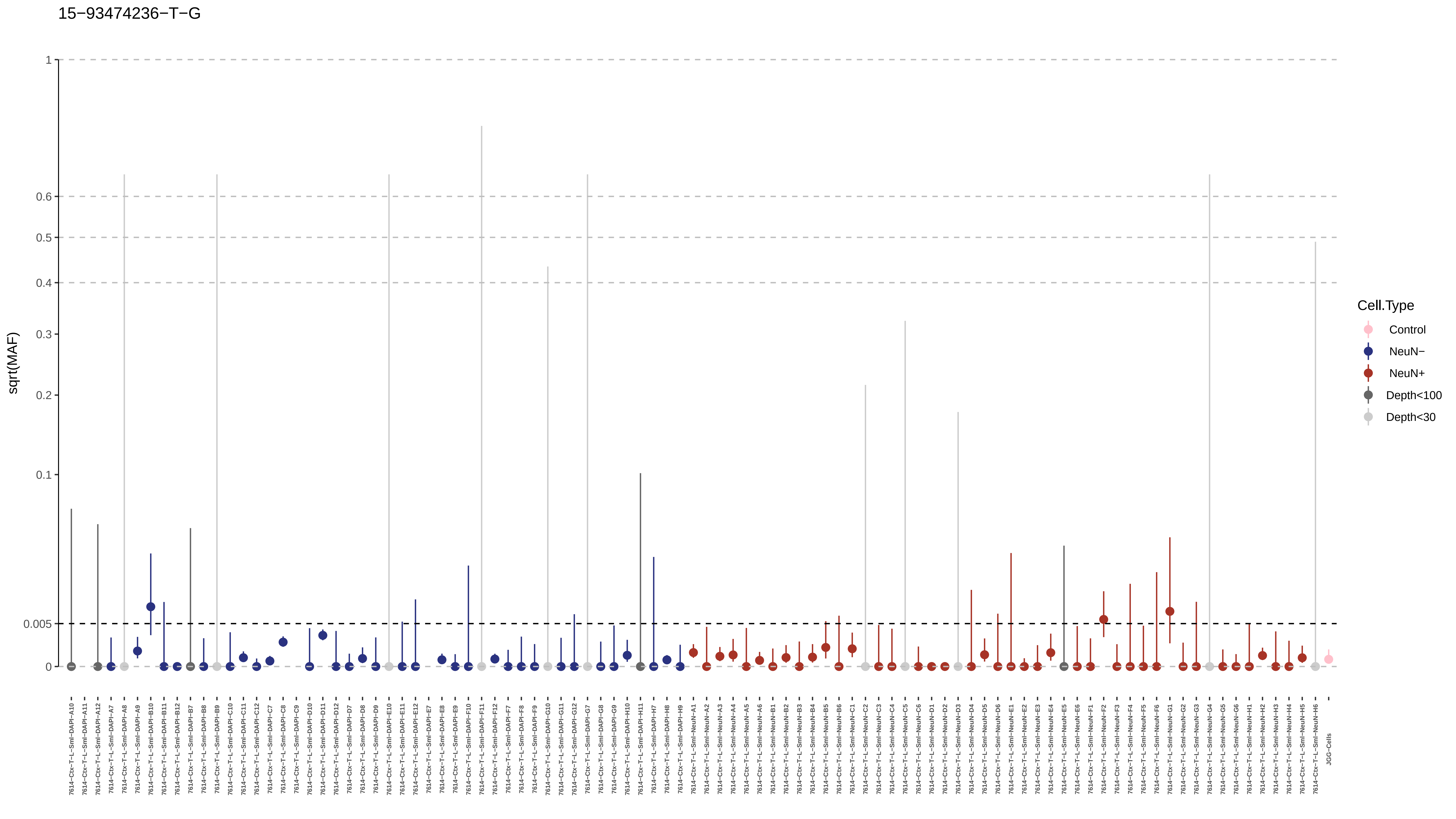
15−93474236−T−G
sqrt(MAF)
1
0.6
0.5
0.4
0.3
0.2
0.1
0.005
0
Cell.Type
Control
NeuN−
NeuN+
Depth<100
Depth<30
7614−Ctx−T−L−Sml−DAPI−A10
7614−Ctx−T−L−Sml−DAPI−A11
7614−Ctx−T−L−Sml−DAPI−A12
7614−Ctx−T−L−Sml−DAPI−A7
7614−Ctx−T−L−Sml−DAPI−A8
7614−Ctx−T−L−Sml−DAPI−A9
7614−Ctx−T−L−Sml−DAPI−B10
7614−Ctx−T−L−Sml−DAPI−B11
7614−Ctx−T−L−Sml−DAPI−B12
7614−Ctx−T−L−Sml−DAPI−B7
7614−Ctx−T−L−Sml−DAPI−B8
7614−Ctx−T−L−Sml−DAPI−B9
7614−Ctx−T−L−Sml−DAPI−C10
7614−Ctx−T−L−Sml−DAPI−C11
7614−Ctx−T−L−Sml−DAPI−C12
7614−Ctx−T−L−Sml−DAPI−C7
7614−Ctx−T−L−Sml−DAPI−C8
7614−Ctx−T−L−Sml−DAPI−C9
7614−Ctx−T−L−Sml−DAPI−D10
7614−Ctx−T−L−Sml−DAPI−D11
7614−Ctx−T−L−Sml−DAPI−D12
7614−Ctx−T−L−Sml−DAPI−D7
7614−Ctx−T−L−Sml−DAPI−D8
7614−Ctx−T−L−Sml−DAPI−D9
7614−Ctx−T−L−Sml−DAPI−E10
7614−Ctx−T−L−Sml−DAPI−E11
7614−Ctx−T−L−Sml−DAPI−E12
7614−Ctx−T−L−Sml−DAPI−E7
7614−Ctx−T−L−Sml−DAPI−E8
7614−Ctx−T−L−Sml−DAPI−E9
7614−Ctx−T−L−Sml−DAPI−F10
7614−Ctx−T−L−Sml−DAPI−F11
7614−Ctx−T−L−Sml−DAPI−F12
7614−Ctx−T−L−Sml−DAPI−F7
7614−Ctx−T−L−Sml−DAPI−F8
7614−Ctx−T−L−Sml−DAPI−F9
7614−Ctx−T−L−Sml−DAPI−G10
7614−Ctx−T−L−Sml−DAPI−G11
7614−Ctx−T−L−Sml−DAPI−G12
7614−Ctx−T−L−Sml−DAPI−G7
7614−Ctx−T−L−Sml−DAPI−G8
7614−Ctx−T−L−Sml−DAPI−G9
7614−Ctx−T−L−Sml−DAPI−H10
7614−Ctx−T−L−Sml−DAPI−H11
7614−Ctx−T−L−Sml−DAPI−H7
7614−Ctx−T−L−Sml−DAPI−H8
7614−Ctx−T−L−Sml−DAPI−H9
7614−Ctx−T−L−Sml−NeuN−A1
7614−Ctx−T−L−Sml−NeuN−A2
7614−Ctx−T−L−Sml−NeuN−A3
7614−Ctx−T−L−Sml−NeuN−A4
7614−Ctx−T−L−Sml−NeuN−A5
7614−Ctx−T−L−Sml−NeuN−A6
7614−Ctx−T−L−Sml−NeuN−B1
7614−Ctx−T−L−Sml−NeuN−B2
7614−Ctx−T−L−Sml−NeuN−B3
7614−Ctx−T−L−Sml−NeuN−B4
7614−Ctx−T−L−Sml−NeuN−B5
7614−Ctx−T−L−Sml−NeuN−B6
7614−Ctx−T−L−Sml−NeuN−C1
7614−Ctx−T−L−Sml−NeuN−C2
7614−Ctx−T−L−Sml−NeuN−C3
7614−Ctx−T−L−Sml−NeuN−C4
7614−Ctx−T−L−Sml−NeuN−C5
7614−Ctx−T−L−Sml−NeuN−C6
7614−Ctx−T−L−Sml−NeuN−D1
7614−Ctx−T−L−Sml−NeuN−D2
7614−Ctx−T−L−Sml−NeuN−D3
7614−Ctx−T−L−Sml−NeuN−D4
7614−Ctx−T−L−Sml−NeuN−D5
7614−Ctx−T−L−Sml−NeuN−D6
7614−Ctx−T−L−Sml−NeuN−E1
7614−Ctx−T−L−Sml−NeuN−E2
7614−Ctx−T−L−Sml−NeuN−E3
7614−Ctx−T−L−Sml−NeuN−E4
7614−Ctx−T−L−Sml−NeuN−E5
7614−Ctx−T−L−Sml−NeuN−E6
7614−Ctx−T−L−Sml−NeuN−F1
7614−Ctx−T−L−Sml−NeuN−F2
7614−Ctx−T−L−Sml−NeuN−F3
7614−Ctx−T−L−Sml−NeuN−F4
7614−Ctx−T−L−Sml−NeuN−F5
7614−Ctx−T−L−Sml−NeuN−F6
7614−Ctx−T−L−Sml−NeuN−G1
7614−Ctx−T−L−Sml−NeuN−G2
7614−Ctx−T−L−Sml−NeuN−G3
7614−Ctx−T−L−Sml−NeuN−G4
7614−Ctx−T−L−Sml−NeuN−G5
7614−Ctx−T−L−Sml−NeuN−G6
7614−Ctx−T−L−Sml−NeuN−H1
7614−Ctx−T−L−Sml−NeuN−H2
7614−Ctx−T−L−Sml−NeuN−H3
7614−Ctx−T−L−Sml−NeuN−H4
7614−Ctx−T−L−Sml−NeuN−H5
7614−Ctx−T−L−Sml−NeuN−H6
JGG-Cells

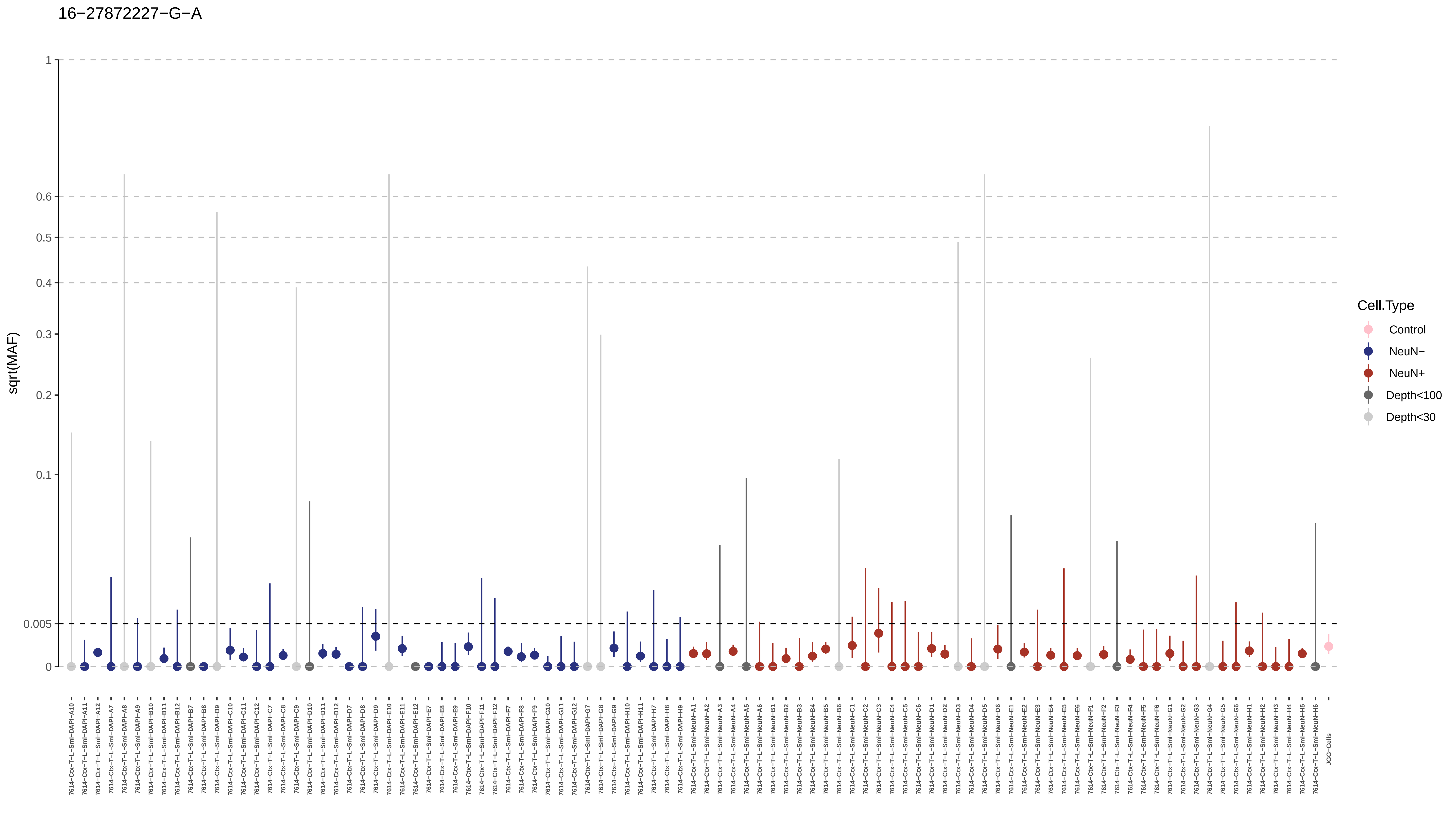
16−27872227−G−A
sqrt(MAF)
1
0.6
0.5
0.4
0.3
0.2
0.1
0.005
0
Cell.Type
Control
NeuN−
NeuN+
Depth<100
Depth<30
7614−Ctx−T−L−Sml−DAPI−A10
7614−Ctx−T−L−Sml−DAPI−A11
7614−Ctx−T−L−Sml−DAPI−A12
7614−Ctx−T−L−Sml−DAPI−A7
7614−Ctx−T−L−Sml−DAPI−A8
7614−Ctx−T−L−Sml−DAPI−A9
7614−Ctx−T−L−Sml−DAPI−B10
7614−Ctx−T−L−Sml−DAPI−B11
7614−Ctx−T−L−Sml−DAPI−B12
7614−Ctx−T−L−Sml−DAPI−B7
7614−Ctx−T−L−Sml−DAPI−B8
7614−Ctx−T−L−Sml−DAPI−B9
7614−Ctx−T−L−Sml−DAPI−C10
7614−Ctx−T−L−Sml−DAPI−C11
7614−Ctx−T−L−Sml−DAPI−C12
7614−Ctx−T−L−Sml−DAPI−C7
7614−Ctx−T−L−Sml−DAPI−C8
7614−Ctx−T−L−Sml−DAPI−C9
7614−Ctx−T−L−Sml−DAPI−D10
7614−Ctx−T−L−Sml−DAPI−D11
7614−Ctx−T−L−Sml−DAPI−D12
7614−Ctx−T−L−Sml−DAPI−D7
7614−Ctx−T−L−Sml−DAPI−D8
7614−Ctx−T−L−Sml−DAPI−D9
7614−Ctx−T−L−Sml−DAPI−E10
7614−Ctx−T−L−Sml−DAPI−E11
7614−Ctx−T−L−Sml−DAPI−E12
7614−Ctx−T−L−Sml−DAPI−E7
7614−Ctx−T−L−Sml−DAPI−E8
7614−Ctx−T−L−Sml−DAPI−E9
7614−Ctx−T−L−Sml−DAPI−F10
7614−Ctx−T−L−Sml−DAPI−F11
7614−Ctx−T−L−Sml−DAPI−F12
7614−Ctx−T−L−Sml−DAPI−F7
7614−Ctx−T−L−Sml−DAPI−F8
7614−Ctx−T−L−Sml−DAPI−F9
7614−Ctx−T−L−Sml−DAPI−G10
7614−Ctx−T−L−Sml−DAPI−G11
7614−Ctx−T−L−Sml−DAPI−G12
7614−Ctx−T−L−Sml−DAPI−G7
7614−Ctx−T−L−Sml−DAPI−G8
7614−Ctx−T−L−Sml−DAPI−G9
7614−Ctx−T−L−Sml−DAPI−H10
7614−Ctx−T−L−Sml−DAPI−H11
7614−Ctx−T−L−Sml−DAPI−H7
7614−Ctx−T−L−Sml−DAPI−H8
7614−Ctx−T−L−Sml−DAPI−H9
7614−Ctx−T−L−Sml−NeuN−A1
7614−Ctx−T−L−Sml−NeuN−A2
7614−Ctx−T−L−Sml−NeuN−A3
7614−Ctx−T−L−Sml−NeuN−A4
7614−Ctx−T−L−Sml−NeuN−A5
7614−Ctx−T−L−Sml−NeuN−A6
7614−Ctx−T−L−Sml−NeuN−B1
7614−Ctx−T−L−Sml−NeuN−B2
7614−Ctx−T−L−Sml−NeuN−B3
7614−Ctx−T−L−Sml−NeuN−B4
7614−Ctx−T−L−Sml−NeuN−B5
7614−Ctx−T−L−Sml−NeuN−B6
7614−Ctx−T−L−Sml−NeuN−C1
7614−Ctx−T−L−Sml−NeuN−C2
7614−Ctx−T−L−Sml−NeuN−C3
7614−Ctx−T−L−Sml−NeuN−C4
7614−Ctx−T−L−Sml−NeuN−C5
7614−Ctx−T−L−Sml−NeuN−C6
7614−Ctx−T−L−Sml−NeuN−D1
7614−Ctx−T−L−Sml−NeuN−D2
7614−Ctx−T−L−Sml−NeuN−D3
7614−Ctx−T−L−Sml−NeuN−D4
7614−Ctx−T−L−Sml−NeuN−D5
7614−Ctx−T−L−Sml−NeuN−D6
7614−Ctx−T−L−Sml−NeuN−E1
7614−Ctx−T−L−Sml−NeuN−E2
7614−Ctx−T−L−Sml−NeuN−E3
7614−Ctx−T−L−Sml−NeuN−E4
7614−Ctx−T−L−Sml−NeuN−E5
7614−Ctx−T−L−Sml−NeuN−E6
7614−Ctx−T−L−Sml−NeuN−F1
7614−Ctx−T−L−Sml−NeuN−F2
7614−Ctx−T−L−Sml−NeuN−F3
7614−Ctx−T−L−Sml−NeuN−F4
7614−Ctx−T−L−Sml−NeuN−F5
7614−Ctx−T−L−Sml−NeuN−F6
7614−Ctx−T−L−Sml−NeuN−G1
7614−Ctx−T−L−Sml−NeuN−G2
7614−Ctx−T−L−Sml−NeuN−G3
7614−Ctx−T−L−Sml−NeuN−G4
7614−Ctx−T−L−Sml−NeuN−G5
7614−Ctx−T−L−Sml−NeuN−G6
7614−Ctx−T−L−Sml−NeuN−H1
7614−Ctx−T−L−Sml−NeuN−H2
7614−Ctx−T−L−Sml−NeuN−H3
7614−Ctx−T−L−Sml−NeuN−H4
7614−Ctx−T−L−Sml−NeuN−H5
7614−Ctx−T−L−Sml−NeuN−H6
JGG-Cells

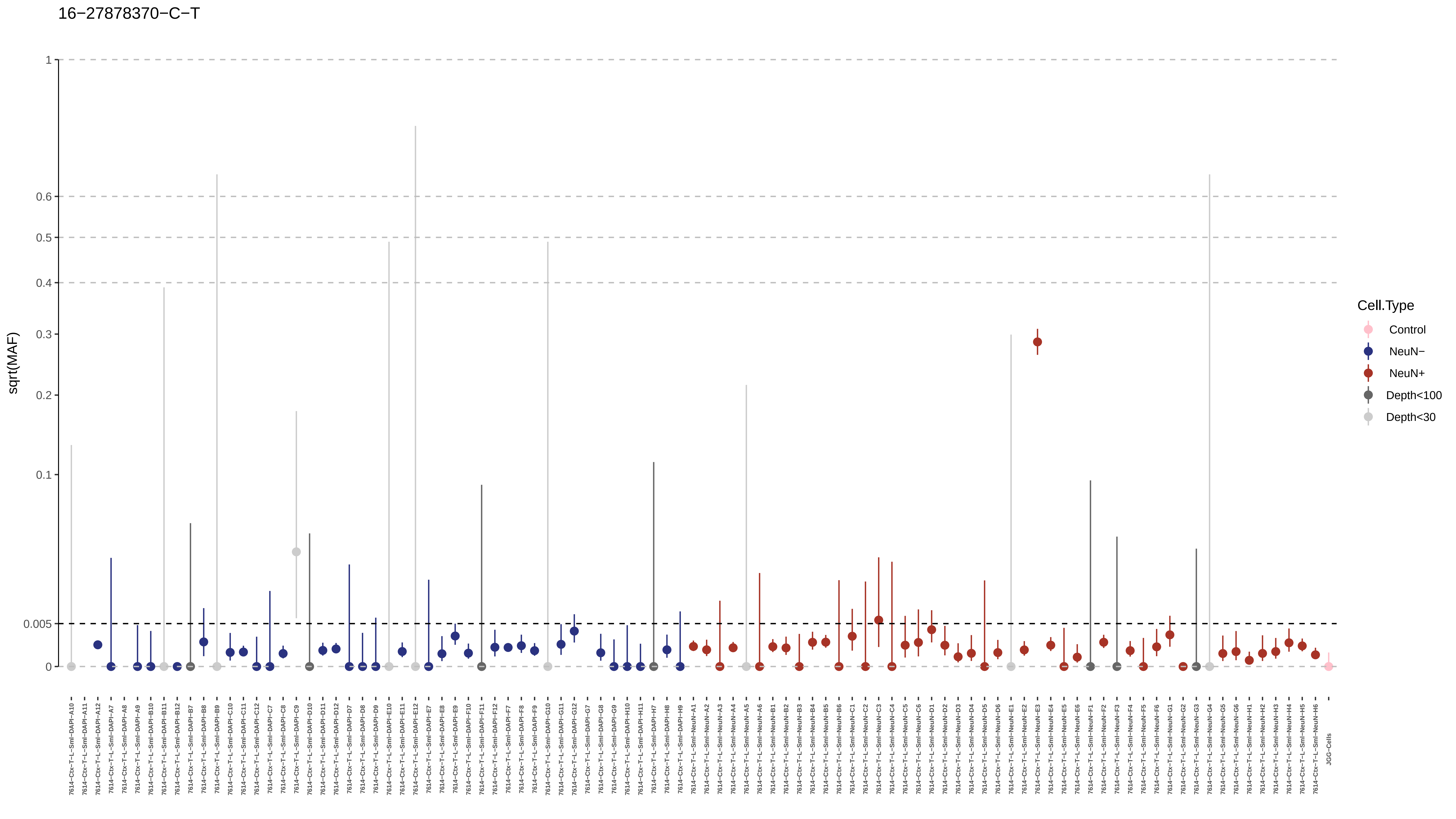
16−27878370−C−T
sqrt(MAF)
1
0.6
0.5
0.4
0.3
0.2
0.1
0.005
0
Cell.Type
Control
NeuN−
NeuN+
Depth<100
Depth<30
7614−Ctx−T−L−Sml−DAPI−A10
7614−Ctx−T−L−Sml−DAPI−A11
7614−Ctx−T−L−Sml−DAPI−A12
7614−Ctx−T−L−Sml−DAPI−A7
7614−Ctx−T−L−Sml−DAPI−A8
7614−Ctx−T−L−Sml−DAPI−A9
7614−Ctx−T−L−Sml−DAPI−B10
7614−Ctx−T−L−Sml−DAPI−B11
7614−Ctx−T−L−Sml−DAPI−B12
7614−Ctx−T−L−Sml−DAPI−B7
7614−Ctx−T−L−Sml−DAPI−B8
7614−Ctx−T−L−Sml−DAPI−B9
7614−Ctx−T−L−Sml−DAPI−C10
7614−Ctx−T−L−Sml−DAPI−C11
7614−Ctx−T−L−Sml−DAPI−C12
7614−Ctx−T−L−Sml−DAPI−C7
7614−Ctx−T−L−Sml−DAPI−C8
7614−Ctx−T−L−Sml−DAPI−C9
7614−Ctx−T−L−Sml−DAPI−D10
7614−Ctx−T−L−Sml−DAPI−D11
7614−Ctx−T−L−Sml−DAPI−D12
7614−Ctx−T−L−Sml−DAPI−D7
7614−Ctx−T−L−Sml−DAPI−D8
7614−Ctx−T−L−Sml−DAPI−D9
7614−Ctx−T−L−Sml−DAPI−E10
7614−Ctx−T−L−Sml−DAPI−E11
7614−Ctx−T−L−Sml−DAPI−E12
7614−Ctx−T−L−Sml−DAPI−E7
7614−Ctx−T−L−Sml−DAPI−E8
7614−Ctx−T−L−Sml−DAPI−E9
7614−Ctx−T−L−Sml−DAPI−F10
7614−Ctx−T−L−Sml−DAPI−F11
7614−Ctx−T−L−Sml−DAPI−F12
7614−Ctx−T−L−Sml−DAPI−F7
7614−Ctx−T−L−Sml−DAPI−F8
7614−Ctx−T−L−Sml−DAPI−F9
7614−Ctx−T−L−Sml−DAPI−G10
7614−Ctx−T−L−Sml−DAPI−G11
7614−Ctx−T−L−Sml−DAPI−G12
7614−Ctx−T−L−Sml−DAPI−G7
7614−Ctx−T−L−Sml−DAPI−G8
7614−Ctx−T−L−Sml−DAPI−G9
7614−Ctx−T−L−Sml−DAPI−H10
7614−Ctx−T−L−Sml−DAPI−H11
7614−Ctx−T−L−Sml−DAPI−H7
7614−Ctx−T−L−Sml−DAPI−H8
7614−Ctx−T−L−Sml−DAPI−H9
7614−Ctx−T−L−Sml−NeuN−A1
7614−Ctx−T−L−Sml−NeuN−A2
7614−Ctx−T−L−Sml−NeuN−A3
7614−Ctx−T−L−Sml−NeuN−A4
7614−Ctx−T−L−Sml−NeuN−A5
7614−Ctx−T−L−Sml−NeuN−A6
7614−Ctx−T−L−Sml−NeuN−B1
7614−Ctx−T−L−Sml−NeuN−B2
7614−Ctx−T−L−Sml−NeuN−B3
7614−Ctx−T−L−Sml−NeuN−B4
7614−Ctx−T−L−Sml−NeuN−B5
7614−Ctx−T−L−Sml−NeuN−B6
7614−Ctx−T−L−Sml−NeuN−C1
7614−Ctx−T−L−Sml−NeuN−C2
7614−Ctx−T−L−Sml−NeuN−C3
7614−Ctx−T−L−Sml−NeuN−C4
7614−Ctx−T−L−Sml−NeuN−C5
7614−Ctx−T−L−Sml−NeuN−C6
7614−Ctx−T−L−Sml−NeuN−D1
7614−Ctx−T−L−Sml−NeuN−D2
7614−Ctx−T−L−Sml−NeuN−D3
7614−Ctx−T−L−Sml−NeuN−D4
7614−Ctx−T−L−Sml−NeuN−D5
7614−Ctx−T−L−Sml−NeuN−D6
7614−Ctx−T−L−Sml−NeuN−E1
7614−Ctx−T−L−Sml−NeuN−E2
7614−Ctx−T−L−Sml−NeuN−E3
7614−Ctx−T−L−Sml−NeuN−E4
7614−Ctx−T−L−Sml−NeuN−E5
7614−Ctx−T−L−Sml−NeuN−E6
7614−Ctx−T−L−Sml−NeuN−F1
7614−Ctx−T−L−Sml−NeuN−F2
7614−Ctx−T−L−Sml−NeuN−F3
7614−Ctx−T−L−Sml−NeuN−F4
7614−Ctx−T−L−Sml−NeuN−F5
7614−Ctx−T−L−Sml−NeuN−F6
7614−Ctx−T−L−Sml−NeuN−G1
7614−Ctx−T−L−Sml−NeuN−G2
7614−Ctx−T−L−Sml−NeuN−G3
7614−Ctx−T−L−Sml−NeuN−G4
7614−Ctx−T−L−Sml−NeuN−G5
7614−Ctx−T−L−Sml−NeuN−G6
7614−Ctx−T−L−Sml−NeuN−H1
7614−Ctx−T−L−Sml−NeuN−H2
7614−Ctx−T−L−Sml−NeuN−H3
7614−Ctx−T−L−Sml−NeuN−H4
7614−Ctx−T−L−Sml−NeuN−H5
7614−Ctx−T−L−Sml−NeuN−H6
JGG−Cells

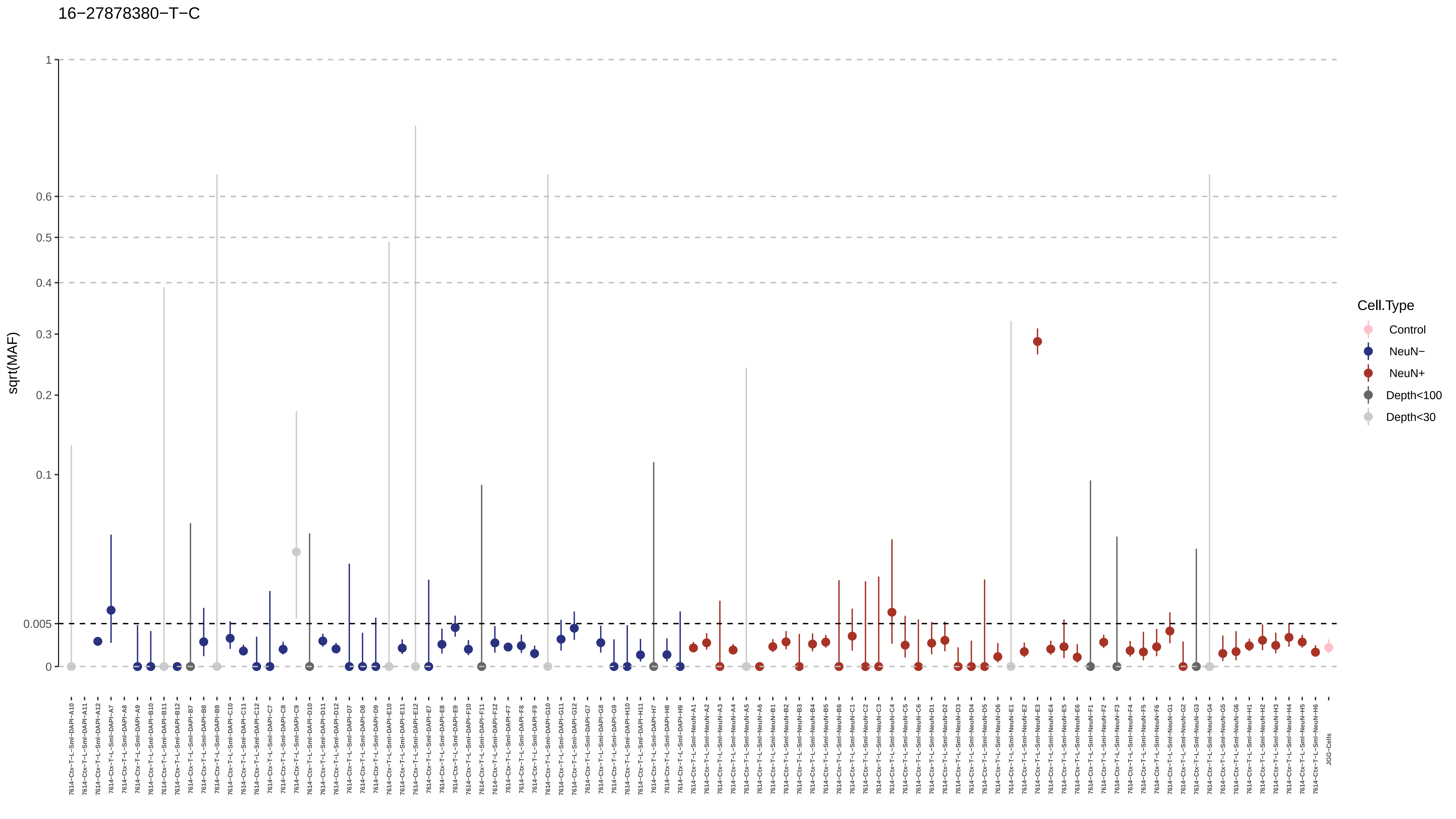
16−27878380−T−C
sqrt(MAF)
1
0.6
0.5
0.4
0.3
0.2
0.1
0.005
0
JGG-Cells
Cell.Type
Control
NeuN−
NeuN+
Depth<100
Depth<30

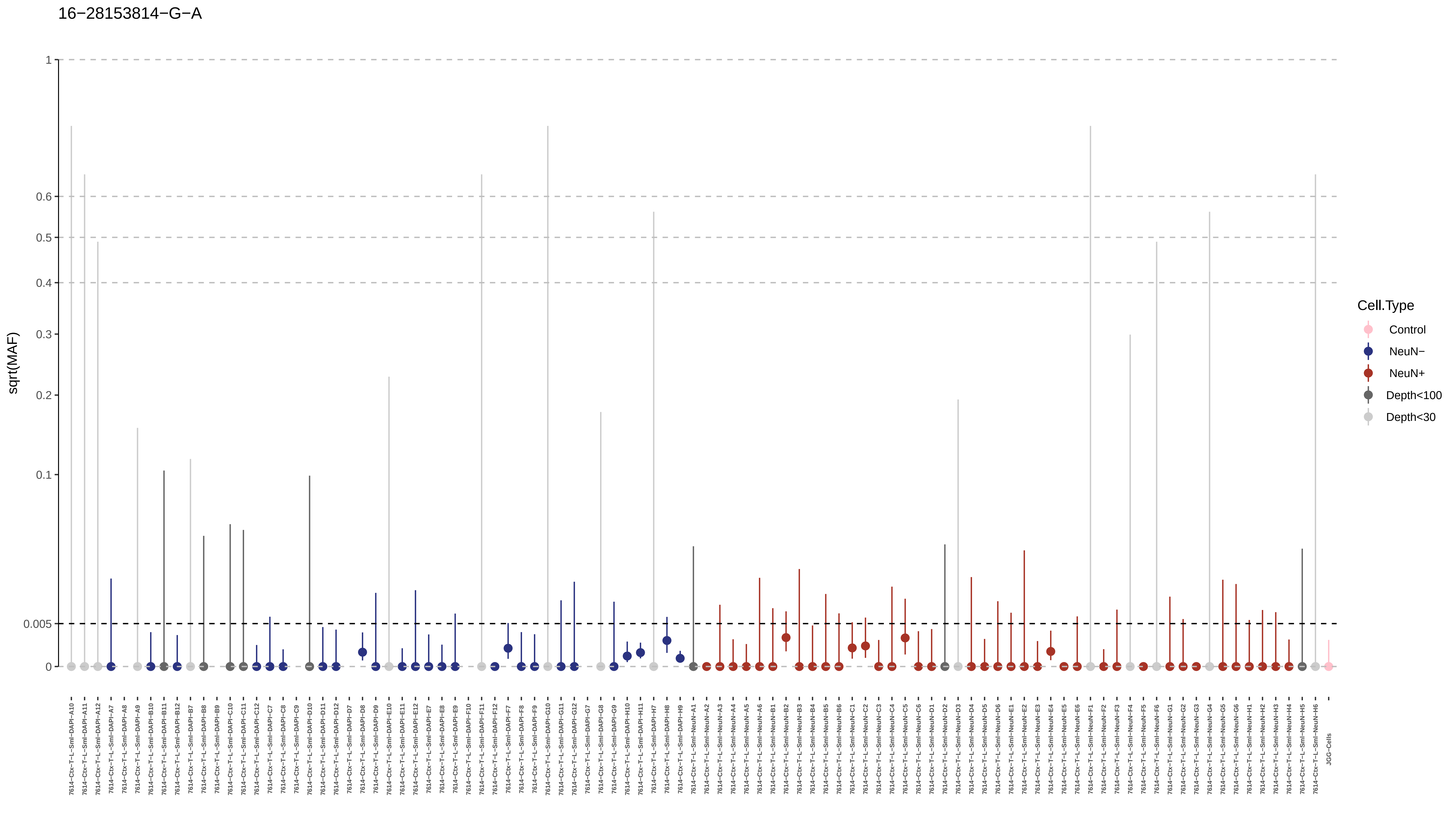

16−28153814−G−A
sqrt(MAF)
1
0.6
0.5
0.4
0.3
0.2
0.1
0.005
0
Cell.Type
Control
NeuN−
NeuN+
Depth<100
Depth<30

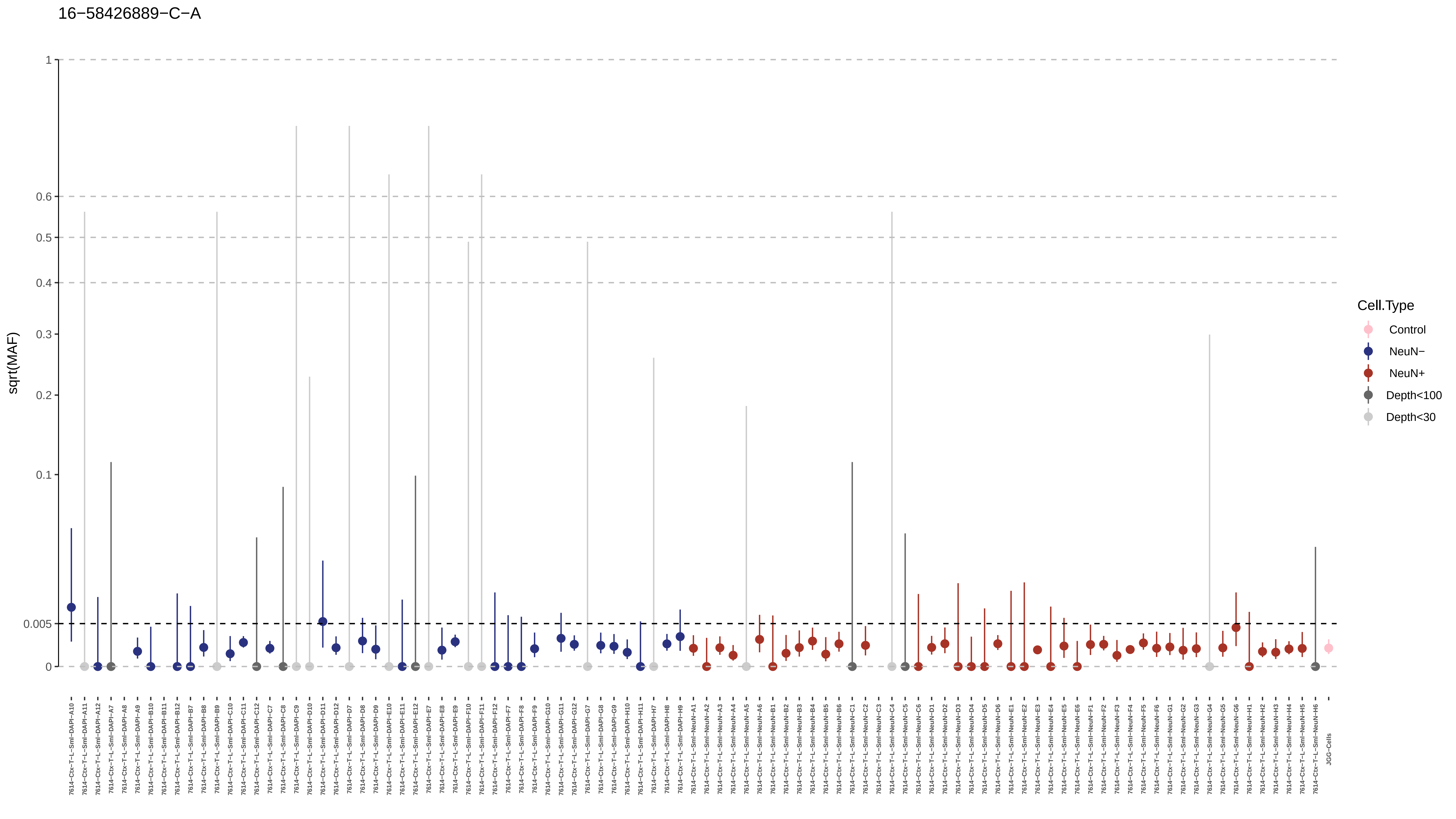
16−58426889−C−A
sqrt(MAF)
1
0.6
0.5
0.4
0.3
0.2
0.1
0.005
0
Cell.Type
Control
NeuN−
NeuN+
Depth<100
Depth<30

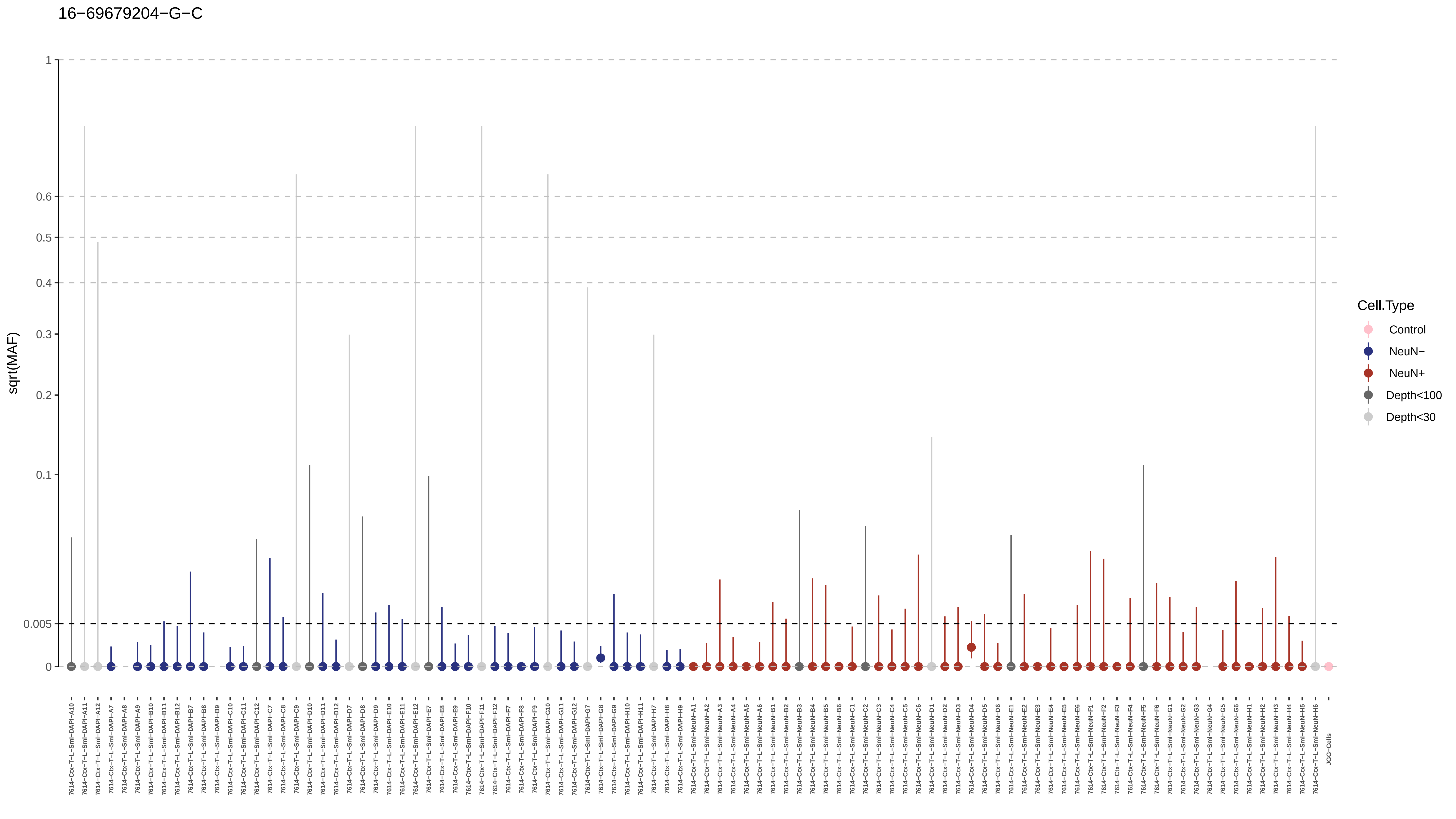

16−69679204−G−C
sqrt(MAF)
1
0.6
0.5
0.4
0.3
0.2
0.1
0.005
0
Cell.Type
Control
NeuN−
NeuN+
Depth<100
Depth<30
JGG-Cells

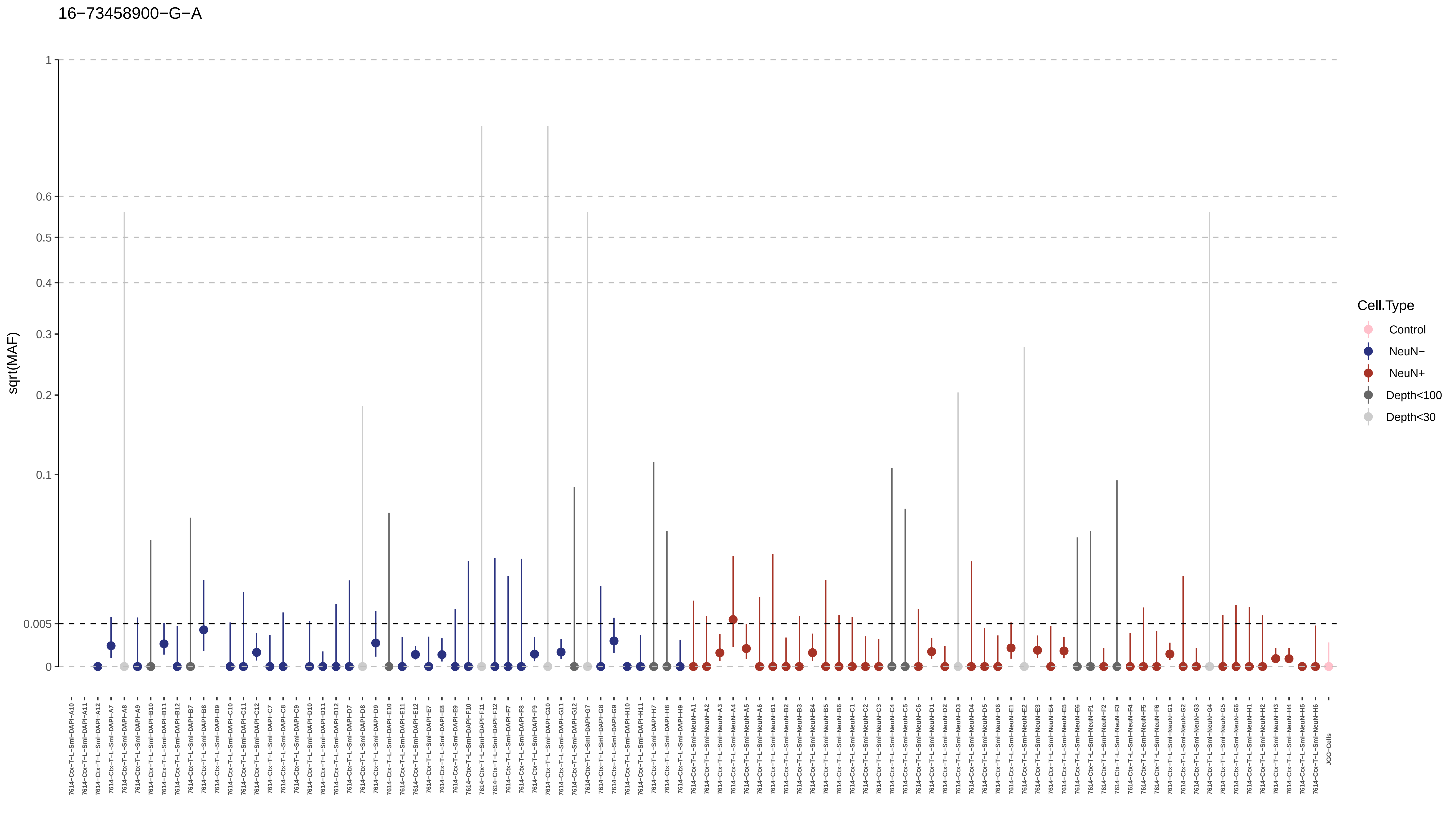
16−73458900−G−A
sqrt(MAF)
1
0.6
0.5
0.4
0.3
0.2
0.1
0.005
0
Cell.Type
Control
NeuN−
NeuN+
Depth<100
Depth<30
7614−Ctx−T−L−Sml−DAPI−A10
7614−Ctx−T−L−Sml−DAPI−A11
7614−Ctx−T−L−Sml−DAPI−A12
7614−Ctx−T−L−Sml−DAPI−A7
7614−Ctx−T−L−Sml−DAPI−A8
7614−Ctx−T−L−Sml−DAPI−A9
7614−Ctx−T−L−Sml−DAPI−B10
7614−Ctx−T−L−Sml−DAPI−B11
7614−Ctx−T−L−Sml−DAPI−B12
7614−Ctx−T−L−Sml−DAPI−B7
7614−Ctx−T−L−Sml−DAPI−B8
7614−Ctx−T−L−Sml−DAPI−B9
7614−Ctx−T−L−Sml−DAPI−C10
7614−Ctx−T−L−Sml−DAPI−C11
7614−Ctx−T−L−Sml−DAPI−C12
7614−Ctx−T−L−Sml−DAPI−C7
7614−Ctx−T−L−Sml−DAPI−C8
7614−Ctx−T−L−Sml−DAPI−C9
7614−Ctx−T−L−Sml−DAPI−D10
7614−Ctx−T−L−Sml−DAPI−D11
7614−Ctx−T−L−Sml−DAPI−D12
7614−Ctx−T−L−Sml−DAPI−D7
7614−Ctx−T−L−Sml−DAPI−D8
7614−Ctx−T−L−Sml−DAPI−D9
7614−Ctx−T−L−Sml−DAPI−E10
7614−Ctx−T−L−Sml−DAPI−E11
7614−Ctx−T−L−Sml−DAPI−E12
7614−Ctx−T−L−Sml−DAPI−E7
7614−Ctx−T−L−Sml−DAPI−E8
7614−Ctx−T−L−Sml−DAPI−E9
7614−Ctx−T−L−Sml−DAPI−F10
7614−Ctx−T−L−Sml−DAPI−F11
7614−Ctx−T−L−Sml−DAPI−F12
7614−Ctx−T−L−Sml−DAPI−F7
7614−Ctx−T−L−Sml−DAPI−F8
7614−Ctx−T−L−Sml−DAPI−F9
7614−Ctx−T−L−Sml−DAPI−G10
7614−Ctx−T−L−Sml−DAPI−G11
7614−Ctx−T−L−Sml−DAPI−G12
7614−Ctx−T−L−Sml−DAPI−G7
7614−Ctx−T−L−Sml−DAPI−G8
7614−Ctx−T−L−Sml−DAPI−G9
7614−Ctx−T−L−Sml−DAPI−H10
7614−Ctx−T−L−Sml−DAPI−H11
7614−Ctx−T−L−Sml−DAPI−H7
7614−Ctx−T−L−Sml−DAPI−H8
7614−Ctx−T−L−Sml−DAPI−H9
7614−Ctx−T−L−Sml−NeuN−A1
7614−Ctx−T−L−Sml−NeuN−A2
7614−Ctx−T−L−Sml−NeuN−A3
7614−Ctx−T−L−Sml−NeuN−A4
7614−Ctx−T−L−Sml−NeuN−A5
7614−Ctx−T−L−Sml−NeuN−A6
7614−Ctx−T−L−Sml−NeuN−B1
7614−Ctx−T−L−Sml−NeuN−B2
7614−Ctx−T−L−Sml−NeuN−B3
7614−Ctx−T−L−Sml−NeuN−B4
7614−Ctx−T−L−Sml−NeuN−B5
7614−Ctx−T−L−Sml−NeuN−B6
7614−Ctx−T−L−Sml−NeuN−C1
7614−Ctx−T−L−Sml−NeuN−C2
7614−Ctx−T−L−Sml−NeuN−C3
7614−Ctx−T−L−Sml−NeuN−C4
7614−Ctx−T−L−Sml−NeuN−C5
7614−Ctx−T−L−Sml−NeuN−C6
7614−Ctx−T−L−Sml−NeuN−D1
7614−Ctx−T−L−Sml−NeuN−D2
7614−Ctx−T−L−Sml−NeuN−D3
7614−Ctx−T−L−Sml−NeuN−D4
7614−Ctx−T−L−Sml−NeuN−D5
7614−Ctx−T−L−Sml−NeuN−D6
7614−Ctx−T−L−Sml−NeuN−E1
7614−Ctx−T−L−Sml−NeuN−E2
7614−Ctx−T−L−Sml−NeuN−E3
7614−Ctx−T−L−Sml−NeuN−E4
7614−Ctx−T−L−Sml−NeuN−E5
7614−Ctx−T−L−Sml−NeuN−E6
7614−Ctx−T−L−Sml−NeuN−F1
7614−Ctx−T−L−Sml−NeuN−F2
7614−Ctx−T−L−Sml−NeuN−F3
7614−Ctx−T−L−Sml−NeuN−F4
7614−Ctx−T−L−Sml−NeuN−F5
7614−Ctx−T−L−Sml−NeuN−F6
7614−Ctx−T−L−Sml−NeuN−G1
7614−Ctx−T−L−Sml−NeuN−G2
7614−Ctx−T−L−Sml−NeuN−G3
7614−Ctx−T−L−Sml−NeuN−G4
7614−Ctx−T−L−Sml−NeuN−G5
7614−Ctx−T−L−Sml−NeuN−G6
7614−Ctx−T−L−Sml−NeuN−H1
7614−Ctx−T−L−Sml−NeuN−H2
7614−Ctx−T−L−Sml−NeuN−H3
7614−Ctx−T−L−Sml−NeuN−H4
7614−Ctx−T−L−Sml−NeuN−H5
7614−Ctx−T−L−Sml−NeuN−H6
JGG−Cells

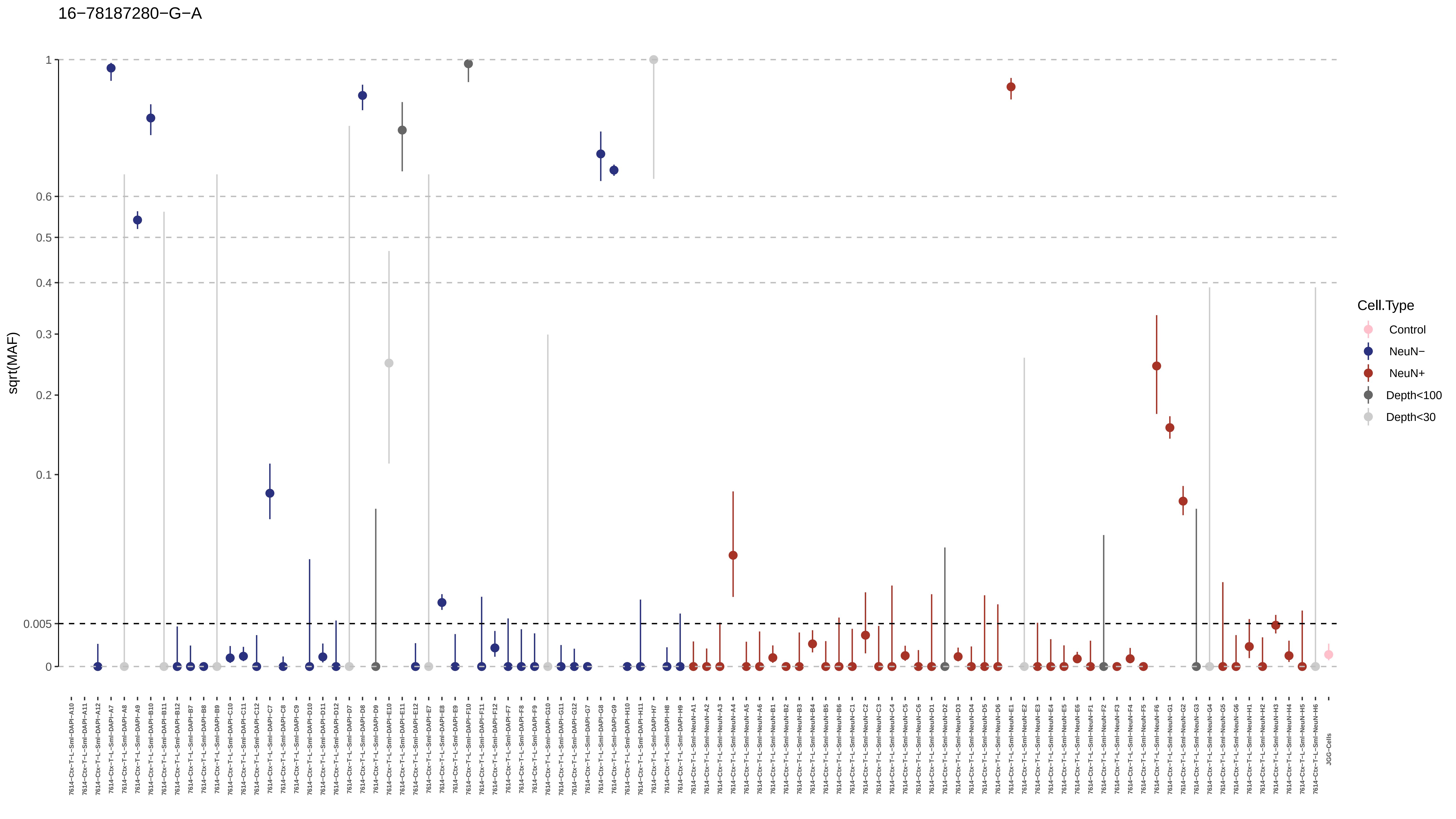
16−78187280−G−A
sqrt(MAF)
1
0.6
0.5
0.4
0.3
0.2
0.1
0.005
0
Cell.Type
Control
NeuN−
NeuN+
Depth<100
Depth<30

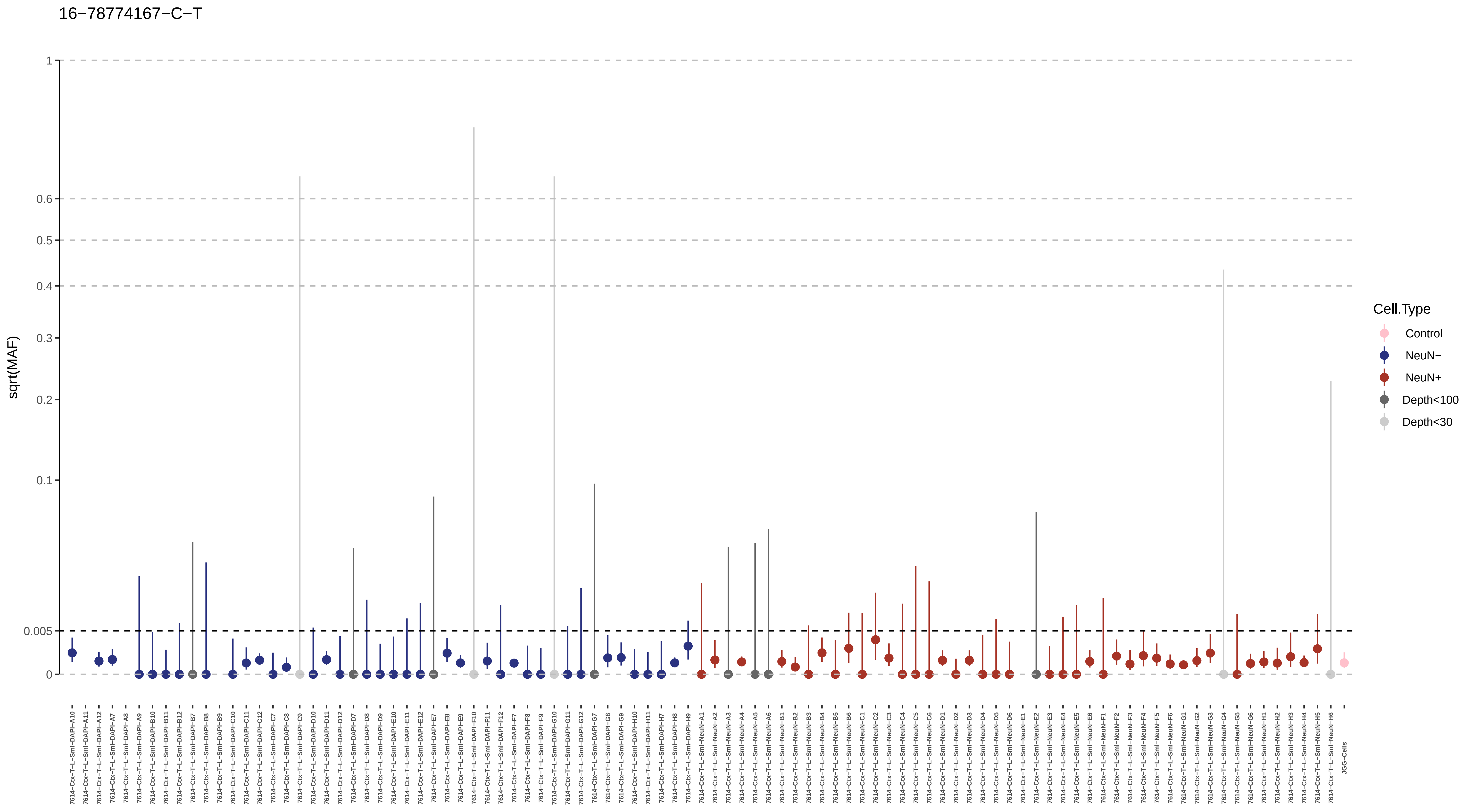

16−78774167−C−T
sqrt(MAF)
1
0.6
0.5
0.4
0.3
0.2
0.1
0.005
0
Cell.Type
Control
NeuN−
NeuN+
Depth<100
Depth<30
JGG-Cells

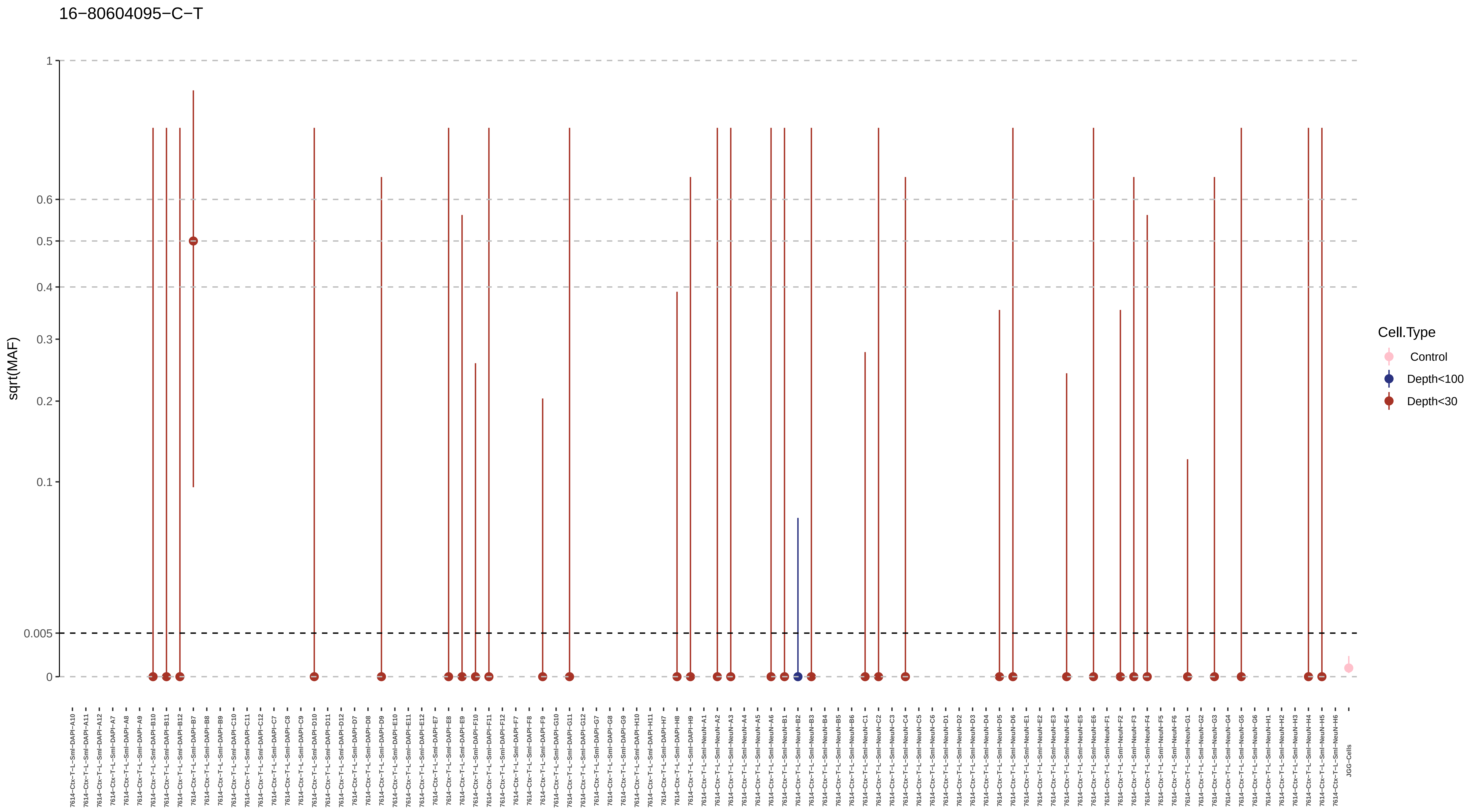
16−80604095−C−T
sqrt(MAF)
1
0.6
0.5
0.4
0.3
0.2
0.1
0.005
0
Cell.Type
Control
Depth<100
Depth<30
7614−Ctx−T−L−Sml−DAPI−A10
7614−Ctx−T−L−Sml−DAPI−A11
7614−Ctx−T−L−Sml−DAPI−A12
7614−Ctx−T−L−Sml−DAPI−A7
7614−Ctx−T−L−Sml−DAPI−A8
7614−Ctx−T−L−Sml−DAPI−A9
7614−Ctx−T−L−Sml−DAPI−B10
7614−Ctx−T−L−Sml−DAPI−B11
7614−Ctx−T−L−Sml−DAPI−B12
7614−Ctx−T−L−Sml−DAPI−B7
7614−Ctx−T−L−Sml−DAPI−B8
7614−Ctx−T−L−Sml−DAPI−B9
7614−Ctx−T−L−Sml−DAPI−C10
7614−Ctx−T−L−Sml−DAPI−C11
7614−Ctx−T−L−Sml−DAPI−C12
7614−Ctx−T−L−Sml−DAPI−C7
7614−Ctx−T−L−Sml−DAPI−C8
7614−Ctx−T−L−Sml−DAPI−C9
7614−Ctx−T−L−Sml−DAPI−D10
7614−Ctx−T−L−Sml−DAPI−D11
7614−Ctx−T−L−Sml−DAPI−D12
7614−Ctx−T−L−Sml−DAPI−D7
7614−Ctx−T−L−Sml−DAPI−D8
7614−Ctx−T−L−Sml−DAPI−D9
7614−Ctx−T−L−Sml−DAPI−E10
7614−Ctx−T−L−Sml−DAPI−E11
7614−Ctx−T−L−Sml−DAPI−E12
7614−Ctx−T−L−Sml−DAPI−E7
7614−Ctx−T−L−Sml−DAPI−E8
7614−Ctx−T−L−Sml−DAPI−E9
7614−Ctx−T−L−Sml−DAPI−F10
7614−Ctx−T−L−Sml−DAPI−F11
7614−Ctx−T−L−Sml−DAPI−F12
7614−Ctx−T−L−Sml−DAPI−F7
7614−Ctx−T−L−Sml−DAPI−F8
7614−Ctx−T−L−Sml−DAPI−F9
7614−Ctx−T−L−Sml−DAPI−G10
7614−Ctx−T−L−Sml−DAPI−G11
7614−Ctx−T−L−Sml−DAPI−G12
7614−Ctx−T−L−Sml−DAPI−G7
7614−Ctx−T−L−Sml−DAPI−G8
7614−Ctx−T−L−Sml−DAPI−G9
7614−Ctx−T−L−Sml−DAPI−H10
7614−Ctx−T−L−Sml−DAPI−H11
7614−Ctx−T−L−Sml−DAPI−H7
7614−Ctx−T−L−Sml−DAPI−H8
7614−Ctx−T−L−Sml−DAPI−H9
7614−Ctx−T−L−Sml−NeuN−A1
7614−Ctx−T−L−Sml−NeuN−A2
7614−Ctx−T−L−Sml−NeuN−A3
7614−Ctx−T−L−Sml−NeuN−A4
7614−Ctx−T−L−Sml−NeuN−A5
7614−Ctx−T−L−Sml−NeuN−A6
7614−Ctx−T−L−Sml−NeuN−B1
7614−Ctx−T−L−Sml−NeuN−B2
7614−Ctx−T−L−Sml−NeuN−B3
7614−Ctx−T−L−Sml−NeuN−B4
7614−Ctx−T−L−Sml−NeuN−B5
7614−Ctx−T−L−Sml−NeuN−B6
7614−Ctx−T−L−Sml−NeuN−C1
7614−Ctx−T−L−Sml−NeuN−C2
7614−Ctx−T−L−Sml−NeuN−C3
7614−Ctx−T−L−Sml−NeuN−C4
7614−Ctx−T−L−Sml−NeuN−C5
7614−Ctx−T−L−Sml−NeuN−C6
7614−Ctx−T−L−Sml−NeuN−D1
7614−Ctx−T−L−Sml−NeuN−D2
7614−Ctx−T−L−Sml−NeuN−D3
7614−Ctx−T−L−Sml−NeuN−D4
7614−Ctx−T−L−Sml−NeuN−D5
7614−Ctx−T−L−Sml−NeuN−D6
7614−Ctx−T−L−Sml−NeuN−E1
7614−Ctx−T−L−Sml−NeuN−E2
7614−Ctx−T−L−Sml−NeuN−E3
7614−Ctx−T−L−Sml−NeuN−E4
7614−Ctx−T−L−Sml−NeuN−E5
7614−Ctx−T−L−Sml−NeuN−E6
7614−Ctx−T−L−Sml−NeuN−F1
7614−Ctx−T−L−Sml−NeuN−F2
7614−Ctx−T−L−Sml−NeuN−F3
7614−Ctx−T−L−Sml−NeuN−F4
7614−Ctx−T−L−Sml−NeuN−F5
7614−Ctx−T−L−Sml−NeuN−F6
7614−Ctx−T−L−Sml−NeuN−G1
7614−Ctx−T−L−Sml−NeuN−G2
7614−Ctx−T−L−Sml−NeuN−G3
7614−Ctx−T−L−Sml−NeuN−G4
7614−Ctx−T−L−Sml−NeuN−G5
7614−Ctx−T−L−Sml−NeuN−G6
7614−Ctx−T−L−Sml−NeuN−H1
7614−Ctx−T−L−Sml−NeuN−H2
7614−Ctx−T−L−Sml−NeuN−H3
7614−Ctx−T−L−Sml−NeuN−H4
7614−Ctx−T−L−Sml−NeuN−H5
7614−Ctx−T−L−Sml−NeuN−H6
JGG-Cells

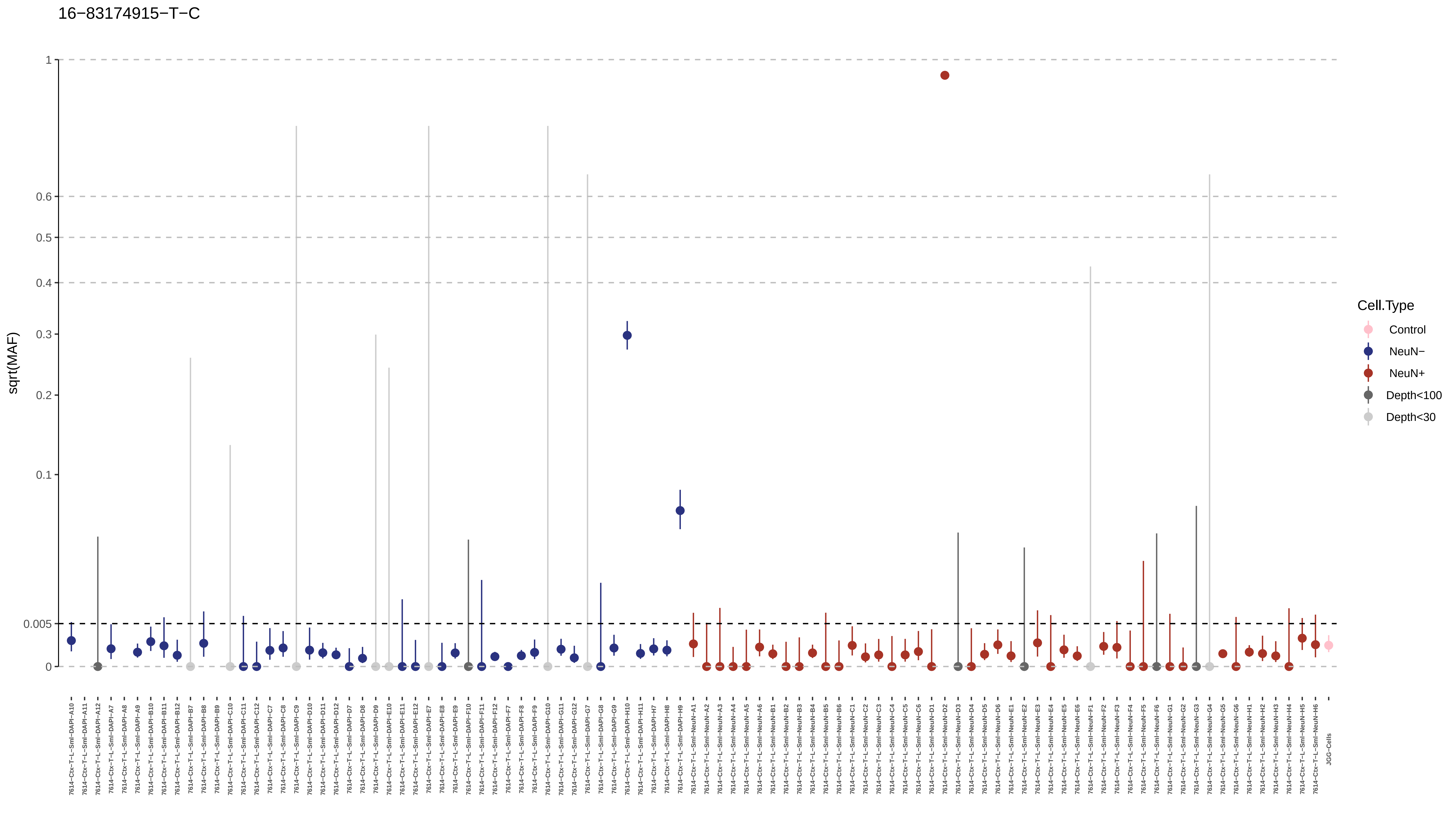
16−83174915−T−C
sqrt(MAF)
1
0.6
0.5
0.4
0.3
0.2
0.1
0.005
0
Cell.Type
Control
NeuN−
NeuN+
Depth<100
Depth<30

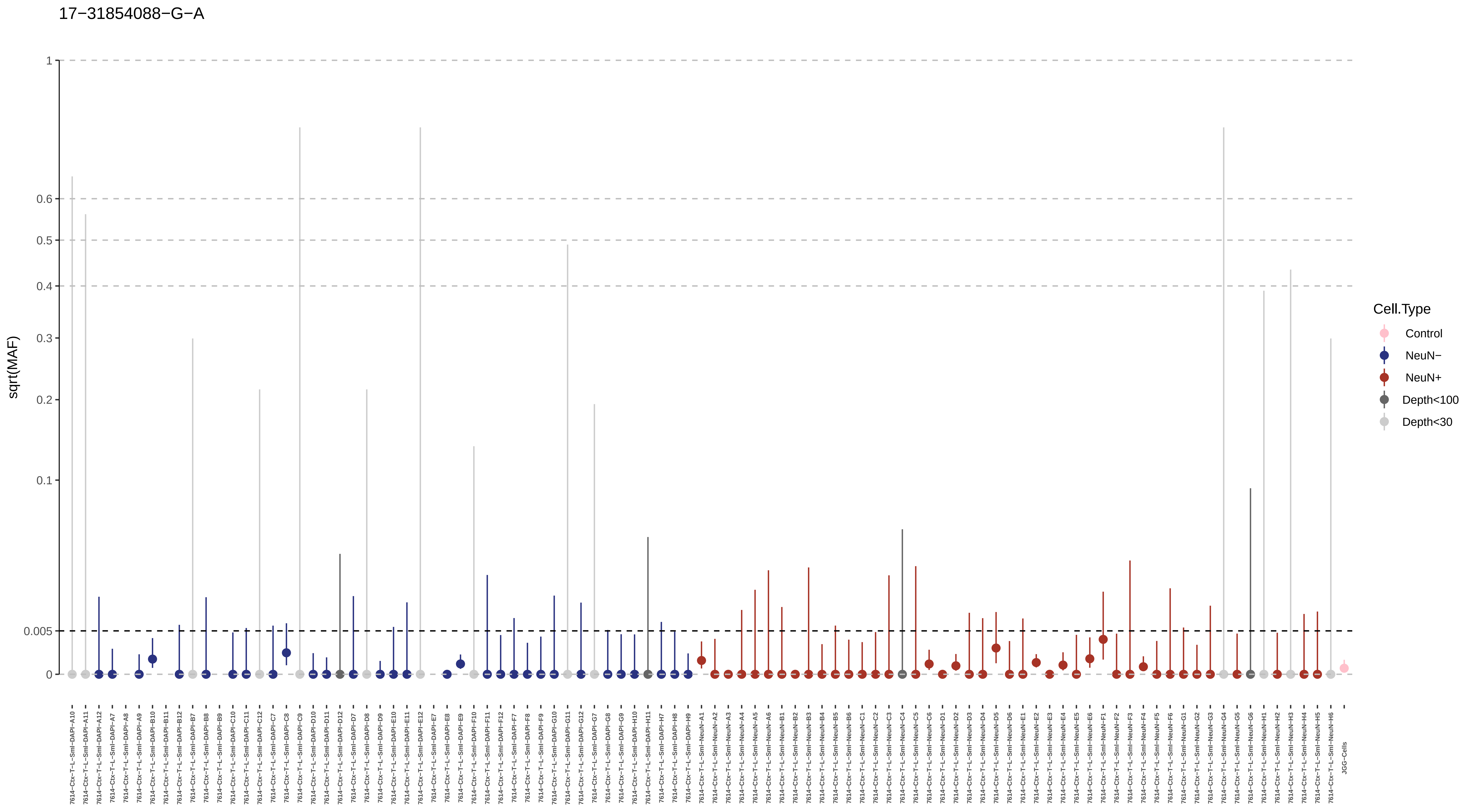

17−31854088−G−A
sqrt(MAF)
1
0.6
0.5
0.4
0.3
0.2
0.1
0.005
0
Cell.Type
Control
NeuN−
NeuN+
Depth<100
Depth<30
JGG-Cells

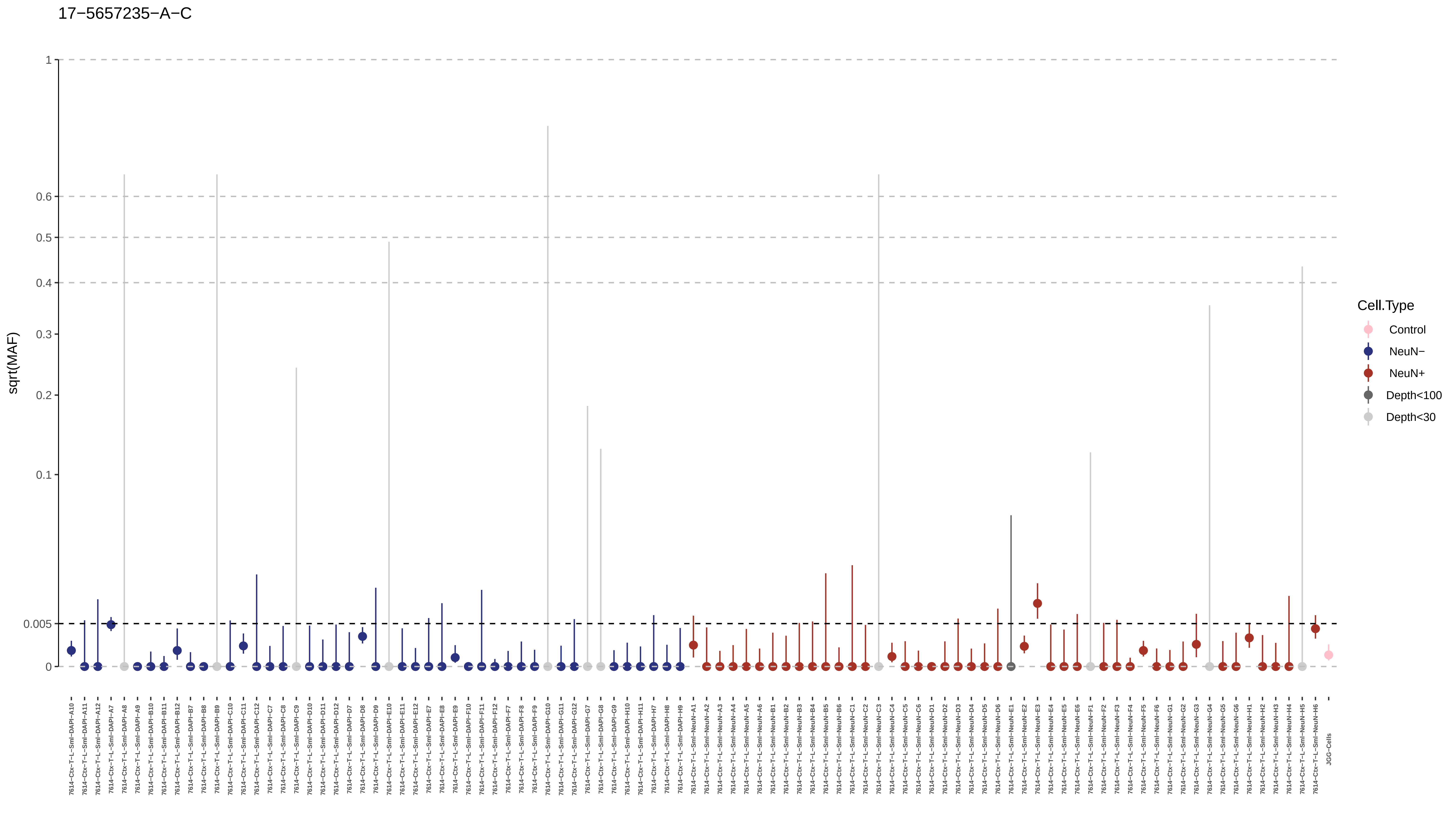

17−5657235−A−C
sqrt(MAF)
1
0.6
0.5
0.4
0.3
0.2
0.1
0.005
0
Cell.Type
Control
NeuN−
NeuN+
Depth<100
Depth<30
JGG-Cells

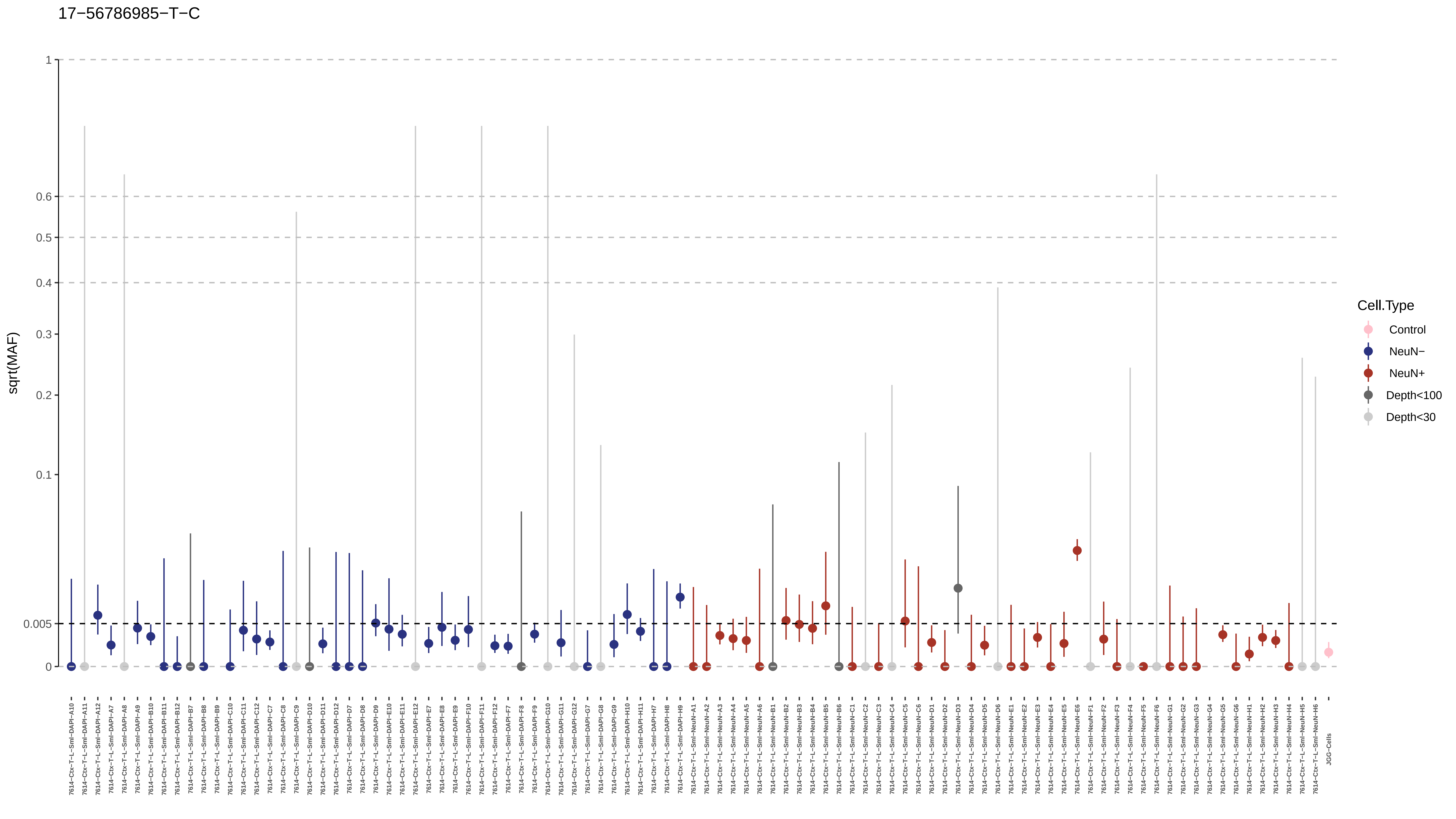

17−56786985−T−C
sqrt(MAF)
1
0.6
0.5
0.4
0.3
0.2
0.1
0.005
0
Cell.Type
Control
NeuN−
NeuN+
Depth<100
Depth<30
7614−Ctx−T−L−Sml−DAPI−A10
7614−Ctx−T−L−Sml−DAPI−A11
7614−Ctx−T−L−Sml−DAPI−A12
7614−Ctx−T−L−Sml−DAPI−A7
7614−Ctx−T−L−Sml−DAPI−A8
7614−Ctx−T−L−Sml−DAPI−A9
7614−Ctx−T−L−Sml−DAPI−B10
7614−Ctx−T−L−Sml−DAPI−B11
7614−Ctx−T−L−Sml−DAPI−B12
7614−Ctx−T−L−Sml−DAPI−B7
7614−Ctx−T−L−Sml−DAPI−B8
7614−Ctx−T−L−Sml−DAPI−B9
7614−Ctx−T−L−Sml−DAPI−C10
7614−Ctx−T−L−Sml−DAPI−C11
7614−Ctx−T−L−Sml−DAPI−C12
7614−Ctx−T−L−Sml−DAPI−C7
7614−Ctx−T−L−Sml−DAPI−C8
7614−Ctx−T−L−Sml−DAPI−C9
7614−Ctx−T−L−Sml−DAPI−D10
7614−Ctx−T−L−Sml−DAPI−D11
7614−Ctx−T−L−Sml−DAPI−D12
7614−Ctx−T−L−Sml−DAPI−D7
7614−Ctx−T−L−Sml−DAPI−D8
7614−Ctx−T−L−Sml−DAPI−D9
7614−Ctx−T−L−Sml−DAPI−E10
7614−Ctx−T−L−Sml−DAPI−E11
7614−Ctx−T−L−Sml−DAPI−E12
7614−Ctx−T−L−Sml−DAPI−E7
7614−Ctx−T−L−Sml−DAPI−E8
7614−Ctx−T−L−Sml−DAPI−E9
7614−Ctx−T−L−Sml−DAPI−F10
7614−Ctx−T−L−Sml−DAPI−F11
7614−Ctx−T−L−Sml−DAPI−F12
7614−Ctx−T−L−Sml−DAPI−F7
7614−Ctx−T−L−Sml−DAPI−F8
7614−Ctx−T−L−Sml−DAPI−F9
7614−Ctx−T−L−Sml−DAPI−G10
7614−Ctx−T−L−Sml−DAPI−G11
7614−Ctx−T−L−Sml−DAPI−G12
7614−Ctx−T−L−Sml−DAPI−G7
7614−Ctx−T−L−Sml−DAPI−G8
7614−Ctx−T−L−Sml−DAPI−G9
7614−Ctx−T−L−Sml−DAPI−H10
7614−Ctx−T−L−Sml−DAPI−H11
7614−Ctx−T−L−Sml−DAPI−H7
7614−Ctx−T−L−Sml−DAPI−H8
7614−Ctx−T−L−Sml−DAPI−H9
7614−Ctx−T−L−Sml−NeuN−A1
7614−Ctx−T−L−Sml−NeuN−A2
7614−Ctx−T−L−Sml−NeuN−A3
7614−Ctx−T−L−Sml−NeuN−A4
7614−Ctx−T−L−Sml−NeuN−A5
7614−Ctx−T−L−Sml−NeuN−A6
7614−Ctx−T−L−Sml−NeuN−B1
7614−Ctx−T−L−Sml−NeuN−B2
7614−Ctx−T−L−Sml−NeuN−B3
7614−Ctx−T−L−Sml−NeuN−B4
7614−Ctx−T−L−Sml−NeuN−B5
7614−Ctx−T−L−Sml−NeuN−B6
7614−Ctx−T−L−Sml−NeuN−C1
7614−Ctx−T−L−Sml−NeuN−C2
7614−Ctx−T−L−Sml−NeuN−C3
7614−Ctx−T−L−Sml−NeuN−C4
7614−Ctx−T−L−Sml−NeuN−C5
7614−Ctx−T−L−Sml−NeuN−C6
7614−Ctx−T−L−Sml−NeuN−D1
7614−Ctx−T−L−Sml−NeuN−D2
7614−Ctx−T−L−Sml−NeuN−D3
7614−Ctx−T−L−Sml−NeuN−D4
7614−Ctx−T−L−Sml−NeuN−D5
7614−Ctx−T−L−Sml−NeuN−D6
7614−Ctx−T−L−Sml−NeuN−E1
7614−Ctx−T−L−Sml−NeuN−E2
7614−Ctx−T−L−Sml−NeuN−E3
7614−Ctx−T−L−Sml−NeuN−E4
7614−Ctx−T−L−Sml−NeuN−E5
7614−Ctx−T−L−Sml−NeuN−E6
7614−Ctx−T−L−Sml−NeuN−F1
7614−Ctx−T−L−Sml−NeuN−F2
7614−Ctx−T−L−Sml−NeuN−F3
7614−Ctx−T−L−Sml−NeuN−F4
7614−Ctx−T−L−Sml−NeuN−F5
7614−Ctx−T−L−Sml−NeuN−F6
7614−Ctx−T−L−Sml−NeuN−G1
7614−Ctx−T−L−Sml−NeuN−G2
7614−Ctx−T−L−Sml−NeuN−G3
7614−Ctx−T−L−Sml−NeuN−G4
7614−Ctx−T−L−Sml−NeuN−G5
7614−Ctx−T−L−Sml−NeuN−G6
7614−Ctx−T−L−Sml−NeuN−H1
7614−Ctx−T−L−Sml−NeuN−H2
7614−Ctx−T−L−Sml−NeuN−H3
7614−Ctx−T−L−Sml−NeuN−H4
7614−Ctx−T−L−Sml−NeuN−H5
7614−Ctx−T−L−Sml−NeuN−H6
JGG-Cells

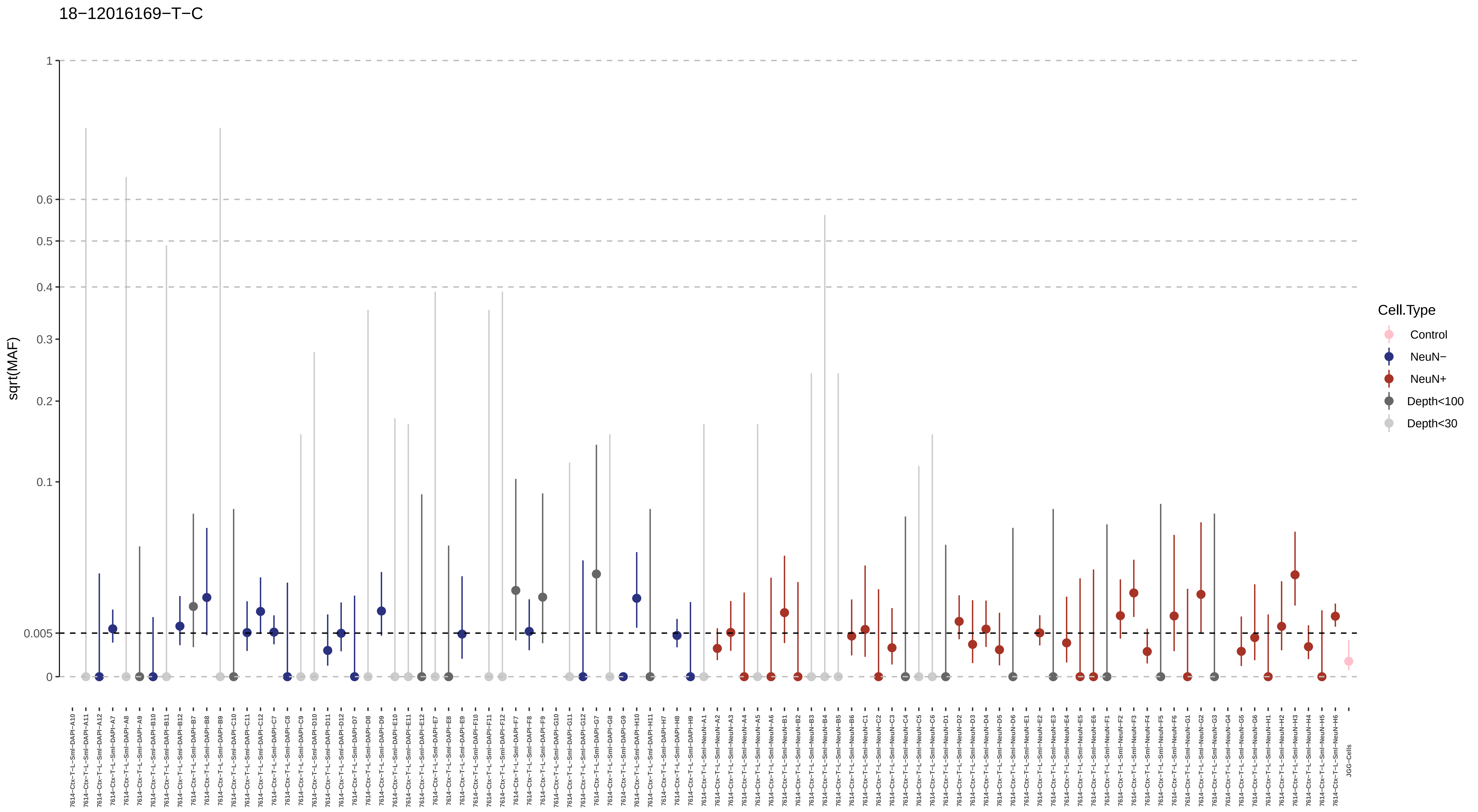

18−12016169−T−C
sqrt(MAF)
Cell.Type
Control
NeuN−
NeuN+
Depth<100
Depth<30

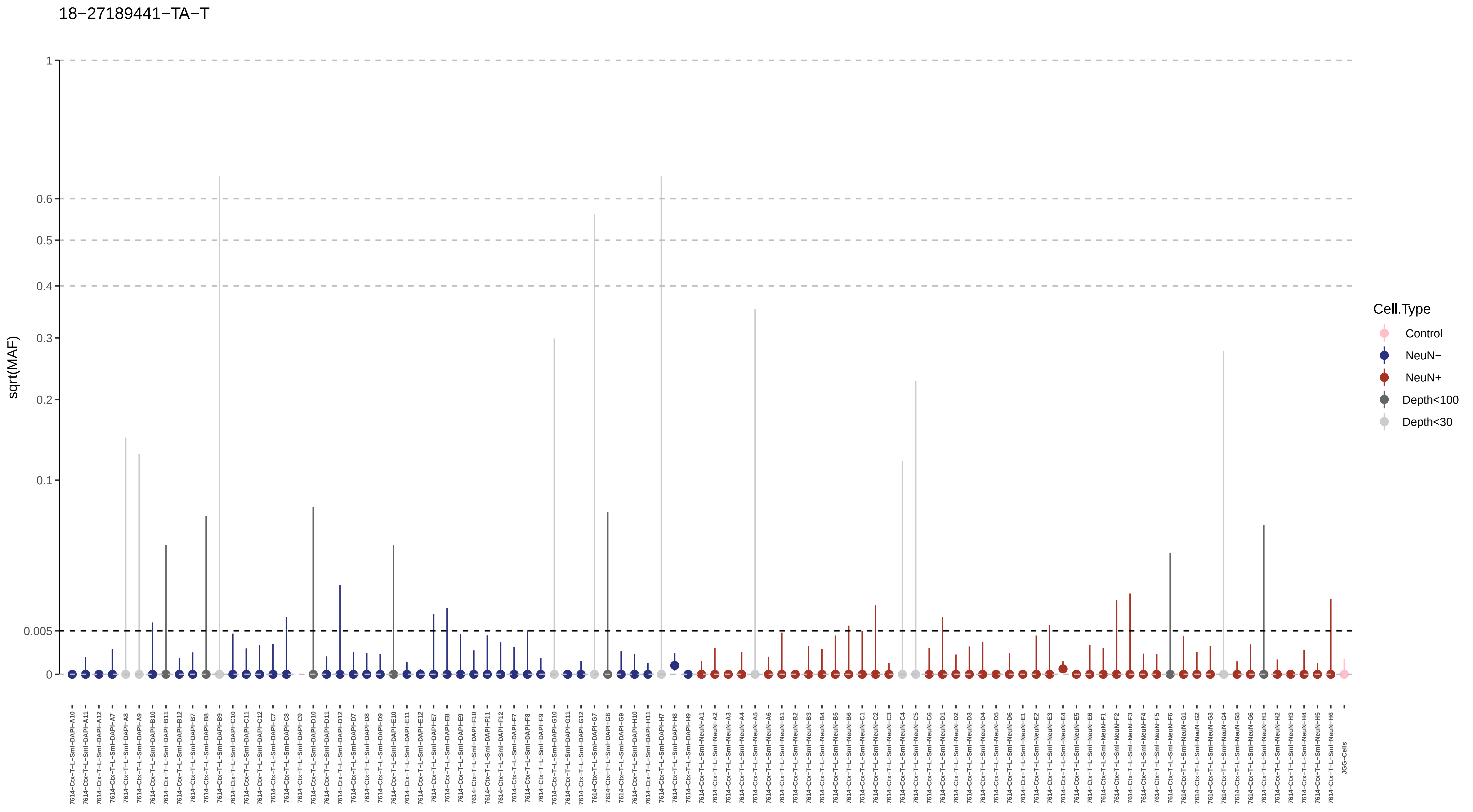

18−27189441−TA−T
sqrt(MAF)
1
0.6
0.5
0.4
0.3
0.2
0.1
0.005
0
Cell.Type
Control
NeuN−
NeuN+
Depth<100
Depth<30
JGG-Cells

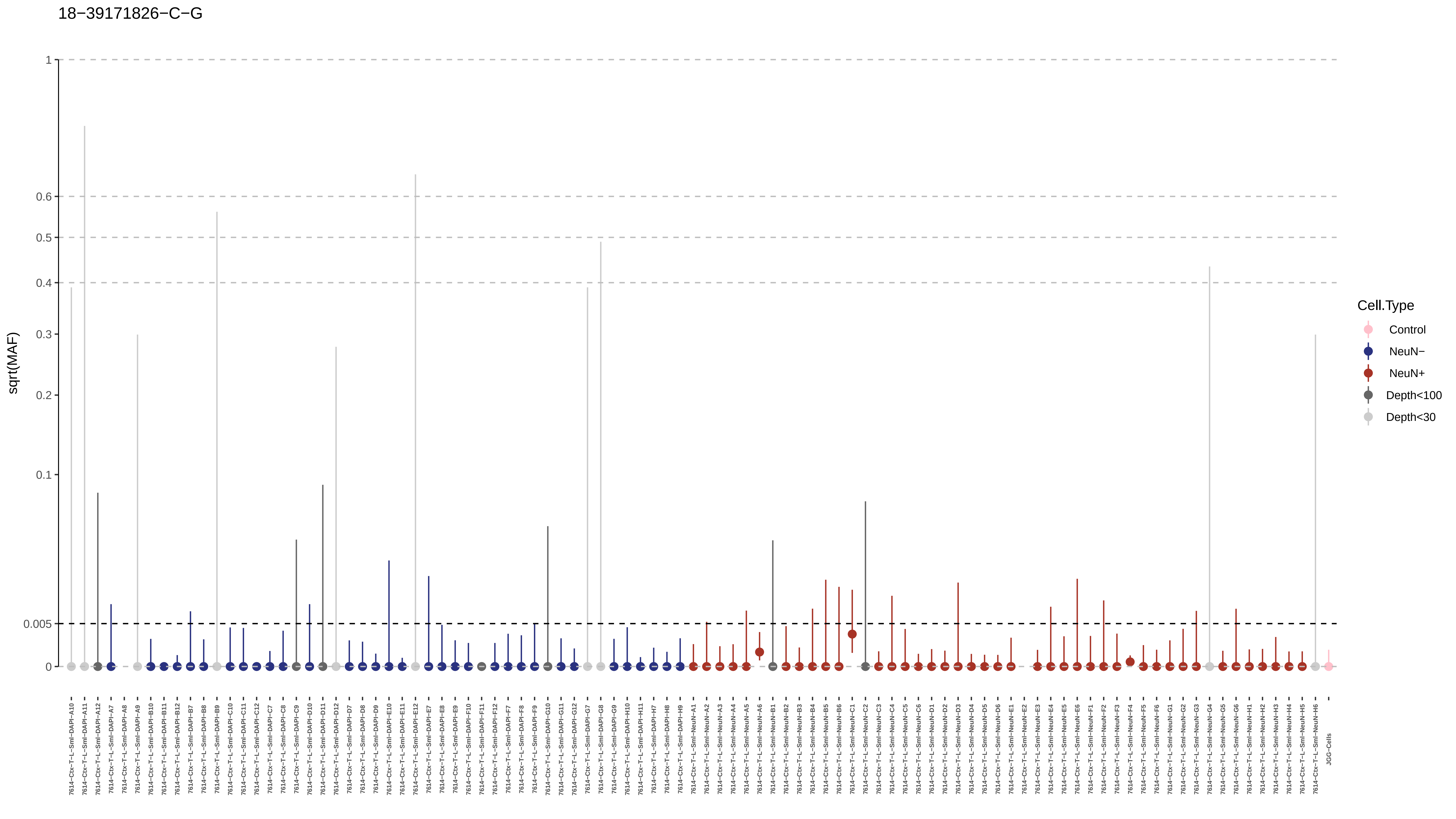

18−39171826−C−G
sqrt(MAF)
1
0.6
0.5
0.4
0.3
0.2
0.1
0.005
0
Cell.Type
Control
NeuN−
NeuN+
Depth<100
Depth<30

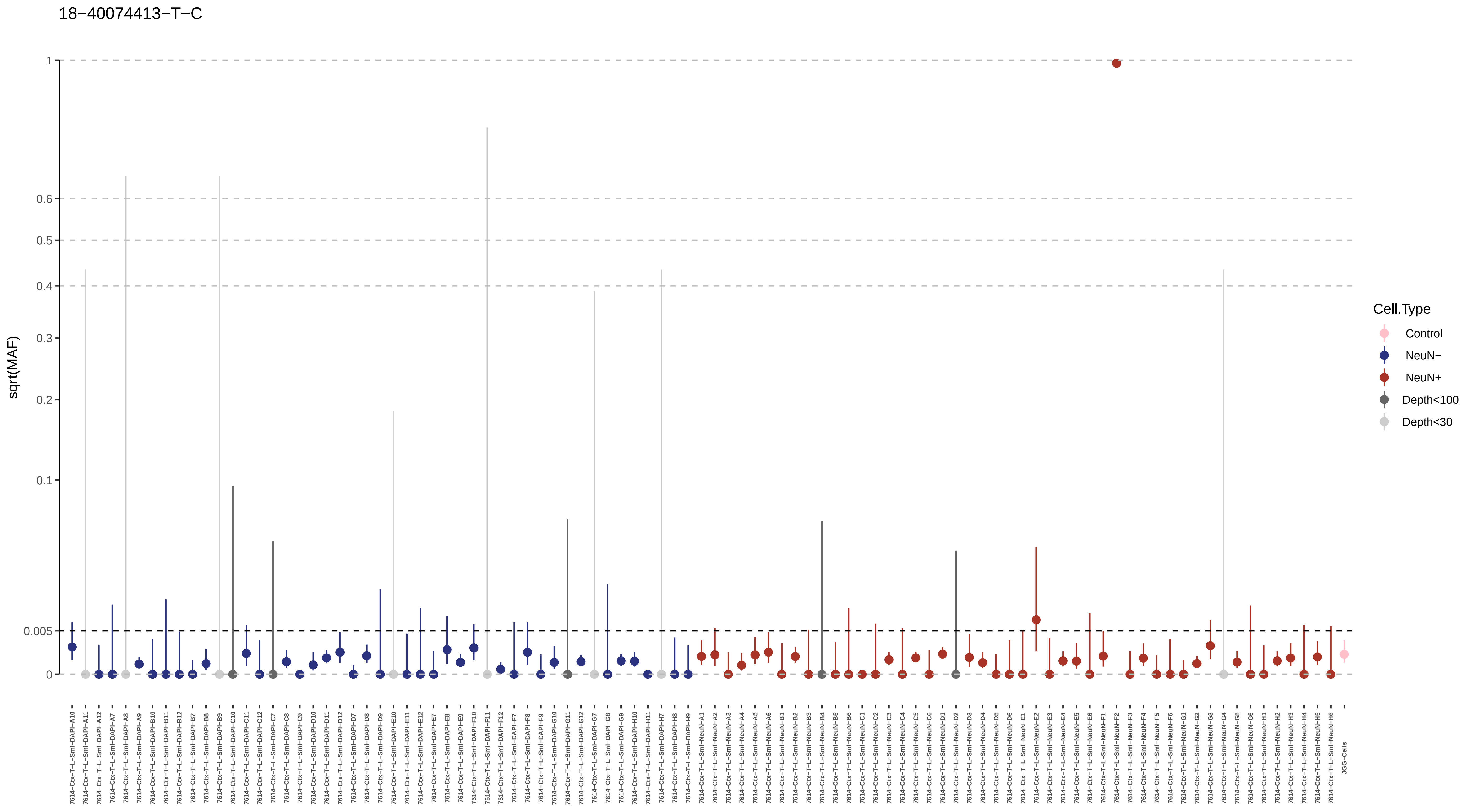
18−40074413−T−C
sqrt(MAF)
1
0.6
0.5
0.4
0.3
0.2
0.1
0.005
0
Cell.Type
Control
NeuN−
NeuN+
Depth<100
Depth<30

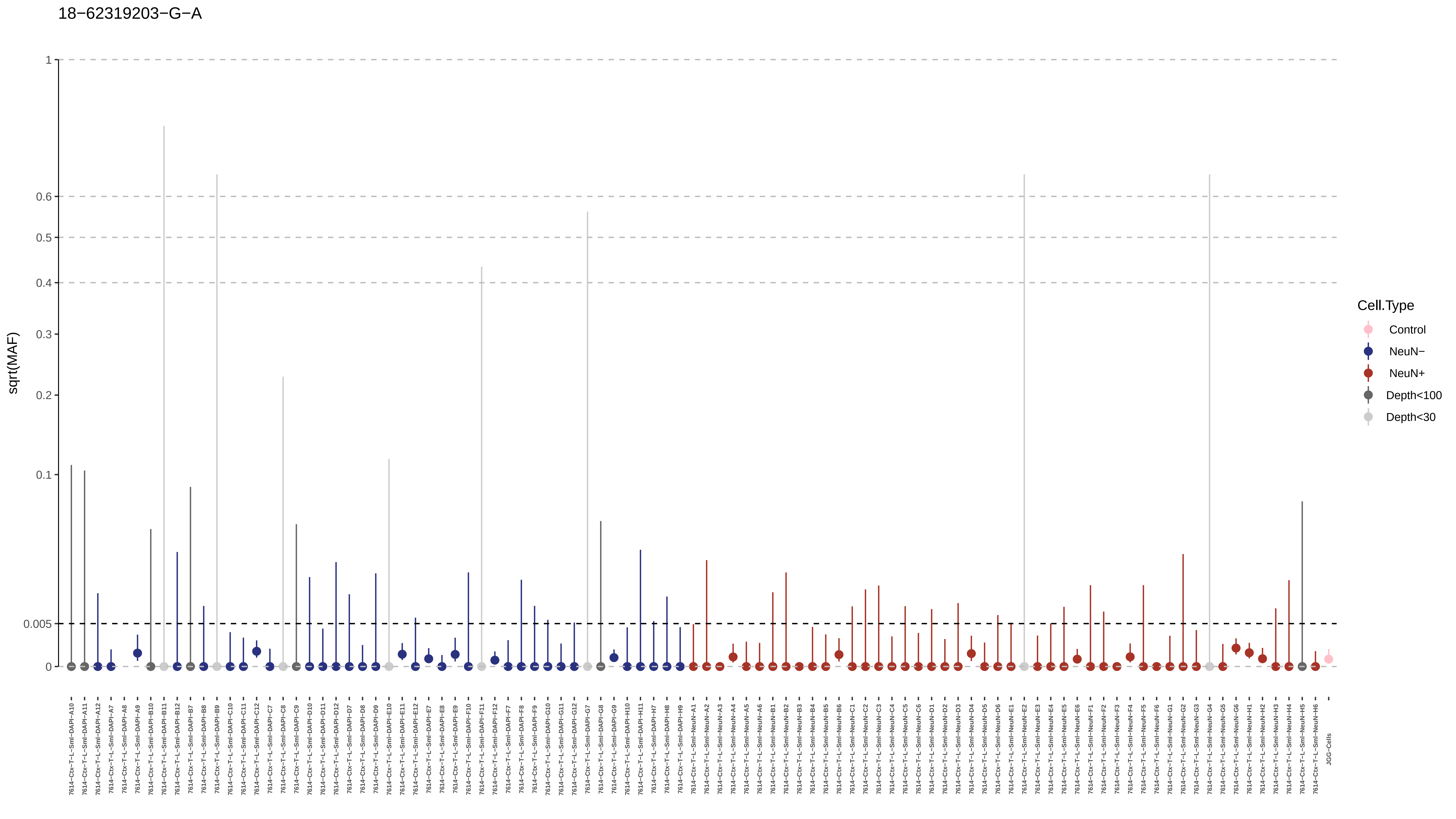
18−62319203−G−A
sqrt(MAF)
1
0.6
0.5
0.4
0.3
0.2
0.1
0.005
0
Cell.Type
Control
NeuN−
NeuN+
Depth<100
Depth<30

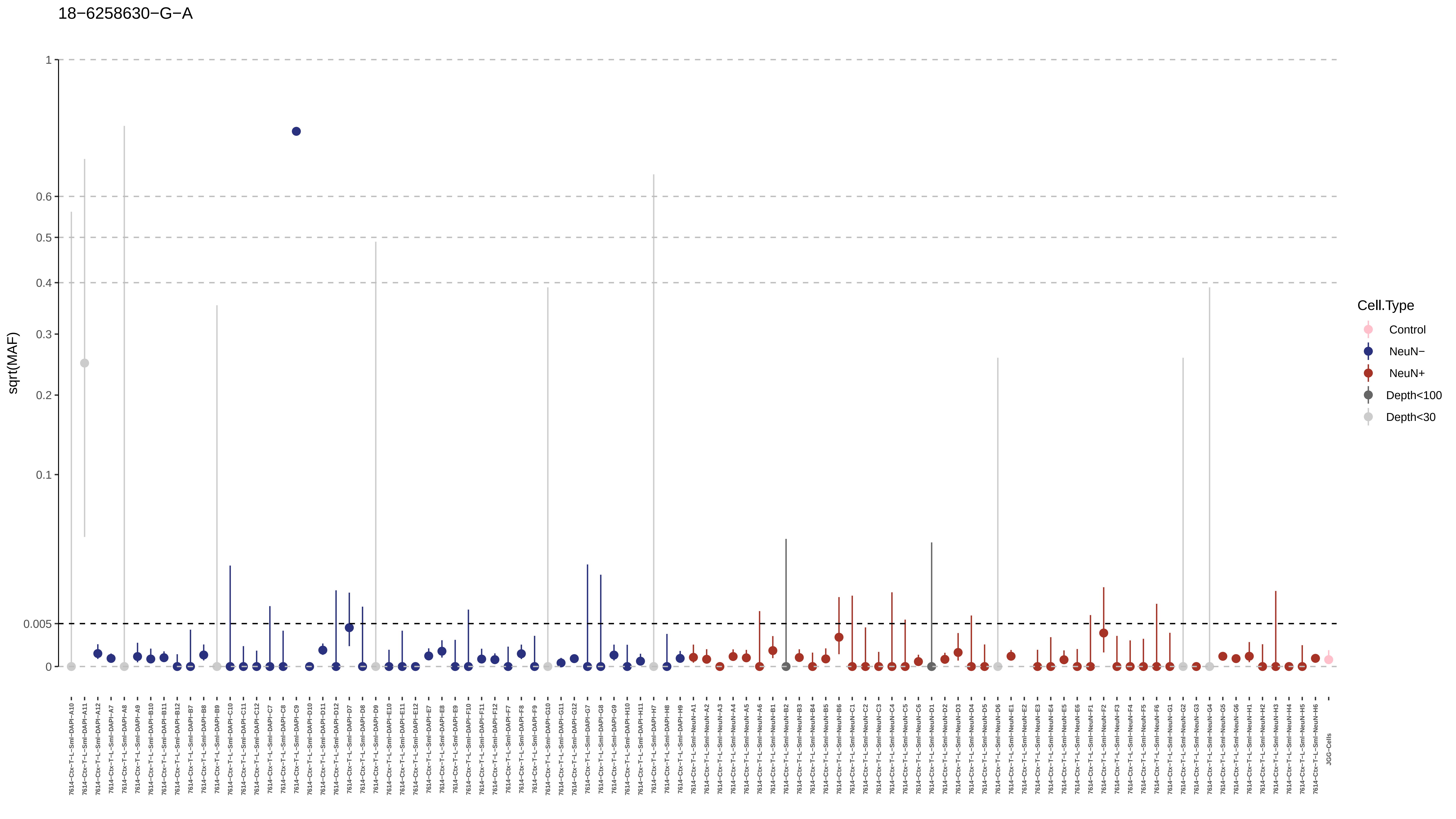
18−6258630−G−A
sqrt(MAF)
1
0.6
0.5
0.4
0.3
0.2
0.1
0.005
0
Cell.Type
Control
NeuN−
NeuN+
Depth<100
Depth<30
7614−Ctx−T−L−Sml−DAPI−A10
7614−Ctx−T−L−Sml−DAPI−A11
7614−Ctx−T−L−Sml−DAPI−A12
7614−Ctx−T−L−Sml−DAPI−A7
7614−Ctx−T−L−Sml−DAPI−A8
7614−Ctx−T−L−Sml−DAPI−A9
7614−Ctx−T−L−Sml−DAPI−B10
7614−Ctx−T−L−Sml−DAPI−B11
7614−Ctx−T−L−Sml−DAPI−B12
7614−Ctx−T−L−Sml−DAPI−B7
7614−Ctx−T−L−Sml−DAPI−B8
7614−Ctx−T−L−Sml−DAPI−B9
7614−Ctx−T−L−Sml−DAPI−C10
7614−Ctx−T−L−Sml−DAPI−C11
7614−Ctx−T−L−Sml−DAPI−C12
7614−Ctx−T−L−Sml−DAPI−C7
7614−Ctx−T−L−Sml−DAPI−C8
7614−Ctx−T−L−Sml−DAPI−C9
7614−Ctx−T−L−Sml−DAPI−D10
7614−Ctx−T−L−Sml−DAPI−D11
7614−Ctx−T−L−Sml−DAPI−D12
7614−Ctx−T−L−Sml−DAPI−D7
7614−Ctx−T−L−Sml−DAPI−D8
7614−Ctx−T−L−Sml−DAPI−D9
7614−Ctx−T−L−Sml−DAPI−E10
7614−Ctx−T−L−Sml−DAPI−E11
7614−Ctx−T−L−Sml−DAPI−E12
7614−Ctx−T−L−Sml−DAPI−E7
7614−Ctx−T−L−Sml−DAPI−E8
7614−Ctx−T−L−Sml−DAPI−E9
7614−Ctx−T−L−Sml−DAPI−F10
7614−Ctx−T−L−Sml−DAPI−F11
7614−Ctx−T−L−Sml−DAPI−F12
7614−Ctx−T−L−Sml−DAPI−F7
7614−Ctx−T−L−Sml−DAPI−F8
7614−Ctx−T−L−Sml−DAPI−F9
7614−Ctx−T−L−Sml−DAPI−G10
7614−Ctx−T−L−Sml−DAPI−G11
7614−Ctx−T−L−Sml−DAPI−G12
7614−Ctx−T−L−Sml−DAPI−G7
7614−Ctx−T−L−Sml−DAPI−G8
7614−Ctx−T−L−Sml−DAPI−G9
7614−Ctx−T−L−Sml−DAPI−H10
7614−Ctx−T−L−Sml−DAPI−H11
7614−Ctx−T−L−Sml−DAPI−H7
7614−Ctx−T−L−Sml−DAPI−H8
7614−Ctx−T−L−Sml−DAPI−H9
7614−Ctx−T−L−Sml−NeuN−A1
7614−Ctx−T−L−Sml−NeuN−A2
7614−Ctx−T−L−Sml−NeuN−A3
7614−Ctx−T−L−Sml−NeuN−A4
7614−Ctx−T−L−Sml−NeuN−A5
7614−Ctx−T−L−Sml−NeuN−A6
7614−Ctx−T−L−Sml−NeuN−B1
7614−Ctx−T−L−Sml−NeuN−B2
7614−Ctx−T−L−Sml−NeuN−B3
7614−Ctx−T−L−Sml−NeuN−B4
7614−Ctx−T−L−Sml−NeuN−B5
7614−Ctx−T−L−Sml−NeuN−B6
7614−Ctx−T−L−Sml−NeuN−C1
7614−Ctx−T−L−Sml−NeuN−C2
7614−Ctx−T−L−Sml−NeuN−C3
7614−Ctx−T−L−Sml−NeuN−C4
7614−Ctx−T−L−Sml−NeuN−C5
7614−Ctx−T−L−Sml−NeuN−C6
7614−Ctx−T−L−Sml−NeuN−D1
7614−Ctx−T−L−Sml−NeuN−D2
7614−Ctx−T−L−Sml−NeuN−D3
7614−Ctx−T−L−Sml−NeuN−D4
7614−Ctx−T−L−Sml−NeuN−D5
7614−Ctx−T−L−Sml−NeuN−D6
7614−Ctx−T−L−Sml−NeuN−E1
7614−Ctx−T−L−Sml−NeuN−E2
7614−Ctx−T−L−Sml−NeuN−E3
7614−Ctx−T−L−Sml−NeuN−E4
7614−Ctx−T−L−Sml−NeuN−E5
7614−Ctx−T−L−Sml−NeuN−E6
7614−Ctx−T−L−Sml−NeuN−F1
7614−Ctx−T−L−Sml−NeuN−F2
7614−Ctx−T−L−Sml−NeuN−F3
7614−Ctx−T−L−Sml−NeuN−F4
7614−Ctx−T−L−Sml−NeuN−F5
7614−Ctx−T−L−Sml−NeuN−F6
7614−Ctx−T−L−Sml−NeuN−G1
7614−Ctx−T−L−Sml−NeuN−G2
7614−Ctx−T−L−Sml−NeuN−G3
7614−Ctx−T−L−Sml−NeuN−G4
7614−Ctx−T−L−Sml−NeuN−G5
7614−Ctx−T−L−Sml−NeuN−G6
7614−Ctx−T−L−Sml−NeuN−H1
7614−Ctx−T−L−Sml−NeuN−H2
7614−Ctx−T−L−Sml−NeuN−H3
7614−Ctx−T−L−Sml−NeuN−H4
7614−Ctx−T−L−Sml−NeuN−H5
7614−Ctx−T−L−Sml−NeuN−H6
JGG-Cells

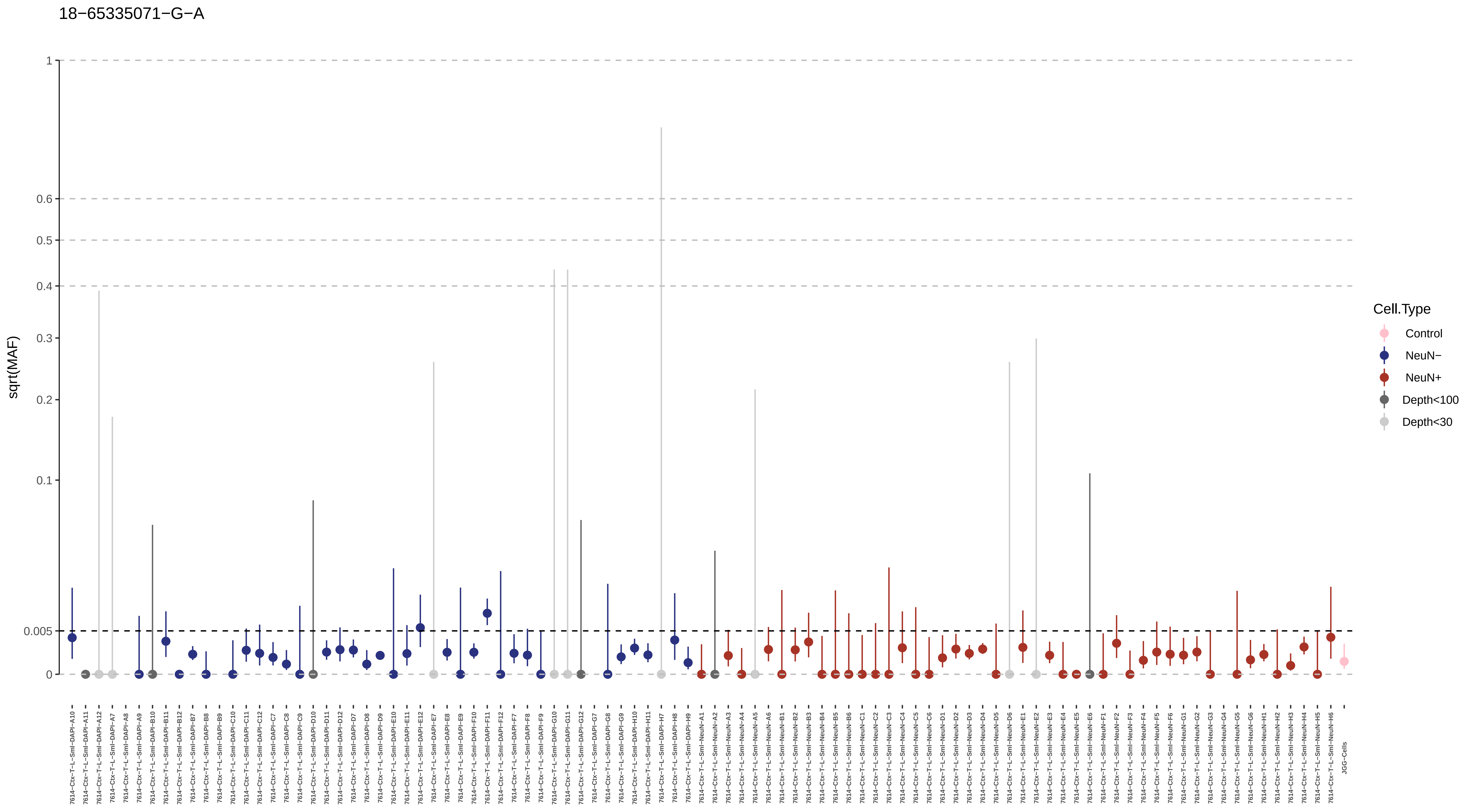
18−65335071−G−A
sqrt(MAF)
1
0.6
0.5
0.4
0.3
0.2
0.1
0.005
0
Cell.Type
Control
NeuN−
NeuN+
Depth<100
Depth<30
JGG-Cells

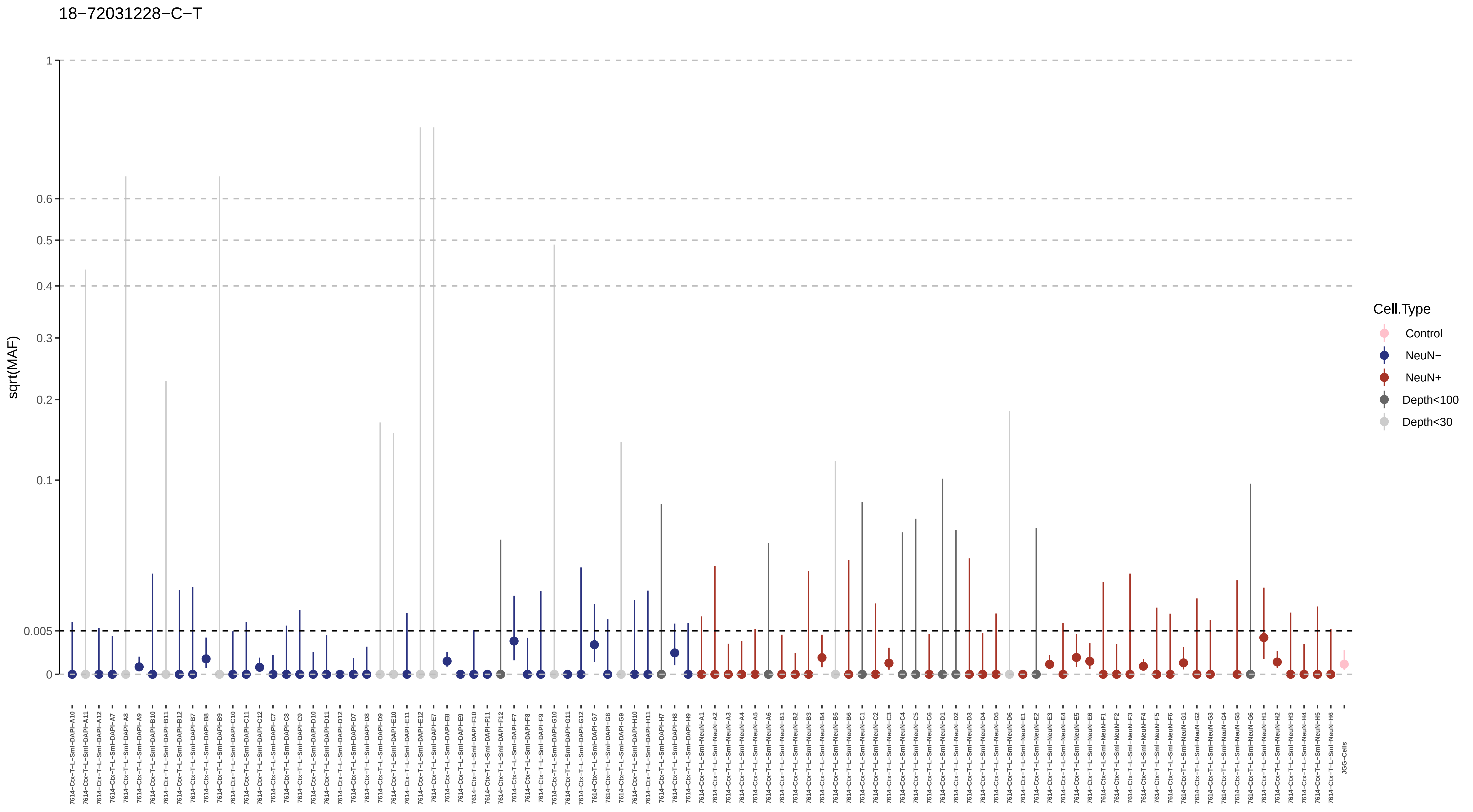

18−72031228−C−T
sqrt(MAF)
1
0.6
0.5
0.4
0.3
0.2
0.1
0.005
0
Cell.Type
Control
NeuN−
NeuN+
Depth<100
Depth<30
JGG-Cells

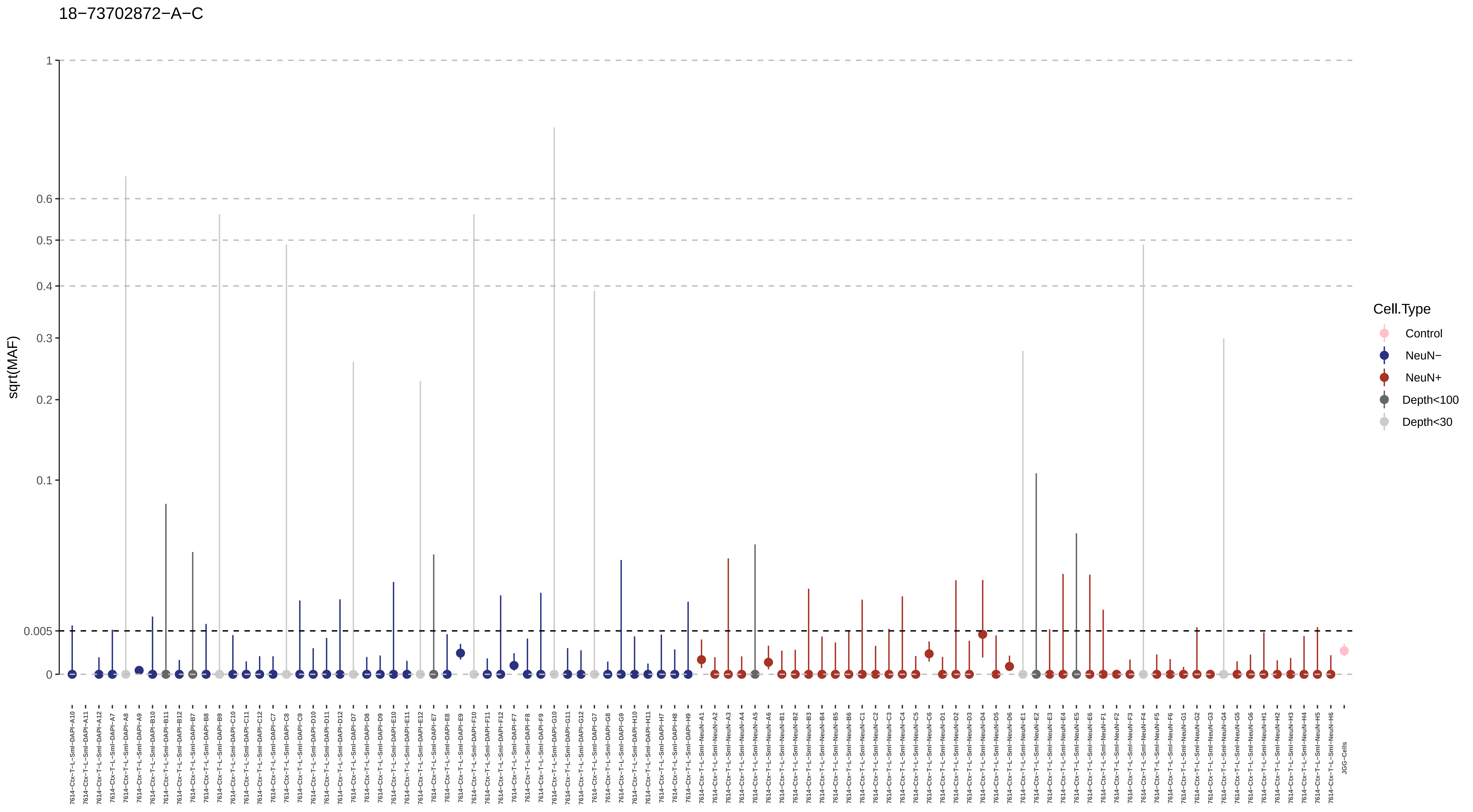
18−73702872−A−C
sqrt(MAF)
1
0.6
0.5
0.4
0.3
0.2
0.1
0.005
0
Cell.Type
Control
NeuN−
NeuN+
Depth<100
Depth<30
7614−Ctx−T−L−Sml−DAPI−A10
7614−Ctx−T−L−Sml−DAPI−A11
7614−Ctx−T−L−Sml−DAPI−A12
7614−Ctx−T−L−Sml−DAPI−A7
7614−Ctx−T−L−Sml−DAPI−A8
7614−Ctx−T−L−Sml−DAPI−A9
7614−Ctx−T−L−Sml−DAPI−B10
7614−Ctx−T−L−Sml−DAPI−B11
7614−Ctx−T−L−Sml−DAPI−B12
7614−Ctx−T−L−Sml−DAPI−B7
7614−Ctx−T−L−Sml−DAPI−B8
7614−Ctx−T−L−Sml−DAPI−B9
7614−Ctx−T−L−Sml−DAPI−C10
7614−Ctx−T−L−Sml−DAPI−C11
7614−Ctx−T−L−Sml−DAPI−C12
7614−Ctx−T−L−Sml−DAPI−C7
7614−Ctx−T−L−Sml−DAPI−C8
7614−Ctx−T−L−Sml−DAPI−C9
7614−Ctx−T−L−Sml−DAPI−D10
7614−Ctx−T−L−Sml−DAPI−D11
7614−Ctx−T−L−Sml−DAPI−D12
7614−Ctx−T−L−Sml−DAPI−D7
7614−Ctx−T−L−Sml−DAPI−D8
7614−Ctx−T−L−Sml−DAPI−D9
7614−Ctx−T−L−Sml−DAPI−E10
7614−Ctx−T−L−Sml−DAPI−E11
7614−Ctx−T−L−Sml−DAPI−E12
7614−Ctx−T−L−Sml−DAPI−E7
7614−Ctx−T−L−Sml−DAPI−E8
7614−Ctx−T−L−Sml−DAPI−E9
7614−Ctx−T−L−Sml−DAPI−F10
7614−Ctx−T−L−Sml−DAPI−F11
7614−Ctx−T−L−Sml−DAPI−F12
7614−Ctx−T−L−Sml−DAPI−F7
7614−Ctx−T−L−Sml−DAPI−F8
7614−Ctx−T−L−Sml−DAPI−F9
7614−Ctx−T−L−Sml−DAPI−G10
7614−Ctx−T−L−Sml−DAPI−G11
7614−Ctx−T−L−Sml−DAPI−G12
7614−Ctx−T−L−Sml−DAPI−G7
7614−Ctx−T−L−Sml−DAPI−G8
7614−Ctx−T−L−Sml−DAPI−G9
7614−Ctx−T−L−Sml−DAPI−H10
7614−Ctx−T−L−Sml−DAPI−H11
7614−Ctx−T−L−Sml−DAPI−H7
7614−Ctx−T−L−Sml−DAPI−H8
7614−Ctx−T−L−Sml−DAPI−H9
7614−Ctx−T−L−Sml−NeuN−A1
7614−Ctx−T−L−Sml−NeuN−A2
7614−Ctx−T−L−Sml−NeuN−A3
7614−Ctx−T−L−Sml−NeuN−A4
7614−Ctx−T−L−Sml−NeuN−A5
7614−Ctx−T−L−Sml−NeuN−A6
7614−Ctx−T−L−Sml−NeuN−B1
7614−Ctx−T−L−Sml−NeuN−B2
7614−Ctx−T−L−Sml−NeuN−B3
7614−Ctx−T−L−Sml−NeuN−B4
7614−Ctx−T−L−Sml−NeuN−B5
7614−Ctx−T−L−Sml−NeuN−B6
7614−Ctx−T−L−Sml−NeuN−C1
7614−Ctx−T−L−Sml−NeuN−C2
7614−Ctx−T−L−Sml−NeuN−C3
7614−Ctx−T−L−Sml−NeuN−C4
7614−Ctx−T−L−Sml−NeuN−C5
7614−Ctx−T−L−Sml−NeuN−C6
7614−Ctx−T−L−Sml−NeuN−D1
7614−Ctx−T−L−Sml−NeuN−D2
7614−Ctx−T−L−Sml−NeuN−D3
7614−Ctx−T−L−Sml−NeuN−D4
7614−Ctx−T−L−Sml−NeuN−D5
7614−Ctx−T−L−Sml−NeuN−D6
7614−Ctx−T−L−Sml−NeuN−E1
7614−Ctx−T−L−Sml−NeuN−E2
7614−Ctx−T−L−Sml−NeuN−E3
7614−Ctx−T−L−Sml−NeuN−E4
7614−Ctx−T−L−Sml−NeuN−E5
7614−Ctx−T−L−Sml−NeuN−E6
7614−Ctx−T−L−Sml−NeuN−F1
7614−Ctx−T−L−Sml−NeuN−F2
7614−Ctx−T−L−Sml−NeuN−F3
7614−Ctx−T−L−Sml−NeuN−F4
7614−Ctx−T−L−Sml−NeuN−F5
7614−Ctx−T−L−Sml−NeuN−F6
7614−Ctx−T−L−Sml−NeuN−G1
7614−Ctx−T−L−Sml−NeuN−G2
7614−Ctx−T−L−Sml−NeuN−G3
7614−Ctx−T−L−Sml−NeuN−G4
7614−Ctx−T−L−Sml−NeuN−G5
7614−Ctx−T−L−Sml−NeuN−G6
7614−Ctx−T−L−Sml−NeuN−H1
7614−Ctx−T−L−Sml−NeuN−H2
7614−Ctx−T−L−Sml−NeuN−H3
7614−Ctx−T−L−Sml−NeuN−H4
7614−Ctx−T−L−Sml−NeuN−H5
7614−Ctx−T−L−Sml−NeuN−H6
JGG-Cells

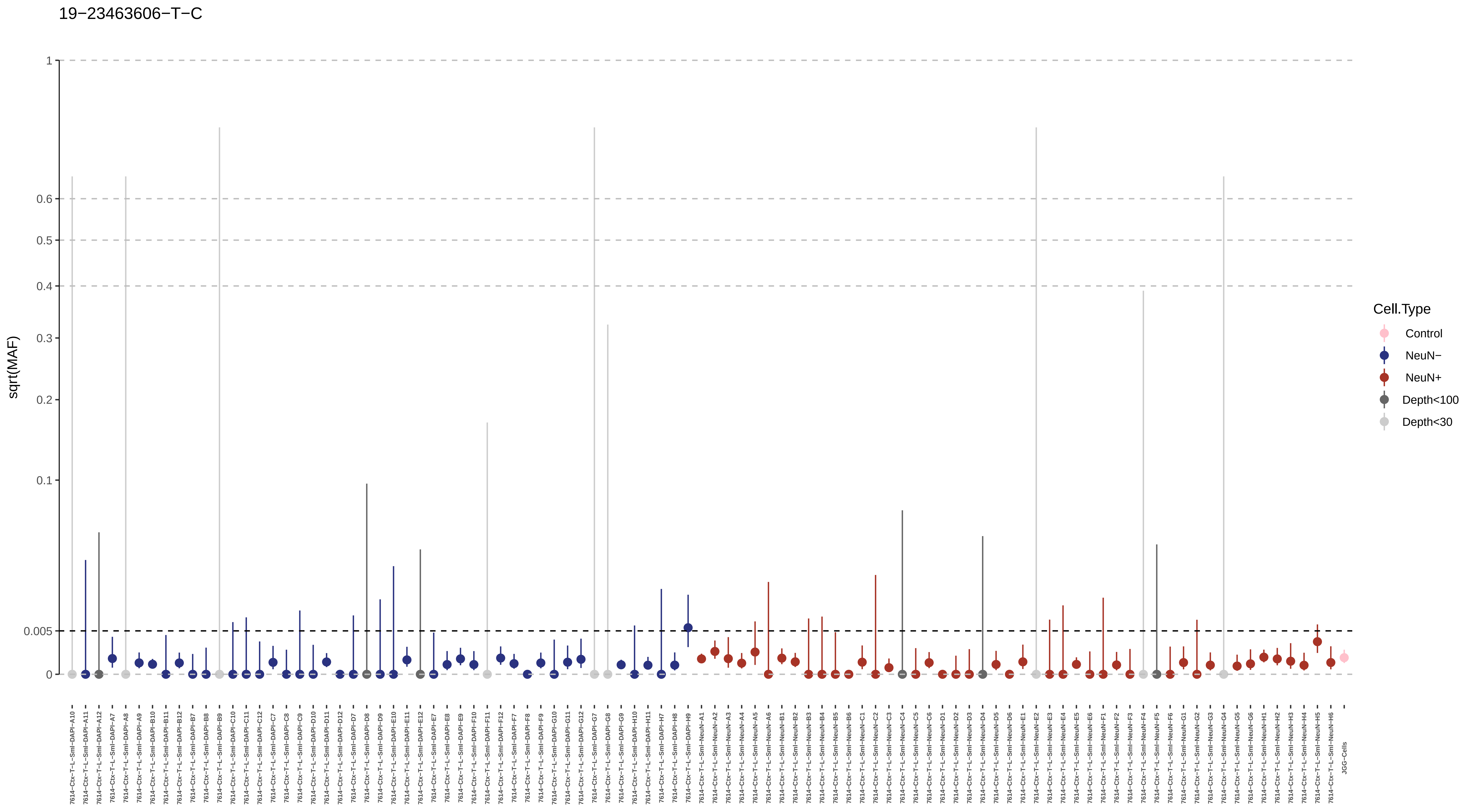

19−23463606−T−C
sqrt(MAF)
1
0.6
0.5
0.4
0.3
0.2
0.1
0.005
0
Cell.Type
Control
NeuN−
NeuN+
Depth<100
Depth<30

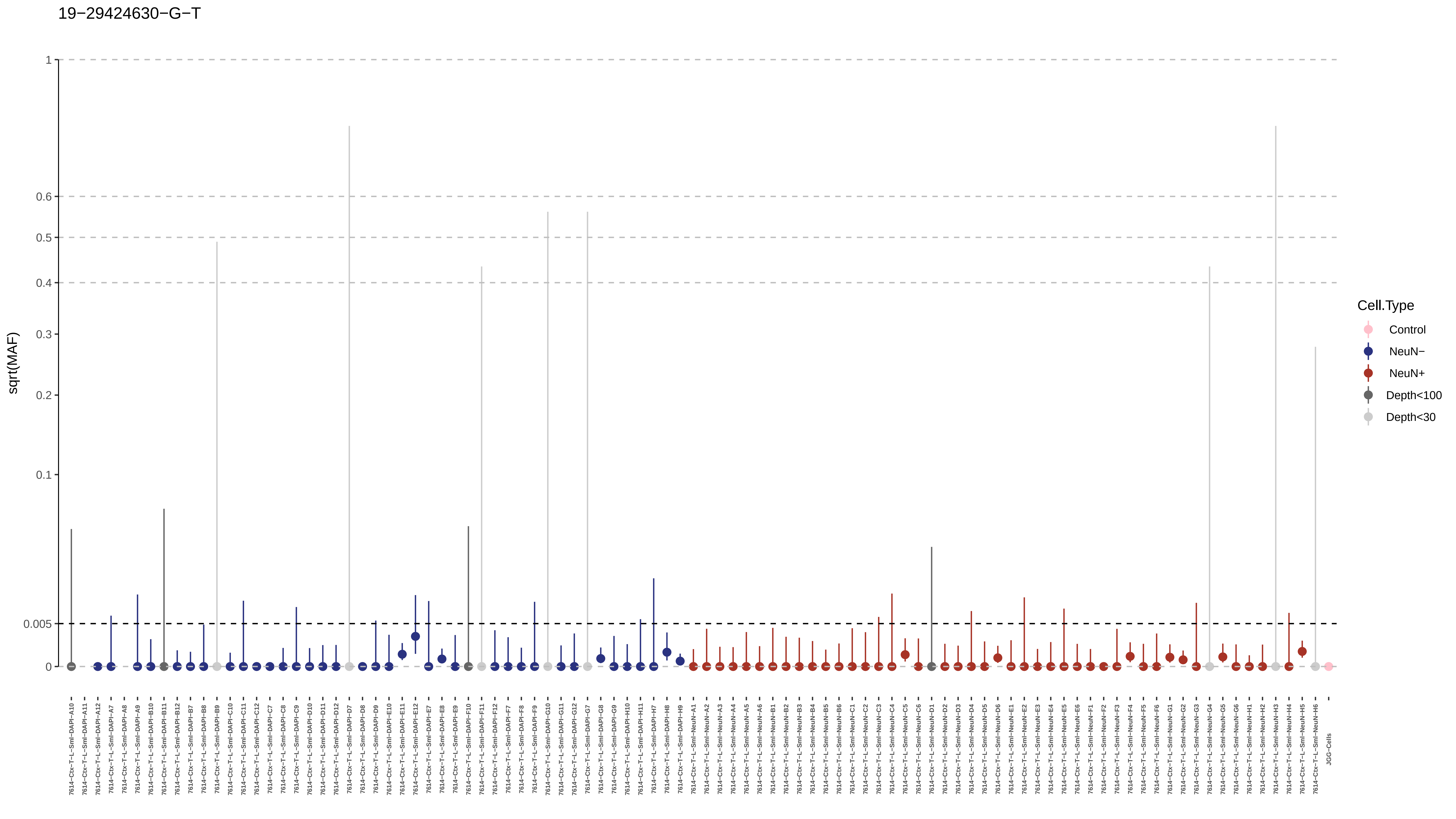
19−29424630−G−T
sqrt(MAF)
1
0.6
0.5
0.4
0.3
0.2
0.1
0.005
0
Cell.Type
Control
NeuN−
NeuN+
Depth<100
Depth<30
JGG-Cells

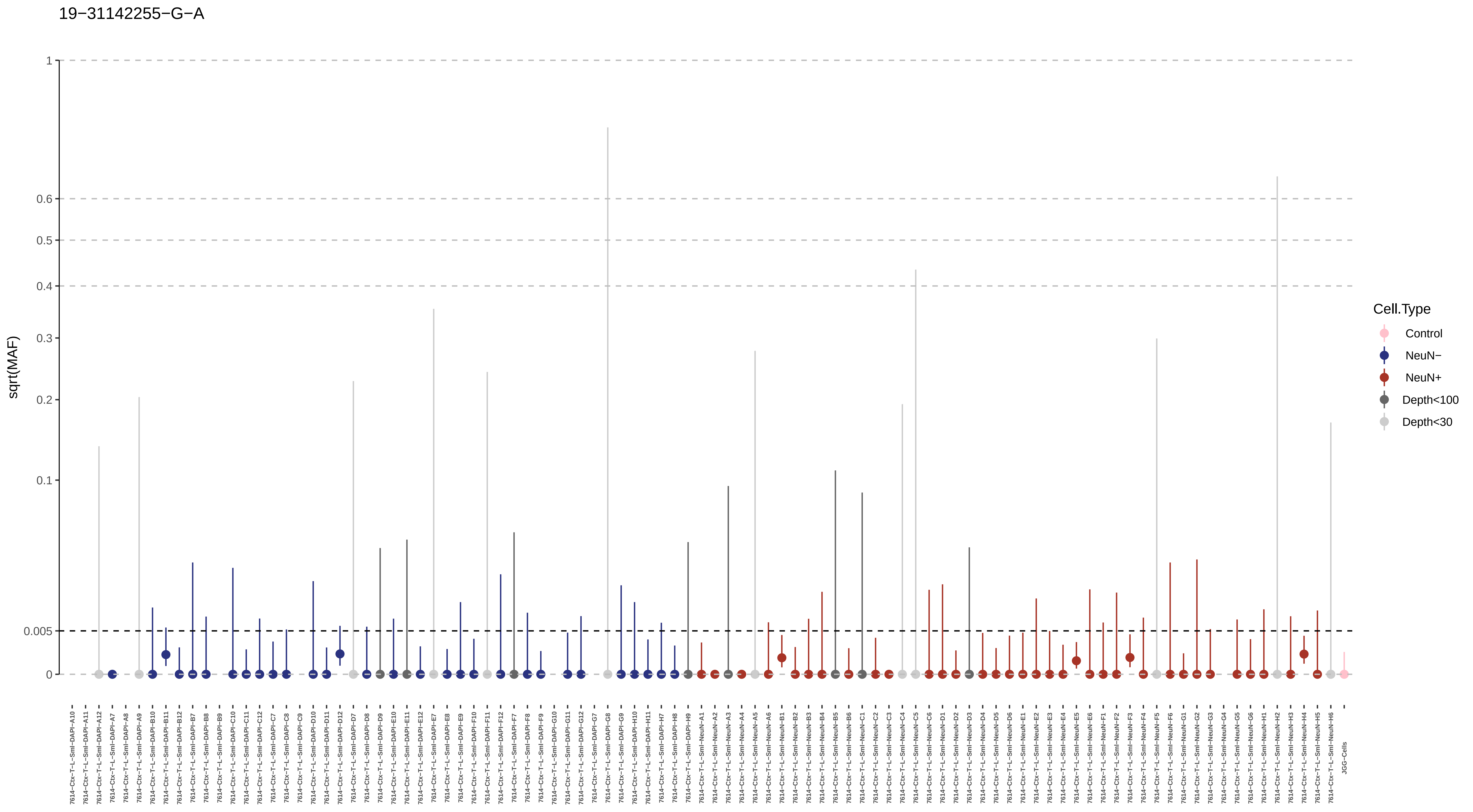

19−31142255−G−A
sqrt(MAF)
1
0.6
0.5
0.4
0.3
0.2
0.1
0.005
0
Cell.Type
Control
NeuN−
NeuN+
Depth<100
Depth<30

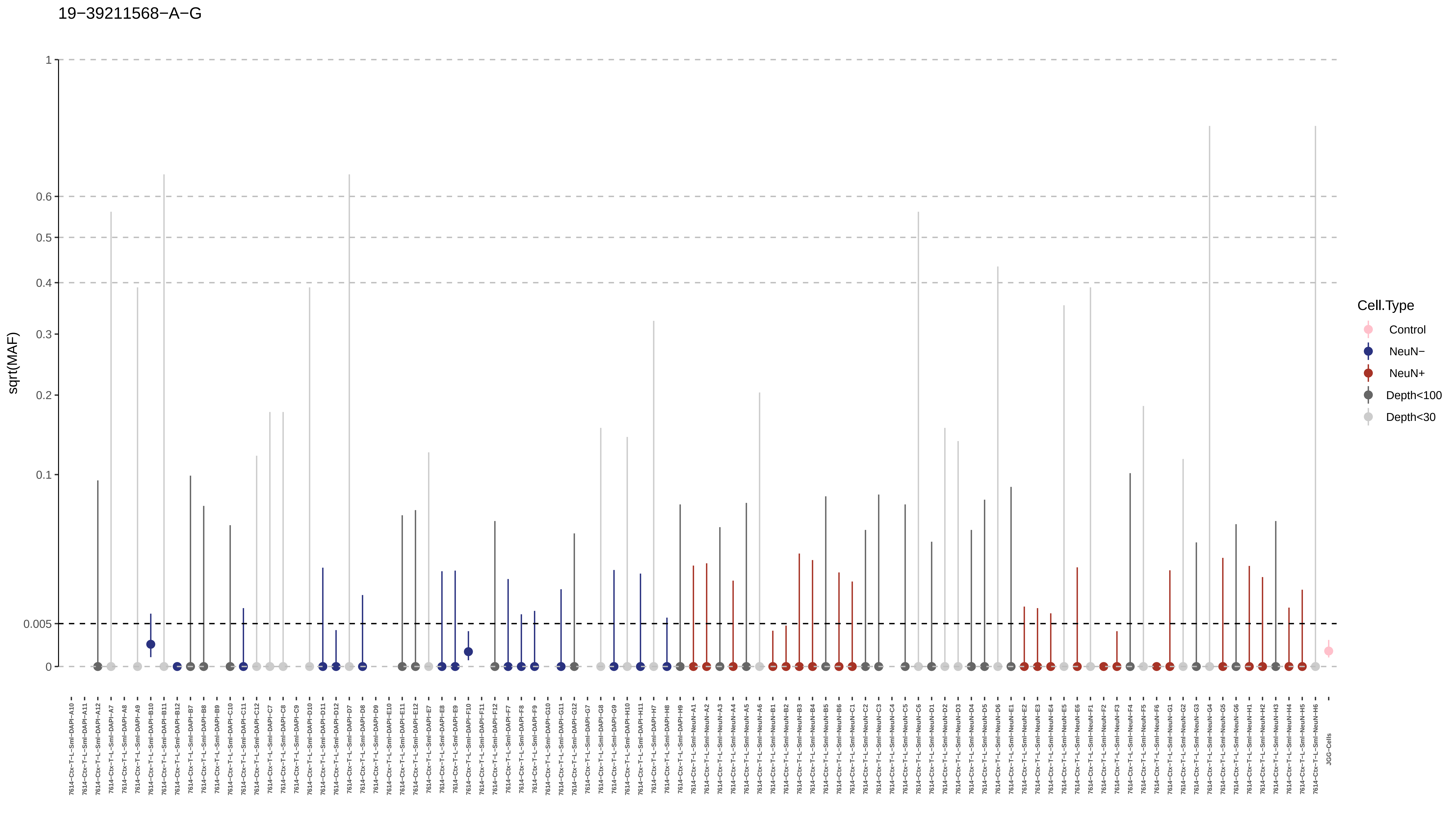

19−39211568−A−G
sqrt(MAF)
1
0.6
0.5
0.4
0.3
0.2
0.1
0.005
0
Cell.Type
Control
NeuN−
NeuN+
Depth<100
Depth<30
JGG-Cells

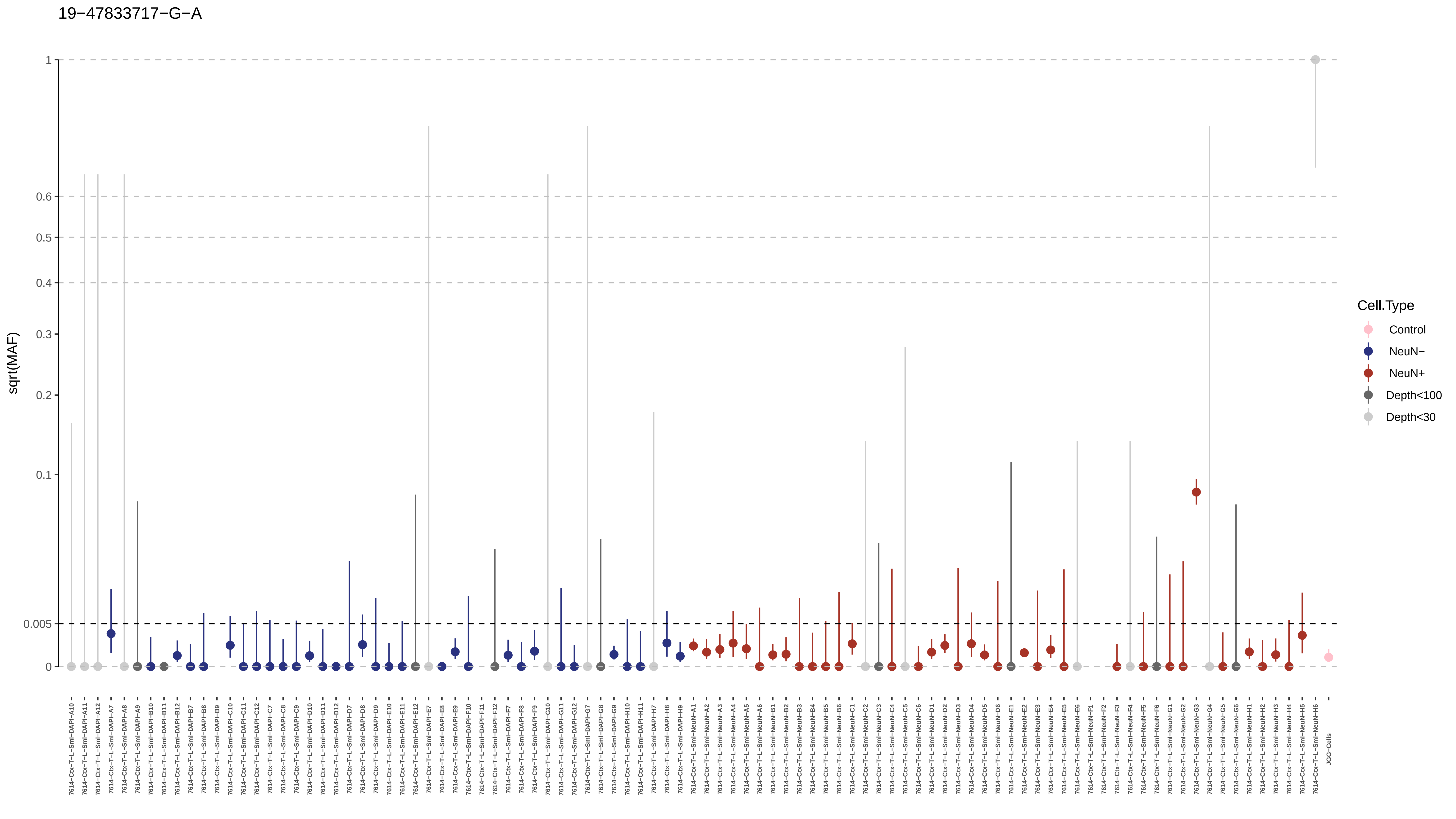
19−47833717−G−A
sqrt(MAF)
1
0.6
0.5
0.4
0.3
0.2
0.1
0.005
0
Cell.Type
Control
NeuN−
NeuN+
Depth<100
Depth<30

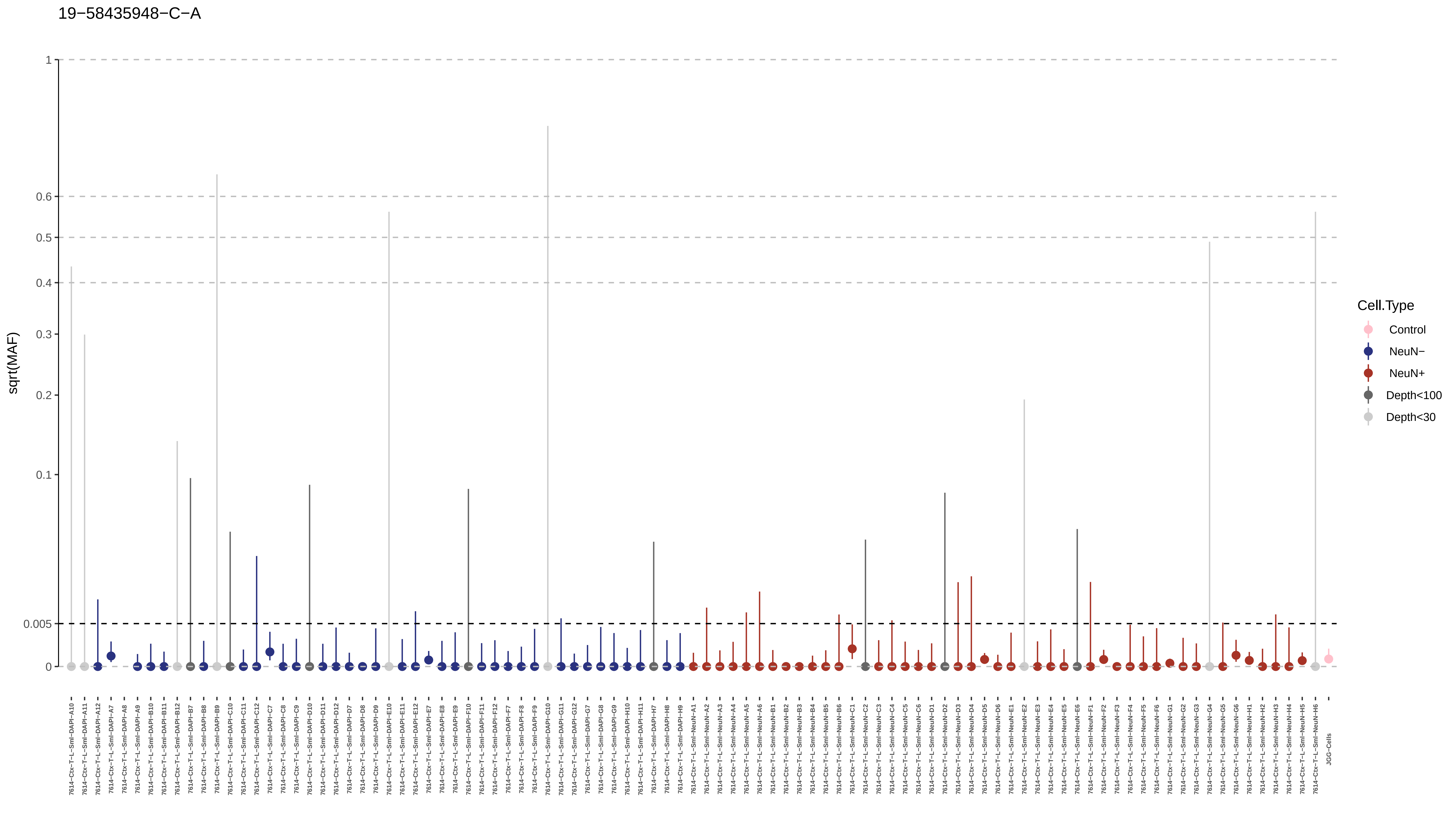
19−58435948−C−A
sqrt(MAF)
1
0.6
0.5
0.4
0.3
0.2
0.1
0.005
0
Cell.Type
Control
NeuN−
NeuN+
Depth<100
Depth<30
JGG-Cells

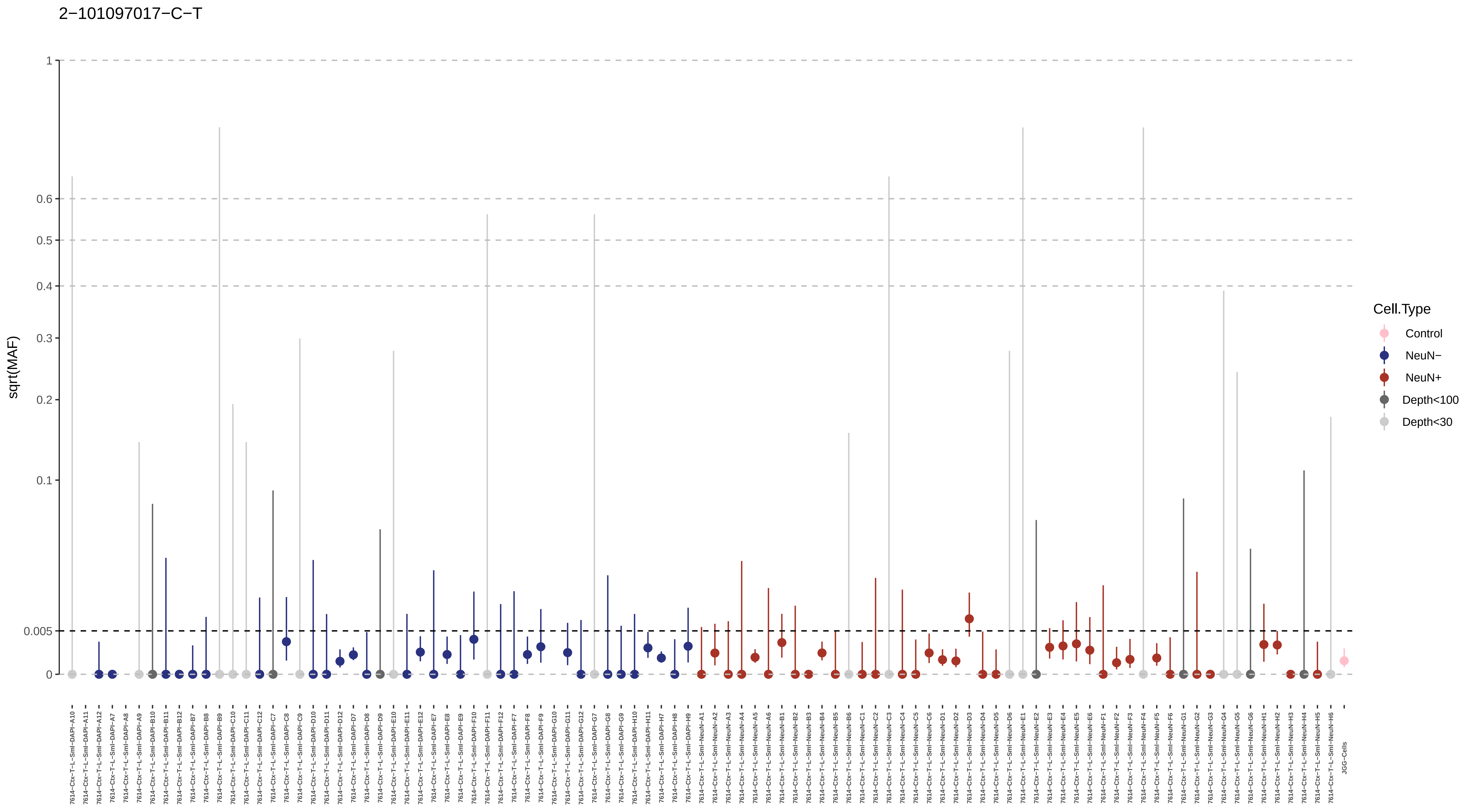

2−101097017−C−T
sqrt(MAF)
1
0.6
0.5
0.4
0.3
0.2
0.1
0.005
0
Cell.Type
Control
NeuN−
NeuN+
Depth<100
Depth<30
7614−Ctx−T−L−Sml−DAPI−A10
7614−Ctx−T−L−Sml−DAPI−A11
7614−Ctx−T−L−Sml−DAPI−A12
7614−Ctx−T−L−Sml−DAPI−A7
7614−Ctx−T−L−Sml−DAPI−A8
7614−Ctx−T−L−Sml−DAPI−A9
7614−Ctx−T−L−Sml−DAPI−B10
7614−Ctx−T−L−Sml−DAPI−B11
7614−Ctx−T−L−Sml−DAPI−B12
7614−Ctx−T−L−Sml−DAPI−B7
7614−Ctx−T−L−Sml−DAPI−B8
7614−Ctx−T−L−Sml−DAPI−B9
7614−Ctx−T−L−Sml−DAPI−C10
7614−Ctx−T−L−Sml−DAPI−C11
7614−Ctx−T−L−Sml−DAPI−C12
7614−Ctx−T−L−Sml−DAPI−C7
7614−Ctx−T−L−Sml−DAPI−C8
7614−Ctx−T−L−Sml−DAPI−C9
7614−Ctx−T−L−Sml−DAPI−D10
7614−Ctx−T−L−Sml−DAPI−D11
7614−Ctx−T−L−Sml−DAPI−D12
7614−Ctx−T−L−Sml−DAPI−D7
7614−Ctx−T−L−Sml−DAPI−D8
7614−Ctx−T−L−Sml−DAPI−D9
7614−Ctx−T−L−Sml−DAPI−E10
7614−Ctx−T−L−Sml−DAPI−E11
7614−Ctx−T−L−Sml−DAPI−E12
7614−Ctx−T−L−Sml−DAPI−E7
7614−Ctx−T−L−Sml−DAPI−E8
7614−Ctx−T−L−Sml−DAPI−E9
7614−Ctx−T−L−Sml−DAPI−F10
7614−Ctx−T−L−Sml−DAPI−F11
7614−Ctx−T−L−Sml−DAPI−F12
7614−Ctx−T−L−Sml−DAPI−F7
7614−Ctx−T−L−Sml−DAPI−F8
7614−Ctx−T−L−Sml−DAPI−F9
7614−Ctx−T−L−Sml−DAPI−G10
7614−Ctx−T−L−Sml−DAPI−G11
7614−Ctx−T−L−Sml−DAPI−G12
7614−Ctx−T−L−Sml−DAPI−G7
7614−Ctx−T−L−Sml−DAPI−G8
7614−Ctx−T−L−Sml−DAPI−G9
7614−Ctx−T−L−Sml−DAPI−H10
7614−Ctx−T−L−Sml−DAPI−H11
7614−Ctx−T−L−Sml−DAPI−H7
7614−Ctx−T−L−Sml−DAPI−H8
7614−Ctx−T−L−Sml−DAPI−H9
7614−Ctx−T−L−Sml−NeuN−A1
7614−Ctx−T−L−Sml−NeuN−A2
7614−Ctx−T−L−Sml−NeuN−A3
7614−Ctx−T−L−Sml−NeuN−A4
7614−Ctx−T−L−Sml−NeuN−A5
7614−Ctx−T−L−Sml−NeuN−A6
7614−Ctx−T−L−Sml−NeuN−B1
7614−Ctx−T−L−Sml−NeuN−B2
7614−Ctx−T−L−Sml−NeuN−B3
7614−Ctx−T−L−Sml−NeuN−B4
7614−Ctx−T−L−Sml−NeuN−B5
7614−Ctx−T−L−Sml−NeuN−B6
7614−Ctx−T−L−Sml−NeuN−C1
7614−Ctx−T−L−Sml−NeuN−C2
7614−Ctx−T−L−Sml−NeuN−C3
7614−Ctx−T−L−Sml−NeuN−C4
7614−Ctx−T−L−Sml−NeuN−C5
7614−Ctx−T−L−Sml−NeuN−C6
7614−Ctx−T−L−Sml−NeuN−D1
7614−Ctx−T−L−Sml−NeuN−D2
7614−Ctx−T−L−Sml−NeuN−D3
7614−Ctx−T−L−Sml−NeuN−D4
7614−Ctx−T−L−Sml−NeuN−D5
7614−Ctx−T−L−Sml−NeuN−D6
7614−Ctx−T−L−Sml−NeuN−E1
7614−Ctx−T−L−Sml−NeuN−E2
7614−Ctx−T−L−Sml−NeuN−E3
7614−Ctx−T−L−Sml−NeuN−E4
7614−Ctx−T−L−Sml−NeuN−E5
7614−Ctx−T−L−Sml−NeuN−E6
7614−Ctx−T−L−Sml−NeuN−F1
7614−Ctx−T−L−Sml−NeuN−F2
7614−Ctx−T−L−Sml−NeuN−F3
7614−Ctx−T−L−Sml−NeuN−F4
7614−Ctx−T−L−Sml−NeuN−F5
7614−Ctx−T−L−Sml−NeuN−F6
7614−Ctx−T−L−Sml−NeuN−G1
7614−Ctx−T−L−Sml−NeuN−G2
7614−Ctx−T−L−Sml−NeuN−G3
7614−Ctx−T−L−Sml−NeuN−G4
7614−Ctx−T−L−Sml−NeuN−G5
7614−Ctx−T−L−Sml−NeuN−G6
7614−Ctx−T−L−Sml−NeuN−H1
7614−Ctx−T−L−Sml−NeuN−H2
7614−Ctx−T−L−Sml−NeuN−H3
7614−Ctx−T−L−Sml−NeuN−H4
7614−Ctx−T−L−Sml−NeuN−H5
7614−Ctx−T−L−Sml−NeuN−H6
JGG-Cells

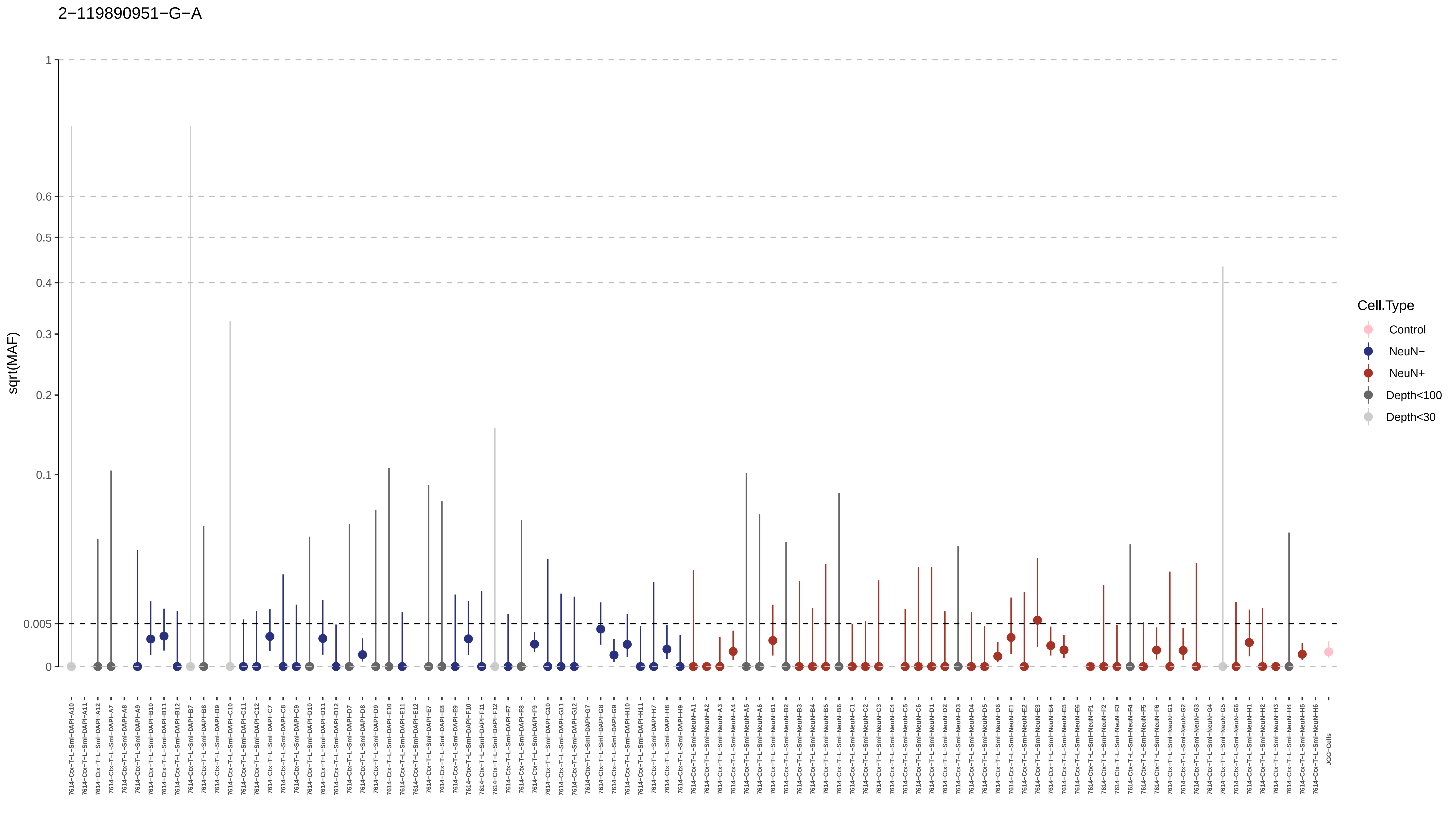

2−119890951−G−A
sqrt(MAF)
1
0.6
0.5
0.4
0.3
0.2
0.1
0.005
0
Cell.Type
Control
NeuN−
NeuN+
Depth<100
Depth<30
JGG-Cells

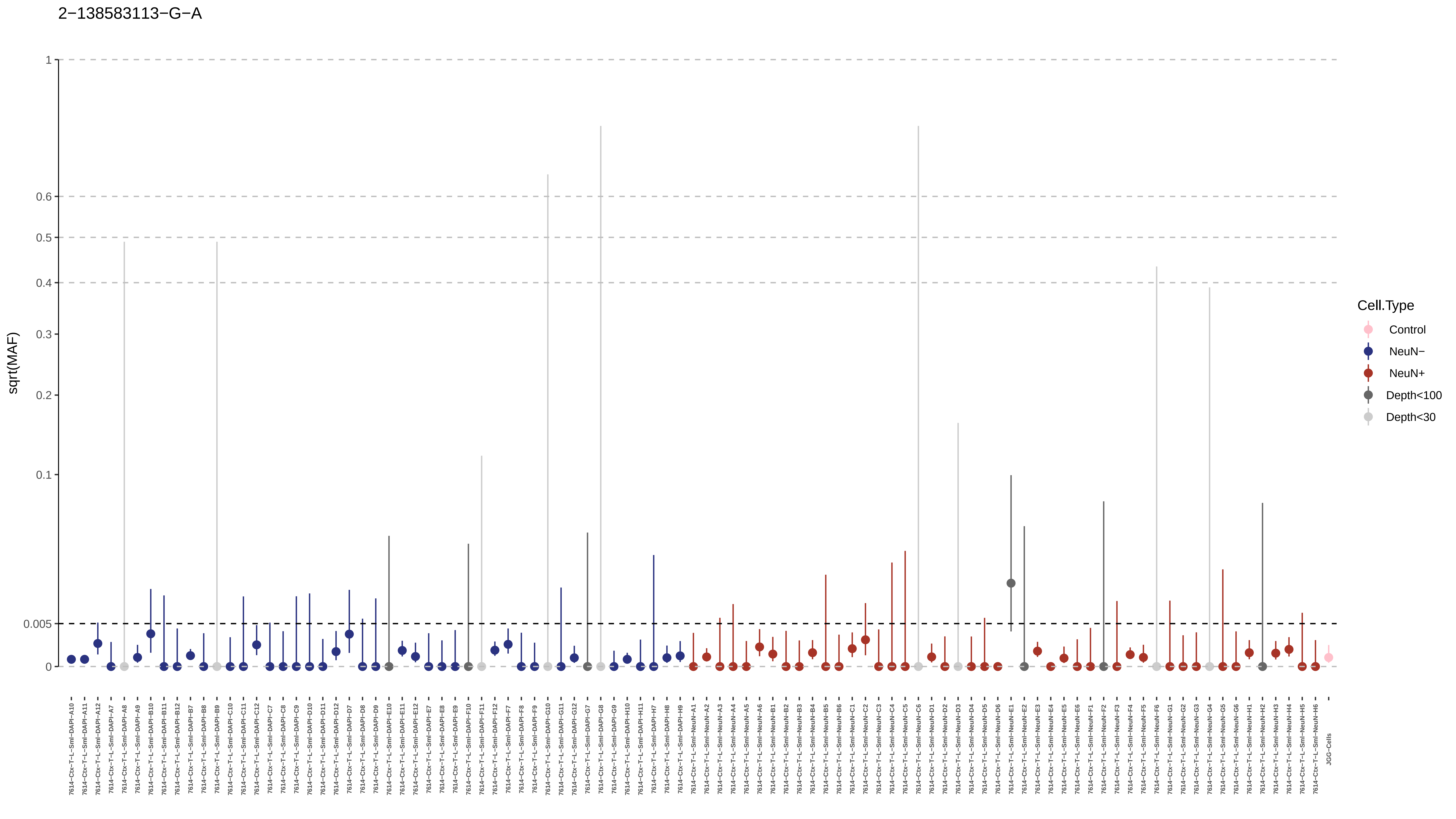
2−138583113−G−A
sqrt(MAF)
1
0.6
0.5
0.4
0.3
0.2
0.1
0.005
0
Cell.Type
Control
NeuN−
NeuN+
Depth<100
Depth<30

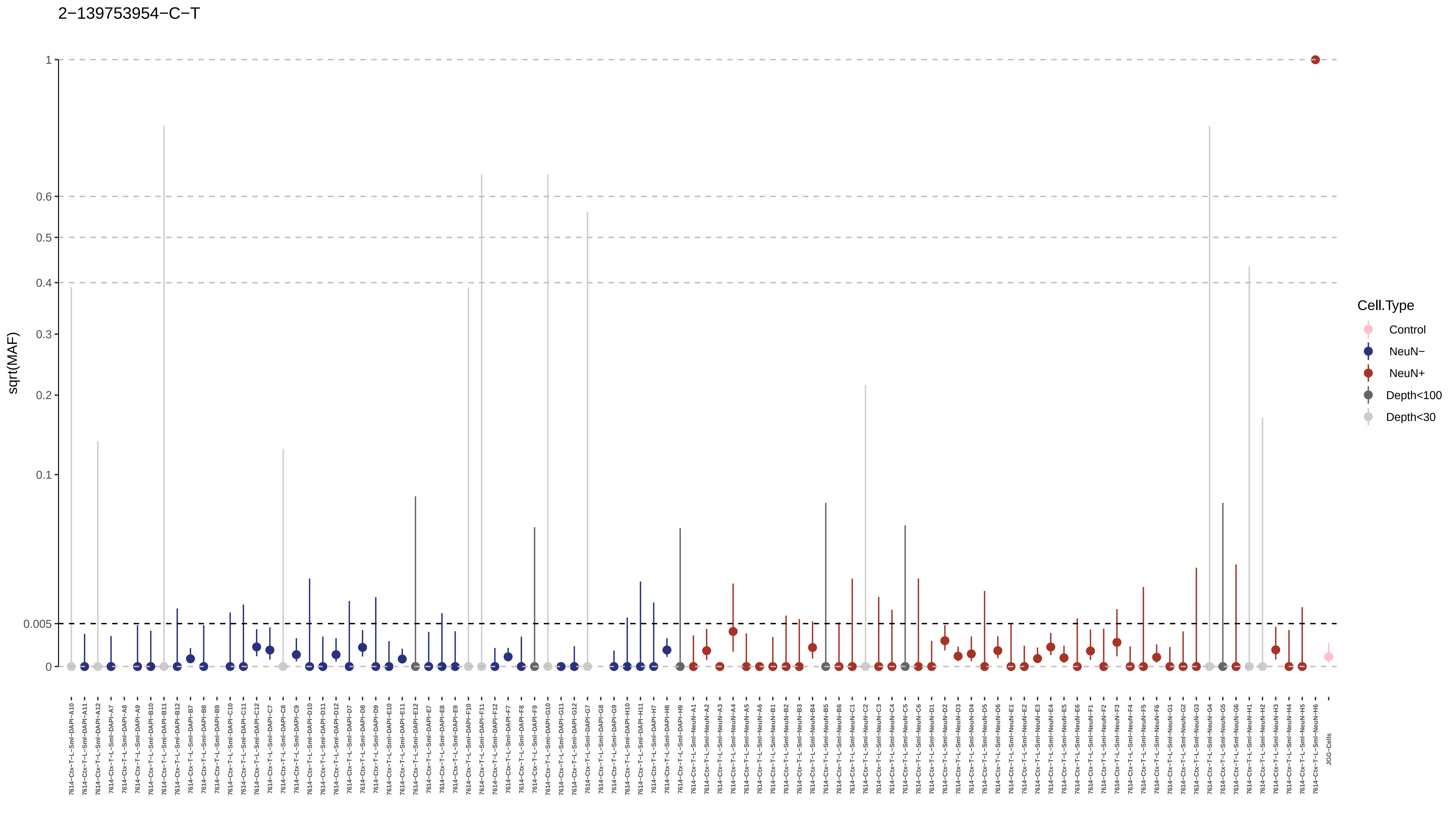

2−139753954−C−T
sqrt(MAF)
1
0.6
0.5
0.4
0.3
0.2
0.1
0.005
0
Cell.Type
Control
NeuN−
NeuN+
Depth<100
Depth<30
JGG-Cells

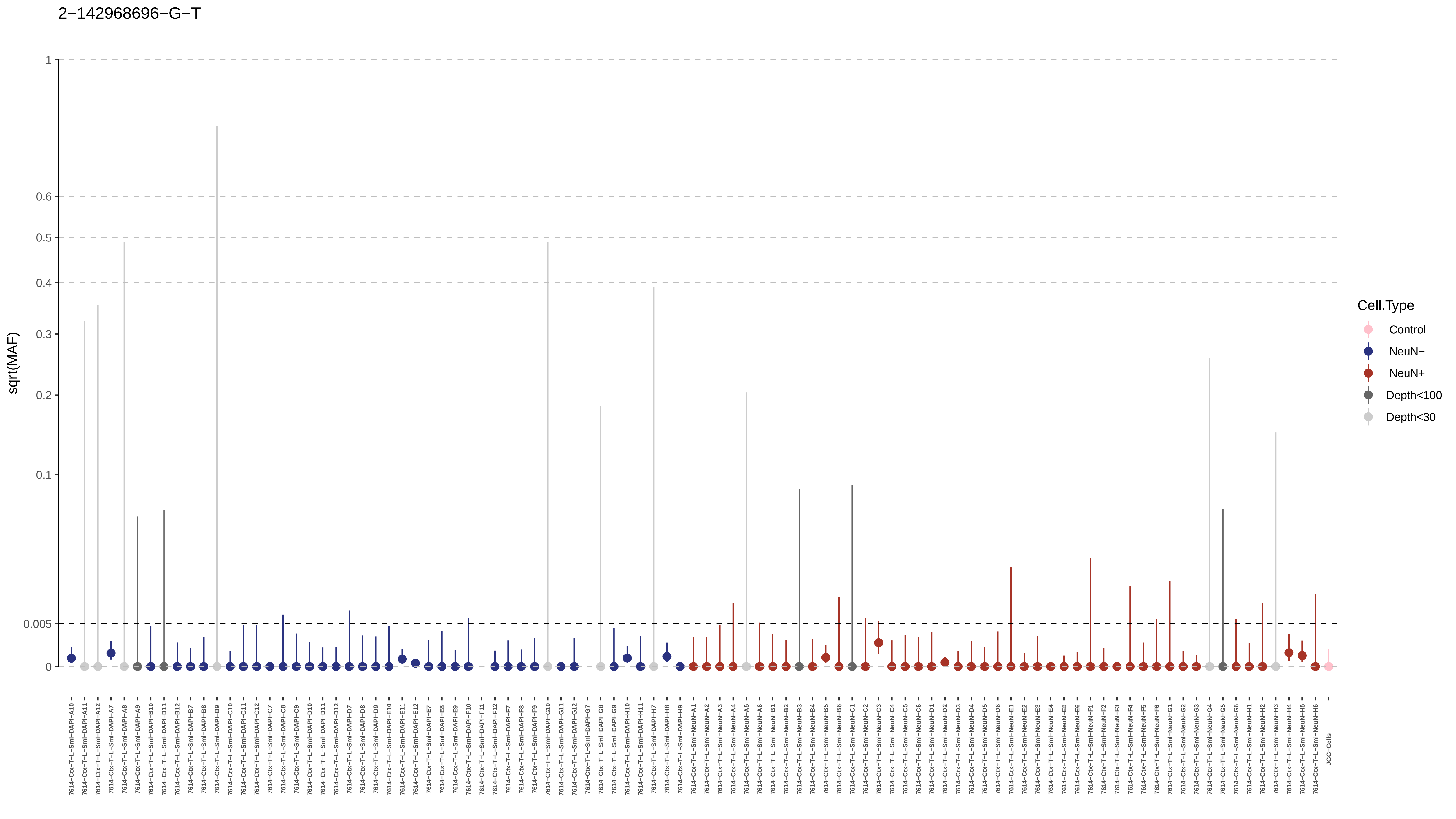
2−142968696−G−T
sqrt(MAF)
1
0.6
0.5
0.4
0.3
0.2
0.1
0.005
0
Cell.Type
Control
NeuN−
NeuN+
Depth<100
Depth<30
7614−Ctx−T−L−Sml−DAPI−A10
7614−Ctx−T−L−Sml−DAPI−A11
7614−Ctx−T−L−Sml−DAPI−A12
7614−Ctx−T−L−Sml−DAPI−A7
7614−Ctx−T−L−Sml−DAPI−A8
7614−Ctx−T−L−Sml−DAPI−A9
7614−Ctx−T−L−Sml−DAPI−B10
7614−Ctx−T−L−Sml−DAPI−B11
7614−Ctx−T−L−Sml−DAPI−B12
7614−Ctx−T−L−Sml−DAPI−B7
7614−Ctx−T−L−Sml−DAPI−B8
7614−Ctx−T−L−Sml−DAPI−B9
7614−Ctx−T−L−Sml−DAPI−C10
7614−Ctx−T−L−Sml−DAPI−C11
7614−Ctx−T−L−Sml−DAPI−C12
7614−Ctx−T−L−Sml−DAPI−C7
7614−Ctx−T−L−Sml−DAPI−C8
7614−Ctx−T−L−Sml−DAPI−C9
7614−Ctx−T−L−Sml−DAPI−D10
7614−Ctx−T−L−Sml−DAPI−D11
7614−Ctx−T−L−Sml−DAPI−D12
7614−Ctx−T−L−Sml−DAPI−D7
7614−Ctx−T−L−Sml−DAPI−D8
7614−Ctx−T−L−Sml−DAPI−D9
7614−Ctx−T−L−Sml−DAPI−E10
7614−Ctx−T−L−Sml−DAPI−E11
7614−Ctx−T−L−Sml−DAPI−E12
7614−Ctx−T−L−Sml−DAPI−E7
7614−Ctx−T−L−Sml−DAPI−E8
7614−Ctx−T−L−Sml−DAPI−E9
7614−Ctx−T−L−Sml−DAPI−F10
7614−Ctx−T−L−Sml−DAPI−F11
7614−Ctx−T−L−Sml−DAPI−F12
7614−Ctx−T−L−Sml−DAPI−F7
7614−Ctx−T−L−Sml−DAPI−F8
7614−Ctx−T−L−Sml−DAPI−F9
7614−Ctx−T−L−Sml−DAPI−G10
7614−Ctx−T−L−Sml−DAPI−G11
7614−Ctx−T−L−Sml−DAPI−G12
7614−Ctx−T−L−Sml−DAPI−G7
7614−Ctx−T−L−Sml−DAPI−G8
7614−Ctx−T−L−Sml−DAPI−G9
7614−Ctx−T−L−Sml−DAPI−H10
7614−Ctx−T−L−Sml−DAPI−H11
7614−Ctx−T−L−Sml−DAPI−H7
7614−Ctx−T−L−Sml−DAPI−H8
7614−Ctx−T−L−Sml−DAPI−H9
7614−Ctx−T−L−Sml−NeuN−A1
7614−Ctx−T−L−Sml−NeuN−A2
7614−Ctx−T−L−Sml−NeuN−A3
7614−Ctx−T−L−Sml−NeuN−A4
7614−Ctx−T−L−Sml−NeuN−A5
7614−Ctx−T−L−Sml−NeuN−A6
7614−Ctx−T−L−Sml−NeuN−B1
7614−Ctx−T−L−Sml−NeuN−B2
7614−Ctx−T−L−Sml−NeuN−B3
7614−Ctx−T−L−Sml−NeuN−B4
7614−Ctx−T−L−Sml−NeuN−B5
7614−Ctx−T−L−Sml−NeuN−B6
7614−Ctx−T−L−Sml−NeuN−C1
7614−Ctx−T−L−Sml−NeuN−C2
7614−Ctx−T−L−Sml−NeuN−C3
7614−Ctx−T−L−Sml−NeuN−C4
7614−Ctx−T−L−Sml−NeuN−C5
7614−Ctx−T−L−Sml−NeuN−C6
7614−Ctx−T−L−Sml−NeuN−D1
7614−Ctx−T−L−Sml−NeuN−D2
7614−Ctx−T−L−Sml−NeuN−D3
7614−Ctx−T−L−Sml−NeuN−D4
7614−Ctx−T−L−Sml−NeuN−D5
7614−Ctx−T−L−Sml−NeuN−D6
7614−Ctx−T−L−Sml−NeuN−E1
7614−Ctx−T−L−Sml−NeuN−E2
7614−Ctx−T−L−Sml−NeuN−E3
7614−Ctx−T−L−Sml−NeuN−E4
7614−Ctx−T−L−Sml−NeuN−E5
7614−Ctx−T−L−Sml−NeuN−E6
7614−Ctx−T−L−Sml−NeuN−F1
7614−Ctx−T−L−Sml−NeuN−F2
7614−Ctx−T−L−Sml−NeuN−F3
7614−Ctx−T−L−Sml−NeuN−F4
7614−Ctx−T−L−Sml−NeuN−F5
7614−Ctx−T−L−Sml−NeuN−F6
7614−Ctx−T−L−Sml−NeuN−G1
7614−Ctx−T−L−Sml−NeuN−G2
7614−Ctx−T−L−Sml−NeuN−G3
7614−Ctx−T−L−Sml−NeuN−G4
7614−Ctx−T−L−Sml−NeuN−G5
7614−Ctx−T−L−Sml−NeuN−G6
7614−Ctx−T−L−Sml−NeuN−H1
7614−Ctx−T−L−Sml−NeuN−H2
7614−Ctx−T−L−Sml−NeuN−H3
7614−Ctx−T−L−Sml−NeuN−H4
7614−Ctx−T−L−Sml−NeuN−H5
7614−Ctx−T−L−Sml−NeuN−H6
JGG−Cells

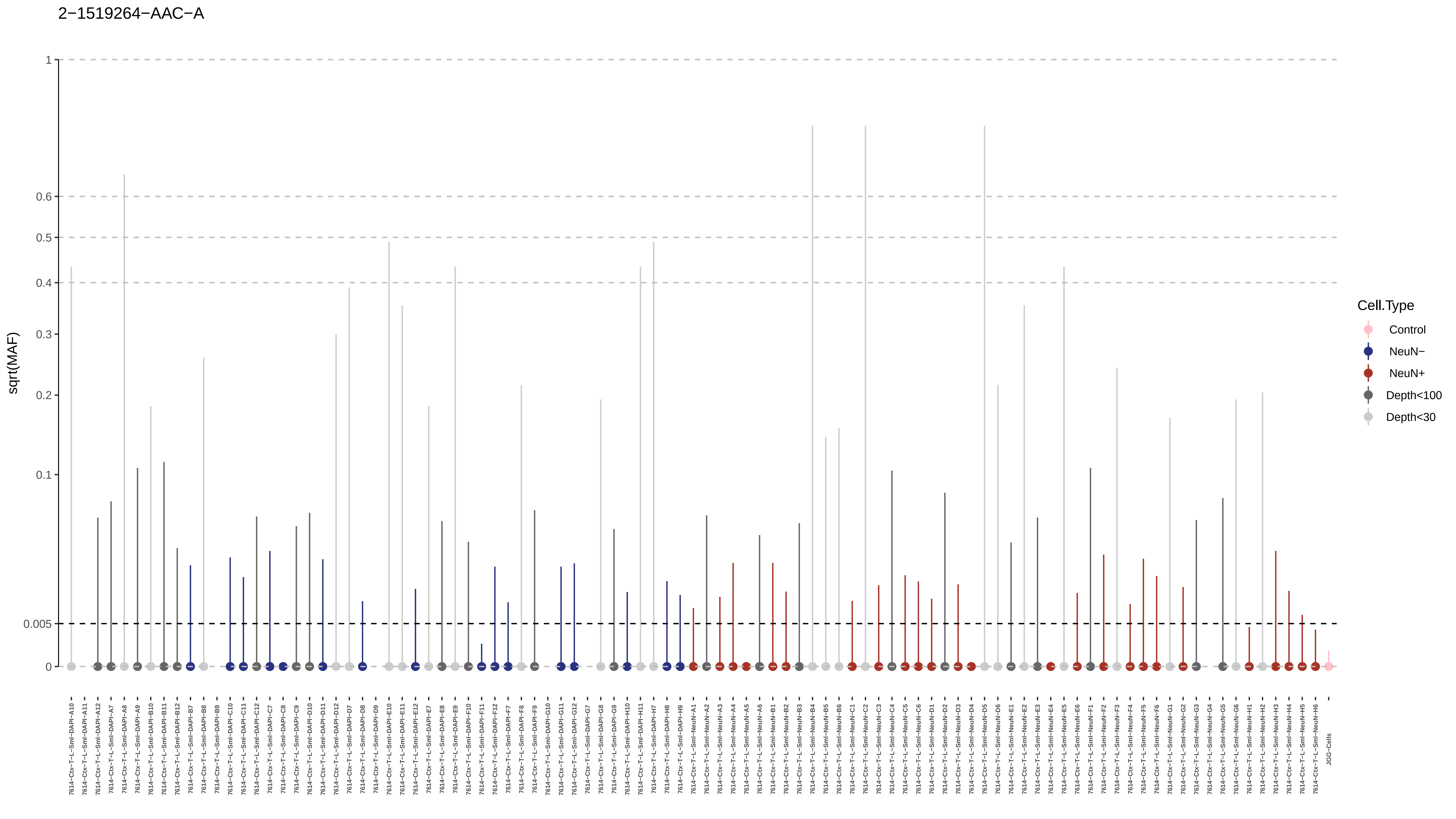
2−1519264−AAC−A
sqrt(MAF)
1
0.6
0.5
0.4
0.3
0.2
0.1
0.005
0
Cell.Type
Control
NeuN−
NeuN+
Depth<100
Depth<30
7614−Ctx−T−L−Sml−DAPI−A10
7614−Ctx−T−L−Sml−DAPI−A11
7614−Ctx−T−L−Sml−DAPI−A12
7614−Ctx−T−L−Sml−DAPI−A7
7614−Ctx−T−L−Sml−DAPI−A8
7614−Ctx−T−L−Sml−DAPI−A9
7614−Ctx−T−L−Sml−DAPI−B10
7614−Ctx−T−L−Sml−DAPI−B11
7614−Ctx−T−L−Sml−DAPI−B12
7614−Ctx−T−L−Sml−DAPI−B7
7614−Ctx−T−L−Sml−DAPI−B8
7614−Ctx−T−L−Sml−DAPI−B9
7614−Ctx−T−L−Sml−DAPI−C10
7614−Ctx−T−L−Sml−DAPI−C11
7614−Ctx−T−L−Sml−DAPI−C12
7614−Ctx−T−L−Sml−DAPI−C7
7614−Ctx−T−L−Sml−DAPI−C8
7614−Ctx−T−L−Sml−DAPI−C9
7614−Ctx−T−L−Sml−DAPI−D10
7614−Ctx−T−L−Sml−DAPI−D11
7614−Ctx−T−L−Sml−DAPI−D12
7614−Ctx−T−L−Sml−DAPI−D7
7614−Ctx−T−L−Sml−DAPI−D8
7614−Ctx−T−L−Sml−DAPI−D9
7614−Ctx−T−L−Sml−DAPI−E10
7614−Ctx−T−L−Sml−DAPI−E11
7614−Ctx−T−L−Sml−DAPI−E12
7614−Ctx−T−L−Sml−DAPI−E7
7614−Ctx−T−L−Sml−DAPI−E8
7614−Ctx−T−L−Sml−DAPI−E9
7614−Ctx−T−L−Sml−DAPI−F10
7614−Ctx−T−L−Sml−DAPI−F11
7614−Ctx−T−L−Sml−DAPI−F12
7614−Ctx−T−L−Sml−DAPI−F7
7614−Ctx−T−L−Sml−DAPI−F8
7614−Ctx−T−L−Sml−DAPI−F9
7614−Ctx−T−L−Sml−DAPI−G10
7614−Ctx−T−L−Sml−DAPI−G11
7614−Ctx−T−L−Sml−DAPI−G12
7614−Ctx−T−L−Sml−DAPI−G7
7614−Ctx−T−L−Sml−DAPI−G8
7614−Ctx−T−L−Sml−DAPI−G9
7614−Ctx−T−L−Sml−DAPI−H10
7614−Ctx−T−L−Sml−DAPI−H11
7614−Ctx−T−L−Sml−DAPI−H7
7614−Ctx−T−L−Sml−DAPI−H8
7614−Ctx−T−L−Sml−DAPI−H9
7614−Ctx−T−L−Sml−NeuN−A1
7614−Ctx−T−L−Sml−NeuN−A2
7614−Ctx−T−L−Sml−NeuN−A3
7614−Ctx−T−L−Sml−NeuN−A4
7614−Ctx−T−L−Sml−NeuN−A5
7614−Ctx−T−L−Sml−NeuN−A6
7614−Ctx−T−L−Sml−NeuN−B1
7614−Ctx−T−L−Sml−NeuN−B2
7614−Ctx−T−L−Sml−NeuN−B3
7614−Ctx−T−L−Sml−NeuN−B4
7614−Ctx−T−L−Sml−NeuN−B5
7614−Ctx−T−L−Sml−NeuN−B6
7614−Ctx−T−L−Sml−NeuN−C1
7614−Ctx−T−L−Sml−NeuN−C2
7614−Ctx−T−L−Sml−NeuN−C3
7614−Ctx−T−L−Sml−NeuN−C4
7614−Ctx−T−L−Sml−NeuN−C5
7614−Ctx−T−L−Sml−NeuN−C6
7614−Ctx−T−L−Sml−NeuN−D1
7614−Ctx−T−L−Sml−NeuN−D2
7614−Ctx−T−L−Sml−NeuN−D3
7614−Ctx−T−L−Sml−NeuN−D4
7614−Ctx−T−L−Sml−NeuN−D5
7614−Ctx−T−L−Sml−NeuN−D6
7614−Ctx−T−L−Sml−NeuN−E1
7614−Ctx−T−L−Sml−NeuN−E2
7614−Ctx−T−L−Sml−NeuN−E3
7614−Ctx−T−L−Sml−NeuN−E4
7614−Ctx−T−L−Sml−NeuN−E5
7614−Ctx−T−L−Sml−NeuN−E6
7614−Ctx−T−L−Sml−NeuN−F1
7614−Ctx−T−L−Sml−NeuN−F2
7614−Ctx−T−L−Sml−NeuN−F3
7614−Ctx−T−L−Sml−NeuN−F4
7614−Ctx−T−L−Sml−NeuN−F5
7614−Ctx−T−L−Sml−NeuN−F6
7614−Ctx−T−L−Sml−NeuN−G1
7614−Ctx−T−L−Sml−NeuN−G2
7614−Ctx−T−L−Sml−NeuN−G3
7614−Ctx−T−L−Sml−NeuN−G4
7614−Ctx−T−L−Sml−NeuN−G5
7614−Ctx−T−L−Sml−NeuN−G6
7614−Ctx−T−L−Sml−NeuN−H1
7614−Ctx−T−L−Sml−NeuN−H2
7614−Ctx−T−L−Sml−NeuN−H3
7614−Ctx−T−L−Sml−NeuN−H4
7614−Ctx−T−L−Sml−NeuN−H5
7614−Ctx−T−L−Sml−NeuN−H6
JGG−Cells

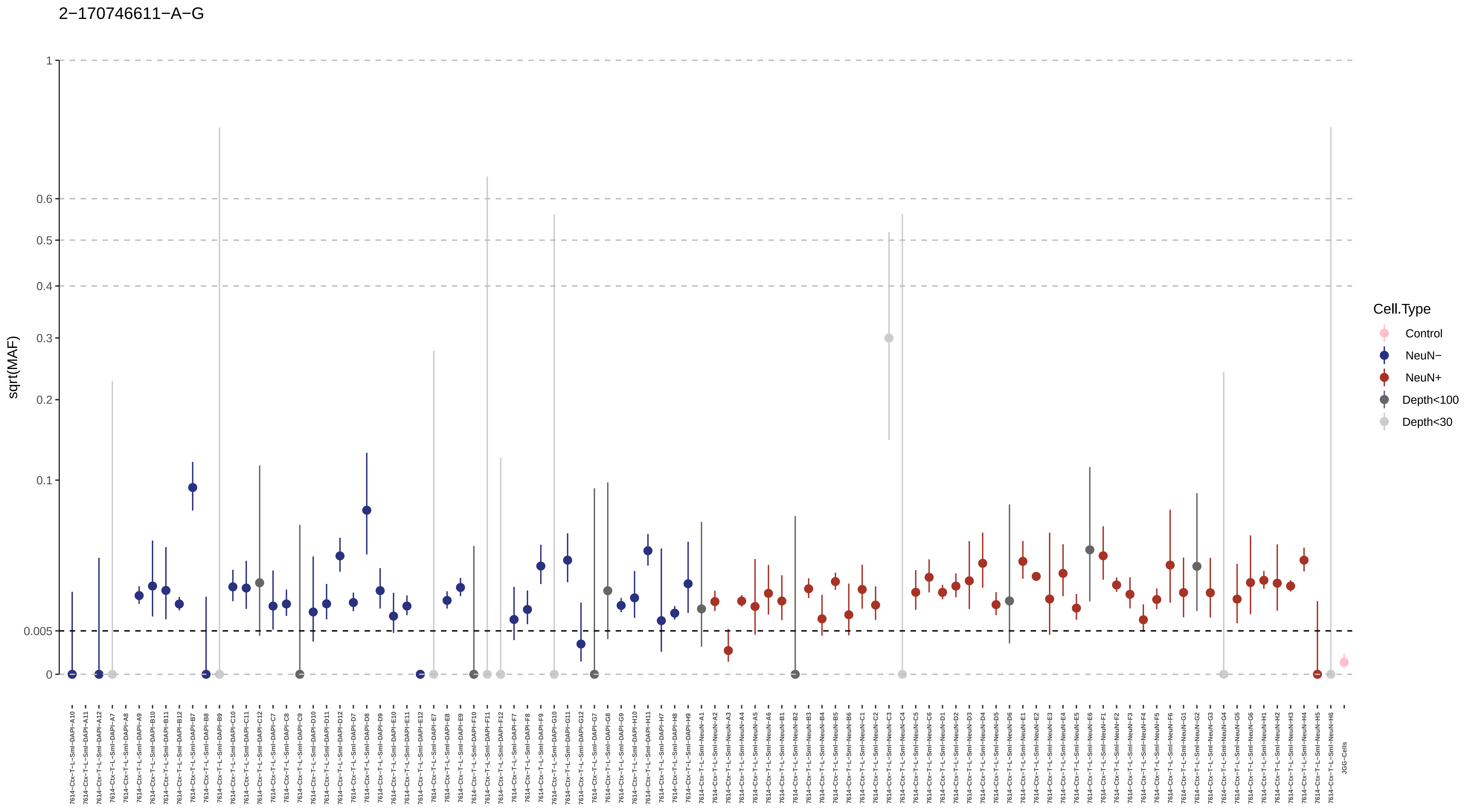
2−170746611−A−G
sqrt(MAF)
1
0.6
0.5
0.4
0.3
0.2
0.1
0.005
0
Cell.Type
Control
NeuN−
NeuN+
Depth<100
Depth<30
7614−Ctx−T−L−Sml−DAPI−A10
7614−Ctx−T−L−Sml−DAPI−A11
7614−Ctx−T−L−Sml−DAPI−A12
7614−Ctx−T−L−Sml−DAPI−A7
7614−Ctx−T−L−Sml−DAPI−A8
7614−Ctx−T−L−Sml−DAPI−A9
7614−Ctx−T−L−Sml−DAPI−B10
7614−Ctx−T−L−Sml−DAPI−B11
7614−Ctx−T−L−Sml−DAPI−B12
7614−Ctx−T−L−Sml−DAPI−B7
7614−Ctx−T−L−Sml−DAPI−B8
7614−Ctx−T−L−Sml−DAPI−B9
7614−Ctx−T−L−Sml−DAPI−C10
7614−Ctx−T−L−Sml−DAPI−C11
7614−Ctx−T−L−Sml−DAPI−C12
7614−Ctx−T−L−Sml−DAPI−C7
7614−Ctx−T−L−Sml−DAPI−C8
7614−Ctx−T−L−Sml−DAPI−C9
7614−Ctx−T−L−Sml−DAPI−D10
7614−Ctx−T−L−Sml−DAPI−D11
7614−Ctx−T−L−Sml−DAPI−D12
7614−Ctx−T−L−Sml−DAPI−D7
7614−Ctx−T−L−Sml−DAPI−D8
7614−Ctx−T−L−Sml−DAPI−D9
7614−Ctx−T−L−Sml−DAPI−E10
7614−Ctx−T−L−Sml−DAPI−E11
7614−Ctx−T−L−Sml−DAPI−E12
7614−Ctx−T−L−Sml−DAPI−E7
7614−Ctx−T−L−Sml−DAPI−E8
7614−Ctx−T−L−Sml−DAPI−E9
7614−Ctx−T−L−Sml−DAPI−F10
7614−Ctx−T−L−Sml−DAPI−F11
7614−Ctx−T−L−Sml−DAPI−F12
7614−Ctx−T−L−Sml−DAPI−F7
7614−Ctx−T−L−Sml−DAPI−F8
7614−Ctx−T−L−Sml−DAPI−F9
7614−Ctx−T−L−Sml−DAPI−G10
7614−Ctx−T−L−Sml−DAPI−G11
7614−Ctx−T−L−Sml−DAPI−G12
7614−Ctx−T−L−Sml−DAPI−G7
7614−Ctx−T−L−Sml−DAPI−G8
7614−Ctx−T−L−Sml−DAPI−G9
7614−Ctx−T−L−Sml−DAPI−H10
7614−Ctx−T−L−Sml−DAPI−H11
7614−Ctx−T−L−Sml−DAPI−H7
7614−Ctx−T−L−Sml−DAPI−H8
7614−Ctx−T−L−Sml−DAPI−H9
7614−Ctx−T−L−Sml−NeuN−A1
7614−Ctx−T−L−Sml−NeuN−A2
7614−Ctx−T−L−Sml−NeuN−A3
7614−Ctx−T−L−Sml−NeuN−A4
7614−Ctx−T−L−Sml−NeuN−A5
7614−Ctx−T−L−Sml−NeuN−A6
7614−Ctx−T−L−Sml−NeuN−B1
7614−Ctx−T−L−Sml−NeuN−B2
7614−Ctx−T−L−Sml−NeuN−B3
7614−Ctx−T−L−Sml−NeuN−B4
7614−Ctx−T−L−Sml−NeuN−B5
7614−Ctx−T−L−Sml−NeuN−B6
7614−Ctx−T−L−Sml−NeuN−C1
7614−Ctx−T−L−Sml−NeuN−C2
7614−Ctx−T−L−Sml−NeuN−C3
7614−Ctx−T−L−Sml−NeuN−C4
7614−Ctx−T−L−Sml−NeuN−C5
7614−Ctx−T−L−Sml−NeuN−C6
7614−Ctx−T−L−Sml−NeuN−D1
7614−Ctx−T−L−Sml−NeuN−D2
7614−Ctx−T−L−Sml−NeuN−D3
7614−Ctx−T−L−Sml−NeuN−D4
7614−Ctx−T−L−Sml−NeuN−D5
7614−Ctx−T−L−Sml−NeuN−D6
7614−Ctx−T−L−Sml−NeuN−E1
7614−Ctx−T−L−Sml−NeuN−E2
7614−Ctx−T−L−Sml−NeuN−E3
7614−Ctx−T−L−Sml−NeuN−E4
7614−Ctx−T−L−Sml−NeuN−E5
7614−Ctx−T−L−Sml−NeuN−E6
7614−Ctx−T−L−Sml−NeuN−F1
7614−Ctx−T−L−Sml−NeuN−F2
7614−Ctx−T−L−Sml−NeuN−F3
7614−Ctx−T−L−Sml−NeuN−F4
7614−Ctx−T−L−Sml−NeuN−F5
7614−Ctx−T−L−Sml−NeuN−F6
7614−Ctx−T−L−Sml−NeuN−G1
7614−Ctx−T−L−Sml−NeuN−G2
7614−Ctx−T−L−Sml−NeuN−G3
7614−Ctx−T−L−Sml−NeuN−G4
7614−Ctx−T−L−Sml−NeuN−G5
7614−Ctx−T−L−Sml−NeuN−G6
7614−Ctx−T−L−Sml−NeuN−H1
7614−Ctx−T−L−Sml−NeuN−H2
7614−Ctx−T−L−Sml−NeuN−H3
7614−Ctx−T−L−Sml−NeuN−H4
7614−Ctx−T−L−Sml−NeuN−H5
7614−Ctx−T−L−Sml−NeuN−H6
JGG−Cells

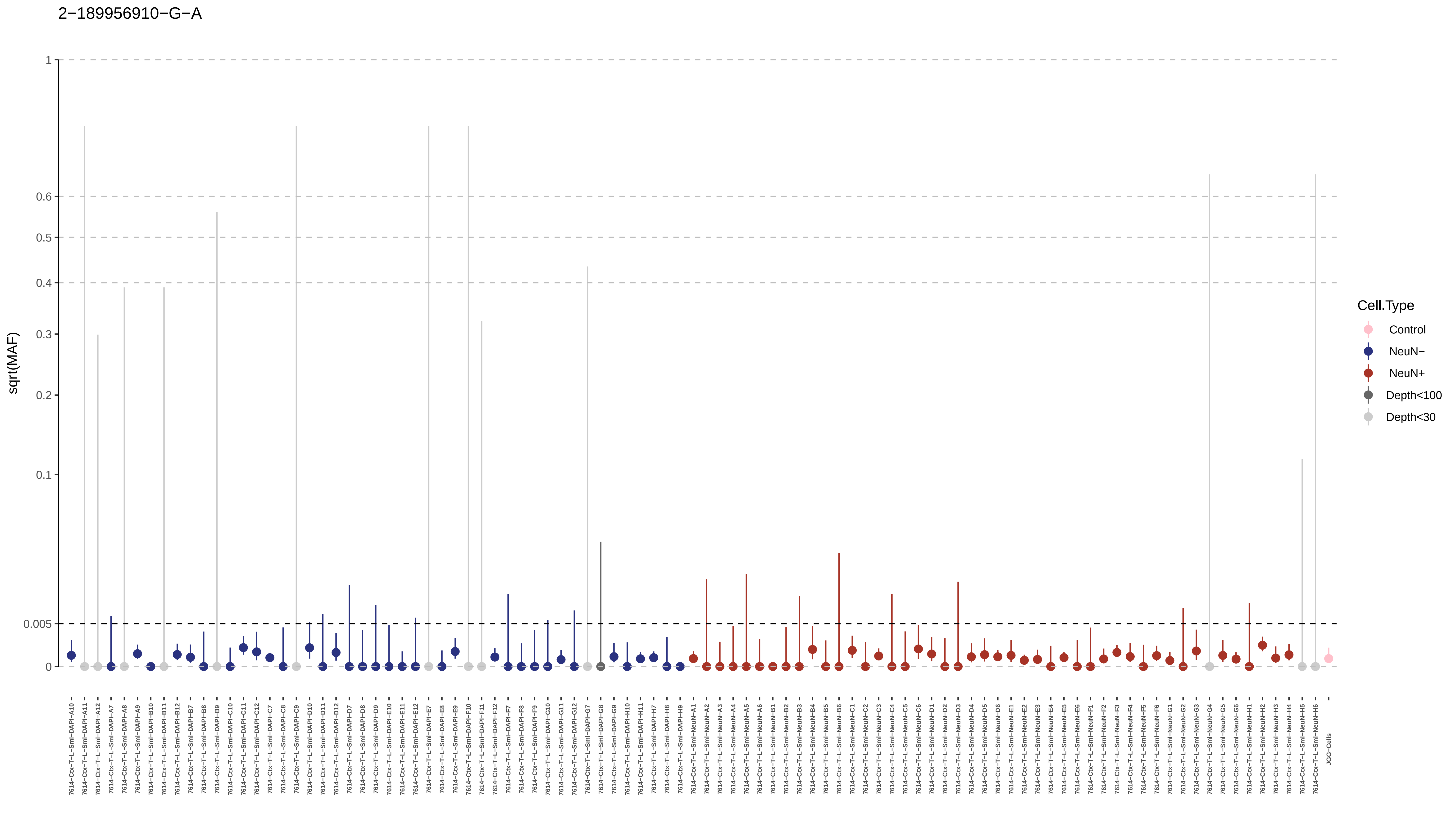
2−189956910−G−A
sqrt(MAF)
1
0.6
0.5
0.4
0.3
0.2
0.1
0.005
0
Cell.Type
Control
NeuN−
NeuN+
Depth<100
Depth<30

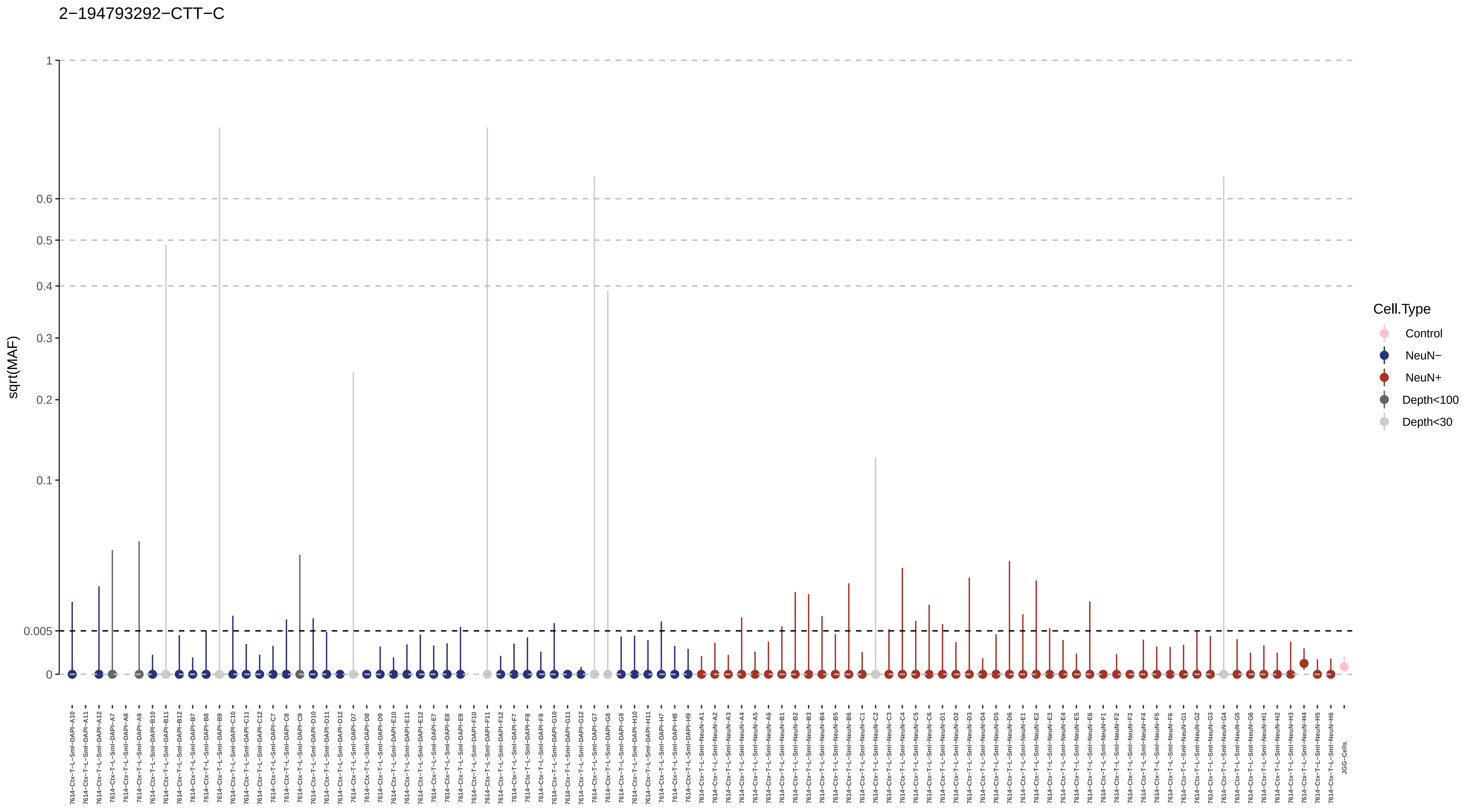

2−194793292−CTT−C
sqrt(MAF)
1
0.6
0.5
0.4
0.3
0.2
0.1
0.005
0
Cell.Type
Control
NeuN−
NeuN+
Depth<100
Depth<30
7614−Ctx−T−L−Sml−DAPI−A10
7614−Ctx−T−L−Sml−DAPI−A11
7614−Ctx−T−L−Sml−DAPI−A12
7614−Ctx−T−L−Sml−DAPI−A7
7614−Ctx−T−L−Sml−DAPI−A8
7614−Ctx−T−L−Sml−DAPI−A9
7614−Ctx−T−L−Sml−DAPI−B10
7614−Ctx−T−L−Sml−DAPI−B11
7614−Ctx−T−L−Sml−DAPI−B12
7614−Ctx−T−L−Sml−DAPI−B7
7614−Ctx−T−L−Sml−DAPI−B8
7614−Ctx−T−L−Sml−DAPI−B9
7614−Ctx−T−L−Sml−DAPI−C10
7614−Ctx−T−L−Sml−DAPI−C11
7614−Ctx−T−L−Sml−DAPI−C12
7614−Ctx−T−L−Sml−DAPI−C7
7614−Ctx−T−L−Sml−DAPI−C8
7614−Ctx−T−L−Sml−DAPI−C9
7614−Ctx−T−L−Sml−DAPI−D10
7614−Ctx−T−L−Sml−DAPI−D11
7614−Ctx−T−L−Sml−DAPI−D12
7614−Ctx−T−L−Sml−DAPI−D7
7614−Ctx−T−L−Sml−DAPI−D8
7614−Ctx−T−L−Sml−DAPI−D9
7614−Ctx−T−L−Sml−DAPI−E10
7614−Ctx−T−L−Sml−DAPI−E11
7614−Ctx−T−L−Sml−DAPI−E12
7614−Ctx−T−L−Sml−DAPI−E7
7614−Ctx−T−L−Sml−DAPI−E8
7614−Ctx−T−L−Sml−DAPI−E9
7614−Ctx−T−L−Sml−DAPI−F10
7614−Ctx−T−L−Sml−DAPI−F11
7614−Ctx−T−L−Sml−DAPI−F12
7614−Ctx−T−L−Sml−DAPI−F7
7614−Ctx−T−L−Sml−DAPI−F8
7614−Ctx−T−L−Sml−DAPI−F9
7614−Ctx−T−L−Sml−DAPI−G10
7614−Ctx−T−L−Sml−DAPI−G11
7614−Ctx−T−L−Sml−DAPI−G12
7614−Ctx−T−L−Sml−DAPI−G7
7614−Ctx−T−L−Sml−DAPI−G8
7614−Ctx−T−L−Sml−DAPI−G9
7614−Ctx−T−L−Sml−DAPI−H10
7614−Ctx−T−L−Sml−DAPI−H11
7614−Ctx−T−L−Sml−DAPI−H7
7614−Ctx−T−L−Sml−DAPI−H8
7614−Ctx−T−L−Sml−DAPI−H9
7614−Ctx−T−L−Sml−NeuN−A1
7614−Ctx−T−L−Sml−NeuN−A2
7614−Ctx−T−L−Sml−NeuN−A3
7614−Ctx−T−L−Sml−NeuN−A4
7614−Ctx−T−L−Sml−NeuN−A5
7614−Ctx−T−L−Sml−NeuN−A6
7614−Ctx−T−L−Sml−NeuN−B1
7614−Ctx−T−L−Sml−NeuN−B2
7614−Ctx−T−L−Sml−NeuN−B3
7614−Ctx−T−L−Sml−NeuN−B4
7614−Ctx−T−L−Sml−NeuN−B5
7614−Ctx−T−L−Sml−NeuN−B6
7614−Ctx−T−L−Sml−NeuN−C1
7614−Ctx−T−L−Sml−NeuN−C2
7614−Ctx−T−L−Sml−NeuN−C3
7614−Ctx−T−L−Sml−NeuN−C4
7614−Ctx−T−L−Sml−NeuN−C5
7614−Ctx−T−L−Sml−NeuN−C6
7614−Ctx−T−L−Sml−NeuN−D1
7614−Ctx−T−L−Sml−NeuN−D2
7614−Ctx−T−L−Sml−NeuN−D3
7614−Ctx−T−L−Sml−NeuN−D4
7614−Ctx−T−L−Sml−NeuN−D5
7614−Ctx−T−L−Sml−NeuN−D6
7614−Ctx−T−L−Sml−NeuN−E1
7614−Ctx−T−L−Sml−NeuN−E2
7614−Ctx−T−L−Sml−NeuN−E3
7614−Ctx−T−L−Sml−NeuN−E4
7614−Ctx−T−L−Sml−NeuN−E5
7614−Ctx−T−L−Sml−NeuN−E6
7614−Ctx−T−L−Sml−NeuN−F1
7614−Ctx−T−L−Sml−NeuN−F2
7614−Ctx−T−L−Sml−NeuN−F3
7614−Ctx−T−L−Sml−NeuN−F4
7614−Ctx−T−L−Sml−NeuN−F5
7614−Ctx−T−L−Sml−NeuN−F6
7614−Ctx−T−L−Sml−NeuN−G1
7614−Ctx−T−L−Sml−NeuN−G2
7614−Ctx−T−L−Sml−NeuN−G3
7614−Ctx−T−L−Sml−NeuN−G4
7614−Ctx−T−L−Sml−NeuN−G5
7614−Ctx−T−L−Sml−NeuN−G6
7614−Ctx−T−L−Sml−NeuN−H1
7614−Ctx−T−L−Sml−NeuN−H2
7614−Ctx−T−L−Sml−NeuN−H3
7614−Ctx−T−L−Sml−NeuN−H4
7614−Ctx−T−L−Sml−NeuN−H5
7614−Ctx−T−L−Sml−NeuN−H6
JGG−Cells

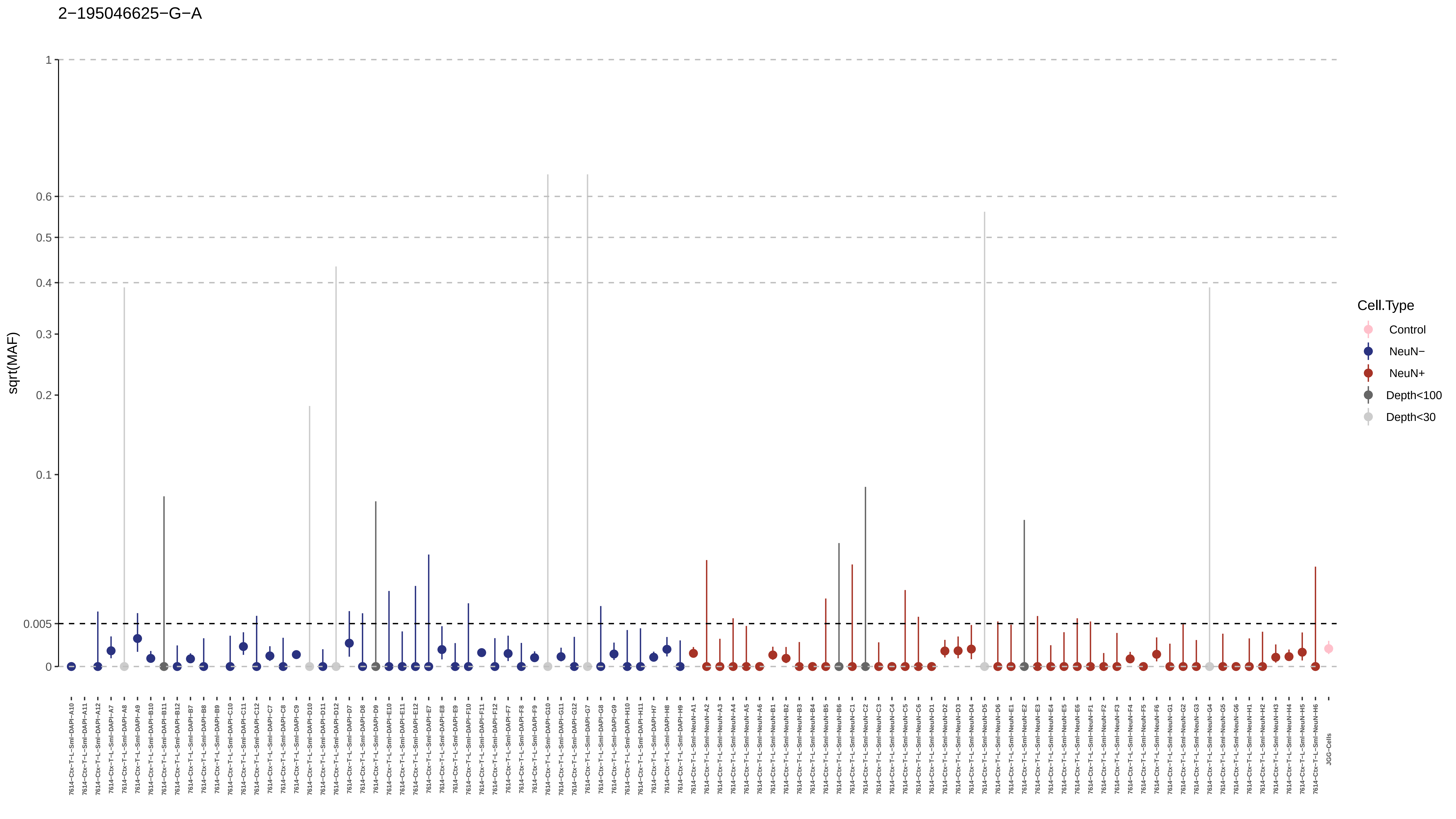

2−195046625−G−A
sqrt(MAF)
1
0.6
0.5
0.4
0.3
0.2
0.1
0.005
0
Cell.Type
Control
NeuN−
NeuN+
Depth<100
Depth<30
7614−Ctx−T−L−Sml−DAPI−A10
7614−Ctx−T−L−Sml−DAPI−A11
7614−Ctx−T−L−Sml−DAPI−A12
7614−Ctx−T−L−Sml−DAPI−A7
7614−Ctx−T−L−Sml−DAPI−A8
7614−Ctx−T−L−Sml−DAPI−A9
7614−Ctx−T−L−Sml−DAPI−B10
7614−Ctx−T−L−Sml−DAPI−B11
7614−Ctx−T−L−Sml−DAPI−B12
7614−Ctx−T−L−Sml−DAPI−B7
7614−Ctx−T−L−Sml−DAPI−B8
7614−Ctx−T−L−Sml−DAPI−B9
7614−Ctx−T−L−Sml−DAPI−C10
7614−Ctx−T−L−Sml−DAPI−C11
7614−Ctx−T−L−Sml−DAPI−C12
7614−Ctx−T−L−Sml−DAPI−C7
7614−Ctx−T−L−Sml−DAPI−C8
7614−Ctx−T−L−Sml−DAPI−C9
7614−Ctx−T−L−Sml−DAPI−D10
7614−Ctx−T−L−Sml−DAPI−D11
7614−Ctx−T−L−Sml−DAPI−D12
7614−Ctx−T−L−Sml−DAPI−D7
7614−Ctx−T−L−Sml−DAPI−D8
7614−Ctx−T−L−Sml−DAPI−D9
7614−Ctx−T−L−Sml−DAPI−E10
7614−Ctx−T−L−Sml−DAPI−E11
7614−Ctx−T−L−Sml−DAPI−E12
7614−Ctx−T−L−Sml−DAPI−E7
7614−Ctx−T−L−Sml−DAPI−E8
7614−Ctx−T−L−Sml−DAPI−E9
7614−Ctx−T−L−Sml−DAPI−F10
7614−Ctx−T−L−Sml−DAPI−F11
7614−Ctx−T−L−Sml−DAPI−F12
7614−Ctx−T−L−Sml−DAPI−F7
7614−Ctx−T−L−Sml−DAPI−F8
7614−Ctx−T−L−Sml−DAPI−F9
7614−Ctx−T−L−Sml−DAPI−G10
7614−Ctx−T−L−Sml−DAPI−G11
7614−Ctx−T−L−Sml−DAPI−G12
7614−Ctx−T−L−Sml−DAPI−G7
7614−Ctx−T−L−Sml−DAPI−G8
7614−Ctx−T−L−Sml−DAPI−G9
7614−Ctx−T−L−Sml−DAPI−H10
7614−Ctx−T−L−Sml−DAPI−H11
7614−Ctx−T−L−Sml−DAPI−H7
7614−Ctx−T−L−Sml−DAPI−H8
7614−Ctx−T−L−Sml−DAPI−H9
7614−Ctx−T−L−Sml−NeuN−A1
7614−Ctx−T−L−Sml−NeuN−A2
7614−Ctx−T−L−Sml−NeuN−A3
7614−Ctx−T−L−Sml−NeuN−A4
7614−Ctx−T−L−Sml−NeuN−A5
7614−Ctx−T−L−Sml−NeuN−A6
7614−Ctx−T−L−Sml−NeuN−B1
7614−Ctx−T−L−Sml−NeuN−B2
7614−Ctx−T−L−Sml−NeuN−B3
7614−Ctx−T−L−Sml−NeuN−B4
7614−Ctx−T−L−Sml−NeuN−B5
7614−Ctx−T−L−Sml−NeuN−B6
7614−Ctx−T−L−Sml−NeuN−C1
7614−Ctx−T−L−Sml−NeuN−C2
7614−Ctx−T−L−Sml−NeuN−C3
7614−Ctx−T−L−Sml−NeuN−C4
7614−Ctx−T−L−Sml−NeuN−C5
7614−Ctx−T−L−Sml−NeuN−C6
7614−Ctx−T−L−Sml−NeuN−D1
7614−Ctx−T−L−Sml−NeuN−D2
7614−Ctx−T−L−Sml−NeuN−D3
7614−Ctx−T−L−Sml−NeuN−D4
7614−Ctx−T−L−Sml−NeuN−D5
7614−Ctx−T−L−Sml−NeuN−D6
7614−Ctx−T−L−Sml−NeuN−E1
7614−Ctx−T−L−Sml−NeuN−E2
7614−Ctx−T−L−Sml−NeuN−E3
7614−Ctx−T−L−Sml−NeuN−E4
7614−Ctx−T−L−Sml−NeuN−E5
7614−Ctx−T−L−Sml−NeuN−E6
7614−Ctx−T−L−Sml−NeuN−F1
7614−Ctx−T−L−Sml−NeuN−F2
7614−Ctx−T−L−Sml−NeuN−F3
7614−Ctx−T−L−Sml−NeuN−F4
7614−Ctx−T−L−Sml−NeuN−F5
7614−Ctx−T−L−Sml−NeuN−F6
7614−Ctx−T−L−Sml−NeuN−G1
7614−Ctx−T−L−Sml−NeuN−G2
7614−Ctx−T−L−Sml−NeuN−G3
7614−Ctx−T−L−Sml−NeuN−G4
7614−Ctx−T−L−Sml−NeuN−G5
7614−Ctx−T−L−Sml−NeuN−G6
7614−Ctx−T−L−Sml−NeuN−H1
7614−Ctx−T−L−Sml−NeuN−H2
7614−Ctx−T−L−Sml−NeuN−H3
7614−Ctx−T−L−Sml−NeuN−H4
7614−Ctx−T−L−Sml−NeuN−H5
7614−Ctx−T−L−Sml−NeuN−H6
JGG-Cells

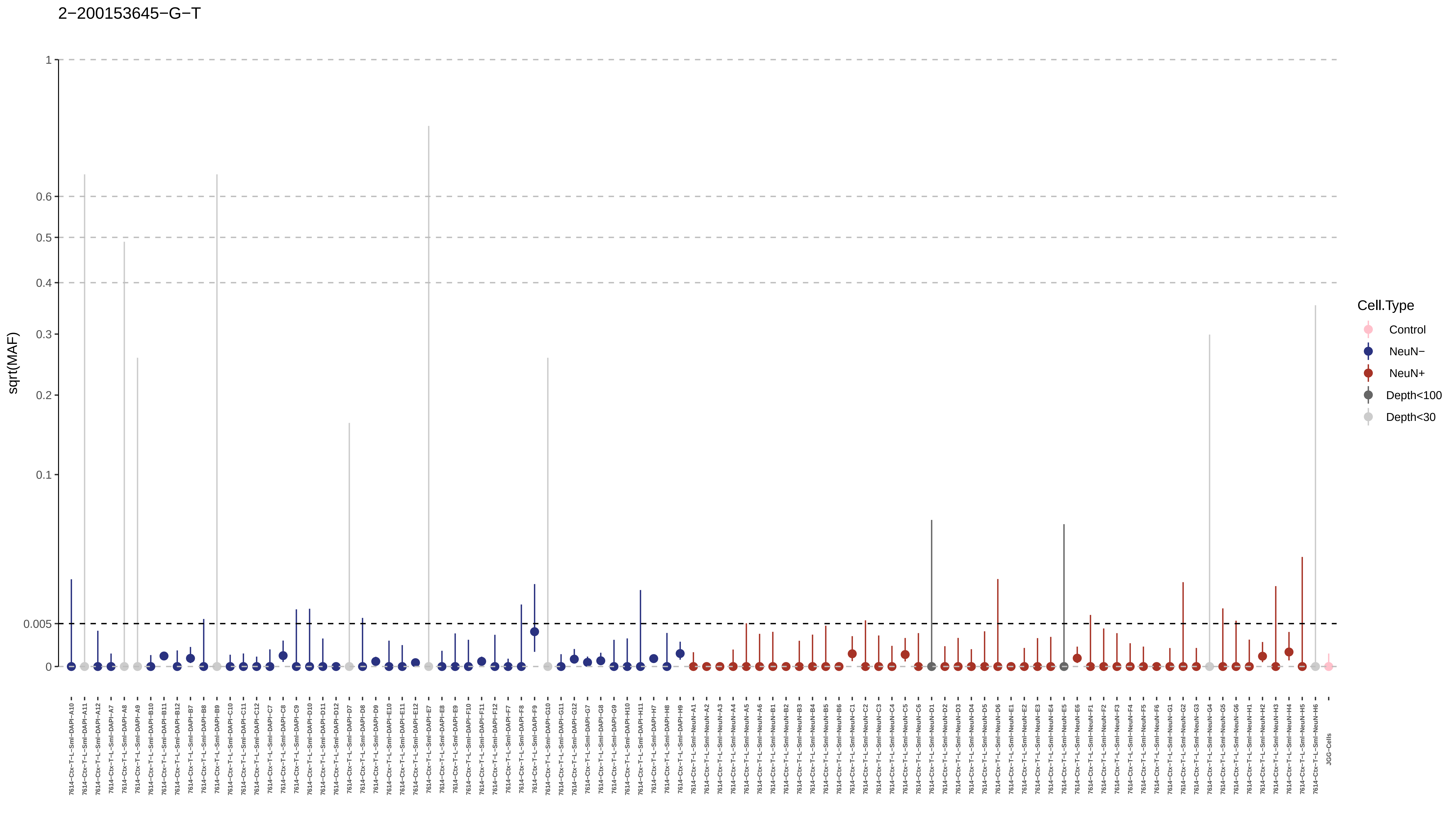

2−200153645−G−T
sqrt(MAF)
1
0.6
0.5
0.4
0.3
0.2
0.1
0.005
0
Cell.Type
Control
NeuN−
NeuN+
Depth<100
Depth<30

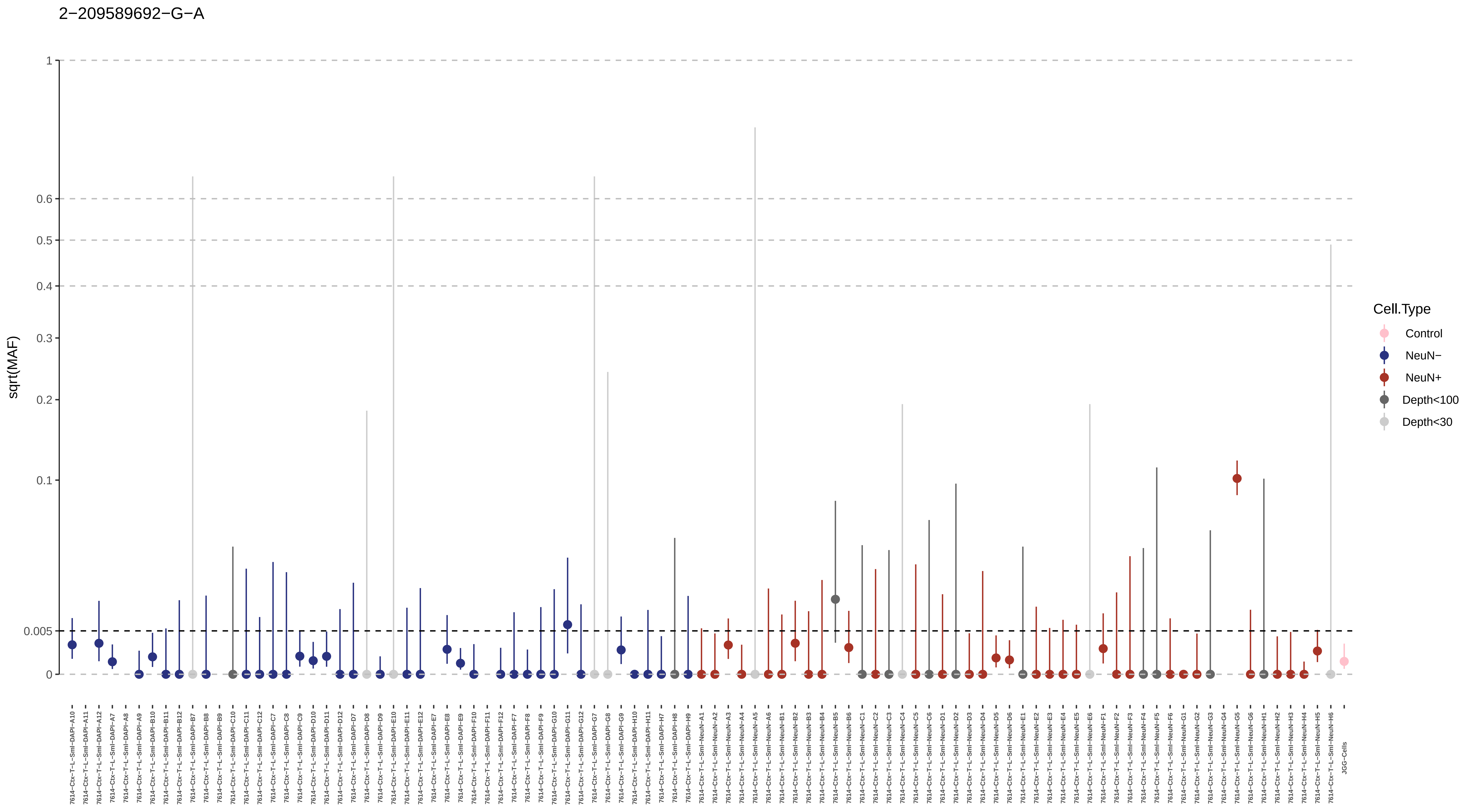

2−209589692−G−A
sqrt(MAF)
1
0.6
0.5
0.4
0.3
0.2
0.1
0.005
0
Cell.Type
Control
NeuN−
NeuN+
Depth<100
Depth<30
7614−Ctx−T−L−Sml−DAPI−A10
7614−Ctx−T−L−Sml−DAPI−A11
7614−Ctx−T−L−Sml−DAPI−A12
7614−Ctx−T−L−Sml−DAPI−A7
7614−Ctx−T−L−Sml−DAPI−A8
7614−Ctx−T−L−Sml−DAPI−A9
7614−Ctx−T−L−Sml−DAPI−B10
7614−Ctx−T−L−Sml−DAPI−B11
7614−Ctx−T−L−Sml−DAPI−B12
7614−Ctx−T−L−Sml−DAPI−B7
7614−Ctx−T−L−Sml−DAPI−B8
7614−Ctx−T−L−Sml−DAPI−B9
7614−Ctx−T−L−Sml−DAPI−C10
7614−Ctx−T−L−Sml−DAPI−C11
7614−Ctx−T−L−Sml−DAPI−C12
7614−Ctx−T−L−Sml−DAPI−C7
7614−Ctx−T−L−Sml−DAPI−C8
7614−Ctx−T−L−Sml−DAPI−C9
7614−Ctx−T−L−Sml−DAPI−D10
7614−Ctx−T−L−Sml−DAPI−D11
7614−Ctx−T−L−Sml−DAPI−D12
7614−Ctx−T−L−Sml−DAPI−D7
7614−Ctx−T−L−Sml−DAPI−D8
7614−Ctx−T−L−Sml−DAPI−D9
7614−Ctx−T−L−Sml−DAPI−E10
7614−Ctx−T−L−Sml−DAPI−E11
7614−Ctx−T−L−Sml−DAPI−E12
7614−Ctx−T−L−Sml−DAPI−E7
7614−Ctx−T−L−Sml−DAPI−E8
7614−Ctx−T−L−Sml−DAPI−E9
7614−Ctx−T−L−Sml−DAPI−F10
7614−Ctx−T−L−Sml−DAPI−F11
7614−Ctx−T−L−Sml−DAPI−F12
7614−Ctx−T−L−Sml−DAPI−F7
7614−Ctx−T−L−Sml−DAPI−F8
7614−Ctx−T−L−Sml−DAPI−F9
7614−Ctx−T−L−Sml−DAPI−G10
7614−Ctx−T−L−Sml−DAPI−G11
7614−Ctx−T−L−Sml−DAPI−G12
7614−Ctx−T−L−Sml−DAPI−G7
7614−Ctx−T−L−Sml−DAPI−G8
7614−Ctx−T−L−Sml−DAPI−G9
7614−Ctx−T−L−Sml−DAPI−H10
7614−Ctx−T−L−Sml−DAPI−H11
7614−Ctx−T−L−Sml−DAPI−H7
7614−Ctx−T−L−Sml−DAPI−H8
7614−Ctx−T−L−Sml−DAPI−H9
7614−Ctx−T−L−Sml−NeuN−A1
7614−Ctx−T−L−Sml−NeuN−A2
7614−Ctx−T−L−Sml−NeuN−A3
7614−Ctx−T−L−Sml−NeuN−A4
7614−Ctx−T−L−Sml−NeuN−A5
7614−Ctx−T−L−Sml−NeuN−A6
7614−Ctx−T−L−Sml−NeuN−B1
7614−Ctx−T−L−Sml−NeuN−B2
7614−Ctx−T−L−Sml−NeuN−B3
7614−Ctx−T−L−Sml−NeuN−B4
7614−Ctx−T−L−Sml−NeuN−B5
7614−Ctx−T−L−Sml−NeuN−B6
7614−Ctx−T−L−Sml−NeuN−C1
7614−Ctx−T−L−Sml−NeuN−C2
7614−Ctx−T−L−Sml−NeuN−C3
7614−Ctx−T−L−Sml−NeuN−C4
7614−Ctx−T−L−Sml−NeuN−C5
7614−Ctx−T−L−Sml−NeuN−C6
7614−Ctx−T−L−Sml−NeuN−D1
7614−Ctx−T−L−Sml−NeuN−D2
7614−Ctx−T−L−Sml−NeuN−D3
7614−Ctx−T−L−Sml−NeuN−D4
7614−Ctx−T−L−Sml−NeuN−D5
7614−Ctx−T−L−Sml−NeuN−D6
7614−Ctx−T−L−Sml−NeuN−E1
7614−Ctx−T−L−Sml−NeuN−E2
7614−Ctx−T−L−Sml−NeuN−E3
7614−Ctx−T−L−Sml−NeuN−E4
7614−Ctx−T−L−Sml−NeuN−E5
7614−Ctx−T−L−Sml−NeuN−E6
7614−Ctx−T−L−Sml−NeuN−F1
7614−Ctx−T−L−Sml−NeuN−F2
7614−Ctx−T−L−Sml−NeuN−F3
7614−Ctx−T−L−Sml−NeuN−F4
7614−Ctx−T−L−Sml−NeuN−F5
7614−Ctx−T−L−Sml−NeuN−F6
7614−Ctx−T−L−Sml−NeuN−G1
7614−Ctx−T−L−Sml−NeuN−G2
7614−Ctx−T−L−Sml−NeuN−G3
7614−Ctx−T−L−Sml−NeuN−G4
7614−Ctx−T−L−Sml−NeuN−G5
7614−Ctx−T−L−Sml−NeuN−G6
7614−Ctx−T−L−Sml−NeuN−H1
7614−Ctx−T−L−Sml−NeuN−H2
7614−Ctx−T−L−Sml−NeuN−H3
7614−Ctx−T−L−Sml−NeuN−H4
7614−Ctx−T−L−Sml−NeuN−H5
7614−Ctx−T−L−Sml−NeuN−H6
JGG-Cells

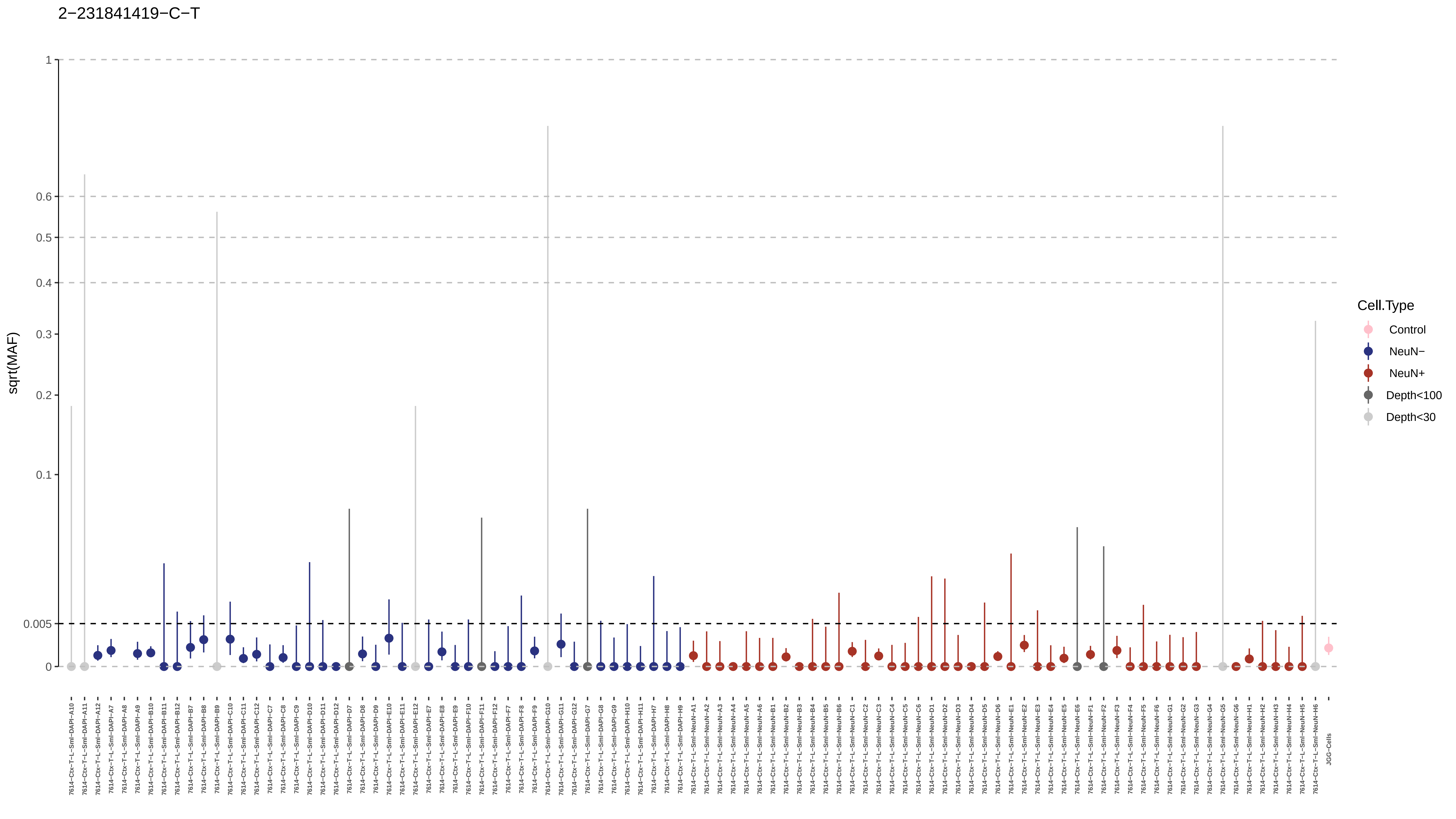

2−231841419−C−T
sqrt(MAF)
1
0.6
0.5
0.4
0.3
0.2
0.1
0.005
0
Cell.Type
Control
NeuN−
NeuN+
Depth<100
Depth<30
JGG-Cells

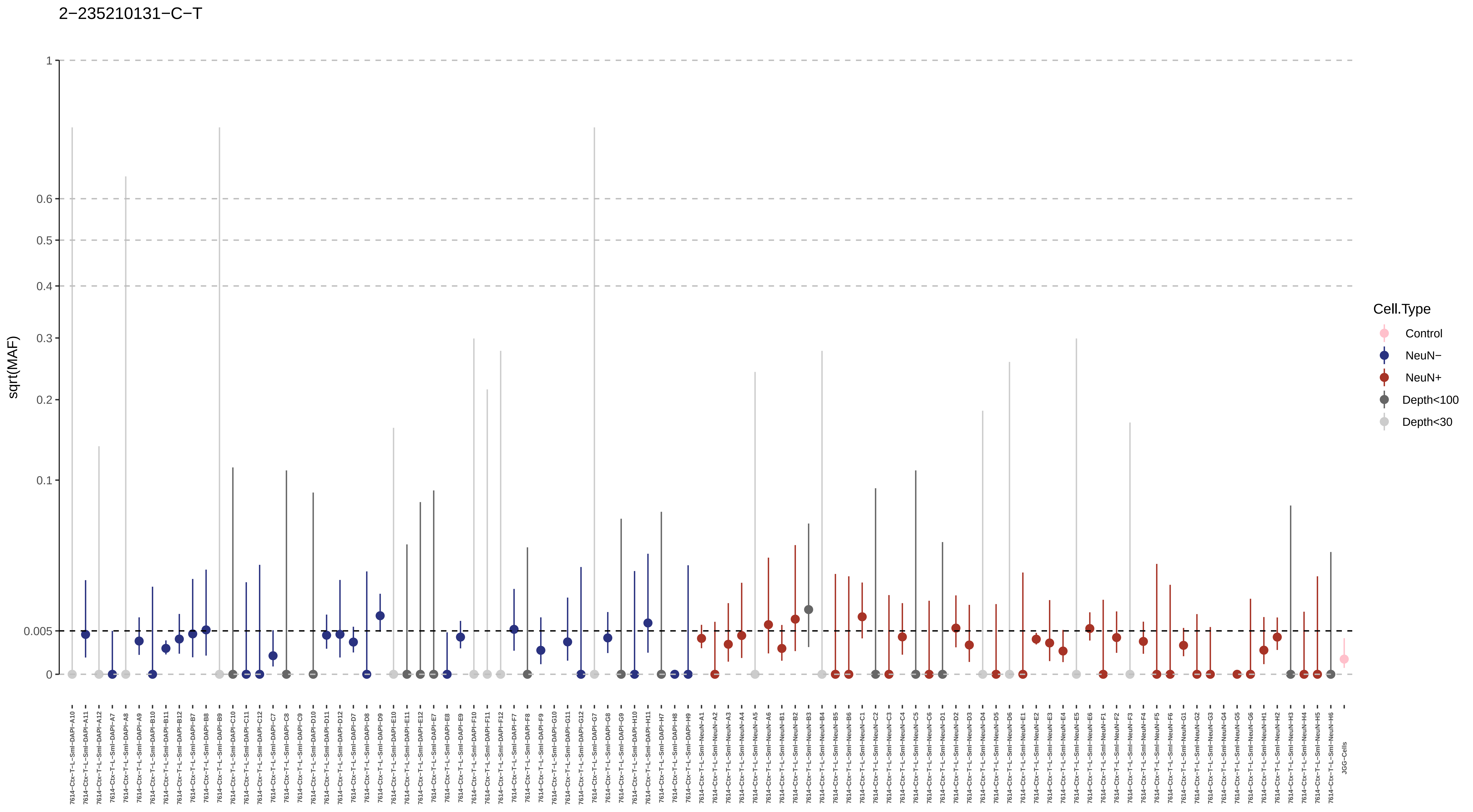
2−235210131−C−T
sqrt(MAF)
1
0.6
0.5
0.4
0.3
0.2
0.1
0.005
0
Cell.Type
Control
NeuN−
NeuN+
Depth<100
Depth<30

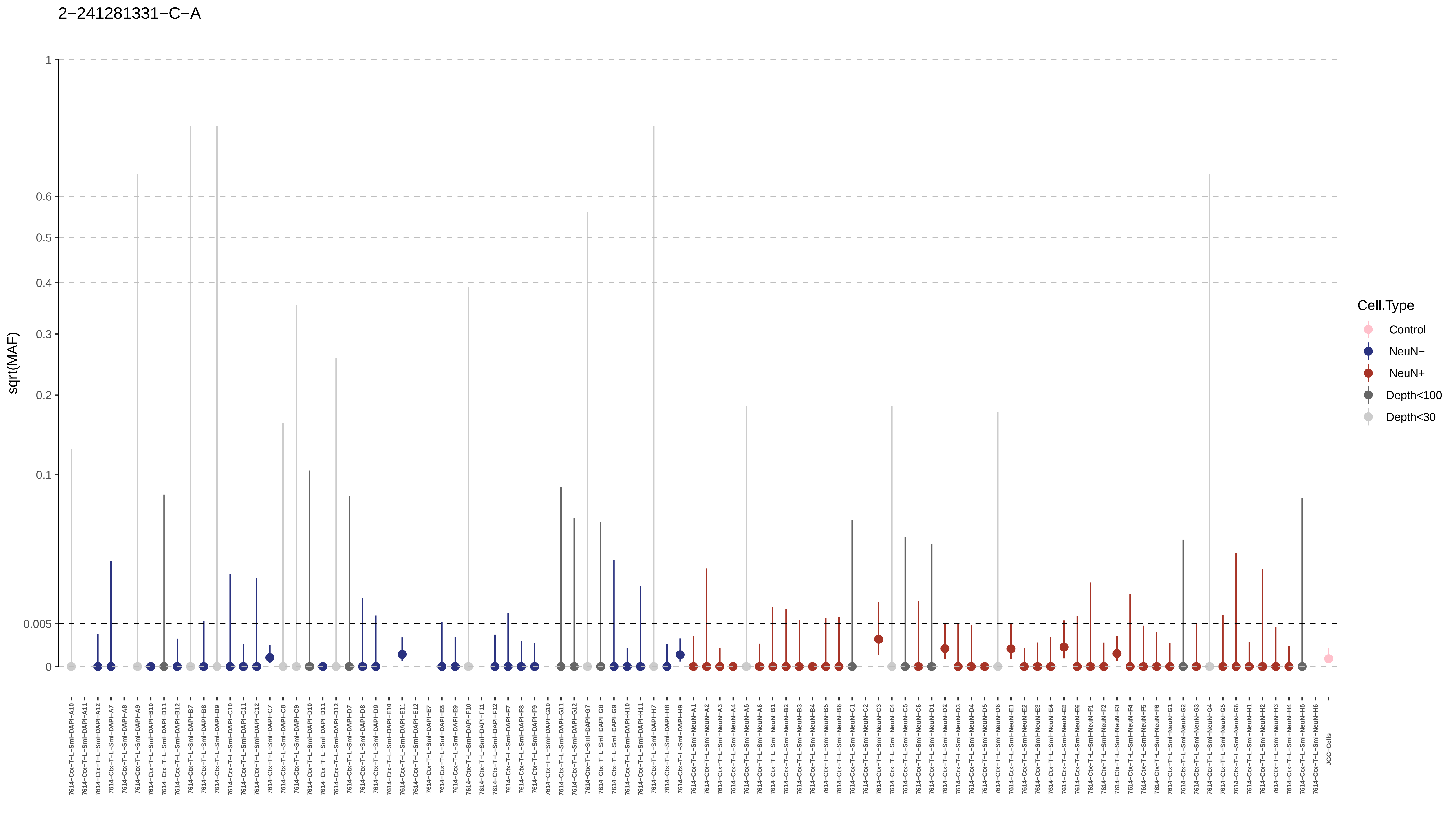

2−241281331−C−A
sqrt(MAF)
1
0.6
0.5
0.4
0.3
0.2
0.1
0.005
0
Cell.Type
Control
NeuN−
NeuN+
Depth<100
Depth<30
JGG-Cells

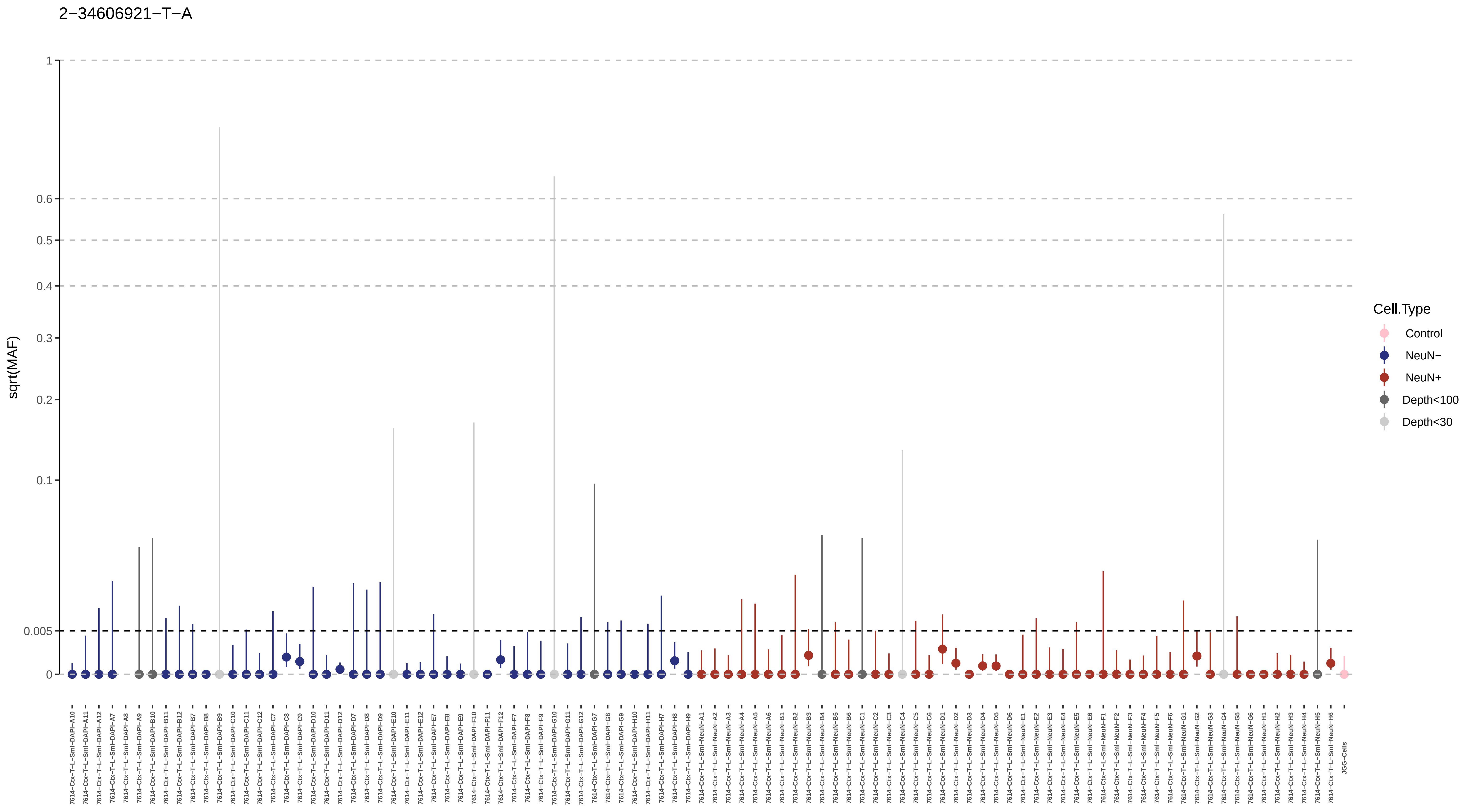
2−34606921−T−A
sqrt(MAF)
1
0.6
0.5
0.4
0.3
0.2
0.1
0.005
0
7614−Ctx−T−L−Sml−DAPI−A10
7614−Ctx−T−L−Sml−DAPI−A11
7614−Ctx−T−L−Sml−DAPI−A12
7614−Ctx−T−L−Sml−DAPI−A7
7614−Ctx−T−L−Sml−DAPI−A8
7614−Ctx−T−L−Sml−DAPI−A9
7614−Ctx−T−L−Sml−DAPI−B10
7614−Ctx−T−L−Sml−DAPI−B11
7614−Ctx−T−L−Sml−DAPI−B12
7614−Ctx−T−L−Sml−DAPI−B7
7614−Ctx−T−L−Sml−DAPI−B8
7614−Ctx−T−L−Sml−DAPI−B9
7614−Ctx−T−L−Sml−DAPI−C10
7614−Ctx−T−L−Sml−DAPI−C11
7614−Ctx−T−L−Sml−DAPI−C12
7614−Ctx−T−L−Sml−DAPI−C7
7614−Ctx−T−L−Sml−DAPI−C8
7614−Ctx−T−L−Sml−DAPI−C9
7614−Ctx−T−L−Sml−DAPI−D10
7614−Ctx−T−L−Sml−DAPI−D11
7614−Ctx−T−L−Sml−DAPI−D12
7614−Ctx−T−L−Sml−DAPI−D7
7614−Ctx−T−L−Sml−DAPI−D8
7614−Ctx−T−L−Sml−DAPI−D9
7614−Ctx−T−L−Sml−DAPI−E10
7614−Ctx−T−L−Sml−DAPI−E11
7614−Ctx−T−L−Sml−DAPI−E12
7614−Ctx−T−L−Sml−DAPI−E7
7614−Ctx−T−L−Sml−DAPI−E8
7614−Ctx−T−L−Sml−DAPI−E9
7614−Ctx−T−L−Sml−DAPI−F10
7614−Ctx−T−L−Sml−DAPI−F11
7614−Ctx−T−L−Sml−DAPI−F12
7614−Ctx−T−L−Sml−DAPI−F7
7614−Ctx−T−L−Sml−DAPI−F8
7614−Ctx−T−L−Sml−DAPI−F9
7614−Ctx−T−L−Sml−DAPI−G10
7614−Ctx−T−L−Sml−DAPI−G11
7614−Ctx−T−L−Sml−DAPI−G12
7614−Ctx−T−L−Sml−DAPI−G7
7614−Ctx−T−L−Sml−DAPI−G8
7614−Ctx−T−L−Sml−DAPI−G9
7614−Ctx−T−L−Sml−DAPI−H10
7614−Ctx−T−L−Sml−DAPI−H11
7614−Ctx−T−L−Sml−DAPI−H7
7614−Ctx−T−L−Sml−DAPI−H8
7614−Ctx−T−L−Sml−DAPI−H9
7614−Ctx−T−L−Sml−NeuN−A1
7614−Ctx−T−L−Sml−NeuN−A2
7614−Ctx−T−L−Sml−NeuN−A3
7614−Ctx−T−L−Sml−NeuN−A4
7614−Ctx−T−L−Sml−NeuN−A5
7614−Ctx−T−L−Sml−NeuN−A6
7614−Ctx−T−L−Sml−NeuN−B1
7614−Ctx−T−L−Sml−NeuN−B2
7614−Ctx−T−L−Sml−NeuN−B3
7614−Ctx−T−L−Sml−NeuN−B4
7614−Ctx−T−L−Sml−NeuN−B5
7614−Ctx−T−L−Sml−NeuN−B6
7614−Ctx−T−L−Sml−NeuN−C1
7614−Ctx−T−L−Sml−NeuN−C2
7614−Ctx−T−L−Sml−NeuN−C3
7614−Ctx−T−L−Sml−NeuN−C4
7614−Ctx−T−L−Sml−NeuN−C5
7614−Ctx−T−L−Sml−NeuN−C6
7614−Ctx−T−L−Sml−NeuN−D1
7614−Ctx−T−L−Sml−NeuN−D2
7614−Ctx−T−L−Sml−NeuN−D3
7614−Ctx−T−L−Sml−NeuN−D4
7614−Ctx−T−L−Sml−NeuN−D5
7614−Ctx−T−L−Sml−NeuN−D6
7614−Ctx−T−L−Sml−NeuN−E1
7614−Ctx−T−L−Sml−NeuN−E2
7614−Ctx−T−L−Sml−NeuN−E3
7614−Ctx−T−L−Sml−NeuN−E4
7614−Ctx−T−L−Sml−NeuN−E5
7614−Ctx−T−L−Sml−NeuN−E6
7614−Ctx−T−L−Sml−NeuN−F1
7614−Ctx−T−L−Sml−NeuN−F2
7614−Ctx−T−L−Sml−NeuN−F3
7614−Ctx−T−L−Sml−NeuN−F4
7614−Ctx−T−L−Sml−NeuN−F5
7614−Ctx−T−L−Sml−NeuN−F6
7614−Ctx−T−L−Sml−NeuN−G1
7614−Ctx−T−L−Sml−NeuN−G2
7614−Ctx−T−L−Sml−NeuN−G3
7614−Ctx−T−L−Sml−NeuN−G4
7614−Ctx−T−L−Sml−NeuN−G5
7614−Ctx−T−L−Sml−NeuN−G6
7614−Ctx−T−L−Sml−NeuN−H1
7614−Ctx−T−L−Sml−NeuN−H2
7614−Ctx−T−L−Sml−NeuN−H3
7614−Ctx−T−L−Sml−NeuN−H4
7614−Ctx−T−L−Sml−NeuN−H5
7614−Ctx−T−L−Sml−NeuN−H6
JGG−Cells
Cell.Type
Control
NeuN−
NeuN+
Depth<100
Depth<30

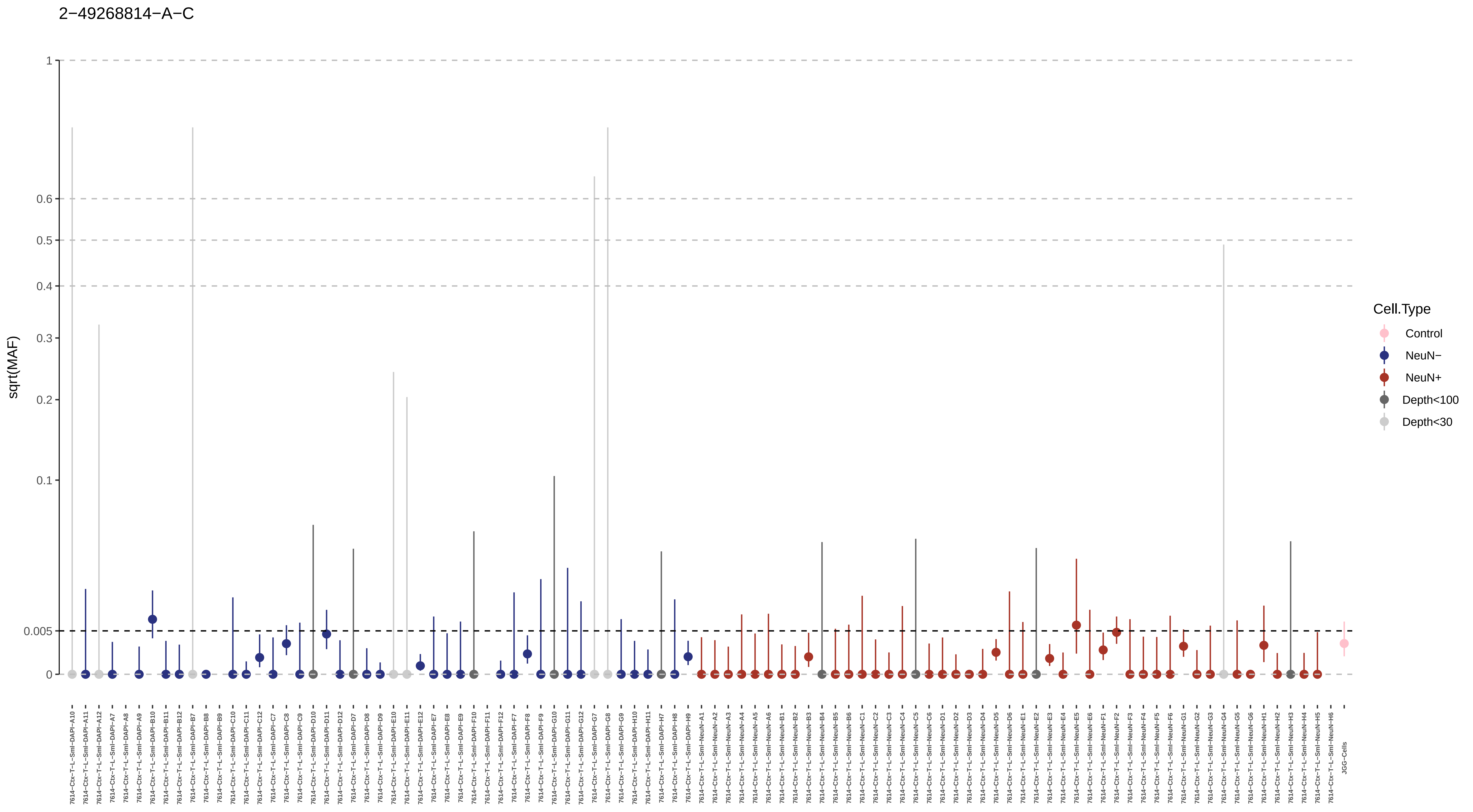

2−49268814−A−C
sqrt(MAF)
1
0.6
0.5
0.4
0.3
0.2
0.1
0.005
0
Cell.Type
Control
NeuN−
NeuN+
Depth<100
Depth<30

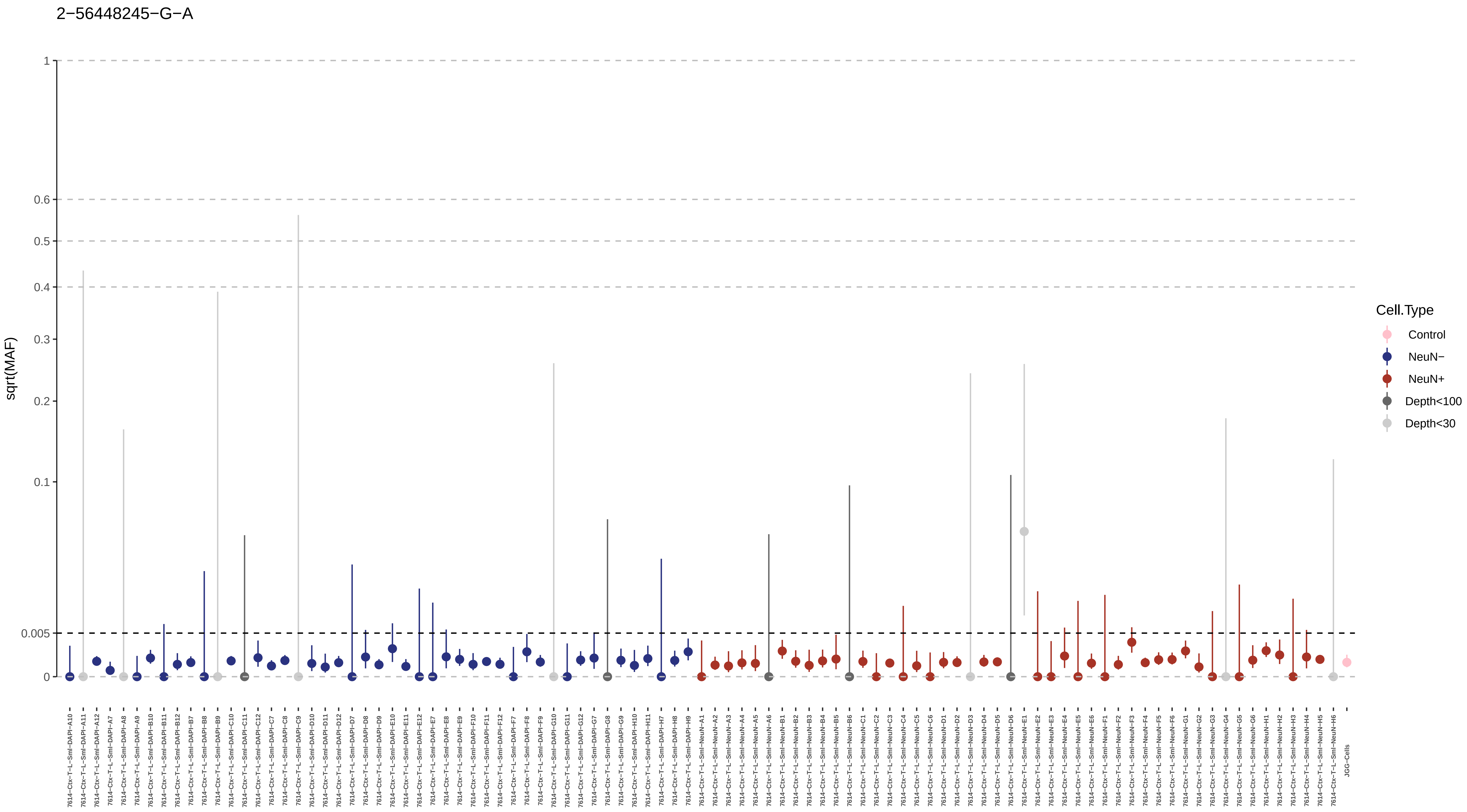

2−56448245−G−A
sqrt(MAF)
1
0.6
0.5
0.4
0.3
0.2
0.1
0.005
0
Cell.Type
Control
NeuN−
NeuN+
Depth<100
Depth<30
7614−Ctx−T−L−Sml−DAPI−A10
7614−Ctx−T−L−Sml−DAPI−A11
7614−Ctx−T−L−Sml−DAPI−A12
7614−Ctx−T−L−Sml−DAPI−A7
7614−Ctx−T−L−Sml−DAPI−A8
7614−Ctx−T−L−Sml−DAPI−A9
7614−Ctx−T−L−Sml−DAPI−B10
7614−Ctx−T−L−Sml−DAPI−B11
7614−Ctx−T−L−Sml−DAPI−B12
7614−Ctx−T−L−Sml−DAPI−B7
7614−Ctx−T−L−Sml−DAPI−B8
7614−Ctx−T−L−Sml−DAPI−B9
7614−Ctx−T−L−Sml−DAPI−C10
7614−Ctx−T−L−Sml−DAPI−C11
7614−Ctx−T−L−Sml−DAPI−C12
7614−Ctx−T−L−Sml−DAPI−C7
7614−Ctx−T−L−Sml−DAPI−C8
7614−Ctx−T−L−Sml−DAPI−C9
7614−Ctx−T−L−Sml−DAPI−D10
7614−Ctx−T−L−Sml−DAPI−D11
7614−Ctx−T−L−Sml−DAPI−D12
7614−Ctx−T−L−Sml−DAPI−D7
7614−Ctx−T−L−Sml−DAPI−D8
7614−Ctx−T−L−Sml−DAPI−D9
7614−Ctx−T−L−Sml−DAPI−E10
7614−Ctx−T−L−Sml−DAPI−E11
7614−Ctx−T−L−Sml−DAPI−E12
7614−Ctx−T−L−Sml−DAPI−E7
7614−Ctx−T−L−Sml−DAPI−E8
7614−Ctx−T−L−Sml−DAPI−E9
7614−Ctx−T−L−Sml−DAPI−F10
7614−Ctx−T−L−Sml−DAPI−F11
7614−Ctx−T−L−Sml−DAPI−F12
7614−Ctx−T−L−Sml−DAPI−F7
7614−Ctx−T−L−Sml−DAPI−F8
7614−Ctx−T−L−Sml−DAPI−F9
7614−Ctx−T−L−Sml−DAPI−G10
7614−Ctx−T−L−Sml−DAPI−G11
7614−Ctx−T−L−Sml−DAPI−G12
7614−Ctx−T−L−Sml−DAPI−G7
7614−Ctx−T−L−Sml−DAPI−G8
7614−Ctx−T−L−Sml−DAPI−G9
7614−Ctx−T−L−Sml−DAPI−H10
7614−Ctx−T−L−Sml−DAPI−H11
7614−Ctx−T−L−Sml−DAPI−H7
7614−Ctx−T−L−Sml−DAPI−H8
7614−Ctx−T−L−Sml−DAPI−H9
7614−Ctx−T−L−Sml−NeuN−A1
7614−Ctx−T−L−Sml−NeuN−A2
7614−Ctx−T−L−Sml−NeuN−A3
7614−Ctx−T−L−Sml−NeuN−A4
7614−Ctx−T−L−Sml−NeuN−A5
7614−Ctx−T−L−Sml−NeuN−A6
7614−Ctx−T−L−Sml−NeuN−B1
7614−Ctx−T−L−Sml−NeuN−B2
7614−Ctx−T−L−Sml−NeuN−B3
7614−Ctx−T−L−Sml−NeuN−B4
7614−Ctx−T−L−Sml−NeuN−B5
7614−Ctx−T−L−Sml−NeuN−B6
7614−Ctx−T−L−Sml−NeuN−C1
7614−Ctx−T−L−Sml−NeuN−C2
7614−Ctx−T−L−Sml−NeuN−C3
7614−Ctx−T−L−Sml−NeuN−C4
7614−Ctx−T−L−Sml−NeuN−C5
7614−Ctx−T−L−Sml−NeuN−C6
7614−Ctx−T−L−Sml−NeuN−D1
7614−Ctx−T−L−Sml−NeuN−D2
7614−Ctx−T−L−Sml−NeuN−D3
7614−Ctx−T−L−Sml−NeuN−D4
7614−Ctx−T−L−Sml−NeuN−D5
7614−Ctx−T−L−Sml−NeuN−D6
7614−Ctx−T−L−Sml−NeuN−E1
7614−Ctx−T−L−Sml−NeuN−E2
7614−Ctx−T−L−Sml−NeuN−E3
7614−Ctx−T−L−Sml−NeuN−E4
7614−Ctx−T−L−Sml−NeuN−E5
7614−Ctx−T−L−Sml−NeuN−E6
7614−Ctx−T−L−Sml−NeuN−F1
7614−Ctx−T−L−Sml−NeuN−F2
7614−Ctx−T−L−Sml−NeuN−F3
7614−Ctx−T−L−Sml−NeuN−F4
7614−Ctx−T−L−Sml−NeuN−F5
7614−Ctx−T−L−Sml−NeuN−F6
7614−Ctx−T−L−Sml−NeuN−G1
7614−Ctx−T−L−Sml−NeuN−G2
7614−Ctx−T−L−Sml−NeuN−G3
7614−Ctx−T−L−Sml−NeuN−G4
7614−Ctx−T−L−Sml−NeuN−G5
7614−Ctx−T−L−Sml−NeuN−G6
7614−Ctx−T−L−Sml−NeuN−H1
7614−Ctx−T−L−Sml−NeuN−H2
7614−Ctx−T−L−Sml−NeuN−H3
7614−Ctx−T−L−Sml−NeuN−H4
7614−Ctx−T−L−Sml−NeuN−H5
7614−Ctx−T−L−Sml−NeuN−H6
JGG−Cells

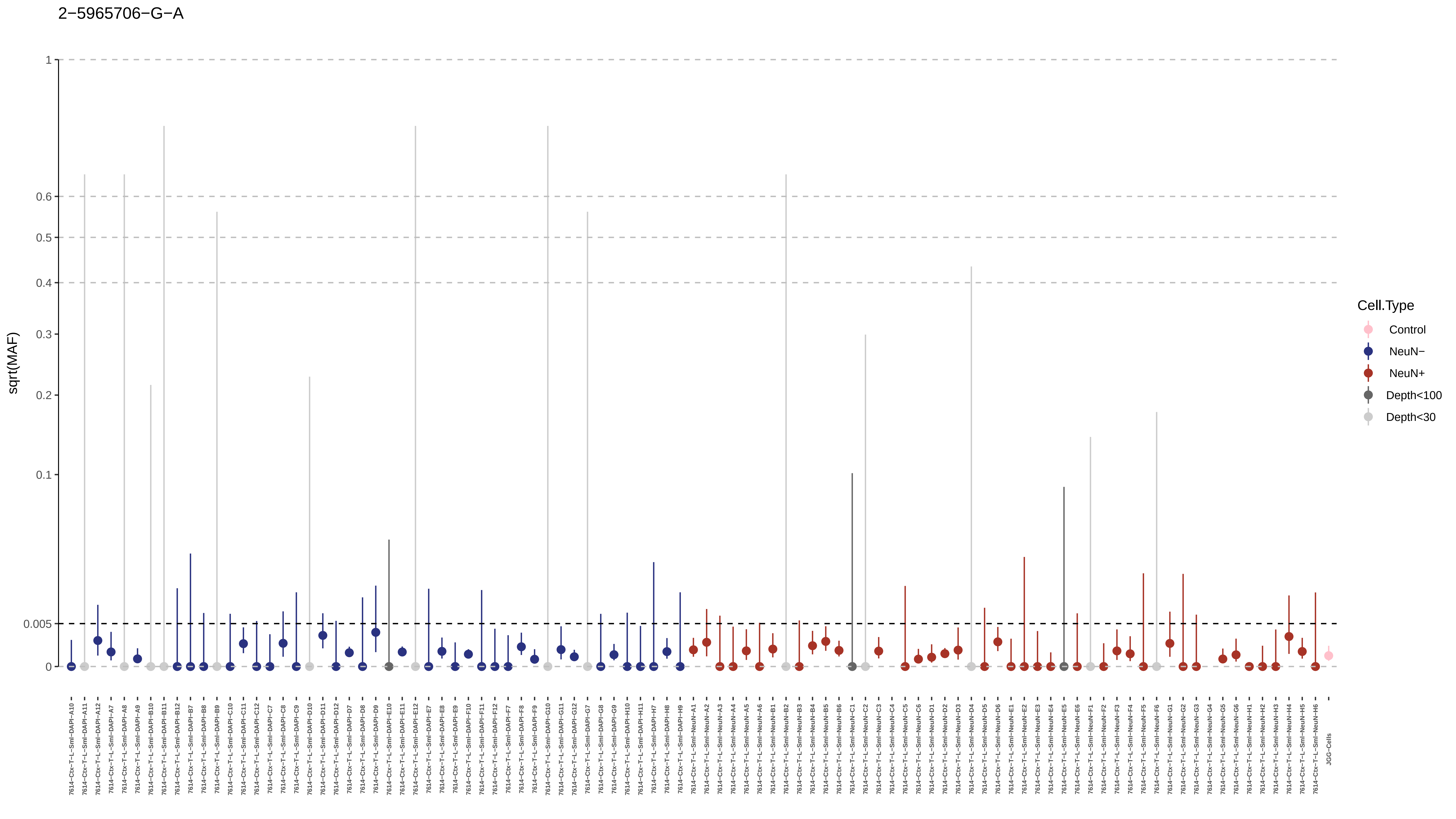

2−5965706−G−A
sqrt(MAF)
1
0.6
0.5
0.4
0.3
0.2
0.1
0.005
0
Cell.Type
Control
NeuN−
NeuN+
Depth<100
Depth<30
JGG-Cells

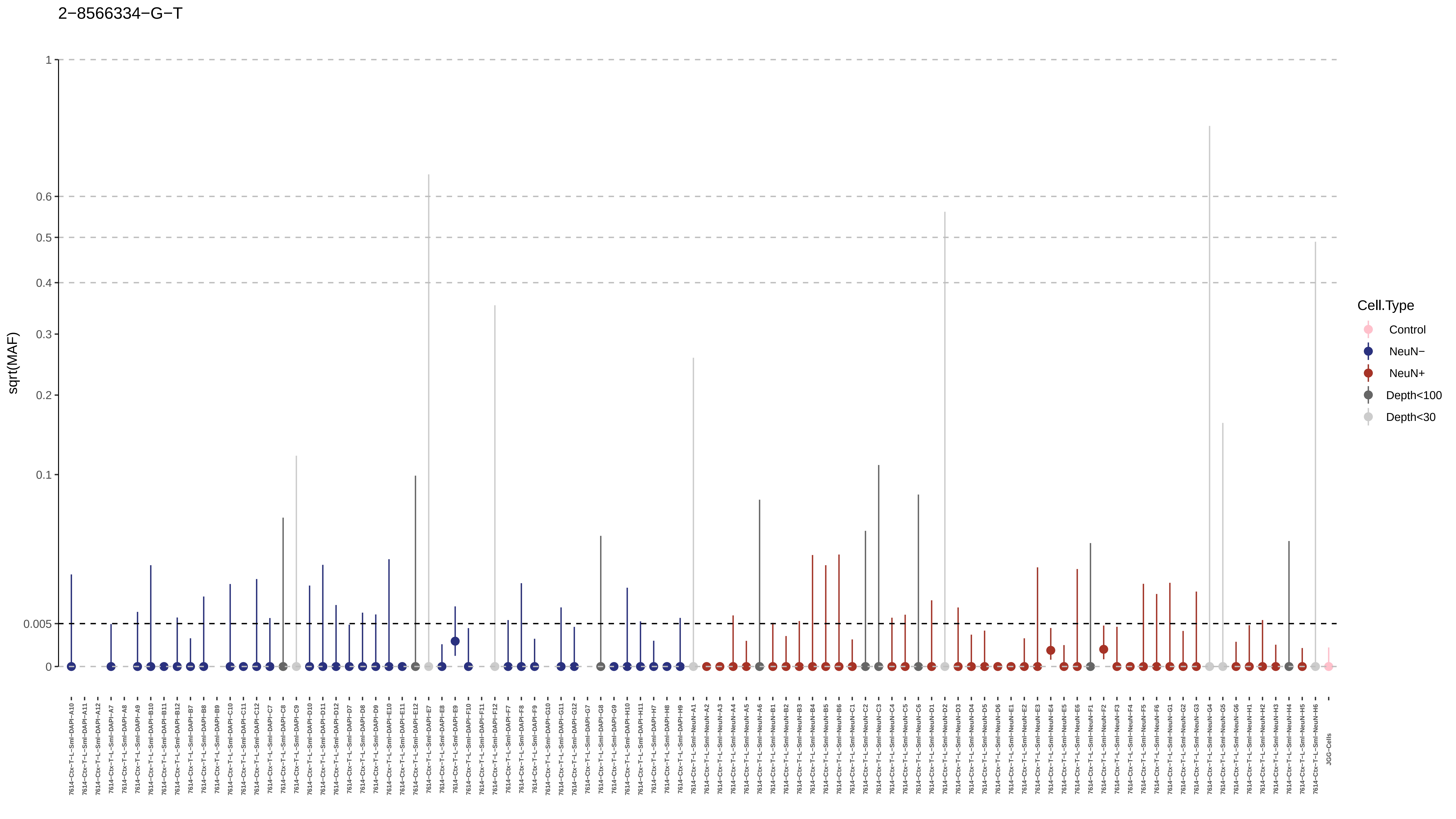

2−8566334−G−T
sqrt(MAF)
1
0.6
0.5
0.4
0.3
0.2
0.1
0.005
0
Cell.Type
Control
NeuN−
NeuN+
Depth<100
Depth<30

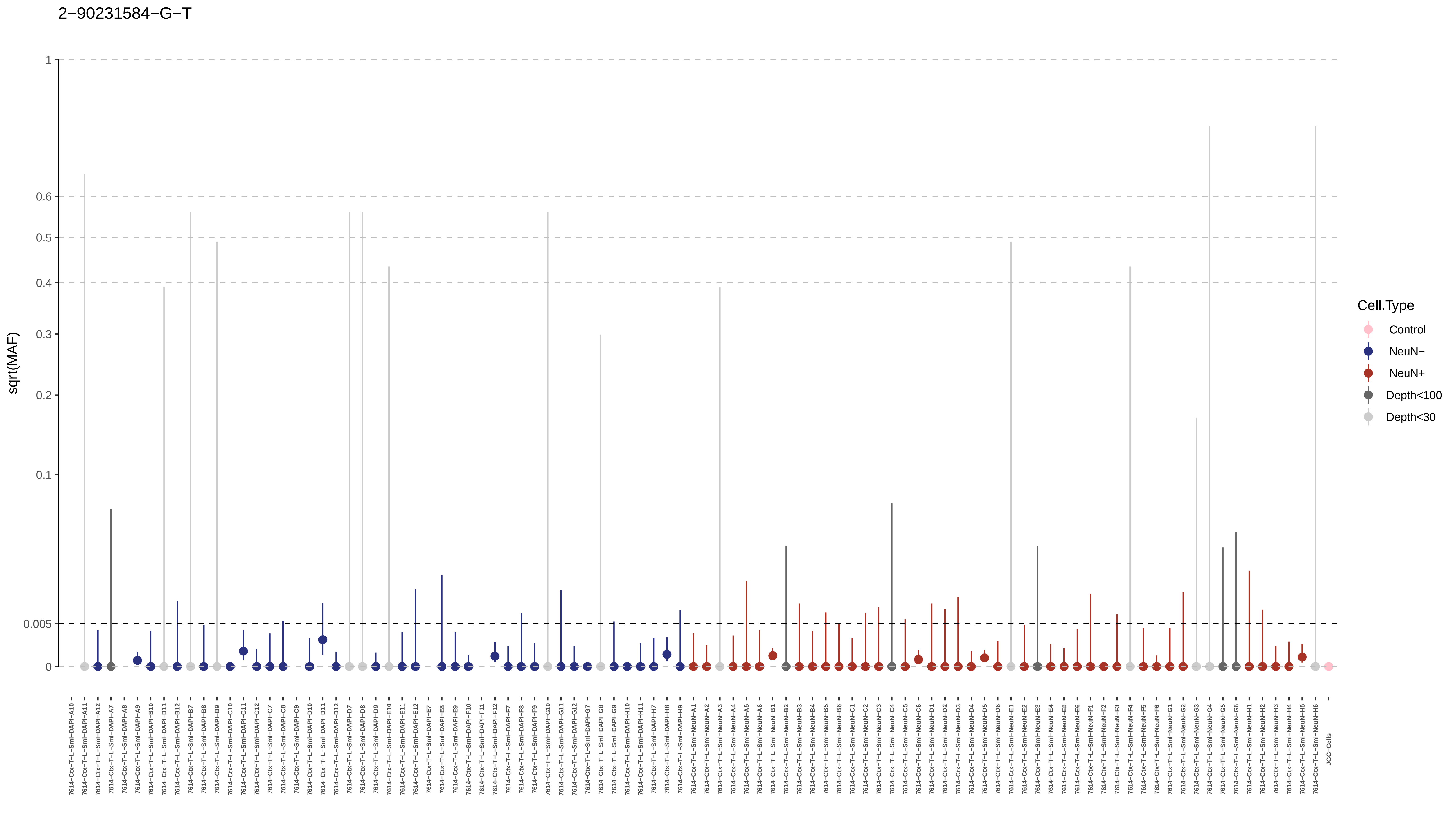
2−90231584−G−T
sqrt(MAF)
1
0.6
0.5
0.4
0.3
0.2
0.1
0.005
0
Cell.Type
Control
NeuN−
NeuN+
Depth<100
Depth<30

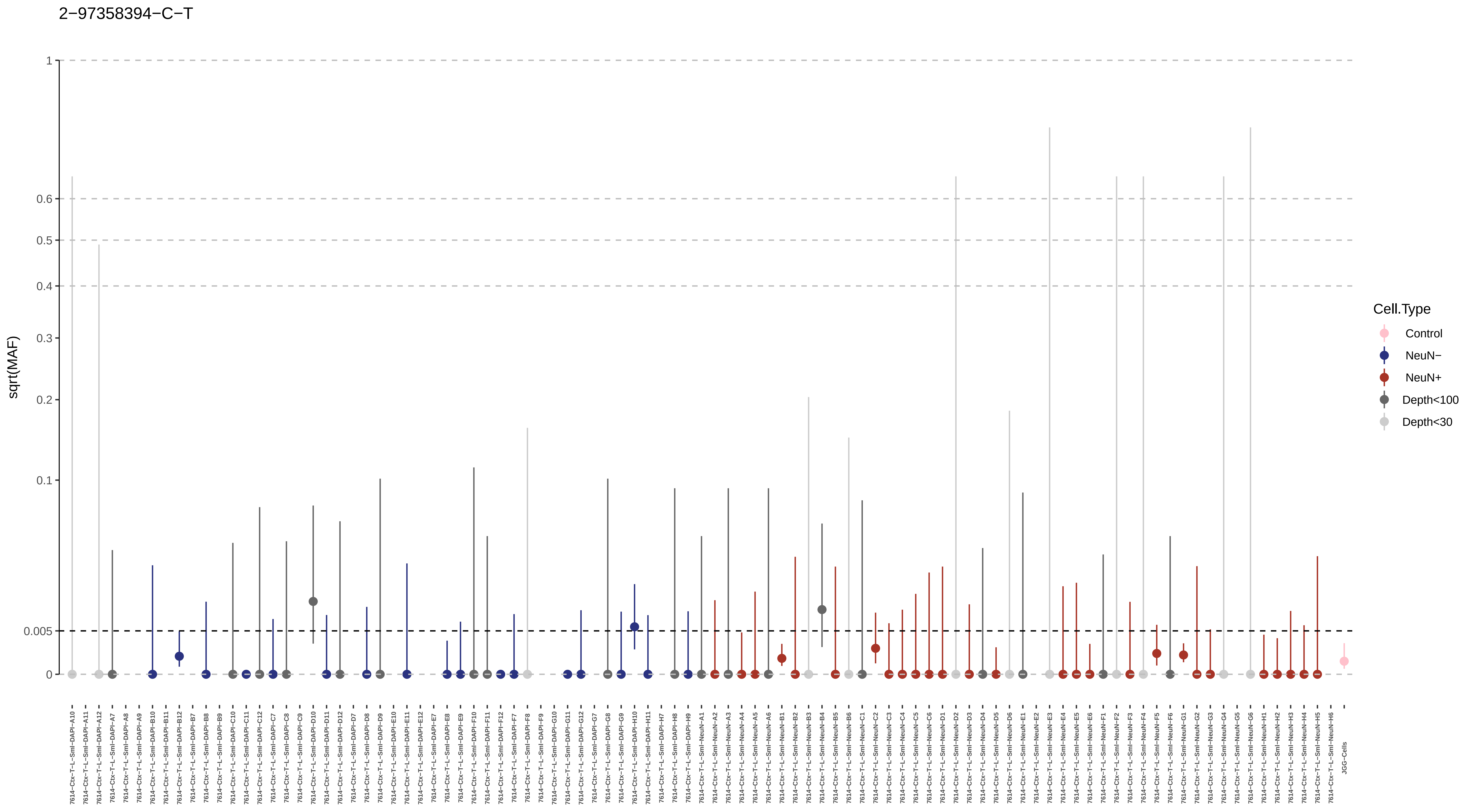

2−97358394−C−T
sqrt(MAF)
1
0.6
0.5
0.4
0.3
0.2
0.1
0.005
0
7614−Ctx−T−L−Sml−DAPI−A10
7614−Ctx−T−L−Sml−DAPI−A11
7614−Ctx−T−L−Sml−DAPI−A12
7614−Ctx−T−L−Sml−DAPI−A7
7614−Ctx−T−L−Sml−DAPI−A8
7614−Ctx−T−L−Sml−DAPI−A9
7614−Ctx−T−L−Sml−DAPI−B10
7614−Ctx−T−L−Sml−DAPI−B11
7614−Ctx−T−L−Sml−DAPI−B12
7614−Ctx−T−L−Sml−DAPI−B7
7614−Ctx−T−L−Sml−DAPI−B8
7614−Ctx−T−L−Sml−DAPI−B9
7614−Ctx−T−L−Sml−DAPI−C10
7614−Ctx−T−L−Sml−DAPI−C11
7614−Ctx−T−L−Sml−DAPI−C12
7614−Ctx−T−L−Sml−DAPI−C7
7614−Ctx−T−L−Sml−DAPI−C8
7614−Ctx−T−L−Sml−DAPI−C9
7614−Ctx−T−L−Sml−DAPI−D10
7614−Ctx−T−L−Sml−DAPI−D11
7614−Ctx−T−L−Sml−DAPI−D12
7614−Ctx−T−L−Sml−DAPI−D7
7614−Ctx−T−L−Sml−DAPI−D8
7614−Ctx−T−L−Sml−DAPI−D9
7614−Ctx−T−L−Sml−DAPI−E10
7614−Ctx−T−L−Sml−DAPI−E11
7614−Ctx−T−L−Sml−DAPI−E12
7614−Ctx−T−L−Sml−DAPI−E7
7614−Ctx−T−L−Sml−DAPI−E8
7614−Ctx−T−L−Sml−DAPI−E9
7614−Ctx−T−L−Sml−DAPI−F10
7614−Ctx−T−L−Sml−DAPI−F11
7614−Ctx−T−L−Sml−DAPI−F12
7614−Ctx−T−L−Sml−DAPI−F7
7614−Ctx−T−L−Sml−DAPI−F8
7614−Ctx−T−L−Sml−DAPI−F9
7614−Ctx−T−L−Sml−DAPI−G10
7614−Ctx−T−L−Sml−DAPI−G11
7614−Ctx−T−L−Sml−DAPI−G12
7614−Ctx−T−L−Sml−DAPI−G7
7614−Ctx−T−L−Sml−DAPI−G8
7614−Ctx−T−L−Sml−DAPI−G9
7614−Ctx−T−L−Sml−DAPI−H10
7614−Ctx−T−L−Sml−DAPI−H11
7614−Ctx−T−L−Sml−DAPI−H7
7614−Ctx−T−L−Sml−DAPI−H8
7614−Ctx−T−L−Sml−DAPI−H9
7614−Ctx−T−L−Sml−NeuN−A1
7614−Ctx−T−L−Sml−NeuN−A2
7614−Ctx−T−L−Sml−NeuN−A3
7614−Ctx−T−L−Sml−NeuN−A4
7614−Ctx−T−L−Sml−NeuN−A5
7614−Ctx−T−L−Sml−NeuN−A6
7614−Ctx−T−L−Sml−NeuN−B1
7614−Ctx−T−L−Sml−NeuN−B2
7614−Ctx−T−L−Sml−NeuN−B3
7614−Ctx−T−L−Sml−NeuN−B4
7614−Ctx−T−L−Sml−NeuN−B5
7614−Ctx−T−L−Sml−NeuN−B6
7614−Ctx−T−L−Sml−NeuN−C1
7614−Ctx−T−L−Sml−NeuN−C2
7614−Ctx−T−L−Sml−NeuN−C3
7614−Ctx−T−L−Sml−NeuN−C4
7614−Ctx−T−L−Sml−NeuN−C5
7614−Ctx−T−L−Sml−NeuN−C6
7614−Ctx−T−L−Sml−NeuN−D1
7614−Ctx−T−L−Sml−NeuN−D2
7614−Ctx−T−L−Sml−NeuN−D3
7614−Ctx−T−L−Sml−NeuN−D4
7614−Ctx−T−L−Sml−NeuN−D5
7614−Ctx−T−L−Sml−NeuN−D6
7614−Ctx−T−L−Sml−NeuN−E1
7614−Ctx−T−L−Sml−NeuN−E2
7614−Ctx−T−L−Sml−NeuN−E3
7614−Ctx−T−L−Sml−NeuN−E4
7614−Ctx−T−L−Sml−NeuN−E5
7614−Ctx−T−L−Sml−NeuN−E6
7614−Ctx−T−L−Sml−NeuN−F1
7614−Ctx−T−L−Sml−NeuN−F2
7614−Ctx−T−L−Sml−NeuN−F3
7614−Ctx−T−L−Sml−NeuN−F4
7614−Ctx−T−L−Sml−NeuN−F5
7614−Ctx−T−L−Sml−NeuN−F6
7614−Ctx−T−L−Sml−NeuN−G1
7614−Ctx−T−L−Sml−NeuN−G2
7614−Ctx−T−L−Sml−NeuN−G3
7614−Ctx−T−L−Sml−NeuN−G4
7614−Ctx−T−L−Sml−NeuN−G5
7614−Ctx−T−L−Sml−NeuN−G6
7614−Ctx−T−L−Sml−NeuN−H1
7614−Ctx−T−L−Sml−NeuN−H2
7614−Ctx−T−L−Sml−NeuN−H3
7614−Ctx−T−L−Sml−NeuN−H4
7614−Ctx−T−L−Sml−NeuN−H5
7614−Ctx−T−L−Sml−NeuN−H6
JGG-Cells
Cell.Type
Control
NeuN−
NeuN+
Depth<100
Depth<30

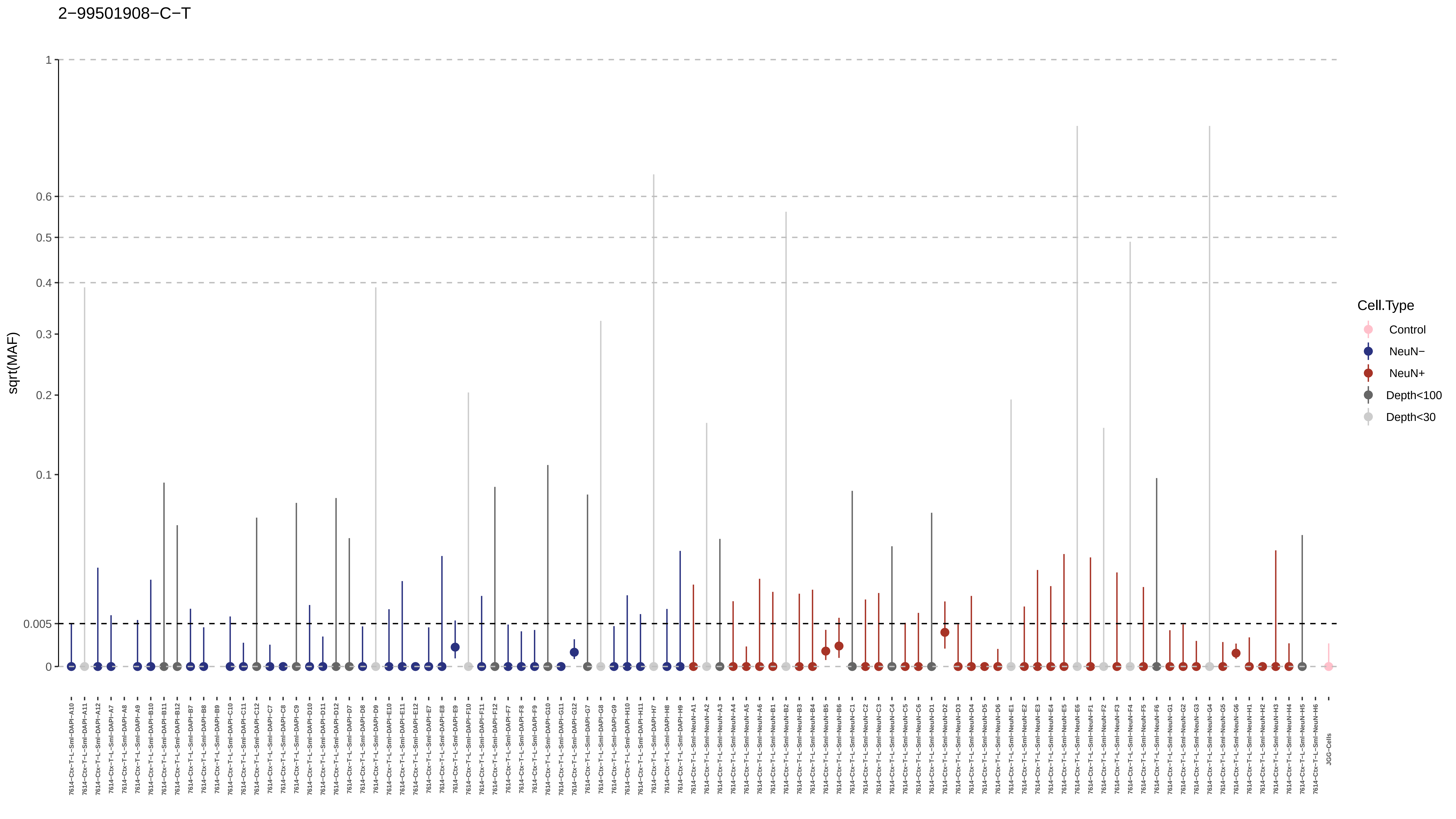
2−99501908−C−T
sqrt(MAF)
1
0.6
0.5
0.4
0.3
0.2
0.1
0.005
0
Cell.Type
Control
NeuN−
NeuN+
Depth<100
Depth<30
JGG-Cells

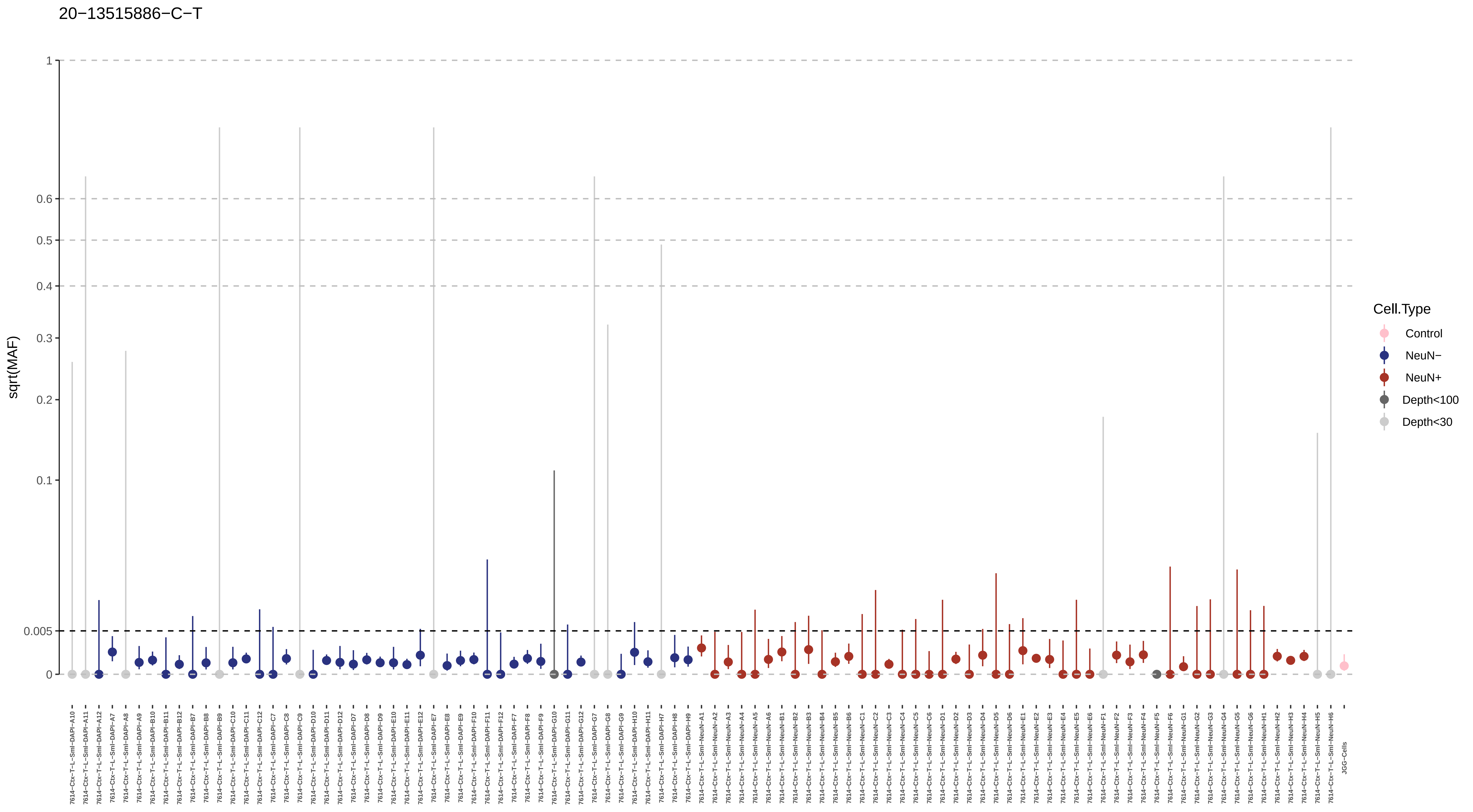
20−13515886−C−T
sqrt(MAF)
1
0.6
0.5
0.4
0.3
0.2
0.1
0.005
0
Cell.Type
Control
NeuN−
NeuN+
Depth<100
Depth<30

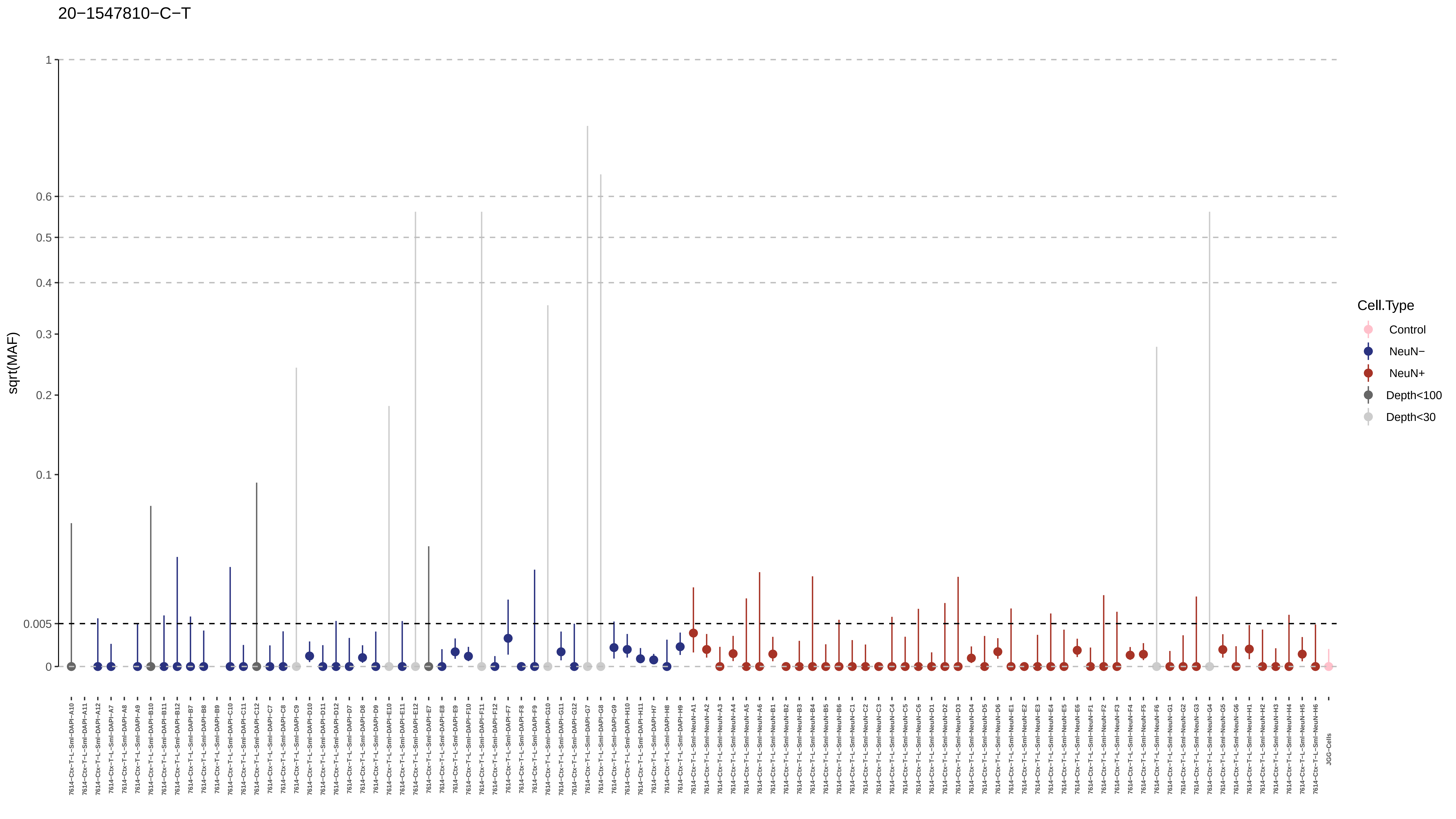
20−1547810−C−T
sqrt(MAF)
1
0.6
0.5
0.4
0.3
0.2
0.1
0.005
0
Cell.Type
Control
NeuN−
NeuN+
Depth<100
Depth<30
JGG-Cells

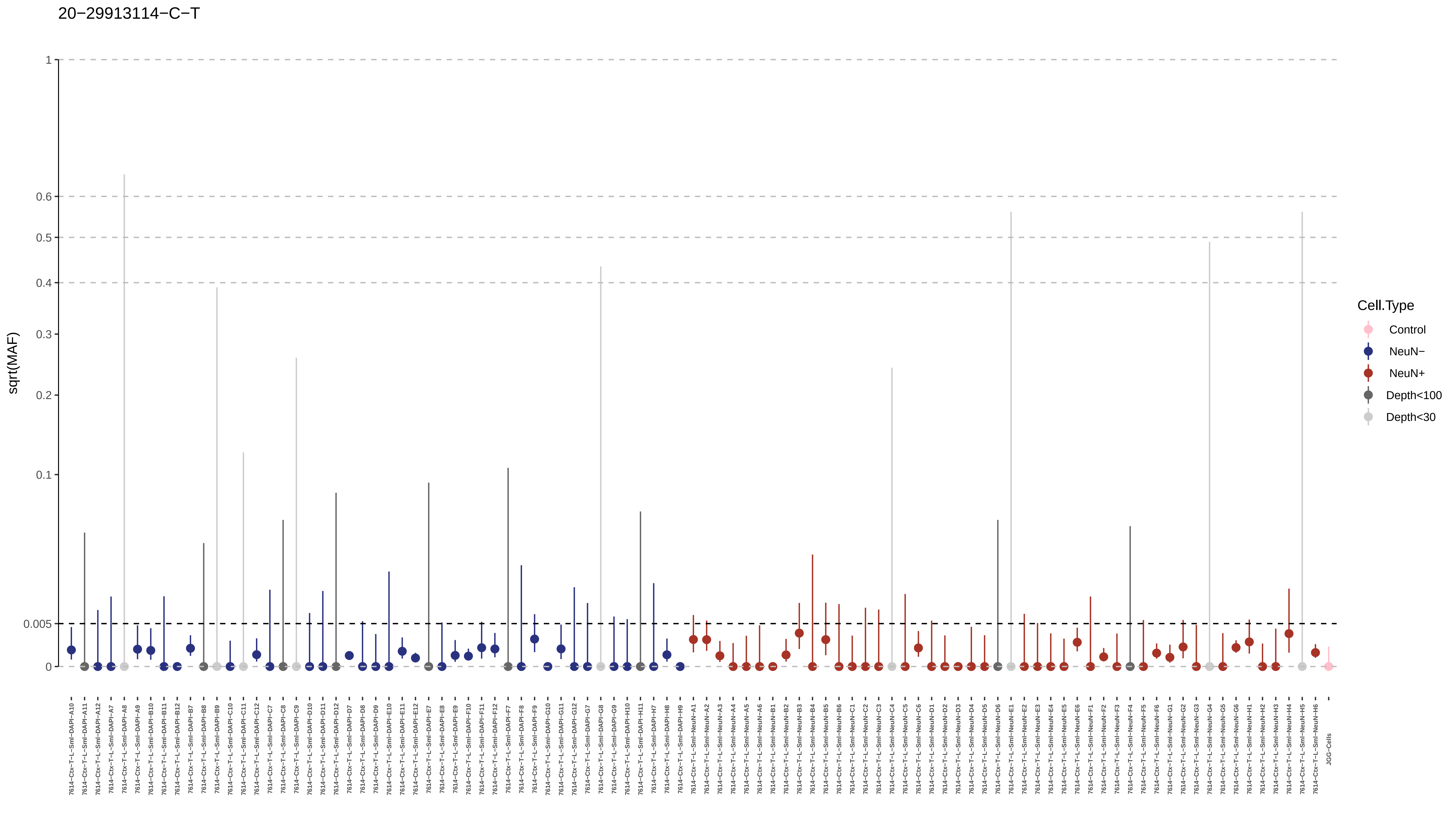

20−29913114−C−T
sqrt(MAF)
1
0.6
0.5
0.4
0.3
0.2
0.1
0.005
0
7614−Ctx−T−L−Sml−DAPI−A10
7614−Ctx−T−L−Sml−DAPI−A11
7614−Ctx−T−L−Sml−DAPI−A12
7614−Ctx−T−L−Sml−DAPI−A7
7614−Ctx−T−L−Sml−DAPI−A8
7614−Ctx−T−L−Sml−DAPI−A9
7614−Ctx−T−L−Sml−DAPI−B10
7614−Ctx−T−L−Sml−DAPI−B11
7614−Ctx−T−L−Sml−DAPI−B12
7614−Ctx−T−L−Sml−DAPI−B7
7614−Ctx−T−L−Sml−DAPI−B8
7614−Ctx−T−L−Sml−DAPI−B9
7614−Ctx−T−L−Sml−DAPI−C10
7614−Ctx−T−L−Sml−DAPI−C11
7614−Ctx−T−L−Sml−DAPI−C12
7614−Ctx−T−L−Sml−DAPI−C7
7614−Ctx−T−L−Sml−DAPI−C8
7614−Ctx−T−L−Sml−DAPI−C9
7614−Ctx−T−L−Sml−DAPI−D10
7614−Ctx−T−L−Sml−DAPI−D11
7614−Ctx−T−L−Sml−DAPI−D12
7614−Ctx−T−L−Sml−DAPI−D7
7614−Ctx−T−L−Sml−DAPI−D8
7614−Ctx−T−L−Sml−DAPI−D9
7614−Ctx−T−L−Sml−DAPI−E10
7614−Ctx−T−L−Sml−DAPI−E11
7614−Ctx−T−L−Sml−DAPI−E12
7614−Ctx−T−L−Sml−DAPI−E7
7614−Ctx−T−L−Sml−DAPI−E8
7614−Ctx−T−L−Sml−DAPI−E9
7614−Ctx−T−L−Sml−DAPI−F10
7614−Ctx−T−L−Sml−DAPI−F11
7614−Ctx−T−L−Sml−DAPI−F12
7614−Ctx−T−L−Sml−DAPI−F7
7614−Ctx−T−L−Sml−DAPI−F8
7614−Ctx−T−L−Sml−DAPI−F9
7614−Ctx−T−L−Sml−DAPI−G10
7614−Ctx−T−L−Sml−DAPI−G11
7614−Ctx−T−L−Sml−DAPI−G12
7614−Ctx−T−L−Sml−DAPI−G7
7614−Ctx−T−L−Sml−DAPI−G8
7614−Ctx−T−L−Sml−DAPI−G9
7614−Ctx−T−L−Sml−DAPI−H10
7614−Ctx−T−L−Sml−DAPI−H11
7614−Ctx−T−L−Sml−DAPI−H7
7614−Ctx−T−L−Sml−DAPI−H8
7614−Ctx−T−L−Sml−DAPI−H9
7614−Ctx−T−L−Sml−NeuN−A1
7614−Ctx−T−L−Sml−NeuN−A2
7614−Ctx−T−L−Sml−NeuN−A3
7614−Ctx−T−L−Sml−NeuN−A4
7614−Ctx−T−L−Sml−NeuN−A5
7614−Ctx−T−L−Sml−NeuN−A6
7614−Ctx−T−L−Sml−NeuN−B1
7614−Ctx−T−L−Sml−NeuN−B2
7614−Ctx−T−L−Sml−NeuN−B3
7614−Ctx−T−L−Sml−NeuN−B4
7614−Ctx−T−L−Sml−NeuN−B5
7614−Ctx−T−L−Sml−NeuN−B6
7614−Ctx−T−L−Sml−NeuN−C1
7614−Ctx−T−L−Sml−NeuN−C2
7614−Ctx−T−L−Sml−NeuN−C3
7614−Ctx−T−L−Sml−NeuN−C4
7614−Ctx−T−L−Sml−NeuN−C5
7614−Ctx−T−L−Sml−NeuN−C6
7614−Ctx−T−L−Sml−NeuN−D1
7614−Ctx−T−L−Sml−NeuN−D2
7614−Ctx−T−L−Sml−NeuN−D3
7614−Ctx−T−L−Sml−NeuN−D4
7614−Ctx−T−L−Sml−NeuN−D5
7614−Ctx−T−L−Sml−NeuN−D6
7614−Ctx−T−L−Sml−NeuN−E1
7614−Ctx−T−L−Sml−NeuN−E2
7614−Ctx−T−L−Sml−NeuN−E3
7614−Ctx−T−L−Sml−NeuN−E4
7614−Ctx−T−L−Sml−NeuN−E5
7614−Ctx−T−L−Sml−NeuN−E6
7614−Ctx−T−L−Sml−NeuN−F1
7614−Ctx−T−L−Sml−NeuN−F2
7614−Ctx−T−L−Sml−NeuN−F3
7614−Ctx−T−L−Sml−NeuN−F4
7614−Ctx−T−L−Sml−NeuN−F5
7614−Ctx−T−L−Sml−NeuN−F6
7614−Ctx−T−L−Sml−NeuN−G1
7614−Ctx−T−L−Sml−NeuN−G2
7614−Ctx−T−L−Sml−NeuN−G3
7614−Ctx−T−L−Sml−NeuN−G4
7614−Ctx−T−L−Sml−NeuN−G5
7614−Ctx−T−L−Sml−NeuN−G6
7614−Ctx−T−L−Sml−NeuN−H1
7614−Ctx−T−L−Sml−NeuN−H2
7614−Ctx−T−L−Sml−NeuN−H3
7614−Ctx−T−L−Sml−NeuN−H4
7614−Ctx−T−L−Sml−NeuN−H5
7614−Ctx−T−L−Sml−NeuN−H6
JGG−Cells
Cell.Type
Control
NeuN−
NeuN+
Depth<100
Depth<30

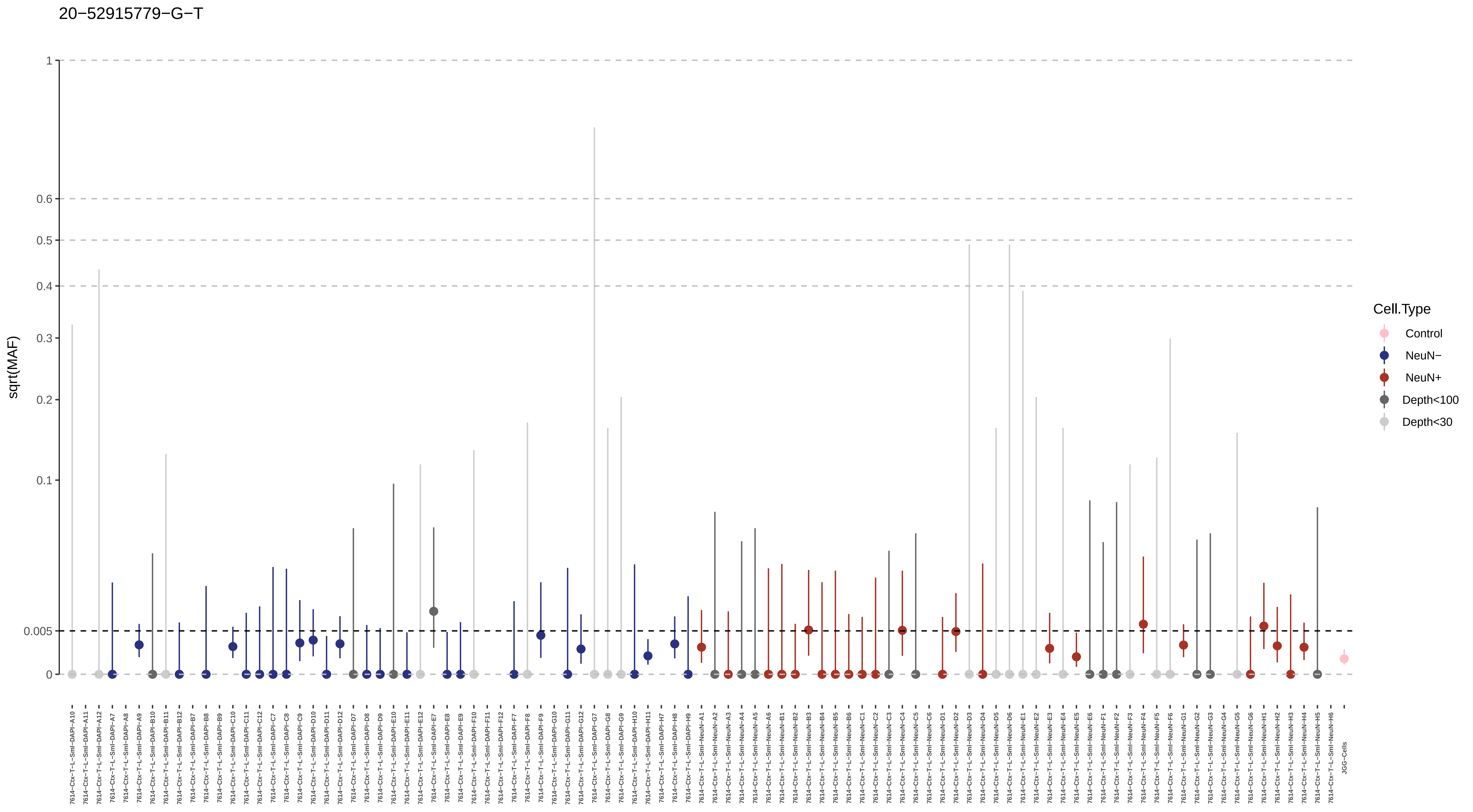
20−52915779−G−T
sqrt(MAF)
1
0.6
0.5
0.4
0.3
0.2
0.1
0.005
0
Cell.Type
Control
NeuN−
NeuN+
Depth<100
Depth<30
7614−Ctx−T−L−Sml−DAPI−A10
7614−Ctx−T−L−Sml−DAPI−A11
7614−Ctx−T−L−Sml−DAPI−A12
7614−Ctx−T−L−Sml−DAPI−A7
7614−Ctx−T−L−Sml−DAPI−A8
7614−Ctx−T−L−Sml−DAPI−A9
7614−Ctx−T−L−Sml−DAPI−B10
7614−Ctx−T−L−Sml−DAPI−B11
7614−Ctx−T−L−Sml−DAPI−B12
7614−Ctx−T−L−Sml−DAPI−B7
7614−Ctx−T−L−Sml−DAPI−B8
7614−Ctx−T−L−Sml−DAPI−B9
7614−Ctx−T−L−Sml−DAPI−C10
7614−Ctx−T−L−Sml−DAPI−C11
7614−Ctx−T−L−Sml−DAPI−C12
7614−Ctx−T−L−Sml−DAPI−C7
7614−Ctx−T−L−Sml−DAPI−C8
7614−Ctx−T−L−Sml−DAPI−C9
7614−Ctx−T−L−Sml−DAPI−D10
7614−Ctx−T−L−Sml−DAPI−D11
7614−Ctx−T−L−Sml−DAPI−D12
7614−Ctx−T−L−Sml−DAPI−D7
7614−Ctx−T−L−Sml−DAPI−D8
7614−Ctx−T−L−Sml−DAPI−D9
7614−Ctx−T−L−Sml−DAPI−E10
7614−Ctx−T−L−Sml−DAPI−E11
7614−Ctx−T−L−Sml−DAPI−E12
7614−Ctx−T−L−Sml−DAPI−E7
7614−Ctx−T−L−Sml−DAPI−E8
7614−Ctx−T−L−Sml−DAPI−E9
7614−Ctx−T−L−Sml−DAPI−F10
7614−Ctx−T−L−Sml−DAPI−F11
7614−Ctx−T−L−Sml−DAPI−F12
7614−Ctx−T−L−Sml−DAPI−F7
7614−Ctx−T−L−Sml−DAPI−F8
7614−Ctx−T−L−Sml−DAPI−F9
7614−Ctx−T−L−Sml−DAPI−G10
7614−Ctx−T−L−Sml−DAPI−G11
7614−Ctx−T−L−Sml−DAPI−G12
7614−Ctx−T−L−Sml−DAPI−G7
7614−Ctx−T−L−Sml−DAPI−G8
7614−Ctx−T−L−Sml−DAPI−G9
7614−Ctx−T−L−Sml−DAPI−H10
7614−Ctx−T−L−Sml−DAPI−H11
7614−Ctx−T−L−Sml−DAPI−H7
7614−Ctx−T−L−Sml−DAPI−H8
7614−Ctx−T−L−Sml−DAPI−H9
7614−Ctx−T−L−Sml−NeuN−A1
7614−Ctx−T−L−Sml−NeuN−A2
7614−Ctx−T−L−Sml−NeuN−A3
7614−Ctx−T−L−Sml−NeuN−A4
7614−Ctx−T−L−Sml−NeuN−A5
7614−Ctx−T−L−Sml−NeuN−A6
7614−Ctx−T−L−Sml−NeuN−B1
7614−Ctx−T−L−Sml−NeuN−B2
7614−Ctx−T−L−Sml−NeuN−B3
7614−Ctx−T−L−Sml−NeuN−B4
7614−Ctx−T−L−Sml−NeuN−B5
7614−Ctx−T−L−Sml−NeuN−B6
7614−Ctx−T−L−Sml−NeuN−C1
7614−Ctx−T−L−Sml−NeuN−C2
7614−Ctx−T−L−Sml−NeuN−C3
7614−Ctx−T−L−Sml−NeuN−C4
7614−Ctx−T−L−Sml−NeuN−C5
7614−Ctx−T−L−Sml−NeuN−C6
7614−Ctx−T−L−Sml−NeuN−D1
7614−Ctx−T−L−Sml−NeuN−D2
7614−Ctx−T−L−Sml−NeuN−D3
7614−Ctx−T−L−Sml−NeuN−D4
7614−Ctx−T−L−Sml−NeuN−D5
7614−Ctx−T−L−Sml−NeuN−D6
7614−Ctx−T−L−Sml−NeuN−E1
7614−Ctx−T−L−Sml−NeuN−E2
7614−Ctx−T−L−Sml−NeuN−E3
7614−Ctx−T−L−Sml−NeuN−E4
7614−Ctx−T−L−Sml−NeuN−E5
7614−Ctx−T−L−Sml−NeuN−E6
7614−Ctx−T−L−Sml−NeuN−F1
7614−Ctx−T−L−Sml−NeuN−F2
7614−Ctx−T−L−Sml−NeuN−F3
7614−Ctx−T−L−Sml−NeuN−F4
7614−Ctx−T−L−Sml−NeuN−F5
7614−Ctx−T−L−Sml−NeuN−F6
7614−Ctx−T−L−Sml−NeuN−G1
7614−Ctx−T−L−Sml−NeuN−G2
7614−Ctx−T−L−Sml−NeuN−G3
7614−Ctx−T−L−Sml−NeuN−G4
7614−Ctx−T−L−Sml−NeuN−G5
7614−Ctx−T−L−Sml−NeuN−G6
7614−Ctx−T−L−Sml−NeuN−H1
7614−Ctx−T−L−Sml−NeuN−H2
7614−Ctx−T−L−Sml−NeuN−H3
7614−Ctx−T−L−Sml−NeuN−H4
7614−Ctx−T−L−Sml−NeuN−H5
7614−Ctx−T−L−Sml−NeuN−H6
JGG−Cells

20−52916352−A−C

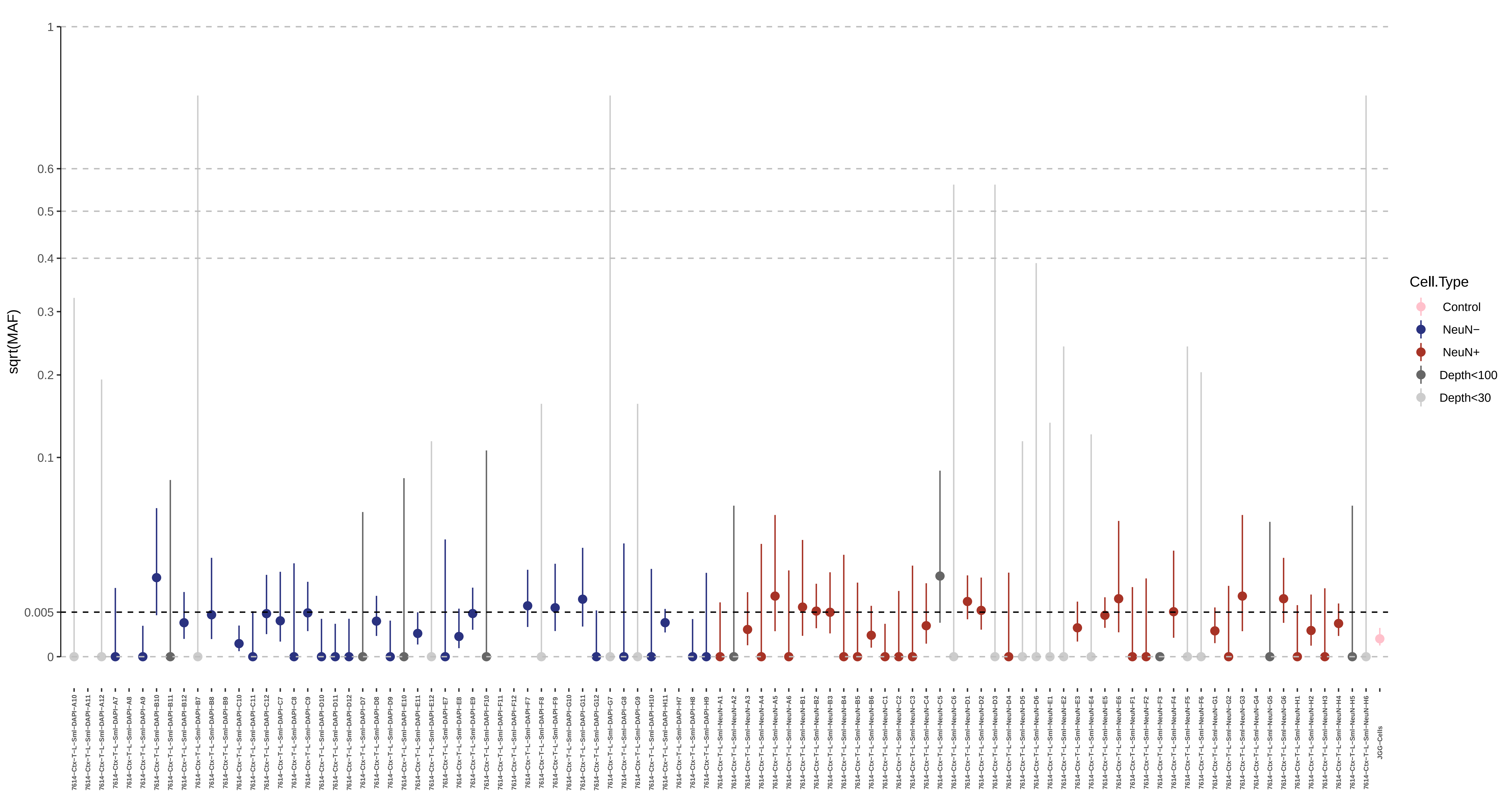

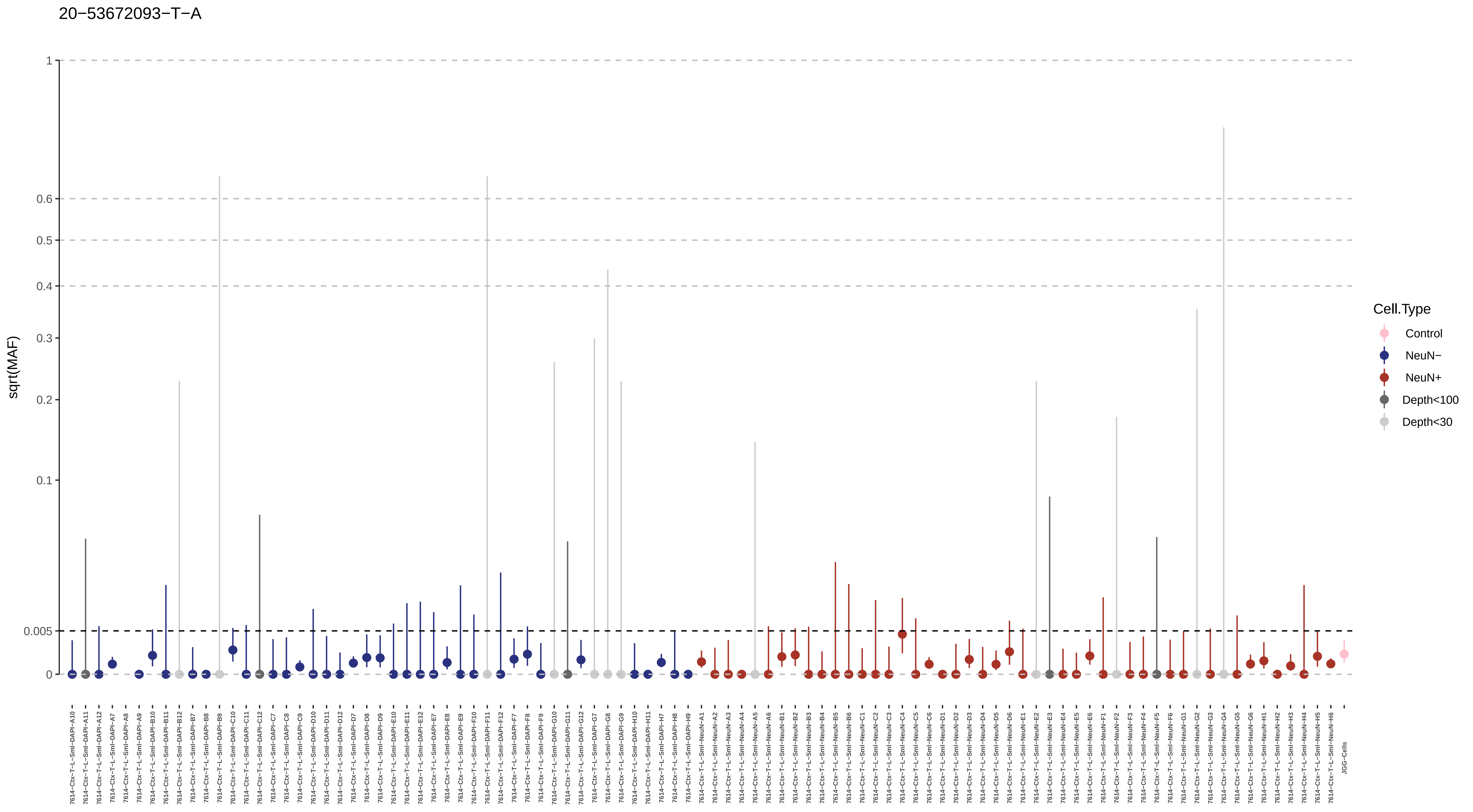

20−53672093−T−A
sqrt(MAF)
1
0.6
0.5
0.4
0.3
0.2
0.1
0.005
0
Cell.Type
Control
NeuN−
NeuN+
Depth<100
Depth<30

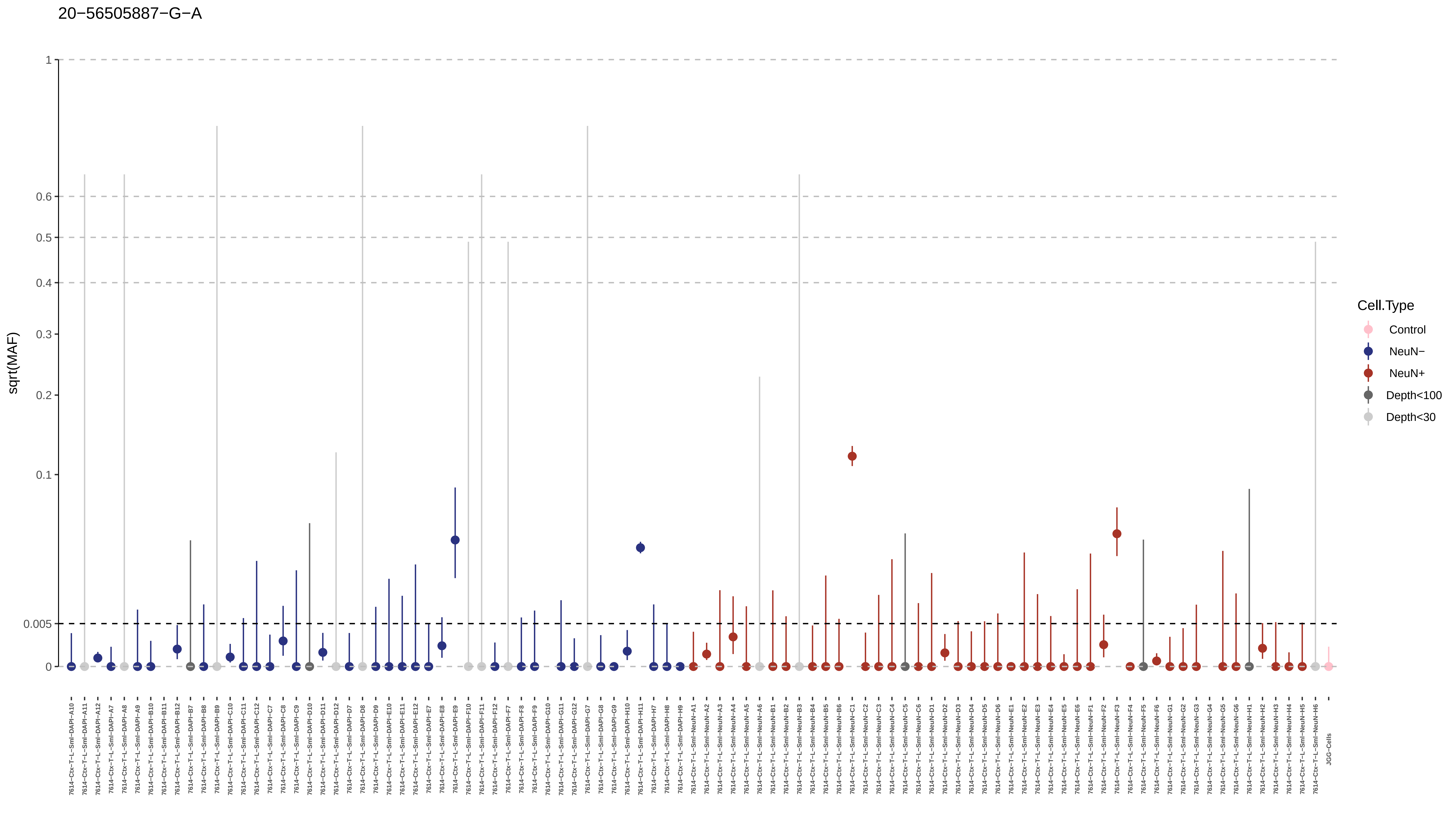

20−56505887−G−A
sqrt(MAF)
1
0.6
0.5
0.4
0.3
0.2
0.1
0.005
0
Cell.Type
Control
NeuN−
NeuN+
Depth<100
Depth<30
7614−Ctx−T−L−Sml−DAPI−A10
7614−Ctx−T−L−Sml−DAPI−A11
7614−Ctx−T−L−Sml−DAPI−A12
7614−Ctx−T−L−Sml−DAPI−A7
7614−Ctx−T−L−Sml−DAPI−A8
7614−Ctx−T−L−Sml−DAPI−A9
7614−Ctx−T−L−Sml−DAPI−B10
7614−Ctx−T−L−Sml−DAPI−B11
7614−Ctx−T−L−Sml−DAPI−B12
7614−Ctx−T−L−Sml−DAPI−B7
7614−Ctx−T−L−Sml−DAPI−B8
7614−Ctx−T−L−Sml−DAPI−B9
7614−Ctx−T−L−Sml−DAPI−C10
7614−Ctx−T−L−Sml−DAPI−C11
7614−Ctx−T−L−Sml−DAPI−C12
7614−Ctx−T−L−Sml−DAPI−C7
7614−Ctx−T−L−Sml−DAPI−C8
7614−Ctx−T−L−Sml−DAPI−C9
7614−Ctx−T−L−Sml−DAPI−D10
7614−Ctx−T−L−Sml−DAPI−D11
7614−Ctx−T−L−Sml−DAPI−D12
7614−Ctx−T−L−Sml−DAPI−D7
7614−Ctx−T−L−Sml−DAPI−D8
7614−Ctx−T−L−Sml−DAPI−D9
7614−Ctx−T−L−Sml−DAPI−E10
7614−Ctx−T−L−Sml−DAPI−E11
7614−Ctx−T−L−Sml−DAPI−E12
7614−Ctx−T−L−Sml−DAPI−E7
7614−Ctx−T−L−Sml−DAPI−E8
7614−Ctx−T−L−Sml−DAPI−E9
7614−Ctx−T−L−Sml−DAPI−F10
7614−Ctx−T−L−Sml−DAPI−F11
7614−Ctx−T−L−Sml−DAPI−F12
7614−Ctx−T−L−Sml−DAPI−F7
7614−Ctx−T−L−Sml−DAPI−F8
7614−Ctx−T−L−Sml−DAPI−F9
7614−Ctx−T−L−Sml−DAPI−G10
7614−Ctx−T−L−Sml−DAPI−G11
7614−Ctx−T−L−Sml−DAPI−G12
7614−Ctx−T−L−Sml−DAPI−G7
7614−Ctx−T−L−Sml−DAPI−G8
7614−Ctx−T−L−Sml−DAPI−G9
7614−Ctx−T−L−Sml−DAPI−H10
7614−Ctx−T−L−Sml−DAPI−H11
7614−Ctx−T−L−Sml−DAPI−H7
7614−Ctx−T−L−Sml−DAPI−H8
7614−Ctx−T−L−Sml−DAPI−H9
7614−Ctx−T−L−Sml−NeuN−A1
7614−Ctx−T−L−Sml−NeuN−A2
7614−Ctx−T−L−Sml−NeuN−A3
7614−Ctx−T−L−Sml−NeuN−A4
7614−Ctx−T−L−Sml−NeuN−A5
7614−Ctx−T−L−Sml−NeuN−A6
7614−Ctx−T−L−Sml−NeuN−B1
7614−Ctx−T−L−Sml−NeuN−B2
7614−Ctx−T−L−Sml−NeuN−B3
7614−Ctx−T−L−Sml−NeuN−B4
7614−Ctx−T−L−Sml−NeuN−B5
7614−Ctx−T−L−Sml−NeuN−B6
7614−Ctx−T−L−Sml−NeuN−C1
7614−Ctx−T−L−Sml−NeuN−C2
7614−Ctx−T−L−Sml−NeuN−C3
7614−Ctx−T−L−Sml−NeuN−C4
7614−Ctx−T−L−Sml−NeuN−C5
7614−Ctx−T−L−Sml−NeuN−C6
7614−Ctx−T−L−Sml−NeuN−D1
7614−Ctx−T−L−Sml−NeuN−D2
7614−Ctx−T−L−Sml−NeuN−D3
7614−Ctx−T−L−Sml−NeuN−D4
7614−Ctx−T−L−Sml−NeuN−D5
7614−Ctx−T−L−Sml−NeuN−D6
7614−Ctx−T−L−Sml−NeuN−E1
7614−Ctx−T−L−Sml−NeuN−E2
7614−Ctx−T−L−Sml−NeuN−E3
7614−Ctx−T−L−Sml−NeuN−E4
7614−Ctx−T−L−Sml−NeuN−E5
7614−Ctx−T−L−Sml−NeuN−E6
7614−Ctx−T−L−Sml−NeuN−F1
7614−Ctx−T−L−Sml−NeuN−F2
7614−Ctx−T−L−Sml−NeuN−F3
7614−Ctx−T−L−Sml−NeuN−F4
7614−Ctx−T−L−Sml−NeuN−F5
7614−Ctx−T−L−Sml−NeuN−F6
7614−Ctx−T−L−Sml−NeuN−G1
7614−Ctx−T−L−Sml−NeuN−G2
7614−Ctx−T−L−Sml−NeuN−G3
7614−Ctx−T−L−Sml−NeuN−G4
7614−Ctx−T−L−Sml−NeuN−G5
7614−Ctx−T−L−Sml−NeuN−G6
7614−Ctx−T−L−Sml−NeuN−H1
7614−Ctx−T−L−Sml−NeuN−H2
7614−Ctx−T−L−Sml−NeuN−H3
7614−Ctx−T−L−Sml−NeuN−H4
7614−Ctx−T−L−Sml−NeuN−H5
7614−Ctx−T−L−Sml−NeuN−H6
JGG−Cells

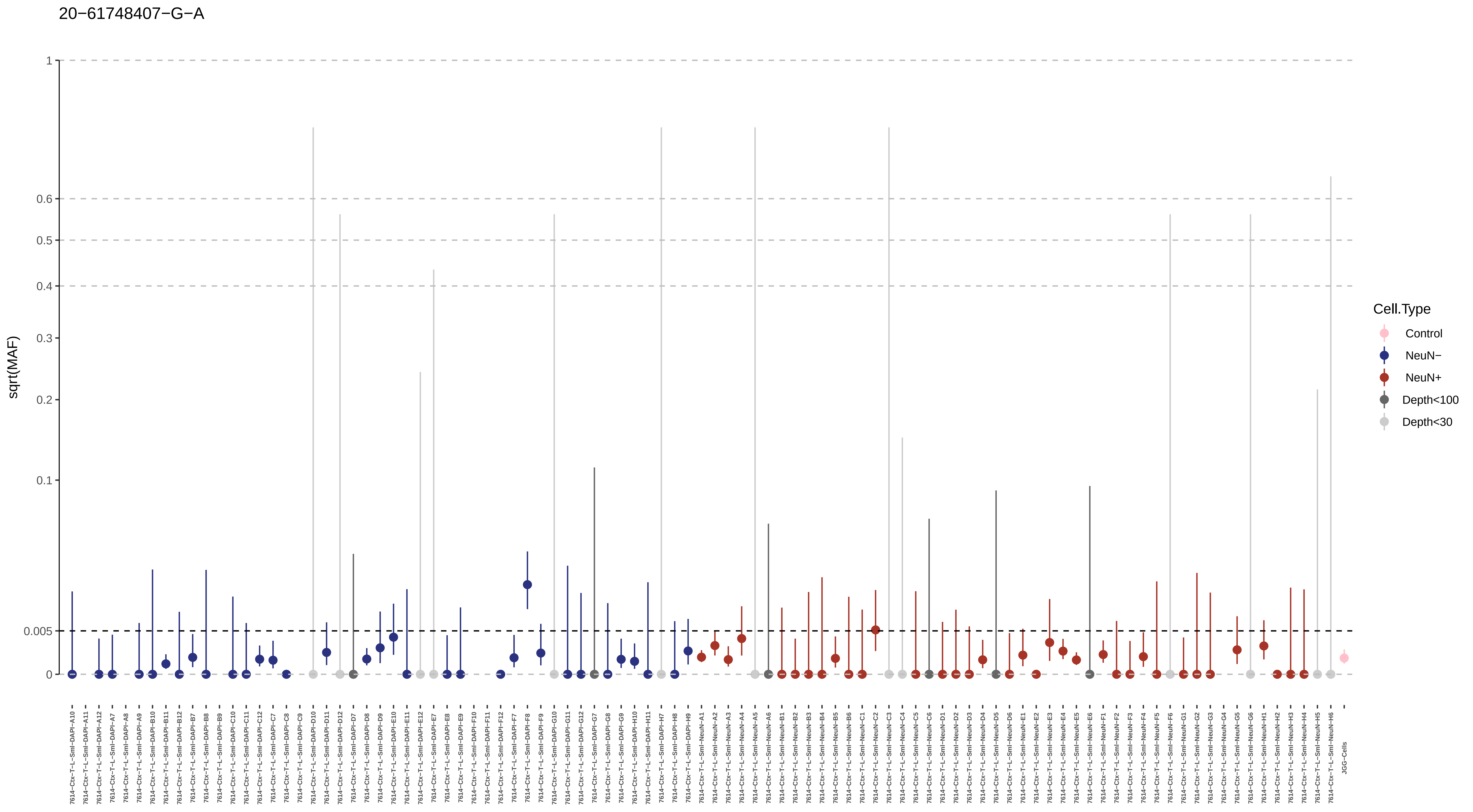
20−61748407−G−A
sqrt(MAF)
1
0.6
0.5
0.4
0.3
0.2
0.1
0.005
0
Cell.Type
Control
NeuN−
NeuN+
Depth<100
Depth<30
7614−Ctx−T−L−Sml−DAPI−A10
7614−Ctx−T−L−Sml−DAPI−A11
7614−Ctx−T−L−Sml−DAPI−A12
7614−Ctx−T−L−Sml−DAPI−A7
7614−Ctx−T−L−Sml−DAPI−A8
7614−Ctx−T−L−Sml−DAPI−A9
7614−Ctx−T−L−Sml−DAPI−B10
7614−Ctx−T−L−Sml−DAPI−B11
7614−Ctx−T−L−Sml−DAPI−B12
7614−Ctx−T−L−Sml−DAPI−B7
7614−Ctx−T−L−Sml−DAPI−B8
7614−Ctx−T−L−Sml−DAPI−B9
7614−Ctx−T−L−Sml−DAPI−C10
7614−Ctx−T−L−Sml−DAPI−C11
7614−Ctx−T−L−Sml−DAPI−C12
7614−Ctx−T−L−Sml−DAPI−C7
7614−Ctx−T−L−Sml−DAPI−C8
7614−Ctx−T−L−Sml−DAPI−C9
7614−Ctx−T−L−Sml−DAPI−D10
7614−Ctx−T−L−Sml−DAPI−D11
7614−Ctx−T−L−Sml−DAPI−D12
7614−Ctx−T−L−Sml−DAPI−D7
7614−Ctx−T−L−Sml−DAPI−D8
7614−Ctx−T−L−Sml−DAPI−D9
7614−Ctx−T−L−Sml−DAPI−E10
7614−Ctx−T−L−Sml−DAPI−E11
7614−Ctx−T−L−Sml−DAPI−E12
7614−Ctx−T−L−Sml−DAPI−E7
7614−Ctx−T−L−Sml−DAPI−E8
7614−Ctx−T−L−Sml−DAPI−E9
7614−Ctx−T−L−Sml−DAPI−F10
7614−Ctx−T−L−Sml−DAPI−F11
7614−Ctx−T−L−Sml−DAPI−F12
7614−Ctx−T−L−Sml−DAPI−F7
7614−Ctx−T−L−Sml−DAPI−F8
7614−Ctx−T−L−Sml−DAPI−F9
7614−Ctx−T−L−Sml−DAPI−G10
7614−Ctx−T−L−Sml−DAPI−G11
7614−Ctx−T−L−Sml−DAPI−G12
7614−Ctx−T−L−Sml−DAPI−G7
7614−Ctx−T−L−Sml−DAPI−G8
7614−Ctx−T−L−Sml−DAPI−G9
7614−Ctx−T−L−Sml−DAPI−H10
7614−Ctx−T−L−Sml−DAPI−H11
7614−Ctx−T−L−Sml−DAPI−H7
7614−Ctx−T−L−Sml−DAPI−H8
7614−Ctx−T−L−Sml−DAPI−H9
7614−Ctx−T−L−Sml−NeuN−A1
7614−Ctx−T−L−Sml−NeuN−A2
7614−Ctx−T−L−Sml−NeuN−A3
7614−Ctx−T−L−Sml−NeuN−A4
7614−Ctx−T−L−Sml−NeuN−A5
7614−Ctx−T−L−Sml−NeuN−A6
7614−Ctx−T−L−Sml−NeuN−B1
7614−Ctx−T−L−Sml−NeuN−B2
7614−Ctx−T−L−Sml−NeuN−B3
7614−Ctx−T−L−Sml−NeuN−B4
7614−Ctx−T−L−Sml−NeuN−B5
7614−Ctx−T−L−Sml−NeuN−B6
7614−Ctx−T−L−Sml−NeuN−C1
7614−Ctx−T−L−Sml−NeuN−C2
7614−Ctx−T−L−Sml−NeuN−C3
7614−Ctx−T−L−Sml−NeuN−C4
7614−Ctx−T−L−Sml−NeuN−C5
7614−Ctx−T−L−Sml−NeuN−C6
7614−Ctx−T−L−Sml−NeuN−D1
7614−Ctx−T−L−Sml−NeuN−D2
7614−Ctx−T−L−Sml−NeuN−D3
7614−Ctx−T−L−Sml−NeuN−D4
7614−Ctx−T−L−Sml−NeuN−D5
7614−Ctx−T−L−Sml−NeuN−D6
7614−Ctx−T−L−Sml−NeuN−E1
7614−Ctx−T−L−Sml−NeuN−E2
7614−Ctx−T−L−Sml−NeuN−E3
7614−Ctx−T−L−Sml−NeuN−E4
7614−Ctx−T−L−Sml−NeuN−E5
7614−Ctx−T−L−Sml−NeuN−E6
7614−Ctx−T−L−Sml−NeuN−F1
7614−Ctx−T−L−Sml−NeuN−F2
7614−Ctx−T−L−Sml−NeuN−F3
7614−Ctx−T−L−Sml−NeuN−F4
7614−Ctx−T−L−Sml−NeuN−F5
7614−Ctx−T−L−Sml−NeuN−F6
7614−Ctx−T−L−Sml−NeuN−G1
7614−Ctx−T−L−Sml−NeuN−G2
7614−Ctx−T−L−Sml−NeuN−G3
7614−Ctx−T−L−Sml−NeuN−G4
7614−Ctx−T−L−Sml−NeuN−G5
7614−Ctx−T−L−Sml−NeuN−G6
7614−Ctx−T−L−Sml−NeuN−H1
7614−Ctx−T−L−Sml−NeuN−H2
7614−Ctx−T−L−Sml−NeuN−H3
7614−Ctx−T−L−Sml−NeuN−H4
7614−Ctx−T−L−Sml−NeuN−H5
7614−Ctx−T−L−Sml−NeuN−H6
JGG-Cells

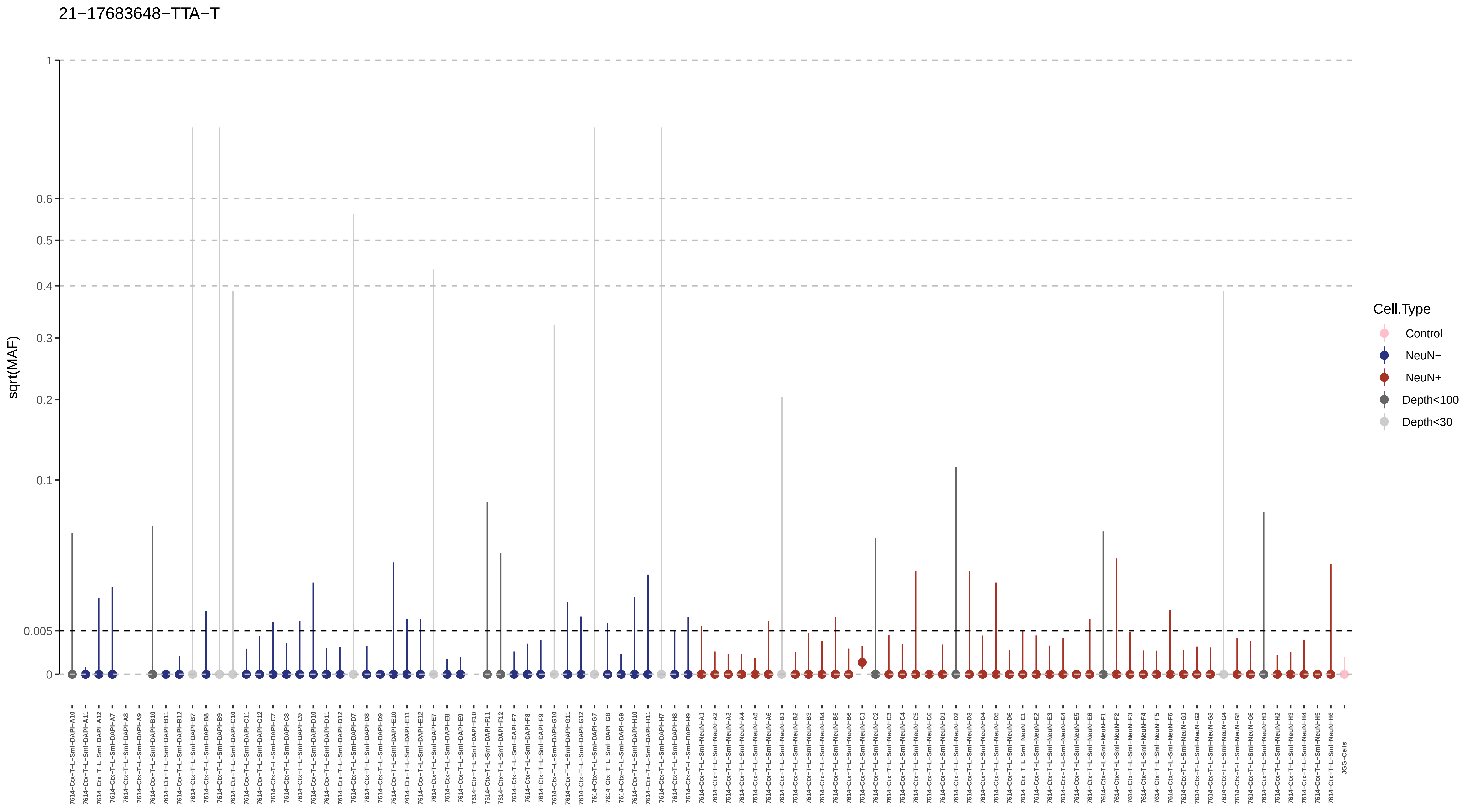

21−17683648−TTA−T
sqrt(MAF)
1
0.6
0.5
0.4
0.3
0.2
0.1
0.005
0
Cell.Type
Control
NeuN−
NeuN+
Depth<100
Depth<30
JGG-Cells

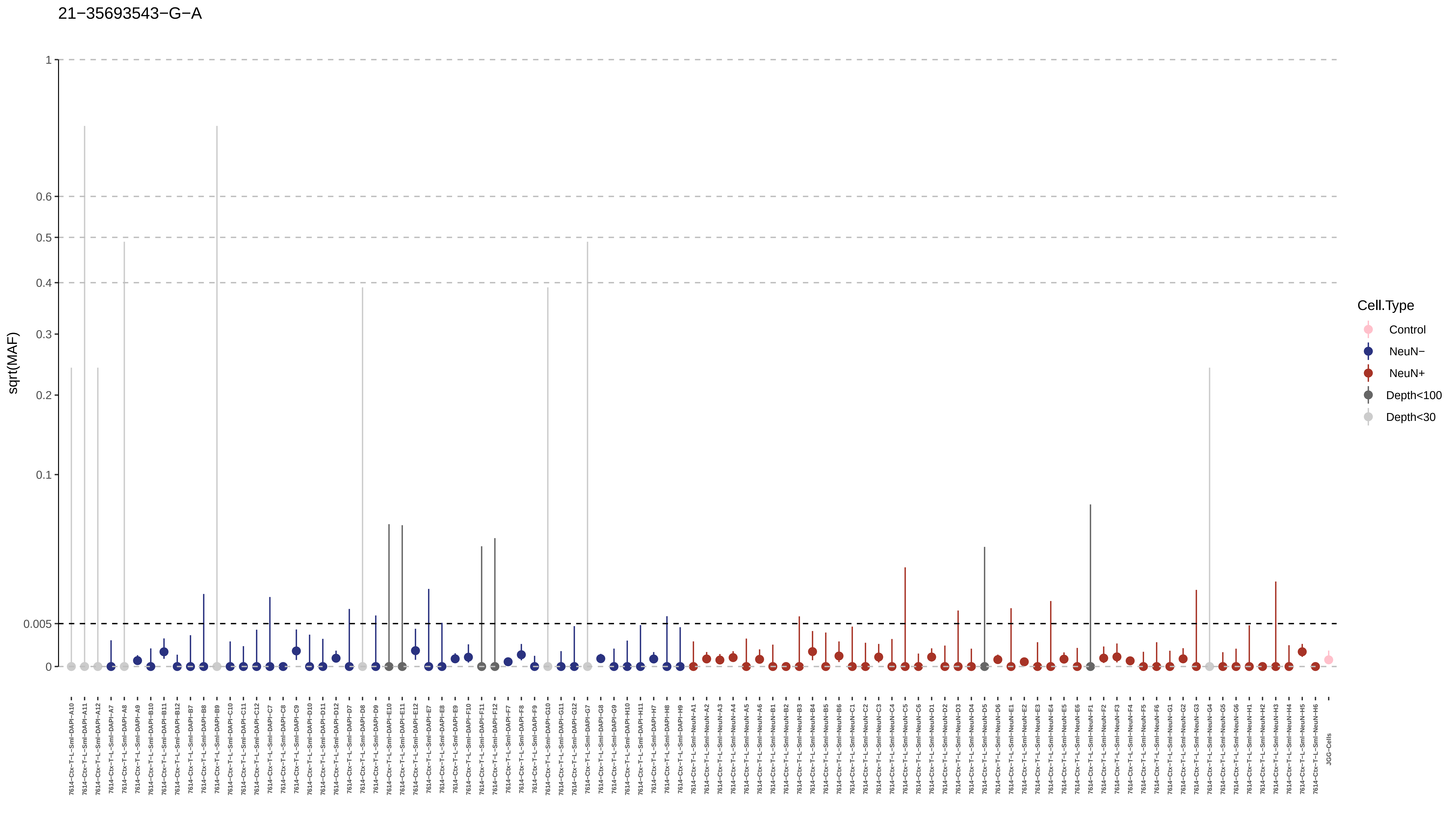

21−35693543−G−A
sqrt(MAF)
1
0.6
0.5
0.4
0.3
0.2
0.1
0.005
0
Cell.Type
Control
NeuN−
NeuN+
Depth<100
Depth<30
7614−Ctx−T−L−Sml−DAPI−A10
7614−Ctx−T−L−Sml−DAPI−A11
7614−Ctx−T−L−Sml−DAPI−A12
7614−Ctx−T−L−Sml−DAPI−A7
7614−Ctx−T−L−Sml−DAPI−A8
7614−Ctx−T−L−Sml−DAPI−A9
7614−Ctx−T−L−Sml−DAPI−B10
7614−Ctx−T−L−Sml−DAPI−B11
7614−Ctx−T−L−Sml−DAPI−B12
7614−Ctx−T−L−Sml−DAPI−B7
7614−Ctx−T−L−Sml−DAPI−B8
7614−Ctx−T−L−Sml−DAPI−B9
7614−Ctx−T−L−Sml−DAPI−C10
7614−Ctx−T−L−Sml−DAPI−C11
7614−Ctx−T−L−Sml−DAPI−C12
7614−Ctx−T−L−Sml−DAPI−C7
7614−Ctx−T−L−Sml−DAPI−C8
7614−Ctx−T−L−Sml−DAPI−C9
7614−Ctx−T−L−Sml−DAPI−D10
7614−Ctx−T−L−Sml−DAPI−D11
7614−Ctx−T−L−Sml−DAPI−D12
7614−Ctx−T−L−Sml−DAPI−D7
7614−Ctx−T−L−Sml−DAPI−D8
7614−Ctx−T−L−Sml−DAPI−D9
7614−Ctx−T−L−Sml−DAPI−E10
7614−Ctx−T−L−Sml−DAPI−E11
7614−Ctx−T−L−Sml−DAPI−E12
7614−Ctx−T−L−Sml−DAPI−E7
7614−Ctx−T−L−Sml−DAPI−E8
7614−Ctx−T−L−Sml−DAPI−E9
7614−Ctx−T−L−Sml−DAPI−F10
7614−Ctx−T−L−Sml−DAPI−F11
7614−Ctx−T−L−Sml−DAPI−F12
7614−Ctx−T−L−Sml−DAPI−F7
7614−Ctx−T−L−Sml−DAPI−F8
7614−Ctx−T−L−Sml−DAPI−F9
7614−Ctx−T−L−Sml−DAPI−G10
7614−Ctx−T−L−Sml−DAPI−G11
7614−Ctx−T−L−Sml−DAPI−G12
7614−Ctx−T−L−Sml−DAPI−G7
7614−Ctx−T−L−Sml−DAPI−G8
7614−Ctx−T−L−Sml−DAPI−G9
7614−Ctx−T−L−Sml−DAPI−H10
7614−Ctx−T−L−Sml−DAPI−H11
7614−Ctx−T−L−Sml−DAPI−H7
7614−Ctx−T−L−Sml−DAPI−H8
7614−Ctx−T−L−Sml−DAPI−H9
7614−Ctx−T−L−Sml−NeuN−A1
7614−Ctx−T−L−Sml−NeuN−A2
7614−Ctx−T−L−Sml−NeuN−A3
7614−Ctx−T−L−Sml−NeuN−A4
7614−Ctx−T−L−Sml−NeuN−A5
7614−Ctx−T−L−Sml−NeuN−A6
7614−Ctx−T−L−Sml−NeuN−B1
7614−Ctx−T−L−Sml−NeuN−B2
7614−Ctx−T−L−Sml−NeuN−B3
7614−Ctx−T−L−Sml−NeuN−B4
7614−Ctx−T−L−Sml−NeuN−B5
7614−Ctx−T−L−Sml−NeuN−B6
7614−Ctx−T−L−Sml−NeuN−C1
7614−Ctx−T−L−Sml−NeuN−C2
7614−Ctx−T−L−Sml−NeuN−C3
7614−Ctx−T−L−Sml−NeuN−C4
7614−Ctx−T−L−Sml−NeuN−C5
7614−Ctx−T−L−Sml−NeuN−C6
7614−Ctx−T−L−Sml−NeuN−D1
7614−Ctx−T−L−Sml−NeuN−D2
7614−Ctx−T−L−Sml−NeuN−D3
7614−Ctx−T−L−Sml−NeuN−D4
7614−Ctx−T−L−Sml−NeuN−D5
7614−Ctx−T−L−Sml−NeuN−D6
7614−Ctx−T−L−Sml−NeuN−E1
7614−Ctx−T−L−Sml−NeuN−E2
7614−Ctx−T−L−Sml−NeuN−E3
7614−Ctx−T−L−Sml−NeuN−E4
7614−Ctx−T−L−Sml−NeuN−E5
7614−Ctx−T−L−Sml−NeuN−E6
7614−Ctx−T−L−Sml−NeuN−F1
7614−Ctx−T−L−Sml−NeuN−F2
7614−Ctx−T−L−Sml−NeuN−F3
7614−Ctx−T−L−Sml−NeuN−F4
7614−Ctx−T−L−Sml−NeuN−F5
7614−Ctx−T−L−Sml−NeuN−F6
7614−Ctx−T−L−Sml−NeuN−G1
7614−Ctx−T−L−Sml−NeuN−G2
7614−Ctx−T−L−Sml−NeuN−G3
7614−Ctx−T−L−Sml−NeuN−G4
7614−Ctx−T−L−Sml−NeuN−G5
7614−Ctx−T−L−Sml−NeuN−G6
7614−Ctx−T−L−Sml−NeuN−H1
7614−Ctx−T−L−Sml−NeuN−H2
7614−Ctx−T−L−Sml−NeuN−H3
7614−Ctx−T−L−Sml−NeuN−H4
7614−Ctx−T−L−Sml−NeuN−H5
7614−Ctx−T−L−Sml−NeuN−H6
JGG-Cells

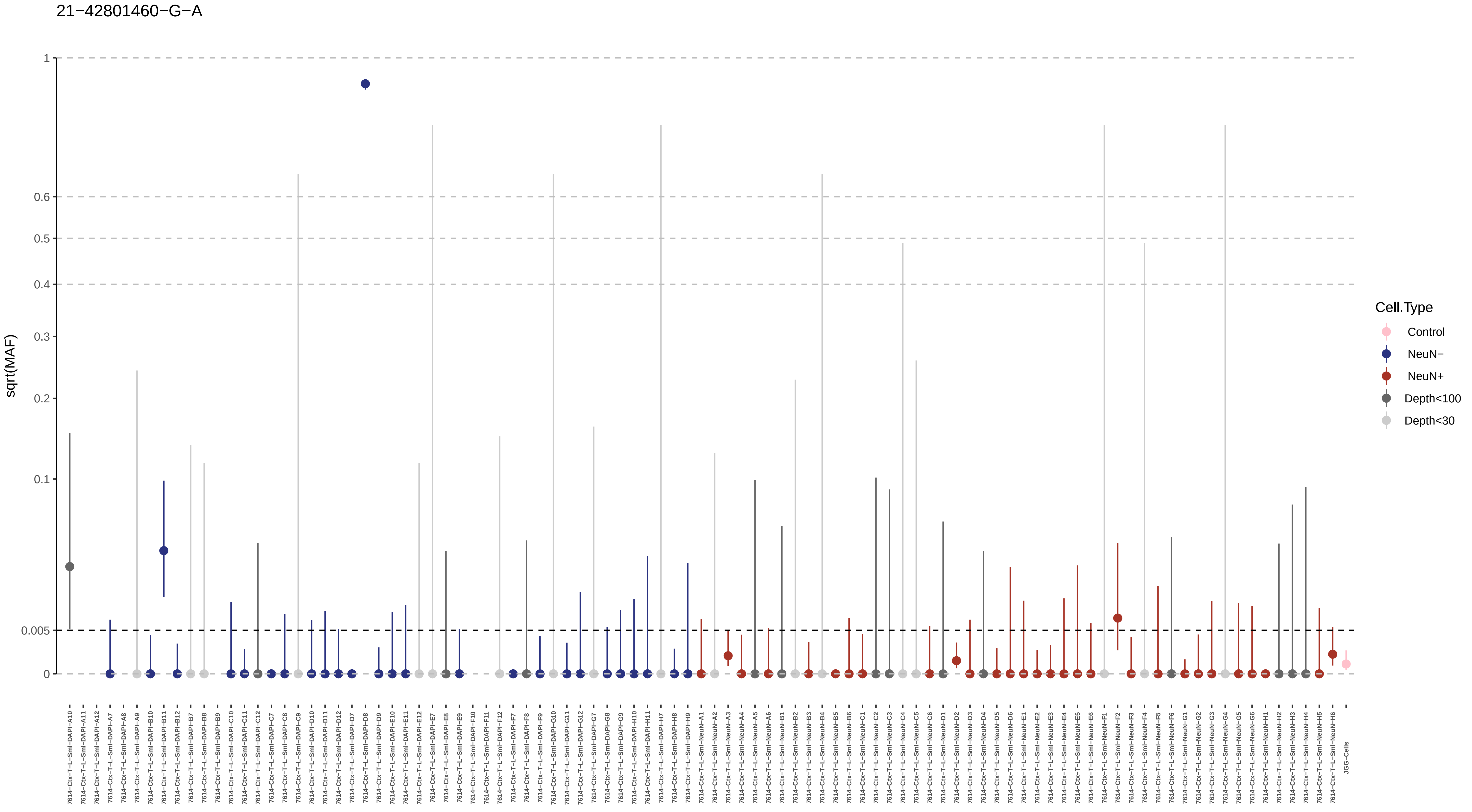

21−42801460−G−A
sqrt(MAF)
1
0.6
0.5
0.4
0.3
0.2
0.1
0.005
0
Cell.Type
Control
NeuN−
NeuN+
Depth<100
Depth<30
7614−Ctx−T−L−Sml−DAPI−A10
7614−Ctx−T−L−Sml−DAPI−A11
7614−Ctx−T−L−Sml−DAPI−A12
7614−Ctx−T−L−Sml−DAPI−A7
7614−Ctx−T−L−Sml−DAPI−A8
7614−Ctx−T−L−Sml−DAPI−A9
7614−Ctx−T−L−Sml−DAPI−B10
7614−Ctx−T−L−Sml−DAPI−B11
7614−Ctx−T−L−Sml−DAPI−B12
7614−Ctx−T−L−Sml−DAPI−B7
7614−Ctx−T−L−Sml−DAPI−B8
7614−Ctx−T−L−Sml−DAPI−B9
7614−Ctx−T−L−Sml−DAPI−C10
7614−Ctx−T−L−Sml−DAPI−C11
7614−Ctx−T−L−Sml−DAPI−C12
7614−Ctx−T−L−Sml−DAPI−C7
7614−Ctx−T−L−Sml−DAPI−C8
7614−Ctx−T−L−Sml−DAPI−C9
7614−Ctx−T−L−Sml−DAPI−D10
7614−Ctx−T−L−Sml−DAPI−D11
7614−Ctx−T−L−Sml−DAPI−D12
7614−Ctx−T−L−Sml−DAPI−D7
7614−Ctx−T−L−Sml−DAPI−D8
7614−Ctx−T−L−Sml−DAPI−D9
7614−Ctx−T−L−Sml−DAPI−E10
7614−Ctx−T−L−Sml−DAPI−E11
7614−Ctx−T−L−Sml−DAPI−E12
7614−Ctx−T−L−Sml−DAPI−E7
7614−Ctx−T−L−Sml−DAPI−E8
7614−Ctx−T−L−Sml−DAPI−E9
7614−Ctx−T−L−Sml−DAPI−F10
7614−Ctx−T−L−Sml−DAPI−F11
7614−Ctx−T−L−Sml−DAPI−F12
7614−Ctx−T−L−Sml−DAPI−F7
7614−Ctx−T−L−Sml−DAPI−F8
7614−Ctx−T−L−Sml−DAPI−F9
7614−Ctx−T−L−Sml−DAPI−G10
7614−Ctx−T−L−Sml−DAPI−G11
7614−Ctx−T−L−Sml−DAPI−G12
7614−Ctx−T−L−Sml−DAPI−G7
7614−Ctx−T−L−Sml−DAPI−G8
7614−Ctx−T−L−Sml−DAPI−G9
7614−Ctx−T−L−Sml−DAPI−H10
7614−Ctx−T−L−Sml−DAPI−H11
7614−Ctx−T−L−Sml−DAPI−H7
7614−Ctx−T−L−Sml−DAPI−H8
7614−Ctx−T−L−Sml−DAPI−H9
7614−Ctx−T−L−Sml−NeuN−A1
7614−Ctx−T−L−Sml−NeuN−A2
7614−Ctx−T−L−Sml−NeuN−A3
7614−Ctx−T−L−Sml−NeuN−A4
7614−Ctx−T−L−Sml−NeuN−A5
7614−Ctx−T−L−Sml−NeuN−A6
7614−Ctx−T−L−Sml−NeuN−B1
7614−Ctx−T−L−Sml−NeuN−B2
7614−Ctx−T−L−Sml−NeuN−B3
7614−Ctx−T−L−Sml−NeuN−B4
7614−Ctx−T−L−Sml−NeuN−B5
7614−Ctx−T−L−Sml−NeuN−B6
7614−Ctx−T−L−Sml−NeuN−C1
7614−Ctx−T−L−Sml−NeuN−C2
7614−Ctx−T−L−Sml−NeuN−C3
7614−Ctx−T−L−Sml−NeuN−C4
7614−Ctx−T−L−Sml−NeuN−C5
7614−Ctx−T−L−Sml−NeuN−C6
7614−Ctx−T−L−Sml−NeuN−D1
7614−Ctx−T−L−Sml−NeuN−D2
7614−Ctx−T−L−Sml−NeuN−D3
7614−Ctx−T−L−Sml−NeuN−D4
7614−Ctx−T−L−Sml−NeuN−D5
7614−Ctx−T−L−Sml−NeuN−D6
7614−Ctx−T−L−Sml−NeuN−E1
7614−Ctx−T−L−Sml−NeuN−E2
7614−Ctx−T−L−Sml−NeuN−E3
7614−Ctx−T−L−Sml−NeuN−E4
7614−Ctx−T−L−Sml−NeuN−E5
7614−Ctx−T−L−Sml−NeuN−E6
7614−Ctx−T−L−Sml−NeuN−F1
7614−Ctx−T−L−Sml−NeuN−F2
7614−Ctx−T−L−Sml−NeuN−F3
7614−Ctx−T−L−Sml−NeuN−F4
7614−Ctx−T−L−Sml−NeuN−F5
7614−Ctx−T−L−Sml−NeuN−F6
7614−Ctx−T−L−Sml−NeuN−G1
7614−Ctx−T−L−Sml−NeuN−G2
7614−Ctx−T−L−Sml−NeuN−G3
7614−Ctx−T−L−Sml−NeuN−G4
7614−Ctx−T−L−Sml−NeuN−G5
7614−Ctx−T−L−Sml−NeuN−G6
7614−Ctx−T−L−Sml−NeuN−H1
7614−Ctx−T−L−Sml−NeuN−H2
7614−Ctx−T−L−Sml−NeuN−H3
7614−Ctx−T−L−Sml−NeuN−H4
7614−Ctx−T−L−Sml−NeuN−H5
7614−Ctx−T−L−Sml−NeuN−H6
JGG-Cells

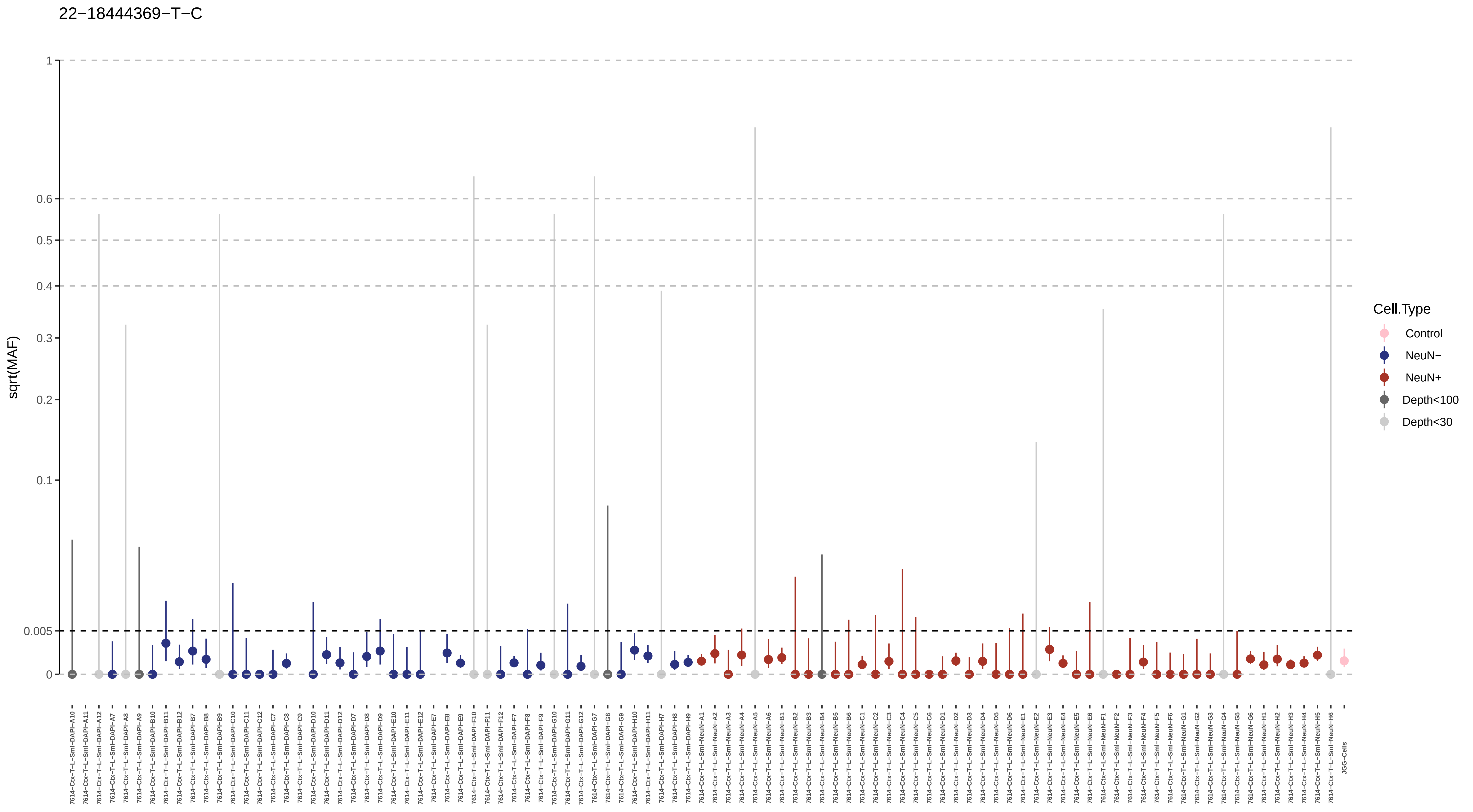

22−18444369−T−C
sqrt(MAF)
1
0.6
0.5
0.4
0.3
0.2
0.1
0.005
0
Cell.Type
Control
NeuN−
NeuN+
Depth<100
Depth<30
7614−Ctx−T−L−Sml−DAPI−A10
7614−Ctx−T−L−Sml−DAPI−A11
7614−Ctx−T−L−Sml−DAPI−A12
7614−Ctx−T−L−Sml−DAPI−A7
7614−Ctx−T−L−Sml−DAPI−A8
7614−Ctx−T−L−Sml−DAPI−A9
7614−Ctx−T−L−Sml−DAPI−B10
7614−Ctx−T−L−Sml−DAPI−B11
7614−Ctx−T−L−Sml−DAPI−B12
7614−Ctx−T−L−Sml−DAPI−B7
7614−Ctx−T−L−Sml−DAPI−B8
7614−Ctx−T−L−Sml−DAPI−B9
7614−Ctx−T−L−Sml−DAPI−C10
7614−Ctx−T−L−Sml−DAPI−C11
7614−Ctx−T−L−Sml−DAPI−C12
7614−Ctx−T−L−Sml−DAPI−C7
7614−Ctx−T−L−Sml−DAPI−C8
7614−Ctx−T−L−Sml−DAPI−C9
7614−Ctx−T−L−Sml−DAPI−D10
7614−Ctx−T−L−Sml−DAPI−D11
7614−Ctx−T−L−Sml−DAPI−D12
7614−Ctx−T−L−Sml−DAPI−D7
7614−Ctx−T−L−Sml−DAPI−D8
7614−Ctx−T−L−Sml−DAPI−D9
7614−Ctx−T−L−Sml−DAPI−E10
7614−Ctx−T−L−Sml−DAPI−E11
7614−Ctx−T−L−Sml−DAPI−E12
7614−Ctx−T−L−Sml−DAPI−E7
7614−Ctx−T−L−Sml−DAPI−E8
7614−Ctx−T−L−Sml−DAPI−E9
7614−Ctx−T−L−Sml−DAPI−F10
7614−Ctx−T−L−Sml−DAPI−F11
7614−Ctx−T−L−Sml−DAPI−F12
7614−Ctx−T−L−Sml−DAPI−F7
7614−Ctx−T−L−Sml−DAPI−F8
7614−Ctx−T−L−Sml−DAPI−F9
7614−Ctx−T−L−Sml−DAPI−G10
7614−Ctx−T−L−Sml−DAPI−G11
7614−Ctx−T−L−Sml−DAPI−G12
7614−Ctx−T−L−Sml−DAPI−G7
7614−Ctx−T−L−Sml−DAPI−G8
7614−Ctx−T−L−Sml−DAPI−G9
7614−Ctx−T−L−Sml−DAPI−H10
7614−Ctx−T−L−Sml−DAPI−H11
7614−Ctx−T−L−Sml−DAPI−H7
7614−Ctx−T−L−Sml−DAPI−H8
7614−Ctx−T−L−Sml−DAPI−H9
7614−Ctx−T−L−Sml−NeuN−A1
7614−Ctx−T−L−Sml−NeuN−A2
7614−Ctx−T−L−Sml−NeuN−A3
7614−Ctx−T−L−Sml−NeuN−A4
7614−Ctx−T−L−Sml−NeuN−A5
7614−Ctx−T−L−Sml−NeuN−A6
7614−Ctx−T−L−Sml−NeuN−B1
7614−Ctx−T−L−Sml−NeuN−B2
7614−Ctx−T−L−Sml−NeuN−B3
7614−Ctx−T−L−Sml−NeuN−B4
7614−Ctx−T−L−Sml−NeuN−B5
7614−Ctx−T−L−Sml−NeuN−B6
7614−Ctx−T−L−Sml−NeuN−C1
7614−Ctx−T−L−Sml−NeuN−C2
7614−Ctx−T−L−Sml−NeuN−C3
7614−Ctx−T−L−Sml−NeuN−C4
7614−Ctx−T−L−Sml−NeuN−C5
7614−Ctx−T−L−Sml−NeuN−C6
7614−Ctx−T−L−Sml−NeuN−D1
7614−Ctx−T−L−Sml−NeuN−D2
7614−Ctx−T−L−Sml−NeuN−D3
7614−Ctx−T−L−Sml−NeuN−D4
7614−Ctx−T−L−Sml−NeuN−D5
7614−Ctx−T−L−Sml−NeuN−D6
7614−Ctx−T−L−Sml−NeuN−E1
7614−Ctx−T−L−Sml−NeuN−E2
7614−Ctx−T−L−Sml−NeuN−E3
7614−Ctx−T−L−Sml−NeuN−E4
7614−Ctx−T−L−Sml−NeuN−E5
7614−Ctx−T−L−Sml−NeuN−E6
7614−Ctx−T−L−Sml−NeuN−F1
7614−Ctx−T−L−Sml−NeuN−F2
7614−Ctx−T−L−Sml−NeuN−F3
7614−Ctx−T−L−Sml−NeuN−F4
7614−Ctx−T−L−Sml−NeuN−F5
7614−Ctx−T−L−Sml−NeuN−F6
7614−Ctx−T−L−Sml−NeuN−G1
7614−Ctx−T−L−Sml−NeuN−G2
7614−Ctx−T−L−Sml−NeuN−G3
7614−Ctx−T−L−Sml−NeuN−G4
7614−Ctx−T−L−Sml−NeuN−G5
7614−Ctx−T−L−Sml−NeuN−G6
7614−Ctx−T−L−Sml−NeuN−H1
7614−Ctx−T−L−Sml−NeuN−H2
7614−Ctx−T−L−Sml−NeuN−H3
7614−Ctx−T−L−Sml−NeuN−H4
7614−Ctx−T−L−Sml−NeuN−H5
7614−Ctx−T−L−Sml−NeuN−H6
JGG-Cells

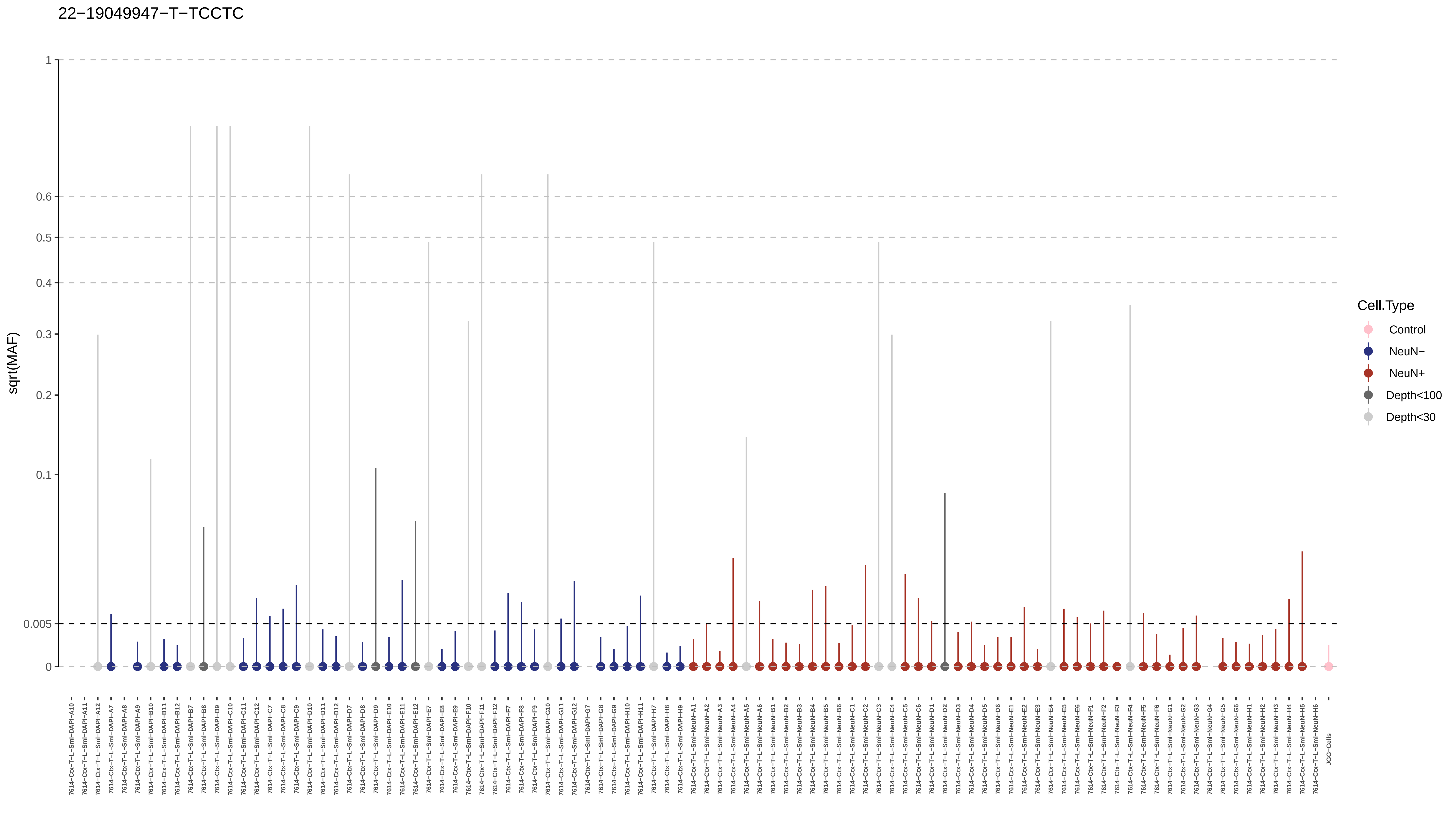

22−19049947−T−TCCTC
sqrt(MAF)
1
0.6
0.5
0.4
0.3
0.2
0.1
0.005
0
Cell.Type
Control
NeuN−
NeuN+
Depth<100
Depth<30
7614−Ctx−T−L−Sml−DAPI−A10
7614−Ctx−T−L−Sml−DAPI−A11
7614−Ctx−T−L−Sml−DAPI−A12
7614−Ctx−T−L−Sml−DAPI−A7
7614−Ctx−T−L−Sml−DAPI−A8
7614−Ctx−T−L−Sml−DAPI−A9
7614−Ctx−T−L−Sml−DAPI−B10
7614−Ctx−T−L−Sml−DAPI−B11
7614−Ctx−T−L−Sml−DAPI−B12
7614−Ctx−T−L−Sml−DAPI−B7
7614−Ctx−T−L−Sml−DAPI−B8
7614−Ctx−T−L−Sml−DAPI−B9
7614−Ctx−T−L−Sml−DAPI−C10
7614−Ctx−T−L−Sml−DAPI−C11
7614−Ctx−T−L−Sml−DAPI−C12
7614−Ctx−T−L−Sml−DAPI−C7
7614−Ctx−T−L−Sml−DAPI−C8
7614−Ctx−T−L−Sml−DAPI−C9
7614−Ctx−T−L−Sml−DAPI−D10
7614−Ctx−T−L−Sml−DAPI−D11
7614−Ctx−T−L−Sml−DAPI−D12
7614−Ctx−T−L−Sml−DAPI−D7
7614−Ctx−T−L−Sml−DAPI−D8
7614−Ctx−T−L−Sml−DAPI−D9
7614−Ctx−T−L−Sml−DAPI−E10
7614−Ctx−T−L−Sml−DAPI−E11
7614−Ctx−T−L−Sml−DAPI−E12
7614−Ctx−T−L−Sml−DAPI−E7
7614−Ctx−T−L−Sml−DAPI−E8
7614−Ctx−T−L−Sml−DAPI−E9
7614−Ctx−T−L−Sml−DAPI−F10
7614−Ctx−T−L−Sml−DAPI−F11
7614−Ctx−T−L−Sml−DAPI−F12
7614−Ctx−T−L−Sml−DAPI−F7
7614−Ctx−T−L−Sml−DAPI−F8
7614−Ctx−T−L−Sml−DAPI−F9
7614−Ctx−T−L−Sml−DAPI−G10
7614−Ctx−T−L−Sml−DAPI−G11
7614−Ctx−T−L−Sml−DAPI−G12
7614−Ctx−T−L−Sml−DAPI−G7
7614−Ctx−T−L−Sml−DAPI−G8
7614−Ctx−T−L−Sml−DAPI−G9
7614−Ctx−T−L−Sml−DAPI−H10
7614−Ctx−T−L−Sml−DAPI−H11
7614−Ctx−T−L−Sml−DAPI−H7
7614−Ctx−T−L−Sml−DAPI−H8
7614−Ctx−T−L−Sml−DAPI−H9
7614−Ctx−T−L−Sml−NeuN−A1
7614−Ctx−T−L−Sml−NeuN−A2
7614−Ctx−T−L−Sml−NeuN−A3
7614−Ctx−T−L−Sml−NeuN−A4
7614−Ctx−T−L−Sml−NeuN−A5
7614−Ctx−T−L−Sml−NeuN−A6
7614−Ctx−T−L−Sml−NeuN−B1
7614−Ctx−T−L−Sml−NeuN−B2
7614−Ctx−T−L−Sml−NeuN−B3
7614−Ctx−T−L−Sml−NeuN−B4
7614−Ctx−T−L−Sml−NeuN−B5
7614−Ctx−T−L−Sml−NeuN−B6
7614−Ctx−T−L−Sml−NeuN−C1
7614−Ctx−T−L−Sml−NeuN−C2
7614−Ctx−T−L−Sml−NeuN−C3
7614−Ctx−T−L−Sml−NeuN−C4
7614−Ctx−T−L−Sml−NeuN−C5
7614−Ctx−T−L−Sml−NeuN−C6
7614−Ctx−T−L−Sml−NeuN−D1
7614−Ctx−T−L−Sml−NeuN−D2
7614−Ctx−T−L−Sml−NeuN−D3
7614−Ctx−T−L−Sml−NeuN−D4
7614−Ctx−T−L−Sml−NeuN−D5
7614−Ctx−T−L−Sml−NeuN−D6
7614−Ctx−T−L−Sml−NeuN−E1
7614−Ctx−T−L−Sml−NeuN−E2
7614−Ctx−T−L−Sml−NeuN−E3
7614−Ctx−T−L−Sml−NeuN−E4
7614−Ctx−T−L−Sml−NeuN−E5
7614−Ctx−T−L−Sml−NeuN−E6
7614−Ctx−T−L−Sml−NeuN−F1
7614−Ctx−T−L−Sml−NeuN−F2
7614−Ctx−T−L−Sml−NeuN−F3
7614−Ctx−T−L−Sml−NeuN−F4
7614−Ctx−T−L−Sml−NeuN−F5
7614−Ctx−T−L−Sml−NeuN−F6
7614−Ctx−T−L−Sml−NeuN−G1
7614−Ctx−T−L−Sml−NeuN−G2
7614−Ctx−T−L−Sml−NeuN−G3
7614−Ctx−T−L−Sml−NeuN−G4
7614−Ctx−T−L−Sml−NeuN−G5
7614−Ctx−T−L−Sml−NeuN−G6
7614−Ctx−T−L−Sml−NeuN−H1
7614−Ctx−T−L−Sml−NeuN−H2
7614−Ctx−T−L−Sml−NeuN−H3
7614−Ctx−T−L−Sml−NeuN−H4
7614−Ctx−T−L−Sml−NeuN−H5
7614−Ctx−T−L−Sml−NeuN−H6
JGG-Cells

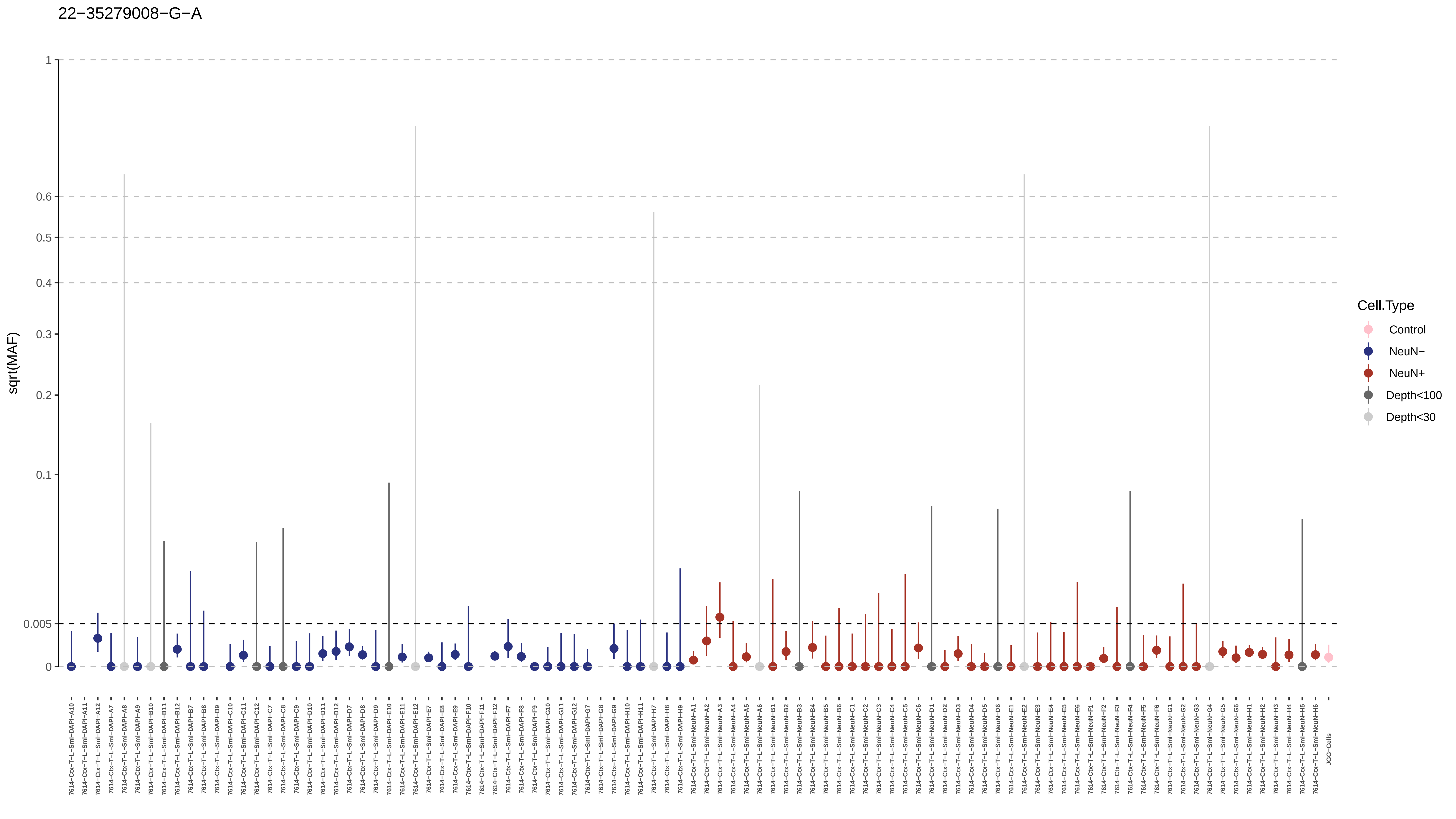

22−35279008−G−A
sqrt(MAF)
1
0.6
0.5
0.4
0.3
0.2
0.1
0.005
0
Cell.Type
Control
NeuN−
NeuN+
Depth<100
Depth<30
JGG-Cells

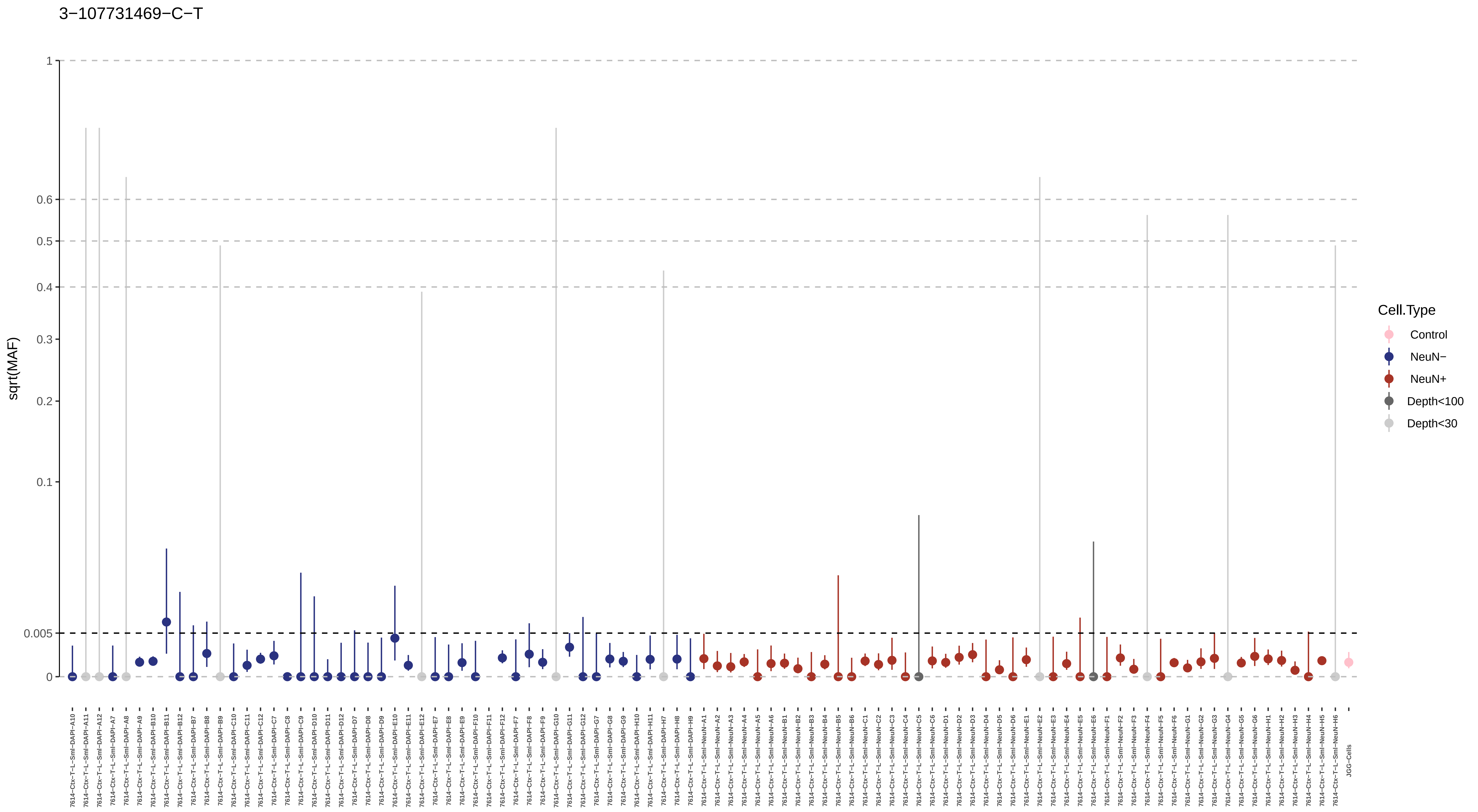

3−107731469−C−T
sqrt(MAF)
1
0.6
0.5
0.4
0.3
0.2
0.1
0.005
0
Cell.Type
Control
NeuN−
NeuN+
Depth<100
Depth<30
7614−Ctx−T−L−Sml−DAPI−A10
7614−Ctx−T−L−Sml−DAPI−A11
7614−Ctx−T−L−Sml−DAPI−A12
7614−Ctx−T−L−Sml−DAPI−A7
7614−Ctx−T−L−Sml−DAPI−A8
7614−Ctx−T−L−Sml−DAPI−A9
7614−Ctx−T−L−Sml−DAPI−B10
7614−Ctx−T−L−Sml−DAPI−B11
7614−Ctx−T−L−Sml−DAPI−B12
7614−Ctx−T−L−Sml−DAPI−B7
7614−Ctx−T−L−Sml−DAPI−B8
7614−Ctx−T−L−Sml−DAPI−B9
7614−Ctx−T−L−Sml−DAPI−C10
7614−Ctx−T−L−Sml−DAPI−C11
7614−Ctx−T−L−Sml−DAPI−C12
7614−Ctx−T−L−Sml−DAPI−C7
7614−Ctx−T−L−Sml−DAPI−C8
7614−Ctx−T−L−Sml−DAPI−C9
7614−Ctx−T−L−Sml−DAPI−D10
7614−Ctx−T−L−Sml−DAPI−D11
7614−Ctx−T−L−Sml−DAPI−D12
7614−Ctx−T−L−Sml−DAPI−D7
7614−Ctx−T−L−Sml−DAPI−D8
7614−Ctx−T−L−Sml−DAPI−D9
7614−Ctx−T−L−Sml−DAPI−E10
7614−Ctx−T−L−Sml−DAPI−E11
7614−Ctx−T−L−Sml−DAPI−E12
7614−Ctx−T−L−Sml−DAPI−E7
7614−Ctx−T−L−Sml−DAPI−E8
7614−Ctx−T−L−Sml−DAPI−E9
7614−Ctx−T−L−Sml−DAPI−F10
7614−Ctx−T−L−Sml−DAPI−F11
7614−Ctx−T−L−Sml−DAPI−F12
7614−Ctx−T−L−Sml−DAPI−F7
7614−Ctx−T−L−Sml−DAPI−F8
7614−Ctx−T−L−Sml−DAPI−F9
7614−Ctx−T−L−Sml−DAPI−G10
7614−Ctx−T−L−Sml−DAPI−G11
7614−Ctx−T−L−Sml−DAPI−G12
7614−Ctx−T−L−Sml−DAPI−G7
7614−Ctx−T−L−Sml−DAPI−G8
7614−Ctx−T−L−Sml−DAPI−G9
7614−Ctx−T−L−Sml−DAPI−H10
7614−Ctx−T−L−Sml−DAPI−H11
7614−Ctx−T−L−Sml−DAPI−H7
7614−Ctx−T−L−Sml−DAPI−H8
7614−Ctx−T−L−Sml−DAPI−H9
7614−Ctx−T−L−Sml−NeuN−A1
7614−Ctx−T−L−Sml−NeuN−A2
7614−Ctx−T−L−Sml−NeuN−A3
7614−Ctx−T−L−Sml−NeuN−A4
7614−Ctx−T−L−Sml−NeuN−A5
7614−Ctx−T−L−Sml−NeuN−A6
7614−Ctx−T−L−Sml−NeuN−B1
7614−Ctx−T−L−Sml−NeuN−B2
7614−Ctx−T−L−Sml−NeuN−B3
7614−Ctx−T−L−Sml−NeuN−B4
7614−Ctx−T−L−Sml−NeuN−B5
7614−Ctx−T−L−Sml−NeuN−B6
7614−Ctx−T−L−Sml−NeuN−C1
7614−Ctx−T−L−Sml−NeuN−C2
7614−Ctx−T−L−Sml−NeuN−C3
7614−Ctx−T−L−Sml−NeuN−C4
7614−Ctx−T−L−Sml−NeuN−C5
7614−Ctx−T−L−Sml−NeuN−C6
7614−Ctx−T−L−Sml−NeuN−D1
7614−Ctx−T−L−Sml−NeuN−D2
7614−Ctx−T−L−Sml−NeuN−D3
7614−Ctx−T−L−Sml−NeuN−D4
7614−Ctx−T−L−Sml−NeuN−D5
7614−Ctx−T−L−Sml−NeuN−D6
7614−Ctx−T−L−Sml−NeuN−E1
7614−Ctx−T−L−Sml−NeuN−E2
7614−Ctx−T−L−Sml−NeuN−E3
7614−Ctx−T−L−Sml−NeuN−E4
7614−Ctx−T−L−Sml−NeuN−E5
7614−Ctx−T−L−Sml−NeuN−E6
7614−Ctx−T−L−Sml−NeuN−F1
7614−Ctx−T−L−Sml−NeuN−F2
7614−Ctx−T−L−Sml−NeuN−F3
7614−Ctx−T−L−Sml−NeuN−F4
7614−Ctx−T−L−Sml−NeuN−F5
7614−Ctx−T−L−Sml−NeuN−F6
7614−Ctx−T−L−Sml−NeuN−G1
7614−Ctx−T−L−Sml−NeuN−G2
7614−Ctx−T−L−Sml−NeuN−G3
7614−Ctx−T−L−Sml−NeuN−G4
7614−Ctx−T−L−Sml−NeuN−G5
7614−Ctx−T−L−Sml−NeuN−G6
7614−Ctx−T−L−Sml−NeuN−H1
7614−Ctx−T−L−Sml−NeuN−H2
7614−Ctx−T−L−Sml−NeuN−H3
7614−Ctx−T−L−Sml−NeuN−H4
7614−Ctx−T−L−Sml−NeuN−H5
7614−Ctx−T−L−Sml−NeuN−H6
JGG−Cells

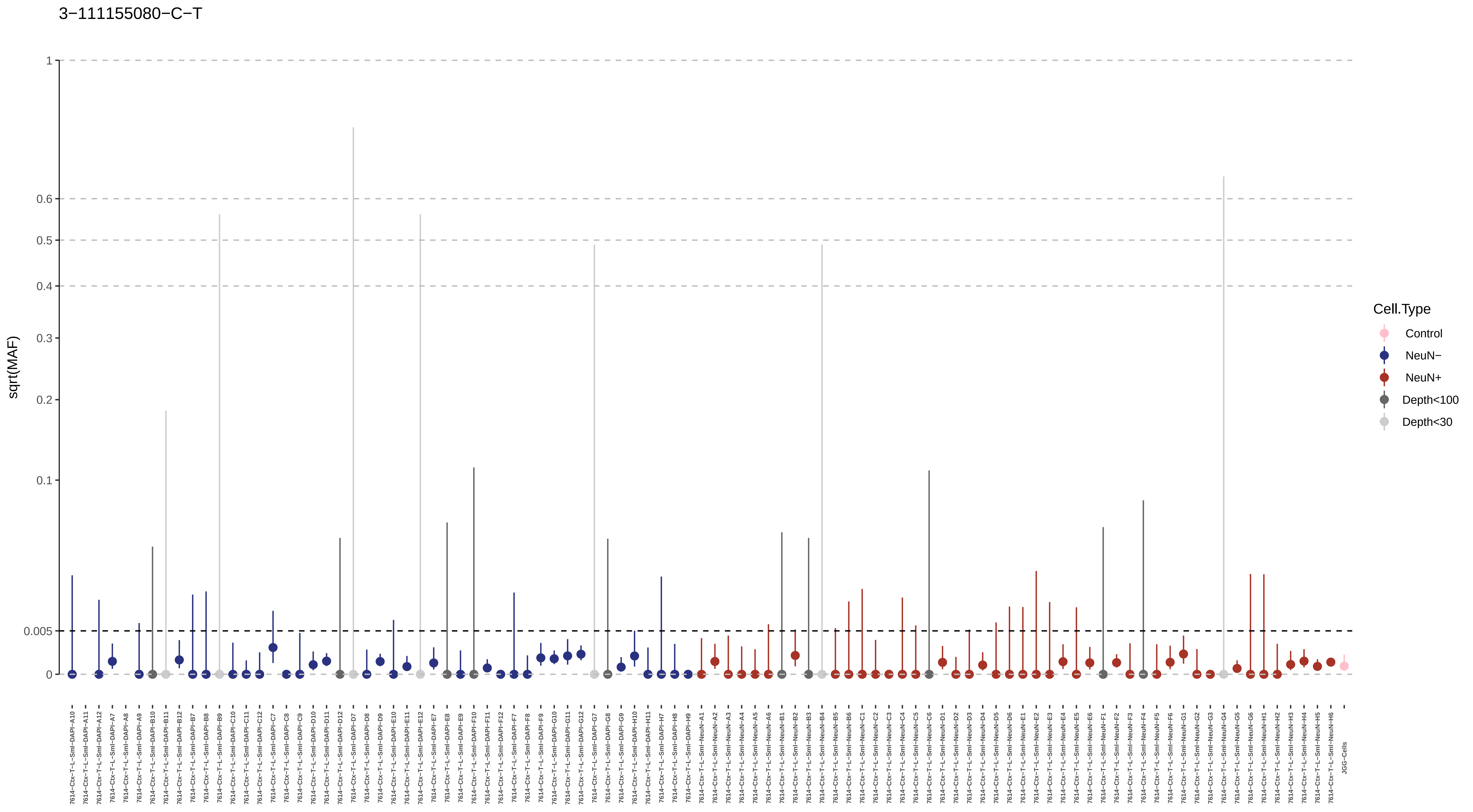

3−111155080−C−T
sqrt(MAF)
1
0.6
0.5
0.4
0.3
0.2
0.1
0.005
0
Cell.Type
Control
NeuN−
NeuN+
Depth<100
Depth<30
JGG-Cells

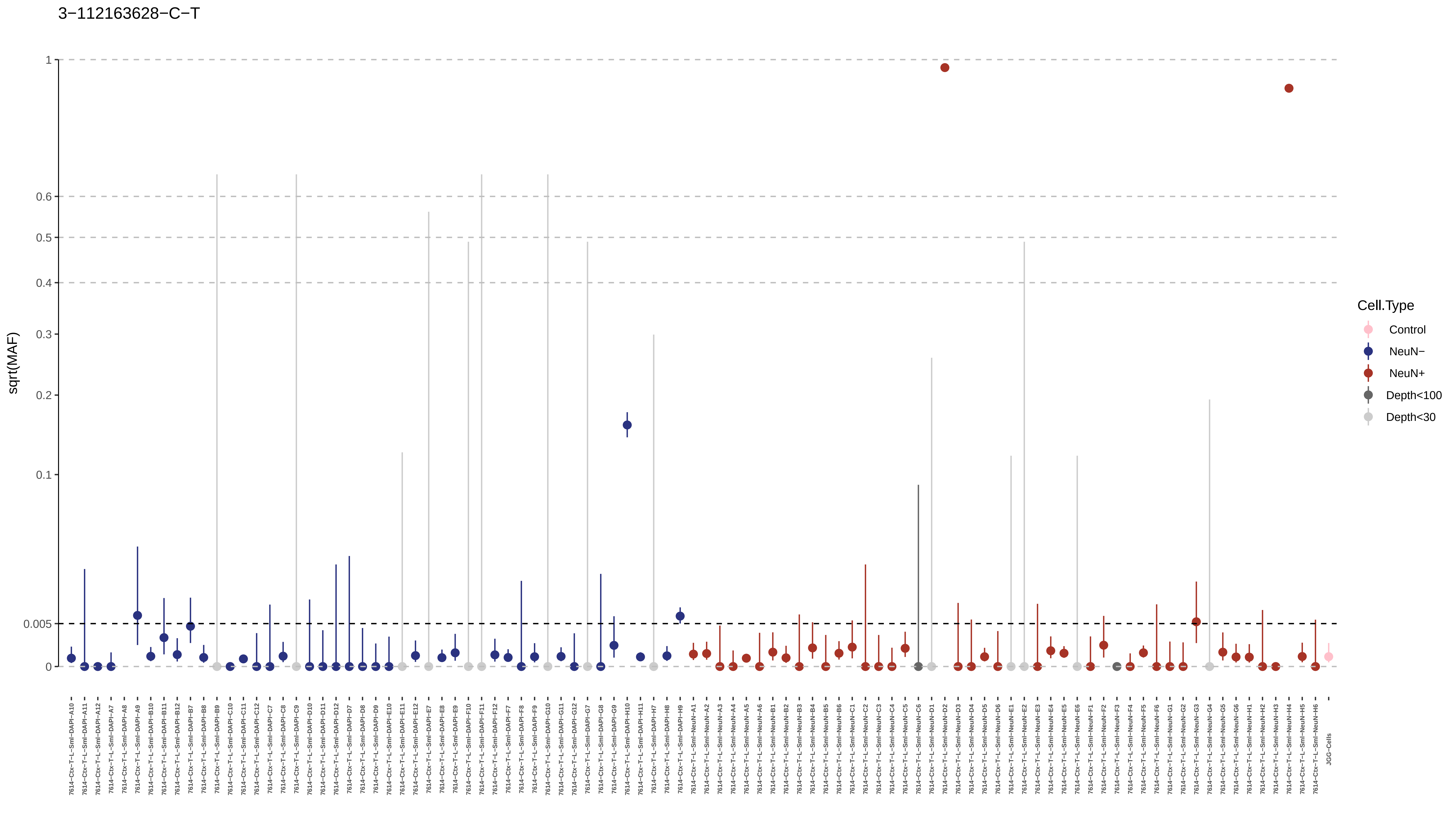

3−112163628−C−T
sqrt(MAF)
1
0.6
0.5
0.4
0.3
0.2
0.1
0.005
0
Cell.Type
Control
NeuN−
NeuN+
Depth<100
Depth<30
7614−Ctx−T−L−Sml−DAPI−A10
7614−Ctx−T−L−Sml−DAPI−A11
7614−Ctx−T−L−Sml−DAPI−A12
7614−Ctx−T−L−Sml−DAPI−A7
7614−Ctx−T−L−Sml−DAPI−A8
7614−Ctx−T−L−Sml−DAPI−A9
7614−Ctx−T−L−Sml−DAPI−B10
7614−Ctx−T−L−Sml−DAPI−B11
7614−Ctx−T−L−Sml−DAPI−B12
7614−Ctx−T−L−Sml−DAPI−B7
7614−Ctx−T−L−Sml−DAPI−B8
7614−Ctx−T−L−Sml−DAPI−B9
7614−Ctx−T−L−Sml−DAPI−C10
7614−Ctx−T−L−Sml−DAPI−C11
7614−Ctx−T−L−Sml−DAPI−C12
7614−Ctx−T−L−Sml−DAPI−C7
7614−Ctx−T−L−Sml−DAPI−C8
7614−Ctx−T−L−Sml−DAPI−C9
7614−Ctx−T−L−Sml−DAPI−D10
7614−Ctx−T−L−Sml−DAPI−D11
7614−Ctx−T−L−Sml−DAPI−D12
7614−Ctx−T−L−Sml−DAPI−D7
7614−Ctx−T−L−Sml−DAPI−D8
7614−Ctx−T−L−Sml−DAPI−D9
7614−Ctx−T−L−Sml−DAPI−E10
7614−Ctx−T−L−Sml−DAPI−E11
7614−Ctx−T−L−Sml−DAPI−E12
7614−Ctx−T−L−Sml−DAPI−E7
7614−Ctx−T−L−Sml−DAPI−E8
7614−Ctx−T−L−Sml−DAPI−E9
7614−Ctx−T−L−Sml−DAPI−F10
7614−Ctx−T−L−Sml−DAPI−F11
7614−Ctx−T−L−Sml−DAPI−F12
7614−Ctx−T−L−Sml−DAPI−F7
7614−Ctx−T−L−Sml−DAPI−F8
7614−Ctx−T−L−Sml−DAPI−F9
7614−Ctx−T−L−Sml−DAPI−G10
7614−Ctx−T−L−Sml−DAPI−G11
7614−Ctx−T−L−Sml−DAPI−G12
7614−Ctx−T−L−Sml−DAPI−G7
7614−Ctx−T−L−Sml−DAPI−G8
7614−Ctx−T−L−Sml−DAPI−G9
7614−Ctx−T−L−Sml−DAPI−H10
7614−Ctx−T−L−Sml−DAPI−H11
7614−Ctx−T−L−Sml−DAPI−H7
7614−Ctx−T−L−Sml−DAPI−H8
7614−Ctx−T−L−Sml−DAPI−H9
7614−Ctx−T−L−Sml−NeuN−A1
7614−Ctx−T−L−Sml−NeuN−A2
7614−Ctx−T−L−Sml−NeuN−A3
7614−Ctx−T−L−Sml−NeuN−A4
7614−Ctx−T−L−Sml−NeuN−A5
7614−Ctx−T−L−Sml−NeuN−A6
7614−Ctx−T−L−Sml−NeuN−B1
7614−Ctx−T−L−Sml−NeuN−B2
7614−Ctx−T−L−Sml−NeuN−B3
7614−Ctx−T−L−Sml−NeuN−B4
7614−Ctx−T−L−Sml−NeuN−B5
7614−Ctx−T−L−Sml−NeuN−B6
7614−Ctx−T−L−Sml−NeuN−C1
7614−Ctx−T−L−Sml−NeuN−C2
7614−Ctx−T−L−Sml−NeuN−C3
7614−Ctx−T−L−Sml−NeuN−C4
7614−Ctx−T−L−Sml−NeuN−C5
7614−Ctx−T−L−Sml−NeuN−C6
7614−Ctx−T−L−Sml−NeuN−D1
7614−Ctx−T−L−Sml−NeuN−D2
7614−Ctx−T−L−Sml−NeuN−D3
7614−Ctx−T−L−Sml−NeuN−D4
7614−Ctx−T−L−Sml−NeuN−D5
7614−Ctx−T−L−Sml−NeuN−D6
7614−Ctx−T−L−Sml−NeuN−E1
7614−Ctx−T−L−Sml−NeuN−E2
7614−Ctx−T−L−Sml−NeuN−E3
7614−Ctx−T−L−Sml−NeuN−E4
7614−Ctx−T−L−Sml−NeuN−E5
7614−Ctx−T−L−Sml−NeuN−E6
7614−Ctx−T−L−Sml−NeuN−F1
7614−Ctx−T−L−Sml−NeuN−F2
7614−Ctx−T−L−Sml−NeuN−F3
7614−Ctx−T−L−Sml−NeuN−F4
7614−Ctx−T−L−Sml−NeuN−F5
7614−Ctx−T−L−Sml−NeuN−F6
7614−Ctx−T−L−Sml−NeuN−G1
7614−Ctx−T−L−Sml−NeuN−G2
7614−Ctx−T−L−Sml−NeuN−G3
7614−Ctx−T−L−Sml−NeuN−G4
7614−Ctx−T−L−Sml−NeuN−G5
7614−Ctx−T−L−Sml−NeuN−G6
7614−Ctx−T−L−Sml−NeuN−H1
7614−Ctx−T−L−Sml−NeuN−H2
7614−Ctx−T−L−Sml−NeuN−H3
7614−Ctx−T−L−Sml−NeuN−H4
7614−Ctx−T−L−Sml−NeuN−H5
7614−Ctx−T−L−Sml−NeuN−H6
JGG-Cells

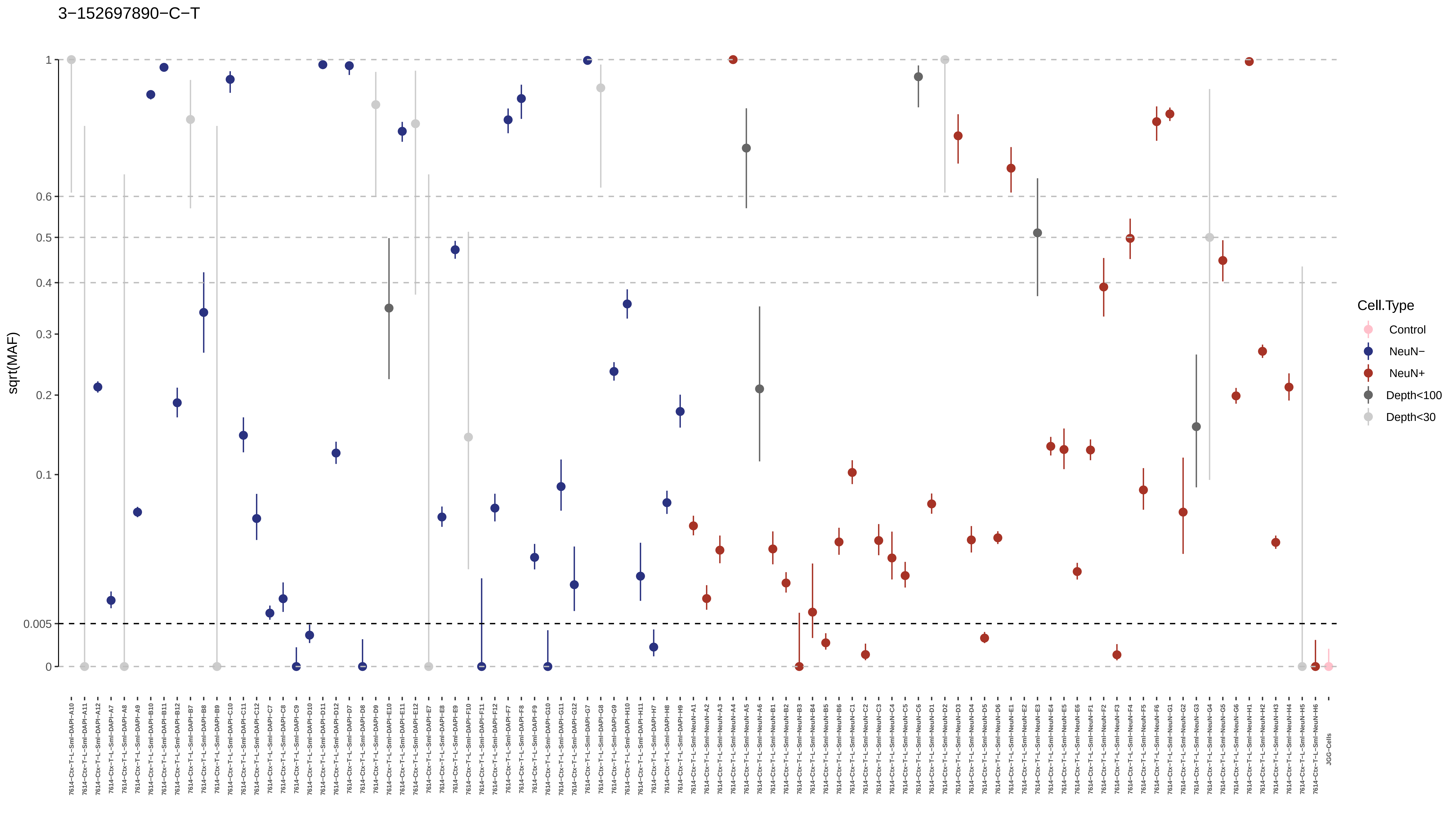
3−152697890−C−T
sqrt(MAF)
1
0.6
0.5
0.4
0.3
0.2
0.1
0.005
0
Cell.Type
Control
NeuN−
NeuN+
Depth<100
Depth<30

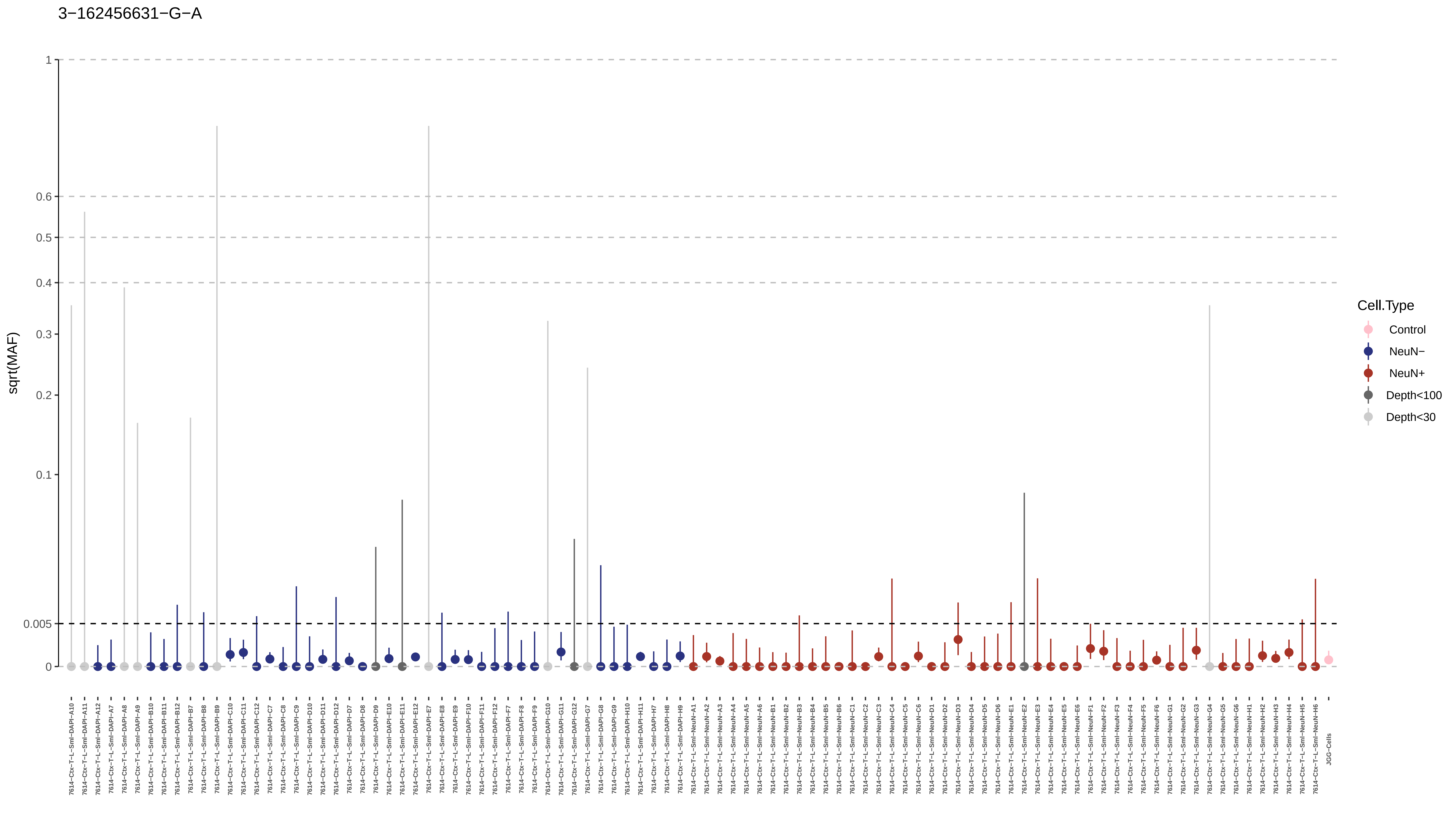
3−162456631−G−A
sqrt(MAF)
1
0.6
0.5
0.4
0.3
0.2
0.1
0.005
0
Cell.Type
Control
NeuN−
NeuN+
Depth<100
Depth<30
7614−Ctx−T−L−Sml−DAPI−A10
7614−Ctx−T−L−Sml−DAPI−A11
7614−Ctx−T−L−Sml−DAPI−A12
7614−Ctx−T−L−Sml−DAPI−A7
7614−Ctx−T−L−Sml−DAPI−A8
7614−Ctx−T−L−Sml−DAPI−A9
7614−Ctx−T−L−Sml−DAPI−B10
7614−Ctx−T−L−Sml−DAPI−B11
7614−Ctx−T−L−Sml−DAPI−B12
7614−Ctx−T−L−Sml−DAPI−B7
7614−Ctx−T−L−Sml−DAPI−B8
7614−Ctx−T−L−Sml−DAPI−B9
7614−Ctx−T−L−Sml−DAPI−C10
7614−Ctx−T−L−Sml−DAPI−C11
7614−Ctx−T−L−Sml−DAPI−C12
7614−Ctx−T−L−Sml−DAPI−C7
7614−Ctx−T−L−Sml−DAPI−C8
7614−Ctx−T−L−Sml−DAPI−C9
7614−Ctx−T−L−Sml−DAPI−D10
7614−Ctx−T−L−Sml−DAPI−D11
7614−Ctx−T−L−Sml−DAPI−D12
7614−Ctx−T−L−Sml−DAPI−D7
7614−Ctx−T−L−Sml−DAPI−D8
7614−Ctx−T−L−Sml−DAPI−D9
7614−Ctx−T−L−Sml−DAPI−E10
7614−Ctx−T−L−Sml−DAPI−E11
7614−Ctx−T−L−Sml−DAPI−E12
7614−Ctx−T−L−Sml−DAPI−E7
7614−Ctx−T−L−Sml−DAPI−E8
7614−Ctx−T−L−Sml−DAPI−E9
7614−Ctx−T−L−Sml−DAPI−F10
7614−Ctx−T−L−Sml−DAPI−F11
7614−Ctx−T−L−Sml−DAPI−F12
7614−Ctx−T−L−Sml−DAPI−F7
7614−Ctx−T−L−Sml−DAPI−F8
7614−Ctx−T−L−Sml−DAPI−F9
7614−Ctx−T−L−Sml−DAPI−G10
7614−Ctx−T−L−Sml−DAPI−G11
7614−Ctx−T−L−Sml−DAPI−G12
7614−Ctx−T−L−Sml−DAPI−G7
7614−Ctx−T−L−Sml−DAPI−G8
7614−Ctx−T−L−Sml−DAPI−G9
7614−Ctx−T−L−Sml−DAPI−H10
7614−Ctx−T−L−Sml−DAPI−H11
7614−Ctx−T−L−Sml−DAPI−H7
7614−Ctx−T−L−Sml−DAPI−H8
7614−Ctx−T−L−Sml−DAPI−H9
7614−Ctx−T−L−Sml−NeuN−A1
7614−Ctx−T−L−Sml−NeuN−A2
7614−Ctx−T−L−Sml−NeuN−A3
7614−Ctx−T−L−Sml−NeuN−A4
7614−Ctx−T−L−Sml−NeuN−A5
7614−Ctx−T−L−Sml−NeuN−A6
7614−Ctx−T−L−Sml−NeuN−B1
7614−Ctx−T−L−Sml−NeuN−B2
7614−Ctx−T−L−Sml−NeuN−B3
7614−Ctx−T−L−Sml−NeuN−B4
7614−Ctx−T−L−Sml−NeuN−B5
7614−Ctx−T−L−Sml−NeuN−B6
7614−Ctx−T−L−Sml−NeuN−C1
7614−Ctx−T−L−Sml−NeuN−C2
7614−Ctx−T−L−Sml−NeuN−C3
7614−Ctx−T−L−Sml−NeuN−C4
7614−Ctx−T−L−Sml−NeuN−C5
7614−Ctx−T−L−Sml−NeuN−C6
7614−Ctx−T−L−Sml−NeuN−D1
7614−Ctx−T−L−Sml−NeuN−D2
7614−Ctx−T−L−Sml−NeuN−D3
7614−Ctx−T−L−Sml−NeuN−D4
7614−Ctx−T−L−Sml−NeuN−D5
7614−Ctx−T−L−Sml−NeuN−D6
7614−Ctx−T−L−Sml−NeuN−E1
7614−Ctx−T−L−Sml−NeuN−E2
7614−Ctx−T−L−Sml−NeuN−E3
7614−Ctx−T−L−Sml−NeuN−E4
7614−Ctx−T−L−Sml−NeuN−E5
7614−Ctx−T−L−Sml−NeuN−E6
7614−Ctx−T−L−Sml−NeuN−F1
7614−Ctx−T−L−Sml−NeuN−F2
7614−Ctx−T−L−Sml−NeuN−F3
7614−Ctx−T−L−Sml−NeuN−F4
7614−Ctx−T−L−Sml−NeuN−F5
7614−Ctx−T−L−Sml−NeuN−F6
7614−Ctx−T−L−Sml−NeuN−G1
7614−Ctx−T−L−Sml−NeuN−G2
7614−Ctx−T−L−Sml−NeuN−G3
7614−Ctx−T−L−Sml−NeuN−G4
7614−Ctx−T−L−Sml−NeuN−G5
7614−Ctx−T−L−Sml−NeuN−G6
7614−Ctx−T−L−Sml−NeuN−H1
7614−Ctx−T−L−Sml−NeuN−H2
7614−Ctx−T−L−Sml−NeuN−H3
7614−Ctx−T−L−Sml−NeuN−H4
7614−Ctx−T−L−Sml−NeuN−H5
7614−Ctx−T−L−Sml−NeuN−H6
JGG-Cells

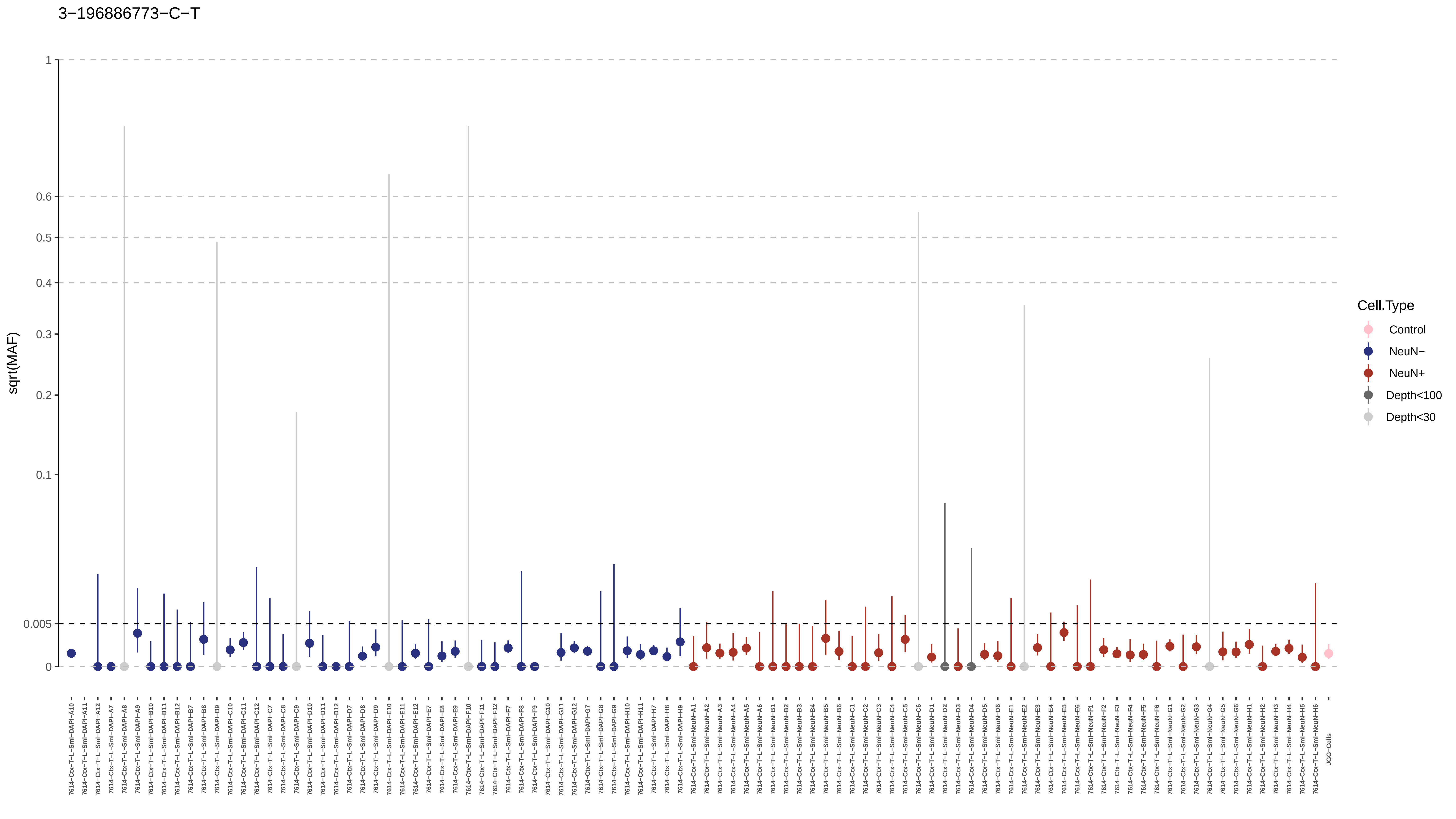
3−196886773−C−T
sqrt(MAF)
1
0.6
0.5
0.4
0.3
0.2
0.1
0.005
0
Cell.Type
Control
NeuN−
NeuN+
Depth<100
Depth<30

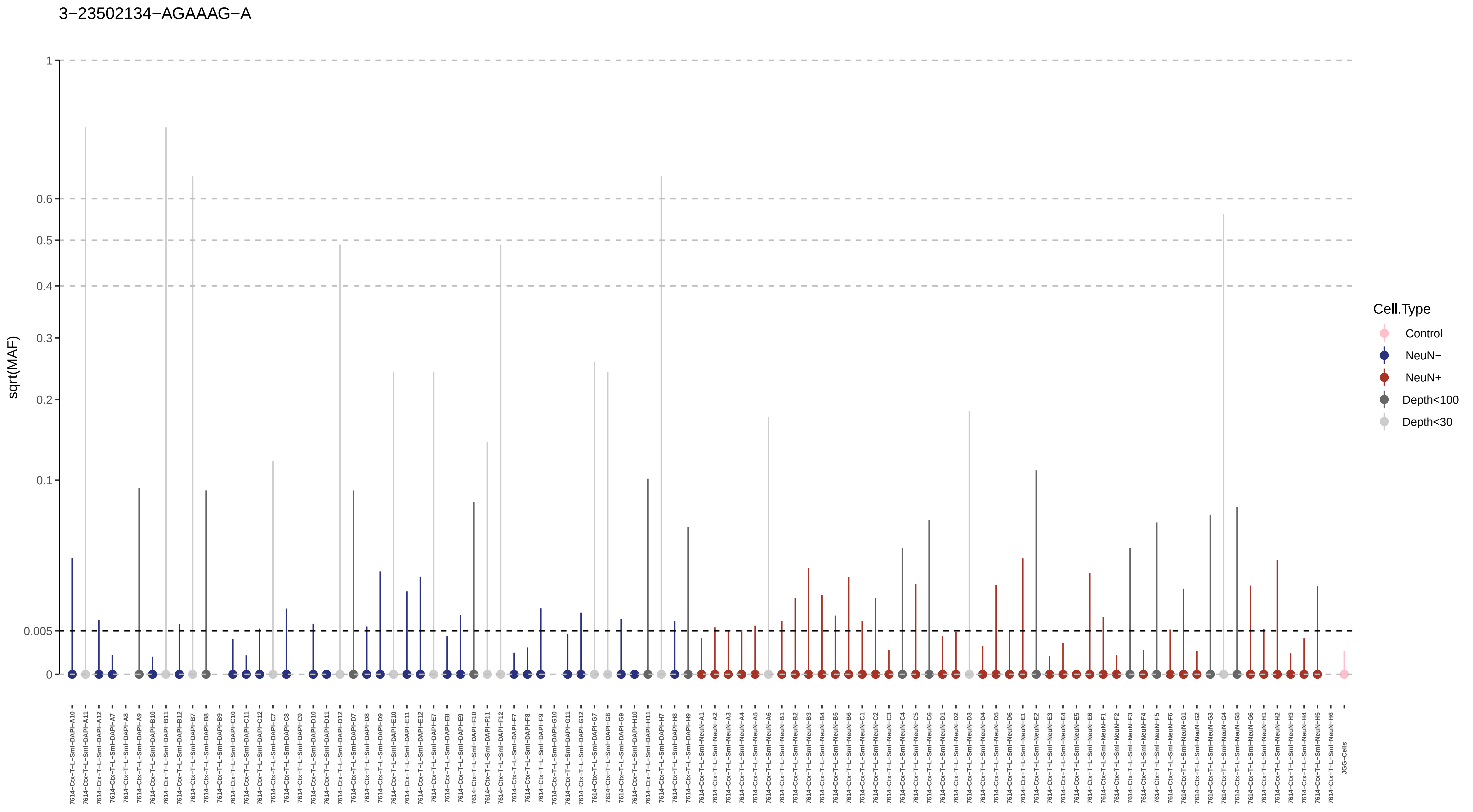
3−23502134−AGAAAG−A
sqrt(MAF)
1
0.6
0.5
0.4
0.3
0.2
0.1
0.005
0
Cell.Type
Control
NeuN−
NeuN+
Depth<100
Depth<30
JGG-Cells

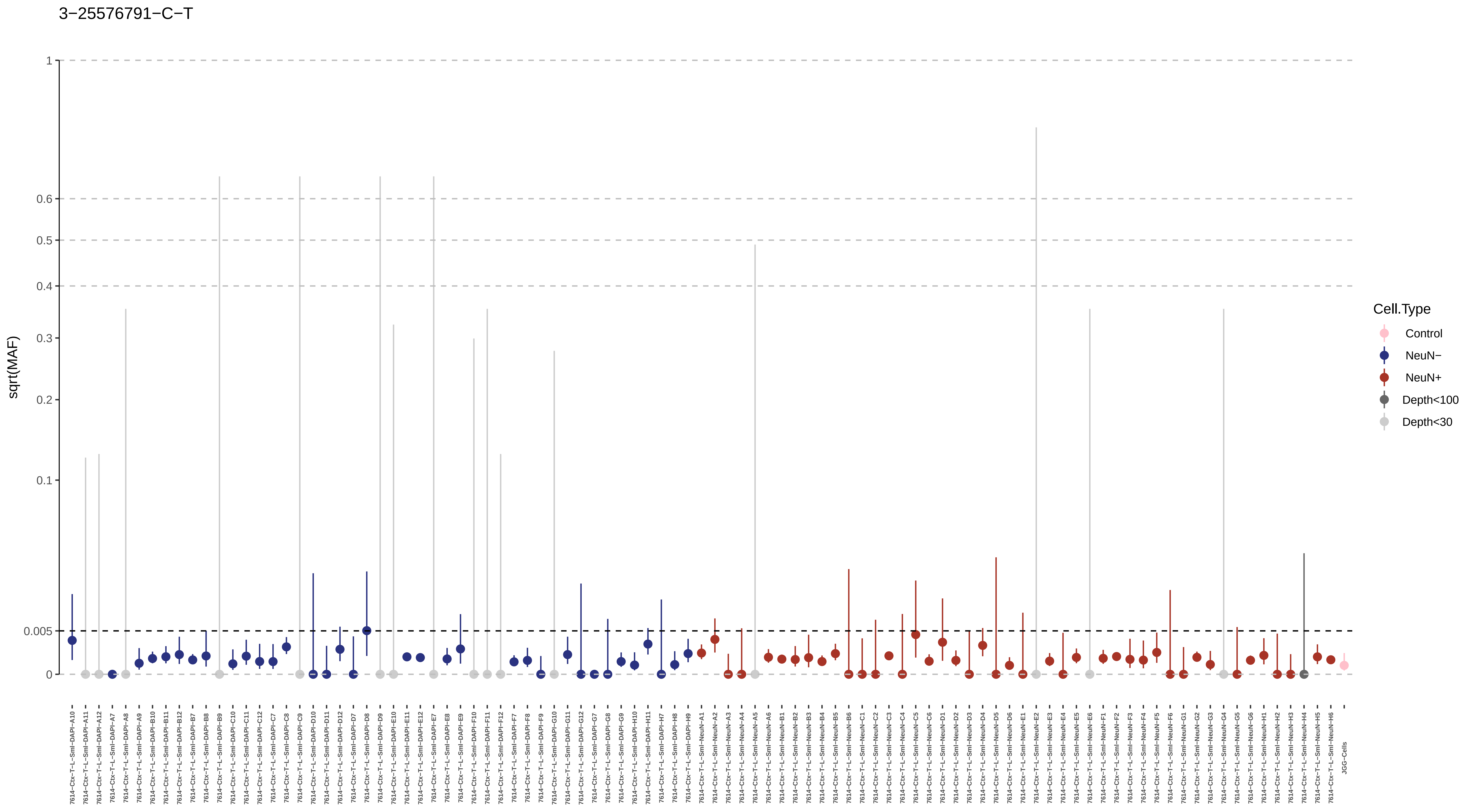
3−25576791−C−T
sqrt(MAF)
1
0.6
0.5
0.4
0.3
0.2
0.1
0.005
0
Cell.Type
Control
NeuN−
NeuN+
Depth<100
Depth<30
JGG-Cells

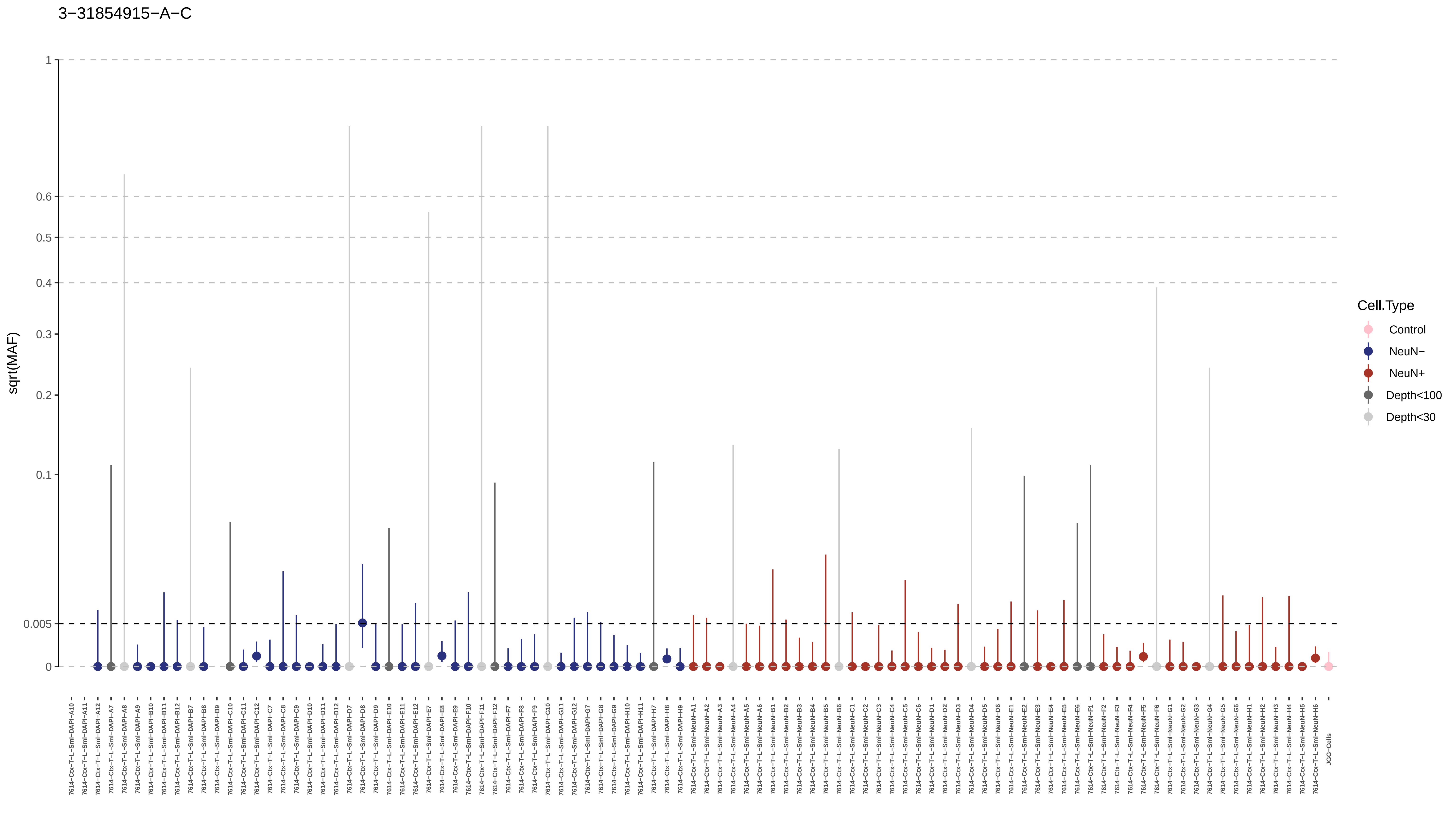
3−31854915−A−C
sqrt(MAF)
1
0.6
0.5
0.4
0.3
0.2
0.1
0.005
0
Cell.Type
Control
NeuN−
NeuN+
Depth<100
Depth<30
7614−Ctx−T−L−Sml−DAPI−A10
7614−Ctx−T−L−Sml−DAPI−A11
7614−Ctx−T−L−Sml−DAPI−A12
7614−Ctx−T−L−Sml−DAPI−A7
7614−Ctx−T−L−Sml−DAPI−A8
7614−Ctx−T−L−Sml−DAPI−A9
7614−Ctx−T−L−Sml−DAPI−B10
7614−Ctx−T−L−Sml−DAPI−B11
7614−Ctx−T−L−Sml−DAPI−B12
7614−Ctx−T−L−Sml−DAPI−B7
7614−Ctx−T−L−Sml−DAPI−B8
7614−Ctx−T−L−Sml−DAPI−B9
7614−Ctx−T−L−Sml−DAPI−C10
7614−Ctx−T−L−Sml−DAPI−C11
7614−Ctx−T−L−Sml−DAPI−C12
7614−Ctx−T−L−Sml−DAPI−C7
7614−Ctx−T−L−Sml−DAPI−C8
7614−Ctx−T−L−Sml−DAPI−C9
7614−Ctx−T−L−Sml−DAPI−D10
7614−Ctx−T−L−Sml−DAPI−D11
7614−Ctx−T−L−Sml−DAPI−D12
7614−Ctx−T−L−Sml−DAPI−D7
7614−Ctx−T−L−Sml−DAPI−D8
7614−Ctx−T−L−Sml−DAPI−D9
7614−Ctx−T−L−Sml−DAPI−E10
7614−Ctx−T−L−Sml−DAPI−E11
7614−Ctx−T−L−Sml−DAPI−E12
7614−Ctx−T−L−Sml−DAPI−E7
7614−Ctx−T−L−Sml−DAPI−E8
7614−Ctx−T−L−Sml−DAPI−E9
7614−Ctx−T−L−Sml−DAPI−F10
7614−Ctx−T−L−Sml−DAPI−F11
7614−Ctx−T−L−Sml−DAPI−F12
7614−Ctx−T−L−Sml−DAPI−F7
7614−Ctx−T−L−Sml−DAPI−F8
7614−Ctx−T−L−Sml−DAPI−F9
7614−Ctx−T−L−Sml−DAPI−G10
7614−Ctx−T−L−Sml−DAPI−G11
7614−Ctx−T−L−Sml−DAPI−G12
7614−Ctx−T−L−Sml−DAPI−G7
7614−Ctx−T−L−Sml−DAPI−G8
7614−Ctx−T−L−Sml−DAPI−G9
7614−Ctx−T−L−Sml−DAPI−H10
7614−Ctx−T−L−Sml−DAPI−H11
7614−Ctx−T−L−Sml−DAPI−H7
7614−Ctx−T−L−Sml−DAPI−H8
7614−Ctx−T−L−Sml−DAPI−H9
7614−Ctx−T−L−Sml−NeuN−A1
7614−Ctx−T−L−Sml−NeuN−A2
7614−Ctx−T−L−Sml−NeuN−A3
7614−Ctx−T−L−Sml−NeuN−A4
7614−Ctx−T−L−Sml−NeuN−A5
7614−Ctx−T−L−Sml−NeuN−A6
7614−Ctx−T−L−Sml−NeuN−B1
7614−Ctx−T−L−Sml−NeuN−B2
7614−Ctx−T−L−Sml−NeuN−B3
7614−Ctx−T−L−Sml−NeuN−B4
7614−Ctx−T−L−Sml−NeuN−B5
7614−Ctx−T−L−Sml−NeuN−B6
7614−Ctx−T−L−Sml−NeuN−C1
7614−Ctx−T−L−Sml−NeuN−C2
7614−Ctx−T−L−Sml−NeuN−C3
7614−Ctx−T−L−Sml−NeuN−C4
7614−Ctx−T−L−Sml−NeuN−C5
7614−Ctx−T−L−Sml−NeuN−C6
7614−Ctx−T−L−Sml−NeuN−D1
7614−Ctx−T−L−Sml−NeuN−D2
7614−Ctx−T−L−Sml−NeuN−D3
7614−Ctx−T−L−Sml−NeuN−D4
7614−Ctx−T−L−Sml−NeuN−D5
7614−Ctx−T−L−Sml−NeuN−D6
7614−Ctx−T−L−Sml−NeuN−E1
7614−Ctx−T−L−Sml−NeuN−E2
7614−Ctx−T−L−Sml−NeuN−E3
7614−Ctx−T−L−Sml−NeuN−E4
7614−Ctx−T−L−Sml−NeuN−E5
7614−Ctx−T−L−Sml−NeuN−E6
7614−Ctx−T−L−Sml−NeuN−F1
7614−Ctx−T−L−Sml−NeuN−F2
7614−Ctx−T−L−Sml−NeuN−F3
7614−Ctx−T−L−Sml−NeuN−F4
7614−Ctx−T−L−Sml−NeuN−F5
7614−Ctx−T−L−Sml−NeuN−F6
7614−Ctx−T−L−Sml−NeuN−G1
7614−Ctx−T−L−Sml−NeuN−G2
7614−Ctx−T−L−Sml−NeuN−G3
7614−Ctx−T−L−Sml−NeuN−G4
7614−Ctx−T−L−Sml−NeuN−G5
7614−Ctx−T−L−Sml−NeuN−G6
7614−Ctx−T−L−Sml−NeuN−H1
7614−Ctx−T−L−Sml−NeuN−H2
7614−Ctx−T−L−Sml−NeuN−H3
7614−Ctx−T−L−Sml−NeuN−H4
7614−Ctx−T−L−Sml−NeuN−H5
7614−Ctx−T−L−Sml−NeuN−H6
JGG−Cells

3−34583283−C−T

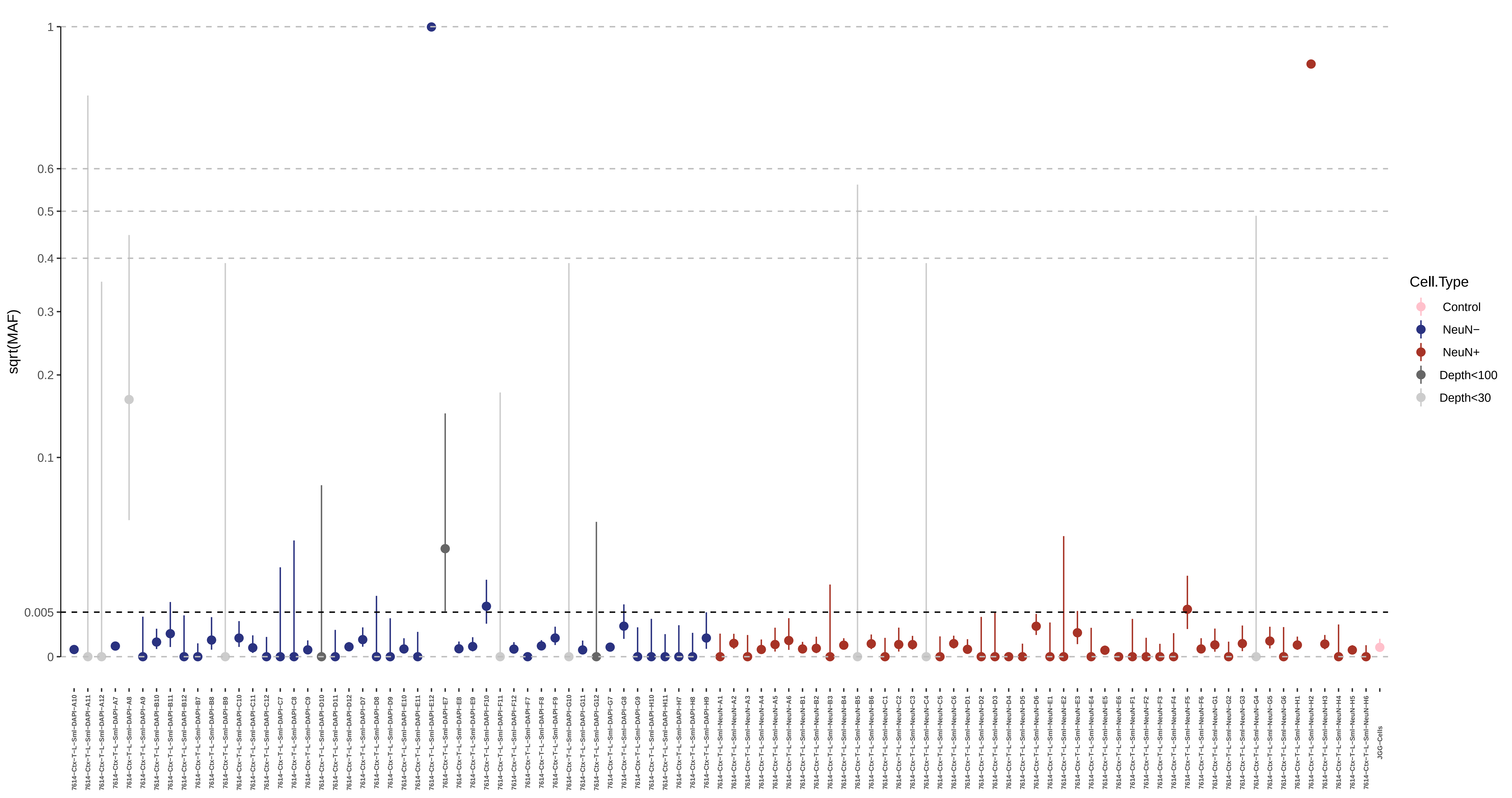

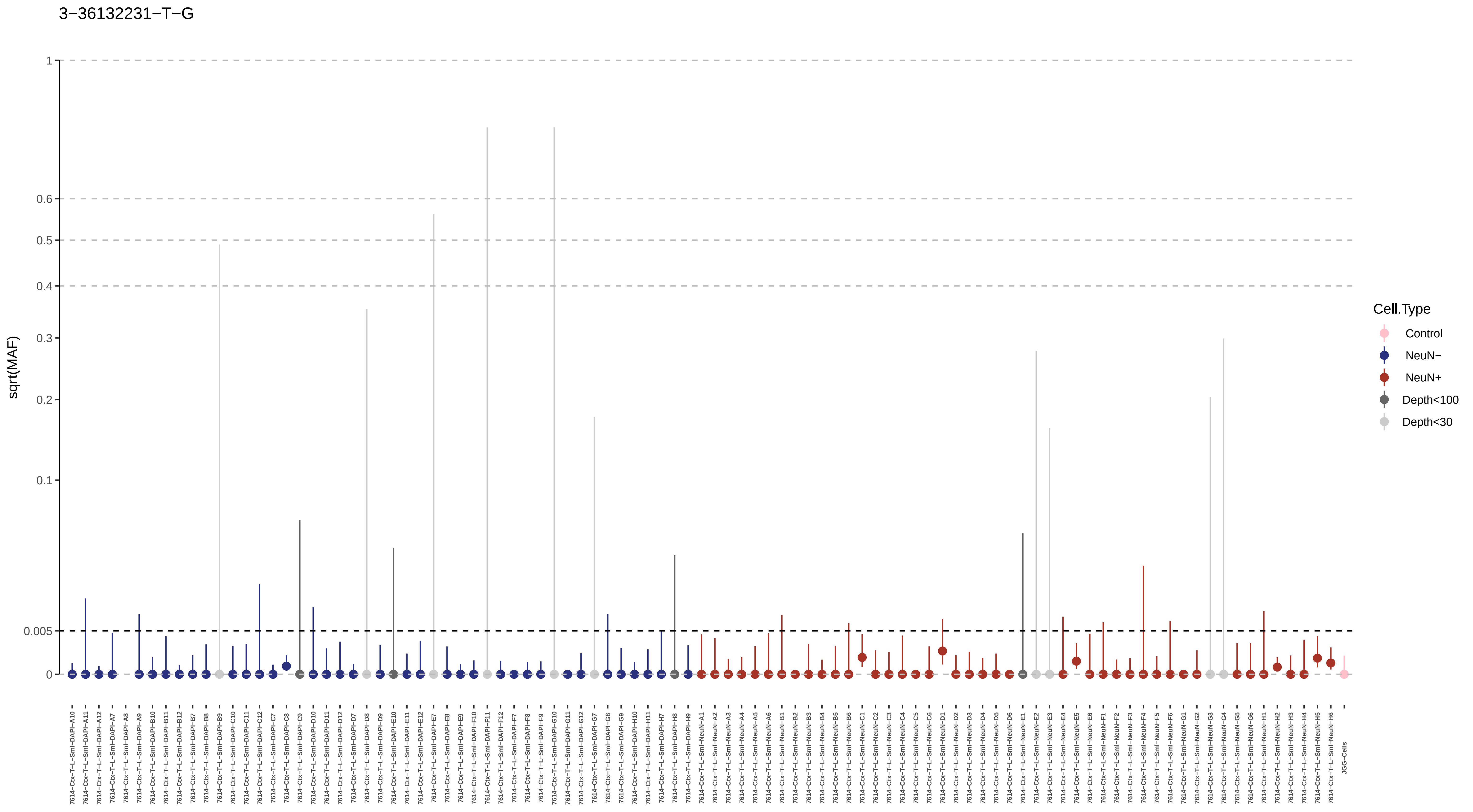

3−36132231−T−G
sqrt(MAF)
1
0.6
0.5
0.4
0.3
0.2
0.1
0.005
0
Cell.Type
Control
NeuN−
NeuN+
Depth<100
Depth<30
7614−Ctx−T−L−Sml−DAPI−A10
7614−Ctx−T−L−Sml−DAPI−A11
7614−Ctx−T−L−Sml−DAPI−A12
7614−Ctx−T−L−Sml−DAPI−A7
7614−Ctx−T−L−Sml−DAPI−A8
7614−Ctx−T−L−Sml−DAPI−A9
7614−Ctx−T−L−Sml−DAPI−B10
7614−Ctx−T−L−Sml−DAPI−B11
7614−Ctx−T−L−Sml−DAPI−B12
7614−Ctx−T−L−Sml−DAPI−B7
7614−Ctx−T−L−Sml−DAPI−B8
7614−Ctx−T−L−Sml−DAPI−B9
7614−Ctx−T−L−Sml−DAPI−C10
7614−Ctx−T−L−Sml−DAPI−C11
7614−Ctx−T−L−Sml−DAPI−C12
7614−Ctx−T−L−Sml−DAPI−C7
7614−Ctx−T−L−Sml−DAPI−C8
7614−Ctx−T−L−Sml−DAPI−C9
7614−Ctx−T−L−Sml−DAPI−D10
7614−Ctx−T−L−Sml−DAPI−D11
7614−Ctx−T−L−Sml−DAPI−D12
7614−Ctx−T−L−Sml−DAPI−D7
7614−Ctx−T−L−Sml−DAPI−D8
7614−Ctx−T−L−Sml−DAPI−D9
7614−Ctx−T−L−Sml−DAPI−E10
7614−Ctx−T−L−Sml−DAPI−E11
7614−Ctx−T−L−Sml−DAPI−E12
7614−Ctx−T−L−Sml−DAPI−E7
7614−Ctx−T−L−Sml−DAPI−E8
7614−Ctx−T−L−Sml−DAPI−E9
7614−Ctx−T−L−Sml−DAPI−F10
7614−Ctx−T−L−Sml−DAPI−F11
7614−Ctx−T−L−Sml−DAPI−F12
7614−Ctx−T−L−Sml−DAPI−F7
7614−Ctx−T−L−Sml−DAPI−F8
7614−Ctx−T−L−Sml−DAPI−F9
7614−Ctx−T−L−Sml−DAPI−G10
7614−Ctx−T−L−Sml−DAPI−G11
7614−Ctx−T−L−Sml−DAPI−G12
7614−Ctx−T−L−Sml−DAPI−G7
7614−Ctx−T−L−Sml−DAPI−G8
7614−Ctx−T−L−Sml−DAPI−G9
7614−Ctx−T−L−Sml−DAPI−H10
7614−Ctx−T−L−Sml−DAPI−H11
7614−Ctx−T−L−Sml−DAPI−H7
7614−Ctx−T−L−Sml−DAPI−H8
7614−Ctx−T−L−Sml−DAPI−H9
7614−Ctx−T−L−Sml−NeuN−A1
7614−Ctx−T−L−Sml−NeuN−A2
7614−Ctx−T−L−Sml−NeuN−A3
7614−Ctx−T−L−Sml−NeuN−A4
7614−Ctx−T−L−Sml−NeuN−A5
7614−Ctx−T−L−Sml−NeuN−A6
7614−Ctx−T−L−Sml−NeuN−B1
7614−Ctx−T−L−Sml−NeuN−B2
7614−Ctx−T−L−Sml−NeuN−B3
7614−Ctx−T−L−Sml−NeuN−B4
7614−Ctx−T−L−Sml−NeuN−B5
7614−Ctx−T−L−Sml−NeuN−B6
7614−Ctx−T−L−Sml−NeuN−C1
7614−Ctx−T−L−Sml−NeuN−C2
7614−Ctx−T−L−Sml−NeuN−C3
7614−Ctx−T−L−Sml−NeuN−C4
7614−Ctx−T−L−Sml−NeuN−C5
7614−Ctx−T−L−Sml−NeuN−C6
7614−Ctx−T−L−Sml−NeuN−D1
7614−Ctx−T−L−Sml−NeuN−D2
7614−Ctx−T−L−Sml−NeuN−D3
7614−Ctx−T−L−Sml−NeuN−D4
7614−Ctx−T−L−Sml−NeuN−D5
7614−Ctx−T−L−Sml−NeuN−D6
7614−Ctx−T−L−Sml−NeuN−E1
7614−Ctx−T−L−Sml−NeuN−E2
7614−Ctx−T−L−Sml−NeuN−E3
7614−Ctx−T−L−Sml−NeuN−E4
7614−Ctx−T−L−Sml−NeuN−E5
7614−Ctx−T−L−Sml−NeuN−E6
7614−Ctx−T−L−Sml−NeuN−F1
7614−Ctx−T−L−Sml−NeuN−F2
7614−Ctx−T−L−Sml−NeuN−F3
7614−Ctx−T−L−Sml−NeuN−F4
7614−Ctx−T−L−Sml−NeuN−F5
7614−Ctx−T−L−Sml−NeuN−F6
7614−Ctx−T−L−Sml−NeuN−G1
7614−Ctx−T−L−Sml−NeuN−G2
7614−Ctx−T−L−Sml−NeuN−G3
7614−Ctx−T−L−Sml−NeuN−G4
7614−Ctx−T−L−Sml−NeuN−G5
7614−Ctx−T−L−Sml−NeuN−G6
7614−Ctx−T−L−Sml−NeuN−H1
7614−Ctx−T−L−Sml−NeuN−H2
7614−Ctx−T−L−Sml−NeuN−H3
7614−Ctx−T−L−Sml−NeuN−H4
7614−Ctx−T−L−Sml−NeuN−H5
7614−Ctx−T−L−Sml−NeuN−H6
JGG−Cells

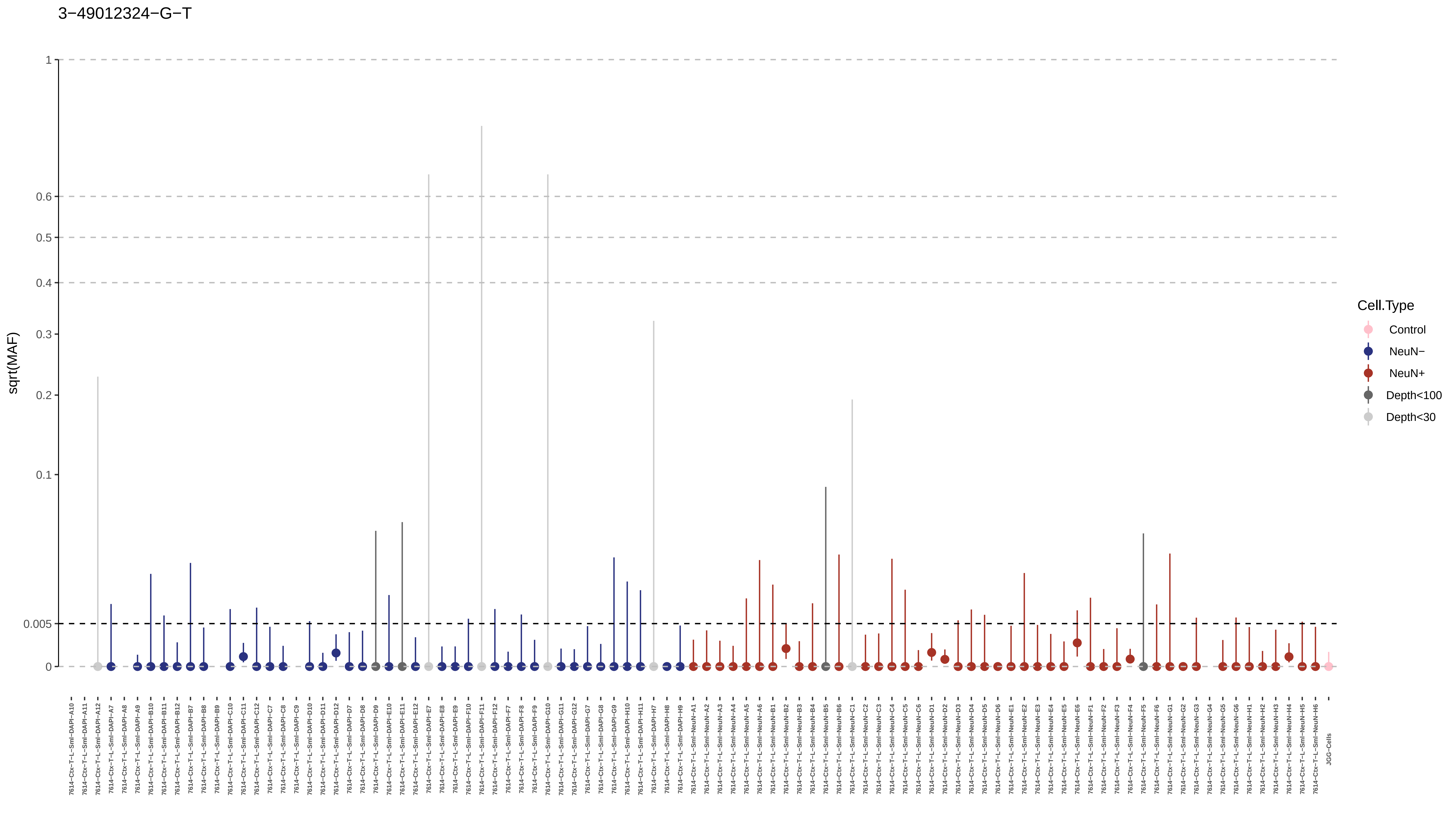

3−49012324−G−T
sqrt(MAF)
1
0.6
0.5
0.4
0.3
0.2
0.1
0.005
0
Cell.Type
Control
NeuN−
NeuN+
Depth<100
Depth<30
7614−Ctx−T−L−Sml−DAPI−A10
7614−Ctx−T−L−Sml−DAPI−A11
7614−Ctx−T−L−Sml−DAPI−A12
7614−Ctx−T−L−Sml−DAPI−A7
7614−Ctx−T−L−Sml−DAPI−A8
7614−Ctx−T−L−Sml−DAPI−A9
7614−Ctx−T−L−Sml−DAPI−B10
7614−Ctx−T−L−Sml−DAPI−B11
7614−Ctx−T−L−Sml−DAPI−B12
7614−Ctx−T−L−Sml−DAPI−B7
7614−Ctx−T−L−Sml−DAPI−B8
7614−Ctx−T−L−Sml−DAPI−B9
7614−Ctx−T−L−Sml−DAPI−C10
7614−Ctx−T−L−Sml−DAPI−C11
7614−Ctx−T−L−Sml−DAPI−C12
7614−Ctx−T−L−Sml−DAPI−C7
7614−Ctx−T−L−Sml−DAPI−C8
7614−Ctx−T−L−Sml−DAPI−C9
7614−Ctx−T−L−Sml−DAPI−D10
7614−Ctx−T−L−Sml−DAPI−D11
7614−Ctx−T−L−Sml−DAPI−D12
7614−Ctx−T−L−Sml−DAPI−D7
7614−Ctx−T−L−Sml−DAPI−D8
7614−Ctx−T−L−Sml−DAPI−D9
7614−Ctx−T−L−Sml−DAPI−E10
7614−Ctx−T−L−Sml−DAPI−E11
7614−Ctx−T−L−Sml−DAPI−E12
7614−Ctx−T−L−Sml−DAPI−E7
7614−Ctx−T−L−Sml−DAPI−E8
7614−Ctx−T−L−Sml−DAPI−E9
7614−Ctx−T−L−Sml−DAPI−F10
7614−Ctx−T−L−Sml−DAPI−F11
7614−Ctx−T−L−Sml−DAPI−F12
7614−Ctx−T−L−Sml−DAPI−F7
7614−Ctx−T−L−Sml−DAPI−F8
7614−Ctx−T−L−Sml−DAPI−F9
7614−Ctx−T−L−Sml−DAPI−G10
7614−Ctx−T−L−Sml−DAPI−G11
7614−Ctx−T−L−Sml−DAPI−G12
7614−Ctx−T−L−Sml−DAPI−G7
7614−Ctx−T−L−Sml−DAPI−G8
7614−Ctx−T−L−Sml−DAPI−G9
7614−Ctx−T−L−Sml−DAPI−H10
7614−Ctx−T−L−Sml−DAPI−H11
7614−Ctx−T−L−Sml−DAPI−H7
7614−Ctx−T−L−Sml−DAPI−H8
7614−Ctx−T−L−Sml−DAPI−H9
7614−Ctx−T−L−Sml−NeuN−A1
7614−Ctx−T−L−Sml−NeuN−A2
7614−Ctx−T−L−Sml−NeuN−A3
7614−Ctx−T−L−Sml−NeuN−A4
7614−Ctx−T−L−Sml−NeuN−A5
7614−Ctx−T−L−Sml−NeuN−A6
7614−Ctx−T−L−Sml−NeuN−B1
7614−Ctx−T−L−Sml−NeuN−B2
7614−Ctx−T−L−Sml−NeuN−B3
7614−Ctx−T−L−Sml−NeuN−B4
7614−Ctx−T−L−Sml−NeuN−B5
7614−Ctx−T−L−Sml−NeuN−B6
7614−Ctx−T−L−Sml−NeuN−C1
7614−Ctx−T−L−Sml−NeuN−C2
7614−Ctx−T−L−Sml−NeuN−C3
7614−Ctx−T−L−Sml−NeuN−C4
7614−Ctx−T−L−Sml−NeuN−C5
7614−Ctx−T−L−Sml−NeuN−C6
7614−Ctx−T−L−Sml−NeuN−D1
7614−Ctx−T−L−Sml−NeuN−D2
7614−Ctx−T−L−Sml−NeuN−D3
7614−Ctx−T−L−Sml−NeuN−D4
7614−Ctx−T−L−Sml−NeuN−D5
7614−Ctx−T−L−Sml−NeuN−D6
7614−Ctx−T−L−Sml−NeuN−E1
7614−Ctx−T−L−Sml−NeuN−E2
7614−Ctx−T−L−Sml−NeuN−E3
7614−Ctx−T−L−Sml−NeuN−E4
7614−Ctx−T−L−Sml−NeuN−E5
7614−Ctx−T−L−Sml−NeuN−E6
7614−Ctx−T−L−Sml−NeuN−F1
7614−Ctx−T−L−Sml−NeuN−F2
7614−Ctx−T−L−Sml−NeuN−F3
7614−Ctx−T−L−Sml−NeuN−F4
7614−Ctx−T−L−Sml−NeuN−F5
7614−Ctx−T−L−Sml−NeuN−F6
7614−Ctx−T−L−Sml−NeuN−G1
7614−Ctx−T−L−Sml−NeuN−G2
7614−Ctx−T−L−Sml−NeuN−G3
7614−Ctx−T−L−Sml−NeuN−G4
7614−Ctx−T−L−Sml−NeuN−G5
7614−Ctx−T−L−Sml−NeuN−G6
7614−Ctx−T−L−Sml−NeuN−H1
7614−Ctx−T−L−Sml−NeuN−H2
7614−Ctx−T−L−Sml−NeuN−H3
7614−Ctx−T−L−Sml−NeuN−H4
7614−Ctx−T−L−Sml−NeuN−H5
7614−Ctx−T−L−Sml−NeuN−H6
JGG−Cells

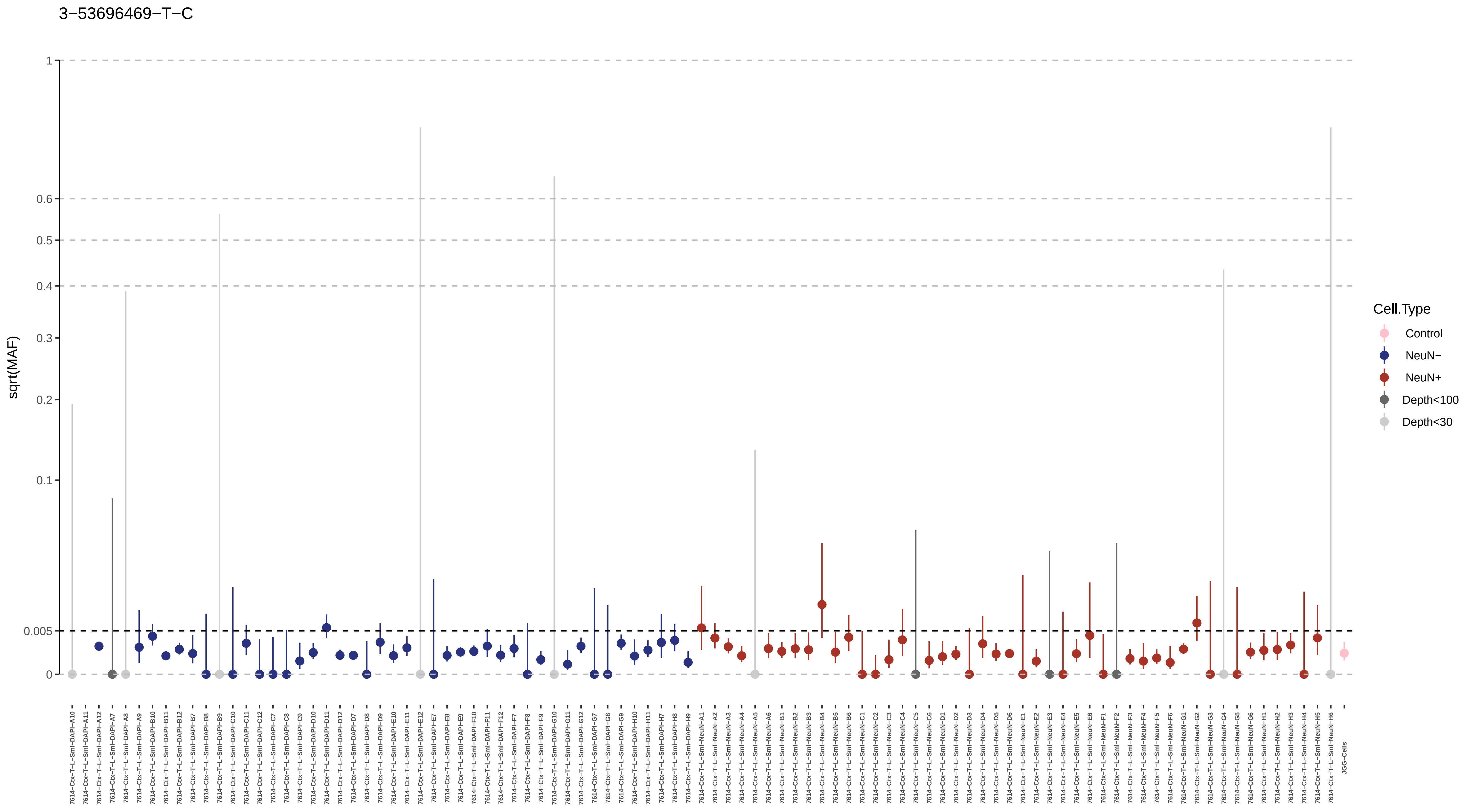
3−53696469−T−C
sqrt(MAF)
1
0.6
0.5
0.4
0.3
0.2
0.1
0.005
0
Cell.Type
Control
NeuN−
NeuN+
Depth<100
Depth<30
7614−Ctx−T−L−Sml−DAPI−A10
7614−Ctx−T−L−Sml−DAPI−A11
7614−Ctx−T−L−Sml−DAPI−A12
7614−Ctx−T−L−Sml−DAPI−A7
7614−Ctx−T−L−Sml−DAPI−A8
7614−Ctx−T−L−Sml−DAPI−A9
7614−Ctx−T−L−Sml−DAPI−B10
7614−Ctx−T−L−Sml−DAPI−B11
7614−Ctx−T−L−Sml−DAPI−B12
7614−Ctx−T−L−Sml−DAPI−B7
7614−Ctx−T−L−Sml−DAPI−B8
7614−Ctx−T−L−Sml−DAPI−B9
7614−Ctx−T−L−Sml−DAPI−C10
7614−Ctx−T−L−Sml−DAPI−C11
7614−Ctx−T−L−Sml−DAPI−C12
7614−Ctx−T−L−Sml−DAPI−C7
7614−Ctx−T−L−Sml−DAPI−C8
7614−Ctx−T−L−Sml−DAPI−C9
7614−Ctx−T−L−Sml−DAPI−D10
7614−Ctx−T−L−Sml−DAPI−D11
7614−Ctx−T−L−Sml−DAPI−D12
7614−Ctx−T−L−Sml−DAPI−D7
7614−Ctx−T−L−Sml−DAPI−D8
7614−Ctx−T−L−Sml−DAPI−D9
7614−Ctx−T−L−Sml−DAPI−E10
7614−Ctx−T−L−Sml−DAPI−E11
7614−Ctx−T−L−Sml−DAPI−E12
7614−Ctx−T−L−Sml−DAPI−E7
7614−Ctx−T−L−Sml−DAPI−E8
7614−Ctx−T−L−Sml−DAPI−E9
7614−Ctx−T−L−Sml−DAPI−F10
7614−Ctx−T−L−Sml−DAPI−F11
7614−Ctx−T−L−Sml−DAPI−F12
7614−Ctx−T−L−Sml−DAPI−F7
7614−Ctx−T−L−Sml−DAPI−F8
7614−Ctx−T−L−Sml−DAPI−F9
7614−Ctx−T−L−Sml−DAPI−G10
7614−Ctx−T−L−Sml−DAPI−G11
7614−Ctx−T−L−Sml−DAPI−G12
7614−Ctx−T−L−Sml−DAPI−G7
7614−Ctx−T−L−Sml−DAPI−G8
7614−Ctx−T−L−Sml−DAPI−G9
7614−Ctx−T−L−Sml−DAPI−H10
7614−Ctx−T−L−Sml−DAPI−H11
7614−Ctx−T−L−Sml−DAPI−H7
7614−Ctx−T−L−Sml−DAPI−H8
7614−Ctx−T−L−Sml−DAPI−H9
7614−Ctx−T−L−Sml−NeuN−A1
7614−Ctx−T−L−Sml−NeuN−A2
7614−Ctx−T−L−Sml−NeuN−A3
7614−Ctx−T−L−Sml−NeuN−A4
7614−Ctx−T−L−Sml−NeuN−A5
7614−Ctx−T−L−Sml−NeuN−A6
7614−Ctx−T−L−Sml−NeuN−B1
7614−Ctx−T−L−Sml−NeuN−B2
7614−Ctx−T−L−Sml−NeuN−B3
7614−Ctx−T−L−Sml−NeuN−B4
7614−Ctx−T−L−Sml−NeuN−B5
7614−Ctx−T−L−Sml−NeuN−B6
7614−Ctx−T−L−Sml−NeuN−C1
7614−Ctx−T−L−Sml−NeuN−C2
7614−Ctx−T−L−Sml−NeuN−C3
7614−Ctx−T−L−Sml−NeuN−C4
7614−Ctx−T−L−Sml−NeuN−C5
7614−Ctx−T−L−Sml−NeuN−C6
7614−Ctx−T−L−Sml−NeuN−D1
7614−Ctx−T−L−Sml−NeuN−D2
7614−Ctx−T−L−Sml−NeuN−D3
7614−Ctx−T−L−Sml−NeuN−D4
7614−Ctx−T−L−Sml−NeuN−D5
7614−Ctx−T−L−Sml−NeuN−D6
7614−Ctx−T−L−Sml−NeuN−E1
7614−Ctx−T−L−Sml−NeuN−E2
7614−Ctx−T−L−Sml−NeuN−E3
7614−Ctx−T−L−Sml−NeuN−E4
7614−Ctx−T−L−Sml−NeuN−E5
7614−Ctx−T−L−Sml−NeuN−E6
7614−Ctx−T−L−Sml−NeuN−F1
7614−Ctx−T−L−Sml−NeuN−F2
7614−Ctx−T−L−Sml−NeuN−F3
7614−Ctx−T−L−Sml−NeuN−F4
7614−Ctx−T−L−Sml−NeuN−F5
7614−Ctx−T−L−Sml−NeuN−F6
7614−Ctx−T−L−Sml−NeuN−G1
7614−Ctx−T−L−Sml−NeuN−G2
7614−Ctx−T−L−Sml−NeuN−G3
7614−Ctx−T−L−Sml−NeuN−G4
7614−Ctx−T−L−Sml−NeuN−G5
7614−Ctx−T−L−Sml−NeuN−G6
7614−Ctx−T−L−Sml−NeuN−H1
7614−Ctx−T−L−Sml−NeuN−H2
7614−Ctx−T−L−Sml−NeuN−H3
7614−Ctx−T−L−Sml−NeuN−H4
7614−Ctx−T−L−Sml−NeuN−H5
7614−Ctx−T−L−Sml−NeuN−H6
JGG-Cells

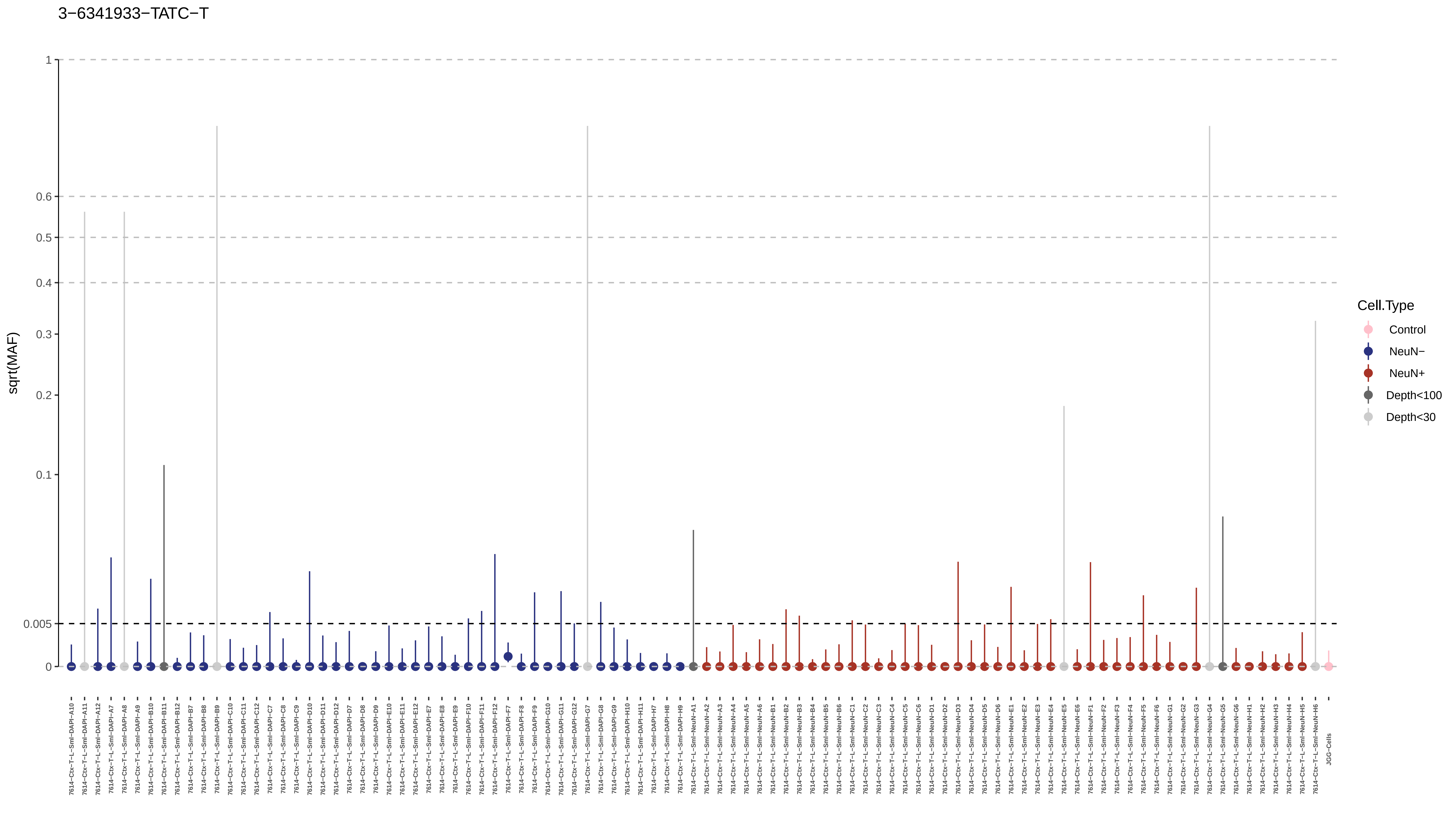
3−6341933−TATC−T
sqrt(MAF)
1
0.6
0.5
0.4
0.3
0.2
0.1
0.005
0
Cell.Type
Control
NeuN−
NeuN+
Depth<100
Depth<30

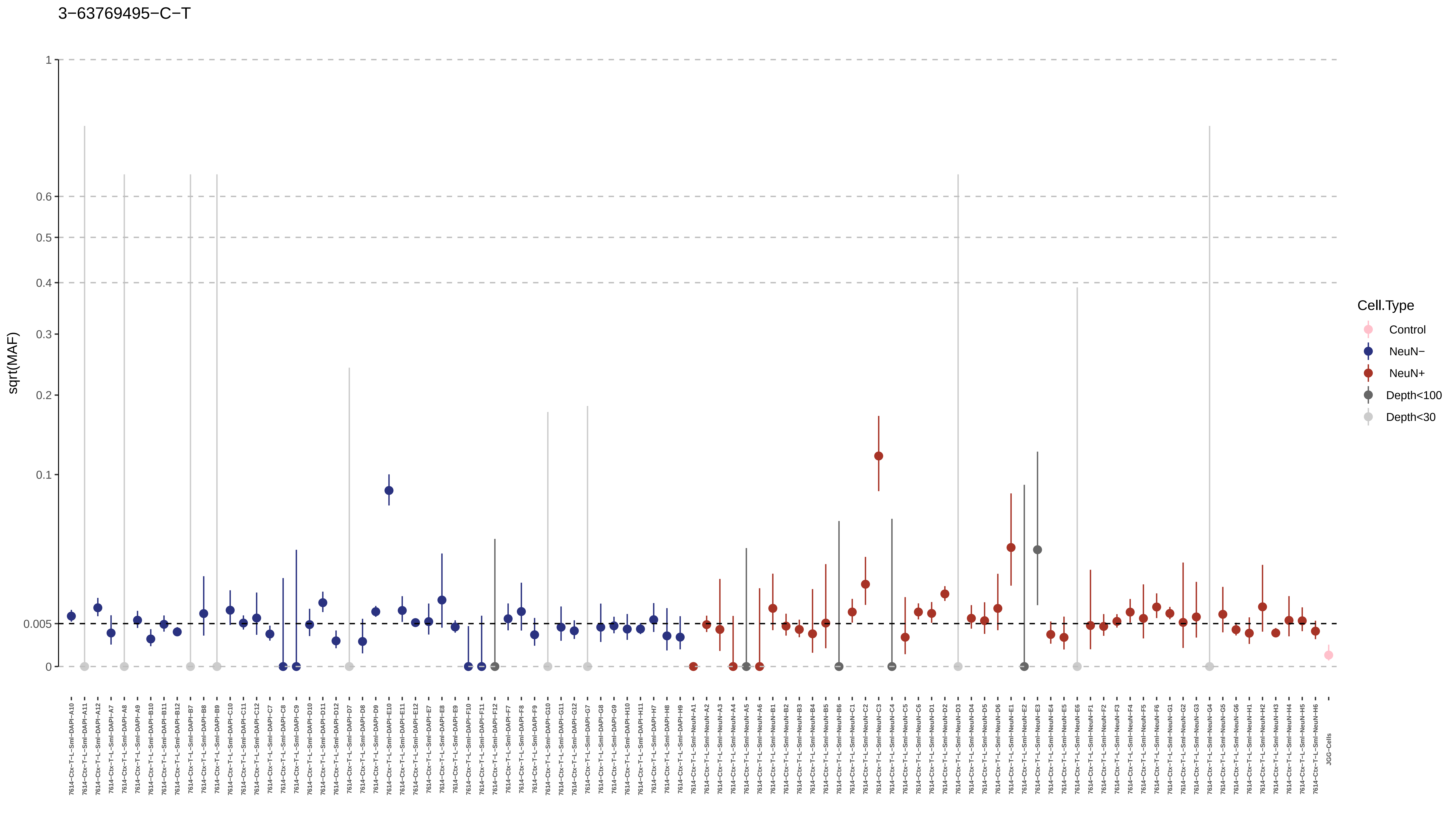

3−63769495−C−T
sqrt(MAF)
1
0.6
0.5
0.4
0.3
0.2
0.1
0.005
0
Cell.Type
Control
NeuN−
NeuN+
Depth<100
Depth<30
JGG-Cells

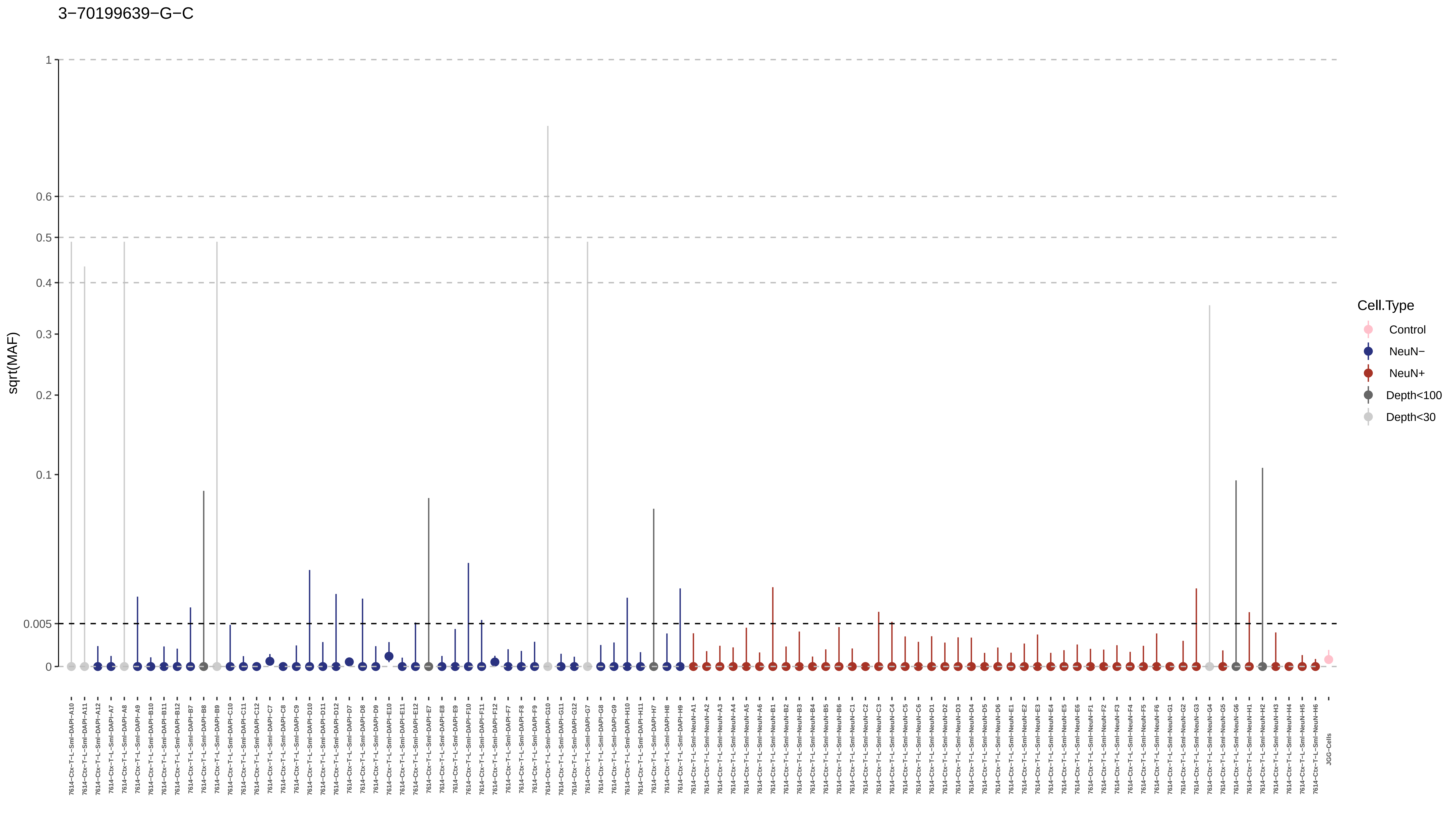
3−70199639−G−C
sqrt(MAF)
1
0.6
0.5
0.4
0.3
0.2
0.1
0.005
0
7614−Ctx−T−L−Sml−DAPI−A10
7614−Ctx−T−L−Sml−DAPI−A11
7614−Ctx−T−L−Sml−DAPI−A12
7614−Ctx−T−L−Sml−DAPI−A7
7614−Ctx−T−L−Sml−DAPI−A8
7614−Ctx−T−L−Sml−DAPI−A9
7614−Ctx−T−L−Sml−DAPI−B10
7614−Ctx−T−L−Sml−DAPI−B11
7614−Ctx−T−L−Sml−DAPI−B12
7614−Ctx−T−L−Sml−DAPI−B7
7614−Ctx−T−L−Sml−DAPI−B8
7614−Ctx−T−L−Sml−DAPI−B9
7614−Ctx−T−L−Sml−DAPI−C10
7614−Ctx−T−L−Sml−DAPI−C11
7614−Ctx−T−L−Sml−DAPI−C12
7614−Ctx−T−L−Sml−DAPI−C7
7614−Ctx−T−L−Sml−DAPI−C8
7614−Ctx−T−L−Sml−DAPI−C9
7614−Ctx−T−L−Sml−DAPI−D10
7614−Ctx−T−L−Sml−DAPI−D11
7614−Ctx−T−L−Sml−DAPI−D12
7614−Ctx−T−L−Sml−DAPI−D7
7614−Ctx−T−L−Sml−DAPI−D8
7614−Ctx−T−L−Sml−DAPI−D9
7614−Ctx−T−L−Sml−DAPI−E10
7614−Ctx−T−L−Sml−DAPI−E11
7614−Ctx−T−L−Sml−DAPI−E12
7614−Ctx−T−L−Sml−DAPI−E7
7614−Ctx−T−L−Sml−DAPI−E8
7614−Ctx−T−L−Sml−DAPI−E9
7614−Ctx−T−L−Sml−DAPI−F10
7614−Ctx−T−L−Sml−DAPI−F11
7614−Ctx−T−L−Sml−DAPI−F12
7614−Ctx−T−L−Sml−DAPI−F7
7614−Ctx−T−L−Sml−DAPI−F8
7614−Ctx−T−L−Sml−DAPI−F9
7614−Ctx−T−L−Sml−DAPI−G10
7614−Ctx−T−L−Sml−DAPI−G11
7614−Ctx−T−L−Sml−DAPI−G12
7614−Ctx−T−L−Sml−DAPI−G7
7614−Ctx−T−L−Sml−DAPI−G8
7614−Ctx−T−L−Sml−DAPI−G9
7614−Ctx−T−L−Sml−DAPI−H10
7614−Ctx−T−L−Sml−DAPI−H11
7614−Ctx−T−L−Sml−DAPI−H7
7614−Ctx−T−L−Sml−DAPI−H8
7614−Ctx−T−L−Sml−DAPI−H9
7614−Ctx−T−L−Sml−NeuN−A1
7614−Ctx−T−L−Sml−NeuN−A2
7614−Ctx−T−L−Sml−NeuN−A3
7614−Ctx−T−L−Sml−NeuN−A4
7614−Ctx−T−L−Sml−NeuN−A5
7614−Ctx−T−L−Sml−NeuN−A6
7614−Ctx−T−L−Sml−NeuN−B1
7614−Ctx−T−L−Sml−NeuN−B2
7614−Ctx−T−L−Sml−NeuN−B3
7614−Ctx−T−L−Sml−NeuN−B4
7614−Ctx−T−L−Sml−NeuN−B5
7614−Ctx−T−L−Sml−NeuN−B6
7614−Ctx−T−L−Sml−NeuN−C1
7614−Ctx−T−L−Sml−NeuN−C2
7614−Ctx−T−L−Sml−NeuN−C3
7614−Ctx−T−L−Sml−NeuN−C4
7614−Ctx−T−L−Sml−NeuN−C5
7614−Ctx−T−L−Sml−NeuN−C6
7614−Ctx−T−L−Sml−NeuN−D1
7614−Ctx−T−L−Sml−NeuN−D2
7614−Ctx−T−L−Sml−NeuN−D3
7614−Ctx−T−L−Sml−NeuN−D4
7614−Ctx−T−L−Sml−NeuN−D5
7614−Ctx−T−L−Sml−NeuN−D6
7614−Ctx−T−L−Sml−NeuN−E1
7614−Ctx−T−L−Sml−NeuN−E2
7614−Ctx−T−L−Sml−NeuN−E3
7614−Ctx−T−L−Sml−NeuN−E4
7614−Ctx−T−L−Sml−NeuN−E5
7614−Ctx−T−L−Sml−NeuN−E6
7614−Ctx−T−L−Sml−NeuN−F1
7614−Ctx−T−L−Sml−NeuN−F2
7614−Ctx−T−L−Sml−NeuN−F3
7614−Ctx−T−L−Sml−NeuN−F4
7614−Ctx−T−L−Sml−NeuN−F5
7614−Ctx−T−L−Sml−NeuN−F6
7614−Ctx−T−L−Sml−NeuN−G1
7614−Ctx−T−L−Sml−NeuN−G2
7614−Ctx−T−L−Sml−NeuN−G3
7614−Ctx−T−L−Sml−NeuN−G4
7614−Ctx−T−L−Sml−NeuN−G5
7614−Ctx−T−L−Sml−NeuN−G6
7614−Ctx−T−L−Sml−NeuN−H1
7614−Ctx−T−L−Sml−NeuN−H2
7614−Ctx−T−L−Sml−NeuN−H3
7614−Ctx−T−L−Sml−NeuN−H4
7614−Ctx−T−L−Sml−NeuN−H5
7614−Ctx−T−L−Sml−NeuN−H6
JGG−Cells
Cell.Type
Control
NeuN−
NeuN+
Depth<100
Depth<30

3−83951534−A−G

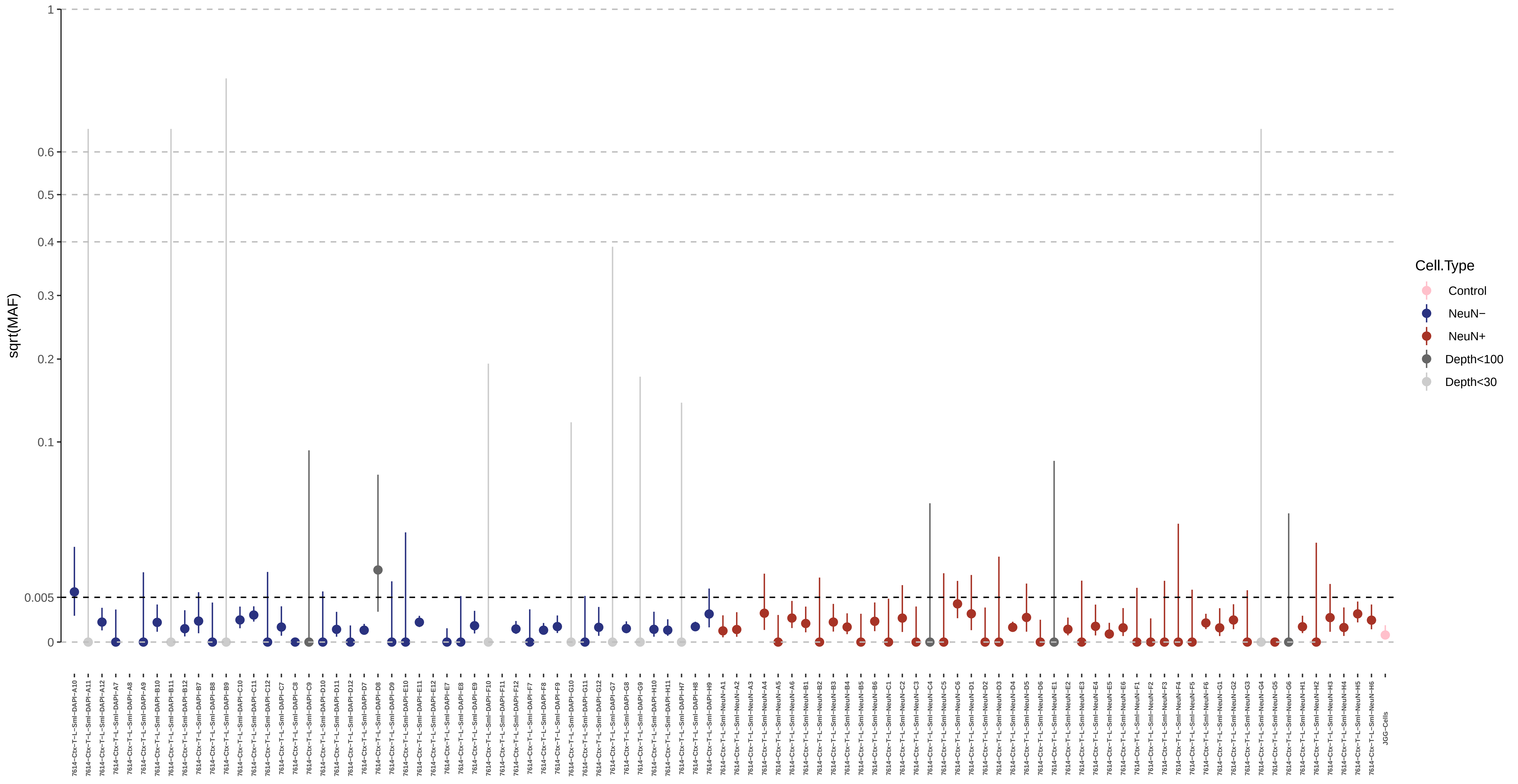

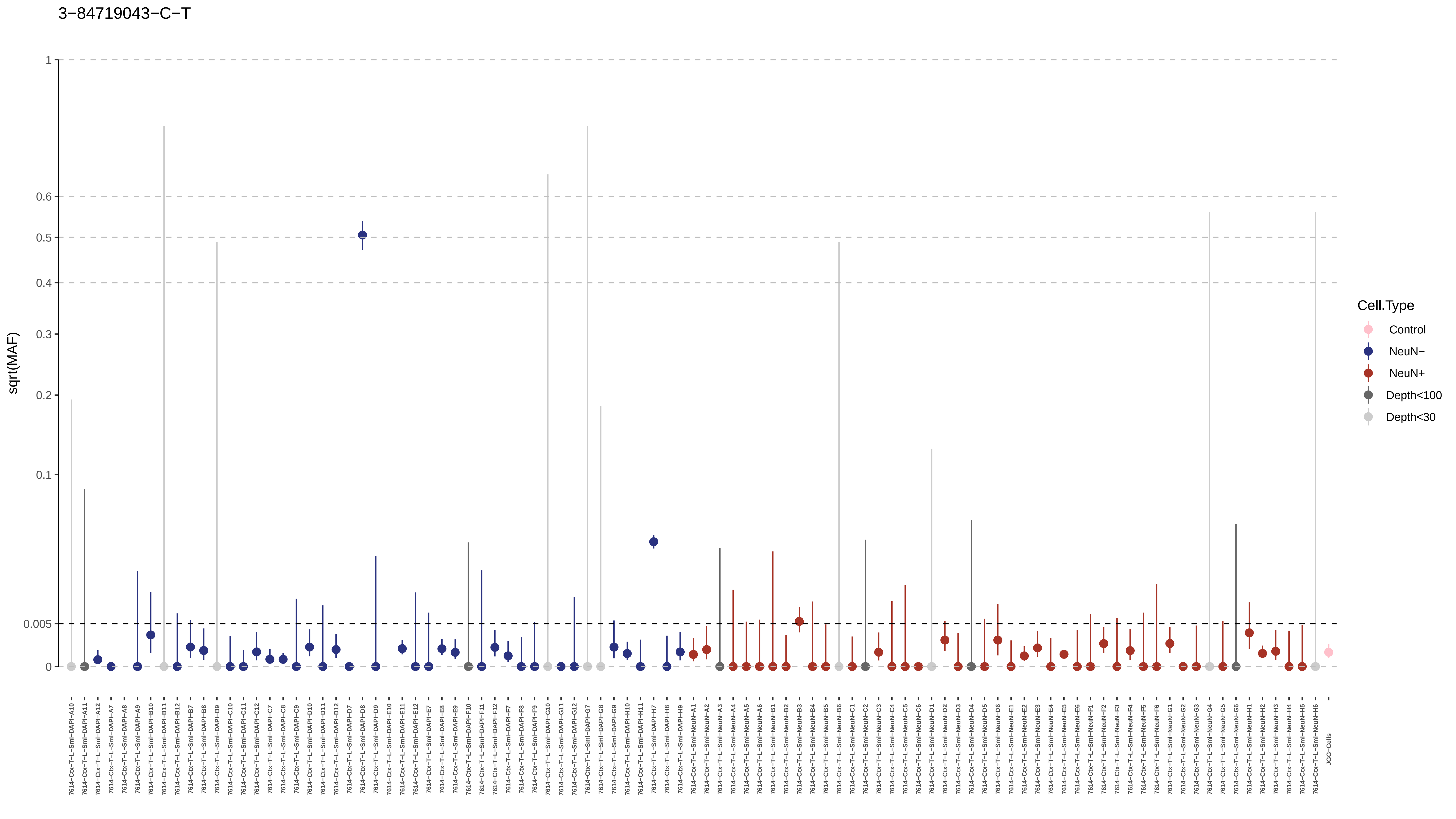

3−84719043−C−T
sqrt(MAF)
1
0.6
0.5
0.4
0.3
0.2
0.1
0.005
0
Cell.Type
Control
NeuN−
NeuN+
Depth<100
Depth<30

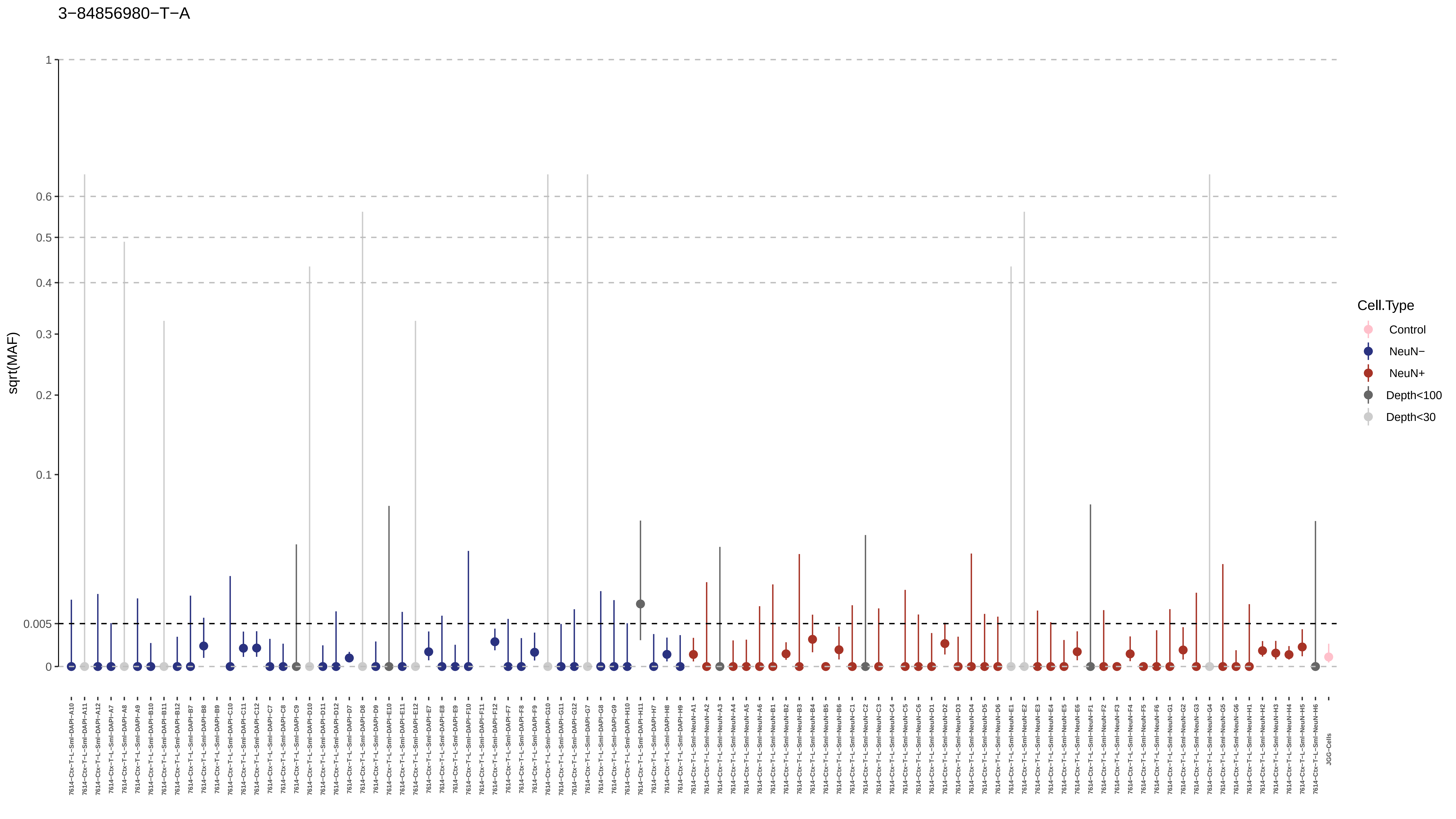
3−84856980−T−A
sqrt(MAF)
1
0.6
0.5
0.4
0.3
0.2
0.1
0.005
0
Cell.Type
Control
NeuN−
NeuN+
Depth<100
Depth<30

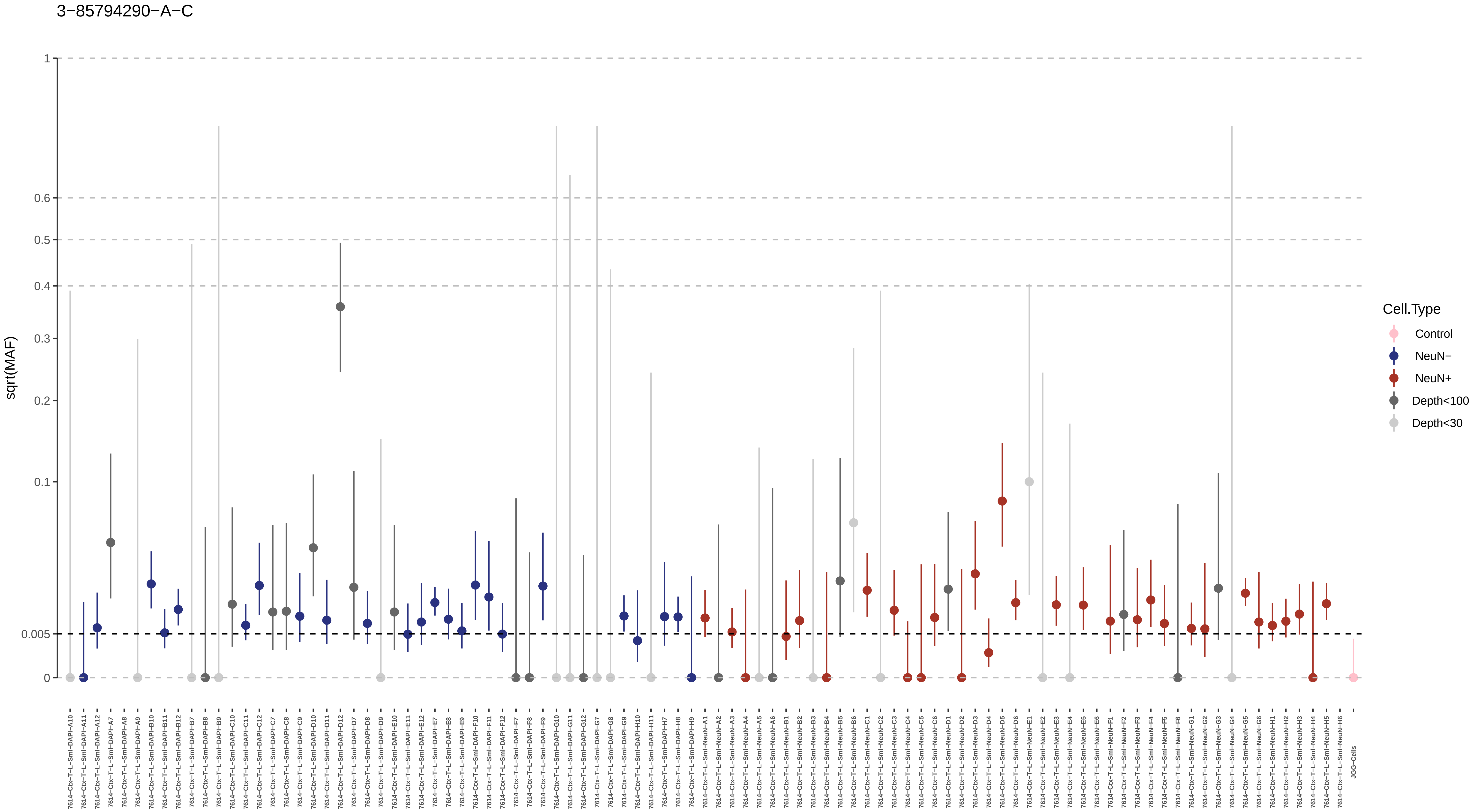
3−85794290−A−C
sqrt(MAF)
1
0.6
0.5
0.4
0.3
0.2
0.1
0.005
0
Cell.Type
Control
NeuN−
NeuN+
Depth<100
Depth<30

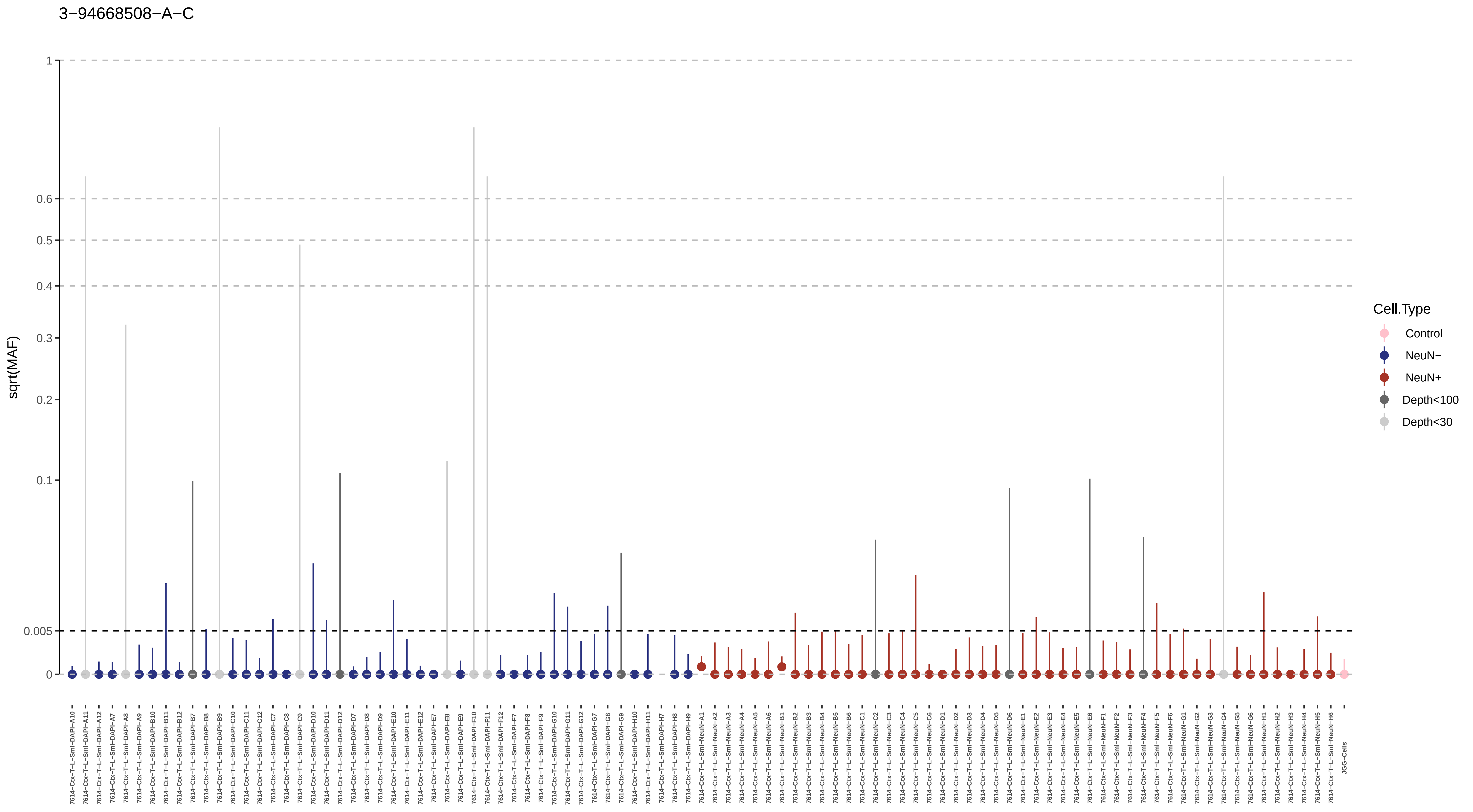
3−94668508−A−C
sqrt(MAF)
1
0.6
0.5
0.4
0.3
0.2
0.1
0.005
0
Cell.Type
Control
NeuN−
NeuN+
Depth<100
Depth<30
JGG-Cells

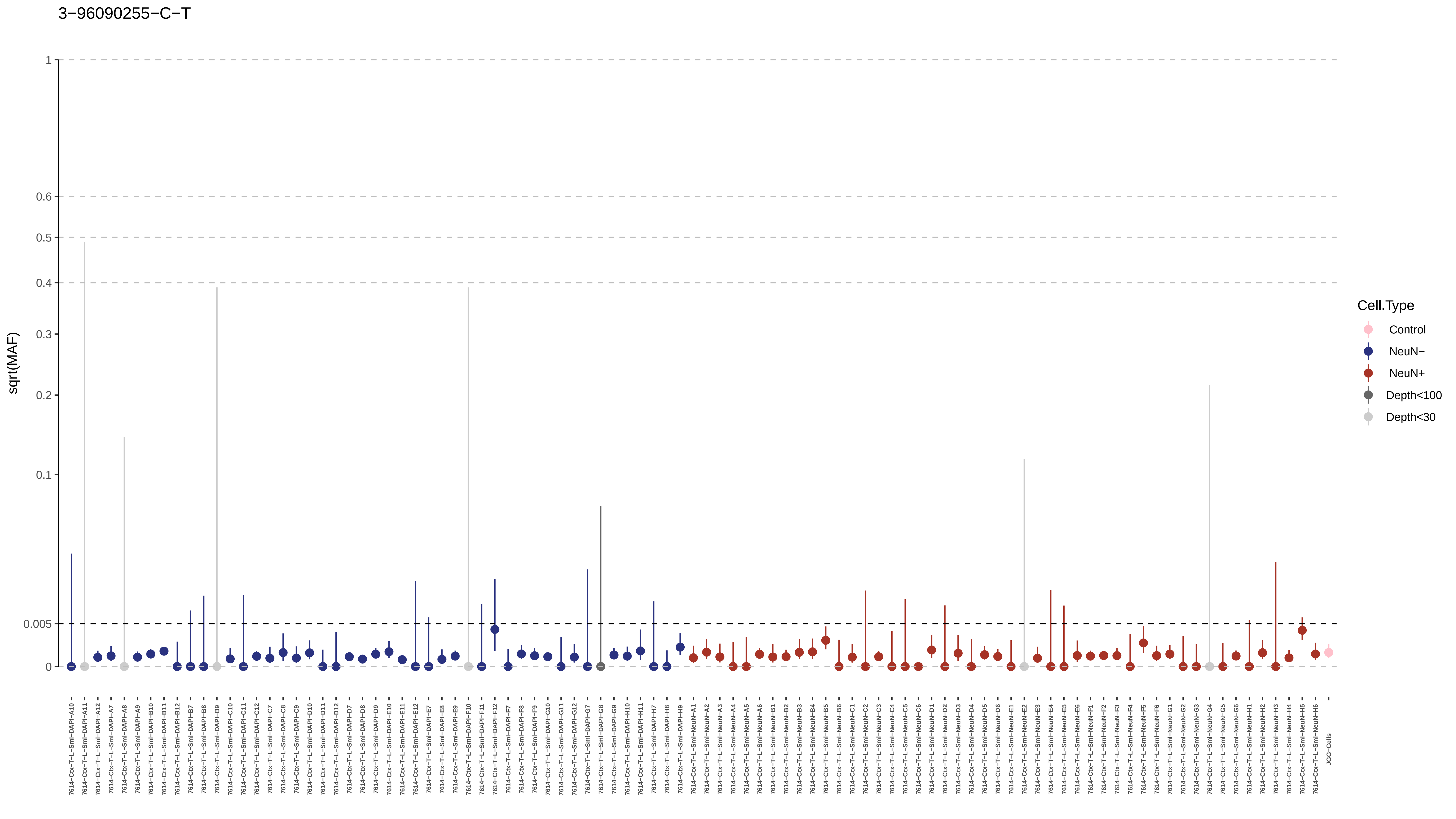

3-96090255-C-T
sqrt(MAF)
1
0.6
0.5
0.4
0.3
0.2
0.1
0.005
0
Cell.Type
Control
NeuN-
NeuN+
Depth<100
Depth<30
7614-Ctx-T-L-Sml-DAPI-A10
7614-Ctx-T-L-Sml-DAPI-A11
7614-Ctx-T-L-Sml-DAPI-A12
7614-Ctx-T-L-Sml-DAPI-A7
7614-Ctx-T-L-Sml-DAPI-A8
7614-Ctx-T-L-Sml-DAPI-A9
7614-Ctx-T-L-Sml-DAPI-B10
7614-Ctx-T-L-Sml-DAPI-B11
7614-Ctx-T-L-Sml-DAPI-B12
7614-Ctx-T-L-Sml-DAPI-B7
7614-Ctx-T-L-Sml-DAPI-B8
7614-Ctx-T-L-Sml-DAPI-B9
7614-Ctx-T-L-Sml-DAPI-C10
7614-Ctx-T-L-Sml-DAPI-C11
7614-Ctx-T-L-Sml-DAPI-C12
7614-Ctx-T-L-Sml-DAPI-C7
7614-Ctx-T-L-Sml-DAPI-C8
7614-Ctx-T-L-Sml-DAPI-C9
7614-Ctx-T-L-Sml-DAPI-D10
7614-Ctx-T-L-Sml-DAPI-D11
7614-Ctx-T-L-Sml-DAPI-D12
7614-Ctx-T-L-Sml-DAPI-D7
7614-Ctx-T-L-Sml-DAPI-D8
7614-Ctx-T-L-Sml-DAPI-D9
7614-Ctx-T-L-Sml-DAPI-E10
7614-Ctx-T-L-Sml-DAPI-E11
7614-Ctx-T-L-Sml-DAPI-E12
7614-Ctx-T-L-Sml-DAPI-E7
7614-Ctx-T-L-Sml-DAPI-E8
7614-Ctx-T-L-Sml-DAPI-E9
7614-Ctx-T-L-Sml-DAPI-F10
7614-Ctx-T-L-Sml-DAPI-F11
7614-Ctx-T-L-Sml-DAPI-F12
7614-Ctx-T-L-Sml-DAPI-F7
7614-Ctx-T-L-Sml-DAPI-F8
7614-Ctx-T-L-Sml-DAPI-F9
7614-Ctx-T-L-Sml-DAPI-G10
7614-Ctx-T-L-Sml-DAPI-G11
7614-Ctx-T-L-Sml-DAPI-G12
7614-Ctx-T-L-Sml-DAPI-G7
7614-Ctx-T-L-Sml-DAPI-G8
7614-Ctx-T-L-Sml-DAPI-G9
7614-Ctx-T-L-Sml-DAPI-H10
7614-Ctx-T-L-Sml-DAPI-H11
7614-Ctx-T-L-Sml-DAPI-H7
7614-Ctx-T-L-Sml-DAPI-H8
7614-Ctx-T-L-Sml-DAPI-H9
7614-Ctx-T-L-Sml-NeuN-A1
7614-Ctx-T-L-Sml-NeuN-A2
7614-Ctx-T-L-Sml-NeuN-A3
7614-Ctx-T-L-Sml-NeuN-A4
7614-Ctx-T-L-Sml-NeuN-A5
7614-Ctx-T-L-Sml-NeuN-A6
7614-Ctx-T-L-Sml-NeuN-B1
7614-Ctx-T-L-Sml-NeuN-B2
7614-Ctx-T-L-Sml-NeuN-B3
7614-Ctx-T-L-Sml-NeuN-B4
7614-Ctx-T-L-Sml-NeuN-B5
7614-Ctx-T-L-Sml-NeuN-B6
7614-Ctx-T-L-Sml-NeuN-C1
7614-Ctx-T-L-Sml-NeuN-C2
7614-Ctx-T-L-Sml-NeuN-C3
7614-Ctx-T-L-Sml-NeuN-C4
7614-Ctx-T-L-Sml-NeuN-C5
7614-Ctx-T-L-Sml-NeuN-C6
7614-Ctx-T-L-Sml-NeuN-D1
7614-Ctx-T-L-Sml-NeuN-D2
7614-Ctx-T-L-Sml-NeuN-D3
7614-Ctx-T-L-Sml-NeuN-D4
7614-Ctx-T-L-Sml-NeuN-D5
7614-Ctx-T-L-Sml-NeuN-D6
7614-Ctx-T-L-Sml-NeuN-E1
7614-Ctx-T-L-Sml-NeuN-E2
7614-Ctx-T-L-Sml-NeuN-E3
7614-Ctx-T-L-Sml-NeuN-E4
7614-Ctx-T-L-Sml-NeuN-E5
7614-Ctx-T-L-Sml-NeuN-E6
7614-Ctx-T-L-Sml-NeuN-F1
7614-Ctx-T-L-Sml-NeuN-F2
7614-Ctx-T-L-Sml-NeuN-F3
7614-Ctx-T-L-Sml-NeuN-F4
7614-Ctx-T-L-Sml-NeuN-F5
7614-Ctx-T-L-Sml-NeuN-F6
7614-Ctx-T-L-Sml-NeuN-G1
7614-Ctx-T-L-Sml-NeuN-G2
7614-Ctx-T-L-Sml-NeuN-G3
7614-Ctx-T-L-Sml-NeuN-G4
7614-Ctx-T-L-Sml-NeuN-G5
7614-Ctx-T-L-Sml-NeuN-G6
7614-Ctx-T-L-Sml-NeuN-H1
7614-Ctx-T-L-Sml-NeuN-H2
7614-Ctx-T-L-Sml-NeuN-H3
7614-Ctx-T-L-Sml-NeuN-H4
7614-Ctx-T-L-Sml-NeuN-H5
7614-Ctx-T-L-Sml-NeuN-H6
JGG-Cells

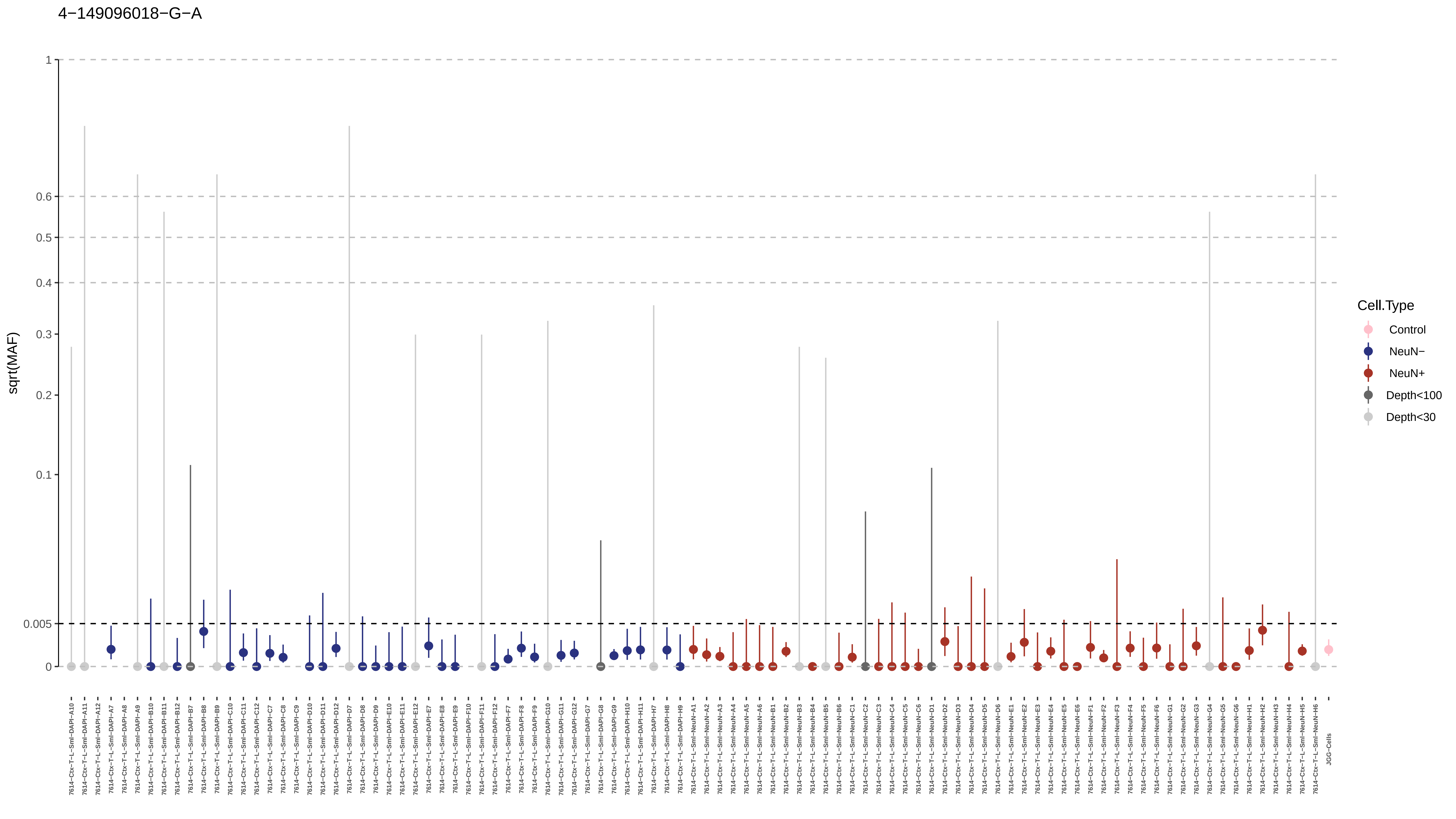
4−149096018−G−A
sqrt(MAF)
1
0.6
0.5
0.4
0.3
0.2
0.1
0.005
0
Cell.Type
Control
NeuN−
NeuN+
Depth<100
Depth<30
7614−Ctx−T−L−Sml−DAPI−A10
7614−Ctx−T−L−Sml−DAPI−A11
7614−Ctx−T−L−Sml−DAPI−A12
7614−Ctx−T−L−Sml−DAPI−A7
7614−Ctx−T−L−Sml−DAPI−A8
7614−Ctx−T−L−Sml−DAPI−A9
7614−Ctx−T−L−Sml−DAPI−B10
7614−Ctx−T−L−Sml−DAPI−B11
7614−Ctx−T−L−Sml−DAPI−B12
7614−Ctx−T−L−Sml−DAPI−B7
7614−Ctx−T−L−Sml−DAPI−B8
7614−Ctx−T−L−Sml−DAPI−B9
7614−Ctx−T−L−Sml−DAPI−C10
7614−Ctx−T−L−Sml−DAPI−C11
7614−Ctx−T−L−Sml−DAPI−C12
7614−Ctx−T−L−Sml−DAPI−C7
7614−Ctx−T−L−Sml−DAPI−C8
7614−Ctx−T−L−Sml−DAPI−C9
7614−Ctx−T−L−Sml−DAPI−D10
7614−Ctx−T−L−Sml−DAPI−D11
7614−Ctx−T−L−Sml−DAPI−D12
7614−Ctx−T−L−Sml−DAPI−D7
7614−Ctx−T−L−Sml−DAPI−D8
7614−Ctx−T−L−Sml−DAPI−D9
7614−Ctx−T−L−Sml−DAPI−E10
7614−Ctx−T−L−Sml−DAPI−E11
7614−Ctx−T−L−Sml−DAPI−E12
7614−Ctx−T−L−Sml−DAPI−E7
7614−Ctx−T−L−Sml−DAPI−E8
7614−Ctx−T−L−Sml−DAPI−E9
7614−Ctx−T−L−Sml−DAPI−F10
7614−Ctx−T−L−Sml−DAPI−F11
7614−Ctx−T−L−Sml−DAPI−F12
7614−Ctx−T−L−Sml−DAPI−F7
7614−Ctx−T−L−Sml−DAPI−F8
7614−Ctx−T−L−Sml−DAPI−F9
7614−Ctx−T−L−Sml−DAPI−G10
7614−Ctx−T−L−Sml−DAPI−G11
7614−Ctx−T−L−Sml−DAPI−G12
7614−Ctx−T−L−Sml−DAPI−G7
7614−Ctx−T−L−Sml−DAPI−G8
7614−Ctx−T−L−Sml−DAPI−G9
7614−Ctx−T−L−Sml−DAPI−H10
7614−Ctx−T−L−Sml−DAPI−H11
7614−Ctx−T−L−Sml−DAPI−H7
7614−Ctx−T−L−Sml−DAPI−H8
7614−Ctx−T−L−Sml−DAPI−H9
7614−Ctx−T−L−Sml−NeuN−A1
7614−Ctx−T−L−Sml−NeuN−A2
7614−Ctx−T−L−Sml−NeuN−A3
7614−Ctx−T−L−Sml−NeuN−A4
7614−Ctx−T−L−Sml−NeuN−A5
7614−Ctx−T−L−Sml−NeuN−A6
7614−Ctx−T−L−Sml−NeuN−B1
7614−Ctx−T−L−Sml−NeuN−B2
7614−Ctx−T−L−Sml−NeuN−B3
7614−Ctx−T−L−Sml−NeuN−B4
7614−Ctx−T−L−Sml−NeuN−B5
7614−Ctx−T−L−Sml−NeuN−B6
7614−Ctx−T−L−Sml−NeuN−C1
7614−Ctx−T−L−Sml−NeuN−C2
7614−Ctx−T−L−Sml−NeuN−C3
7614−Ctx−T−L−Sml−NeuN−C4
7614−Ctx−T−L−Sml−NeuN−C5
7614−Ctx−T−L−Sml−NeuN−C6
7614−Ctx−T−L−Sml−NeuN−D1
7614−Ctx−T−L−Sml−NeuN−D2
7614−Ctx−T−L−Sml−NeuN−D3
7614−Ctx−T−L−Sml−NeuN−D4
7614−Ctx−T−L−Sml−NeuN−D5
7614−Ctx−T−L−Sml−NeuN−D6
7614−Ctx−T−L−Sml−NeuN−E1
7614−Ctx−T−L−Sml−NeuN−E2
7614−Ctx−T−L−Sml−NeuN−E3
7614−Ctx−T−L−Sml−NeuN−E4
7614−Ctx−T−L−Sml−NeuN−E5
7614−Ctx−T−L−Sml−NeuN−E6
7614−Ctx−T−L−Sml−NeuN−F1
7614−Ctx−T−L−Sml−NeuN−F2
7614−Ctx−T−L−Sml−NeuN−F3
7614−Ctx−T−L−Sml−NeuN−F4
7614−Ctx−T−L−Sml−NeuN−F5
7614−Ctx−T−L−Sml−NeuN−F6
7614−Ctx−T−L−Sml−NeuN−G1
7614−Ctx−T−L−Sml−NeuN−G2
7614−Ctx−T−L−Sml−NeuN−G3
7614−Ctx−T−L−Sml−NeuN−G4
7614−Ctx−T−L−Sml−NeuN−G5
7614−Ctx−T−L−Sml−NeuN−G6
7614−Ctx−T−L−Sml−NeuN−H1
7614−Ctx−T−L−Sml−NeuN−H2
7614−Ctx−T−L−Sml−NeuN−H3
7614−Ctx−T−L−Sml−NeuN−H4
7614−Ctx−T−L−Sml−NeuN−H5
7614−Ctx−T−L−Sml−NeuN−H6
JGG-Cells

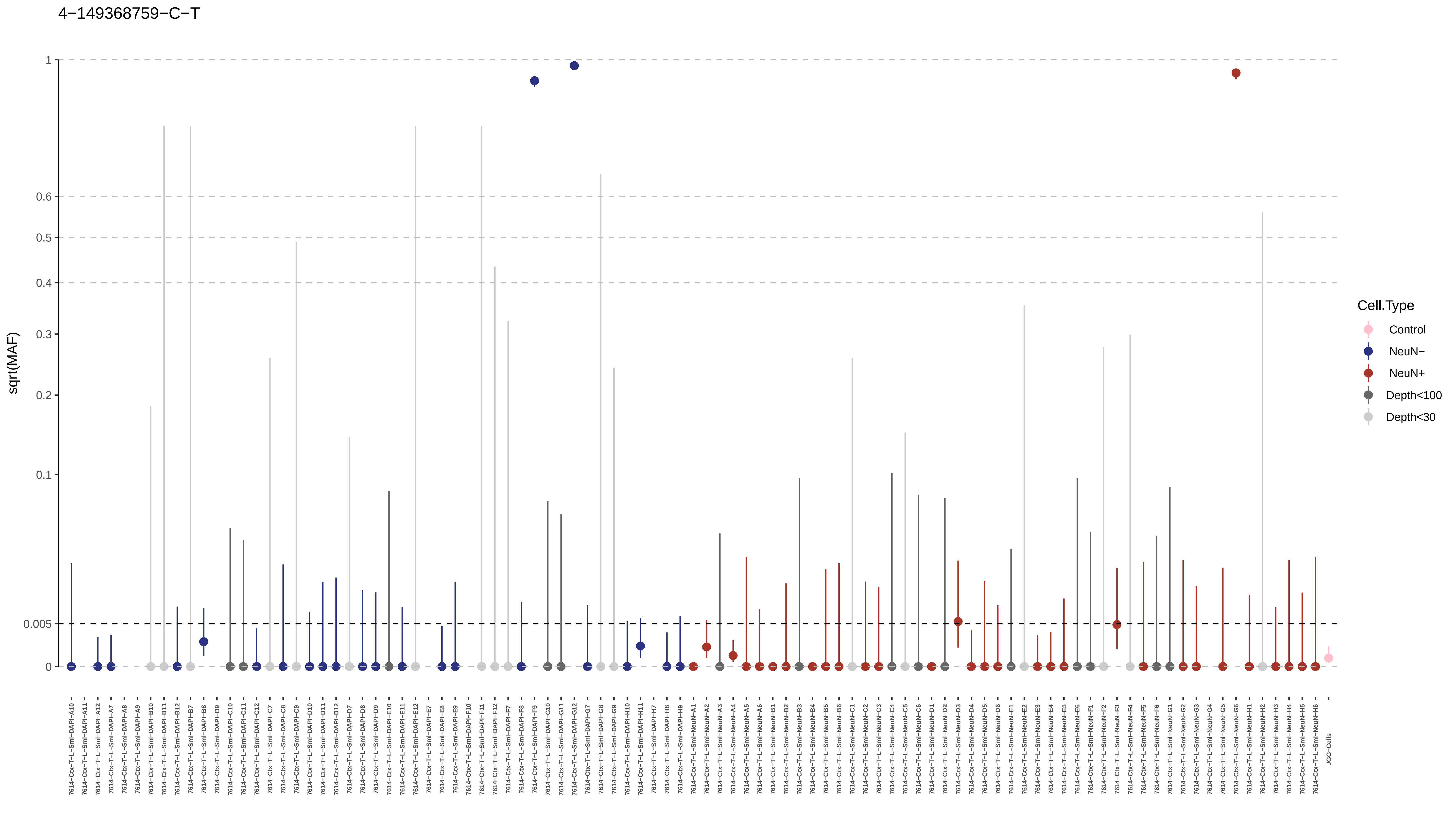

4−149368759−C−T
sqrt(MAF)
1
0.6
0.5
0.4
0.3
0.2
0.1
0.005
0
Cell.Type
Control
NeuN−
NeuN+
Depth<100
Depth<30

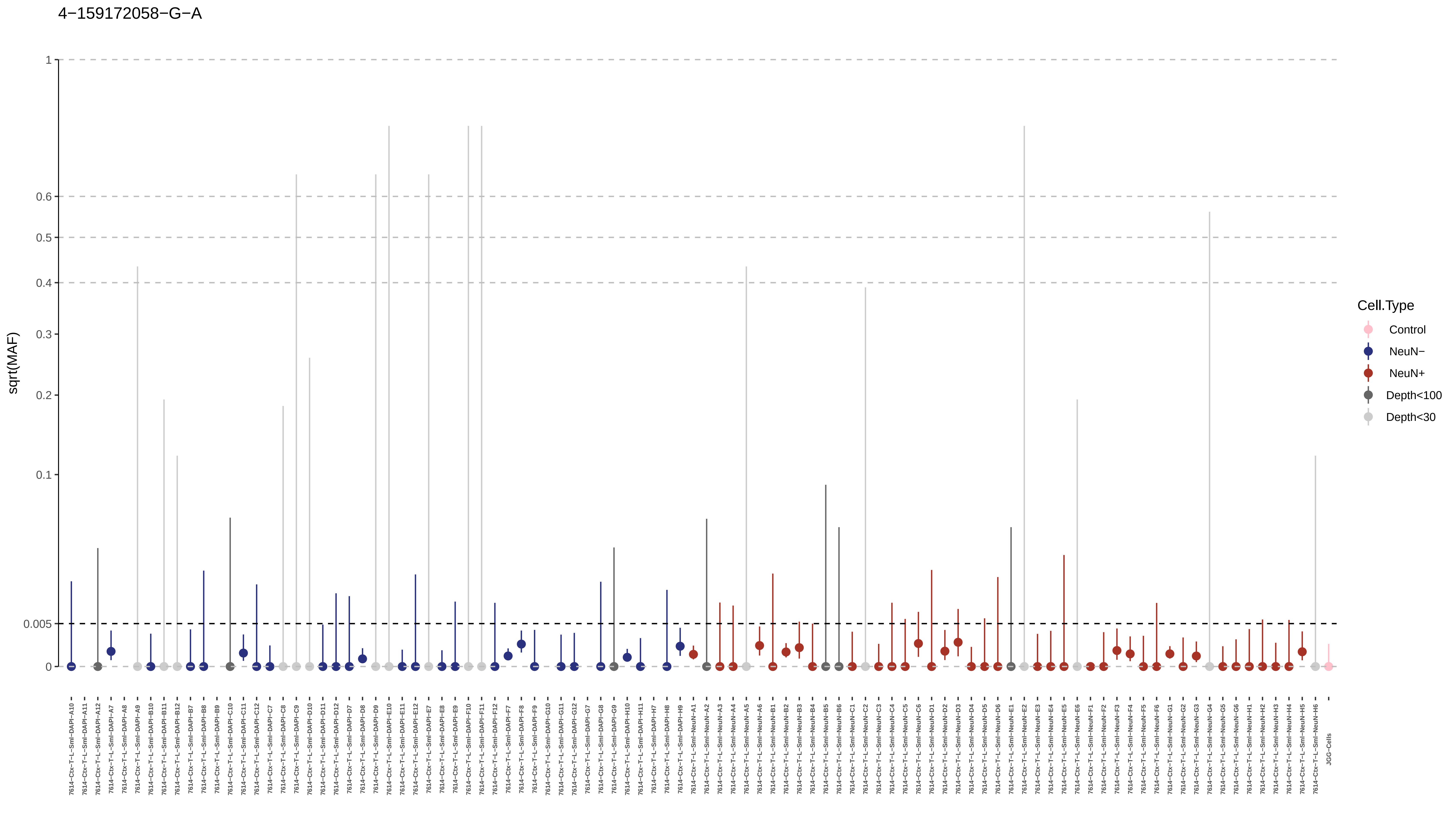
4−159172058−G−A
sqrt(MAF)
1
0.6
0.5
0.4
0.3
0.2
0.1
0.005
0
Cell.Type
Control
NeuN−
NeuN+
Depth<100
Depth<30
7614−Ctx−T−L−Sml−DAPI−A10
7614−Ctx−T−L−Sml−DAPI−A11
7614−Ctx−T−L−Sml−DAPI−A12
7614−Ctx−T−L−Sml−DAPI−A7
7614−Ctx−T−L−Sml−DAPI−A8
7614−Ctx−T−L−Sml−DAPI−A9
7614−Ctx−T−L−Sml−DAPI−B10
7614−Ctx−T−L−Sml−DAPI−B11
7614−Ctx−T−L−Sml−DAPI−B12
7614−Ctx−T−L−Sml−DAPI−B7
7614−Ctx−T−L−Sml−DAPI−B8
7614−Ctx−T−L−Sml−DAPI−B9
7614−Ctx−T−L−Sml−DAPI−C10
7614−Ctx−T−L−Sml−DAPI−C11
7614−Ctx−T−L−Sml−DAPI−C12
7614−Ctx−T−L−Sml−DAPI−C7
7614−Ctx−T−L−Sml−DAPI−C8
7614−Ctx−T−L−Sml−DAPI−C9
7614−Ctx−T−L−Sml−DAPI−D10
7614−Ctx−T−L−Sml−DAPI−D11
7614−Ctx−T−L−Sml−DAPI−D12
7614−Ctx−T−L−Sml−DAPI−D7
7614−Ctx−T−L−Sml−DAPI−D8
7614−Ctx−T−L−Sml−DAPI−D9
7614−Ctx−T−L−Sml−DAPI−E10
7614−Ctx−T−L−Sml−DAPI−E11
7614−Ctx−T−L−Sml−DAPI−E12
7614−Ctx−T−L−Sml−DAPI−E7
7614−Ctx−T−L−Sml−DAPI−E8
7614−Ctx−T−L−Sml−DAPI−E9
7614−Ctx−T−L−Sml−DAPI−F10
7614−Ctx−T−L−Sml−DAPI−F11
7614−Ctx−T−L−Sml−DAPI−F12
7614−Ctx−T−L−Sml−DAPI−F7
7614−Ctx−T−L−Sml−DAPI−F8
7614−Ctx−T−L−Sml−DAPI−F9
7614−Ctx−T−L−Sml−DAPI−G10
7614−Ctx−T−L−Sml−DAPI−G11
7614−Ctx−T−L−Sml−DAPI−G12
7614−Ctx−T−L−Sml−DAPI−G7
7614−Ctx−T−L−Sml−DAPI−G8
7614−Ctx−T−L−Sml−DAPI−G9
7614−Ctx−T−L−Sml−DAPI−H10
7614−Ctx−T−L−Sml−DAPI−H11
7614−Ctx−T−L−Sml−DAPI−H7
7614−Ctx−T−L−Sml−DAPI−H8
7614−Ctx−T−L−Sml−DAPI−H9
7614−Ctx−T−L−Sml−NeuN−A1
7614−Ctx−T−L−Sml−NeuN−A2
7614−Ctx−T−L−Sml−NeuN−A3
7614−Ctx−T−L−Sml−NeuN−A4
7614−Ctx−T−L−Sml−NeuN−A5
7614−Ctx−T−L−Sml−NeuN−A6
7614−Ctx−T−L−Sml−NeuN−B1
7614−Ctx−T−L−Sml−NeuN−B2
7614−Ctx−T−L−Sml−NeuN−B3
7614−Ctx−T−L−Sml−NeuN−B4
7614−Ctx−T−L−Sml−NeuN−B5
7614−Ctx−T−L−Sml−NeuN−B6
7614−Ctx−T−L−Sml−NeuN−C1
7614−Ctx−T−L−Sml−NeuN−C2
7614−Ctx−T−L−Sml−NeuN−C3
7614−Ctx−T−L−Sml−NeuN−C4
7614−Ctx−T−L−Sml−NeuN−C5
7614−Ctx−T−L−Sml−NeuN−C6
7614−Ctx−T−L−Sml−NeuN−D1
7614−Ctx−T−L−Sml−NeuN−D2
7614−Ctx−T−L−Sml−NeuN−D3
7614−Ctx−T−L−Sml−NeuN−D4
7614−Ctx−T−L−Sml−NeuN−D5
7614−Ctx−T−L−Sml−NeuN−D6
7614−Ctx−T−L−Sml−NeuN−E1
7614−Ctx−T−L−Sml−NeuN−E2
7614−Ctx−T−L−Sml−NeuN−E3
7614−Ctx−T−L−Sml−NeuN−E4
7614−Ctx−T−L−Sml−NeuN−E5
7614−Ctx−T−L−Sml−NeuN−E6
7614−Ctx−T−L−Sml−NeuN−F1
7614−Ctx−T−L−Sml−NeuN−F2
7614−Ctx−T−L−Sml−NeuN−F3
7614−Ctx−T−L−Sml−NeuN−F4
7614−Ctx−T−L−Sml−NeuN−F5
7614−Ctx−T−L−Sml−NeuN−F6
7614−Ctx−T−L−Sml−NeuN−G1
7614−Ctx−T−L−Sml−NeuN−G2
7614−Ctx−T−L−Sml−NeuN−G3
7614−Ctx−T−L−Sml−NeuN−G4
7614−Ctx−T−L−Sml−NeuN−G5
7614−Ctx−T−L−Sml−NeuN−G6
7614−Ctx−T−L−Sml−NeuN−H1
7614−Ctx−T−L−Sml−NeuN−H2
7614−Ctx−T−L−Sml−NeuN−H3
7614−Ctx−T−L−Sml−NeuN−H4
7614−Ctx−T−L−Sml−NeuN−H5
7614−Ctx−T−L−Sml−NeuN−H6
JGG−Cells

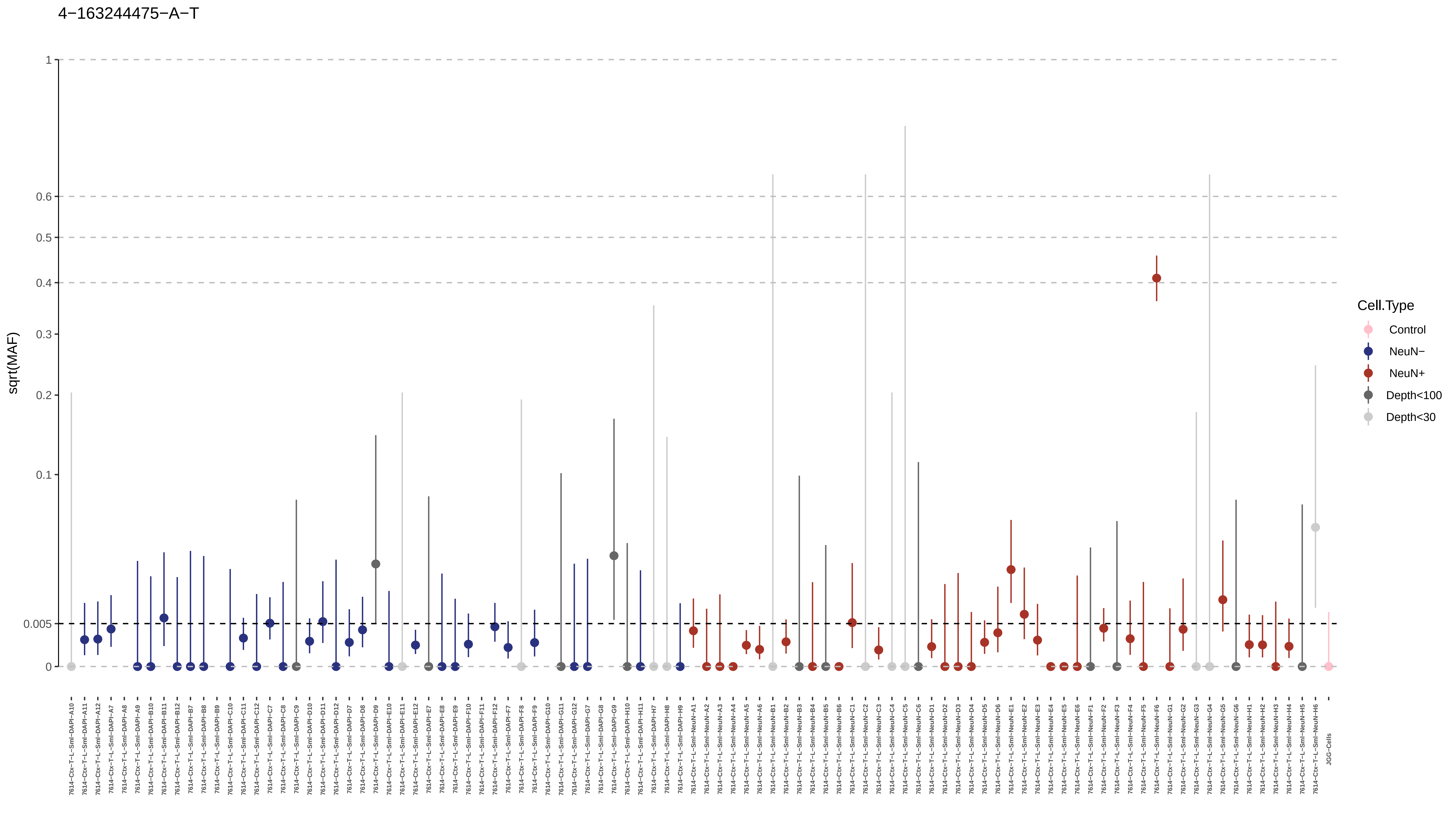
4−163244475−A−T
sqrt(MAF)
1
0.6
0.5
0.4
0.3
0.2
0.1
0.005
0
Cell.Type
Control
NeuN−
NeuN+
Depth<100
Depth<30
7614−Ctx−T−L−Sml−DAPI−A10
7614−Ctx−T−L−Sml−DAPI−A11
7614−Ctx−T−L−Sml−DAPI−A12
7614−Ctx−T−L−Sml−DAPI−A7
7614−Ctx−T−L−Sml−DAPI−A8
7614−Ctx−T−L−Sml−DAPI−A9
7614−Ctx−T−L−Sml−DAPI−B10
7614−Ctx−T−L−Sml−DAPI−B11
7614−Ctx−T−L−Sml−DAPI−B12
7614−Ctx−T−L−Sml−DAPI−B7
7614−Ctx−T−L−Sml−DAPI−B8
7614−Ctx−T−L−Sml−DAPI−B9
7614−Ctx−T−L−Sml−DAPI−C10
7614−Ctx−T−L−Sml−DAPI−C11
7614−Ctx−T−L−Sml−DAPI−C12
7614−Ctx−T−L−Sml−DAPI−C7
7614−Ctx−T−L−Sml−DAPI−C8
7614−Ctx−T−L−Sml−DAPI−C9
7614−Ctx−T−L−Sml−DAPI−D10
7614−Ctx−T−L−Sml−DAPI−D11
7614−Ctx−T−L−Sml−DAPI−D12
7614−Ctx−T−L−Sml−DAPI−D7
7614−Ctx−T−L−Sml−DAPI−D8
7614−Ctx−T−L−Sml−DAPI−D9
7614−Ctx−T−L−Sml−DAPI−E10
7614−Ctx−T−L−Sml−DAPI−E11
7614−Ctx−T−L−Sml−DAPI−E12
7614−Ctx−T−L−Sml−DAPI−E7
7614−Ctx−T−L−Sml−DAPI−E8
7614−Ctx−T−L−Sml−DAPI−E9
7614−Ctx−T−L−Sml−DAPI−F10
7614−Ctx−T−L−Sml−DAPI−F11
7614−Ctx−T−L−Sml−DAPI−F12
7614−Ctx−T−L−Sml−DAPI−F7
7614−Ctx−T−L−Sml−DAPI−F8
7614−Ctx−T−L−Sml−DAPI−F9
7614−Ctx−T−L−Sml−DAPI−G10
7614−Ctx−T−L−Sml−DAPI−G11
7614−Ctx−T−L−Sml−DAPI−G12
7614−Ctx−T−L−Sml−DAPI−G7
7614−Ctx−T−L−Sml−DAPI−G8
7614−Ctx−T−L−Sml−DAPI−G9
7614−Ctx−T−L−Sml−DAPI−H10
7614−Ctx−T−L−Sml−DAPI−H11
7614−Ctx−T−L−Sml−DAPI−H7
7614−Ctx−T−L−Sml−DAPI−H8
7614−Ctx−T−L−Sml−DAPI−H9
7614−Ctx−T−L−Sml−NeuN−A1
7614−Ctx−T−L−Sml−NeuN−A2
7614−Ctx−T−L−Sml−NeuN−A3
7614−Ctx−T−L−Sml−NeuN−A4
7614−Ctx−T−L−Sml−NeuN−A5
7614−Ctx−T−L−Sml−NeuN−A6
7614−Ctx−T−L−Sml−NeuN−B1
7614−Ctx−T−L−Sml−NeuN−B2
7614−Ctx−T−L−Sml−NeuN−B3
7614−Ctx−T−L−Sml−NeuN−B4
7614−Ctx−T−L−Sml−NeuN−B5
7614−Ctx−T−L−Sml−NeuN−B6
7614−Ctx−T−L−Sml−NeuN−C1
7614−Ctx−T−L−Sml−NeuN−C2
7614−Ctx−T−L−Sml−NeuN−C3
7614−Ctx−T−L−Sml−NeuN−C4
7614−Ctx−T−L−Sml−NeuN−C5
7614−Ctx−T−L−Sml−NeuN−C6
7614−Ctx−T−L−Sml−NeuN−D1
7614−Ctx−T−L−Sml−NeuN−D2
7614−Ctx−T−L−Sml−NeuN−D3
7614−Ctx−T−L−Sml−NeuN−D4
7614−Ctx−T−L−Sml−NeuN−D5
7614−Ctx−T−L−Sml−NeuN−D6
7614−Ctx−T−L−Sml−NeuN−E1
7614−Ctx−T−L−Sml−NeuN−E2
7614−Ctx−T−L−Sml−NeuN−E3
7614−Ctx−T−L−Sml−NeuN−E4
7614−Ctx−T−L−Sml−NeuN−E5
7614−Ctx−T−L−Sml−NeuN−E6
7614−Ctx−T−L−Sml−NeuN−F1
7614−Ctx−T−L−Sml−NeuN−F2
7614−Ctx−T−L−Sml−NeuN−F3
7614−Ctx−T−L−Sml−NeuN−F4
7614−Ctx−T−L−Sml−NeuN−F5
7614−Ctx−T−L−Sml−NeuN−F6
7614−Ctx−T−L−Sml−NeuN−G1
7614−Ctx−T−L−Sml−NeuN−G2
7614−Ctx−T−L−Sml−NeuN−G3
7614−Ctx−T−L−Sml−NeuN−G4
7614−Ctx−T−L−Sml−NeuN−G5
7614−Ctx−T−L−Sml−NeuN−G6
7614−Ctx−T−L−Sml−NeuN−H1
7614−Ctx−T−L−Sml−NeuN−H2
7614−Ctx−T−L−Sml−NeuN−H3
7614−Ctx−T−L−Sml−NeuN−H4
7614−Ctx−T−L−Sml−NeuN−H5
7614−Ctx−T−L−Sml−NeuN−H6
JGG−Cells

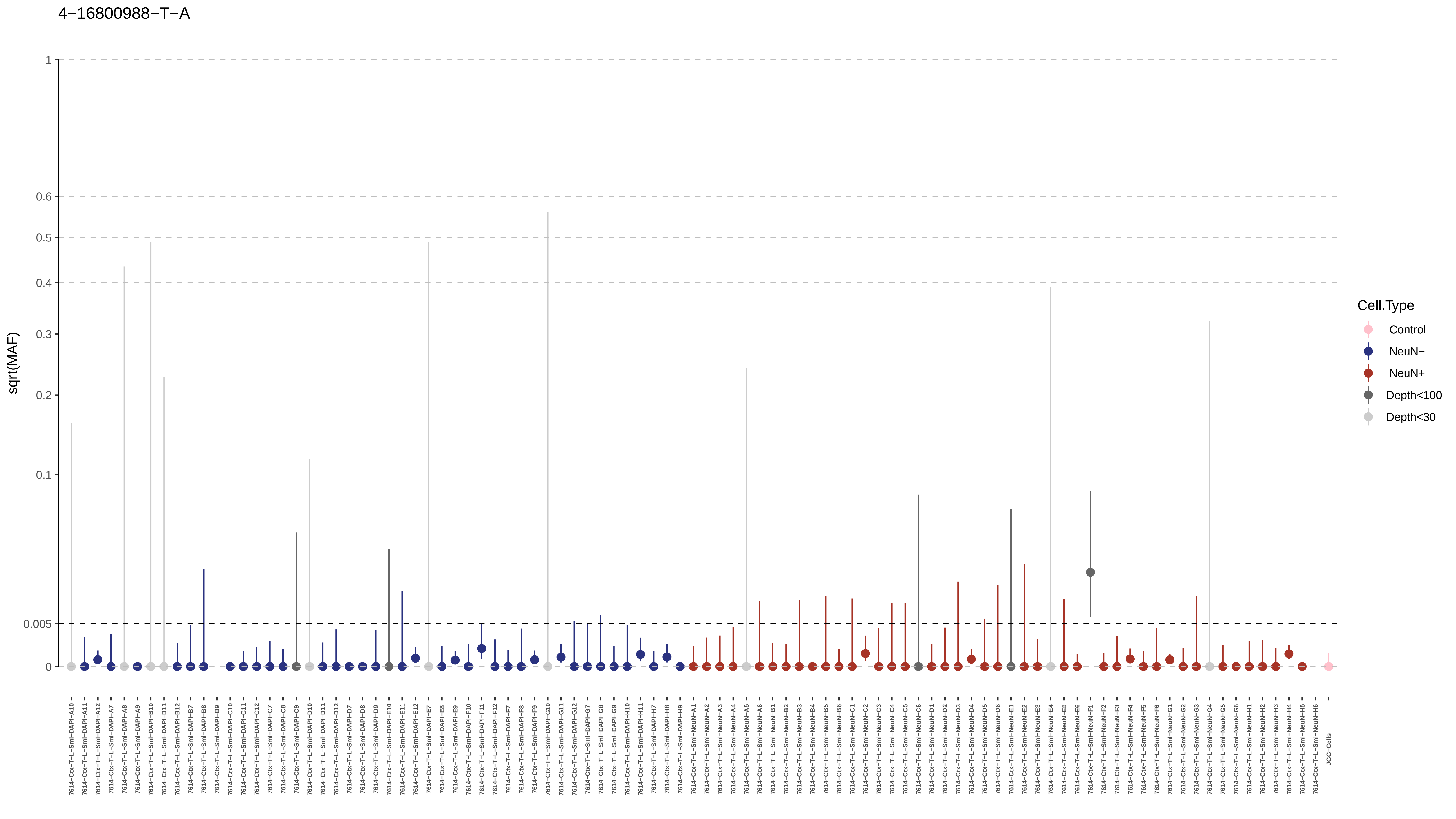
4−16800988−T−A
sqrt(MAF)
1
0.6
0.5
0.4
0.3
0.2
0.1
0.005
0
Cell.Type
Control
NeuN−
NeuN+
Depth<100
Depth<30
7614−Ctx−T−L−Sml−DAPI−A10
7614−Ctx−T−L−Sml−DAPI−A11
7614−Ctx−T−L−Sml−DAPI−A12
7614−Ctx−T−L−Sml−DAPI−A7
7614−Ctx−T−L−Sml−DAPI−A8
7614−Ctx−T−L−Sml−DAPI−A9
7614−Ctx−T−L−Sml−DAPI−B10
7614−Ctx−T−L−Sml−DAPI−B11
7614−Ctx−T−L−Sml−DAPI−B12
7614−Ctx−T−L−Sml−DAPI−B7
7614−Ctx−T−L−Sml−DAPI−B8
7614−Ctx−T−L−Sml−DAPI−B9
7614−Ctx−T−L−Sml−DAPI−C10
7614−Ctx−T−L−Sml−DAPI−C11
7614−Ctx−T−L−Sml−DAPI−C12
7614−Ctx−T−L−Sml−DAPI−C7
7614−Ctx−T−L−Sml−DAPI−C8
7614−Ctx−T−L−Sml−DAPI−C9
7614−Ctx−T−L−Sml−DAPI−D10
7614−Ctx−T−L−Sml−DAPI−D11
7614−Ctx−T−L−Sml−DAPI−D12
7614−Ctx−T−L−Sml−DAPI−D7
7614−Ctx−T−L−Sml−DAPI−D8
7614−Ctx−T−L−Sml−DAPI−D9
7614−Ctx−T−L−Sml−DAPI−E10
7614−Ctx−T−L−Sml−DAPI−E11
7614−Ctx−T−L−Sml−DAPI−E12
7614−Ctx−T−L−Sml−DAPI−E7
7614−Ctx−T−L−Sml−DAPI−E8
7614−Ctx−T−L−Sml−DAPI−E9
7614−Ctx−T−L−Sml−DAPI−F10
7614−Ctx−T−L−Sml−DAPI−F11
7614−Ctx−T−L−Sml−DAPI−F12
7614−Ctx−T−L−Sml−DAPI−F7
7614−Ctx−T−L−Sml−DAPI−F8
7614−Ctx−T−L−Sml−DAPI−F9
7614−Ctx−T−L−Sml−DAPI−G10
7614−Ctx−T−L−Sml−DAPI−G11
7614−Ctx−T−L−Sml−DAPI−G12
7614−Ctx−T−L−Sml−DAPI−G7
7614−Ctx−T−L−Sml−DAPI−G8
7614−Ctx−T−L−Sml−DAPI−G9
7614−Ctx−T−L−Sml−DAPI−H10
7614−Ctx−T−L−Sml−DAPI−H11
7614−Ctx−T−L−Sml−DAPI−H7
7614−Ctx−T−L−Sml−DAPI−H8
7614−Ctx−T−L−Sml−DAPI−H9
7614−Ctx−T−L−Sml−NeuN−A1
7614−Ctx−T−L−Sml−NeuN−A2
7614−Ctx−T−L−Sml−NeuN−A3
7614−Ctx−T−L−Sml−NeuN−A4
7614−Ctx−T−L−Sml−NeuN−A5
7614−Ctx−T−L−Sml−NeuN−A6
7614−Ctx−T−L−Sml−NeuN−B1
7614−Ctx−T−L−Sml−NeuN−B2
7614−Ctx−T−L−Sml−NeuN−B3
7614−Ctx−T−L−Sml−NeuN−B4
7614−Ctx−T−L−Sml−NeuN−B5
7614−Ctx−T−L−Sml−NeuN−B6
7614−Ctx−T−L−Sml−NeuN−C1
7614−Ctx−T−L−Sml−NeuN−C2
7614−Ctx−T−L−Sml−NeuN−C3
7614−Ctx−T−L−Sml−NeuN−C4
7614−Ctx−T−L−Sml−NeuN−C5
7614−Ctx−T−L−Sml−NeuN−C6
7614−Ctx−T−L−Sml−NeuN−D1
7614−Ctx−T−L−Sml−NeuN−D2
7614−Ctx−T−L−Sml−NeuN−D3
7614−Ctx−T−L−Sml−NeuN−D4
7614−Ctx−T−L−Sml−NeuN−D5
7614−Ctx−T−L−Sml−NeuN−D6
7614−Ctx−T−L−Sml−NeuN−E1
7614−Ctx−T−L−Sml−NeuN−E2
7614−Ctx−T−L−Sml−NeuN−E3
7614−Ctx−T−L−Sml−NeuN−E4
7614−Ctx−T−L−Sml−NeuN−E5
7614−Ctx−T−L−Sml−NeuN−E6
7614−Ctx−T−L−Sml−NeuN−F1
7614−Ctx−T−L−Sml−NeuN−F2
7614−Ctx−T−L−Sml−NeuN−F3
7614−Ctx−T−L−Sml−NeuN−F4
7614−Ctx−T−L−Sml−NeuN−F5
7614−Ctx−T−L−Sml−NeuN−F6
7614−Ctx−T−L−Sml−NeuN−G1
7614−Ctx−T−L−Sml−NeuN−G2
7614−Ctx−T−L−Sml−NeuN−G3
7614−Ctx−T−L−Sml−NeuN−G4
7614−Ctx−T−L−Sml−NeuN−G5
7614−Ctx−T−L−Sml−NeuN−G6
7614−Ctx−T−L−Sml−NeuN−H1
7614−Ctx−T−L−Sml−NeuN−H2
7614−Ctx−T−L−Sml−NeuN−H3
7614−Ctx−T−L−Sml−NeuN−H4
7614−Ctx−T−L−Sml−NeuN−H5
7614−Ctx−T−L−Sml−NeuN−H6
JGG−Cells

4−176659497−G−A

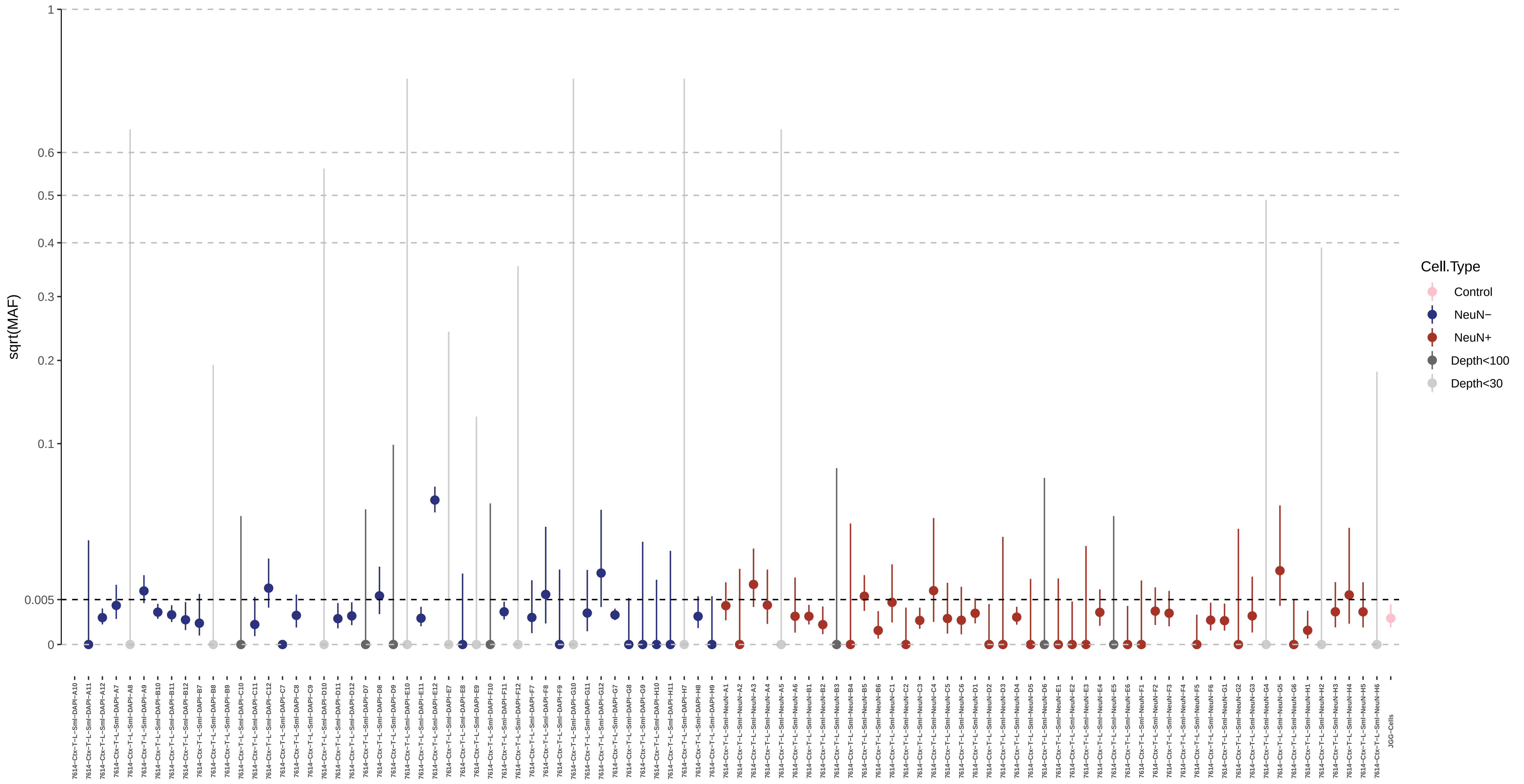

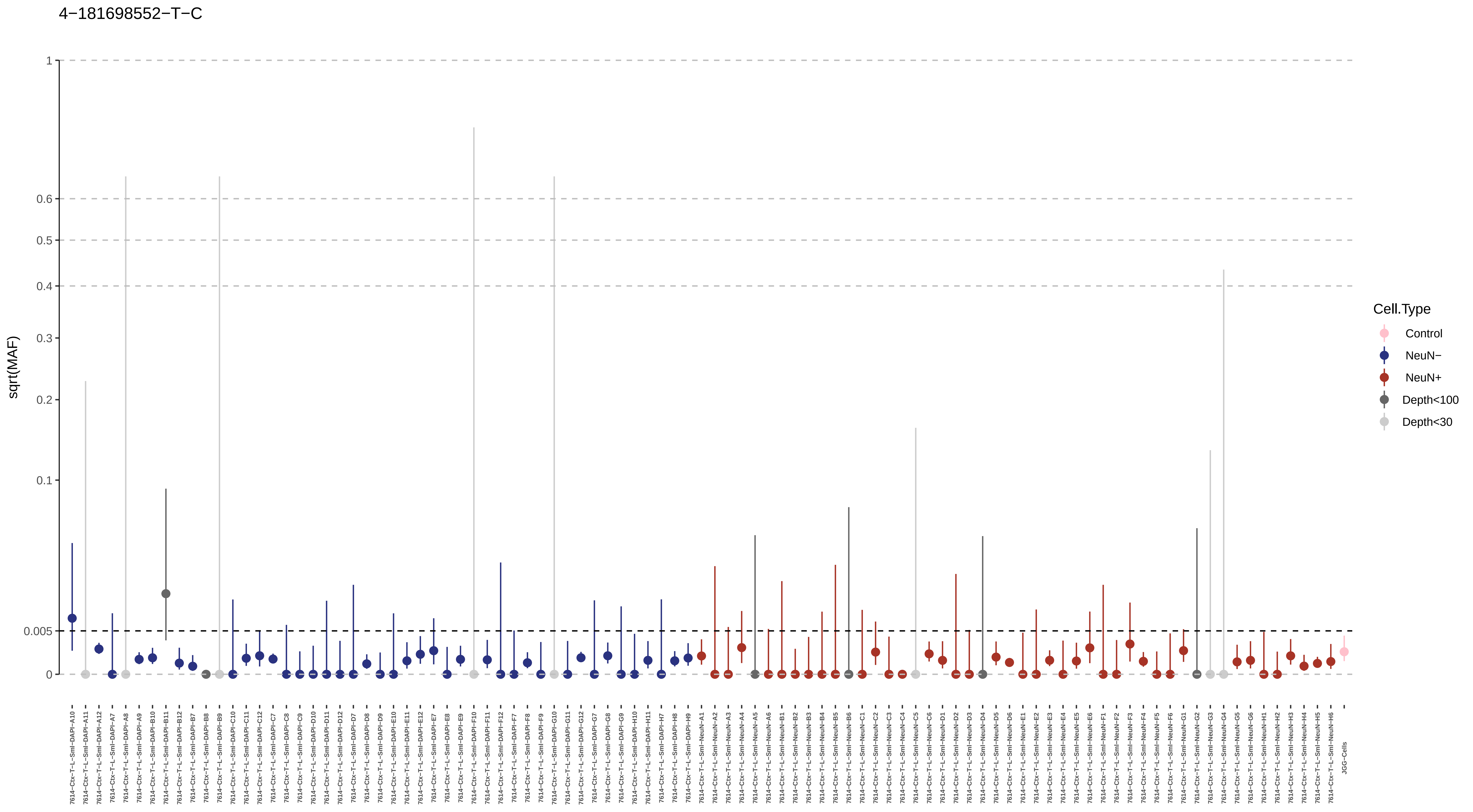
4−181698552−T−C
sqrt(MAF)
1
0.6
0.5
0.4
0.3
0.2
0.1
0.005
0
Cell.Type
Control
NeuN−
NeuN+
Depth<100
Depth<30

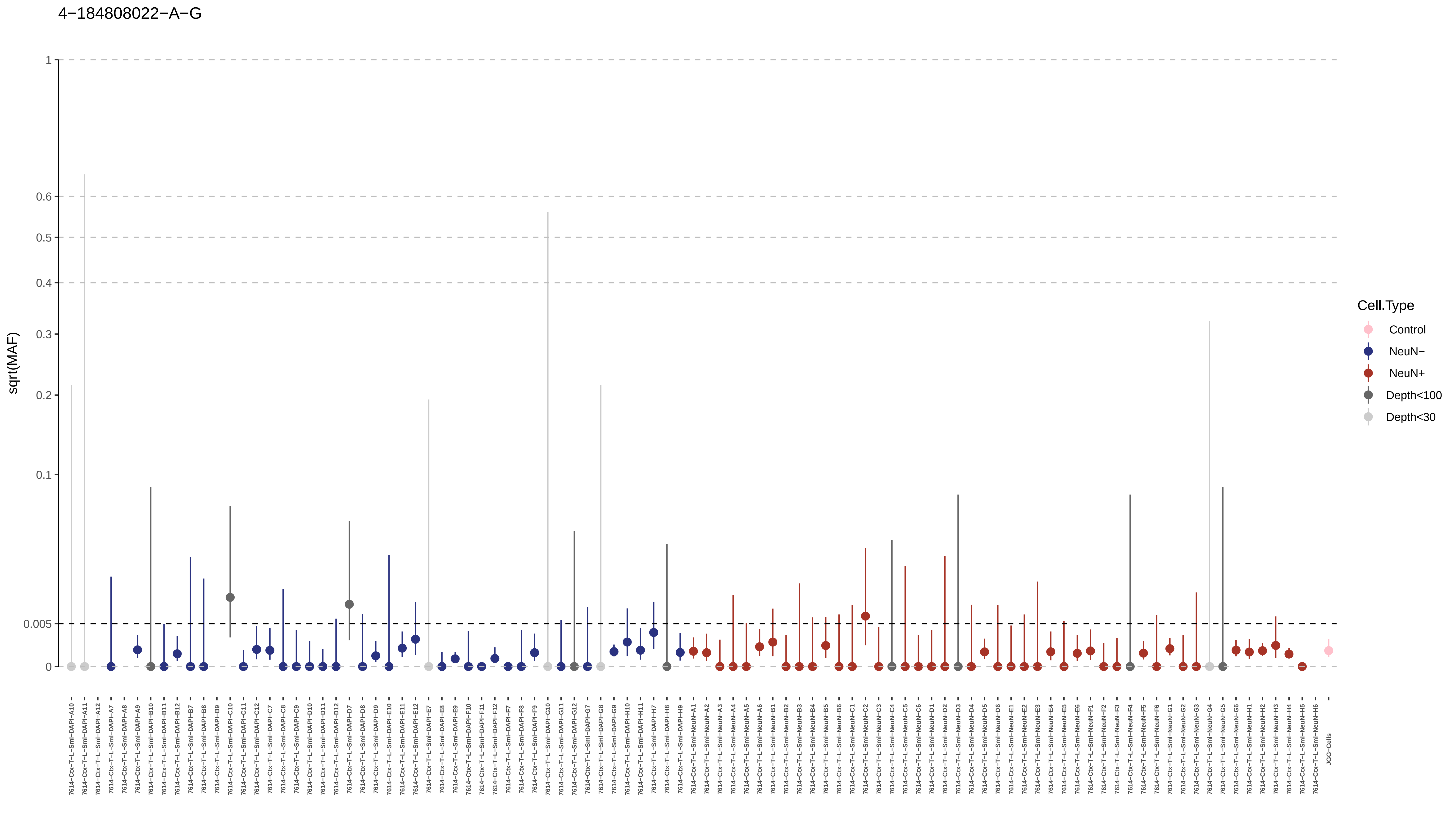
4−184808022−A−G
sqrt(MAF)
1
0.6
0.5
0.4
0.3
0.2
0.1
0.005
0
Cell.Type
Control
NeuN−
NeuN+
Depth<100
Depth<30
7614−Ctx−T−L−Sml−DAPI−A10
7614−Ctx−T−L−Sml−DAPI−A11
7614−Ctx−T−L−Sml−DAPI−A12
7614−Ctx−T−L−Sml−DAPI−A7
7614−Ctx−T−L−Sml−DAPI−A8
7614−Ctx−T−L−Sml−DAPI−A9
7614−Ctx−T−L−Sml−DAPI−B10
7614−Ctx−T−L−Sml−DAPI−B11
7614−Ctx−T−L−Sml−DAPI−B12
7614−Ctx−T−L−Sml−DAPI−B7
7614−Ctx−T−L−Sml−DAPI−B8
7614−Ctx−T−L−Sml−DAPI−B9
7614−Ctx−T−L−Sml−DAPI−C10
7614−Ctx−T−L−Sml−DAPI−C11
7614−Ctx−T−L−Sml−DAPI−C12
7614−Ctx−T−L−Sml−DAPI−C7
7614−Ctx−T−L−Sml−DAPI−C8
7614−Ctx−T−L−Sml−DAPI−C9
7614−Ctx−T−L−Sml−DAPI−D10
7614−Ctx−T−L−Sml−DAPI−D11
7614−Ctx−T−L−Sml−DAPI−D12
7614−Ctx−T−L−Sml−DAPI−D7
7614−Ctx−T−L−Sml−DAPI−D8
7614−Ctx−T−L−Sml−DAPI−D9
7614−Ctx−T−L−Sml−DAPI−E10
7614−Ctx−T−L−Sml−DAPI−E11
7614−Ctx−T−L−Sml−DAPI−E12
7614−Ctx−T−L−Sml−DAPI−E7
7614−Ctx−T−L−Sml−DAPI−E8
7614−Ctx−T−L−Sml−DAPI−E9
7614−Ctx−T−L−Sml−DAPI−F10
7614−Ctx−T−L−Sml−DAPI−F11
7614−Ctx−T−L−Sml−DAPI−F12
7614−Ctx−T−L−Sml−DAPI−F7
7614−Ctx−T−L−Sml−DAPI−F8
7614−Ctx−T−L−Sml−DAPI−F9
7614−Ctx−T−L−Sml−DAPI−G10
7614−Ctx−T−L−Sml−DAPI−G11
7614−Ctx−T−L−Sml−DAPI−G12
7614−Ctx−T−L−Sml−DAPI−G7
7614−Ctx−T−L−Sml−DAPI−G8
7614−Ctx−T−L−Sml−DAPI−G9
7614−Ctx−T−L−Sml−DAPI−H10
7614−Ctx−T−L−Sml−DAPI−H11
7614−Ctx−T−L−Sml−DAPI−H7
7614−Ctx−T−L−Sml−DAPI−H8
7614−Ctx−T−L−Sml−DAPI−H9
7614−Ctx−T−L−Sml−NeuN−A1
7614−Ctx−T−L−Sml−NeuN−A2
7614−Ctx−T−L−Sml−NeuN−A3
7614−Ctx−T−L−Sml−NeuN−A4
7614−Ctx−T−L−Sml−NeuN−A5
7614−Ctx−T−L−Sml−NeuN−A6
7614−Ctx−T−L−Sml−NeuN−B1
7614−Ctx−T−L−Sml−NeuN−B2
7614−Ctx−T−L−Sml−NeuN−B3
7614−Ctx−T−L−Sml−NeuN−B4
7614−Ctx−T−L−Sml−NeuN−B5
7614−Ctx−T−L−Sml−NeuN−B6
7614−Ctx−T−L−Sml−NeuN−C1
7614−Ctx−T−L−Sml−NeuN−C2
7614−Ctx−T−L−Sml−NeuN−C3
7614−Ctx−T−L−Sml−NeuN−C4
7614−Ctx−T−L−Sml−NeuN−C5
7614−Ctx−T−L−Sml−NeuN−C6
7614−Ctx−T−L−Sml−NeuN−D1
7614−Ctx−T−L−Sml−NeuN−D2
7614−Ctx−T−L−Sml−NeuN−D3
7614−Ctx−T−L−Sml−NeuN−D4
7614−Ctx−T−L−Sml−NeuN−D5
7614−Ctx−T−L−Sml−NeuN−D6
7614−Ctx−T−L−Sml−NeuN−E1
7614−Ctx−T−L−Sml−NeuN−E2
7614−Ctx−T−L−Sml−NeuN−E3
7614−Ctx−T−L−Sml−NeuN−E4
7614−Ctx−T−L−Sml−NeuN−E5
7614−Ctx−T−L−Sml−NeuN−E6
7614−Ctx−T−L−Sml−NeuN−F1
7614−Ctx−T−L−Sml−NeuN−F2
7614−Ctx−T−L−Sml−NeuN−F3
7614−Ctx−T−L−Sml−NeuN−F4
7614−Ctx−T−L−Sml−NeuN−F5
7614−Ctx−T−L−Sml−NeuN−F6
7614−Ctx−T−L−Sml−NeuN−G1
7614−Ctx−T−L−Sml−NeuN−G2
7614−Ctx−T−L−Sml−NeuN−G3
7614−Ctx−T−L−Sml−NeuN−G4
7614−Ctx−T−L−Sml−NeuN−G5
7614−Ctx−T−L−Sml−NeuN−G6
7614−Ctx−T−L−Sml−NeuN−H1
7614−Ctx−T−L−Sml−NeuN−H2
7614−Ctx−T−L−Sml−NeuN−H3
7614−Ctx−T−L−Sml−NeuN−H4
7614−Ctx−T−L−Sml−NeuN−H5
7614−Ctx−T−L−Sml−NeuN−H6
JGG-Cells

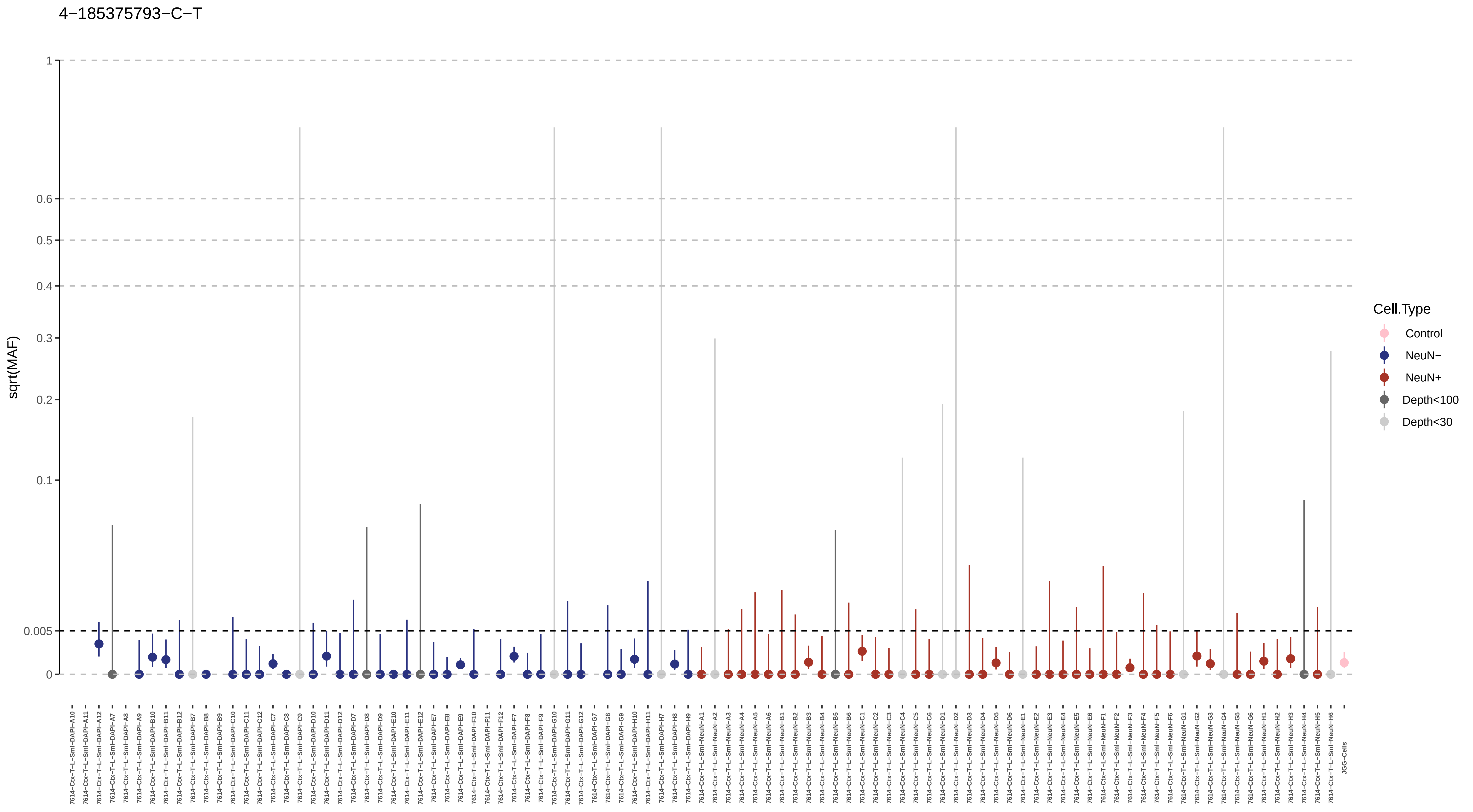
4−185375793−C−T
sqrt(MAF)
1
0.6
0.5
0.4
0.3
0.2
0.1
0.005
0
Cell.Type
Control
NeuN−
NeuN+
Depth<100
Depth<30

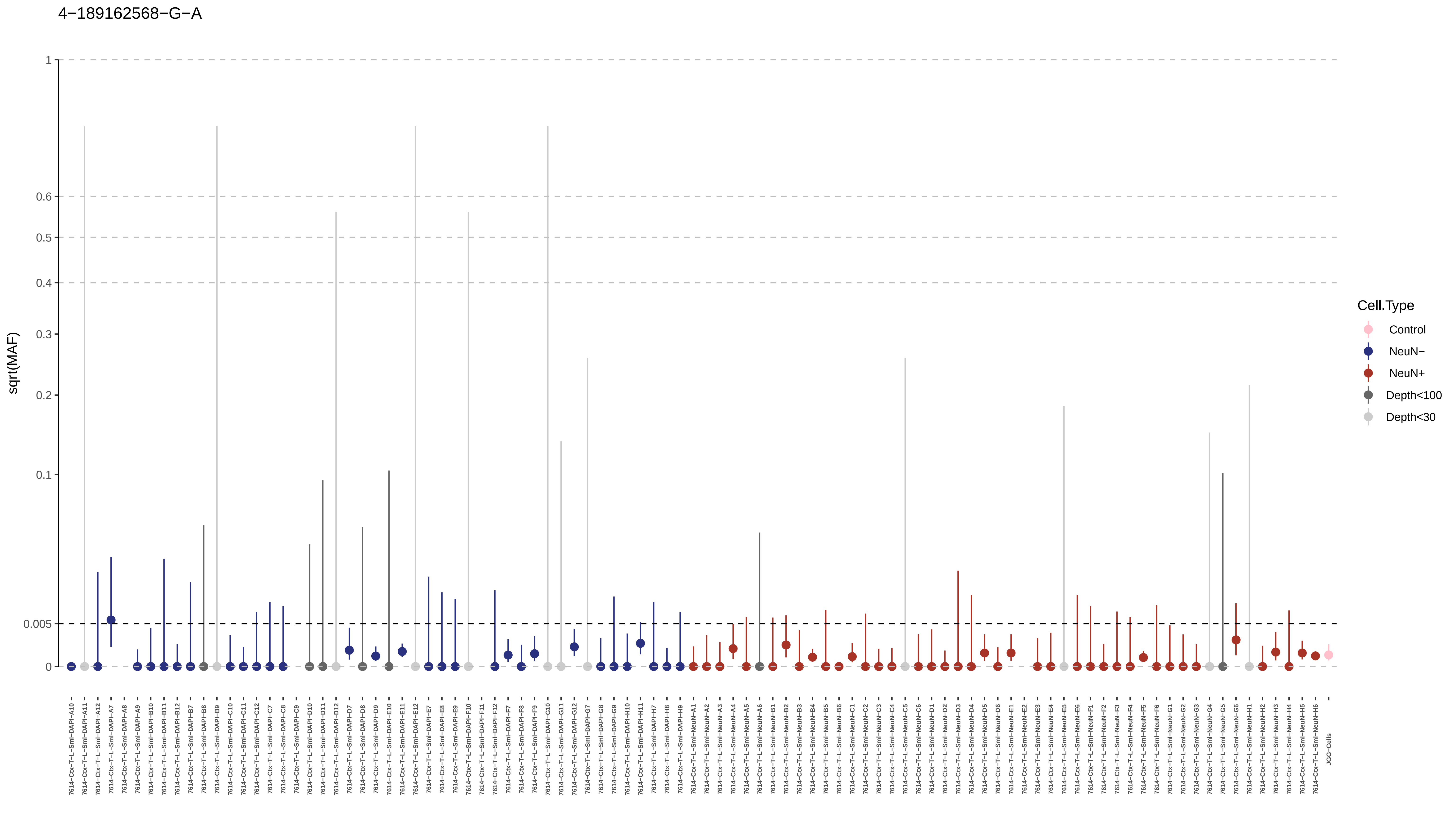
4−189162568−G−A
sqrt(MAF)
1
0.6
0.5
0.4
0.3
0.2
0.1
0.005
0
Cell.Type
Control
NeuN−
NeuN+
Depth<100
Depth<30
JGG-Cells

4−28629401−C−T

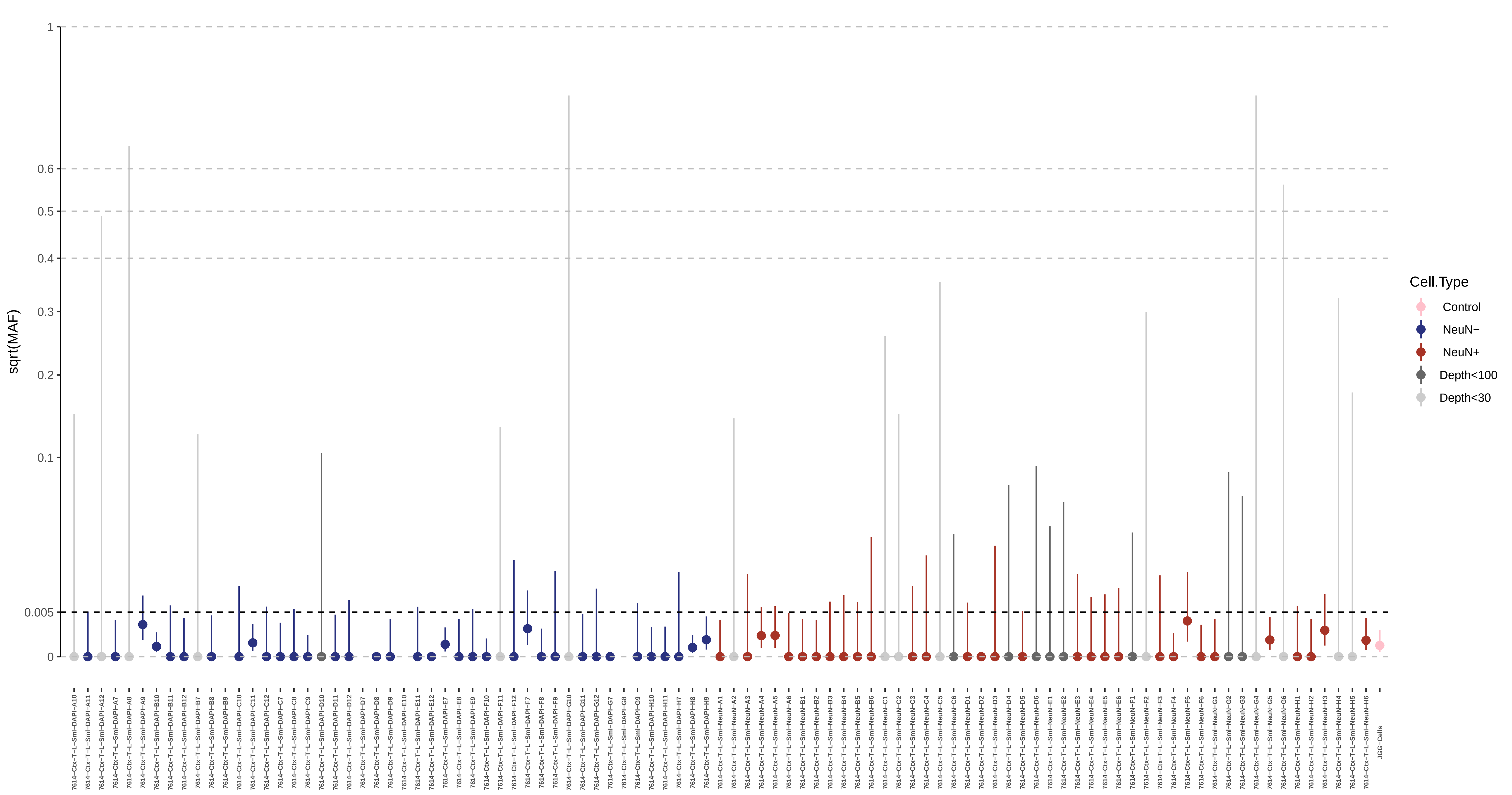

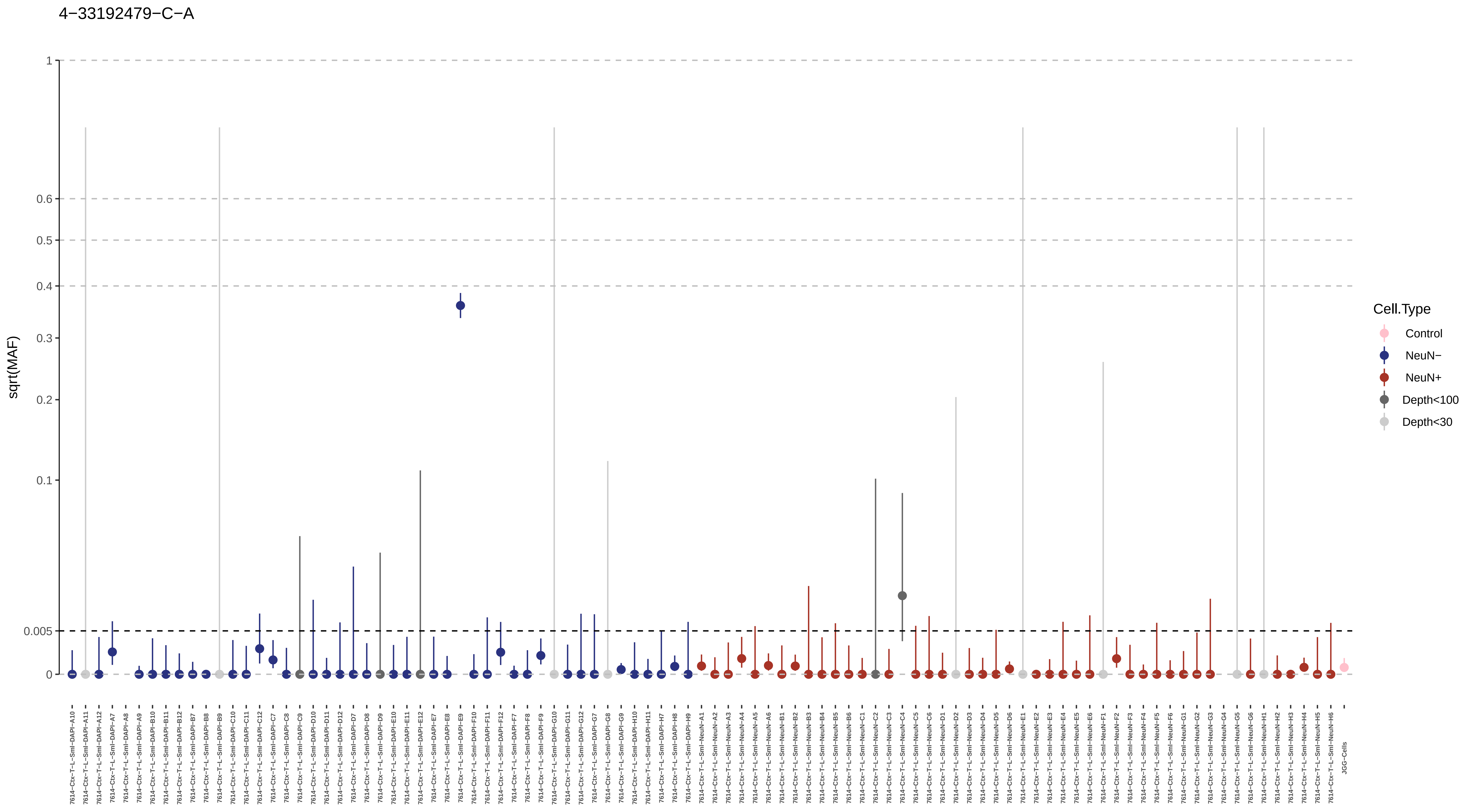

4−33192479−C−A
sqrt(MAF)
1
0.6
0.5
0.4
0.3
0.2
0.1
0.005
0
Cell.Type
Control
NeuN−
NeuN+
Depth<100
Depth<30
7614−Ctx−T−L−SmI−DAPI−A10
7614−Ctx−T−L−SmI−DAPI−A11
7614−Ctx−T−L−SmI−DAPI−A12
7614−Ctx−T−L−SmI−DAPI−A7
7614−Ctx−T−L−SmI−DAPI−A8
7614−Ctx−T−L−SmI−DAPI−A9
7614−Ctx−T−L−SmI−DAPI−B10
7614−Ctx−T−L−SmI−DAPI−B11
7614−Ctx−T−L−SmI−DAPI−B12
7614−Ctx−T−L−SmI−DAPI−B7
7614−Ctx−T−L−SmI−DAPI−B8
7614−Ctx−T−L−SmI−DAPI−B9
7614−Ctx−T−L−SmI−DAPI−C10
7614−Ctx−T−L−SmI−DAPI−C11
7614−Ctx−T−L−SmI−DAPI−C12
7614−Ctx−T−L−SmI−DAPI−C7
7614−Ctx−T−L−SmI−DAPI−C8
7614−Ctx−T−L−SmI−DAPI−C9
7614−Ctx−T−L−SmI−DAPI−D10
7614−Ctx−T−L−SmI−DAPI−D11
7614−Ctx−T−L−SmI−DAPI−D12
7614−Ctx−T−L−SmI−DAPI−D7
7614−Ctx−T−L−SmI−DAPI−D8
7614−Ctx−T−L−SmI−DAPI−D9
7614−Ctx−T−L−SmI−DAPI−E10
7614−Ctx−T−L−SmI−DAPI−E11
7614−Ctx−T−L−SmI−DAPI−E12
7614−Ctx−T−L−SmI−DAPI−E7
7614−Ctx−T−L−SmI−DAPI−E8
7614−Ctx−T−L−SmI−DAPI−E9
7614−Ctx−T−L−SmI−DAPI−F10
7614−Ctx−T−L−SmI−DAPI−F11
7614−Ctx−T−L−SmI−DAPI−F12
7614−Ctx−T−L−SmI−DAPI−F7
7614−Ctx−T−L−SmI−DAPI−F8
7614−Ctx−T−L−SmI−DAPI−F9
7614−Ctx−T−L−SmI−DAPI−G10
7614−Ctx−T−L−SmI−DAPI−G11
7614−Ctx−T−L−SmI−DAPI−G12
7614−Ctx−T−L−SmI−DAPI−G7
7614−Ctx−T−L−SmI−DAPI−G8
7614−Ctx−T−L−SmI−DAPI−G9
7614−Ctx−T−L−SmI−DAPI−H10
7614−Ctx−T−L−SmI−DAPI−H11
7614−Ctx−T−L−SmI−DAPI−H7
7614−Ctx−T−L−SmI−DAPI−H8
7614−Ctx−T−L−SmI−DAPI−H9
7614−Ctx−T−L−SmI−NeuN−A1
7614−Ctx−T−L−SmI−NeuN−A2
7614−Ctx−T−L−SmI−NeuN−A3
7614−Ctx−T−L−SmI−NeuN−A4
7614−Ctx−T−L−SmI−NeuN−A5
7614−Ctx−T−L−SmI−NeuN−A6
7614−Ctx−T−L−SmI−NeuN−B1
7614−Ctx−T−L−SmI−NeuN−B2
7614−Ctx−T−L−SmI−NeuN−B3
7614−Ctx−T−L−SmI−NeuN−B4
7614−Ctx−T−L−SmI−NeuN−B5
7614−Ctx−T−L−SmI−NeuN−B6
7614−Ctx−T−L−SmI−NeuN−C1
7614−Ctx−T−L−SmI−NeuN−C2
7614−Ctx−T−L−SmI−NeuN−C3
7614−Ctx−T−L−SmI−NeuN−C4
7614−Ctx−T−L−SmI−NeuN−C5
7614−Ctx−T−L−SmI−NeuN−C6
7614−Ctx−T−L−SmI−NeuN−D1
7614−Ctx−T−L−SmI−NeuN−D2
7614−Ctx−T−L−SmI−NeuN−D3
7614−Ctx−T−L−SmI−NeuN−D4
7614−Ctx−T−L−SmI−NeuN−D5
7614−Ctx−T−L−SmI−NeuN−D6
7614−Ctx−T−L−SmI−NeuN−E1
7614−Ctx−T−L−SmI−NeuN−E2
7614−Ctx−T−L−SmI−NeuN−E3
7614−Ctx−T−L−SmI−NeuN−E4
7614−Ctx−T−L−SmI−NeuN−E5
7614−Ctx−T−L−SmI−NeuN−E6
7614−Ctx−T−L−SmI−NeuN−F1
7614−Ctx−T−L−SmI−NeuN−F2
7614−Ctx−T−L−SmI−NeuN−F3
7614−Ctx−T−L−SmI−NeuN−F4
7614−Ctx−T−L−SmI−NeuN−F5
7614−Ctx−T−L−SmI−NeuN−F6
7614−Ctx−T−L−SmI−NeuN−G1
7614−Ctx−T−L−SmI−NeuN−G2
7614−Ctx−T−L−SmI−NeuN−G3
7614−Ctx−T−L−SmI−NeuN−G4
7614−Ctx−T−L−SmI−NeuN−G5
7614−Ctx−T−L−SmI−NeuN−G6
7614−Ctx−T−L−SmI−NeuN−H1
7614−Ctx−T−L−SmI−NeuN−H2
7614−Ctx−T−L−SmI−NeuN−H3
7614−Ctx−T−L−SmI−NeuN−H4
7614−Ctx−T−L−SmI−NeuN−H5
7614−Ctx−T−L−SmI−NeuN−H6
JGG−Cells

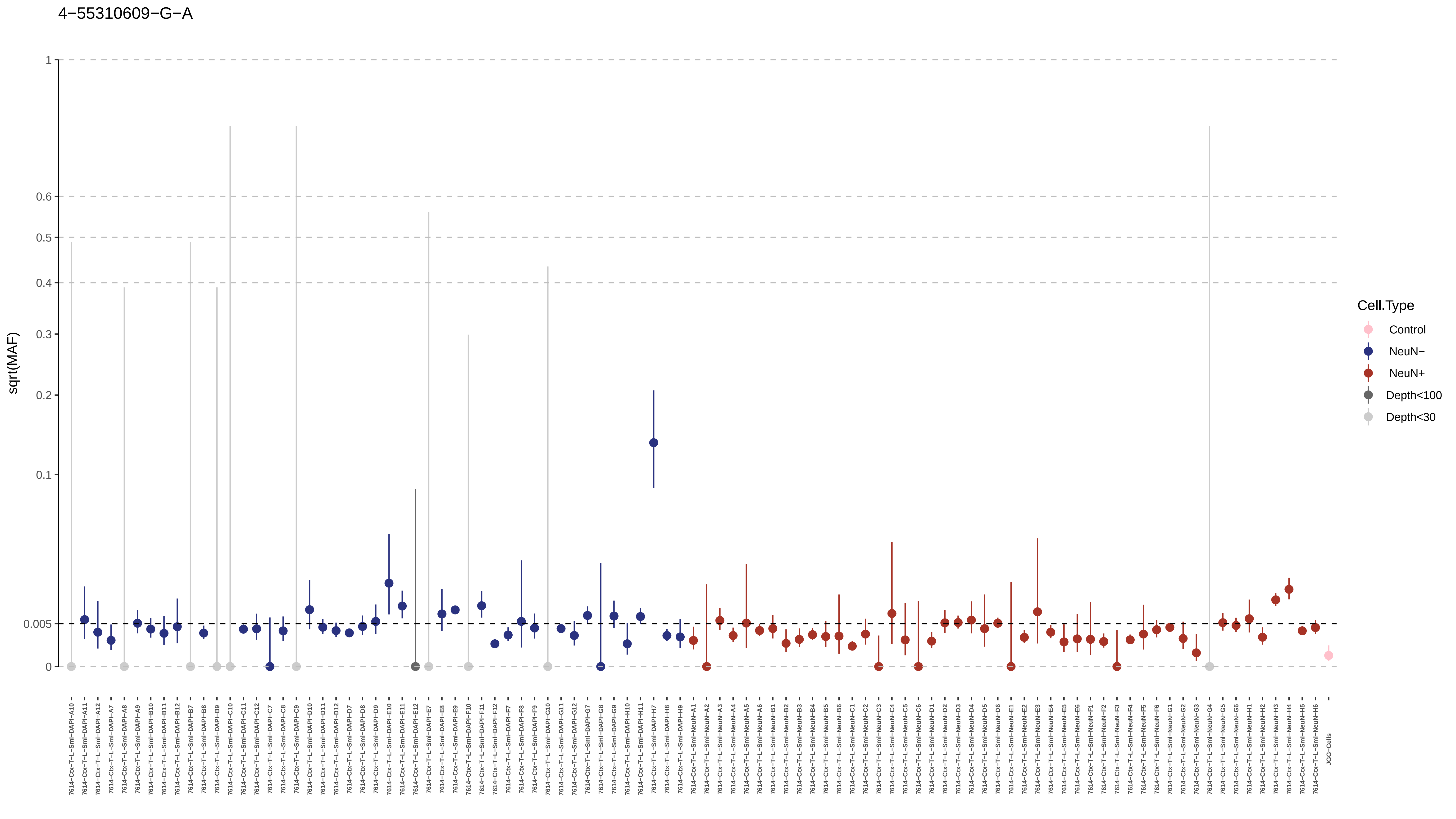

4−55310609−G−A
sqrt(MAF)
1
0.6
0.5
0.4
0.3
0.2
0.1
0.005
0
Cell.Type
Control
NeuN−
NeuN+
Depth<100
Depth<30

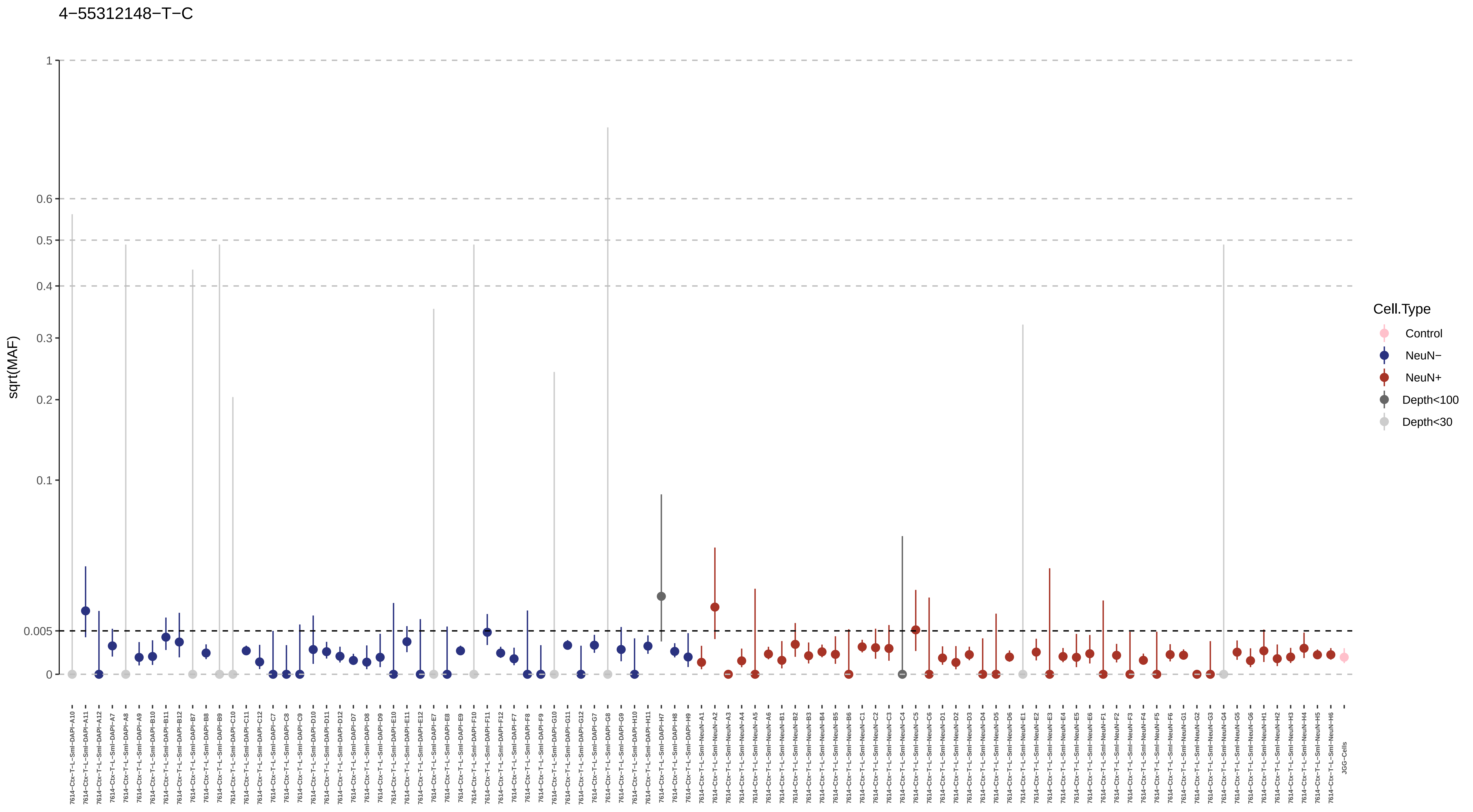

4−55312148−T−C
sqrt(MAF)
1
0.6
0.5
0.4
0.3
0.2
0.1
0.005
0
Cell.Type
Control
NeuN−
NeuN+
Depth<100
Depth<30

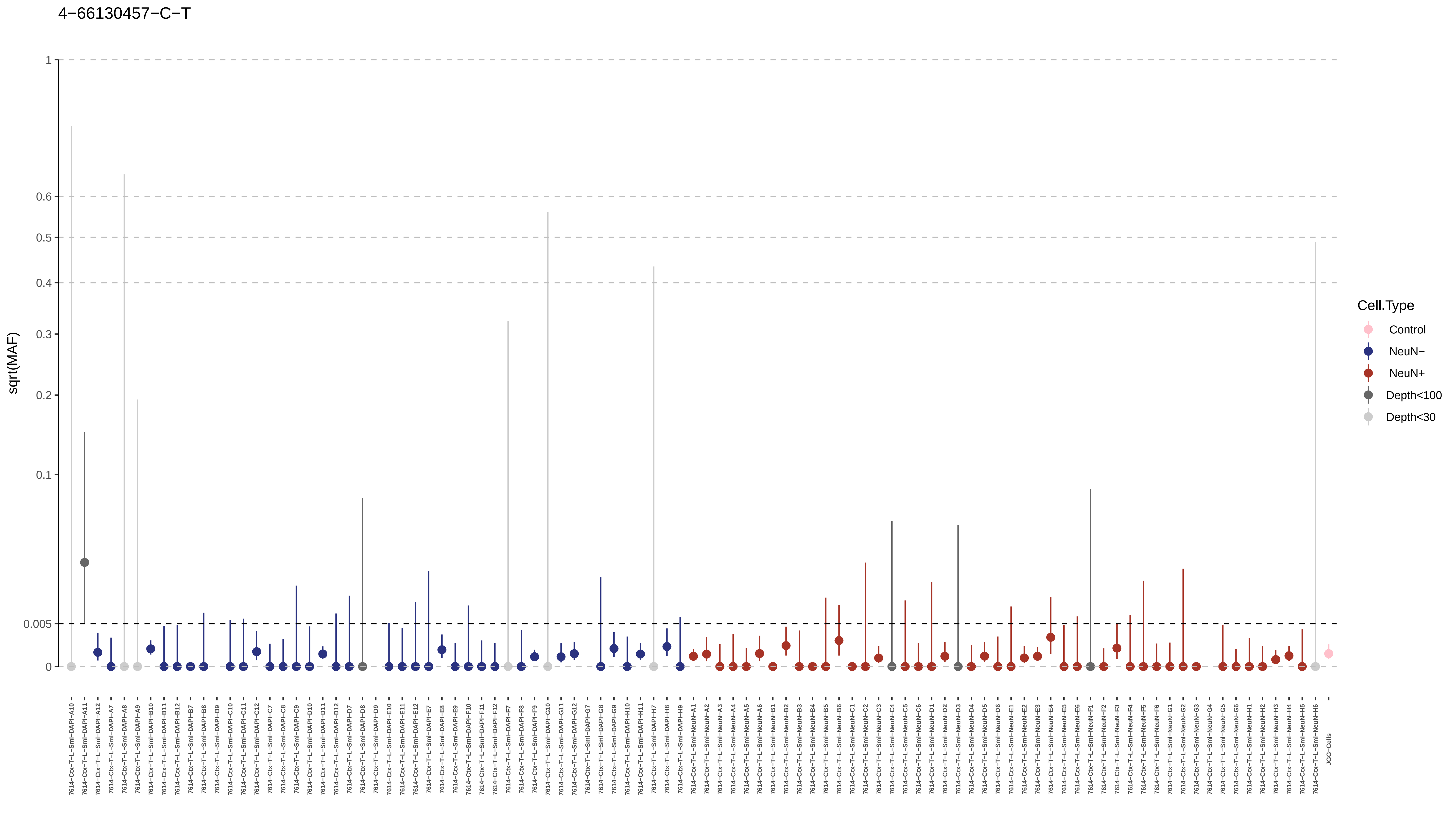
4−66130457−C−T
sqrt(MAF)
1
0.6
0.5
0.4
0.3
0.2
0.1
0.005
0
Cell.Type
Control
NeuN−
NeuN+
Depth<100
Depth<30
JGG-Cells

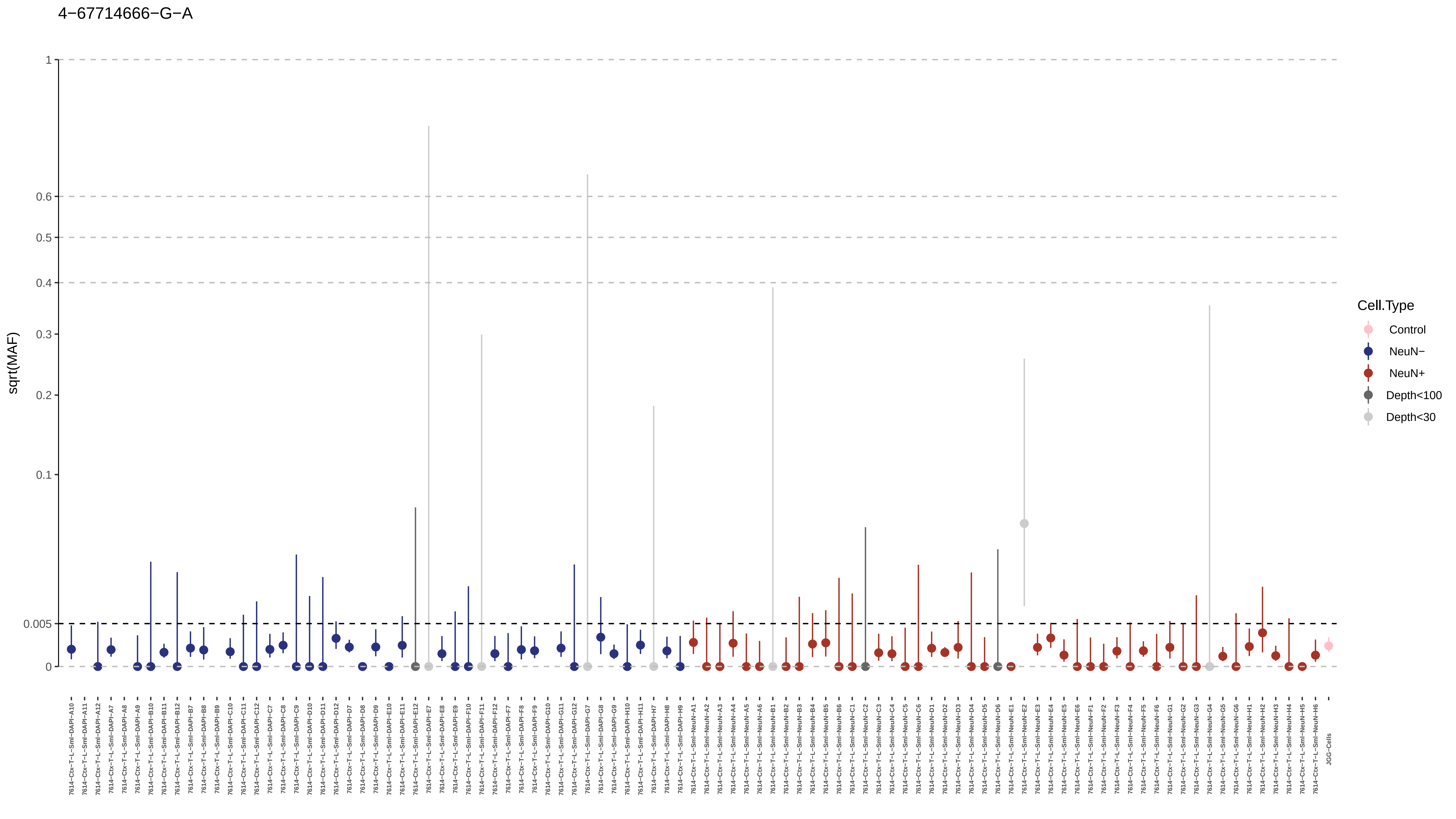
4−67714666−G−A
sqrt(MAF)
1
0.6
0.5
0.4
0.3
0.2
0.1
0.005
0
Cell.Type
Control
NeuN−
NeuN+
Depth<100
Depth<30
JGG-Cells

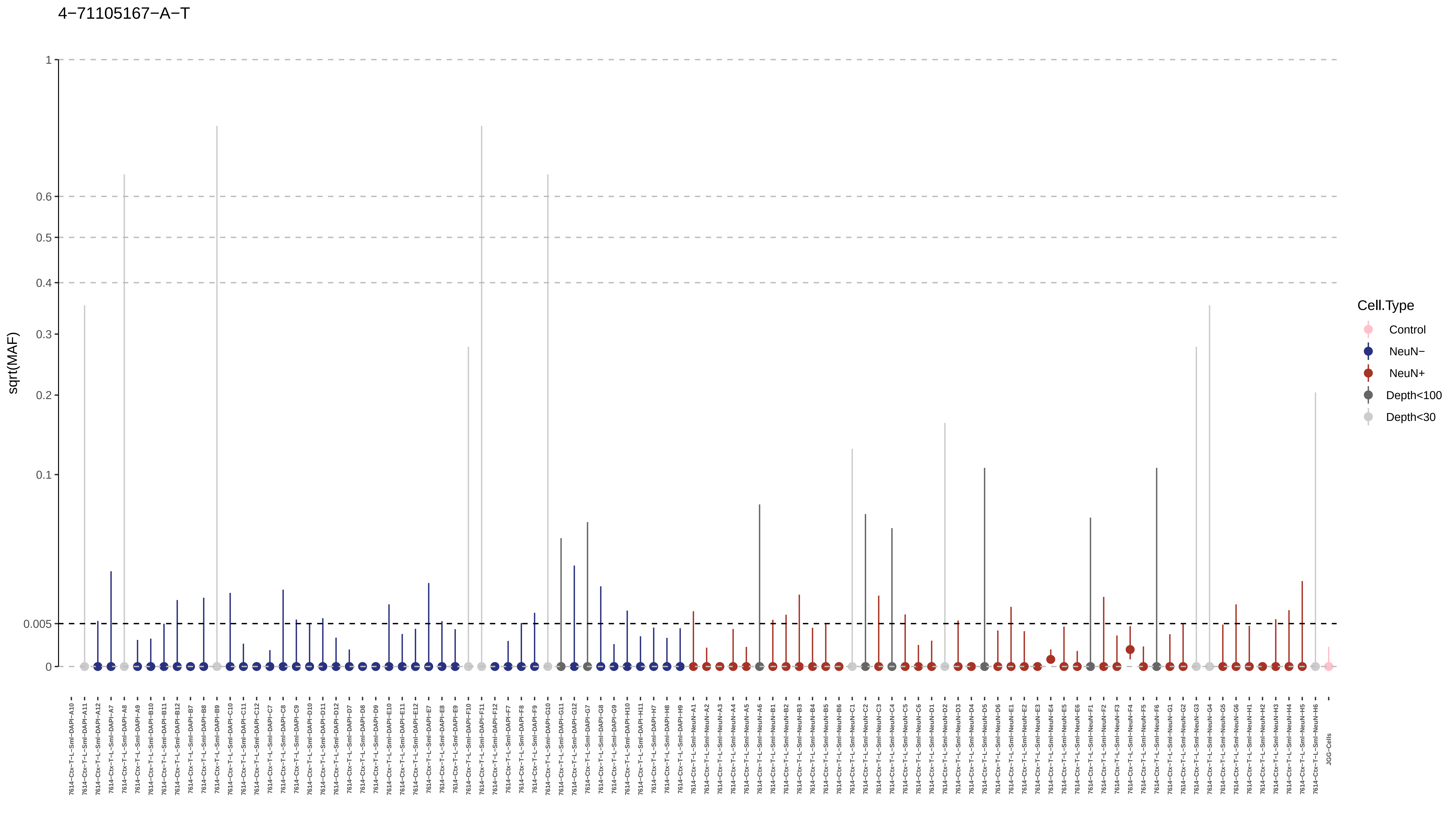

4−71105167−A−T
sqrt(MAF)
1
0.6
0.5
0.4
0.3
0.2
0.1
0.005
0
Cell.Type
Control
NeuN−
NeuN+
Depth<100
Depth<30

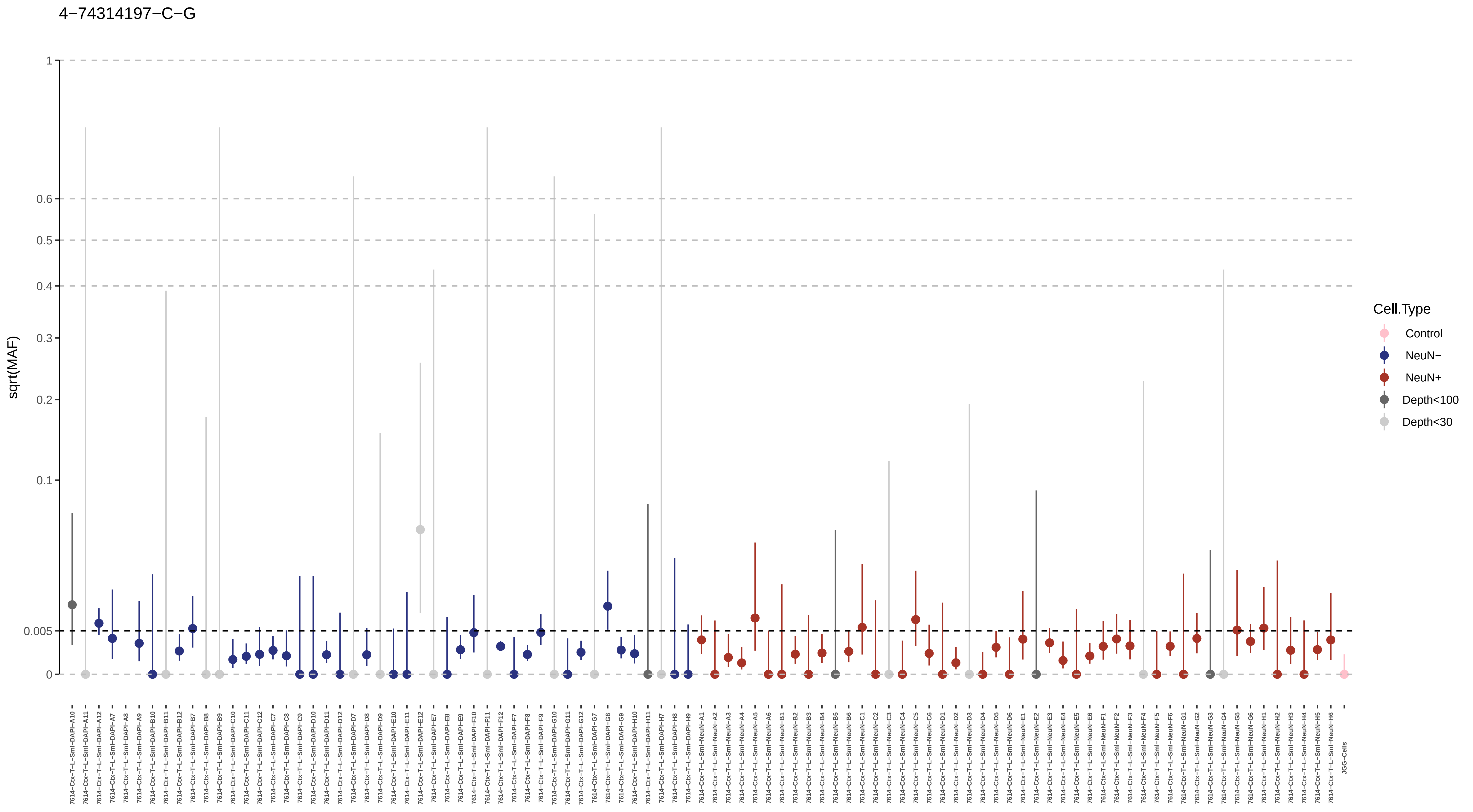
4−74314197−C−G
sqrt(MAF)
1
0.6
0.5
0.4
0.3
0.2
0.1
0.005
0
Cell.Type
Control
NeuN−
NeuN+
Depth<100
Depth<30

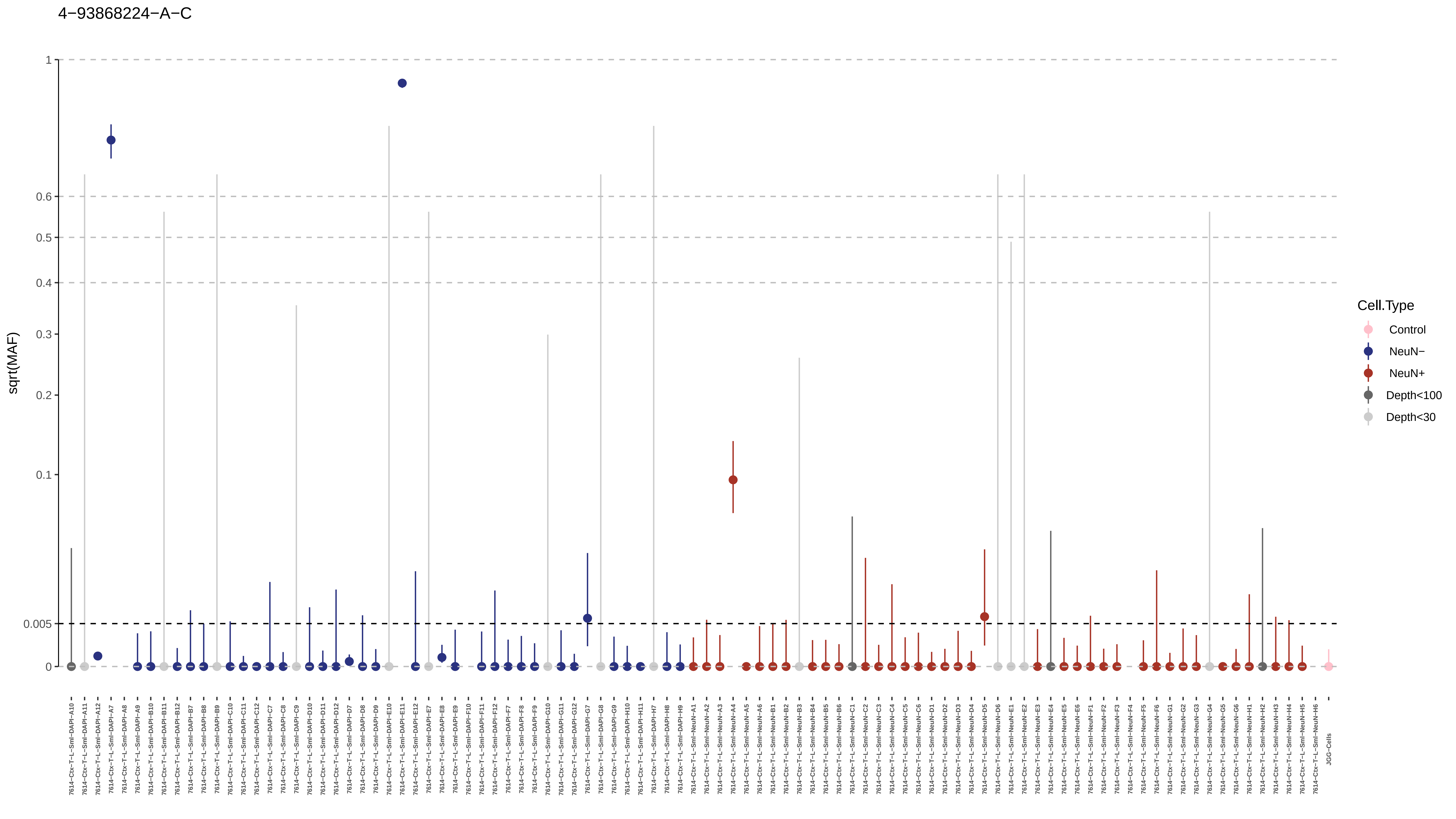
4−93868224−A−C
sqrt(MAF)
1
0.6
0.5
0.4
0.3
0.2
0.1
0.005
0
Cell.Type
Control
NeuN−
NeuN+
Depth<100
Depth<30

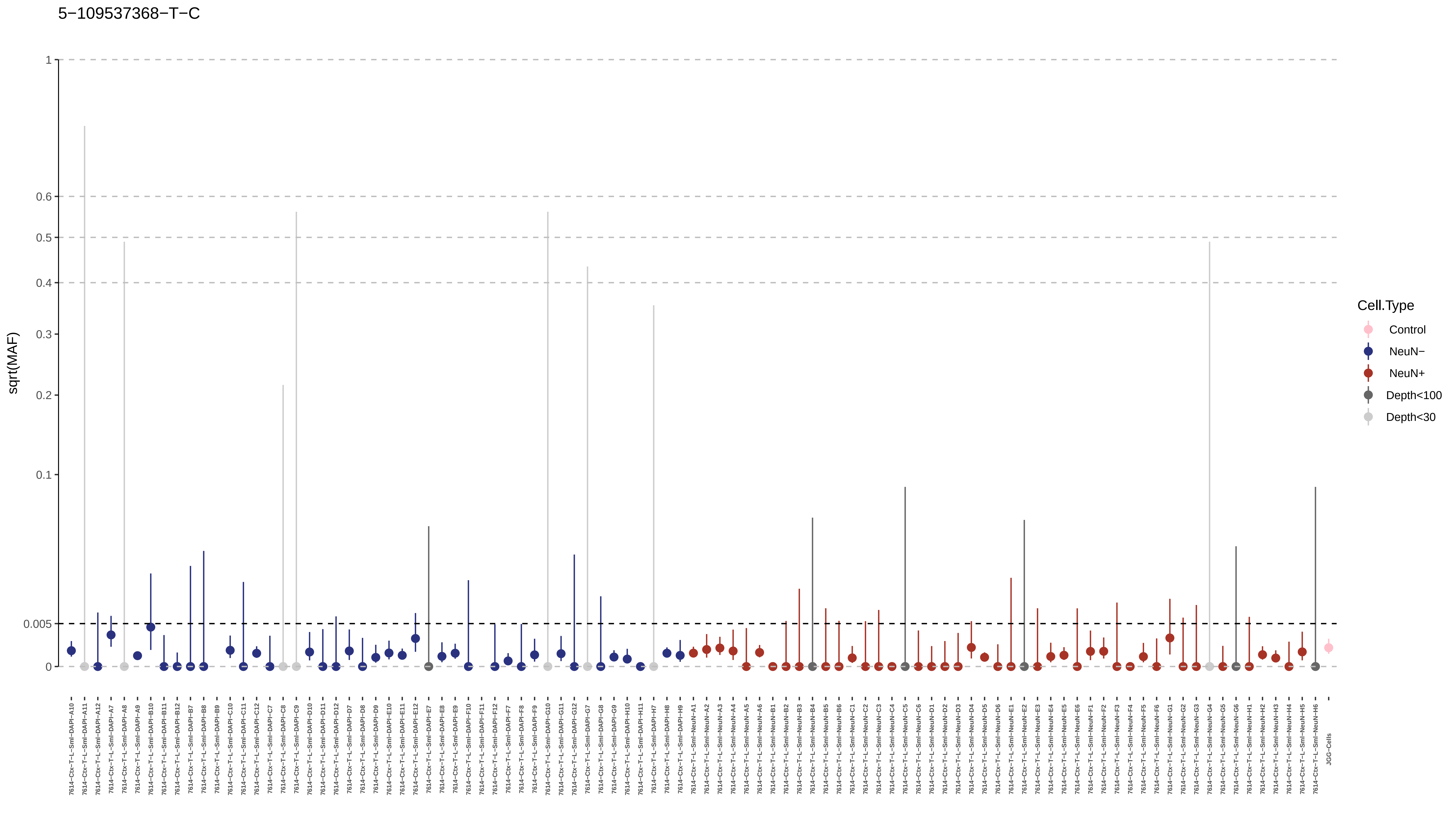

5−109537368−T−C
sqrt(MAF)
1
0.6
0.5
0.4
0.3
0.2
0.1
0.005
0
Cell.Type
Control
NeuN−
NeuN+
Depth<100
Depth<30

5−132659937−G−A

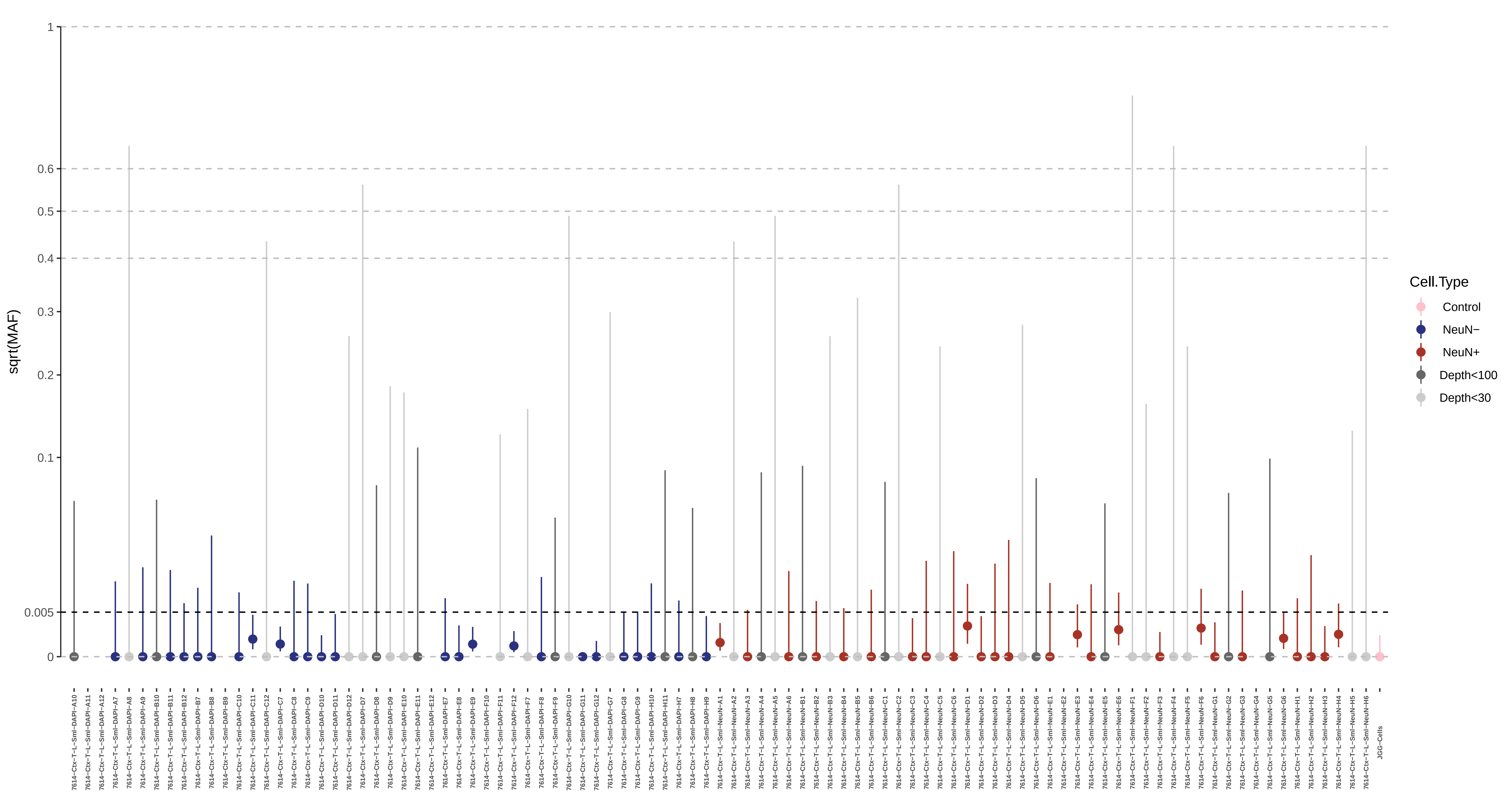

5−146867769−A−G

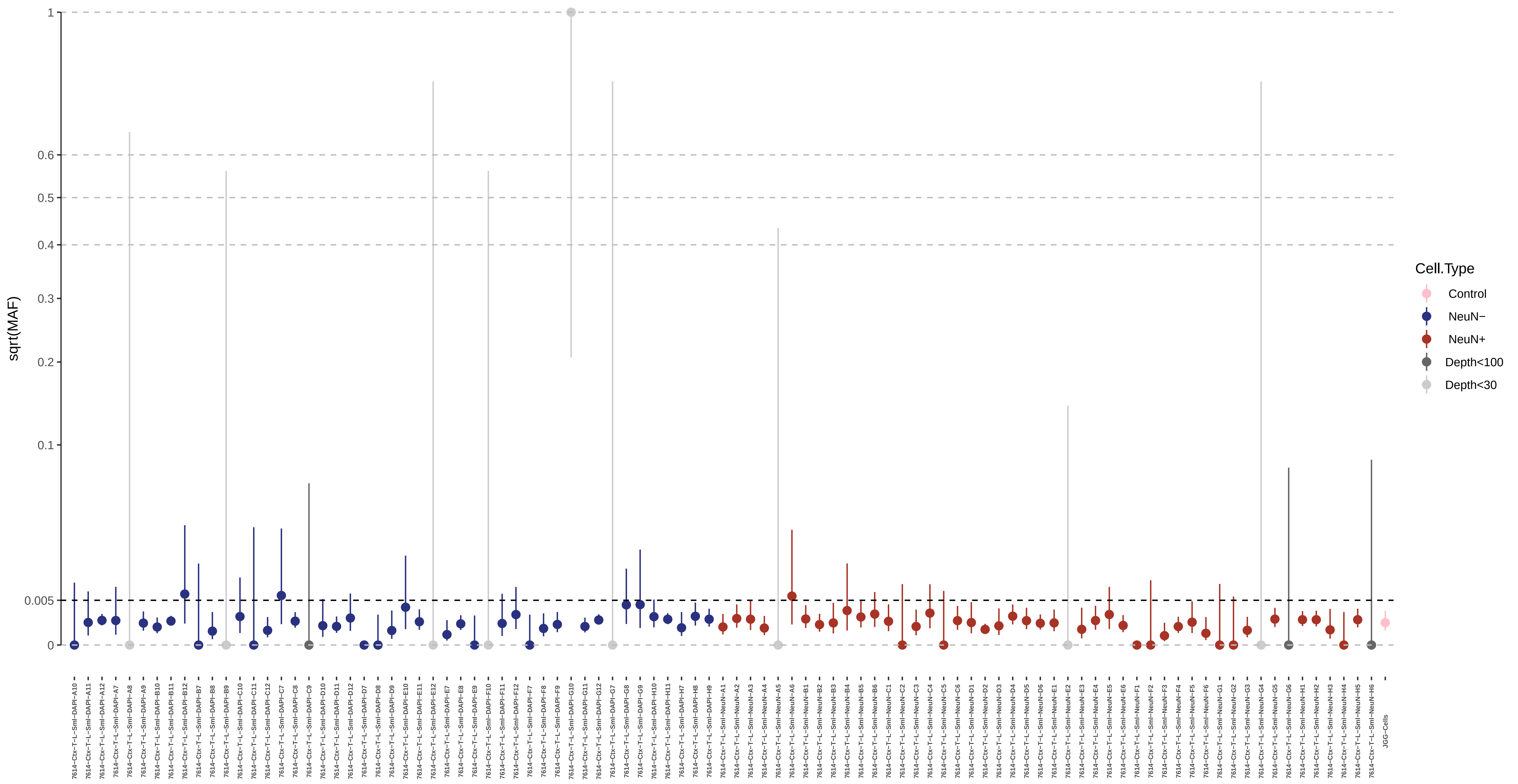

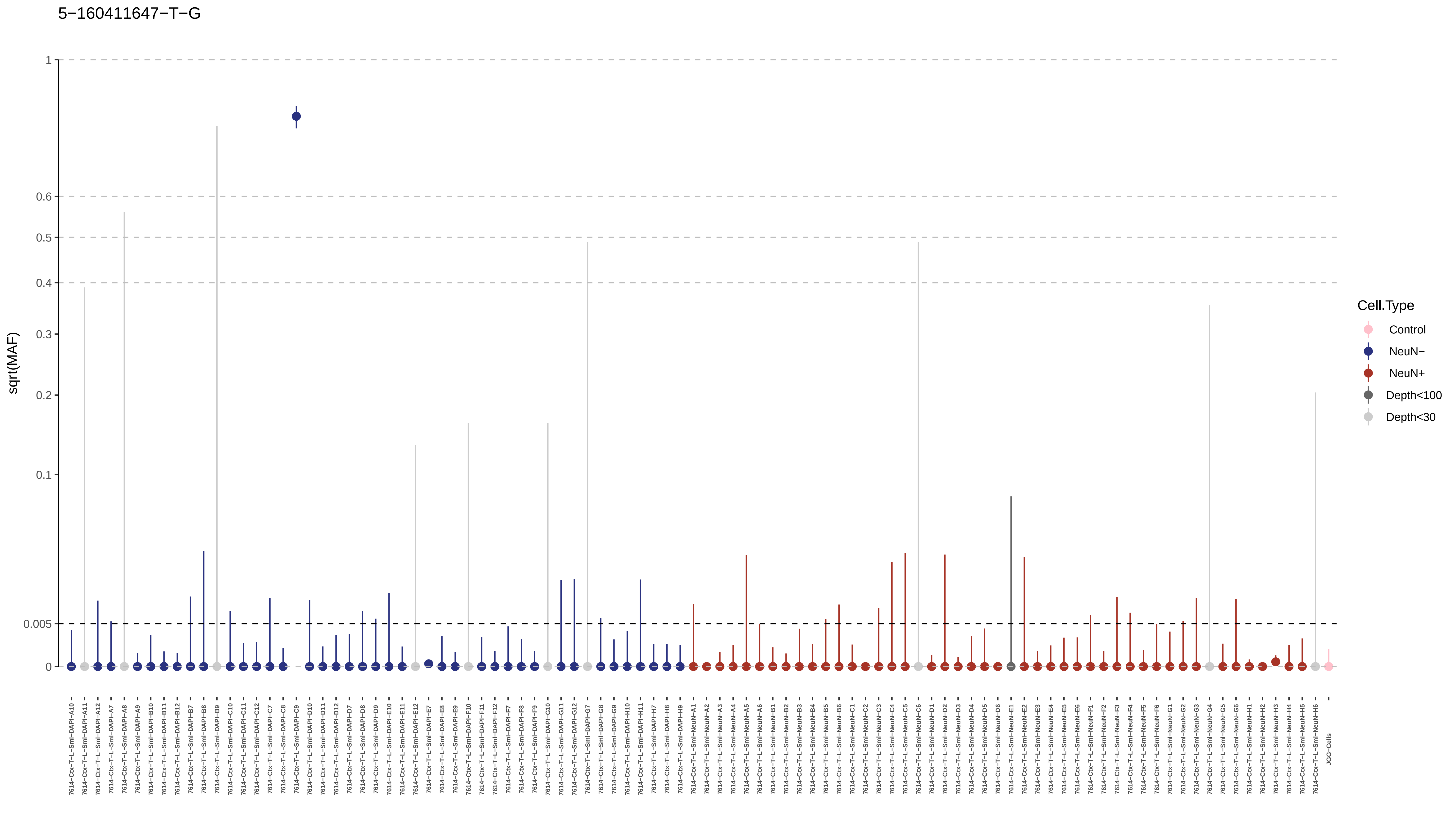
5−160411647−T−G
sqrt(MAF)
1
0.6
0.5
0.4
0.3
0.2
0.1
0.005
0
Cell.Type
Control
NeuN−
NeuN+
Depth<100
Depth<30

5−17226551−A−G

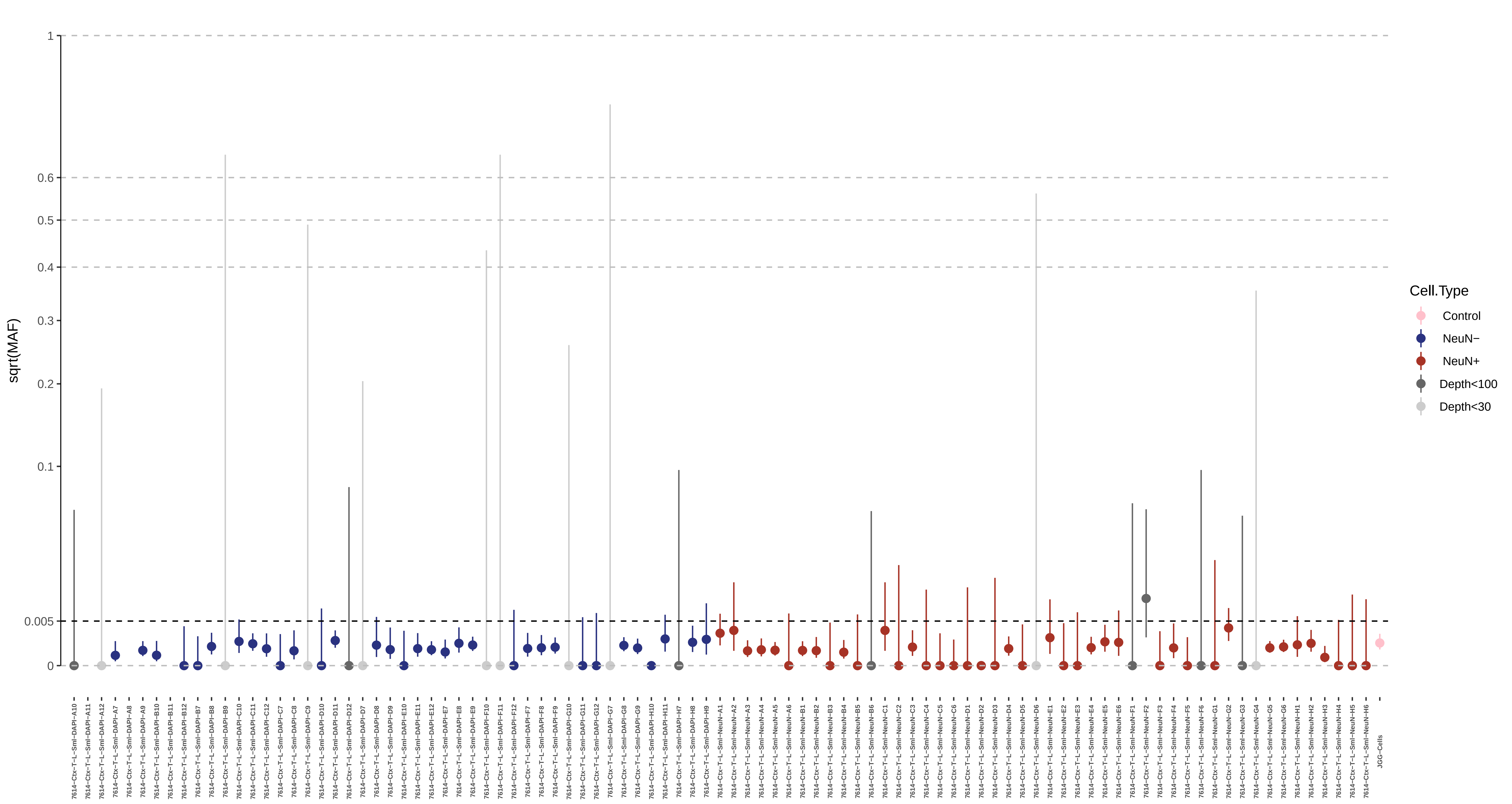

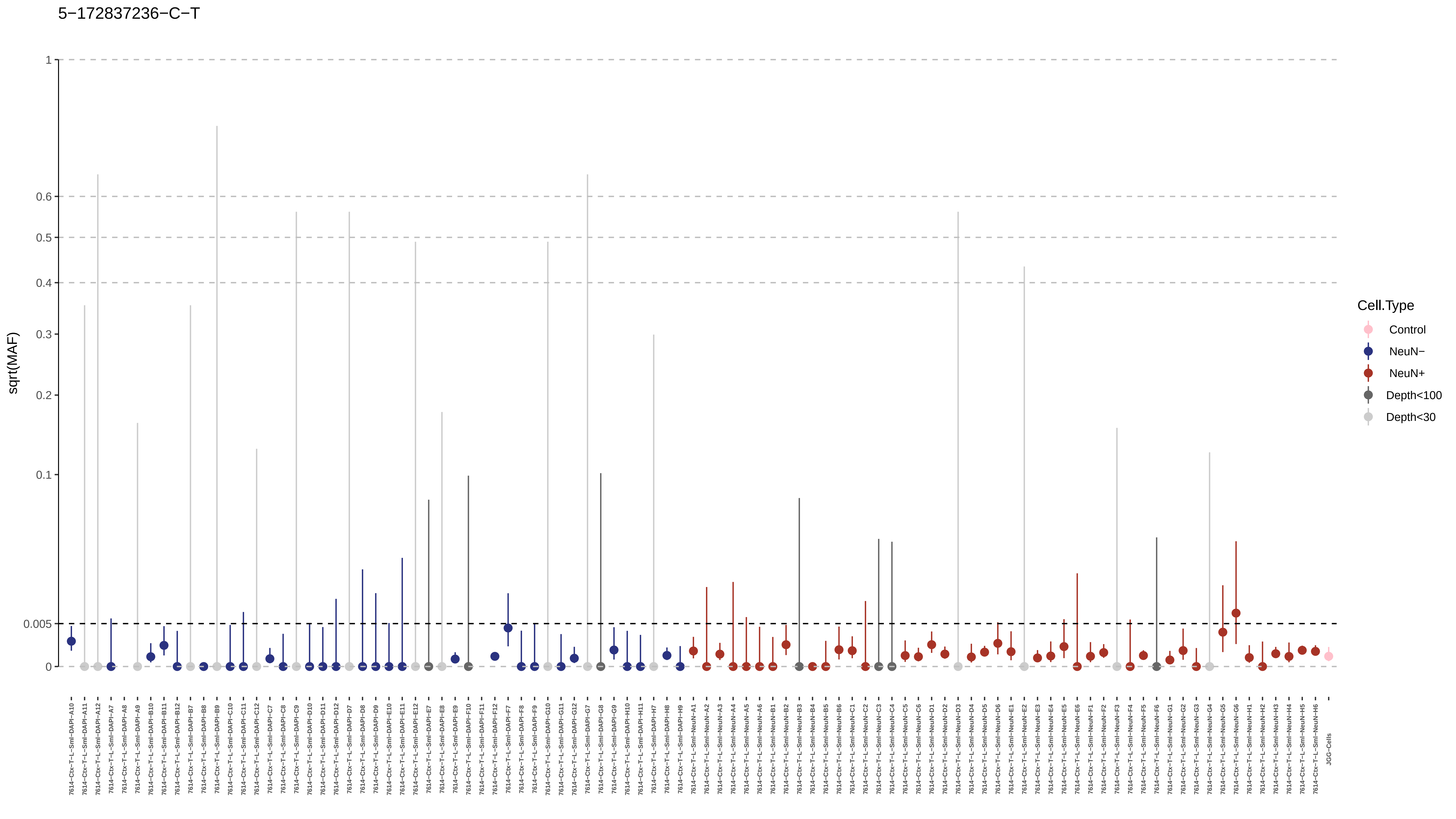

5−172837236−C−T
sqrt(MAF)
1
0.6
0.5
0.4
0.3
0.2
0.1
0.005
0
Cell.Type
Control
NeuN−
NeuN+
Depth<100
Depth<30
JGG-Cells

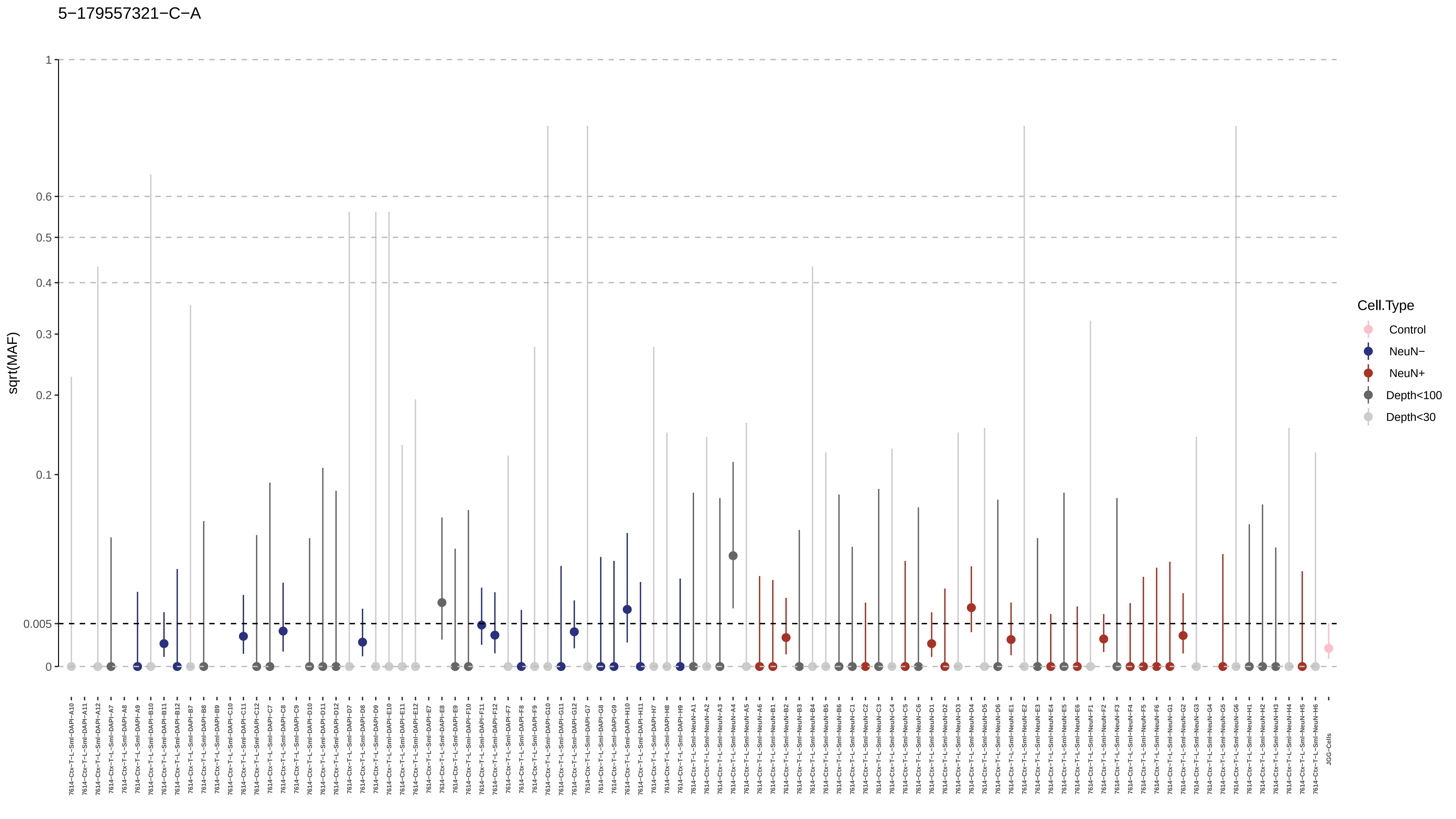
5−179557321−C−A
sqrt(MAF)
1
0.6
0.5
0.4
0.3
0.2
0.1
0.005
0
Cell.Type
Control
NeuN−
NeuN+
Depth<100
Depth<30
JGG-Cells

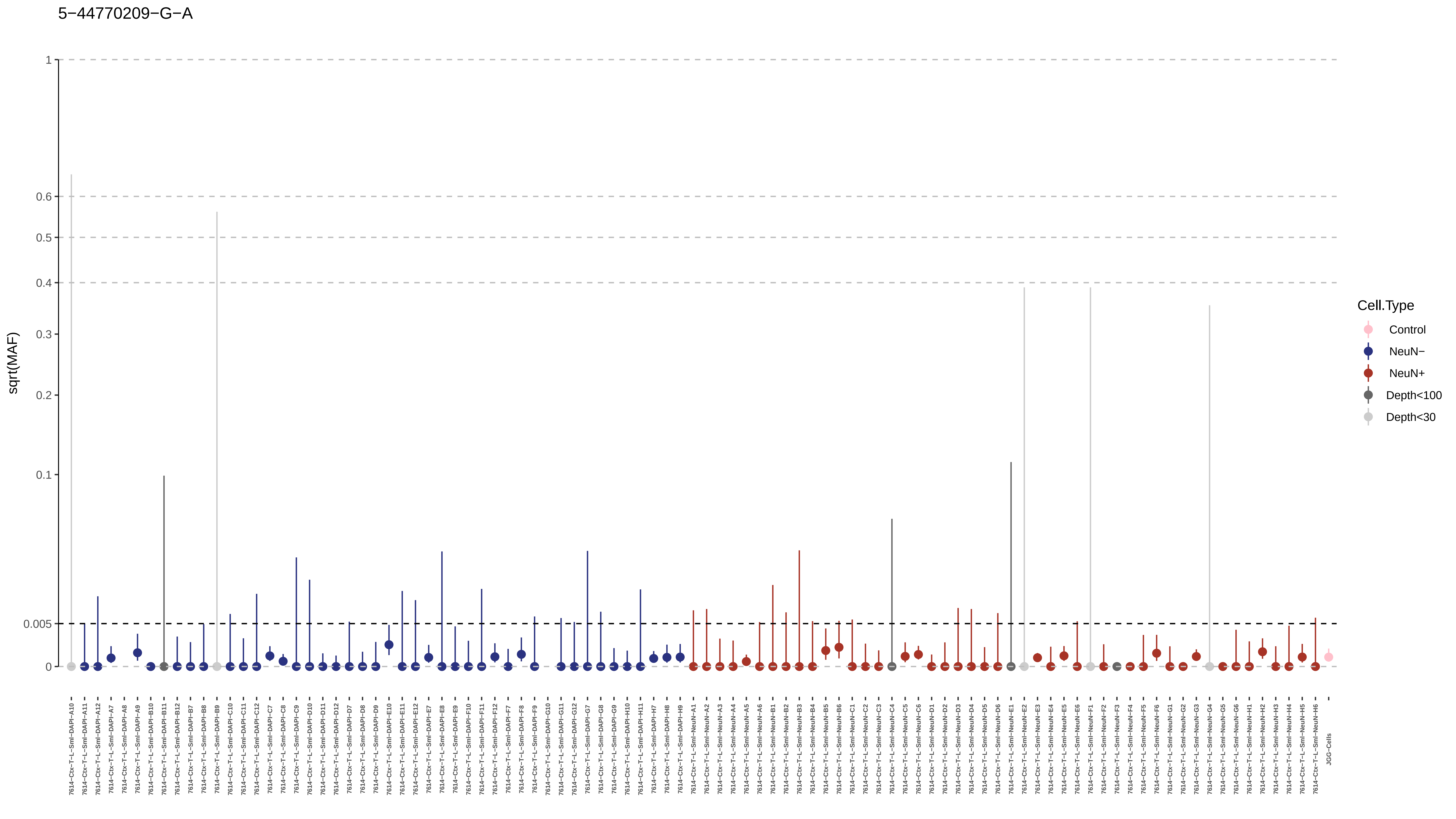
5−44770209−G−A
sqrt(MAF)
1
0.6
0.5
0.4
0.3
0.2
0.1
0.005
0
Cell.Type
Control
NeuN−
NeuN+
Depth<100
Depth<30
JGG-Cells

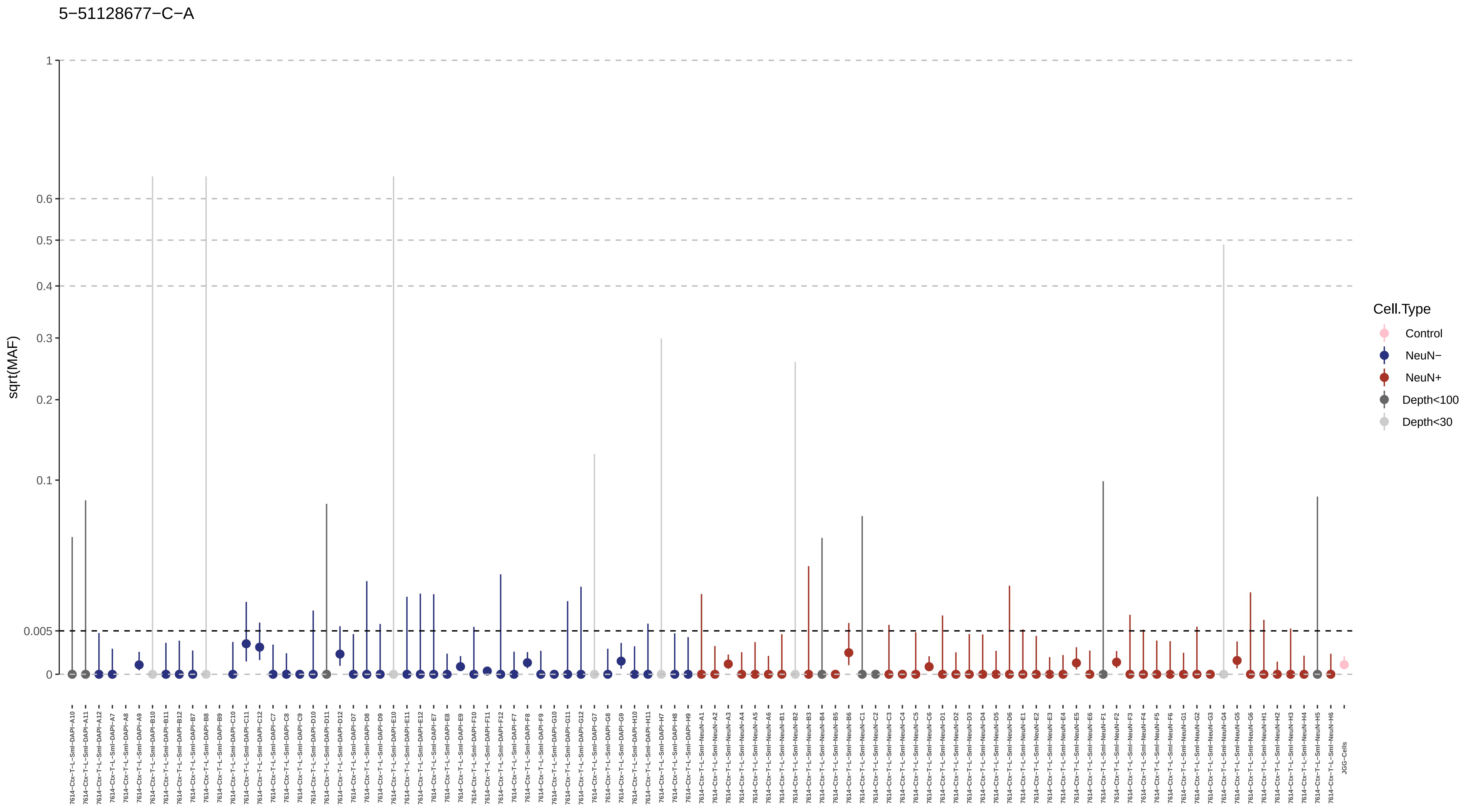
5−51128677−C−A
sqrt(MAF)
1
0.6
0.5
0.4
0.3
0.2
0.1
0.005
0
Cell.Type
Control
NeuN−
NeuN+
Depth<100
Depth<30

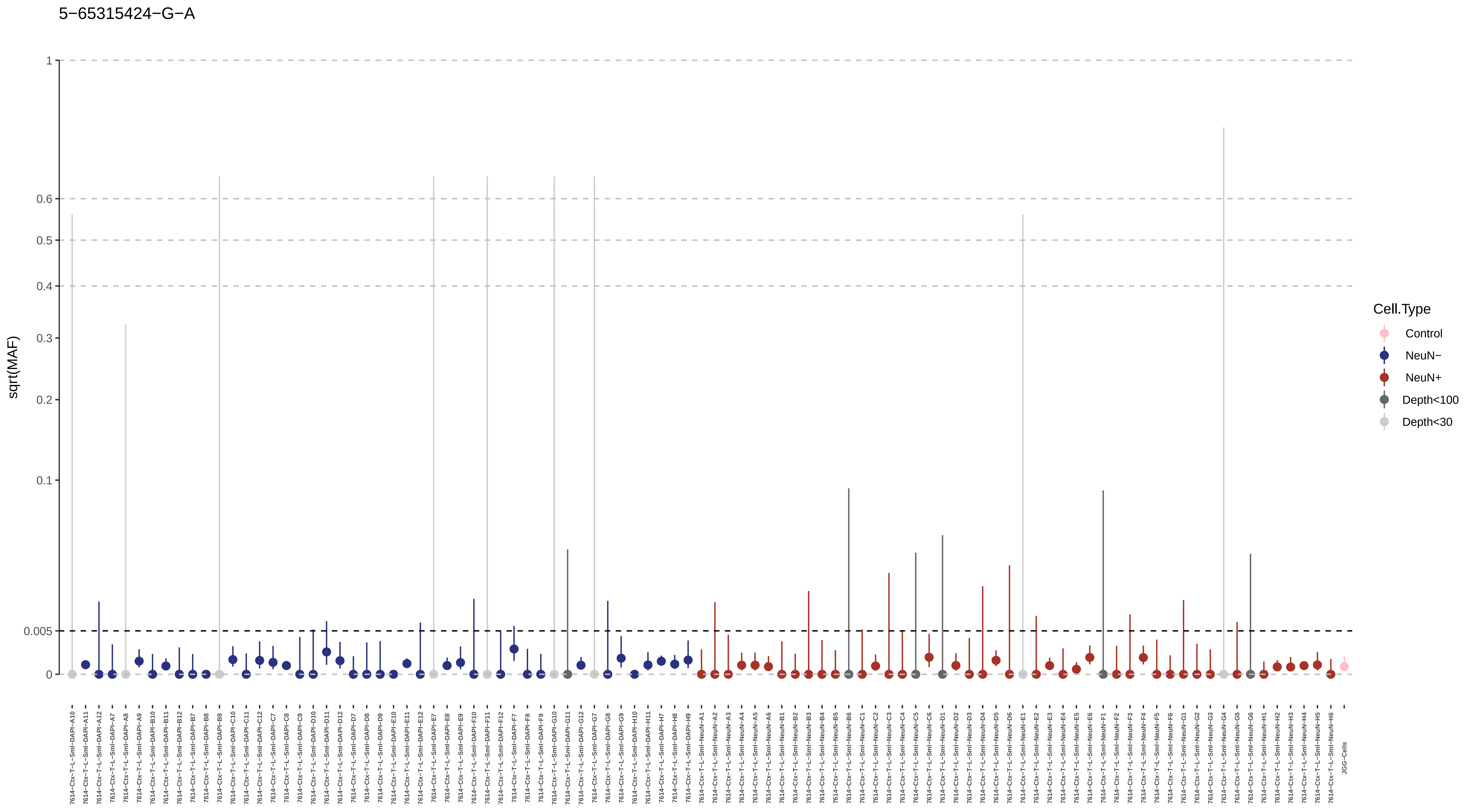
5−65315424−G−A
sqrt(MAF)
1
0.6
0.5
0.4
0.3
0.2
0.1
0.005
0
Cell.Type
Control
NeuN−
NeuN+
Depth<100
Depth<30

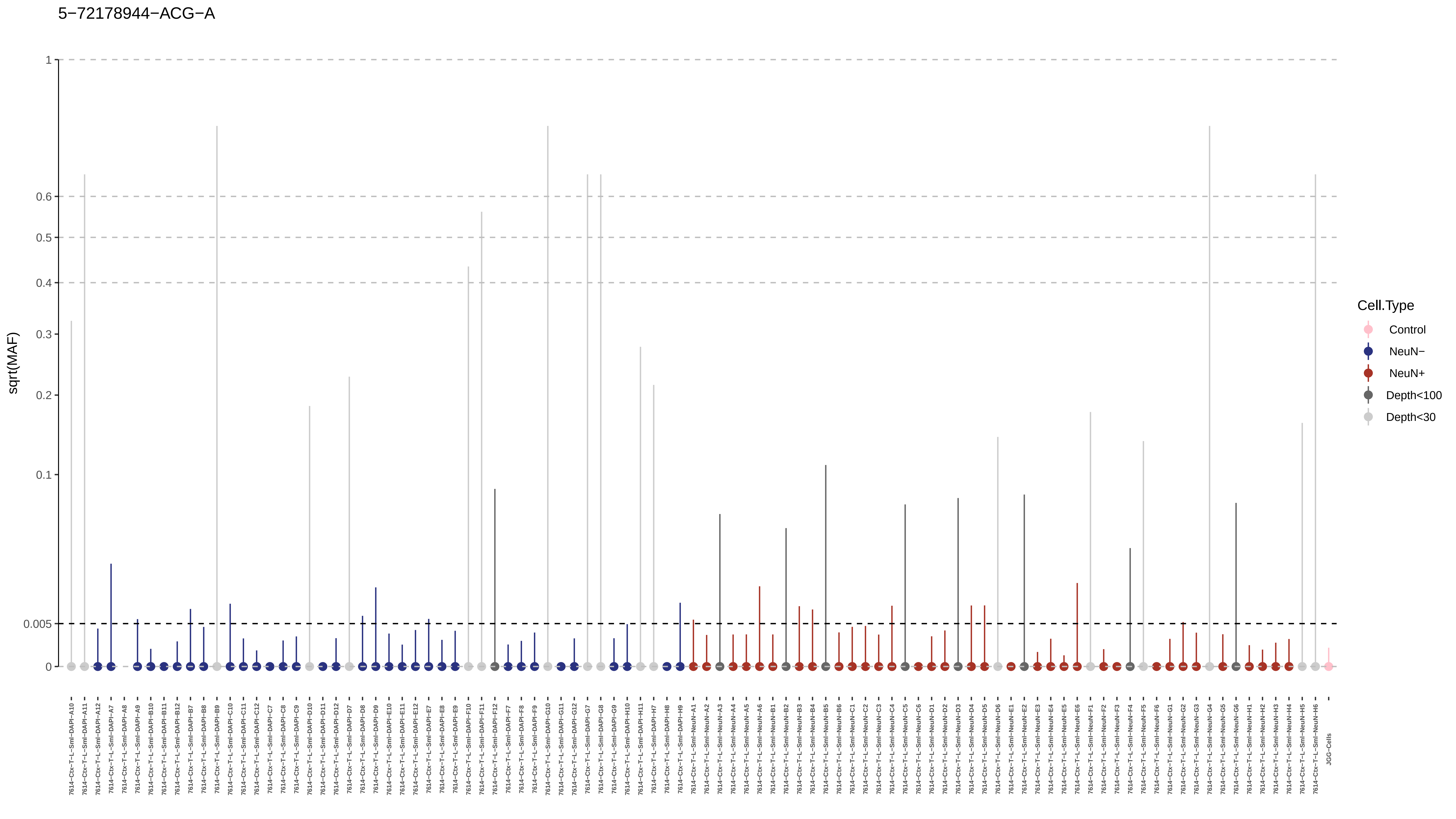

5−72178944−ACG−A
sqrt(MAF)
1
0.6
0.5
0.4
0.3
0.2
0.1
0.005
0
Cell.Type
Control
NeuN−
NeuN+
Depth<100
Depth<30
7614−Ctx−T−L−Sml−DAPI−A10
7614−Ctx−T−L−Sml−DAPI−A11
7614−Ctx−T−L−Sml−DAPI−A12
7614−Ctx−T−L−Sml−DAPI−A7
7614−Ctx−T−L−Sml−DAPI−A8
7614−Ctx−T−L−Sml−DAPI−A9
7614−Ctx−T−L−Sml−DAPI−B10
7614−Ctx−T−L−Sml−DAPI−B11
7614−Ctx−T−L−Sml−DAPI−B12
7614−Ctx−T−L−Sml−DAPI−B7
7614−Ctx−T−L−Sml−DAPI−B8
7614−Ctx−T−L−Sml−DAPI−B9
7614−Ctx−T−L−Sml−DAPI−C10
7614−Ctx−T−L−Sml−DAPI−C11
7614−Ctx−T−L−Sml−DAPI−C12
7614−Ctx−T−L−Sml−DAPI−C7
7614−Ctx−T−L−Sml−DAPI−C8
7614−Ctx−T−L−Sml−DAPI−C9
7614−Ctx−T−L−Sml−DAPI−D10
7614−Ctx−T−L−Sml−DAPI−D11
7614−Ctx−T−L−Sml−DAPI−D12
7614−Ctx−T−L−Sml−DAPI−D7
7614−Ctx−T−L−Sml−DAPI−D8
7614−Ctx−T−L−Sml−DAPI−D9
7614−Ctx−T−L−Sml−DAPI−E10
7614−Ctx−T−L−Sml−DAPI−E11
7614−Ctx−T−L−Sml−DAPI−E12
7614−Ctx−T−L−Sml−DAPI−E7
7614−Ctx−T−L−Sml−DAPI−E8
7614−Ctx−T−L−Sml−DAPI−E9
7614−Ctx−T−L−Sml−DAPI−F10
7614−Ctx−T−L−Sml−DAPI−F11
7614−Ctx−T−L−Sml−DAPI−F12
7614−Ctx−T−L−Sml−DAPI−F7
7614−Ctx−T−L−Sml−DAPI−F8
7614−Ctx−T−L−Sml−DAPI−F9
7614−Ctx−T−L−Sml−DAPI−G10
7614−Ctx−T−L−Sml−DAPI−G11
7614−Ctx−T−L−Sml−DAPI−G12
7614−Ctx−T−L−Sml−DAPI−G7
7614−Ctx−T−L−Sml−DAPI−G8
7614−Ctx−T−L−Sml−DAPI−G9
7614−Ctx−T−L−Sml−DAPI−H10
7614−Ctx−T−L−Sml−DAPI−H11
7614−Ctx−T−L−Sml−DAPI−H7
7614−Ctx−T−L−Sml−DAPI−H8
7614−Ctx−T−L−Sml−DAPI−H9
7614−Ctx−T−L−Sml−NeuN−A1
7614−Ctx−T−L−Sml−NeuN−A2
7614−Ctx−T−L−Sml−NeuN−A3
7614−Ctx−T−L−Sml−NeuN−A4
7614−Ctx−T−L−Sml−NeuN−A5
7614−Ctx−T−L−Sml−NeuN−A6
7614−Ctx−T−L−Sml−NeuN−B1
7614−Ctx−T−L−Sml−NeuN−B2
7614−Ctx−T−L−Sml−NeuN−B3
7614−Ctx−T−L−Sml−NeuN−B4
7614−Ctx−T−L−Sml−NeuN−B5
7614−Ctx−T−L−Sml−NeuN−B6
7614−Ctx−T−L−Sml−NeuN−C1
7614−Ctx−T−L−Sml−NeuN−C2
7614−Ctx−T−L−Sml−NeuN−C3
7614−Ctx−T−L−Sml−NeuN−C4
7614−Ctx−T−L−Sml−NeuN−C5
7614−Ctx−T−L−Sml−NeuN−C6
7614−Ctx−T−L−Sml−NeuN−D1
7614−Ctx−T−L−Sml−NeuN−D2
7614−Ctx−T−L−Sml−NeuN−D3
7614−Ctx−T−L−Sml−NeuN−D4
7614−Ctx−T−L−Sml−NeuN−D5
7614−Ctx−T−L−Sml−NeuN−D6
7614−Ctx−T−L−Sml−NeuN−E1
7614−Ctx−T−L−Sml−NeuN−E2
7614−Ctx−T−L−Sml−NeuN−E3
7614−Ctx−T−L−Sml−NeuN−E4
7614−Ctx−T−L−Sml−NeuN−E5
7614−Ctx−T−L−Sml−NeuN−E6
7614−Ctx−T−L−Sml−NeuN−F1
7614−Ctx−T−L−Sml−NeuN−F2
7614−Ctx−T−L−Sml−NeuN−F3
7614−Ctx−T−L−Sml−NeuN−F4
7614−Ctx−T−L−Sml−NeuN−F5
7614−Ctx−T−L−Sml−NeuN−F6
7614−Ctx−T−L−Sml−NeuN−G1
7614−Ctx−T−L−Sml−NeuN−G2
7614−Ctx−T−L−Sml−NeuN−G3
7614−Ctx−T−L−Sml−NeuN−G4
7614−Ctx−T−L−Sml−NeuN−G5
7614−Ctx−T−L−Sml−NeuN−G6
7614−Ctx−T−L−Sml−NeuN−H1
7614−Ctx−T−L−Sml−NeuN−H2
7614−Ctx−T−L−Sml−NeuN−H3
7614−Ctx−T−L−Sml−NeuN−H4
7614−Ctx−T−L−Sml−NeuN−H5
7614−Ctx−T−L−Sml−NeuN−H6
JGG−Cells

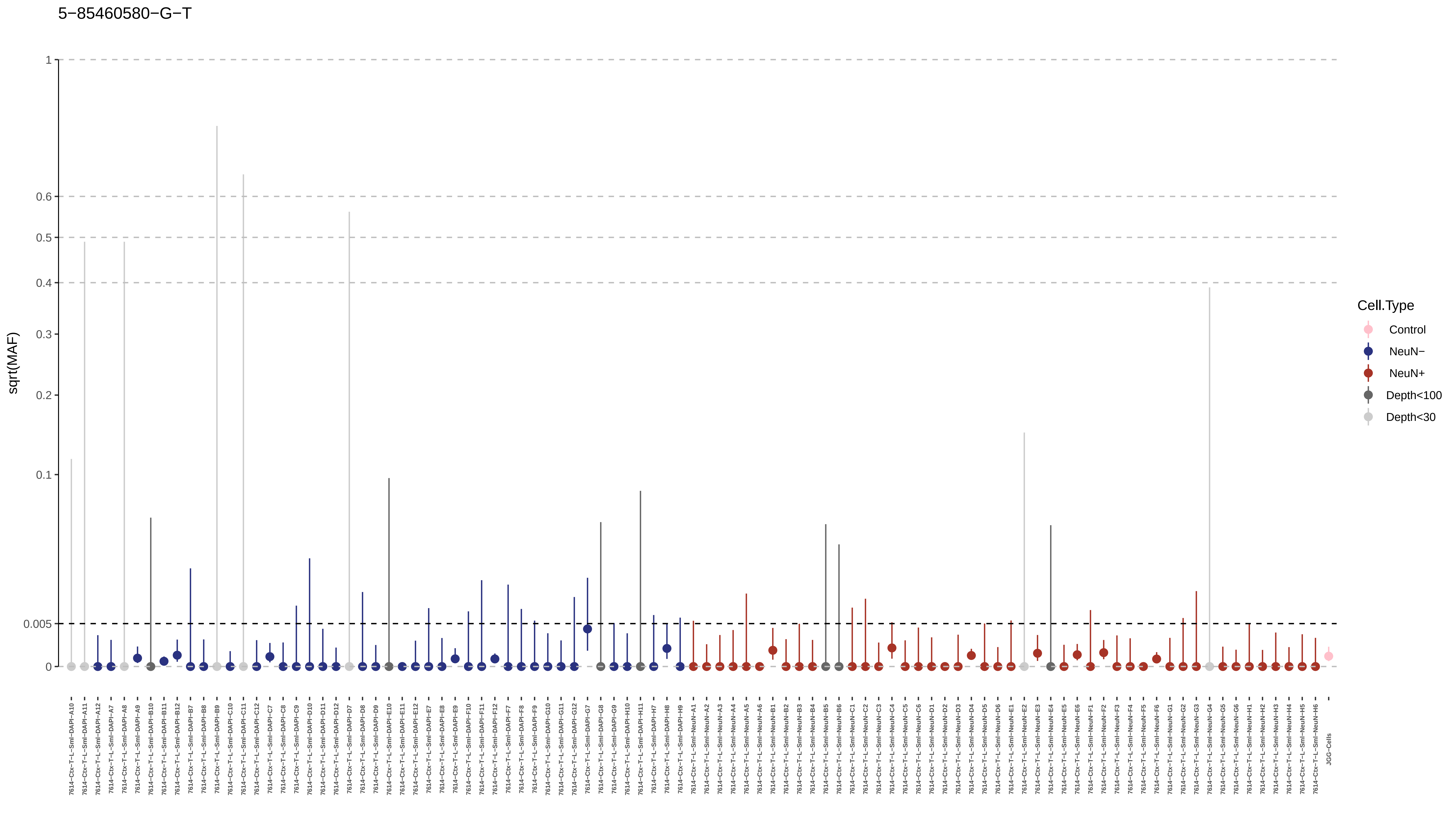

5−85460580−G−T
sqrt(MAF)
1
0.6
0.5
0.4
0.3
0.2
0.1
0.005
0
Cell.Type
Control
NeuN−
NeuN+
Depth<100
Depth<30

6−101898717−G−A

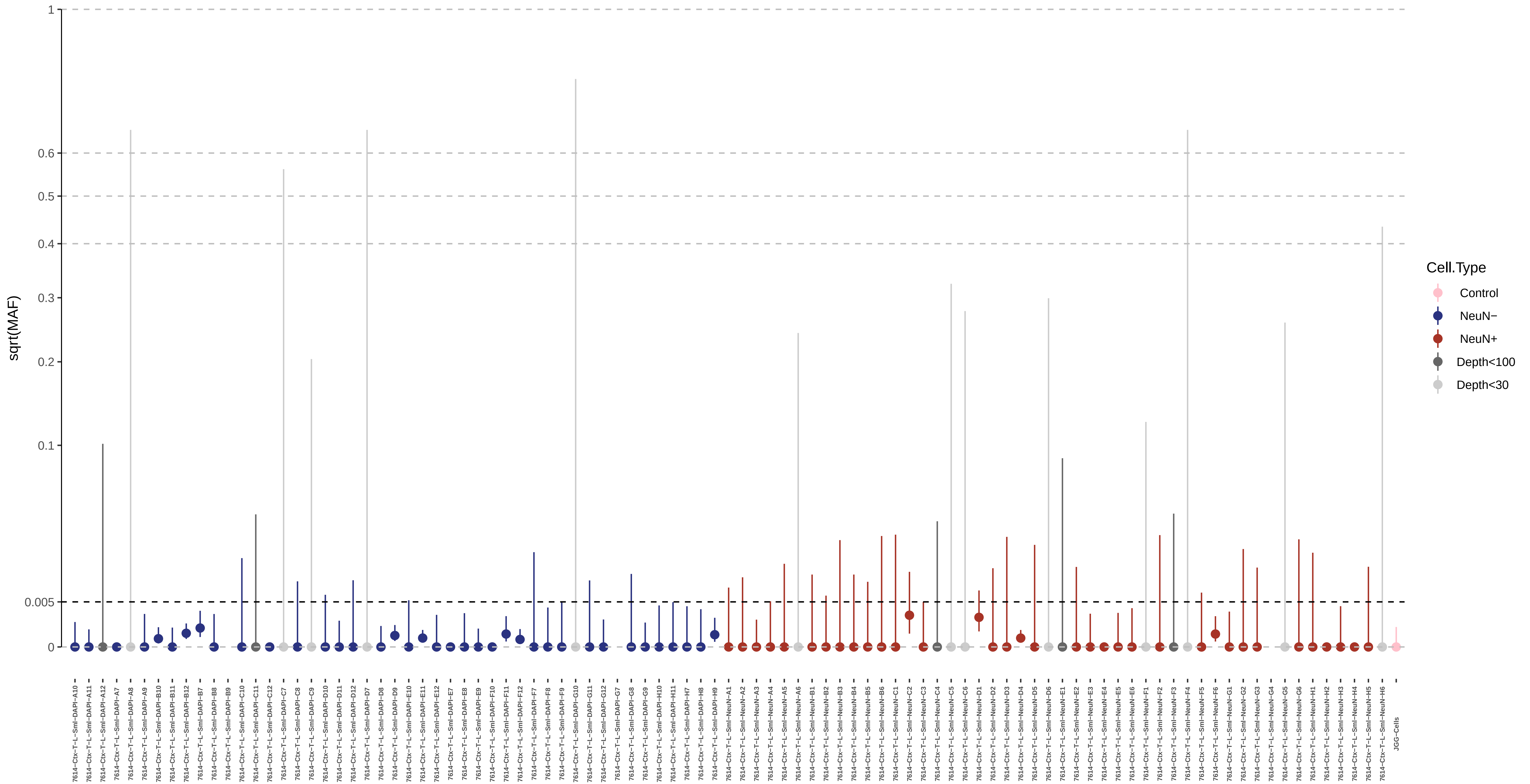

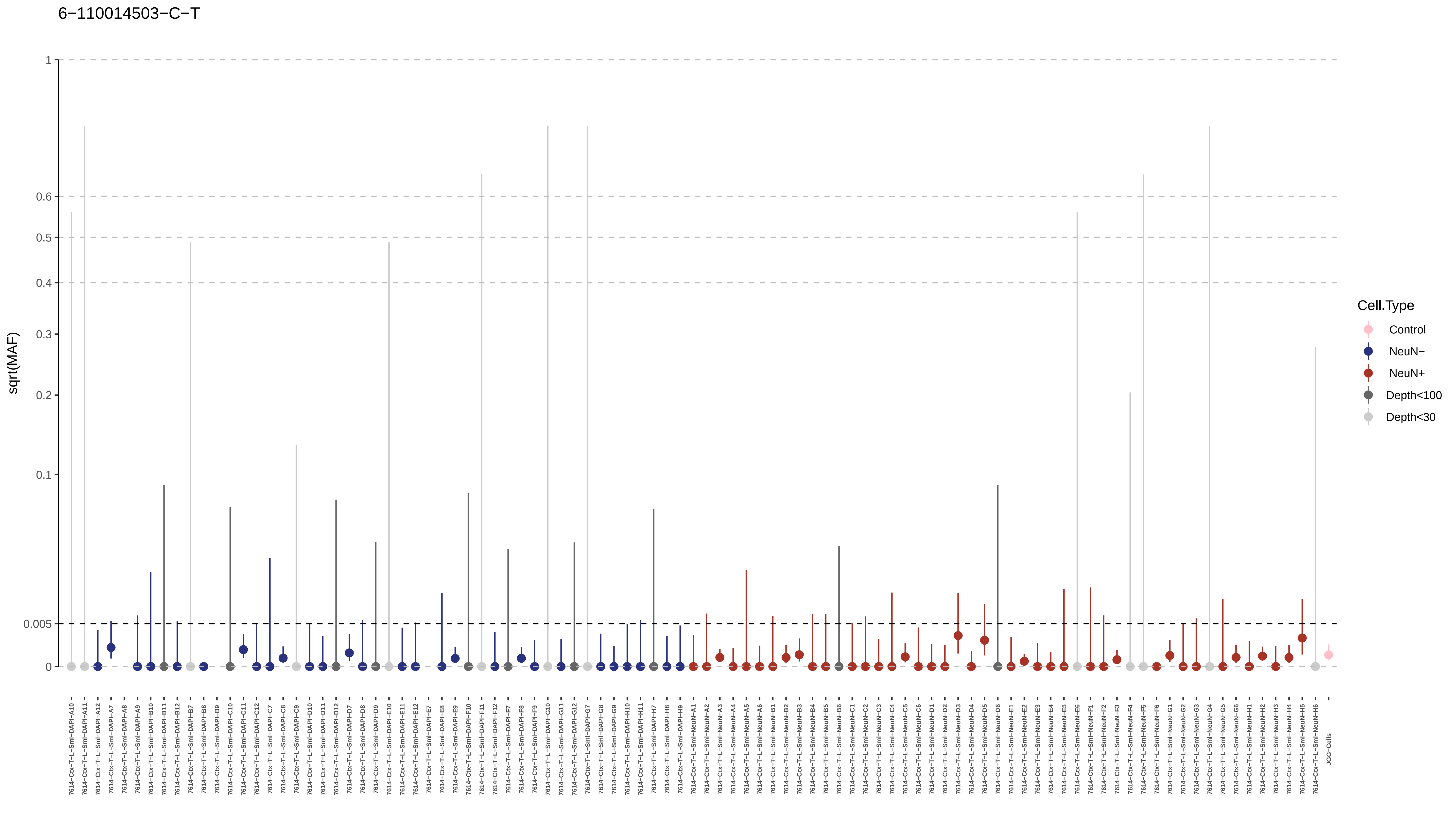
6−110014503−C−T
sqrt(MAF)
1
0.6
0.5
0.4
0.3
0.2
0.1
0.005
0
Cell.Type
Control
NeuN−
NeuN+
Depth<100
Depth<30
JGG-Cells

6−114610552−GAACA−G

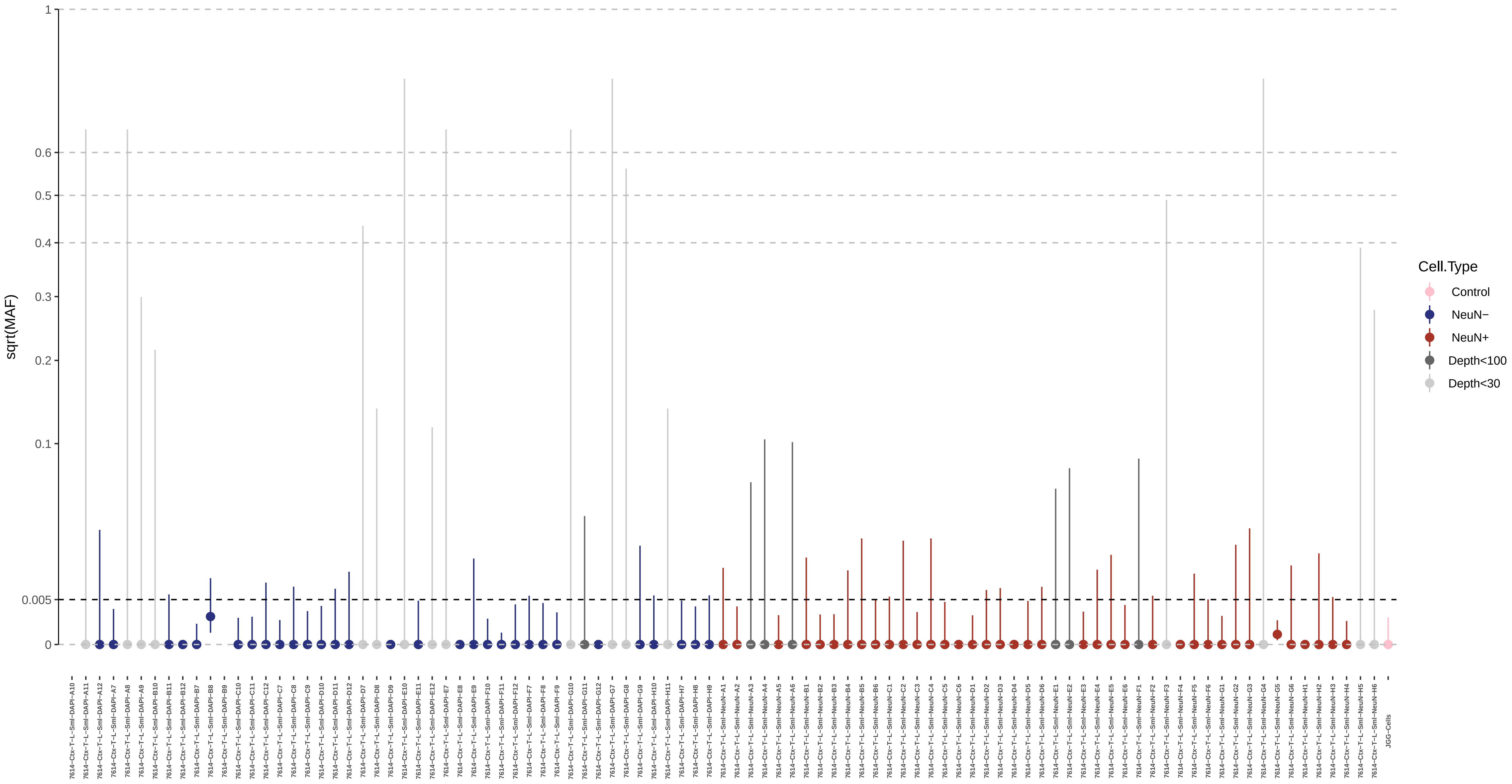

6−131367555−G−A

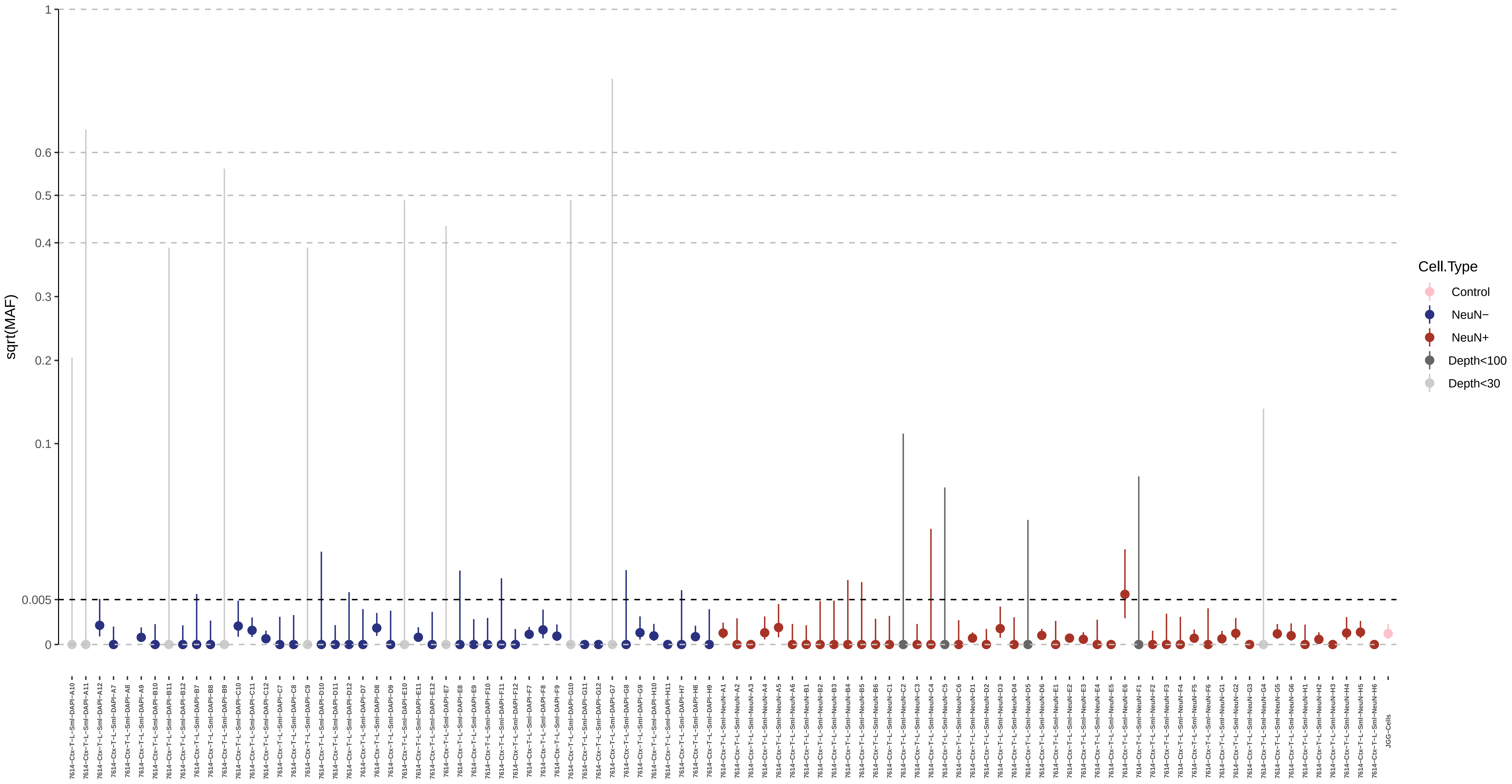

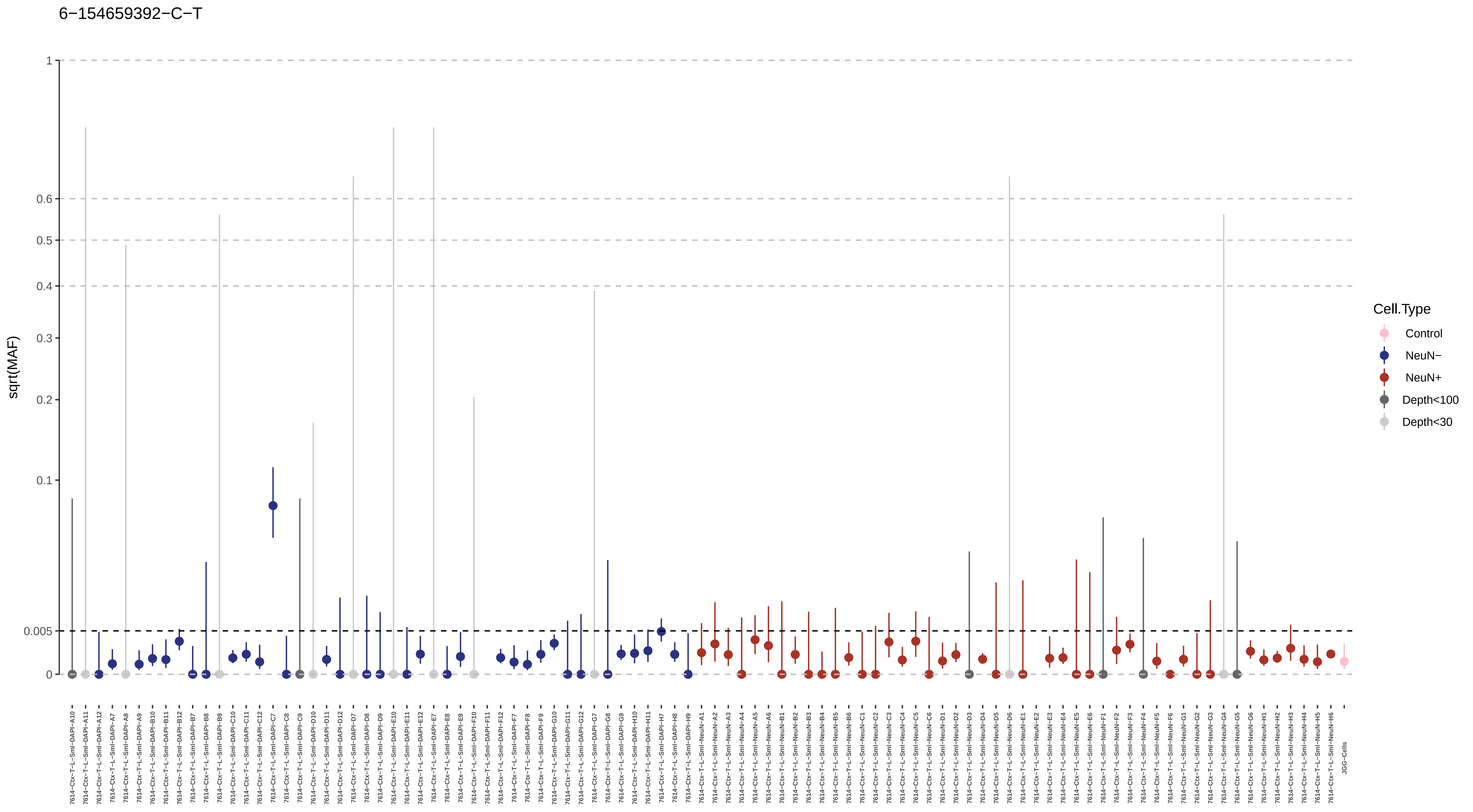
6−154659392−C−T
sqrt(MAF)
1
0.6
0.5
0.4
0.3
0.2
0.1
0.005
0
Cell.Type
Control
NeuN−
NeuN+
Depth<100
Depth<30

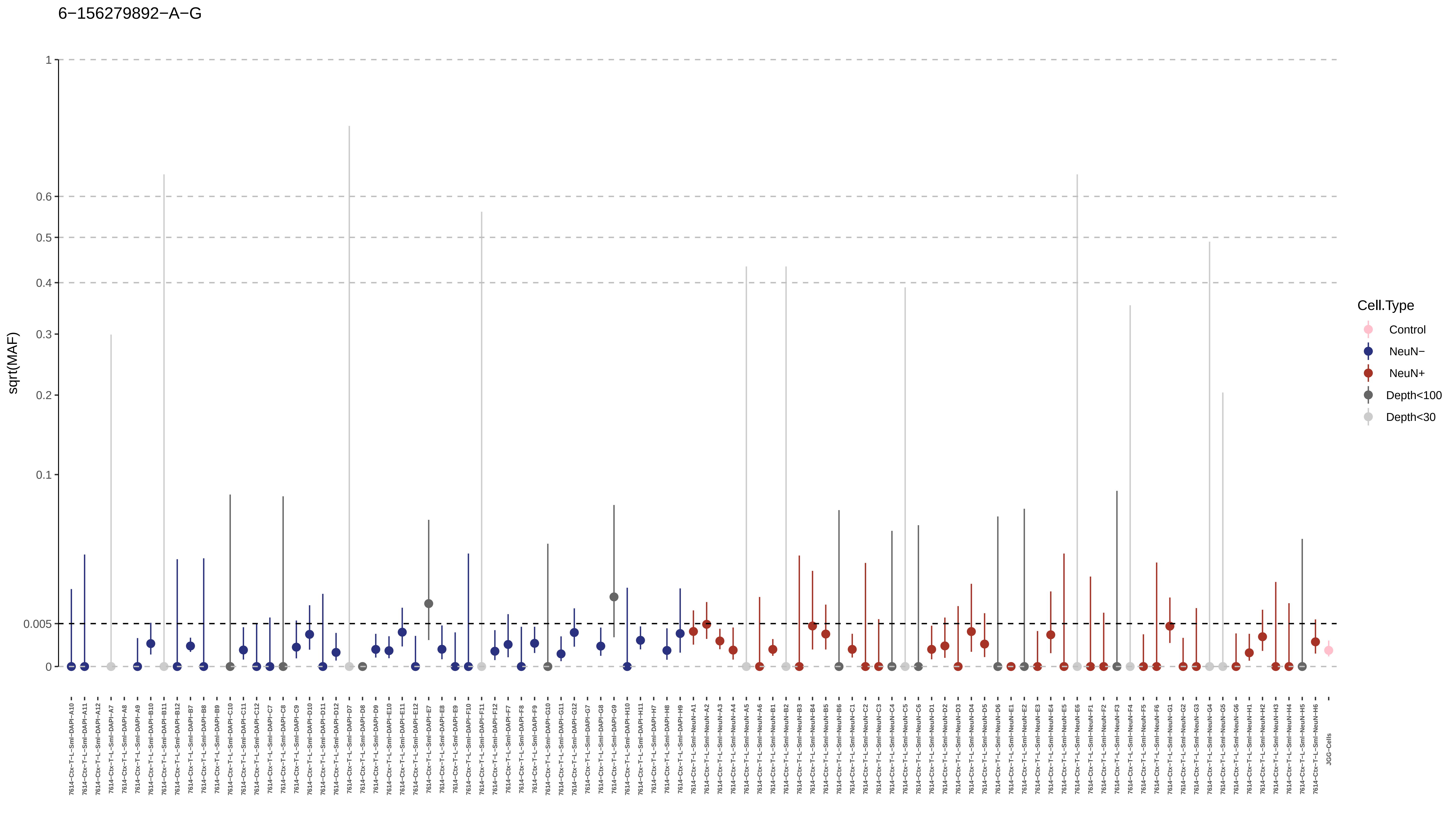

6−156279892−A−G
sqrt(MAF)
1
0.6
0.5
0.4
0.3
0.2
0.1
0.005
0
Cell.Type
Control
NeuN−
NeuN+
Depth<100
Depth<30

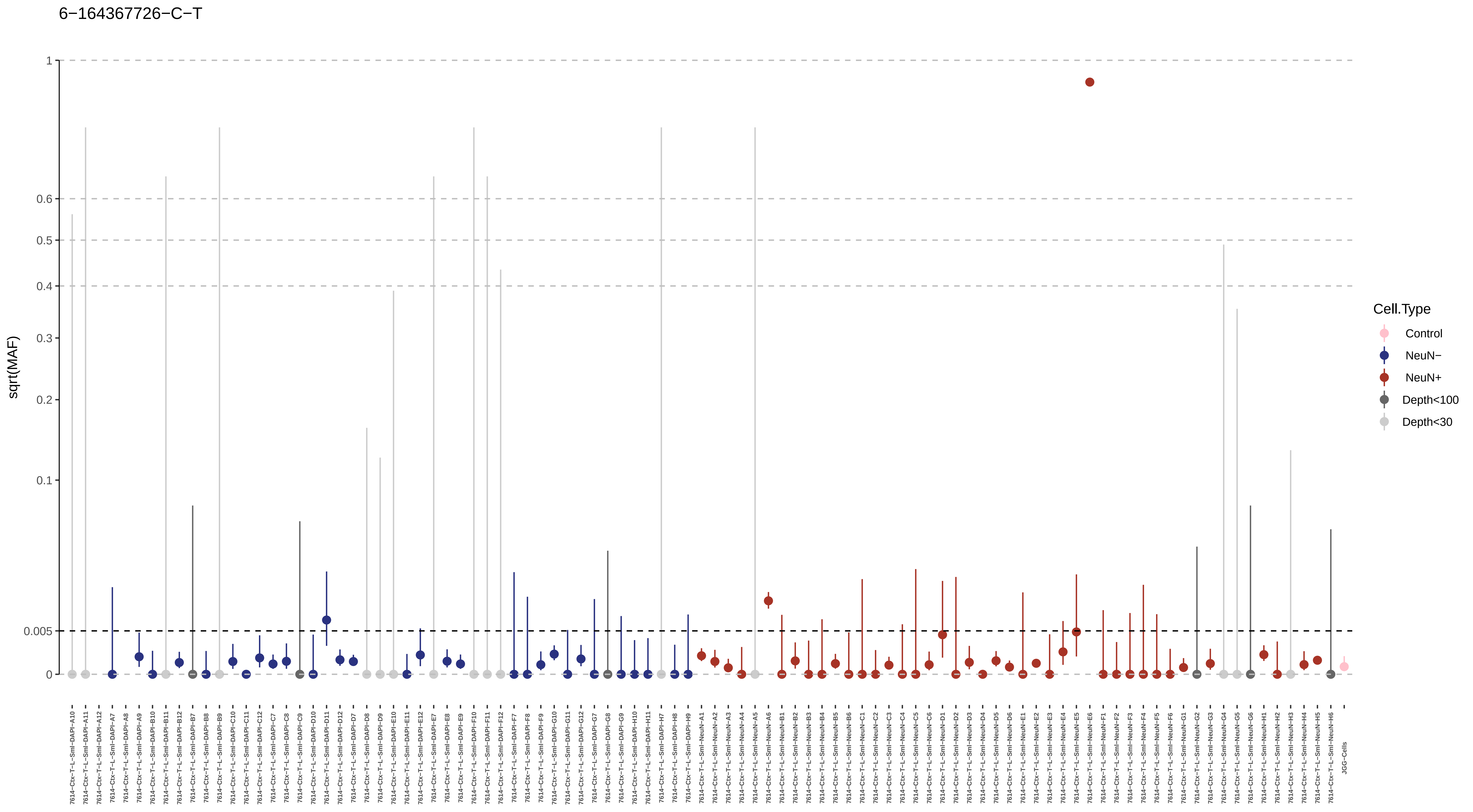

6−164367726−C−T
sqrt(MAF)
1
0.6
0.5
0.4
0.3
0.2
0.1
0.005
0
Cell.Type
Control
NeuN−
NeuN+
Depth<100
Depth<30
JGG-Cells

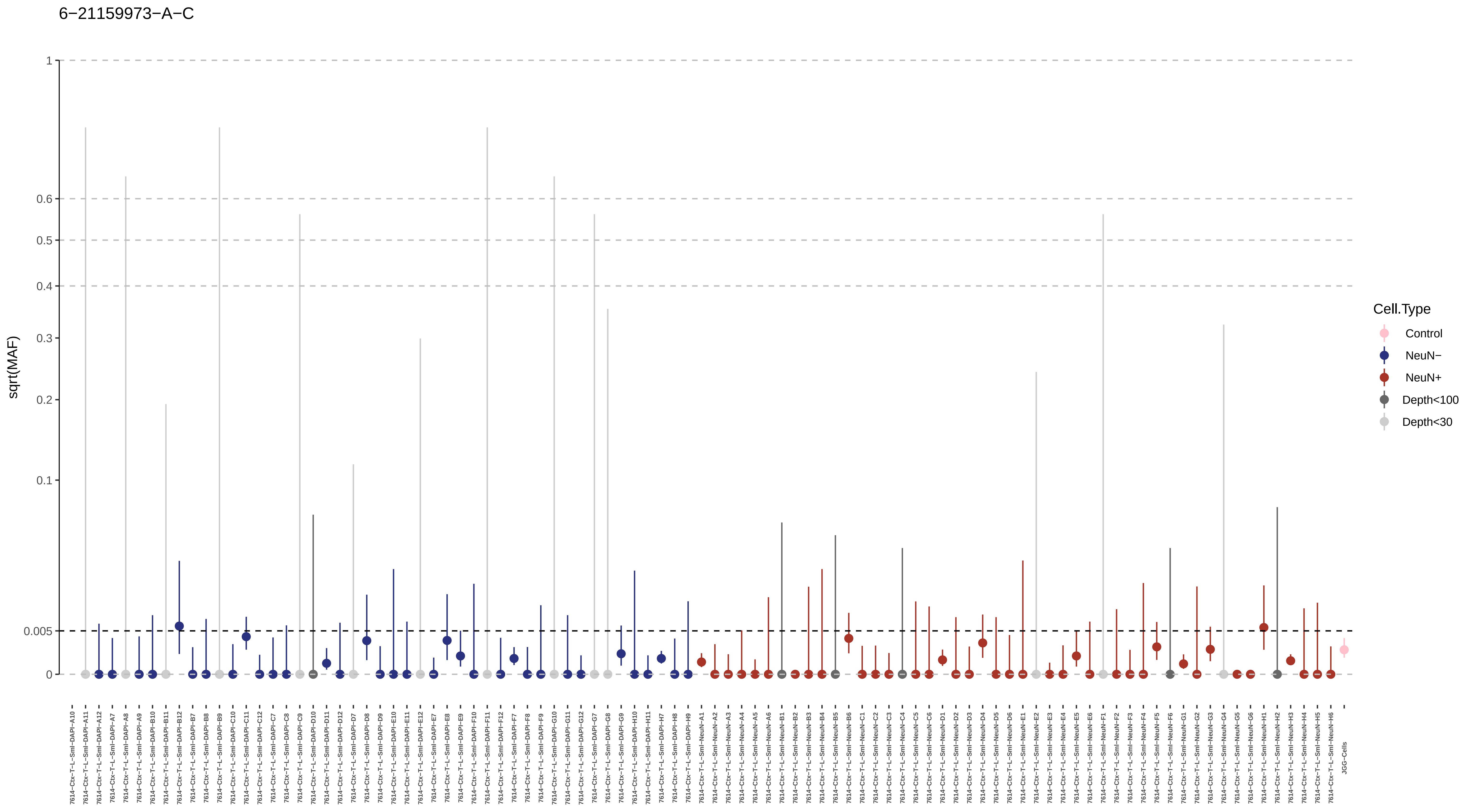
6−21159973−A−C
sqrt(MAF)
1
0.6
0.5
0.4
0.3
0.2
0.1
0.005
0
Cell.Type
Control
NeuN−
NeuN+
Depth<100
Depth<30

6−23944712−G−T

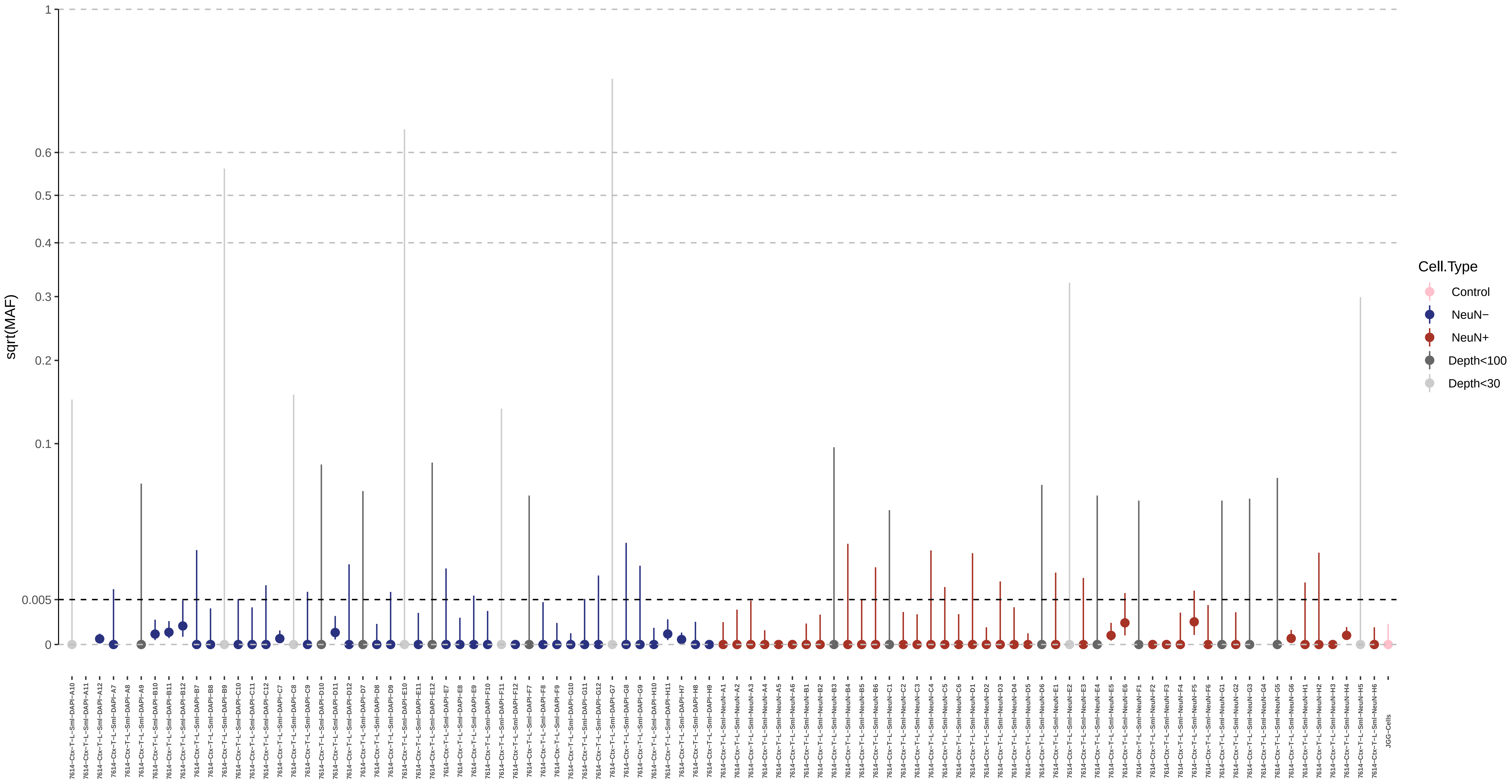

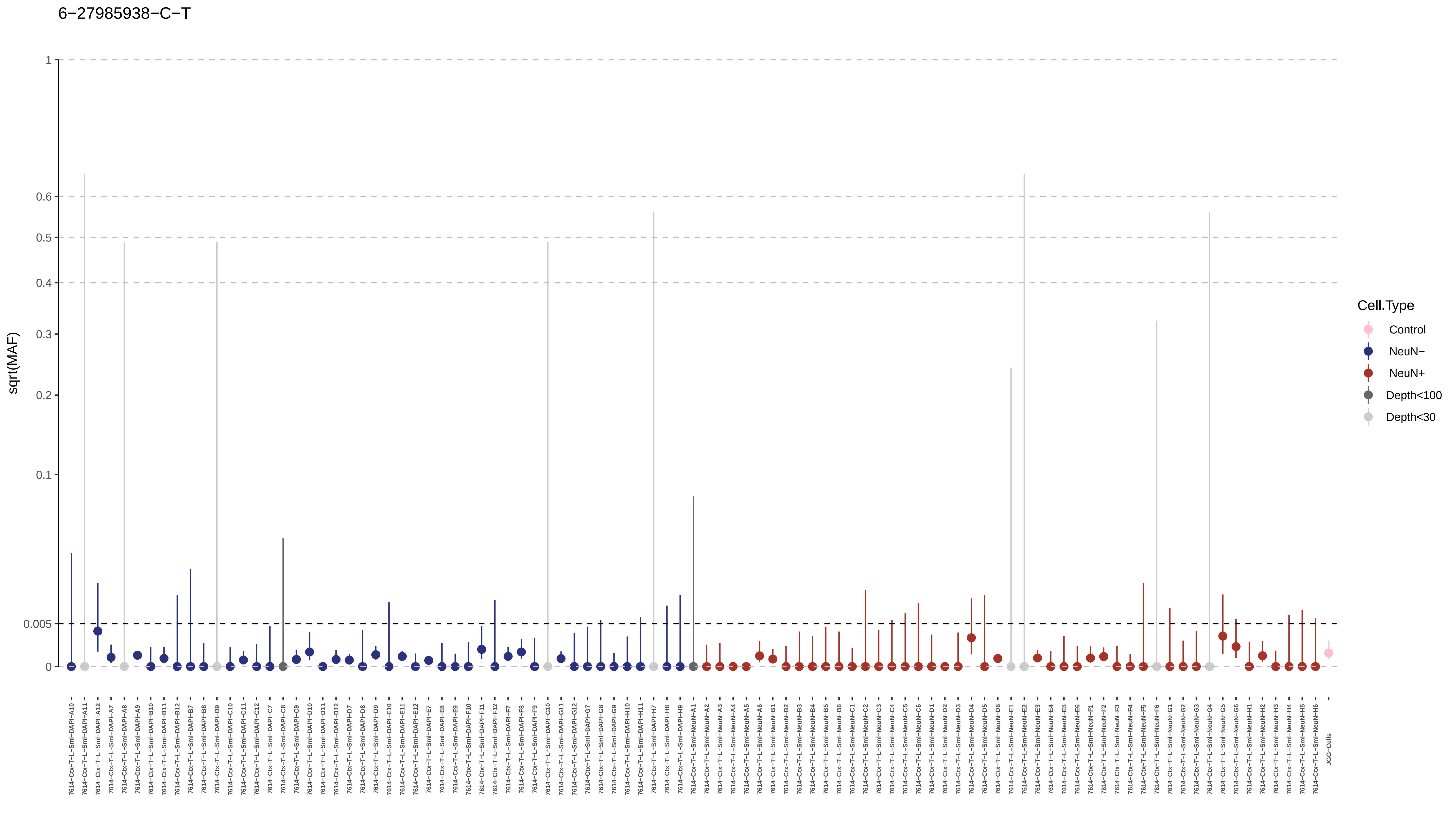
6−27985938−C−T
sqrt(MAF)
1
0.6
0.5
0.4
0.3
0.2
0.1
0.005
0
Cell.Type
Control
NeuN−
NeuN+
Depth<100
Depth<30
7614−Ctx−T−L−Sml−DAPI−A10
7614−Ctx−T−L−Sml−DAPI−A11
7614−Ctx−T−L−Sml−DAPI−A12
7614−Ctx−T−L−Sml−DAPI−A7
7614−Ctx−T−L−Sml−DAPI−A8
7614−Ctx−T−L−Sml−DAPI−A9
7614−Ctx−T−L−Sml−DAPI−B10
7614−Ctx−T−L−Sml−DAPI−B11
7614−Ctx−T−L−Sml−DAPI−B12
7614−Ctx−T−L−Sml−DAPI−B7
7614−Ctx−T−L−Sml−DAPI−B8
7614−Ctx−T−L−Sml−DAPI−B9
7614−Ctx−T−L−Sml−DAPI−C10
7614−Ctx−T−L−Sml−DAPI−C11
7614−Ctx−T−L−Sml−DAPI−C12
7614−Ctx−T−L−Sml−DAPI−C7
7614−Ctx−T−L−Sml−DAPI−C8
7614−Ctx−T−L−Sml−DAPI−C9
7614−Ctx−T−L−Sml−DAPI−D10
7614−Ctx−T−L−Sml−DAPI−D11
7614−Ctx−T−L−Sml−DAPI−D12
7614−Ctx−T−L−Sml−DAPI−D7
7614−Ctx−T−L−Sml−DAPI−D8
7614−Ctx−T−L−Sml−DAPI−D9
7614−Ctx−T−L−Sml−DAPI−E10
7614−Ctx−T−L−Sml−DAPI−E11
7614−Ctx−T−L−Sml−DAPI−E12
7614−Ctx−T−L−Sml−DAPI−E7
7614−Ctx−T−L−Sml−DAPI−E8
7614−Ctx−T−L−Sml−DAPI−E9
7614−Ctx−T−L−Sml−DAPI−F10
7614−Ctx−T−L−Sml−DAPI−F11
7614−Ctx−T−L−Sml−DAPI−F12
7614−Ctx−T−L−Sml−DAPI−F7
7614−Ctx−T−L−Sml−DAPI−F8
7614−Ctx−T−L−Sml−DAPI−F9
7614−Ctx−T−L−Sml−DAPI−G10
7614−Ctx−T−L−Sml−DAPI−G11
7614−Ctx−T−L−Sml−DAPI−G12
7614−Ctx−T−L−Sml−DAPI−G7
7614−Ctx−T−L−Sml−DAPI−G8
7614−Ctx−T−L−Sml−DAPI−G9
7614−Ctx−T−L−Sml−DAPI−H10
7614−Ctx−T−L−Sml−DAPI−H11
7614−Ctx−T−L−Sml−DAPI−H7
7614−Ctx−T−L−Sml−DAPI−H8
7614−Ctx−T−L−Sml−DAPI−H9
7614−Ctx−T−L−Sml−NeuN−A1
7614−Ctx−T−L−Sml−NeuN−A2
7614−Ctx−T−L−Sml−NeuN−A3
7614−Ctx−T−L−Sml−NeuN−A4
7614−Ctx−T−L−Sml−NeuN−A5
7614−Ctx−T−L−Sml−NeuN−A6
7614−Ctx−T−L−Sml−NeuN−B1
7614−Ctx−T−L−Sml−NeuN−B2
7614−Ctx−T−L−Sml−NeuN−B3
7614−Ctx−T−L−Sml−NeuN−B4
7614−Ctx−T−L−Sml−NeuN−B5
7614−Ctx−T−L−Sml−NeuN−B6
7614−Ctx−T−L−Sml−NeuN−C1
7614−Ctx−T−L−Sml−NeuN−C2
7614−Ctx−T−L−Sml−NeuN−C3
7614−Ctx−T−L−Sml−NeuN−C4
7614−Ctx−T−L−Sml−NeuN−C5
7614−Ctx−T−L−Sml−NeuN−C6
7614−Ctx−T−L−Sml−NeuN−D1
7614−Ctx−T−L−Sml−NeuN−D2
7614−Ctx−T−L−Sml−NeuN−D3
7614−Ctx−T−L−Sml−NeuN−D4
7614−Ctx−T−L−Sml−NeuN−D5
7614−Ctx−T−L−Sml−NeuN−D6
7614−Ctx−T−L−Sml−NeuN−E1
7614−Ctx−T−L−Sml−NeuN−E2
7614−Ctx−T−L−Sml−NeuN−E3
7614−Ctx−T−L−Sml−NeuN−E4
7614−Ctx−T−L−Sml−NeuN−E5
7614−Ctx−T−L−Sml−NeuN−E6
7614−Ctx−T−L−Sml−NeuN−F1
7614−Ctx−T−L−Sml−NeuN−F2
7614−Ctx−T−L−Sml−NeuN−F3
7614−Ctx−T−L−Sml−NeuN−F4
7614−Ctx−T−L−Sml−NeuN−F5
7614−Ctx−T−L−Sml−NeuN−F6
7614−Ctx−T−L−Sml−NeuN−G1
7614−Ctx−T−L−Sml−NeuN−G2
7614−Ctx−T−L−Sml−NeuN−G3
7614−Ctx−T−L−Sml−NeuN−G4
7614−Ctx−T−L−Sml−NeuN−G5
7614−Ctx−T−L−Sml−NeuN−G6
7614−Ctx−T−L−Sml−NeuN−H1
7614−Ctx−T−L−Sml−NeuN−H2
7614−Ctx−T−L−Sml−NeuN−H3
7614−Ctx−T−L−Sml−NeuN−H4
7614−Ctx−T−L−Sml−NeuN−H5
7614−Ctx−T−L−Sml−NeuN−H6
JGG-Cells

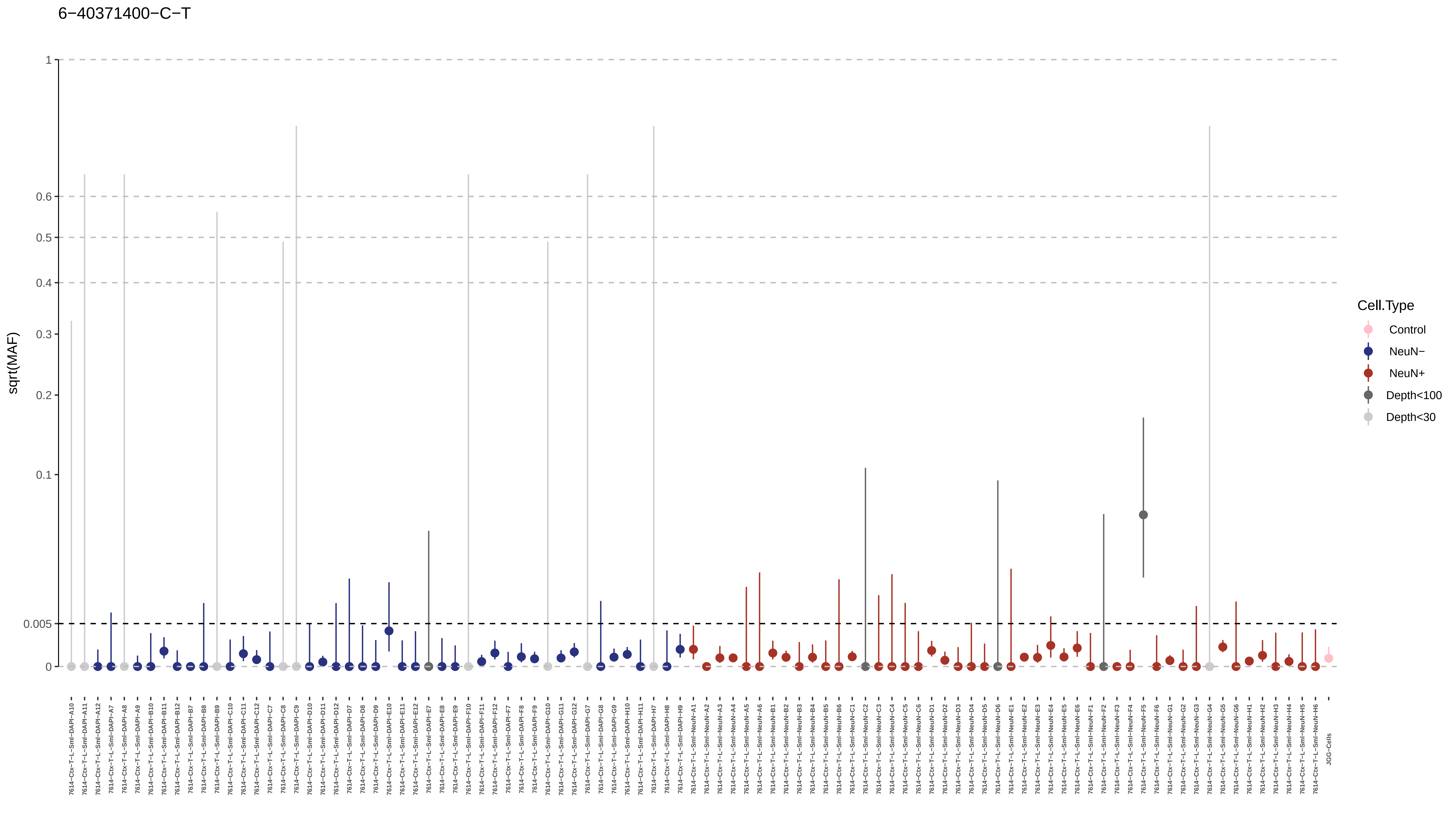

6−40371400−C−T
sqrt(MAF)
1
0.6
0.5
0.4
0.3
0.2
0.1
0.005
0
Cell.Type
Control
NeuN−
NeuN+
Depth<100
Depth<30

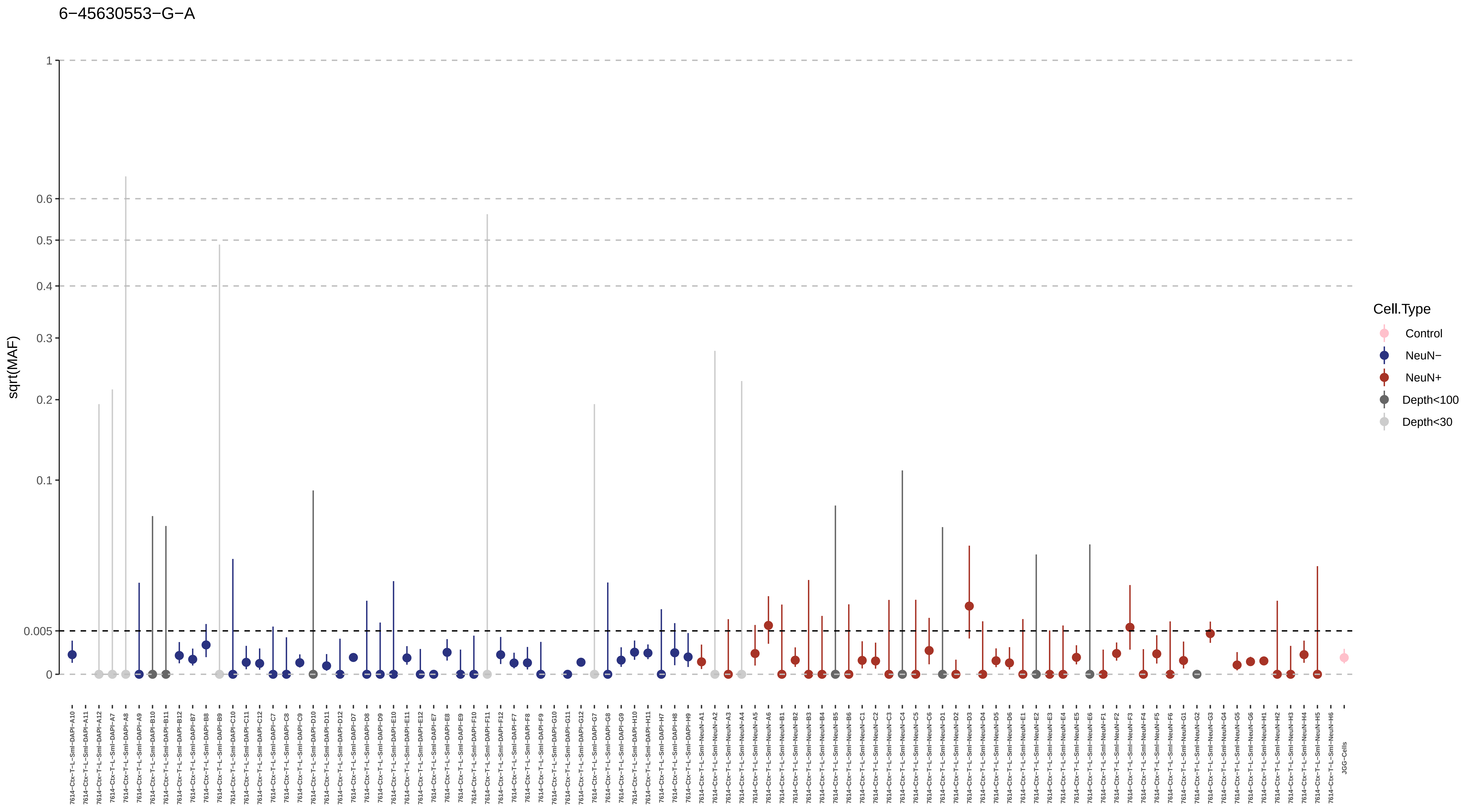
6−45630553−G−A
sqrt(MAF)
1
0.6
0.5
0.4
0.3
0.2
0.1
0.005
0
Cell.Type
Control
NeuN−
NeuN+
Depth<100
Depth<30
JGG-Cells

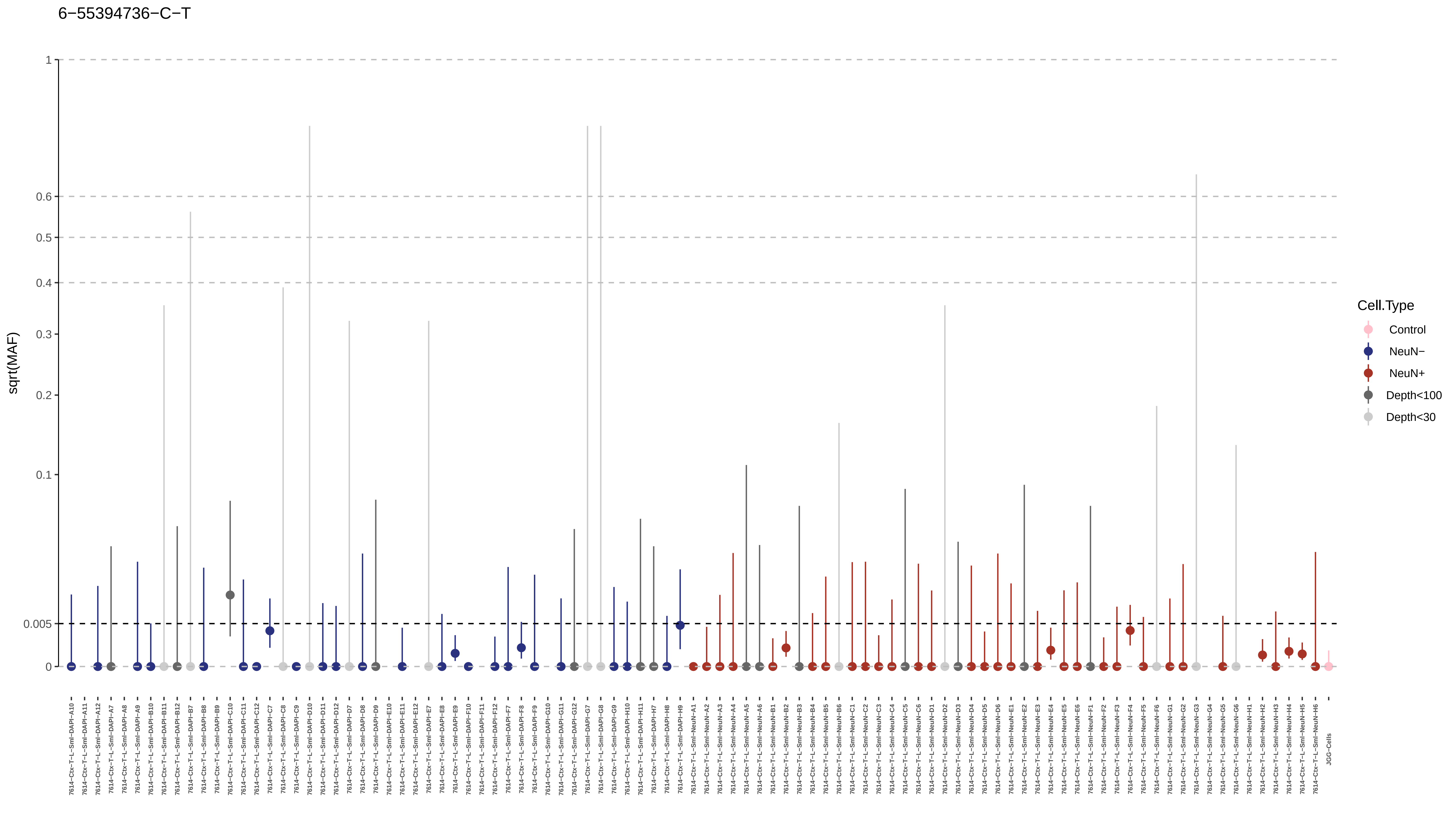
6−55394736−C−T
sqrt(MAF)
1
0.6
0.5
0.4
0.3
0.2
0.1
0.005
0
Cell.Type
Control
NeuN−
NeuN+
Depth<100
Depth<30
JGG-Cells

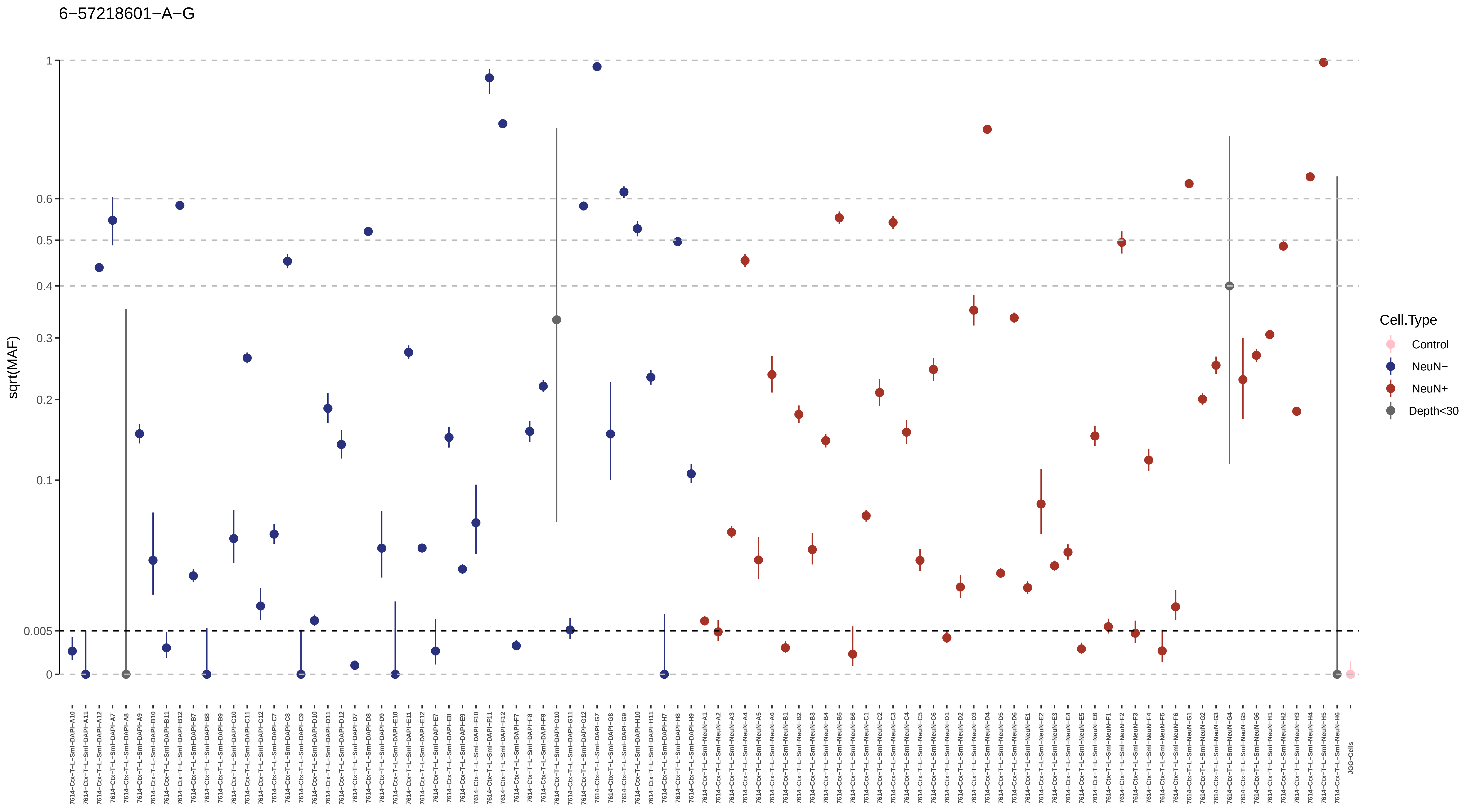
6−57218601−A−G
sqrt(MAF)
1
0.6
0.5
0.4
0.3
0.2
0.1
0.005
0
Cell.Type
Control
NeuN−
NeuN+
Depth<30
7614−Ctx−T−L−Sml−DAPI−A10
7614−Ctx−T−L−Sml−DAPI−A11
7614−Ctx−T−L−Sml−DAPI−A12
7614−Ctx−T−L−Sml−DAPI−A7
7614−Ctx−T−L−Sml−DAPI−A8
7614−Ctx−T−L−Sml−DAPI−A9
7614−Ctx−T−L−Sml−DAPI−B10
7614−Ctx−T−L−Sml−DAPI−B11
7614−Ctx−T−L−Sml−DAPI−B12
7614−Ctx−T−L−Sml−DAPI−B7
7614−Ctx−T−L−Sml−DAPI−B8
7614−Ctx−T−L−Sml−DAPI−B9
7614−Ctx−T−L−Sml−DAPI−C10
7614−Ctx−T−L−Sml−DAPI−C11
7614−Ctx−T−L−Sml−DAPI−C12
7614−Ctx−T−L−Sml−DAPI−C7
7614−Ctx−T−L−Sml−DAPI−C8
7614−Ctx−T−L−Sml−DAPI−C9
7614−Ctx−T−L−Sml−DAPI−D10
7614−Ctx−T−L−Sml−DAPI−D11
7614−Ctx−T−L−Sml−DAPI−D12
7614−Ctx−T−L−Sml−DAPI−D7
7614−Ctx−T−L−Sml−DAPI−D8
7614−Ctx−T−L−Sml−DAPI−D9
7614−Ctx−T−L−Sml−DAPI−E10
7614−Ctx−T−L−Sml−DAPI−E11
7614−Ctx−T−L−Sml−DAPI−E12
7614−Ctx−T−L−Sml−DAPI−E7
7614−Ctx−T−L−Sml−DAPI−E8
7614−Ctx−T−L−Sml−DAPI−E9
7614−Ctx−T−L−Sml−DAPI−F10
7614−Ctx−T−L−Sml−DAPI−F11
7614−Ctx−T−L−Sml−DAPI−F12
7614−Ctx−T−L−Sml−DAPI−F7
7614−Ctx−T−L−Sml−DAPI−F8
7614−Ctx−T−L−Sml−DAPI−F9
7614−Ctx−T−L−Sml−DAPI−G10
7614−Ctx−T−L−Sml−DAPI−G11
7614−Ctx−T−L−Sml−DAPI−G12
7614−Ctx−T−L−Sml−DAPI−G7
7614−Ctx−T−L−Sml−DAPI−G8
7614−Ctx−T−L−Sml−DAPI−G9
7614−Ctx−T−L−Sml−DAPI−H10
7614−Ctx−T−L−Sml−DAPI−H11
7614−Ctx−T−L−Sml−DAPI−H7
7614−Ctx−T−L−Sml−DAPI−H8
7614−Ctx−T−L−Sml−DAPI−H9
7614−Ctx−T−L−Sml−NeuN−A1
7614−Ctx−T−L−Sml−NeuN−A2
7614−Ctx−T−L−Sml−NeuN−A3
7614−Ctx−T−L−Sml−NeuN−A4
7614−Ctx−T−L−Sml−NeuN−A5
7614−Ctx−T−L−Sml−NeuN−A6
7614−Ctx−T−L−Sml−NeuN−B1
7614−Ctx−T−L−Sml−NeuN−B2
7614−Ctx−T−L−Sml−NeuN−B3
7614−Ctx−T−L−Sml−NeuN−B4
7614−Ctx−T−L−Sml−NeuN−B5
7614−Ctx−T−L−Sml−NeuN−B6
7614−Ctx−T−L−Sml−NeuN−C1
7614−Ctx−T−L−Sml−NeuN−C2
7614−Ctx−T−L−Sml−NeuN−C3
7614−Ctx−T−L−Sml−NeuN−C4
7614−Ctx−T−L−Sml−NeuN−C5
7614−Ctx−T−L−Sml−NeuN−C6
7614−Ctx−T−L−Sml−NeuN−D1
7614−Ctx−T−L−Sml−NeuN−D2
7614−Ctx−T−L−Sml−NeuN−D3
7614−Ctx−T−L−Sml−NeuN−D4
7614−Ctx−T−L−Sml−NeuN−D5
7614−Ctx−T−L−Sml−NeuN−D6
7614−Ctx−T−L−Sml−NeuN−E1
7614−Ctx−T−L−Sml−NeuN−E2
7614−Ctx−T−L−Sml−NeuN−E3
7614−Ctx−T−L−Sml−NeuN−E4
7614−Ctx−T−L−Sml−NeuN−E5
7614−Ctx−T−L−Sml−NeuN−E6
7614−Ctx−T−L−Sml−NeuN−F1
7614−Ctx−T−L−Sml−NeuN−F2
7614−Ctx−T−L−Sml−NeuN−F3
7614−Ctx−T−L−Sml−NeuN−F4
7614−Ctx−T−L−Sml−NeuN−F5
7614−Ctx−T−L−Sml−NeuN−F6
7614−Ctx−T−L−Sml−NeuN−G1
7614−Ctx−T−L−Sml−NeuN−G2
7614−Ctx−T−L−Sml−NeuN−G3
7614−Ctx−T−L−Sml−NeuN−G4
7614−Ctx−T−L−Sml−NeuN−G5
7614−Ctx−T−L−Sml−NeuN−G6
7614−Ctx−T−L−Sml−NeuN−H1
7614−Ctx−T−L−Sml−NeuN−H2
7614−Ctx−T−L−Sml−NeuN−H3
7614−Ctx−T−L−Sml−NeuN−H4
7614−Ctx−T−L−Sml−NeuN−H5
7614−Ctx−T−L−Sml−NeuN−H6
JGG−Cells

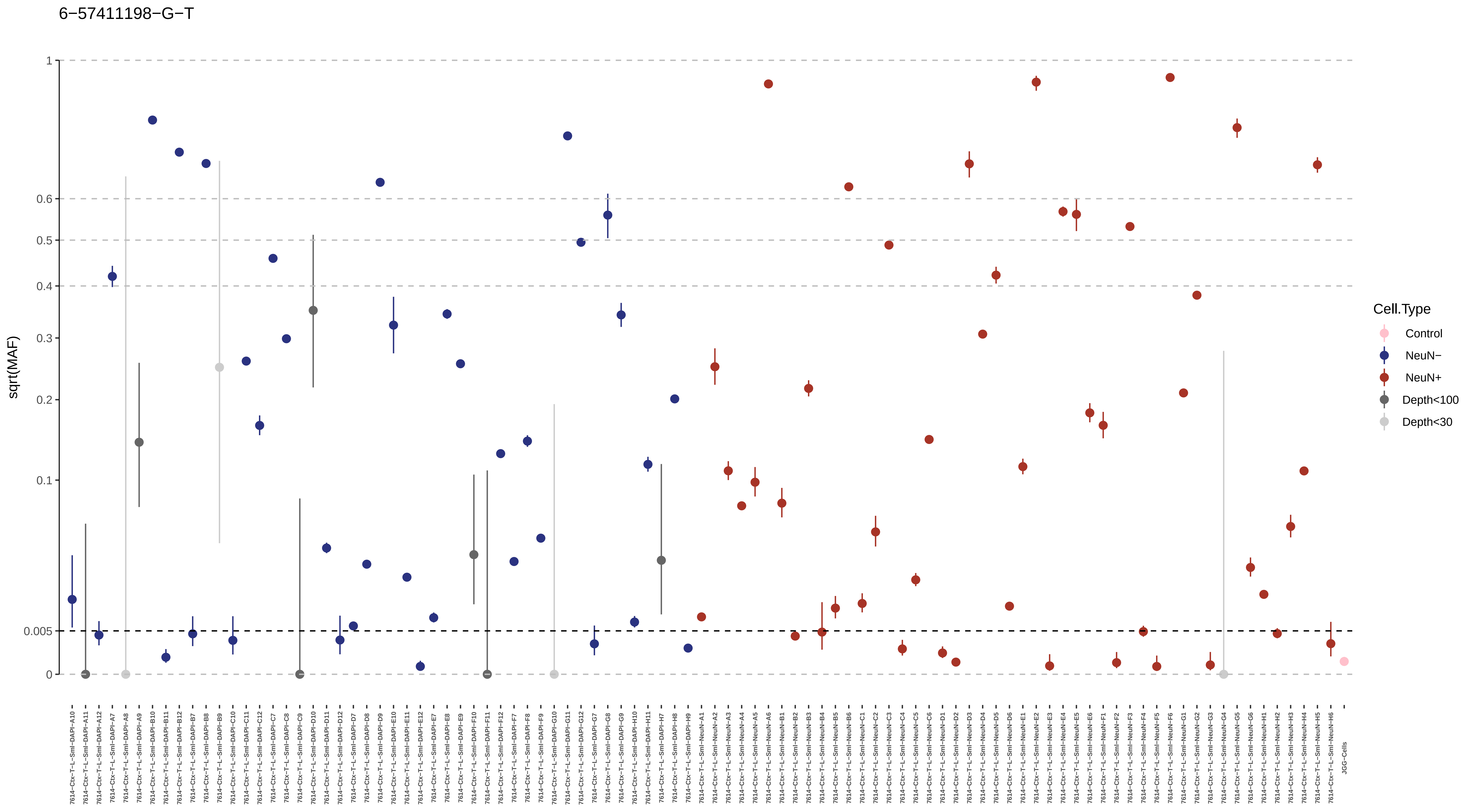
6−57411198−G−T
sqrt(MAF)
1
0.6
0.5
0.4
0.3
0.2
0.1
0.005
0
Cell.Type
Control
NeuN−
NeuN+
Depth<100
Depth<30
JGG-Cells

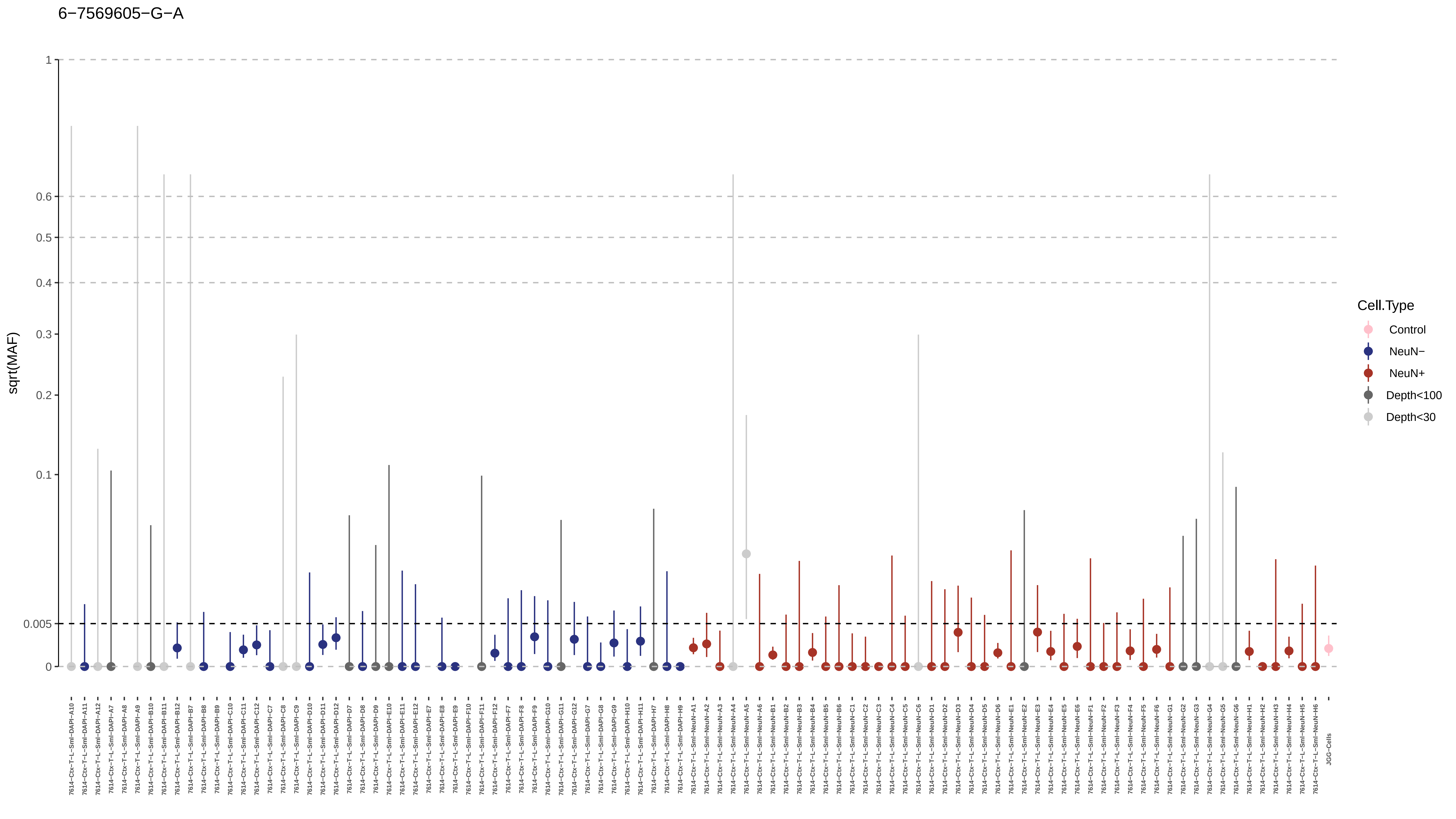
6−7569605−G−A
sqrt(MAF)
1
0.6
0.5
0.4
0.3
0.2
0.1
0.005
0
7614−Ctx−T−L−Sml−DAPI−A10
7614−Ctx−T−L−Sml−DAPI−A11
7614−Ctx−T−L−Sml−DAPI−A12
7614−Ctx−T−L−Sml−DAPI−A7
7614−Ctx−T−L−Sml−DAPI−A8
7614−Ctx−T−L−Sml−DAPI−A9
7614−Ctx−T−L−Sml−DAPI−B10
7614−Ctx−T−L−Sml−DAPI−B11
7614−Ctx−T−L−Sml−DAPI−B12
7614−Ctx−T−L−Sml−DAPI−B7
7614−Ctx−T−L−Sml−DAPI−B8
7614−Ctx−T−L−Sml−DAPI−B9
7614−Ctx−T−L−Sml−DAPI−C10
7614−Ctx−T−L−Sml−DAPI−C11
7614−Ctx−T−L−Sml−DAPI−C12
7614−Ctx−T−L−Sml−DAPI−C7
7614−Ctx−T−L−Sml−DAPI−C8
7614−Ctx−T−L−Sml−DAPI−C9
7614−Ctx−T−L−Sml−DAPI−D10
7614−Ctx−T−L−Sml−DAPI−D11
7614−Ctx−T−L−Sml−DAPI−D12
7614−Ctx−T−L−Sml−DAPI−D7
7614−Ctx−T−L−Sml−DAPI−D8
7614−Ctx−T−L−Sml−DAPI−D9
7614−Ctx−T−L−Sml−DAPI−E10
7614−Ctx−T−L−Sml−DAPI−E11
7614−Ctx−T−L−Sml−DAPI−E12
7614−Ctx−T−L−Sml−DAPI−E7
7614−Ctx−T−L−Sml−DAPI−E8
7614−Ctx−T−L−Sml−DAPI−E9
7614−Ctx−T−L−Sml−DAPI−F10
7614−Ctx−T−L−Sml−DAPI−F11
7614−Ctx−T−L−Sml−DAPI−F12
7614−Ctx−T−L−Sml−DAPI−F7
7614−Ctx−T−L−Sml−DAPI−F8
7614−Ctx−T−L−Sml−DAPI−F9
7614−Ctx−T−L−Sml−DAPI−G10
7614−Ctx−T−L−Sml−DAPI−G11
7614−Ctx−T−L−Sml−DAPI−G12
7614−Ctx−T−L−Sml−DAPI−G7
7614−Ctx−T−L−Sml−DAPI−G8
7614−Ctx−T−L−Sml−DAPI−G9
7614−Ctx−T−L−Sml−DAPI−H10
7614−Ctx−T−L−Sml−DAPI−H11
7614−Ctx−T−L−Sml−DAPI−H7
7614−Ctx−T−L−Sml−DAPI−H8
7614−Ctx−T−L−Sml−DAPI−H9
7614−Ctx−T−L−Sml−NeuN−A1
7614−Ctx−T−L−Sml−NeuN−A2
7614−Ctx−T−L−Sml−NeuN−A3
7614−Ctx−T−L−Sml−NeuN−A4
7614−Ctx−T−L−Sml−NeuN−A5
7614−Ctx−T−L−Sml−NeuN−A6
7614−Ctx−T−L−Sml−NeuN−B1
7614−Ctx−T−L−Sml−NeuN−B2
7614−Ctx−T−L−Sml−NeuN−B3
7614−Ctx−T−L−Sml−NeuN−B4
7614−Ctx−T−L−Sml−NeuN−B5
7614−Ctx−T−L−Sml−NeuN−B6
7614−Ctx−T−L−Sml−NeuN−C1
7614−Ctx−T−L−Sml−NeuN−C2
7614−Ctx−T−L−Sml−NeuN−C3
7614−Ctx−T−L−Sml−NeuN−C4
7614−Ctx−T−L−Sml−NeuN−C5
7614−Ctx−T−L−Sml−NeuN−C6
7614−Ctx−T−L−Sml−NeuN−D1
7614−Ctx−T−L−Sml−NeuN−D2
7614−Ctx−T−L−Sml−NeuN−D3
7614−Ctx−T−L−Sml−NeuN−D4
7614−Ctx−T−L−Sml−NeuN−D5
7614−Ctx−T−L−Sml−NeuN−D6
7614−Ctx−T−L−Sml−NeuN−E1
7614−Ctx−T−L−Sml−NeuN−E2
7614−Ctx−T−L−Sml−NeuN−E3
7614−Ctx−T−L−Sml−NeuN−E4
7614−Ctx−T−L−Sml−NeuN−E5
7614−Ctx−T−L−Sml−NeuN−E6
7614−Ctx−T−L−Sml−NeuN−F1
7614−Ctx−T−L−Sml−NeuN−F2
7614−Ctx−T−L−Sml−NeuN−F3
7614−Ctx−T−L−Sml−NeuN−F4
7614−Ctx−T−L−Sml−NeuN−F5
7614−Ctx−T−L−Sml−NeuN−F6
7614−Ctx−T−L−Sml−NeuN−G1
7614−Ctx−T−L−Sml−NeuN−G2
7614−Ctx−T−L−Sml−NeuN−G3
7614−Ctx−T−L−Sml−NeuN−G4
7614−Ctx−T−L−Sml−NeuN−G5
7614−Ctx−T−L−Sml−NeuN−G6
7614−Ctx−T−L−Sml−NeuN−H1
7614−Ctx−T−L−Sml−NeuN−H2
7614−Ctx−T−L−Sml−NeuN−H3
7614−Ctx−T−L−Sml−NeuN−H4
7614−Ctx−T−L−Sml−NeuN−H5
7614−Ctx−T−L−Sml−NeuN−H6
JGG−Cells
Cell.Type
Control
NeuN−
NeuN+
Depth<100
Depth<30

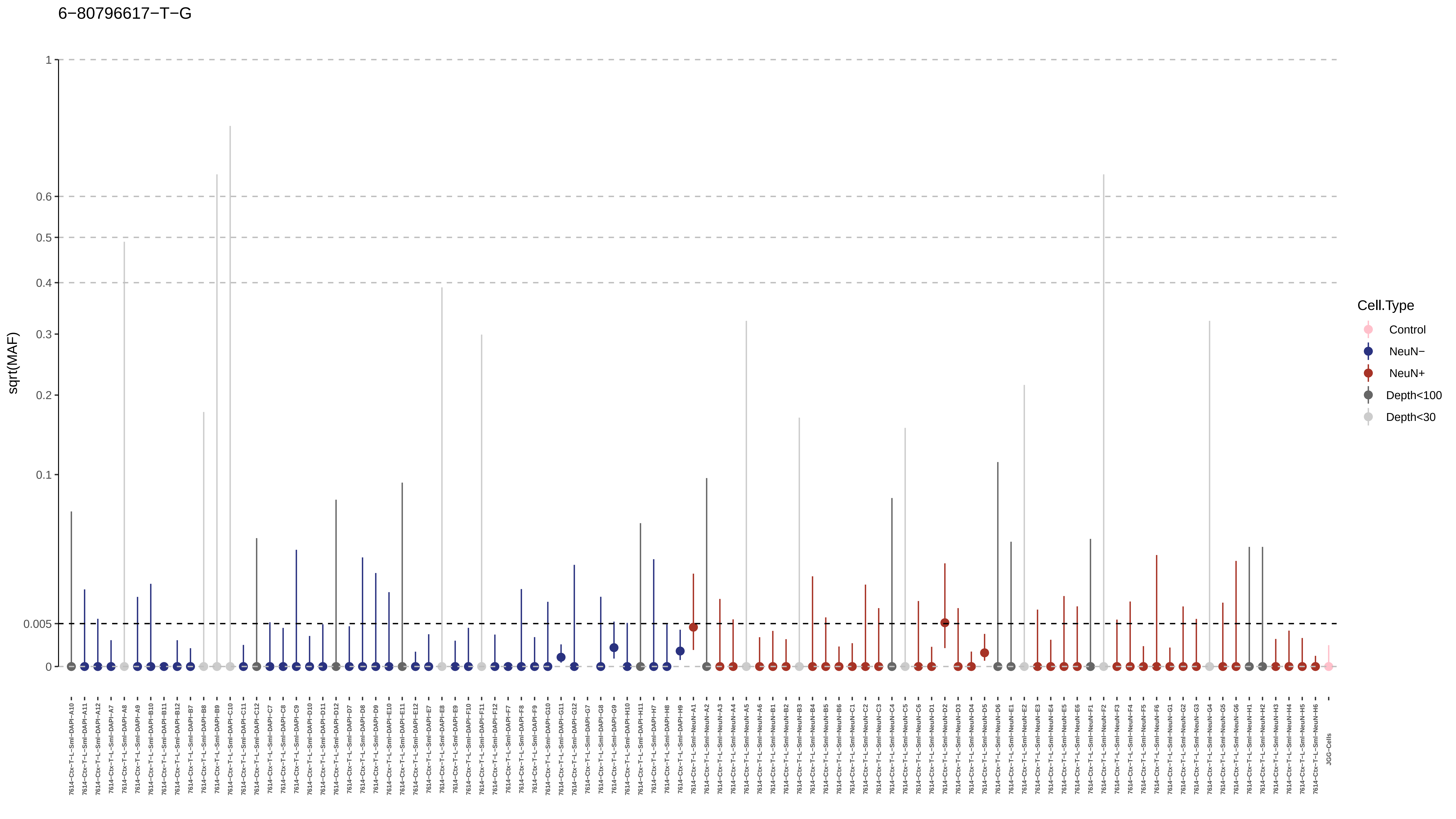
6−80796617−T−G
sqrt(MAF)
1
0.6
0.5
0.4
0.3
0.2
0.1
0.005
0
Cell.Type
Control
NeuN−
NeuN+
Depth<100
Depth<30
7614−Ctx−T−L−Sml−DAPI−A10
7614−Ctx−T−L−Sml−DAPI−A11
7614−Ctx−T−L−Sml−DAPI−A12
7614−Ctx−T−L−Sml−DAPI−A7
7614−Ctx−T−L−Sml−DAPI−A8
7614−Ctx−T−L−Sml−DAPI−A9
7614−Ctx−T−L−Sml−DAPI−B10
7614−Ctx−T−L−Sml−DAPI−B11
7614−Ctx−T−L−Sml−DAPI−B12
7614−Ctx−T−L−Sml−DAPI−B7
7614−Ctx−T−L−Sml−DAPI−B8
7614−Ctx−T−L−Sml−DAPI−B9
7614−Ctx−T−L−Sml−DAPI−C10
7614−Ctx−T−L−Sml−DAPI−C11
7614−Ctx−T−L−Sml−DAPI−C12
7614−Ctx−T−L−Sml−DAPI−C7
7614−Ctx−T−L−Sml−DAPI−C8
7614−Ctx−T−L−Sml−DAPI−C9
7614−Ctx−T−L−Sml−DAPI−D10
7614−Ctx−T−L−Sml−DAPI−D11
7614−Ctx−T−L−Sml−DAPI−D12
7614−Ctx−T−L−Sml−DAPI−D7
7614−Ctx−T−L−Sml−DAPI−D8
7614−Ctx−T−L−Sml−DAPI−D9
7614−Ctx−T−L−Sml−DAPI−E10
7614−Ctx−T−L−Sml−DAPI−E11
7614−Ctx−T−L−Sml−DAPI−E12
7614−Ctx−T−L−Sml−DAPI−E7
7614−Ctx−T−L−Sml−DAPI−E8
7614−Ctx−T−L−Sml−DAPI−E9
7614−Ctx−T−L−Sml−DAPI−F10
7614−Ctx−T−L−Sml−DAPI−F11
7614−Ctx−T−L−Sml−DAPI−F12
7614−Ctx−T−L−Sml−DAPI−F7
7614−Ctx−T−L−Sml−DAPI−F8
7614−Ctx−T−L−Sml−DAPI−F9
7614−Ctx−T−L−Sml−DAPI−G10
7614−Ctx−T−L−Sml−DAPI−G11
7614−Ctx−T−L−Sml−DAPI−G12
7614−Ctx−T−L−Sml−DAPI−G7
7614−Ctx−T−L−Sml−DAPI−G8
7614−Ctx−T−L−Sml−DAPI−G9
7614−Ctx−T−L−Sml−DAPI−H10
7614−Ctx−T−L−Sml−DAPI−H11
7614−Ctx−T−L−Sml−DAPI−H7
7614−Ctx−T−L−Sml−DAPI−H8
7614−Ctx−T−L−Sml−DAPI−H9
7614−Ctx−T−L−Sml−NeuN−A1
7614−Ctx−T−L−Sml−NeuN−A2
7614−Ctx−T−L−Sml−NeuN−A3
7614−Ctx−T−L−Sml−NeuN−A4
7614−Ctx−T−L−Sml−NeuN−A5
7614−Ctx−T−L−Sml−NeuN−A6
7614−Ctx−T−L−Sml−NeuN−B1
7614−Ctx−T−L−Sml−NeuN−B2
7614−Ctx−T−L−Sml−NeuN−B3
7614−Ctx−T−L−Sml−NeuN−B4
7614−Ctx−T−L−Sml−NeuN−B5
7614−Ctx−T−L−Sml−NeuN−B6
7614−Ctx−T−L−Sml−NeuN−C1
7614−Ctx−T−L−Sml−NeuN−C2
7614−Ctx−T−L−Sml−NeuN−C3
7614−Ctx−T−L−Sml−NeuN−C4
7614−Ctx−T−L−Sml−NeuN−C5
7614−Ctx−T−L−Sml−NeuN−C6
7614−Ctx−T−L−Sml−NeuN−D1
7614−Ctx−T−L−Sml−NeuN−D2
7614−Ctx−T−L−Sml−NeuN−D3
7614−Ctx−T−L−Sml−NeuN−D4
7614−Ctx−T−L−Sml−NeuN−D5
7614−Ctx−T−L−Sml−NeuN−D6
7614−Ctx−T−L−Sml−NeuN−E1
7614−Ctx−T−L−Sml−NeuN−E2
7614−Ctx−T−L−Sml−NeuN−E3
7614−Ctx−T−L−Sml−NeuN−E4
7614−Ctx−T−L−Sml−NeuN−E5
7614−Ctx−T−L−Sml−NeuN−E6
7614−Ctx−T−L−Sml−NeuN−F1
7614−Ctx−T−L−Sml−NeuN−F2
7614−Ctx−T−L−Sml−NeuN−F3
7614−Ctx−T−L−Sml−NeuN−F4
7614−Ctx−T−L−Sml−NeuN−F5
7614−Ctx−T−L−Sml−NeuN−F6
7614−Ctx−T−L−Sml−NeuN−G1
7614−Ctx−T−L−Sml−NeuN−G2
7614−Ctx−T−L−Sml−NeuN−G3
7614−Ctx−T−L−Sml−NeuN−G4
7614−Ctx−T−L−Sml−NeuN−G5
7614−Ctx−T−L−Sml−NeuN−G6
7614−Ctx−T−L−Sml−NeuN−H1
7614−Ctx−T−L−Sml−NeuN−H2
7614−Ctx−T−L−Sml−NeuN−H3
7614−Ctx−T−L−Sml−NeuN−H4
7614−Ctx−T−L−Sml−NeuN−H5
7614−Ctx−T−L−Sml−NeuN−H6
JGG−Cells

7−121347585−T−G

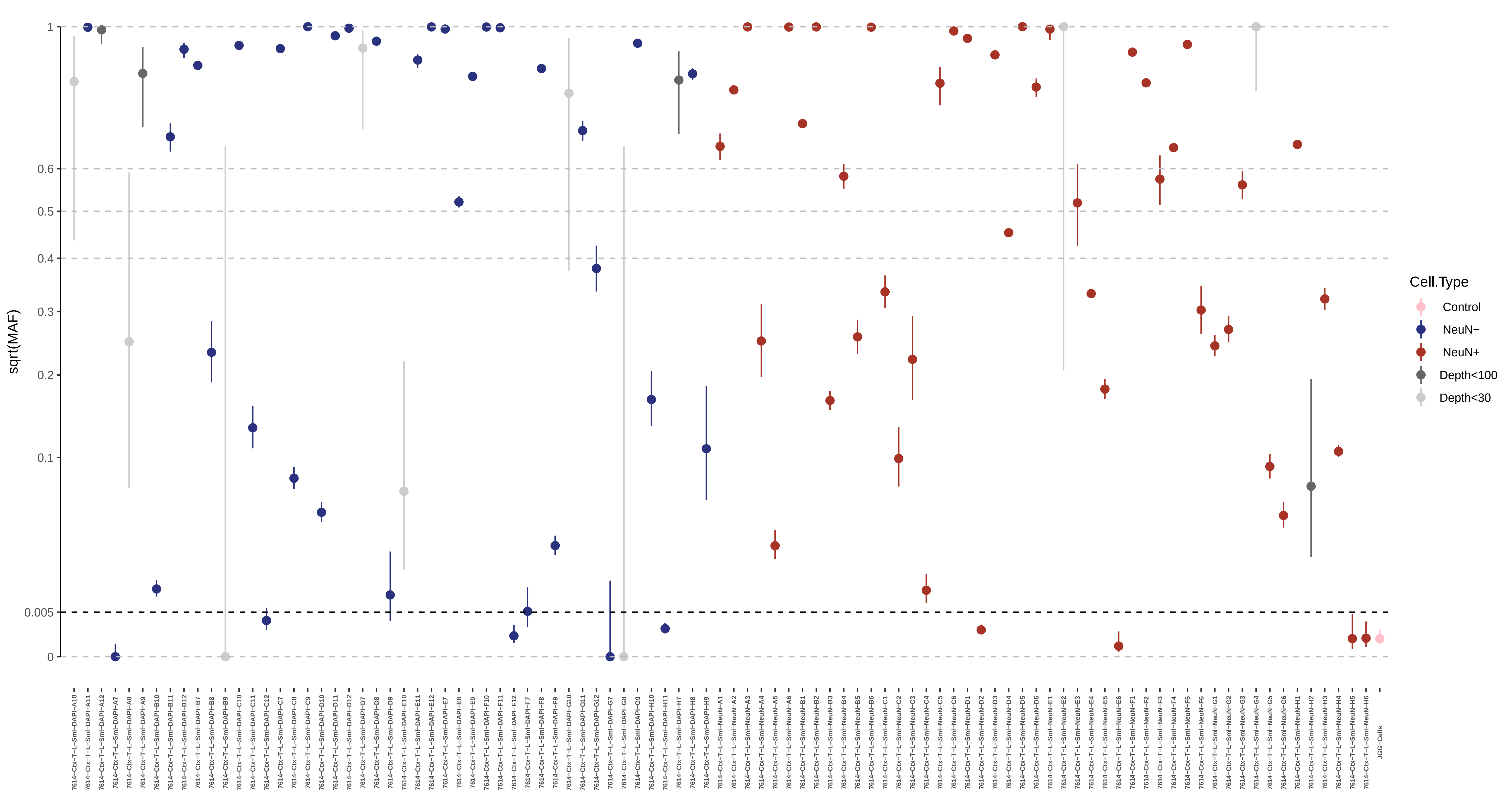

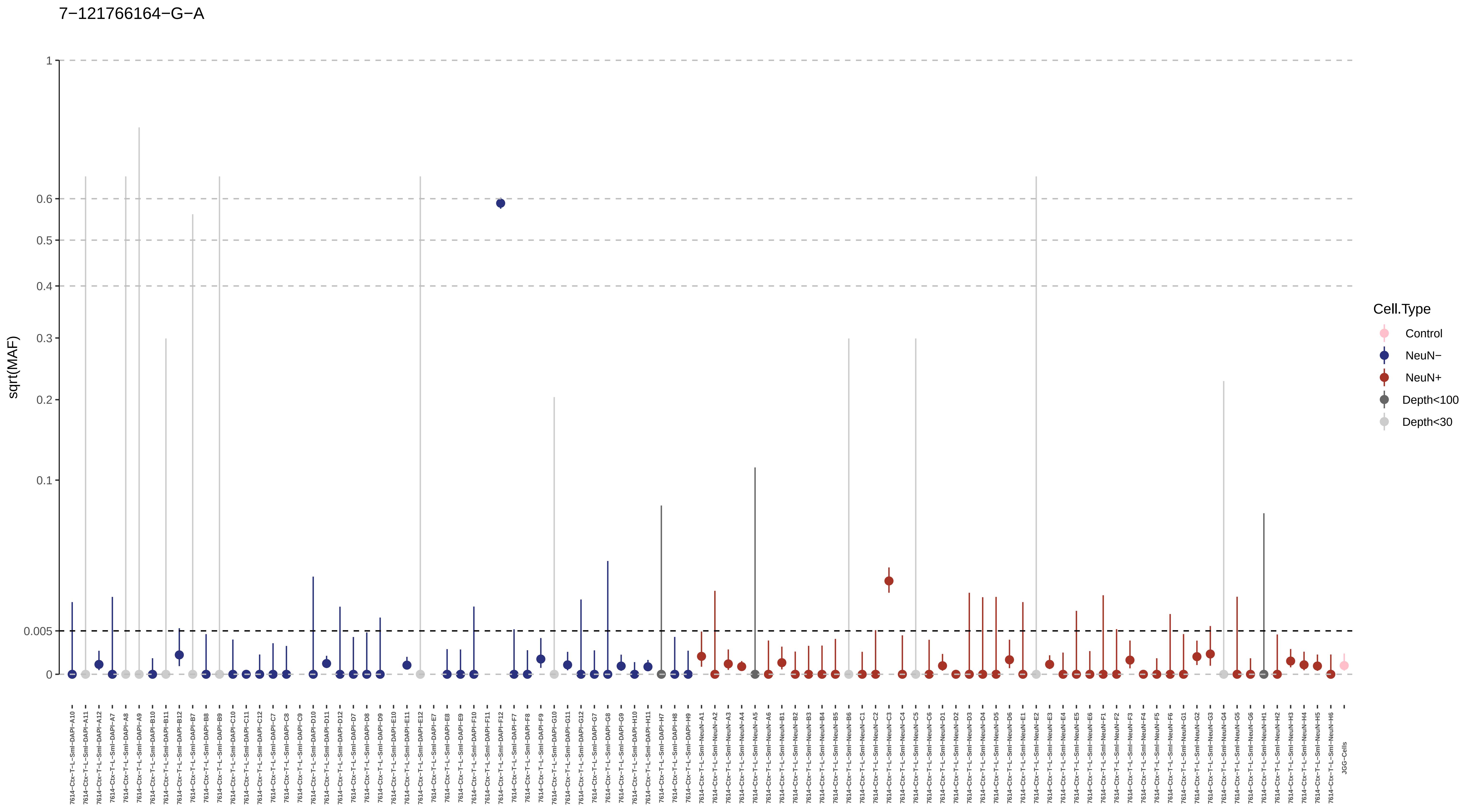
7−121766164−G−A
sqrt(MAF)
1
0.6
0.5
0.4
0.3
0.2
0.1
0.005
0
Cell.Type
Control
NeuN−
NeuN+
Depth<100
Depth<30

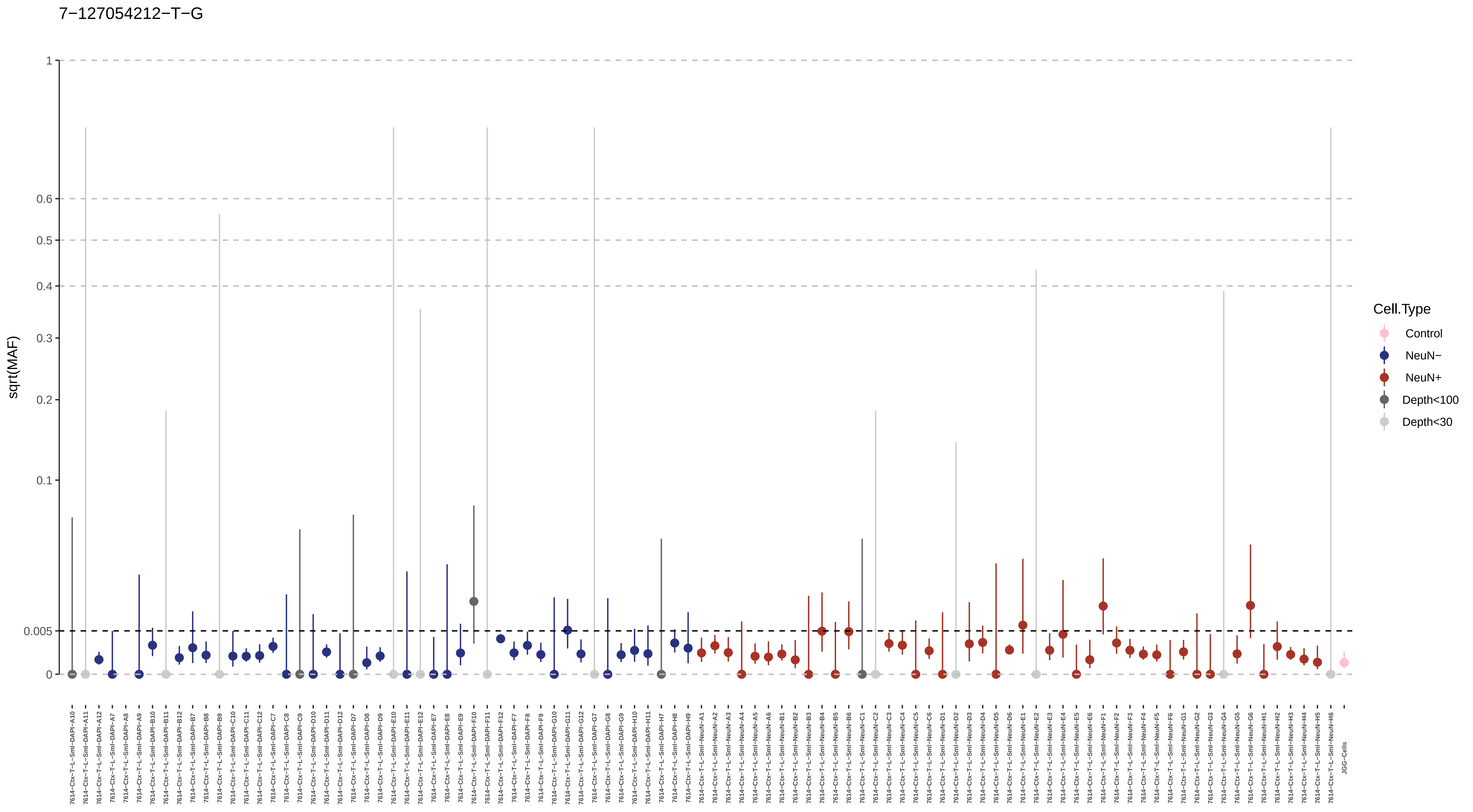

7−127054212−T−G
sqrt(MAF)
1
0.6
0.5
0.4
0.3
0.2
0.1
0.005
0
Cell.Type
Control
NeuN−
NeuN+
Depth<100
Depth<30
7614−Ctx−T−L−Sml−DAPI−A10
7614−Ctx−T−L−Sml−DAPI−A11
7614−Ctx−T−L−Sml−DAPI−A12
7614−Ctx−T−L−Sml−DAPI−A7
7614−Ctx−T−L−Sml−DAPI−A8
7614−Ctx−T−L−Sml−DAPI−A9
7614−Ctx−T−L−Sml−DAPI−B10
7614−Ctx−T−L−Sml−DAPI−B11
7614−Ctx−T−L−Sml−DAPI−B12
7614−Ctx−T−L−Sml−DAPI−B7
7614−Ctx−T−L−Sml−DAPI−B8
7614−Ctx−T−L−Sml−DAPI−B9
7614−Ctx−T−L−Sml−DAPI−C10
7614−Ctx−T−L−Sml−DAPI−C11
7614−Ctx−T−L−Sml−DAPI−C12
7614−Ctx−T−L−Sml−DAPI−C7
7614−Ctx−T−L−Sml−DAPI−C8
7614−Ctx−T−L−Sml−DAPI−C9
7614−Ctx−T−L−Sml−DAPI−D10
7614−Ctx−T−L−Sml−DAPI−D11
7614−Ctx−T−L−Sml−DAPI−D12
7614−Ctx−T−L−Sml−DAPI−D7
7614−Ctx−T−L−Sml−DAPI−D8
7614−Ctx−T−L−Sml−DAPI−D9
7614−Ctx−T−L−Sml−DAPI−E10
7614−Ctx−T−L−Sml−DAPI−E11
7614−Ctx−T−L−Sml−DAPI−E12
7614−Ctx−T−L−Sml−DAPI−E7
7614−Ctx−T−L−Sml−DAPI−E8
7614−Ctx−T−L−Sml−DAPI−E9
7614−Ctx−T−L−Sml−DAPI−F10
7614−Ctx−T−L−Sml−DAPI−F11
7614−Ctx−T−L−Sml−DAPI−F12
7614−Ctx−T−L−Sml−DAPI−F7
7614−Ctx−T−L−Sml−DAPI−F8
7614−Ctx−T−L−Sml−DAPI−F9
7614−Ctx−T−L−Sml−DAPI−G10
7614−Ctx−T−L−Sml−DAPI−G11
7614−Ctx−T−L−Sml−DAPI−G12
7614−Ctx−T−L−Sml−DAPI−G7
7614−Ctx−T−L−Sml−DAPI−G8
7614−Ctx−T−L−Sml−DAPI−G9
7614−Ctx−T−L−Sml−DAPI−H10
7614−Ctx−T−L−Sml−DAPI−H11
7614−Ctx−T−L−Sml−DAPI−H7
7614−Ctx−T−L−Sml−DAPI−H8
7614−Ctx−T−L−Sml−DAPI−H9
7614−Ctx−T−L−Sml−NeuN−A1
7614−Ctx−T−L−Sml−NeuN−A2
7614−Ctx−T−L−Sml−NeuN−A3
7614−Ctx−T−L−Sml−NeuN−A4
7614−Ctx−T−L−Sml−NeuN−A5
7614−Ctx−T−L−Sml−NeuN−A6
7614−Ctx−T−L−Sml−NeuN−B1
7614−Ctx−T−L−Sml−NeuN−B2
7614−Ctx−T−L−Sml−NeuN−B3
7614−Ctx−T−L−Sml−NeuN−B4
7614−Ctx−T−L−Sml−NeuN−B5
7614−Ctx−T−L−Sml−NeuN−B6
7614−Ctx−T−L−Sml−NeuN−C1
7614−Ctx−T−L−Sml−NeuN−C2
7614−Ctx−T−L−Sml−NeuN−C3
7614−Ctx−T−L−Sml−NeuN−C4
7614−Ctx−T−L−Sml−NeuN−C5
7614−Ctx−T−L−Sml−NeuN−C6
7614−Ctx−T−L−Sml−NeuN−D1
7614−Ctx−T−L−Sml−NeuN−D2
7614−Ctx−T−L−Sml−NeuN−D3
7614−Ctx−T−L−Sml−NeuN−D4
7614−Ctx−T−L−Sml−NeuN−D5
7614−Ctx−T−L−Sml−NeuN−D6
7614−Ctx−T−L−Sml−NeuN−E1
7614−Ctx−T−L−Sml−NeuN−E2
7614−Ctx−T−L−Sml−NeuN−E3
7614−Ctx−T−L−Sml−NeuN−E4
7614−Ctx−T−L−Sml−NeuN−E5
7614−Ctx−T−L−Sml−NeuN−E6
7614−Ctx−T−L−Sml−NeuN−F1
7614−Ctx−T−L−Sml−NeuN−F2
7614−Ctx−T−L−Sml−NeuN−F3
7614−Ctx−T−L−Sml−NeuN−F4
7614−Ctx−T−L−Sml−NeuN−F5
7614−Ctx−T−L−Sml−NeuN−F6
7614−Ctx−T−L−Sml−NeuN−G1
7614−Ctx−T−L−Sml−NeuN−G2
7614−Ctx−T−L−Sml−NeuN−G3
7614−Ctx−T−L−Sml−NeuN−G4
7614−Ctx−T−L−Sml−NeuN−G5
7614−Ctx−T−L−Sml−NeuN−G6
7614−Ctx−T−L−Sml−NeuN−H1
7614−Ctx−T−L−Sml−NeuN−H2
7614−Ctx−T−L−Sml−NeuN−H3
7614−Ctx−T−L−Sml−NeuN−H4
7614−Ctx−T−L−Sml−NeuN−H5
7614−Ctx−T−L−Sml−NeuN−H6
JGG-Cells

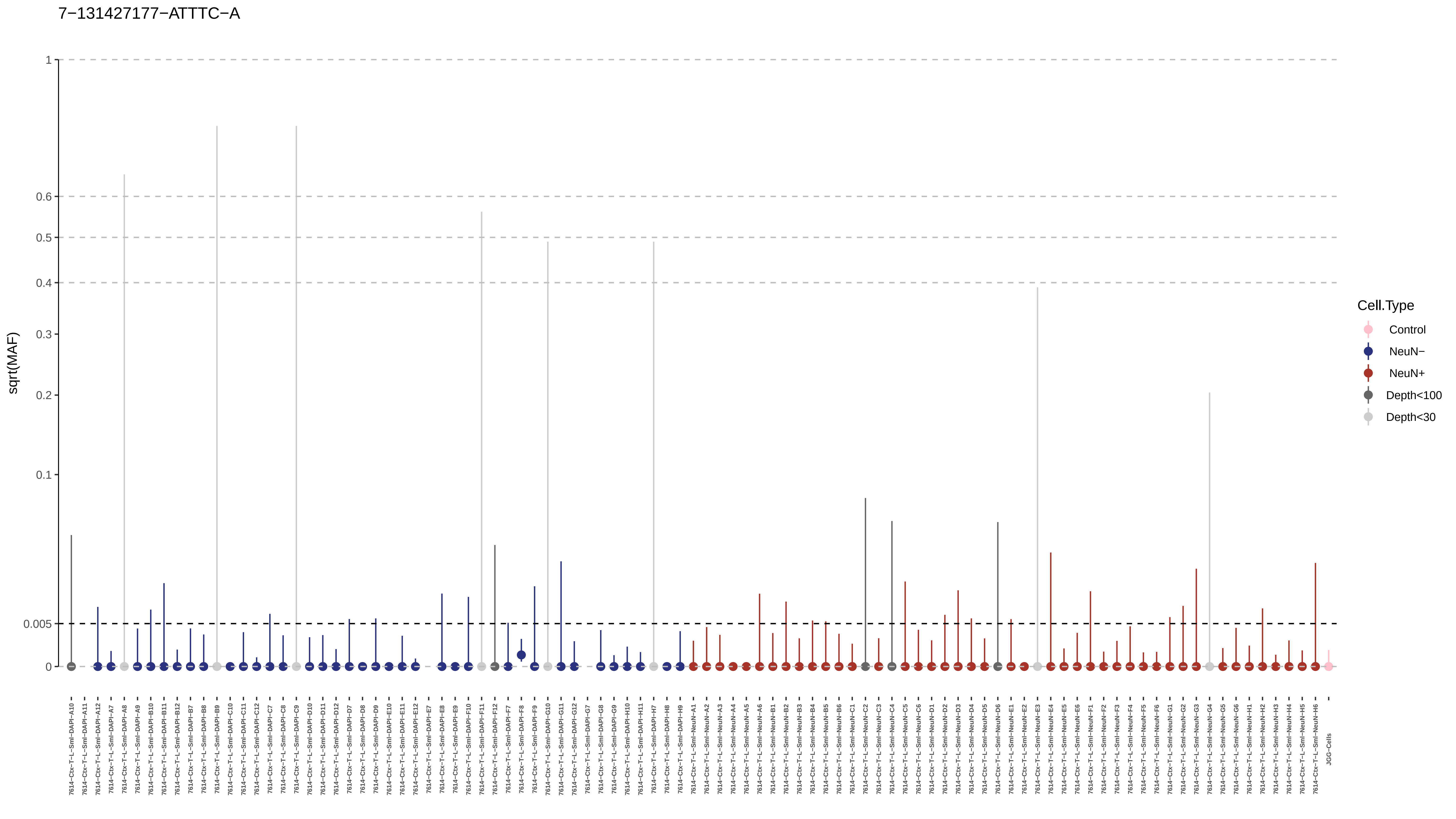
7−131427177−ATTTC−A
sqrt(MAF)
1
0.6
0.5
0.4
0.3
0.2
0.1
0.005
0
Cell.Type
Control
NeuN−
NeuN+
Depth<100
Depth<30
JGG-Cells

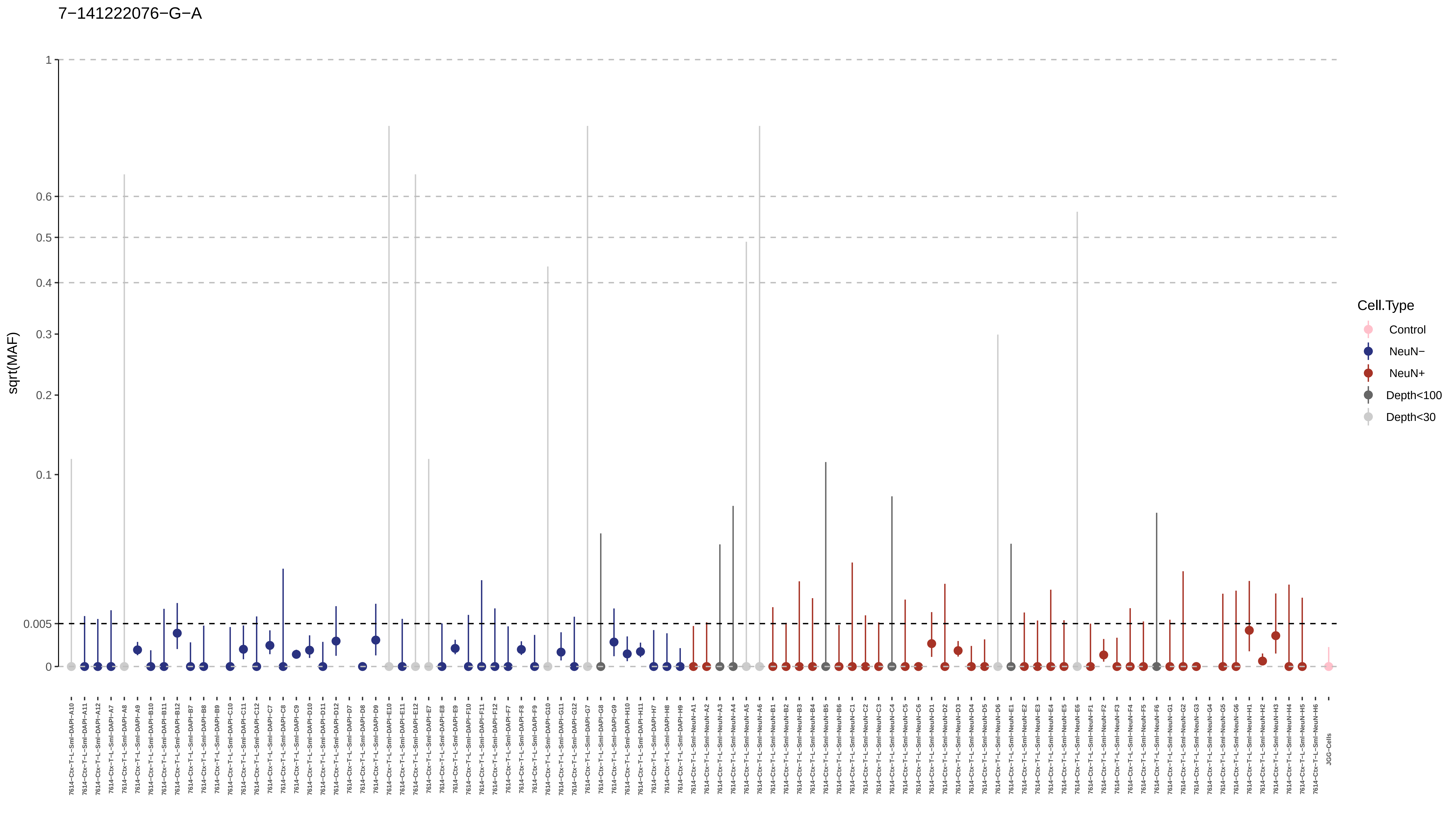
7−141222076−G−A
sqrt(MAF)
1
0.6
0.5
0.4
0.3
0.2
0.1
0.005
0
7614−Ctx−T−L−Sml−DAPI−A10
7614−Ctx−T−L−Sml−DAPI−A11
7614−Ctx−T−L−Sml−DAPI−A12
7614−Ctx−T−L−Sml−DAPI−A7
7614−Ctx−T−L−Sml−DAPI−A8
7614−Ctx−T−L−Sml−DAPI−A9
7614−Ctx−T−L−Sml−DAPI−B10
7614−Ctx−T−L−Sml−DAPI−B11
7614−Ctx−T−L−Sml−DAPI−B12
7614−Ctx−T−L−Sml−DAPI−B7
7614−Ctx−T−L−Sml−DAPI−B8
7614−Ctx−T−L−Sml−DAPI−B9
7614−Ctx−T−L−Sml−DAPI−C10
7614−Ctx−T−L−Sml−DAPI−C11
7614−Ctx−T−L−Sml−DAPI−C12
7614−Ctx−T−L−Sml−DAPI−C7
7614−Ctx−T−L−Sml−DAPI−C8
7614−Ctx−T−L−Sml−DAPI−C9
7614−Ctx−T−L−Sml−DAPI−D10
7614−Ctx−T−L−Sml−DAPI−D11
7614−Ctx−T−L−Sml−DAPI−D12
7614−Ctx−T−L−Sml−DAPI−D7
7614−Ctx−T−L−Sml−DAPI−D8
7614−Ctx−T−L−Sml−DAPI−D9
7614−Ctx−T−L−Sml−DAPI−E10
7614−Ctx−T−L−Sml−DAPI−E11
7614−Ctx−T−L−Sml−DAPI−E12
7614−Ctx−T−L−Sml−DAPI−E7
7614−Ctx−T−L−Sml−DAPI−E8
7614−Ctx−T−L−Sml−DAPI−E9
7614−Ctx−T−L−Sml−DAPI−F10
7614−Ctx−T−L−Sml−DAPI−F11
7614−Ctx−T−L−Sml−DAPI−F12
7614−Ctx−T−L−Sml−DAPI−F7
7614−Ctx−T−L−Sml−DAPI−F8
7614−Ctx−T−L−Sml−DAPI−F9
7614−Ctx−T−L−Sml−DAPI−G10
7614−Ctx−T−L−Sml−DAPI−G11
7614−Ctx−T−L−Sml−DAPI−G12
7614−Ctx−T−L−Sml−DAPI−G7
7614−Ctx−T−L−Sml−DAPI−G8
7614−Ctx−T−L−Sml−DAPI−G9
7614−Ctx−T−L−Sml−DAPI−H10
7614−Ctx−T−L−Sml−DAPI−H11
7614−Ctx−T−L−Sml−DAPI−H7
7614−Ctx−T−L−Sml−DAPI−H8
7614−Ctx−T−L−Sml−DAPI−H9
7614−Ctx−T−L−Sml−NeuN−A1
7614−Ctx−T−L−Sml−NeuN−A2
7614−Ctx−T−L−Sml−NeuN−A3
7614−Ctx−T−L−Sml−NeuN−A4
7614−Ctx−T−L−Sml−NeuN−A5
7614−Ctx−T−L−Sml−NeuN−A6
7614−Ctx−T−L−Sml−NeuN−B1
7614−Ctx−T−L−Sml−NeuN−B2
7614−Ctx−T−L−Sml−NeuN−B3
7614−Ctx−T−L−Sml−NeuN−B4
7614−Ctx−T−L−Sml−NeuN−B5
7614−Ctx−T−L−Sml−NeuN−B6
7614−Ctx−T−L−Sml−NeuN−C1
7614−Ctx−T−L−Sml−NeuN−C2
7614−Ctx−T−L−Sml−NeuN−C3
7614−Ctx−T−L−Sml−NeuN−C4
7614−Ctx−T−L−Sml−NeuN−C5
7614−Ctx−T−L−Sml−NeuN−C6
7614−Ctx−T−L−Sml−NeuN−D1
7614−Ctx−T−L−Sml−NeuN−D2
7614−Ctx−T−L−Sml−NeuN−D3
7614−Ctx−T−L−Sml−NeuN−D4
7614−Ctx−T−L−Sml−NeuN−D5
7614−Ctx−T−L−Sml−NeuN−D6
7614−Ctx−T−L−Sml−NeuN−E1
7614−Ctx−T−L−Sml−NeuN−E2
7614−Ctx−T−L−Sml−NeuN−E3
7614−Ctx−T−L−Sml−NeuN−E4
7614−Ctx−T−L−Sml−NeuN−E5
7614−Ctx−T−L−Sml−NeuN−E6
7614−Ctx−T−L−Sml−NeuN−F1
7614−Ctx−T−L−Sml−NeuN−F2
7614−Ctx−T−L−Sml−NeuN−F3
7614−Ctx−T−L−Sml−NeuN−F4
7614−Ctx−T−L−Sml−NeuN−F5
7614−Ctx−T−L−Sml−NeuN−F6
7614−Ctx−T−L−Sml−NeuN−G1
7614−Ctx−T−L−Sml−NeuN−G2
7614−Ctx−T−L−Sml−NeuN−G3
7614−Ctx−T−L−Sml−NeuN−G4
7614−Ctx−T−L−Sml−NeuN−G5
7614−Ctx−T−L−Sml−NeuN−G6
7614−Ctx−T−L−Sml−NeuN−H1
7614−Ctx−T−L−Sml−NeuN−H2
7614−Ctx−T−L−Sml−NeuN−H3
7614−Ctx−T−L−Sml−NeuN−H4
7614−Ctx−T−L−Sml−NeuN−H5
7614−Ctx−T−L−Sml−NeuN−H6
JGG−Cells
Cell.Type
Control
NeuN−
NeuN+
Depth<100
Depth<30

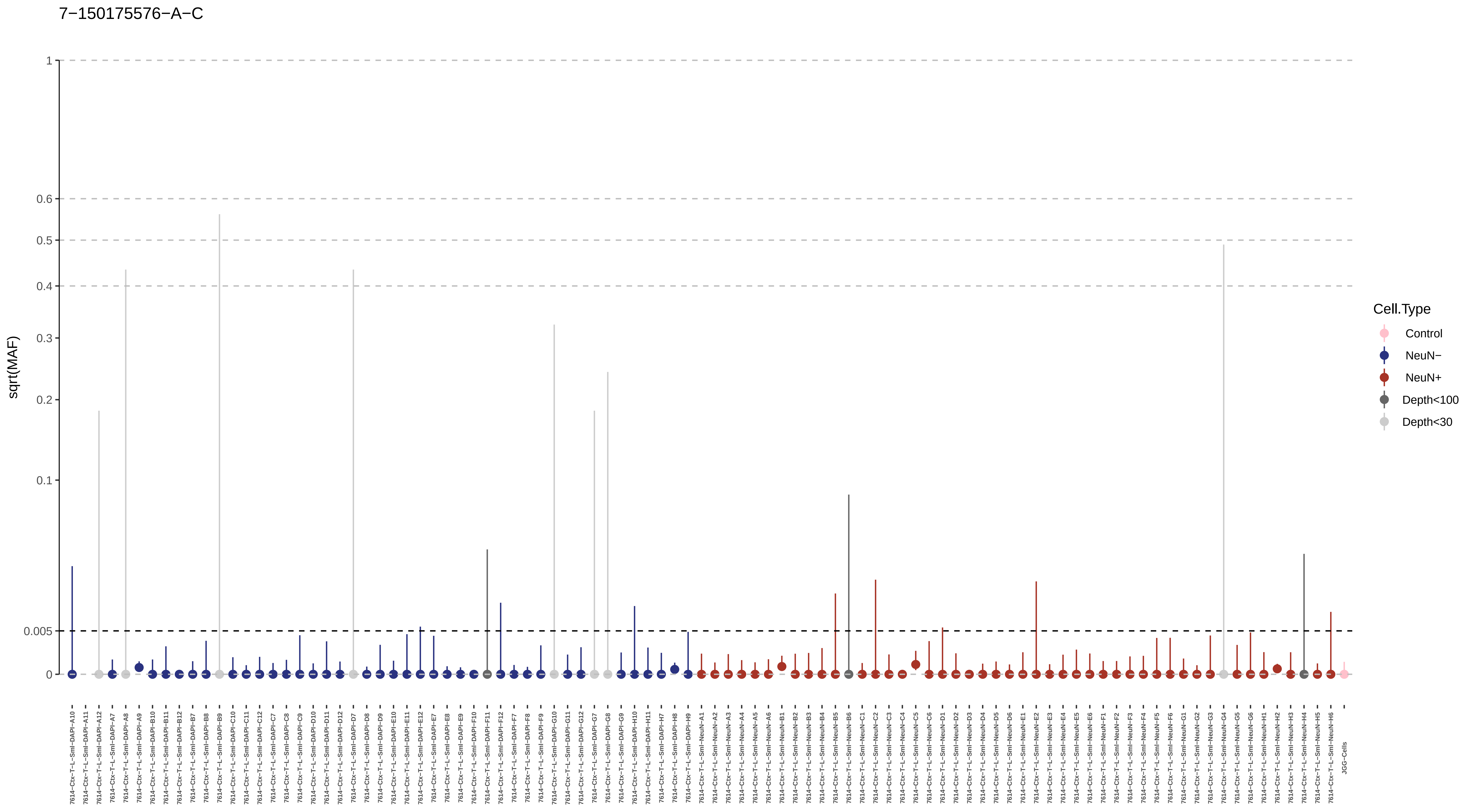
7−150175576−A−C
sqrt(MAF)
1
0.6
0.5
0.4
0.3
0.2
0.1
0.005
0
Cell.Type
Control
NeuN−
NeuN+
Depth<100
Depth<30
7614−Ctx−T−L−Sml−DAPI−A10
7614−Ctx−T−L−Sml−DAPI−A11
7614−Ctx−T−L−Sml−DAPI−A12
7614−Ctx−T−L−Sml−DAPI−A7
7614−Ctx−T−L−Sml−DAPI−A8
7614−Ctx−T−L−Sml−DAPI−A9
7614−Ctx−T−L−Sml−DAPI−B10
7614−Ctx−T−L−Sml−DAPI−B11
7614−Ctx−T−L−Sml−DAPI−B12
7614−Ctx−T−L−Sml−DAPI−B7
7614−Ctx−T−L−Sml−DAPI−B8
7614−Ctx−T−L−Sml−DAPI−B9
7614−Ctx−T−L−Sml−DAPI−C10
7614−Ctx−T−L−Sml−DAPI−C11
7614−Ctx−T−L−Sml−DAPI−C12
7614−Ctx−T−L−Sml−DAPI−C7
7614−Ctx−T−L−Sml−DAPI−C8
7614−Ctx−T−L−Sml−DAPI−C9
7614−Ctx−T−L−Sml−DAPI−D10
7614−Ctx−T−L−Sml−DAPI−D11
7614−Ctx−T−L−Sml−DAPI−D12
7614−Ctx−T−L−Sml−DAPI−D7
7614−Ctx−T−L−Sml−DAPI−D8
7614−Ctx−T−L−Sml−DAPI−D9
7614−Ctx−T−L−Sml−DAPI−E10
7614−Ctx−T−L−Sml−DAPI−E11
7614−Ctx−T−L−Sml−DAPI−E12
7614−Ctx−T−L−Sml−DAPI−E7
7614−Ctx−T−L−Sml−DAPI−E8
7614−Ctx−T−L−Sml−DAPI−E9
7614−Ctx−T−L−Sml−DAPI−F10
7614−Ctx−T−L−Sml−DAPI−F11
7614−Ctx−T−L−Sml−DAPI−F12
7614−Ctx−T−L−Sml−DAPI−F7
7614−Ctx−T−L−Sml−DAPI−F8
7614−Ctx−T−L−Sml−DAPI−F9
7614−Ctx−T−L−Sml−DAPI−G10
7614−Ctx−T−L−Sml−DAPI−G11
7614−Ctx−T−L−Sml−DAPI−G12
7614−Ctx−T−L−Sml−DAPI−G7
7614−Ctx−T−L−Sml−DAPI−G8
7614−Ctx−T−L−Sml−DAPI−G9
7614−Ctx−T−L−Sml−DAPI−H10
7614−Ctx−T−L−Sml−DAPI−H11
7614−Ctx−T−L−Sml−DAPI−H7
7614−Ctx−T−L−Sml−DAPI−H8
7614−Ctx−T−L−Sml−DAPI−H9
7614−Ctx−T−L−Sml−NeuN−A1
7614−Ctx−T−L−Sml−NeuN−A2
7614−Ctx−T−L−Sml−NeuN−A3
7614−Ctx−T−L−Sml−NeuN−A4
7614−Ctx−T−L−Sml−NeuN−A5
7614−Ctx−T−L−Sml−NeuN−A6
7614−Ctx−T−L−Sml−NeuN−B1
7614−Ctx−T−L−Sml−NeuN−B2
7614−Ctx−T−L−Sml−NeuN−B3
7614−Ctx−T−L−Sml−NeuN−B4
7614−Ctx−T−L−Sml−NeuN−B5
7614−Ctx−T−L−Sml−NeuN−B6
7614−Ctx−T−L−Sml−NeuN−C1
7614−Ctx−T−L−Sml−NeuN−C2
7614−Ctx−T−L−Sml−NeuN−C3
7614−Ctx−T−L−Sml−NeuN−C4
7614−Ctx−T−L−Sml−NeuN−C5
7614−Ctx−T−L−Sml−NeuN−C6
7614−Ctx−T−L−Sml−NeuN−D1
7614−Ctx−T−L−Sml−NeuN−D2
7614−Ctx−T−L−Sml−NeuN−D3
7614−Ctx−T−L−Sml−NeuN−D4
7614−Ctx−T−L−Sml−NeuN−D5
7614−Ctx−T−L−Sml−NeuN−D6
7614−Ctx−T−L−Sml−NeuN−E1
7614−Ctx−T−L−Sml−NeuN−E2
7614−Ctx−T−L−Sml−NeuN−E3
7614−Ctx−T−L−Sml−NeuN−E4
7614−Ctx−T−L−Sml−NeuN−E5
7614−Ctx−T−L−Sml−NeuN−E6
7614−Ctx−T−L−Sml−NeuN−F1
7614−Ctx−T−L−Sml−NeuN−F2
7614−Ctx−T−L−Sml−NeuN−F3
7614−Ctx−T−L−Sml−NeuN−F4
7614−Ctx−T−L−Sml−NeuN−F5
7614−Ctx−T−L−Sml−NeuN−F6
7614−Ctx−T−L−Sml−NeuN−G1
7614−Ctx−T−L−Sml−NeuN−G2
7614−Ctx−T−L−Sml−NeuN−G3
7614−Ctx−T−L−Sml−NeuN−G4
7614−Ctx−T−L−Sml−NeuN−G5
7614−Ctx−T−L−Sml−NeuN−G6
7614−Ctx−T−L−Sml−NeuN−H1
7614−Ctx−T−L−Sml−NeuN−H2
7614−Ctx−T−L−Sml−NeuN−H3
7614−Ctx−T−L−Sml−NeuN−H4
7614−Ctx−T−L−Sml−NeuN−H5
7614−Ctx−T−L−Sml−NeuN−H6
JGG−Cells

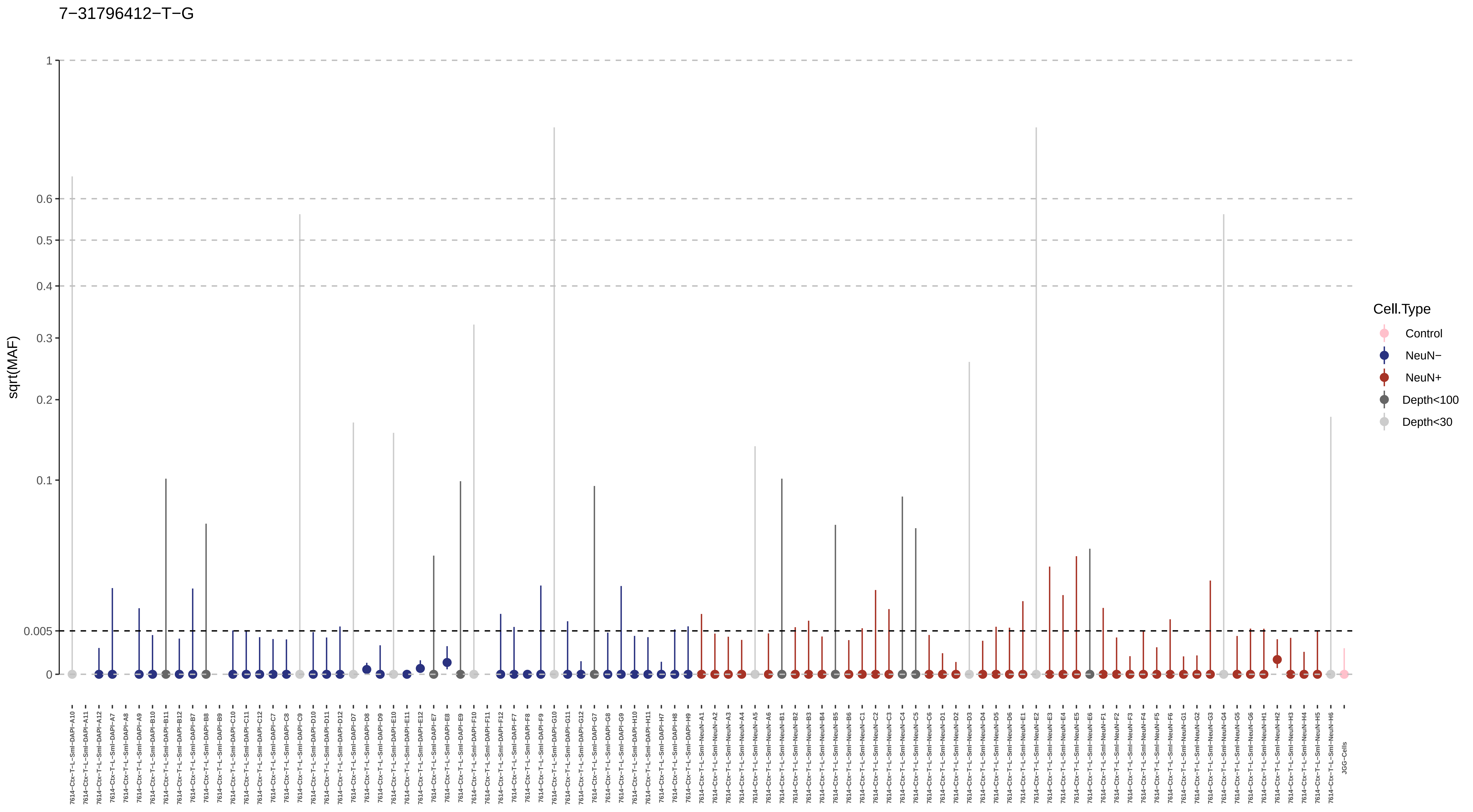

7−31796412−T−G
sqrt(MAF)
1
0.6
0.5
0.4
0.3
0.2
0.1
0.005
0
Cell.Type
Control
NeuN−
NeuN+
Depth<100
Depth<30

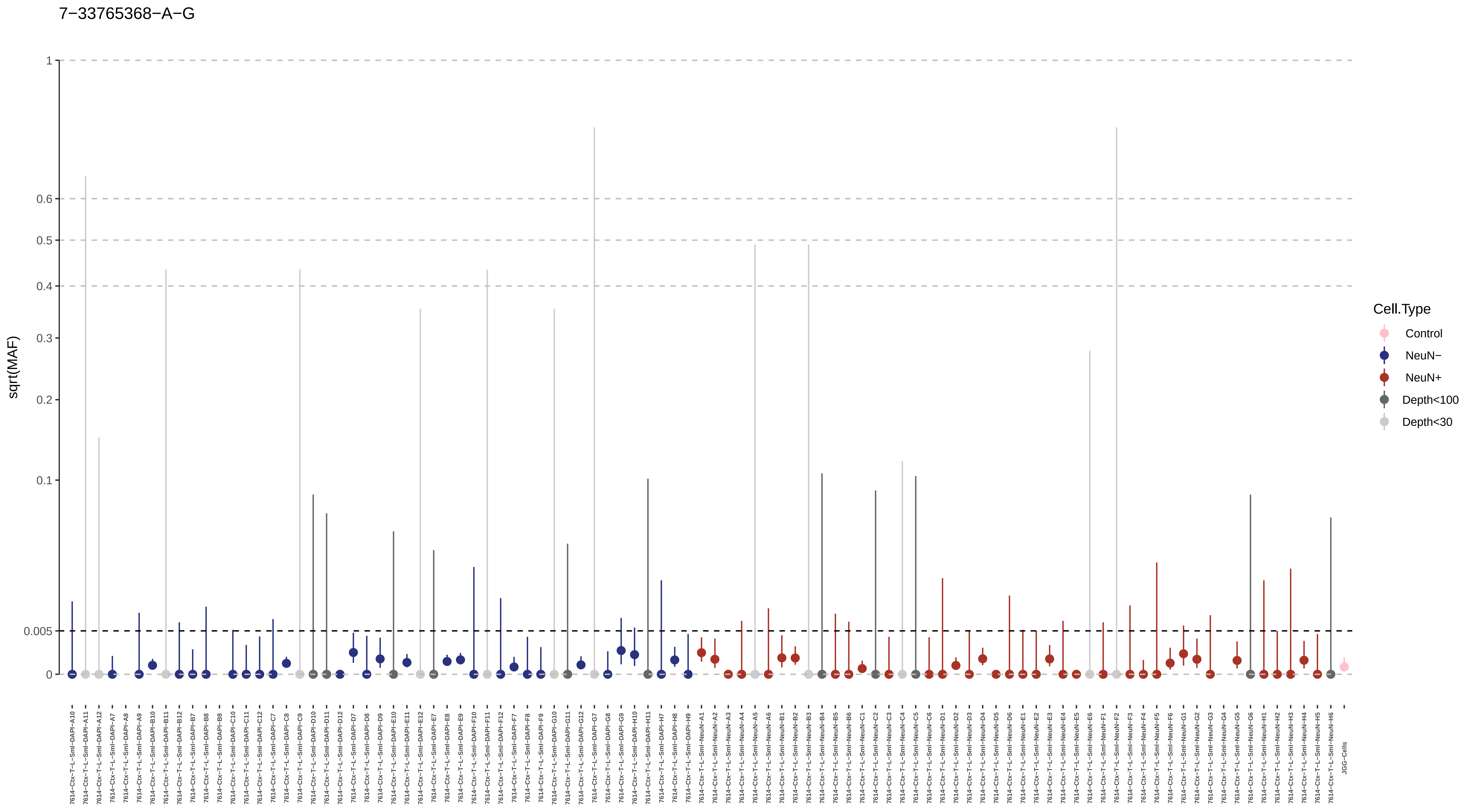
7−33765368−A−G
sqrt(MAF)
1
0.6
0.5
0.4
0.3
0.2
0.1
0.005
0
Cell.Type
Control
NeuN−
NeuN+
Depth<100
Depth<30

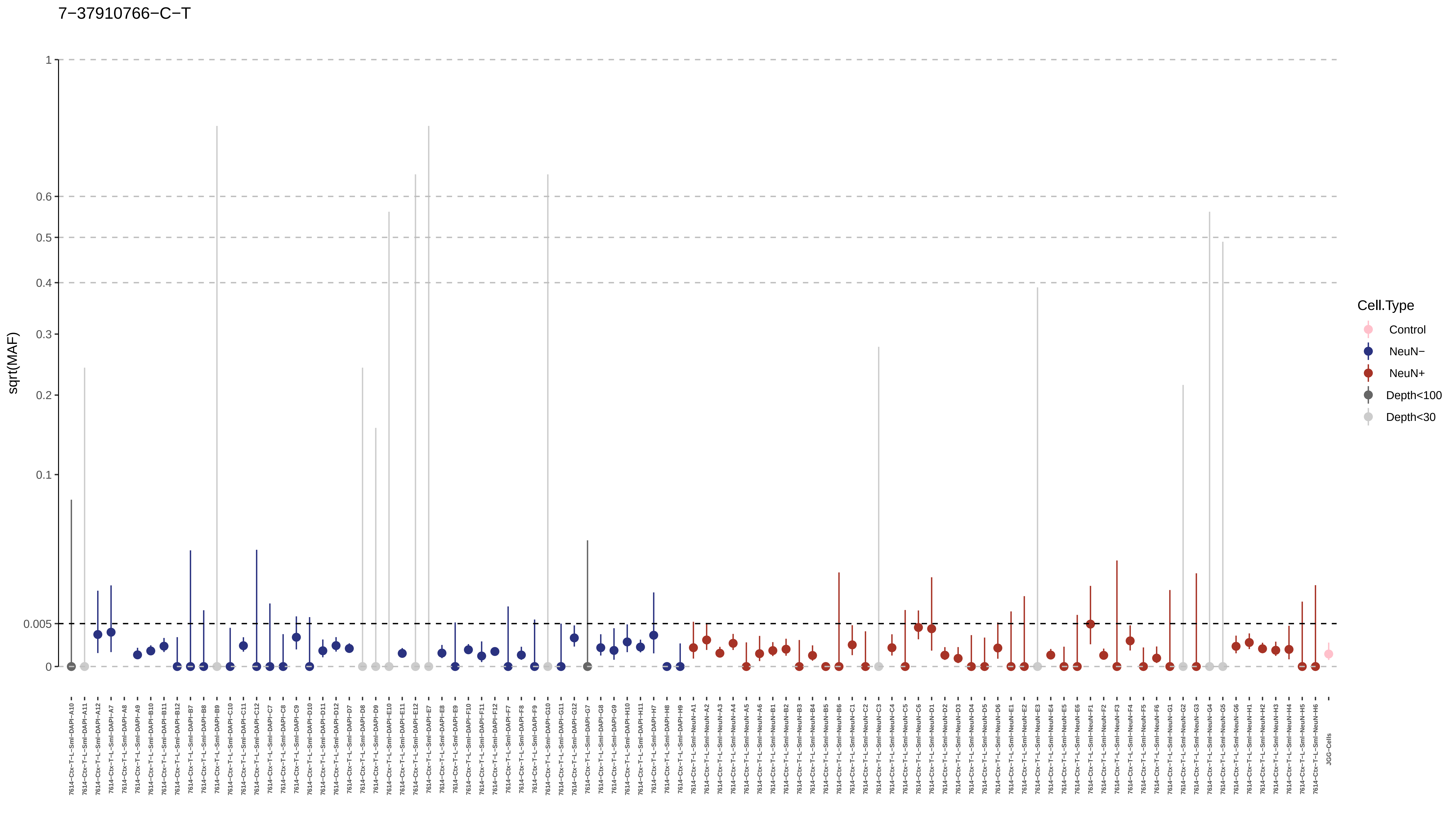

7−37910766−C−T
sqrt(MAF)
1
0.6
0.5
0.4
0.3
0.2
0.1
0.005
0
Cell.Type
Control
NeuN−
NeuN+
Depth<100
Depth<30

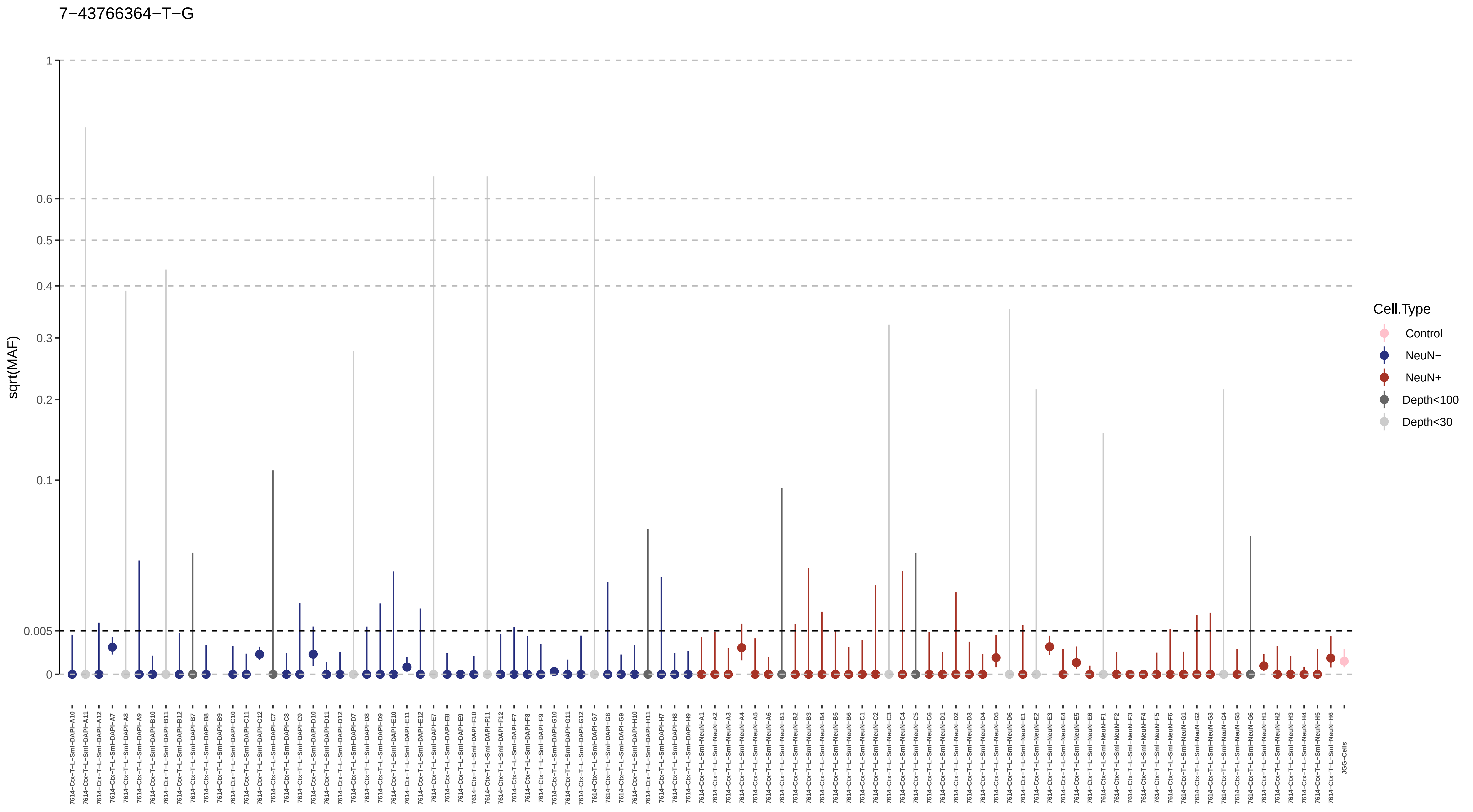
7−43766364−T−G
sqrt(MAF)
1
0.6
0.5
0.4
0.3
0.2
0.1
0.005
0
Cell.Type
Control
NeuN−
NeuN+
Depth<100
Depth<30
JGG-Cells

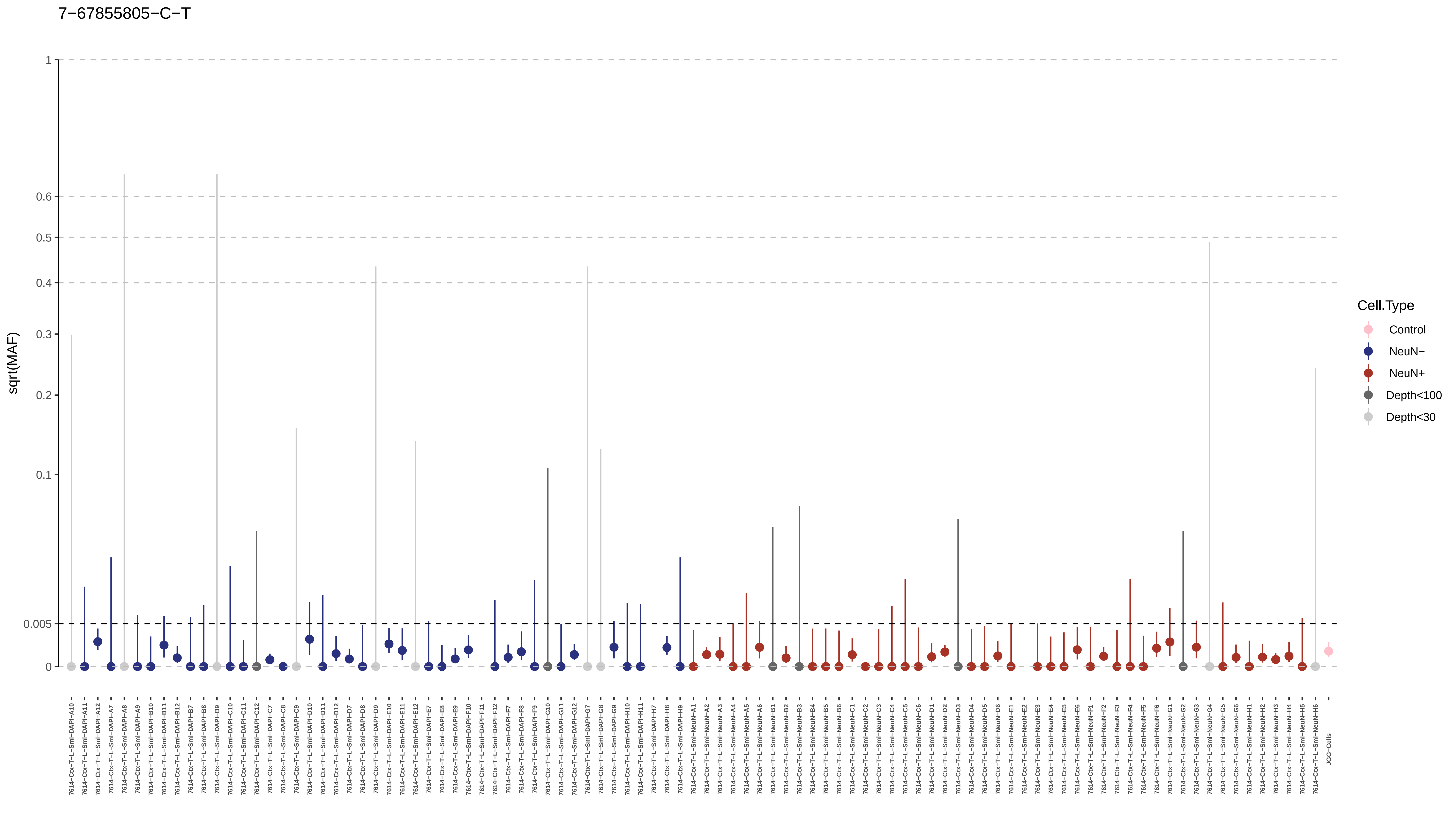
7−67855805−C−T
sqrt(MAF)
1
0.6
0.5
0.4
0.3
0.2
0.1
0.005
0
Cell.Type
Control
NeuN−
NeuN+
Depth<100
Depth<30

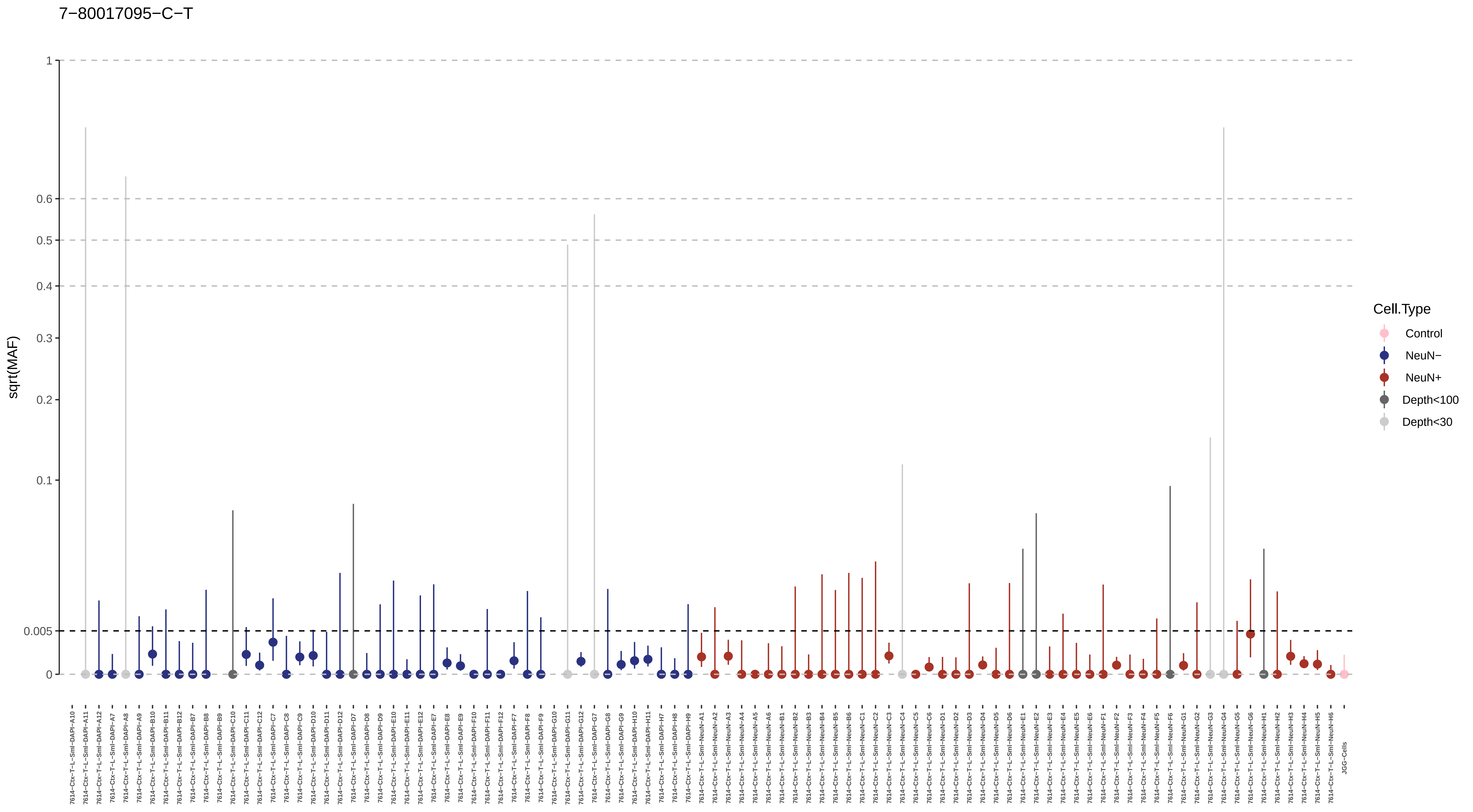

7−80017095−C−T
sqrt(MAF)
1
0.6
0.5
0.4
0.3
0.2
0.1
0.005
0
Cell.Type
Control
NeuN−
NeuN+
Depth<100
Depth<30

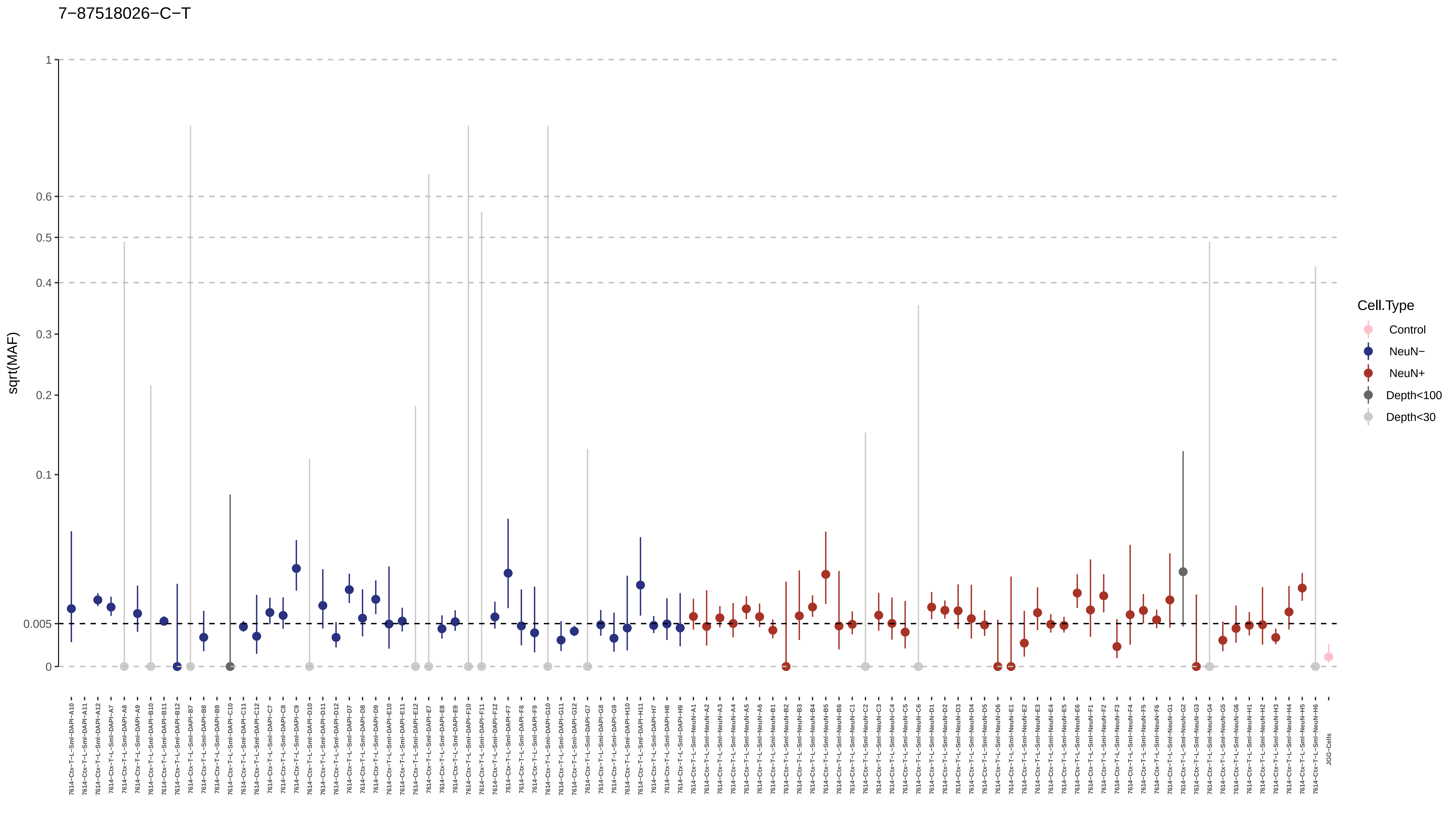

7−87518026−C−T
sqrt(MAF)
1
0.6
0.5
0.4
0.3
0.2
0.1
0.005
0
Cell.Type
Control
NeuN−
NeuN+
Depth<100
Depth<30

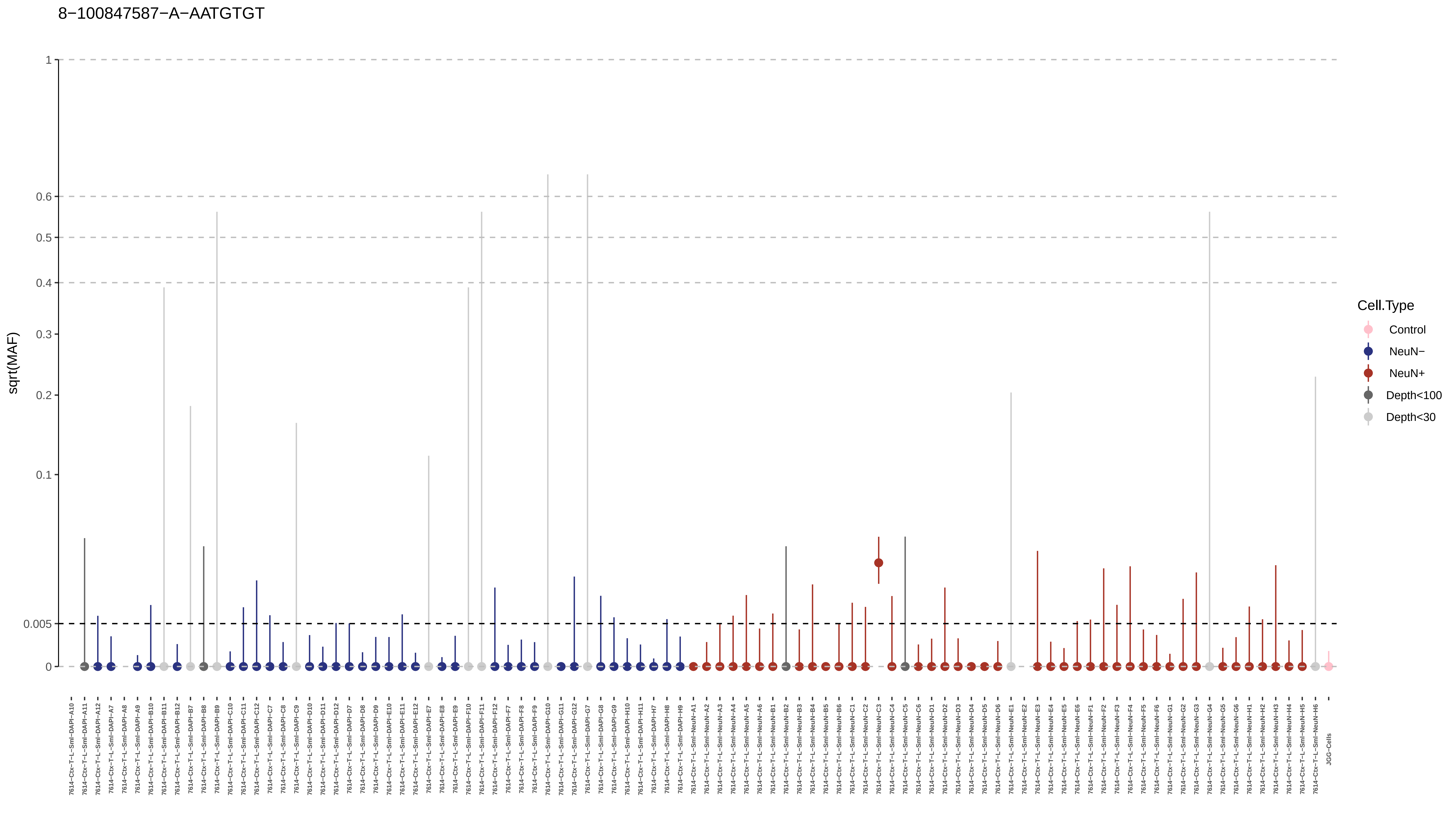
8−100847587−A−AATGTGT
sqrt(MAF)
1
0.6
0.5
0.4
0.3
0.2
0.1
0.005
0
Cell.Type
Control
NeuN−
NeuN+
Depth<100
Depth<30
JGG-Cells

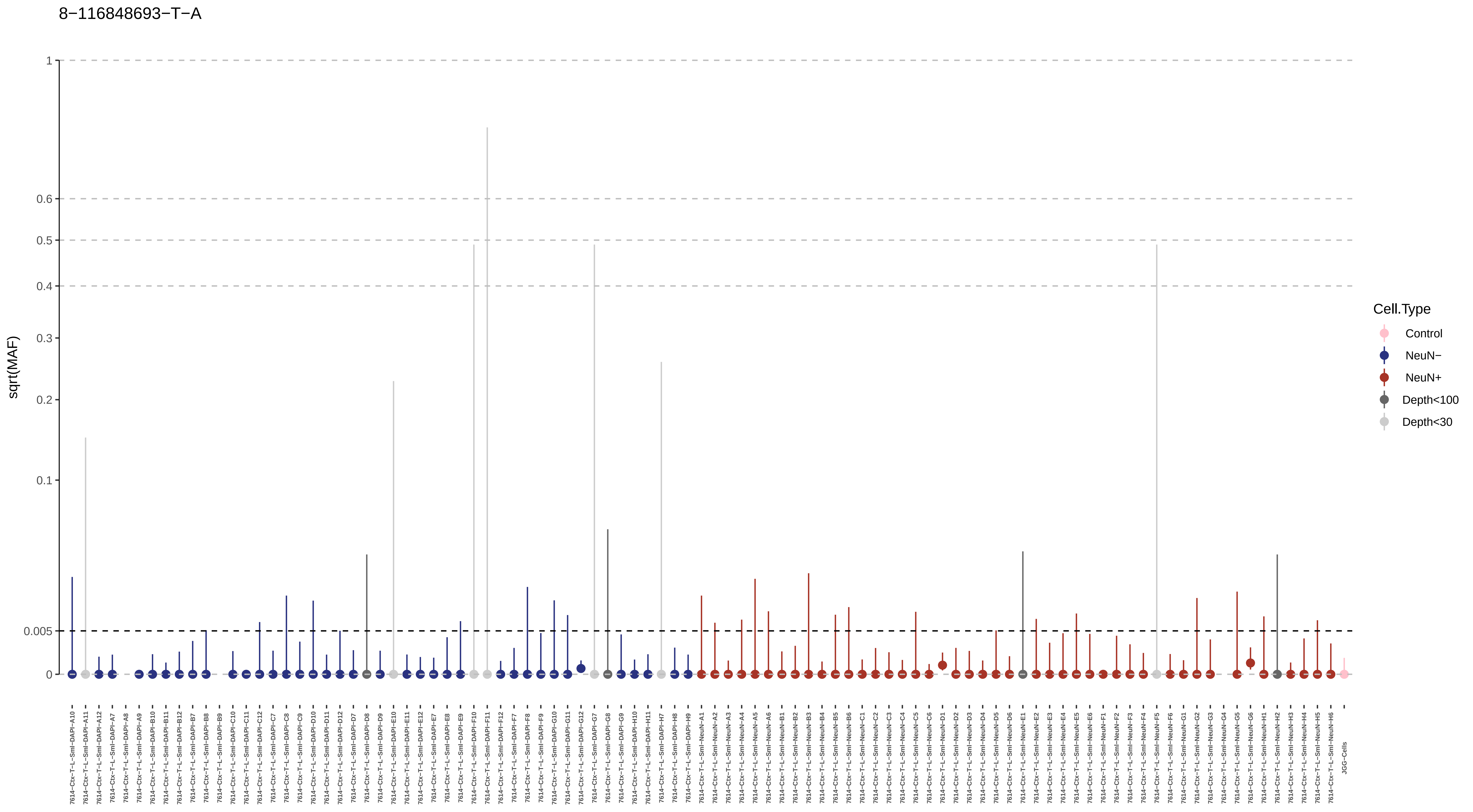

8−116848693−T−A
sqrt(MAF)
1
0.6
0.5
0.4
0.3
0.2
0.1
0.005
0
Cell.Type
Control
NeuN−
NeuN+
Depth<100
Depth<30

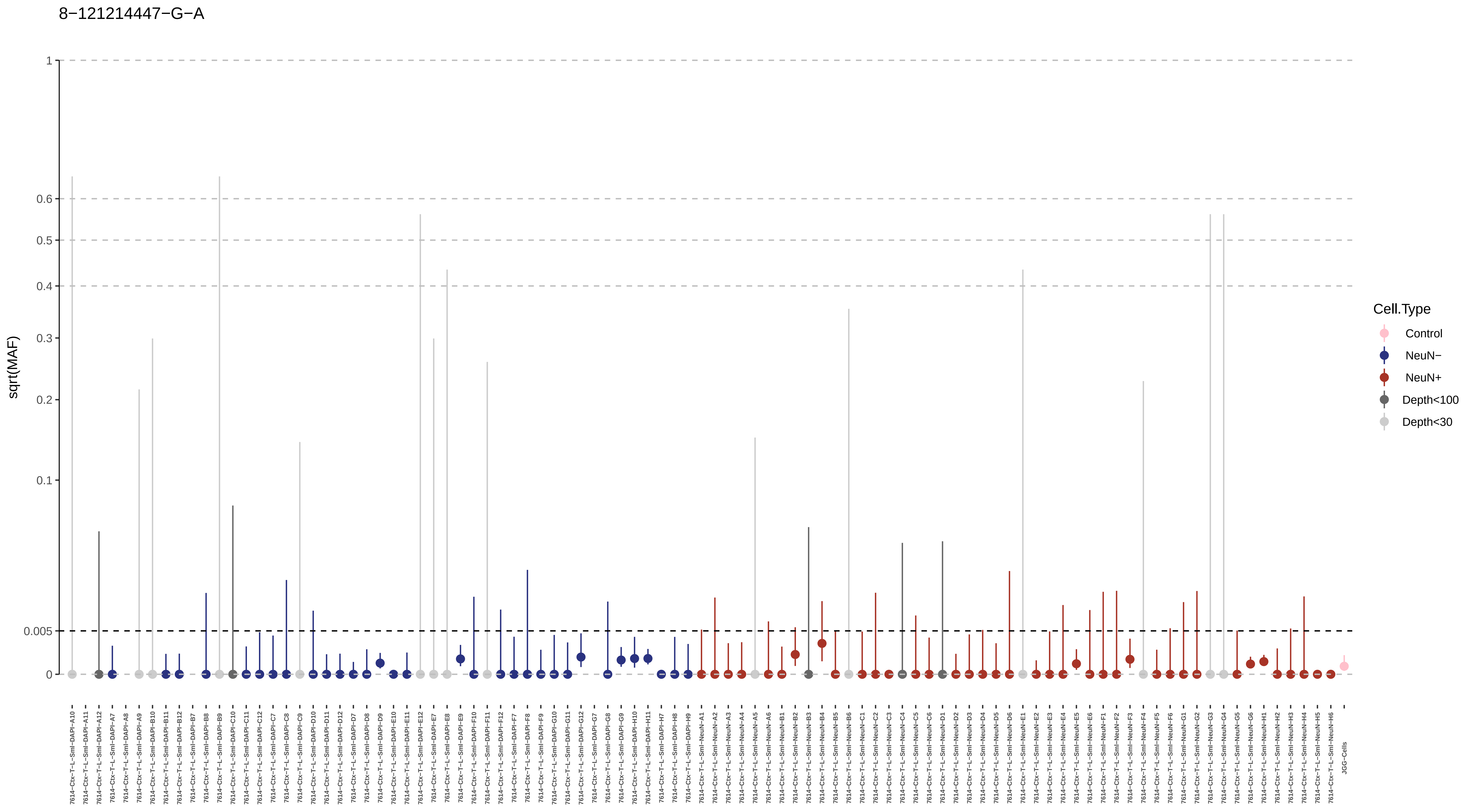
8−121214447−G−A
sqrt(MAF)
1
0.6
0.5
0.4
0.3
0.2
0.1
0.005
0
Cell.Type
Control
NeuN−
NeuN+
Depth<100
Depth<30

8−122651370−C−A

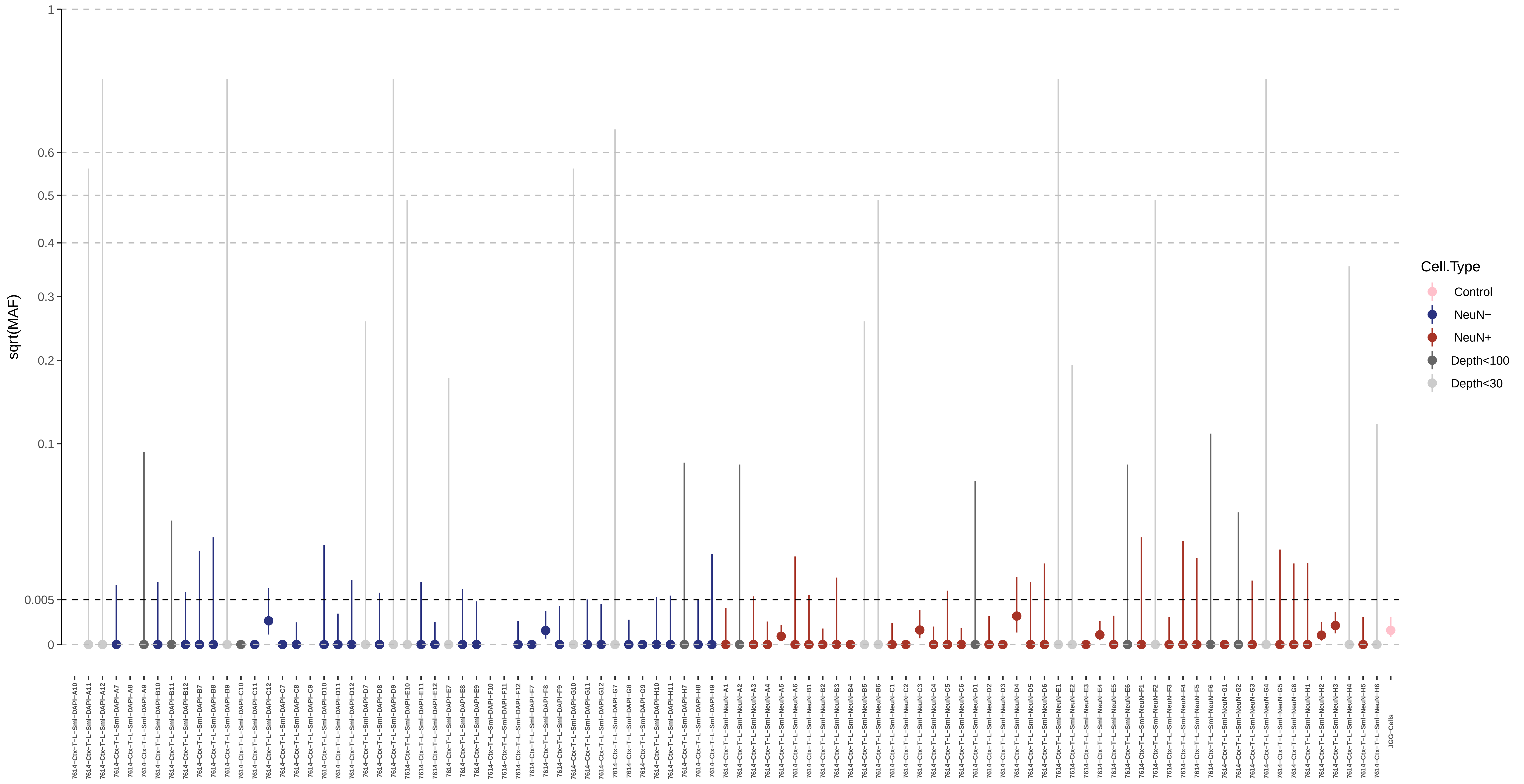

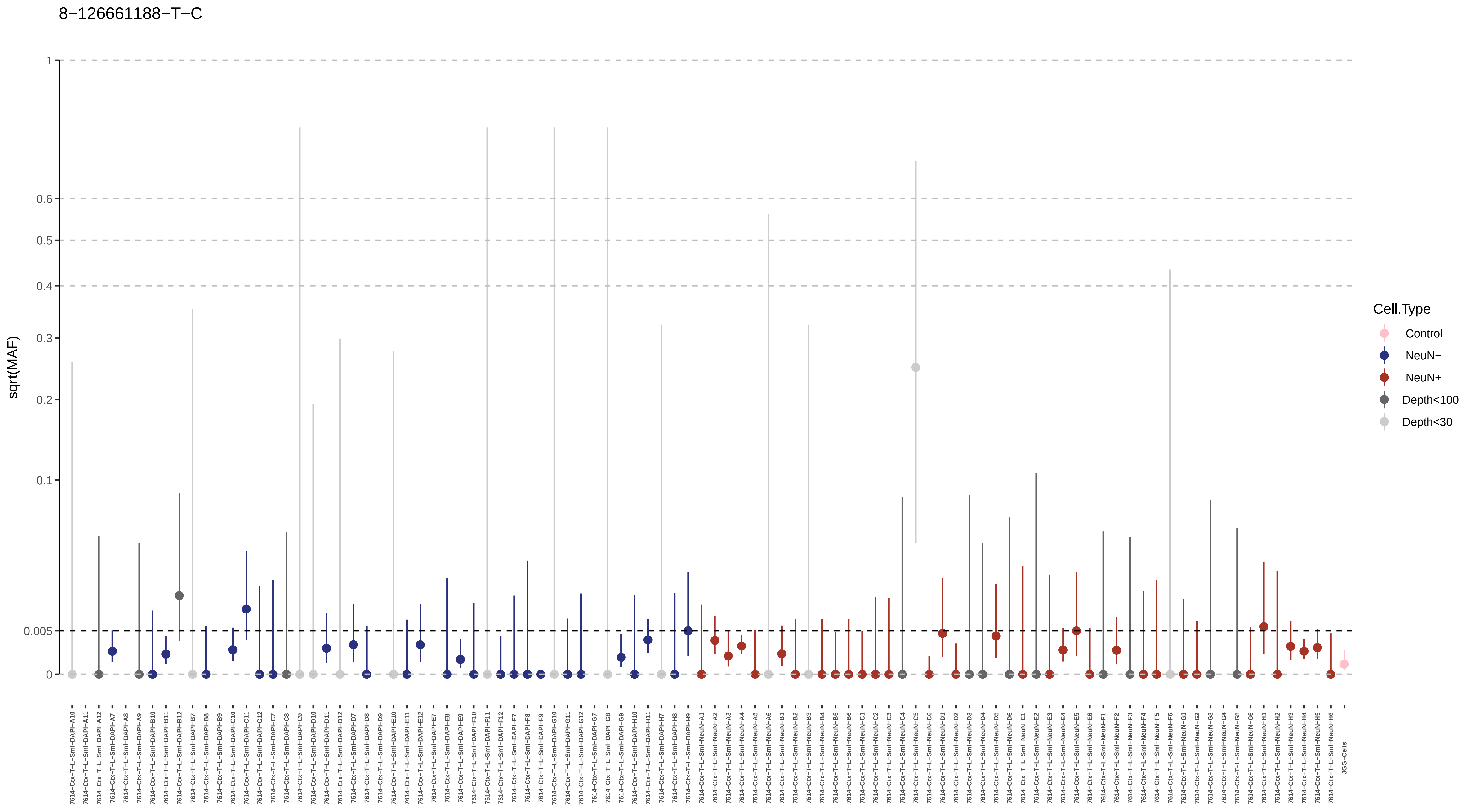
8−126661188−T−C
sqrt(MAF)
1
0.6
0.5
0.4
0.3
0.2
0.1
0.005
0
Cell.Type
Control
NeuN−
NeuN+
Depth<100
Depth<30
7614−Ctx−T−L−Sml−DAPI−A10
7614−Ctx−T−L−Sml−DAPI−A11
7614−Ctx−T−L−Sml−DAPI−A12
7614−Ctx−T−L−Sml−DAPI−A7
7614−Ctx−T−L−Sml−DAPI−A8
7614−Ctx−T−L−Sml−DAPI−A9
7614−Ctx−T−L−Sml−DAPI−B10
7614−Ctx−T−L−Sml−DAPI−B11
7614−Ctx−T−L−Sml−DAPI−B12
7614−Ctx−T−L−Sml−DAPI−B7
7614−Ctx−T−L−Sml−DAPI−B8
7614−Ctx−T−L−Sml−DAPI−B9
7614−Ctx−T−L−Sml−DAPI−C10
7614−Ctx−T−L−Sml−DAPI−C11
7614−Ctx−T−L−Sml−DAPI−C12
7614−Ctx−T−L−Sml−DAPI−C7
7614−Ctx−T−L−Sml−DAPI−C8
7614−Ctx−T−L−Sml−DAPI−C9
7614−Ctx−T−L−Sml−DAPI−D10
7614−Ctx−T−L−Sml−DAPI−D11
7614−Ctx−T−L−Sml−DAPI−D12
7614−Ctx−T−L−Sml−DAPI−D7
7614−Ctx−T−L−Sml−DAPI−D8
7614−Ctx−T−L−Sml−DAPI−D9
7614−Ctx−T−L−Sml−DAPI−E10
7614−Ctx−T−L−Sml−DAPI−E11
7614−Ctx−T−L−Sml−DAPI−E12
7614−Ctx−T−L−Sml−DAPI−E7
7614−Ctx−T−L−Sml−DAPI−E8
7614−Ctx−T−L−Sml−DAPI−E9
7614−Ctx−T−L−Sml−DAPI−F10
7614−Ctx−T−L−Sml−DAPI−F11
7614−Ctx−T−L−Sml−DAPI−F12
7614−Ctx−T−L−Sml−DAPI−F7
7614−Ctx−T−L−Sml−DAPI−F8
7614−Ctx−T−L−Sml−DAPI−F9
7614−Ctx−T−L−Sml−DAPI−G10
7614−Ctx−T−L−Sml−DAPI−G11
7614−Ctx−T−L−Sml−DAPI−G12
7614−Ctx−T−L−Sml−DAPI−G7
7614−Ctx−T−L−Sml−DAPI−G8
7614−Ctx−T−L−Sml−DAPI−G9
7614−Ctx−T−L−Sml−DAPI−H10
7614−Ctx−T−L−Sml−DAPI−H11
7614−Ctx−T−L−Sml−DAPI−H7
7614−Ctx−T−L−Sml−DAPI−H8
7614−Ctx−T−L−Sml−DAPI−H9
7614−Ctx−T−L−Sml−NeuN−A1
7614−Ctx−T−L−Sml−NeuN−A2
7614−Ctx−T−L−Sml−NeuN−A3
7614−Ctx−T−L−Sml−NeuN−A4
7614−Ctx−T−L−Sml−NeuN−A5
7614−Ctx−T−L−Sml−NeuN−A6
7614−Ctx−T−L−Sml−NeuN−B1
7614−Ctx−T−L−Sml−NeuN−B2
7614−Ctx−T−L−Sml−NeuN−B3
7614−Ctx−T−L−Sml−NeuN−B4
7614−Ctx−T−L−Sml−NeuN−B5
7614−Ctx−T−L−Sml−NeuN−B6
7614−Ctx−T−L−Sml−NeuN−C1
7614−Ctx−T−L−Sml−NeuN−C2
7614−Ctx−T−L−Sml−NeuN−C3
7614−Ctx−T−L−Sml−NeuN−C4
7614−Ctx−T−L−Sml−NeuN−C5
7614−Ctx−T−L−Sml−NeuN−C6
7614−Ctx−T−L−Sml−NeuN−D1
7614−Ctx−T−L−Sml−NeuN−D2
7614−Ctx−T−L−Sml−NeuN−D3
7614−Ctx−T−L−Sml−NeuN−D4
7614−Ctx−T−L−Sml−NeuN−D5
7614−Ctx−T−L−Sml−NeuN−D6
7614−Ctx−T−L−Sml−NeuN−E1
7614−Ctx−T−L−Sml−NeuN−E2
7614−Ctx−T−L−Sml−NeuN−E3
7614−Ctx−T−L−Sml−NeuN−E4
7614−Ctx−T−L−Sml−NeuN−E5
7614−Ctx−T−L−Sml−NeuN−E6
7614−Ctx−T−L−Sml−NeuN−F1
7614−Ctx−T−L−Sml−NeuN−F2
7614−Ctx−T−L−Sml−NeuN−F3
7614−Ctx−T−L−Sml−NeuN−F4
7614−Ctx−T−L−Sml−NeuN−F5
7614−Ctx−T−L−Sml−NeuN−F6
7614−Ctx−T−L−Sml−NeuN−G1
7614−Ctx−T−L−Sml−NeuN−G2
7614−Ctx−T−L−Sml−NeuN−G3
7614−Ctx−T−L−Sml−NeuN−G4
7614−Ctx−T−L−Sml−NeuN−G5
7614−Ctx−T−L−Sml−NeuN−G6
7614−Ctx−T−L−Sml−NeuN−H1
7614−Ctx−T−L−Sml−NeuN−H2
7614−Ctx−T−L−Sml−NeuN−H3
7614−Ctx−T−L−Sml−NeuN−H4
7614−Ctx−T−L−Sml−NeuN−H5
7614−Ctx−T−L−Sml−NeuN−H6
JGG−Cells

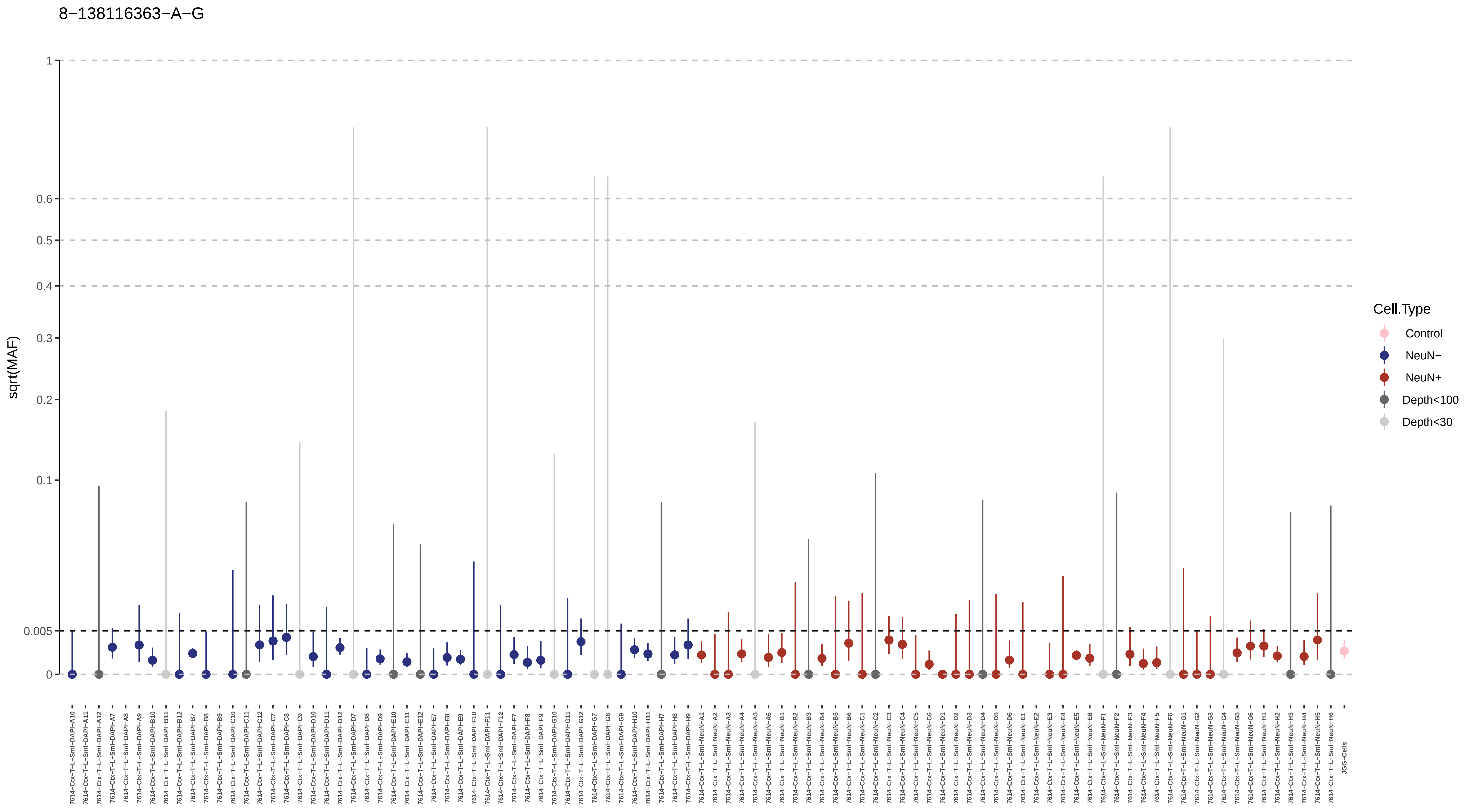

8−138116363−A−G
sqrt(MAF)
1
0.6
0.5
0.4
0.3
0.2
0.1
0.005
0
Cell.Type
Control
NeuN−
NeuN+
Depth<100
Depth<30

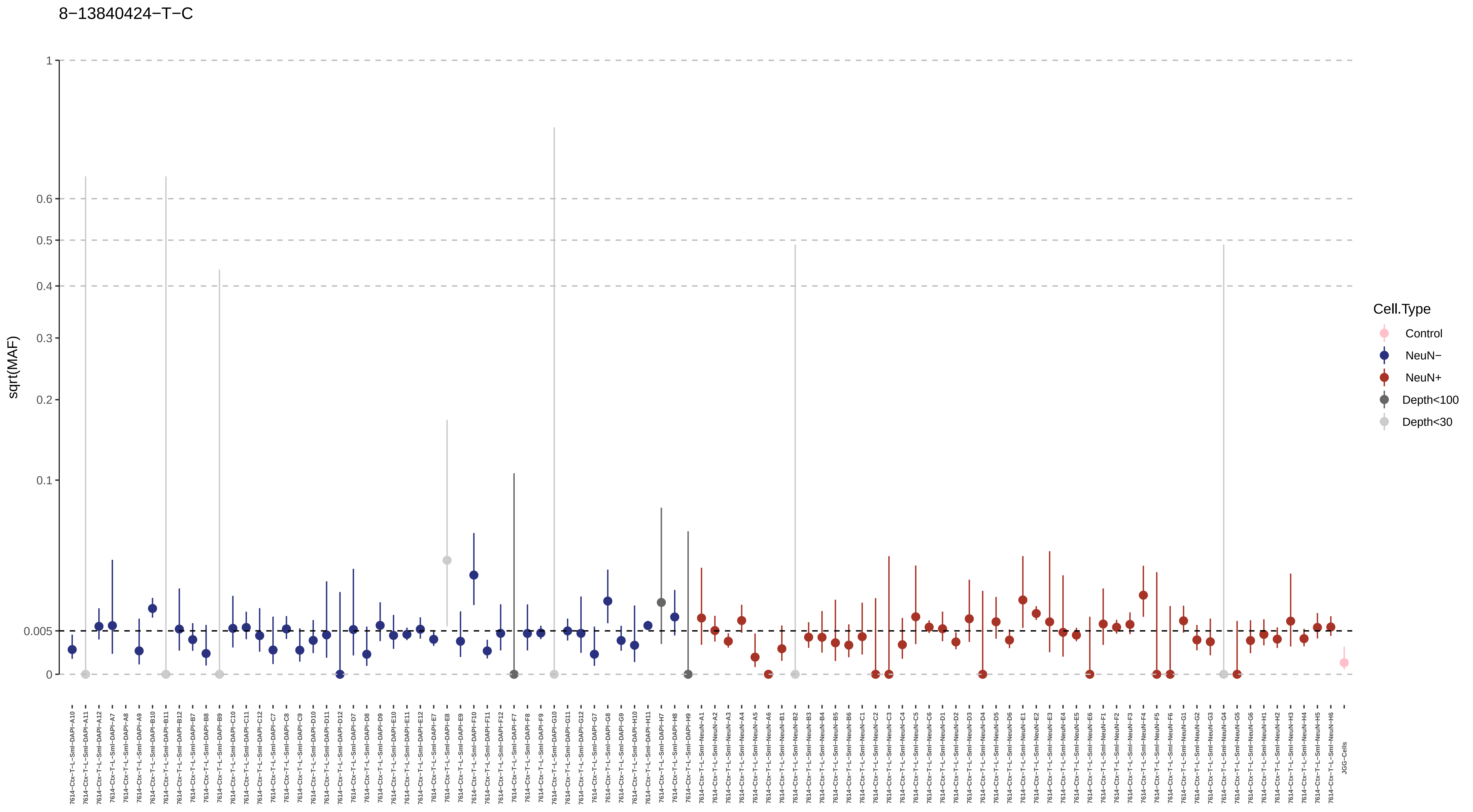
8−13840424−T−C
sqrt(MAF)
1
0.6
0.5
0.4
0.3
0.2
0.1
0.005
0
Cell.Type
Control
NeuN−
NeuN+
Depth<100
Depth<30
7614−Ctx−T−L−Sml−DAPI−A10
7614−Ctx−T−L−Sml−DAPI−A11
7614−Ctx−T−L−Sml−DAPI−A12
7614−Ctx−T−L−Sml−DAPI−A7
7614−Ctx−T−L−Sml−DAPI−A8
7614−Ctx−T−L−Sml−DAPI−A9
7614−Ctx−T−L−Sml−DAPI−B10
7614−Ctx−T−L−Sml−DAPI−B11
7614−Ctx−T−L−Sml−DAPI−B12
7614−Ctx−T−L−Sml−DAPI−B7
7614−Ctx−T−L−Sml−DAPI−B8
7614−Ctx−T−L−Sml−DAPI−B9
7614−Ctx−T−L−Sml−DAPI−C10
7614−Ctx−T−L−Sml−DAPI−C11
7614−Ctx−T−L−Sml−DAPI−C12
7614−Ctx−T−L−Sml−DAPI−C7
7614−Ctx−T−L−Sml−DAPI−C8
7614−Ctx−T−L−Sml−DAPI−C9
7614−Ctx−T−L−Sml−DAPI−D10
7614−Ctx−T−L−Sml−DAPI−D11
7614−Ctx−T−L−Sml−DAPI−D12
7614−Ctx−T−L−Sml−DAPI−D7
7614−Ctx−T−L−Sml−DAPI−D8
7614−Ctx−T−L−Sml−DAPI−D9
7614−Ctx−T−L−Sml−DAPI−E10
7614−Ctx−T−L−Sml−DAPI−E11
7614−Ctx−T−L−Sml−DAPI−E12
7614−Ctx−T−L−Sml−DAPI−E7
7614−Ctx−T−L−Sml−DAPI−E8
7614−Ctx−T−L−Sml−DAPI−E9
7614−Ctx−T−L−Sml−DAPI−F10
7614−Ctx−T−L−Sml−DAPI−F11
7614−Ctx−T−L−Sml−DAPI−F12
7614−Ctx−T−L−Sml−DAPI−F7
7614−Ctx−T−L−Sml−DAPI−F8
7614−Ctx−T−L−Sml−DAPI−F9
7614−Ctx−T−L−Sml−DAPI−G10
7614−Ctx−T−L−Sml−DAPI−G11
7614−Ctx−T−L−Sml−DAPI−G12
7614−Ctx−T−L−Sml−DAPI−G7
7614−Ctx−T−L−Sml−DAPI−G8
7614−Ctx−T−L−Sml−DAPI−G9
7614−Ctx−T−L−Sml−DAPI−H10
7614−Ctx−T−L−Sml−DAPI−H11
7614−Ctx−T−L−Sml−DAPI−H7
7614−Ctx−T−L−Sml−DAPI−H8
7614−Ctx−T−L−Sml−DAPI−H9
7614−Ctx−T−L−Sml−NeuN−A1
7614−Ctx−T−L−Sml−NeuN−A2
7614−Ctx−T−L−Sml−NeuN−A3
7614−Ctx−T−L−Sml−NeuN−A4
7614−Ctx−T−L−Sml−NeuN−A5
7614−Ctx−T−L−Sml−NeuN−A6
7614−Ctx−T−L−Sml−NeuN−B1
7614−Ctx−T−L−Sml−NeuN−B2
7614−Ctx−T−L−Sml−NeuN−B3
7614−Ctx−T−L−Sml−NeuN−B4
7614−Ctx−T−L−Sml−NeuN−B5
7614−Ctx−T−L−Sml−NeuN−B6
7614−Ctx−T−L−Sml−NeuN−C1
7614−Ctx−T−L−Sml−NeuN−C2
7614−Ctx−T−L−Sml−NeuN−C3
7614−Ctx−T−L−Sml−NeuN−C4
7614−Ctx−T−L−Sml−NeuN−C5
7614−Ctx−T−L−Sml−NeuN−C6
7614−Ctx−T−L−Sml−NeuN−D1
7614−Ctx−T−L−Sml−NeuN−D2
7614−Ctx−T−L−Sml−NeuN−D3
7614−Ctx−T−L−Sml−NeuN−D4
7614−Ctx−T−L−Sml−NeuN−D5
7614−Ctx−T−L−Sml−NeuN−D6
7614−Ctx−T−L−Sml−NeuN−E1
7614−Ctx−T−L−Sml−NeuN−E2
7614−Ctx−T−L−Sml−NeuN−E3
7614−Ctx−T−L−Sml−NeuN−E4
7614−Ctx−T−L−Sml−NeuN−E5
7614−Ctx−T−L−Sml−NeuN−E6
7614−Ctx−T−L−Sml−NeuN−F1
7614−Ctx−T−L−Sml−NeuN−F2
7614−Ctx−T−L−Sml−NeuN−F3
7614−Ctx−T−L−Sml−NeuN−F4
7614−Ctx−T−L−Sml−NeuN−F5
7614−Ctx−T−L−Sml−NeuN−F6
7614−Ctx−T−L−Sml−NeuN−G1
7614−Ctx−T−L−Sml−NeuN−G2
7614−Ctx−T−L−Sml−NeuN−G3
7614−Ctx−T−L−Sml−NeuN−G4
7614−Ctx−T−L−Sml−NeuN−G5
7614−Ctx−T−L−Sml−NeuN−G6
7614−Ctx−T−L−Sml−NeuN−H1
7614−Ctx−T−L−Sml−NeuN−H2
7614−Ctx−T−L−Sml−NeuN−H3
7614−Ctx−T−L−Sml−NeuN−H4
7614−Ctx−T−L−Sml−NeuN−H5
7614−Ctx−T−L−Sml−NeuN−H6
JGG-Cells

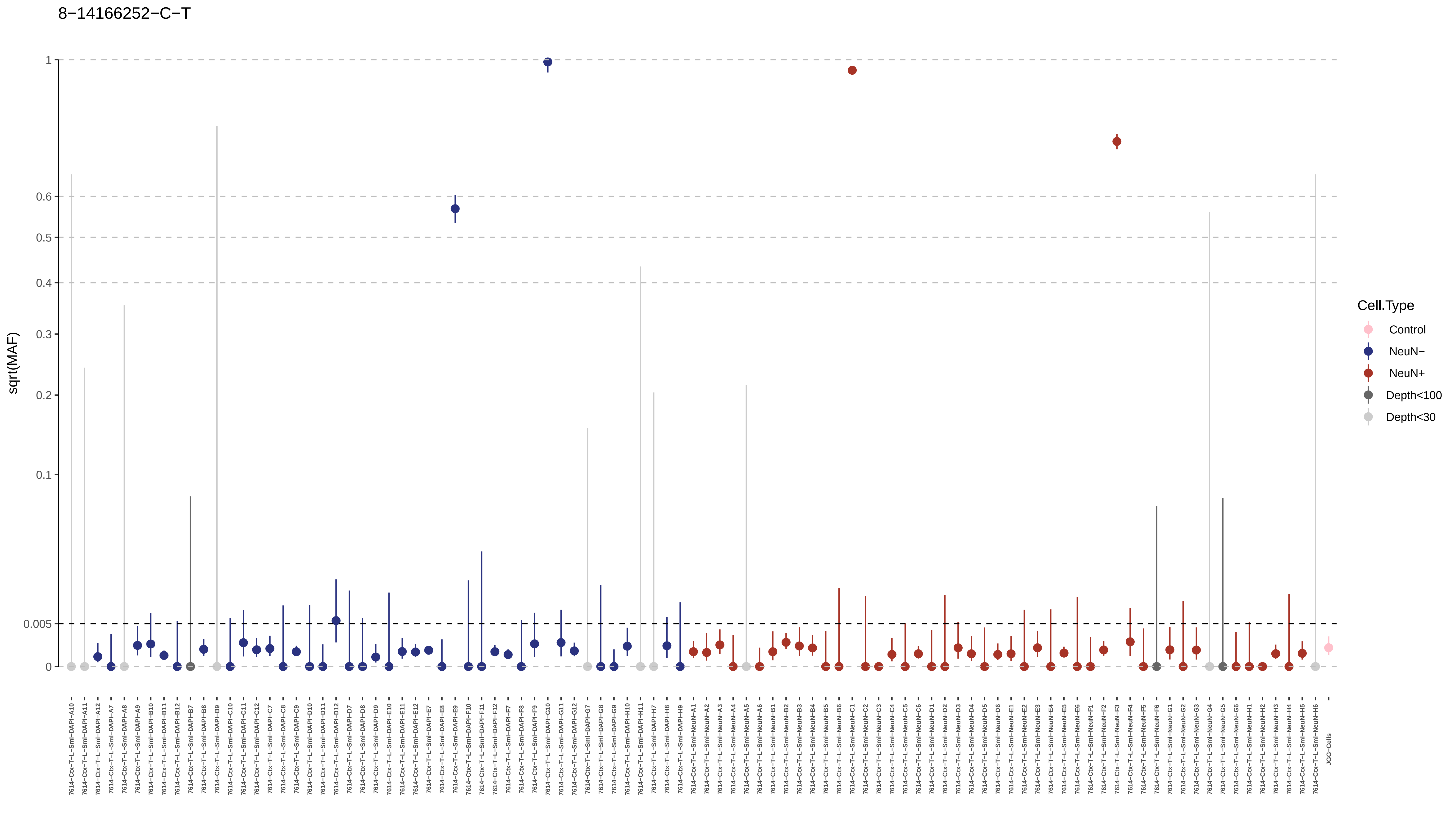
8−14166252−C−T
sqrt(MAF)
1
0.6
0.5
0.4
0.3
0.2
0.1
0.005
0
Cell.Type
Control
NeuN−
NeuN+
Depth<100
Depth<30

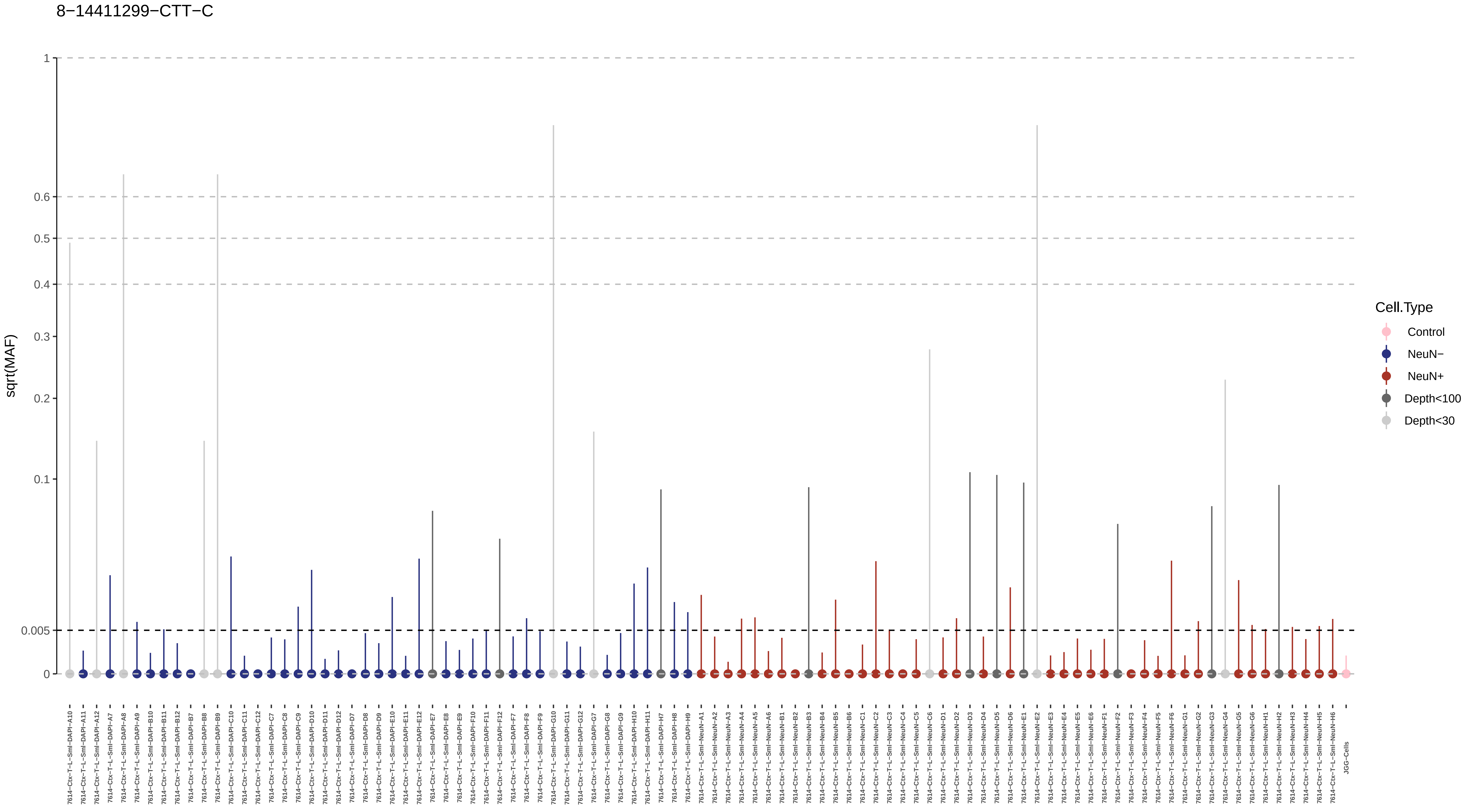
8−14411299−CTT−C
sqrt(MAF)
1
0.6
0.5
0.4
0.3
0.2
0.1
0.005
0
Cell.Type
Control
NeuN−
NeuN+
Depth<100
Depth<30

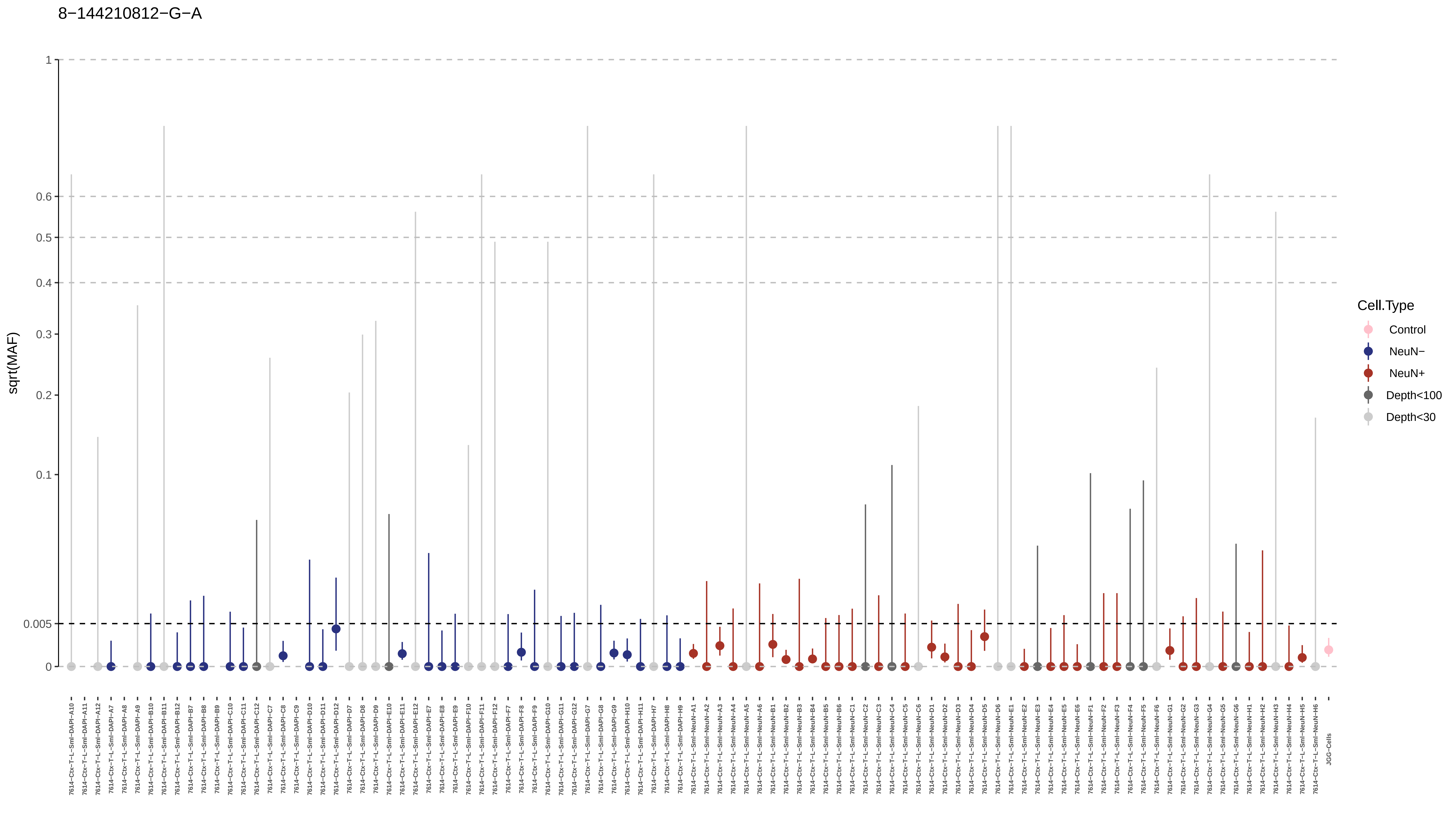
8−144210812−G−A
sqrt(MAF)
1
0.6
0.5
0.4
0.3
0.2
0.1
0.005
0
Cell.Type
Control
NeuN−
NeuN+
Depth<100
Depth<30

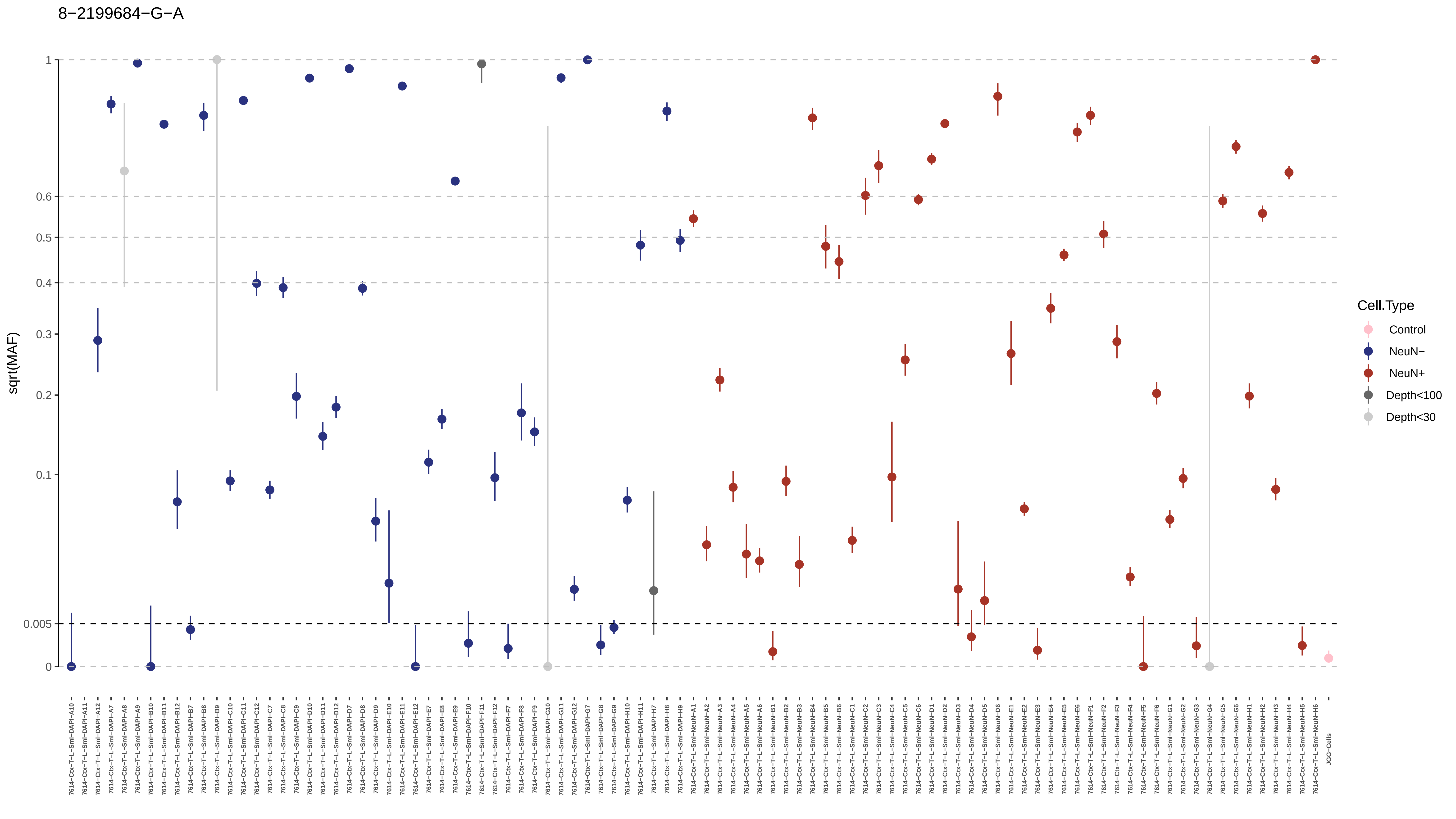

8−2199684−G−A
sqrt(MAF)
1
0.6
0.5
0.4
0.3
0.2
0.1
0.005
0
Cell.Type
Control
NeuN−
NeuN+
Depth<100
Depth<30
7614−Ctx−T−L−Sml−DAPI−A10
7614−Ctx−T−L−Sml−DAPI−A11
7614−Ctx−T−L−Sml−DAPI−A12
7614−Ctx−T−L−Sml−DAPI−A7
7614−Ctx−T−L−Sml−DAPI−A8
7614−Ctx−T−L−Sml−DAPI−A9
7614−Ctx−T−L−Sml−DAPI−B10
7614−Ctx−T−L−Sml−DAPI−B11
7614−Ctx−T−L−Sml−DAPI−B12
7614−Ctx−T−L−Sml−DAPI−B7
7614−Ctx−T−L−Sml−DAPI−B8
7614−Ctx−T−L−Sml−DAPI−B9
7614−Ctx−T−L−Sml−DAPI−C10
7614−Ctx−T−L−Sml−DAPI−C11
7614−Ctx−T−L−Sml−DAPI−C12
7614−Ctx−T−L−Sml−DAPI−C7
7614−Ctx−T−L−Sml−DAPI−C8
7614−Ctx−T−L−Sml−DAPI−C9
7614−Ctx−T−L−Sml−DAPI−D10
7614−Ctx−T−L−Sml−DAPI−D11
7614−Ctx−T−L−Sml−DAPI−D12
7614−Ctx−T−L−Sml−DAPI−D7
7614−Ctx−T−L−Sml−DAPI−D8
7614−Ctx−T−L−Sml−DAPI−D9
7614−Ctx−T−L−Sml−DAPI−E10
7614−Ctx−T−L−Sml−DAPI−E11
7614−Ctx−T−L−Sml−DAPI−E12
7614−Ctx−T−L−Sml−DAPI−E7
7614−Ctx−T−L−Sml−DAPI−E8
7614−Ctx−T−L−Sml−DAPI−E9
7614−Ctx−T−L−Sml−DAPI−F10
7614−Ctx−T−L−Sml−DAPI−F11
7614−Ctx−T−L−Sml−DAPI−F12
7614−Ctx−T−L−Sml−DAPI−F7
7614−Ctx−T−L−Sml−DAPI−F8
7614−Ctx−T−L−Sml−DAPI−F9
7614−Ctx−T−L−Sml−DAPI−G10
7614−Ctx−T−L−Sml−DAPI−G11
7614−Ctx−T−L−Sml−DAPI−G12
7614−Ctx−T−L−Sml−DAPI−G7
7614−Ctx−T−L−Sml−DAPI−G8
7614−Ctx−T−L−Sml−DAPI−G9
7614−Ctx−T−L−Sml−DAPI−H10
7614−Ctx−T−L−Sml−DAPI−H11
7614−Ctx−T−L−Sml−DAPI−H7
7614−Ctx−T−L−Sml−DAPI−H8
7614−Ctx−T−L−Sml−DAPI−H9
7614−Ctx−T−L−Sml−NeuN−A1
7614−Ctx−T−L−Sml−NeuN−A2
7614−Ctx−T−L−Sml−NeuN−A3
7614−Ctx−T−L−Sml−NeuN−A4
7614−Ctx−T−L−Sml−NeuN−A5
7614−Ctx−T−L−Sml−NeuN−A6
7614−Ctx−T−L−Sml−NeuN−B1
7614−Ctx−T−L−Sml−NeuN−B2
7614−Ctx−T−L−Sml−NeuN−B3
7614−Ctx−T−L−Sml−NeuN−B4
7614−Ctx−T−L−Sml−NeuN−B5
7614−Ctx−T−L−Sml−NeuN−B6
7614−Ctx−T−L−Sml−NeuN−C1
7614−Ctx−T−L−Sml−NeuN−C2
7614−Ctx−T−L−Sml−NeuN−C3
7614−Ctx−T−L−Sml−NeuN−C4
7614−Ctx−T−L−Sml−NeuN−C5
7614−Ctx−T−L−Sml−NeuN−C6
7614−Ctx−T−L−Sml−NeuN−D1
7614−Ctx−T−L−Sml−NeuN−D2
7614−Ctx−T−L−Sml−NeuN−D3
7614−Ctx−T−L−Sml−NeuN−D4
7614−Ctx−T−L−Sml−NeuN−D5
7614−Ctx−T−L−Sml−NeuN−D6
7614−Ctx−T−L−Sml−NeuN−E1
7614−Ctx−T−L−Sml−NeuN−E2
7614−Ctx−T−L−Sml−NeuN−E3
7614−Ctx−T−L−Sml−NeuN−E4
7614−Ctx−T−L−Sml−NeuN−E5
7614−Ctx−T−L−Sml−NeuN−E6
7614−Ctx−T−L−Sml−NeuN−F1
7614−Ctx−T−L−Sml−NeuN−F2
7614−Ctx−T−L−Sml−NeuN−F3
7614−Ctx−T−L−Sml−NeuN−F4
7614−Ctx−T−L−Sml−NeuN−F5
7614−Ctx−T−L−Sml−NeuN−F6
7614−Ctx−T−L−Sml−NeuN−G1
7614−Ctx−T−L−Sml−NeuN−G2
7614−Ctx−T−L−Sml−NeuN−G3
7614−Ctx−T−L−Sml−NeuN−G4
7614−Ctx−T−L−Sml−NeuN−G5
7614−Ctx−T−L−Sml−NeuN−G6
7614−Ctx−T−L−Sml−NeuN−H1
7614−Ctx−T−L−Sml−NeuN−H2
7614−Ctx−T−L−Sml−NeuN−H3
7614−Ctx−T−L−Sml−NeuN−H4
7614−Ctx−T−L−Sml−NeuN−H5
7614−Ctx−T−L−Sml−NeuN−H6
JGG−Cells

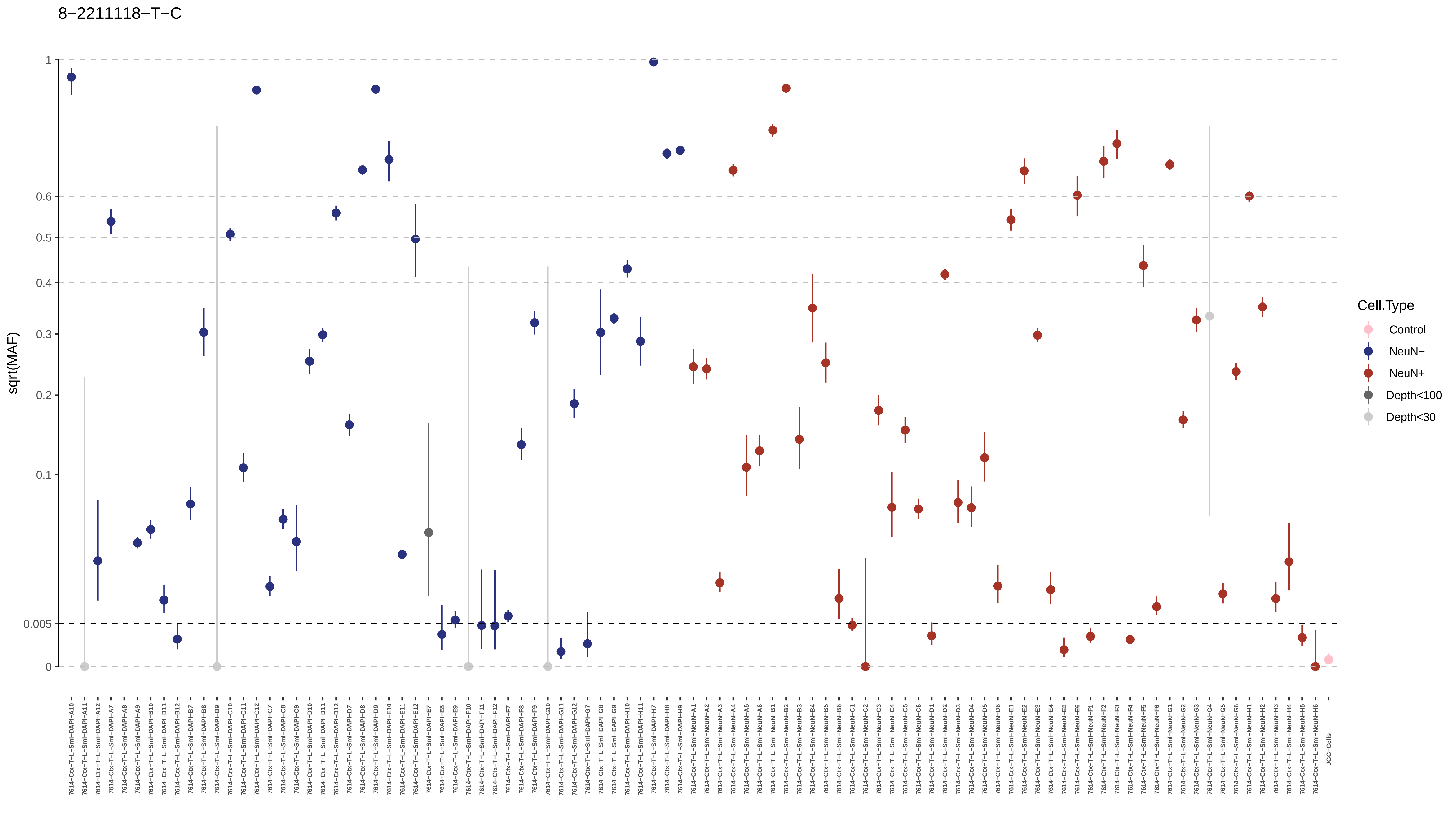
8−2211118−T−C
1
0.6
0.5
0.4
0.3
0.2
0.1
0.005
0
sqrt(MAF)
Cell.Type
Control
NeuN−
NeuN+
Depth<100
Depth<30

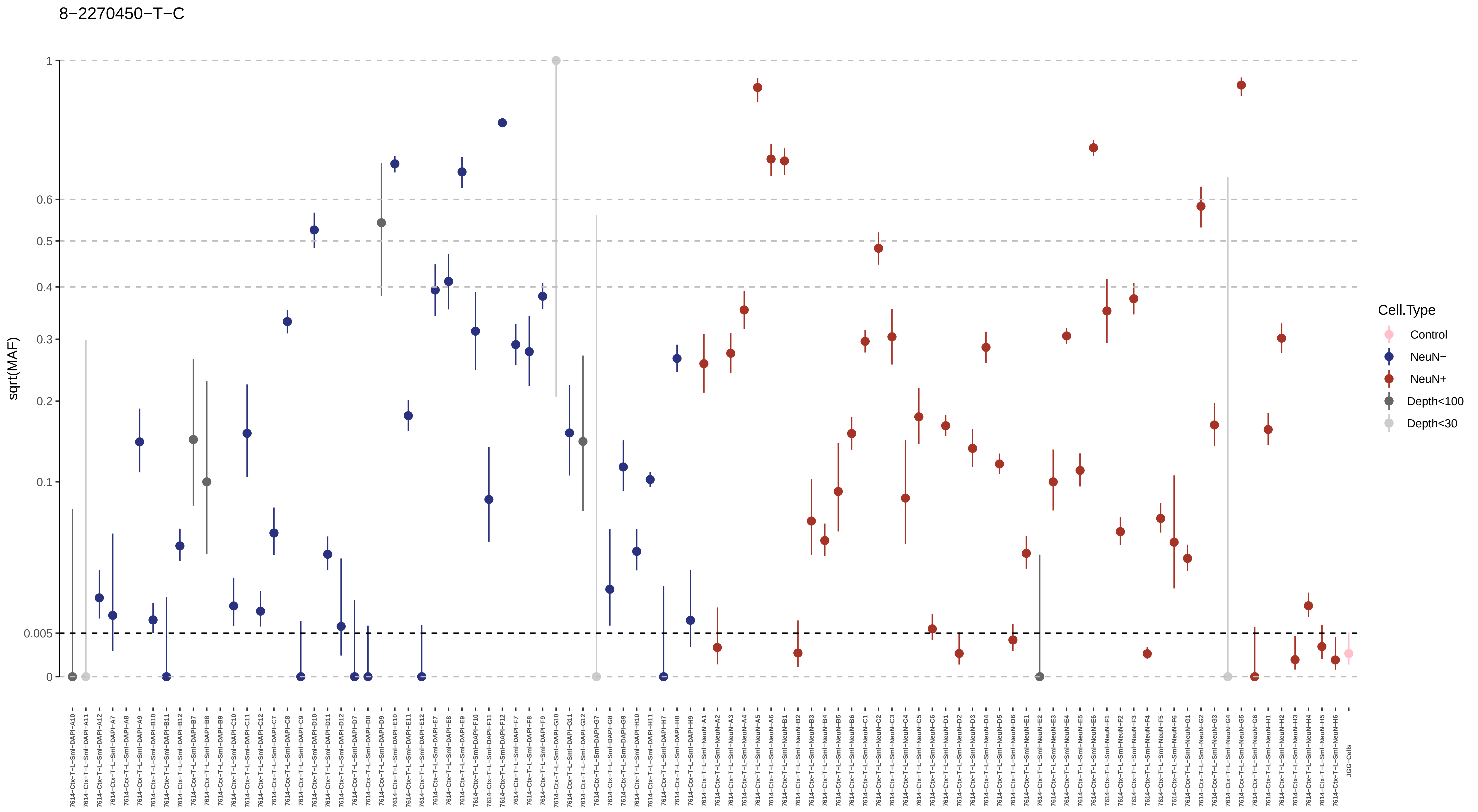
8−2270450−T−C
sqrt(MAF)
1
0.6
0.5
0.4
0.3
0.2
0.1
0.005
0
Cell.Type
Control
NeuN−
NeuN+
Depth<100
Depth<30
7614−Ctx−T−L−Sml−DAPI−A10
7614−Ctx−T−L−Sml−DAPI−A11
7614−Ctx−T−L−Sml−DAPI−A12
7614−Ctx−T−L−Sml−DAPI−A7
7614−Ctx−T−L−Sml−DAPI−A8
7614−Ctx−T−L−Sml−DAPI−A9
7614−Ctx−T−L−Sml−DAPI−B10
7614−Ctx−T−L−Sml−DAPI−B11
7614−Ctx−T−L−Sml−DAPI−B12
7614−Ctx−T−L−Sml−DAPI−B7
7614−Ctx−T−L−Sml−DAPI−B8
7614−Ctx−T−L−Sml−DAPI−B9
7614−Ctx−T−L−Sml−DAPI−C10
7614−Ctx−T−L−Sml−DAPI−C11
7614−Ctx−T−L−Sml−DAPI−C12
7614−Ctx−T−L−Sml−DAPI−C7
7614−Ctx−T−L−Sml−DAPI−C8
7614−Ctx−T−L−Sml−DAPI−C9
7614−Ctx−T−L−Sml−DAPI−D10
7614−Ctx−T−L−Sml−DAPI−D11
7614−Ctx−T−L−Sml−DAPI−D12
7614−Ctx−T−L−Sml−DAPI−D7
7614−Ctx−T−L−Sml−DAPI−D8
7614−Ctx−T−L−Sml−DAPI−D9
7614−Ctx−T−L−Sml−DAPI−E10
7614−Ctx−T−L−Sml−DAPI−E11
7614−Ctx−T−L−Sml−DAPI−E12
7614−Ctx−T−L−Sml−DAPI−E7
7614−Ctx−T−L−Sml−DAPI−E8
7614−Ctx−T−L−Sml−DAPI−E9
7614−Ctx−T−L−Sml−DAPI−F10
7614−Ctx−T−L−Sml−DAPI−F11
7614−Ctx−T−L−Sml−DAPI−F12
7614−Ctx−T−L−Sml−DAPI−F7
7614−Ctx−T−L−Sml−DAPI−F8
7614−Ctx−T−L−Sml−DAPI−F9
7614−Ctx−T−L−Sml−DAPI−G10
7614−Ctx−T−L−Sml−DAPI−G11
7614−Ctx−T−L−Sml−DAPI−G12
7614−Ctx−T−L−Sml−DAPI−G7
7614−Ctx−T−L−Sml−DAPI−G8
7614−Ctx−T−L−Sml−DAPI−G9
7614−Ctx−T−L−Sml−DAPI−H10
7614−Ctx−T−L−Sml−DAPI−H11
7614−Ctx−T−L−Sml−DAPI−H7
7614−Ctx−T−L−Sml−DAPI−H8
7614−Ctx−T−L−Sml−DAPI−H9
7614−Ctx−T−L−Sml−NeuN−A1
7614−Ctx−T−L−Sml−NeuN−A2
7614−Ctx−T−L−Sml−NeuN−A3
7614−Ctx−T−L−Sml−NeuN−A4
7614−Ctx−T−L−Sml−NeuN−A5
7614−Ctx−T−L−Sml−NeuN−A6
7614−Ctx−T−L−Sml−NeuN−B1
7614−Ctx−T−L−Sml−NeuN−B2
7614−Ctx−T−L−Sml−NeuN−B3
7614−Ctx−T−L−Sml−NeuN−B4
7614−Ctx−T−L−Sml−NeuN−B5
7614−Ctx−T−L−Sml−NeuN−B6
7614−Ctx−T−L−Sml−NeuN−C1
7614−Ctx−T−L−Sml−NeuN−C2
7614−Ctx−T−L−Sml−NeuN−C3
7614−Ctx−T−L−Sml−NeuN−C4
7614−Ctx−T−L−Sml−NeuN−C5
7614−Ctx−T−L−Sml−NeuN−C6
7614−Ctx−T−L−Sml−NeuN−D1
7614−Ctx−T−L−Sml−NeuN−D2
7614−Ctx−T−L−Sml−NeuN−D3
7614−Ctx−T−L−Sml−NeuN−D4
7614−Ctx−T−L−Sml−NeuN−D5
7614−Ctx−T−L−Sml−NeuN−D6
7614−Ctx−T−L−Sml−NeuN−E1
7614−Ctx−T−L−Sml−NeuN−E2
7614−Ctx−T−L−Sml−NeuN−E3
7614−Ctx−T−L−Sml−NeuN−E4
7614−Ctx−T−L−Sml−NeuN−E5
7614−Ctx−T−L−Sml−NeuN−E6
7614−Ctx−T−L−Sml−NeuN−F1
7614−Ctx−T−L−Sml−NeuN−F2
7614−Ctx−T−L−Sml−NeuN−F3
7614−Ctx−T−L−Sml−NeuN−F4
7614−Ctx−T−L−Sml−NeuN−F5
7614−Ctx−T−L−Sml−NeuN−F6
7614−Ctx−T−L−Sml−NeuN−G1
7614−Ctx−T−L−Sml−NeuN−G2
7614−Ctx−T−L−Sml−NeuN−G3
7614−Ctx−T−L−Sml−NeuN−G4
7614−Ctx−T−L−Sml−NeuN−G5
7614−Ctx−T−L−Sml−NeuN−G6
7614−Ctx−T−L−Sml−NeuN−H1
7614−Ctx−T−L−Sml−NeuN−H2
7614−Ctx−T−L−Sml−NeuN−H3
7614−Ctx−T−L−Sml−NeuN−H4
7614−Ctx−T−L−Sml−NeuN−H5
7614−Ctx−T−L−Sml−NeuN−H6
JGG-Cells

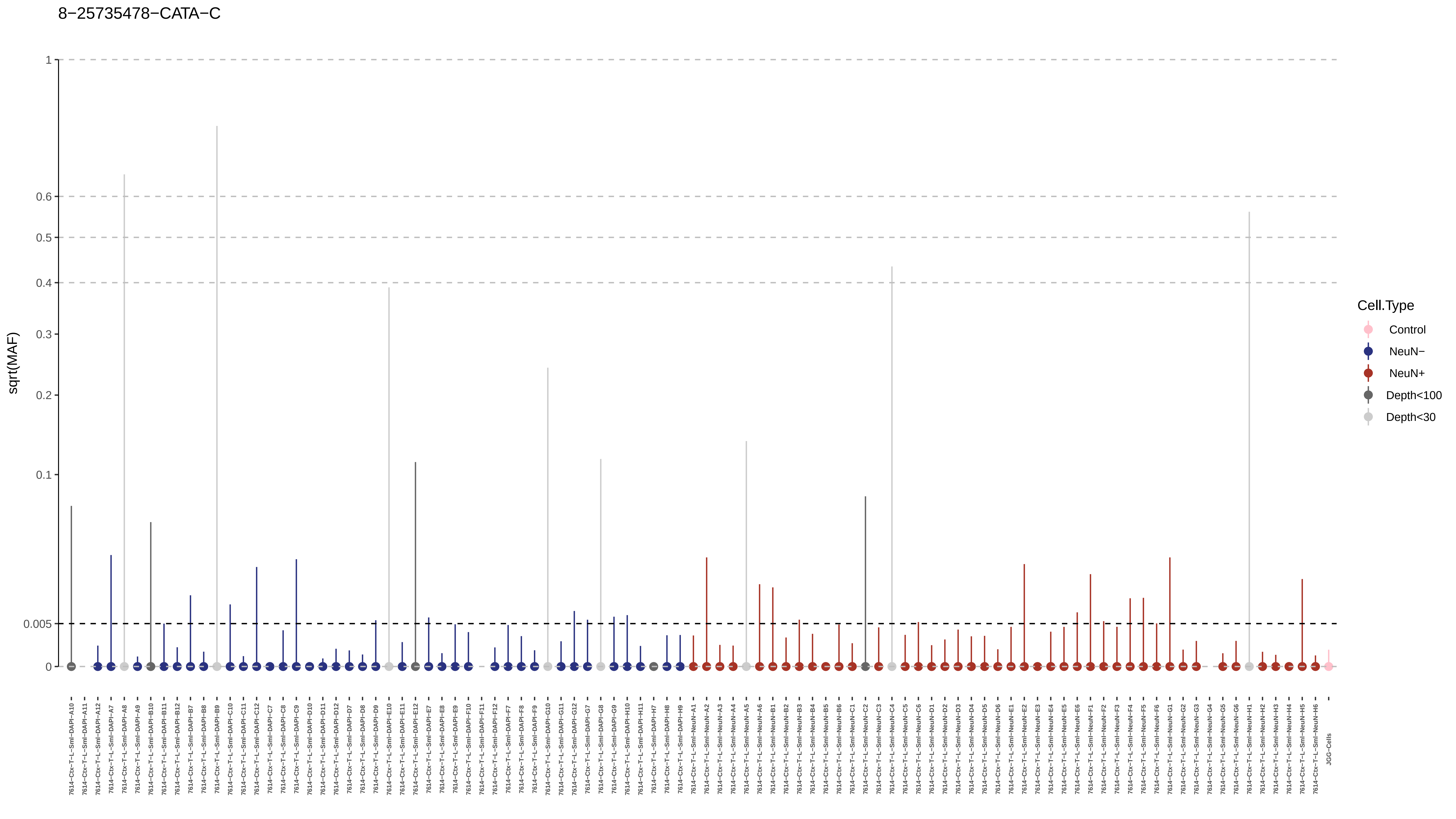
8−25735478−CATA−C
sqrt(MAF)
1
0.6
0.5
0.4
0.3
0.2
0.1
0.005
0
7614−Ctx−T−L−Sml−DAPI−A10
7614−Ctx−T−L−Sml−DAPI−A11
7614−Ctx−T−L−Sml−DAPI−A12
7614−Ctx−T−L−Sml−DAPI−A7
7614−Ctx−T−L−Sml−DAPI−A8
7614−Ctx−T−L−Sml−DAPI−A9
7614−Ctx−T−L−Sml−DAPI−B10
7614−Ctx−T−L−Sml−DAPI−B11
7614−Ctx−T−L−Sml−DAPI−B12
7614−Ctx−T−L−Sml−DAPI−B7
7614−Ctx−T−L−Sml−DAPI−B8
7614−Ctx−T−L−Sml−DAPI−B9
7614−Ctx−T−L−Sml−DAPI−C10
7614−Ctx−T−L−Sml−DAPI−C11
7614−Ctx−T−L−Sml−DAPI−C12
7614−Ctx−T−L−Sml−DAPI−C7
7614−Ctx−T−L−Sml−DAPI−C8
7614−Ctx−T−L−Sml−DAPI−C9
7614−Ctx−T−L−Sml−DAPI−D10
7614−Ctx−T−L−Sml−DAPI−D11
7614−Ctx−T−L−Sml−DAPI−D12
7614−Ctx−T−L−Sml−DAPI−D7
7614−Ctx−T−L−Sml−DAPI−D8
7614−Ctx−T−L−Sml−DAPI−D9
7614−Ctx−T−L−Sml−DAPI−E10
7614−Ctx−T−L−Sml−DAPI−E11
7614−Ctx−T−L−Sml−DAPI−E12
7614−Ctx−T−L−Sml−DAPI−E7
7614−Ctx−T−L−Sml−DAPI−E8
7614−Ctx−T−L−Sml−DAPI−E9
7614−Ctx−T−L−Sml−DAPI−F10
7614−Ctx−T−L−Sml−DAPI−F11
7614−Ctx−T−L−Sml−DAPI−F12
7614−Ctx−T−L−Sml−DAPI−F7
7614−Ctx−T−L−Sml−DAPI−F8
7614−Ctx−T−L−Sml−DAPI−F9
7614−Ctx−T−L−Sml−DAPI−G10
7614−Ctx−T−L−Sml−DAPI−G11
7614−Ctx−T−L−Sml−DAPI−G12
7614−Ctx−T−L−Sml−DAPI−G7
7614−Ctx−T−L−Sml−DAPI−G8
7614−Ctx−T−L−Sml−DAPI−G9
7614−Ctx−T−L−Sml−DAPI−H10
7614−Ctx−T−L−Sml−DAPI−H11
7614−Ctx−T−L−Sml−DAPI−H7
7614−Ctx−T−L−Sml−DAPI−H8
7614−Ctx−T−L−Sml−DAPI−H9
7614−Ctx−T−L−Sml−NeuN−A1
7614−Ctx−T−L−Sml−NeuN−A2
7614−Ctx−T−L−Sml−NeuN−A3
7614−Ctx−T−L−Sml−NeuN−A4
7614−Ctx−T−L−Sml−NeuN−A5
7614−Ctx−T−L−Sml−NeuN−A6
7614−Ctx−T−L−Sml−NeuN−B1
7614−Ctx−T−L−Sml−NeuN−B2
7614−Ctx−T−L−Sml−NeuN−B3
7614−Ctx−T−L−Sml−NeuN−B4
7614−Ctx−T−L−Sml−NeuN−B5
7614−Ctx−T−L−Sml−NeuN−B6
7614−Ctx−T−L−Sml−NeuN−C1
7614−Ctx−T−L−Sml−NeuN−C2
7614−Ctx−T−L−Sml−NeuN−C3
7614−Ctx−T−L−Sml−NeuN−C4
7614−Ctx−T−L−Sml−NeuN−C5
7614−Ctx−T−L−Sml−NeuN−C6
7614−Ctx−T−L−Sml−NeuN−D1
7614−Ctx−T−L−Sml−NeuN−D2
7614−Ctx−T−L−Sml−NeuN−D3
7614−Ctx−T−L−Sml−NeuN−D4
7614−Ctx−T−L−Sml−NeuN−D5
7614−Ctx−T−L−Sml−NeuN−D6
7614−Ctx−T−L−Sml−NeuN−E1
7614−Ctx−T−L−Sml−NeuN−E2
7614−Ctx−T−L−Sml−NeuN−E3
7614−Ctx−T−L−Sml−NeuN−E4
7614−Ctx−T−L−Sml−NeuN−E5
7614−Ctx−T−L−Sml−NeuN−E6
7614−Ctx−T−L−Sml−NeuN−F1
7614−Ctx−T−L−Sml−NeuN−F2
7614−Ctx−T−L−Sml−NeuN−F3
7614−Ctx−T−L−Sml−NeuN−F4
7614−Ctx−T−L−Sml−NeuN−F5
7614−Ctx−T−L−Sml−NeuN−F6
7614−Ctx−T−L−Sml−NeuN−G1
7614−Ctx−T−L−Sml−NeuN−G2
7614−Ctx−T−L−Sml−NeuN−G3
7614−Ctx−T−L−Sml−NeuN−G4
7614−Ctx−T−L−Sml−NeuN−G5
7614−Ctx−T−L−Sml−NeuN−G6
7614−Ctx−T−L−Sml−NeuN−H1
7614−Ctx−T−L−Sml−NeuN−H2
7614−Ctx−T−L−Sml−NeuN−H3
7614−Ctx−T−L−Sml−NeuN−H4
7614−Ctx−T−L−Sml−NeuN−H5
7614−Ctx−T−L−Sml−NeuN−H6
JGG-Cells
Cell.Type
Control
NeuN−
NeuN+
Depth<100
Depth<30

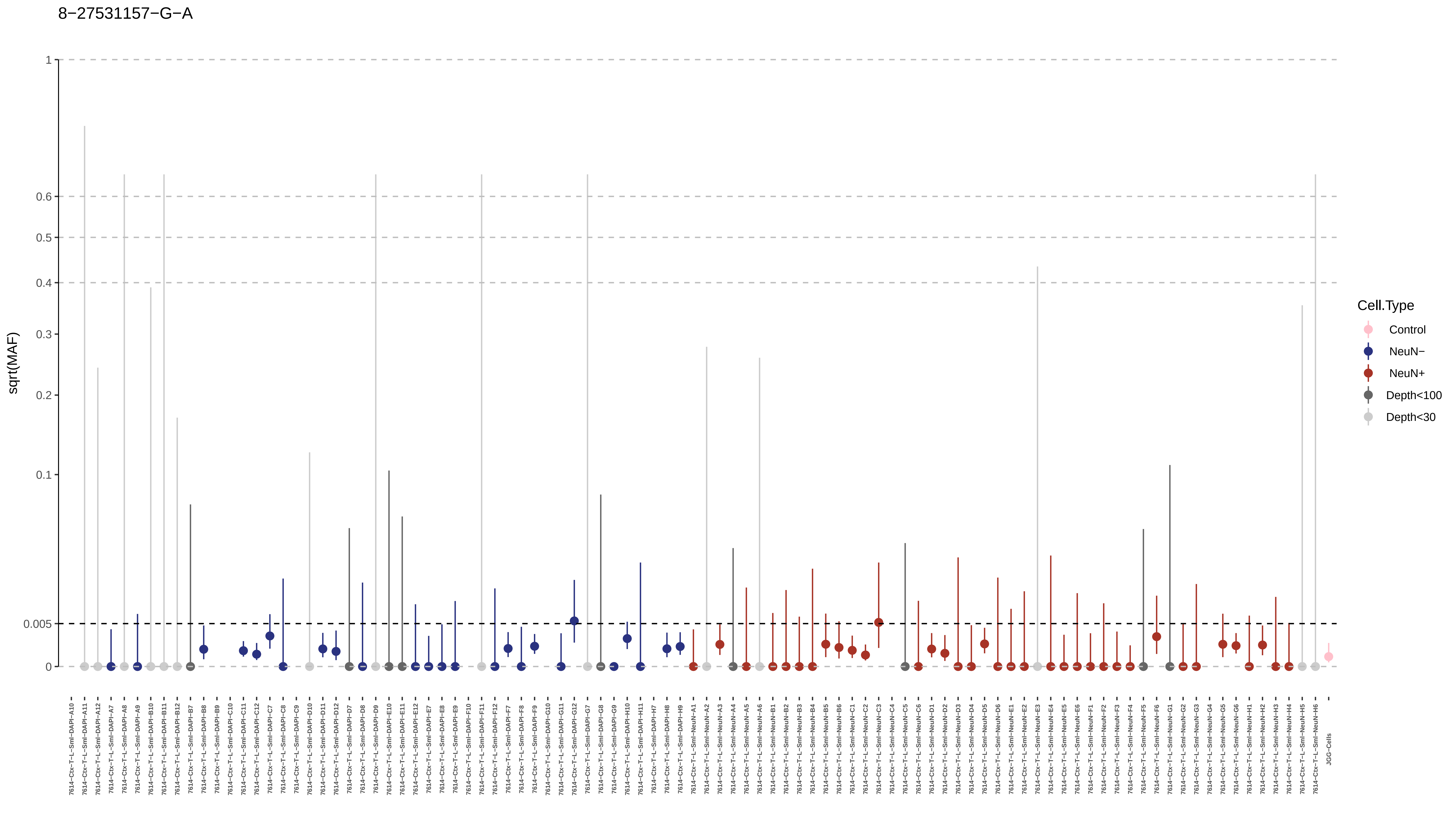

8−27531157−G−A
sqrt(MAF)
1
0.6
0.5
0.4
0.3
0.2
0.1
0.005
0
Cell.Type
Control
NeuN−
NeuN+
Depth<100
Depth<30
JGG-Cells

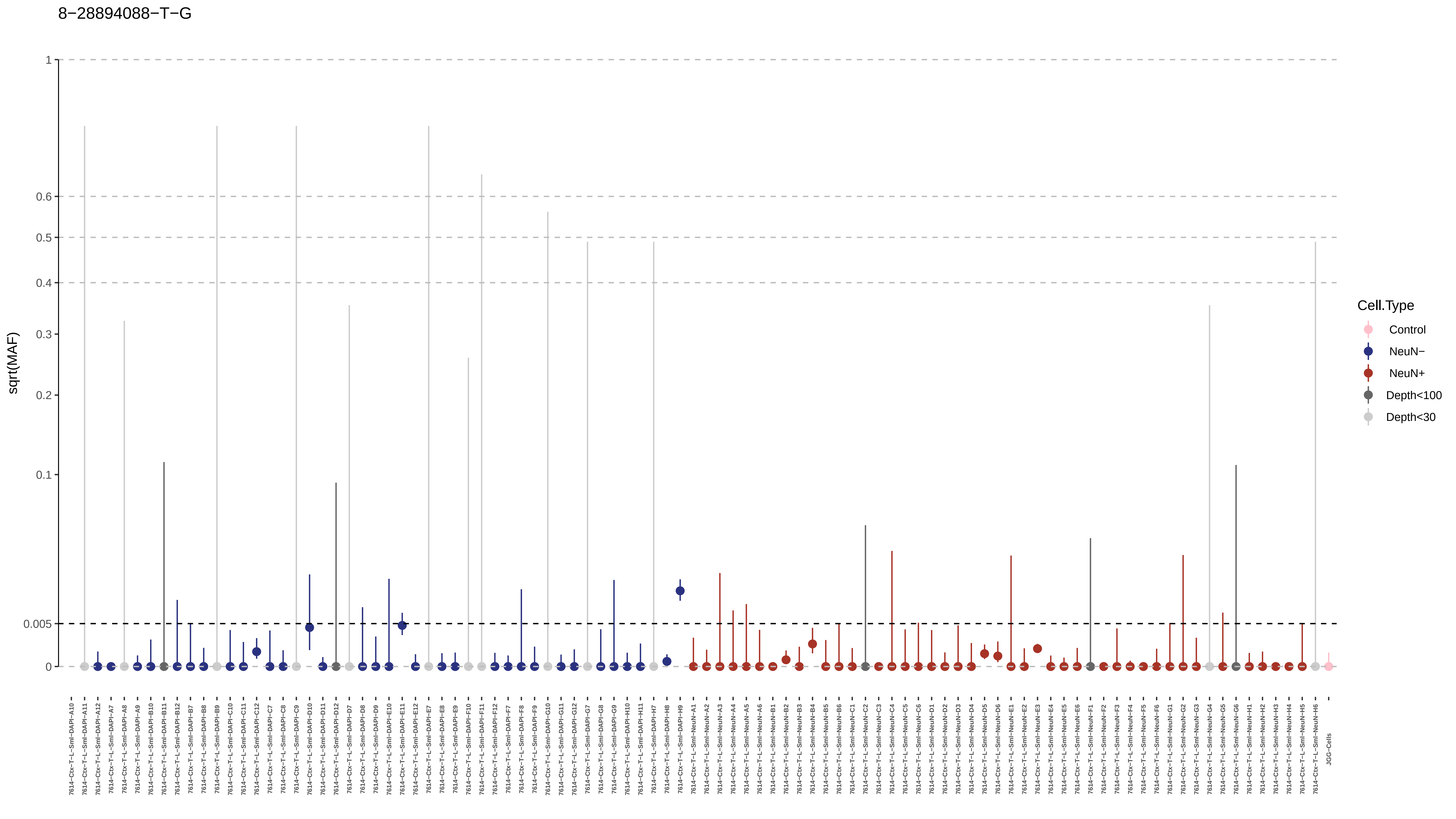

8−28894088−T−G
sqrt(MAF)
1
0.6
0.5
0.4
0.3
0.2
0.1
0.005
0
Cell.Type
Control
NeuN−
NeuN+
Depth<100
Depth<30

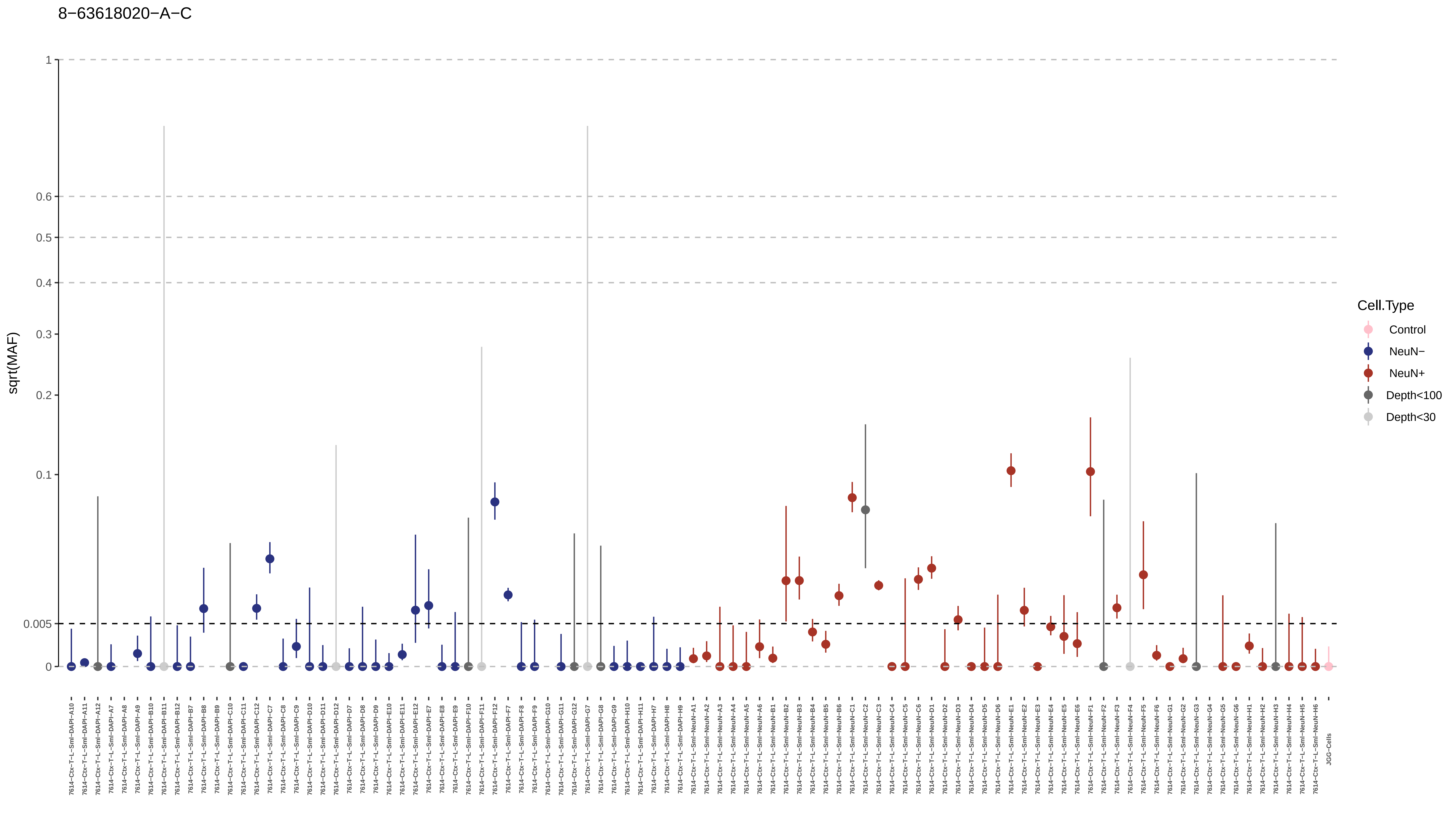
8−63618020−A−C
sqrt(MAF)
1
0.6
0.5
0.4
0.3
0.2
0.1
0.005
0
Cell.Type
Control
NeuN−
NeuN+
Depth<100
Depth<30
7614−Ctx−T−L−Sml−DAPI−A10
7614−Ctx−T−L−Sml−DAPI−A11
7614−Ctx−T−L−Sml−DAPI−A12
7614−Ctx−T−L−Sml−DAPI−A7
7614−Ctx−T−L−Sml−DAPI−A8
7614−Ctx−T−L−Sml−DAPI−A9
7614−Ctx−T−L−Sml−DAPI−B10
7614−Ctx−T−L−Sml−DAPI−B11
7614−Ctx−T−L−Sml−DAPI−B12
7614−Ctx−T−L−Sml−DAPI−B7
7614−Ctx−T−L−Sml−DAPI−B8
7614−Ctx−T−L−Sml−DAPI−B9
7614−Ctx−T−L−Sml−DAPI−C10
7614−Ctx−T−L−Sml−DAPI−C11
7614−Ctx−T−L−Sml−DAPI−C12
7614−Ctx−T−L−Sml−DAPI−C7
7614−Ctx−T−L−Sml−DAPI−C8
7614−Ctx−T−L−Sml−DAPI−C9
7614−Ctx−T−L−Sml−DAPI−D10
7614−Ctx−T−L−Sml−DAPI−D11
7614−Ctx−T−L−Sml−DAPI−D12
7614−Ctx−T−L−Sml−DAPI−D7
7614−Ctx−T−L−Sml−DAPI−D8
7614−Ctx−T−L−Sml−DAPI−D9
7614−Ctx−T−L−Sml−DAPI−E10
7614−Ctx−T−L−Sml−DAPI−E11
7614−Ctx−T−L−Sml−DAPI−E12
7614−Ctx−T−L−Sml−DAPI−E7
7614−Ctx−T−L−Sml−DAPI−E8
7614−Ctx−T−L−Sml−DAPI−E9
7614−Ctx−T−L−Sml−DAPI−F10
7614−Ctx−T−L−Sml−DAPI−F11
7614−Ctx−T−L−Sml−DAPI−F12
7614−Ctx−T−L−Sml−DAPI−F7
7614−Ctx−T−L−Sml−DAPI−F8
7614−Ctx−T−L−Sml−DAPI−F9
7614−Ctx−T−L−Sml−DAPI−G10
7614−Ctx−T−L−Sml−DAPI−G11
7614−Ctx−T−L−Sml−DAPI−G12
7614−Ctx−T−L−Sml−DAPI−G7
7614−Ctx−T−L−Sml−DAPI−G8
7614−Ctx−T−L−Sml−DAPI−G9
7614−Ctx−T−L−Sml−DAPI−H10
7614−Ctx−T−L−Sml−DAPI−H11
7614−Ctx−T−L−Sml−DAPI−H7
7614−Ctx−T−L−Sml−DAPI−H8
7614−Ctx−T−L−Sml−DAPI−H9
7614−Ctx−T−L−Sml−NeuN−A1
7614−Ctx−T−L−Sml−NeuN−A2
7614−Ctx−T−L−Sml−NeuN−A3
7614−Ctx−T−L−Sml−NeuN−A4
7614−Ctx−T−L−Sml−NeuN−A5
7614−Ctx−T−L−Sml−NeuN−A6
7614−Ctx−T−L−Sml−NeuN−B1
7614−Ctx−T−L−Sml−NeuN−B2
7614−Ctx−T−L−Sml−NeuN−B3
7614−Ctx−T−L−Sml−NeuN−B4
7614−Ctx−T−L−Sml−NeuN−B5
7614−Ctx−T−L−Sml−NeuN−B6
7614−Ctx−T−L−Sml−NeuN−C1
7614−Ctx−T−L−Sml−NeuN−C2
7614−Ctx−T−L−Sml−NeuN−C3
7614−Ctx−T−L−Sml−NeuN−C4
7614−Ctx−T−L−Sml−NeuN−C5
7614−Ctx−T−L−Sml−NeuN−C6
7614−Ctx−T−L−Sml−NeuN−D1
7614−Ctx−T−L−Sml−NeuN−D2
7614−Ctx−T−L−Sml−NeuN−D3
7614−Ctx−T−L−Sml−NeuN−D4
7614−Ctx−T−L−Sml−NeuN−D5
7614−Ctx−T−L−Sml−NeuN−D6
7614−Ctx−T−L−Sml−NeuN−E1
7614−Ctx−T−L−Sml−NeuN−E2
7614−Ctx−T−L−Sml−NeuN−E3
7614−Ctx−T−L−Sml−NeuN−E4
7614−Ctx−T−L−Sml−NeuN−E5
7614−Ctx−T−L−Sml−NeuN−E6
7614−Ctx−T−L−Sml−NeuN−F1
7614−Ctx−T−L−Sml−NeuN−F2
7614−Ctx−T−L−Sml−NeuN−F3
7614−Ctx−T−L−Sml−NeuN−F4
7614−Ctx−T−L−Sml−NeuN−F5
7614−Ctx−T−L−Sml−NeuN−F6
7614−Ctx−T−L−Sml−NeuN−G1
7614−Ctx−T−L−Sml−NeuN−G2
7614−Ctx−T−L−Sml−NeuN−G3
7614−Ctx−T−L−Sml−NeuN−G4
7614−Ctx−T−L−Sml−NeuN−G5
7614−Ctx−T−L−Sml−NeuN−G6
7614−Ctx−T−L−Sml−NeuN−H1
7614−Ctx−T−L−Sml−NeuN−H2
7614−Ctx−T−L−Sml−NeuN−H3
7614−Ctx−T−L−Sml−NeuN−H4
7614−Ctx−T−L−Sml−NeuN−H5
7614−Ctx−T−L−Sml−NeuN−H6
JGG−Cells

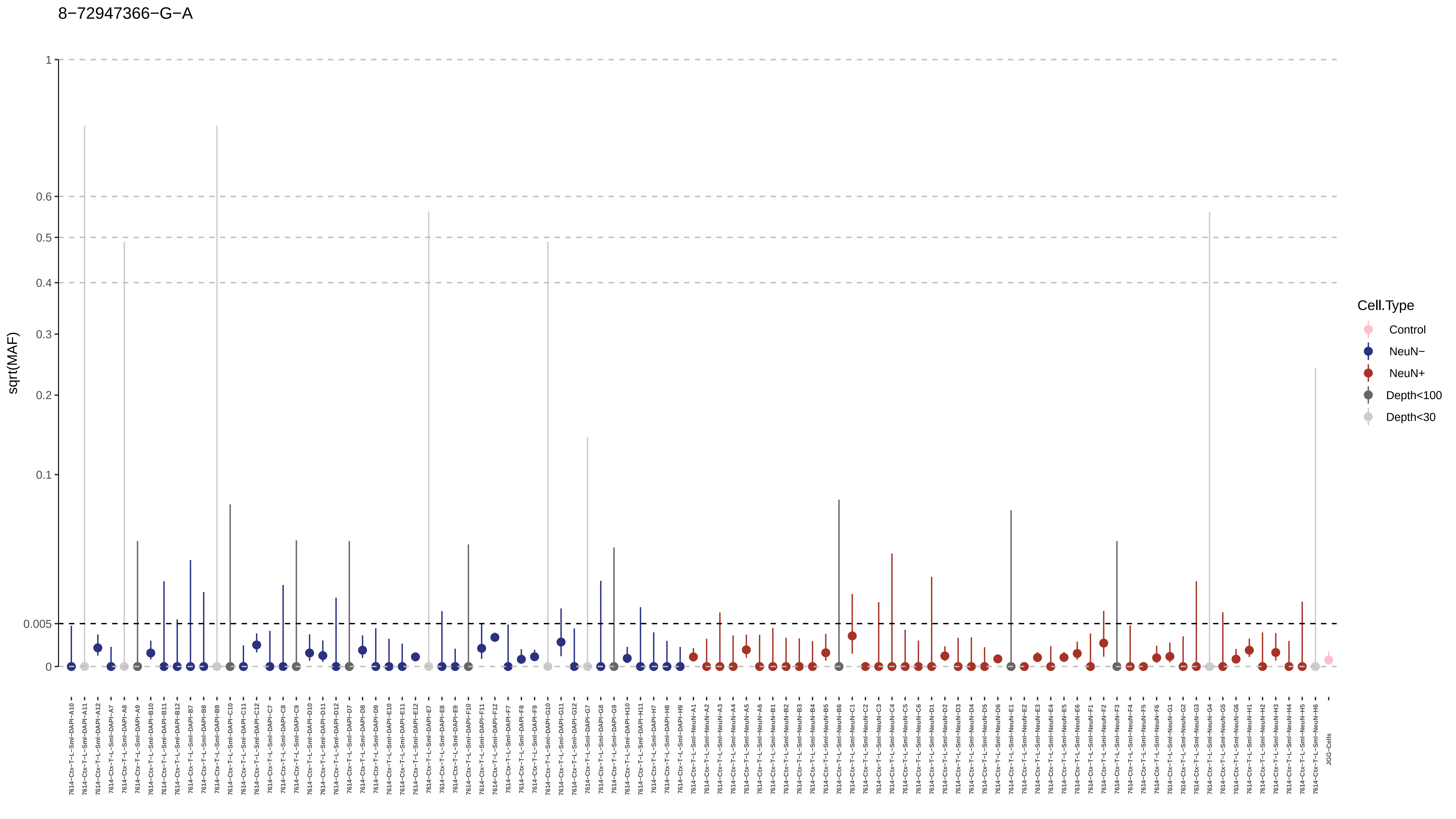
8−72947366−G−A
sqrt(MAF)
1
0.6
0.5
0.4
0.3
0.2
0.1
0.005
0
Cell.Type
Control
NeuN−
NeuN+
Depth<100
Depth<30
JGG-Cells

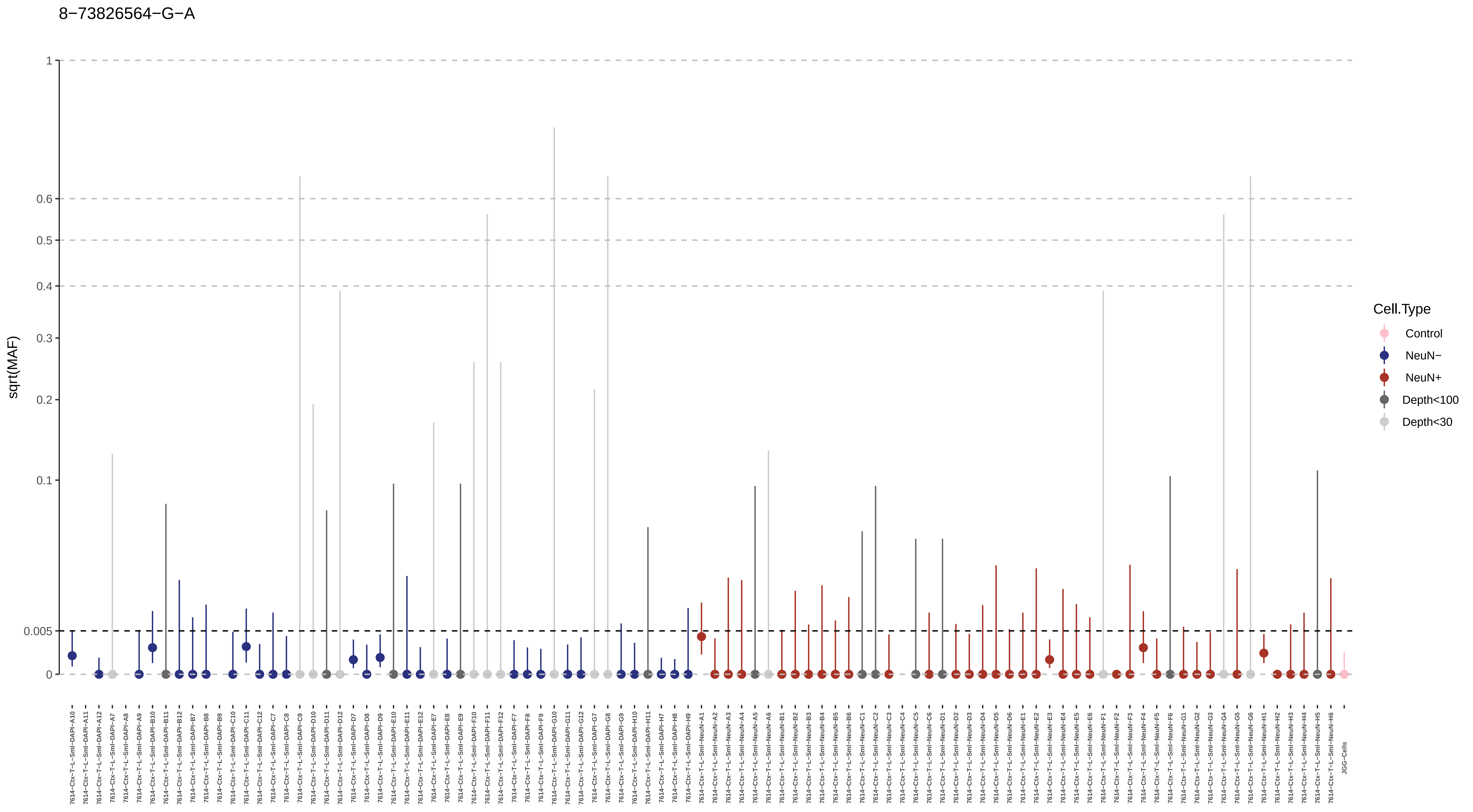
8−73826564−G−A
sqrt(MAF)
1
0.6
0.5
0.4
0.3
0.2
0.1
0.005
0
Cell.Type
Control
NeuN−
NeuN+
Depth<100
Depth<30
7614−Ctx−T−L−Sml−DAPI−A10
7614−Ctx−T−L−Sml−DAPI−A11
7614−Ctx−T−L−Sml−DAPI−A12
7614−Ctx−T−L−Sml−DAPI−A7
7614−Ctx−T−L−Sml−DAPI−A8
7614−Ctx−T−L−Sml−DAPI−A9
7614−Ctx−T−L−Sml−DAPI−B10
7614−Ctx−T−L−Sml−DAPI−B11
7614−Ctx−T−L−Sml−DAPI−B12
7614−Ctx−T−L−Sml−DAPI−B7
7614−Ctx−T−L−Sml−DAPI−B8
7614−Ctx−T−L−Sml−DAPI−B9
7614−Ctx−T−L−Sml−DAPI−C10
7614−Ctx−T−L−Sml−DAPI−C11
7614−Ctx−T−L−Sml−DAPI−C12
7614−Ctx−T−L−Sml−DAPI−C7
7614−Ctx−T−L−Sml−DAPI−C8
7614−Ctx−T−L−Sml−DAPI−C9
7614−Ctx−T−L−Sml−DAPI−D10
7614−Ctx−T−L−Sml−DAPI−D11
7614−Ctx−T−L−Sml−DAPI−D12
7614−Ctx−T−L−Sml−DAPI−D7
7614−Ctx−T−L−Sml−DAPI−D8
7614−Ctx−T−L−Sml−DAPI−D9
7614−Ctx−T−L−Sml−DAPI−E10
7614−Ctx−T−L−Sml−DAPI−E11
7614−Ctx−T−L−Sml−DAPI−E12
7614−Ctx−T−L−Sml−DAPI−E7
7614−Ctx−T−L−Sml−DAPI−E8
7614−Ctx−T−L−Sml−DAPI−E9
7614−Ctx−T−L−Sml−DAPI−F10
7614−Ctx−T−L−Sml−DAPI−F11
7614−Ctx−T−L−Sml−DAPI−F12
7614−Ctx−T−L−Sml−DAPI−F7
7614−Ctx−T−L−Sml−DAPI−F8
7614−Ctx−T−L−Sml−DAPI−F9
7614−Ctx−T−L−Sml−DAPI−G10
7614−Ctx−T−L−Sml−DAPI−G11
7614−Ctx−T−L−Sml−DAPI−G12
7614−Ctx−T−L−Sml−DAPI−G7
7614−Ctx−T−L−Sml−DAPI−G8
7614−Ctx−T−L−Sml−DAPI−G9
7614−Ctx−T−L−Sml−DAPI−H10
7614−Ctx−T−L−Sml−DAPI−H11
7614−Ctx−T−L−Sml−DAPI−H7
7614−Ctx−T−L−Sml−DAPI−H8
7614−Ctx−T−L−Sml−DAPI−H9
7614−Ctx−T−L−Sml−NeuN−A1
7614−Ctx−T−L−Sml−NeuN−A2
7614−Ctx−T−L−Sml−NeuN−A3
7614−Ctx−T−L−Sml−NeuN−A4
7614−Ctx−T−L−Sml−NeuN−A5
7614−Ctx−T−L−Sml−NeuN−A6
7614−Ctx−T−L−Sml−NeuN−B1
7614−Ctx−T−L−Sml−NeuN−B2
7614−Ctx−T−L−Sml−NeuN−B3
7614−Ctx−T−L−Sml−NeuN−B4
7614−Ctx−T−L−Sml−NeuN−B5
7614−Ctx−T−L−Sml−NeuN−B6
7614−Ctx−T−L−Sml−NeuN−C1
7614−Ctx−T−L−Sml−NeuN−C2
7614−Ctx−T−L−Sml−NeuN−C3
7614−Ctx−T−L−Sml−NeuN−C4
7614−Ctx−T−L−Sml−NeuN−C5
7614−Ctx−T−L−Sml−NeuN−C6
7614−Ctx−T−L−Sml−NeuN−D1
7614−Ctx−T−L−Sml−NeuN−D2
7614−Ctx−T−L−Sml−NeuN−D3
7614−Ctx−T−L−Sml−NeuN−D4
7614−Ctx−T−L−Sml−NeuN−D5
7614−Ctx−T−L−Sml−NeuN−D6
7614−Ctx−T−L−Sml−NeuN−E1
7614−Ctx−T−L−Sml−NeuN−E2
7614−Ctx−T−L−Sml−NeuN−E3
7614−Ctx−T−L−Sml−NeuN−E4
7614−Ctx−T−L−Sml−NeuN−E5
7614−Ctx−T−L−Sml−NeuN−E6
7614−Ctx−T−L−Sml−NeuN−F1
7614−Ctx−T−L−Sml−NeuN−F2
7614−Ctx−T−L−Sml−NeuN−F3
7614−Ctx−T−L−Sml−NeuN−F4
7614−Ctx−T−L−Sml−NeuN−F5
7614−Ctx−T−L−Sml−NeuN−F6
7614−Ctx−T−L−Sml−NeuN−G1
7614−Ctx−T−L−Sml−NeuN−G2
7614−Ctx−T−L−Sml−NeuN−G3
7614−Ctx−T−L−Sml−NeuN−G4
7614−Ctx−T−L−Sml−NeuN−G5
7614−Ctx−T−L−Sml−NeuN−G6
7614−Ctx−T−L−Sml−NeuN−H1
7614−Ctx−T−L−Sml−NeuN−H2
7614−Ctx−T−L−Sml−NeuN−H3
7614−Ctx−T−L−Sml−NeuN−H4
7614−Ctx−T−L−Sml−NeuN−H5
7614−Ctx−T−L−Sml−NeuN−H6
JGG-Cells

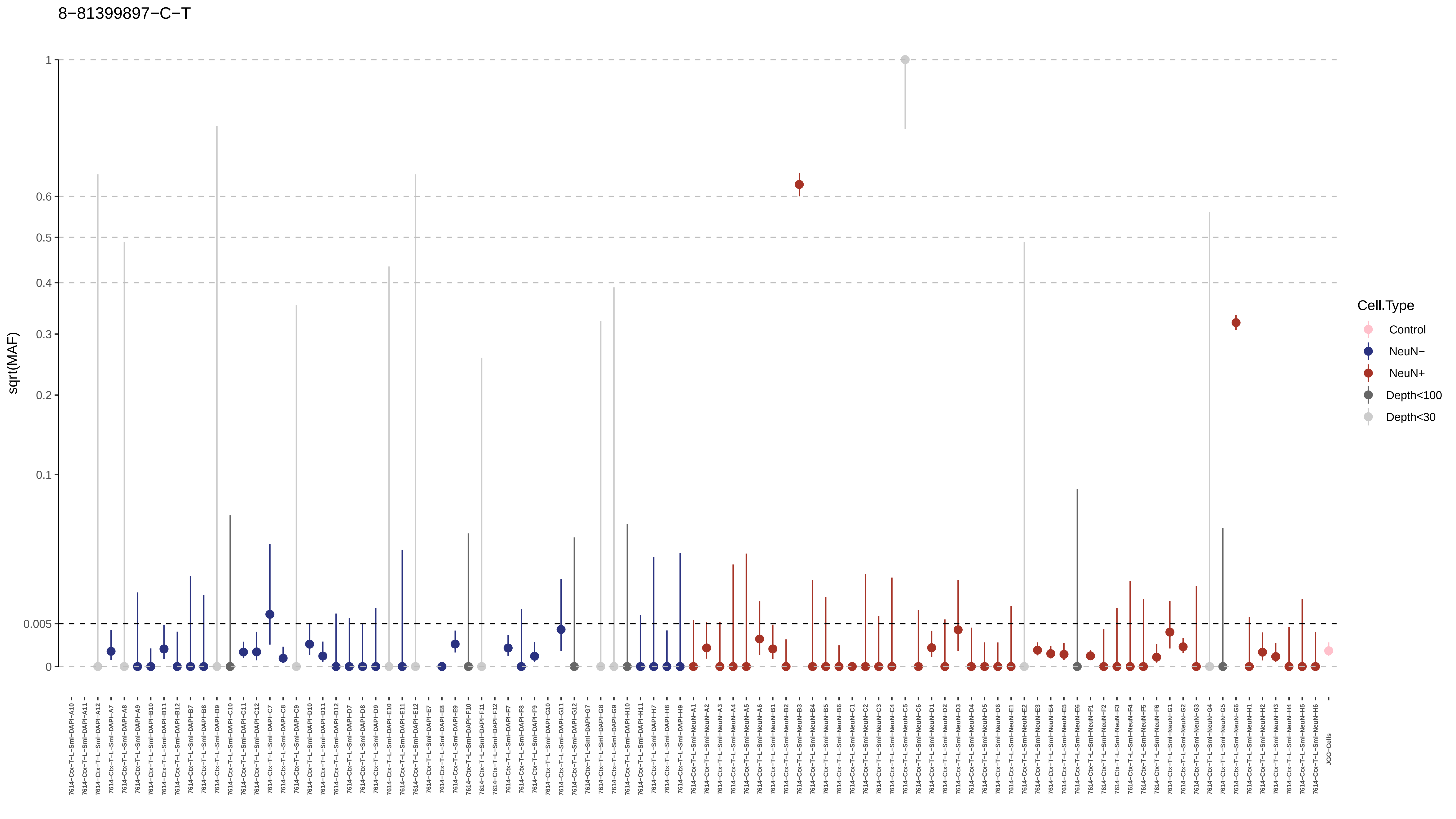
8−81399897−C−T
sqrt(MAF)
1
0.6
0.5
0.4
0.3
0.2
0.1
0.005
0
Cell.Type
Control
NeuN−
NeuN+
Depth<100
Depth<30

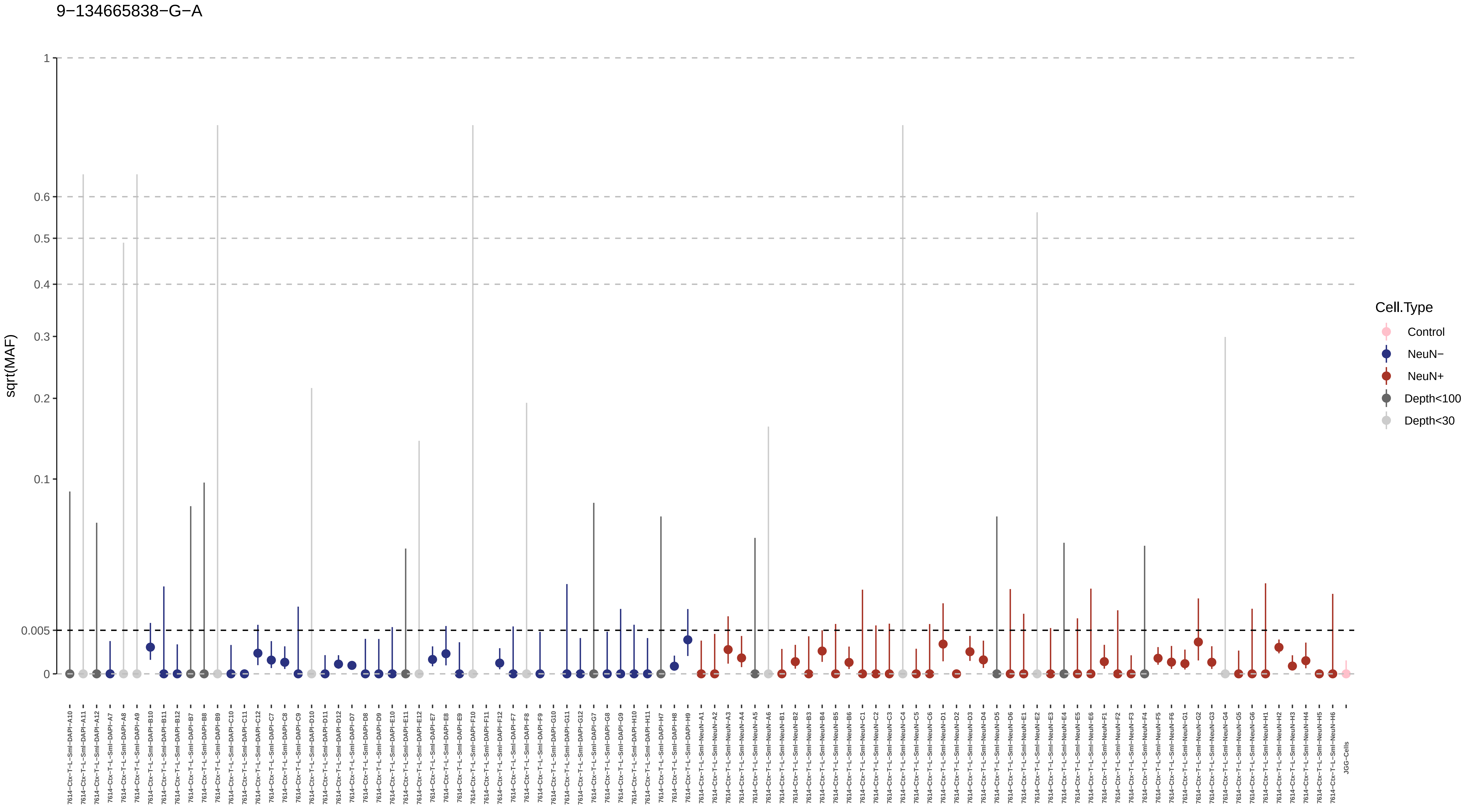
9−134665838−G−A
sqrt(MAF)
1
0.6
0.5
0.4
0.3
0.2
0.1
0.005
0
Cell.Type
Control
NeuN−
NeuN+
Depth<100
Depth<30
7614−Ctx−T−L−Sml−DAPI−A10
7614−Ctx−T−L−Sml−DAPI−A11
7614−Ctx−T−L−Sml−DAPI−A12
7614−Ctx−T−L−Sml−DAPI−A7
7614−Ctx−T−L−Sml−DAPI−A8
7614−Ctx−T−L−Sml−DAPI−A9
7614−Ctx−T−L−Sml−DAPI−B10
7614−Ctx−T−L−Sml−DAPI−B11
7614−Ctx−T−L−Sml−DAPI−B12
7614−Ctx−T−L−Sml−DAPI−B7
7614−Ctx−T−L−Sml−DAPI−B8
7614−Ctx−T−L−Sml−DAPI−B9
7614−Ctx−T−L−Sml−DAPI−C10
7614−Ctx−T−L−Sml−DAPI−C11
7614−Ctx−T−L−Sml−DAPI−C12
7614−Ctx−T−L−Sml−DAPI−C7
7614−Ctx−T−L−Sml−DAPI−C8
7614−Ctx−T−L−Sml−DAPI−C9
7614−Ctx−T−L−Sml−DAPI−D10
7614−Ctx−T−L−Sml−DAPI−D11
7614−Ctx−T−L−Sml−DAPI−D12
7614−Ctx−T−L−Sml−DAPI−D7
7614−Ctx−T−L−Sml−DAPI−D8
7614−Ctx−T−L−Sml−DAPI−D9
7614−Ctx−T−L−Sml−DAPI−E10
7614−Ctx−T−L−Sml−DAPI−E11
7614−Ctx−T−L−Sml−DAPI−E12
7614−Ctx−T−L−Sml−DAPI−E7
7614−Ctx−T−L−Sml−DAPI−E8
7614−Ctx−T−L−Sml−DAPI−E9
7614−Ctx−T−L−Sml−DAPI−F10
7614−Ctx−T−L−Sml−DAPI−F11
7614−Ctx−T−L−Sml−DAPI−F12
7614−Ctx−T−L−Sml−DAPI−F7
7614−Ctx−T−L−Sml−DAPI−F8
7614−Ctx−T−L−Sml−DAPI−F9
7614−Ctx−T−L−Sml−DAPI−G10
7614−Ctx−T−L−Sml−DAPI−G11
7614−Ctx−T−L−Sml−DAPI−G12
7614−Ctx−T−L−Sml−DAPI−G7
7614−Ctx−T−L−Sml−DAPI−G8
7614−Ctx−T−L−Sml−DAPI−G9
7614−Ctx−T−L−Sml−DAPI−H10
7614−Ctx−T−L−Sml−DAPI−H11
7614−Ctx−T−L−Sml−DAPI−H7
7614−Ctx−T−L−Sml−DAPI−H8
7614−Ctx−T−L−Sml−DAPI−H9
7614−Ctx−T−L−Sml−NeuN−A1
7614−Ctx−T−L−Sml−NeuN−A2
7614−Ctx−T−L−Sml−NeuN−A3
7614−Ctx−T−L−Sml−NeuN−A4
7614−Ctx−T−L−Sml−NeuN−A5
7614−Ctx−T−L−Sml−NeuN−A6
7614−Ctx−T−L−Sml−NeuN−B1
7614−Ctx−T−L−Sml−NeuN−B2
7614−Ctx−T−L−Sml−NeuN−B3
7614−Ctx−T−L−Sml−NeuN−B4
7614−Ctx−T−L−Sml−NeuN−B5
7614−Ctx−T−L−Sml−NeuN−B6
7614−Ctx−T−L−Sml−NeuN−C1
7614−Ctx−T−L−Sml−NeuN−C2
7614−Ctx−T−L−Sml−NeuN−C3
7614−Ctx−T−L−Sml−NeuN−C4
7614−Ctx−T−L−Sml−NeuN−C5
7614−Ctx−T−L−Sml−NeuN−C6
7614−Ctx−T−L−Sml−NeuN−D1
7614−Ctx−T−L−Sml−NeuN−D2
7614−Ctx−T−L−Sml−NeuN−D3
7614−Ctx−T−L−Sml−NeuN−D4
7614−Ctx−T−L−Sml−NeuN−D5
7614−Ctx−T−L−Sml−NeuN−D6
7614−Ctx−T−L−Sml−NeuN−E1
7614−Ctx−T−L−Sml−NeuN−E2
7614−Ctx−T−L−Sml−NeuN−E3
7614−Ctx−T−L−Sml−NeuN−E4
7614−Ctx−T−L−Sml−NeuN−E5
7614−Ctx−T−L−Sml−NeuN−E6
7614−Ctx−T−L−Sml−NeuN−F1
7614−Ctx−T−L−Sml−NeuN−F2
7614−Ctx−T−L−Sml−NeuN−F3
7614−Ctx−T−L−Sml−NeuN−F4
7614−Ctx−T−L−Sml−NeuN−F5
7614−Ctx−T−L−Sml−NeuN−F6
7614−Ctx−T−L−Sml−NeuN−G1
7614−Ctx−T−L−Sml−NeuN−G2
7614−Ctx−T−L−Sml−NeuN−G3
7614−Ctx−T−L−Sml−NeuN−G4
7614−Ctx−T−L−Sml−NeuN−G5
7614−Ctx−T−L−Sml−NeuN−G6
7614−Ctx−T−L−Sml−NeuN−H1
7614−Ctx−T−L−Sml−NeuN−H2
7614−Ctx−T−L−Sml−NeuN−H3
7614−Ctx−T−L−Sml−NeuN−H4
7614−Ctx−T−L−Sml−NeuN−H5
7614−Ctx−T−L−Sml−NeuN−H6
JGG−Cells

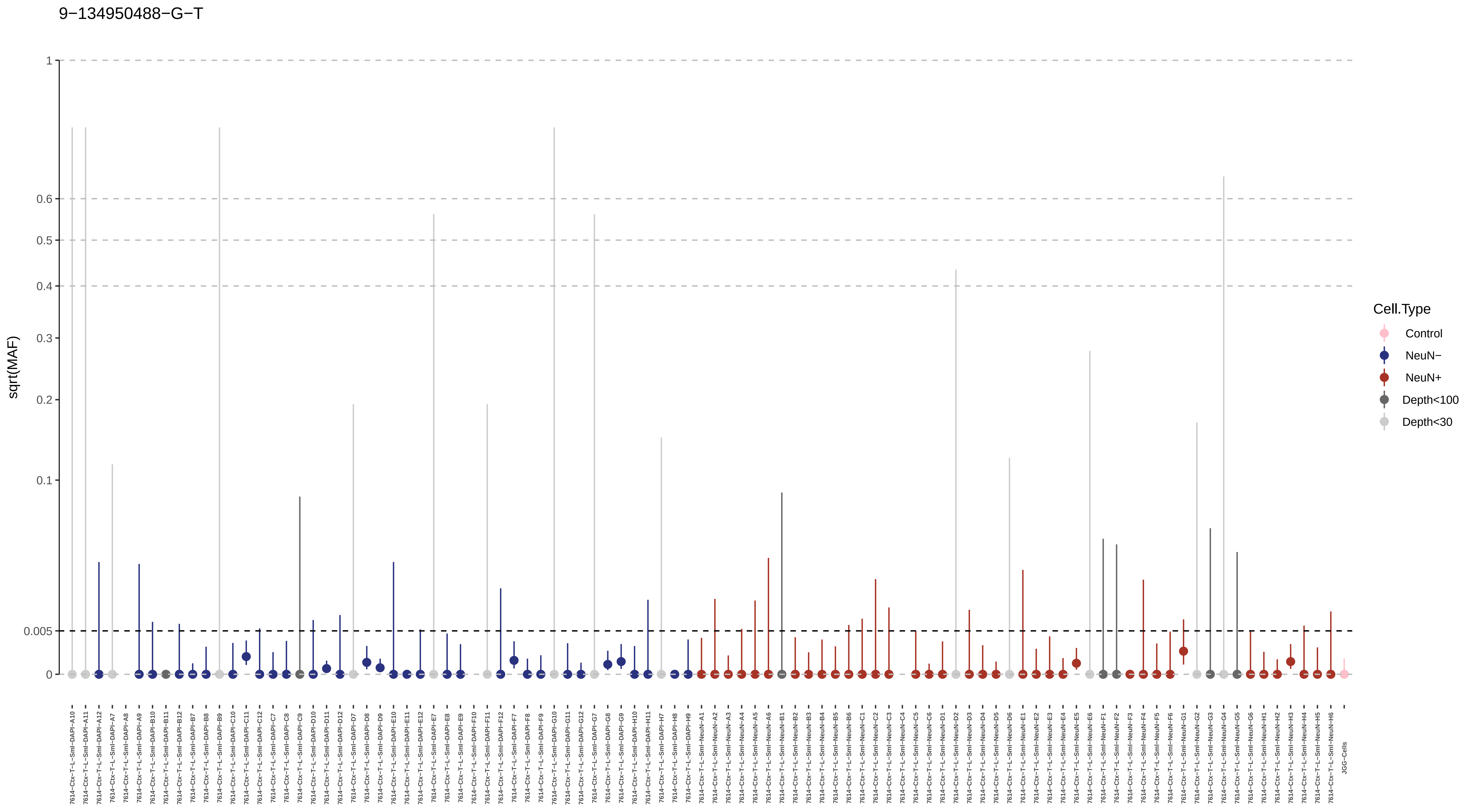
9−134950488−G−T
sqrt(MAF)
1
0.6
0.5
0.4
0.3
0.2
0.1
0.005
0
Cell.Type
Control
NeuN−
NeuN+
Depth<100
Depth<30
7614−Ctx−T−L−Sml−DAPI−A10
7614−Ctx−T−L−Sml−DAPI−A11
7614−Ctx−T−L−Sml−DAPI−A12
7614−Ctx−T−L−Sml−DAPI−A7
7614−Ctx−T−L−Sml−DAPI−A8
7614−Ctx−T−L−Sml−DAPI−A9
7614−Ctx−T−L−Sml−DAPI−B10
7614−Ctx−T−L−Sml−DAPI−B11
7614−Ctx−T−L−Sml−DAPI−B12
7614−Ctx−T−L−Sml−DAPI−B7
7614−Ctx−T−L−Sml−DAPI−B8
7614−Ctx−T−L−Sml−DAPI−B9
7614−Ctx−T−L−Sml−DAPI−C10
7614−Ctx−T−L−Sml−DAPI−C11
7614−Ctx−T−L−Sml−DAPI−C12
7614−Ctx−T−L−Sml−DAPI−C7
7614−Ctx−T−L−Sml−DAPI−C8
7614−Ctx−T−L−Sml−DAPI−C9
7614−Ctx−T−L−Sml−DAPI−D10
7614−Ctx−T−L−Sml−DAPI−D11
7614−Ctx−T−L−Sml−DAPI−D12
7614−Ctx−T−L−Sml−DAPI−D7
7614−Ctx−T−L−Sml−DAPI−D8
7614−Ctx−T−L−Sml−DAPI−D9
7614−Ctx−T−L−Sml−DAPI−E10
7614−Ctx−T−L−Sml−DAPI−E11
7614−Ctx−T−L−Sml−DAPI−E12
7614−Ctx−T−L−Sml−DAPI−E7
7614−Ctx−T−L−Sml−DAPI−E8
7614−Ctx−T−L−Sml−DAPI−E9
7614−Ctx−T−L−Sml−DAPI−F10
7614−Ctx−T−L−Sml−DAPI−F11
7614−Ctx−T−L−Sml−DAPI−F12
7614−Ctx−T−L−Sml−DAPI−F7
7614−Ctx−T−L−Sml−DAPI−F8
7614−Ctx−T−L−Sml−DAPI−F9
7614−Ctx−T−L−Sml−DAPI−G10
7614−Ctx−T−L−Sml−DAPI−G11
7614−Ctx−T−L−Sml−DAPI−G12
7614−Ctx−T−L−Sml−DAPI−G7
7614−Ctx−T−L−Sml−DAPI−G8
7614−Ctx−T−L−Sml−DAPI−G9
7614−Ctx−T−L−Sml−DAPI−H10
7614−Ctx−T−L−Sml−DAPI−H11
7614−Ctx−T−L−Sml−DAPI−H7
7614−Ctx−T−L−Sml−DAPI−H8
7614−Ctx−T−L−Sml−DAPI−H9
7614−Ctx−T−L−Sml−NeuN−A1
7614−Ctx−T−L−Sml−NeuN−A2
7614−Ctx−T−L−Sml−NeuN−A3
7614−Ctx−T−L−Sml−NeuN−A4
7614−Ctx−T−L−Sml−NeuN−A5
7614−Ctx−T−L−Sml−NeuN−A6
7614−Ctx−T−L−Sml−NeuN−B1
7614−Ctx−T−L−Sml−NeuN−B2
7614−Ctx−T−L−Sml−NeuN−B3
7614−Ctx−T−L−Sml−NeuN−B4
7614−Ctx−T−L−Sml−NeuN−B5
7614−Ctx−T−L−Sml−NeuN−B6
7614−Ctx−T−L−Sml−NeuN−C1
7614−Ctx−T−L−Sml−NeuN−C2
7614−Ctx−T−L−Sml−NeuN−C3
7614−Ctx−T−L−Sml−NeuN−C4
7614−Ctx−T−L−Sml−NeuN−C5
7614−Ctx−T−L−Sml−NeuN−C6
7614−Ctx−T−L−Sml−NeuN−D1
7614−Ctx−T−L−Sml−NeuN−D2
7614−Ctx−T−L−Sml−NeuN−D3
7614−Ctx−T−L−Sml−NeuN−D4
7614−Ctx−T−L−Sml−NeuN−D5
7614−Ctx−T−L−Sml−NeuN−D6
7614−Ctx−T−L−Sml−NeuN−E1
7614−Ctx−T−L−Sml−NeuN−E2
7614−Ctx−T−L−Sml−NeuN−E3
7614−Ctx−T−L−Sml−NeuN−E4
7614−Ctx−T−L−Sml−NeuN−E5
7614−Ctx−T−L−Sml−NeuN−E6
7614−Ctx−T−L−Sml−NeuN−F1
7614−Ctx−T−L−Sml−NeuN−F2
7614−Ctx−T−L−Sml−NeuN−F3
7614−Ctx−T−L−Sml−NeuN−F4
7614−Ctx−T−L−Sml−NeuN−F5
7614−Ctx−T−L−Sml−NeuN−F6
7614−Ctx−T−L−Sml−NeuN−G1
7614−Ctx−T−L−Sml−NeuN−G2
7614−Ctx−T−L−Sml−NeuN−G3
7614−Ctx−T−L−Sml−NeuN−G4
7614−Ctx−T−L−Sml−NeuN−G5
7614−Ctx−T−L−Sml−NeuN−G6
7614−Ctx−T−L−Sml−NeuN−H1
7614−Ctx−T−L−Sml−NeuN−H2
7614−Ctx−T−L−Sml−NeuN−H3
7614−Ctx−T−L−Sml−NeuN−H4
7614−Ctx−T−L−Sml−NeuN−H5
7614−Ctx−T−L−Sml−NeuN−H6
JGG−Cells

9−1722340−A−G

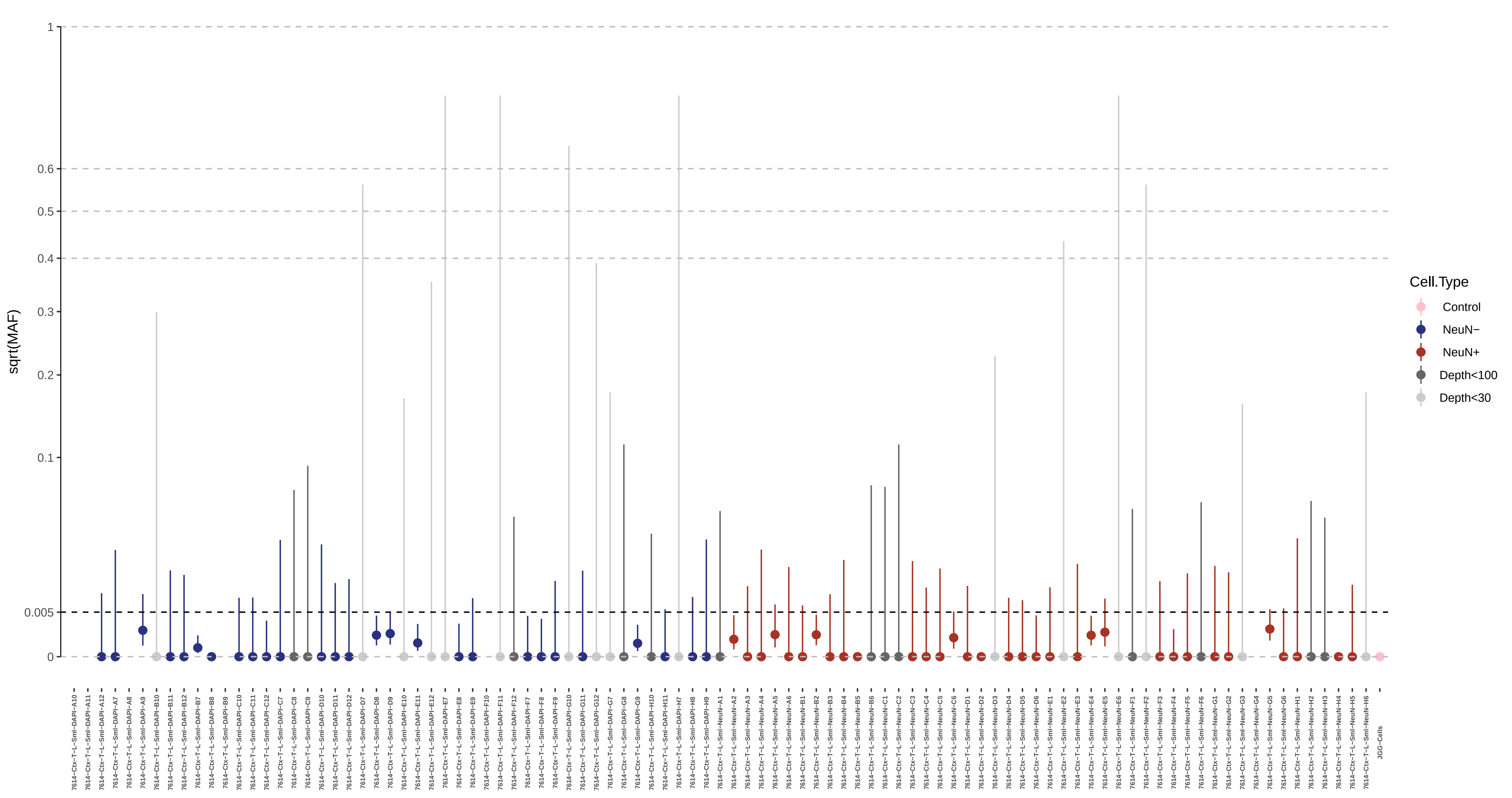

sqrt(MAF)
1
0.6
0.5
0.4
0.3
0.2
0.1
0.005
0
Cell.Type
Control
NeuN−
NeuN+
Depth<100
Depth<30
7614−Ctx−T−L−Sml−DAPI−A10
7614−Ctx−T−L−Sml−DAPI−A11
7614−Ctx−T−L−Sml−DAPI−A12
7614−Ctx−T−L−Sml−DAPI−A7
7614−Ctx−T−L−Sml−DAPI−A8
7614−Ctx−T−L−Sml−DAPI−A9
7614−Ctx−T−L−Sml−DAPI−B10
7614−Ctx−T−L−Sml−DAPI−B11
7614−Ctx−T−L−Sml−DAPI−B12
7614−Ctx−T−L−Sml−DAPI−B7
7614−Ctx−T−L−Sml−DAPI−B8
7614−Ctx−T−L−Sml−DAPI−B9
7614−Ctx−T−L−Sml−DAPI−C10
7614−Ctx−T−L−Sml−DAPI−C11
7614−Ctx−T−L−Sml−DAPI−C12
7614−Ctx−T−L−Sml−DAPI−C7
7614−Ctx−T−L−Sml−DAPI−C8
7614−Ctx−T−L−Sml−DAPI−C9
7614−Ctx−T−L−Sml−DAPI−D10
7614−Ctx−T−L−Sml−DAPI−D11
7614−Ctx−T−L−Sml−DAPI−D12
7614−Ctx−T−L−Sml−DAPI−D7
7614−Ctx−T−L−Sml−DAPI−D8
7614−Ctx−T−L−Sml−DAPI−D9
7614−Ctx−T−L−Sml−DAPI−E10
7614−Ctx−T−L−Sml−DAPI−E11
7614−Ctx−T−L−Sml−DAPI−E12
7614−Ctx−T−L−Sml−DAPI−E7
7614−Ctx−T−L−Sml−DAPI−E8
7614−Ctx−T−L−Sml−DAPI−E9
7614−Ctx−T−L−Sml−DAPI−F10
7614−Ctx−T−L−Sml−DAPI−F11
7614−Ctx−T−L−Sml−DAPI−F12
7614−Ctx−T−L−Sml−DAPI−F7
7614−Ctx−T−L−Sml−DAPI−F8
7614−Ctx−T−L−Sml−DAPI−F9
7614−Ctx−T−L−Sml−DAPI−G10
7614−Ctx−T−L−Sml−DAPI−G11
7614−Ctx−T−L−Sml−DAPI−G12
7614−Ctx−T−L−Sml−DAPI−G7
7614−Ctx−T−L−Sml−DAPI−G8
7614−Ctx−T−L−Sml−DAPI−G9
7614−Ctx−T−L−Sml−DAPI−H10
7614−Ctx−T−L−Sml−DAPI−H11
7614−Ctx−T−L−Sml−DAPI−H7
7614−Ctx−T−L−Sml−DAPI−H8
7614−Ctx−T−L−Sml−DAPI−H9
7614−Ctx−T−L−Sml−NeuN−A1
7614−Ctx−T−L−Sml−NeuN−A2
7614−Ctx−T−L−Sml−NeuN−A3
7614−Ctx−T−L−Sml−NeuN−A4
7614−Ctx−T−L−Sml−NeuN−A5
7614−Ctx−T−L−Sml−NeuN−A6
7614−Ctx−T−L−Sml−NeuN−B1
7614−Ctx−T−L−Sml−NeuN−B2
7614−Ctx−T−L−Sml−NeuN−B3
7614−Ctx−T−L−Sml−NeuN−B4
7614−Ctx−T−L−Sml−NeuN−B5
7614−Ctx−T−L−Sml−NeuN−B6
7614−Ctx−T−L−Sml−NeuN−C1
7614−Ctx−T−L−Sml−NeuN−C2
7614−Ctx−T−L−Sml−NeuN−C3
7614−Ctx−T−L−Sml−NeuN−C4
7614−Ctx−T−L−Sml−NeuN−C5
7614−Ctx−T−L−Sml−NeuN−C6
7614−Ctx−T−L−Sml−NeuN−D1
7614−Ctx−T−L−Sml−NeuN−D2
7614−Ctx−T−L−Sml−NeuN−D3
7614−Ctx−T−L−Sml−NeuN−D4
7614−Ctx−T−L−Sml−NeuN−D5
7614−Ctx−T−L−Sml−NeuN−D6
7614−Ctx−T−L−Sml−NeuN−E1
7614−Ctx−T−L−Sml−NeuN−E2
7614−Ctx−T−L−Sml−NeuN−E3
7614−Ctx−T−L−Sml−NeuN−E4
7614−Ctx−T−L−Sml−NeuN−E5
7614−Ctx−T−L−Sml−NeuN−E6
7614−Ctx−T−L−Sml−NeuN−F1
7614−Ctx−T−L−Sml−NeuN−F2
7614−Ctx−T−L−Sml−NeuN−F3
7614−Ctx−T−L−Sml−NeuN−F4
7614−Ctx−T−L−Sml−NeuN−F5
7614−Ctx−T−L−Sml−NeuN−F6
7614−Ctx−T−L−Sml−NeuN−G1
7614−Ctx−T−L−Sml−NeuN−G2
7614−Ctx−T−L−Sml−NeuN−G3
7614−Ctx−T−L−Sml−NeuN−G4
7614−Ctx−T−L−Sml−NeuN−G5
7614−Ctx−T−L−Sml−NeuN−G6
7614−Ctx−T−L−Sml−NeuN−H1
7614−Ctx−T−L−Sml−NeuN−H2
7614−Ctx−T−L−Sml−NeuN−H3
7614−Ctx−T−L−Sml−NeuN−H4
7614−Ctx−T−L−Sml−NeuN−H5
7614−Ctx−T−L−Sml−NeuN−H6
JGG−Cells

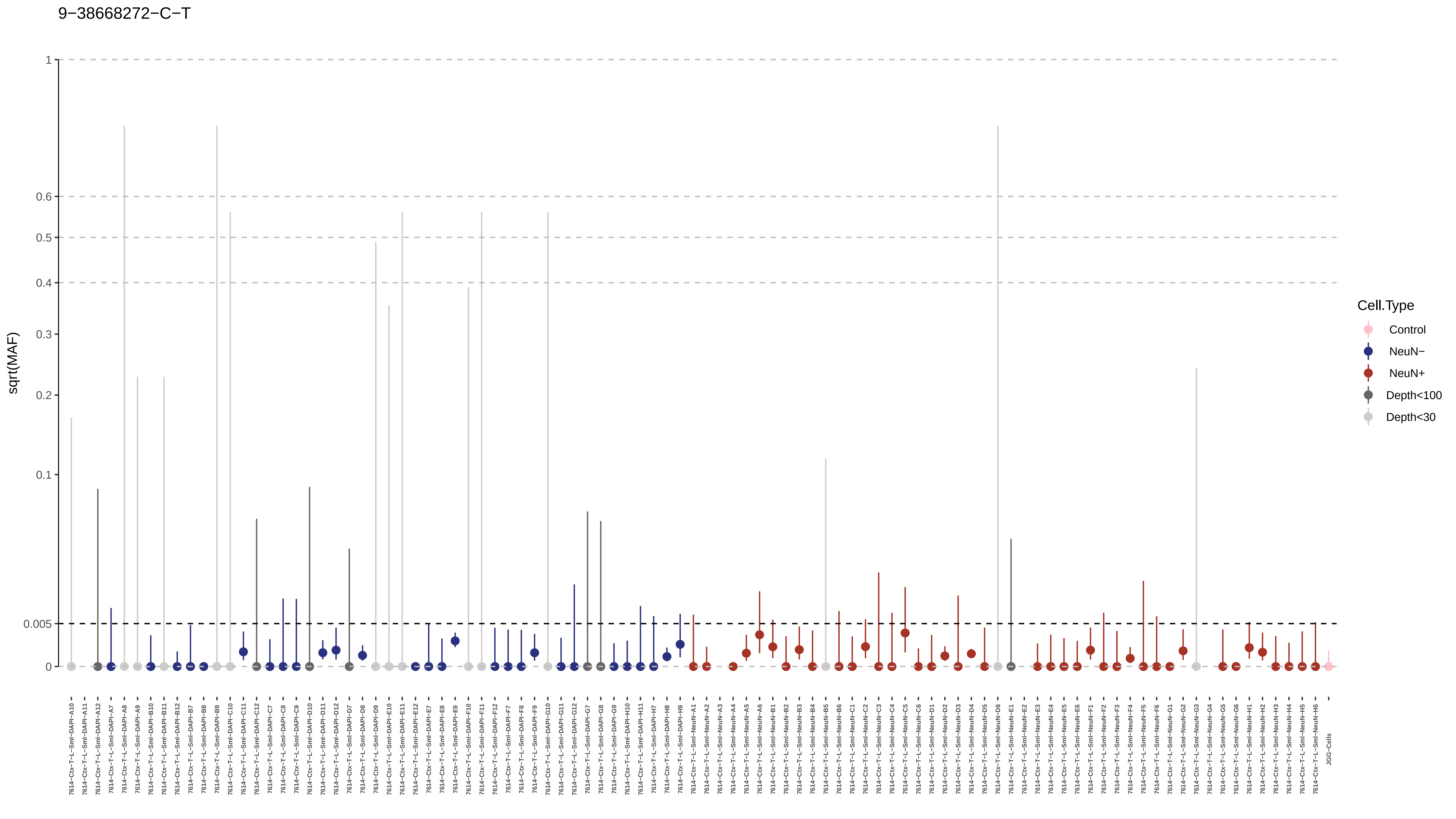
9−38668272−C−T
sqrt(MAF)
1
0.6
0.5
0.4
0.3
0.2
0.1
0.005
0
Cell.Type
Control
NeuN−
NeuN+
Depth<100
Depth<30
JGG-Cells

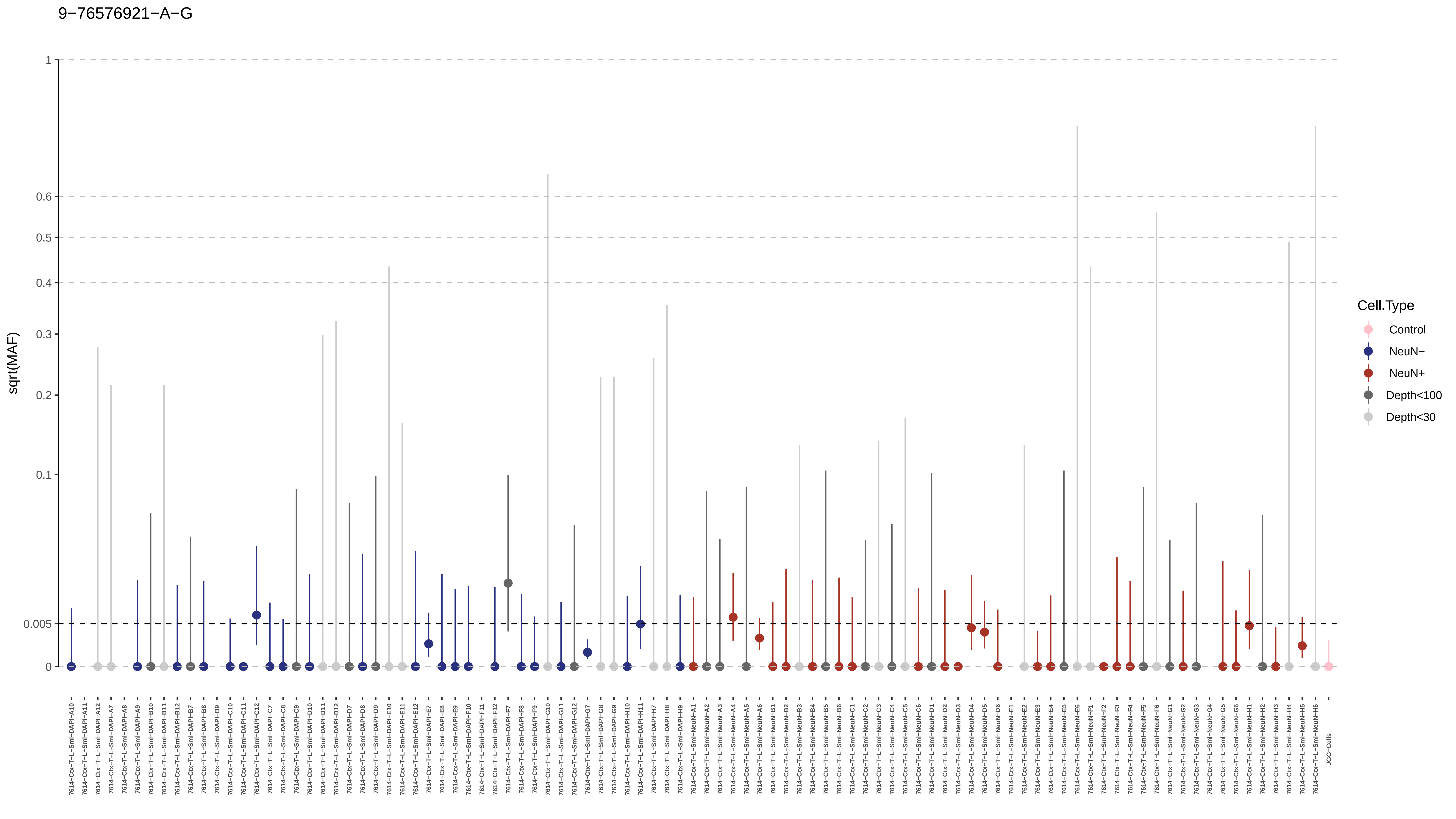
9−76576921−A−G
sqrt(MAF)
1
0.6
0.5
0.4
0.3
0.2
0.1
0.005
0
Cell.Type
Control
NeuN−
NeuN+
Depth<100
Depth<30

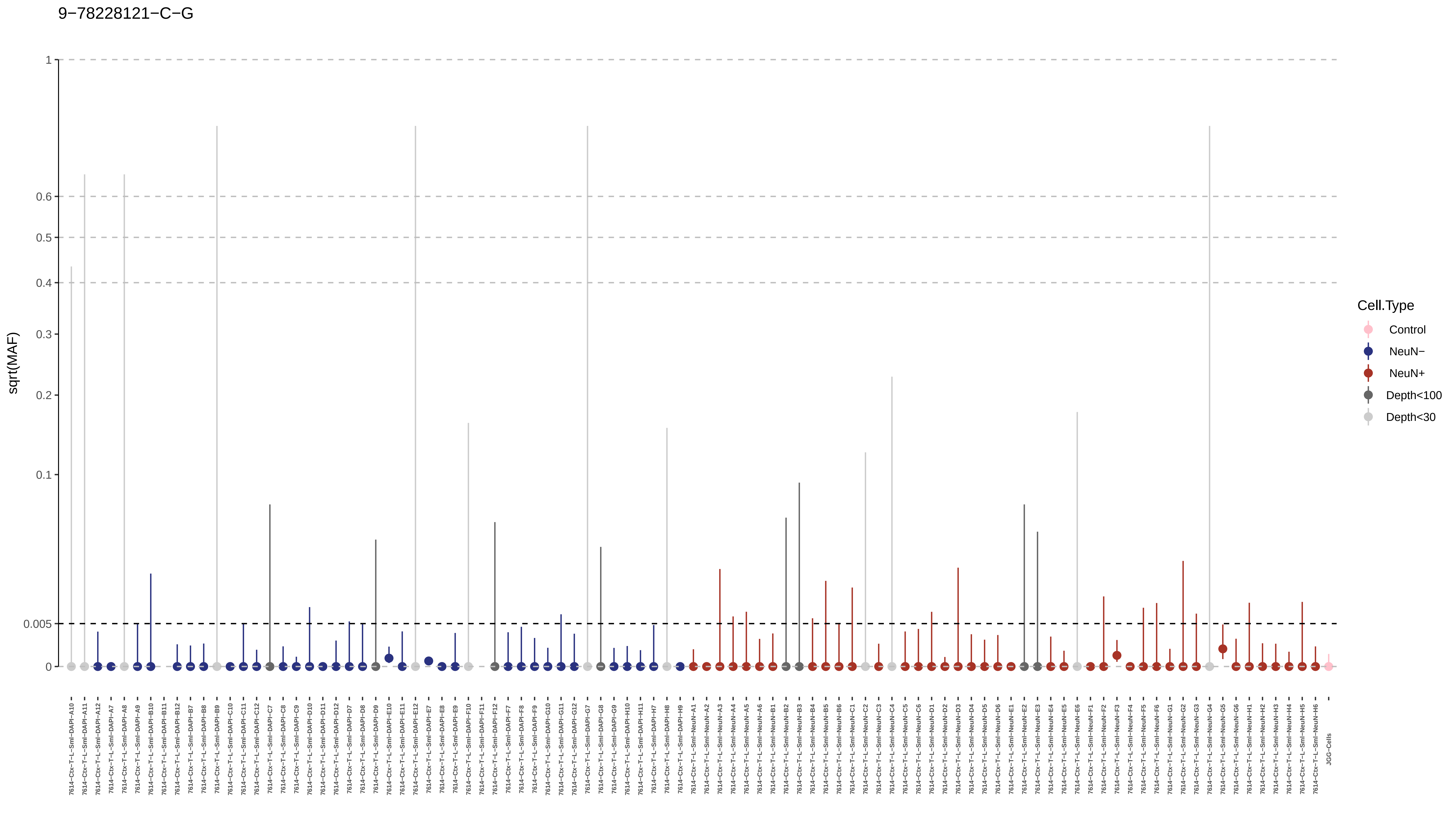
9−78228121−C−G
sqrt(MAF)
1
0.6
0.5
0.4
0.3
0.2
0.1
0.005
0
Cell.Type
Control
NeuN−
NeuN+
Depth<100
Depth<30
JGG-Cells

9−7860755−C−T

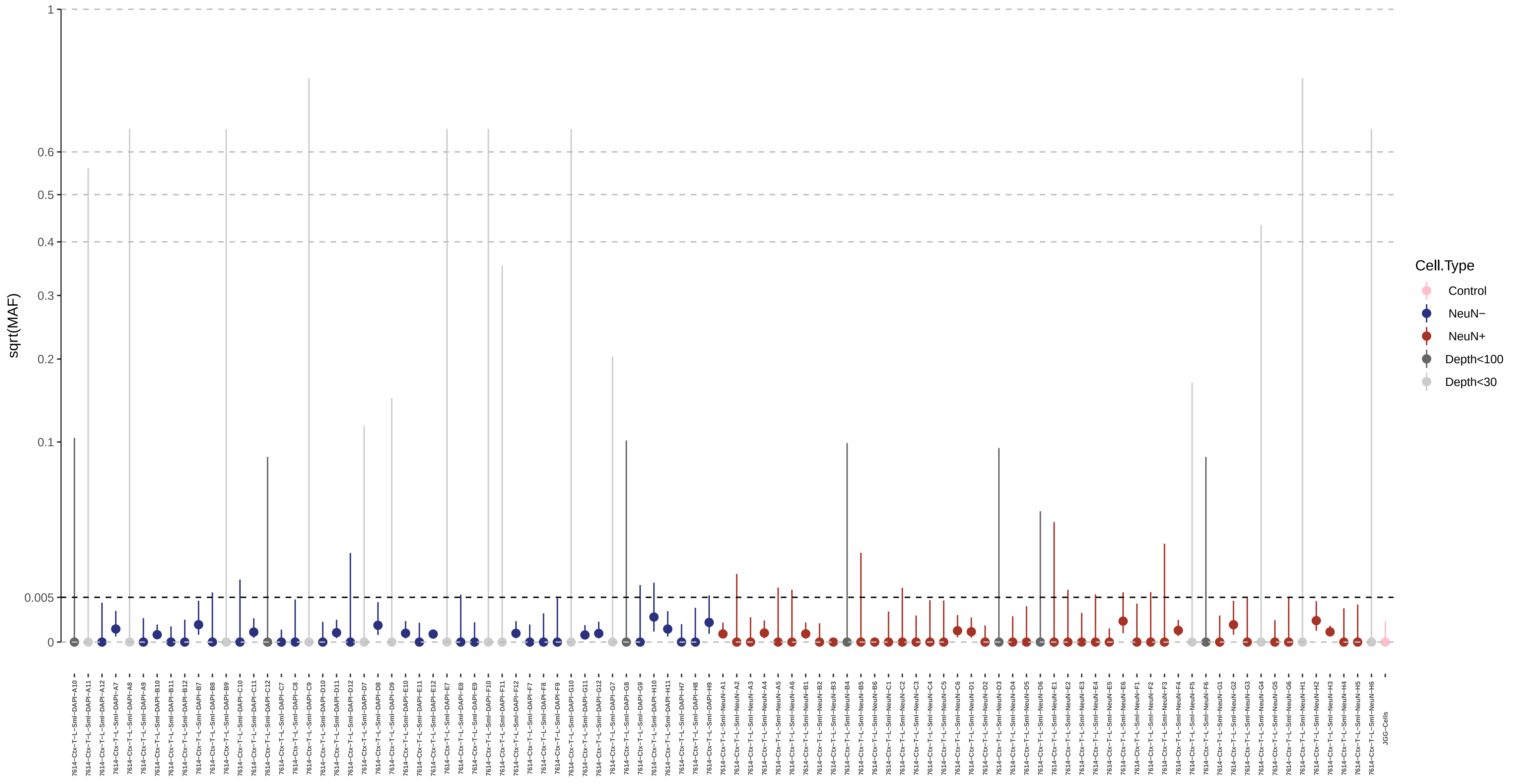

sqrt(MAF)
1
0.6
0.5
0.4
0.3
0.2
0.1
0.005
0
Cell.Type
Control
NeuN−
NeuN+
Depth<100
Depth<30

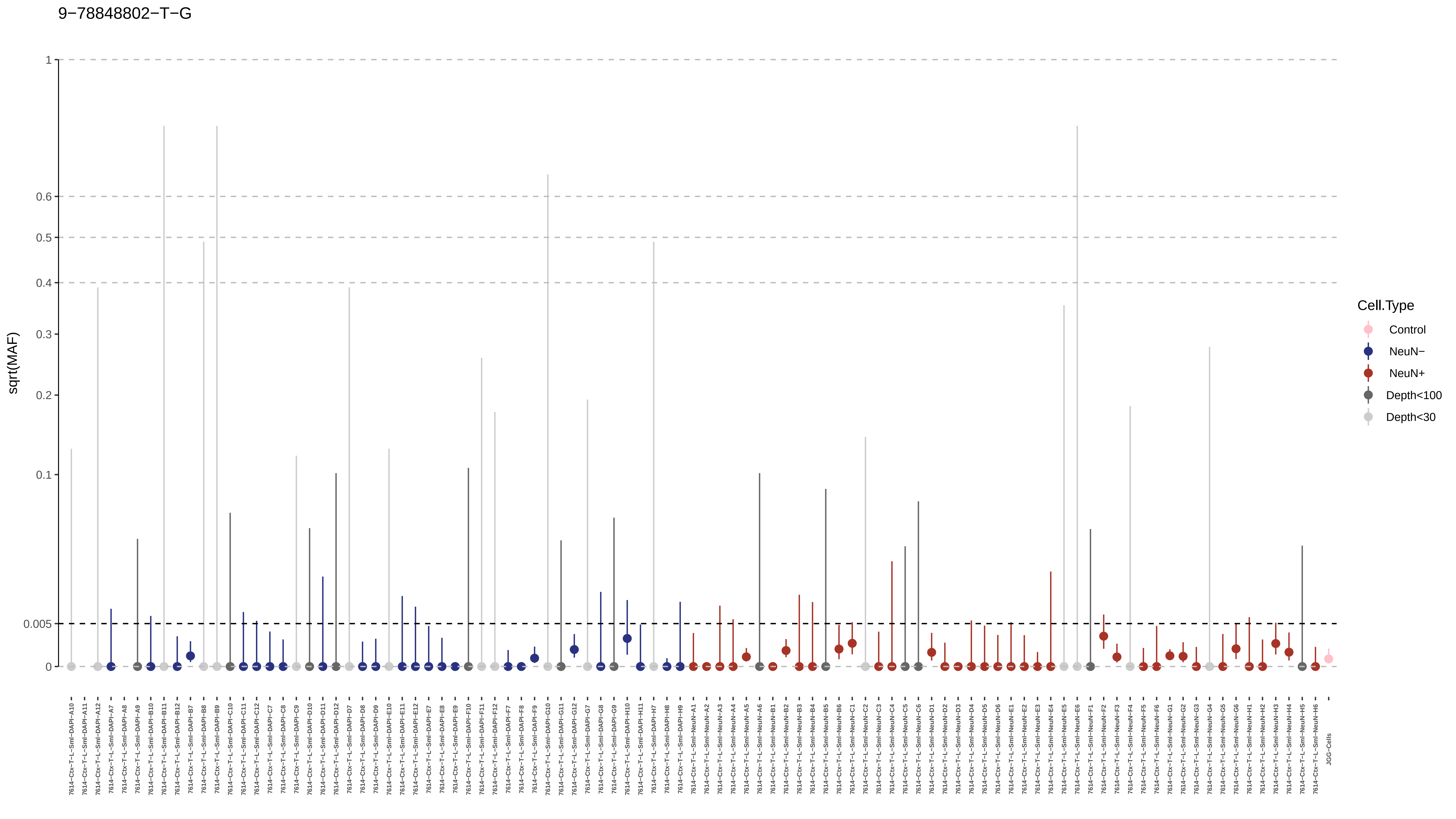

9−78848802−T−G
sqrt(MAF)
1
0.6
0.5
0.4
0.3
0.2
0.1
0.005
0
Cell.Type
Control
NeuN−
NeuN+
Depth<100
Depth<30

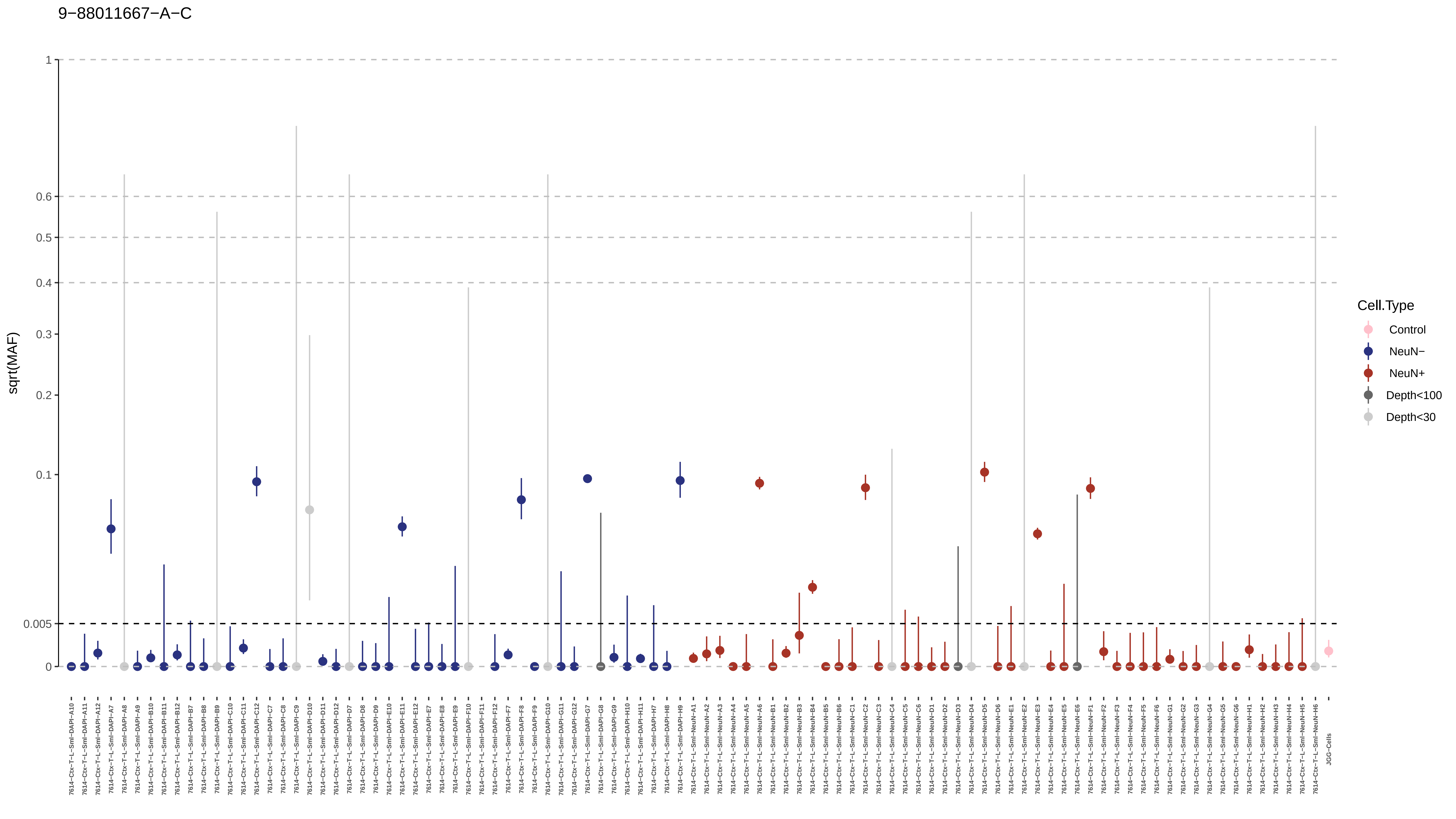
9−88011667−A−C
sqrt(MAF)
1
0.6
0.5
0.4
0.3
0.2
0.1
0.005
0
Cell.Type
Control
NeuN−
NeuN+
Depth<100
Depth<30

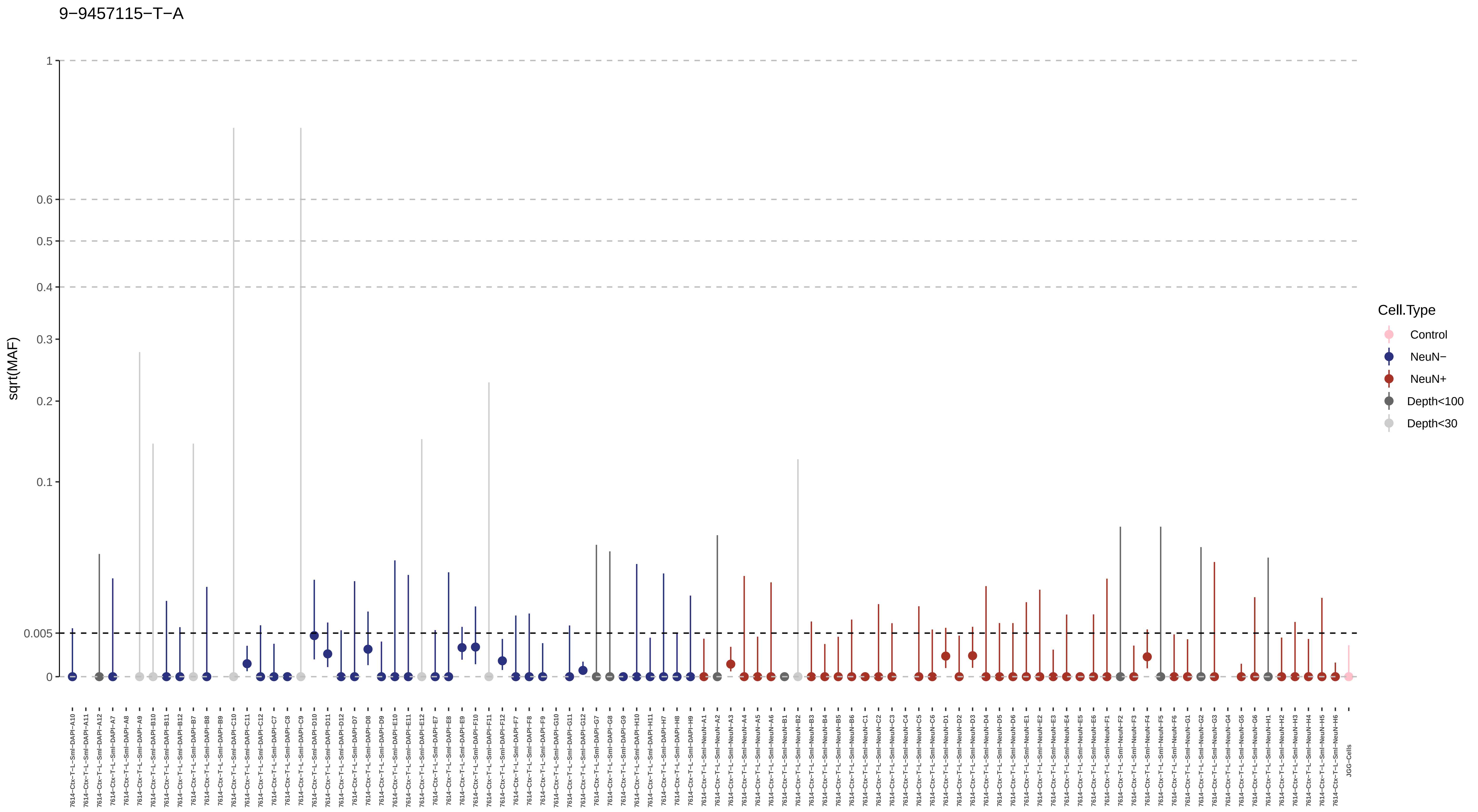
9−9457115−T−A
sqrt(MAF)
1
0.6
0.5
0.4
0.3
0.2
0.1
0.005
0
Cell.Type
Control
NeuN−
NeuN+
Depth<100
Depth<30
7614−Ctx−T−L−Sml−DAPI−A10
7614−Ctx−T−L−Sml−DAPI−A11
7614−Ctx−T−L−Sml−DAPI−A12
7614−Ctx−T−L−Sml−DAPI−A7
7614−Ctx−T−L−Sml−DAPI−A8
7614−Ctx−T−L−Sml−DAPI−A9
7614−Ctx−T−L−Sml−DAPI−B10
7614−Ctx−T−L−Sml−DAPI−B11
7614−Ctx−T−L−Sml−DAPI−B12
7614−Ctx−T−L−Sml−DAPI−B7
7614−Ctx−T−L−Sml−DAPI−B8
7614−Ctx−T−L−Sml−DAPI−B9
7614−Ctx−T−L−Sml−DAPI−C10
7614−Ctx−T−L−Sml−DAPI−C11
7614−Ctx−T−L−Sml−DAPI−C12
7614−Ctx−T−L−Sml−DAPI−C7
7614−Ctx−T−L−Sml−DAPI−C8
7614−Ctx−T−L−Sml−DAPI−C9
7614−Ctx−T−L−Sml−DAPI−D10
7614−Ctx−T−L−Sml−DAPI−D11
7614−Ctx−T−L−Sml−DAPI−D12
7614−Ctx−T−L−Sml−DAPI−D7
7614−Ctx−T−L−Sml−DAPI−D8
7614−Ctx−T−L−Sml−DAPI−D9
7614−Ctx−T−L−Sml−DAPI−E10
7614−Ctx−T−L−Sml−DAPI−E11
7614−Ctx−T−L−Sml−DAPI−E12
7614−Ctx−T−L−Sml−DAPI−E7
7614−Ctx−T−L−Sml−DAPI−E8
7614−Ctx−T−L−Sml−DAPI−E9
7614−Ctx−T−L−Sml−DAPI−F10
7614−Ctx−T−L−Sml−DAPI−F11
7614−Ctx−T−L−Sml−DAPI−F12
7614−Ctx−T−L−Sml−DAPI−F7
7614−Ctx−T−L−Sml−DAPI−F8
7614−Ctx−T−L−Sml−DAPI−F9
7614−Ctx−T−L−Sml−DAPI−G10
7614−Ctx−T−L−Sml−DAPI−G11
7614−Ctx−T−L−Sml−DAPI−G12
7614−Ctx−T−L−Sml−DAPI−G7
7614−Ctx−T−L−Sml−DAPI−G8
7614−Ctx−T−L−Sml−DAPI−G9
7614−Ctx−T−L−Sml−DAPI−H10
7614−Ctx−T−L−Sml−DAPI−H11
7614−Ctx−T−L−Sml−DAPI−H7
7614−Ctx−T−L−Sml−DAPI−H8
7614−Ctx−T−L−Sml−DAPI−H9
7614−Ctx−T−L−Sml−NeuN−A1
7614−Ctx−T−L−Sml−NeuN−A2
7614−Ctx−T−L−Sml−NeuN−A3
7614−Ctx−T−L−Sml−NeuN−A4
7614−Ctx−T−L−Sml−NeuN−A5
7614−Ctx−T−L−Sml−NeuN−A6
7614−Ctx−T−L−Sml−NeuN−B1
7614−Ctx−T−L−Sml−NeuN−B2
7614−Ctx−T−L−Sml−NeuN−B3
7614−Ctx−T−L−Sml−NeuN−B4
7614−Ctx−T−L−Sml−NeuN−B5
7614−Ctx−T−L−Sml−NeuN−B6
7614−Ctx−T−L−Sml−NeuN−C1
7614−Ctx−T−L−Sml−NeuN−C2
7614−Ctx−T−L−Sml−NeuN−C3
7614−Ctx−T−L−Sml−NeuN−C4
7614−Ctx−T−L−Sml−NeuN−C5
7614−Ctx−T−L−Sml−NeuN−C6
7614−Ctx−T−L−Sml−NeuN−D1
7614−Ctx−T−L−Sml−NeuN−D2
7614−Ctx−T−L−Sml−NeuN−D3
7614−Ctx−T−L−Sml−NeuN−D4
7614−Ctx−T−L−Sml−NeuN−D5
7614−Ctx−T−L−Sml−NeuN−D6
7614−Ctx−T−L−Sml−NeuN−E1
7614−Ctx−T−L−Sml−NeuN−E2
7614−Ctx−T−L−Sml−NeuN−E3
7614−Ctx−T−L−Sml−NeuN−E4
7614−Ctx−T−L−Sml−NeuN−E5
7614−Ctx−T−L−Sml−NeuN−E6
7614−Ctx−T−L−Sml−NeuN−F1
7614−Ctx−T−L−Sml−NeuN−F2
7614−Ctx−T−L−Sml−NeuN−F3
7614−Ctx−T−L−Sml−NeuN−F4
7614−Ctx−T−L−Sml−NeuN−F5
7614−Ctx−T−L−Sml−NeuN−F6
7614−Ctx−T−L−Sml−NeuN−G1
7614−Ctx−T−L−Sml−NeuN−G2
7614−Ctx−T−L−Sml−NeuN−G3
7614−Ctx−T−L−Sml−NeuN−G4
7614−Ctx−T−L−Sml−NeuN−G5
7614−Ctx−T−L−Sml−NeuN−G6
7614−Ctx−T−L−Sml−NeuN−H1
7614−Ctx−T−L−Sml−NeuN−H2
7614−Ctx−T−L−Sml−NeuN−H3
7614−Ctx−T−L−Sml−NeuN−H4
7614−Ctx−T−L−Sml−NeuN−H5
7614−Ctx−T−L−Sml−NeuN−H6
JGG-Cells

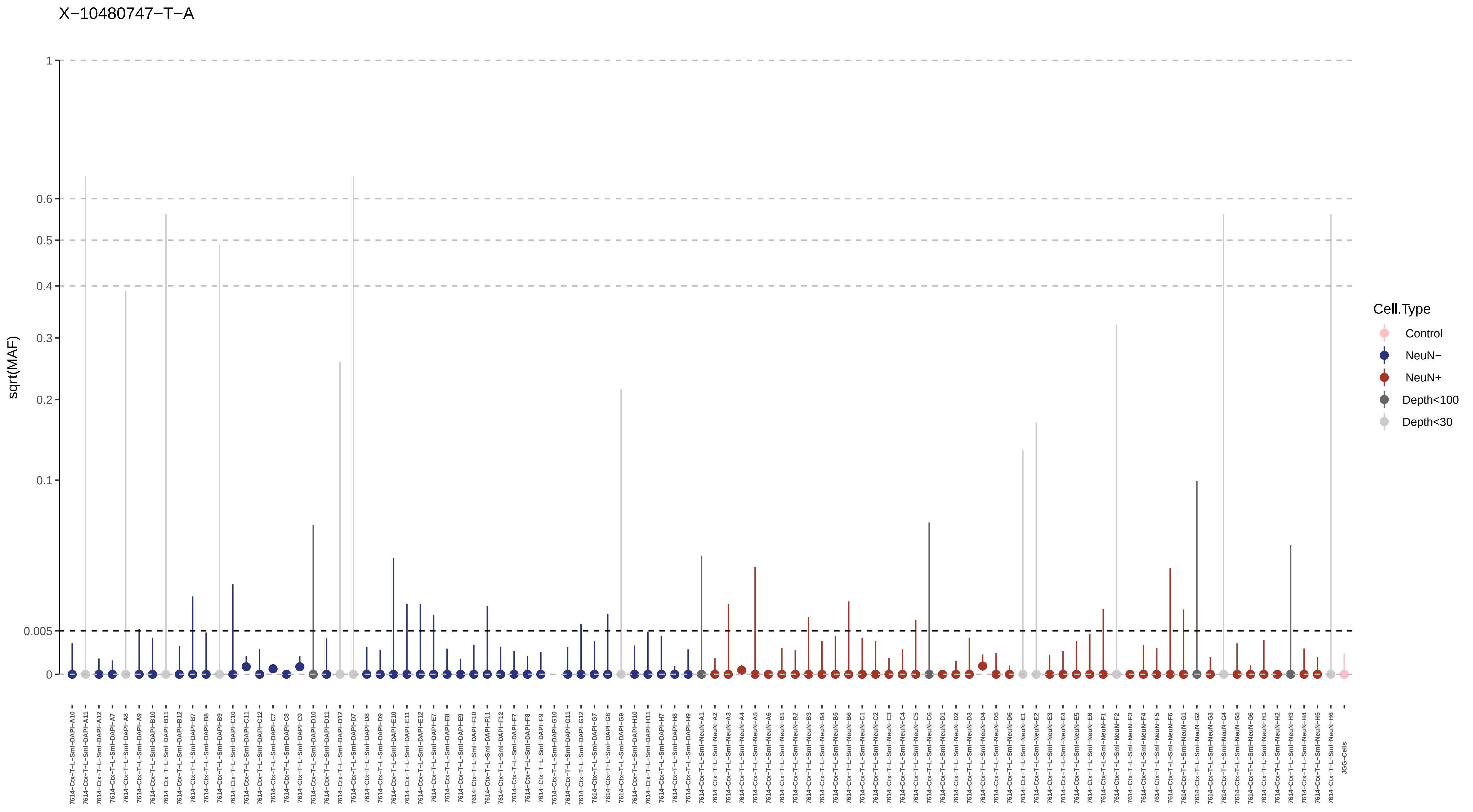

X−10480747−T−A
sqrt(MAF)
1
0.6
0.5
0.4
0.3
0.2
0.1
0.005
0
Cell.Type
Control
NeuN−
NeuN+
Depth<100
Depth<30

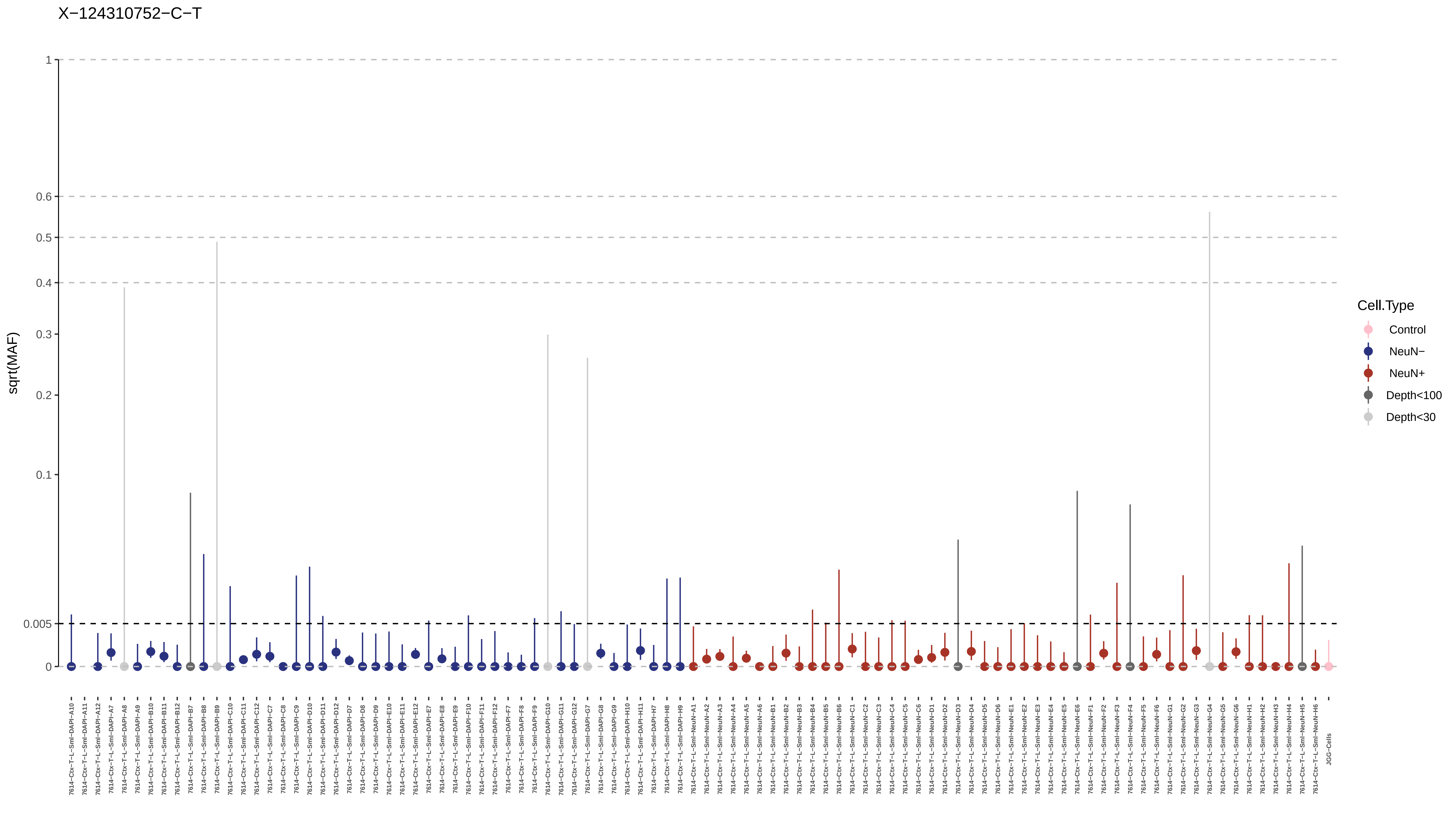
X−124310752−C−T
sqrt(MAF)
1
0.6
0.5
0.4
0.3
0.2
0.1
0.005
0
Cell.Type
Control
NeuN−
NeuN+
Depth<100
Depth<30
JGG-Cells

X−138175923−C−T

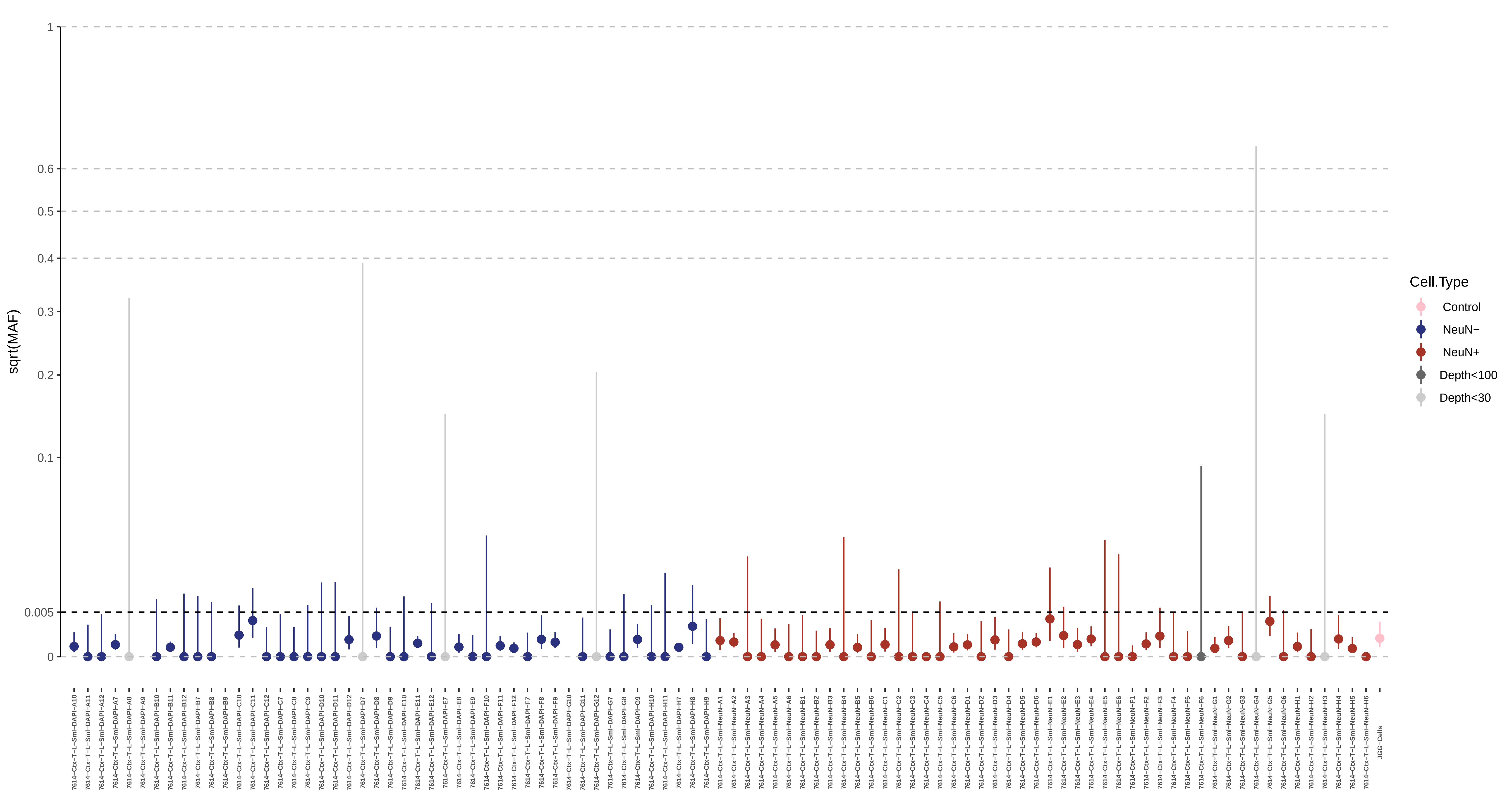

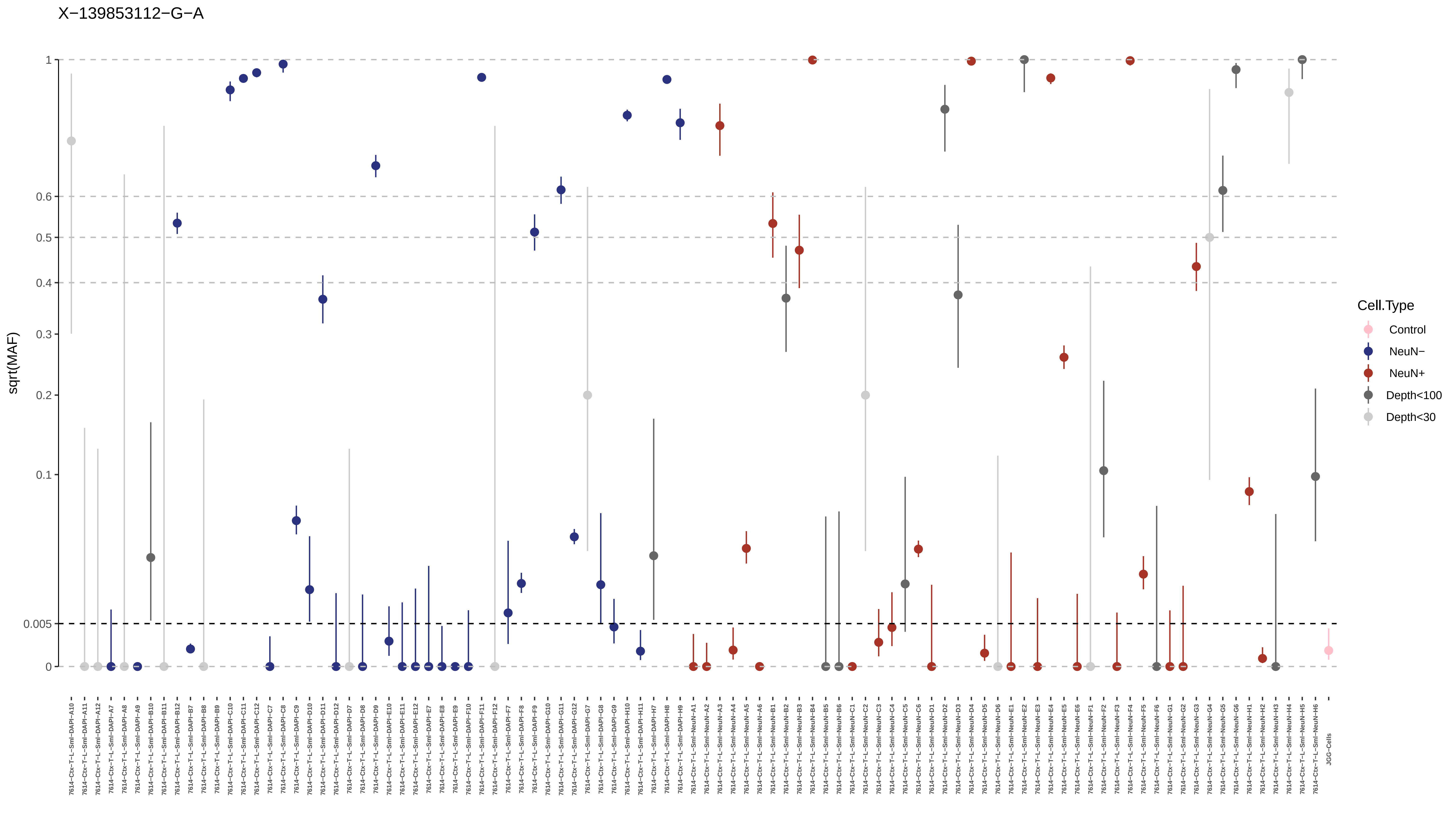
X−139853112−G−A
sqrt(MAF)
1
0.6
0.5
0.4
0.3
0.2
0.1
0.005
0
Cell.Type
Control
NeuN−
NeuN+
Depth<100
Depth<30

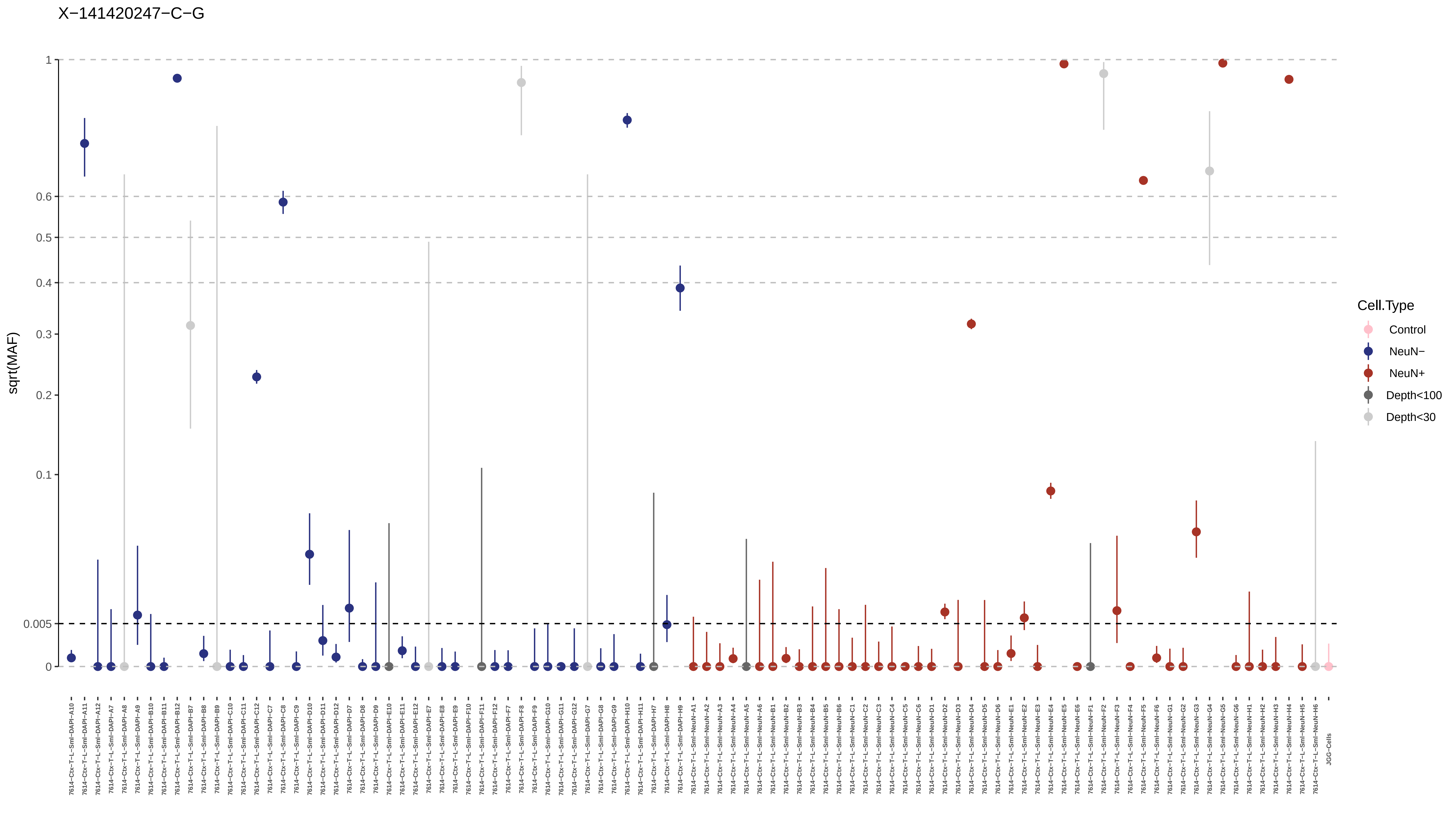
X−141420247−C−G
sqrt(MAF)
1
0.6
0.5
0.4
0.3
0.2
0.1
0.005
0
Cell.Type
Control
NeuN−
NeuN+
Depth<100
Depth<30
7614−Ctx−T−L−Sml−DAPI−A10
7614−Ctx−T−L−Sml−DAPI−A11
7614−Ctx−T−L−Sml−DAPI−A12
7614−Ctx−T−L−Sml−DAPI−A7
7614−Ctx−T−L−Sml−DAPI−A8
7614−Ctx−T−L−Sml−DAPI−A9
7614−Ctx−T−L−Sml−DAPI−B10
7614−Ctx−T−L−Sml−DAPI−B11
7614−Ctx−T−L−Sml−DAPI−B12
7614−Ctx−T−L−Sml−DAPI−B7
7614−Ctx−T−L−Sml−DAPI−B8
7614−Ctx−T−L−Sml−DAPI−B9
7614−Ctx−T−L−Sml−DAPI−C10
7614−Ctx−T−L−Sml−DAPI−C11
7614−Ctx−T−L−Sml−DAPI−C12
7614−Ctx−T−L−Sml−DAPI−C7
7614−Ctx−T−L−Sml−DAPI−C8
7614−Ctx−T−L−Sml−DAPI−C9
7614−Ctx−T−L−Sml−DAPI−D10
7614−Ctx−T−L−Sml−DAPI−D11
7614−Ctx−T−L−Sml−DAPI−D12
7614−Ctx−T−L−Sml−DAPI−D7
7614−Ctx−T−L−Sml−DAPI−D8
7614−Ctx−T−L−Sml−DAPI−D9
7614−Ctx−T−L−Sml−DAPI−E10
7614−Ctx−T−L−Sml−DAPI−E11
7614−Ctx−T−L−Sml−DAPI−E12
7614−Ctx−T−L−Sml−DAPI−E7
7614−Ctx−T−L−Sml−DAPI−E8
7614−Ctx−T−L−Sml−DAPI−E9
7614−Ctx−T−L−Sml−DAPI−F10
7614−Ctx−T−L−Sml−DAPI−F11
7614−Ctx−T−L−Sml−DAPI−F12
7614−Ctx−T−L−Sml−DAPI−F7
7614−Ctx−T−L−Sml−DAPI−F8
7614−Ctx−T−L−Sml−DAPI−F9
7614−Ctx−T−L−Sml−DAPI−G10
7614−Ctx−T−L−Sml−DAPI−G11
7614−Ctx−T−L−Sml−DAPI−G12
7614−Ctx−T−L−Sml−DAPI−G7
7614−Ctx−T−L−Sml−DAPI−G8
7614−Ctx−T−L−Sml−DAPI−G9
7614−Ctx−T−L−Sml−DAPI−H10
7614−Ctx−T−L−Sml−DAPI−H11
7614−Ctx−T−L−Sml−DAPI−H7
7614−Ctx−T−L−Sml−DAPI−H8
7614−Ctx−T−L−Sml−DAPI−H9
7614−Ctx−T−L−Sml−NeuN−A1
7614−Ctx−T−L−Sml−NeuN−A2
7614−Ctx−T−L−Sml−NeuN−A3
7614−Ctx−T−L−Sml−NeuN−A4
7614−Ctx−T−L−Sml−NeuN−A5
7614−Ctx−T−L−Sml−NeuN−A6
7614−Ctx−T−L−Sml−NeuN−B1
7614−Ctx−T−L−Sml−NeuN−B2
7614−Ctx−T−L−Sml−NeuN−B3
7614−Ctx−T−L−Sml−NeuN−B4
7614−Ctx−T−L−Sml−NeuN−B5
7614−Ctx−T−L−Sml−NeuN−B6
7614−Ctx−T−L−Sml−NeuN−C1
7614−Ctx−T−L−Sml−NeuN−C2
7614−Ctx−T−L−Sml−NeuN−C3
7614−Ctx−T−L−Sml−NeuN−C4
7614−Ctx−T−L−Sml−NeuN−C5
7614−Ctx−T−L−Sml−NeuN−C6
7614−Ctx−T−L−Sml−NeuN−D1
7614−Ctx−T−L−Sml−NeuN−D2
7614−Ctx−T−L−Sml−NeuN−D3
7614−Ctx−T−L−Sml−NeuN−D4
7614−Ctx−T−L−Sml−NeuN−D5
7614−Ctx−T−L−Sml−NeuN−D6
7614−Ctx−T−L−Sml−NeuN−E1
7614−Ctx−T−L−Sml−NeuN−E2
7614−Ctx−T−L−Sml−NeuN−E3
7614−Ctx−T−L−Sml−NeuN−E4
7614−Ctx−T−L−Sml−NeuN−E5
7614−Ctx−T−L−Sml−NeuN−E6
7614−Ctx−T−L−Sml−NeuN−F1
7614−Ctx−T−L−Sml−NeuN−F2
7614−Ctx−T−L−Sml−NeuN−F3
7614−Ctx−T−L−Sml−NeuN−F4
7614−Ctx−T−L−Sml−NeuN−F5
7614−Ctx−T−L−Sml−NeuN−F6
7614−Ctx−T−L−Sml−NeuN−G1
7614−Ctx−T−L−Sml−NeuN−G2
7614−Ctx−T−L−Sml−NeuN−G3
7614−Ctx−T−L−Sml−NeuN−G4
7614−Ctx−T−L−Sml−NeuN−G5
7614−Ctx−T−L−Sml−NeuN−G6
7614−Ctx−T−L−Sml−NeuN−H1
7614−Ctx−T−L−Sml−NeuN−H2
7614−Ctx−T−L−Sml−NeuN−H3
7614−Ctx−T−L−Sml−NeuN−H4
7614−Ctx−T−L−Sml−NeuN−H5
7614−Ctx−T−L−Sml−NeuN−H6
JGG-Cells

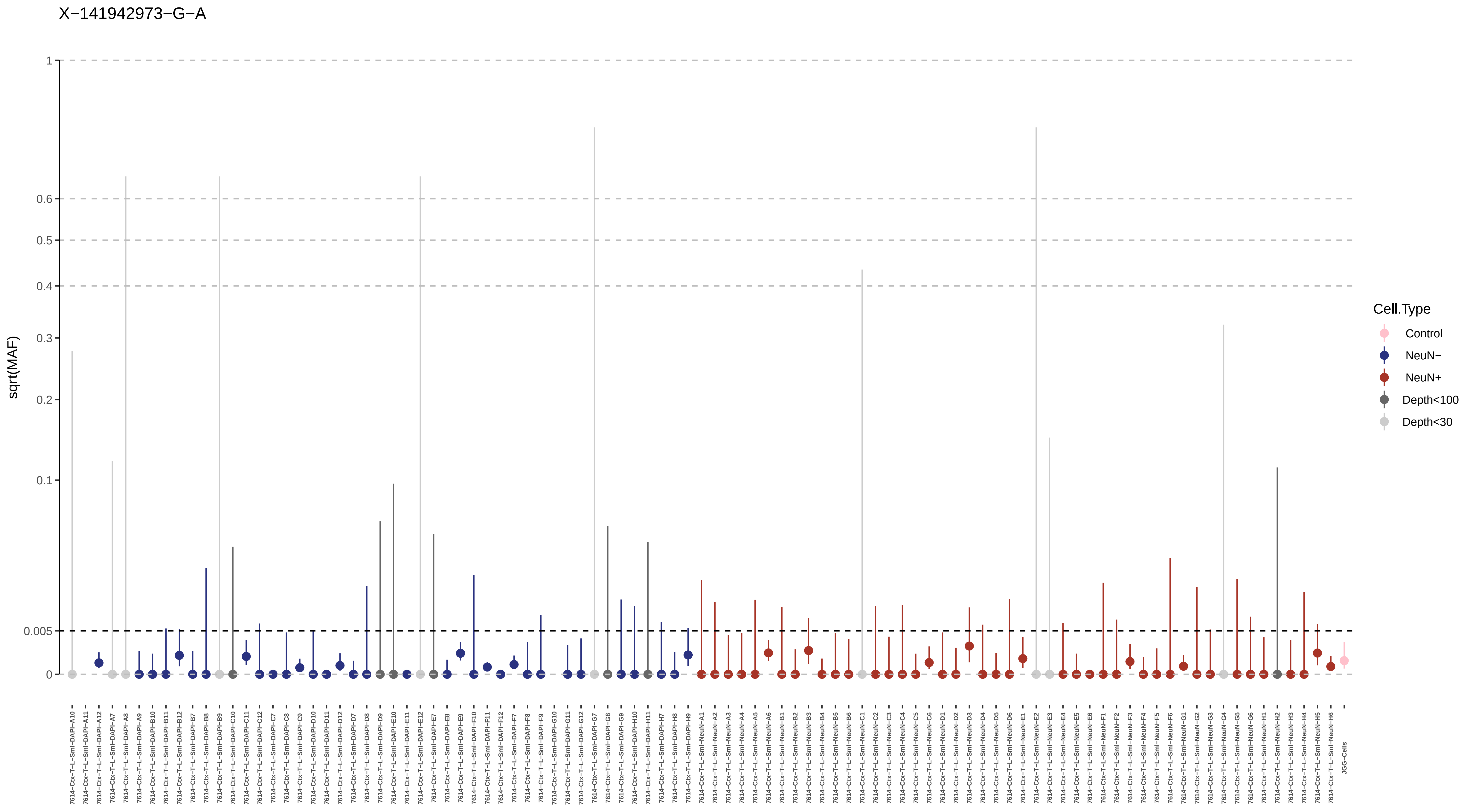
X−141942973−G−A
sqrt(MAF)
1
0.6
0.5
0.4
0.3
0.2
0.1
0.005
0
Cell.Type
Control
NeuN−
NeuN+
Depth<100
Depth<30

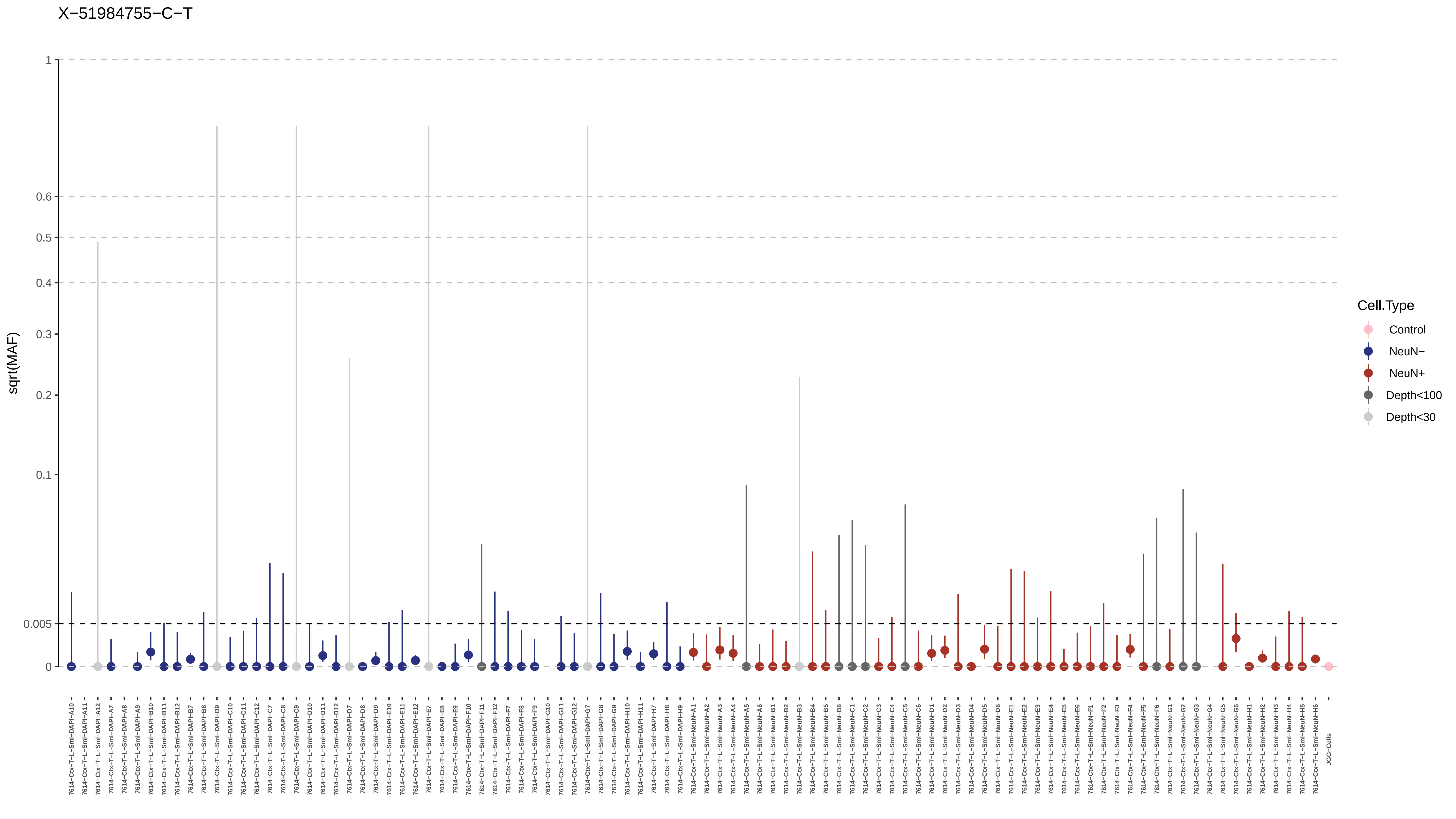

X−51984755−C−T
sqrt(MAF)
1
0.6
0.5
0.4
0.3
0.2
0.1
0.005
0
Cell.Type
Control
NeuN−
NeuN+
Depth<100
Depth<30
7614−Ctx−T−L−Sml−DAPI−A10
7614−Ctx−T−L−Sml−DAPI−A11
7614−Ctx−T−L−Sml−DAPI−A12
7614−Ctx−T−L−Sml−DAPI−A7
7614−Ctx−T−L−Sml−DAPI−A8
7614−Ctx−T−L−Sml−DAPI−A9
7614−Ctx−T−L−Sml−DAPI−B10
7614−Ctx−T−L−Sml−DAPI−B11
7614−Ctx−T−L−Sml−DAPI−B12
7614−Ctx−T−L−Sml−DAPI−B7
7614−Ctx−T−L−Sml−DAPI−B8
7614−Ctx−T−L−Sml−DAPI−B9
7614−Ctx−T−L−Sml−DAPI−C10
7614−Ctx−T−L−Sml−DAPI−C11
7614−Ctx−T−L−Sml−DAPI−C12
7614−Ctx−T−L−Sml−DAPI−C7
7614−Ctx−T−L−Sml−DAPI−C8
7614−Ctx−T−L−Sml−DAPI−C9
7614−Ctx−T−L−Sml−DAPI−D10
7614−Ctx−T−L−Sml−DAPI−D11
7614−Ctx−T−L−Sml−DAPI−D12
7614−Ctx−T−L−Sml−DAPI−D7
7614−Ctx−T−L−Sml−DAPI−D8
7614−Ctx−T−L−Sml−DAPI−D9
7614−Ctx−T−L−Sml−DAPI−E10
7614−Ctx−T−L−Sml−DAPI−E11
7614−Ctx−T−L−Sml−DAPI−E12
7614−Ctx−T−L−Sml−DAPI−E7
7614−Ctx−T−L−Sml−DAPI−E8
7614−Ctx−T−L−Sml−DAPI−E9
7614−Ctx−T−L−Sml−DAPI−F10
7614−Ctx−T−L−Sml−DAPI−F11
7614−Ctx−T−L−Sml−DAPI−F12
7614−Ctx−T−L−Sml−DAPI−F7
7614−Ctx−T−L−Sml−DAPI−F8
7614−Ctx−T−L−Sml−DAPI−F9
7614−Ctx−T−L−Sml−DAPI−G10
7614−Ctx−T−L−Sml−DAPI−G11
7614−Ctx−T−L−Sml−DAPI−G12
7614−Ctx−T−L−Sml−DAPI−G7
7614−Ctx−T−L−Sml−DAPI−G8
7614−Ctx−T−L−Sml−DAPI−G9
7614−Ctx−T−L−Sml−DAPI−H10
7614−Ctx−T−L−Sml−DAPI−H11
7614−Ctx−T−L−Sml−DAPI−H7
7614−Ctx−T−L−Sml−DAPI−H8
7614−Ctx−T−L−Sml−DAPI−H9
7614−Ctx−T−L−Sml−NeuN−A1
7614−Ctx−T−L−Sml−NeuN−A2
7614−Ctx−T−L−Sml−NeuN−A3
7614−Ctx−T−L−Sml−NeuN−A4
7614−Ctx−T−L−Sml−NeuN−A5
7614−Ctx−T−L−Sml−NeuN−A6
7614−Ctx−T−L−Sml−NeuN−B1
7614−Ctx−T−L−Sml−NeuN−B2
7614−Ctx−T−L−Sml−NeuN−B3
7614−Ctx−T−L−Sml−NeuN−B4
7614−Ctx−T−L−Sml−NeuN−B5
7614−Ctx−T−L−Sml−NeuN−B6
7614−Ctx−T−L−Sml−NeuN−C1
7614−Ctx−T−L−Sml−NeuN−C2
7614−Ctx−T−L−Sml−NeuN−C3
7614−Ctx−T−L−Sml−NeuN−C4
7614−Ctx−T−L−Sml−NeuN−C5
7614−Ctx−T−L−Sml−NeuN−C6
7614−Ctx−T−L−Sml−NeuN−D1
7614−Ctx−T−L−Sml−NeuN−D2
7614−Ctx−T−L−Sml−NeuN−D3
7614−Ctx−T−L−Sml−NeuN−D4
7614−Ctx−T−L−Sml−NeuN−D5
7614−Ctx−T−L−Sml−NeuN−D6
7614−Ctx−T−L−Sml−NeuN−E1
7614−Ctx−T−L−Sml−NeuN−E2
7614−Ctx−T−L−Sml−NeuN−E3
7614−Ctx−T−L−Sml−NeuN−E4
7614−Ctx−T−L−Sml−NeuN−E5
7614−Ctx−T−L−Sml−NeuN−E6
7614−Ctx−T−L−Sml−NeuN−F1
7614−Ctx−T−L−Sml−NeuN−F2
7614−Ctx−T−L−Sml−NeuN−F3
7614−Ctx−T−L−Sml−NeuN−F4
7614−Ctx−T−L−Sml−NeuN−F5
7614−Ctx−T−L−Sml−NeuN−F6
7614−Ctx−T−L−Sml−NeuN−G1
7614−Ctx−T−L−Sml−NeuN−G2
7614−Ctx−T−L−Sml−NeuN−G3
7614−Ctx−T−L−Sml−NeuN−G4
7614−Ctx−T−L−Sml−NeuN−G5
7614−Ctx−T−L−Sml−NeuN−G6
7614−Ctx−T−L−Sml−NeuN−H1
7614−Ctx−T−L−Sml−NeuN−H2
7614−Ctx−T−L−Sml−NeuN−H3
7614−Ctx−T−L−Sml−NeuN−H4
7614−Ctx−T−L−Sml−NeuN−H5
7614−Ctx−T−L−Sml−NeuN−H6
JGG−Cells

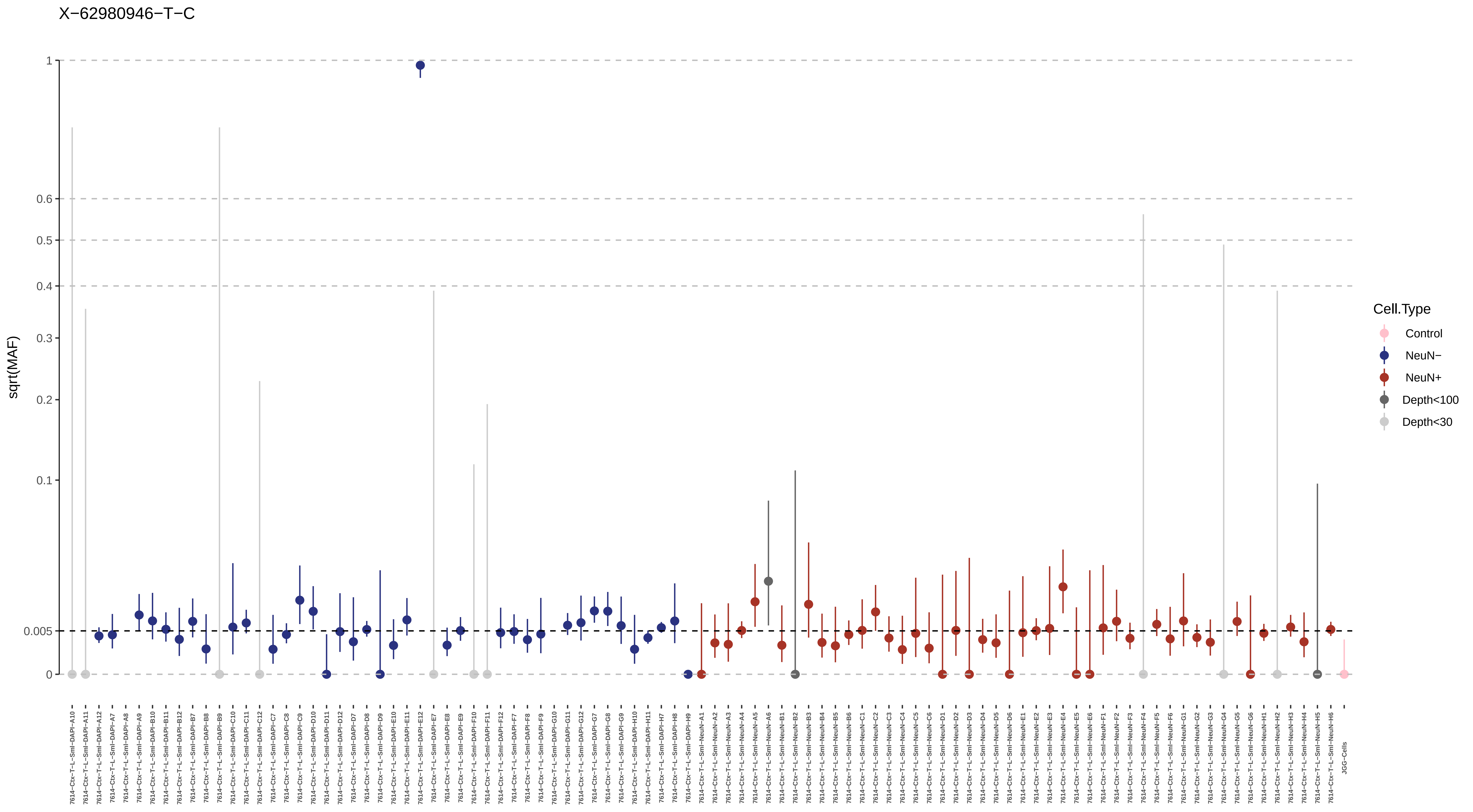
X−62980946−T−C
sqrt(MAF)
1
0.6
0.5
0.4
0.3
0.2
0.1
0.005
0
Cell.Type
Control
NeuN−
NeuN+
Depth<100
Depth<30
7614−Ctx−T−L−Sml−DAPI−A10
7614−Ctx−T−L−Sml−DAPI−A11
7614−Ctx−T−L−Sml−DAPI−A12
7614−Ctx−T−L−Sml−DAPI−A7
7614−Ctx−T−L−Sml−DAPI−A8
7614−Ctx−T−L−Sml−DAPI−A9
7614−Ctx−T−L−Sml−DAPI−B10
7614−Ctx−T−L−Sml−DAPI−B11
7614−Ctx−T−L−Sml−DAPI−B12
7614−Ctx−T−L−Sml−DAPI−B7
7614−Ctx−T−L−Sml−DAPI−B8
7614−Ctx−T−L−Sml−DAPI−B9
7614−Ctx−T−L−Sml−DAPI−C10
7614−Ctx−T−L−Sml−DAPI−C11
7614−Ctx−T−L−Sml−DAPI−C12
7614−Ctx−T−L−Sml−DAPI−C7
7614−Ctx−T−L−Sml−DAPI−C8
7614−Ctx−T−L−Sml−DAPI−C9
7614−Ctx−T−L−Sml−DAPI−D10
7614−Ctx−T−L−Sml−DAPI−D11
7614−Ctx−T−L−Sml−DAPI−D12
7614−Ctx−T−L−Sml−DAPI−D7
7614−Ctx−T−L−Sml−DAPI−D8
7614−Ctx−T−L−Sml−DAPI−D9
7614−Ctx−T−L−Sml−DAPI−E10
7614−Ctx−T−L−Sml−DAPI−E11
7614−Ctx−T−L−Sml−DAPI−E12
7614−Ctx−T−L−Sml−DAPI−E7
7614−Ctx−T−L−Sml−DAPI−E8
7614−Ctx−T−L−Sml−DAPI−E9
7614−Ctx−T−L−Sml−DAPI−F10
7614−Ctx−T−L−Sml−DAPI−F11
7614−Ctx−T−L−Sml−DAPI−F12
7614−Ctx−T−L−Sml−DAPI−F7
7614−Ctx−T−L−Sml−DAPI−F8
7614−Ctx−T−L−Sml−DAPI−F9
7614−Ctx−T−L−Sml−DAPI−G10
7614−Ctx−T−L−Sml−DAPI−G11
7614−Ctx−T−L−Sml−DAPI−G12
7614−Ctx−T−L−Sml−DAPI−G7
7614−Ctx−T−L−Sml−DAPI−G8
7614−Ctx−T−L−Sml−DAPI−G9
7614−Ctx−T−L−Sml−DAPI−H10
7614−Ctx−T−L−Sml−DAPI−H11
7614−Ctx−T−L−Sml−DAPI−H7
7614−Ctx−T−L−Sml−DAPI−H8
7614−Ctx−T−L−Sml−DAPI−H9
7614−Ctx−T−L−Sml−NeuN−A1
7614−Ctx−T−L−Sml−NeuN−A2
7614−Ctx−T−L−Sml−NeuN−A3
7614−Ctx−T−L−Sml−NeuN−A4
7614−Ctx−T−L−Sml−NeuN−A5
7614−Ctx−T−L−Sml−NeuN−A6
7614−Ctx−T−L−Sml−NeuN−B1
7614−Ctx−T−L−Sml−NeuN−B2
7614−Ctx−T−L−Sml−NeuN−B3
7614−Ctx−T−L−Sml−NeuN−B4
7614−Ctx−T−L−Sml−NeuN−B5
7614−Ctx−T−L−Sml−NeuN−B6
7614−Ctx−T−L−Sml−NeuN−C1
7614−Ctx−T−L−Sml−NeuN−C2
7614−Ctx−T−L−Sml−NeuN−C3
7614−Ctx−T−L−Sml−NeuN−C4
7614−Ctx−T−L−Sml−NeuN−C5
7614−Ctx−T−L−Sml−NeuN−C6
7614−Ctx−T−L−Sml−NeuN−D1
7614−Ctx−T−L−Sml−NeuN−D2
7614−Ctx−T−L−Sml−NeuN−D3
7614−Ctx−T−L−Sml−NeuN−D4
7614−Ctx−T−L−Sml−NeuN−D5
7614−Ctx−T−L−Sml−NeuN−D6
7614−Ctx−T−L−Sml−NeuN−E1
7614−Ctx−T−L−Sml−NeuN−E2
7614−Ctx−T−L−Sml−NeuN−E3
7614−Ctx−T−L−Sml−NeuN−E4
7614−Ctx−T−L−Sml−NeuN−E5
7614−Ctx−T−L−Sml−NeuN−E6
7614−Ctx−T−L−Sml−NeuN−F1
7614−Ctx−T−L−Sml−NeuN−F2
7614−Ctx−T−L−Sml−NeuN−F3
7614−Ctx−T−L−Sml−NeuN−F4
7614−Ctx−T−L−Sml−NeuN−F5
7614−Ctx−T−L−Sml−NeuN−F6
7614−Ctx−T−L−Sml−NeuN−G1
7614−Ctx−T−L−Sml−NeuN−G2
7614−Ctx−T−L−Sml−NeuN−G3
7614−Ctx−T−L−Sml−NeuN−G4
7614−Ctx−T−L−Sml−NeuN−G5
7614−Ctx−T−L−Sml−NeuN−G6
7614−Ctx−T−L−Sml−NeuN−H1
7614−Ctx−T−L−Sml−NeuN−H2
7614−Ctx−T−L−Sml−NeuN−H3
7614−Ctx−T−L−Sml−NeuN−H4
7614−Ctx−T−L−Sml−NeuN−H5
7614−Ctx−T−L−Sml−NeuN−H6
JGG-Cells

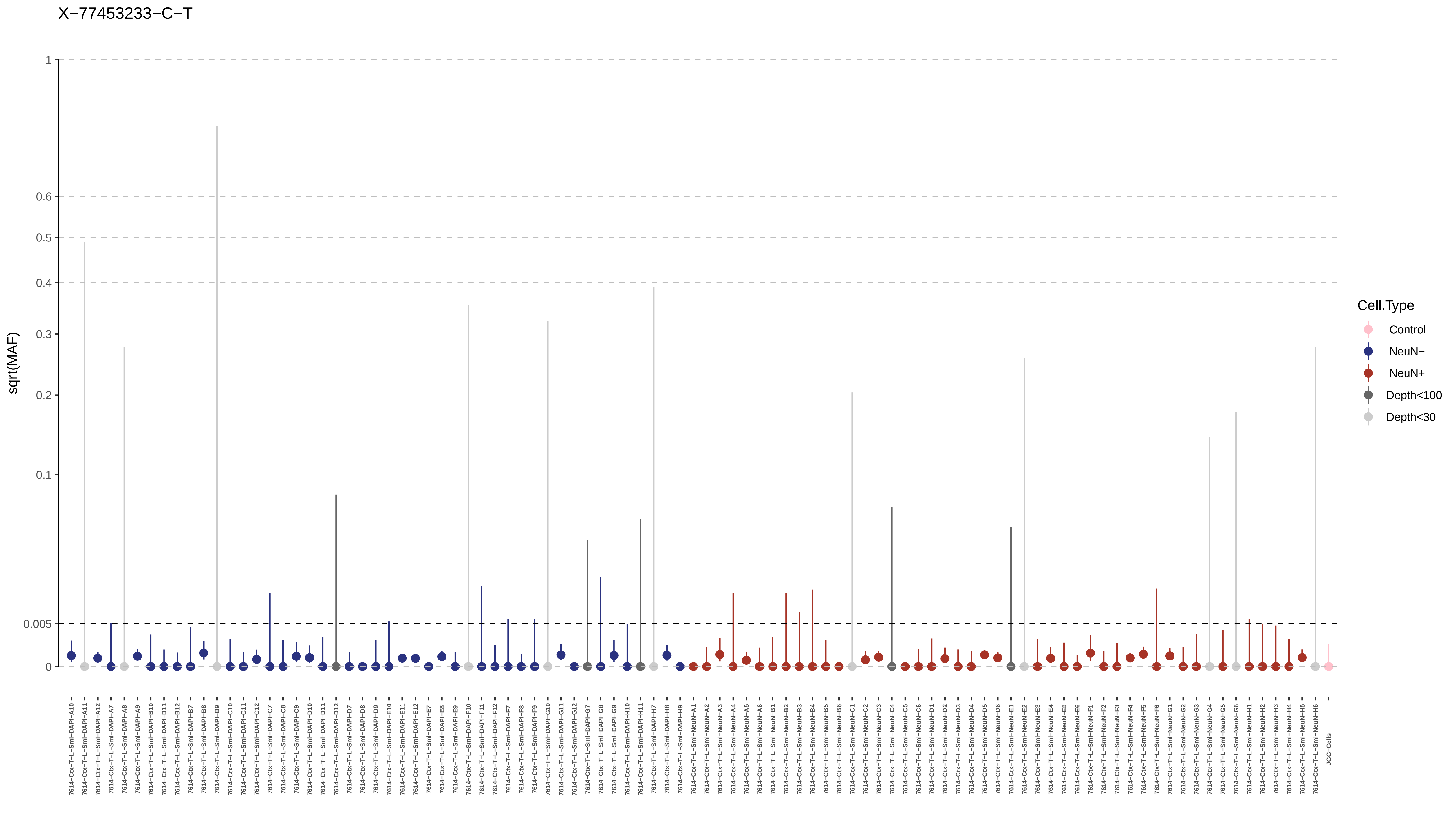
X−77453233−C−T
sqrt(MAF)
1
0.6
0.5
0.4
0.3
0.2
0.1
0.005
0
Cell.Type
Control
NeuN−
NeuN+
Depth<100
Depth<30
JGG-Cells
