## Supplementary Materials for "Somatic mosaicism in the mature brain reveals clonal cellular distributions during cortical development"

⁑Full membership of the Brain Somatic Mosaicism Network is listed in the Supplementary Text.

**This PDF file include s:**

Materials and Methods
Supplementary Text
Figs. S1 to S20
Data S1 to S4 (Legends)

**Other Supplementary Materials for this manuscript include the following:**

Data S1 to S4
Data S1. Raw mosaic SNV/INDEL calls from the 300× WGS.
Data S2. MPAS and snMPAS genotyping and quantification results.
Data S3. Detailed visual representation for each of the 259 variants.
Data S4. Visual representation of snMPAS results for each of the 259 variants.

### Materials and Methods

Sample recruitment and tissue dissection

Whole brain, heart, liver and both kidneys were provided by the UC San Diego Anatomical Material Program (Case Number UCSD-19-110). Organs were donated by a 70-year-old Caucasian female. Prior medical history showed no signs of neurological diseases. Noted cause of death was ‘global geriatric decline’ with a contributing cause of ‘post-surgical malabsorption’. Organs were collected within a ~24-hour postmortem interval. Briefly, after the removal of the meninges, diencephalon regions, and brain stem the cerebral cortical regions including prefrontal lobes, frontal lobes, parietal lobes, occipital lobes, temporal lobes, and cerebellum were dissected by a pathologist. 13 subsamples were collected from each lobe with 8 mm diameter disposable punches and disposable scalpels; the thickness of each biopsy was <1 cm. Two punches were collected from each of the non-neocortical tissues: two from each hemisphere of the cerebellum, two from each side of the heart, two from the liver, and two from each side of the cortex region of each kidney. After dissection, subsamples and the remnants of the large pieces were immediately put on dry ice and stored at -80°C.

Tissue homogenization and nuclei extraction

Frozen brain lobar samples (after the removal of the 8 mm biopsies; i.e. Lrg) were ground up in liquid nitrogen, then homogenized in 1% formaldehyde in Dulbecco’s phosphate buffered saline (PBS, Corning) using a motorized homogenizer (Fisherbrand PowerGen 125), and finally incubated on a rocker at room temperature for 10 minutes. Fixed homogenates were quenched with 0.125 M glycine at room temperature on a rocker for 5 minutes. Next, homogenates were centrifuged at 1,100 xg in a swinging bucket centrifuge. The following steps were all performed on ice except where indicated. Homogenates were washed twice with NF1 buffer (10 mM Tris-HCl pH 8.0, 1 mM EDTA, 5mM $MgCl_2$, 0.1M sucrose, 0.5% Triton X-100 in UltraPure water) and centrifuged at 1,100 xg for 5 minutes at 4°C in a swinging bucket centrifuge. Next, pellets were resuspended in 5 ml NF1 buffer and dounced five times in a 7 ml Wheaton Dounce Tissue Grinder (DWK Life Sciences) using a ‘loose’ pestle. After 30 minutes incubation on ice, homogenates were dounced 20 times with a ‘tight’ pestle and filtered through a 70 µm strainer. To remove myelin debris, homogenates were underlaid with a sucrose cushion (1.2M sucrose, 1 M Tris-HCl pH 8.0, 1 mM $MgCl_2$, 0.1 M DTT) and centrifuged at 3,200 xg for 30 minutes with acceleration and brakes on ‘low’. Pellets of nuclei were washed with NF1 buffer and centrifuged at 1,600 xg for 5 minutes and stored at -80°C.

DNA extraction of bulk tissue and nuclear fractions

Small cortical biopsies were first cut in half on dry ice. Half of the biopsy was stored as backup and partly used for single nuclei fluorescence-activated nuclei sorting (FANS). The other half of cortical biopsies was homogenized with a Pellet Pestle Motor (Kimble, #749540-0000) and resuspended with 450 µL RLT buffer (Qiagen, #40724) in a 1.5 ml microcentrifuge tube (USA Scientific, #1615-5500). The same experimental procedure was carried out on both punches from cerebellum, heart, liver, and both kidneys. Nuclear preparations were pelleted at 1,000 xg for 5 minutes and resuspended with 450 µL RLT buffer in a 1.5 ml microcentrifuge tube. Both homogenates and nuclear preps were then treated with the same protocol: following vortexing for 1 minute, samples were incubated at 70°C for 30 minutes. 50 µl Bond-Breaker TCEP solution (Thermo Scientific, #77720) and 120 mg stainless steel beads with 0.2 mm diameter (Next Advance, #SSB02) were added and cellular/nuclear disruption was performed for

5 minutes on a DisruptorGenie (Scientific industries). Supernatant was transferred to a DNA Mini Column from an AllPrep DNA/RNA Mini Kit (Qiagen, #80204) and centrifuged at 8500 xg for 30 seconds. The column was then washed with Buffer AW1 (kit-supplied), centrifuged at 8500 xg for 30 seconds and washed again with Buffer AW2 (kit-supplied), and then centrifuged at full speed for 2 minutes. The DNA was eluted two times with 50 µl of pre-heated (70°C) EB (kit-supplied) through centrifugation at 8,500 xg for 1 minute.

Whole-genome library preparation and sequencing

A total of 1.0 µg of extracted DNA was used as the starting material for PCR-free library construction using the KAPA HyperPrep PCR-Free Library Prep kit (Roche, #KK8505). Mechanical shearing using the Covaris microtube system (Covaris, # SKU 520053) was performed to generate fragments with peak size ~400 base pairs (bp). Each fragmented DNA sample went through multiple enzymatic reactions to generate a library in which an Illumina dual index adapter would be ligated to the DNA fragments. Beads-based double size selection was performed to ensure the fragment size of each sample was between 300-600 bp as measured by an Agilent DNA High Sensitivity NGS Fragment Analysis Kit (Agilent, #DNF-474-0500). The concentration of ligated fragments in each library was quantified with the KAPA Library Quantification Kits for Illumina platforms (Roche/KAPA Biosystems, # KK4824) on a Roche LightCycler 480 Instrument (Roche). Libraries with concentrations of more than 3 nM and fragments with peak size 400 bp were sequenced on an Illumina Novaseq 6000 S4 and/or S2 Flow Cell (FC). Each library was sequenced in 6-8 independent pools. For each sequencing run, 24 WGS libraries were normalized to obtain a final concentration of 2 nM using 10 mM Tris-HCl (pH 8 or 8.5; Fisher Scientific, # 50-190-8153). 0.5 to 1% Phix library was spiked into the library pool as a positive control. The normalized libraries in a pool with a total of 311 µl libraries were incubated with 77 µl of 0.2 N Sodium Hydroxyl (NaOH) (VWR, #82023-092) at room temperature for 8 minutes in order to denature double stranded DNA. 78 µl of 400 mM Tris-HCl were used to terminate the denaturing process. The denatured library with a final loading concentration of 400 pM in a pool was loaded on the S4 FC using Illumina SBS kits (Illumina, #20012866) with the following setting on the Novaseq 6000: PE150:S4 FC, dual Index, Read 1:151, Index_Read2:8; Index_Read3:8; Read 4:151. The target for whole genome sequencing with high quality sequencing raw data was 120 GB or greater with a Q30 >90% per library per sequencing run. In case the first sequencing run generated less than that, additional sequencing was performed by sequencing the same library on a Novaseq 6000 S2 FC with 2x101 read length. Raw data was processed through the DRAGEN platform to generate BAM files.

Whole-genome sequencing (WGS) data processing

Due to reference genome differences, FASTQ files were first extracted from BAM files generated by the DRAGEN platform by Picard's (v 2.20.7) *SamToFastq* command. FASTQ files were then aligned to the human_g1k_v37_decoy genome by BWA's (v 0.7.17) *mem* with *-K 100000000 -Y* parameters. SAM files were compressed to BAM files via SAMtools's (v 1.7) *view* command. BAM files were subsequently sorted by SAMBAMBA's (v 0.7.0) *sort* command and duplicated reads marked by its *markdup* command. Reads aligned to the INDEL regions were realigned with GATK's (v 3.8-1) *RealignerTargetCreator* and *IndelRealigner* following best practice. Base qualities scores were recalibrated using GATK's (v 3.8-1) *BaseRecalibrator* and *PrintReads*. Germline heterozygous variants were called by GATK's (v 3.8.1) *HaplotypeCaller*. The distribution of library DNA insertion sizes for each sample was summarized by Picard's (v 2.20.7) *CollectInsertSizeMetrics*. The depth of coverage of each sample was calculated by BEDTools's (v2.27.1) *coverage* command.

Mosaic SNV/INDEL detection in WGS data

Mosaic single nucleotide variants/mosaic small (typically below 20 bp) INDELs were called by using a combination of four different computational methods: the intersection of variants from the paired-mode of GATK's (v.4.0-4) Mutect2 and Strelka2 (v 2.9.2) (set on "pass" for all variant filter criteria) for sample-specific variants; MosaicHunter (single-mode, v 1.0) with a posterior mosaic probability >0.05 (*16*) for sample-specific or tissue-shared variants; or single-mode of Mutect2 (with an in-house panel of normal) followed by MosaicForecast (v 0.0.1) for sample-specific or tissue-shared variants (*15*). Variants were excluded if 1) residing in segmental duplication regions as annotated in the UCSC genome browser (UCSC SegDup) or RepeatMasker regions, 2) residing within a homopolymer or dinucleotide repeat with more than 3 units, or 3) overlapped with annotated germline INDELs. We further removed any variants with a population allele frequency (AF) higher than 0.001 in gnomAD (v 2.1.1) (*26*). Finally, variants with an upper confidence interval (CI) of AF > 0.45 in more than half of the tissues were considered likely germline variants and removed. Variants with a lower CI of AF < 0.001 were also removed. Fractions of mutant alleles for variants called in one sample were calculated in all the other samples together with the exact binomial confidence intervals using scripts described below for MPAS analysis. Scripts for variant filtering are provided on GitHub (https://github.com/shishenyxx/Adult_brain_somatic_mosaicism).

Fluorescence-activated nuclei sorting

Pellets of brain nuclei were washed twice in staining buffer (HBSS without magnesium and calcium, 5% BSA, 1mM EDTA) and then re-suspended in 0.2 ml staining buffer and incubated overnight at 4°C. The following antibodies were used: NeuN Alexa Fluor 488 (1:2,500; Millipore Sigma, MAB377), TBR1 unconjugated (1:1,000; Abcam, #ab31940), OLIG2 unconjugated (1:1,000; Abcam, #ab1091986), LHX2 unconjugated (1:500; Abcam, #ab2199883), PU.1 Alexa Fluor 647 (1:100; BioLegend, #658004). The following day, nuclei were washed with staining buffer and in case an unconjugated antibody was used, nuclei were stained subsequently for 30 minutes with goat anti-rabbit Alexa 647 (1:4,000; ThermoFisher Scientific, #A21244) for TBR1 or LHX2, and goat anti-rabbit Alexa 555 (1:4,000; ThermoFisher Scientific, #A32732) for OLIG2. Stained nuclei were washed one more time with staining buffer and passed through a 70 µm strainer. Immediately before the sort, nuclei were stained with 0.5 µg/ml DAPI. Nuclei for cell type of origin were sorted either on a MoFlo Astrio EQ sorter (Beckman Coulter) or on a BD InFlux Cytometer (Becton-Dickinson). Sorted nuclei were pelleted in staining buffer at 1,600 xg for 10 minutes. Nuclei for DNA extraction and H3K27ac ChIP-seq were stored at -80°C. FANS data was visualized using FlowJo software (Ashland, Oregon). Following MPAS (see below) sorted populations were deemed to be of sufficient overall quality (Fig. 3B) if at least 95% variants were sequenced above >1,000×.

Single nuclei fluorescence-activated nuclei sorting

Frozen, non-fixed brain tissue from the left temporal cortex was homogenized in 1 ml ice-cold NIB (0.25M sucrose, 25 mMKCl, 5mM $MgCl_2$, 10 mM Tris pH 7.5, 100 mM DTT, and 0.1% Triton X-100) and dounced five times in a 2 ml Wheaton Dounce Tissue Grinder (DWK Life Sciences). The homogenate was incubated on a rocker for 5 minutes at 4°C and centrifuged at 1000 xg using the 'soft' setting in a swinging bucket centrifuge. Supernatant was removed and the pellet was resuspended in 0.5 ml sorting buffer and filtered through a 70 um strainer. Pellets were stained for 30 minutes using NeuN Alexa Fluor 488 (1:2,500; Millipore Sigma, #MAB377). After washing, nuclei were stained with 0.5 µg/ml DAPI. $DAPI^+/NeuN^+$ and $DAPI^+/NeuN^-$ nuclei were sorted on a BD InFlux Cytometer (Becton-Dickinson) into a 96-well plate pre-filled with PBS. The 96-well plate with single nuclei in each well was quickly spun down and stored at -80°C until further processing for snMPAS. Single nuclei FANS data was visualized using FlowJo software.

<u>H3K27ac ChIP-seq of sorted nuclei for cell-type of origin</u>

Chromatin immunoprecipitation (ChIP) for H3K27ac was performed as previously described (*27*). Fixed, sorted nuclei (~200,000 nuclei per sample) were resuspended in 130 µl ice-cold LB3 (10 mM Tris/HCl pH 7.5, 100 mM NaCl, 1 mM EDTA, 0.5 mM EGTA, 0.1% Na-deoxycholate, 0.5% N-lauroylsarcosine, 1 X protease inhibitor cocktail). Chromatin was sheared by sonication using a Covaris E220 focused-ultrasonicator (Covaris, MA) with the following setting: time, 240 seconds; duty, 5.0; PIP, 140; cycles, 200; amplitude, 0.0; velocity, 0.0; dwell, 0.0). The lysates were adjusted to 250 µl with LB3 and further diluted with 25 µl 10% Triton X-100 (final concentration 1%). Samples were spun down at maximum speed at 4°C for 10 minutes. For DNA input control, 3 µl of the lysate were taken and volume adjusted to 25 µl with TT (10 mM Tris-HCl pH 8, 0.05% Tween-20) and stored at 4°C until library preparation. For immunoprecipitation, 20 µl of Dynabeads Protein A (ThermoFisher Scientific, #10001D) and H3K27ac antibody (2 µl serum; ActiveMotif, #39135) was added to the diluted lysates and rotated overnight at 4°C. Beads were collected on a magnet and washed three times each with wash buffer I (20 mM Tris/HCl pH 7.5, 150 mM NaCl, 1% Triton X-100, 0.1% SDS, 2 mM EDTA), wash buffer III (10 mM Tris/HCl pH 7.4, 250 mM LiCl, 1% Triton X-100, 0.7% Na-Deoxycholate, 1 mM EDTA), twice with ice-cold TET (10 mM Tris/HCl pH7.5, 1 mM EDTA, 0.2% Tween-20), once with TE-NaCl (10 mM Tris-HCl pH8, 1 mM EDTA, 50 mM NaCl) and finally resuspended in 25 µl TT. Libraries from ChIP and DNA input samples were prepared with the NEBNext Ultra II DNA library prep kit (NEB) reagents according to the manufacturer's protocol on the beads suspended in 25 µL TT (10 mM Tris/HCl pH7.5, 0.05% Tween-20), with reagent volumes reduced by half. Next, DNA was eluted and crosslinks reversed by adding 4 µl 10% SDS, 4.5 µl 5 M NaCl, 3 µl EDTA, 1 µl proteinase K (20 mg/ml), 20 µl water, incubating for 1 hour at 55°C , then 30 minutes to overnight at 65°C. DNA was purified using 2 µL of SpeedBeads (GE Healthcare), diluted with 20% PEG8000, 1.5 M NaCl to a final of 12% PEG, eluted with 12.5 µl TT. DNA contained in the eluate was then amplified for 14 cycles in 25 µl PCR reactions using NEBNext High-Fidelity 2X PCR Master Mix (NEB) and 0.5 mM each of primers Solexa 1GA and Solexa 1GB. Resulting libraries were size selected by gel excision to 225-350 bp, purified, and single-end sequenced using a HiSeq 4000 or a NextSeq 500 (Illumina).

<u>H3K27ac ChIP-seq data processing and data visualization</u>

FASTQ-files were obtained from the Illumina Studio pipeline and mapped and aligned to hg19 with Bowtie2 (v2.2.9). Quantification of H3K27ac ChIP-seq was performed using HOMER (v4.9.1) (*28*). First, HOMER tag directories were generated using HOMER's 'findPeaks' command with the following parameters: "style histone -size 1000 -minDist 2500 -region". Next, H3K27ac signals at peaks were merged for all cell populations followed by annotation using HOMER's 'annotatePeaks' function with the following parameter: '-norm 1e7'. Heat maps were generated using the seaborn package in Python. H3K27ac ChIP-seq obtained NeuN, TBR1, OLIG2, NeuN/LHX2, and PU.1 populations were compared to H3K27ac CHIP-seq data derived from cell populations from pediatric brain tissue (*18*). PCA was generated using the Python (v3.7.1) packages scipy (v1.5.1) and sklearn (v0.20.1). Browser images were generated from the UCSC genome browser and can be found at the following address: https://genome.ucsc.edu/s/jschlachetzki/Schlachetzki_7614_celltypes_H3K27ac.

Whole-genome amplification (WGA) of DNA from sorted single nuclei

Following single nuclei fluorescence-activated sorting, single nuclei WGA was performed using the REPLI-g Single Cell Kit according to the manufacturer's protocol (Qiagen, #150345).

Massive parallel amplicon sequencing (MPAS) and single nuclei MPAS (snMPAS) design and procedure

A customized AmpliSeq Custom DNA Panel for Illumina (#20020495, Illumina, San Diego, CA, USA) was used for MPAS and snMPAS. Designed genomic regions are provided in Data S1. A list of 1455 candidate mosaic variants from the mosaic variant detection pipeline described above were subjected to the AmpliSeq design system. We randomly selected 120 high-confidence heterozygous variants as positive controls. These heterozygous variants presented with estimated AF between 48-52% for all the 25 sequenced bulk tissues, and with read depths between 270-330×. Of the 120 variants, 45 were private variants and 75 were present in gnomAD at different population allele frequencies. We also randomly selected 40 reference homozygous variants as negative controls. These reference homozygous variants presented with ~0% AF across all sequenced samples, with average depth 270-330×, and gnomAD (v 2.1.1) AF >0.5 to exclude any potential contamination or amplification bias. The AmpliSeq design software determined ~1400 pairs of primers suitable for multiplex PCR reaction in a single pool after optimization. DNA from extracted tissue, nuclei, amplified single nuclei, and a duplicate unrelated control sample was diluted to 5 ng/µl in low TE provided in AmpliSeq Library PLUS (384 Reactions) kit (Illumina, #20019103). AmpliSeq was carried out following the manufacturer's protocol (document #1000000036408v07). For amplification, 14 cycles each with 8 minutes were used. After amplification and FUPA treatment, libraries were barcoded with AmpliSeq CD Indexes (Illumina, #20031676) and pooled with similar molecular numbers based on measurements made with a Qubit dsDNA High Sensitivity kit (Thermo Fisher Scientific, #Q32854) and a plate reader (Eppendorf, PlateReader AF2200). To avoid index hopping, the two library pools were sequenced on separate lanes on different NovaSeq 6000 runs. 87.4 GB of FASTQ data were obtained from the MPAS libraries, aiming for an average of 5000× coverage for each variant; and 27.5 GB of FASTQ data were obtained from the snMPAS libraries, aiming for an average of 1500× for each variant.

Data analysis for MPAS and snMPAS

Raw reads from MPAS and snMPAS were mapped to the human_g1k_v37_decoy genome with BWA's (v0.7.17) *mem* command. BAM files were processed without marking PCR duplicates. Reads near insertion/deletions were re-aligned with GATK's (v3.8-1) *IndelRealigner* and base qualities scores were recalibrated with GATK's (v3.8-1) *BaseRecalibrator*. The final BAM files were parsed by SAMtools's (v1.7) *mpileup* and the 95% confidence intervals (CIs) of the real allelic fractions of all the candidate mosaic variants, together with the reference homozygous (negative control) and heterozygous (positive control) variants were estimated based on an exact binomial estimation (https://github.com/shishenyxx/Adult_brain_somatic_mosaicism). Following depth calculation, regions of 1349 mosaic candidates, 113 heterozygous variants (positive controls) and 27 reference homozygous variants (negative controls) were detected and subjected to the next genotyping steps. The genotypes of candidate mosaic variants from MPAS were determined by comparing them to the AF distribution of the reference homozygous and heterozygous variants. The exact binomial lower bounds of all reference homozygous variants with >30 read depth were estimated (Fig. S2B, left) and the 95% single-tail confidence threshold for the lower bound was calculated to be 1.397e-3. Distribution of the exact binomial upper bond of all heterozygous variants were calculated (Fig. S2B, right) and 0.4 was considered to be the threshold for the upper bond based on ~5% FDR and manual inspection. Mosaic candidates from WGS were considered positive if 1) the 95% exact binomial lower bound was >1.397e-3 and above the upper CI of the unrelated control sample, 2) the sequencing depth was >30, and 3) the assessed alternative allele was supported by ≥3 reads. These criteria ensured the FDR for each variant was under 5%. If mosaic candidates were detected with an upper CI >0.4 in more than half (≥13) of the original 25 samples that underwent WGS they were considered as likely heterozygous variants and removed from the mosaic variant list. Due to the high allelic dropout rate and imbalanced amplification of different alleles, we carried out a stricter genotyping strategy for snMPAS. For determination of positively detected variants in single nuclei, the cut-off for read depth was above 30, and the lower CI of the calculated allelic fraction >0.05 and above the upper CI of the negative control sample; this resulted in a ~0.01 false discover rate based on the analysis of reference homozygous controls. To determine the lateralization of a variant, first, the number of the original 25 samples, in which the variant was detected in the left and right hemisphere were calculated, then, the lateralization was determined as:

1. 'Not lateralized' if not present in any of the lateralized tissues;
2. 'Left only' if variant only presented in the left tissues;
3. 'Right only' if variant only presented in the right tissues;
4. 'Left enriched' if $\frac{\sum Number_{left}}{\sum Number_{right}} \geq 1.5$ or also present in non-lateralized tissues;
5. 'Right enriched' if $\frac{\sum Number_{right}}{\sum Number_{left}} \geq 1.5$ or also present in non-lateralized tissues; and
6. 'Both sides' if $\frac{\sum Number_{left}}{\sum Number_{right}} < 1.5\ and\ \frac{\sum Number_{right}}{\sum Number_{left}} < 1.5$

Due to the rate of genotyping errors and the variability among heterozygous variants, the following criteria had to be fulfilled for a variant to be considered for the lineage reconstruction in single cells: a variant had to be detectable in any of the original samples (large or small) from the left temporal cortex, i.e. the same brain region from which single nuclei were isolated; a variant had to be present in ≤20 cells, to avoid genotyping errors; and a variant had to be present in more than 1 cell to be informative. Likewise, only cells that harbored more than one variant were included. Following a double-ranked plot for variants and cells, clades were determined manually. 'Non-informative' variants were labeled as such, if they distributed among other major clades, even in case their overall AFs were inconsistent with their abundance in snMPAS. These variants were excluded from subsequent lineage tree analyses. Details and codes for the data processing and annotation are provided on GitHub (https://github.com/shishenyxx/Adult_brain_somatic_mosaicism).

Analysis of mosaic variant overlap with different genomic features

Annotations were sourced as follows. Whole-genome histone modifications data for *H3k27ac, H3k27me3, H3k4me1, and H3k4me3* were downloaded from the UCSC genome browser (http://hgdownload.soe.ucsc.edu/goldenPath/hg19/database/). To compare the somatic variants density detected in this study with the Encode v3 features, we calculated the overlap of the variants with peaks called from the H1 human embryonic cell line (H1), and with peaks merged from 9 different cell lines (Mrg; Gm12878, H1hesc, Hmec, Hsmm, Huvec, K562, Nha, Nhek, and Nhlf). Gene region, intronic, and exonic regions were downloaded from NCBI RefSeqGene (http://hgdownload.soe.ucsc.edu/goldenPath/hg19/database/refGene.txt.gz); Topoisomerase 2A/2B (*Top2a/b*) sensitive regions from ChIP-seq data (Samples: GSM2635602, GSM2635603, GSM2635606, and GSM2635607) (*29*); *CpG islands*: data from the UCSC genome browser (http://hgdownload.soe.ucsc.edu/goldenPath/hg19/database/); *genomic regions with annotated early and late replication timing*: areas as described (*30*); *enhancer* genomic regions from the VISTA Enhancer Browser (https://enhancer.lbl.gov/); *DNase I hypersensitive regions* and *transcription factor binding sites* from Encode v3 tracks from the UCSC genome browser (wgEncodeRegDnaseClusteredV3 and wgEncodeRegTfbsClusteredV3, respectively). For analysis, all single nucleotide mutations from gnomAD (v 2.1.1; https://gnomad.broadinstitute.org/) were first intersected with the callable regions described above to make sure that all selected variants have the same distribution on genome as the mosaic candidates. Genomic features described above were annotated to those gnomAD (v 2.1.1) variants as well as mosaic variants detected in this study by using BEDTools (v2.27.1) *annotate*. 10,000 permutations were carried out for those variants by selecting the number of variants equal to each variant categories randomly by using the bash command *shuf*. The fraction of variants within each annotated region was then calculated for the 10,000 independent samples, and the 95% confidence intervals of the mutation densities were calculated across all permutations. Together this set up the null distribution of variant overlap with the annotated genomic features.

Analysis of asymmetric variant distributions and estimation of the starting cell population during left-right split

To determine the brain-specific mean in each hemisphere on the left (L) and right (R), we considered variants in all sorted samples that fulfilled the following criteria: coverage >1,000×; and originated developmentally from the brain (i.e. not PU.1$^{+}$). Across these samples the overall average AF was determined for each variant. Likewise, for the anterior-posterior assessment the sorted populations from PF and F (anterior; A) and P, O, T (posterior, P) cortical areas from both hemispheres were used. For both analyses, a normalized $\Delta_{LR}$ was determined by subtracting the mean left from the mean right.

$$\text{For any variant, } \Delta_{LR} = \frac{\sum_i^n AF_{Sorted\_population\_left}}{n_{Sample\ from\ the\ left}} - \frac{\sum_i^n AF_{Sorted\_population\_right}}{n_{Sample\ from\ the\ righr}},$$

and the normalized $\Delta_{AP}$ was the mean posterior from the mean anterior,

$$\text{For any variant, } \Delta_{AP} = \frac{\sum_i^n AF_{Sorted\_populations\_PF\_and\_F}}{n_{Sample\_from\_PF\_and\_F}} - \frac{\sum_i^n AF_{Sorted\_population\_P\_O\_T}}{n_{Sample\_from\_P\_O\_T}},$$

and then normalizing this difference with the larger of the two values.

Estimation of the maximum number of starting population before lateralization based on variants shared in both hemispheres

According to the law of large numbers, the CI of sampling will become closer to the expected value with increasing sample number. Assuming that the starting population N is split in two hemisphere populations N/2, and that the AF observed in the adult tissue is directly related to the AF at the time of the split, we can estimate the maximally supported N based on the observed difference in AF between the two hemispheres. Based on this idea, the upper limit of the starting cell population during the left-right split is determined by using the observed AF distribution and inferred cell fraction in the hemisphere as extreme values of a hypergeometric distribution using the function *hyperCI* from the *FSA* (v0.8.30) package in R (v3.5.1) for each individual variant. We chose the 95% CI as a threshold to determine the maximally supported N, and we considered all variants that were present across hemispheres, present in at least one non-cortical tissue, or both. This assumed that such shared variants arose prior to the split and were subject to the left-right split.

Estimation of the minimum number of progenitors in the starting population after lateralization based on hemisphere-specific variants

Theoretically, variants that were detected in one hemisphere only in all the sorted, brain-specific populations must have occurred after the left right split, or at least after the lateralization was determined. In the most extreme case, the highest AF measured for those variants is due to one cell carrying the mutation immediately after the split (e.g. if one cell out of a population of 10 cells has a heterozygous mutation, we would observe an AF of 5% across the hemisphere). We calculated the most extreme number of the hemisphere-specific variants as the lower bound to estimate the starting cell population during the left-right split. Note that this assumes that the used variants are present in only one cell, as they arise at or after the left-right split.

Lineage determination for genotyped single nuclei using BEAST

We reconstructed lineages for genotyped single nuclei using BEAST (Bayesian Evolutionary Analysis by Sampling Trees V1.10.4). Input for BEAST was a constructed multiple alignment of 33 base pairs, representing the 33 mutations genotyped in 71 sampled nuclei. We recorded an alternate nucleotide if the MAF was above 0 and if the variant was not flagged as noise, while recording the reference nucleotide if the variant for a given sample failed to meet these conditions. We implemented the Jukes Cantor (JC69) base substitution model since it assumes equal base substitution rates. An exponential model was chosen for determining the coalescent given that cells divide likely exponentially during early (*31*). We then assumed a strict molecular clock and propagated the Markov chain for one million iterations. Lineage and clade membership was visualized using the maximum clade credibility tree after 100,000 burin states.

Computational deconvolution of the contribution of cell lineages validated from snMPAS

The cellular lineage and topological structures were determined using the high-confidence snMPAS data of variants as described before. The information of allelic fractions quantified from MPAS from bulk tissues were likewise considered for these variants and a combined matrix was estimated through ‘principles’ modified from LICHeE (*32*). For any collection of AFs estimated from bulk tissues and sorted populations (denoted *i*) and the collection of genotypes

from snMPAS (denoted $j$) for each variant, the following principles must be fulfilled to determine the parent-child relationship between any pair of variants (denoted $u$ and $v$):
1. $Parent\ \overline{AF_i} \geq Child\ \overline{AF_i}$;
2. If $Child_{genotype_j} = 1$, then has to fulfill $Parent_{genotype_j} = 1$;
3. A direct edge between $Variant_u$ and $Variant_v$ does not exist if Pearson's correlation ($AF_{i,Variant_u}$, $AF_{j,Variant_v}$) coefficient $< 0$ and P-value $< 0.05$.

We started to construct the tree by connecting variants in the same levels and adjacent levels (levels were defined according to the Hamming weight or the number of ones in the single cell genotyping profile). Variants without direct parent after the first round of reconstruction were further investigated whether they could be connected to variants in other levels. A potential root was assumed to be the genotype of the zygote without any of the detected somatic variants, and it was set as the parent node of all detected clade founders. Mutations were then encoded for each resulting lineages at each level in "0, 1" sequences:

$L_i$ = (0, 1, 0, ..., 0, 1) (1 if assessed variant was present)

Thus, presence and absence of each variant in the tree structure was defined as a vector $L_i$ for each node in each lineage. The tree structure was then represented as matrix **L**:

$$\mathbf{L} = \begin{pmatrix} L_1 \\ L_2 \\ L_3 \\ \dots \\ L_i \end{pmatrix}$$

The relative contribution (weight) of each lineage was defined as vector $\mathbf{w}^{\boldsymbol{T}}$. $\mathbf{w}^{\boldsymbol{T}}$ was estimated through a LASSO regression model and the mean squared error method was used to minimize the sum of residual squares between $\mathbf{L}\mathbf{w}^{\boldsymbol{T}}$ and the AF matrix **E** detected by MPAS:

$$||\mathbf{L}\mathbf{w}^{\boldsymbol{T}} - \boldsymbol{E}||$$

Statistical tests and packages for customized plots

One-way ANOVA was performed using Python (3.6.8) with the pingouin (0.3.5) package on pandas (0.24.2) dataframes. Exact binomial CI of AFs were calculated in R (version 3.5.1) with *binom.test()*. Spearman correlation coefficients were estimated using Python with the scipy (v.1.3.1) package. Unless otherwise noted, data analysis and processing was done using Python with pandas and numpy (v1.16.2); customized plots were generated by Python using seaborn (0.9.0) and matplotlib (3.1.1). Details and codes for the data processing and annotation are provided on GitHub (https://github.com/shishenyxx/Adult_brain_somatic_mosaicism).

### Supplementary Text

<u>Members of *The Brain Somatic Mosaicism Network*</u>

Boston Children's Hospital: August Yue Huang, Alissa D'Gama, Caroline Dias, Christopher A. Walsh, Javier Ganz, Michael Lodato, Michael Miller, Pengpeng Li, Rachel Rodin, Robert Hill, Sara Bizzotto, Sattar Khoshkhoo, Zinan Zhou

Harvard University: Alice Lee, Alison Barton, Alon Galor, Chong Chu, Craig Bohrson, Doga Gulhan, Eduardo Maury, Elaine Lim, Euncheon Lim, Giorgio Melloni, Isidro Cortes, Jake Lee, Joe Luquette, Lixing Yang, Maxwell Sherman, Michael Coulter, Minseok Kwon, Peter J. Park, Rebeca Borges-Monroy, Semin Lee, Sonia Kim, Soo Lee, Vinary Viswanadham, Yanmei Dou

Icahn School of Medicine at Mt. Sinai: Andrew J. Chess, Attila Jones, Chaggai Rosenbluh, Schahram Akbarian

Kennedy Krieger Institute: Ben Langmead, Jeremy Thorpe, Jonathan Pevsner, Sean Cho

Lieber Institute for Brain Development: Andrew Jaffe, Apua Paquola, Daniel Weinberger, Jennifer Erwin, Jooheon Shin, Michael McConnell, Richard Straub, Rujuta Narurkar

Mayo Clinic: Alexej Abyzov, Taejeong Bae, Yeongjun Jang, Yifan Wang

NIMH: Anjene Addington, Geetha Senthil

Sage Bionetworks: Cindy Molitor, Mette Peters

Salk Institute for Biological Studies: Fred H. Gage, Meiyan Wang, Patrick Reed, Sara Linker

Stanford University: Alexander Urban, Bo Zhou, Xiaowei Zhu

Universitat Pompeu Fabra: Aitor Serres Amero, David Juan, Inna Povolotskaya, Irene Lobon, Manuel Solis Moruno, Raquel Garcia Perez, Tomas Marques-Bonet

University of Barcelona: Eduardo Soriano

University of California, Los Angeles: Gary Mathern

University of California, San Diego: Danny Antaki, Dan Averbuj, Eric Courchesne, Joseph Gleeson, Laurel Ball, Martin Breuss, Subhojit Roy, Xiaoxu Yang

University of Michigan: Diane Flasch, Frisbie Trenton, Huira Kopera, Jeffrey Kidd, John Moldovan, John V. Moran, Kenneth Kwan, Ryan Mills, Sarah Emery, Weichen Zhou, Xuefang Zhao

University of Virginia: Aakrosh Ratan

Yale University: Alexandre Jourdon, Flora M. Vaccarino, Liana Fasching, Nenad Sestan, Sirisha Pochareddy, Soraya Scuderi

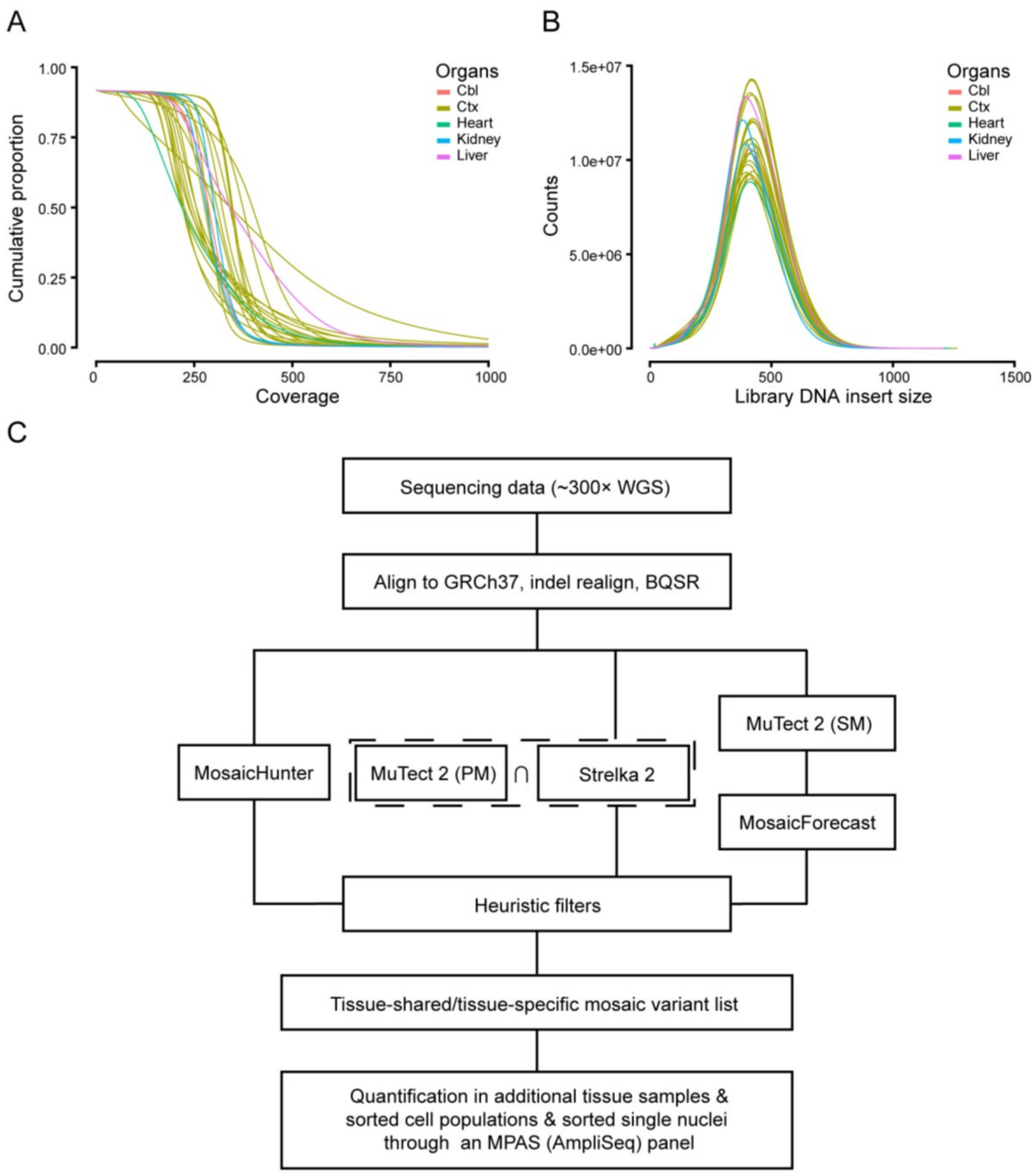


**Fig. S1. Quality assessment of the WGS data and the analytical workflow.** (**A**) Cumulative proportion of coverage depth for each whole genome sequencing (WGS) sample. Colors were used to distinguish different organs. Cbl: cerebellum, Ctx: neocortex. The majority of the sequenced samples reached at least 300×. (**B**) Library DNA insertion size for each sample. All the samples had a consistent single peak at ~400 bp. Colors are the same as in A. (**C**) Workflow for sequencing data processing, mosaic variant calling, and variant quantification. Details are described in Materials and Methods. INDEL realign: realignment of reads near detected insertion/deletions, BQSR: base quality score recalibration, PM: paired mode, SM: single mode.

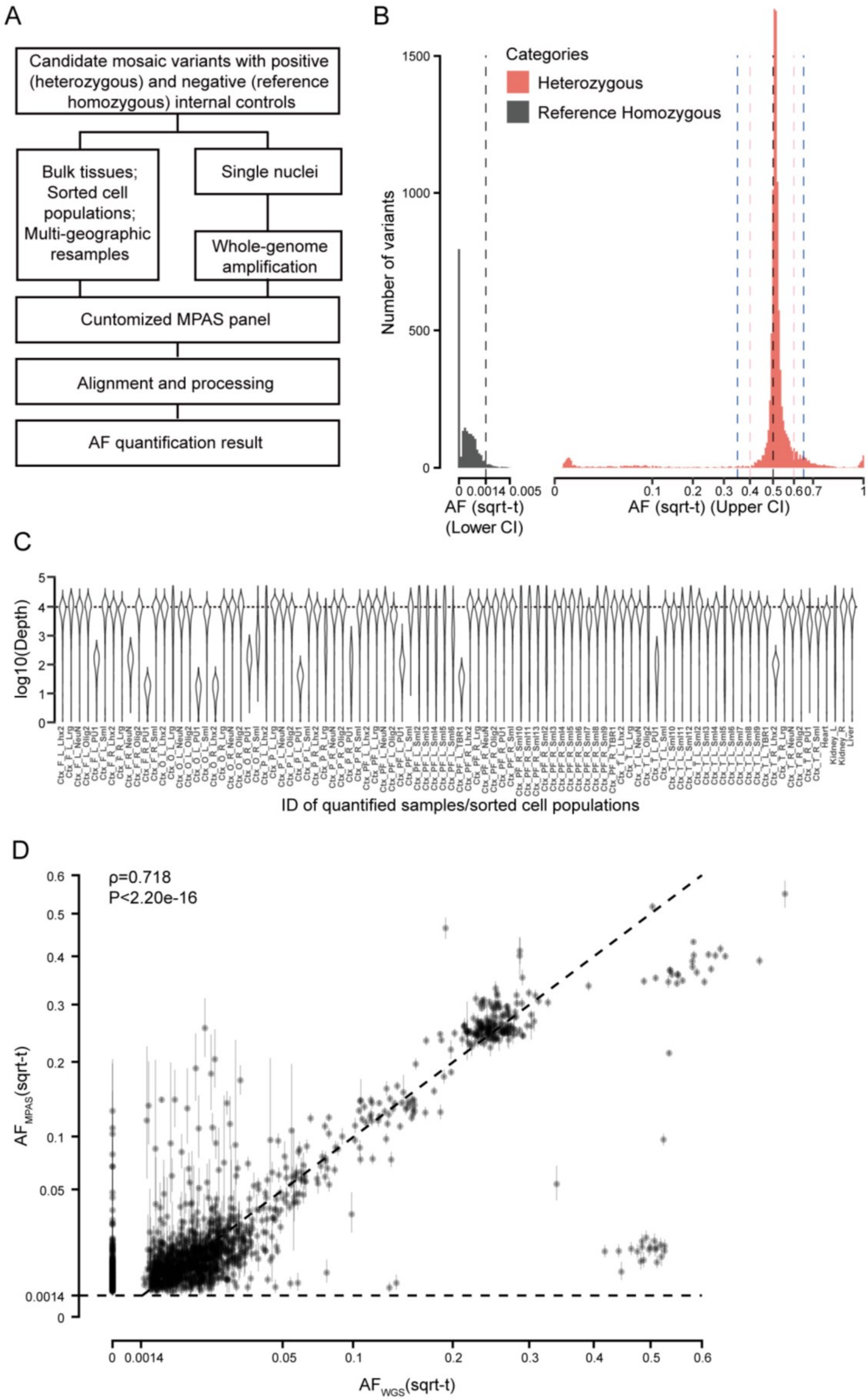
A
Candidate mosaic variants with positive (heterozygous) and negative (reference homozygous) internal controls
Bulk tissues; Sorted cell populations; Multi-geographic resamples
Single nuclei
Whole-genome amplification
Cuntomized MPAS panel
Alignment and processing
AF quantification result
B
Categories
Heterozygous
Reference Homozygous
Number of variants
1500
1000
500
0
0 0.0014 0.005
AF (sqrt-t) (Lower CI)
0 0.1 0.2 0.3 0.4 0.5 0.60.7 1
AF (sqrt-t) (Upper CI)
C
log10(Depth)
5
4
3
2
1
0
ID of quantified samples/sorted cell populations
D
ρ=0.718
P<2.20e-16
AF_MPAS(sqrt-t)
0.6
0.5
0.4
0.3
0.2
0.1
0.05
0.0014
0
0 0.0014 0.05 0.1 0.2 0.3 0.4 0.5 0.6
AF_WGS(sqrt-t)

**Fig. S2. MPAS workflow and quality assessment of the data.** (**A**) Workflow for massive parallel amplicon sequencing (MPAS) and single nuclei MPAS (snMPAS). Reference homozygous variants (negative control) and heterozygous variants (positive control) from dbSNP were added to the MPAS panel. AF: Allelic fraction. (**B**) AF distribution of the exact binomial lower bound of MPAS results for 27 reference homozygous variants (grey) and the exact binomial upper bound of 113 heterozygous variants (red) are shown. x-axis is square root transformed. 0.00139 was identified as the cutoff for the exact binomial lower bound and 0.350 as the upper bond for MPAS detection with a false discovery rate of 5%. (**C**) Depth distribution of all the 1349 candidate genomic positions designed for MPAS for each sequenced sample form from the donor. The depth is $\log_{10}$ transformed. PF_L and PF_R, prefrontal cortex from left and right hemisphere; F_L and F_R, frontal cortex from left and right; O_R and O_R, occipital cortex from left and right; P_L and P_R, parietal cortex from left and right; T_L and T_R, temporal cortex from left and right. (**D**) Correlation of AF estimated in all the samples by AmpliSeq based MPAS and by Whole Genome sequencing. Spearman's correlation ($\rho=0.718$) showed a significant correlation with $P<2.20e-16$. Error bars showed 95% exact binomial confidence intervals measured in snMPAS.

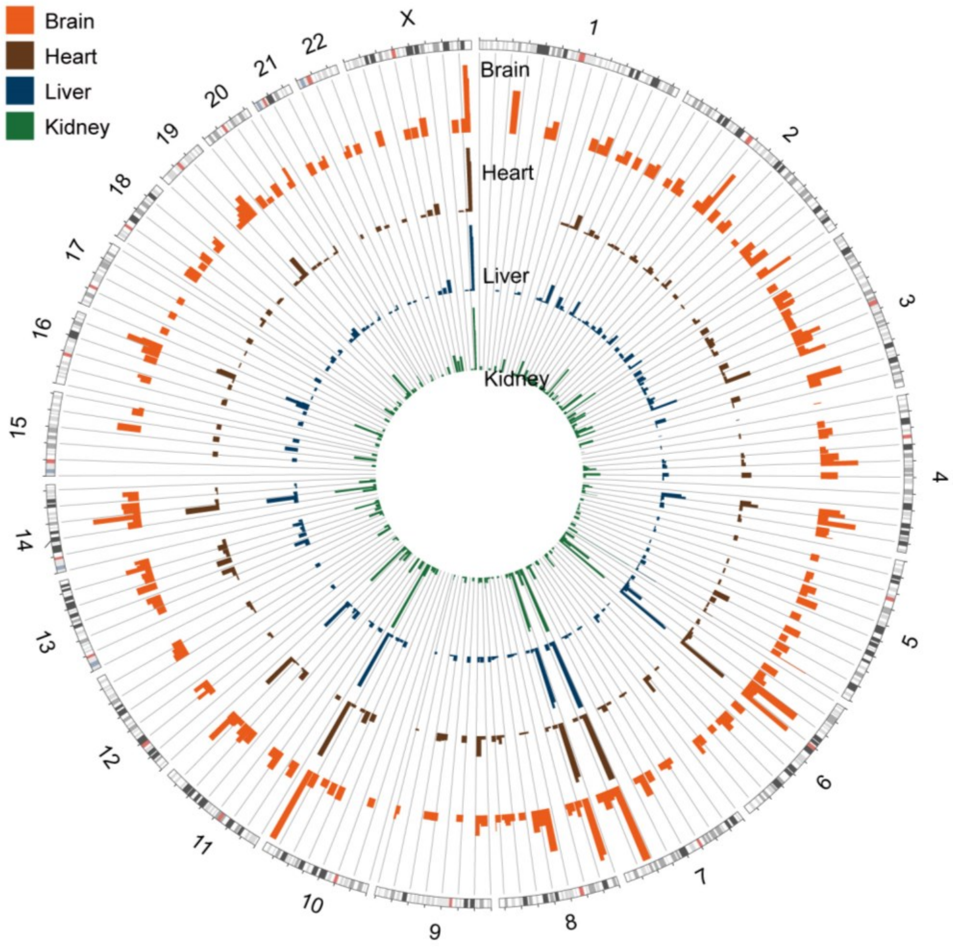


**Fig. S3. Mosaic variants were distributed across the genome.** Circos plot of the genomic positions (hg19) of all detected and quantified positive variants. Different colors were used to distinguish AF from different organs, the highest AF from all sequenced bulk brain regions is shown for each variant in the brain track. The higher AF of both kidneys is plotted for the kidney track if present in left and right. Height of each category reflects the square root transformed AF from 0.0 to 0.5. Chromosomes are indicated by number or with ‘X’.

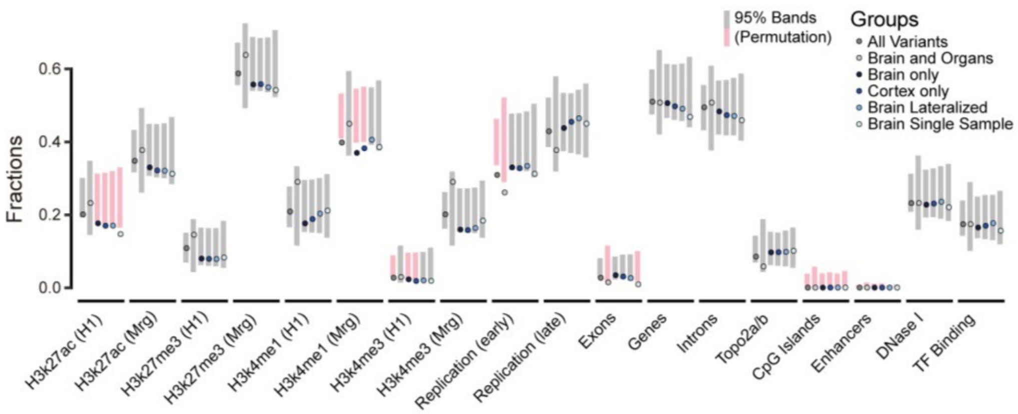


**Fig. S4. Putative early embryonic variants were enriched in distinct genomic regions.** Fraction of variants located in different genomic regions for the six categories based on tissue distribution. H3k27ac/H3k27me3/H3K4me1 (H1): H3k27ac/H3k27me3/H3K4me1 acetylation peak regions measured in human H1esc from Encode v3; H3k27ac/H3k27me3/H3K4me1(Mrg): H3k27ac/H3k27me3/H3K4me1 peak regions merged from 9 different cell lines from Encode v3 (see Materials and Methods); Exons/Gene/Introns: annotated from UCSC human RefSeqGene; Top2a/b: topoisomerase hypersensitive regions from ChIP-seq data; Early and Late replication: measured DNA replication timing; DNase I: DNase I hypersensitive regions from Encode v3; TF Binding: Transcription factor binding sites from Encode v3. 95% permutation intervals were calculated from 10,000 random permutations of the same number of variants as for each mutation category from gnomAD (v2.1.1). If the detected variant category was outside of the permutation band, the band was labeled pink.

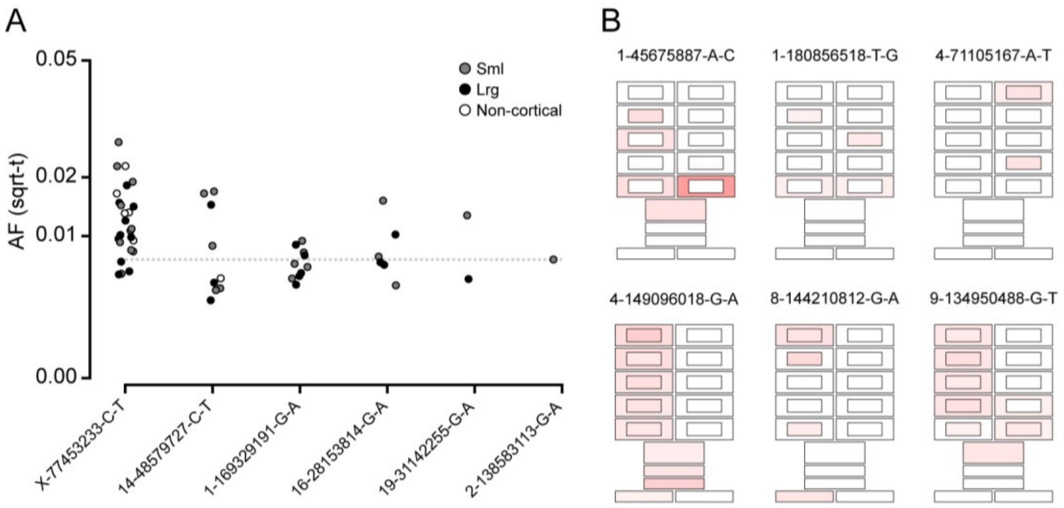


**Fig. S5. One sample biopsies have limited predictive value to determine the stochastic distribution of variants.** (**A**) AF from all tissues where a given mosaic variant shown in Fig. 2, B to G, was detected. Horizontal line indicates the observed AF for the variant only detected in one biopsy (Fig. 2G). Note that all variants have observed AF(s) in small biopsies at or below the level of the single-tissue variant, suggesting limited predictive value from sequencing of a single small biopsy. (**B**) Additional examples of geoclones with patterns that are suggestive of stochastic distribution of variants across and within hemispheres. Variants were only detectable in a subset of discontinuous tissues.

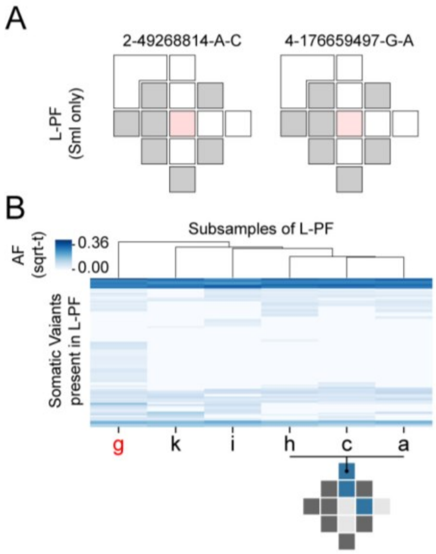


**Fig. S6. Allelic fraction of variants across subsamples in the left prefrontal cortex.** (**A**) Examples of two variants that were originally only called in the Sml central biopsy in L-PF and showed no spread across the cortical lobe within the sampled areas. (**B**) Hierarchical clustering of variants and tissues based on the AF found in the subsamples in L-PF. Shown is a schematic marking the location of the areas within the region that were clustered together (blue: present; light grey: not detected; dark grey: No Data).

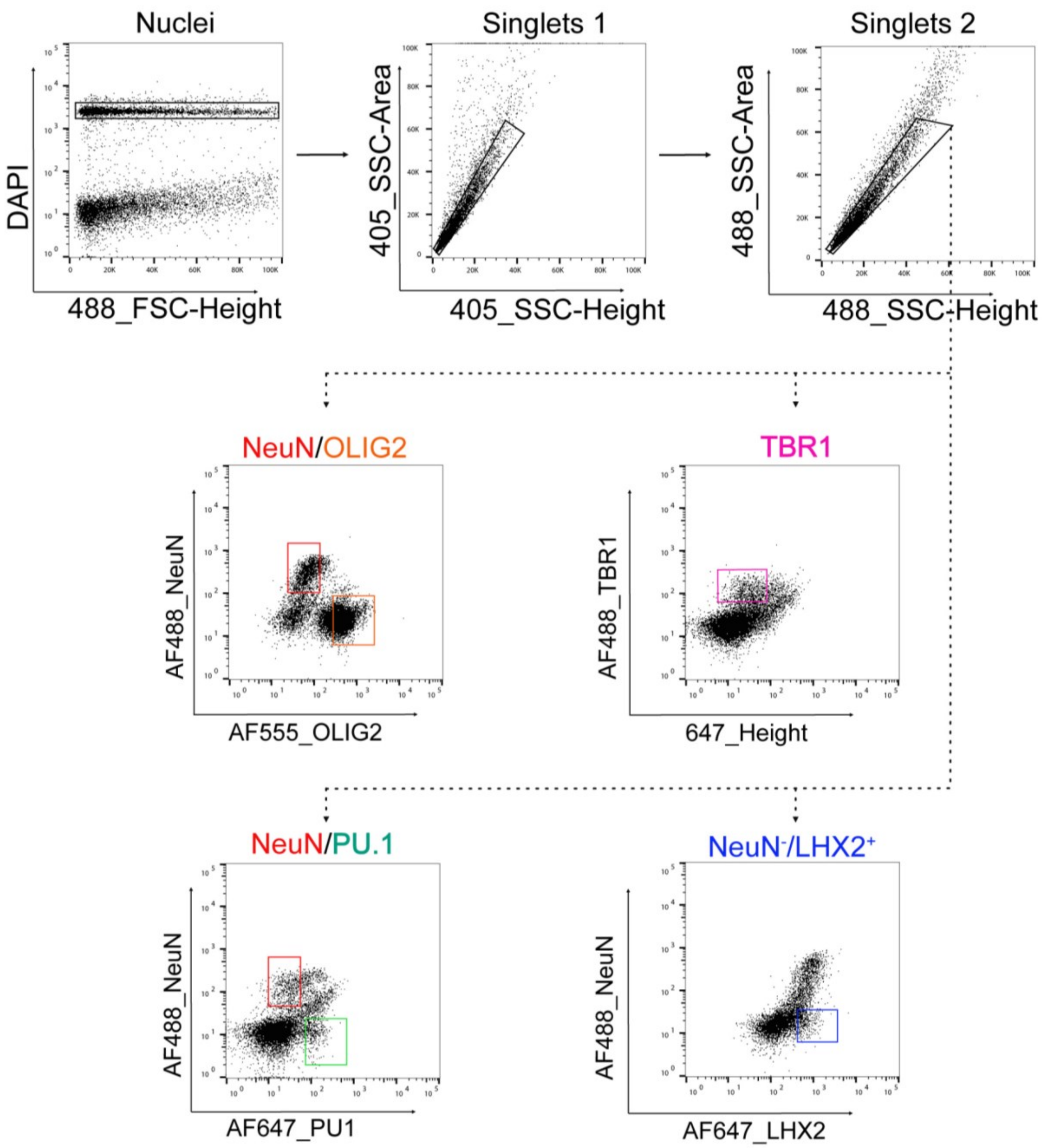


**Fig. S7. FANS for cell-type of origin from postmortem brain tissue.** Selection of DAPI positive nuclei and gating on singlets. Neurons, OPCs/oligodendrocytes, and microglia were sorted by staining against NeuN, OLIG2, and PU.1, respectively. Excitatory neurons were sorted by using the excitatory neuronal marker TBR1. Astrocytes were sorted by gating on $NeuN^-$/$LHX2^+$ nuclei.

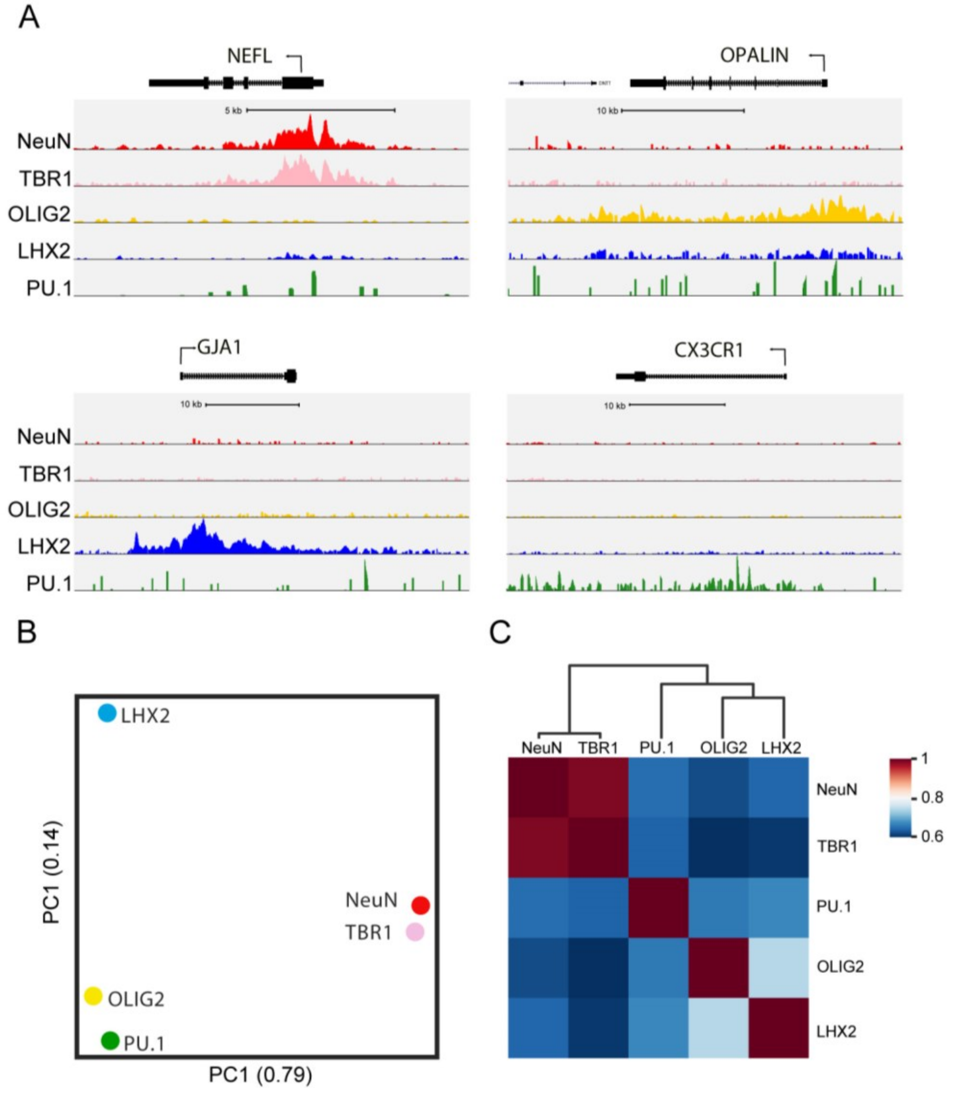


**Fig. S8. H3K27ac ChIP-seq of nuclei populations from the *postmortem* brain shows that they are distinct and specific.** (**A**) UCSC genome browser tracks of H3K27ac for brain cell-type nuclei populations. Representatives genes for neurons including excitatory neurons (*NEFL* encoding Neurofilament Light), OPCs/Oligodendrocytes (*OPALIN* encoding for Oligodendrocytic Myelin Paranodal And Inner Loop Protein), astrocytes (*GJA1* for Gap Junction Protein Alpha 1), and microglia (*CX3CR1* for Fractalkine Receptor). (**B**) PCA of H3K27ac in nuclei from NeuN$^+$, TBR1$^+$, OLIG2$^+$, NeuN$^-$/LHX2$^+$, and PU.1$^+$ brain populations. (**C**) Heatmap of Pearson's correlation of H3K27ac ChIP-seq $\log_2$(Normalized tags+1) in NeuN$^+$, TBR1$^+$, OLIG2$^+$, NeuN$^-$/LHX2$^+$, and PU.1$^+$ brain populations.

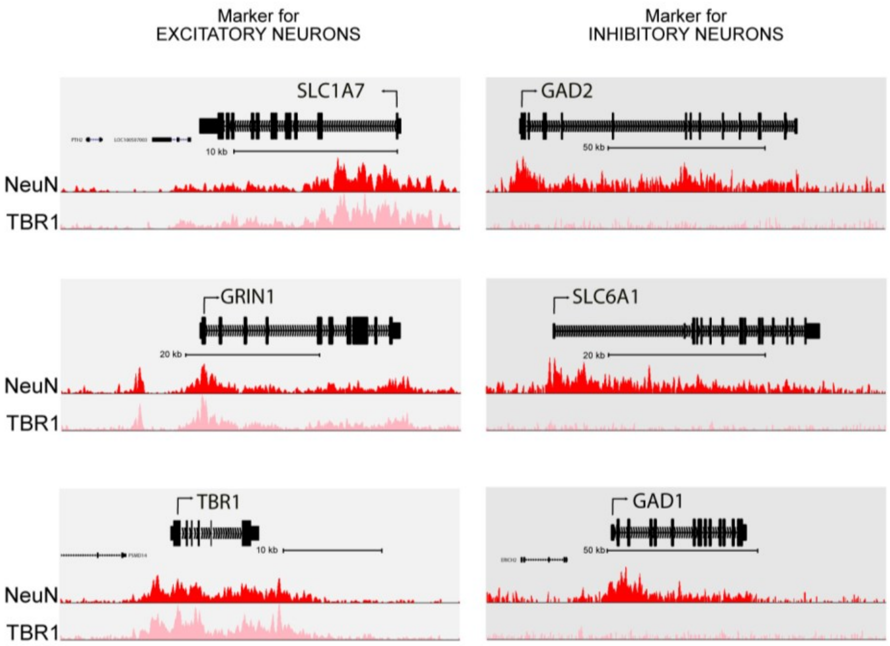


**Fig. S9. H3K27ac ChIP-seq from NeuN$^+$ and TBR1$^+$ neurons shows the enrichment of all and excitatory neurons.** TBR1 sorted nuclei show acetylation of H3K27 at promoters specific for excitatory neurons but for inhibitory neurons. UCSC genome browser track for H3K27ac in NeuN$^+$ and TBR1$^+$ populations at loci for excitatory and inhibitory neuronal markers (*SLC1A7* – Excitatory amino acid transporter 5, *GRIN1* – N-Methyl-D-Aspartate Receptor Channel, Subunit Zeta-1, *TBR1* – T-box Brain Transcription Factor 1; *GAD2* – Glutamate Decarboxlase 2, *SLC6A1* – GABA-Transporter 1, *GAD1* - Glutamate Decarboxlase 1).

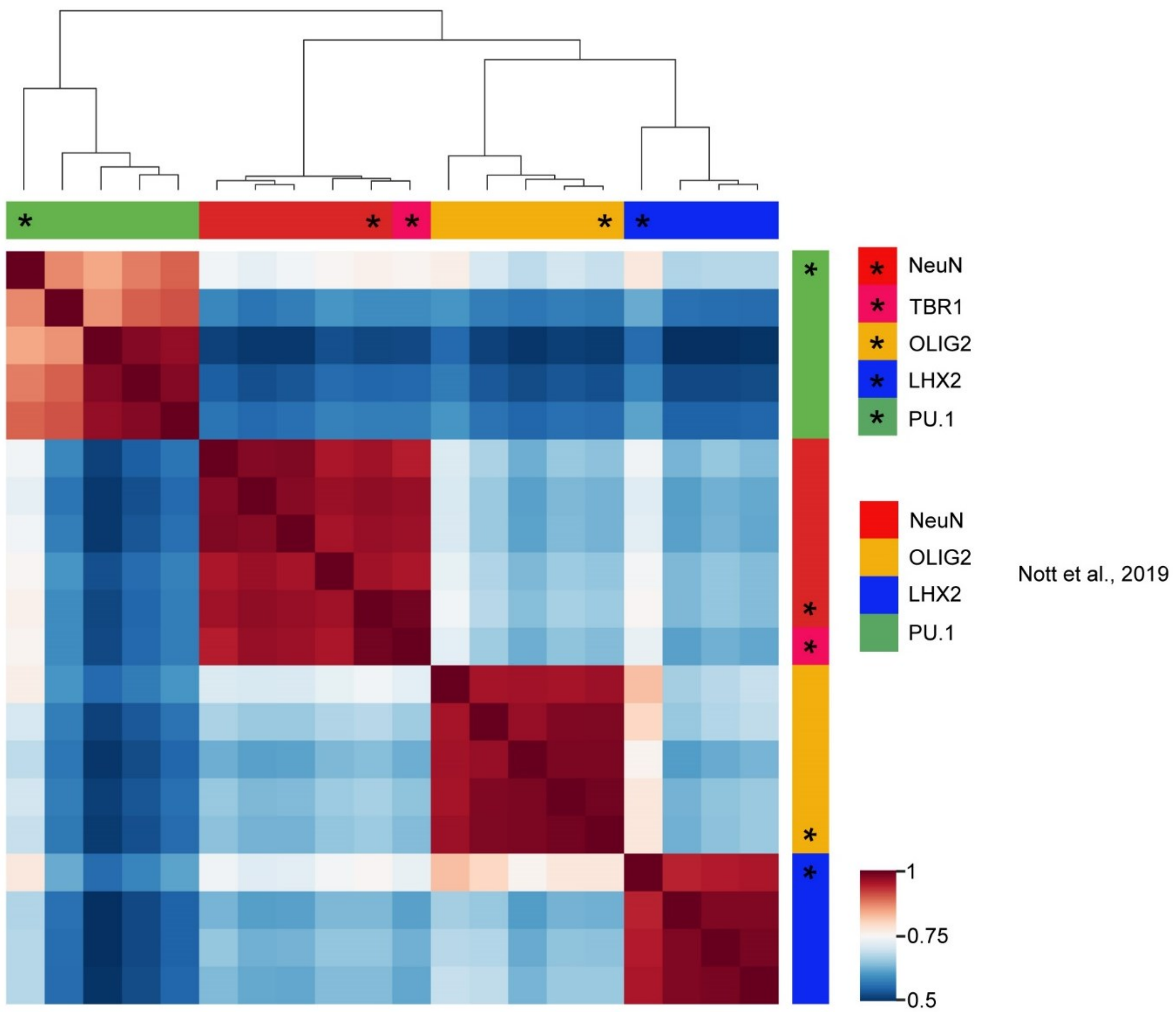


**Fig. S10. Comparison of H3K27ac ChIP-seq of brain nuclei populations from the *postmortem*, adult brain with nuclei populations from surgically resected, pediatric brain.** Heatmap of Pearson's correlation of all H3K27ac ChIP-seq $log_2$(Normalized tags+1) values from cell types in the *postmortem* tissue (marked with an asterisk) compared to H3K27ac ChIP-seq data sets from surgically resected brain tissue of pediatric patients (*18*).

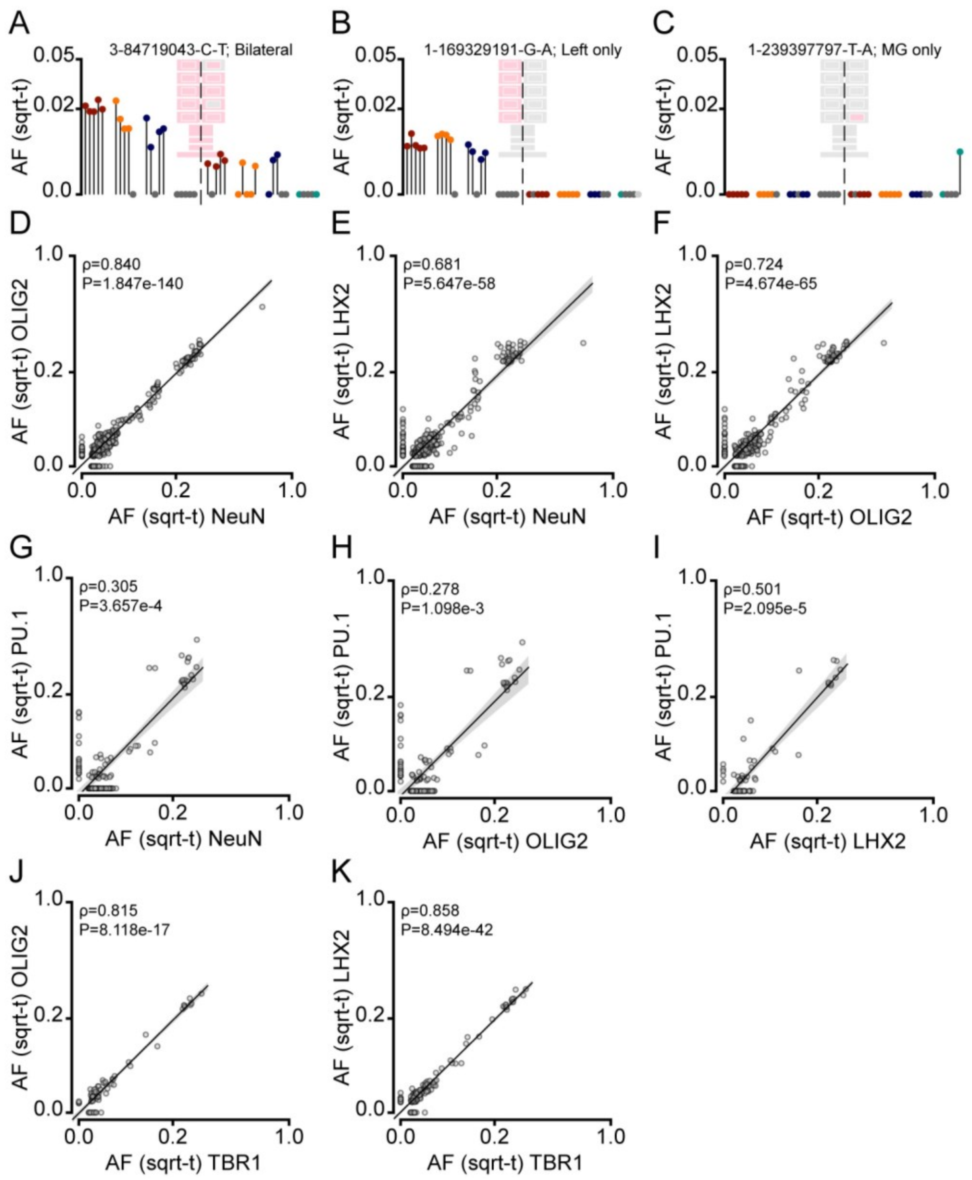


**Fig. S11. Variants exhibited differences within cell types that are reflected across the sorted populations.** (**A** to **C**) Lolliplots of examples showing a bilateral, unequally distributed (A), left lateralized (B), and a microglia-enriched population-only (C) variant. (**D** to **K**) Correlation plots of all sorted populations for each variant and region where high-quality data was available (i.e. included population and read depth >1,000×). Spearman correlation's ρ and P-value are shown for each pair-wise comparison.

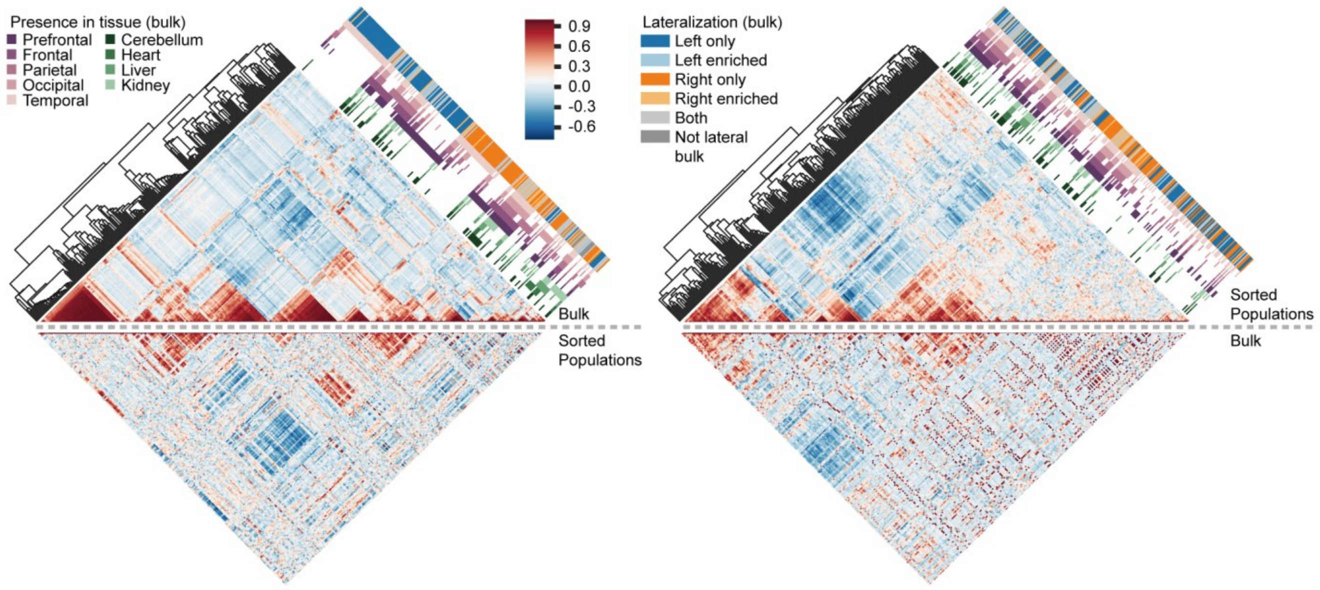


**Fig. S12. Correlations of AFs in bulk tissues and sorted populations highlight different features of the mosaic variants.** Correlation plots with hierarchical clustering based on the Pearson correlation coefficients between AFs measured in different bulk tissues (Bulk) or sorted cellular fractions (Sorted Populations). AFs were assessed by MPAS and calculated between all possible combinations from the 259 detected variants, as described in Figure 1K. Color codes show the left-right distribution of the variant and in which tissue the variants were detected on the level of bulk tissues. Upper half of the diamond is the correlation used to determine the order in the lower half of the diamond. The two correlations show that bulk tissue analysis and sorted cellular fraction analysis contain overlapping but distinct information.

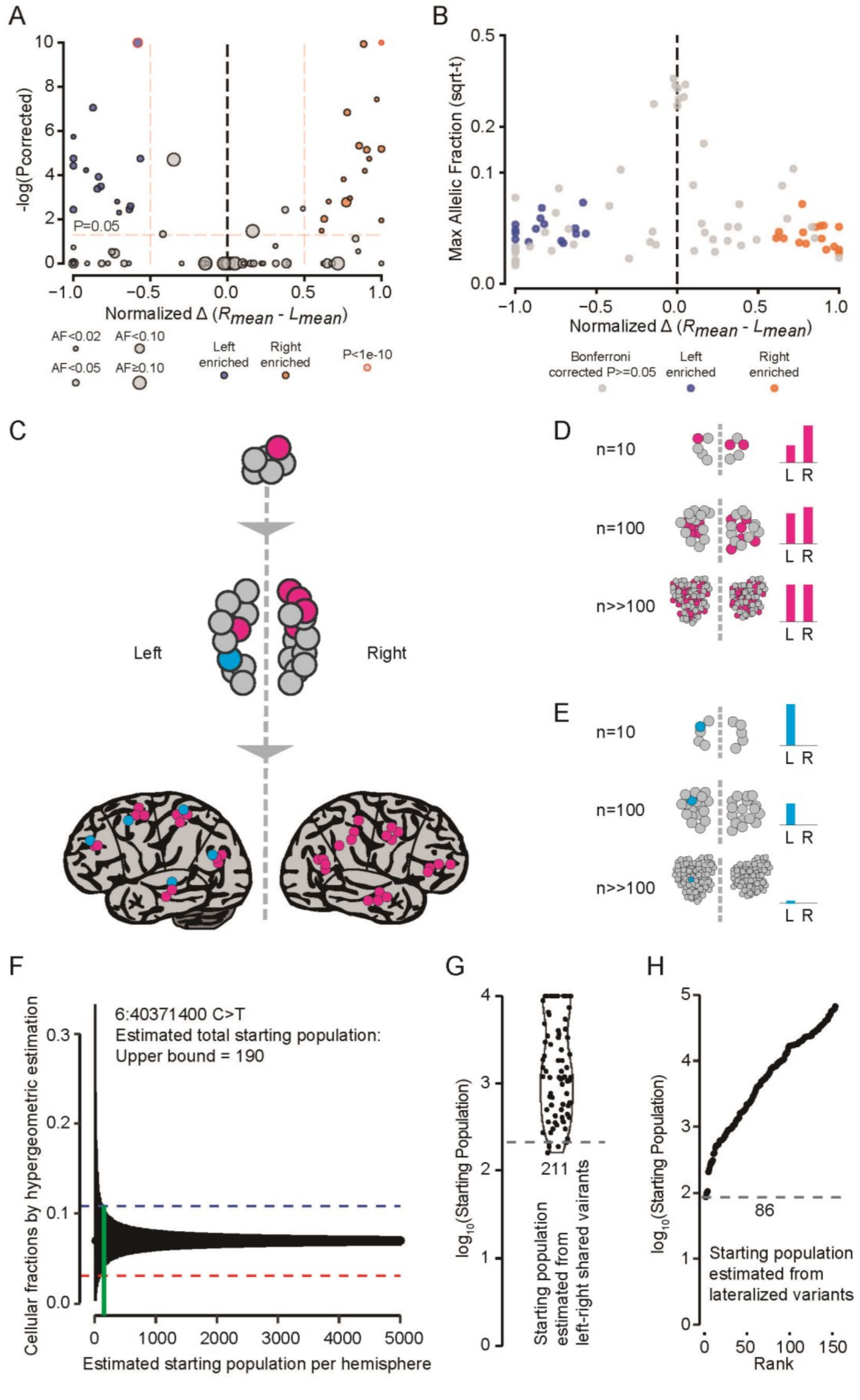
A
-log(Pcorrected)
P=0.05
Normalized Δ (Rmean - Lmean)
AF<0.02
AF<0.10
AF<0.05
AF≥0.10
Left enriched
Right enriched
P<1e-10
B
Max Allelic Fraction (sqrt-t)
Normalized Δ (Rmean - Lmean)
Bonferroni corrected P>=0.05
Left enriched
Right enriched
C
Left
Right
D
n=10
n=100
n>>100
L R
E
n=10
n=100
n>>100
L R
F
6:40371400 C>T
Estimated total starting population:
Upper bound = 190
Cellular fractions by hypergeometric estimation
Estimated starting population per hemisphere
G
log10(Starting Population)
211
Starting population estimated from left-right shared vairants
H
log10(Starting Population)
86
Starting population estimated from lateralized variants
Rank

**Fig. S13. Statistical modeling estimates an effective population size of ~90-200 progenitors immediately upstream of the left-right partition.** (**A**) Normalized difference of mosaic variants between their average AF ($R_{mean}$ and $L_{mean}$) of sorted brain-derived cells (i.e. non-PU.1$^{+}$) on the left and right hemisphere (Normalized Δ; see Materials and Methods) and their negative $\log_{10}$ P-value when comparing the individual values on both sides (Two-way ANOVA for side, using side and sorted cell type as two independent variables; Bonferroni-corrected). Size of markers, fill-color, and edge-color indicate a variant's $AF_{max}$, significant lateralization, and $P<10^{-10}$, respectively and as indicated. Enrichment is determined by a $P<0.05$ and a Normalized Δ of below -0.5 or above 0.5. (**B**) Allelic fractions of variants enriched in either hemisphere. X-axis is the same as in A and Y-axis is the $AF_{max}$ of a variant. Color indicates enrichment as in A. (**C**) Schematic of variants occurring during development, before and after lateralization. Red: cells with variants occurring during very early development stages before brain lateralization. These variants will be distributed differently in both hemispheres and might also be shared by non-brain tissues. Blue: bells with variants that occur after the left-right split will be detected only in one hemisphere. These two types of variants can be used for the estimation of the effective starting populations. For these analyses the sorted populations of neurons and neuroglia have to be used to exclude signal from cell types that do not originate from the brain. (**D**) AF quantified from the left and right hemisphere of the red variants: the larger the predicted starting population at time of the left-right separation is, the smaller the expected AF differences will be. (**E**) AF quantified for one-hemisphere-specific variants; the smaller the population immediately after the left-right separation, the higher AF will be observed for lateralized variants. (**F**) An example variant used for the estimation of the maximal effective population size supported by the observed difference between left and right (95% bands of a hypergeometric distribution are plotted in black). Blue and red dash line: average AF measured in both hemispheres. Green line: upper bound of the estimated starting population. (**G**) Upper bound of the starting population estimated from all variants shared in both hemispheres, by non-brain organs, or both, suggesting that they were present before the left-right split. The 5% percentile for all the estimated variants was 211, the lowest estimation was 160. (**H**) Minimum Starting population estimated from all variants unique to one hemisphere; the smallest estimated number was 86. Together, this allows to estimate that the effective founder population prior to the left-right partition was ~86-211 progenitors.

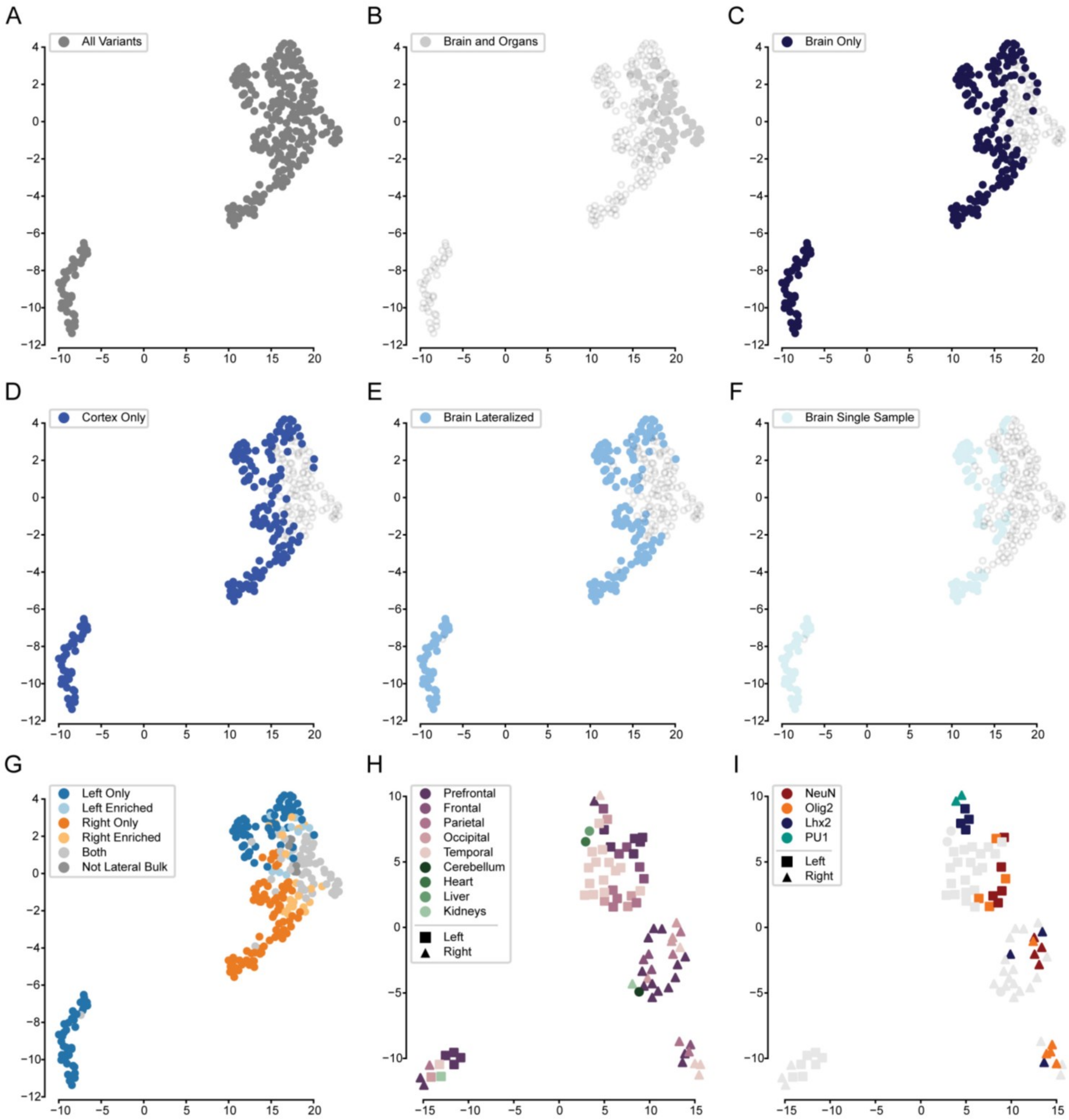


**Fig. S14. UMAP embeddings of mosaic variants and tissue samples.** (**A** to **G**) UMAP embeddings of mosaic variants (n=259) across 79 samples using the considered AF for tissues. Variants are colored according to the presence of the variant across the body (B), in the brain or cortex only (C and D), if the variant is restricted to one side of the brain (E), and if the variant was localized to one brain sample (F). Lateralized variants cluster together as shown in G. (**H** and **I**) When performing dimensionality reduction with respect to samples (i.e. transposing the input matrix for A to G), samples cluster according to lateralization (left body: square points; right body: triangle points; no lateralization: circle points) as well as anatomical location. When coloring points according to the cell marker used for nuclear sorting (I), microglia (PU.1) and astrocytes (LHX2), in contrast to oligodendrocytes (OLIG2) and neurons (NeuN), cluster among themselves.

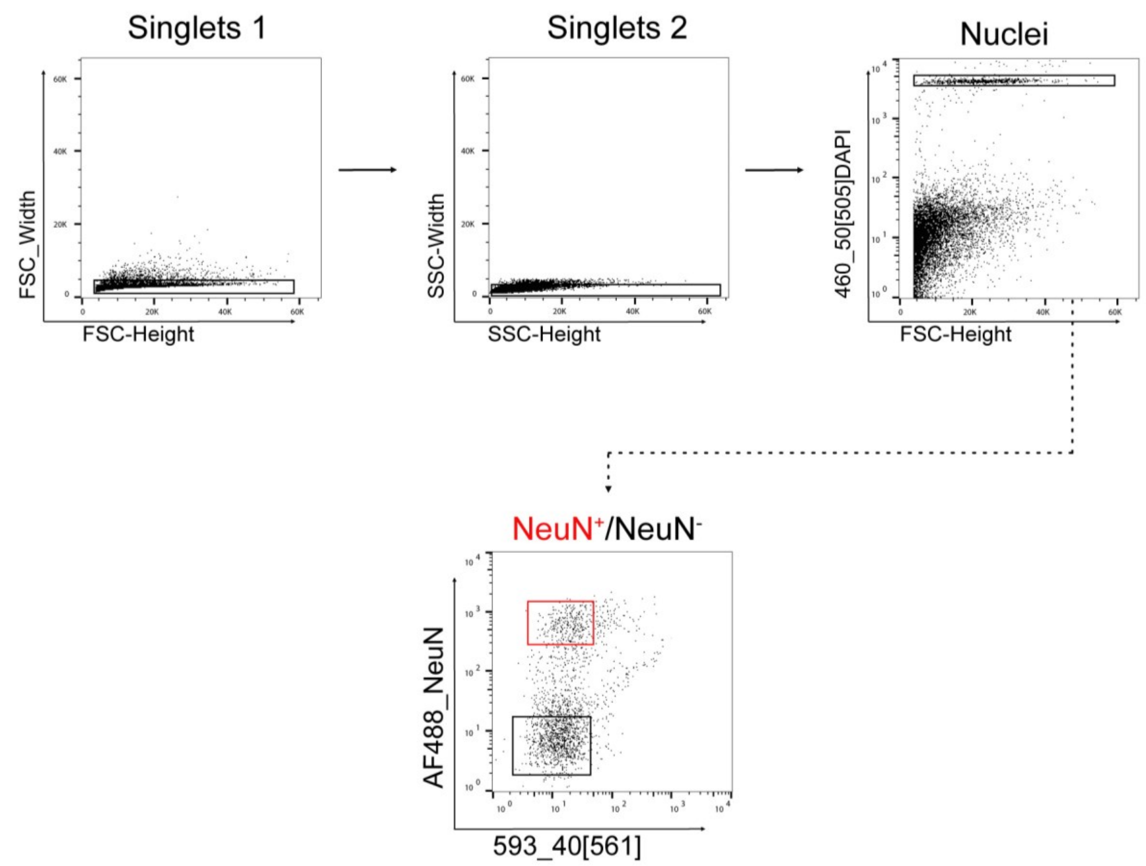


**Fig. S15. Sorting of neuronal and non-neuronal single nuclei for snMPAS.** Gating strategy. Doublets were excluded and nuclei positive for DAPI were further gated into NeuN$^+$ and NeuN$^-$ populations. 48 DAPI$^+$/NeuN$^+$ and 47 DAPI$^+$/NeuN$^-$ nuclei were sorted into a 96-well plate.

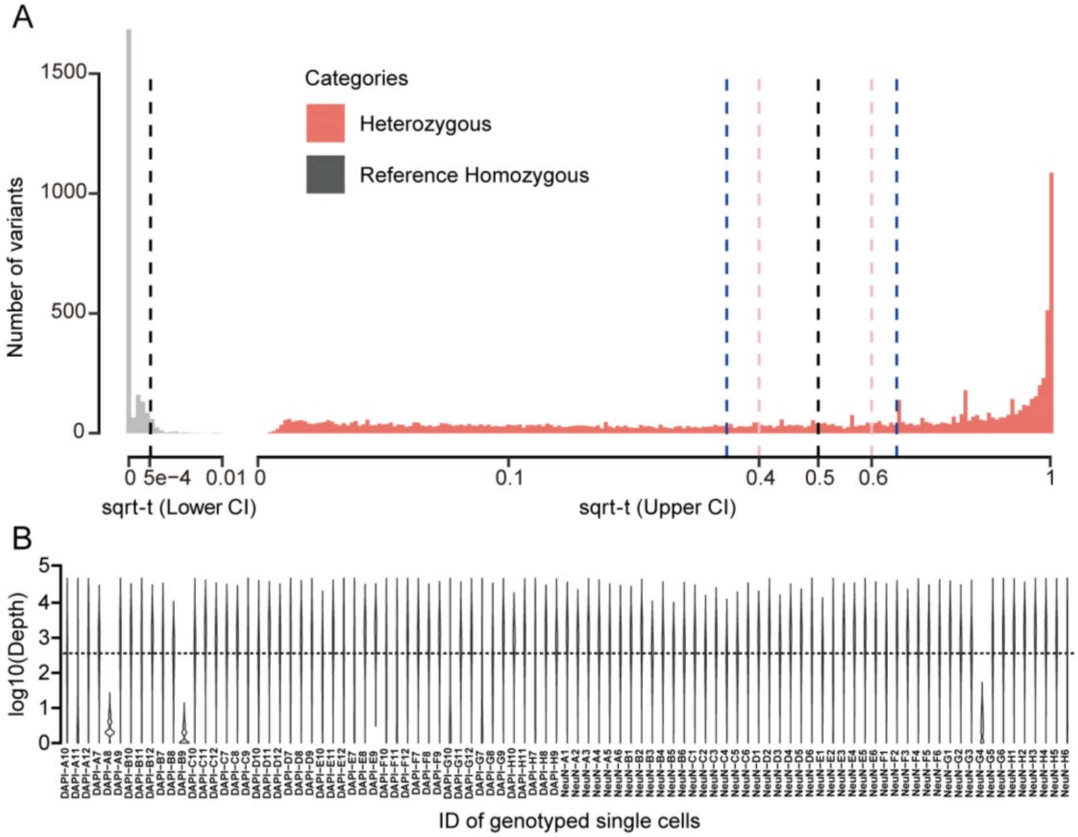


**Fig. S16. Quality control for snMPAS.** (**A**) Distribution of estimated exact binomial lower bounds of allelic fractions from 27 reference homozygous variants and upper bounds from 113 heterozygous variants in the snMPAS panel; the single-tail 95% cut off for reference homozygous variants is 5.43e-4 (black), AF 0.4 and 0.6 cutoff for heterozygous variants were labeled pink and 0.3% and 0.65 cutoff for heterozygous variants were labeled blue. The AF is square root transformed. (**B**) Depth distribution of all the 1349 candidate genomic positions designed for snMPAS for each sequenced single nuclei from the donor. The depth is $\log_{10}$ transformed. NeuN: NeuN+ nuclei; DAPI: DAPI+/NeuN- nuclei.

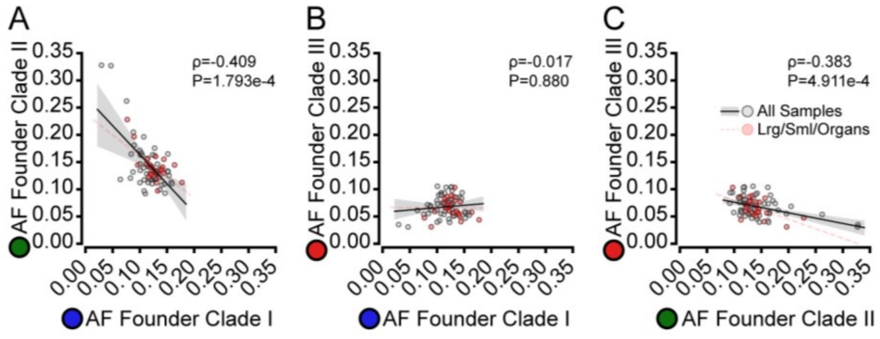


**Fig. S17. Founder variants from Clades I and II are significantly anti-correlated.** (**A to C**) Correlation of the founder variants from clades I to III with each other for all samples (grey) or only the 25 bulk tissues (red). Spearman correlation's ρ and P-value across all samples are shown for each pair-wise comparison.

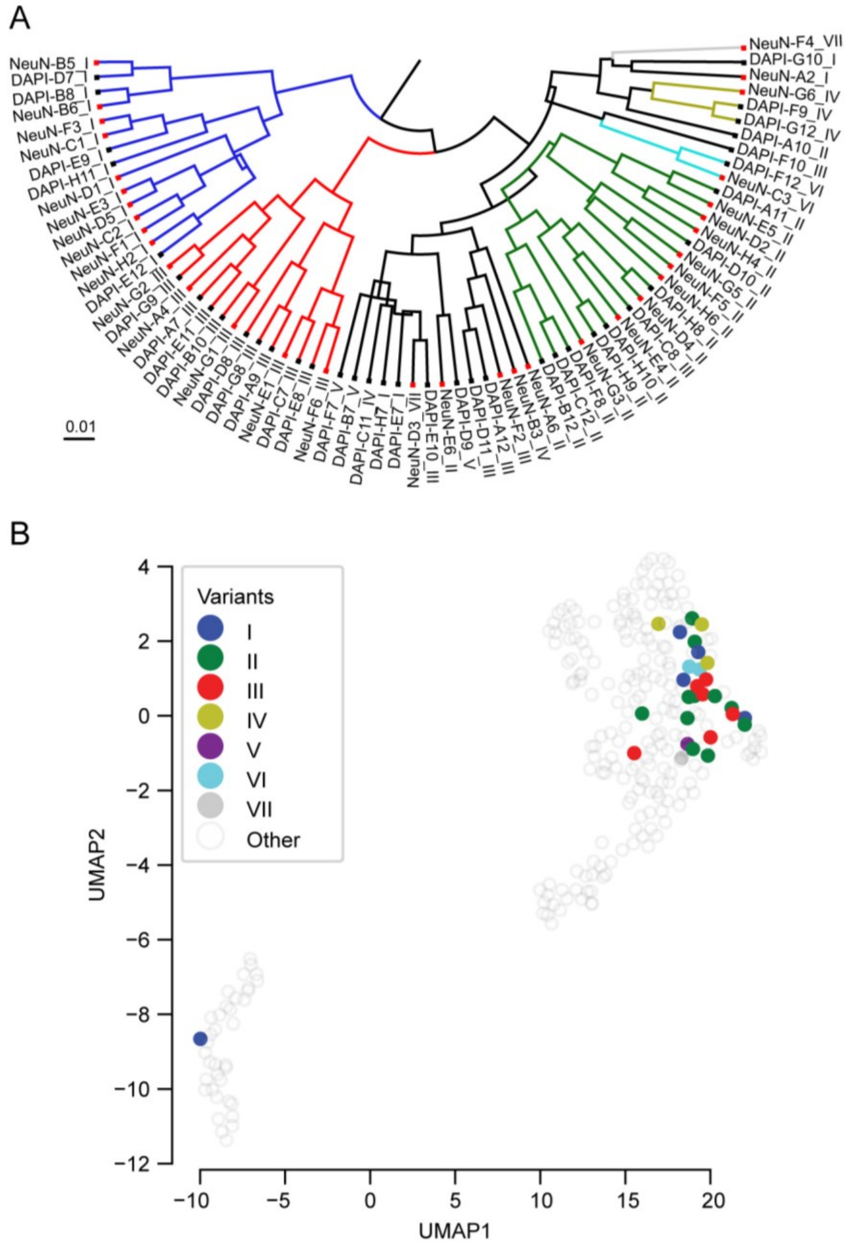


**Fig. S18. BEAST analysis and UMAP embeddings of clades defined in snMPAS.** (**A**) Lineage tree for all considered cells (n=71) and variants (n=33) showing relatedness of cells to each other. Representative tree was constructed using the maximum clade credibility method while branch colors represent inferred clades shown in Fig. 4B. Scale bar represents the expected substitutions per site as a function of branch length. (**B**) UMAP embeddings constructed in the same manner as Fig. S14A-G. Variants are colored according to the major clade determined from single nuclei samples.

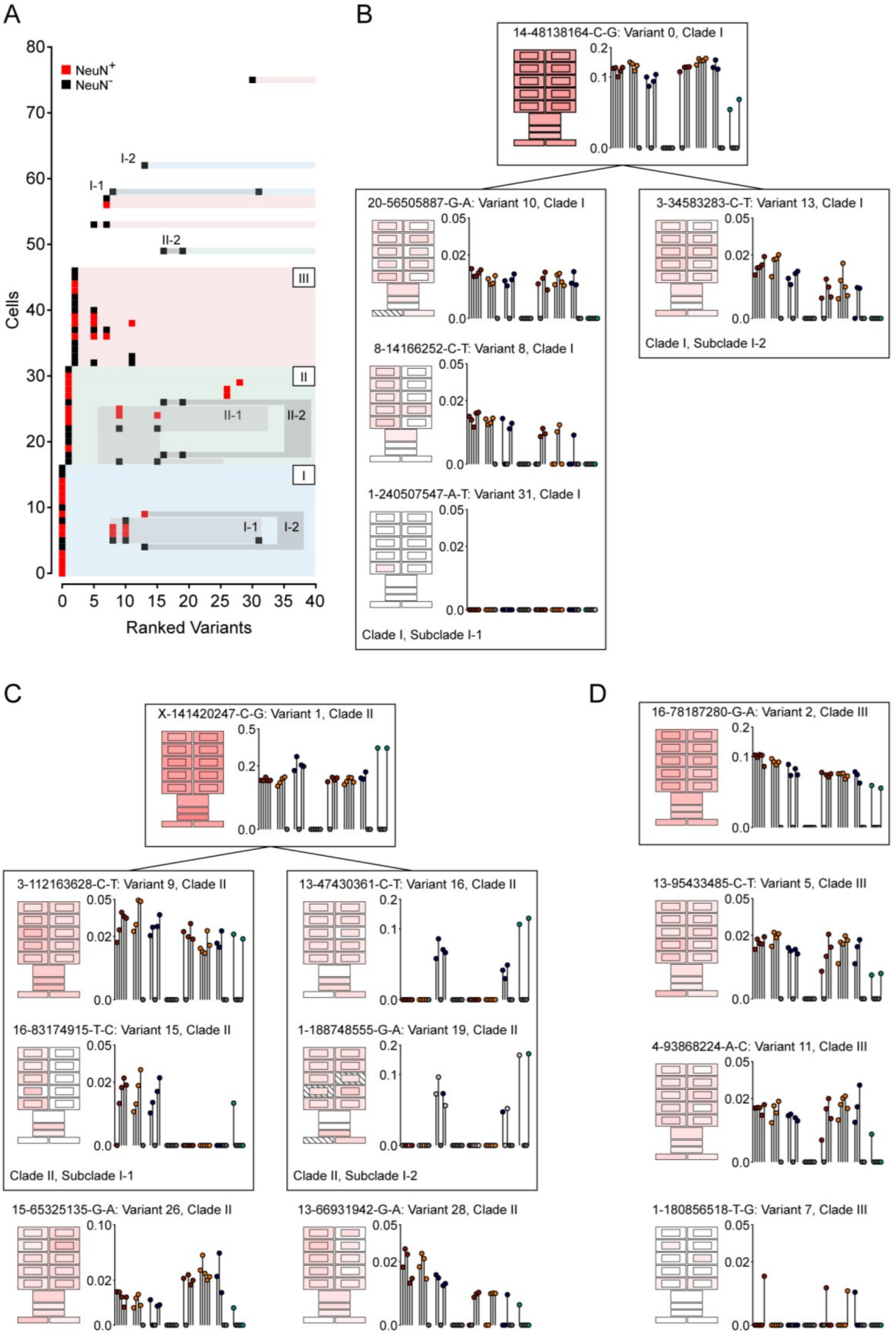
A
B
C
D
NeuN+
NeuN-
Cells
Ranked Variants
I-1
I-2
II-1
II-2
I
II
III
14-48138164-C-G: Variant 0, Clade I
20-56505887-G-A: Variant 10, Clade I
3-34583283-C-T: Variant 13, Clade I
8-14166252-C-T: Variant 8, Clade I
1-240507547-A-T: Variant 31, Clade I
Clade I, Subclade I-1
Clade I, Subclade I-2
X-141420247-C-G: Variant 1, Clade II
3-112163628-C-T: Variant 9, Clade II
13-47430361-C-T: Variant 16, Clade II
16-83174915-T-C: Variant 15, Clade II
1-188748555-G-A: Variant 19, Clade II
Clade II, Subclade I-1
Clade II, Subclade I-2
15-65325135-G-A: Variant 26, Clade II
13-66931942-G-A: Variant 28, Clade II
16-78187280-G-A: Variant 2, Clade III
13-95433485-C-T: Variant 5, Clade III
4-93868224-A-C: Variant 11, Clade III
1-180856518-T-G: Variant 7, Clade III

**Fig. S19. The three major clades and their variants are widely distributed and contain restricted subclones.** (**A**) Replica of the ranked plot of filtered, mosaic variants in Fig. 4B highlighting only those variants visualized in this figure. (**B to D**) Founder variants of the three major clades and example variants. Also shown are suspected subclades (B and C) for Clade I and II. Geoclones and lolliplots represent data as shown in Fig. 2A and Fig. 3C.

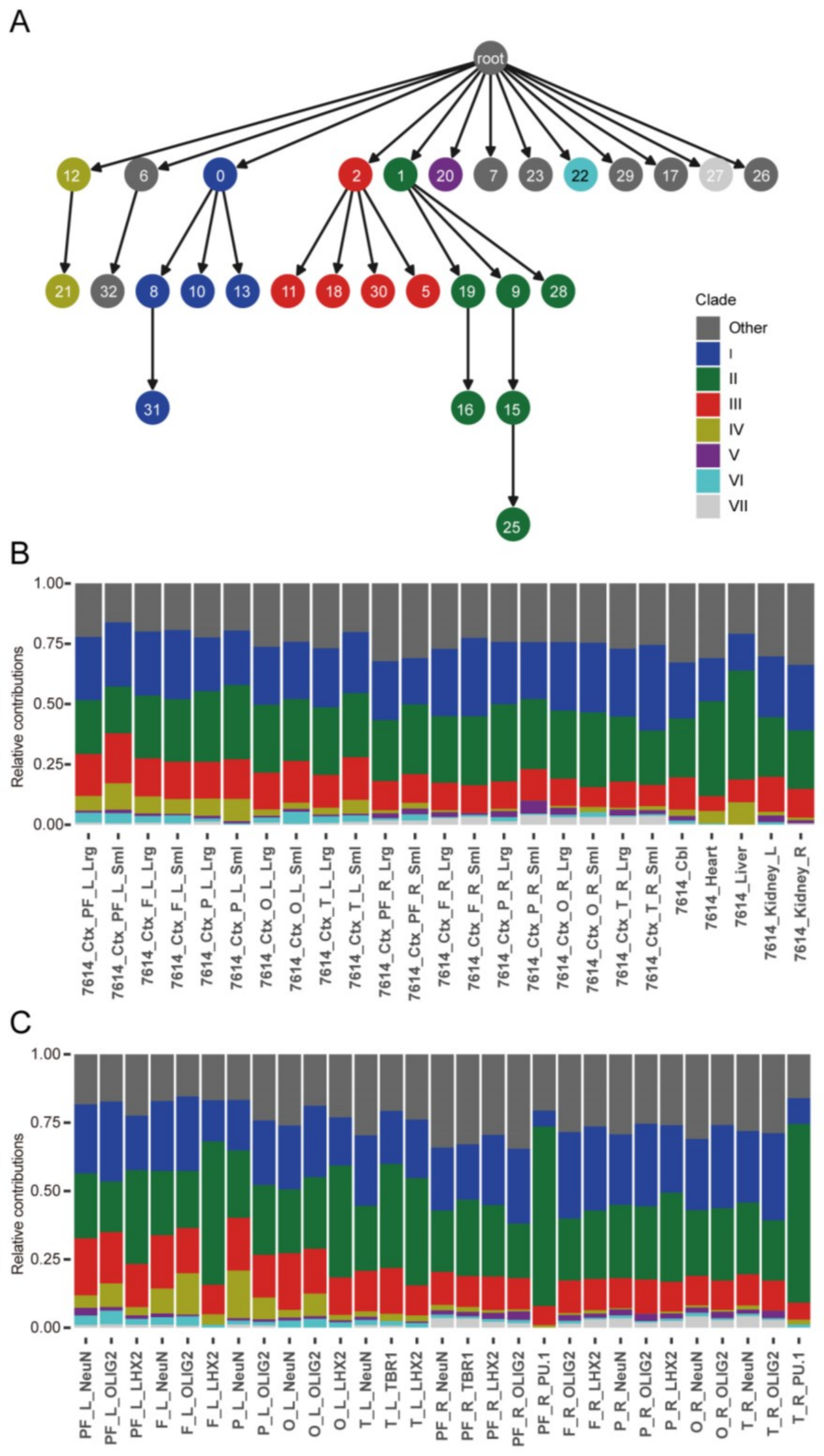


**Fig. S20. Reconstructing of cellular lineages and the deconvolution of contributions from each major lineage.** (**A**) Somatic variants correctly genotyped by snMPAS as well as the AF information from bulk MPAS were used to reconstruct lineages, which were colored according to Fig. 4C. (**B**) Relative contribution of variants labeled in each lineage group presented in A were calculated through a linear regression model. A mean squared error method was used to optimize the estimation so that the weighted sum of all predicted lineages reflected the allelic fractions measured in the 25 bulk tissues. (**C**) Relative contribution of lineages from each clade for all sorted populations.

**Data S1. Raw mosaic SNV/INDEL calls from the 300× WGS. (Separate file)**
Raw mosaic SNV/INDEL calls from the 300× WGS. Information includes the genomic position, reference and alternative alleles, as well as caller agreement on the specific variant. The candidate variant list subjected to MPAS and snMPAS panel design and the considered regions are included as separate sheets.

**Data S2. MPAS and snMPAS genotyping and quantification results. (Separate file)**
MPAS and snMPAS genotyping and quantification results.

**Data S3. Detailed visual representation for each of the 259 variants. (Separate file)**
Geoclones for bulk samples and geographic subsamples as well as lolliplot representations for all of the 259 positively detected mosaic variants based on MPAS results. Plots are further explained in Figure 2 and 3.

**Data S4. Visual representation of snMPAS results for each of the 259 variants. (Separate file)**
Visualized AF for all 259 positively detected mosaic variants in the 95 sorted single nuclei based on snMPAS results. Y-axis shows the square root transformed AF. Dots and error bars show the calculated AF and an exact binomial 95% CI. Cellular ID shows the individual's ID (7614), its origin, and whether or not a cell was $NeuN^+$ (NeuN) or $NeuN^-$ (DAPI). Each cell is further identified by its well position (A01-H11). Title of each page represents the variant ID (chromosome-position-reference-alternative).
